# Characterizing the Emergence of Liver and Gallbladder from the Embryonic Endoderm through Single-Cell RNA-Seq

**DOI:** 10.1101/718775

**Authors:** Tianhao Mu, Liqin Xu, Yu Zhong, Xinyu Liu, Zhikun Zhao, Chaoben Huang, Xiaofeng Lan, Chengchen Lufei, Yi Zhou, Yixun Su, Luang Xu, Miaomiao Jiang, Hongpo Zhou, Xinxin Lin, Liang Wu, Siqi Peng, Shiping Liu, Susanne Brix, Michael Dean, Norris R. Dunn, Kenneth S. Zaret, Xin-Yuan Fu, Yong Hou

## Abstract

The liver and gallbladder are among the most important internal organs derived from the endoderm. Several inductive signals regulate liver development, yet the pure nascent hepatic and gallbladder cells are unable to be isolated due to limited cell markers and cell numbers. The transcriptome networks of the hepatic lineage in the endoderm, and how the gallbladder differentiates from the adjacent endoderm population, are not fully understood. Using a transgenic Foxa2^eGFP^ reporter mouse line, we performed deep single-cell RNA sequencing on 1,966 individual cells, including nascent hepatic and gallbladder cells, isolated from the endoderm and hepatic regions from ten embryonic stages, ranging from day E7.5 to E15.5. We identified the embryonic liver developmental trajectory from primitive streak to hepatoblasts and characterized the transcriptome of the hepatic lineage. During pre-hepatogenesis (5-6 somite stage), we have identified two groups of foregut endoderm cells, one derived from visceral endoderm and the second derived from primitive streak via a mesenchymal-epithelial transition (MET). During the liver specification stages, liver primordium was identified to share both foregut and liver features. We also documented dynamic gene expression during the epithelial-hepatic transition (EHT). Six gene groups were found to switch on or off at different stages during liver specification. Importantly, we found that RXR complex signaling and newly identified transcription factors associated with liver specification. Moreover, we revealed the gallbladder primordium cells at E9.5 and found genes that transcriptionally distinguish them from the liver primordium. The present data provides a high-resolution resource and critical insights for understanding the emergence of the endoderm, liver and gallbladder development.

## Introduction

The liver is the largest internal organ and provides many essential metabolic, exocrine and endocrine functions including the production of bile, the metabolism of dietary compounds, detoxification, regulation of glucose levels and control of blood homeostasis through secretion of clotting factors and serum proteins such as Albumin (*Alb*) ^1^. After gastrulation, the foregut endoderm is derived from the primitive streak (PS) at mouse embryonic day 7.5 of gestation (E7.5) ^2^. The liver is derived from the foregut endoderm, and the hepatic marker *Alb* is first detected in the nascent hepatic endoderm within the 7-8 somite stage at E8.5 ^3, 4^. *Foxa2* has been considered as an endoderm marker at E6.5 and is expressed in all the differentiated endoderm-derived organs including the liver ^5^. FOXA2 acts as a “pioneer” factor in liver development and serves to de-compact chromatin at its target sites ^6^. Disruption of FOX factors (*Foxa2, Foxh1*), GATA factors, *Sox17*, *Mixl1* or SMAD signaling all lead to defects in gut tube and liver morphogenesis ^7–12^. During liver specification, a portion of the gut tube cells receives fibroblast growth factor (FGF) signals from the developing heart ^3^ and bone morphogenetic protein (BMP) from the septum transversum mesenchyme (STM)^13^. This leads to differentiation of the hepatoblast, which constitutes the liver primordium or liver bud at E10.5 ^14, 15^. Several transcription factors (TFs) have shown to be essential for liver specification including TBX3, HNF4A, and PROX1 ^16–18^. Primarily, the program of hepatogenesis has been studied by conventional immunohistochemistry and analysis of tissue explants, however, a complete pattern of transcriptional dynamics of the process remains to be unveiled, due to the difficulties of isolation of pure nascent hepatic progenitors.

The liver primordium, primitive gallbladder, and primitive pancreas arise from the foregut endoderm at almost the same time at E9.5 ^19–21^. The PDX1+ and SOX17+ pancreatobiliary progenitor cells segregate into a PDX1+/SOX17-ventral pancreas and a SOX17+/PDX1-biliary primordium ^22^. In another study, *Lgr4* has been shown to be significant for gallbladder development, since *Lgr4* depletion affects the elongation of the gallbladder but has no effect on the liver bud and ventral pancreas ^23^. Apart from such studies, the molecular features and drivers of gallbladder development are unexplored.

Recently, single-cell RNA sequencing has been used to study hepatic tissue differentiated from the induced pluripotent stem cells (iPSC) ^24^. In that study, iPSCs were made into endoderm and liver bud, but gallbladder development was absent, leaving unclear the patterning discrimination between the tissues. Two other studies focused on liver differentiation from E10.5 or 11.5 onwards, and discerned the split between the hepatocyte and cholangiocyte lineages ^25, 26^. However, early developmental stages of liver differentiation from the foregut endoderm has not been described. In this study, we constructed a transgenic Foxa2^eGFP^ reporter mouse line to trace the endodermal and hepatic cells in the early stages of development. By applying single-cell RNA-sequencing of 1,966 single-cells from endodermal and hepatic regions from E7.5 to E15.5, we have identified the endoderm and liver developmental lineages and characterized the key networks and transcription factors responsible for endoderm morphogenesis and liver development. We also identified the gallbladder primordium at E9.5 and found it could be distinguished transcriptionally from liver primordium. Our data provides a resource for further research into endodermal differentiation and liver development, which could potentially lead to therapeutically useful tissue for liver transplantation.

## Results

### Foxa2^eGFP^ tracing of endoderm and hepatic cells and scRNA-sequencing

To access purified endodermal and hepatic-related cells, we generated a transgenic Foxa2^eGFP^ reporter mouse line (Figure S1). In this mouse model, enhanced green fluorescent protein (*eGFP*) is linked to the third exon of *Foxa2* (Figure 1A). Homozygous transgenic mice develop normally and do not show an abnormal phenotype. As expected for the endogenous *Foxa2* gene^27–29^, we found eGFP to be expressed in the mouse embryo labeling the endoderm, neural system and endoderm-derived organs including the liver (Figure 1B, 1C). The fluorescence in the liver was impaired because of the perfusion of hematopoietic cells from E11.5, but fluorescence was evident upon liver dissection. Immunofluorescence assay showed that hepatoblasts expressed FOXA2 and DLK1 at E14.5 (Figure 1D). We dissected the distal half part of the whole embryo at E7.5 and the foregut endoderm at E8.5, the hepatic region from E9.5, E10.0 and E10.5, and the whole liver from E11.5, E12.5, E13.5, E14.5 and E15.5, including two replicates at E12.5, E13.5 and E14.5. (Figure 1B). At E11.5, the liver was precisely dissected excluding the pancreas, lung, and stomach (Figure 1C).

**Figure.1.**
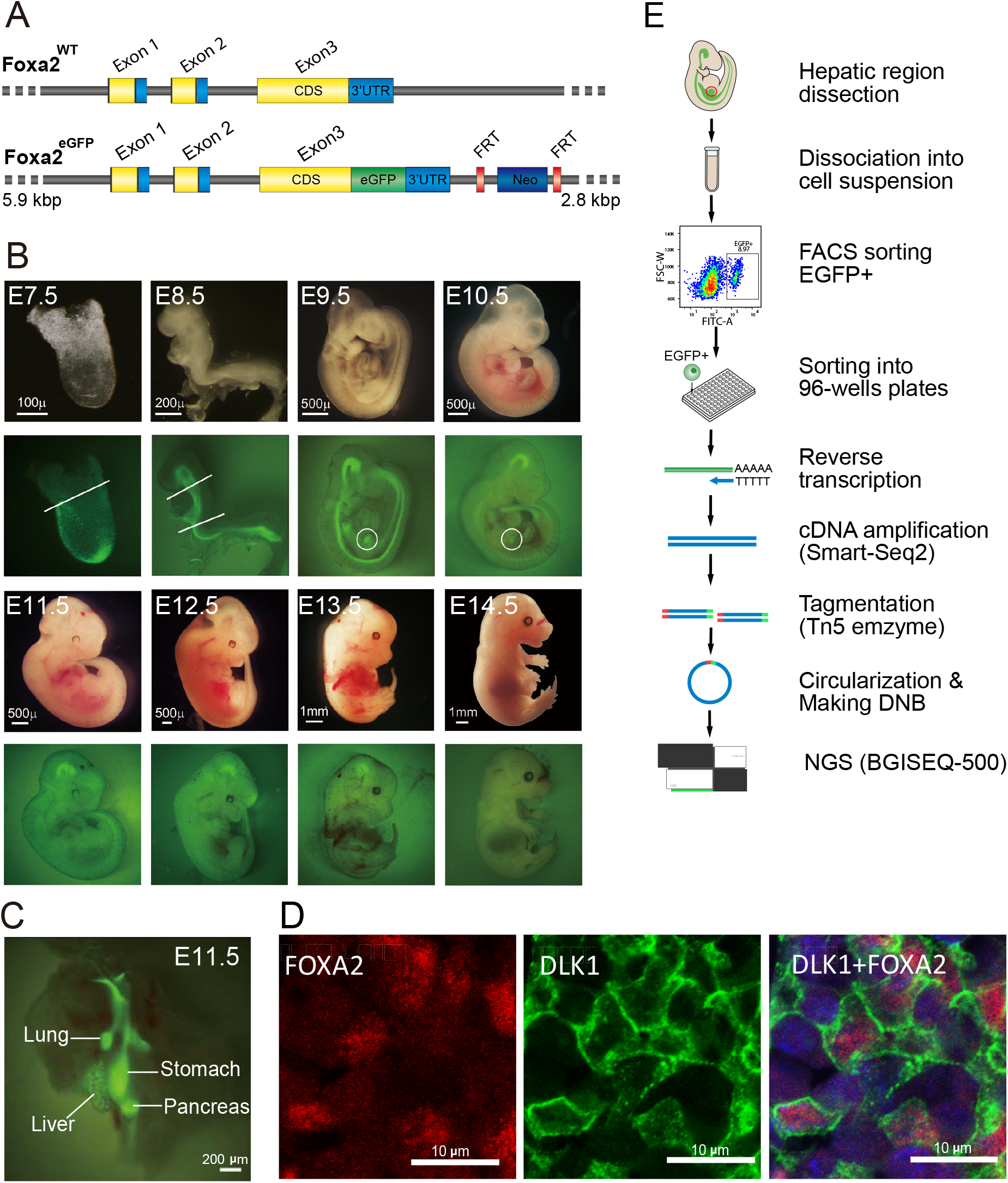
Single-cell RNA-seq to analyze liver development during E7.5-E15.5 by using a Foxa2^eGFP^ mouse model. A, Vector design for Foxa2^eGFP^ mouse. eGFP is linked to the third exon of *Foxa2*. B, eGFP labeled mouse embryos from E7.5-E14.5. The endoderm, neural system and endoderm-derived organs including liver expressed eGFP. The general dissection strategies are shown (white lines or circles). C, Precise dissection for the endodermal organs at E11.5. Lung, liver, stomach and pancreas are shown. D, Immunofluorescence analysis of paraffin-sectioned mouse embryo at E14.5, showing co-expression of Foxa2 (red), DLK1 (green) and DAPI (blue) in hepatoblasts. E, Workflow of single-cell RNA sequencing of Foxa2^eGFP^+ cells.

To characterize endoderm and liver development, we performed single-cell RNA-Seq experiments on the Foxa2^eGFP^+ cells isolated from E7.5-E15.5 (Figure 1E). The tissues were dissociated into a single-cell suspension and Foxa2^eGFP^+ cells were sorted into 96 well plates with one cell in each well using a FACSAria III. Doublets and multiplets were excluded by analysis of side scatter (SSC) and forward scatter (FSC) (Figure S2). The amplified cDNA was assessed by agarose gel and qPCR of *Afp*, a hepatic marker, before library generation (Figure S3). In total, 1,246 individual cells were collected and the mRNA amplified following the SMART-seq2 protocol. In addition, we generated libraries from 720 cells from E11.5, E12.5 and E13.5 by MIRALCS (microwell full-length mRNA amplification and library construction system)^30^. Altogether, the transcriptomes of 1,966 individual cells as well as bulk control samples from 10 embryo stages were sequenced.

The 1,246 SMART-seq2 cells were used to identify cell populations during liver development. After filtering unqualified reads, gene expression levels were characterized by RPKM (Reads Per Kilobase per Million mapped reads) with RPKM>1 as the threshold (Figure S4). To obtain high-quality cells for subsequent analysis, we removed cells with fewer than 6,000 expressed genes (RPKM>1), since most of those cells express low levels of *Gapdh* (Figure S5B). We obtained 922 cells with an average of 9,378 genes with RPKM>1 and an average of 9.5 million mapped sequencing reads (Figure S5A, S5C). Then we excluded 321 (35.9%) cells that did not express *Foxa2* or *eGFP* mRNA, to minimize the issues of protein perdurance from a prior cell state. Finally, 601 cells that were both *Foxa2* and *eGFP* positive (RPKM>1) with high quality data were used in the cell clustering (Table S1).

Technical noise was assessed by bulk sample sequencing, experimental replicates, and sequencing batch effect analysis (Figure S6A-D), confirming that the final dataset was of high quality and reliable. As *eGFP* and *Foxa2* were co-expressed in our mouse model, a high correlation was detected (Pearson r=0.95) between these two genes (Figure S6E).

### Identification of primitive streak and heterogeneity of foregut endoderm

To identify the primitive streak in E7.5 and the foregut endoderm in E8.5, we clustered Foxa2^eGFP^+ (RPKM>1) cells from E7.5 and E8.5 and visualized them by t-SNE (T-distributed Stochastic Neighbor Embedding) (Figure 2A). A total of five groups of cells were identified as primitive streak (PS), foregut endoderm (FG), visceral endoderm (VE), neural tube (NT) and notochord (NC) based on marker expression (Figure 2B, 2C). The primitive streak cells were identified at E7.5 based on the expression of genes related to gastrulation, including *Pou5f1, Mixl1, Lefty2, Cer1, Cyp26a1, Lhx1, Fgf5, Hesx1* and *Snail1* (Figure 2B). We validated the primitive streak cells using the iTranscriptome database ^31^, and all of these cells were predicted to be located at the posterior side of the embryo endoderm, which corresponded to the PS region, hence supporting our cell taxonomy (Figure 2D).

**Figure 2.**
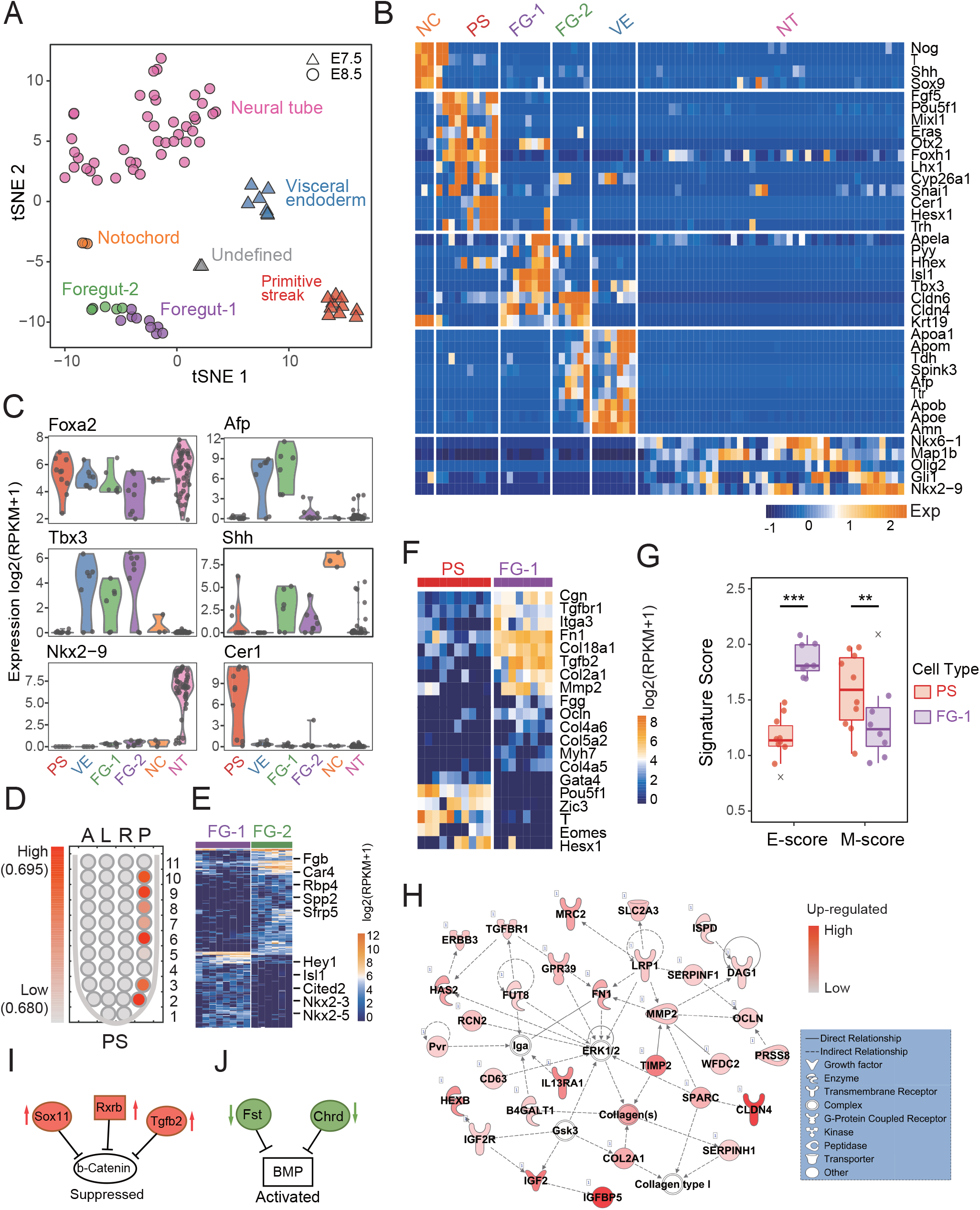
Extracellular matrix molecules and MET process during foregut morphogenesis at E7.5-E8.5 A, Single-cells from E7.5 and E8.5 were clustered by t-SNE into 5 groups, Primitive Streak (PS), Visceral endoderm (VE), Foregut1/2 (FG), Neural tube (NT) and Notochord (NC). B, Specific markers for each cell group in Figure 2a. C, Violin plots of the gene expression levels of specific markers, *Foxa2, Afp, Tbx3, Shh, Nkx2-9* and *Cer1* for each cell group are shown. D, Genes in Primitive streak group were validated by iTranscriptome database. A, anterior; P, posterior; L, left; R, right; E, Differentially expressed genes between foregut-1 and foregut-2 group. FG-1, foregut-1; FG-2, foregut-2. F, Differentially expressed genes between Primitive streak and foregut-1group. PS, Primitive Streak; FG-1, foregut-1. G, The mesenchymal features (M score) and epithelial features (E score) of primitive streak and foregut-1 in mesenchymal-epithelial transition (MET) progress during E7.5-E8.5. H, Up-regulated genes enriched in the network related with cellular motility and connective tissue development in Foregut-1 group, with ERK1/2 located in the center of the network. I, The Wnt/β-catenin signaling pathway was identified to be repressed by *Sox11, Rarb* and *Tgfb2* in the foregut. J, BMP signaling was activated by repressing *Fst* and *Chrd.*

A unique group of cells from E8.5 (5-6 somite stage) expressing *Pyy, Apela* and *Cldn4* was defined as foregut endoderm (Figure 2A, 2B). We noted that this stage preceded the 7-8 somite stage at which hepatic specification occurs ^4^. To study cell heterogeneity, we then re-clustered the 14 foregut endoderm cells and found they could be divided into two cell groups, the foregut-1(FG-1) and foregut-2(FG-2) (Figure 2A, 2B). Foregut-1 cells expressed high levels of *Tbx3* and *Hhex,* which are essential transcription factors for liver development, as *Hhex*−/− mice fail to develop the liver bud ^15^. Therefore, we conclude that foregut-1 cells differentiate into liver bud at later stages. By differential expression analysis, we found a group of genes related to embryonic development including *Hey2, Isl1, Cited2, Nkx2-3* and *Nkx2-5* to be enriched in the foregut-1 cells (Figure 2E). The foregut-2 cells expressed high levels of *Afp, Ttr* and apolipoprotein genes which together are characteristic of visceral endoderm. Approximately half of the foregut-2 cells expressed *Amn*, which is known to be expressed in the extraembryonic visceral endoderm layer during gastrulation. These results indicate that foregut-2 cells are derived from visceral endoderm. Thus, as previously reported, some of the descendants of visceral endoderm could be incorporated into the gut tube ^32^. Other genes including *Fgb, Car4, Spp2, Sfrp5* and *Rbp4* were enriched in the foregut-2 cells (Figure 2E, Table S4). However, further studies are needed to determine the cell fate of the foregut-2 cells and the roles they play during foregut development.

In addition to the primitive streak and foregut endoderm, we identified visceral endoderm, neural tube and notochord in E7.5 and E8.5 (Figure 2A, S7B) as they also express *Foxa2* ^27, 28^. Visceral endoderm was defined based on the expression of *Amn, Afp, Ttr, Tdh* and apolipoprotein genes (*Apob, Apoe, Apom* and *Apoa1*). Interestingly, 28 genes of the solute carrier family (*Slc* family) were highly expressed in visceral endoderm (Figure S7B). Neural tube was defined based on the expression of *Nkx2-9* and *Olig2*. As expected, notochord cells expressed *Nog, T, Shh* and *Sox9* (Figure 2B, 2B). In addition to these known cell marker genes, we have identified differentially expressed genes of these cell groups and characterized their respective functions (Figure S7A, S8, Table S3). To our knowledge, this is the first time that the single-cell transcriptome profile of these early embryonic cell types has been elucidated.

### Extracellular matrix molecules and MET progression in foregut morphogenesis

To characterize gene regulation during foregut morphogenesis, we compared the expression pattern of primitive streak and foregut-1 cells (Figure 2F, Table S5). Differentially expressed genes were analyzed by Ingenuity pathway analysis (IPA), indicating that the Wnt/β-catenin signaling pathway was repressed by *Sox11, Rxrb* and *Tgfb2* in the foregut, while BMP signaling was activated by down-regulation of the BMP suppressor *Fst* and *Chrd* (Figure 2I, 2J). These results are consistent with previous studies, which found that Wnt signaling initially suppresses mammalian liver induction ^33^, while BMP signaling helps induce it ^13, 34^. Moreover, we found that 6 transcription factor genes (*Pou5f1*, *T*, *Gata4*, *Eomes*, *Hesx1*, and *Zic3*) related with pluripotency had decreased expression in foregut endoderm, compared to primitive streak (Figure 2F), indicating that cell pluripotency decreases during gut morphogenesis. Interestingly, we found many epithelial factors and extracellular matrix (ECM) molecules including collagens (*Col18a1, Col2A1, Col4a5, Col4a6, Col5a2*), fibronectin (*Fn1, Mmp2*), fibrinogen (*Fgg*), integrin (*Itga3*) and tight junction proteins (*Cgn, Myh7, Ocln, Tgfb2, Tgfbr1*) to be up-regulated in the foregut endoderm compared to the primitive streak (Figure 2F). This result indicates that the primitive streak cells with a mesenchymal morphology are fated to become foregut endoderm with epithelium features through a mesenchymal-epithelial transition (MET) process. To support this hypothesis, we analyzed the mesenchymal features (M score) and epithelial features (E score) of the PS cells and FG-1 cells (Table S2). We found the PS cells displayed a high M score and a low E score, while the FG-1 cells displayed a low M score and a high E score (Figure 2G). By further analyzing the FG-1 cells, we found 35 up-regulated genes enriched in a network related to cellular motility and connective tissue development (Figure 2H). The ECM including MMP2, collagens and tight junction genes were present in this gene network, and ERK1/2 was found to be a central regulator of the network. ERK activation propagates in epithelial cell sheets and regulates migration ^35^. In summary, the results indicate that foregut morphogenesis is dependent on the MET signaling pathway by activation of ERK.

### An epithelial-hepatic transition (EHT) in nascent hepatoblasts

To characterize liver specification and budding processes, we re-clustered and visualized all the Foxa2^eGFP^+ cells from E9.5, E10.0 and E10.5 by t-SNE. The differentially expressed genes between each group were then identified (Figure S9, Table S6). Undifferentiated gut tube (GT) with epithelial features were identified to express high levels of *Epcam, Gata4* and *Shh* (Fig 3A, 3B). Differentiated hepatic cells were identified based on the expression of well-known marker genes like *Alb, Afp, Hnf4a, Hhex, Prox1* and *Dlk1*. *Epcam* decreased during the differentiation of hepatic cells, while *Shh* and *Gata4* were almost completely silenced during liver specification (Figure 3B), consistent with previous reports ^36, 37^. We found that the hepatic cells could be clustered into two groups and we defined them as liver primordium (LP) and liver bud (LB) by two criteria (Figure 3A). Firstly, most of the LP cells were found in E9.5 and E10.0, while the LB cells were mainly found at a later stage E10.5. Secondly, LP cells have higher expression of the epithelial marker *Epcam* but lower expression of the hepatic marker *Alb* compared with LB (Figure 3B). For confirmation, we quantified the hepatic features with a hepatic score using the expression of a set of genes related to hepatic functions (Table S2). By analyzing the hepatic and epithelial score of gut tube, liver primordium and liver bud, we found the epithelial score decreases while the hepatic score increases during liver specification. Liver primordium exhibited an intermediate state between the undifferentiated gut tube and differentiated liver bud (Figure 3D). The transitional process from endoderm with epithelial characteristics to the liver bud with hepatic characteristics was termed epithelial-hepatic transition (EHT), and is remarkably consistent with cytological changes reported during this period ^15^. During the EHT, LP presented both epithelial features and hepatic features, indicating that LP cells were the nascent hepatic cells that differentiated from the gut tube.

**Figure.3.**
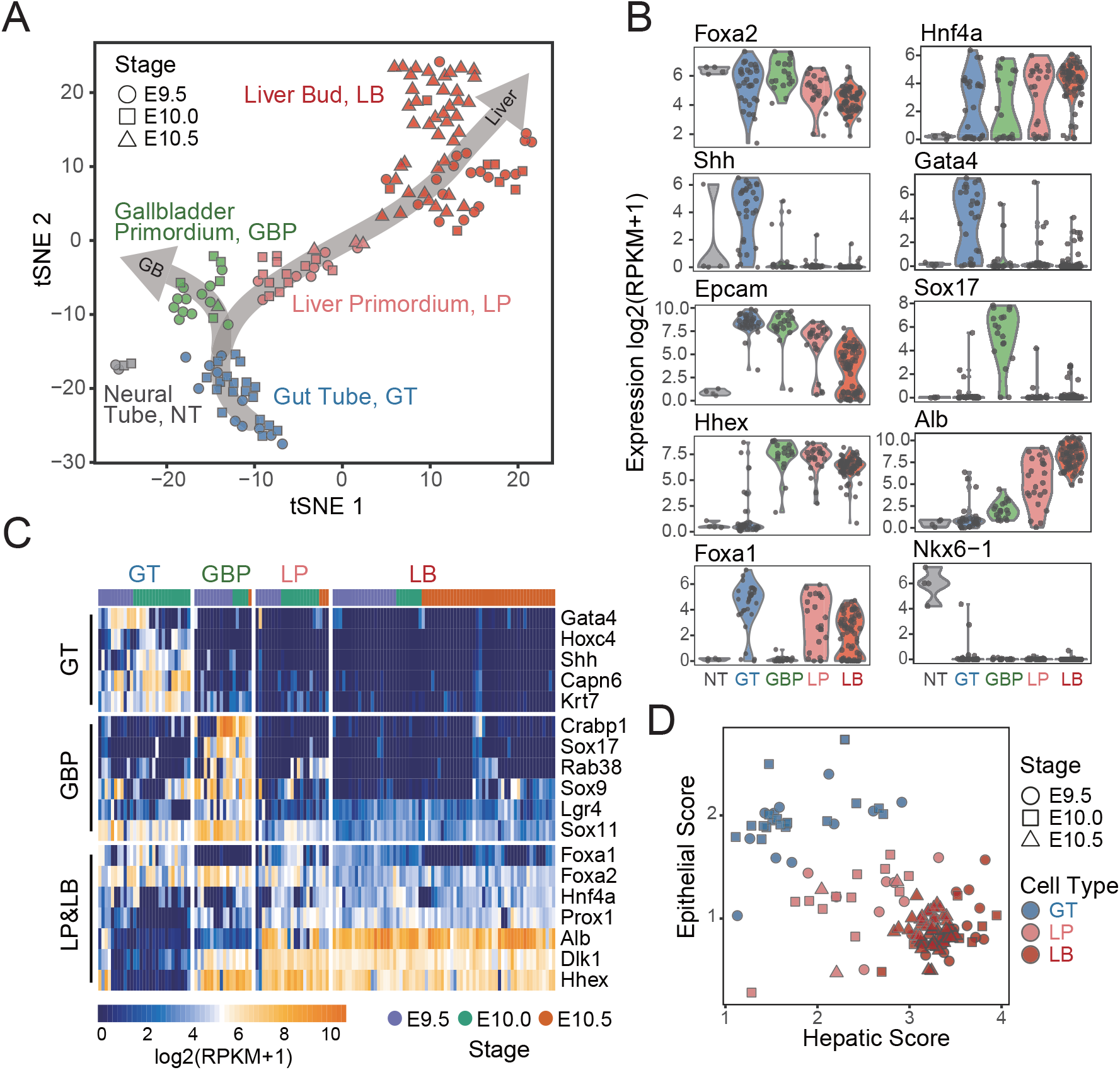
Liver primordium cells at E9.5 identified as nascent hepatoblasts A, Single-cells from E9.5 and E10.5 were clustered by t-SNE into 4 groups, gut tube (GT), liver primordium (LP), liver bud (LB) and gallbladder primordium (GBP). B, Violin plots of the gene expression levels of specific markers, *Foxa2, Hnf4a, Shh, Gata4, Epcam, Sox17, Hhex, Alb, Foxa1* and *Nkx6-1* in each cell group. C, Differentially expressed genes of GT, GBP, LP, LB groups identified during E9.5-E10.5. D, Liver primordium exhibited an intermediate state between the undifferentiated gut tube and differentiated liver bud by analyzing hepatic score and epithelial score.

### Gallbladder primordium at E9.5 expresses hepatic and non-hepatic genes

In addition to the hepatic lineage, the *Foxa2+* neural tube (NT) cells expressing *Nkx6-1* were spatially close to the gut tube and were detected at E9.5 and E10.0 (Figure 3A, 3B). Three pancreas-like cells that expressed *Pdx1* but not *Sox17* were identified and were excluded from further analysis, due to the limited cell number. Moreover, we identified a group of gallbladder primordium (GBP) cells mainly at E9.5 that expressed *Sox17* and *Lgr4*, but not *Pdx1* (Figure 3A, 3B). Taken together with the gut tube and liver primordium cells, a two-direction developmental trajectory of the gut tube was identified (Figure 3A). Interestingly, the gallbladder primordium cells express many hepatic genes including *Alb* and *Dlk1* but not as high as liver primordium (Figure 3C). Moreover, *Foxa1* was expressed in the gut tube and liver primordium but not in the gallbladder primordium, while the expression of *Foxa2* and *Foxa3* was positive in both liver primordium and gallbladder primordium, indicating that *Foxa1* is selectively suppressed during gallbladder development. By differentially expressed genes analysis, 411 genes were found to be up-regulated in the gallbladder primordium compared with the gut tube from E9.5-E10.5 (Figure S10B, Table S7). This 411 genes group included the *Sox* family genes *Sox9*, *Sox11* and *Sox17. Sox9* has been reported to be related to cholangiocyte differentiation ^38^. In addition, *Crabp1, Rab38, Flt1, Slco5a1, Ptpn5, Vstm2b, Ntrk2, Ets1 and Ypel4* were identified as potential new markers in the gallbladder primordium, while barely expressed in the gut tube and hepatic cells (Figure 3C, Figure S11). By contrast with gut tube and liver primordium, *Junb, Hpx, Mt2, Lrrc3, Dkk3, Apob, Acss1 and Dhrs3* were negative in the gallbladder (Figure S11).

### Major gene expression dynamics during the epithelial-hepatic transition (EHT)

To characterize the epithelial-hepatic transition process, we analyzed the differentially expressed genes between the gut tube (GT), liver primordium (LP), liver bud (LB) and hepatic cells from E11.5 (E11.5 Hep) by RaceID ^39^. The heatmap of differentially expressed genes demonstrated a programmed change of gene expression from the gut tube to the hepatoblast (Figure 4A). These genes could be clustered into 6 gene groups: L1, L2, L3, G1, G2 and G3 (representing the following gene groups: Liver 1, Liver 2, Liver 3, Gut tube 1, Gut tube 2, Gut tube 3, respectively), based on temporal order and biological functions (Figure 4A, Table S8). These gene groups were dynamically regulated by the developmental axis GT-LP-LB-Liver (Figure 4B). During the liver development, L1 genes were firstly switched on in liver primordium, followed by L2 in the liver bud and L3 in E11.5 liver. Meanwhile, G1, G2 and G3 genes were down-regulated or switched off in the liver primordium, liver bud and liver in E11.5, respectively. The L1 genes were enriched in blood coagulation and hemostasis (*F10*, *F12*, *Fga*, *Serpina6* and *Serpind1*) and lipid metabolic processes (*Apoc2, C3*) (Figure 4C). L2 genes were related to oxidation reduction (such as *Cyp2d10*, *Sord* and *Hsd17b2*) and triglyceride catabolism (*Aadac* and *Apoh*). The L3 genes are involved in the glucose metabolic process and fatty acid oxidation (*Pdk4*, *Cpt1a*, *Slc27a5*). Meanwhile, in the G1, G2 and G3 gene groups that decreased during the liver development, we have identified many epithelial feature genes (including collagen, claudin, and laminin). The *Grhl2* gene was firstly down-regulated in the liver primordium, the *Kit*, *Krt19* and *Col2a1* were then down-regulated in the liver bud, and finally, the *Epcam* and *Cldn6* almost disappeared in the liver of E11.5. In summary, extensive change in gene expression patterns was a dominant feature during the EHT.

**Figure 4.**
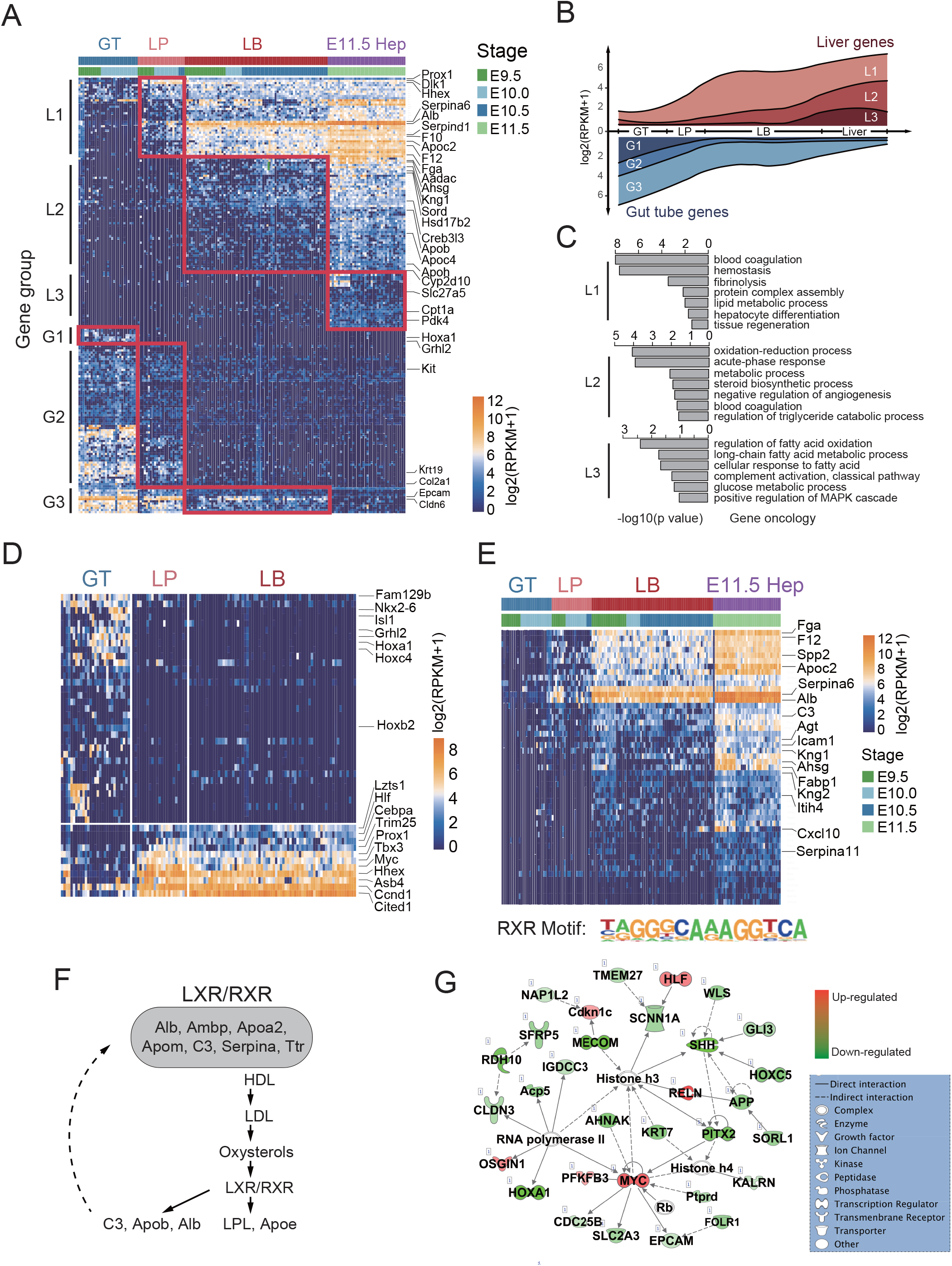
The gene expression dynamics during the epithelial-hepatic transition (EHT). A, Differentially expressed genes demonstrated a dynamic change of the gene expression from the gut tube to the hepatoblast. Six groups of differentially expressed genes were identified to be switched on or off, including three live gene groups: L1, L2 and L3, and three gut tube gene groups: G1, G2, and G3. B, The gene expression levels of 6 gene groups (L1, L2, L3, G1, G2, G3) were identified followed by the development axis GT-LP-LB-Liver. C, Gene ontology of the gene groups L1, L2, L3 were identified. D, Transcription factors that differentially expressed between the gut tube and liver primordium were identified including some new TFs such as *Lzts1, Hlf, Trim25, Myc, Asb4, Ccnd1* and *Cited1.* E, RXR motif was identified in the genes highly expressed in liver primordium. The 49 targets of RXR motif were shown. F, LXR/RXR signaling pathway was significantly up-regulated in the liver primordium compared with the gut tube and formed a positive-feedback loop. G, The networks ‘Cellular Development’, ‘Cell Growth and Proliferation’, ‘Connective Tissue Development and Function’, ‘Embryonic Development’ and ‘Organismal Development’ identified in liver primordium.

### Significant transcription factors and RXR complex signaling regulate liver specification

To identify the genes that trigger the hepatic fate, we focused on the 548 genes encoding transcription factors, enzymes, cytokines, transporters and kinases that were differentially expressed between the gut tube and liver primordium (Table S9, Figure S10A). In total, 49 TF genes were found to be significantly activated in the liver primordium, including *Cebpa, Prox1, Tbx3* and *Hhex* as expected. Moreover, we have identified several new up-regulated TF genes including *Lzts1, Hlf, Trim25, Myc, Asb4, Ccnd1* and *Cited1* (Figure 4D). Mutation of the mouse *Lzts1* gene has been reported to result in hepatocellular carcinoma ^40^. We also identified TF genes down-regulated in the liver primordium including *Hoxa1, Hoxb2, Hoxc4, Grhl2, Isl1, Nkx2-6* and *Fam129b*. Target genes (*Cdh1, Cldn4, Sema3c, Sema3b, Rfx2, Nrp2*) of *Grhl2* were also found to be down-regulated (Figure S10C).

The gene networks and signaling pathways of the 548 differentially expressed genes were enriched in ‘Cellular Development’, ‘Cell Growth and Proliferation’, ‘Connective Tissue Development and Function’, ‘Embryonic Development’ and ‘Organismal Development’ (Figure 4G). More importantly, we found the liver X receptors/retinoid X receptors (LXR/RXR) pathway was significantly up-regulated in the liver primordium compared with the gut tube, including *Alb, Ambp, ApoA1, ApoA2, ApoE, ApoF, ApoM, C3, Ttr, SerpinA1, SerpinF1, SerpinF2* and others (Figure S12). To validate the role of the LXR/RXR pathway, we analyzed the promoters of the differentially expressed genes between the gut tube and hepatoblasts by motif analysis. The promoters of 49 genes (including *Alb, C3, Apo* and *Serpin* family members) highly expressed in the hepatoblasts had putative RXRA elements (Table S10). The expression of these target genes increased in the liver primordium and peaked within the liver bud (Figure 4E). Combined with IPA analysis, the *Alb* and *Serpin* family served as both the ligands and the targets in the LXR/RXR pathway, which implies a positive-feedback loop during liver specification (Figure 4D). Beside RXRA, genes up-regulated in hepatoblasts were found to be targets of HNF4A and PPARG, while the targets of SOX2 and TEAD1 was found in the down-regulated genes (Figure S13). In conclusion, activation of the RXR complex signaling pathway and several new TF genes including *Lzts1* are concomitant with liver specification.

### Transient transcription factor gene expression during hepatoblast maturation into hepatocytes

To study the dynamics of hepatoblast maturation into hepatocytes, we retrieved the Alb+ hepatoblast/hepatocytes in E11.5-E15.5 after excluding the erythroblast cells expressing *Gata1* and monocytes expressing *Ptprc* (CD45) (Figure S14A, S14B). The liver primordium and liver bud cells from E9.5-E10.5 and hepatoblast/hepatocytes cells from E11.5-E15.5 were reordered by Monocle ^41^, and a trajectory of hepatic development was determined (Figure 5A). Notably, there were no branches on the trajectory and the predicted pseudotime of the trajectory agreed with the gestation day. During this timeline, we examined specific genes and gene sets defining the ‘hepatic score’ (liver metabolic function genes including *Alb*), ‘stemness score’ (stem markers including *Nanog*), and ‘proliferation score’ (cell cycle genes including *Mki67*). The metabolic function of hepatoblasts/hepatocytes increased while the cell pluripotency and the proliferation rate decreased during liver maturation (Figure 5B, 5C). These results were also validated by the 720 cells generated by the MIRALCS method from the E11.5, E12.5 and E13.5 stages (Figure S14C).

**Figure 5.**
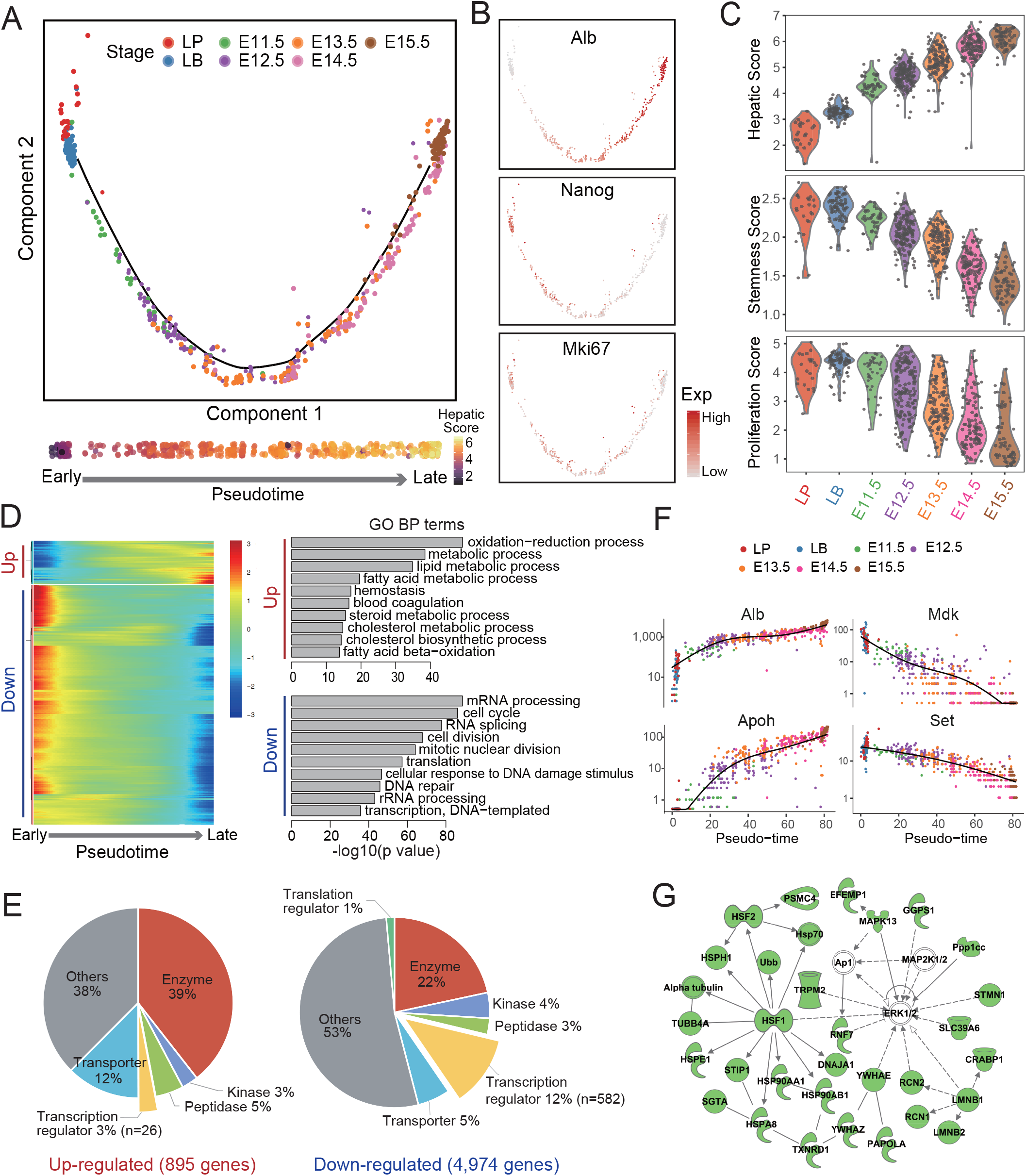
Dynamic gene expression during hepatoblast maturation into hepatocytes. A, A trajectory of hepatic development was determined during E9.5-E15.5 by Monocle analysis. No branches on the trajectory were found and the predicted pseudotime of the trajectory agreed with the gestation day. B, The expressions of *Alb, Nanog, Mki67* were shown in the hepatic trajectory. C, The metabolic function of hepatoblasts/hepatocytes increased while the cell pluripotency and the proliferation rate decreased during liver development based on ‘hepatic score’, ‘stemness score’ and ‘proliferation score’ during E9.5-E15.5. D, Genes dynamically regulated during hepatoblast maturation and the respective function of up/down-regulated genes were identified. E, The composition of up-regulated or down-regulated genes during E9.5-E15.5 were shown. F, Genes metabolism function of the liver (such as *Alb* and *Apoh*) were up-regulated, while the down-regulated genes were enriched in cell cycles, RNA splicing, cell division and translation (such as *Mdk* and *Set*). G, Genes network that were down-regulated during the hepatoblasts maturation. Genes labeled by green were down-regulated, while by white were not detected.

With the Monocle analysis, we found 5,869 genes dynamically regulated during hepatoblast maturation (q value< 0.01) (Table S11). Interestingly, 85% (4,974 genes) of these genes were down-regulated while only 15% (895 genes) were up-regulated (Figure 5D). The down-regulated genes consisted of 12% TFs (582 genes), while the up-regulated genes consisted of only 3% TFs (26 genes) (Figure 5E). These results indicate that a large number of TFs play transient roles in liver specification and are decreased afterward. The up-regulated genes were mostly related to the metabolic function of the liver (such as *Alb* and *Apoh*), while the down-regulated genes were enriched in the cell cycle, RNA splicing, cell division and translation (such as *Mdk* and *Set*) (Figure 5D, 5F). Moreover, we found *Ubb* that regulate the protein ubiquitination process was down-regulated during hepatoblasts maturation, perhaps to protect the metabolic enzymes and other proteins produced by the hepatocytes (Figure 5G). The heat shock response (HSR) pathway (genes including *Hsf1*, *Hsf2,* Hs*p70, Hsph1, Hspe1, Hspa8, Hsp90ab1* and *Hsp90aa1*) that control the protein folding process was also found to be down-regulated (Figure 5G).

## Discussion

Single-Cell RNA-Seq is a powerful tool in developmental biology. Although the throughput of the method has been increased dramatically in the recent few years, careful quality control of the data is essential. Due to the rare cell number of organ progenitors, it is not feasible to directly perform high-throughput scRNA-Seq to study the early liver development from bulk tissue, due to the requirement for a large number of loading cells and the low cell capture rate of current methods. To circumvent these limitations in the study of specific organ development, we therefore constructed a Foxa2^eGFP^ mouse model and used FACS to isolate hepatic-related single-cells to obtain high-quality data. This allowed high-coverage transcriptome profiling (10,000 detected genes/cell) to identify low-expression genes, such as transcription factors. Besides, the Foxa2^eGFP^ mouse model also had the advantages of co-expression of *Foxa2* and *eGFP*. The high correlation of *Foxa2* and *eGFP* (>0.95) gave high confidence in the transcriptome profiling. Another advantage of the Foxa2^eGFP^ mouse model is that *Foxa2* is an endoderm marker, which can track liver development from the endoderm stage to hepatoblasts. By using this model, we performed precise embryonic dissection and excluded irrelevant cells by FACS.

By combining scRNA-seq, the Foxa2^eGFP^ mouse model, precise microdissection and annotation, we acquired high quality and full-length transcriptome data of primitive streak, foregut endoderm, liver primordium, liver bud and early fetal liver at the single-cell level. We found that the transcriptional dynamics of embryonic liver development passed through a “three-step” process. Firstly, the E8.5 foregut endoderm develops from the primitive streak; secondly, hepatoblasts are specified from the foregut tube at E9.5; and finally, the hepatoblasts mature into hepatocytes (Figure 6). These processes nicely agree with previous morphological studies ^15^ and reveal that the cells at each stage have markedly distinct transcriptional programs.

**Figure 6.**
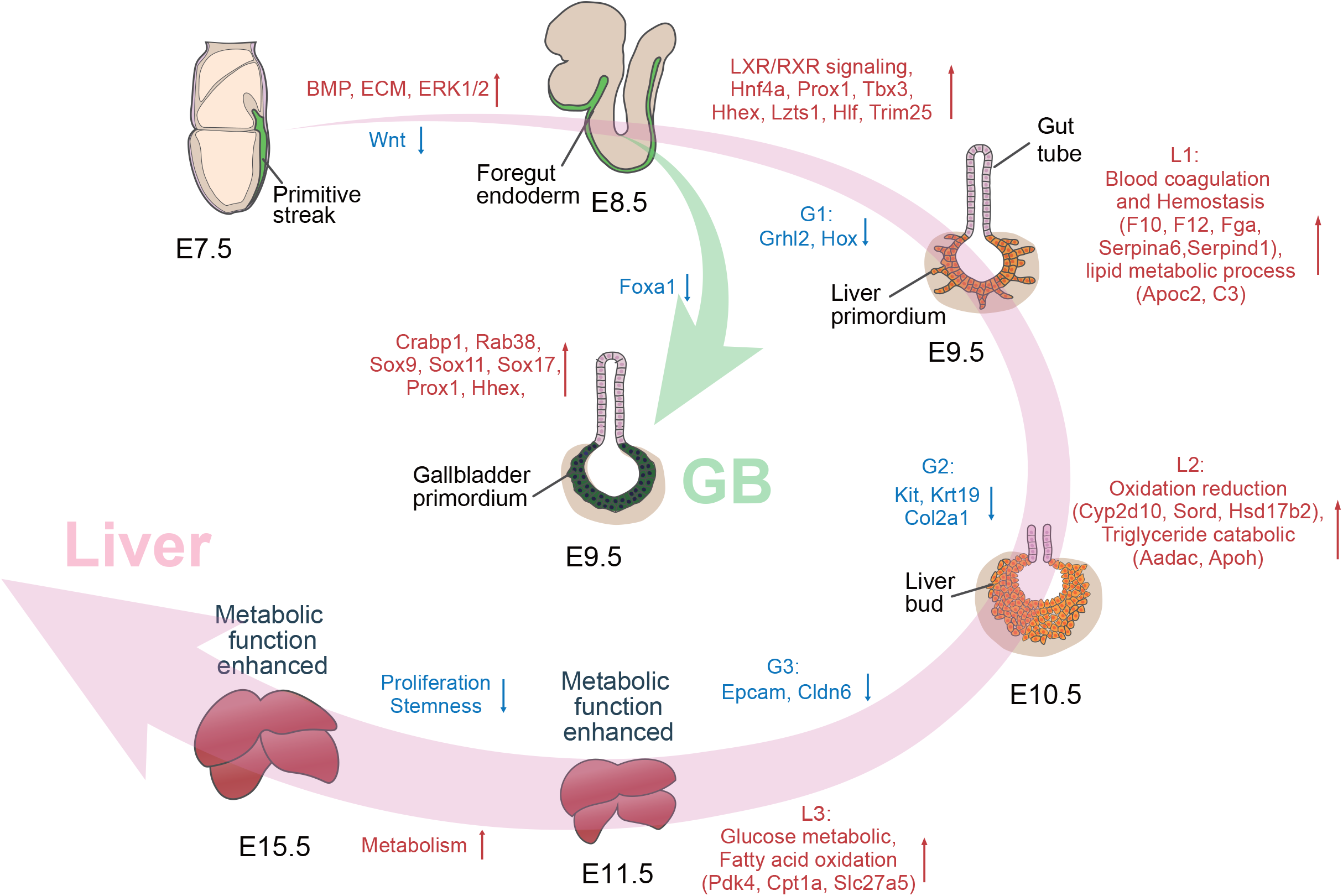
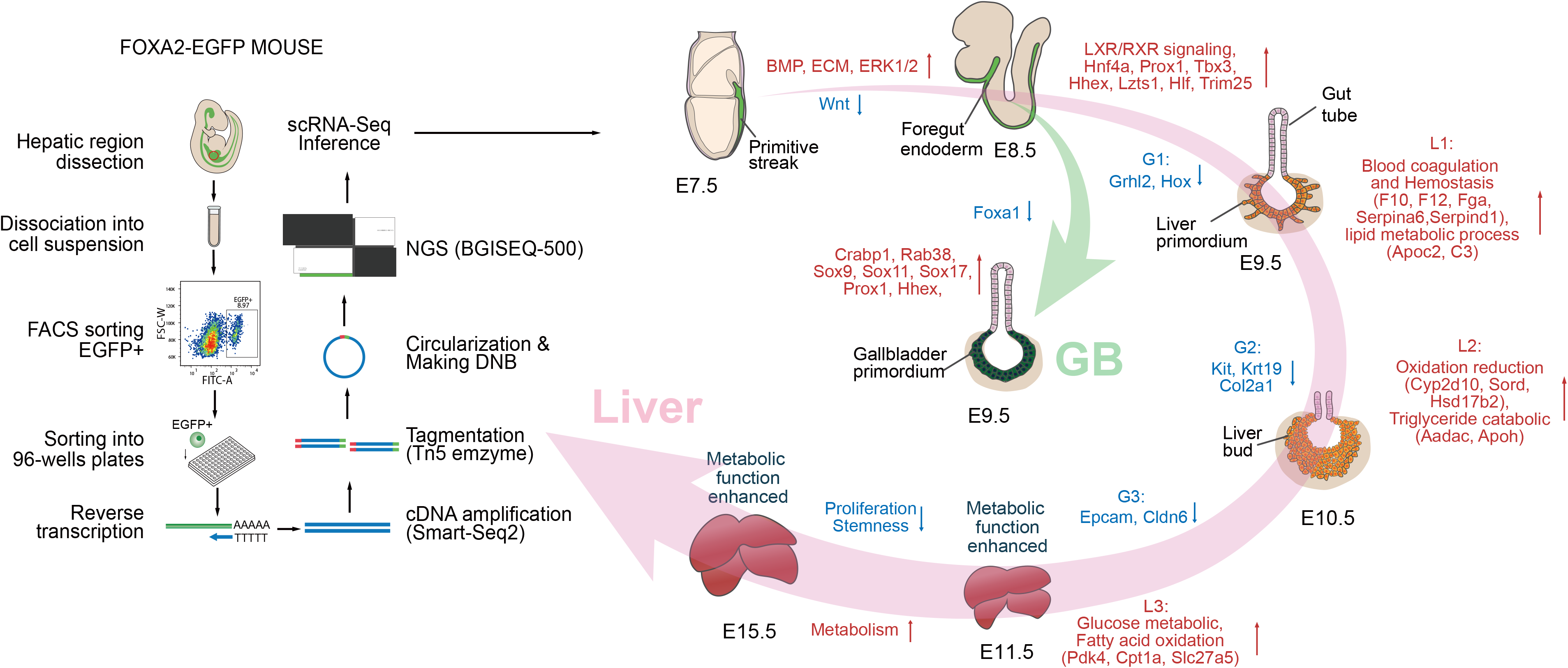
Dynamic transcriptome of liver and gall bladder development from the embryonic endoderm during E7.5-E15.5 by scRNA-Seq

During the endoderm morphogenesis, we found a subgroup of the foregut endoderm cells (Foregut-1) at E8.5 that were the descendants of visceral endoderm in E7.5 ^32^. The second group of foregut endoderm epithelium (Foregut-2) at E8.5 differentiated from the E7.5 primitive streak cells with mesenchymal morphology throughout a MET progress. Our data are consistent with BMP signaling being activated by inhibiting *Fst* and *Chrd*, while Wnt/β-catenin signaling was repressed by *Sox11, Rarb* and *Tgfb2* in the foregut. We also found expression of ECM genes, including collagens fibrinogen, integrin, and tight junction protein genes, to be up-regulated in the foregut, perhaps through activation of ERK1/2.

During the liver specification progress, we identified the liver primordium as nascent hepatoblasts that derived from the gut tube at E9.5. Liver primordium exhibited an intermediate state between the undifferentiated gut tube and differentiated liver bud, and had increased hepatic features and reduced epithelial features through the EHT process. We found that during the EHT process, the dynamically expressed genes could be clustered into 6 groups (L1, L2, L3, G1, G2, G3), which were fated to be switched on or off following the developmental axis GT-LP-LB-Liver. Different functions of each gene group may provide clues about the sequence of significant events during liver development. Moreover, we found the EHT process to be associated with the activation of the LXR/RXR signaling pathway and TFs such as *Lzts1*, *Hlf*, *Trim25, Myc, Asb4, Ccnd1* and *Cited1*. In contrast, *Hoxa1, Hoxb2, Hoxc4, Grhl2, Isl1, Nkx2-6* and *Fam129b* were inhibited in the liver primordium. A positive-feedback loop of the LXR/RXR signaling pathway could explain why the expression levels of *Alb* and *Serpin* family genes were increased over 1,000-fold in a short time compared with the gut tube. RAR deficient mice are not lethal, but display abnormal liver development ^42^. Among 49 TFs that were differentially expressed between the liver primordium and gut tube, only 11 TFs were up-regulated while 38 TFs were down-regulated, which implies that suppression of specific TFs in the gut tube is critical for liver specification.

Together with the liver primordium, we identified a group of gallbladder primordium cells. Interestingly, we found the gallbladder primordium expressed many hepatic marker genes including *Hnf4a*, *Prox1*, *Foxa2/3, Dlk1* and *Alb*, but did not express *Foxa1*. Furthermore, we also identified some potential new markers and effectors for the gallbladder, such as *Crabp1, Rab38, Flt1, Slco5a1, Ptpn5, Vstm2b, Ntrk2, Ets1 and Ypel4.*

During the maturation of hepatoblasts into hepatocytes between E11.5-E15.5, the transcriptome was relatively stable and changed gradually during this process, implying that the majority of liver specification occurs during E9.5-E10.5. In addition, the hepatic features increased, while stemness features and cell proliferation decreased. Interestingly, 85% of the dynamically expressed genes were down-regulated including many TFs. This suggests that many genes including TFs are necessary for organ specification and are turned off after the specification is completed.

Our study is the first to reveal the transcriptome profile of the embryonic liver from the emergence of nascent hepatoblasts to the stable formation of liver cellular structures by tracing the Foxa2 lineage with single-cell resolution. We have identified numerous key networks critical for liver development, features that distinguish the gallbladder, and potential clues leading to therapeutically useful tissues for transplantation. Moreover, this Foxa2^eGFP^ mouse model could be used to study the development of other endodermal organs, such as the lung, pancreas and stomach, which would provide insight into the mechanisms of endodermal organ development.

## Acknowledgements

We thank Dr. Zakir Hossain for mouse embryo electroporation; Dr. Tang Fuchou for comments on the manuscript; Dr. Nicolas Plachta for the 3D-imaging of mouse embryo culture; Dr. Xu Chengran for demonstration of mouse manipulation; Dr. Eseng Lai, Robert H. Costa, and James E. Darnell Jr. for intellectual contributions to the Foxa2 study. This is a prerint of an article published in COMMUNICATIONS BIOLOGY. The final authenticated version is available online at https://doi.org/10.1038/s42003-020-01364-8.

## Abbreviations used in this paper

MET: mesenchymal-epithelial transition
EHT: epithelial-hepatic transition
E7.5: Embryonic day 7.5
MIRALCS: microwell full-length mRNA amplification and library construction system
RPKM: Reads Per Kilobase per Million mapped reads
t-SNE: T-distributed Stochastic Neighbor Embedding
PS: primitive streak
FG: foregut endoderm
VE: visceral endoderm
NT: neural tube
NC: notochord
IPA: Ingenuity pathway analysis
ECM: extracellular matrix
M score: mesenchymal score
E score: epithelial score
GT: gut tube
LP: liver primordium
LB: liver bud
GBP: gallbladder primordium
RaceID: rare cell type identification
L1: gene group Liver 1
L2: gene group Liver 2
L3: gene group Liver 3
G1: gene group Gut tube 1
G2: gene group Gut tube 2
G3: gene group Gut tube 3
LXR/RXR: liver X receptors/retinoid X receptors
TF: transcription factor
FACS: Fluorescence-activated cell sorting

