## supplementary method for "Characterizing the Emergence of Liver and Gallbladder from the Embryonic Endoderm through Single-Cell RNA-Seq"

### Supplementary Methods

**Targeting vector construction:**

BAC clone RP23-469P2 containing mouse *Foxa2* locus was obtained from BACPAC as the template. Exon3 of *Foxa2* (Chr2: 147869253-147870662) was amplified by HiFi-PCR and inserted into pEYFPC1 by XhoI /EcoRI, then *eGFP* coding region was amplified from pLEGFPC1 vector to fuse in-frame with Exon3 of *Foxa2* by linker CTA- GGA-ATT-CTA (Leu-Gly-Ile-Leu). Next 3'UTR of *Foxa2* (Chr2: 147867876-147869250) was amplified from BAC clone and inserted after *Foxa2-eGFP* fusion by KpnI/BamHI on pEYFPC1 backbone. The whole region of Exon3 of *Foxa2*-eGFP-3'UTR of *Foxa2* was retrieved and inserted into pEASY-Flox vector by XbaI/SalI which was flanked by the latter two LoxP sites. The first LoxP site and Neo cassette were removed and restriction sites were introduced to facilitate downstream construction. 4.5kb *Foxa2* genomic region (Chr2: 147870690-147875176) before Exon3 was amplified from BAC and inserted into pEASY-flox by ClaI/BamHI. 2.8kb genomic region after 3'UTR of *Foxa2* (Chr2: 147865027-147867871) was amplified from BAC and inserted by HindIII/XhoI. The PGK (phosphoglycerate kinase) promoter-driven Neomycin expression cassette flanked by FRT sequence was inserted at the reverse direction by HindIII. The vector was partially verified by sequencing of ligated regions. Genotyping was done by PCR using DNA extracted from tail tips from 3-week-old mice. The sequence of primer pairs used are provided:

Foxa2-eGFP-1: 5'-CTTTGGGGCCCAGAGGACTTGGTG-3';

Foxa2-eGFP-2: 5’-GTATGTGTTCATGCCATTCATCCCCAGG-3’.

Foxa2-linker-eGFP:

5’-TATGAACTCATCCCTAGGAATTCTAGTGAGCAAGGGCGAG-3’.

**Mouse breeding and dissection/timed mating and embryos collection:**

The Foxa2$^{eGFP}$ mice have been backcrossed to C57BL/6 background for more than 20 generations and do not show any abnormal phenotypes. All procedures were performed under the strict instruction of Comparative Medicine of National University of Singapore. All the mice used as the sample for Single-cell RNA-Seq were homozygous for Foxa2$^{eGFP}$. Timed-pregnant Foxa2$^{eGFP}$ females were used for the sequencing.

**FACS:**

Flow cytometry (FACSAria III cell sorter) was used to analyze and sort target cells. Cells collected from the wild-type mouse liver were used as control and set the eGFP negative gate on the cell sorter. Multiplets were excluded in our sorting. Single cells were sorted onto a glass slide and checked under a microscope before sorted into each well of 96-well plates. Each well of the plates was pre-loaded with 4ul of cell lysis buffer. 100-200 cells were sorted into 1-3 wells as bulk control, and 1 negative well with no cells was designed to evaluate the contamination of mRNA amplification.

**Single cell mRNA amplification**

Message RNA from single cells were amplified using Smart-seq2 technology as previously described (Picelli et al., 2014) with modifications. Briefly, cells were lysed at 65°C for 3 min and then subject to reverse transcription using an oligo(dT) primer and a locked nucleic acid (LNA)-containing template-switching oligonucleotide (Exiqon). Full-length cDNAs were amplified by 20 cycles of PCR using KAPA HiFi DNA polymerase (KAPA biosystems) and IS primer. The products from randomly selected 3 wells and the negative well were run on the agarose gel. Only the plates that were successfully amplified without contamination were processed to library construction. All the products were purified with 0.8× Ampure XP beads (Beckman) and quantified with AccuBlue High Sensitivity dsDNA Quantitation Kit (Biotium). The amplified cDNA was assessed by agarose gel and qPCR before library generation to ensure the sequencing quality (Figure S2).

Oligo-dT: 5'-AAGCAGTGGTATCAACGCAGAGTACT (30) VN-3' (V=G/A/C, N= G/A/C/T)

TSO: 5'-AAGCAGTGGTATCAACGCAGAGTACrGrG+G-3' (rG=RNA Guanine, +G=LNA modified Guanine)

IS primer: 5'-AAGCAGTGGTATCAACGCAGAGTAC-3'

**Real-time PCR**

Real-time PCR was used to test the success of Single cell cDNA amplification. The cDNA products were subjected to RT–PCR with KAPA SYBR® FAST Universal 2X qPCR Master Mix40 (KK4600) using a 7300 or 7500 Real-Time PCR machine (Applied Biosystems). All mRNA expression values were normalized against the internal housekeeping gene GAPDH.

**Embryo fixation**

The whole embryo was fixed in 4% paraformaldehyde for 1-2 hours at room temperature. After fixation, tissue was rinsed with PBS until fixative is completely removed. Tissue was dehydrated by using different series of Ethanol, Citrisolve and paraffin: 50% Ethanol for 10 min; 70% Ethanol for 10 min; 80% Ethanol for 10 min; 95% Ethanol for 10 min; 100% Ethanol for 10 min; 100% Ethanol for 10 min; 100% Ethanol for 10 min; 2:1 Ethanol : Citrisolve for 10-15 min; 1:1 Ethanol : Citrisolve for 10-15 min; 1:2 Ethanol : Citrisolve for 10-15 min; 100% Citrisolve for 10-15 min; 100% Citrisolve for 10-15 min; 100% Citrisolve for 10-15 min; 2:1 Citrisolve : Paraffin for 30 min; 1:1 Citrisolve : Paraffin for 30 min; 1:2 Citrisolve : Paraffin for 30 min; 100% Paraffin for 1-2 hr; 100% Paraffin for 1-2 hr or overnight.

**Immunohistochemistry**

Tissues were either fixed overnight and embedded in paraffin. Sections were cut 4-5 mm thick. Paraffin sections were deparaffinized, dehydrated, and we performed antigen retrieval by steaming slides in sodium citrate buffer for 30 min. Sections were blocked in the blocking serum buffer (5% serum in 13 PBS + 0.5% Triton X-100) for 30 min. Primary antibodies were diluted in blocking buffer and incubated on tissue sections overnight at 4 . Slides were washed and incubated in secondary antibody in blocking buffer for 2 hr at room temperature. Slides were washed and mounted using Fluormount-G, and observed under microscope. Antibody information: Foxa2 (Cell Signaling, #3143); DLK1 (Abcam, ab21682).

**Library preparation and sequencing with BGISEQ-500**

All the cDNAs were converted into libraries and sequenced on the BGISEQ-500 sequencer. BGISEQ-500 is an industry leading high throughput sequencing solution, powered by combinatorial Probe-Anchor Synthesis (cPAS) and improved DNA Nanoballs (DNB) technology (Drmanac et al., 2010; Huang et al., 2017).

2 ng of the cDNA was fragmented using the Tn5 enzyme-adaptor compound. 15 cycles of PCR were then carried out with barcoded primers compatible with the BGISEQ-500. The 300-500 bp DNA fragments will be selected and purified. The fragments will be heat-denatured and one of the single strands will be circularized with DNA ligase to obtain a single-strand circular DNA library. The remaining single strand was digested with the exonuclease. The sequencing process was conducted according to the BGISEQ-500 protocol as described (Huang et al., 2017).

**Public dataset access**

Mouse (Mus musculus) reference genome (mm10) (Waterston et al., 2002) was downloaded from http://genome.ucsc.edu/. The transcriptome reference annotation GTF file (Ensembl GRCm38)(Aken et al., 2016) was downloaded from http://www.ensembl.org/. The sequence of *eGFP* was inserted into the *Foxa2* in mm10 reference file based on the structure of the transgenic vector. The annotation of eGFP was also added to the GTF file.

**RNA-seq data processing**

The raw sequencing data were accessed by filtering reads with adapters/poly-A, N rate > 0.05 or low-quality base rate > 0.5 using SOAPnuke (v1.5.6) (Chen et al., 2018). Clean reads were mapped to the reference by TopHat (v2.1.0) (Trapnell et al., 2012) using Bowtie (v0.12.9.0) (Langmead et al., 2009) with parameters "–-bowtie1 –p 4 -g 1 --solexa1.3-quals --fusion-search --fusion-min-dist 100000". The Bowtie index of mm10 was built on autosomes and chrX.

**Quantification of gene expression levels**

Gene expression levels were quantified as reads per kilobase of gene per million mapped reads (RPKM). Read counts were calculated by Rsubread (Liao et al., 2013) (v1.16.1), and RPKM values were calculated using edgeR (Robinson et al., 2010) (v2.6.12) with the edited GTF file (GRCm38). Cells with mapping reads < 1 million or mapping rate < 40% were discarded. Cells in 96-wells plates with detected gene number (RPKM > 1) ≥ 6,000 were defined as qualified cells, and the threshold of gene number was set as 4,000 for cells generated by MIRALCS. Cells with $RPKM_{Foxa2} \geq 1$ and $RPKM_{eGFP} \geq 1$ were defined as Foxa2+ cells.

**Sequencing data assessment**

We devised a pseudobulk by pooling all single-cells from the same stage and compared their gene expression levels with that of bulk sample (Figure S5a). The correlations were extremely high (Pearson r >0.9), indicating that our method was accurate and sensitive. Additionally, the amplification bias was assessed by comparing the expression level of the *Foxa2* and *eGFP* since they were expected to be transcribed at the same time. We found a high correlation between *eGFP* and *Foxa2* (r>0.9) in the Foxa2+ cells, indicating a low bias in amplification (Figure S4b, S4c). Moreover, we assessed the systematic error within a batch of repeat experiment using E12.5, E13.5 and E14.5 livers. The results showed high correlations between the two batches of data from the same developmental stage (Figure S5c). To assess the sequencing batch effect, we pooled the libraries from E9.5 and E10.5 and sequenced them on two separate sequencing chips (Figure S5d). Besides, libraries from E7.5 were also sequenced on two chips (Figure S5e). All these results showed a high correlation and low sequencing batch effect.

**Cell clustering and marker genes identification**

922 cells from E7.5 to E15.5 were clustered by Seurat (v2.1.0) (Satija et al., 2015). The read count matrix was firstly column-normalized and log-transformed. The high

variable genes identified by “Find Variable Genes” function were used for PCA analysis. The appropriate PCs were selected for clustering with the specific resolution parameters. Then t-distributed stochastic neighbor embedding (t-SNE) was used with the same number of PCs to visualize the clustering results. To detect cluster-specific expressed genes (marker genes), clusters were compared pairwise using “Find All Markers” function. Genes with at least 0.5-fold difference (log-scale) and a detectable expression in more than 50% of cells in either population were identified as candidate marker genes. For cells from E7.5 and E8.5, Foxa2+ cells were involved in the analysis, and the top 12 PCs were selected with resolution parameter equal to 1. For cells from E9.5 and E10.5, Foxa2+ cells were used and the top 5 PCs were selected with resolution parameter equal to 0.8. For cells from E11.5 to E15.5, all cells were used and the top 8 PCs were selected with resolution parameter equal to 0.7.

**Differential expression analysis**

We performed differential expression analysis using SCDE (Kharchenko et al., 2014) (v1.99.1), which adopted a Bayesian approach fitting individual error models. We selected the genes whose value of the most likely fold expression difference more than 1 (ce >1) as differential expression genes for subsequent analysis.

For cells from E9.5 and E10.5, the cells were separated into different groups based on the cell types identified by Seurat. We performed the Kruskal-Wallis test (KW test) and retained the genes with adjusted q value < 1e-4. RaceID (Grun et al., 2015) was applied to analyze the dataset with default model parameters. The cluster-specific marker genes were selected using the “clustdiffgenes” of RaceID with p-value equal to 0.05.

**Pseudo-temporal analysis**

We construct pseudo-temporal analysis using Monocle2 (Qiu et al., 2017) (v2.6.4) in cells from E9.5 to E11.5. Only cells identified as "Liver primordium", "Liver bud" or "Hepatoblast" from E9.5 to E15.5 were included. The genes expressed in at least 10 cells with RPKM > 1 were retained. The differentially expressed genes across different cell stages with q values < 0.001 were used for pseudo-temporal analysis. After constructing the cell trajectories, differentially expressed genes along the pseudotime were detected using “differential Gene Test” function, and genes with q values < 0.01 were retained.

**Ingenuity Pathway Analysis (IPA)**

Ingenuity Pathway Analysis was performed to interrogate the biological functions during the development. The differentially expressed genes between different cell types were uploaded into the IPA software for the core analysis, and identified the canonical pathways, diseases and functions, upstream regulators and gene networks.

**ITranscriptome analysis**

lTranscriptome (Peng et al., 2016) was used to identify the potential spatial locations of cells defined as primitive streak (PS). The zipcode genes were downloaded from http://www.picb.ac.cn/hanlab/media/lmdseq/static/zipcodegenes.txt. The expression matrix of zipcode genes of PS cells was uploaded to perform Zipcode Mapping analysis.

**Motif analysis**

HOMER (Hypergeometric Optimization of Motif EnRichment, v4.8.3, http://homer.ucsd.edu/homer/motif/) (Heinz et al., 2010) was applied for searching promoters of genes and motifs enriched in target gene promoters as well. The "findMotifs.pl" script was downloaded from the website to analyze the regulators involved in up/down regulation of differential expression genes. Cluster-specific expressed genes were used to search for motifs with length 8, 10 or 12 from -300 to +100 relative to the transcription start site (TSS).

**Gene ontology analysis and feature score quantification**

We performed gene ontology analysis using the DAVID Bioinformatics Resource v.6.8 (https://david.ncifcrf.gov/). The feature scores were calculated by the average expression (log2-transformed) of each feature gene set (Table S2).
