## supplementary figure for "Characterizing the Emergence of Liver and Gallbladder from the Embryonic Endoderm through Single-Cell RNA-Seq"

### Figure S1.

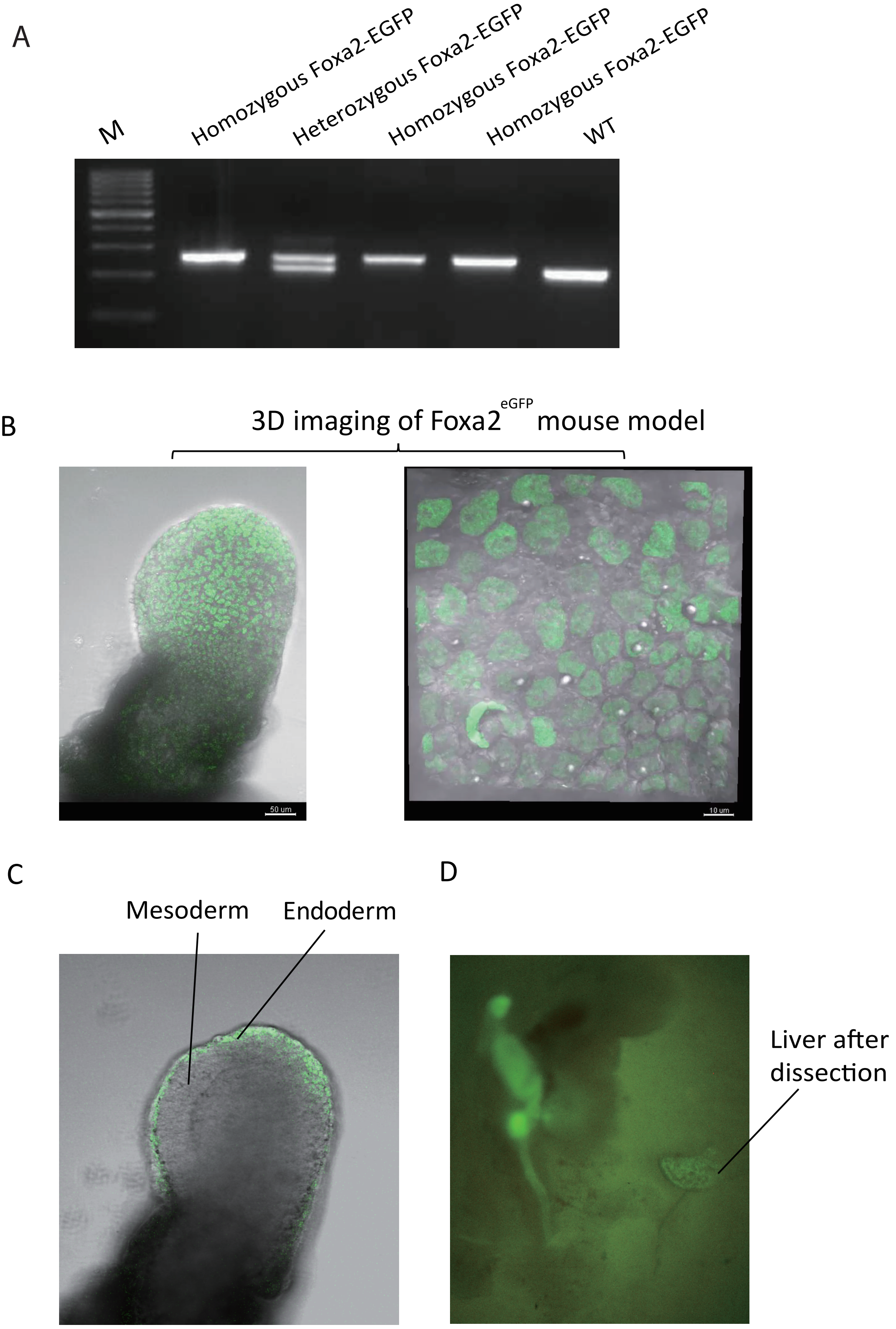

A
M
Homozygous Foxa2-EGFP
Heterozygous Foxa2-EGFP
Homozygous Foxa2-EGFP
Homozygous Foxa2-EGFP
WT
B
3D imaging of Foxa2 eGFP mouse model
50 um
10 um
C
Mesoderm
Endoderm
D
Liver after dissection

### Figure S2.

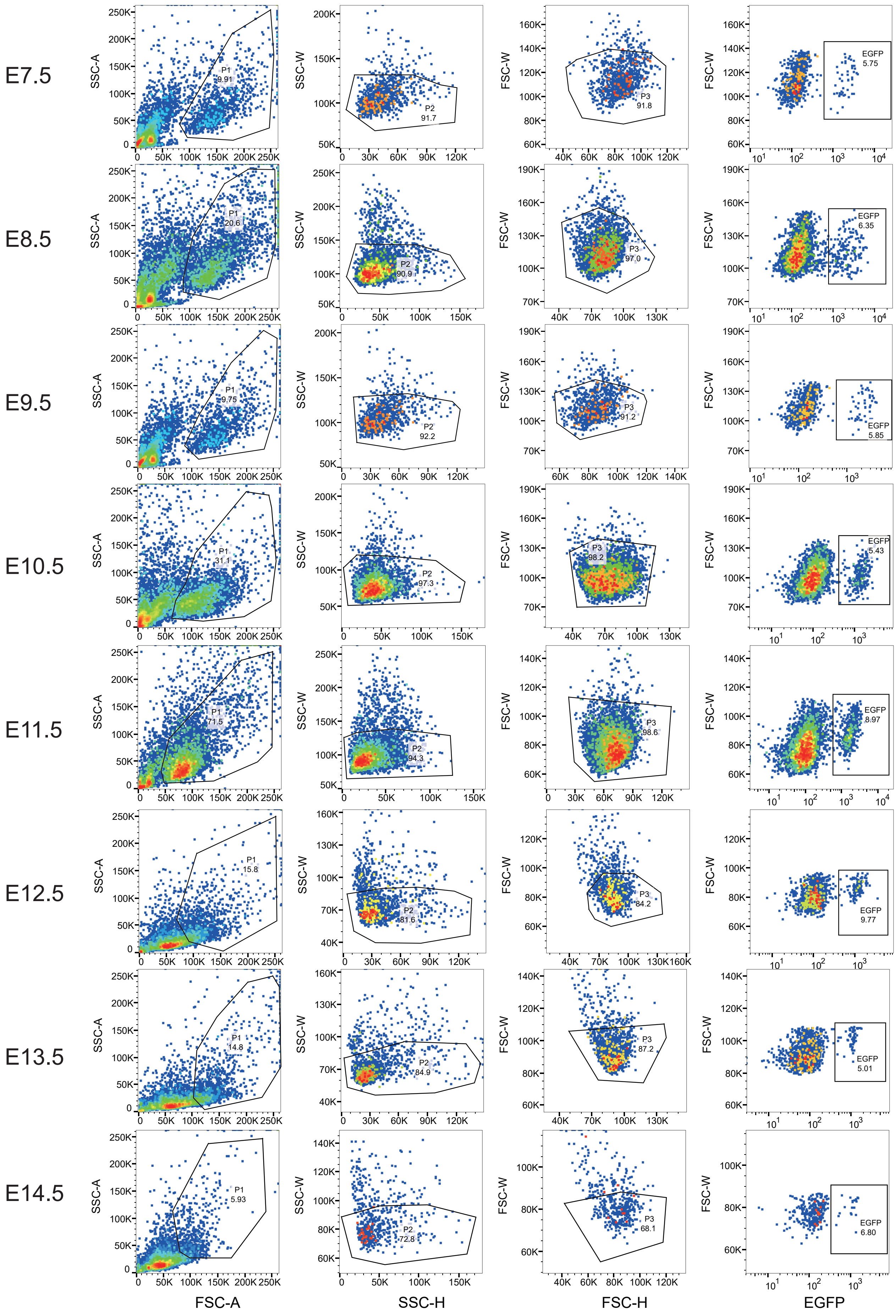
E7.5
E8.5
E9.5
E10.5
E11.5
E12.5
E13.5
E14.5
FSC-A
SSC-H
FSC-H
EGFP
SSC-A
SSC-W
FSC-W
P1
P2
P3
EGFP
9.91
91.7
91.8
5.75
20.6
90.9
97.0
6.35
9.75
92.2
91.2
5.85
31.1
97.3
98.2
5.43
71.5
94.3
98.6
8.97
15.8
81.6
84.2
9.77
14.8
84.9
87.2
5.01
5.93
72.8
68.1
6.80

Figure S3.

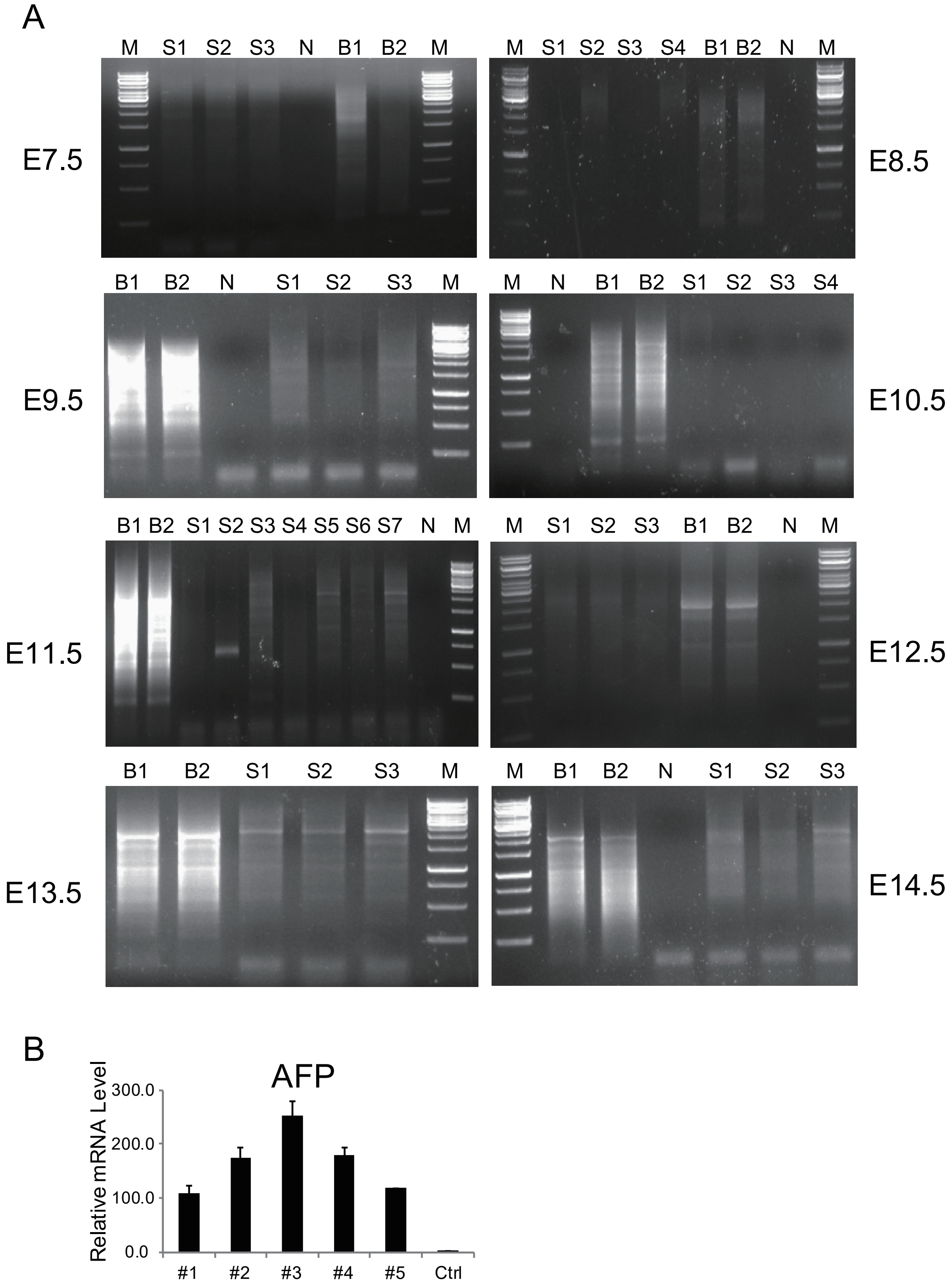

A
M S1 S2 S3 N B1 B2 M
E7.5
M S1 S2 S3 S4 B1 B2 N M
E8.5
B1 B2 N S1 S2 S3 M
E9.5
M N B1 B2 S1 S2 S3 S4
E10.5
B1 B2 S1 S2 S3 S4 S5 S6 S7 N M
E11.5
M S1 S2 S3 B1 B2 N M
E12.5
B1 B2 S1 S2 S3 M
E13.5
M B1 B2 N S1 S2 S3
E14.5
B
AFP
Relative mRNA Level
300.0
200.0
100.0
0.0
#1
#2
#3
#4
#5
Ctrl

Figure S4.

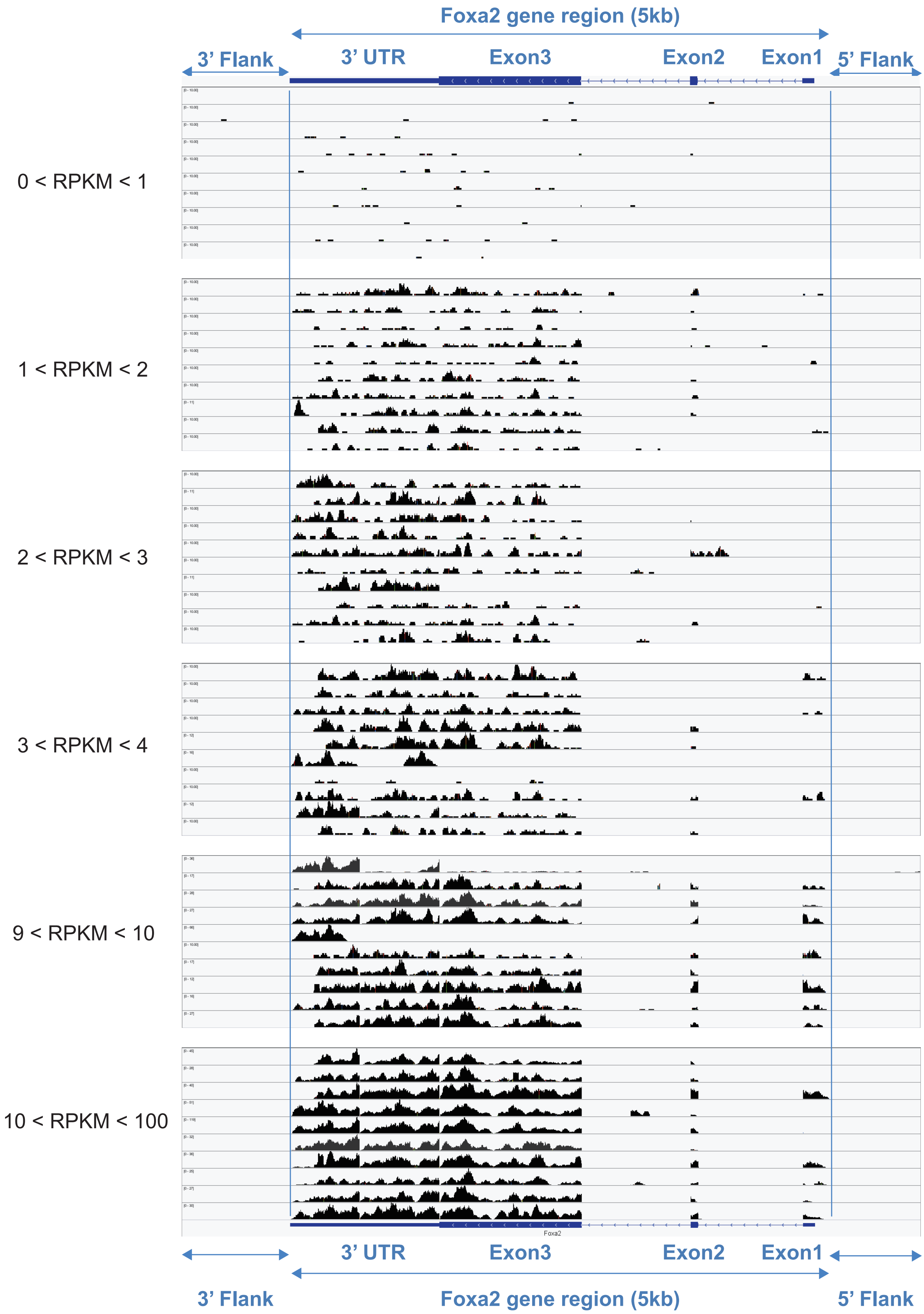

Foxa2 gene region (5kb)
3' Flank
3' UTR
Exon3
Exon2
Exon1
5' Flank
0 < RPKM < 1
1 < RPKM < 2
2 < RPKM < 3
3 < RPKM < 4
9 < RPKM < 10
10 < RPKM < 100
Foxa2
3' UTR
Exon3
Exon2
Exon1
3' Flank
Foxa2 gene region (5kb)
5' Flank

### Figure S5.

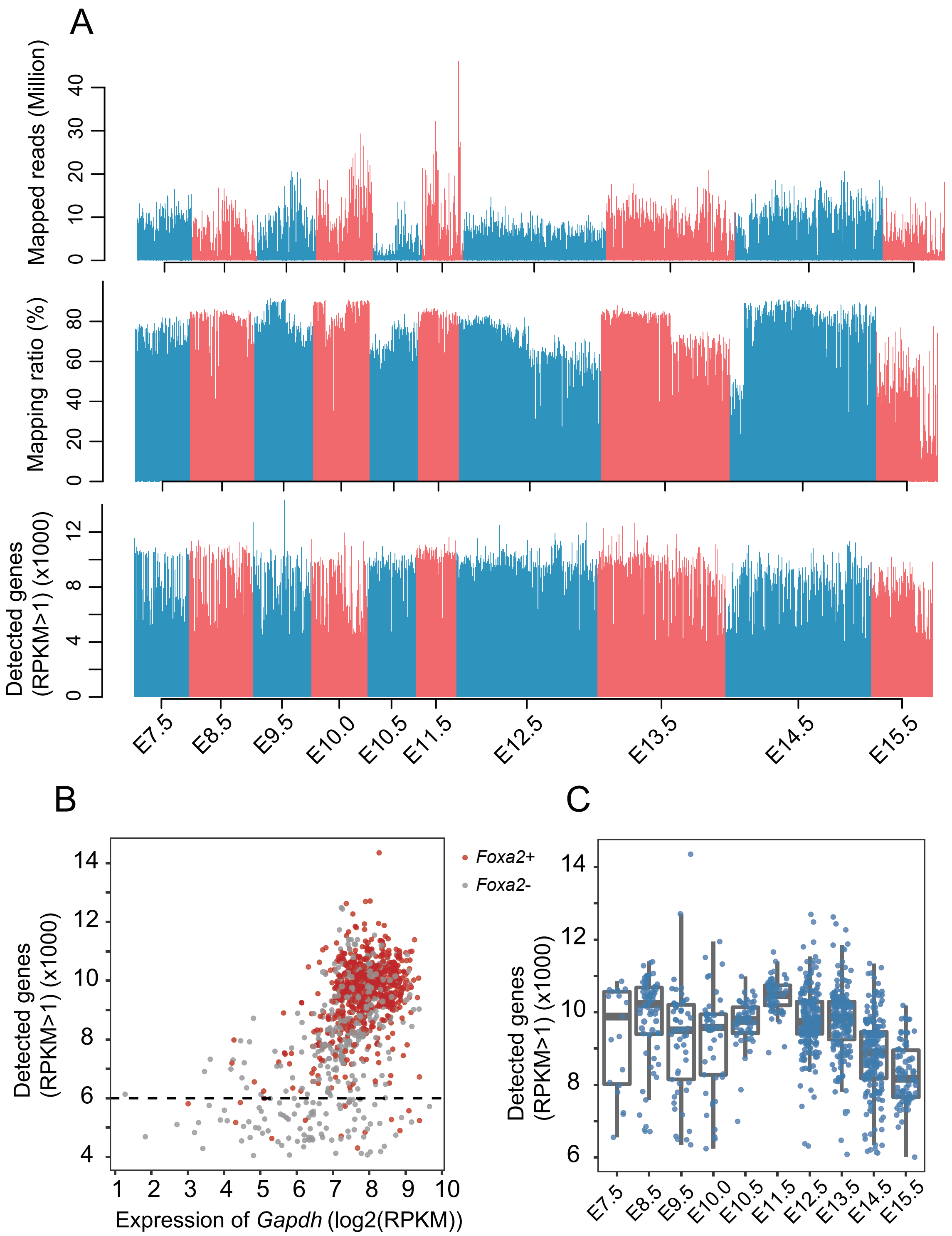
A
Mapped reads (Million)
Mapping ratio (%)
Detected genes (RPKM>1) (x1000)
E7.5
E8.5
E9.5
E10.0
E10.5
E11.5
E12.5
E13.5
E14.5
E15.5
B
Foxa2+
Foxa2-
Detected genes (RPKM>1) (x1000)
Expression of Gapdh (log2(RPKM))
C
Detected genes (RPKM>1) (x1000)

### Figure S6.

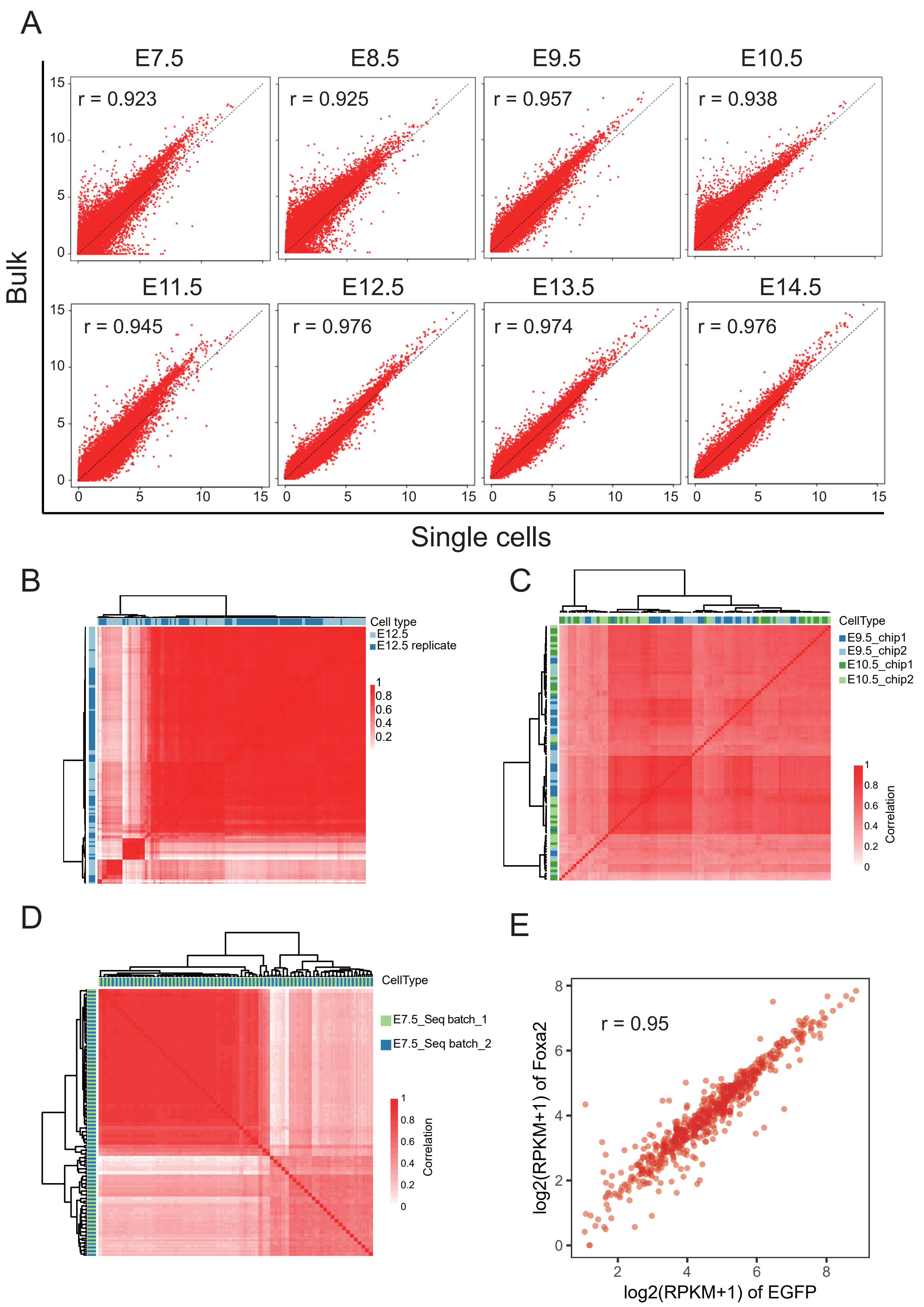
A
E7.5
E8.5
E9.5
E10.5
r = 0.923
r = 0.925
r = 0.957
r = 0.938
E11.5
E12.5
E13.5
E14.5
r = 0.945
r = 0.976
r = 0.974
r = 0.976
Bulk
Single cells
B
Cell type
E12.5
E12.5 replicate
C
CellType
E9.5_chip1
E9.5_chip2
E10.5_chip1
E10.5_chip2
Correlation
D
CellType
E7.5_Seq batch_1
E7.5_Seq batch_2
Correlation
E
r = 0.95
log2(RPKM+1) of Foxa2
log2(RPKM+1) of EGFP

Figure S7.

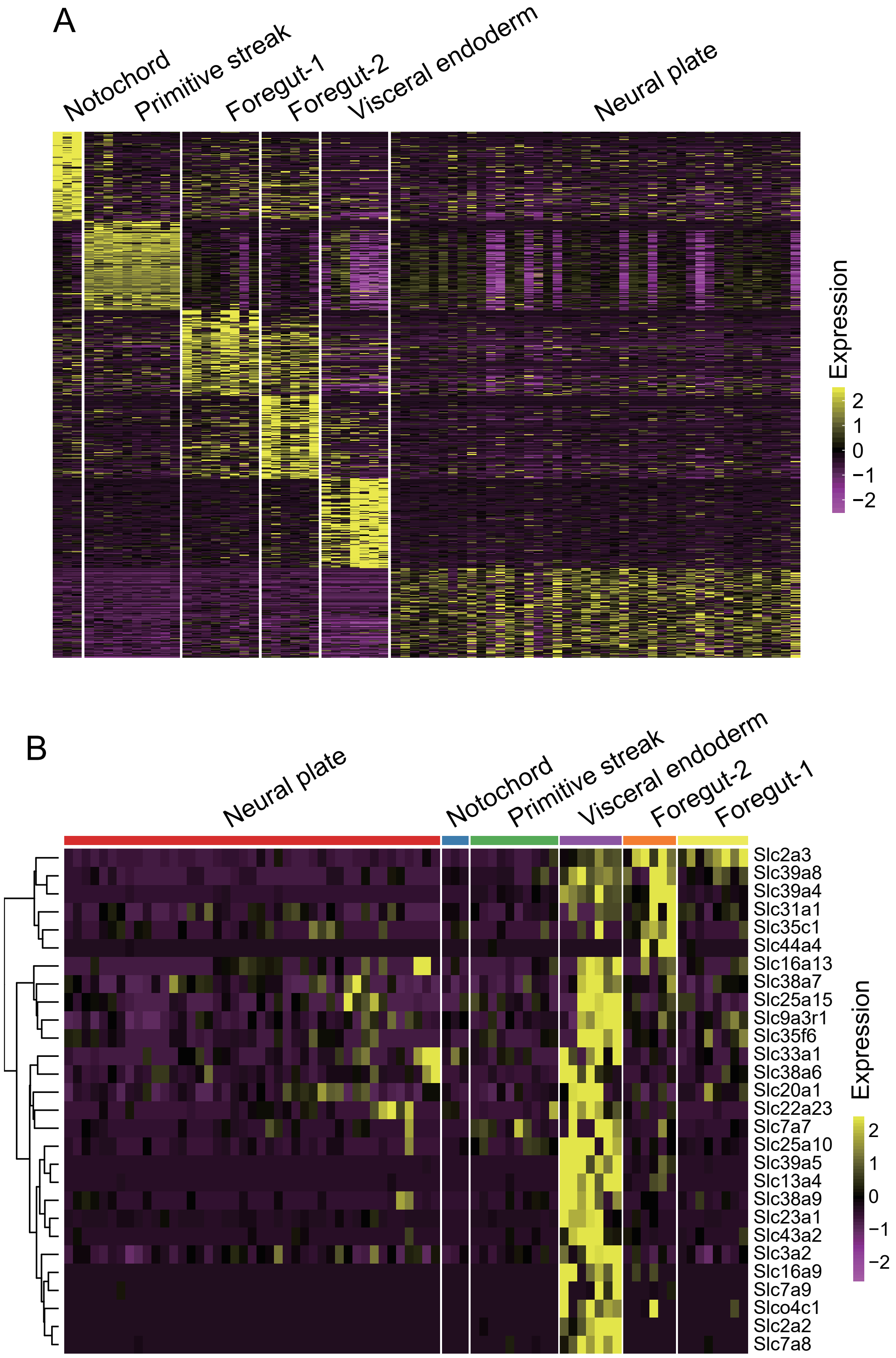
A
Notochord
Primitive streak
Foregut-1
Foregut-2
Visceral endoderm
Neural plate
Expression
2
1
0
−1
−2
B
Neural plate
Notochord
Primitive streak
Visceral endoderm
Foregut-2
Foregut-1
Slc2a3
Slc39a8
Slc39a4
Slc31a1
Slc35c1
Slc44a4
Slc16a13
Slc38a7
Slc25a15
Slc9a3r1
Slc35f6
Slc33a1
Slc38a6
Slc20a1
Slc22a23
Slc7a7
Slc25a10
Slc39a5
Slc13a4
Slc38a9
Slc23a1
Slc43a2
Slc3a2
Slc16a9
Slc7a9
Slco4c1
Slc2a2
Slc7a8
Expression
2
1
0
−1
−2

Figure S8.

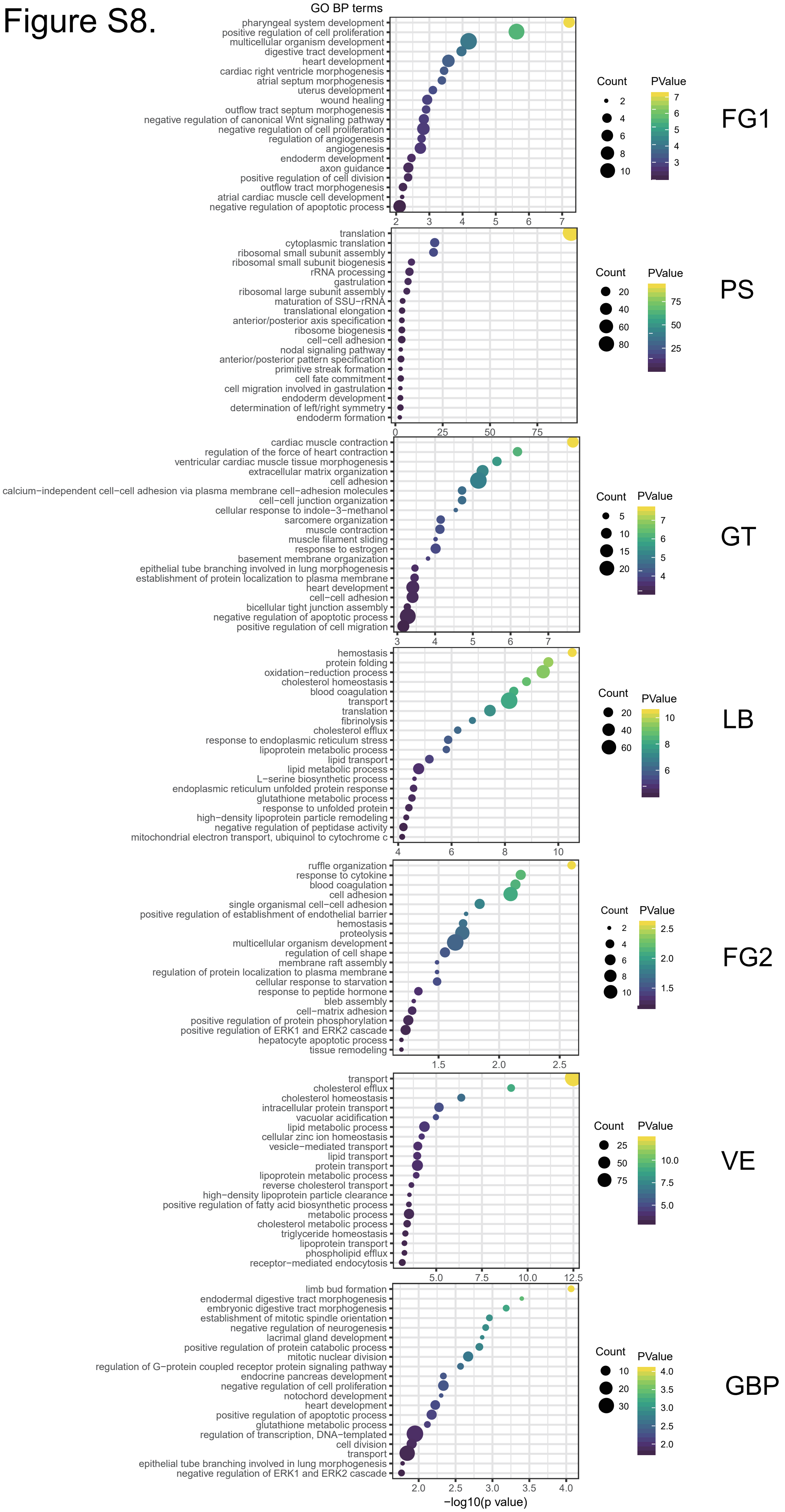

GO BP terms
pharyngeal system development
positive regulation of cell proliferation
multicellular organism development
digestive tract development
heart development
cardiac right ventricle morphogenesis
atrial septum morphogenesis
uterus development
wound healing
outflow tract septum morphogenesis
negative regulation of canonical Wnt signaling pathway
negative regulation of cell proliferation
regulation of angiogenesis
angiogenesis
endoderm development
axon guidance
positive regulation of cell division
outflow tract morphogenesis
atrial cardiac muscle cell development
negative regulation of apoptotic process
Count
PValue
FG1
translation
cytoplasmic translation
ribosomal small subunit assembly
ribosomal small subunit biogenesis
rRNA processing
gastrulation
ribosomal large subunit assembly
maturation of SSU−rRNA
translational elongation
anterior/posterior axis specification
ribosome biogenesis
cell−cell adhesion
nodal signaling pathway
anterior/posterior pattern specification
primitive streak formation
cell fate commitment
cell migration involved in gastrulation
endoderm development
determination of left/right symmetry
endoderm formation
PS
cardiac muscle contraction
regulation of the force of heart contraction
ventricular cardiac muscle tissue morphogenesis
extracellular matrix organization
cell adhesion
calcium−independent cell−cell adhesion via plasma membrane cell−adhesion molecules
cell−cell junction organization
cellular response to indole−3−methanol
sarcomere organization
muscle contraction
muscle filament sliding
response to estrogen
basement membrane organization
epithelial tube branching involved in lung morphogenesis
establishment of protein localization to plasma membrane
heart development
cell−cell adhesion
bicellular tight junction assembly
negative regulation of apoptotic process
positive regulation of cell migration
GT
hemostasis
protein folding
oxidation−reduction process
cholesterol homeostasis
blood coagulation
transport
translation
fibrinolysis
cholesterol efflux
response to endoplasmic reticulum stress
lipoprotein metabolic process
lipid transport
lipid metabolic process
L−serine biosynthetic process
endoplasmic reticulum unfolded protein response
glutathione metabolic process
response to unfolded protein
high−density lipoprotein particle remodeling
negative regulation of peptidase activity
mitochondrial electron transport, ubiquinol to cytochrome c
LB
ruffle organization
response to cytokine
blood coagulation
cell adhesion
single organismal cell−cell adhesion
positive regulation of establishment of endothelial barrier
hemostasis
proteolysis
multicellular organism development
regulation of cell shape
membrane raft assembly
regulation of protein localization to plasma membrane
cellular response to starvation
response to peptide hormone
bleb assembly
cell−matrix adhesion
positive regulation of protein phosphorylation
positive regulation of ERK1 and ERK2 cascade
hepatocyte apoptotic process
tissue remodeling
FG2
transport
cholesterol efflux
cholesterol homeostasis
intracellular protein transport
vacuolar acidification
lipid metabolic process
cellular zinc ion homeostasis
vesicle−mediated transport
lipid transport
protein transport
lipoprotein metabolic process
reverse cholesterol transport
high−density lipoprotein particle clearance
positive regulation of fatty acid biosynthetic process
metabolic process
cholesterol metabolic process
triglyceride homeostasis
lipoprotein transport
phospholipid efflux
receptor−mediated endocytosis
VE
limb bud formation
endodermal digestive tract morphogenesis
embryonic digestive tract morphogenesis
establishment of mitotic spindle orientation
negative regulation of neurogenesis
lacrimal gland development
positive regulation of protein catabolic process
mitotic nuclear division
regulation of G−protein coupled receptor protein signaling pathway
endocrine pancreas development
negative regulation of cell proliferation
notochord development
heart development
positive regulation of apoptotic process
glutathione metabolic process
regulation of transcription, DNA−templated
cell division
transport
epithelial tube branching involved in lung morphogenesis
negative regulation of ERK1 and ERK2 cascade
GBP
−log10(p value)

### Figure S9.

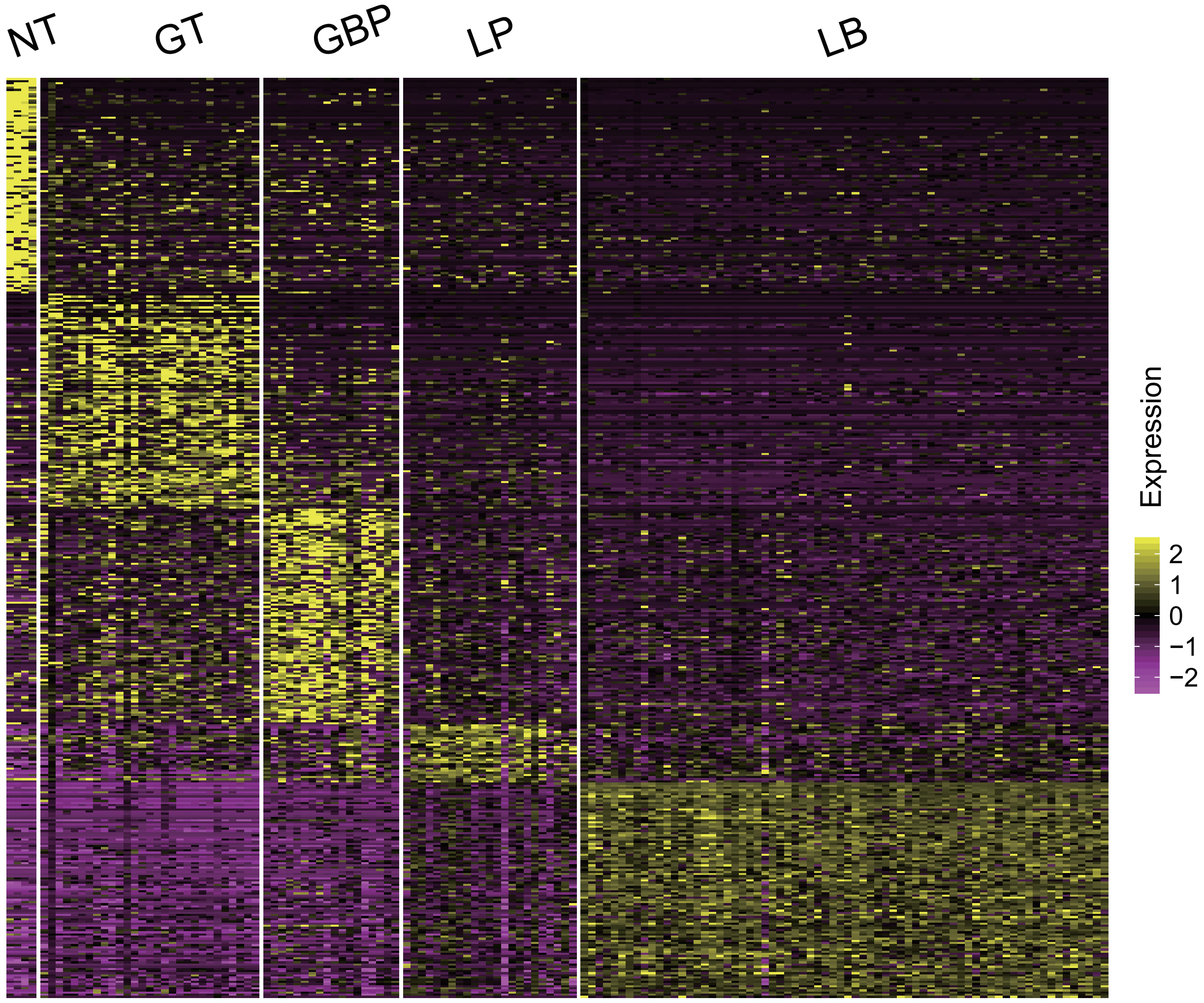

Figure S10.

A

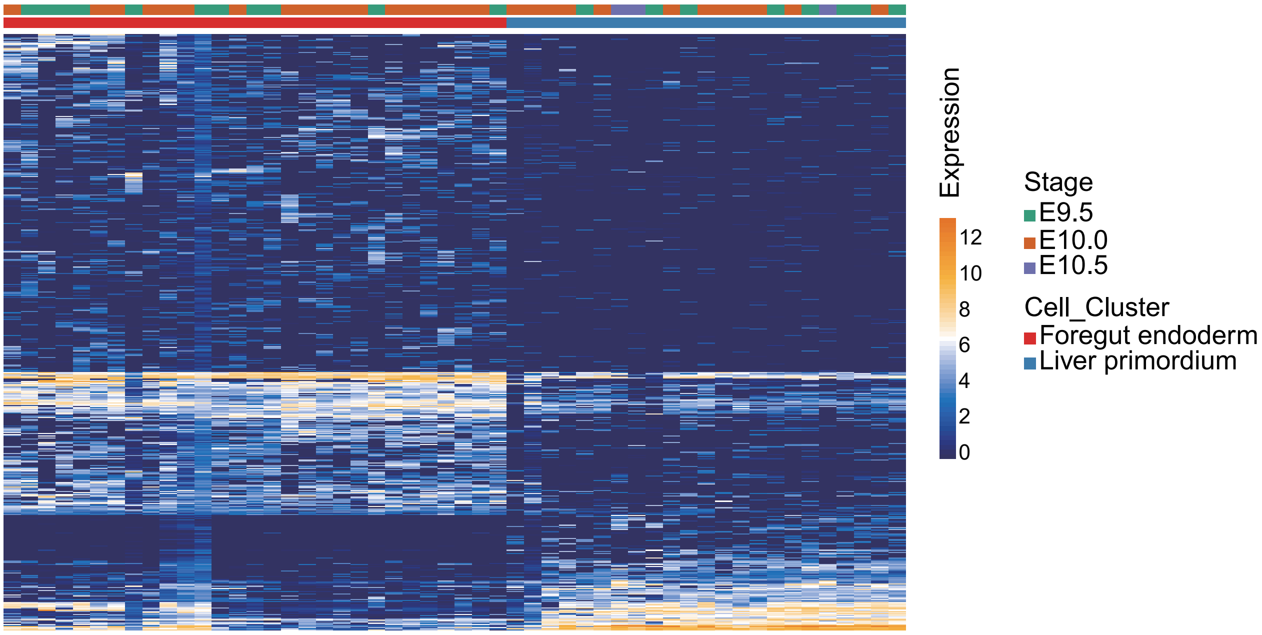


B

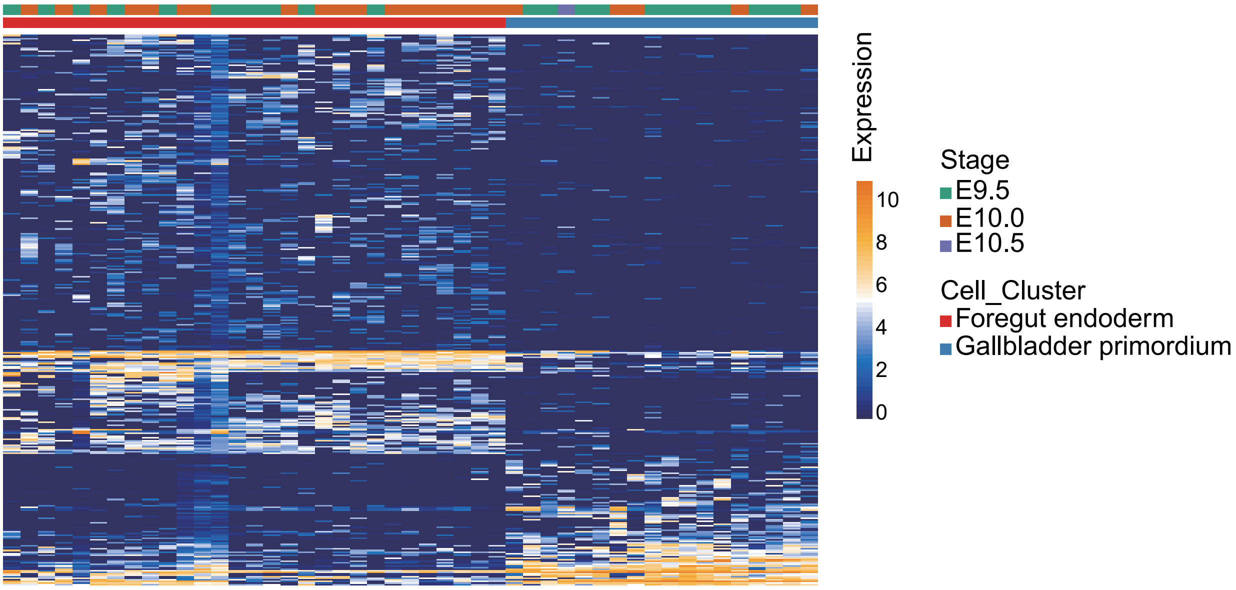


C

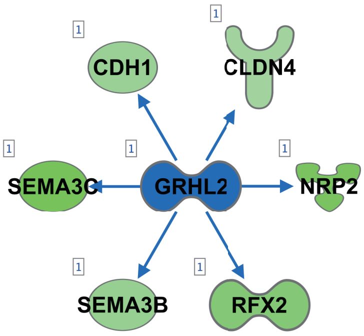

### Figure S11.

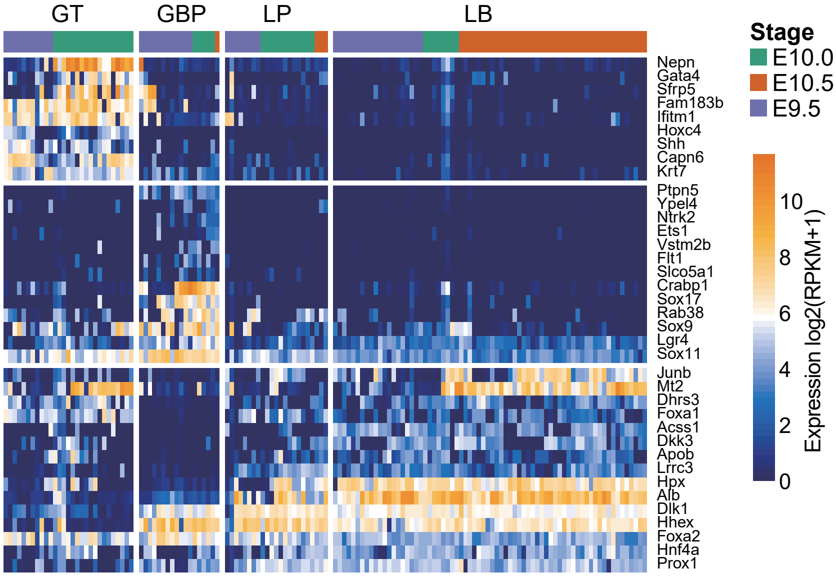
GT
GBP
LP
LB
Nepn
Gata4
Sfrp5
Fam183b
Ifitm1
Hoxc4
Shh
Capn6
Krt7
Ptpn5
Ypel4
Ntrk2
Ets1
Vstm2b
Flt1
Slco5a1
Crabp1
Sox17
Rab38
Sox9
Lgr4
Sox11
Junb
Mt2
Dhrs3
Foxa1
Acss1
Dkk3
Apob
Lrrc3
Hpx
Alb
Dlk1
Hhex
Foxa2
Hnf4a
Prox1
Stage
E10.0
E10.5
E9.5
10
8
6
4
2
0
Expression log2(RPKM+1)

### Figure S12.

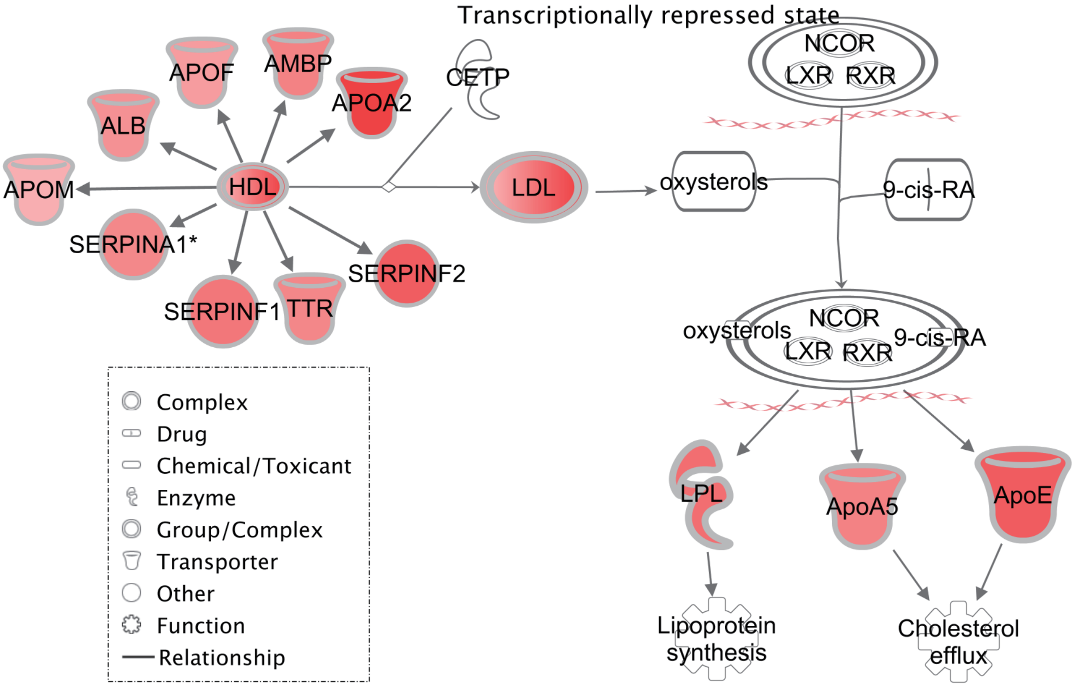


Figure legend

### Figure S13.

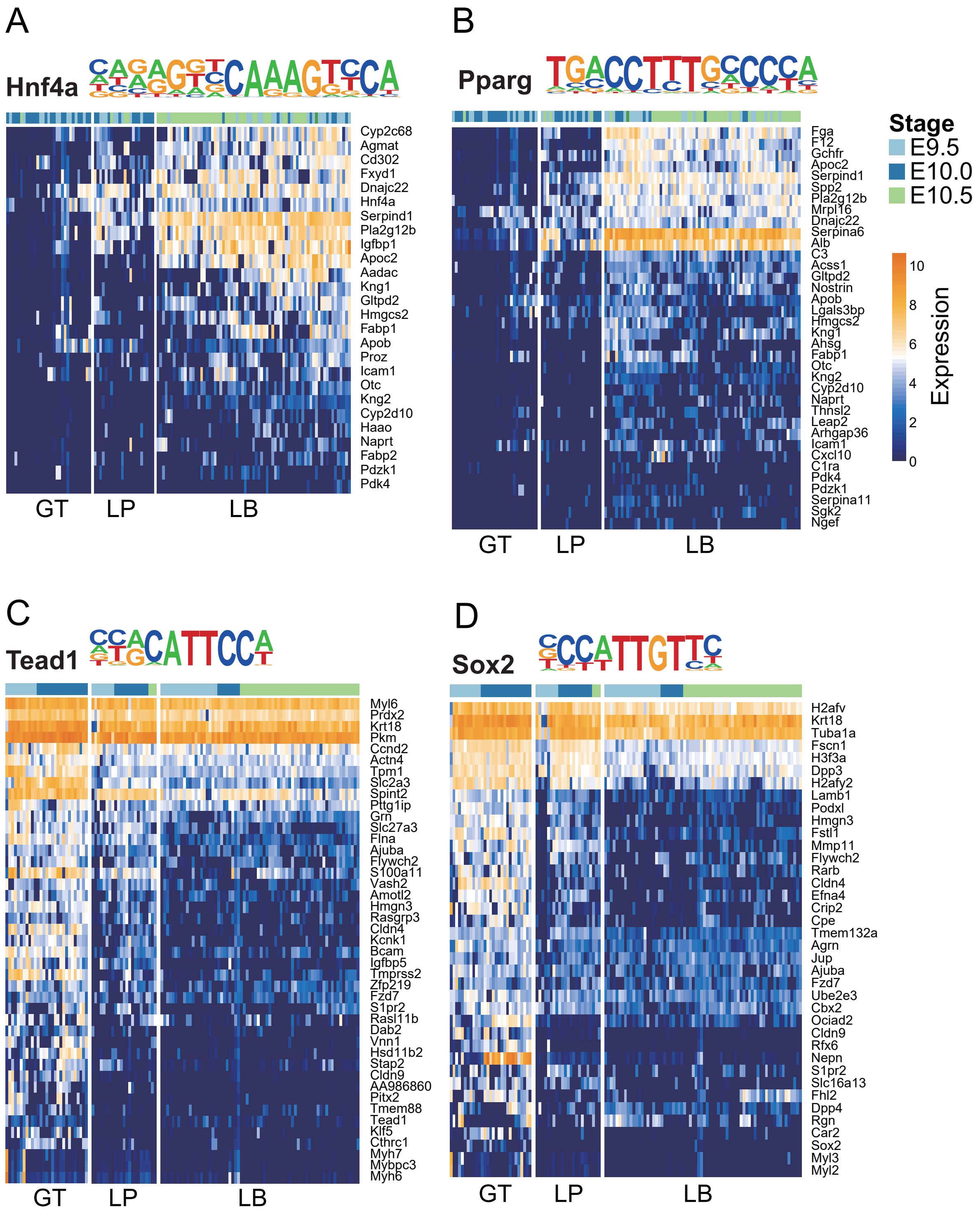
A
Hnf4a
Cyp2c68
Agmat
Cd302
Fxyd1
Dnajc22
Hnf4a
Serpind1
Pla2g12b
Igfbp1
Apoc2
Aadac
Kng1
Gltpd2
Hmgcs2
Fabp1
Apob
Proz
Icam1
Otc
Kng2
Cyp2d10
Haao
Naprt
Fabp2
Pdzk1
Pdk4
GT
LP
LB
B
Pparg
Fga
F12
Gchfr
Apoc2
Serpind1
Spp2
Pla2g12b
Mrpl16
Dnajc22
Serpina6
Alb
C3
Acss1
Gltpd2
Nostrin
Apob
Lgals3bp
Hmgcs2
Kng1
Ahsg
Fabp1
Otc
Kng2
Cyp2d10
Naprt
Thnsl2
Leap2
Arhgap36
Icam1
Cxcl10
C1ra
Pdk4
Pdzk1
Serpina11
Sgk2
Ngef
GT
LP
LB
Stage
E9.5
E10.0
E10.5
Expression
10
8
6
4
2
0
C
Tead1
Myl6
Prdx2
Krt18
Pkm
Ccnd2
Actn4
Tpm1
Slc2a3
Spint2
Pttg1ip
Grn
Slc27a3
Flna
Ajuba
Flywch2
S100a11
Vash2
Amotl2
Hmgn3
Rasgrp3
Cldn4
Kcnk1
Bcam
Igfbp5
Tmprss2
Zfp219
Fzd7
S1pr2
Rasl11b
Dab2
Vnn1
Hsd11b2
Stap2
Cldn9
AA986860
Pitx2
Tmem88
Tead1
Klf5
Cthrc1
Myh7
Mybpc3
Myh6
GT
LP
LB
D
Sox2
H2afv
Krt18
Tuba1a
Fscn1
H3f3a
Dpp3
H2afy2
Lamb1
Podxl
Hmgn3
Fstl1
Mmp11
Flywch2
Rarb
Cldn4
Efna4
Crip2
Cpe
Tmem132a
Agrn
Jup
Ajuba
Fzd7
Ube2e3
Cbx2
Ociad2
Cldn9
Rfx6
Nepn
S1pr2
Slc16a13
Fhl2
Dpp4
Rgn
Car2
Sox2
Myl3
Myl2
GT
LP
LB

### Figure S14.

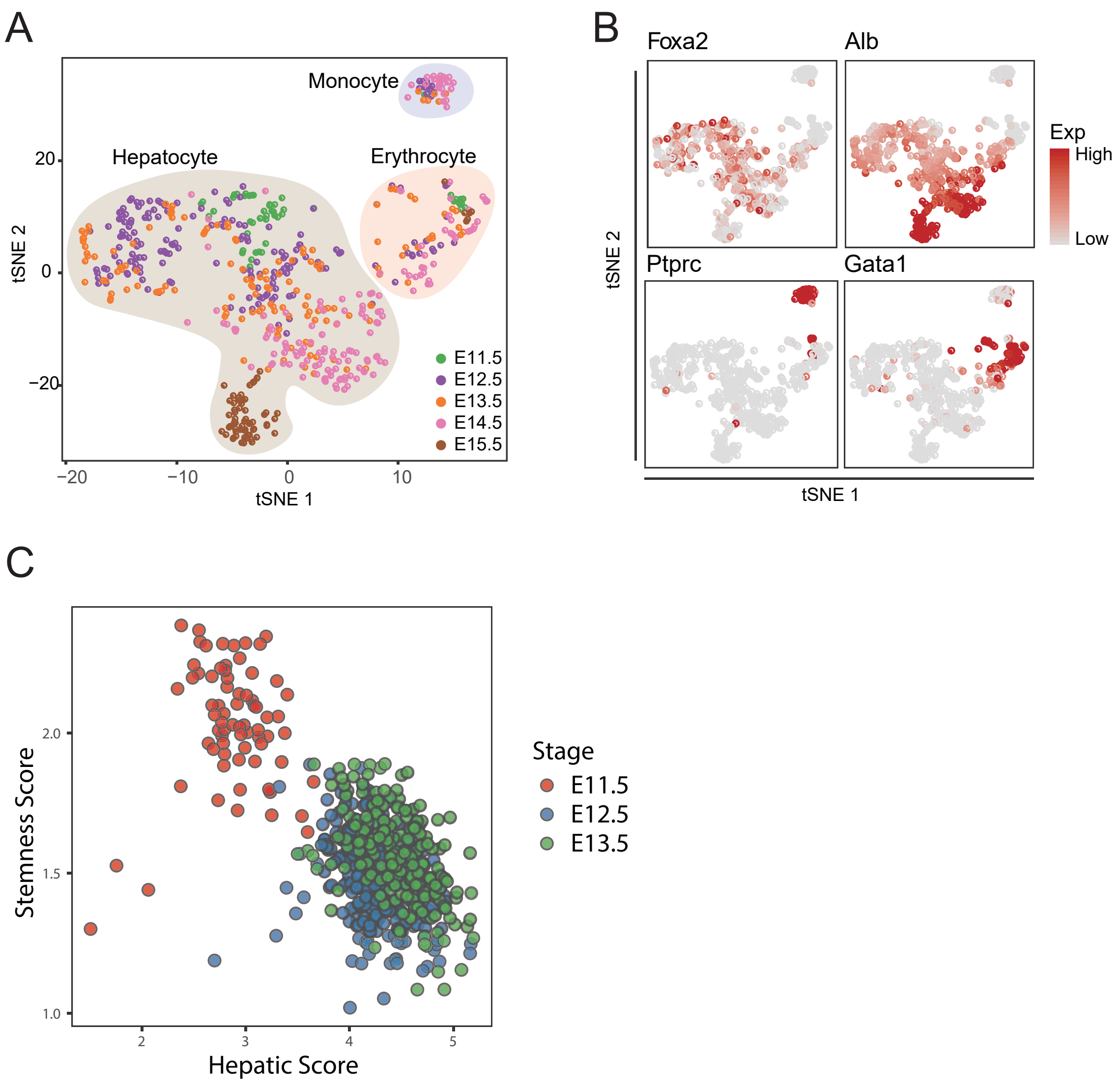
A
Monocyte
Hepatocyte
Erythrocyte
20
0
−20
tSNE 2
−20
−10
0
10
tSNE 1
E11.5
E12.5
E13.5
E14.5
E15.5
B
Foxa2
Alb
Ptprc
Gata1
tSNE 2
tSNE 1
Exp
High
Low
C
Stemness Score
2.0
1.5
1.0
2
3
4
5
Hepatic Score
Stage
E11.5
E12.5
E13.5
