## supplementary figure legend for "Characterizing the Emergence of Liver and Gallbladder from the Embryonic Endoderm through Single-Cell RNA-Seq"

Figure S1. The transgenic Foxa2$^{eGFP}$ reporter mouse model

a, Genotyping of homozygous Foxa2$^{eGFP}$ (269bp), heterozygous Foxa2$^{eGFP}$ (230/269bp) and Wild type (230bp) mice was shown.

b, 3D mouse imaging showed 6hs embryo culture of Foxa2$^{eGFP}$ at E6.5.

c, Endoderm and mesoderm of Foxa2$^{eGFP}$ mouse model.

d, Demonstration of microdissection of liver at E11.5.

Figure S2. FACS sorting data of Foxa2$^{eGFP}$ mice by eGFP during E7.5-E14.5. Doublets and multiplets were excluded by analysis of side scatter (SSC) and forward scatter (FSC).

Figure S3. The amplified cDNA was assessed by agarose gel (a) and qPCR of Afp, a hepatic marker (b), before library generation to ensure the sequencing quality. M, Marker; S1-S7, Single cell 1-7; N, negative control. B1-B2, bulk 1-2.

Figure S4. The gene expression level was characterized by RPKM (Reads Per Kilobase per Million mapped reads) and RPKM>1 was used as the threshold.

Figure S5. Sequencing data of scRNA-seq

a, the number of mapping reads, mapping ratio and detected genes of each single cell were shown.

b, the filtering threshold of detected genes was set as 6,000.

c, detected genes of each single cell during E7.5 to E15.5 after filtering.

S6. Quality control of single-cell RNA-seq.

a, correlation of average gene expression between single cells and bulk samples.

b-d, technical noise was assessed by calculating the Pearson correlation between experimental replicates (b), chips pooled different embryonic day (c) and sequencing batches (d).

e, as eGFP and Foxa2 were co-expressed in our mouse model, a high correlation was detected (r=0.95) between these two genes

Figure S7. Differentially expressed genes during E7.5-E8.5.

a, differentially expressed genes in Notochord, Primitive streak, Foregut-1, Foregut-2, visceral endoderm and Neural plate during E7.5-E8.5.

b, 28 genes of *Slc* family were highly expressed in visceral endoderm.

Figure S8. Gene Ontology analysis identified related functions of differentially expressed genes for each cell group, including Foregut-1 (FG1), Primitive Streak (PS), Gut tube (GT), Liver bud (LB), Foregut-2 (FG2), Visceral Endoderm (VE), Gallbladder Primordium (GBP).

Figure S9. Differentially expressed genes in Neural Tube, Gut tube, Gallbladder primordium (GBP), Liver primordium (LP), and Liver bud (LB) during E9.5-E11.5.

Figure S10. Differentially expressed genes among gut tube, liver primordium and gallbladder primordium.

a, 548 genes encoding transcription factors, enzymes, cytokines, transporters and kinases were differentially expressed between the gut tube and liver primordium.

b, 411 genes were found to be up-regulated in the gallbladder primordium compared with gut tube and hepatic cells from E9.5-E10.5.

c. *Grhl2* and its downstream targets (*Cdh1, Cldn4, Sema3c, Sema3b, Rfx2, Nrp2*) were found to be down-regulated in liver primordium, compared with gut tube.

Figure S11. Potential new markers were identified in gallbladder primordium.

*Crabp1, Rab38, Flt1, Slco5a1, Ptpn5, Vstm2b, Ntrk2, Ets1* and *Ypel4* were identified as potential new markers in the gallbladder primordium, while barely expressed in the gut tube and hepatic cells. By contrast, *Junb, Hpx, Mt2, Irrc3, Dkk3, Apob, Acss1* and *Dhrs3* were negative in the gallbladder.

Figure S12. The liver X receptors/retinoid X receptors (LXR/RXR) pathway was significantly up-regulated in the liver primordium compared with the gut tube.

Figure S13. Motif analysis showed targets of HNF4A and PPARG were up-regulated in hepatoblasts, while targets of SOX2 and TEAD1 were found to be down-regualted.

Figure S14. Dynamic gene expression of hepatoblast maturation during E11.5-E15.5.

a, t-SNE analysis identified cell clustering of single-cells from E11.5-E15.5.

b, the gene expression of specific markers, such as *Foxa2, Alb, Ptprc, Gata1,* were shown.

c, 720 cells generated by MIRALCS method validated that the hepatic score of hepatoblasts/hepatocytes increased, while the stemness score cell decreased during liver maturation.
