## supplementary table for "Characterizing the Emergence of Liver and Gallbladder from the Embryonic Endoderm through Single-Cell RNA-Seq"

### Cells statistic

| | Total | Qualified | Foxa2_EGFP >0 | Foxa2_EGFP >1 |
|---|---|---|---|---|
| E7.5 | 96 | 61 (64.89%) | 29 (47.54%) | 19 (31.15%) |
| E8.5 | 94 | 73 (77.66%) | 65 (89.04%) | 60 (82.19%) |
| E9.5 | 84 | 66 (78.57%) | 61 (92.42%) | 53 (80.30%) |
| E10.5 | 80 | 60 (75%) | 47 (78.33%) | 46 (76.67%) |
| E10.0 | 74 | 56 (75.68%) | 53 (94.64%) | 45 (80.36%) |
| E11.5 | 65 | 51 (78.46%) | 37 (72.55%) | 36 (70.59%) |
| E12.5 | 94 | 89 (94.68%) | 76 (85.39%) | 71 (79.78%) |
| E12.5_B2 | 94 | 90 (95.74%) | 66 (73.33%) | 64 (71.11%) |
| E13.5 | 94 | 85 (90.43%) | 70 (82.35%) | 67 (78.82%) |
| E13.5_B2 | 94 | 61 (64.89%) | 36 (59.02%) | 32 (52.46%) |
| E14.5 | 94 | 78 (82.98%) | 59 (75.64%) | 46 (58.97%) |
| E14.5_GFPn | 96 | 13 (13.54%) | 0 (0%) | 0 (0%) |
| E14.5_GFPp | 94 | 74 (78.72%) | 48 (64.86%) | 39 (52.7%) |
| E15.5 | 95 | 66 (69.47%) | 27 (40.91%) | 23 (34.85%) |
| Total | 1246 | 922 (74.00%) | 674 (73.10%) | 601 (65.18%) |

(Qualified standard: Mapped reads > 1M; Detected genes (RPKM>1) > 6000)

### Feature genes list

| Hepatic_gene | Epithelial_gene | Mesenchymal_gene | Stemness_g | Proliferation_gene |
|---|---|---|---|---|
| Alb | Epcam | Vim | Fzd7 | Clspn |
| Ttr | Cdh1 | S100a4 | Klf4 | Blm |
| Afp | Ocln | Acta2 | Tbx3 | Ung |
| Apoc1 | Tjp1 | Fn1 | Tpt1 | E2f8 |
| Apoc2 | Tjp2 | Cdh2 | Rif1 | Pola1 |
| Apoa4 | Tjp3 | Col1a1 | Med28 | Tipin |
| Apoa2 | Col4a1 | Col1a2 | Med27 | Prim1 |
| Apob | Col4a2 | Col3a1 | Med30 | Ccne2 |
| Apoa1 | Col4a3 | | Nanog | Rpa2 |
| Apod | Col4a4 | | Rtf1 | Tyms |
| Apoe | Col4a5 | | Leo1 | Cdc45 |
| Apof | Col4a6 | | Padi4 | Cdca7 |
| Apoc4 | Cldn1 | | Stag2 | Casp8ap2 |
| Apoc3 | Cldn10 | | Ctr9 | Wdr76 |
| Apoa5 | Cldn11 | | Esrrb | Fen1 |
| Apoh | Cldn12 | | Med14 | Hells |
| Apol6 | Cldn13 | | Nkap | Exo1 |
| Apom | Cldn14 | | Mettl14 | Gins2 |
| Apoo | Cldn15 | | Zcchc11 | Cdc6 |
| F11 | Cldn16 | | Setd1a | Rad51ap1 |
| F2rl2 | Cldn17 | | Panct2 | Msh2 |
| F12 | Cldn18 | | Pramel7 | Dtl |
| F2rl3 | Cldn19 | | Ctnnb1 | Gmnn |
| F10 | Cldn2 | | Ddx6 | Nasp |
| F13a1 | Cldn20 | | Pou5f1 | Usp1 |
| F8 | Cldn22 | | Phf19 | Brip1 |
| F9 | Cldn23 | | Eif4enif1 | Atad2 |
| F7 | Cldn24 | | Pelo | Mcm2 |
| F13b | Cldn3 | | Mtf2 | Mcm4 |
| Fgg | Cldn34a | | Sox2 | Mcm5 |
| Fga | Cldn34b1 | | Stat3 | Mcm6 |
| F5 | Cldn34b2 | | Tcl1 | Rad51 |
| Fgb | Cldn34b3 | | Setd6 | Slbp |
| F3 | Cldn34b4 | | Prdm16 | Pold3 |
| F2 | Cldn34c1 | | Trim8 | Uhrf1 |
| F2r | Cldn34c4 | | Arid1a | Ubr7 |
| Gsta1 | Cldn34d | | Sall4 | Rfc2 |
| Gsta2 | Cldn4 | | Zfp358 | Rrm2 |
| Gsta3 | Cldn5 | | Dicer1 | Rrm1 |
| Gsta4 | Cldn6 | | Miat | Pcna |
| Gstt1 | Cldn7 | | Prdm14 | Chaf1b |
| Gstt2 | Cldn8 | | Bmpr1a | Dscc1 |
| Gstm5 | Cldn9 | | Smc3 | Kif23 |
| Gstm1 | Cldnd1 | | Eif4e | Gas2l3 |
| Gstm2 | Cldnd2 | | Npm1 | Ttk |
| Gstm3 | Krt1 | | Smarca4 | Aurka |
| Gstm4 | Krt10 | | Tfap2c | Ctcf |
| Gstk1 | Krt12 | | Med7 | Rangap1 |
| Gstz1 | Krt13 | | Lsm1 | Aurkb |
| Gsto2 | Krt14 | | Dis3l2 | Gtse1 |
| Gsto1 | Krt15 | | Fgf4 | Cbx5 |
| Gstp1 | Krt16 | | Fgfr1 | Kif2c |
| Serpina10 | Krt17 | | Med10 | G2e3 |
| Serpina11 | Krt18 | | Mettl3 | Cdca8 |

| | | | |
|---|---|---|---|
| Serpina12 | Krt19 | Dppa2 | Cdca2 |
| Serpinh1 | Krt2 | Med15 | Top2a |
| Serpina9 | Krt20 | Med21 | Lbr |
| Serpina7 | Krt222 | Med24 | Cdca3 |
| Serpine2 | Krt23 | Smc1a | Cdk1 |
| Serpine3 | Krt24 | Tet1 | Kif11 |
| Serpina6 | Krt25 | Paf1 | Anp32e |
| Serpina5 | Krt26 | Med6 | Tpx2 |
| Serpine1 | Krt27 | Med17 | Nusap1 |
| Serpinc1 | Krt28 | Apc | Hn1 |
| Serpinb10 | Krt31 | Eomes | Tacc3 |
| Serpinb11 | Krt32 | Lif | Ube2c |
| Serpinb12 | Krt33a | Nodal | Ect2 |
| Serpinb13 | Krt33b | Med12 | Ncapd2 |
| Serping1 | Krt34 | Nipbl | Ckap2l |
| Serpini1 | Krt35 | Lin28a | Cks1b |
| Serpini2 | Krt36 | Cdc73 | Hmgb2 |
| Serpinb9 | Krt39 | | Nek2 |
| Serpinf1 | Krt4 | | Anln |
| Serpinf2 | Krt40 | | Hmmr |
| Serpinb5 | Krt42 | | Cenpa |
| Serpinb8 | Krt5 | | Hjurp |
| Serpinb7 | Krt6a | | Bub1 |
| Serpinb2 | Krt6b | | Tubb4b |
| Serpind1 | Krt7 | | Ckap2 |
| | Krt71 | | Mki67 |
| | Krt72 | | Ckap5 |
| | Krt73 | | Dlgap5 |
| | Krt74 | | Psrc1 |
| | Krt75 | | Nuf2 |
| | Krt76 | | Cenpf |
| | Krt77 | | Cenpe |
| | Krt78 | | Ndc80 |
| | Krt79 | | Birc5 |
| | Krt8 | | Cdc20 |
| | Krt80 | | Cdc25c |
| | Krt81 | | Smc4 |
| | Krt82 | | Fam64a |
| | Krt83 | | Ccnb2 |
| | Krt84 | | Cks2 |
| | Krt85 | | Kif20b |
| | Krt86 | | Tmpo |
| | Krt9 | | |

### Differentially expressed genes (DEGs) of cells from E7.5-E8.

**Instructions:**

- **p_val :** p-value (unadjusted)
- **avg_logFC :** log fold-chage of the average expression between the given cell cluster and the
- **pct.1 :** The ratio of expression in the given cell cluster
- **pct.2 :** The ratio of expression in the left cell clusters from E7.5-E8.5
- **p_val_adj :** Adjusted p-value, based on bonferroni correction using all genes in the dataset.

| | p_val | avg_logFC | pct.1 | pct.2 | p_val_adj | cell cluster |
|---|---|---|---|---|---|---|
| Nkx2-3 | 3.13E-83 | 1.089190235 | 0.714 | 0.508 | 3.93E-79 | Foregut |
| Cldn4 | 2.61E-77 | 1.159223817 | 1 | 0.415 | 3.28E-73 | Foregut |
| Atp2c2 | 9.25E-47 | 0.467218724 | 0.643 | 0.277 | 1.16E-42 | Foregut |
| Akap5 | 1.68E-43 | 0.358529113 | 1 | 0.923 | 2.12E-39 | Foregut |
| Krt7 | 3.72E-43 | 0.463636044 | 1 | 1 | 4.68E-39 | Foregut |
| Pyy | 4.46E-39 | 1.887131314 | 0.786 | 0.231 | 5.61E-35 | Foregut |
| Cpm | 7.15E-35 | 1.151741523 | 1 | 0.954 | 8.99E-31 | Foregut |
| Nkx2-6 | 1.08E-31 | 0.28712096 | 0.643 | 0.2 | 1.36E-27 | Foregut |
| Isl1 | 1.13E-31 | 0.536125482 | 0.786 | 0.446 | 1.42E-27 | Foregut |
| Dok5 | 6.64E-31 | 0.376664906 | 0.786 | 0.477 | 8.35E-27 | Foregut |
| Nepn | 8.64E-31 | 3.073694455 | 0.929 | 0.492 | 1.09E-26 | Foregut |
| Bcam | 7.08E-27 | 0.901870106 | 1 | 0.738 | 8.90E-23 | Foregut |
| Npnt | 1.44E-26 | 0.454850311 | 0.929 | 0.462 | 1.82E-22 | Foregut |
| Angpt2 | 7.59E-25 | 0.51872691 | 0.714 | 0.477 | 9.55E-21 | Foregut |
| Igf2 | 6.38E-24 | 0.988357385 | 1 | 0.985 | 8.02E-20 | Foregut |
| Fblim1 | 5.07E-22 | 0.468982913 | 1 | 0.815 | 6.37E-18 | Foregut |
| Nrp1 | 2.10E-21 | 0.501194126 | 0.857 | 0.569 | 2.64E-17 | Foregut |
| Cldn3 | 1.87E-18 | 0.475004072 | 0.786 | 0.062 | 2.35E-14 | Foregut |
| Tes | 2.27E-18 | 0.845716737 | 1 | 0.846 | 2.86E-14 | Foregut |
| Ripply3 | 2.72E-18 | 0.33771761 | 1 | 1 | 3.42E-14 | Foregut |
| Efna1 | 4.75E-18 | 0.719042656 | 0.786 | 0.308 | 5.97E-14 | Foregut |
| Mmp11 | 6.79E-18 | 0.766888968 | 1 | 0.508 | 8.54E-14 | Foregut |
| Ldoc1 | 1.62E-17 | 0.307825694 | 0.571 | 0.062 | 2.04E-13 | Foregut |
| Hs3st3b1 | 1.05E-16 | 0.339912815 | 0.929 | 0.508 | 1.32E-12 | Foregut |
| Wbp5 | 2.19E-16 | 1.506903105 | 1 | 1 | 2.75E-12 | Foregut |
| Epcam | 4.61E-16 | 1.850988326 | 1 | 1 | 5.80E-12 | Foregut |
| Tpm1 | 8.40E-16 | 1.088363697 | 1 | 1 | 1.06E-11 | Foregut |
| Wfdc2 | 1.09E-15 | 0.916468662 | 0.929 | 0.738 | 1.37E-11 | Foregut |
| Cd55 | 2.96E-15 | 0.275548025 | 0.714 | 0.308 | 3.73E-11 | Foregut |
| Jup | 8.20E-15 | 0.324319686 | 1 | 0.923 | 1.03E-10 | Foregut |
| Gm8113 | 8.39E-15 | 1.061756381 | 1 | 0.954 | 1.06E-10 | Foregut |
| Spint1 | 1.08E-14 | 0.852698326 | 1 | 0.892 | 1.36E-10 | Foregut |
| F11r | 1.85E-14 | 1.194996259 | 1 | 0.969 | 2.33E-10 | Foregut |
| Cpn1 | 3.67E-14 | 0.621389242 | 1 | 0.862 | 4.62E-10 | Foregut |
| St14 | 4.73E-14 | 0.523692061 | 1 | 0.631 | 5.95E-10 | Foregut |
| Arrdc1 | 6.64E-14 | 0.340250488 | 1 | 0.569 | 8.35E-10 | Foregut |
| Tmem59l | 8.94E-14 | 0.438588937 | 0.786 | 0.446 | 1.12E-09 | Foregut |
| Krt18 | 1.05E-13 | 2.027708049 | 1 | 1 | 1.32E-09 | Foregut |
| Spint2 | 1.48E-13 | 1.355289598 | 1 | 0.969 | 1.87E-09 | Foregut |
| Itga3 | 1.78E-13 | 0.467298214 | 1 | 0.738 | 2.23E-09 | Foregut |
| Dsc2 | 3.25E-13 | 0.310761169 | 0.929 | 0.631 | 4.09E-09 | Foregut |
| Mapk13 | 6.26E-13 | 0.548335354 | 0.857 | 0.4 | 7.87E-09 | Foregut |
| Lpar3 | 1.24E-12 | 0.728578749 | 0.857 | 0.538 | 1.56E-08 | Foregut |
| Slc2a3 | 1.50E-12 | 1.766512969 | 1 | 1 | 1.88E-08 | Foregut |
| Rasgrp3 | 2.31E-12 | 0.259128325 | 1 | 0.662 | 2.91E-08 | Foregut |

| | | | | | | |
|---|---|---|---|---|---|---|
| Kitl | 3.10E-12 | 0.274954832 | 0.857 | 0.662 | 3.90E-08 | Foregut |
| Mterf3 | 3.91E-12 | 0.317282629 | 1 | 1 | 4.92E-08 | Foregut |
| Krt8 | 4.64E-12 | 1.58634582 | 1 | 1 | 5.84E-08 | Foregut |
| Capn6 | 5.23E-12 | 0.478992087 | 1 | 0.892 | 6.57E-08 | Foregut |
| Cldn7 | 6.89E-12 | 0.956975701 | 1 | 0.862 | 8.67E-08 | Foregut |
| Dusp9 | 7.47E-12 | 0.642289897 | 1 | 0.692 | 9.40E-08 | Foregut |
| Cldn6 | 8.99E-12 | 2.196791131 | 1 | 0.985 | 1.13E-07 | Foregut |
| Tspan7 | 1.25E-11 | 0.507294915 | 1 | 1 | 1.57E-07 | Foregut |
| Gas6 | 1.78E-11 | 0.69601228 | 0.643 | 0.369 | 2.23E-07 | Foregut |
| Fn1 | 2.61E-11 | 0.635115458 | 1 | 1 | 3.29E-07 | Foregut |
| Tfpi | 1.04E-10 | 0.308504751 | 0.786 | 0.662 | 1.31E-06 | Foregut |
| Sh3glb1 | 1.33E-10 | 0.265351709 | 1 | 0.846 | 1.67E-06 | Foregut |
| Bex1 | 1.43E-10 | 1.346725261 | 1 | 1 | 1.80E-06 | Foregut |
| Tspan12 | 1.71E-10 | 0.896824101 | 1 | 0.954 | 2.15E-06 | Foregut |
| Igfbp3 | 1.79E-10 | 0.399714969 | 0.786 | 0.692 | 2.25E-06 | Foregut |
| Gfpt2 | 1.94E-10 | 0.467510824 | 1 | 0.785 | 2.44E-06 | Foregut |
| Apela | 1.94E-10 | 0.889470331 | 1 | 0.877 | 2.45E-06 | Foregut |
| Afap1l1 | 1.96E-10 | 0.417801032 | 0.857 | 0.462 | 2.47E-06 | Foregut |
| Amph | 2.09E-10 | 0.267581238 | 0.643 | 0.446 | 2.63E-06 | Foregut |
| Ngfrap1 | 2.91E-10 | 1.195093283 | 1 | 1 | 3.66E-06 | Foregut |
| Pvrl2 | 4.04E-10 | 0.8585494 | 1 | 1 | 5.08E-06 | Foregut |
| Parm1 | 4.33E-10 | 0.285114464 | 0.929 | 0.415 | 5.44E-06 | Foregut |
| Tmem102 | 6.12E-10 | 0.254668341 | 0.714 | 0.169 | 7.70E-06 | Foregut |
| Rcan2 | 6.71E-10 | 0.287448896 | 0.786 | 0.415 | 8.43E-06 | Foregut |
| Vamp8 | 7.08E-10 | 0.906538094 | 1 | 0.923 | 8.90E-06 | Foregut |
| Lad1 | 7.12E-10 | 0.540799983 | 0.857 | 0.538 | 8.96E-06 | Foregut |
| Cmtm8 | 7.88E-10 | 0.786438197 | 1 | 0.846 | 9.91E-06 | Foregut |
| Spin2c | 9.73E-10 | 0.548561287 | 1 | 0.631 | 1.22E-05 | Foregut |
| Gpc3 | 1.20E-09 | 1.231701554 | 1 | 1 | 1.51E-05 | Foregut |
| Kcnk1 | 1.47E-09 | 0.577179129 | 1 | 0.708 | 1.85E-05 | Foregut |
| Lipa | 1.65E-09 | 0.544039005 | 1 | 0.815 | 2.08E-05 | Foregut |
| Peg3 | 1.99E-09 | 0.761022361 | 1 | 1 | 2.50E-05 | Foregut |
| Gprc5c | 2.21E-09 | 0.723932434 | 1 | 0.985 | 2.78E-05 | Foregut |
| Slc16a1 | 2.39E-09 | 1.241839315 | 1 | 1 | 3.01E-05 | Foregut |
| Tmsb10 | 3.02E-09 | 0.860153214 | 1 | 1 | 3.80E-05 | Foregut |
| Frmd4b | 3.12E-09 | 0.250442933 | 0.857 | 0.938 | 3.92E-05 | Foregut |
| Rbp1 | 3.45E-09 | 0.783137436 | 1 | 0.923 | 4.34E-05 | Foregut |
| Ctxn1 | 3.74E-09 | 0.77146138 | 1 | 0.892 | 4.70E-05 | Foregut |
| Dkk3 | 3.77E-09 | 0.556313934 | 0.929 | 0.492 | 4.74E-05 | Foregut |
| Krt19 | 3.83E-09 | 1.053323678 | 1 | 0.8 | 4.81E-05 | Foregut |
| Cdc42ep1 | 4.07E-09 | 0.55120445 | 1 | 0.708 | 5.12E-05 | Foregut |
| Podxl | 6.50E-09 | 0.84238999 | 1 | 1 | 8.18E-05 | Foregut |
| Cdh1 | 7.02E-09 | 0.863359311 | 1 | 0.862 | 8.83E-05 | Foregut |
| Gata3 | 8.46E-09 | 0.363738104 | 0.929 | 0.523 | 0.000106 | Foregut |
| Crip2 | 9.37E-09 | 0.750387006 | 1 | 0.923 | 0.000118 | Foregut |
| Marveld2 | 1.08E-08 | 0.299781278 | 0.857 | 0.369 | 0.000135 | Foregut |
| Cdh3 | 1.38E-08 | 0.573004381 | 0.929 | 0.708 | 0.000173 | Foregut |
| Csrp2 | 1.54E-08 | 1.176478532 | 1 | 1 | 0.000194 | Foregut |
| Prss8 | 1.55E-08 | 0.99099104 | 1 | 0.738 | 0.000194 | Foregut |
| Smoc2 | 2.25E-08 | 0.433752691 | 0.571 | 0.369 | 0.000283 | Foregut |
| Tmem254c | 2.59E-08 | 0.425188833 | 1 | 0.892 | 0.000325 | Foregut |
| Tspan9 | 2.74E-08 | 0.417951971 | 0.929 | 0.738 | 0.000345 | Foregut |
| Mboat1 | 3.39E-08 | 0.394484319 | 0.857 | 0.554 | 0.000426 | Foregut |
| Ube2h | 3.44E-08 | 0.404220455 | 1 | 0.908 | 0.000433 | Foregut |
| Tmem254a | 3.53E-08 | 0.427413678 | 1 | 0.877 | 0.000444 | Foregut |
| Tmem254b | 4.05E-08 | 0.415931532 | 1 | 0.954 | 0.000509 | Foregut |
| Igfbp2 | 5.11E-08 | 1.084725353 | 1 | 1 | 0.000642 | Foregut |
| 1700047I1 | 5.70E-08 | 0.313848947 | 0.929 | 0.677 | 0.000717 | Foregut |

| | | | | | | |
|---|---|---|---|---|---|---|
| Raet1d | 5.89E-08 | 0.300144908 | 0.929 | 0.462 | 0.000741 | Foregut |
| Serpinb6c | 6.22E-08 | 0.280352711 | 0.929 | 0.631 | 0.000782 | Foregut |
| Krt20 | 6.51E-08 | 0.437442139 | 0.714 | 0.092 | 0.000819 | Foregut |
| Vnn1 | 7.33E-08 | 0.469777428 | 0.857 | 0.169 | 0.000922 | Foregut |
| Lsr | 7.89E-08 | 0.459083838 | 1 | 0.969 | 0.000992 | Foregut |
| Cox4i2 | 9.61E-08 | 0.68753126 | 0.786 | 0.338 | 0.001209 | Foregut |
| Fam107b | 1.46E-07 | 0.489543714 | 1 | 0.892 | 0.001842 | Foregut |
| Habp2 | 1.69E-07 | 0.480489 | 0.643 | 0.462 | 0.002131 | Foregut |
| Tgfb2 | 1.86E-07 | 0.652573402 | 1 | 0.677 | 0.002339 | Foregut |
| Ddah2 | 1.95E-07 | 0.792216175 | 1 | 0.985 | 0.002453 | Foregut |
| Kctd12b | 2.26E-07 | 0.253906718 | 0.643 | 0.508 | 0.002838 | Foregut |
| 0610040J0 | 3.20E-07 | 0.359422301 | 0.929 | 0.4 | 0.004028 | Foregut |
| Sfrp5 | 3.72E-07 | 1.392350035 | 0.643 | 0.523 | 0.004675 | Foregut |
| Tuft1 | 3.85E-07 | 0.333077095 | 0.571 | 0.369 | 0.004839 | Foregut |
| Actb | 4.71E-07 | 0.317964522 | 1 | 1 | 0.00592 | Foregut |
| Calcoco1 | 5.06E-07 | 0.263429885 | 0.929 | 0.815 | 0.006361 | Foregut |
| Car4 | 5.11E-07 | 2.204875225 | 1 | 1 | 0.006427 | Foregut |
| Nudt11 | 5.41E-07 | 0.396667246 | 0.929 | 0.723 | 0.00681 | Foregut |
| Bex4 | 5.50E-07 | 1.068334526 | 1 | 1 | 0.006921 | Foregut |
| Crb3 | 5.78E-07 | 0.625961718 | 1 | 0.615 | 0.007265 | Foregut |
| Cpxm1 | 6.09E-07 | 0.39140601 | 0.929 | 0.738 | 0.007664 | Foregut |
| Ctnna1 | 8.97E-07 | 0.561092557 | 1 | 1 | 0.011279 | Foregut |
| Bsg | 9.08E-07 | 0.833465527 | 1 | 1 | 0.011414 | Foregut |
| Gpx2 | 9.42E-07 | 0.407348089 | 0.786 | 0.154 | 0.011848 | Foregut |
| Emb | 9.63E-07 | 1.469137552 | 1 | 1 | 0.012107 | Foregut |
| S100a11 | 1.10E-06 | 1.001447815 | 1 | 0.862 | 0.013895 | Foregut |
| Cd276 | 1.34E-06 | 0.448277221 | 1 | 0.846 | 0.016843 | Foregut |
| Samd10 | 1.47E-06 | 0.304061261 | 1 | 0.431 | 0.018529 | Foregut |
| Fam210b | 1.55E-06 | 0.299249208 | 0.786 | 0.554 | 0.019434 | Foregut |
| Hkdc1 | 1.57E-06 | 0.300247548 | 0.857 | 0.677 | 0.019776 | Foregut |
| Tmem9 | 1.71E-06 | 0.437524339 | 1 | 0.969 | 0.021461 | Foregut |
| Cers5 | 2.06E-06 | 0.278906427 | 0.929 | 1 | 0.025959 | Foregut |
| Casp8 | 2.14E-06 | 0.270732911 | 0.786 | 0.754 | 0.026889 | Foregut |
| S100a10 | 2.37E-06 | 0.895057818 | 1 | 0.908 | 0.029753 | Foregut |
| Hacd2 | 2.48E-06 | 0.360419359 | 1 | 0.969 | 0.031196 | Foregut |
| Trap1a | 2.52E-06 | 1.496613036 | 0.429 | 0.508 | 0.031667 | Foregut |
| Cnn3 | 2.90E-06 | 0.482006033 | 1 | 0.985 | 0.036492 | Foregut |
| Frzb | 3.02E-06 | 0.513122022 | 0.929 | 0.723 | 0.037981 | Foregut |
| App | 3.18E-06 | 0.675249152 | 1 | 0.969 | 0.039959 | Foregut |
| Igsf21 | 3.23E-06 | 0.28496613 | 0.714 | 0.185 | 0.040563 | Foregut |
| Nucb1 | 3.34E-06 | 0.32865695 | 1 | 0.985 | 0.041948 | Foregut |
| Slc38a4 | 3.85E-06 | 0.491870105 | 1 | 0.908 | 0.048365 | Foregut |
| Dpp4 | 4.20E-06 | 0.391013668 | 0.786 | 0.877 | 0.052823 | Foregut |
| Kdelr2 | 4.38E-06 | 0.494705771 | 1 | 1 | 0.055056 | Foregut |
| Myl6 | 4.49E-06 | 0.705022775 | 1 | 1 | 0.056477 | Foregut |
| Erlin2 | 5.25E-06 | 0.434162339 | 1 | 0.938 | 0.066058 | Foregut |
| Mdk | 5.73E-06 | 0.489154134 | 1 | 1 | 0.072056 | Foregut |
| Tax1bp3 | 6.07E-06 | 0.437362319 | 1 | 1 | 0.076313 | Foregut |
| Pygl | 6.96E-06 | 0.331495333 | 1 | 0.738 | 0.087565 | Foregut |
| Pdpn | 6.98E-06 | 1.039940491 | 0.929 | 1 | 0.08772 | Foregut |
| Krtcap3 | 8.06E-06 | 0.483386053 | 0.857 | 0.431 | 0.101317 | Foregut |
| Cgn | 8.12E-06 | 0.513866124 | 1 | 1 | 0.10211 | Foregut |
| Gpx8 | 8.62E-06 | 0.390679666 | 1 | 0.615 | 0.108413 | Foregut |
| Cd24a | 9.16E-06 | 0.543691937 | 1 | 1 | 0.115137 | Foregut |
| Carhsp1 | 9.49E-06 | 0.534620959 | 1 | 1 | 0.119389 | Foregut |
| Laptm4a | 1.01E-05 | 0.666878297 | 1 | 0.985 | 0.126503 | Foregut |
| Gbas | 1.08E-05 | 0.266449304 | 1 | 0.969 | 0.135828 | Foregut |
| Pgrmc1 | 1.30E-05 | 0.616867468 | 1 | 1 | 0.163514 | Foregut |

| | | | | | | |
|---|---|---|---|---|---|---|
| Sdcbp | 1.31E-05 | 0.330893638 | 1 | 1 | 0.164608 | Foregut |
| Npc2 | 1.32E-05 | 0.471109409 | 1 | 1 | 0.165603 | Foregut |
| Anxa6 | 1.56E-05 | 0.56319929 | 1 | 0.8 | 0.196458 | Foregut |
| Pigs | 1.73E-05 | 0.443187786 | 1 | 1 | 0.217223 | Foregut |
| Dstn | 1.82E-05 | 0.514129482 | 1 | 1 | 0.228282 | Foregut |
| Fam3c | 1.84E-05 | 0.292463625 | 1 | 0.923 | 0.231481 | Foregut |
| Fam213a | 1.91E-05 | 0.447699572 | 1 | 0.938 | 0.240775 | Foregut |
| S1pr2 | 2.01E-05 | 0.402850062 | 0.929 | 0.846 | 0.253028 | Foregut |
| Afp | 2.12E-05 | 1.567828045 | 0.929 | 0.738 | 0.266712 | Foregut |
| Plp2 | 2.32E-05 | 0.408166002 | 1 | 0.938 | 0.291281 | Foregut |
| Clic1 | 2.37E-05 | 0.50398053 | 1 | 1 | 0.298351 | Foregut |
| Rnase4 | 2.51E-05 | 0.476433059 | 0.643 | 0.262 | 0.315135 | Foregut |
| 9530068E( | 2.57E-05 | 0.468525033 | 1 | 0.954 | 0.323252 | Foregut |
| Fam174b | 2.61E-05 | 0.464713176 | 0.929 | 0.585 | 0.327647 | Foregut |
| Pdia3 | 2.76E-05 | 0.518626883 | 1 | 1 | 0.347148 | Foregut |
| Prmt2 | 3.01E-05 | 0.308202686 | 0.929 | 0.815 | 0.378948 | Foregut |
| Smim22 | 3.14E-05 | 0.358232113 | 0.929 | 0.262 | 0.394303 | Foregut |
| Mapre2 | 3.25E-05 | 0.258041113 | 1 | 0.969 | 0.408254 | Foregut |
| Tmco3 | 3.36E-05 | 0.257736043 | 1 | 1 | 0.422721 | Foregut |
| Ap1m2 | 3.49E-05 | 0.567872944 | 0.929 | 0.538 | 0.438936 | Foregut |
| Dap | 3.50E-05 | 0.463075233 | 1 | 0.831 | 0.440035 | Foregut |
| Gm6169 | 3.51E-05 | 0.361484412 | 1 | 0.938 | 0.441903 | Foregut |
| Tmed2 | 3.65E-05 | 0.478112741 | 1 | 1 | 0.459479 | Foregut |
| Ctsd | 3.65E-05 | 0.544172298 | 1 | 1 | 0.459588 | Foregut |
| Wbp1 | 3.82E-05 | 0.291806819 | 1 | 0.892 | 0.479902 | Foregut |
| Csrp1 | 3.82E-05 | 0.562294093 | 1 | 0.846 | 0.480011 | Foregut |
| Atpif1 | 3.98E-05 | 0.443235761 | 1 | 1 | 0.500267 | Foregut |
| Mpzl1 | 4.37E-05 | 0.53948105 | 1 | 0.985 | 0.549704 | Foregut |
| Serinc2 | 4.40E-05 | 0.330734509 | 0.929 | 0.385 | 0.552785 | Foregut |
| Pttg1ip | 4.62E-05 | 0.459336831 | 1 | 0.954 | 0.580438 | Foregut |
| Pld3 | 4.66E-05 | 0.476310341 | 0.929 | 0.877 | 0.586321 | Foregut |
| Cystm1 | 4.72E-05 | 0.254332932 | 0.929 | 0.585 | 0.593674 | Foregut |
| Ddost | 4.74E-05 | 0.387835152 | 1 | 1 | 0.596097 | Foregut |
| Ywhaz | 4.75E-05 | 0.309211234 | 1 | 1 | 0.596879 | Foregut |
| Ywhaq | 5.56E-05 | 0.363727522 | 1 | 1 | 0.698769 | Foregut |
| Arg1 | 5.63E-05 | 1.120719366 | 0.929 | 0.677 | 0.707895 | Foregut |
| Ppib | 5.64E-05 | 0.430233127 | 1 | 1 | 0.709708 | Foregut |
| Lasp1 | 5.72E-05 | 0.417665757 | 1 | 0.954 | 0.719086 | Foregut |
| P4hb | 5.89E-05 | 0.486342417 | 1 | 1 | 0.740176 | Foregut |
| Calr | 6.10E-05 | 0.566798279 | 1 | 1 | 0.767162 | Foregut |
| B4galt1 | 6.17E-05 | 0.34561537 | 1 | 0.954 | 0.776227 | Foregut |
| Aktip | 6.20E-05 | 0.268559875 | 1 | 0.877 | 0.779664 | Foregut |
| Tmem54 | 6.85E-05 | 0.385936347 | 0.643 | 0.138 | 0.861227 | Foregut |
| Igsf9 | 6.98E-05 | 0.310461597 | 1 | 0.892 | 0.877415 | Foregut |
| Hsp90b1 | 7.02E-05 | 0.577974483 | 1 | 1 | 0.882521 | Foregut |
| Suox | 7.33E-05 | 0.284929719 | 0.929 | 0.569 | 0.921779 | Foregut |
| Grn | 8.24E-05 | 0.344687574 | 1 | 0.969 | 1 | Foregut |
| Gdpd1 | 8.91E-05 | 0.349580848 | 1 | 0.646 | 1 | Foregut |
| Ocln | 9.69E-05 | 0.256831871 | 0.929 | 0.492 | 1 | Foregut |
| Tmprss2 | 9.89E-05 | 0.871327467 | 0.929 | 0.708 | 1 | Foregut |
| Pdlim1 | 9.99E-05 | 0.324663785 | 1 | 0.738 | 1 | Foregut |
| Laptm4b | 0.000103 | 0.456125167 | 1 | 0.985 | 1 | Foregut |
| Tbx3 | 0.000103 | 0.374455489 | 0.857 | 0.508 | 1 | Foregut |
| Ndfip1 | 0.000114 | 0.449899832 | 1 | 1 | 1 | Foregut |
| Gadd45b | 0.000128 | 0.288425375 | 0.714 | 0.185 | 1 | Foregut |
| Spink3 | 0.00013 | 1.232120186 | 1 | 0.923 | 1 | Foregut |
| Tmed9 | 0.00013 | 0.553002834 | 1 | 0.923 | 1 | Foregut |
| Rnf128 | 0.000133 | 0.444219331 | 0.929 | 0.785 | 1 | Foregut |

| | | | | | | |
|---|---|---|---|---|---|---|
| Gng11 | 0.000146 | 0.37823713 | 0.714 | 0.308 | 1 | Foregut |
| Klk8 | 0.000165 | 0.359072008 | 0.786 | 0.385 | 1 | Foregut |
| M6pr | 0.000178 | 0.385585601 | 1 | 1 | 1 | Foregut |
| Clu | 0.000181 | 0.381408146 | 1 | 0.615 | 1 | Foregut |
| Cst3 | 0.000185 | 0.774477862 | 1 | 1 | 1 | Foregut |
| Pacsin2 | 0.000203 | 0.38501491 | 1 | 0.938 | 1 | Foregut |
| Atp6v0e | 0.000203 | 0.293645217 | 1 | 1 | 1 | Foregut |
| Tma7 | 0.000204 | 0.279194571 | 1 | 1 | 1 | Foregut |
| Fam129b | 0.000224 | 0.294495066 | 1 | 0.708 | 1 | Foregut |
| Slc31a1 | 0.000228 | 0.458721505 | 1 | 0.954 | 1 | Foregut |
| Cd63 | 0.000234 | 0.658429684 | 1 | 0.985 | 1 | Foregut |
| Arf4 | 0.000245 | 0.282381248 | 1 | 1 | 1 | Foregut |
| Ttr | 0.000251 | 1.504483615 | 1 | 0.923 | 1 | Foregut |
| Serf1 | 0.000265 | 0.25678776 | 1 | 0.8 | 1 | Foregut |
| Tmed4 | 0.000276 | 0.516717862 | 1 | 0.938 | 1 | Foregut |
| Id2 | 0.000288 | 0.73749884 | 1 | 1 | 1 | Foregut |
| Rbpms | 0.000309 | 0.324668305 | 1 | 0.985 | 1 | Foregut |
| Tceal8 | 0.00032 | 0.568420554 | 1 | 1 | 1 | Foregut |
| Cnn2 | 0.000327 | 0.653955077 | 1 | 0.969 | 1 | Foregut |
| Rcn1 | 0.000344 | 0.609087525 | 1 | 0.985 | 1 | Foregut |
| Tbca | 0.000358 | 0.475312446 | 1 | 0.985 | 1 | Foregut |
| Txn1 | 0.000359 | 0.26234686 | 1 | 1 | 1 | Foregut |
| Itgb1 | 0.000359 | 0.425734846 | 1 | 1 | 1 | Foregut |
| Nudt14 | 0.000381 | 0.324057385 | 1 | 0.846 | 1 | Foregut |
| Hmgn1 | 0.000383 | 0.315528167 | 1 | 1 | 1 | Foregut |
| Tmem88 | 0.000392 | 0.504581075 | 0.929 | 0.754 | 1 | Foregut |
| Myl12b | 0.000422 | 0.434088747 | 1 | 1 | 1 | Foregut |
| Dhrs3 | 0.000426 | 0.371412871 | 0.857 | 0.492 | 1 | Foregut |
| Chrnb1 | 0.000434 | 0.260875633 | 0.643 | 0.215 | 1 | Foregut |
| Surf4 | 0.000447 | 0.417022287 | 1 | 1 | 1 | Foregut |
| Tpm3-rs7 | 0.000453 | 0.286841484 | 1 | 1 | 1 | Foregut |
| Slc9a3r2 | 0.00046 | 0.266730577 | 0.786 | 0.446 | 1 | Foregut |
| Mum1l1 | 0.000479 | 0.276434339 | 1 | 0.831 | 1 | Foregut |
| Cdc37 | 0.000507 | 0.299518582 | 1 | 1 | 1 | Foregut |
| Slc2a1 | 0.000514 | 0.31773734 | 1 | 1 | 1 | Foregut |
| Itm2b | 0.000542 | 0.577449515 | 1 | 0.969 | 1 | Foregut |
| Rcn2 | 0.000552 | 0.347998712 | 1 | 0.985 | 1 | Foregut |
| Myl12a | 0.000575 | 0.304787228 | 1 | 1 | 1 | Foregut |
| Tmed10 | 0.000588 | 0.494444995 | 1 | 1 | 1 | Foregut |
| Slc39a8 | 0.000589 | 0.263559502 | 1 | 0.877 | 1 | Foregut |
| Bambi | 0.00062 | 0.269336956 | 0.857 | 0.692 | 1 | Foregut |
| Ssr3 | 0.000655 | 0.418172776 | 1 | 1 | 1 | Foregut |
| Pgm2 | 0.000677 | 0.279229358 | 1 | 0.892 | 1 | Foregut |
| Ndufa4 | 0.000687 | 0.327841175 | 1 | 1 | 1 | Foregut |
| Cdkn1c | 0.000692 | 0.615721002 | 1 | 0.723 | 1 | Foregut |
| Tpm3 | 0.000702 | 0.278906933 | 1 | 1 | 1 | Foregut |
| Cldn11 | 0.000711 | 0.356499107 | 0.643 | 0.308 | 1 | Foregut |
| Cdo1 | 0.000718 | 0.325335572 | 0.714 | 0.2 | 1 | Foregut |
| Sod1 | 0.000744 | 0.432482844 | 1 | 1 | 1 | Foregut |
| Klf10 | 0.000747 | 0.276906717 | 0.857 | 0.815 | 1 | Foregut |
| Tpm4 | 0.000778 | 0.414026995 | 1 | 1 | 1 | Foregut |
| Rbbp7 | 0.000785 | 0.575820317 | 1 | 1 | 1 | Foregut |
| Hexb | 0.00083 | 0.320030193 | 0.929 | 0.692 | 1 | Foregut |
| Paip1 | 0.000886 | 0.30172183 | 1 | 1 | 1 | Foregut |
| Lman2 | 0.000927 | 0.339001259 | 1 | 1 | 1 | Foregut |
| Atox1 | 0.000942 | 0.458212947 | 1 | 0.954 | 1 | Foregut |
| Bcap31 | 0.000954 | 0.673713084 | 1 | 1 | 1 | Foregut |
| Cyba | 0.001028 | 0.447977798 | 1 | 0.985 | 1 | Foregut |

| | | | | | | |
|---|---|---|---|---|---|---|
| Pdia6 | 0.001028 | 0.437925828 | 1 | 1 | 1 | Foregut |
| Serpinf2 | 0.001066 | 0.262866358 | 0.714 | 0.185 | 1 | Foregut |
| B4galnt2 | 0.001125 | 0.367423228 | 0.714 | 0.231 | 1 | Foregut |
| Atxn10 | 0.001221 | 0.261239331 | 1 | 0.985 | 1 | Foregut |
| Bnip3 | 0.001247 | 0.777813844 | 1 | 0.985 | 1 | Foregut |
| Dbn1 | 0.001325 | 0.38697617 | 1 | 1 | 1 | Foregut |
| Hspa5 | 0.001386 | 0.40391051 | 1 | 1 | 1 | Foregut |
| Copz1 | 0.001441 | 0.350629106 | 1 | 1 | 1 | Foregut |
| Nucb2 | 0.001447 | 0.352876498 | 0.857 | 0.677 | 1 | Foregut |
| Nudt16l1 | 0.001497 | 0.313058525 | 1 | 0.969 | 1 | Foregut |
| Cep19 | 0.001749 | 0.266265408 | 1 | 0.769 | 1 | Foregut |
| Prdx6 | 0.001755 | 0.449021363 | 1 | 1 | 1 | Foregut |
| Oaz2 | 0.001877 | 0.398506622 | 1 | 0.985 | 1 | Foregut |
| Clic6 | 0.002087 | 0.407298277 | 0.857 | 0.615 | 1 | Foregut |
| Ssr2 | 0.002087 | 0.397367049 | 1 | 1 | 1 | Foregut |
| Spg21 | 0.002104 | 0.314440677 | 1 | 0.862 | 1 | Foregut |
| Cttn | 0.00215 | 0.368123411 | 1 | 1 | 1 | Foregut |
| Rp9 | 0.002172 | 0.295735712 | 1 | 0.831 | 1 | Foregut |
| Rbbp9 | 0.002398 | 0.3135955 | 1 | 0.877 | 1 | Foregut |
| Spcs1 | 0.00268 | 0.252018568 | 1 | 1 | 1 | Foregut |
| Etv5 | 0.002693 | 0.31118169 | 1 | 0.862 | 1 | Foregut |
| Arpc5 | 0.002697 | 0.297025349 | 1 | 0.985 | 1 | Foregut |
| Fscn1 | 0.0027 | 0.328768235 | 1 | 1 | 1 | Foregut |
| Tmem50a | 0.002828 | 0.345997862 | 1 | 0.985 | 1 | Foregut |
| 1110012L1 | 0.002892 | 0.263768574 | 1 | 0.708 | 1 | Foregut |
| Rpn1 | 0.002939 | 0.370060153 | 1 | 1 | 1 | Foregut |
| Reep5 | 0.002996 | 0.408635177 | 1 | 0.923 | 1 | Foregut |
| Phlda2 | 0.00301 | 0.627278387 | 0.929 | 0.462 | 1 | Foregut |
| Slc39a4 | 0.003131 | 0.552586981 | 0.929 | 0.662 | 1 | Foregut |
| Hyal2 | 0.003381 | 0.292092819 | 1 | 0.969 | 1 | Foregut |
| Tmem59 | 0.003392 | 0.444735323 | 1 | 1 | 1 | Foregut |
| Vps36 | 0.003625 | 0.25760974 | 1 | 0.954 | 1 | Foregut |
| Akr1a1 | 0.003625 | 0.347053678 | 1 | 1 | 1 | Foregut |
| Mcfd2 | 0.003665 | 0.285115028 | 1 | 0.985 | 1 | Foregut |
| Rasl11a | 0.003779 | 0.31581533 | 0.857 | 0.677 | 1 | Foregut |
| Maged1 | 0.003831 | 0.353971149 | 1 | 1 | 1 | Foregut |
| 5730507C( | 0.004137 | 0.374041325 | 1 | 0.985 | 1 | Foregut |
| Papss1 | 0.004182 | 0.306277825 | 1 | 0.985 | 1 | Foregut |
| Cops5 | 0.004234 | 0.258682895 | 1 | 1 | 1 | Foregut |
| Prelid2 | 0.004371 | 0.448122877 | 0.929 | 0.631 | 1 | Foregut |
| Slc25a4 | 0.004525 | 0.293822941 | 1 | 1 | 1 | Foregut |
| Twsg1 | 0.004534 | 0.319540337 | 1 | 0.969 | 1 | Foregut |
| Slc44a2 | 0.004697 | 0.282697109 | 0.929 | 0.831 | 1 | Foregut |
| Nrbp1 | 0.004756 | 0.270027561 | 1 | 0.985 | 1 | Foregut |
| Myadm | 0.004777 | 0.312846176 | 1 | 0.8 | 1 | Foregut |
| Cd81 | 0.004881 | 0.325471969 | 1 | 1 | 1 | Foregut |
| Arpc2 | 0.004897 | 0.291873298 | 1 | 1 | 1 | Foregut |
| Nradd | 0.004909 | 0.311126507 | 0.929 | 0.738 | 1 | Foregut |
| Enpp3 | 0.005012 | 0.252919049 | 0.857 | 0.4 | 1 | Foregut |
| Sdc1 | 0.005256 | 0.414938936 | 1 | 0.908 | 1 | Foregut |
| Hif1a | 0.005332 | 0.278079897 | 1 | 0.985 | 1 | Foregut |
| Zfand6 | 0.005429 | 0.314150969 | 1 | 0.985 | 1 | Foregut |
| Stmn1 | 0.005495 | 0.521512509 | 1 | 1 | 1 | Foregut |
| Chmp2b | 0.00559 | 0.351555565 | 1 | 0.938 | 1 | Foregut |
| Tmub2 | 0.005875 | 0.289844424 | 1 | 0.769 | 1 | Foregut |
| Tubb4a | 0.005892 | 0.5011792 | 1 | 0.985 | 1 | Foregut |
| Lman1 | 0.006035 | 0.269323339 | 1 | 0.985 | 1 | Foregut |
| Snx6 | 0.006087 | 0.382638815 | 1 | 1 | 1 | Foregut |

| | | | | | | |
|---|---|---|---|---|---|---|
| Cstb | 0.006262 | 0.34513381 | 1 | 0.969 | 1 | Foregut |
| Myl9 | 0.006321 | 0.271335396 | 1 | 0.692 | 1 | Foregut |
| F2r | 0.006811 | 0.363836358 | 1 | 0.954 | 1 | Foregut |
| Smpd1 | 0.006944 | 0.263327669 | 1 | 0.815 | 1 | Foregut |
| Smtnl2 | 0.007123 | 0.257044093 | 1 | 1 | 1 | Foregut |
| Cnpy3 | 0.007162 | 0.264642449 | 1 | 1 | 1 | Foregut |
| Sh3bgrl3 | 0.00717 | 0.294724368 | 0.929 | 0.923 | 1 | Foregut |
| Rbp4 | 0.007283 | 0.867320211 | 1 | 0.846 | 1 | Foregut |
| Ifi30 | 0.007403 | 0.444368938 | 1 | 0.769 | 1 | Foregut |
| Anxa5 | 0.007494 | 0.412875825 | 1 | 0.831 | 1 | Foregut |
| Cd164 | 0.007618 | 0.326739526 | 1 | 1 | 1 | Foregut |
| Dazap2 | 0.00772 | 0.340066939 | 1 | 0.985 | 1 | Foregut |
| Stx7 | 0.008369 | 0.472602574 | 0.929 | 0.923 | 1 | Foregut |
| Tmbim6 | 0.008523 | 0.361203159 | 1 | 1 | 1 | Foregut |
| Txndc12 | 0.008694 | 0.356301127 | 1 | 0.985 | 1 | Foregut |
| C1galt1c1 | 0.008998 | 0.296690014 | 0.786 | 0.615 | 1 | Foregut |
| Plat | 0.009554 | 0.42351575 | 1 | 0.754 | 1 | Foregut |
| Zfp125 | 0.009749 | 0.378070138 | 1 | 0.954 | 1 | Foregut |
| Slit2 | 8.91E-31 | 0.322328311 | 1 | 0.908 | 1.12E-26 | Notochord |
| Lmx1a | 1.28E-19 | 1.184135831 | 1 | 0.329 | 1.61E-15 | Notochord |
| Tmem255a | 2.65E-18 | 0.97558456 | 1 | 0.224 | 3.34E-14 | Notochord |
| Aplnr | 8.64E-18 | 2.490989173 | 1 | 0.816 | 1.09E-13 | Notochord |
| Col9a1 | 5.40E-17 | 1.372743388 | 1 | 0.289 | 6.79E-13 | Notochord |
| Htra1 | 8.85E-16 | 1.453941019 | 1 | 0.342 | 1.11E-11 | Notochord |
| Car3 | 3.08E-14 | 1.470281011 | 1 | 0.763 | 3.88E-10 | Notochord |
| Tgm2 | 3.11E-14 | 1.622440903 | 1 | 0.737 | 3.91E-10 | Notochord |
| Shh | 8.33E-14 | 1.910652405 | 1 | 0.461 | 1.05E-09 | Notochord |
| B3gnt7 | 9.24E-14 | 1.094926079 | 1 | 0.737 | 1.16E-09 | Notochord |
| Gng4 | 1.37E-13 | 0.563778893 | 1 | 0.145 | 1.72E-09 | Notochord |
| Cthrc1 | 9.43E-13 | 1.917733163 | 1 | 0.763 | 1.19E-08 | Notochord |
| Arhgef16 | 3.17E-12 | 0.725232717 | 1 | 0.461 | 3.99E-08 | Notochord |
| Samd3 | 3.60E-12 | 1.493004284 | 1 | 0.434 | 4.53E-08 | Notochord |
| Kirrel3 | 9.16E-12 | 0.311590927 | 1 | 0.25 | 1.15E-07 | Notochord |
| Nlgn3 | 9.39E-12 | 0.26426431 | 1 | 0.737 | 1.18E-07 | Notochord |
| Gnai1 | 7.73E-11 | 0.502108283 | 1 | 0.579 | 9.72E-07 | Notochord |
| Wwp2 | 9.71E-11 | 1.485238919 | 1 | 0.947 | 1.22E-06 | Notochord |
| Tmem100 | 1.23E-10 | 1.251635635 | 1 | 0.237 | 1.55E-06 | Notochord |
| Alcam | 1.24E-10 | 0.314480337 | 1 | 0.329 | 1.56E-06 | Notochord |
| Prickle1 | 1.34E-10 | 1.083628501 | 1 | 0.908 | 1.68E-06 | Notochord |
| Sept6 | 1.61E-10 | 0.590503487 | 1 | 0.921 | 2.02E-06 | Notochord |
| Moxd1 | 5.18E-10 | 0.793743182 | 1 | 0.237 | 6.51E-06 | Notochord |
| Susd5 | 1.50E-09 | 0.666138489 | 1 | 0.224 | 1.89E-05 | Notochord |
| Vgll4 | 6.25E-09 | 0.499602032 | 1 | 0.803 | 7.86E-05 | Notochord |
| Krt19 | 8.42E-09 | 2.367221124 | 1 | 0.829 | 0.000106 | Notochord |
| Sox9 | 8.58E-09 | 1.799031061 | 1 | 0.895 | 0.000108 | Notochord |
| Rnd3 | 1.14E-08 | 1.412534236 | 1 | 1 | 0.000143 | Notochord |
| Egflam | 1.37E-08 | 0.432403274 | 1 | 0.526 | 0.000172 | Notochord |
| Rab38 | 1.84E-08 | 0.934952665 | 1 | 0.395 | 0.000231 | Notochord |
| Sema3d | 2.06E-08 | 0.290299735 | 1 | 0.289 | 0.000259 | Notochord |
| Cd36 | 3.64E-08 | 1.082656182 | 1 | 0.329 | 0.000458 | Notochord |
| St6galnac3 | 4.99E-08 | 0.374449169 | 1 | 0.658 | 0.000628 | Notochord |
| Tmod2 | 5.17E-08 | 0.310230873 | 1 | 0.592 | 0.000651 | Notochord |
| Mturn | 5.97E-08 | 0.294772867 | 0.667 | 0.342 | 0.000751 | Notochord |
| Plekhb1 | 6.40E-08 | 0.756189013 | 1 | 0.303 | 0.000805 | Notochord |
| Nkd1 | 1.02E-07 | 0.814470199 | 1 | 0.776 | 0.001284 | Notochord |
| Lypd6 | 1.04E-07 | 0.316480259 | 0.667 | 0.368 | 0.00131 | Notochord |
| Sulf1 | 1.18E-07 | 0.876541549 | 1 | 0.934 | 0.00148 | Notochord |
| Gpm6b | 2.13E-07 | 0.436994308 | 1 | 0.526 | 0.002684 | Notochord |

| | | | | | | |
|---|---|---|---|---|---|---|
| Sox6 | 2.47E-07 | 0.251866823 | 1 | 0.382 | 0.003101 | Notochord |
| T | 3.27E-07 | 1.933115998 | 1 | 0.566 | 0.004118 | Notochord |
| Arid3b | 3.75E-07 | 0.952664076 | 1 | 1 | 0.004716 | Notochord |
| Epb4.1l3 | 3.87E-07 | 1.388827056 | 1 | 0.934 | 0.004864 | Notochord |
| Col5a1 | 4.15E-07 | 0.488960126 | 1 | 0.789 | 0.005214 | Notochord |
| Anxa3 | 4.85E-07 | 0.358821422 | 1 | 0.224 | 0.006101 | Notochord |
| Scara3 | 5.11E-07 | 0.946548091 | 1 | 0.474 | 0.006422 | Notochord |
| Nap1l5 | 6.30E-07 | 0.802138767 | 1 | 0.276 | 0.007927 | Notochord |
| Myl9 | 6.34E-07 | 1.279094694 | 1 | 0.737 | 0.007971 | Notochord |
| Isg20 | 8.96E-07 | 0.526071682 | 1 | 0.145 | 0.011267 | Notochord |
| Tpm2 | 1.11E-06 | 0.865731462 | 1 | 0.816 | 0.013981 | Notochord |
| Gstm5 | 1.69E-06 | 1.02066552 | 1 | 0.921 | 0.02123 | Notochord |
| Usp18 | 1.71E-06 | 0.648124398 | 1 | 0.158 | 0.021503 | Notochord |
| Smox | 1.76E-06 | 0.367235655 | 1 | 0.5 | 0.022161 | Notochord |
| Clu | 1.95E-06 | 1.152723835 | 1 | 0.671 | 0.024546 | Notochord |
| Mnx1 | 1.98E-06 | 0.298942388 | 1 | 0.224 | 0.024879 | Notochord |
| Psme2b | 3.74E-06 | 0.426443554 | 1 | 0.882 | 0.046995 | Notochord |
| Angptl2 | 3.81E-06 | 0.535776293 | 1 | 0.658 | 0.047873 | Notochord |
| Gcnt1 | 4.31E-06 | 0.625265694 | 1 | 0.789 | 0.054194 | Notochord |
| Vtn | 4.79E-06 | 1.171259608 | 1 | 0.566 | 0.060213 | Notochord |
| Cnn2 | 5.16E-06 | 1.823740699 | 1 | 0.974 | 0.064886 | Notochord |
| Lrig1 | 7.17E-06 | 0.436624866 | 1 | 0.842 | 0.090131 | Notochord |
| Mmd | 9.14E-06 | 0.544931814 | 1 | 0.882 | 0.114895 | Notochord |
| Gpt2 | 1.15E-05 | 0.322236425 | 1 | 0.855 | 0.145173 | Notochord |
| Col2a1 | 1.17E-05 | 0.85914495 | 1 | 0.803 | 0.147325 | Notochord |
| Guca1a | 1.34E-05 | 0.406914599 | 1 | 0.079 | 0.169037 | Notochord |
| Tapbp | 1.87E-05 | 0.560160709 | 1 | 0.934 | 0.235442 | Notochord |
| Rarg | 2.10E-05 | 0.309223679 | 1 | 0.553 | 0.26448 | Notochord |
| Ap1s2 | 2.30E-05 | 0.671078079 | 1 | 0.711 | 0.289526 | Notochord |
| Leprot | 3.13E-05 | 0.808659052 | 1 | 0.882 | 0.393061 | Notochord |
| Kank1 | 3.29E-05 | 0.686616442 | 1 | 0.934 | 0.414107 | Notochord |
| Gulp1 | 4.08E-05 | 0.577566911 | 1 | 0.632 | 0.512497 | Notochord |
| Krt8 | 4.37E-05 | 1.602999044 | 1 | 1 | 0.549412 | Notochord |
| Irx3 | 4.39E-05 | 0.445691866 | 1 | 0.513 | 0.552343 | Notochord |
| Dffa | 5.22E-05 | 0.352552679 | 1 | 0.974 | 0.656699 | Notochord |
| Ugdh | 6.76E-05 | 0.569322577 | 1 | 0.974 | 0.850071 | Notochord |
| Nid2 | 7.59E-05 | 0.795815478 | 1 | 0.987 | 0.954567 | Notochord |
| Mfap4 | 9.67E-05 | 0.433000082 | 1 | 0.395 | 1 | Notochord |
| Pctp | 0.000143 | 0.574187895 | 1 | 0.934 | 1 | Notochord |
| Ddah1 | 0.000163 | 0.595232372 | 1 | 0.882 | 1 | Notochord |
| B3gat3 | 0.000171 | 0.410694712 | 1 | 0.855 | 1 | Notochord |
| Hs6st2 | 0.000189 | 0.270597089 | 1 | 0.737 | 1 | Notochord |
| Bcar3 | 0.000218 | 0.408481743 | 1 | 0.987 | 1 | Notochord |
| Rcn1 | 0.000238 | 0.864547159 | 1 | 0.987 | 1 | Notochord |
| Smoc1 | 0.000247 | 1.080582889 | 1 | 0.724 | 1 | Notochord |
| Ppfibp1 | 0.00026 | 0.358991304 | 1 | 0.934 | 1 | Notochord |
| Sh3bgr | 0.000287 | 0.353878291 | 1 | 0.289 | 1 | Notochord |
| Six1 | 0.000331 | 0.388518277 | 1 | 0.408 | 1 | Notochord |
| Arl4a | 0.00036 | 0.476730987 | 1 | 1 | 1 | Notochord |
| Krt18 | 0.000399 | 1.009381643 | 1 | 1 | 1 | Notochord |
| Prdx5 | 0.00041 | 0.319230271 | 1 | 0.961 | 1 | Notochord |
| Ube2k | 0.000424 | 0.380662179 | 1 | 1 | 1 | Notochord |
| Slc39a10 | 0.000442 | 0.64640244 | 1 | 1 | 1 | Notochord |
| Rcn2 | 0.00049 | 0.694843921 | 1 | 0.987 | 1 | Notochord |
| Reep5 | 0.000498 | 0.859664204 | 1 | 0.934 | 1 | Notochord |
| Fkbp9 | 0.000553 | 1.051362836 | 1 | 0.974 | 1 | Notochord |
| Slc12a4 | 0.000745 | 0.349852183 | 1 | 0.75 | 1 | Notochord |
| Dynll1 | 0.000755 | 0.716732256 | 1 | 1 | 1 | Notochord |

| | | | | | | |
|---|---|---|---|---|---|---|
| Efcab11 | 0.000819 | 0.398825356 | 1 | 0.553 | 1 | Notochord |
| Pltp | 0.000854 | 0.335474599 | 1 | 0.566 | 1 | Notochord |
| Iqcc | 0.000924 | 0.332097395 | 1 | 0.803 | 1 | Notochord |
| Mgst1 | 0.001082 | 0.411329029 | 1 | 0.618 | 1 | Notochord |
| Serf2 | 0.001148 | 0.497127238 | 1 | 1 | 1 | Notochord |
| Arsi | 0.001152 | 0.262744468 | 0.667 | 0.224 | 1 | Notochord |
| Psme2 | 0.00117 | 0.685246839 | 1 | 0.947 | 1 | Notochord |
| Spsb1 | 0.001175 | 0.425215545 | 0.667 | 0.605 | 1 | Notochord |
| Hsbp1 | 0.00121 | 0.428420731 | 1 | 1 | 1 | Notochord |
| Klk7 | 0.001316 | 0.287411574 | 1 | 0.092 | 1 | Notochord |
| Lysmd2 | 0.001373 | 0.565623629 | 1 | 0.342 | 1 | Notochord |
| Rasgef1b | 0.001392 | 0.277452811 | 1 | 0.421 | 1 | Notochord |
| Rbms1 | 0.001451 | 0.459448696 | 1 | 1 | 1 | Notochord |
| Bmp5 | 0.001455 | 0.259592834 | 1 | 0.145 | 1 | Notochord |
| Tceal8 | 0.001485 | 0.428860983 | 1 | 1 | 1 | Notochord |
| Nog | 0.001583 | 0.304608754 | 1 | 0.211 | 1 | Notochord |
| Fam89a | 0.00172 | 0.518638073 | 1 | 0.118 | 1 | Notochord |
| BC023829 | 0.001861 | 0.300658547 | 0.333 | 0.803 | 1 | Notochord |
| 2700029M | 0.001866 | 0.350575919 | 1 | 1 | 1 | Notochord |
| Fkbp10 | 0.001869 | 0.586842598 | 1 | 0.671 | 1 | Notochord |
| Marcksl1 | 0.001981 | 0.587057575 | 1 | 1 | 1 | Notochord |
| Anxa5 | 0.002037 | 0.974038856 | 1 | 0.855 | 1 | Notochord |
| Hapln1 | 0.002091 | 0.272845012 | 1 | 0.145 | 1 | Notochord |
| Sept9 | 0.002092 | 0.426870943 | 1 | 1 | 1 | Notochord |
| Litaf | 0.002271 | 0.685189472 | 1 | 1 | 1 | Notochord |
| Nrep | 0.002294 | 0.898715044 | 1 | 0.974 | 1 | Notochord |
| Dusp4 | 0.002391 | 0.902579582 | 1 | 0.763 | 1 | Notochord |
| Dpm2 | 0.002567 | 0.338005402 | 1 | 0.908 | 1 | Notochord |
| Nudt16l1 | 0.002846 | 0.263443473 | 1 | 0.974 | 1 | Notochord |
| Snrk | 0.002886 | 0.261237343 | 1 | 0.737 | 1 | Notochord |
| Cirbp | 0.002897 | 0.342462338 | 1 | 0.868 | 1 | Notochord |
| Fam213b | 0.002971 | 0.271548624 | 1 | 0.421 | 1 | Notochord |
| Rhoc | 0.003232 | 0.681343211 | 1 | 0.842 | 1 | Notochord |
| Tmem106a | 0.003339 | 0.258515162 | 0.667 | 0.382 | 1 | Notochord |
| Tmem246 | 0.003526 | 0.283744495 | 1 | 0.592 | 1 | Notochord |
| Zfp758 | 0.003804 | 0.345911403 | 1 | 0.855 | 1 | Notochord |
| Fabp7 | 0.004192 | 0.540334456 | 1 | 0.368 | 1 | Notochord |
| Actb | 0.004219 | 0.4213859 | 1 | 1 | 1 | Notochord |
| Brk1 | 0.004252 | 0.573234682 | 1 | 1 | 1 | Notochord |
| Armc1 | 0.004303 | 0.486884016 | 1 | 0.974 | 1 | Notochord |
| Hoxa1 | 0.004338 | 0.828714539 | 0.667 | 0.776 | 1 | Notochord |
| Spats2l | 0.00436 | 0.260176513 | 1 | 0.382 | 1 | Notochord |
| Ncstn | 0.00449 | 0.375894081 | 1 | 1 | 1 | Notochord |
| Creb3l2 | 0.004494 | 0.322280903 | 1 | 0.724 | 1 | Notochord |
| Gabarap | 0.004547 | 0.387355705 | 1 | 0.987 | 1 | Notochord |
| Sms | 0.004607 | 0.433959467 | 1 | 1 | 1 | Notochord |
| Snap47 | 0.005064 | 0.592425828 | 1 | 0.947 | 1 | Notochord |
| Tmsb10 | 0.005132 | 0.982741706 | 1 | 1 | 1 | Notochord |
| Ube2n | 0.005367 | 0.315142135 | 1 | 1 | 1 | Notochord |
| Maged1 | 0.005432 | 0.726430957 | 1 | 1 | 1 | Notochord |
| Itpa | 0.005502 | 0.340273501 | 1 | 0.961 | 1 | Notochord |
| Ubap1 | 0.005529 | 0.28225769 | 1 | 0.947 | 1 | Notochord |
| Tex2 | 0.005837 | 0.260207668 | 1 | 0.711 | 1 | Notochord |
| Tagln | 0.006353 | 0.400457422 | 1 | 0.237 | 1 | Notochord |
| S100a11 | 0.006726 | 0.601293592 | 1 | 0.882 | 1 | Notochord |
| Manbal | 0.007037 | 0.401412318 | 1 | 0.855 | 1 | Notochord |
| Mical1 | 0.007165 | 0.27754978 | 1 | 0.882 | 1 | Notochord |
| 9530068E( | 0.007216 | 0.412679237 | 1 | 0.961 | 1 | Notochord |

| | | | | | | |
|---|---|---|---|---|---|---|
| Apitd1 | 0.007358 | 0.605625955 | 1 | 0.868 | 1 | Notochord |
| Dnajc10 | 0.007395 | 0.418355323 | 1 | 0.987 | 1 | Notochord |
| Rpl7l1 | 0.007562 | 0.251812508 | 1 | 1 | 1 | Notochord |
| Mpzl1 | 0.007637 | 0.434882267 | 1 | 0.987 | 1 | Notochord |
| Copa | 0.008047 | 0.361388988 | 1 | 1 | 1 | Notochord |
| Dguok | 0.008047 | 0.312740636 | 1 | 0.947 | 1 | Notochord |
| Sp5 | 0.008449 | 0.796257716 | 1 | 0.566 | 1 | Notochord |
| Fermt2 | 0.008498 | 0.599650217 | 1 | 0.987 | 1 | Notochord |
| Twsg1 | 0.008543 | 0.265778694 | 1 | 0.974 | 1 | Notochord |
| Tbc1d7 | 0.008578 | 0.288464298 | 0.667 | 0.724 | 1 | Notochord |
| Gpatch1 | 0.008628 | 0.25407407 | 1 | 0.75 | 1 | Notochord |
| Tgfb2 | 0.009069 | 0.520823681 | 1 | 0.724 | 1 | Notochord |
| Ubb | 0.009327 | 0.279308897 | 1 | 1 | 1 | Notochord |
| Foxd4 | 0.009613 | 0.898801472 | 0.667 | 0.197 | 1 | Notochord |
| Sf3a3 | 0.009647 | 0.32317614 | 1 | 1 | 1 | Notochord |
| Pxk | 0.009715 | 0.448407175 | 1 | 0.803 | 1 | Notochord |
| Rcn3 | 0.009787 | 0.570935782 | 1 | 1 | 1 | Notochord |
| Nkx2-9 | 8.82E-45 | 1.615938853 | 1 | 0.389 | 1.11E-40 | Neural Tube |
| Phox2b | 4.64E-33 | 0.724756584 | 0.86 | 0.361 | 5.83E-29 | Neural Tube |
| Rfx4 | 2.93E-32 | 0.984166233 | 1 | 0.556 | 3.69E-28 | Neural Tube |
| Olig2 | 2.00E-30 | 0.409292443 | 0.791 | 0.306 | 2.51E-26 | Neural Tube |
| Nkx6-1 | 1.08E-25 | 0.518319702 | 1 | 0.333 | 1.36E-21 | Neural Tube |
| Map1b | 3.98E-24 | 1.027662434 | 1 | 0.944 | 5.00E-20 | Neural Tube |
| Fez1 | 5.51E-24 | 0.395943025 | 1 | 0.25 | 6.93E-20 | Neural Tube |
| Timp3 | 4.36E-23 | 0.433542461 | 1 | 0.861 | 5.48E-19 | Neural Tube |
| Dtx4 | 3.58E-20 | 0.283701038 | 0.953 | 0.583 | 4.50E-16 | Neural Tube |
| Apba2 | 1.56E-19 | 0.291977275 | 0.86 | 0.194 | 1.96E-15 | Neural Tube |
| Crabp1 | 9.46E-19 | 2.861388142 | 1 | 1 | 1.19E-14 | Neural Tube |
| Vangl2 | 1.74E-18 | 0.703656303 | 1 | 0.972 | 2.19E-14 | Neural Tube |
| Emilin2 | 6.06E-18 | 0.304492145 | 0.884 | 0.528 | 7.62E-14 | Neural Tube |
| Nrcam | 6.47E-18 | 0.368065931 | 1 | 0.861 | 8.14E-14 | Neural Tube |
| Crmp1 | 6.97E-18 | 0.552604089 | 0.953 | 0.694 | 8.77E-14 | Neural Tube |
| Msi1 | 2.19E-17 | 0.302593336 | 1 | 0.889 | 2.75E-13 | Neural Tube |
| Ptch1 | 3.04E-17 | 0.911595283 | 1 | 0.917 | 3.82E-13 | Neural Tube |
| Fam181a | 6.20E-17 | 0.366269019 | 0.767 | 0.194 | 7.80E-13 | Neural Tube |
| Mafb | 9.46E-17 | 0.317292749 | 0.651 | 0.389 | 1.19E-12 | Neural Tube |
| Nkx6-2 | 1.11E-16 | 0.289012556 | 0.721 | 0.083 | 1.40E-12 | Neural Tube |
| Crabp2 | 1.28E-16 | 2.233817459 | 1 | 0.917 | 1.61E-12 | Neural Tube |
| Tmem47 | 2.23E-16 | 0.90212273 | 0.977 | 0.917 | 2.80E-12 | Neural Tube |
| Ednrb | 4.45E-16 | 0.456279021 | 0.93 | 0.361 | 5.59E-12 | Neural Tube |
| Mapk11 | 5.98E-16 | 0.43018603 | 0.884 | 0.389 | 7.52E-12 | Neural Tube |
| Nr6a1 | 6.84E-16 | 0.397135409 | 1 | 1 | 8.60E-12 | Neural Tube |
| Mn1 | 7.12E-16 | 0.405905343 | 1 | 0.972 | 8.95E-12 | Neural Tube |
| Rnf220 | 2.67E-15 | 0.398807989 | 1 | 0.889 | 3.36E-11 | Neural Tube |
| Nell2 | 2.73E-15 | 0.291192403 | 0.86 | 0.389 | 3.44E-11 | Neural Tube |
| Peg12 | 2.77E-15 | 0.625310721 | 1 | 0.806 | 3.49E-11 | Neural Tube |
| Olfml3 | 3.42E-15 | 0.405605519 | 1 | 0.694 | 4.31E-11 | Neural Tube |
| Ncam1 | 4.85E-15 | 0.49258733 | 1 | 0.806 | 6.09E-11 | Neural Tube |
| Igdcc3 | 5.11E-15 | 0.378342838 | 1 | 0.694 | 6.43E-11 | Neural Tube |
| Sfrp1 | 8.58E-15 | 0.764455712 | 1 | 0.944 | 1.08E-10 | Neural Tube |
| Ptn | 9.93E-15 | 1.036399491 | 0.977 | 0.472 | 1.25E-10 | Neural Tube |
| Fgfr2 | 2.08E-14 | 0.361052106 | 1 | 0.694 | 2.62E-10 | Neural Tube |
| Sparcl1 | 6.46E-14 | 0.781179996 | 0.93 | 0.333 | 8.12E-10 | Neural Tube |
| Zfp811 | 1.42E-13 | 0.357270836 | 0.907 | 0.167 | 1.78E-09 | Neural Tube |
| Aldh1a2 | 1.55E-13 | 0.319395097 | 0.651 | 0.583 | 1.94E-09 | Neural Tube |
| Sept3 | 2.13E-13 | 0.329131228 | 0.977 | 0.639 | 2.68E-09 | Neural Tube |
| Meis1 | 2.52E-13 | 0.333735705 | 1 | 0.722 | 3.16E-09 | Neural Tube |
| Cdh2 | 3.94E-13 | 0.660594514 | 1 | 1 | 4.95E-09 | Neural Tube |

| | | | | | | |
|---|---|---|---|---|---|---|
| Crb2 | 1.04E-12 | 0.263352485 | 1 | 0.556 | 1.31E-08 | Neural Tube |
| Camkv | 1.87E-12 | 0.28055737 | 0.884 | 0.306 | 2.35E-08 | Neural Tube |
| Sfrp2 | 2.93E-12 | 0.392397179 | 0.884 | 0.417 | 3.68E-08 | Neural Tube |
| Prtg | 3.11E-12 | 0.768406768 | 1 | 1 | 3.91E-08 | Neural Tube |
| Galnt16 | 7.70E-12 | 0.264856431 | 0.791 | 0.417 | 9.69E-08 | Neural Tube |
| Hoxa2 | 1.13E-11 | 0.255230346 | 0.767 | 0.222 | 1.42E-07 | Neural Tube |
| Spsb4 | 1.41E-11 | 0.342303128 | 0.977 | 0.639 | 1.78E-07 | Neural Tube |
| Fgfbp3 | 1.57E-11 | 0.503852273 | 0.86 | 0.333 | 1.97E-07 | Neural Tube |
| Net1 | 1.91E-11 | 0.694855918 | 1 | 1 | 2.40E-07 | Neural Tube |
| Meox1 | 2.28E-11 | 0.425423737 | 0.837 | 0.694 | 2.86E-07 | Neural Tube |
| Spon1 | 4.19E-11 | 0.332858421 | 1 | 0.667 | 5.27E-07 | Neural Tube |
| Rai14 | 6.99E-11 | 0.463377 | 0.977 | 0.944 | 8.79E-07 | Neural Tube |
| Efnb3 | 7.66E-11 | 0.516342839 | 1 | 0.694 | 9.63E-07 | Neural Tube |
| Zmynd8 | 8.04E-11 | 0.316969963 | 1 | 1 | 1.01E-06 | Neural Tube |
| Meis2 | 9.56E-11 | 0.251384016 | 1 | 0.806 | 1.20E-06 | Neural Tube |
| Stmn3 | 1.70E-10 | 0.434415247 | 0.814 | 0.222 | 2.14E-06 | Neural Tube |
| Phf6 | 1.97E-10 | 0.371912158 | 0.977 | 0.944 | 2.48E-06 | Neural Tube |
| Gli1 | 2.13E-10 | 0.25283194 | 0.977 | 0.528 | 2.68E-06 | Neural Tube |
| Idi1 | 2.76E-10 | 0.569310103 | 1 | 0.889 | 3.47E-06 | Neural Tube |
| Acly | 3.19E-10 | 0.617705343 | 1 | 1 | 4.02E-06 | Neural Tube |
| Anp32e | 5.24E-10 | 0.573978945 | 1 | 1 | 6.59E-06 | Neural Tube |
| Zfp41 | 6.88E-10 | 0.360022425 | 0.953 | 0.861 | 8.65E-06 | Neural Tube |
| Igf1r | 7.09E-10 | 0.335029497 | 0.977 | 0.75 | 8.92E-06 | Neural Tube |
| Mycl | 7.13E-10 | 0.395934532 | 0.953 | 0.722 | 8.97E-06 | Neural Tube |
| Dek | 8.04E-10 | 0.795034433 | 1 | 1 | 1.01E-05 | Neural Tube |
| Egr2 | 8.19E-10 | 0.348032473 | 0.651 | 0.472 | 1.03E-05 | Neural Tube |
| Lin28b | 8.37E-10 | 0.312833628 | 1 | 0.861 | 1.05E-05 | Neural Tube |
| Pknox2 | 1.21E-09 | 0.374696532 | 1 | 0.806 | 1.52E-05 | Neural Tube |
| Egr1 | 1.24E-09 | 1.24434849 | 1 | 0.944 | 1.56E-05 | Neural Tube |
| Pygb | 1.36E-09 | 0.575125204 | 0.977 | 0.972 | 1.71E-05 | Neural Tube |
| Pcsk9 | 1.38E-09 | 0.384030607 | 0.884 | 0.417 | 1.74E-05 | Neural Tube |
| Qk | 1.83E-09 | 0.278788843 | 1 | 1 | 2.30E-05 | Neural Tube |
| Llgl1 | 2.16E-09 | 0.367010277 | 1 | 0.944 | 2.72E-05 | Neural Tube |
| Tagln3 | 2.18E-09 | 0.353952566 | 0.721 | 0.278 | 2.74E-05 | Neural Tube |
| Nsg1 | 2.20E-09 | 0.269085946 | 0.767 | 0.194 | 2.77E-05 | Neural Tube |
| Vezf1 | 2.45E-09 | 0.349104758 | 1 | 0.917 | 3.07E-05 | Neural Tube |
| Dpysl2 | 2.70E-09 | 0.586619802 | 1 | 1 | 3.40E-05 | Neural Tube |
| Dusp1 | 2.71E-09 | 0.485697784 | 0.953 | 0.972 | 3.41E-05 | Neural Tube |
| 1700011H | 3.87E-09 | 1.153551988 | 0.953 | 0.861 | 4.87E-05 | Neural Tube |
| Msn | 4.94E-09 | 0.400920822 | 0.953 | 0.972 | 6.21E-05 | Neural Tube |
| Hpcal1 | 5.15E-09 | 0.442364456 | 0.953 | 0.972 | 6.48E-05 | Neural Tube |
| Grb10 | 6.11E-09 | 0.484653001 | 1 | 1 | 7.69E-05 | Neural Tube |
| Slc39a6 | 6.27E-09 | 0.337910583 | 1 | 0.722 | 7.89E-05 | Neural Tube |
| Ppp1r10 | 1.15E-08 | 0.570951601 | 1 | 0.972 | 0.000144 | Neural Tube |
| Cenpf | 1.61E-08 | 0.255840343 | 1 | 1 | 0.000202 | Neural Tube |
| Mki67 | 2.04E-08 | 0.395435238 | 1 | 0.944 | 0.000257 | Neural Tube |
| Hoxb1 | 2.08E-08 | 0.560065472 | 0.86 | 0.667 | 0.000262 | Neural Tube |
| Cenpe | 2.52E-08 | 0.363267875 | 1 | 1 | 0.000317 | Neural Tube |
| Klc1 | 2.59E-08 | 0.369721208 | 1 | 0.972 | 0.000325 | Neural Tube |
| Dnmt1 | 2.79E-08 | 0.513822121 | 1 | 1 | 0.000351 | Neural Tube |
| Rad50 | 4.43E-08 | 0.264411039 | 0.953 | 0.917 | 0.000557 | Neural Tube |
| Gpr19 | 4.90E-08 | 0.278426935 | 0.884 | 0.389 | 0.000616 | Neural Tube |
| Lpar6 | 5.80E-08 | 0.422918831 | 1 | 0.972 | 0.000729 | Neural Tube |
| Arvcf | 5.98E-08 | 0.39708021 | 1 | 0.833 | 0.000753 | Neural Tube |
| Nasp | 6.64E-08 | 0.430737055 | 1 | 1 | 0.000835 | Neural Tube |
| Dpysl5 | 9.86E-08 | 0.359087335 | 1 | 0.972 | 0.00124 | Neural Tube |
| Kcne1l | 1.44E-07 | 0.35326953 | 0.698 | 0.25 | 0.001813 | Neural Tube |
| Rarb | 1.46E-07 | 0.452315548 | 0.977 | 0.556 | 0.001833 | Neural Tube |

| | | | | | | |
|---|---|---|---|---|---|---|
| Col18a1 | 1.66E-07 | 0.878613741 | 1 | 1 | 0.00209 | Neural Tube |
| Kif15 | 1.84E-07 | 0.387507096 | 1 | 0.972 | 0.002316 | Neural Tube |
| Gtse1 | 1.87E-07 | 0.34761896 | 0.907 | 0.806 | 0.002354 | Neural Tube |
| Armcx1 | 1.90E-07 | 0.380285805 | 1 | 0.833 | 0.002396 | Neural Tube |
| Hmgcs1 | 1.91E-07 | 0.657338482 | 1 | 0.944 | 0.002401 | Neural Tube |
| Sh3bgrl | 2.15E-07 | 0.613212958 | 1 | 0.944 | 0.002701 | Neural Tube |
| Arhgap19 | 2.69E-07 | 0.259039156 | 1 | 0.972 | 0.003377 | Neural Tube |
| Vangl1 | 2.73E-07 | 0.365516462 | 1 | 0.944 | 0.003437 | Neural Tube |
| Gm10184 | 3.06E-07 | 0.544147119 | 0.977 | 1 | 0.003845 | Neural Tube |
| Sept11 | 3.63E-07 | 0.370415195 | 1 | 1 | 0.00457 | Neural Tube |
| Rbbp5 | 4.09E-07 | 0.301342114 | 0.953 | 1 | 0.005143 | Neural Tube |
| Mcm7 | 4.25E-07 | 0.440037831 | 1 | 1 | 0.005346 | Neural Tube |
| Ccnd2 | 4.33E-07 | 1.024748449 | 1 | 1 | 0.005447 | Neural Tube |
| Zfml | 4.36E-07 | 0.303276511 | 1 | 0.972 | 0.005478 | Neural Tube |
| Rcc2 | 5.14E-07 | 0.316035824 | 1 | 1 | 0.006465 | Neural Tube |
| Pald1 | 5.86E-07 | 0.304938234 | 1 | 0.889 | 0.007366 | Neural Tube |
| Rtn3 | 5.88E-07 | 0.519980871 | 1 | 1 | 0.007392 | Neural Tube |
| Gm4076 | 6.89E-07 | 0.57291107 | 1 | 1 | 0.008663 | Neural Tube |
| Lig1 | 7.08E-07 | 0.524154096 | 1 | 1 | 0.008904 | Neural Tube |
| Hmgb3 | 7.10E-07 | 0.424776988 | 1 | 1 | 0.008933 | Neural Tube |
| Cep55 | 7.77E-07 | 0.326917013 | 0.907 | 0.694 | 0.009776 | Neural Tube |
| Myh10 | 8.39E-07 | 0.373696583 | 1 | 0.972 | 0.010546 | Neural Tube |
| Gsk3b | 8.42E-07 | 0.283500556 | 1 | 1 | 0.010586 | Neural Tube |
| Thrap3 | 9.21E-07 | 0.320572674 | 1 | 1 | 0.011577 | Neural Tube |
| Cdca7 | 9.50E-07 | 0.686284977 | 1 | 1 | 0.011953 | Neural Tube |
| Hnrnph1 | 1.09E-06 | 0.339192525 | 1 | 1 | 0.013754 | Neural Tube |
| Nusap1 | 1.19E-06 | 0.436476299 | 0.86 | 0.889 | 0.014968 | Neural Tube |
| Plekha5 | 1.25E-06 | 0.264057497 | 0.977 | 0.917 | 0.015696 | Neural Tube |
| Gpx3 | 1.28E-06 | 0.743634437 | 1 | 1 | 0.016036 | Neural Tube |
| Ccdc160 | 1.32E-06 | 0.336447207 | 0.651 | 0.194 | 0.016606 | Neural Tube |
| Mdk | 1.39E-06 | 0.337086977 | 1 | 1 | 0.017535 | Neural Tube |
| Top2a | 1.60E-06 | 0.53747056 | 1 | 1 | 0.020178 | Neural Tube |
| Dll1 | 1.63E-06 | 0.274545112 | 0.674 | 0.611 | 0.020553 | Neural Tube |
| Ipo5 | 1.81E-06 | 0.499265148 | 1 | 1 | 0.02281 | Neural Tube |
| Rad51ap1 | 1.88E-06 | 0.272991243 | 0.907 | 0.972 | 0.023649 | Neural Tube |
| Ilf3 | 1.95E-06 | 0.453807179 | 1 | 1 | 0.024565 | Neural Tube |
| Fam64a | 2.02E-06 | 0.509734826 | 0.953 | 0.944 | 0.025393 | Neural Tube |
| Ncaph | 2.05E-06 | 0.334190096 | 1 | 0.917 | 0.02572 | Neural Tube |
| Sfpq | 2.17E-06 | 0.367381693 | 1 | 1 | 0.027308 | Neural Tube |
| Meis3 | 2.33E-06 | 0.257678486 | 0.977 | 0.778 | 0.029267 | Neural Tube |
| Prpf3 | 2.34E-06 | 0.276294171 | 1 | 0.944 | 0.029441 | Neural Tube |
| Cdh6 | 2.78E-06 | 0.377016073 | 0.953 | 0.528 | 0.034972 | Neural Tube |
| Fndc3c1 | 2.93E-06 | 0.268013068 | 1 | 0.917 | 0.036885 | Neural Tube |
| Fubp1 | 3.16E-06 | 0.256427573 | 1 | 1 | 0.039793 | Neural Tube |
| Brd2 | 3.26E-06 | 0.276474705 | 1 | 1 | 0.040992 | Neural Tube |
| Apex1 | 3.88E-06 | 0.404397248 | 1 | 1 | 0.04879 | Neural Tube |
| Klf13 | 4.09E-06 | 0.279711797 | 1 | 1 | 0.051376 | Neural Tube |
| Dkc1 | 4.13E-06 | 0.52029247 | 1 | 1 | 0.051945 | Neural Tube |
| Arl5c | 4.26E-06 | 0.312160045 | 1 | 0.972 | 0.053542 | Neural Tube |
| Snrnp200 | 4.35E-06 | 0.268819294 | 1 | 1 | 0.05466 | Neural Tube |
| Gsg2 | 4.38E-06 | 0.264835206 | 0.744 | 0.472 | 0.055058 | Neural Tube |
| Rfc1 | 4.49E-06 | 0.304889104 | 1 | 1 | 0.056434 | Neural Tube |
| Ndn | 4.76E-06 | 0.503258329 | 0.977 | 0.806 | 0.05984 | Neural Tube |
| Efna5 | 5.44E-06 | 0.431610102 | 1 | 0.806 | 0.068453 | Neural Tube |
| Kpna2 | 5.63E-06 | 0.6978043 | 1 | 1 | 0.070857 | Neural Tube |
| Prc1 | 6.70E-06 | 0.339825091 | 1 | 1 | 0.084301 | Neural Tube |
| Elovl2 | 7.47E-06 | 0.315780421 | 0.93 | 0.75 | 0.093946 | Neural Tube |
| Jam3 | 8.31E-06 | 0.419636974 | 1 | 0.889 | 0.104464 | Neural Tube |

| | | | | | | |
|---|---|---|---|---|---|---|
| Tsc22d1 | 8.71E-06 | 0.375188779 | 1 | 1 | 0.109538 | Neural Tube |
| Trim35 | 8.84E-06 | 0.393861072 | 0.977 | 0.972 | 0.111181 | Neural Tube |
| Leo1 | 8.99E-06 | 0.488052225 | 0.953 | 1 | 0.113118 | Neural Tube |
| Rsrc2 | 1.06E-05 | 0.368373593 | 1 | 1 | 0.132925 | Neural Tube |
| Agk | 1.15E-05 | 0.268571491 | 0.884 | 0.778 | 0.144866 | Neural Tube |
| Eif5 | 1.21E-05 | 0.494286341 | 1 | 1 | 0.151788 | Neural Tube |
| Aurkb | 1.40E-05 | 0.46930805 | 0.953 | 0.917 | 0.176148 | Neural Tube |
| Kpnb1 | 1.45E-05 | 0.27420643 | 1 | 1 | 0.182745 | Neural Tube |
| Serbp1 | 1.69E-05 | 0.399046903 | 1 | 1 | 0.212824 | Neural Tube |
| Ccnf | 2.40E-05 | 0.36738876 | 0.953 | 0.972 | 0.301364 | Neural Tube |
| Skap2 | 2.53E-05 | 0.425519887 | 0.907 | 0.889 | 0.318777 | Neural Tube |
| Polr2b | 2.62E-05 | 0.331777252 | 1 | 1 | 0.329734 | Neural Tube |
| Tubb2b | 2.75E-05 | 0.253898565 | 1 | 0.972 | 0.346148 | Neural Tube |
| Ddx5 | 3.03E-05 | 0.405021308 | 1 | 1 | 0.381345 | Neural Tube |
| Sf3b3 | 3.09E-05 | 0.416901523 | 1 | 1 | 0.388516 | Neural Tube |
| Rbm39 | 3.52E-05 | 0.367418054 | 1 | 1 | 0.443021 | Neural Tube |
| Fos | 3.76E-05 | 0.773655905 | 0.953 | 0.917 | 0.473052 | Neural Tube |
| EGFP | 4.22E-05 | 0.562506269 | 1 | 1 | 0.530698 | Neural Tube |
| Fxyd6 | 4.30E-05 | 0.569335225 | 1 | 0.833 | 0.540696 | Neural Tube |
| Zfp422 | 4.41E-05 | 0.337609931 | 1 | 1 | 0.554248 | Neural Tube |
| Nucks1 | 4.95E-05 | 0.291393018 | 1 | 1 | 0.623037 | Neural Tube |
| Nnat | 4.97E-05 | 0.427126753 | 1 | 1 | 0.624973 | Neural Tube |
| Ift74 | 5.16E-05 | 0.422037322 | 0.93 | 0.917 | 0.648637 | Neural Tube |
| Hnrnpa2b: | 5.50E-05 | 0.464353814 | 1 | 1 | 0.691684 | Neural Tube |
| Hmgn2 | 6.50E-05 | 0.529910816 | 1 | 1 | 0.818066 | Neural Tube |
| Pkm | 6.83E-05 | 0.397864431 | 1 | 1 | 0.859548 | Neural Tube |
| Ptp4a2 | 9.80E-05 | 0.305483848 | 1 | 1 | 1 | Neural Tube |
| Nap1l1 | 9.83E-05 | 0.399423838 | 1 | 1 | 1 | Neural Tube |
| Sf3b2 | 0.000101 | 0.472592358 | 1 | 1 | 1 | Neural Tube |
| Scoc | 0.000101 | 0.385977366 | 1 | 0.944 | 1 | Neural Tube |
| Marcksl1 | 0.000105 | 0.2915567 | 1 | 1 | 1 | Neural Tube |
| Smc3 | 0.000105 | 0.32082772 | 1 | 1 | 1 | Neural Tube |
| Incenp | 0.000106 | 0.551851019 | 1 | 0.944 | 1 | Neural Tube |
| Fgf15 | 0.000107 | 0.412744367 | 0.744 | 0.389 | 1 | Neural Tube |
| Sox11 | 0.000109 | 0.442222144 | 1 | 0.972 | 1 | Neural Tube |
| Hnrnpm | 0.000117 | 0.441627144 | 1 | 1 | 1 | Neural Tube |
| Mrfap1 | 0.000124 | 0.341669215 | 1 | 1 | 1 | Neural Tube |
| Wdr6 | 0.000131 | 0.374427774 | 1 | 1 | 1 | Neural Tube |
| Trp53i11 | 0.000131 | 0.347793785 | 0.977 | 0.917 | 1 | Neural Tube |
| Btg2 | 0.000137 | 0.434741489 | 0.93 | 0.889 | 1 | Neural Tube |
| Jun | 0.000138 | 0.704132101 | 1 | 0.972 | 1 | Neural Tube |
| Map2k6 | 0.000143 | 0.310469403 | 0.977 | 0.944 | 1 | Neural Tube |
| Zfp930 | 0.000147 | 0.274018157 | 0.907 | 0.972 | 1 | Neural Tube |
| Rad9a | 0.00015 | 0.360362402 | 1 | 0.972 | 1 | Neural Tube |
| Lmnb1 | 0.00017 | 0.348151634 | 1 | 1 | 1 | Neural Tube |
| Ckb | 0.000175 | 0.760550981 | 0.977 | 0.806 | 1 | Neural Tube |
| Cep57 | 0.000179 | 0.304566966 | 1 | 1 | 1 | Neural Tube |
| Dars | 0.000184 | 0.279121898 | 1 | 1 | 1 | Neural Tube |
| 2610318N | 0.000187 | 0.302087051 | 1 | 1 | 1 | Neural Tube |
| Eif4b | 0.000198 | 0.336874832 | 1 | 1 | 1 | Neural Tube |
| Akap8 | 0.000206 | 0.342768652 | 1 | 1 | 1 | Neural Tube |
| Azin1 | 0.000214 | 0.261762524 | 1 | 1 | 1 | Neural Tube |
| Gsta4 | 0.000221 | 0.422152684 | 0.953 | 0.944 | 1 | Neural Tube |
| Spop | 0.000229 | 0.33703462 | 1 | 1 | 1 | Neural Tube |
| Trim59 | 0.000235 | 0.439984962 | 1 | 0.972 | 1 | Neural Tube |
| Blcap | 0.000239 | 0.264326157 | 0.977 | 0.917 | 1 | Neural Tube |
| Atp6v0e2 | 0.00024 | 0.29937477 | 0.884 | 0.667 | 1 | Neural Tube |
| Hsd11b2 | 0.00024 | 0.425193645 | 0.953 | 0.833 | 1 | Neural Tube |

| | | | | | | |
|---|---|---|---|---|---|---|
| Dnajc9 | 0.000264 | 0.549781094 | 1 | 1 | 1 | Neural Tube |
| Kif20b | 0.000284 | 0.260404459 | 0.977 | 0.944 | 1 | Neural Tube |
| Abcf1 | 0.000288 | 0.311592736 | 1 | 1 | 1 | Neural Tube |
| Kdm3a | 0.000292 | 0.256132237 | 1 | 0.889 | 1 | Neural Tube |
| Ptma | 0.000292 | 0.372239614 | 1 | 1 | 1 | Neural Tube |
| Ndc1 | 0.000306 | 0.264587772 | 1 | 1 | 1 | Neural Tube |
| Rbm14 | 0.000308 | 0.265891076 | 1 | 1 | 1 | Neural Tube |
| Ankrd10 | 0.000316 | 0.275649906 | 1 | 1 | 1 | Neural Tube |
| Mns1 | 0.000323 | 0.255167128 | 1 | 0.944 | 1 | Neural Tube |
| H1f0 | 0.000332 | 0.659444751 | 1 | 1 | 1 | Neural Tube |
| Ndc80 | 0.000383 | 0.38863701 | 0.907 | 0.972 | 1 | Neural Tube |
| Igfbp5 | 0.000394 | 0.389770053 | 1 | 0.75 | 1 | Neural Tube |
| Rab31 | 0.0004 | 0.281355099 | 0.953 | 0.778 | 1 | Neural Tube |
| Igf2bp1 | 0.000401 | 0.313630334 | 1 | 1 | 1 | Neural Tube |
| Bub1b | 0.000417 | 0.376238356 | 1 | 0.972 | 1 | Neural Tube |
| Sf3b1 | 0.000429 | 0.275565452 | 1 | 1 | 1 | Neural Tube |
| Dnajb9 | 0.000447 | 0.395246021 | 0.907 | 0.972 | 1 | Neural Tube |
| Nudt9 | 0.000451 | 0.321362392 | 1 | 1 | 1 | Neural Tube |
| Set | 0.000464 | 0.378922754 | 1 | 1 | 1 | Neural Tube |
| Tuba1a | 0.000513 | 0.318681456 | 1 | 1 | 1 | Neural Tube |
| Fstl1 | 0.000516 | 0.385207002 | 1 | 1 | 1 | Neural Tube |
| Srsf10 | 0.00054 | 0.299640756 | 1 | 1 | 1 | Neural Tube |
| Ppat | 0.000559 | 0.309449254 | 1 | 1 | 1 | Neural Tube |
| Ap1m1 | 0.000582 | 0.317975458 | 1 | 1 | 1 | Neural Tube |
| Bub3 | 0.000588 | 0.411395343 | 1 | 1 | 1 | Neural Tube |
| Kif22 | 0.000592 | 0.472630435 | 1 | 1 | 1 | Neural Tube |
| Fbxo5 | 0.000596 | 0.409391333 | 0.977 | 0.972 | 1 | Neural Tube |
| Bora | 0.00061 | 0.275857335 | 1 | 0.944 | 1 | Neural Tube |
| Ska2 | 0.000627 | 0.29815316 | 1 | 1 | 1 | Neural Tube |
| Ckap2l | 0.000627 | 0.434718776 | 0.907 | 0.889 | 1 | Neural Tube |
| Kif11 | 0.000632 | 0.308649029 | 0.977 | 0.944 | 1 | Neural Tube |
| Cdc42se2 | 0.000641 | 0.37631208 | 1 | 0.972 | 1 | Neural Tube |
| Mybbp1a | 0.000722 | 0.322213601 | 1 | 1 | 1 | Neural Tube |
| Smc4 | 0.000729 | 0.407536826 | 1 | 1 | 1 | Neural Tube |
| Gm10282 | 0.000733 | 0.451285957 | 1 | 1 | 1 | Neural Tube |
| Foxa2 | 0.000746 | 0.330895852 | 1 | 1 | 1 | Neural Tube |
| Sde2 | 0.000754 | 0.275602237 | 0.953 | 0.972 | 1 | Neural Tube |
| Nap1l4 | 0.000756 | 0.361547232 | 1 | 1 | 1 | Neural Tube |
| 2810417H | 0.000764 | 0.358020997 | 1 | 1 | 1 | Neural Tube |
| Ccar1 | 0.000863 | 0.267410471 | 1 | 1 | 1 | Neural Tube |
| Slc38a2 | 0.000924 | 0.386200465 | 1 | 1 | 1 | Neural Tube |
| Hnrnph2 | 0.000981 | 0.316120112 | 1 | 1 | 1 | Neural Tube |
| Mcm2 | 0.000993 | 0.402469645 | 1 | 1 | 1 | Neural Tube |
| Pnrc2 | 0.001026 | 0.366432502 | 0.977 | 1 | 1 | Neural Tube |
| Chtop | 0.001074 | 0.340779704 | 1 | 1 | 1 | Neural Tube |
| Hmgn5 | 0.001103 | 0.318301053 | 0.977 | 0.889 | 1 | Neural Tube |
| Plk1 | 0.001109 | 0.624609859 | 1 | 1 | 1 | Neural Tube |
| Kif2c | 0.00121 | 0.33788301 | 1 | 0.972 | 1 | Neural Tube |
| Pafah1b2 | 0.001219 | 0.323354095 | 1 | 1 | 1 | Neural Tube |
| Pebp1 | 0.001271 | 0.276999374 | 1 | 1 | 1 | Neural Tube |
| Parp1 | 0.001296 | 0.300642805 | 1 | 1 | 1 | Neural Tube |
| Hnrnpc | 0.001296 | 0.411229047 | 1 | 1 | 1 | Neural Tube |
| 1110004F1 | 0.001327 | 0.298494447 | 1 | 1 | 1 | Neural Tube |
| Fgfr1op | 0.001393 | 0.265494349 | 0.977 | 0.972 | 1 | Neural Tube |
| Elovl5 | 0.001411 | 0.308287176 | 1 | 0.972 | 1 | Neural Tube |
| Hmmr | 0.001452 | 0.278615001 | 1 | 1 | 1 | Neural Tube |
| Rfc4 | 0.001476 | 0.292159317 | 1 | 0.917 | 1 | Neural Tube |
| Ddx21 | 0.001588 | 0.329025038 | 1 | 1 | 1 | Neural Tube |

| | | | | | | |
|---|---|---|---|---|---|---|
| Eno1b | 0.001632 | 0.277566319 | 1 | 1 | 1 | Neural Tube |
| Fads2 | 0.001727 | 0.349879635 | 1 | 0.889 | 1 | Neural Tube |
| Fam60a | 0.001763 | 0.351491943 | 1 | 1 | 1 | Neural Tube |
| Golm1 | 0.001806 | 0.399701976 | 1 | 1 | 1 | Neural Tube |
| Slc2a1 | 0.001883 | 0.62322719 | 1 | 1 | 1 | Neural Tube |
| Kif23 | 0.001887 | 0.284082386 | 0.93 | 1 | 1 | Neural Tube |
| Nedd4 | 0.0021 | 0.302744183 | 1 | 1 | 1 | Neural Tube |
| Mcm3 | 0.002151 | 0.401188245 | 1 | 1 | 1 | Neural Tube |
| Ssb | 0.002262 | 0.434221271 | 1 | 1 | 1 | Neural Tube |
| Ccnd1 | 0.002327 | 0.284084756 | 1 | 0.944 | 1 | Neural Tube |
| Fads1 | 0.002343 | 0.274110913 | 1 | 0.944 | 1 | Neural Tube |
| Glyr1 | 0.00241 | 0.2851222 | 1 | 0.972 | 1 | Neural Tube |
| Cbx3 | 0.002556 | 0.319256464 | 1 | 1 | 1 | Neural Tube |
| Bzw2 | 0.00276 | 0.361075692 | 1 | 1 | 1 | Neural Tube |
| Mcm6 | 0.002791 | 0.319560032 | 1 | 1 | 1 | Neural Tube |
| Ckap2 | 0.002825 | 0.258000075 | 0.93 | 0.917 | 1 | Neural Tube |
| Scd2 | 0.002885 | 0.387864393 | 1 | 1 | 1 | Neural Tube |
| Ncbp1 | 0.003045 | 0.275755561 | 0.977 | 1 | 1 | Neural Tube |
| Hspd1 | 0.003096 | 0.388136808 | 1 | 1 | 1 | Neural Tube |
| Hnrnpk | 0.003143 | 0.348221471 | 1 | 1 | 1 | Neural Tube |
| H2afy2 | 0.003202 | 0.317796038 | 0.977 | 1 | 1 | Neural Tube |
| Hsp90aa1 | 0.00343 | 0.291952434 | 1 | 1 | 1 | Neural Tube |
| Sap30 | 0.003505 | 0.273576151 | 0.837 | 0.75 | 1 | Neural Tube |
| Mcm5 | 0.003701 | 0.311711693 | 1 | 1 | 1 | Neural Tube |
| Ppme1 | 0.003987 | 0.269545968 | 0.977 | 1 | 1 | Neural Tube |
| Polr2m | 0.004269 | 0.265877577 | 1 | 1 | 1 | Neural Tube |
| Hnrnph3 | 0.004386 | 0.283154101 | 1 | 1 | 1 | Neural Tube |
| Abce1 | 0.004524 | 0.318833538 | 1 | 1 | 1 | Neural Tube |
| Xrcc1 | 0.004684 | 0.338103626 | 1 | 1 | 1 | Neural Tube |
| Tpx2 | 0.00501 | 0.419062266 | 1 | 0.972 | 1 | Neural Tube |
| Lin7c | 0.005367 | 0.271278095 | 1 | 1 | 1 | Neural Tube |
| Gm3839 | 0.005587 | 0.313867403 | 1 | 1 | 1 | Neural Tube |
| Hmgb1 | 0.00561 | 0.314613373 | 1 | 1 | 1 | Neural Tube |
| Hnrnpa1 | 0.005684 | 0.332416884 | 1 | 1 | 1 | Neural Tube |
| Pcbp4 | 0.005789 | 0.278846976 | 1 | 1 | 1 | Neural Tube |
| Ddx1 | 0.006408 | 0.34962834 | 1 | 1 | 1 | Neural Tube |
| Adprh | 0.006886 | 0.312448996 | 1 | 0.972 | 1 | Neural Tube |
| Rrm1 | 0.007741 | 0.297956315 | 1 | 1 | 1 | Neural Tube |
| Prps1 | 0.007875 | 0.335551533 | 1 | 0.972 | 1 | Neural Tube |
| Pold3 | 0.00825 | 0.272985316 | 1 | 1 | 1 | Neural Tube |
| Parp2 | 0.009262 | 0.314157573 | 0.953 | 0.972 | 1 | Neural Tube |
| Ptcd3 | 0.009388 | 0.289208694 | 1 | 1 | 1 | Neural Tube |
| Gm7964 | 0.009477 | 0.289537998 | 1 | 1 | 1 | Neural Tube |
| Cer1 | 1.55E-57 | 2.83592159 | 1 | 0.58 | 1.95E-53 | Primitive Streak |
| Tdgf1 | 1.04E-41 | 1.684705721 | 1 | 0.203 | 1.30E-37 | Primitive Streak |
| 2410004P( | 1.13E-39 | 0.373000619 | 0.9 | 0.957 | 1.42E-35 | Primitive Streak |
| Fgf8 | 3.12E-29 | 0.94025276 | 0.8 | 0.145 | 3.93E-25 | Primitive Streak |
| Pou5f1 | 8.98E-23 | 0.806663526 | 1 | 0.449 | 1.13E-18 | Primitive Streak |
| Lhx1 | 2.46E-21 | 1.260169754 | 1 | 0.333 | 3.10E-17 | Primitive Streak |
| Tppp3 | 9.49E-20 | 0.796424336 | 0.6 | 0.725 | 1.19E-15 | Primitive Streak |
| Fgf5 | 2.13E-18 | 1.292315624 | 1 | 0.275 | 2.67E-14 | Primitive Streak |
| Ndrg1 | 1.66E-17 | 0.452578292 | 0.9 | 0.739 | 2.08E-13 | Primitive Streak |
| Eomes | 1.41E-16 | 0.259502134 | 1 | 0.232 | 1.77E-12 | Primitive Streak |
| Rpl24 | 5.92E-16 | 0.541769672 | 1 | 0.986 | 7.45E-12 | Primitive Streak |
| Rps5 | 8.89E-16 | 1.001939405 | 1 | 1 | 1.12E-11 | Primitive Streak |
| Foxj1 | 4.48E-15 | 0.391295316 | 0.8 | 0.217 | 5.64E-11 | Primitive Streak |
| Gm10073 | 8.44E-15 | 1.010935579 | 1 | 1 | 1.06E-10 | Primitive Streak |
| Gnb2l1 | 8.50E-15 | 0.740248344 | 1 | 1 | 1.07E-10 | Primitive Streak |

| | | | | | | |
|---|---|---|---|---|---|---|
| Gm17669 | 1.23E-14 | 0.489602694 | 1 | 0.986 | 1.55E-10 | Primitive Streak |
| Rps21 | 1.51E-14 | 0.984520154 | 1 | 1 | 1.90E-10 | Primitive Streak |
| Gm17511 | 6.01E-14 | 0.655967411 | 1 | 1 | 7.56E-10 | Primitive Streak |
| Pcsk1n | 6.23E-14 | 0.278939891 | 1 | 0.348 | 7.83E-10 | Primitive Streak |
| Gm10076 | 7.03E-14 | 1.000108274 | 1 | 1 | 8.84E-10 | Primitive Streak |
| Gm10126 | 7.52E-14 | 0.734169024 | 1 | 0.986 | 9.45E-10 | Primitive Streak |
| Rps17 | 8.72E-14 | 0.909276572 | 1 | 1 | 1.10E-09 | Primitive Streak |
| Gm10263 | 1.21E-13 | 0.50717827 | 1 | 0.986 | 1.53E-09 | Primitive Streak |
| Rps26-ps1 | 4.17E-13 | 1.025842875 | 1 | 1 | 5.24E-09 | Primitive Streak |
| Nodal | 4.21E-13 | 0.56443185 | 0.7 | 0.058 | 5.29E-09 | Primitive Streak |
| Gpc4 | 4.53E-13 | 0.597284908 | 1 | 0.507 | 5.70E-09 | Primitive Streak |
| Rps12-ps3 | 5.97E-13 | 0.540834219 | 1 | 0.957 | 7.50E-09 | Primitive Streak |
| Gm17541 | 6.30E-13 | 0.377997677 | 1 | 1 | 7.92E-09 | Primitive Streak |
| Rps15a | 7.68E-13 | 0.452091206 | 1 | 1 | 9.66E-09 | Primitive Streak |
| Gm10709 | 1.15E-12 | 0.503216891 | 1 | 1 | 1.45E-08 | Primitive Streak |
| Polr2h | 1.26E-12 | 0.976747854 | 1 | 0.971 | 1.58E-08 | Primitive Streak |
| Gm5160 | 1.89E-12 | 0.570671701 | 1 | 1 | 2.37E-08 | Primitive Streak |
| Vrtn | 2.25E-12 | 0.630748804 | 1 | 0.319 | 2.82E-08 | Primitive Streak |
| Cfc1 | 2.48E-12 | 0.799470953 | 0.8 | 0.174 | 3.11E-08 | Primitive Streak |
| Gm5786 | 3.84E-12 | 0.596718197 | 1 | 1 | 4.83E-08 | Primitive Streak |
| Rps20 | 9.07E-12 | 0.680772922 | 1 | 1 | 1.14E-07 | Primitive Streak |
| Rpl12 | 1.41E-11 | 0.551427216 | 1 | 1 | 1.77E-07 | Primitive Streak |
| Rplp1 | 1.90E-11 | 0.853595537 | 1 | 1 | 2.39E-07 | Primitive Streak |
| Rps15 | 2.29E-11 | 0.8481995 | 1 | 1 | 2.88E-07 | Primitive Streak |
| Chrd | 2.90E-11 | 0.372585341 | 0.9 | 0.319 | 3.65E-07 | Primitive Streak |
| Rpl27 | 3.61E-11 | 0.819610204 | 1 | 1 | 4.54E-07 | Primitive Streak |
| Gm4950 | 4.36E-11 | 0.372763275 | 1 | 0.986 | 5.49E-07 | Primitive Streak |
| Rps29 | 4.76E-11 | 1.004129284 | 1 | 1 | 5.99E-07 | Primitive Streak |
| Rpl41 | 5.22E-11 | 0.625278779 | 1 | 1 | 6.56E-07 | Primitive Streak |
| Eras | 6.48E-11 | 0.695670282 | 1 | 0.13 | 8.15E-07 | Primitive Streak |
| Gm9843 | 7.80E-11 | 0.725281991 | 1 | 1 | 9.81E-07 | Primitive Streak |
| Gm6576 | 9.22E-11 | 0.409537895 | 1 | 1 | 1.16E-06 | Primitive Streak |
| Upp1 | 1.01E-10 | 1.437308204 | 0.9 | 0.754 | 1.27E-06 | Primitive Streak |
| Rps23 | 1.08E-10 | 0.399058426 | 1 | 1 | 1.36E-06 | Primitive Streak |
| Rpl34 | 1.51E-10 | 0.588820328 | 1 | 1 | 1.91E-06 | Primitive Streak |
| Rpl37a | 1.52E-10 | 0.892364113 | 1 | 1 | 1.91E-06 | Primitive Streak |
| Mixl1 | 1.55E-10 | 0.413745408 | 0.8 | 0.145 | 1.95E-06 | Primitive Streak |
| Gm18025 | 1.63E-10 | 0.275027362 | 1 | 1 | 2.05E-06 | Primitive Streak |
| Gm10036 | 1.82E-10 | 0.548692754 | 1 | 1 | 2.29E-06 | Primitive Streak |
| Rpl8 | 1.90E-10 | 0.793269591 | 1 | 1 | 2.39E-06 | Primitive Streak |
| Eci1 | 1.99E-10 | 0.489529091 | 1 | 0.768 | 2.50E-06 | Primitive Streak |
| Rps3 | 2.08E-10 | 0.667991069 | 1 | 1 | 2.61E-06 | Primitive Streak |
| Rps10 | 2.70E-10 | 0.682932739 | 1 | 1 | 3.39E-06 | Primitive Streak |
| Timm13 | 2.82E-10 | 0.738442221 | 1 | 1 | 3.55E-06 | Primitive Streak |
| Rps2 | 3.25E-10 | 0.67544126 | 1 | 1 | 4.09E-06 | Primitive Streak |
| Rpl36-ps3 | 3.33E-10 | 0.610535279 | 1 | 1 | 4.19E-06 | Primitive Streak |
| Rpl9-ps6 | 3.52E-10 | 0.509050073 | 1 | 1 | 4.42E-06 | Primitive Streak |
| Rpl28 | 3.55E-10 | 0.696148982 | 1 | 1 | 4.47E-06 | Primitive Streak |
| Rpl37 | 4.19E-10 | 0.68235528 | 1 | 1 | 5.27E-06 | Primitive Streak |
| Rps27rt | 4.96E-10 | 0.550578334 | 1 | 1 | 6.23E-06 | Primitive Streak |
| Rpl27-ps3 | 5.18E-10 | 0.712178339 | 1 | 0.986 | 6.51E-06 | Primitive Streak |
| Gm8225 | 1.10E-09 | 0.386899347 | 1 | 1 | 1.39E-05 | Primitive Streak |
| Rplp2 | 1.18E-09 | 0.664039426 | 1 | 1 | 1.48E-05 | Primitive Streak |
| Rpl27a | 1.20E-09 | 0.59393714 | 1 | 1 | 1.51E-05 | Primitive Streak |
| Rps12 | 1.26E-09 | 0.786382696 | 1 | 1 | 1.58E-05 | Primitive Streak |
| Eif3k | 1.30E-09 | 0.778796718 | 1 | 1 | 1.64E-05 | Primitive Streak |
| Psmb3 | 1.34E-09 | 0.613920334 | 1 | 1 | 1.69E-05 | Primitive Streak |
| Rps2-ps6 | 1.37E-09 | 0.439723041 | 1 | 1 | 1.73E-05 | Primitive Streak |

| | | | | | | |
|---|---|---|---|---|---|---|
| Rps27 | 1.77E-09 | 0.597459022 | 1 | 1 | 2.23E-05 | Primitive Streak |
| Rpl26 | 2.37E-09 | 0.523217767 | 1 | 1 | 2.98E-05 | Primitive Streak |
| Gm8730 | 2.58E-09 | 0.72284645 | 1 | 1 | 3.24E-05 | Primitive Streak |
| Rps28 | 2.83E-09 | 0.803629862 | 1 | 1 | 3.56E-05 | Primitive Streak |
| Gm12728 | 2.95E-09 | 0.458710758 | 1 | 1 | 3.71E-05 | Primitive Streak |
| Rpl10a | 3.07E-09 | 0.605598009 | 1 | 1 | 3.86E-05 | Primitive Streak |
| Rpl10-ps3 | 3.28E-09 | 0.608435912 | 1 | 1 | 4.13E-05 | Primitive Streak |
| Rps3a1 | 3.34E-09 | 0.35143234 | 1 | 1 | 4.20E-05 | Primitive Streak |
| Sp5 | 3.71E-09 | 0.926073196 | 1 | 0.522 | 4.67E-05 | Primitive Streak |
| Rps13 | 3.83E-09 | 0.777906153 | 1 | 1 | 4.82E-05 | Primitive Streak |
| Dnmt3b | 4.01E-09 | 0.774506648 | 1 | 0.884 | 5.05E-05 | Primitive Streak |
| Rplp0 | 4.47E-09 | 0.848507225 | 1 | 1 | 5.62E-05 | Primitive Streak |
| Foxh1 | 4.91E-09 | 0.817159099 | 1 | 0.768 | 6.17E-05 | Primitive Streak |
| Rps14 | 5.34E-09 | 0.618310413 | 1 | 1 | 6.71E-05 | Primitive Streak |
| Gata1 | 5.61E-09 | 0.260655521 | 0.7 | 0.072 | 7.05E-05 | Primitive Streak |
| Pmaip1 | 6.05E-09 | 0.539655463 | 1 | 0.29 | 7.61E-05 | Primitive Streak |
| Rps18 | 6.62E-09 | 0.447604099 | 1 | 1 | 8.32E-05 | Primitive Streak |
| Rps26 | 6.80E-09 | 0.585561282 | 1 | 1 | 8.55E-05 | Primitive Streak |
| Rpl18 | 7.53E-09 | 0.58451792 | 1 | 1 | 9.47E-05 | Primitive Streak |
| Rpl18a | 1.01E-08 | 0.597084297 | 1 | 1 | 0.000127 | Primitive Streak |
| Trh | 1.08E-08 | 1.785645183 | 1 | 0.754 | 0.000136 | Primitive Streak |
| Pim2 | 1.09E-08 | 0.772541428 | 1 | 0.754 | 0.000137 | Primitive Streak |
| Rpl7a | 1.17E-08 | 0.659620998 | 1 | 1 | 0.000147 | Primitive Streak |
| Rpl29 | 1.35E-08 | 0.443133252 | 1 | 1 | 0.00017 | Primitive Streak |
| Tspo | 1.39E-08 | 0.528777581 | 1 | 0.797 | 0.000174 | Primitive Streak |
| Zbed5 | 1.48E-08 | 0.322558498 | 1 | 1 | 0.000186 | Primitive Streak |
| Rpl31 | 1.63E-08 | 0.583800185 | 1 | 1 | 0.000205 | Primitive Streak |
| Rps11 | 1.77E-08 | 0.888909687 | 1 | 1 | 0.000223 | Primitive Streak |
| Prdx1 | 1.80E-08 | 0.633033907 | 1 | 1 | 0.000226 | Primitive Streak |
| Rpl32 | 1.93E-08 | 0.598091949 | 1 | 1 | 0.000243 | Primitive Streak |
| Pfn1 | 2.02E-08 | 0.492325869 | 1 | 1 | 0.000254 | Primitive Streak |
| Gm11273 | 2.04E-08 | 0.448857505 | 1 | 0.986 | 0.000257 | Primitive Streak |
| Fau | 2.33E-08 | 0.464451172 | 1 | 1 | 0.000293 | Primitive Streak |
| Eif3i | 3.04E-08 | 0.470362191 | 1 | 1 | 0.000382 | Primitive Streak |
| Rpl30 | 3.24E-08 | 0.442834773 | 1 | 1 | 0.000407 | Primitive Streak |
| Mrpl34 | 3.62E-08 | 0.802300659 | 1 | 0.942 | 0.000456 | Primitive Streak |
| Cox8a | 4.41E-08 | 0.623687601 | 1 | 1 | 0.000555 | Primitive Streak |
| Gm10146 | 4.93E-08 | 0.520135142 | 1 | 0.957 | 0.00062 | Primitive Streak |
| Tceb2 | 5.00E-08 | 0.731711432 | 1 | 1 | 0.000629 | Primitive Streak |
| Rpl23a-ps | 5.35E-08 | 0.455546126 | 1 | 1 | 0.000673 | Primitive Streak |
| Isyna1 | 6.05E-08 | 0.669175754 | 1 | 0.957 | 0.000761 | Primitive Streak |
| Rps9 | 6.92E-08 | 0.662414765 | 1 | 1 | 0.00087 | Primitive Streak |
| Ppp1r1a | 7.14E-08 | 0.860696185 | 1 | 0.739 | 0.000898 | Primitive Streak |
| Gm9847 | 7.31E-08 | 0.299823538 | 0.9 | 0.42 | 0.000919 | Primitive Streak |
| Gm9493 | 8.03E-08 | 0.463425521 | 1 | 1 | 0.00101 | Primitive Streak |
| Rpl11 | 8.47E-08 | 0.449657608 | 1 | 1 | 0.001065 | Primitive Streak |
| Rpl36 | 8.54E-08 | 0.624194948 | 1 | 1 | 0.001074 | Primitive Streak |
| H2afz | 8.59E-08 | 0.418000716 | 1 | 1 | 0.001081 | Primitive Streak |
| Rpl10 | 9.19E-08 | 0.526972051 | 1 | 1 | 0.001155 | Primitive Streak |
| Siva1 | 9.87E-08 | 0.637043722 | 1 | 0.971 | 0.001241 | Primitive Streak |
| Rps27a | 1.03E-07 | 0.561834218 | 1 | 1 | 0.0013 | Primitive Streak |
| Rpl39 | 1.11E-07 | 0.349551257 | 1 | 1 | 0.001396 | Primitive Streak |
| Tomm7 | 1.14E-07 | 0.757250566 | 1 | 1 | 0.001433 | Primitive Streak |
| Rps16 | 1.27E-07 | 0.667623245 | 1 | 1 | 0.001592 | Primitive Streak |
| Hint1 | 1.32E-07 | 0.607397425 | 1 | 1 | 0.001656 | Primitive Streak |
| Atp5j2 | 1.34E-07 | 0.322673275 | 1 | 1 | 0.001684 | Primitive Streak |
| Gm10020 | 1.40E-07 | 0.593551675 | 1 | 1 | 0.001759 | Primitive Streak |
| Rpl23 | 1.56E-07 | 0.476285026 | 1 | 1 | 0.001964 | Primitive Streak |

| | | | | | | |
|---|---|---|---|---|---|---|
| Rps18-ps3 | 1.68E-07 | 0.324249772 | 1 | 1 | 0.002109 | Primitive Streak |
| Rps19 | 1.69E-07 | 0.327950628 | 1 | 1 | 0.002131 | Primitive Streak |
| Ppia | 1.80E-07 | 0.408199805 | 1 | 1 | 0.002266 | Primitive Streak |
| Rpl19 | 1.96E-07 | 0.596009414 | 1 | 1 | 0.00247 | Primitive Streak |
| Uqcr11 | 2.15E-07 | 0.636946654 | 1 | 1 | 0.0027 | Primitive Streak |
| Snrpg | 2.61E-07 | 0.747893035 | 1 | 1 | 0.003285 | Primitive Streak |
| Otx2 | 2.67E-07 | 0.518251839 | 1 | 0.304 | 0.003359 | Primitive Streak |
| Rpl7 | 2.69E-07 | 0.475868804 | 1 | 1 | 0.003386 | Primitive Streak |
| Rpl9 | 3.15E-07 | 0.418515057 | 1 | 1 | 0.003962 | Primitive Streak |
| Rpsa | 3.46E-07 | 0.534235232 | 1 | 1 | 0.004353 | Primitive Streak |
| Pycr2 | 3.55E-07 | 0.48322153 | 1 | 1 | 0.004469 | Primitive Streak |
| Atp5l | 3.93E-07 | 0.637911914 | 1 | 0.986 | 0.004946 | Primitive Streak |
| Sssca1 | 4.28E-07 | 0.270398837 | 1 | 0.942 | 0.005388 | Primitive Streak |
| Gm5428 | 4.72E-07 | 0.267266512 | 1 | 1 | 0.00594 | Primitive Streak |
| Myl7 | 4.96E-07 | 0.330914799 | 1 | 0.507 | 0.006237 | Primitive Streak |
| Gm5239 | 5.80E-07 | 0.264236965 | 1 | 1 | 0.007297 | Primitive Streak |
| Cox7a2 | 5.81E-07 | 0.494528747 | 1 | 1 | 0.007311 | Primitive Streak |
| Mif | 5.82E-07 | 0.545639871 | 1 | 1 | 0.007321 | Primitive Streak |
| Rps25 | 5.90E-07 | 0.399983124 | 1 | 1 | 0.007415 | Primitive Streak |
| Ndufb4 | 6.34E-07 | 0.487465949 | 1 | 0.928 | 0.007973 | Primitive Streak |
| Trmt112 | 6.59E-07 | 0.451993845 | 1 | 0.986 | 0.008285 | Primitive Streak |
| Eef1a1 | 8.99E-07 | 0.314025153 | 1 | 1 | 0.011312 | Primitive Streak |
| Usmg5 | 9.56E-07 | 0.576476719 | 1 | 1 | 0.012017 | Primitive Streak |
| Ldha | 1.12E-06 | 0.512214543 | 1 | 1 | 0.014137 | Primitive Streak |
| Ndufa2 | 1.18E-06 | 0.690226577 | 1 | 1 | 0.014901 | Primitive Streak |
| Gm5093 | 1.40E-06 | 0.25361151 | 1 | 1 | 0.0176 | Primitive Streak |
| Gm11808 | 1.48E-06 | 0.485584561 | 1 | 1 | 0.018568 | Primitive Streak |
| Psmb5 | 1.49E-06 | 0.586701516 | 1 | 1 | 0.0187 | Primitive Streak |
| Rps4x | 1.60E-06 | 0.660555353 | 1 | 1 | 0.020142 | Primitive Streak |
| Nop10 | 1.66E-06 | 0.672839026 | 1 | 1 | 0.020938 | Primitive Streak |
| Rps6 | 1.70E-06 | 0.2961254 | 1 | 1 | 0.021381 | Primitive Streak |
| Lefty2 | 2.03E-06 | 0.704597864 | 0.9 | 0.246 | 0.025483 | Primitive Streak |
| Gm8186 | 2.09E-06 | 0.591381853 | 1 | 1 | 0.026333 | Primitive Streak |
| Rpl13a | 2.15E-06 | 0.258731782 | 1 | 1 | 0.027062 | Primitive Streak |
| Rpl15 | 2.22E-06 | 0.514481924 | 1 | 1 | 0.027919 | Primitive Streak |
| N6amt2 | 2.22E-06 | 0.592748766 | 1 | 0.899 | 0.027948 | Primitive Streak |
| Grrp1 | 2.30E-06 | 0.565704533 | 0.9 | 0.362 | 0.028893 | Primitive Streak |
| Ran | 2.43E-06 | 0.641386051 | 1 | 1 | 0.030498 | Primitive Streak |
| Atp5a1 | 2.48E-06 | 0.45808129 | 1 | 1 | 0.031197 | Primitive Streak |
| Psmb6 | 2.50E-06 | 0.583983283 | 1 | 1 | 0.031494 | Primitive Streak |
| Cyc1 | 2.51E-06 | 0.633823443 | 1 | 1 | 0.031512 | Primitive Streak |
| Lars2 | 2.57E-06 | 0.442751485 | 1 | 1 | 0.032375 | Primitive Streak |
| Rpl21 | 2.82E-06 | 0.328335162 | 1 | 1 | 0.035501 | Primitive Streak |
| Rpl13 | 3.23E-06 | 0.399679609 | 1 | 1 | 0.040645 | Primitive Streak |
| Eef2 | 3.43E-06 | 0.387510858 | 1 | 1 | 0.043169 | Primitive Streak |
| Snrpd2 | 3.91E-06 | 0.267806247 | 1 | 1 | 0.049185 | Primitive Streak |
| Rps8 | 4.06E-06 | 0.43436831 | 1 | 1 | 0.051112 | Primitive Streak |
| Ndufa7 | 4.73E-06 | 0.48741953 | 1 | 1 | 0.059439 | Primitive Streak |
| Ifitm2 | 4.73E-06 | 0.402853495 | 1 | 0.986 | 0.059479 | Primitive Streak |
| Rps4l | 4.73E-06 | 0.652573551 | 1 | 0.971 | 0.059494 | Primitive Streak |
| 2700094K1 | 5.25E-06 | 0.575202648 | 1 | 1 | 0.06605 | Primitive Streak |
| Gpx1 | 5.62E-06 | 0.566561126 | 1 | 1 | 0.070656 | Primitive Streak |
| Rpl17 | 5.79E-06 | 0.294141368 | 1 | 1 | 0.07277 | Primitive Streak |
| Asns | 5.94E-06 | 0.530587263 | 1 | 0.87 | 0.074681 | Primitive Streak |
| Gm15013 | 6.12E-06 | 0.29454383 | 1 | 0.986 | 0.076958 | Primitive Streak |
| Ndufa11 | 6.44E-06 | 0.327957133 | 1 | 0.971 | 0.081036 | Primitive Streak |
| Psma6 | 6.54E-06 | 0.451127559 | 1 | 1 | 0.082267 | Primitive Streak |
| Slc25a3 | 7.11E-06 | 0.471004049 | 1 | 1 | 0.089411 | Primitive Streak |

| | | | | | | |
|---|---|---|---|---|---|---|
| Atp6v1f | 7.21E-06 | 0.537881674 | 1 | 1 | 0.090703 | Primitive Streak |
| Car2 | 7.95E-06 | 0.84507983 | 1 | 0.899 | 0.099979 | Primitive Streak |
| Mrps6 | 8.13E-06 | 0.485500498 | 1 | 0.899 | 0.102273 | Primitive Streak |
| Gm9803 | 9.42E-06 | 0.300860273 | 1 | 0.812 | 0.118488 | Primitive Streak |
| Polr2e | 9.80E-06 | 0.433972834 | 1 | 1 | 0.123264 | Primitive Streak |
| Atp5d | 1.10E-05 | 0.479375381 | 1 | 0.986 | 0.137741 | Primitive Streak |
| Rpl13-ps3 | 1.19E-05 | 0.289772789 | 1 | 1 | 0.149656 | Primitive Streak |
| Phc1 | 1.25E-05 | 0.295133117 | 1 | 0.913 | 0.157199 | Primitive Streak |
| Dll3 | 1.39E-05 | 0.378248095 | 1 | 0.406 | 0.175036 | Primitive Streak |
| Lsm4 | 1.44E-05 | 0.479100684 | 1 | 1 | 0.180671 | Primitive Streak |
| Atp5c1 | 1.46E-05 | 0.461510142 | 1 | 1 | 0.184165 | Primitive Streak |
| Zic3 | 1.49E-05 | 0.379916164 | 1 | 0.696 | 0.187212 | Primitive Streak |
| Rpl38 | 1.69E-05 | 0.545427107 | 1 | 1 | 0.213162 | Primitive Streak |
| Eif1 | 1.89E-05 | 0.388869422 | 1 | 1 | 0.238167 | Primitive Streak |
| Polr2c | 1.99E-05 | 0.494904428 | 1 | 1 | 0.249684 | Primitive Streak |
| 2010107E( | 2.02E-05 | 0.34139475 | 1 | 1 | 0.254554 | Primitive Streak |
| Ldhb | 2.12E-05 | 0.616676409 | 1 | 0.913 | 0.26719 | Primitive Streak |
| Uqcrb | 2.36E-05 | 0.333102045 | 1 | 1 | 0.296294 | Primitive Streak |
| Eif3g | 2.46E-05 | 0.494529954 | 1 | 1 | 0.309203 | Primitive Streak |
| Rpl36al | 2.48E-05 | 0.491928567 | 1 | 1 | 0.312432 | Primitive Streak |
| Gsc | 2.50E-05 | 0.493161005 | 0.8 | 0.087 | 0.313795 | Primitive Streak |
| Rpl35a | 2.63E-05 | 0.289821151 | 1 | 1 | 0.331083 | Primitive Streak |
| Snrpe | 2.82E-05 | 0.361858709 | 1 | 0.986 | 0.354518 | Primitive Streak |
| Atp5b | 3.59E-05 | 0.365659852 | 1 | 1 | 0.451622 | Primitive Streak |
| Hesx1 | 3.85E-05 | 0.486305002 | 0.7 | 0.072 | 0.483615 | Primitive Streak |
| Cxx1c | 3.87E-05 | 0.573854464 | 0.8 | 0.377 | 0.486294 | Primitive Streak |
| Ndufc2 | 3.88E-05 | 0.436979123 | 1 | 1 | 0.488485 | Primitive Streak |
| Chchd10 | 3.93E-05 | 0.387873301 | 1 | 0.42 | 0.494612 | Primitive Streak |
| Hmga1 | 4.41E-05 | 0.256847513 | 1 | 1 | 0.554054 | Primitive Streak |
| Hprt | 4.42E-05 | 0.578839078 | 1 | 0.928 | 0.555927 | Primitive Streak |
| 2410015M | 4.50E-05 | 0.423639665 | 1 | 1 | 0.566223 | Primitive Streak |
| Pfdn6 | 4.63E-05 | 0.416338256 | 1 | 1 | 0.58245 | Primitive Streak |
| Ina | 4.97E-05 | 0.267195147 | 0.7 | 0.203 | 0.624732 | Primitive Streak |
| Hcfc1r1 | 5.08E-05 | 0.497114888 | 1 | 0.739 | 0.639387 | Primitive Streak |
| Hmga1-rs | 5.51E-05 | 0.29388381 | 1 | 1 | 0.692589 | Primitive Streak |
| Eif3m | 5.59E-05 | 0.312370793 | 1 | 1 | 0.702504 | Primitive Streak |
| Samd3 | 5.69E-05 | 0.397652549 | 1 | 0.377 | 0.714992 | Primitive Streak |
| Ppp1ca | 5.72E-05 | 0.308526712 | 1 | 1 | 0.719419 | Primitive Streak |
| Btf3 | 5.93E-05 | 0.434854816 | 1 | 1 | 0.745222 | Primitive Streak |
| Cox5b | 5.94E-05 | 0.376460978 | 1 | 1 | 0.747353 | Primitive Streak |
| Ubb | 6.24E-05 | 0.382640147 | 1 | 1 | 0.785288 | Primitive Streak |
| Snrpd1 | 6.28E-05 | 0.327238049 | 1 | 1 | 0.790193 | Primitive Streak |
| H3f3a | 7.40E-05 | 0.299645883 | 1 | 1 | 0.930642 | Primitive Streak |
| Cenpw | 7.65E-05 | 0.378046459 | 1 | 0.971 | 0.962471 | Primitive Streak |
| Tpt1 | 7.93E-05 | 0.308567839 | 1 | 1 | 0.996801 | Primitive Streak |
| Emb | 8.00E-05 | 0.528256081 | 1 | 1 | 1 | Primitive Streak |
| Gm5616 | 8.87E-05 | 0.35160483 | 1 | 0.986 | 1 | Primitive Streak |
| Eif3f | 9.29E-05 | 0.266107035 | 1 | 0.986 | 1 | Primitive Streak |
| Rpl22l1 | 0.000101 | 0.382364537 | 1 | 1 | 1 | Primitive Streak |
| Psmb7 | 0.000102 | 0.261179508 | 1 | 1 | 1 | Primitive Streak |
| Mrpl28 | 0.000111 | 0.444411845 | 1 | 1 | 1 | Primitive Streak |
| Rpl5 | 0.000112 | 0.27891636 | 1 | 1 | 1 | Primitive Streak |
| Prdx2 | 0.000115 | 0.271544923 | 1 | 1 | 1 | Primitive Streak |
| Thop1 | 0.000118 | 0.270933224 | 1 | 1 | 1 | Primitive Streak |
| Rpl22 | 0.000121 | 0.305191596 | 1 | 1 | 1 | Primitive Streak |
| Mdh2 | 0.000131 | 0.296777815 | 1 | 1 | 1 | Primitive Streak |
| Rpl3 | 0.000132 | 0.309007901 | 1 | 1 | 1 | Primitive Streak |
| Psmb4 | 0.000134 | 0.451101251 | 1 | 1 | 1 | Primitive Streak |

| | | | | | | |
|---|---|---|---|---|---|---|
| T | 0.000143 | 0.96164756 | 1 | 0.522 | 1 | Primitive Streak |
| Psma5 | 0.000143 | 0.349931214 | 1 | 1 | 1 | Primitive Streak |
| Bmp7 | 0.000145 | 0.34218587 | 1 | 0.812 | 1 | Primitive Streak |
| Mlf1 | 0.000151 | 0.379432142 | 0.8 | 0.623 | 1 | Primitive Streak |
| Bola2 | 0.000153 | 0.337242147 | 1 | 1 | 1 | Primitive Streak |
| Znhit2 | 0.000154 | 0.250491713 | 1 | 0.783 | 1 | Primitive Streak |
| Csnk2b | 0.000166 | 0.307706304 | 1 | 1 | 1 | Primitive Streak |
| Lsm7 | 0.000169 | 0.414190421 | 1 | 0.971 | 1 | Primitive Streak |
| Ndufs7 | 0.000171 | 0.365174328 | 1 | 0.986 | 1 | Primitive Streak |
| Eif5a | 0.000172 | 0.31020828 | 1 | 1 | 1 | Primitive Streak |
| Cops6 | 0.000184 | 0.366866805 | 1 | 1 | 1 | Primitive Streak |
| Mrps28 | 0.000185 | 0.338161432 | 1 | 0.986 | 1 | Primitive Streak |
| Polr1c | 0.000191 | 0.308073858 | 1 | 1 | 1 | Primitive Streak |
| Cox5a | 0.000196 | 0.434487721 | 1 | 1 | 1 | Primitive Streak |
| Pomp | 0.000197 | 0.461814713 | 1 | 1 | 1 | Primitive Streak |
| Pgp | 0.0002 | 0.415065038 | 1 | 0.855 | 1 | Primitive Streak |
| Mrpl20 | 0.000203 | 0.397820866 | 1 | 1 | 1 | Primitive Streak |
| Polr2k | 0.000206 | 0.298278792 | 1 | 0.971 | 1 | Primitive Streak |
| Slc25a5 | 0.000209 | 0.581788178 | 1 | 1 | 1 | Primitive Streak |
| Pam16 | 0.000217 | 0.349539961 | 1 | 0.913 | 1 | Primitive Streak |
| Cct7 | 0.000222 | 0.27509994 | 1 | 1 | 1 | Primitive Streak |
| Uqcrq | 0.000226 | 0.314976752 | 1 | 0.986 | 1 | Primitive Streak |
| Wdr55 | 0.000244 | 0.279130861 | 0.9 | 0.942 | 1 | Primitive Streak |
| Sec61b | 0.000244 | 0.310299481 | 1 | 0.986 | 1 | Primitive Streak |
| Cyp26a1 | 0.000247 | 0.73798153 | 1 | 0.362 | 1 | Primitive Streak |
| Penk | 0.000252 | 0.622119785 | 0.7 | 0.348 | 1 | Primitive Streak |
| Plp2 | 0.000255 | 0.332192051 | 1 | 0.942 | 1 | Primitive Streak |
| Gng5 | 0.000263 | 0.366805009 | 1 | 1 | 1 | Primitive Streak |
| Psmb2 | 0.000266 | 0.282809571 | 1 | 1 | 1 | Primitive Streak |
| Arl6 | 0.000269 | 0.30165722 | 0.9 | 0.899 | 1 | Primitive Streak |
| Dctpp1 | 0.000281 | 0.606740083 | 1 | 0.971 | 1 | Primitive Streak |
| Noto | 0.000286 | 0.965060111 | 0.6 | 0.058 | 1 | Primitive Streak |
| Psmb10 | 0.000292 | 0.334480595 | 1 | 0.739 | 1 | Primitive Streak |
| Cox7b | 0.000299 | 0.467521691 | 1 | 1 | 1 | Primitive Streak |
| Pcbp1 | 0.000317 | 0.457968734 | 1 | 1 | 1 | Primitive Streak |
| Ahcy | 0.000321 | 0.408091621 | 1 | 1 | 1 | Primitive Streak |
| Polr2g | 0.000323 | 0.430048608 | 1 | 0.971 | 1 | Primitive Streak |
| Aurkaip1 | 0.000335 | 0.336051539 | 1 | 0.986 | 1 | Primitive Streak |
| Psmd8 | 0.000357 | 0.322468173 | 1 | 0.971 | 1 | Primitive Streak |
| Zscan10 | 0.000364 | 0.274570337 | 1 | 0.478 | 1 | Primitive Streak |
| Cks1b | 0.00037 | 0.304294493 | 1 | 1 | 1 | Primitive Streak |
| Lefty1 | 0.000372 | 0.400642507 | 0.6 | 0.072 | 1 | Primitive Streak |
| Dohh | 0.000376 | 0.31987697 | 1 | 0.986 | 1 | Primitive Streak |
| Med31 | 0.000388 | 0.26283288 | 1 | 0.913 | 1 | Primitive Streak |
| Gm561 | 0.000396 | 0.315682026 | 0.8 | 0.58 | 1 | Primitive Streak |
| Supt4a | 0.000419 | 0.520343381 | 1 | 0.971 | 1 | Primitive Streak |
| Psma7 | 0.000419 | 0.27317904 | 1 | 1 | 1 | Primitive Streak |
| Psmb1 | 0.000427 | 0.497032492 | 1 | 1 | 1 | Primitive Streak |
| Psmc5 | 0.000456 | 0.341113112 | 1 | 1 | 1 | Primitive Streak |
| Snrpn | 0.000465 | 0.537746987 | 1 | 0.986 | 1 | Primitive Streak |
| Ndufs8 | 0.000471 | 0.443244916 | 1 | 0.957 | 1 | Primitive Streak |
| Uqcc2 | 0.000495 | 0.340831519 | 1 | 0.957 | 1 | Primitive Streak |
| Atp5k | 0.000508 | 0.383879828 | 1 | 0.986 | 1 | Primitive Streak |
| Trappc4 | 0.000529 | 0.367987975 | 1 | 0.986 | 1 | Primitive Streak |
| Cox4i1 | 0.000548 | 0.320322126 | 1 | 1 | 1 | Primitive Streak |
| Atp5o | 0.000562 | 0.321563322 | 1 | 1 | 1 | Primitive Streak |
| Chchd1 | 0.000564 | 0.499833057 | 1 | 1 | 1 | Primitive Streak |
| Stoml2 | 0.000591 | 0.331217506 | 1 | 0.971 | 1 | Primitive Streak |

| | | | | | | |
|---|---|---|---|---|---|---|
| Mrpl42 | 0.000664 | 0.394980332 | 1 | 1 | 1 | Primitive Streak |
| Spint2 | 0.000697 | 0.362991237 | 1 | 0.971 | 1 | Primitive Streak |
| Perp | 0.000705 | 0.469150395 | 0.9 | 0.348 | 1 | Primitive Streak |
| Epcam | 0.000721 | 0.27495789 | 1 | 1 | 1 | Primitive Streak |
| Tpgs1 | 0.000728 | 0.274051823 | 1 | 0.87 | 1 | Primitive Streak |
| Oat | 0.000755 | 0.353582127 | 1 | 0.942 | 1 | Primitive Streak |
| Sec61g | 0.000779 | 0.339975076 | 1 | 1 | 1 | Primitive Streak |
| Rpp25l | 0.000782 | 0.301443115 | 1 | 0.899 | 1 | Primitive Streak |
| Sdhd | 0.00082 | 0.343292186 | 1 | 1 | 1 | Primitive Streak |
| Nhp2 | 0.000854 | 0.649033776 | 1 | 1 | 1 | Primitive Streak |
| Ruvbl2 | 0.001015 | 0.327600767 | 1 | 1 | 1 | Primitive Streak |
| Alkbh7 | 0.001025 | 0.257662764 | 1 | 0.783 | 1 | Primitive Streak |
| Ppp4c | 0.001034 | 0.476176635 | 1 | 0.986 | 1 | Primitive Streak |
| Polr2l | 0.001086 | 0.352294533 | 1 | 1 | 1 | Primitive Streak |
| Tmem256 | 0.001158 | 0.330666371 | 1 | 0.957 | 1 | Primitive Streak |
| Clpp | 0.001207 | 0.3869569 | 1 | 0.942 | 1 | Primitive Streak |
| Uqcr10 | 0.001271 | 0.277262558 | 1 | 1 | 1 | Primitive Streak |
| Mrps12 | 0.001292 | 0.3722314 | 1 | 0.986 | 1 | Primitive Streak |
| Gm4737 | 0.001644 | 0.303364799 | 1 | 0.986 | 1 | Primitive Streak |
| Imp3 | 0.001684 | 0.470604391 | 1 | 0.971 | 1 | Primitive Streak |
| Timm23 | 0.001861 | 0.355958045 | 1 | 1 | 1 | Primitive Streak |
| Capsl | 0.001907 | 0.268078344 | 0.7 | 0.188 | 1 | Primitive Streak |
| Ndufa6 | 0.001916 | 0.485298004 | 1 | 0.971 | 1 | Primitive Streak |
| Bambi | 0.002057 | 0.292800908 | 0.9 | 0.696 | 1 | Primitive Streak |
| Sox17 | 0.002097 | 0.513532968 | 0.7 | 0.522 | 1 | Primitive Streak |
| Cox7c | 0.002121 | 0.291444592 | 1 | 1 | 1 | Primitive Streak |
| Mrpl55 | 0.002167 | 0.40115786 | 1 | 0.971 | 1 | Primitive Streak |
| D8Ertd738 | 0.002196 | 0.341495026 | 1 | 1 | 1 | Primitive Streak |
| Dkk1 | 0.002345 | 0.46950935 | 0.9 | 0.783 | 1 | Primitive Streak |
| Atp5g1 | 0.002371 | 0.254744567 | 1 | 1 | 1 | Primitive Streak |
| C1qbp | 0.002372 | 0.395337404 | 1 | 0.986 | 1 | Primitive Streak |
| Ndufa12 | 0.002387 | 0.311829314 | 1 | 0.986 | 1 | Primitive Streak |
| Polr2i | 0.002475 | 0.315040106 | 1 | 0.928 | 1 | Primitive Streak |
| Phf5a | 0.002497 | 0.439863161 | 1 | 0.986 | 1 | Primitive Streak |
| Tubb4b | 0.002577 | 0.479041586 | 1 | 1 | 1 | Primitive Streak |
| Uqcrfs1 | 0.002633 | 0.349002731 | 1 | 0.986 | 1 | Primitive Streak |
| Rps19bp1 | 0.002647 | 0.364874074 | 1 | 0.928 | 1 | Primitive Streak |
| Fkbp11 | 0.002703 | 0.551207443 | 1 | 0.71 | 1 | Primitive Streak |
| Myeov2 | 0.002708 | 0.342515406 | 1 | 0.971 | 1 | Primitive Streak |
| Mrps21 | 0.002757 | 0.361350291 | 1 | 0.986 | 1 | Primitive Streak |
| Phb | 0.00277 | 0.325678611 | 1 | 1 | 1 | Primitive Streak |
| Bax | 0.00283 | 0.318711451 | 1 | 0.986 | 1 | Primitive Streak |
| Urod | 0.002979 | 0.293476742 | 1 | 0.971 | 1 | Primitive Streak |
| Cstb | 0.003091 | 0.34511388 | 1 | 0.971 | 1 | Primitive Streak |
| Rab1b | 0.00312 | 0.263409543 | 1 | 1 | 1 | Primitive Streak |
| Cst3 | 0.003203 | 0.30139244 | 1 | 1 | 1 | Primitive Streak |
| Ap2s1 | 0.003547 | 0.351502349 | 1 | 0.971 | 1 | Primitive Streak |
| Ost4 | 0.00357 | 0.371703202 | 1 | 0.957 | 1 | Primitive Streak |
| Gstp2 | 0.003599 | 0.27341972 | 1 | 1 | 1 | Primitive Streak |
| Uqcrh | 0.003918 | 0.342693121 | 1 | 1 | 1 | Primitive Streak |
| Gm10273 | 0.003965 | 0.267791925 | 1 | 1 | 1 | Primitive Streak |
| Mrpl11 | 0.004035 | 0.274336423 | 1 | 1 | 1 | Primitive Streak |
| Dpcd | 0.00427 | 0.254454451 | 1 | 0.884 | 1 | Primitive Streak |
| Naa38 | 0.004395 | 0.274717431 | 1 | 0.913 | 1 | Primitive Streak |
| Mid1ip1 | 0.004417 | 0.281505557 | 0.9 | 0.565 | 1 | Primitive Streak |
| Eef1d | 0.004466 | 0.32387623 | 1 | 1 | 1 | Primitive Streak |
| Noc4l | 0.004656 | 0.299725798 | 1 | 0.971 | 1 | Primitive Streak |
| Gart | 0.004852 | 0.261053009 | 1 | 1 | 1 | Primitive Streak |

| | | | | | | |
|---|---|---|---|---|---|---|
| Nme1 | 0.004928 | 0.25223185 | 1 | 1 | 1 | Primitive Streak |
| Gstp1 | 0.005685 | 0.349356755 | 1 | 1 | 1 | Primitive Streak |
| Pop4 | 0.005724 | 0.271654038 | 1 | 0.971 | 1 | Primitive Streak |
| Emg1 | 0.005736 | 0.374533198 | 1 | 1 | 1 | Primitive Streak |
| Dusp4 | 0.005776 | 0.337957891 | 1 | 0.739 | 1 | Primitive Streak |
| Tmem107 | 0.005997 | 0.360836099 | 0.9 | 0.841 | 1 | Primitive Streak |
| Bphl | 0.006005 | 0.259348296 | 0.9 | 0.667 | 1 | Primitive Streak |
| Timm8b | 0.006091 | 0.454933502 | 1 | 1 | 1 | Primitive Streak |
| Epha1 | 0.006214 | 0.254788997 | 0.8 | 0.493 | 1 | Primitive Streak |
| Plrg1 | 0.006936 | 0.263322478 | 1 | 0.986 | 1 | Primitive Streak |
| Dynll1 | 0.007338 | 0.27039057 | 1 | 1 | 1 | Primitive Streak |
| Snai1 | 0.007428 | 0.357381147 | 1 | 0.565 | 1 | Primitive Streak |
| Coa3 | 0.00766 | 0.288415707 | 1 | 0.986 | 1 | Primitive Streak |
| Sac3d1 | 0.007755 | 0.277733579 | 1 | 0.884 | 1 | Primitive Streak |
| Rps27l | 0.007844 | 0.496986057 | 1 | 1 | 1 | Primitive Streak |
| Mrpl41 | 0.007895 | 0.270487897 | 1 | 0.928 | 1 | Primitive Streak |
| Hhex | 0.008317 | 0.481719911 | 0.9 | 0.406 | 1 | Primitive Streak |
| Dnajc15 | 0.008599 | 0.423840517 | 1 | 0.913 | 1 | Primitive Streak |
| Mrpl46 | 0.008839 | 0.26984916 | 1 | 0.971 | 1 | Primitive Streak |
| Cct2 | 0.008854 | 0.333166833 | 1 | 1 | 1 | Primitive Streak |
| Nup37 | 0.009118 | 0.403132673 | 1 | 0.928 | 1 | Primitive Streak |
| Mrpl54 | 0.009669 | 0.311415996 | 1 | 0.971 | 1 | Primitive Streak |
| Cubn | 2.70E-69 | 1.962531349 | 1 | 0.597 | 3.39E-65 | Visceral Endoderm |
| Nrk | 6.70E-69 | 0.437025658 | 1 | 0.861 | 8.43E-65 | Visceral Endoderm |
| Abcc2 | 1.21E-58 | 0.518659244 | 1 | 0.264 | 1.53E-54 | Visceral Endoderm |
| Mlxipl | 1.95E-55 | 0.744061366 | 0.857 | 0.708 | 2.46E-51 | Visceral Endoderm |
| Prodh2 | 1.78E-48 | 0.907193502 | 1 | 0.972 | 2.24E-44 | Visceral Endoderm |
| Slc23a1 | 3.38E-46 | 0.974235745 | 1 | 1 | 4.25E-42 | Visceral Endoderm |
| Tpcn1 | 6.27E-43 | 0.871743877 | 1 | 0.778 | 7.88E-39 | Visceral Endoderm |
| Npc1 | 3.64E-38 | 0.564892045 | 1 | 1 | 4.58E-34 | Visceral Endoderm |
| Swt1 | 2.22E-37 | 0.749909396 | 1 | 1 | 2.80E-33 | Visceral Endoderm |
| Dmxl1 | 5.50E-36 | 0.33761215 | 1 | 1 | 6.91E-32 | Visceral Endoderm |
| Soat2 | 2.40E-35 | 1.325083325 | 1 | 0.944 | 3.02E-31 | Visceral Endoderm |
| Tcte2 | 6.98E-35 | 0.452707894 | 1 | 0.903 | 8.78E-31 | Visceral Endoderm |
| Ube2l6 | 3.48E-34 | 1.366846201 | 1 | 0.861 | 4.37E-30 | Visceral Endoderm |
| Sgk3 | 1.03E-32 | 0.298960762 | 1 | 0.903 | 1.30E-28 | Visceral Endoderm |
| Foxo4 | 1.32E-31 | 0.362271214 | 0.857 | 0.361 | 1.66E-27 | Visceral Endoderm |
| Rita1 | 2.08E-31 | 1.61583155 | 0.714 | 0.681 | 2.62E-27 | Visceral Endoderm |
| Rtl1 | 2.41E-31 | 0.702181141 | 1 | 0.5 | 3.03E-27 | Visceral Endoderm |
| Ptpn23 | 3.45E-30 | 0.579580814 | 0.857 | 0.722 | 4.34E-26 | Visceral Endoderm |
| Slc13a4 | 5.05E-30 | 1.563357924 | 1 | 0.764 | 6.35E-26 | Visceral Endoderm |
| Jade1 | 7.65E-30 | 1.201560024 | 1 | 0.889 | 9.63E-26 | Visceral Endoderm |
| Slc7a8 | 8.22E-30 | 1.469950558 | 1 | 0.514 | 1.03E-25 | Visceral Endoderm |
| Mfsd12 | 3.80E-29 | 0.928268247 | 1 | 0.528 | 4.78E-25 | Visceral Endoderm |
| Rpia | 1.61E-28 | 0.252363614 | 1 | 0.972 | 2.02E-24 | Visceral Endoderm |
| Fnip2 | 2.48E-28 | 0.256147482 | 1 | 0.972 | 3.12E-24 | Visceral Endoderm |
| Dpm1 | 3.72E-28 | 0.252165161 | 1 | 0.972 | 4.68E-24 | Visceral Endoderm |
| Trpm2 | 4.79E-28 | 0.368627785 | 1 | 1 | 6.02E-24 | Visceral Endoderm |
| Acy1 | 5.02E-28 | 0.967108842 | 1 | 0.861 | 6.31E-24 | Visceral Endoderm |
| Apoc2 | 2.02E-27 | 3.434650059 | 1 | 0.278 | 2.54E-23 | Visceral Endoderm |
| Abcd4 | 8.28E-27 | 1.171027109 | 1 | 0.861 | 1.04E-22 | Visceral Endoderm |
| Iqsec1 | 1.08E-26 | 0.259062046 | 0.857 | 0.944 | 1.36E-22 | Visceral Endoderm |
| Sfmbt2 | 9.61E-26 | 0.914848838 | 1 | 0.292 | 1.21E-21 | Visceral Endoderm |
| Morc4 | 1.20E-25 | 0.721166847 | 1 | 0.944 | 1.51E-21 | Visceral Endoderm |
| Ctsh | 1.83E-25 | 1.349614228 | 1 | 0.556 | 2.30E-21 | Visceral Endoderm |
| Evi5 | 4.51E-25 | 0.423177451 | 1 | 0.611 | 5.68E-21 | Visceral Endoderm |
| Synj1 | 4.73E-25 | 0.655431898 | 1 | 0.569 | 5.94E-21 | Visceral Endoderm |
| Dennd6b | 4.97E-25 | 0.596800172 | 0.857 | 0.583 | 6.25E-21 | Visceral Endoderm |

| | | | | | | |
|---|---|---|---|---|---|---|
| Npl | 9.02E-25 | 1.184165351 | 1 | 0.847 | 1.13E-20 | Visceral Endoderm |
| Col27a1 | 1.26E-24 | 0.252968328 | 1 | 0.778 | 1.58E-20 | Visceral Endoderm |
| C77080 | 1.81E-24 | 0.28779641 | 1 | 0.528 | 2.28E-20 | Visceral Endoderm |
| Sec24a | 3.12E-24 | 0.570788033 | 1 | 1 | 3.92E-20 | Visceral Endoderm |
| Myo6 | 4.40E-24 | 0.680041316 | 1 | 0.861 | 5.54E-20 | Visceral Endoderm |
| Mt1 | 5.74E-24 | 2.937105755 | 1 | 0.764 | 7.22E-20 | Visceral Endoderm |
| Gcat | 6.12E-24 | 0.950523531 | 1 | 0.958 | 7.69E-20 | Visceral Endoderm |
| Trim6 | 6.96E-24 | 0.588088022 | 1 | 1 | 8.75E-20 | Visceral Endoderm |
| Neu1 | 8.95E-24 | 1.069391956 | 1 | 0.875 | 1.13E-19 | Visceral Endoderm |
| Gga3 | 1.33E-23 | 0.361967338 | 1 | 1 | 1.68E-19 | Visceral Endoderm |
| Sfi1 | 1.83E-23 | 0.491288143 | 1 | 0.986 | 2.31E-19 | Visceral Endoderm |
| Myo5b | 4.99E-23 | 0.256316229 | 1 | 0.583 | 6.27E-19 | Visceral Endoderm |
| Ppcdc | 5.47E-23 | 0.688368481 | 0.571 | 0.639 | 6.89E-19 | Visceral Endoderm |
| Slc43a2 | 6.43E-23 | 0.361111187 | 1 | 0.528 | 8.08E-19 | Visceral Endoderm |
| Apoa1 | 9.32E-23 | 3.46344483 | 1 | 0.722 | 1.17E-18 | Visceral Endoderm |
| Amn | 1.05E-22 | 2.932966125 | 1 | 0.403 | 1.32E-18 | Visceral Endoderm |
| Pcyt2 | 1.32E-22 | 1.310207142 | 1 | 0.986 | 1.66E-18 | Visceral Endoderm |
| Car7 | 1.39E-22 | 1.688170112 | 1 | 0.306 | 1.74E-18 | Visceral Endoderm |
| Snx8 | 1.80E-22 | 1.046110064 | 1 | 0.847 | 2.27E-18 | Visceral Endoderm |
| Acox1 | 2.03E-22 | 0.659733623 | 1 | 0.653 | 2.55E-18 | Visceral Endoderm |
| Slc2a2 | 2.29E-22 | 0.714117891 | 1 | 0.167 | 2.88E-18 | Visceral Endoderm |
| Stat3 | 2.73E-22 | 0.307939832 | 1 | 1 | 3.44E-18 | Visceral Endoderm |
| Gm5424 | 3.37E-22 | 0.820509993 | 1 | 0.222 | 4.24E-18 | Visceral Endoderm |
| Clk2 | 5.90E-22 | 0.411588927 | 0.857 | 1 | 7.42E-18 | Visceral Endoderm |
| Gramd1b | 6.71E-22 | 0.474519932 | 1 | 0.556 | 8.44E-18 | Visceral Endoderm |
| Apoe | 7.72E-22 | 3.195591509 | 1 | 0.986 | 9.71E-18 | Visceral Endoderm |
| Sgpp2 | 1.78E-21 | 0.459976949 | 1 | 0.528 | 2.24E-17 | Visceral Endoderm |
| Setd4 | 1.87E-21 | 0.332878597 | 1 | 1 | 2.36E-17 | Visceral Endoderm |
| Foxk1 | 2.40E-21 | 0.259292062 | 0.857 | 0.736 | 3.01E-17 | Visceral Endoderm |
| Slc39a5 | 2.85E-21 | 1.826405961 | 1 | 0.542 | 3.58E-17 | Visceral Endoderm |
| Med12 | 2.86E-21 | 0.436886164 | 1 | 0.875 | 3.59E-17 | Visceral Endoderm |
| Polq | 3.09E-21 | 0.364079869 | 1 | 0.833 | 3.88E-17 | Visceral Endoderm |
| Prdm2 | 4.59E-21 | 0.472078945 | 1 | 0.819 | 5.77E-17 | Visceral Endoderm |
| Trak2 | 5.38E-21 | 0.520552895 | 1 | 0.583 | 6.77E-17 | Visceral Endoderm |
| Adck5 | 5.71E-21 | 0.609529947 | 1 | 0.819 | 7.18E-17 | Visceral Endoderm |
| Atp6v0a1 | 5.74E-21 | 0.572604599 | 1 | 0.972 | 7.22E-17 | Visceral Endoderm |
| Foxp4 | 6.77E-21 | 0.269370379 | 0.857 | 0.778 | 8.52E-17 | Visceral Endoderm |
| Ass1 | 7.78E-21 | 1.43498294 | 1 | 0.264 | 9.79E-17 | Visceral Endoderm |
| Dvl1 | 9.91E-21 | 0.420826333 | 1 | 0.611 | 1.25E-16 | Visceral Endoderm |
| Slc25a10 | 1.29E-20 | 0.951815398 | 1 | 0.806 | 1.62E-16 | Visceral Endoderm |
| Rbm47 | 1.73E-20 | 0.731850309 | 1 | 0.458 | 2.18E-16 | Visceral Endoderm |
| Neil1 | 2.44E-20 | 0.532655309 | 0.714 | 0.722 | 3.07E-16 | Visceral Endoderm |
| Furin | 3.11E-20 | 0.260161219 | 1 | 1 | 3.91E-16 | Visceral Endoderm |
| Lgmn | 3.46E-20 | 1.862900501 | 1 | 0.972 | 4.35E-16 | Visceral Endoderm |
| Tmem245 | 5.03E-20 | 0.324469575 | 1 | 1 | 6.33E-16 | Visceral Endoderm |
| Irf2bp2 | 9.04E-20 | 0.433575389 | 1 | 0.764 | 1.14E-15 | Visceral Endoderm |
| Rrbp1 | 1.04E-19 | 0.703866411 | 1 | 0.986 | 1.30E-15 | Visceral Endoderm |
| Ctsa | 1.44E-19 | 1.344852359 | 1 | 1 | 1.81E-15 | Visceral Endoderm |
| Apob | 2.43E-19 | 1.327503145 | 1 | 0.625 | 3.06E-15 | Visceral Endoderm |
| B4galt5 | 2.79E-19 | 0.394819917 | 0.857 | 0.681 | 3.51E-15 | Visceral Endoderm |
| Dhrs11 | 3.56E-19 | 0.911661284 | 1 | 0.375 | 4.48E-15 | Visceral Endoderm |
| Pqlc2 | 3.68E-19 | 0.680167731 | 1 | 0.653 | 4.63E-15 | Visceral Endoderm |
| Gipc2 | 3.98E-19 | 0.496150333 | 1 | 0.833 | 5.01E-15 | Visceral Endoderm |
| Gldc | 4.21E-19 | 0.652509097 | 1 | 0.611 | 5.29E-15 | Visceral Endoderm |
| Lpin2 | 9.61E-19 | 0.343099735 | 1 | 0.806 | 1.21E-14 | Visceral Endoderm |
| 3110002H | 9.65E-19 | 0.563354283 | 1 | 1 | 1.21E-14 | Visceral Endoderm |
| Dab2 | 2.57E-18 | 1.798213968 | 1 | 1 | 3.23E-14 | Visceral Endoderm |
| Gga2 | 2.66E-18 | 0.891261901 | 1 | 1 | 3.34E-14 | Visceral Endoderm |

| | | | | | | |
|---|---|---|---|---|---|---|
| Nmrk1 | 4.94E-18 | 0.297477909 | 1 | 0.986 | 6.21E-14 | Visceral Endoderm |
| Zfp692 | 5.91E-18 | 0.895144932 | 1 | 0.861 | 7.43E-14 | Visceral Endoderm |
| Gm21887 | 6.02E-18 | 2.364999621 | 1 | 1 | 7.57E-14 | Visceral Endoderm |
| Atp7b | 6.22E-18 | 0.590077062 | 1 | 0.708 | 7.83E-14 | Visceral Endoderm |
| Ascc3 | 6.33E-18 | 0.337713812 | 0.857 | 0.875 | 7.96E-14 | Visceral Endoderm |
| Triml2 | 6.36E-18 | 0.853725349 | 1 | 0.097 | 8.00E-14 | Visceral Endoderm |
| Cul7 | 1.10E-17 | 0.518791142 | 1 | 0.861 | 1.38E-13 | Visceral Endoderm |
| P2rx4 | 1.20E-17 | 0.279434362 | 1 | 1 | 1.51E-13 | Visceral Endoderm |
| Cited1 | 1.59E-17 | 1.778450251 | 1 | 0.528 | 2.00E-13 | Visceral Endoderm |
| Dpp7 | 2.12E-17 | 1.060890976 | 1 | 0.792 | 2.66E-13 | Visceral Endoderm |
| Nup153 | 2.63E-17 | 0.602559424 | 0.857 | 1 | 3.31E-13 | Visceral Endoderm |
| Fxyd4 | 2.86E-17 | 0.510133677 | 0.429 | 0.514 | 3.60E-13 | Visceral Endoderm |
| Sccpdh | 2.98E-17 | 0.583832264 | 1 | 0.931 | 3.75E-13 | Visceral Endoderm |
| Agap3 | 3.42E-17 | 0.334732279 | 0.714 | 0.736 | 4.30E-13 | Visceral Endoderm |
| Aqp3 | 6.20E-17 | 1.356355832 | 1 | 0.375 | 7.80E-13 | Visceral Endoderm |
| Cluh | 6.60E-17 | 0.622354621 | 0.857 | 0.889 | 8.29E-13 | Visceral Endoderm |
| Hsd17b11 | 6.69E-17 | 0.373474527 | 0.857 | 0.361 | 8.42E-13 | Visceral Endoderm |
| Slc38a9 | 7.93E-17 | 0.41878525 | 0.857 | 0.653 | 9.97E-13 | Visceral Endoderm |
| Creb3l3 | 9.40E-17 | 1.094637392 | 1 | 0.097 | 1.18E-12 | Visceral Endoderm |
| Ttc19 | 1.14E-16 | 0.295068503 | 0.857 | 0.847 | 1.43E-12 | Visceral Endoderm |
| Nktr | 1.32E-16 | 0.264706986 | 1 | 0.986 | 1.65E-12 | Visceral Endoderm |
| Nlrc3 | 2.14E-16 | 0.250473204 | 1 | 1 | 2.69E-12 | Visceral Endoderm |
| Ctsb | 2.48E-16 | 1.373244107 | 1 | 1 | 3.12E-12 | Visceral Endoderm |
| Tmem150a | 3.22E-16 | 1.402503041 | 0.857 | 0.458 | 4.05E-12 | Visceral Endoderm |
| Amot | 5.00E-16 | 1.073064971 | 1 | 0.944 | 6.29E-12 | Visceral Endoderm |
| Dcbld1 | 6.06E-16 | 0.33677893 | 1 | 0.903 | 7.62E-12 | Visceral Endoderm |
| Erdr1 | 9.88E-16 | 2.521773526 | 1 | 1 | 1.24E-11 | Visceral Endoderm |
| Mospd2 | 1.15E-15 | 0.262384856 | 0.571 | 0.444 | 1.45E-11 | Visceral Endoderm |
| Reep6 | 1.21E-15 | 1.635022324 | 1 | 0.514 | 1.53E-11 | Visceral Endoderm |
| Kifc2 | 1.81E-15 | 0.461216657 | 0.714 | 0.708 | 2.27E-11 | Visceral Endoderm |
| 2610528J1 | 2.27E-15 | 1.102970308 | 1 | 0.264 | 2.86E-11 | Visceral Endoderm |
| Nprl2 | 3.84E-15 | 0.653733277 | 0.857 | 0.847 | 4.83E-11 | Visceral Endoderm |
| Cdhr2 | 3.97E-15 | 0.571870337 | 1 | 0.097 | 4.99E-11 | Visceral Endoderm |
| Plekhg5 | 4.41E-15 | 0.301799572 | 1 | 0.514 | 5.55E-11 | Visceral Endoderm |
| Pitpna | 5.50E-15 | 0.41994227 | 0.571 | 0.5 | 6.91E-11 | Visceral Endoderm |
| Aass | 5.67E-15 | 0.54830332 | 1 | 0.236 | 7.13E-11 | Visceral Endoderm |
| G730046D | 5.95E-15 | 0.288610544 | 0.714 | 0.736 | 7.48E-11 | Visceral Endoderm |
| Ap1ar | 6.02E-15 | 0.358524555 | 1 | 0.875 | 7.58E-11 | Visceral Endoderm |
| Ndel1 | 6.49E-15 | 1.162928088 | 1 | 0.986 | 8.16E-11 | Visceral Endoderm |
| Polg | 6.51E-15 | 0.613414577 | 1 | 0.917 | 8.18E-11 | Visceral Endoderm |
| Akap1 | 8.29E-15 | 0.498052344 | 1 | 0.903 | 1.04E-10 | Visceral Endoderm |
| Gna12 | 1.16E-14 | 0.303366932 | 0.857 | 0.708 | 1.46E-10 | Visceral Endoderm |
| Crls1 | 1.23E-14 | 0.262394114 | 0.429 | 0.653 | 1.54E-10 | Visceral Endoderm |
| Ano9 | 1.60E-14 | 0.775379115 | 0.857 | 0.278 | 2.01E-10 | Visceral Endoderm |
| Optn | 1.81E-14 | 0.75191262 | 0.857 | 0.194 | 2.28E-10 | Visceral Endoderm |
| Ppm1a | 1.92E-14 | 0.409561388 | 0.857 | 0.792 | 2.41E-10 | Visceral Endoderm |
| Leng8 | 2.42E-14 | 0.382072719 | 0.857 | 0.917 | 3.04E-10 | Visceral Endoderm |
| Pxn | 2.45E-14 | 0.331402577 | 1 | 0.861 | 3.09E-10 | Visceral Endoderm |
| Sema4b | 2.70E-14 | 0.268548666 | 1 | 0.986 | 3.39E-10 | Visceral Endoderm |
| Uap1l1 | 2.91E-14 | 0.854489425 | 1 | 0.444 | 3.66E-10 | Visceral Endoderm |
| Sgk1 | 3.11E-14 | 0.761024177 | 1 | 0.819 | 3.90E-10 | Visceral Endoderm |
| Apoa4 | 3.16E-14 | 1.181905743 | 1 | 0.125 | 3.98E-10 | Visceral Endoderm |
| Lrp2 | 3.20E-14 | 0.631878181 | 1 | 0.903 | 4.03E-10 | Visceral Endoderm |
| Gpam | 3.23E-14 | 0.529674554 | 1 | 0.708 | 4.06E-10 | Visceral Endoderm |
| Acin1 | 3.24E-14 | 0.325341097 | 1 | 1 | 4.07E-10 | Visceral Endoderm |
| Klhl2 | 3.62E-14 | 0.53551754 | 1 | 0.583 | 4.56E-10 | Visceral Endoderm |
| Rin3 | 4.60E-14 | 0.262260341 | 1 | 0.361 | 5.79E-10 | Visceral Endoderm |
| AI413582 | 5.01E-14 | 0.552922993 | 0.857 | 0.306 | 6.30E-10 | Visceral Endoderm |

| | | | | | | |
|---|---|---|---|---|---|---|
| Qprt | 5.12E-14 | 0.361287675 | 1 | 0.875 | 6.44E-10 | Visceral Endoderm |
| Sirt7 | 6.56E-14 | 0.342049953 | 0.857 | 0.833 | 8.24E-10 | Visceral Endoderm |
| Gba2 | 6.67E-14 | 0.363203511 | 0.429 | 0.75 | 8.39E-10 | Visceral Endoderm |
| Mafk | 6.91E-14 | 0.628484124 | 0.857 | 0.667 | 8.69E-10 | Visceral Endoderm |
| Slc35f6 | 7.72E-14 | 0.270769037 | 1 | 0.847 | 9.71E-10 | Visceral Endoderm |
| Wdr24 | 8.38E-14 | 0.426543386 | 0.571 | 0.764 | 1.05E-09 | Visceral Endoderm |
| Nhlrc3 | 8.81E-14 | 0.276159936 | 1 | 0.681 | 1.11E-09 | Visceral Endoderm |
| Adora2b | 1.04E-13 | 1.053441998 | 1 | 0.153 | 1.30E-09 | Visceral Endoderm |
| 2310035C | 1.06E-13 | 0.269711732 | 0.857 | 1 | 1.34E-09 | Visceral Endoderm |
| Kctd6 | 1.37E-13 | 0.510605718 | 0.571 | 0.347 | 1.72E-09 | Visceral Endoderm |
| Tmem37 | 1.41E-13 | 1.025414812 | 1 | 0.583 | 1.77E-09 | Visceral Endoderm |
| Asl | 1.43E-13 | 0.56199822 | 1 | 0.917 | 1.80E-09 | Visceral Endoderm |
| Rhot1 | 1.45E-13 | 0.456042314 | 1 | 0.986 | 1.82E-09 | Visceral Endoderm |
| Tmem79 | 1.52E-13 | 0.263873772 | 0.571 | 0.403 | 1.91E-09 | Visceral Endoderm |
| Sfxn5 | 1.52E-13 | 0.30169969 | 1 | 0.792 | 1.91E-09 | Visceral Endoderm |
| Mt2 | 1.62E-13 | 2.289109467 | 1 | 0.347 | 2.04E-09 | Visceral Endoderm |
| Slc25a19 | 1.90E-13 | 0.282789222 | 1 | 0.694 | 2.40E-09 | Visceral Endoderm |
| Slc38a6 | 2.26E-13 | 0.556396483 | 1 | 1 | 2.85E-09 | Visceral Endoderm |
| Trim25 | 2.30E-13 | 0.423301756 | 1 | 0.667 | 2.89E-09 | Visceral Endoderm |
| Gigyf2 | 2.82E-13 | 0.426176017 | 1 | 0.986 | 3.55E-09 | Visceral Endoderm |
| Dcaf15 | 3.08E-13 | 0.37280991 | 0.857 | 0.653 | 3.87E-09 | Visceral Endoderm |
| Bach1 | 3.46E-13 | 0.304033003 | 0.571 | 0.75 | 4.35E-09 | Visceral Endoderm |
| Sh3tc1 | 4.18E-13 | 0.43720778 | 1 | 0.278 | 5.26E-09 | Visceral Endoderm |
| Hmox1 | 4.27E-13 | 0.961065741 | 1 | 0.694 | 5.38E-09 | Visceral Endoderm |
| Folr1 | 4.43E-13 | 1.065544004 | 1 | 0.472 | 5.57E-09 | Visceral Endoderm |
| Tfcp2l1 | 5.58E-13 | 0.489863731 | 1 | 0.472 | 7.02E-09 | Visceral Endoderm |
| Trrap | 8.07E-13 | 0.275153774 | 1 | 1 | 1.01E-08 | Visceral Endoderm |
| Zc3h7a | 8.55E-13 | 0.37784773 | 1 | 1 | 1.08E-08 | Visceral Endoderm |
| Mthfr | 9.55E-13 | 0.254359234 | 1 | 0.736 | 1.20E-08 | Visceral Endoderm |
| Atp6v1a | 1.03E-12 | 0.584135068 | 1 | 1 | 1.29E-08 | Visceral Endoderm |
| Stxbp2 | 1.24E-12 | 0.926899447 | 1 | 0.986 | 1.56E-08 | Visceral Endoderm |
| Josd2 | 1.29E-12 | 0.321231308 | 0.429 | 0.861 | 1.62E-08 | Visceral Endoderm |
| Lgals2 | 1.31E-12 | 1.308674916 | 0.714 | 0.111 | 1.65E-08 | Visceral Endoderm |
| Ift46 | 1.42E-12 | 0.593221599 | 1 | 1 | 1.78E-08 | Visceral Endoderm |
| Nadsyn1 | 1.43E-12 | 0.393654867 | 0.857 | 0.542 | 1.80E-08 | Visceral Endoderm |
| Hook2 | 1.72E-12 | 1.045345207 | 1 | 0.736 | 2.16E-08 | Visceral Endoderm |
| Eef2kmt | 1.72E-12 | 0.266063209 | 1 | 0.792 | 2.16E-08 | Visceral Endoderm |
| Fuca2 | 2.23E-12 | 0.323160048 | 1 | 0.917 | 2.81E-08 | Visceral Endoderm |
| Ctsz | 2.83E-12 | 1.606027398 | 1 | 1 | 3.56E-08 | Visceral Endoderm |
| Renbp | 3.56E-12 | 0.657514513 | 1 | 0.722 | 4.47E-08 | Visceral Endoderm |
| Kmt2d | 3.61E-12 | 0.420978901 | 1 | 1 | 4.54E-08 | Visceral Endoderm |
| Tbc1d10a | 4.15E-12 | 0.831671946 | 1 | 0.764 | 5.22E-08 | Visceral Endoderm |
| Txndc11 | 4.56E-12 | 0.295530381 | 1 | 0.986 | 5.74E-08 | Visceral Endoderm |
| Cmtm4 | 4.64E-12 | 0.385400095 | 1 | 0.569 | 5.83E-08 | Visceral Endoderm |
| Muc13 | 4.64E-12 | 0.338858386 | 0.714 | 0.222 | 5.84E-08 | Visceral Endoderm |
| Slc25a22 | 4.76E-12 | 0.765954373 | 0.857 | 0.917 | 5.98E-08 | Visceral Endoderm |
| Hnf4a | 6.63E-12 | 0.489142225 | 0.857 | 0.139 | 8.34E-08 | Visceral Endoderm |
| Xpo6 | 1.11E-11 | 0.265978985 | 1 | 0.903 | 1.39E-07 | Visceral Endoderm |
| Lpar1 | 1.12E-11 | 0.738283891 | 1 | 0.708 | 1.41E-07 | Visceral Endoderm |
| Atp6v0d1 | 1.25E-11 | 0.415760757 | 1 | 0.958 | 1.57E-07 | Visceral Endoderm |
| Wash1 | 1.33E-11 | 0.274029302 | 0.857 | 0.958 | 1.67E-07 | Visceral Endoderm |
| Ctsl | 1.34E-11 | 1.662357133 | 1 | 1 | 1.68E-07 | Visceral Endoderm |
| Pmm2 | 1.42E-11 | 0.473154628 | 1 | 0.972 | 1.78E-07 | Visceral Endoderm |
| Pick1 | 1.49E-11 | 0.3028381 | 1 | 0.986 | 1.88E-07 | Visceral Endoderm |
| Dtx3 | 1.56E-11 | 0.570948734 | 1 | 0.958 | 1.97E-07 | Visceral Endoderm |
| Cltc | 1.93E-11 | 0.799770946 | 1 | 0.986 | 2.43E-07 | Visceral Endoderm |
| Hps5 | 1.95E-11 | 0.303685358 | 1 | 0.931 | 2.45E-07 | Visceral Endoderm |
| Atp6v1h | 2.33E-11 | 0.319609039 | 1 | 1 | 2.93E-07 | Visceral Endoderm |

| | | | | | | |
|---|---|---|---|---|---|---|
| Commd4 | 2.52E-11 | 1.100221388 | 1 | 0.931 | 3.17E-07 | Visceral Endoderm |
| Tuba4a | 2.54E-11 | 0.543412117 | 0.857 | 0.569 | 3.19E-07 | Visceral Endoderm |
| Nfx1 | 2.58E-11 | 0.733086684 | 1 | 1 | 3.25E-07 | Visceral Endoderm |
| Narfl | 2.71E-11 | 0.473408677 | 0.857 | 0.833 | 3.41E-07 | Visceral Endoderm |
| Ninj1 | 2.77E-11 | 0.597262959 | 0.857 | 0.639 | 3.48E-07 | Visceral Endoderm |
| Cgnl1 | 2.88E-11 | 0.431168824 | 1 | 0.681 | 3.62E-07 | Visceral Endoderm |
| Tlcd1 | 3.07E-11 | 0.293826576 | 1 | 0.833 | 3.86E-07 | Visceral Endoderm |
| Ubqln2 | 3.26E-11 | 0.351374586 | 1 | 0.958 | 4.10E-07 | Visceral Endoderm |
| Rpusd3 | 3.29E-11 | 0.369324212 | 0.571 | 0.708 | 4.14E-07 | Visceral Endoderm |
| Uvrag | 3.34E-11 | 0.602371771 | 0.857 | 0.792 | 4.19E-07 | Visceral Endoderm |
| Rogdi | 3.40E-11 | 1.063289806 | 1 | 0.917 | 4.28E-07 | Visceral Endoderm |
| Camsap3 | 3.53E-11 | 0.674206463 | 1 | 0.764 | 4.43E-07 | Visceral Endoderm |
| Immp2l | 4.30E-11 | 0.34448971 | 0.571 | 0.278 | 5.41E-07 | Visceral Endoderm |
| Prrc2a | 4.32E-11 | 0.409559827 | 1 | 1 | 5.43E-07 | Visceral Endoderm |
| Hmgxb3 | 4.36E-11 | 0.315534846 | 1 | 0.931 | 5.48E-07 | Visceral Endoderm |
| Plekhg3 | 4.86E-11 | 0.277300156 | 0.714 | 0.764 | 6.12E-07 | Visceral Endoderm |
| Atp1b1 | 5.10E-11 | 0.895758226 | 1 | 0.625 | 6.42E-07 | Visceral Endoderm |
| Cib1 | 5.32E-11 | 0.483979658 | 0.857 | 0.889 | 6.70E-07 | Visceral Endoderm |
| Smcr8 | 5.37E-11 | 0.364732031 | 0.857 | 0.764 | 6.75E-07 | Visceral Endoderm |
| Glis1 | 6.47E-11 | 0.300563002 | 0.857 | 0.111 | 8.14E-07 | Visceral Endoderm |
| Tcirg1 | 6.68E-11 | 0.506453374 | 1 | 0.819 | 8.40E-07 | Visceral Endoderm |
| Pbld2 | 6.74E-11 | 0.260019782 | 1 | 1 | 8.47E-07 | Visceral Endoderm |
| Ralgps1 | 7.28E-11 | 0.255017864 | 1 | 0.486 | 9.16E-07 | Visceral Endoderm |
| Rbm25 | 8.03E-11 | 0.50129983 | 1 | 1 | 1.01E-06 | Visceral Endoderm |
| Grtp1 | 8.14E-11 | 0.815317181 | 0.857 | 0.556 | 1.02E-06 | Visceral Endoderm |
| Zdhhc12 | 8.33E-11 | 1.018458127 | 0.857 | 0.833 | 1.05E-06 | Visceral Endoderm |
| 2510039O | 9.81E-11 | 0.320025558 | 0.571 | 0.5 | 1.23E-06 | Visceral Endoderm |
| Ghdc | 9.99E-11 | 0.344956855 | 1 | 1 | 1.26E-06 | Visceral Endoderm |
| Golgb1 | 1.05E-10 | 0.251832671 | 1 | 0.917 | 1.32E-06 | Visceral Endoderm |
| B4galt7 | 1.06E-10 | 0.415288096 | 1 | 0.986 | 1.33E-06 | Visceral Endoderm |
| Sigirr | 1.11E-10 | 0.646642503 | 1 | 0.167 | 1.39E-06 | Visceral Endoderm |
| Klc4 | 1.35E-10 | 0.882361646 | 1 | 0.778 | 1.70E-06 | Visceral Endoderm |
| Dgat2 | 1.60E-10 | 0.965011129 | 1 | 0.847 | 2.01E-06 | Visceral Endoderm |
| Lrp10 | 1.60E-10 | 0.50631572 | 1 | 0.764 | 2.01E-06 | Visceral Endoderm |
| Trafd1 | 1.60E-10 | 0.396427243 | 1 | 0.833 | 2.01E-06 | Visceral Endoderm |
| Trip12 | 1.68E-10 | 0.353409847 | 1 | 1 | 2.11E-06 | Visceral Endoderm |
| Pafah2 | 1.72E-10 | 0.548763916 | 0.857 | 0.528 | 2.16E-06 | Visceral Endoderm |
| Zfp948 | 1.79E-10 | 0.969884399 | 0.857 | 0.847 | 2.25E-06 | Visceral Endoderm |
| Rabggta | 1.92E-10 | 0.525152435 | 1 | 1 | 2.41E-06 | Visceral Endoderm |
| Polr1b | 1.99E-10 | 0.485921979 | 1 | 0.972 | 2.51E-06 | Visceral Endoderm |
| Psph | 2.00E-10 | 0.429415826 | 1 | 0.958 | 2.51E-06 | Visceral Endoderm |
| Preb | 2.02E-10 | 0.611382381 | 1 | 0.972 | 2.54E-06 | Visceral Endoderm |
| Ecm1 | 2.04E-10 | 0.338713336 | 0.714 | 0.306 | 2.56E-06 | Visceral Endoderm |
| Fpgs | 2.20E-10 | 0.364073922 | 0.714 | 0.861 | 2.77E-06 | Visceral Endoderm |
| Syvn1 | 2.24E-10 | 0.427317471 | 1 | 0.958 | 2.82E-06 | Visceral Endoderm |
| Gpatch3 | 2.39E-10 | 0.542390162 | 0.714 | 0.722 | 3.01E-06 | Visceral Endoderm |
| Slc12a9 | 2.45E-10 | 0.396048222 | 0.714 | 0.681 | 3.08E-06 | Visceral Endoderm |
| Gjb2 | 2.48E-10 | 0.385403432 | 0.714 | 0.167 | 3.11E-06 | Visceral Endoderm |
| Phyh | 2.48E-10 | 0.917874211 | 0.857 | 0.736 | 3.12E-06 | Visceral Endoderm |
| Hrsp12 | 2.52E-10 | 0.382968811 | 0.857 | 1 | 3.17E-06 | Visceral Endoderm |
| Man2c1 | 2.62E-10 | 0.265454782 | 1 | 0.903 | 3.29E-06 | Visceral Endoderm |
| Cep192 | 2.62E-10 | 0.294570145 | 0.857 | 0.931 | 3.30E-06 | Visceral Endoderm |
| Lama5 | 2.63E-10 | 0.326731291 | 1 | 0.931 | 3.30E-06 | Visceral Endoderm |
| Hpn | 2.82E-10 | 0.671532392 | 1 | 0.403 | 3.55E-06 | Visceral Endoderm |
| Mttp | 2.99E-10 | 0.905108897 | 1 | 0.903 | 3.77E-06 | Visceral Endoderm |
| Tpra1 | 3.18E-10 | 0.626711356 | 0.857 | 0.931 | 4.01E-06 | Visceral Endoderm |
| Ip6k1 | 3.60E-10 | 0.428534816 | 1 | 0.972 | 4.52E-06 | Visceral Endoderm |
| 1600002H | 3.88E-10 | 0.723121948 | 1 | 0.917 | 4.88E-06 | Visceral Endoderm |

| | | | | | | |
|---|---|---|---|---|---|---|
| Selt | 4.14E-10 | 0.399457149 | 1 | 0.889 | 5.21E-06 | Visceral Endoderm |
| Dnmbp | 4.64E-10 | 0.381371181 | 1 | 0.944 | 5.83E-06 | Visceral Endoderm |
| Agfg1 | 4.92E-10 | 0.333806884 | 1 | 0.917 | 6.18E-06 | Visceral Endoderm |
| Slc7a9 | 5.11E-10 | 0.669606245 | 0.857 | 0.139 | 6.42E-06 | Visceral Endoderm |
| Arpc1b | 5.34E-10 | 0.256639899 | 1 | 0.917 | 6.71E-06 | Visceral Endoderm |
| Fam120a | 5.47E-10 | 0.428505869 | 1 | 0.917 | 6.88E-06 | Visceral Endoderm |
| Dlg5 | 5.62E-10 | 0.272061913 | 1 | 0.792 | 7.07E-06 | Visceral Endoderm |
| Socs5 | 5.69E-10 | 0.42911267 | 0.429 | 0.597 | 7.15E-06 | Visceral Endoderm |
| Dram1 | 5.80E-10 | 0.672906402 | 1 | 0.056 | 7.30E-06 | Visceral Endoderm |
| Mgrn1 | 5.94E-10 | 0.572199826 | 1 | 0.958 | 7.47E-06 | Visceral Endoderm |
| Zfp871 | 6.07E-10 | 0.256557029 | 0.857 | 0.833 | 7.63E-06 | Visceral Endoderm |
| Pstk | 6.36E-10 | 0.322588488 | 0.714 | 0.903 | 8.00E-06 | Visceral Endoderm |
| Hexa | 6.67E-10 | 1.042304612 | 1 | 0.861 | 8.39E-06 | Visceral Endoderm |
| Slc27a1 | 6.72E-10 | 0.271166458 | 1 | 0.833 | 8.45E-06 | Visceral Endoderm |
| Nle1 | 7.13E-10 | 0.253423508 | 1 | 0.931 | 8.97E-06 | Visceral Endoderm |
| Tom1l1 | 7.75E-10 | 0.260131713 | 1 | 0.917 | 9.75E-06 | Visceral Endoderm |
| Plekha7 | 8.12E-10 | 0.30714299 | 1 | 0.944 | 1.02E-05 | Visceral Endoderm |
| Gmeb1 | 8.86E-10 | 0.255681532 | 1 | 1 | 1.11E-05 | Visceral Endoderm |
| Syne4 | 9.64E-10 | 0.44516165 | 0.857 | 0.472 | 1.21E-05 | Visceral Endoderm |
| Rbp4 | 1.02E-09 | 2.123384547 | 1 | 0.861 | 1.28E-05 | Visceral Endoderm |
| Fastk | 1.04E-09 | 0.515616599 | 1 | 0.986 | 1.31E-05 | Visceral Endoderm |
| Ptov1 | 1.04E-09 | 0.341243989 | 1 | 1 | 1.31E-05 | Visceral Endoderm |
| AI464131 | 1.06E-09 | 0.326228544 | 0.857 | 0.639 | 1.33E-05 | Visceral Endoderm |
| Rnf111 | 1.07E-09 | 0.361092606 | 0.714 | 0.944 | 1.34E-05 | Visceral Endoderm |
| Zfp69 | 1.08E-09 | 0.368177596 | 0.714 | 0.444 | 1.35E-05 | Visceral Endoderm |
| Prdm6 | 1.09E-09 | 0.490807463 | 1 | 0.389 | 1.37E-05 | Visceral Endoderm |
| Dnajc4 | 1.11E-09 | 0.47868084 | 0.286 | 0.514 | 1.40E-05 | Visceral Endoderm |
| Nxf7 | 1.17E-09 | 0.459440762 | 1 | 0.319 | 1.47E-05 | Visceral Endoderm |
| Thnsl2 | 1.21E-09 | 0.386794011 | 0.571 | 0.264 | 1.52E-05 | Visceral Endoderm |
| Gphn | 1.28E-09 | 0.268698051 | 0.571 | 0.639 | 1.61E-05 | Visceral Endoderm |
| Pdzk1 | 1.31E-09 | 0.489466404 | 0.857 | 0.194 | 1.65E-05 | Visceral Endoderm |
| Nipa2 | 1.37E-09 | 0.436590556 | 1 | 1 | 1.72E-05 | Visceral Endoderm |
| Rhou | 1.41E-09 | 0.362627568 | 1 | 0.75 | 1.77E-05 | Visceral Endoderm |
| Naaa | 1.44E-09 | 0.363503234 | 1 | 0.764 | 1.81E-05 | Visceral Endoderm |
| Sec16a | 1.47E-09 | 0.327469645 | 1 | 0.972 | 1.85E-05 | Visceral Endoderm |
| Xlr4b | 1.49E-09 | 0.442936616 | 1 | 0.292 | 1.88E-05 | Visceral Endoderm |
| Lrrc19 | 1.61E-09 | 0.299545522 | 0.714 | 0.167 | 2.03E-05 | Visceral Endoderm |
| Srrm1 | 1.64E-09 | 0.314022311 | 1 | 1 | 2.06E-05 | Visceral Endoderm |
| Zfp36 | 1.64E-09 | 1.07126589 | 0.857 | 0.472 | 2.06E-05 | Visceral Endoderm |
| Tmem161l | 1.65E-09 | 0.600241009 | 1 | 0.903 | 2.07E-05 | Visceral Endoderm |
| Trpm7 | 1.79E-09 | 0.347242119 | 1 | 1 | 2.25E-05 | Visceral Endoderm |
| Tgm1 | 1.82E-09 | 0.443267862 | 0.714 | 0.167 | 2.29E-05 | Visceral Endoderm |
| Mfsd5 | 1.86E-09 | 0.493947617 | 0.571 | 0.667 | 2.33E-05 | Visceral Endoderm |
| Slc20a1 | 1.90E-09 | 0.520000968 | 1 | 0.931 | 2.39E-05 | Visceral Endoderm |
| Slc25a28 | 1.96E-09 | 0.382713747 | 0.571 | 0.278 | 2.46E-05 | Visceral Endoderm |
| Dqx1 | 1.98E-09 | 0.460090237 | 1 | 0.903 | 2.49E-05 | Visceral Endoderm |
| Fem1c | 2.08E-09 | 0.279826856 | 0.857 | 0.931 | 2.62E-05 | Visceral Endoderm |
| Galt | 2.14E-09 | 0.395347572 | 0.714 | 0.542 | 2.69E-05 | Visceral Endoderm |
| Araf | 2.14E-09 | 0.460447153 | 1 | 1 | 2.69E-05 | Visceral Endoderm |
| Ly75 | 2.36E-09 | 0.41896943 | 1 | 0.778 | 2.96E-05 | Visceral Endoderm |
| Msx1 | 2.55E-09 | 0.829897923 | 1 | 0.417 | 3.21E-05 | Visceral Endoderm |
| Rap1gap2 | 2.56E-09 | 0.455927129 | 0.714 | 0.208 | 3.21E-05 | Visceral Endoderm |
| Vil1 | 2.80E-09 | 0.647872181 | 1 | 0.403 | 3.52E-05 | Visceral Endoderm |
| Mfsd1 | 2.86E-09 | 1.089226286 | 1 | 1 | 3.59E-05 | Visceral Endoderm |
| Gns | 3.04E-09 | 0.827379583 | 1 | 0.972 | 3.83E-05 | Visceral Endoderm |
| Slc3a2 | 3.33E-09 | 0.984994879 | 1 | 1 | 4.19E-05 | Visceral Endoderm |
| Msx2 | 3.56E-09 | 0.634707974 | 1 | 0.5 | 4.48E-05 | Visceral Endoderm |
| Fbxl3 | 3.72E-09 | 0.291893593 | 1 | 1 | 4.68E-05 | Visceral Endoderm |

| | | | | | | |
|---|---|---|---|---|---|---|
| Pkp3 | 3.77E-09 | 0.324216704 | 0.714 | 0.25 | 4.74E-05 | Visceral Endoderm |
| Aqp11 | 3.92E-09 | 0.750212948 | 0.714 | 0.347 | 4.93E-05 | Visceral Endoderm |
| Usp7 | 4.05E-09 | 0.485811375 | 1 | 0.958 | 5.09E-05 | Visceral Endoderm |
| Ndor1 | 4.53E-09 | 0.26820222 | 1 | 0.917 | 5.69E-05 | Visceral Endoderm |
| Mtmr12 | 4.83E-09 | 0.410382632 | 0.857 | 0.792 | 6.08E-05 | Visceral Endoderm |
| Usp36 | 4.89E-09 | 0.259770684 | 1 | 0.931 | 6.16E-05 | Visceral Endoderm |
| Parvb | 4.90E-09 | 0.285472005 | 1 | 0.931 | 6.16E-05 | Visceral Endoderm |
| Eps8l2 | 4.93E-09 | 0.305537608 | 0.857 | 0.153 | 6.19E-05 | Visceral Endoderm |
| Inpp5f | 4.98E-09 | 0.313838315 | 1 | 1 | 6.27E-05 | Visceral Endoderm |
| Lars2 | 5.21E-09 | 1.678857442 | 1 | 1 | 6.55E-05 | Visceral Endoderm |
| Irf6 | 5.67E-09 | 0.461552502 | 0.857 | 0.417 | 7.14E-05 | Visceral Endoderm |
| Mtg1 | 5.72E-09 | 0.315595878 | 0.714 | 0.722 | 7.19E-05 | Visceral Endoderm |
| Uso1 | 5.83E-09 | 0.561059258 | 0.857 | 0.764 | 7.33E-05 | Visceral Endoderm |
| Klf9 | 5.84E-09 | 0.356170122 | 0.714 | 0.319 | 7.34E-05 | Visceral Endoderm |
| Lzts2 | 6.05E-09 | 0.492329465 | 1 | 0.972 | 7.61E-05 | Visceral Endoderm |
| Zswim1 | 6.44E-09 | 0.25818916 | 0.714 | 0.694 | 8.10E-05 | Visceral Endoderm |
| Akr1b10 | 6.83E-09 | 0.259623432 | 0.571 | 0.681 | 8.59E-05 | Visceral Endoderm |
| Edem1 | 7.05E-09 | 0.272099329 | 0.714 | 0.903 | 8.86E-05 | Visceral Endoderm |
| Cars2 | 7.22E-09 | 0.362440499 | 0.714 | 0.792 | 9.08E-05 | Visceral Endoderm |
| Ehhadh | 7.42E-09 | 0.437353754 | 0.571 | 0.208 | 9.33E-05 | Visceral Endoderm |
| Znf512b | 8.05E-09 | 0.272144495 | 0.857 | 0.847 | 0.000101 | Visceral Endoderm |
| Plbd1 | 8.25E-09 | 0.342897435 | 1 | 0.25 | 0.000104 | Visceral Endoderm |
| Abcc5 | 8.28E-09 | 0.480193892 | 1 | 0.958 | 0.000104 | Visceral Endoderm |
| Abhd11 | 8.49E-09 | 0.384902145 | 0.857 | 0.75 | 0.000107 | Visceral Endoderm |
| Ino80 | 8.50E-09 | 0.26575337 | 1 | 0.917 | 0.000107 | Visceral Endoderm |
| Foxm1 | 8.69E-09 | 0.410288088 | 1 | 0.778 | 0.000109 | Visceral Endoderm |
| 1700019D | 8.81E-09 | 0.340779375 | 0.857 | 0.528 | 0.000111 | Visceral Endoderm |
| Nfkbiz | 8.96E-09 | 0.272097122 | 1 | 0.639 | 0.000113 | Visceral Endoderm |
| Utp20 | 1.11E-08 | 0.291242481 | 1 | 0.903 | 0.000139 | Visceral Endoderm |
| B4galt3 | 1.13E-08 | 0.329509796 | 1 | 0.931 | 0.000142 | Visceral Endoderm |
| Epb4.1l3 | 1.18E-08 | 1.039544636 | 1 | 0.931 | 0.000148 | Visceral Endoderm |
| Acot8 | 1.24E-08 | 0.256668917 | 0.714 | 0.597 | 0.000155 | Visceral Endoderm |
| Fastkd2 | 1.25E-08 | 0.333915355 | 1 | 0.944 | 0.000158 | Visceral Endoderm |
| Rbm26 | 1.27E-08 | 0.285940233 | 1 | 1 | 0.00016 | Visceral Endoderm |
| Aplp1 | 1.36E-08 | 0.374095823 | 0.571 | 0.458 | 0.00017 | Visceral Endoderm |
| Ankrd13a | 1.36E-08 | 0.255526971 | 0.571 | 0.583 | 0.000171 | Visceral Endoderm |
| Apom | 1.39E-08 | 1.89594686 | 1 | 0.583 | 0.000175 | Visceral Endoderm |
| Mri1 | 1.40E-08 | 0.328479064 | 1 | 0.972 | 0.000177 | Visceral Endoderm |
| Ttr | 1.48E-08 | 1.803863051 | 1 | 0.931 | 0.000186 | Visceral Endoderm |
| Hmg20b | 1.50E-08 | 0.635284926 | 1 | 0.972 | 0.000189 | Visceral Endoderm |
| Lactb2 | 1.55E-08 | 0.321689577 | 1 | 1 | 0.000195 | Visceral Endoderm |
| Susd6 | 1.79E-08 | 0.350952634 | 0.857 | 0.792 | 0.000225 | Visceral Endoderm |
| Cldn2 | 1.88E-08 | 1.019596104 | 1 | 0.278 | 0.000237 | Visceral Endoderm |
| Atp6v0b | 2.06E-08 | 0.713687804 | 1 | 0.986 | 0.000259 | Visceral Endoderm |
| Ncoa3 | 2.10E-08 | 0.289685316 | 1 | 1 | 0.000264 | Visceral Endoderm |
| Vwce | 2.18E-08 | 0.603173638 | 0.714 | 0.333 | 0.000274 | Visceral Endoderm |
| Tlk2 | 2.26E-08 | 0.269475568 | 0.857 | 0.889 | 0.000284 | Visceral Endoderm |
| Pisd | 2.45E-08 | 0.260986579 | 1 | 0.958 | 0.000308 | Visceral Endoderm |
| Bend7 | 2.47E-08 | 0.286534104 | 0.714 | 0.236 | 0.00031 | Visceral Endoderm |
| Slc22a23 | 2.50E-08 | 0.314590458 | 1 | 0.681 | 0.000314 | Visceral Endoderm |
| Mms19 | 2.52E-08 | 0.277795621 | 1 | 0.986 | 0.000316 | Visceral Endoderm |
| Alg13 | 2.80E-08 | 0.390353065 | 1 | 0.917 | 0.000352 | Visceral Endoderm |
| Senp5 | 2.82E-08 | 0.412413353 | 0.857 | 0.847 | 0.000354 | Visceral Endoderm |
| Slc16a9 | 2.83E-08 | 0.395469515 | 0.857 | 0.097 | 0.000355 | Visceral Endoderm |
| Ccdc120 | 3.16E-08 | 0.333817703 | 1 | 0.639 | 0.000398 | Visceral Endoderm |
| Oat | 3.19E-08 | 1.205595587 | 1 | 0.944 | 0.000401 | Visceral Endoderm |
| Srsf2 | 3.33E-08 | 1.134463398 | 1 | 1 | 0.000418 | Visceral Endoderm |
| Ppp2r3d | 3.38E-08 | 0.349860793 | 1 | 0.931 | 0.000425 | Visceral Endoderm |

| | | | | | | |
|---|---|---|---|---|---|---|
| Pus1 | 3.41E-08 | 0.335213058 | 1 | 0.861 | 0.000429 | Visceral Endoderm |
| Nus1 | 3.44E-08 | 0.301735392 | 1 | 0.819 | 0.000433 | Visceral Endoderm |
| Abhd17b | 3.71E-08 | 0.376451421 | 0.571 | 0.569 | 0.000466 | Visceral Endoderm |
| Sypl | 3.78E-08 | 0.468166705 | 1 | 0.958 | 0.000475 | Visceral Endoderm |
| Ankrd11 | 3.79E-08 | 0.502526746 | 1 | 1 | 0.000477 | Visceral Endoderm |
| Fxr2 | 3.87E-08 | 0.354885844 | 0.857 | 0.917 | 0.000487 | Visceral Endoderm |
| Dnajc22 | 4.02E-08 | 0.819628896 | 1 | 0.389 | 0.000505 | Visceral Endoderm |
| Manba | 4.25E-08 | 0.29997096 | 0.857 | 0.403 | 0.000534 | Visceral Endoderm |
| Slco4c1 | 4.29E-08 | 0.366037128 | 1 | 0.125 | 0.00054 | Visceral Endoderm |
| Rab24 | 4.32E-08 | 0.784204878 | 0.857 | 0.875 | 0.000543 | Visceral Endoderm |
| Abhd16a | 4.34E-08 | 0.335077678 | 0.714 | 0.875 | 0.000546 | Visceral Endoderm |
| Ctsc | 4.39E-08 | 0.896568694 | 1 | 0.778 | 0.000553 | Visceral Endoderm |
| Rab6b | 4.62E-08 | 0.294987107 | 0.714 | 0.639 | 0.000581 | Visceral Endoderm |
| Wbp2 | 5.23E-08 | 0.747748812 | 1 | 1 | 0.000658 | Visceral Endoderm |
| Ptpn6 | 5.26E-08 | 0.30262781 | 0.857 | 0.583 | 0.000661 | Visceral Endoderm |
| Stx4a | 5.58E-08 | 0.365340646 | 0.571 | 0.5 | 0.000702 | Visceral Endoderm |
| Pomt1 | 5.71E-08 | 0.261857615 | 1 | 0.958 | 0.000718 | Visceral Endoderm |
| Ifi30 | 5.71E-08 | 1.274274811 | 1 | 0.792 | 0.000719 | Visceral Endoderm |
| Ddt | 5.98E-08 | 0.441161219 | 0.714 | 0.986 | 0.000752 | Visceral Endoderm |
| Dlg1 | 6.00E-08 | 0.3277586 | 1 | 1 | 0.000755 | Visceral Endoderm |
| Tcn2 | 6.14E-08 | 0.89774333 | 1 | 0.833 | 0.000772 | Visceral Endoderm |
| Cndp2 | 6.50E-08 | 1.040560483 | 1 | 0.986 | 0.000817 | Visceral Endoderm |
| Kdf1 | 6.56E-08 | 0.379114598 | 0.857 | 0.5 | 0.000826 | Visceral Endoderm |
| Hsf2 | 6.81E-08 | 0.543311631 | 1 | 0.986 | 0.000856 | Visceral Endoderm |
| Slc7a7 | 6.89E-08 | 0.608072621 | 1 | 0.736 | 0.000867 | Visceral Endoderm |
| Etv4 | 7.23E-08 | 0.298422204 | 0.714 | 0.417 | 0.000909 | Visceral Endoderm |
| Gcsh | 7.48E-08 | 1.185592024 | 1 | 1 | 0.000941 | Visceral Endoderm |
| Daam1 | 8.16E-08 | 0.253027524 | 0.857 | 0.639 | 0.001026 | Visceral Endoderm |
| Shf | 8.31E-08 | 1.790016088 | 0.571 | 0.208 | 0.001045 | Visceral Endoderm |
| Napa | 8.50E-08 | 0.557308745 | 1 | 1 | 0.001069 | Visceral Endoderm |
| Lnx2 | 8.60E-08 | 0.294565244 | 1 | 0.75 | 0.001082 | Visceral Endoderm |
| Zfp956 | 8.97E-08 | 0.400067249 | 0.571 | 0.361 | 0.001129 | Visceral Endoderm |
| Pkp4 | 9.16E-08 | 0.41791547 | 1 | 0.917 | 0.001152 | Visceral Endoderm |
| Mcee | 9.18E-08 | 0.564187998 | 0.857 | 0.694 | 0.001155 | Visceral Endoderm |
| Far2 | 9.34E-08 | 0.259529782 | 0.571 | 0.347 | 0.001175 | Visceral Endoderm |
| Vps29 | 9.64E-08 | 0.270457832 | 1 | 1 | 0.001213 | Visceral Endoderm |
| Xlr3b | 9.84E-08 | 0.516158449 | 1 | 0.681 | 0.001238 | Visceral Endoderm |
| Xlr3a | 9.94E-08 | 0.571588312 | 1 | 0.542 | 0.00125 | Visceral Endoderm |
| Mrps17 | 1.00E-07 | 0.294005896 | 1 | 1 | 0.001259 | Visceral Endoderm |
| Crtap | 1.01E-07 | 0.882272841 | 1 | 0.778 | 0.001264 | Visceral Endoderm |
| Slc25a15 | 1.14E-07 | 0.406263223 | 1 | 0.819 | 0.001431 | Visceral Endoderm |
| Glrx | 1.26E-07 | 0.968773139 | 0.857 | 0.764 | 0.001583 | Visceral Endoderm |
| Stx3 | 1.28E-07 | 0.363501013 | 1 | 0.75 | 0.001615 | Visceral Endoderm |
| Kidins220 | 1.31E-07 | 0.325780971 | 1 | 0.764 | 0.001654 | Visceral Endoderm |
| Tmem63b | 1.33E-07 | 0.376718737 | 1 | 0.944 | 0.001679 | Visceral Endoderm |
| Emilin1 | 1.34E-07 | 0.354429602 | 0.571 | 0.417 | 0.00168 | Visceral Endoderm |
| E4f1 | 1.38E-07 | 0.433766064 | 0.857 | 0.722 | 0.001737 | Visceral Endoderm |
| Tmem241 | 1.61E-07 | 0.452871962 | 0.857 | 0.208 | 0.002021 | Visceral Endoderm |
| Foxn3 | 1.65E-07 | 0.38863183 | 1 | 0.917 | 0.002073 | Visceral Endoderm |
| Flcn | 1.75E-07 | 0.503278164 | 1 | 0.917 | 0.002197 | Visceral Endoderm |
| Lgalsl | 1.76E-07 | 0.278600687 | 1 | 0.694 | 0.002217 | Visceral Endoderm |
| Yod1 | 1.78E-07 | 0.290431359 | 1 | 0.944 | 0.002241 | Visceral Endoderm |
| Vps11 | 1.80E-07 | 0.335429258 | 0.571 | 0.833 | 0.002265 | Visceral Endoderm |
| Ccdc84 | 1.86E-07 | 0.57779086 | 0.857 | 0.778 | 0.002339 | Visceral Endoderm |
| Sephs2 | 1.87E-07 | 0.872723665 | 1 | 0.986 | 0.002355 | Visceral Endoderm |
| Gpn2 | 1.91E-07 | 0.510487308 | 0.714 | 0.861 | 0.002399 | Visceral Endoderm |
| Snx2 | 1.94E-07 | 0.845055775 | 1 | 0.958 | 0.002436 | Visceral Endoderm |
| Xlr4a | 1.98E-07 | 0.398828857 | 1 | 0.306 | 0.002491 | Visceral Endoderm |

| | | | | | | |
|---|---|---|---|---|---|---|
| Inpp5b | 2.11E-07 | 0.39160304 | 1 | 0.722 | 0.00265 | Visceral Endoderm |
| Lztr1 | 2.32E-07 | 0.456734258 | 1 | 1 | 0.002922 | Visceral Endoderm |
| Cdc14b | 2.34E-07 | 0.255645055 | 1 | 0.917 | 0.002945 | Visceral Endoderm |
| Dlk1 | 2.35E-07 | 0.254988326 | 0.857 | 0.306 | 0.002954 | Visceral Endoderm |
| Glul | 2.41E-07 | 0.769390152 | 1 | 1 | 0.003026 | Visceral Endoderm |
| Gsn | 2.52E-07 | 0.571440061 | 1 | 0.681 | 0.003173 | Visceral Endoderm |
| Efnb1 | 2.58E-07 | 0.36418727 | 1 | 0.736 | 0.003246 | Visceral Endoderm |
| Ap3d1 | 2.66E-07 | 0.537240612 | 1 | 0.972 | 0.003344 | Visceral Endoderm |
| Plekha6 | 2.83E-07 | 0.317276943 | 1 | 0.792 | 0.003562 | Visceral Endoderm |
| Oxsr1 | 2.88E-07 | 0.318511913 | 1 | 0.694 | 0.003628 | Visceral Endoderm |
| Igsf5 | 3.03E-07 | 0.315118407 | 0.857 | 0.069 | 0.003805 | Visceral Endoderm |
| Slc39a4 | 3.62E-07 | 0.94391149 | 1 | 0.681 | 0.004547 | Visceral Endoderm |
| Pth1r | 4.00E-07 | 0.827358375 | 1 | 0.778 | 0.005031 | Visceral Endoderm |
| Rab4a | 4.22E-07 | 0.773951537 | 1 | 0.931 | 0.005306 | Visceral Endoderm |
| 2510002D | 4.73E-07 | 0.363176306 | 0.571 | 0.764 | 0.00595 | Visceral Endoderm |
| Snx3 | 4.79E-07 | 1.130989495 | 1 | 0.986 | 0.006021 | Visceral Endoderm |
| Haus7 | 5.14E-07 | 0.954463633 | 1 | 0.931 | 0.006464 | Visceral Endoderm |
| Taf5l | 5.71E-07 | 0.385433113 | 0.714 | 0.833 | 0.007184 | Visceral Endoderm |
| Pgm3 | 6.00E-07 | 0.343937431 | 1 | 0.875 | 0.007544 | Visceral Endoderm |
| Csnk1g2 | 6.05E-07 | 0.435994382 | 0.857 | 0.833 | 0.007604 | Visceral Endoderm |
| Fxyd2 | 6.29E-07 | 0.884810186 | 0.571 | 0.125 | 0.007914 | Visceral Endoderm |
| Xlr4c | 6.30E-07 | 0.310414032 | 1 | 0.278 | 0.00792 | Visceral Endoderm |
| Twf1 | 6.54E-07 | 0.524208663 | 1 | 0.986 | 0.008228 | Visceral Endoderm |
| Rpe | 6.67E-07 | 0.395902682 | 1 | 0.958 | 0.008392 | Visceral Endoderm |
| Mrps26 | 7.09E-07 | 0.320028874 | 1 | 1 | 0.008921 | Visceral Endoderm |
| Plekha3 | 7.49E-07 | 0.253671782 | 1 | 1 | 0.009423 | Visceral Endoderm |
| Map2k2 | 7.60E-07 | 0.401813245 | 1 | 1 | 0.009557 | Visceral Endoderm |
| D15Ertd62 | 7.91E-07 | 0.351701266 | 0.857 | 0.875 | 0.009949 | Visceral Endoderm |
| Kif12 | 8.03E-07 | 0.845436561 | 0.714 | 0.347 | 0.010102 | Visceral Endoderm |
| Cdk11b | 8.10E-07 | 0.586297757 | 1 | 0.972 | 0.010192 | Visceral Endoderm |
| Scarb1 | 8.29E-07 | 0.269246451 | 1 | 0.944 | 0.010425 | Visceral Endoderm |
| Spry4 | 8.68E-07 | 0.353914816 | 0.714 | 0.389 | 0.010914 | Visceral Endoderm |
| Pcgf5 | 9.16E-07 | 0.631593452 | 0.857 | 0.278 | 0.011523 | Visceral Endoderm |
| Gas2l3 | 9.24E-07 | 0.355485952 | 1 | 1 | 0.011626 | Visceral Endoderm |
| Sh3bp5 | 9.27E-07 | 0.454415382 | 0.857 | 0.514 | 0.011657 | Visceral Endoderm |
| Amt | 9.46E-07 | 0.545796532 | 1 | 0.972 | 0.011895 | Visceral Endoderm |
| Soat1 | 9.74E-07 | 0.585487469 | 1 | 1 | 0.012249 | Visceral Endoderm |
| Tysnd1 | 9.79E-07 | 0.338191191 | 0.857 | 0.917 | 0.012317 | Visceral Endoderm |
| Hdac10 | 9.95E-07 | 0.42171991 | 0.857 | 0.542 | 0.012508 | Visceral Endoderm |
| Sbno1 | 1.00E-06 | 0.314488799 | 1 | 1 | 0.012601 | Visceral Endoderm |
| Rnf6 | 1.05E-06 | 0.410869556 | 0.857 | 0.764 | 0.013153 | Visceral Endoderm |
| Cgn | 1.10E-06 | 0.753150849 | 1 | 1 | 0.013878 | Visceral Endoderm |
| Tm4sf5 | 1.12E-06 | 0.784119215 | 0.571 | 0.097 | 0.014134 | Visceral Endoderm |
| Rictor | 1.18E-06 | 0.26307787 | 1 | 0.972 | 0.014874 | Visceral Endoderm |
| Apoc1 | 1.22E-06 | 0.925418197 | 0.857 | 0.208 | 0.015393 | Visceral Endoderm |
| Strn3 | 1.37E-06 | 0.316176891 | 0.857 | 0.986 | 0.017188 | Visceral Endoderm |
| Galm | 1.39E-06 | 0.402185874 | 0.571 | 0.528 | 0.017501 | Visceral Endoderm |
| Lamtor4 | 1.39E-06 | 0.504981431 | 1 | 0.917 | 0.017519 | Visceral Endoderm |
| Crebbp | 1.42E-06 | 0.256905825 | 1 | 1 | 0.017849 | Visceral Endoderm |
| Ihh | 1.46E-06 | 0.315282742 | 0.571 | 0.222 | 0.01833 | Visceral Endoderm |
| Plekhf2 | 1.47E-06 | 0.588957687 | 1 | 0.681 | 0.018503 | Visceral Endoderm |
| Setd8 | 1.51E-06 | 0.283279443 | 1 | 0.931 | 0.019029 | Visceral Endoderm |
| Rfng | 1.55E-06 | 0.494934856 | 0.857 | 0.819 | 0.01947 | Visceral Endoderm |
| Ap2a2 | 1.62E-06 | 0.453539961 | 1 | 1 | 0.020423 | Visceral Endoderm |
| Aprt | 1.66E-06 | 1.034248752 | 1 | 0.986 | 0.020887 | Visceral Endoderm |
| Spink3 | 1.76E-06 | 2.22917359 | 1 | 0.931 | 0.022072 | Visceral Endoderm |
| Slc39a8 | 1.79E-06 | 0.471140635 | 1 | 0.889 | 0.022513 | Visceral Endoderm |
| Palb2 | 1.84E-06 | 0.353663946 | 0.857 | 0.819 | 0.023123 | Visceral Endoderm |

| | | | | | | |
|---|---|---|---|---|---|---|
| Mvb12a | 1.84E-06 | 0.808392536 | 1 | 0.861 | 0.023139 | Visceral Endoderm |
| Stk24 | 1.84E-06 | 0.394587469 | 1 | 0.806 | 0.023142 | Visceral Endoderm |
| Trim44 | 1.91E-06 | 0.295481194 | 0.857 | 0.917 | 0.023964 | Visceral Endoderm |
| Nat10 | 2.24E-06 | 0.370002082 | 1 | 0.972 | 0.028174 | Visceral Endoderm |
| Dhps | 2.28E-06 | 0.345600261 | 0.714 | 0.889 | 0.028708 | Visceral Endoderm |
| Tjp3 | 2.38E-06 | 0.461955858 | 1 | 0.292 | 0.029887 | Visceral Endoderm |
| Gm21962 | 2.51E-06 | 0.506456158 | 1 | 0.986 | 0.031603 | Visceral Endoderm |
| Hars2 | 2.54E-06 | 0.308066177 | 1 | 0.847 | 0.031889 | Visceral Endoderm |
| Ak3 | 2.70E-06 | 0.401951999 | 1 | 0.75 | 0.033897 | Visceral Endoderm |
| Mogs | 2.84E-06 | 0.336922634 | 0.857 | 0.583 | 0.035727 | Visceral Endoderm |
| Tmed5 | 2.95E-06 | 0.276069815 | 1 | 1 | 0.037082 | Visceral Endoderm |
| Cgref1 | 2.98E-06 | 0.490421162 | 0.571 | 0.097 | 0.037467 | Visceral Endoderm |
| Elovl1 | 3.01E-06 | 0.541223773 | 1 | 0.875 | 0.037856 | Visceral Endoderm |
| Trmt1 | 3.20E-06 | 0.292232754 | 1 | 1 | 0.040228 | Visceral Endoderm |
| Bod1 | 3.48E-06 | 0.314591116 | 0.714 | 0.611 | 0.043789 | Visceral Endoderm |
| Mrps24 | 3.53E-06 | 0.425545825 | 0.857 | 0.917 | 0.044331 | Visceral Endoderm |
| Ric8 | 3.55E-06 | 0.264833637 | 1 | 0.819 | 0.0447 | Visceral Endoderm |
| Setd5 | 3.60E-06 | 0.287175171 | 1 | 1 | 0.045327 | Visceral Endoderm |
| Ripk3 | 3.63E-06 | 0.31756082 | 0.571 | 0.208 | 0.045674 | Visceral Endoderm |
| Ltbr | 3.67E-06 | 0.369824825 | 1 | 0.556 | 0.046203 | Visceral Endoderm |
| Snx27 | 3.90E-06 | 0.315495574 | 1 | 0.958 | 0.048997 | Visceral Endoderm |
| Rock1 | 4.15E-06 | 0.289679363 | 1 | 0.972 | 0.052215 | Visceral Endoderm |
| Cbx7 | 4.16E-06 | 0.505706068 | 1 | 0.694 | 0.052357 | Visceral Endoderm |
| Khsrp | 4.20E-06 | 0.311144991 | 1 | 1 | 0.052808 | Visceral Endoderm |
| Zfand2b | 4.20E-06 | 0.28860743 | 0.571 | 0.667 | 0.052848 | Visceral Endoderm |
| Stx7 | 4.35E-06 | 0.989128434 | 1 | 0.917 | 0.054736 | Visceral Endoderm |
| Fam195a | 4.53E-06 | 0.548623606 | 0.571 | 0.611 | 0.056952 | Visceral Endoderm |
| Scamp5 | 4.54E-06 | 0.469762836 | 1 | 0.778 | 0.057038 | Visceral Endoderm |
| Rhebl1 | 4.66E-06 | 0.507156659 | 0.714 | 0.75 | 0.058565 | Visceral Endoderm |
| Ppan | 4.90E-06 | 0.761143758 | 1 | 1 | 0.061591 | Visceral Endoderm |
| Slc38a7 | 4.94E-06 | 0.421390059 | 1 | 0.861 | 0.062182 | Visceral Endoderm |
| Bop1 | 4.95E-06 | 0.675613014 | 1 | 0.972 | 0.062228 | Visceral Endoderm |
| Glmp | 5.03E-06 | 0.471792543 | 1 | 0.986 | 0.063255 | Visceral Endoderm |
| Bckdk | 5.24E-06 | 0.450557846 | 1 | 0.986 | 0.065944 | Visceral Endoderm |
| Eps8 | 5.30E-06 | 0.293825158 | 0.857 | 0.264 | 0.066691 | Visceral Endoderm |
| Ascc2 | 5.35E-06 | 0.53335228 | 1 | 0.986 | 0.067289 | Visceral Endoderm |
| Stard10 | 5.48E-06 | 0.341997393 | 1 | 0.556 | 0.068857 | Visceral Endoderm |
| Sin3b | 5.71E-06 | 0.265037427 | 1 | 0.972 | 0.071771 | Visceral Endoderm |
| Cnot6 | 6.30E-06 | 0.278917296 | 1 | 1 | 0.079236 | Visceral Endoderm |
| Clcn7 | 6.31E-06 | 0.387519576 | 0.857 | 0.972 | 0.079407 | Visceral Endoderm |
| 2310061I0 | 6.33E-06 | 0.33609605 | 0.857 | 0.806 | 0.079627 | Visceral Endoderm |
| Mtmr7 | 6.38E-06 | 0.289210282 | 1 | 0.889 | 0.08024 | Visceral Endoderm |
| Grb7 | 6.79E-06 | 0.740383223 | 1 | 1 | 0.085437 | Visceral Endoderm |
| Lace1 | 7.00E-06 | 0.320656705 | 0.714 | 0.278 | 0.088028 | Visceral Endoderm |
| Ddhd2 | 7.08E-06 | 0.255214341 | 1 | 0.972 | 0.089067 | Visceral Endoderm |
| Rhox5 | 7.38E-06 | 1.596177818 | 1 | 0.375 | 0.092786 | Visceral Endoderm |
| Ell | 7.39E-06 | 0.434896922 | 0.857 | 0.792 | 0.092976 | Visceral Endoderm |
| Styx | 7.68E-06 | 0.491849054 | 0.857 | 0.944 | 0.096557 | Visceral Endoderm |
| Smyd5 | 8.15E-06 | 0.444193429 | 0.857 | 0.986 | 0.102459 | Visceral Endoderm |
| Arglu1 | 8.18E-06 | 0.429033385 | 1 | 1 | 0.102821 | Visceral Endoderm |
| Arhgap18 | 8.43E-06 | 0.357836073 | 0.857 | 0.389 | 0.105957 | Visceral Endoderm |
| 1700017B( | 8.63E-06 | 0.378534148 | 1 | 0.792 | 0.108557 | Visceral Endoderm |
| Arhgap10 | 8.66E-06 | 0.283616103 | 1 | 0.681 | 0.108878 | Visceral Endoderm |
| Rbm38 | 9.02E-06 | 0.391171289 | 1 | 0.931 | 0.113406 | Visceral Endoderm |
| Slc37a1 | 9.40E-06 | 0.287578862 | 0.571 | 0.222 | 0.118224 | Visceral Endoderm |
| Slc33a1 | 9.46E-06 | 0.332234887 | 1 | 0.694 | 0.119006 | Visceral Endoderm |
| Zfp654 | 9.71E-06 | 0.260368319 | 0.857 | 0.319 | 0.122169 | Visceral Endoderm |
| Gcdh | 1.00E-05 | 0.335137848 | 0.857 | 0.861 | 0.125846 | Visceral Endoderm |

| | | | | | | |
|---|---|---|---|---|---|---|
| Vps16 | 1.00E-05 | 0.447329272 | 0.857 | 0.958 | 0.125896 | Visceral Endoderm |
| Rnf181 | 1.05E-05 | 0.717829379 | 1 | 1 | 0.131936 | Visceral Endoderm |
| Arl2 | 1.06E-05 | 0.282710745 | 0.857 | 0.972 | 0.133492 | Visceral Endoderm |
| Abcg2 | 1.13E-05 | 0.40868164 | 1 | 0.903 | 0.141553 | Visceral Endoderm |
| Lxn | 1.19E-05 | 0.645095973 | 1 | 0.75 | 0.149196 | Visceral Endoderm |
| Aldh7a1 | 1.20E-05 | 0.524811628 | 1 | 0.986 | 0.151069 | Visceral Endoderm |
| Acadm | 1.23E-05 | 0.364491483 | 0.857 | 0.944 | 0.155075 | Visceral Endoderm |
| Agrn | 1.23E-05 | 0.535365337 | 1 | 1 | 0.155257 | Visceral Endoderm |
| Snf8 | 1.25E-05 | 0.253929552 | 1 | 1 | 0.157635 | Visceral Endoderm |
| Rbks | 1.26E-05 | 0.305967301 | 0.571 | 0.319 | 0.157905 | Visceral Endoderm |
| Cog3 | 1.28E-05 | 0.263407596 | 0.714 | 0.819 | 0.161247 | Visceral Endoderm |
| Wee1 | 1.34E-05 | 0.520631083 | 0.857 | 0.681 | 0.168915 | Visceral Endoderm |
| Ube2e1 | 1.38E-05 | 0.290749154 | 1 | 0.986 | 0.174009 | Visceral Endoderm |
| Atxn2l | 1.46E-05 | 0.415893875 | 1 | 1 | 0.183003 | Visceral Endoderm |
| Lpcat4 | 1.46E-05 | 0.385018508 | 0.857 | 0.667 | 0.183327 | Visceral Endoderm |
| Slc2a3 | 1.50E-05 | 0.853043087 | 1 | 1 | 0.188359 | Visceral Endoderm |
| Ptdss2 | 1.53E-05 | 0.291537401 | 1 | 0.972 | 0.192543 | Visceral Endoderm |
| Xlr3c | 1.56E-05 | 0.542670962 | 1 | 0.375 | 0.196496 | Visceral Endoderm |
| Dhrs1 | 1.62E-05 | 0.267568617 | 0.857 | 0.889 | 0.204212 | Visceral Endoderm |
| Cyba | 1.73E-05 | 1.100437226 | 1 | 0.986 | 0.216998 | Visceral Endoderm |
| Tmprss2 | 1.74E-05 | 1.06013731 | 1 | 0.722 | 0.218321 | Visceral Endoderm |
| Itsn2 | 1.78E-05 | 0.329878159 | 1 | 0.667 | 0.224356 | Visceral Endoderm |
| Aifm3 | 1.81E-05 | 0.364429786 | 0.857 | 0.181 | 0.227593 | Visceral Endoderm |
| Abcc4 | 1.83E-05 | 0.305941345 | 0.714 | 0.389 | 0.229714 | Visceral Endoderm |
| Stard4 | 1.84E-05 | 0.271961596 | 1 | 0.889 | 0.231954 | Visceral Endoderm |
| Crlf1 | 1.86E-05 | 0.288966618 | 0.571 | 0.056 | 0.234219 | Visceral Endoderm |
| Lpcat3 | 1.90E-05 | 0.453154093 | 1 | 0.958 | 0.239486 | Visceral Endoderm |
| Tsr1 | 1.93E-05 | 0.307913408 | 0.857 | 1 | 0.242633 | Visceral Endoderm |
| Ank | 1.95E-05 | 0.405563761 | 1 | 0.639 | 0.245142 | Visceral Endoderm |
| Ankzf1 | 1.96E-05 | 0.287631997 | 0.857 | 0.861 | 0.246388 | Visceral Endoderm |
| Klf6 | 1.97E-05 | 0.820469595 | 1 | 0.972 | 0.248119 | Visceral Endoderm |
| Arid3c | 1.98E-05 | 0.411993981 | 0.714 | 0.25 | 0.24869 | Visceral Endoderm |
| Pop5 | 2.01E-05 | 0.381378517 | 1 | 0.944 | 0.25315 | Visceral Endoderm |
| Ezr | 2.02E-05 | 0.790038246 | 1 | 1 | 0.253979 | Visceral Endoderm |
| BC029214 | 2.07E-05 | 0.359787927 | 0.571 | 0.264 | 0.259751 | Visceral Endoderm |
| Cc2d1b | 2.09E-05 | 0.323435328 | 0.857 | 0.764 | 0.262652 | Visceral Endoderm |
| Phgdh | 2.10E-05 | 0.83591953 | 1 | 1 | 0.264378 | Visceral Endoderm |
| Gata4 | 2.14E-05 | 0.839748016 | 1 | 0.375 | 0.26917 | Visceral Endoderm |
| Podxl | 2.20E-05 | 1.048720871 | 1 | 1 | 0.277036 | Visceral Endoderm |
| Gm21833 | 2.25E-05 | 0.337803235 | 1 | 0.972 | 0.282589 | Visceral Endoderm |
| Cd2ap | 2.41E-05 | 0.315498725 | 1 | 0.917 | 0.303645 | Visceral Endoderm |
| Gpr137b | 2.45E-05 | 0.276109049 | 1 | 0.875 | 0.30831 | Visceral Endoderm |
| Clic6 | 2.48E-05 | 0.637075273 | 1 | 0.625 | 0.311709 | Visceral Endoderm |
| Ifrd2 | 2.57E-05 | 0.335968261 | 0.857 | 0.931 | 0.323668 | Visceral Endoderm |
| Mb21d2 | 2.58E-05 | 0.289315329 | 0.571 | 0.611 | 0.323901 | Visceral Endoderm |
| Mon1a | 2.65E-05 | 0.252349223 | 0.857 | 0.5 | 0.33316 | Visceral Endoderm |
| Timm44 | 2.70E-05 | 0.340504359 | 1 | 0.972 | 0.339142 | Visceral Endoderm |
| Gps1 | 2.86E-05 | 0.453754991 | 1 | 1 | 0.359197 | Visceral Endoderm |
| Zfp655 | 2.87E-05 | 0.379086501 | 0.857 | 0.861 | 0.360421 | Visceral Endoderm |
| Fuk | 2.92E-05 | 0.361950148 | 0.857 | 0.542 | 0.367434 | Visceral Endoderm |
| S100g | 2.98E-05 | 1.393525031 | 1 | 0.347 | 0.375296 | Visceral Endoderm |
| Pla2g12b | 3.06E-05 | 0.738514555 | 0.857 | 0.236 | 0.384597 | Visceral Endoderm |
| Tiparp | 3.08E-05 | 0.57315254 | 1 | 1 | 0.387485 | Visceral Endoderm |
| Tbc1d17 | 3.18E-05 | 0.351057178 | 1 | 0.986 | 0.399922 | Visceral Endoderm |
| Igf2r | 3.28E-05 | 0.313104693 | 1 | 1 | 0.412833 | Visceral Endoderm |
| Wibg | 3.28E-05 | 0.3404035 | 1 | 0.931 | 0.41304 | Visceral Endoderm |
| Sap130 | 3.30E-05 | 0.487006028 | 1 | 0.972 | 0.415429 | Visceral Endoderm |
| Srrm2 | 3.39E-05 | 0.401831247 | 1 | 1 | 0.426376 | Visceral Endoderm |

| | | | | | | |
|---|---|---|---|---|---|---|
| Ehd1 | 3.48E-05 | 0.513004504 | 1 | 0.944 | 0.437804 | Visceral Endoderm |
| Prr5 | 3.52E-05 | 0.302289439 | 0.857 | 0.611 | 0.443289 | Visceral Endoderm |
| Btaf1 | 3.89E-05 | 0.264888803 | 1 | 0.972 | 0.489815 | Visceral Endoderm |
| Bag2 | 4.25E-05 | 0.340622938 | 0.857 | 0.875 | 0.534102 | Visceral Endoderm |
| Rabggtb | 4.41E-05 | 0.487455151 | 1 | 0.986 | 0.554618 | Visceral Endoderm |
| Gm5884 | 4.44E-05 | 0.250091732 | 0.714 | 0.722 | 0.558688 | Visceral Endoderm |
| Tdh | 4.61E-05 | 0.498300383 | 1 | 0.431 | 0.579264 | Visceral Endoderm |
| Ftsj2 | 4.71E-05 | 0.266008392 | 1 | 0.736 | 0.59227 | Visceral Endoderm |
| Cobll1 | 4.93E-05 | 0.262651686 | 1 | 0.625 | 0.619491 | Visceral Endoderm |
| Tpd52l1 | 4.97E-05 | 0.302559159 | 0.429 | 0.681 | 0.624494 | Visceral Endoderm |
| Mau2 | 4.97E-05 | 0.368858825 | 1 | 1 | 0.624526 | Visceral Endoderm |
| Ndufs7 | 5.14E-05 | 0.257097148 | 0.857 | 1 | 0.646226 | Visceral Endoderm |
| Man2b1 | 5.20E-05 | 0.360895381 | 1 | 0.931 | 0.653692 | Visceral Endoderm |
| Yipf2 | 5.40E-05 | 0.314500593 | 0.571 | 0.694 | 0.678562 | Visceral Endoderm |
| Apba3 | 5.67E-05 | 0.269487816 | 0.857 | 0.819 | 0.713631 | Visceral Endoderm |
| Tead4 | 5.69E-05 | 0.317287457 | 1 | 0.361 | 0.715832 | Visceral Endoderm |
| Slc9a3r1 | 5.77E-05 | 1.068080511 | 1 | 1 | 0.725903 | Visceral Endoderm |
| Gng12 | 5.85E-05 | 0.329099216 | 1 | 0.625 | 0.73582 | Visceral Endoderm |
| Ppap2c | 5.85E-05 | 0.450640899 | 1 | 0.722 | 0.73607 | Visceral Endoderm |
| Tagap1 | 6.21E-05 | 0.307278662 | 0.714 | 0.958 | 0.780801 | Visceral Endoderm |
| Cenpt | 6.49E-05 | 0.399708035 | 1 | 0.972 | 0.816615 | Visceral Endoderm |
| Sat1 | 6.62E-05 | 0.814071081 | 1 | 0.958 | 0.832189 | Visceral Endoderm |
| Srfbp1 | 6.75E-05 | 0.709673689 | 1 | 0.903 | 0.849434 | Visceral Endoderm |
| Vps35 | 6.84E-05 | 0.737700158 | 1 | 1 | 0.860466 | Visceral Endoderm |
| Abhd17c | 7.14E-05 | 0.330659419 | 1 | 0.861 | 0.897348 | Visceral Endoderm |
| Wdfy3 | 7.16E-05 | 0.396932593 | 1 | 1 | 0.900698 | Visceral Endoderm |
| Pnpla6 | 7.46E-05 | 0.349556176 | 0.857 | 0.847 | 0.938358 | Visceral Endoderm |
| Bri3 | 7.46E-05 | 0.274703396 | 0.857 | 0.694 | 0.938577 | Visceral Endoderm |
| Gm17555 | 7.88E-05 | 0.264530857 | 1 | 0.972 | 0.991189 | Visceral Endoderm |
| Mcts1 | 8.53E-05 | 0.397102357 | 1 | 0.889 | 1 | Visceral Endoderm |
| Smg5 | 8.71E-05 | 0.289618751 | 1 | 1 | 1 | Visceral Endoderm |
| C230052I1 | 8.86E-05 | 0.257086111 | 0.571 | 0.5 | 1 | Visceral Endoderm |
| Exoc6 | 9.38E-05 | 0.414443055 | 1 | 0.722 | 1 | Visceral Endoderm |
| Pabpc1 | 9.52E-05 | 0.386248076 | 1 | 1 | 1 | Visceral Endoderm |
| Phlda2 | 9.66E-05 | 1.271283216 | 1 | 0.5 | 1 | Visceral Endoderm |
| Arfgef2 | 9.72E-05 | 0.25217915 | 0.857 | 0.708 | 1 | Visceral Endoderm |
| Kat2a | 0.000105 | 0.272343171 | 0.857 | 0.986 | 1 | Visceral Endoderm |
| Pepd | 0.000107 | 0.328634824 | 1 | 1 | 1 | Visceral Endoderm |
| N4bp3 | 0.000111 | 0.269399773 | 0.429 | 0.667 | 1 | Visceral Endoderm |
| Limk2 | 0.000112 | 0.308223076 | 1 | 1 | 1 | Visceral Endoderm |
| Pls1 | 0.000113 | 0.350912245 | 0.857 | 0.361 | 1 | Visceral Endoderm |
| Rnf14 | 0.000119 | 0.37057376 | 0.857 | 0.931 | 1 | Visceral Endoderm |
| Rbpms2 | 0.000121 | 0.578521788 | 1 | 0.889 | 1 | Visceral Endoderm |
| Smad2 | 0.000123 | 0.272455634 | 1 | 0.958 | 1 | Visceral Endoderm |
| Lrpprc | 0.000126 | 0.315227362 | 1 | 1 | 1 | Visceral Endoderm |
| Plin3 | 0.000129 | 0.633314899 | 1 | 0.986 | 1 | Visceral Endoderm |
| Vamp2 | 0.000133 | 0.262718289 | 0.857 | 0.889 | 1 | Visceral Endoderm |
| Acsl3 | 0.000143 | 0.345026024 | 1 | 0.958 | 1 | Visceral Endoderm |
| Idh3g | 0.000145 | 0.34613571 | 1 | 0.986 | 1 | Visceral Endoderm |
| Rras2 | 0.000152 | 0.4756059 | 0.857 | 0.653 | 1 | Visceral Endoderm |
| Stub1 | 0.000155 | 0.280713155 | 1 | 1 | 1 | Visceral Endoderm |
| Fga | 0.000156 | 0.307789229 | 0.714 | 0.208 | 1 | Visceral Endoderm |
| Qtrt1 | 0.000157 | 0.373431461 | 0.857 | 0.931 | 1 | Visceral Endoderm |
| Atp1a1 | 0.000159 | 0.3374637 | 1 | 1 | 1 | Visceral Endoderm |
| Gmpr2 | 0.000162 | 0.357768217 | 0.714 | 0.931 | 1 | Visceral Endoderm |
| Cotl1 | 0.000163 | 0.845020163 | 1 | 0.958 | 1 | Visceral Endoderm |
| Tom1 | 0.000165 | 0.350715074 | 1 | 0.792 | 1 | Visceral Endoderm |
| S100a1 | 0.000167 | 0.577850712 | 0.857 | 0.444 | 1 | Visceral Endoderm |

| | | | | | | |
|---|---|---|---|---|---|---|
| Fcgrt | 0.000167 | 0.550924485 | 0.714 | 0.333 | 1 | Visceral Endoderm |
| Rab20 | 0.000169 | 0.32701336 | 0.857 | 0.333 | 1 | Visceral Endoderm |
| Cdca2 | 0.00017 | 0.665445333 | 1 | 1 | 1 | Visceral Endoderm |
| Nckap1 | 0.000178 | 0.330925539 | 1 | 0.986 | 1 | Visceral Endoderm |
| Idh3b | 0.000179 | 0.404226153 | 1 | 0.986 | 1 | Visceral Endoderm |
| Isca2 | 0.00018 | 0.516684961 | 1 | 0.889 | 1 | Visceral Endoderm |
| Vmp1 | 0.000188 | 0.439262052 | 1 | 0.986 | 1 | Visceral Endoderm |
| Grhpr | 0.000189 | 0.506070004 | 1 | 0.944 | 1 | Visceral Endoderm |
| Rrp9 | 0.000196 | 0.472273233 | 1 | 0.986 | 1 | Visceral Endoderm |
| Taf1d | 0.000204 | 0.878696556 | 1 | 1 | 1 | Visceral Endoderm |
| Ndufaf4 | 0.000206 | 0.317040804 | 1 | 0.972 | 1 | Visceral Endoderm |
| Fam134a | 0.000208 | 0.309562924 | 0.714 | 0.708 | 1 | Visceral Endoderm |
| E130012A1 | 0.000209 | 0.33758651 | 1 | 0.611 | 1 | Visceral Endoderm |
| Katnb1 | 0.000213 | 0.316723571 | 0.857 | 0.847 | 1 | Visceral Endoderm |
| Mapkap1 | 0.000217 | 0.25391042 | 0.714 | 0.875 | 1 | Visceral Endoderm |
| Sqstm1 | 0.000217 | 0.693315576 | 1 | 0.986 | 1 | Visceral Endoderm |
| Zfand5 | 0.000226 | 0.365234373 | 1 | 1 | 1 | Visceral Endoderm |
| Zcchc6 | 0.000228 | 0.337725583 | 1 | 0.861 | 1 | Visceral Endoderm |
| E430025E2 | 0.000231 | 0.260699421 | 1 | 0.958 | 1 | Visceral Endoderm |
| Mrpl2 | 0.000241 | 0.495533414 | 0.857 | 0.986 | 1 | Visceral Endoderm |
| Gm21814 | 0.000246 | 0.347153855 | 0.286 | 0.611 | 1 | Visceral Endoderm |
| Nup85 | 0.000248 | 0.541223802 | 1 | 1 | 1 | Visceral Endoderm |
| Tmem51 | 0.00027 | 0.580581363 | 0.857 | 0.5 | 1 | Visceral Endoderm |
| Sord | 0.000279 | 0.480816532 | 0.857 | 0.417 | 1 | Visceral Endoderm |
| Xpot | 0.000282 | 0.309482138 | 1 | 0.986 | 1 | Visceral Endoderm |
| Akr7a5 | 0.000288 | 0.285260636 | 1 | 0.972 | 1 | Visceral Endoderm |
| Rere | 0.000289 | 0.252362284 | 1 | 0.986 | 1 | Visceral Endoderm |
| Fus | 0.00029 | 0.4473339 | 1 | 1 | 1 | Visceral Endoderm |
| Dus1l | 0.000299 | 0.278500189 | 1 | 0.986 | 1 | Visceral Endoderm |
| Ubiad1 | 0.00031 | 0.356703165 | 0.857 | 0.847 | 1 | Visceral Endoderm |
| Atp11b | 0.000329 | 0.270061604 | 1 | 0.972 | 1 | Visceral Endoderm |
| Atp6v1c1 | 0.000333 | 0.4997896 | 1 | 0.972 | 1 | Visceral Endoderm |
| Agpat1 | 0.000334 | 0.312932953 | 1 | 1 | 1 | Visceral Endoderm |
| Emb | 0.000337 | 0.690954722 | 1 | 1 | 1 | Visceral Endoderm |
| Rhpn1 | 0.000337 | 0.491022129 | 1 | 0.903 | 1 | Visceral Endoderm |
| Cabin1 | 0.000346 | 0.264020534 | 0.857 | 0.931 | 1 | Visceral Endoderm |
| Ppp1r14b | 0.000351 | 0.315402876 | 0.714 | 0.847 | 1 | Visceral Endoderm |
| Pip5k1c | 0.000378 | 0.371478131 | 1 | 0.944 | 1 | Visceral Endoderm |
| Txndc17 | 0.00038 | 0.695552996 | 1 | 0.972 | 1 | Visceral Endoderm |
| Ifrd1 | 0.000386 | 0.27342846 | 1 | 1 | 1 | Visceral Endoderm |
| Gmppa | 0.000404 | 0.263381948 | 0.714 | 0.875 | 1 | Visceral Endoderm |
| Sertad1 | 0.000419 | 0.519777793 | 0.714 | 0.681 | 1 | Visceral Endoderm |
| Ep400 | 0.000421 | 0.253581455 | 1 | 1 | 1 | Visceral Endoderm |
| Abcf3 | 0.000427 | 0.318358474 | 0.857 | 0.986 | 1 | Visceral Endoderm |
| Depdc7 | 0.000433 | 0.312327514 | 0.429 | 0.528 | 1 | Visceral Endoderm |
| Id1 | 0.000447 | 1.164567693 | 1 | 0.958 | 1 | Visceral Endoderm |
| Bet1l | 0.000455 | 0.307535526 | 0.714 | 0.806 | 1 | Visceral Endoderm |
| Pcbp2 | 0.000464 | 0.293012675 | 1 | 1 | 1 | Visceral Endoderm |
| Nomo1 | 0.000489 | 0.374075356 | 1 | 0.972 | 1 | Visceral Endoderm |
| Mtif2 | 0.000496 | 0.275301071 | 1 | 0.986 | 1 | Visceral Endoderm |
| Stat6 | 0.000504 | 0.279435668 | 0.857 | 0.556 | 1 | Visceral Endoderm |
| Map1s | 0.000507 | 0.30782363 | 1 | 0.903 | 1 | Visceral Endoderm |
| Flnb | 0.000511 | 0.256640583 | 1 | 1 | 1 | Visceral Endoderm |
| Odc1 | 0.000544 | 0.464941381 | 1 | 1 | 1 | Visceral Endoderm |
| Ap1g2 | 0.000553 | 0.25535448 | 0.714 | 0.556 | 1 | Visceral Endoderm |
| Tdrp | 0.000564 | 0.296877147 | 0.857 | 0.347 | 1 | Visceral Endoderm |
| Ubxn4 | 0.000585 | 0.359472909 | 1 | 0.986 | 1 | Visceral Endoderm |
| Mllt10 | 0.000594 | 0.321223969 | 1 | 1 | 1 | Visceral Endoderm |

| | | | | | | |
|---|---|---|---|---|---|---|
| Afp | 0.000607 | 0.967867002 | 0.857 | 0.764 | 1 | Visceral Endoderm |
| Safb2 | 0.000646 | 0.343409105 | 1 | 0.986 | 1 | Visceral Endoderm |
| Rap1b | 0.000686 | 0.428196239 | 1 | 0.806 | 1 | Visceral Endoderm |
| Hdgfrp2 | 0.000718 | 0.757514619 | 1 | 1 | 1 | Visceral Endoderm |
| Gstm1 | 0.000726 | 0.672374186 | 1 | 0.903 | 1 | Visceral Endoderm |
| Plscr1 | 0.000727 | 0.380249198 | 1 | 0.931 | 1 | Visceral Endoderm |
| 0610011F( | 0.000727 | 0.291598493 | 0.714 | 0.917 | 1 | Visceral Endoderm |
| Lamtor5 | 0.000729 | 0.310068461 | 1 | 0.972 | 1 | Visceral Endoderm |
| Mad2l1bp | 0.000739 | 0.29563563 | 0.571 | 0.833 | 1 | Visceral Endoderm |
| Rragc | 0.000746 | 0.266836528 | 0.857 | 0.514 | 1 | Visceral Endoderm |
| Tpp1 | 0.000747 | 0.41294698 | 0.857 | 1 | 1 | Visceral Endoderm |
| Cmpk1 | 0.000755 | 0.34764301 | 1 | 1 | 1 | Visceral Endoderm |
| Tmem55b | 0.000769 | 0.284899139 | 1 | 0.889 | 1 | Visceral Endoderm |
| Ftl1 | 0.000783 | 0.700046849 | 1 | 1 | 1 | Visceral Endoderm |
| Pdzd3 | 0.000801 | 0.634854145 | 0.714 | 0.139 | 1 | Visceral Endoderm |
| Commd9 | 0.000802 | 0.268336046 | 0.714 | 0.903 | 1 | Visceral Endoderm |
| Rps6kb2 | 0.000809 | 0.289241379 | 0.857 | 0.889 | 1 | Visceral Endoderm |
| Atp6v1d | 0.000811 | 0.620880922 | 1 | 0.986 | 1 | Visceral Endoderm |
| Gab1 | 0.000813 | 0.340025125 | 1 | 0.75 | 1 | Visceral Endoderm |
| Med29 | 0.000822 | 0.252233788 | 1 | 0.917 | 1 | Visceral Endoderm |
| Snx29 | 0.000824 | 0.255799054 | 0.714 | 0.319 | 1 | Visceral Endoderm |
| Pecr | 0.000843 | 0.285529208 | 0.714 | 0.833 | 1 | Visceral Endoderm |
| Chd4 | 0.000876 | 0.400936966 | 1 | 1 | 1 | Visceral Endoderm |
| Gm3550 | 0.000943 | 0.291306585 | 1 | 1 | 1 | Visceral Endoderm |
| Uqcrq | 0.001 | 0.302381227 | 1 | 0.986 | 1 | Visceral Endoderm |
| Gm10116 | 0.001061 | 0.650870044 | 1 | 1 | 1 | Visceral Endoderm |
| Rpl10a | 0.001075 | 0.635436189 | 1 | 1 | 1 | Visceral Endoderm |
| Snx9 | 0.001104 | 0.393291401 | 1 | 0.5 | 1 | Visceral Endoderm |
| Slc7a6 | 0.001218 | 0.275568425 | 1 | 1 | 1 | Visceral Endoderm |
| Serpinf2 | 0.001301 | 0.963714923 | 0.571 | 0.25 | 1 | Visceral Endoderm |
| Scpep1 | 0.001312 | 0.697037668 | 1 | 0.986 | 1 | Visceral Endoderm |
| Tctn3 | 0.001323 | 0.273916458 | 0.571 | 0.472 | 1 | Visceral Endoderm |
| Cdc25c | 0.001416 | 0.316565647 | 0.857 | 0.903 | 1 | Visceral Endoderm |
| Shmt2 | 0.00143 | 0.533147155 | 1 | 0.931 | 1 | Visceral Endoderm |
| Fem1b | 0.001436 | 0.262008704 | 1 | 0.972 | 1 | Visceral Endoderm |
| Jtb | 0.001445 | 0.326886282 | 0.857 | 0.958 | 1 | Visceral Endoderm |
| Cdkn3 | 0.001493 | 0.287854297 | 0.429 | 0.583 | 1 | Visceral Endoderm |
| Rnf128 | 0.0015 | 0.343072669 | 0.571 | 0.833 | 1 | Visceral Endoderm |
| Id2 | 0.001575 | 0.797791365 | 1 | 1 | 1 | Visceral Endoderm |
| Tmem9b | 0.001579 | 0.37388917 | 0.857 | 0.792 | 1 | Visceral Endoderm |
| Tor1aip2 | 0.001743 | 0.290991297 | 1 | 1 | 1 | Visceral Endoderm |
| Acaa2 | 0.00177 | 0.427302855 | 0.857 | 0.778 | 1 | Visceral Endoderm |
| Gm21750 | 0.001774 | 0.285772962 | 1 | 0.986 | 1 | Visceral Endoderm |
| Maz | 0.001857 | 0.447361144 | 1 | 0.986 | 1 | Visceral Endoderm |
| Nhp2 | 0.001916 | 0.364049657 | 1 | 1 | 1 | Visceral Endoderm |
| Rhog | 0.001935 | 0.329800924 | 0.714 | 0.833 | 1 | Visceral Endoderm |
| Zfp280b | 0.001937 | 0.372228173 | 1 | 1 | 1 | Visceral Endoderm |
| Slc16a13 | 0.001949 | 0.373399403 | 1 | 0.653 | 1 | Visceral Endoderm |
| Nrde2 | 0.001972 | 0.262877543 | 0.714 | 0.681 | 1 | Visceral Endoderm |
| Setmar | 0.001972 | 0.328679272 | 0.571 | 0.208 | 1 | Visceral Endoderm |
| Grina | 0.002018 | 0.450724571 | 0.714 | 0.806 | 1 | Visceral Endoderm |
| Eif4g1 | 0.002122 | 0.520448996 | 1 | 1 | 1 | Visceral Endoderm |
| Cd63 | 0.002195 | 1.037827776 | 1 | 0.986 | 1 | Visceral Endoderm |
| 9430015G | 0.002333 | 0.390983278 | 1 | 0.986 | 1 | Visceral Endoderm |
| Asah1 | 0.002343 | 0.308376455 | 0.857 | 0.958 | 1 | Visceral Endoderm |
| Igf2bp2 | 0.002366 | 0.369729666 | 1 | 1 | 1 | Visceral Endoderm |
| Ppfibp2 | 0.002403 | 0.326274378 | 0.714 | 0.347 | 1 | Visceral Endoderm |
| Tnfrsf12a | 0.002434 | 0.768059108 | 0.857 | 0.361 | 1 | Visceral Endoderm |

| | | | | | | |
|---|---|---|---|---|---|---|
| Otub1 | 0.002445 | 0.255448706 | 1 | 1 | 1 | Visceral Endoderm |
| Trappc6b | 0.002465 | 0.303277044 | 0.857 | 0.931 | 1 | Visceral Endoderm |
| Atad3a | 0.002568 | 0.455864362 | 1 | 0.986 | 1 | Visceral Endoderm |
| 2210011C2 | 0.00268 | 0.447853537 | 0.571 | 0.139 | 1 | Visceral Endoderm |
| Phlda1 | 0.0027 | 0.267857596 | 1 | 0.611 | 1 | Visceral Endoderm |
| Fam21 | 0.002911 | 0.43550126 | 1 | 0.972 | 1 | Visceral Endoderm |
| Rbck1 | 0.002925 | 0.266051111 | 0.857 | 1 | 1 | Visceral Endoderm |
| Fth1 | 0.003258 | 1.0470764 | 1 | 1 | 1 | Visceral Endoderm |
| Cdk20 | 0.003346 | 0.332589164 | 0.571 | 0.375 | 1 | Visceral Endoderm |
| Phf13 | 0.003434 | 0.386026401 | 1 | 0.875 | 1 | Visceral Endoderm |
| Ugdh | 0.003705 | 0.267935001 | 1 | 0.972 | 1 | Visceral Endoderm |
| Slc35f5 | 0.003713 | 0.291715011 | 1 | 0.819 | 1 | Visceral Endoderm |
| Mapk6 | 0.003759 | 0.259201306 | 1 | 1 | 1 | Visceral Endoderm |
| Tmem120a | 0.003871 | 0.302411176 | 0.857 | 0.833 | 1 | Visceral Endoderm |
| Pip4k2c | 0.004074 | 0.291478385 | 1 | 0.875 | 1 | Visceral Endoderm |
| Nt5dc2 | 0.004295 | 0.369964829 | 1 | 0.986 | 1 | Visceral Endoderm |
| Prss8 | 0.004304 | 0.885998001 | 1 | 0.764 | 1 | Visceral Endoderm |
| Emg1 | 0.004314 | 0.528388099 | 1 | 1 | 1 | Visceral Endoderm |
| Cyp26a1 | 0.004485 | 0.726816981 | 1 | 0.389 | 1 | Visceral Endoderm |
| Pxdc1 | 0.004563 | 0.377088942 | 0.571 | 0.125 | 1 | Visceral Endoderm |
| Usp15 | 0.00464 | 0.40454862 | 1 | 0.944 | 1 | Visceral Endoderm |
| Ccdc61 | 0.004724 | 0.251012661 | 0.714 | 0.833 | 1 | Visceral Endoderm |
| Arhgap1 | 0.004855 | 0.384521015 | 1 | 0.986 | 1 | Visceral Endoderm |
| Dusp4 | 0.004873 | 0.49873059 | 1 | 0.75 | 1 | Visceral Endoderm |
| Slc35b1 | 0.005095 | 0.425896351 | 1 | 0.986 | 1 | Visceral Endoderm |
| Nop56 | 0.005294 | 0.600013405 | 1 | 1 | 1 | Visceral Endoderm |
| Atp5d | 0.005325 | 0.476388624 | 1 | 0.986 | 1 | Visceral Endoderm |
| Gyltl1b | 0.005366 | 0.251083235 | 1 | 0.833 | 1 | Visceral Endoderm |
| Sdc4 | 0.005414 | 0.574719646 | 1 | 1 | 1 | Visceral Endoderm |
| Fmr1nb | 0.005719 | 0.527793036 | 0.714 | 0.181 | 1 | Visceral Endoderm |
| Noc2l | 0.006063 | 0.300180254 | 0.857 | 0.903 | 1 | Visceral Endoderm |
| Scyl1 | 0.006106 | 0.411400018 | 1 | 0.917 | 1 | Visceral Endoderm |
| Snrnp48 | 0.006129 | 0.384576364 | 1 | 0.972 | 1 | Visceral Endoderm |
| Pigyl | 0.006172 | 0.374786423 | 0.857 | 0.875 | 1 | Visceral Endoderm |
| Tomm40 | 0.006264 | 0.28480318 | 1 | 1 | 1 | Visceral Endoderm |
| Yipf3 | 0.006304 | 0.260168683 | 1 | 0.931 | 1 | Visceral Endoderm |
| Fgg | 0.006344 | 0.309719357 | 0.571 | 0.333 | 1 | Visceral Endoderm |
| Clasrp | 0.006669 | 0.403527175 | 1 | 0.958 | 1 | Visceral Endoderm |
| Tmem180 | 0.00667 | 0.295363233 | 1 | 0.972 | 1 | Visceral Endoderm |
| Pacsin2 | 0.006751 | 0.332920173 | 1 | 0.944 | 1 | Visceral Endoderm |
| Dpp4 | 0.006962 | 0.425286115 | 1 | 0.847 | 1 | Visceral Endoderm |
| Mogat2 | 0.007213 | 0.392790046 | 1 | 0.931 | 1 | Visceral Endoderm |
| Lamp2 | 0.007383 | 0.312420105 | 1 | 1 | 1 | Visceral Endoderm |
| Nubp2 | 0.007398 | 0.340334482 | 1 | 0.986 | 1 | Visceral Endoderm |
| Cldn6 | 0.00776 | 0.424001647 | 1 | 0.986 | 1 | Visceral Endoderm |
| Scaf4 | 0.00805 | 0.319549554 | 1 | 0.958 | 1 | Visceral Endoderm |
| Tusc2 | 0.008101 | 0.267156665 | 0.857 | 0.958 | 1 | Visceral Endoderm |
| Tob1 | 0.008475 | 0.252073247 | 0.857 | 0.875 | 1 | Visceral Endoderm |
| Ybx3 | 0.008695 | 0.356890001 | 1 | 0.944 | 1 | Visceral Endoderm |
| Chchd3 | 0.008759 | 0.370140621 | 1 | 1 | 1 | Visceral Endoderm |
| Deb1 | 0.009003 | 0.383524645 | 0.857 | 0.917 | 1 | Visceral Endoderm |
| Cyb5a | 0.009055 | 0.510017381 | 1 | 0.986 | 1 | Visceral Endoderm |
| Snapin | 0.0091 | 0.387689255 | 0.857 | 0.889 | 1 | Visceral Endoderm |
| Sec31a | 0.009316 | 0.271850796 | 1 | 1 | 1 | Visceral Endoderm |
| Snrnp35 | 0.009474 | 0.276711749 | 0.571 | 0.583 | 1 | Visceral Endoderm |
| Gnpda1 | 0.009606 | 0.256814533 | 1 | 0.917 | 1 | Visceral Endoderm |
| Terf1 | 0.009708 | 0.268777349 | 1 | 1 | 1 | Visceral Endoderm |

5

e left cell clusters. Positive values indicate

| gene | Descriptio | Location |
|---|---|---|
| Nkx2-3 | NK2 home | Nucleus |
| Cldn4 | claudin 4 | Plasma Me |
| Atp2c2 | ATPase se | Cytoplasm |
| Akap5 | A-kinase a | Plasma Me |
| Krt7 | keratin 7 | Cytoplasm |
| Pyy | peptide YY | Extracellula |
| Cpm | carboxype | Plasma Me |
| Nkx2-6 | NK2 home | Nucleus |
| Isl1 | ISL LIM ho | Nucleus |
| Dok5 | docking pr | Plasma Me |
| Nepn | nephrocar | Extracellula |
| Bcam | basal cell a | Plasma Me |
| Npnt | nephronec | Plasma Me |
| Angpt2 | angiopoie | Extracellula |
| Igf2 | insulin like | Extracellula |
| Fblim1 | filamin bin | Plasma Me |
| Nrp1 | neuropilin | Plasma Me |
| Cldn3 | claudin 3 | Plasma Me |
| Tes | testin LIM | Plasma Me |
| Ripply3 | ripply tran | Nucleus |
| Efna1 | ephrin A1 | Plasma Me |
| Mmp11 | matrix met | Extracellula |
| Ldoc1 | LDOC1, re | Nucleus |
| Hs3st3b1 | heparan su | Cytoplasm |
| Wbp5 | - | - |
| Epcam | epithelial c | Plasma Me |
| Tpm1 | tropomyos | Plasma Me |
| Wfdc2 | WAP four- | Extracellula |
| Cd55 | CD55 mole | Plasma Me |
| Jup | junction pl | Plasma Me |
| Gm8113 | - | - |
| Spint1 | serine pep | Extracellula |
| F11r | F11 recept | Plasma Me |
| Cpn1 | carboxype | Extracellula |
| St14 | suppressic | Plasma Me |
| Arrdc1 | arrestin dc | Cytoplasm |
| Tmem59l | transmeml | Cytoplasm |
| Krt18 | keratin 18 | Cytoplasm |
| Spint2 | serine pep | Extracellula |
| Itga3 | integrin su | Plasma Me |
| Dsc2 | desmocoll | Plasma Me |
| Mapk13 | mitogen-a | Cytoplasm |
| Lpar3 | lysophosp | Plasma Me |
| Slc2a3 | solute carr | Plasma Me |
| Rasgrp3 | RAS guany | Cytoplasm |

| | | |
|---|---|---|
| Kitl | KIT ligand | Extracellula |
| Mterf3 | mitochonc | Cytoplasm |
| Krt8 | keratin 8 | Cytoplasm |
| Capn6 | calpain 6 | Cytoplasm |
| Cldn7 | claudin 7 | Plasma Me |
| Dusp9 | dual speci | Nucleus |
| Cldn6 | claudin 6 | Plasma Me |
| Tspan7 | tetraspanii | Plasma Me |
| Gas6 | growth arr | Extracellula |
| Fn1 | fibronectir | Extracellula |
| Tfpi | tissue facto | Extracellula |
| Sh3glb1 | SH3 doma | Cytoplasm |
| Bex1 | brain expr | Other |
| Tspan12 | tetraspanii | Plasma Me |
| Igfbp3 | insulin like | Extracellula |
| Gfpt2 | glutamine- | Cytoplasm |
| Apela | apelin rece | Extracellula |
| Afap1l1 | actin filam | Other |
| Amph | amphiphys | Plasma Me |
| Ngfrap1 | - | - |
| Pvrl2 | - | - |
| Parm1 | prostate ai | Extracellula |
| Tmem102 | transmeml | Plasma Me |
| Rcan2 | regulator c | Other |
| Vamp8 | vesicle ass | Plasma Me |
| Lad1 | ladinin 1 | Extracellula |
| Cmtm8 | CKLF like N | Extracellula |
| Spin2c | spindlin fa | Other |
| Gpc3 | glypican 3 | Plasma Me |
| Kcnk1 | potassium | Plasma Me |
| Lipa | lipase A, ly | Cytoplasm |
| Peg3 | paternally | Nucleus |
| Gprc5c | G protein- | Plasma Me |
| Slc16a1 | solute carr | Plasma Me |
| Tmsb10 | thymosin, | Cytoplasm |
| Frmd4b | FERM dom | Cytoplasm |
| Rbp1 | retinol bin | Extracellula |
| Ctxn1 | cortexin 1 | Other |
| Dkk3 | dickkopf W | Extracellula |
| Krt19 | keratin 19 | Cytoplasm |
| Cdc42ep1 | CDC42 eff | Extracellula |
| Podxl | podocalyx | Plasma Me |
| Cdh1 | cadherin 1 | Plasma Me |
| Gata3 | GATA binc | Nucleus |
| Crip2 | cysteine ri | Plasma Me |
| Marveld2 | MARVEL d | Plasma Me |
| Cdh3 | cadherin 3 | Plasma Me |
| Csrp2 | cysteine ar | Nucleus |
| Prss8 | protease, s | Extracellula |
| Smoc2 | SPARC rela | Extracellula |
| Tmem254c | transmeml | Other |
| Tspan9 | tetraspanii | Plasma Me |
| Mboat1 | membrane | Cytoplasm |
| Ube2h | ubiquitin c | Cytoplasm |
| Tmem254a | transmeml | Other |
| Tmem254b | transmeml | Other |
| Igfbp2 | insulin like | Extracellula |
| 1700047I17Rik2 | family with | Other |

| | | |
|---|---|---|
| Raet1d | retinoic ac | Plasma Me |
| Serpinb6c | serine (or ( | Other |
| Krt20 | keratin 20 | Cytoplasm |
| Vnn1 | vanin 1 | Plasma Me |
| Lsr | lipolysis sti | Nucleus |
| Cox4i2 | cytochrom | Cytoplasm |
| Fam107b | family with | Other |
| Habp2 | hyalurona | Extracellul |
| Tgfb2 | transformi | Extracellul |
| Ddah2 | dimethylar | Cytoplasm |
| Kctd12b | potassium | Plasma Me |
| 0610040J01Rik | chromoso | Nucleus |
| Sfrp5 | secreted fr | Plasma Me |
| Tuft1 | tuftelin 1 | Cytoplasm |
| Actb | actin beta | Cytoplasm |
| Calcoco1 | calcium bi | Nucleus |
| Car4 | carbonic a | Plasma Me |
| Nudt11 | nudix hydr | Cytoplasm |
| Bex4 | brain expr | Cytoplasm |
| Crb3 | crumbs fa | Plasma Me |
| Cpxm1 | carboxype | Extracellul |
| Ctnna1 | catenin alp | Plasma Me |
| Bsg | basigin (O | Plasma Me |
| Gpx2 | glutathion | Cytoplasm |
| Emb | embigin | Plasma Me |
| S100a11 | S100 calciu | Cytoplasm |
| Cd276 | CD276 mc | Plasma Me |
| Samd10 | sterile alpl | Other |
| Fam210b | family with | Extracellul |
| Hkdc1 | hexokinase | Cytoplasm |
| Tmem9 | transmeml | Cytoplasm |
| Cers5 | ceramide s | Cytoplasm |
| Casp8 | caspase 8 | Nucleus |
| S100a10 | S100 calciu | Cytoplasm |
| Hacd2 | 3-hydroxy | Cytoplasm |
| Trap1a | tumor reje | Cytoplasm |
| Cnn3 | calponin 3 | Cytoplasm |
| Frzb | frizzled rel | Extracellul |
| App | amyloid be | Plasma Me |
| Igsf21 | immunogl | Extracellul |
| Nucb1 | nucleobin | Cytoplasm |
| Slc38a4 | solute carr | Plasma Me |
| Dpp4 | dipeptidyl | Plasma Me |
| Kdelr2 | KDEL endo | Cytoplasm |
| Myl6 | myosin lig | Cytoplasm |
| Erlin2 | ER lipid ra | Plasma Me |
| Mdk | midkine | Extracellul |
| Tax1bp3 | Tax1 bindi | Cytoplasm |
| Pygl | glycogen p | Cytoplasm |
| Pdpn | podoplani | Plasma Me |
| Krtcap3 | keratinocy | Other |
| Cgn | cingulin | Plasma Me |
| Gpx8 | glutathion | Cytoplasm |
| Cd24a | CD24a ant | Plasma Me |
| Carhsp1 | calcium re | Cytoplasm |
| Laptm4a | lysosomal | Cytoplasm |
| Gbas | - | - |
| Pgrmc1 | progester | Plasma Me |

| | | |
|---|---|---|
| Sdcbp | syndecan l | Plasma Me |
| Npc2 | NPC intrac | Extracellul |
| Anxa6 | annexin A6 | Plasma Me |
| Pigs | phosphati | Cytoplasm |
| Dstn | destrin, ac | Cytoplasm |
| Fam3c | family with | Extracellul |
| Fam213a | family with | Extracellul |
| S1pr2 | sphingosir | Plasma Me |
| Afp | alpha feto | Extracellul |
| Plp2 | proteolipic | Cytoplasm |
| Clic1 | chloride in | Nucleus |
| Rnase4 | ribonuclea | Extracellul |
| 9530068E07Rik | chromoso | Other |
| Fam174b | family with | Other |
| Pdia3 | protein dis | Cytoplasm |
| Prmt2 | protein arg | Nucleus |
| Smim22 | small integ | Other |
| Mapre2 | microtubu | Cytoplasm |
| Tmco3 | transmeml | Other |
| Ap1m2 | adaptor re | Cytoplasm |
| Dap | death asso | Cytoplasm |
| Gm6169 | proteolipic | Other |
| Tmed2 | transmeml | Cytoplasm |
| Ctsd | cathepsin | Cytoplasm |
| Wbp1 | WW doma | Cytoplasm |
| Csrp1 | cysteine ar | Nucleus |
| Atpif1 | ATP synth | Cytoplasm |
| Mpzl1 | myelin pro | Plasma Me |
| Serinc2 | serine inco | Plasma Me |
| Pttg1ip | PTTG1 inte | Nucleus |
| Pld3 | phospholi | Cytoplasm |
| Cystm1 | cysteine-ri | Cytoplasm |
| Ddost | dolichyl-d | Cytoplasm |
| Ywhaz | tyrosine 3- | Cytoplasm |
| Ywhaq | tyrosine 3- | Cytoplasm |
| Arg1 | arginase 1 | Cytoplasm |
| Ppib | peptidylpr | Cytoplasm |
| Lasp1 | LIM and Sl | Cytoplasm |
| P4hb | prolyl 4-hy | Cytoplasm |
| Calr | calreticulir | Cytoplasm |
| B4galt1 | beta-1,4-g | Cytoplasm |
| Aktip | AKT intera | Cytoplasm |
| Tmem54 | transmeml | Other |
| Igsf9 | immunogl | Plasma Me |
| Hsp90b1 | heat shock | Cytoplasm |
| Suox | sulfite oxic | Cytoplasm |
| Grn | granulin p | Extracellul |
| Gdpd1 | glyceroph | Other |
| Ocln | occludin | Plasma Me |
| Tmprss2 | transmeml | Plasma Me |
| Pdlim1 | PDZ and L | Cytoplasm |
| Laptm4b | lysosomal | Cytoplasm |
| Tbx3 | T-box 3 | Nucleus |
| Ndfip1 | Nedd4 fan | Cytoplasm |
| Gadd45b | growth arr | Cytoplasm |
| Spink3 | - | - |
| Tmed9 | transmeml | Cytoplasm |
| Rnf128 | ring finger | Cytoplasm |

| | | |
|---|---|---|
| Gng11 | G protein s | Plasma Me |
| Klk8 | kallikrein r | Extracellul |
| M6pr | mannose- | Cytoplasm |
| Clu | clusterin | Cytoplasm |
| Cst3 | cystatin C | Extracellul |
| Pacsin2 | protein kir | Cytoplasm |
| Atp6v0e | ATPase H+ | Cytoplasm |
| Tma7 | translation | Other |
| Fam129b | family with | Cytoplasm |
| Slc31a1 | solute carr | Plasma Me |
| Cd63 | CD63 mole | Plasma Me |
| Arf4 | ADP ribosy | Cytoplasm |
| Ttr | transthyret | Extracellul |
| Serf1 | small EDRI | Other |
| Tmed4 | transmeml | Cytoplasm |
| Id2 | inhibitor o | Nucleus |
| Rbpms | RNA bindi | Cytoplasm |
| Tceal8 | transcriptio | Other |
| Cnn2 | calponin 2 | Cytoplasm |
| Rcn1 | reticulocal | Cytoplasm |
| Tbca | tubulin fol | Cytoplasm |
| Txn1 | thioredoxi | Cytoplasm |
| Itgb1 | integrin su | Plasma Me |
| Nudt14 | nudix hydr | Cytoplasm |
| Hmgn1 | high mobi | Nucleus |
| Tmem88 | transmeml | Plasma Me |
| Myl12b | myosin lig | Cytoplasm |
| Dhrs3 | dehydroge | Cytoplasm |
| Chrnb1 | cholinergic | Plasma Me |
| Surf4 | surfeit 4 | Cytoplasm |
| Tpm3-rs7 | tropomyos | Other |
| Slc9a3r2 | SLC9A3 re | Plasma Me |
| Mum1l1 | MUM1 like | Cytoplasm |
| Cdc37 | cell divisio | Cytoplasm |
| Slc2a1 | solute carr | Plasma Me |
| Itm2b | integral m | Plasma Me |
| Rcn2 | reticulocal | Cytoplasm |
| Myl12a | myosin lig | Cytoplasm |
| Tmed10 | transmeml | Cytoplasm |
| Slc39a8 | solute carr | Extracellul |
| Bambi | BMP and a | Plasma Me |
| Ssr3 | signal sequ | Cytoplasm |
| Pgm2 | phosphog | Cytoplasm |
| Ndufa4 | NDUFA4, r | Cytoplasm |
| Cdkn1c | cyclin-dep | Nucleus |
| Tpm3 | tropomyos | Cytoplasm |
| Cldn11 | claudin 11 | Plasma Me |
| Cdo1 | cysteine di | Cytoplasm |
| Sod1 | superoxide | Cytoplasm |
| Klf10 | Kruppel lik | Nucleus |
| Tpm4 | tropomyos | Cytoplasm |
| Rbbp7 | RB binding | Nucleus |
| Hexb | hexosamir | Cytoplasm |
| Paip1 | poly(A) bir | Cytoplasm |
| Lman2 | lectin, mar | Cytoplasm |
| Atox1 | antioxidan | Cytoplasm |
| Bcap31 | B cell rece | Cytoplasm |
| Cyba | cytochrom | Cytoplasm |

| | | |
|---|---|---|
| Pdia6 | protein dis | Cytoplasm |
| Serpinf2 | serpin fam | Extracellul |
| B4galnt2 | beta-1,4-N | Cytoplasm |
| Atxn10 | ataxin 10 | Cytoplasm |
| Bnip3 | BCL2 inter | Cytoplasm |
| Dbn1 | drebrin 1 | Cytoplasm |
| Hspa5 | heat shock | Cytoplasm |
| Copz1 | coatomer | Cytoplasm |
| Nucb2 | nucleobin | Nucleus |
| Nudt16l1 | nudix hydr | Cytoplasm |
| Cep19 | centrosom | Cytoplasm |
| Prdx6 | peroxired | Cytoplasm |
| Oaz2 | ornithine c | Cytoplasm |
| Clic6 | chloride in | Plasma Me |
| Ssr2 | signal seq | Cytoplasm |
| Spg21 | SPG21, ma | Plasma Me |
| Cttn | cortactin | Plasma Me |
| Rp9 | RP9, pre-r | Nucleus |
| Rbbp9 | RB binding | Nucleus |
| Spcs1 | signal pep | Cytoplasm |
| Etv5 | ETS varian | Nucleus |
| Arpc5 | actin relat | Cytoplasm |
| Fscn1 | fascin acti | Cytoplasm |
| Tmem50a | transmem | Plasma Me |
| 1110012L19Rik | chromoso | Other |
| Rpn1 | ribophorin | Cytoplasm |
| Reep5 | receptor a | Extracellul |
| Phlda2 | pleckstrin | Cytoplasm |
| Slc39a4 | solute carr | Plasma Me |
| Hyal2 | hyalurono | Cytoplasm |
| Tmem59 | transmem | Plasma Me |
| Vps36 | vacuolar p | Cytoplasm |
| Akr1a1 | aldo-keto | Cytoplasm |
| Mcfd2 | multiple c | Cytoplasm |
| Rasl11a | RAS like fa | Nucleus |
| Maged1 | MAGE fam | Plasma Me |
| 5730507C01Rik | RIKEN cDN | Other |
| Papss1 | 3'-phosph | Cytoplasm |
| Cops5 | COP9 sign | Nucleus |
| Prelid2 | PRELI dom | Cytoplasm |
| Slc25a4 | solute carr | Cytoplasm |
| Twsg1 | twisted ga | Extracellul |
| Slc44a2 | solute carr | Extracellul |
| Nrbp1 | nuclear re | Nucleus |
| Myadm | myeloid as | Nucleus |
| Cd81 | CD81 mol | Plasma Me |
| Arpc2 | actin relat | Cytoplasm |
| Nradd | neurotrop | Extracellul |
| Enpp3 | ectonucle | Plasma Me |
| Sdc1 | syndecan 1 | Plasma Me |
| Hif1a | hypoxia in | Nucleus |
| Zfand6 | zinc finger | Other |
| Stmn1 | stathmin 1 | Cytoplasm |
| Chmp2b | charged m | Cytoplasm |
| Tmub2 | transmem | Other |
| Tubb4a | tubulin be | Cytoplasm |
| Lman1 | lectin, mar | Cytoplasm |
| Snx6 | sorting ne | Cytoplasm |

| | | |
|---|---|---|
| Cstb | cystatin B | Cytoplasm |
| Myl9 | myosin lig | Cytoplasm |
| F2r | coagulatio | Plasma Me |
| Smpd1 | sphingomy | Cytoplasm |
| Smtnl2 | smoothelir | Other |
| Cnpy3 | canopy FG | Cytoplasm |
| Sh3bgrl3 | SH3 doma | Nucleus |
| Rbp4 | retinol bin | Extracellula |
| Ifi30 | IFI30, lysos | Cytoplasm |
| Anxa5 | annexin A5 | Plasma Me |
| Cd164 | CD164 mc | Plasma Me |
| Dazap2 | DAZ assoc | Nucleus |
| Stx7 | syntaxin 7 | Plasma Me |
| Tmbim6 | transmeml | Nucleus |
| Txndc12 | thioredoxi | Cytoplasm |
| C1galt1c1 | C1GALT1 s | Cytoplasm |
| Plat | plasminog | Extracellula |
| Zfp125 | zinc finger | Nucleus |
| Slit2 | slit guidan | Extracellula |
| Lmx1a | LIM home | Nucleus |
| Tmem255a | transmeml | Extracellula |
| Aplnr | apelin rece | Plasma Me |
| Col9a1 | collagen ty | Extracellula |
| Htra1 | HtrA serin | Extracellula |
| Car3 | carbonic a | Cytoplasm |
| Tgm2 | transgluta | Cytoplasm |
| Shh | sonic hedg | Extracellula |
| B3gnt7 | UDP-GlcN | Cytoplasm |
| Gng4 | G protein s | Plasma Me |
| Cthrc1 | collagen tr | Extracellula |
| Arhgef16 | Rho guani | Cytoplasm |
| Samd3 | sterile alph | Other |
| Kirrel3 | kirre like n | Extracellula |
| Nlgn3 | neuroligin | Plasma Me |
| Gnai1 | G protein s | Plasma Me |
| Wwp2 | WW doma | Cytoplasm |
| Tmem100 | transmeml | Plasma Me |
| Alcam | activated l | Plasma Me |
| Prickle1 | prickle pla | Nucleus |
| Sept6 | septin 6 | Cytoplasm |
| Moxd1 | monooxyg | Cytoplasm |
| Susd5 | sushi dom | Other |
| Vgll4 | vestigial lik | Nucleus |
| Krt19 | keratin 19 | Cytoplasm |
| Sox9 | SRY-box 9 | Nucleus |
| Rnd3 | Rho family | Cytoplasm |
| Egflam | EGF like, fi | Extracellula |
| Rab38 | RAB38, me | Cytoplasm |
| Sema3d | semaphori | Extracellula |
| Cd36 | CD36 mole | Plasma Me |
| St6galnac3 | ST6 N-ace | Cytoplasm |
| Tmod2 | tropomod | Cytoplasm |
| Mturn | maturin, n | Other |
| Plekhb1 | pleckstrin | Cytoplasm |
| Nkd1 | naked cuti | Other |
| Lypd6 | LY6/PLAUI | Extracellula |
| Sulf1 | sulfatase 1 | Cytoplasm |
| Gpm6b | glycoprote | Plasma Me |

| | | |
|---|---|---|
| Sox6 | SRY-box 6 | Nucleus |
| T | T-box trar | Nucleus |
| Arid3b | AT-rich int | Nucleus |
| Epb4.1l3 | - | - |
| Col5a1 | collagen ty | Extracellula |
| Anxa3 | annexin A3 | Cytoplasm |
| Scara3 | scavenger | Plasma Me |
| Nap1l5 | nucleosom | Other |
| Myl9 | myosin lig | Cytoplasm |
| Isg20 | interferon | Nucleus |
| Tpm2 | tropomyos | Cytoplasm |
| Gstm5 | glutathion | Cytoplasm |
| Usp18 | ubiquitin s | Cytoplasm |
| Smox | spermine ( | Cytoplasm |
| Clu | clusterin | Cytoplasm |
| Mnx1 | motor neu | Nucleus |
| Psme2b | protease ( | Other |
| Angptl2 | angiopoie | Extracellula |
| Gcnt1 | glucosami | Cytoplasm |
| Vtn | vitronectin | Extracellula |
| Cnn2 | calponin 2 | Cytoplasm |
| Lrig1 | leucine ric | Extracellula |
| Mmd | monocyte | Plasma Me |
| Gpt2 | glutamic-- | Cytoplasm |
| Col2a1 | collagen ty | Extracellula |
| Guca1a | guanylate | Cytoplasm |
| Tapbp | TAP bindir | Cytoplasm |
| Rarg | retinoic ac | Nucleus |
| Ap1s2 | adaptor re | Cytoplasm |
| Leprot | leptin rece | Plasma Me |
| Kank1 | KN motif a | Nucleus |
| Gulp1 | GULP, eng | Cytoplasm |
| Krt8 | keratin 8 | Cytoplasm |
| Irx3 | iroquois h | Nucleus |
| Dffa | DNA fragn | Nucleus |
| Ugdh | UDP-gluc | Nucleus |
| Nid2 | nidogen 2 | Extracellula |
| Mfap4 | microfibril | Extracellula |
| Pctp | phosphati | Cytoplasm |
| Ddah1 | dimethylar | Cytoplasm |
| B3gat3 | beta-1,3-g | Cytoplasm |
| Hs6st2 | heparan su | Plasma Me |
| Bcar3 | BCAR3, NS | Cytoplasm |
| Rcn1 | reticulocal | Cytoplasm |
| Smoc1 | SPARC rela | Extracellula |
| Ppfibp1 | PPFIA binc | Plasma Me |
| Sh3bgr | SH3-bindi | Cytoplasm |
| Six1 | SIX homec | Nucleus |
| Arl4a | ADP ribosy | Nucleus |
| Krt18 | keratin 18 | Cytoplasm |
| Prdx5 | peroxiredo | Cytoplasm |
| Ube2k | ubiquitin c | Cytoplasm |
| Slc39a10 | solute carr | Extracellula |
| Rcn2 | reticulocal | Cytoplasm |
| Reep5 | receptor a | Extracellula |
| Fkbp9 | FK506 binc | Cytoplasm |
| Slc12a4 | solute carr | Plasma Me |
| Dynll1 | dynein ligh | Cytoplasm |

| | | |
|---|---|---|
| Efcab11 | EF-hand c | Other |
| Pltp | phospholi| | Extracellul: |
| Iqcc | IQ motif c | Other |
| Mgst1 | microsoma | Cytoplasm |
| Serf2 | small EDRI | Other |
| Arsi | arylsulfata: | Extracellul: |
| Psme2 | proteasom | Cytoplasm |
| Spsb1 | splA/ryanc | Cytoplasm |
| Hsbp1 | heat shock | Nucleus |
| Klk7 | kallikrein r | Extracellul: |
| Lysmd2 | LysM dom | Other |
| Rasgef1b | RasGEF dc | Other |
| Rbms1 | RNA bindi | Nucleus |
| Bmp5 | bone morp | Extracellul: |
| Tceal8 | transcripti | Other |
| Nog | noggin | Extracellul: |
| Fam89a | family with | Other |
| BC023829 | - | - |
| 2700029M09Rik | - | - |
| Fkbp10 | FK506 binc | Cytoplasm |
| Marcksl1 | MARCKS li | Cytoplasm |
| Anxa5 | annexin A! | Plasma Me |
| Hapln1 | hyaluronai | Extracellul: |
| Sept9 | septin 9 | Cytoplasm |
| Litaf | lipopolysa | Nucleus |
| Nrep | neuronal r | Cytoplasm |
| Dusp4 | dual speci | Nucleus |
| Dpm2 | dolichyl-p | Cytoplasm |
| Nudt16l1 | nudix hydı | Cytoplasm |
| Snrk | SNF relate | Cytoplasm |
| Cirbp | cold induc | Nucleus |
| Fam213b | family with | Cytoplasm |
| Rhoc | ras homol | Plasma Me |
| Tmem106a | transmeml | Cytoplasm |
| Tmem246 | transmeml | Other |
| Zfp758 | zinc finger | Other |
| Fabp7 | fatty acid I | Cytoplasm |
| Actb | actin beta | Cytoplasm |
| Brk1 | BRICK1, SC | Cytoplasm |
| Armc1 | armadillo | Cytoplasm |
| Hoxa1 | homeobo› | Nucleus |
| Spats2l | spermatog | Nucleus |
| Ncstn | nicastrin | Plasma Me |
| Creb3l2 | cAMP resp | Nucleus |
| Gabarap | GABA type | Cytoplasm |
| Sms | spermine s | Cytoplasm |
| Snap47 | synaptoso | Plasma Me |
| Tmsb10 | thymosin, | Cytoplasm |
| Ube2n | ubiquitin c | Cytoplasm |
| Maged1 | MAGE fam | Plasma Me |
| Itpa | inosine tri| | Cytoplasm |
| Ubap1 | ubiquitin a | Cytoplasm |
| Tex2 | testis expr | Other |
| Tagln | transgelin | Cytoplasm |
| S100a11 | S100 calciı | Cytoplasm |
| Manbal | mannosida | Other |
| Mical1 | microtubu | Cytoplasm |
| 9530068E07Rik | chromoso | Other |

| | | |
|---|---|---|
| Apitd1 | - | - |
| Dnajc10 | DnaJ heat | Cytoplasm |
| Rpl7l1 | ribosomal | Cytoplasm |
| Mpzl1 | myelin prc | Plasma Me |
| Copa | coatomer | Cytoplasm |
| Dguok | deoxyguar | Cytoplasm |
| Sp5 | Sp5 transc | Nucleus |
| Fermt2 | fermitin fa | Cytoplasm |
| Twsg1 | twisted ga | Extracellul |
| Tbc1d7 | TBC1 dom | Cytoplasm |
| Gpatch1 | G-patch d | Extracellul |
| Tgfb2 | transformi | Extracellul |
| Ubb | ubiquitin E | Cytoplasm |
| Foxd4 | forkhead k | Nucleus |
| Sf3a3 | splicing fa | Nucleus |
| Pxk | PX domair | Cytoplasm |
| Rcn3 | reticulocal | Cytoplasm |
| Nkx2-9 | NK2 home | Nucleus |
| Phox2b | paired like | Nucleus |
| Rfx4 | regulatory | Nucleus |
| Olig2 | oligodend | Nucleus |
| Nkx6-1 | NK6 home | Nucleus |
| Map1b | microtubu | Cytoplasm |
| Fez1 | fasciculatic | Cytoplasm |
| Timp3 | TIMP meta | Extracellul |
| Dtx4 | deltex E3 u | Cytoplasm |
| Apba2 | amyloid be | Cytoplasm |
| Crabp1 | cellular ret | Cytoplasm |
| Vangl2 | VANGL pla | Plasma Me |
| Emilin2 | elastin mic | Extracellul |
| Nrcam | neuronal c | Plasma Me |
| Crmp1 | collapsin r | Cytoplasm |
| Msi1 | musashi R | Cytoplasm |
| Ptch1 | patched 1 | Plasma Me |
| Fam181a | family with | Other |
| Mafb | MAF bZIP | Nucleus |
| Nkx6-2 | NK6 home | Nucleus |
| Crabp2 | cellular ret | Cytoplasm |
| Tmem47 | transmeml | Plasma Me |
| Ednrb | endothelir | Plasma Me |
| Mapk11 | mitogen-a | Cytoplasm |
| Nr6a1 | nuclear rec | Nucleus |
| Mn1 | MN1 proto | Nucleus |
| Rnf220 | ring finger | Cytoplasm |
| Nell2 | neural EGI | Extracellul |
| Peg12 | paternally | Other |
| Olfml3 | olfactome | Extracellul |
| Ncam1 | neural cell | Plasma Me |
| Igdcc3 | immunogl | Other |
| Sfrp1 | secreted fr | Plasma Me |
| Ptn | pleiotroph | Extracellul |
| Fgfr2 | fibroblast | Plasma Me |
| Sparcl1 | SPARC like | Extracellul |
| Zfp811 | zinc finger | Other |
| Aldh1a2 | aldehyde c | Cytoplasm |
| Sept3 | septin 3 | Cytoplasm |
| Meis1 | Meis home | Nucleus |
| Cdh2 | cadherin 2 | Plasma Me |

| | | |
|---|---|---|
| Crb2 | crumbs 2, | Extracellula |
| Camkv | CaM kinas | Other |
| Sfrp2 | secreted fr | Plasma Me |
| Prtg | protogenir | Extracellula |
| Galnt16 | polypeptic | Cytoplasm |
| Hoxa2 | homeobox | Nucleus |
| Spsb4 | splA/ryanc | Cytoplasm |
| Fgfbp3 | fibroblast ( | Extracellula |
| Net1 | neuroepith | Nucleus |
| Meox1 | mesenchyr | Nucleus |
| Spon1 | spondin 1 | Extracellula |
| Rai14 | retinoic ac | Nucleus |
| Efnb3 | ephrin B3 | Plasma Me |
| Zmynd8 | zinc finger | Nucleus |
| Meis2 | Meis home | Nucleus |
| Stmn3 | stathmin 3 | Nucleus |
| Phf6 | PHD finge | Nucleus |
| Gli1 | GLI family | Nucleus |
| Idi1 | isopenteny | Cytoplasm |
| Acly | ATP citrate | Cytoplasm |
| Anp32e | acidic (leu | Cytoplasm |
| Zfp41 | ZFP41 zinc | Nucleus |
| Igf1r | insulin like | Plasma Me |
| Mycl | MYCL prot | Nucleus |
| Dek | DEK proto | Nucleus |
| Egr2 | early grow | Nucleus |
| Lin28b | lin-28 hon | Cytoplasm |
| Pknox2 | PBX/knotte | Nucleus |
| Egr1 | early grow | Nucleus |
| Pygb | glycogen p | Cytoplasm |
| Pcsk9 | proprotein | Extracellula |
| Qk | QKI, KH dc | Nucleus |
| Llgl1 | LLGL1, scri | Cytoplasm |
| Tagln3 | transgelin | Extracellula |
| Nsg1 | neuronal v | Cytoplasm |
| Vezf1 | vascular er | Nucleus |
| Dpysl2 | dihydropy | Cytoplasm |
| Dusp1 | dual speci | Nucleus |
| 1700011H14Rik | coiled-coi | Other |
| Msn | moesin | Plasma Me |
| Hpcal1 | hippocalci | Cytoplasm |
| Grb10 | growth fac | Cytoplasm |
| Slc39a6 | solute carr | Plasma Me |
| Ppp1r10 | protein ph | Nucleus |
| Cenpf | centromer | Nucleus |
| Mki67 | marker of | Nucleus |
| Hoxb1 | homeobox | Nucleus |
| Cenpe | centromer | Nucleus |
| Klc1 | kinesin ligl | Cytoplasm |
| Dnmt1 | DNA meth | Nucleus |
| Rad50 | RAD50 do | Nucleus |
| Gpr19 | G protein- | Plasma Me |
| Lpar6 | lysophosp | Plasma Me |
| Arvcf | ARVCF, de | Plasma Me |
| Nasp | nuclear au | Nucleus |
| Dpysl5 | dihydropy | Cytoplasm |
| Kcne1l | potassium | Plasma Me |
| Rarb | retinoic ac | Nucleus |

| | | |
|---|---|---|
| Col18a1 | collagen ty | Extracellula |
| Kif15 | kinesin fam | Nucleus |
| Gtse1 | G2 and S- | Cytoplasm |
| Armcx1 | armadillo r | Other |
| Hmgcs1 | 3-hydroxy | Cytoplasm |
| Sh3bgrl | SH3 doma | Cytoplasm |
| Arhgap19 | Rho GTPas | Cytoplasm |
| Vangl1 | VANGL pla | Cytoplasm |
| Gm10184 | karyopheri | Other |
| Sept11 | septin 11 | Nucleus |
| Rbbp5 | RB binding | Nucleus |
| Mcm7 | minichrom | Nucleus |
| Ccnd2 | cyclin D2 | Nucleus |
| Zfml | - | - |
| Rcc2 | regulator c | Nucleus |
| Pald1 | phosphata | Cytoplasm |
| Rtn3 | reticulon 3 | Cytoplasm |
| Gm4076 | predicted | Other |
| Lig1 | DNA ligase | Nucleus |
| Hmgb3 | high mobi | Nucleus |
| Cep55 | centrosom | Cytoplasm |
| Myh10 | myosin he | Cytoplasm |
| Gsk3b | glycogen s | Nucleus |
| Thrap3 | thyroid ho | Nucleus |
| Cdca7 | cell divisio | Nucleus |
| Hnrnph1 | heterogen | Nucleus |
| Nusap1 | nucleolar a | Nucleus |
| Plekha5 | pleckstrin | Cytoplasm |
| Gpx3 | glutathion | Extracellula |
| Ccdc160 | coiled-coi | Other |
| Mdk | midkine | Extracellula |
| Top2a | DNA topo | Nucleus |
| Dll1 | delta like c | Plasma Me |
| Ipo5 | importin 5 | Nucleus |
| Rad51ap1 | RAD51 ass | Nucleus |
| Ilf3 | interleukin | Nucleus |
| Fam64a | - | - |
| Ncaph | non-SMC | Nucleus |
| Sfpq | splicing fa | Nucleus |
| Meis3 | Meis home | Nucleus |
| Prpf3 | pre-mRNA | Nucleus |
| Cdh6 | cadherin 6 | Plasma Me |
| Fndc3c1 | | Other |
| Fubp1 | far upstrea | Nucleus |
| Brd2 | bromodor | Nucleus |
| Apex1 | apurinic/a | Nucleus |
| Klf13 | Kruppel lik | Nucleus |
| Dkc1 | dyskerin p | Nucleus |
| Arl5c | ADP ribosy | Other |
| Snrnp200 | small nucle | Nucleus |
| Gsg2 | - | - |
| Rfc1 | replication | Nucleus |
| Ndn | necdin, MA | Nucleus |
| Efna5 | ephrin A5 | Plasma Me |
| Kpna2 | karyopheri | Nucleus |
| Prc1 | protein reg | Nucleus |
| Elovl2 | ELOVL fatt | Cytoplasm |
| Jam3 | junctional | Plasma Me |

| | | |
|---|---|---|
| Tsc22d1 | TSC22 dor | Nucleus |
| Trim35 | tripartite n | Cytoplasm |
| Leo1 | LEO1 hom | Nucleus |
| Rsrc2 | arginine/s | Other |
| Agk | acylglycer | Cytoplasm |
| Eif5 | eukaryotic | Cytoplasm |
| Aurkb | aurora kin | Nucleus |
| Kpnb1 | karyopheri | Nucleus |
| Serbp1 | SERPINE1 | Cytoplasm |
| Ccnf | cyclin F | Nucleus |
| Skap2 | src kinase | Cytoplasm |
| Polr2b | RNA polyr | Nucleus |
| Tubb2b | tubulin be | Cytoplasm |
| Ddx5 | DEAD-box | Nucleus |
| Sf3b3 | splicing fa | Nucleus |
| Rbm39 | RNA bindi | Nucleus |
| Fos | Fos proto- | Nucleus |
| EGFP | - | - |
| Fxyd6 | FXYD dom | Plasma Me |
| Zfp422 | zinc finger | Nucleus |
| Nucks1 | nuclear ca | Nucleus |
| Nnat | neuronatir | Plasma Me |
| Ift74 | intraflagell | Cytoplasm |
| Hnrnpa2b1 | heterogen | Nucleus |
| Hmgn2 | high mobi | Nucleus |
| Pkm | pyruvate k | Cytoplasm |
| Ptp4a2 | protein tyr | Cytoplasm |
| Nap1l1 | nucleosom | Nucleus |
| Sf3b2 | splicing fa | Nucleus |
| Scoc | short coile | Cytoplasm |
| Marcksl1 | MARCKS li | Cytoplasm |
| Smc3 | structural | Nucleus |
| Incenp | inner centr | Nucleus |
| Fgf15 | fibroblast | Extracellul |
| Sox11 | SRY-box 1 | Nucleus |
| Hnrnpm | heterogen | Nucleus |
| Mrfap1 | Morf4 fam | Nucleus |
| Wdr6 | WD repeat | Cytoplasm |
| Trp53i11 | tumor pro | Other |
| Btg2 | BTG anti-p | Nucleus |
| Jun | Jun proto- | Nucleus |
| Map2k6 | mitogen-a | Cytoplasm |
| Zfp930 | expressed | Other |
| Rad9a | RAD9 chec | Nucleus |
| Lmnb1 | lamin B1 | Nucleus |
| Ckb | creatine ki | Cytoplasm |
| Cep57 | centrosom | Cytoplasm |
| Dars | aspartyl-tF | Cytoplasm |
| 2610318N02Rik | RIKEN cDN | Other |
| Eif4b | eukaryotic | Cytoplasm |
| Akap8 | A-kinase a | Nucleus |
| Azin1 | antizyme i | Cytoplasm |
| Gsta4 | glutathion | Other |
| Spop | speckle typ | Extracellul |
| Trim59 | tripartite n | Cytoplasm |
| Blcap | bladder ca | Other |
| Atp6v0e2 | ATPase H+ | Cytoplasm |
| Hsd11b2 | hydroxyste | Cytoplasm |

| | | |
|---|---|---|
| Dnajc9 | DnaJ heat | Nucleus |
| Kif20b | kinesin fan | Nucleus |
| Abcf1 | ATP bindir | Cytoplasm |
| Kdm3a | lysine dem | Nucleus |
| Ptma | prothymos | Nucleus |
| Ndc1 | NDC1 tran | Nucleus |
| Rbm14 | RNA bindi | Nucleus |
| Ankrd10 | ankyrin re | Nucleus |
| Mns1 | meiosis sp | Cytoplasm |
| H1f0 | H1 histone | Nucleus |
| Ndc80 | NDC80, ki | Nucleus |
| Igfbp5 | insulin like | Extracellula |
| Rab31 | RAB31, me | Cytoplasm |
| Igf2bp1 | insulin like | Cytoplasm |
| Bub1b | BUB1 mito | Nucleus |
| Sf3b1 | splicing fa | Nucleus |
| Dnajb9 | DnaJ heat | Nucleus |
| Nudt9 | nudix hydr | Cytoplasm |
| Set | SET nuclea | Nucleus |
| Tuba1a | tubulin alp | Cytoplasm |
| Fstl1 | follistatin l | Extracellula |
| Srsf10 | serine and | Nucleus |
| Ppat | phosphori | Cytoplasm |
| Ap1m1 | adaptor re | Cytoplasm |
| Bub3 | BUB3, mito | Nucleus |
| Kif22 | kinesin fan | Nucleus |
| Fbxo5 | F-box pro | Nucleus |
| Bora | bora, auro | Cytoplasm |
| Ska2 | spindle an | Nucleus |
| Ckap2l | cytoskelet | Cytoplasm |
| Kif11 | kinesin fan | Nucleus |
| Cdc42se2 | CDC42 sm | Plasma Me |
| Mybbp1a | MYB bindi | Nucleus |
| Smc4 | structural | Nucleus |
| Gm10282 | high mobi | Other |
| Foxa2 | forkhead b | Nucleus |
| Sde2 | SDE2 telor | Other |
| Nap1l4 | nucleosom | Cytoplasm |
| 2810417H13Rik | - | - |
| Ccar1 | cell divisio | Nucleus |
| Slc38a2 | solute carr | Plasma Me |
| Hnrnph2 | heterogen | Nucleus |
| Mcm2 | minichrom | Nucleus |
| Pnrc2 | proline ricl | Nucleus |
| Chtop | chromatin | Nucleus |
| Hmgn5 | high-mob | Nucleus |
| Plk1 | polo like k | Nucleus |
| Kif2c | kinesin fan | Nucleus |
| Pafah1b2 | platelet ac | Cytoplasm |
| Pebp1 | phosphati | Cytoplasm |
| Parp1 | poly(ADP- | Nucleus |
| Hnrnpc | heterogen | Nucleus |
| 1110004F10Rik | chromoso | Other |
| Fgfr1op | FGFR1 onc | Cytoplasm |
| Elovl5 | ELOVL fatt | Cytoplasm |
| Hmmr | hyaluronai | Plasma Me |
| Rfc4 | replication | Nucleus |
| Ddx21 | DExD-box | Nucleus |

| | | |
|---|---|---|
| Eno1b | enolase 1 | Cytoplasm |
| Fads2 | fatty acid ( | Plasma Me |
| Fam60a | SIN3-HDA | Extracellul |
| Golm1 | golgi mem | Cytoplasm |
| Slc2a1 | solute carr | Plasma Me |
| Kif23 | kinesin fan | Cytoplasm |
| Nedd4 | neural pre | Cytoplasm |
| Mcm3 | minichrom | Nucleus |
| Ssb | Sjogren sy | Nucleus |
| Ccnd1 | cyclin D1 | Nucleus |
| Fads1 | fatty acid ( | Plasma Me |
| Glyr1 | glyoxylate | Nucleus |
| Cbx3 | chromobo | Nucleus |
| Bzw2 | basic leuci | Cytoplasm |
| Mcm6 | minichrom | Nucleus |
| Ckap2 | cytoskelet | Cytoplasm |
| Scd2 | stearoyl-C | Cytoplasm |
| Ncbp1 | nuclear ca | Nucleus |
| Hspd1 | heat shock | Cytoplasm |
| Hnrnpk | heterogen | Nucleus |
| H2afy2 | H2A histor | Nucleus |
| Hsp90aa1 | heat shock | Cytoplasm |
| Sap30 | Sin3A assc | Nucleus |
| Mcm5 | minichrom | Nucleus |
| Ppme1 | protein ph | Nucleus |
| Polr2m | RNA polyr | Nucleus |
| Hnrnph3 | heterogen | Nucleus |
| Abce1 | ATP bindir | Cytoplasm |
| Xrcc1 | X-ray repa | Nucleus |
| Tpx2 | TPX2, micr | Nucleus |
| Lin7c | lin-7 hom | Cytoplasm |
| Gm3839 | glyceralde | Other |
| Hmgb1 | high mobi | Nucleus |
| Hnrnpa1 | heterogen | Nucleus |
| Pcbp4 | poly(rC) bi | Nucleus |
| Ddx1 | DEAD-box | Nucleus |
| Adprh | ADP-ribos | Other |
| Rrm1 | ribonucleo | Nucleus |
| Prps1 | phosphori | Cytoplasm |
| Pold3 | DNA polyr | Nucleus |
| Parp2 | poly(ADP- | Nucleus |
| Ptcd3 | pentatrico | Cytoplasm |
| Gm7964 | heterogen | Other |
| Cer1 | cerberus 1 | Extracellul |
| Tdgf1 | teratocarci | Extracellul |
| 2410004P03Rik | chromosoı | Other |
| Fgf8 | fibroblast ( | Extracellul |
| Pou5f1 | POU class | Nucleus |
| Lhx1 | LIM home | Nucleus |
| Tppp3 | tubulin po | Cytoplasm |
| Fgf5 | fibroblast ( | Extracellul |
| Ndrg1 | N-myc do | Nucleus |
| Eomes | eomesode | Nucleus |
| Rpl24 | ribosomal | Cytoplasm |
| Rps5 | ribosomal | Cytoplasm |
| Foxj1 | forkhead b | Nucleus |
| Gm10073 | ribosomal | Other |
| Gnb2l1 | - | - |

| | | |
|---|---|---|
| Gm17669 | predicted | Other |
| Rps21 | ribosomal | Cytoplasm |
| Gm17511 | ribosomal | Other |
| Pcsk1n | proprotein | Extracellula |
| Gm10076 | ribosomal | Other |
| Gm10126 | ribosomal | Other |
| Rps17 | ribosomal | Cytoplasm |
| Gm10263 | 40S ribosc | Other |
| Rps26-ps1 | ribosomal | Other |
| Nodal | nodal grov | Extracellula |
| Gpc4 | glypican 4 | Plasma Me |
| Rps12-ps3 | ribosomal | Other |
| Gm17541 | ribosomal | Other |
| Rps15a | ribosomal | Cytoplasm |
| Gm10709 | ribosomal | Other |
| Polr2h | RNA polyr | Nucleus |
| Gm5160 | peptidylpr | Other |
| Vrtn | vertebrae | Other |
| Cfc1 | cripto, FRL | Extracellula |
| Gm5786 | ribosomal | Other |
| Rps20 | ribosomal | Cytoplasm |
| Rpl12 | ribosomal | Nucleus |
| Rplp1 | ribosomal | Nucleus |
| Rps15 | ribosomal | Cytoplasm |
| Chrd | chordin | Extracellula |
| Rpl27 | ribosomal | Cytoplasm |
| Gm4950 | proteasom | Other |
| Rps29 | ribosomal | Cytoplasm |
| Rpl41 | ribosomal | Cytoplasm |
| Eras | ES cell exp | Plasma Me |
| Gm9843 | predicted | Other |
| Gm6576 | ribosomal | Other |
| Upp1 | uridine ph | Cytoplasm |
| Rps23 | ribosomal | Cytoplasm |
| Rpl34 | ribosomal | Cytoplasm |
| Rpl37a | ribosomal | Cytoplasm |
| Mixl1 | Mix paired | Nucleus |
| Gm18025 | ribosomal | Other |
| Gm10036 | ribosomal | Other |
| Rpl8 | ribosomal | Cytoplasm |
| Eci1 | enoyl-CoA | Cytoplasm |
| Rps3 | ribosomal | Cytoplasm |
| Rps10 | ribosomal | Cytoplasm |
| Timm13 | translocase | Cytoplasm |
| Rps2 | ribosomal | Cytoplasm |
| Rpl36-ps3 | ribosomal | Other |
| Rpl9-ps6 | ribosomal | Other |
| Rpl28 | ribosomal | Cytoplasm |
| Rpl37 | ribosomal | Cytoplasm |
| Rps27rt | ribosomal | Other |
| Rpl27-ps3 | ribosomal | Other |
| Gm8225 | ribosomal | Other |
| Rplp2 | ribosomal | Cytoplasm |
| Rpl27a | ribosomal | Cytoplasm |
| Rps12 | ribosomal | Cytoplasm |
| Eif3k | eukaryotic | Cytoplasm |
| Psmb3 | proteasom | Cytoplasm |
| Rps2-ps6 | ribosomal | Other |

| | | |
|---|---|---|
| Rps27 | ribosomal | Other |
| Rpl26 | ribosomal | Cytoplasm |
| Gm8730 | ribosomal | Other |
| Rps28 | ribosomal | Cytoplasm |
| Gm12728 | peptidyl-p | Other |
| Rpl10a | ribosomal | Nucleus |
| Rpl10-ps3 | ribosomal | Other |
| Rps3a1 | ribosomal | Cytoplasm |
| Sp5 | Sp5 transc | Nucleus |
| Rps13 | ribosomal | Cytoplasm |
| Dnmt3b | DNA meth | Nucleus |
| Rplp0 | ribosomal | Cytoplasm |
| Foxh1 | forkhead b | Nucleus |
| Rps14 | ribosomal | Cytoplasm |
| Gata1 | GATA binc | Nucleus |
| Pmaip1 | phorbol-1 | Cytoplasm |
| Rps18 | ribosomal | Cytoplasm |
| Rps26 | ribosomal | Cytoplasm |
| Rpl18 | ribosomal | Cytoplasm |
| Rpl18a | ribosomal | Cytoplasm |
| Trh | thyrotropi | Extracellul |
| Pim2 | Pim-2 pro | Nucleus |
| Rpl7a | ribosomal | Cytoplasm |
| Rpl29 | ribosomal | Cytoplasm |
| Tspo | translocat | Cytoplasm |
| Zbed5 | zinc finger | Nucleus |
| Rpl31 | ribosomal | Cytoplasm |
| Rps11 | ribosomal | Cytoplasm |
| Prdx1 | peroxired | Cytoplasm |
| Rpl32 | ribosomal | Cytoplasm |
| Pfn1 | profilin 1 | Cytoplasm |
| Gm11273 | cytochrom | Other |
| Fau | FAU, ubiqu | Cytoplasm |
| Eif3i | eukaryotic | Cytoplasm |
| Rpl30 | ribosomal | Cytoplasm |
| Mrpl34 | mitochonc | Cytoplasm |
| Cox8a | cytochrom | Cytoplasm |
| Gm10146 | LSM7 hom | Other |
| Tceb2 | - | - |
| Rpl23a-ps3 | ribosomal | Other |
| Isyna1 | inositol-3- | Cytoplasm |
| Rps9 | ribosomal | Cytoplasm |
| Ppp1r1a | protein ph | Cytoplasm |
| Gm9847 | suppresso | Other |
| Gm9493 | predicted | Cytoplasm |
| Rpl11 | ribosomal | Cytoplasm |
| Rpl36 | ribosomal | Nucleus |
| H2afz | H2A histor | Nucleus |
| Rpl10 | ribosomal | Cytoplasm |
| Siva1 | SIVA1 apo | Cytoplasm |
| Rps27a | ribosomal | Cytoplasm |
| Rpl39 | ribosomal | Cytoplasm |
| Tomm7 | translocas | Cytoplasm |
| Rps16 | ribosomal | Cytoplasm |
| Hint1 | histidine tr | Nucleus |
| Atp5j2 | ATP synth | Cytoplasm |
| Gm10020 | ribosomal | Other |
| Rpl23 | ribosomal | Cytoplasm |

| | | |
|---|---|---|
| Rps18-ps3 | ribosomal | Other |
| Rps19 | ribosomal | Cytoplasm |
| Ppia | peptidylpr | Cytoplasm |
| Rpl19 | ribosomal | Cytoplasm |
| Uqcr11 | ubiquinol- | Cytoplasm |
| Snrpg | small nucl | Nucleus |
| Otx2 | orthodenti | Nucleus |
| Rpl7 | ribosomal | Nucleus |
| Rpl9 | ribosomal | Nucleus |
| Rpsa | ribosomal | Cytoplasm |
| Pycr2 | pyrroline-! | Cytoplasm |
| Atp5l | ATP synth | Cytoplasm |
| Sssca1 | Sjogren sy | Other |
| Gm5428 | - | - |
| Myl7 | myosin lig | Cytoplasm |
| Gm5239 | ubiquitin A | Other |
| Cox7a2 | cytochrom | Cytoplasm |
| Mif | macropha | Extracellul |
| Rps25 | ribosomal | Cytoplasm |
| Ndufb4 | NADH:ubi | Cytoplasm |
| Trmt112 | tRNA meth | Cytoplasm |
| Eef1a1 | eukaryotic | Cytoplasm |
| Usmg5 | ATP synth | Cytoplasm |
| Ldha | lactate del | Cytoplasm |
| Ndufa2 | NADH:ubi | Cytoplasm |
| Gm5093 | predicted | Other |
| Gm11808 | ubiquitin A | Other |
| Psmb5 | proteasom | Cytoplasm |
| Rps4x | ribosomal | Cytoplasm |
| Nop10 | NOP10 rib | Nucleus |
| Rps6 | ribosomal | Cytoplasm |
| Lefty2 | left-right c | Extracellul |
| Gm8186 | small nucl | Other |
| Rpl13a | ribosomal | Cytoplasm |
| Rpl15 | ribosomal | Cytoplasm |
| N6amt2 | - | - |
| Grrp1 | family with | Other |
| Ran | RAN, mem | Nucleus |
| Atp5a1 | ATP synth | Cytoplasm |
| Psmb6 | proteasom | Nucleus |
| Cyc1 | cytochrom | Cytoplasm |
| Lars2 | leucyl-tRN | Cytoplasm |
| Rpl21 | ribosomal | Cytoplasm |
| Rpl13 | ribosomal | Nucleus |
| Eef2 | eukaryotic | Cytoplasm |
| Snrpd2 | small nucl | Nucleus |
| Rps8 | ribosomal | Cytoplasm |
| Ndufa7 | NADH:ubi | Cytoplasm |
| Ifitm2 | interferon | Cytoplasm |
| Rps4l | ribosomal | Cytoplasm |
| 2700094K13Rik | - | - |
| Gpx1 | glutathion | Cytoplasm |
| Rpl17 | ribosomal | Cytoplasm |
| Asns | asparagine | Cytoplasm |
| Gm15013 | ribosomal | Other |
| Ndufa11 | NADH:ubi | Cytoplasm |
| Psma6 | proteasom | Cytoplasm |
| Slc25a3 | solute carr | Cytoplasm |

| | | |
|---|---|---|
| Atp6v1f | ATPase H+ | Cytoplasm |
| Car2 | carbonic a | Cytoplasm |
| Mrps6 | mitochonc | Cytoplasm |
| Gm9803 | presequen | Other |
| Polr2e | RNA polyr | Nucleus |
| Atp5d | ATP synth | Cytoplasm |
| Rpl13-ps3 | ribosomal | Other |
| Phc1 | polyhome | Nucleus |
| Dll3 | delta like c | Extracellul |
| Lsm4 | LSM4 hom | Nucleus |
| Atp5c1 | ATP synth | Cytoplasm |
| Zic3 | Zic family | Nucleus |
| Rpl38 | ribosomal | Cytoplasm |
| Eif1 | eukaryotic | Other |
| Polr2c | RNA polyr | Nucleus |
| 2010107E04Rik | ATP synth | Cytoplasm |
| Ldhb | lactate del | Cytoplasm |
| Uqcrb | ubiquinol- | Cytoplasm |
| Eif3g | eukaryotic | Cytoplasm |
| Rpl36al | ribosomal | Cytoplasm |
| Gsc | goosecoid | Nucleus |
| Rpl35a | ribosomal | Cytoplasm |
| Snrpe | small nucl | Nucleus |
| Atp5b | ATP synth | Cytoplasm |
| Hesx1 | HESX hom | Nucleus |
| Cxx1c | - | - |
| Ndufc2 | NADH:ubi | Cytoplasm |
| Chchd10 | coiled-coi | Cytoplasm |
| Hmga1 | high mobi | Nucleus |
| Hprt | hypoxanth | Cytoplasm |
| 2410015M20Rik | chromoso | Cytoplasm |
| Pfdn6 | prefoldin s | Cytoplasm |
| Ina | internexin | Cytoplasm |
| Hcfc1r1 | host cell fa | Nucleus |
| Hmga1-rs1 | - | - |
| Eif3m | eukaryotic | Cytoplasm |
| Samd3 | sterile alpl | Other |
| Ppp1ca | protein ph | Cytoplasm |
| Btf3 | basic trans | Nucleus |
| Cox5b | cytochrom | Cytoplasm |
| Ubb | ubiquitin E | Cytoplasm |
| Snrpd1 | small nucl | Nucleus |
| H3f3a | H3 histone | Nucleus |
| Cenpw | centromer | Nucleus |
| Tpt1 | tumor pro | Cytoplasm |
| Emb | embigin | Plasma Me |
| Gm5616 | NHP2 non | Nucleus |
| Eif3f | eukaryotic | Cytoplasm |
| Rpl22l1 | ribosomal | Other |
| Psmb7 | proteasom | Cytoplasm |
| Mrpl28 | mitochonc | Cytoplasm |
| Rpl5 | ribosomal | Cytoplasm |
| Prdx2 | peroxiredc | Cytoplasm |
| Thop1 | thimet olig | Cytoplasm |
| Rpl22 | ribosomal | Nucleus |
| Mdh2 | malate del | Cytoplasm |
| Rpl3 | ribosomal | Nucleus |
| Psmb4 | proteasom | Cytoplasm |

| | | |
|---|---|---|
| T | T-box trar | Nucleus |
| Psma5 | proteasom | Cytoplasm |
| Bmp7 | bone mor | Extracellul |
| Mlf1 | myeloid le | Nucleus |
| Bola2 | bolA-like 2 | Cytoplasm |
| Znhit2 | zinc finger | Other |
| Csnk2b | casein kina | Cytoplasm |
| Lsm7 | LSM7 hom | Nucleus |
| Ndufs7 | NADH:ubi | Cytoplasm |
| Eif5a | eukaryotic | Cytoplasm |
| Cops6 | COP9 sign | Nucleus |
| Mrps28 | mitochonc | Cytoplasm |
| Polr1c | RNA polyr | Nucleus |
| Cox5a | cytochrom | Cytoplasm |
| Pomp | proteasom | Nucleus |
| Pgp | phosphog | Cytoplasm |
| Mrpl20 | mitochonc | Cytoplasm |
| Polr2k | RNA polyr | Nucleus |
| Slc25a5 | solute carr | Cytoplasm |
| Pam16 | presequen | Cytoplasm |
| Cct7 | chaperonii | Cytoplasm |
| Uqcrq | ubiquinol- | Cytoplasm |
| Wdr55 | WD repeat | Nucleus |
| Sec61b | Sec61 tran | Cytoplasm |
| Cyp26a1 | cytochrom | Cytoplasm |
| Penk | proenkeph | Extracellul |
| Plp2 | proteolipic | Cytoplasm |
| Gng5 | G protein s | Plasma Me |
| Psmb2 | proteasom | Cytoplasm |
| Arl6 | ADP ribosy | Cytoplasm |
| Dctpp1 | dCTP pyro | Cytoplasm |
| Noto | notochord | Other |
| Psmb10 | proteasom | Cytoplasm |
| Cox7b | cytochrom | Cytoplasm |
| Pcbp1 | poly(rC) bi | Nucleus |
| Ahcy | adenosylh | Cytoplasm |
| Polr2g | RNA polyr | Nucleus |
| Aurkaip1 | aurora kin | Nucleus |
| Psmd8 | proteasom | Cytoplasm |
| Zscan10 | zinc finger | Nucleus |
| Cks1b | CDC28 pro | Other |
| Lefty1 | left-right c | Extracellul |
| Dohh | deoxyhypu | Cytoplasm |
| Med31 | mediator c | Nucleus |
| Gm561 | - | - |
| Supt4a | SPT4 hom | Nucleus |
| Psma7 | proteasom | Cytoplasm |
| Psmb1 | proteasom | Cytoplasm |
| Psmc5 | proteasom | Nucleus |
| Snrpn | small nucl | Nucleus |
| Ndufs8 | NADH:ubi | Cytoplasm |
| Uqcc2 | ubiquinol- | Cytoplasm |
| Atp5k | ATP synth | Cytoplasm |
| Trappc4 | trafficking | Cytoplasm |
| Cox4i1 | cytochrom | Cytoplasm |
| Atp5o | ATP synth | Cytoplasm |
| Chchd1 | coiled-coi | Nucleus |
| Stoml2 | stomatin li | Plasma Me |

| | | |
|---|---|---|
| Mrpl42 | mitochonc | Cytoplasm |
| Spint2 | serine pep | Extracellul: |
| Perp | PERP, TP5: | Plasma Me |
| Epcam | epithelial c | Plasma Me |
| Tpgs1 | tubulin po | Cytoplasm |
| Oat | ornithine a | Cytoplasm |
| Sec61g | Sec61 tran | Plasma Me |
| Rpp25l | ribonuclea | Other |
| Sdhd | succinate c | Cytoplasm |
| Nhp2 | NHP2 ribo | Nucleus |
| Ruvbl2 | RuvB like / | Nucleus |
| Alkbh7 | alkB homc | Cytoplasm |
| Ppp4c | protein ph | Cytoplasm |
| Polr2l | RNA polyr | Nucleus |
| Tmem256 | transmeml | Cytoplasm |
| Clpp | caseinolyti | Cytoplasm |
| Uqcr10 | ubiquinol- | Cytoplasm |
| Mrps12 | mitochonc | Cytoplasm |
| Gm4737 | predicted | Other |
| Imp3 | IMP3, U3 s | Cytoplasm |
| Timm23 | translocas | Cytoplasm |
| Capsl | calcyphosi | Other |
| Ndufa6 | NADH:ubi | Cytoplasm |
| Bambi | BMP and a | Plasma Me |
| Sox17 | SRY-box 1 | Nucleus |
| Cox7c | cytochrom | Cytoplasm |
| Mrpl55 | mitochonc | Cytoplasm |
| D8Ertd738e | chromosoı | Nucleus |
| Dkk1 | dickkopf V | Extracellul: |
| Atp5g1 | ATP synth: | Cytoplasm |
| C1qbp | compleme | Cytoplasm |
| Ndufa12 | NADH:ubi | Cytoplasm |
| Polr2i | RNA polyr | Nucleus |
| Phf5a | PHD finge | Nucleus |
| Tubb4b | tubulin be | Cytoplasm |
| Uqcrfs1 | ubiquinol- | Cytoplasm |
| Rps19bp1 | ribosomal | Nucleus |
| Fkbp11 | FK506 bin | Cytoplasm |
| Myeov2 | - | - |
| Mrps21 | mitochonc | Cytoplasm |
| Phb | prohibitin | Nucleus |
| Bax | BCL2 asso | Cytoplasm |
| Urod | uroporphy | Cytoplasm |
| Cstb | cystatin B | Cytoplasm |
| Rab1b | RAB1B, me | Cytoplasm |
| Cst3 | cystatin C | Extracellul: |
| Ap2s1 | adaptor re | Cytoplasm |
| Ost4 | oligosacch | Other |
| Gstp2 | glutathion | Cytoplasm |
| Uqcrh | ubiquinol- | Cytoplasm |
| Gm10273 | translocas | Other |
| Mrpl11 | mitochonc | Cytoplasm |
| Dpcd | deleted in | Nucleus |
| Naa38 | N(alpha)-a | Nucleus |
| Mid1ip1 | MID1 inter | Cytoplasm |
| Eef1d | eukaryotic | Cytoplasm |
| Noc4l | nucleolar c | Nucleus |
| Gart | phosphori | Cytoplasm |

| | | |
|---|---|---|
| Nme1 | NME/NM2 | Cytoplasm |
| Gstp1 | glutathion | Cytoplasm |
| Pop4 | POP4 hom | Nucleus |
| Emg1 | EMG1, N1- | Nucleus |
| Dusp4 | dual speci | Nucleus |
| Tmem107 | transmeml | Extracellul |
| Bphl | biphenyl h | Cytoplasm |
| Timm8b | translocas | Cytoplasm |
| Epha1 | EPH recep | Plasma Me |
| Plrg1 | pleiotropic | Nucleus |
| Dynll1 | dynein ligl | Cytoplasm |
| Snai1 | snail family | Nucleus |
| Coa3 | cytochrom | Cytoplasm |
| Sac3d1 | SAC3 dom | Cytoplasm |
| Rps27l | ribosomal | Cytoplasm |
| Mrpl41 | mitochond | Cytoplasm |
| Hhex | hematopo | Nucleus |
| Dnajc15 | DnaJ heat | Cytoplasm |
| Mrpl46 | mitochond | Cytoplasm |
| Cct2 | chaperonii | Cytoplasm |
| Nup37 | nucleopor | Nucleus |
| Mrpl54 | mitochond | Cytoplasm |
| Cubn | cubilin | Plasma Me |
| Nrk | Nik relatec | Other |
| Abcc2 | ATP bindir | Plasma Me |
| Mlxipl | MLX intera | Nucleus |
| Prodh2 | proline de | Cytoplasm |
| Slc23a1 | solute carr | Plasma Me |
| Tpcn1 | two pore s | Plasma Me |
| Npc1 | NPC intrac | Cytoplasm |
| Swt1 | SWT1, RN/ | Extracellul |
| Dmxl1 | Dmx like 1 | Extracellul |
| Soat2 | sterol O-a | Cytoplasm |
| Tcte2 | t-complex | Other |
| Ube2l6 | ubiquitin c | Cytoplasm |
| Sgk3 | serum/glu | Cytoplasm |
| Foxo4 | forkhead b | Nucleus |
| Rita1 | RBPJ intera | Cytoplasm |
| Rtl1 | retrotransp | Other |
| Ptpn23 | protein tyr | Cytoplasm |
| Slc13a4 | solute carr | Plasma Me |
| Jade1 | jade family | Nucleus |
| Slc7a8 | solute carr | Plasma Me |
| Mfsd12 | major facil | Cytoplasm |
| Rpia | ribose 5-p | Cytoplasm |
| Fnip2 | folliculin ir | Cytoplasm |
| Dpm1 | dolichyl-p | Cytoplasm |
| Trpm2 | transient r | Plasma Me |
| Acy1 | aminoacyla | Cytoplasm |
| Apoc2 | apolipopro | Extracellul |
| Abcd4 | ATP bindir | Cytoplasm |
| Iqsec1 | IQ motif ar | Cytoplasm |
| Sfmbt2 | Scm like w | Nucleus |
| Morc4 | MORC fam | Nucleus |
| Ctsh | cathepsin | Cytoplasm |
| Evi5 | ecotropic v | Cytoplasm |
| Synj1 | synaptojar | Cytoplasm |
| Dennd6b | DENN dor | Cytoplasm |

| | | |
|---|---|---|
| Npl | N-acetylne | Cytoplasm |
| Col27a1 | collagen ty | Extracellula |
| C77080 | KIAA1522 | Other |
| Sec24a | SEC24 hor | Cytoplasm |
| Myo6 | myosin VI | Cytoplasm |
| Mt1 | metallothi | Cytoplasm |
| Gcat | glycine C- | Cytoplasm |
| Trim6 | tripartite m | Cytoplasm |
| Neu1 | neuramini | Cytoplasm |
| Gga3 | golgi asso | Cytoplasm |
| Sfi1 | SFI1 centri | Cytoplasm |
| Myo5b | myosin VB | Cytoplasm |
| Ppcdc | phosphop | Cytoplasm |
| Slc43a2 | solute carr | Plasma Me |
| Apoa1 | apolipopro | Extracellula |
| Amn | amnion as | Plasma Me |
| Pcyt2 | phosphate | Cytoplasm |
| Car7 | carbonic a | Cytoplasm |
| Snx8 | sorting ne | Cytoplasm |
| Acox1 | acyl-CoA c | Cytoplasm |
| Slc2a2 | solute carr | Plasma Me |
| Stat3 | signal tran | Nucleus |
| Gm5424 | argininosu | Other |
| Clk2 | CDC like k | Nucleus |
| Gramd1b | GRAM dor | Other |
| Apoe | apolipopro | Extracellula |
| Sgpp2 | sphingosir | Cytoplasm |
| Setd4 | SET domai | Other |
| Foxk1 | forkhead b | Nucleus |
| Slc39a5 | solute carr | Plasma Me |
| Med12 | mediator c | Nucleus |
| Polq | DNA polyr | Nucleus |
| Prdm2 | PR/SET do | Nucleus |
| Trak2 | trafficking | Plasma Me |
| Adck5 | aarF doma | Cytoplasm |
| Atp6v0a1 | ATPase H+ | Cytoplasm |
| Foxp4 | forkhead b | Nucleus |
| Ass1 | argininosu | Cytoplasm |
| Dvl1 | dishevelle | Cytoplasm |
| Slc25a10 | solute carr | Cytoplasm |
| Rbm47 | RNA bindi | Nucleus |
| Neil1 | nei like DN | Nucleus |
| Furin | furin, paire | Cytoplasm |
| Lgmn | legumain | Cytoplasm |
| Tmem245 | transmeml | Other |
| Irf2bp2 | interferon | Nucleus |
| Rrbp1 | ribosome l | Cytoplasm |
| Ctsa | cathepsin . | Cytoplasm |
| Apob | apolipopro | Extracellula |
| B4galt5 | beta-1,4-g | Cytoplasm |
| Dhrs11 | dehydroge | Other |
| Pqlc2 | PQ loop re | Cytoplasm |
| Gipc2 | GIPC PDZ | Cytoplasm |
| Gldc | glycine de | Cytoplasm |
| Lpin2 | lipin 2 | Nucleus |
| 3110002H16Rik | regulator c | Cytoplasm |
| Dab2 | DAB2, clat | Plasma Me |
| Gga2 | golgi asso | Cytoplasm |

| | | |
|---|---|---|
| Nmrk1 | nicotinami | Cytoplasm |
| Zfp692 | zinc finger | Nucleus |
| Gm21887 | - | - |
| Atp7b | ATPase co | Cytoplasm |
| Ascc3 | activating | Cytoplasm |
| Triml2 | tripartite n | Other |
| Cul7 | cullin 7 | Cytoplasm |
| P2rx4 | purinergic | Plasma Me |
| Cited1 | Cbp/p300 | Nucleus |
| Dpp7 | dipeptidyl | Cytoplasm |
| Nup153 | nucleopor | Nucleus |
| Fxyd4 | FXYD dom | Plasma Me |
| Sccpdh | saccharop | Cytoplasm |
| Agap3 | ArfGAP wi | Nucleus |
| Aqp3 | aquaporin | Plasma Me |
| Cluh | clustered r | Cytoplasm |
| Hsd17b11 | hydroxyste | Cytoplasm |
| Slc38a9 | solute carr | Extracellula |
| Creb3l3 | cAMP resp | Cytoplasm |
| Ttc19 | tetratricop | Cytoplasm |
| Nktr | natural kill | Plasma Me |
| Nlrc3 | NLR family | Cytoplasm |
| Ctsb | cathepsin | Cytoplasm |
| Tmem150a | transmeml | Plasma Me |
| Amot | angiomoti | Plasma Me |
| Dcbld1 | discoidin, | Extracellula |
| Erdr1 | erythroid c | Other |
| Mospd2 | motile spe | Plasma Me |
| Reep6 | receptor a | Plasma Me |
| Kifc2 | kinesin fan | Cytoplasm |
| 2610528J11Rik | chromosoı | Other |
| Nprl2 | NPR2 like, | Cytoplasm |
| Cdhr2 | cadherin r | Plasma Me |
| Plekhg5 | pleckstrin | Cytoplasm |
| Pitpna | phosphati | Cytoplasm |
| Aass | aminoadip | Cytoplasm |
| G730046D07Rik | RIKEN cDN | Other |
| Ap1ar | adaptor re | Cytoplasm |
| Ndel1 | nudE neur | Nucleus |
| Polg | DNA polyr | Cytoplasm |
| Akap1 | A-kinase a | Cytoplasm |
| Gna12 | G protein s | Plasma Me |
| Crls1 | cardiolipin | Cytoplasm |
| Ano9 | anoctamin | Plasma Me |
| Optn | optineurin | Cytoplasm |
| Ppm1a | protein ph | Cytoplasm |
| Leng8 | leukocyte | Other |
| Pxn | paxillin | Cytoplasm |
| Sema4b | semaphori | Plasma Me |
| Uap1l1 | UDP-N-ac | Other |
| Sgk1 | serum/glu | Cytoplasm |
| Apoa4 | apolipopro | Extracellula |
| Lrp2 | LDL recept | Plasma Me |
| Gpam | glycerol-3 | Cytoplasm |
| Acin1 | apoptotic | Nucleus |
| Klhl2 | kelch like f | Cytoplasm |
| Rin3 | Ras and Ra | Cytoplasm |
| AI413582 | expressed | Other |

| | | |
|---|---|---|
| Qprt | quinolinat | Cytoplasm |
| Sirt7 | sirtuin 7 | Nucleus |
| Gba2 | glucosylce | Cytoplasm |
| Mafk | MAF bZIP | Nucleus |
| Slc35f6 | solute carr | Cytoplasm |
| Wdr24 | WD repeat | Cytoplasm |
| Nhlrc3 | NHL repea | Cytoplasm |
| Adora2b | adenosine | Plasma Me |
| 2310035C23Rik | KIAA1468 | Other |
| Kctd6 | potassium | Plasma Me |
| Tmem37 | transmeml | Plasma Me |
| Asl | argininosu | Cytoplasm |
| Rhot1 | ras homolo | Cytoplasm |
| Tmem79 | transmeml | Cytoplasm |
| Sfxn5 | sideroflexi | Cytoplasm |
| Mt2 | metallothi | Other |
| Slc25a19 | solute carr | Cytoplasm |
| Slc38a6 | solute carr | Plasma Me |
| Trim25 | tripartite n | Cytoplasm |
| Gigyf2 | GRB10 inte | Cytoplasm |
| Dcaf15 | DDB1 and | Other |
| Bach1 | BTB doma | Nucleus |
| Sh3tc1 | SH3 doma | Extracellul |
| Hmox1 | heme oxyg | Cytoplasm |
| Folr1 | folate rece | Plasma Me |
| Tfcp2l1 | transcriptio | Nucleus |
| Trrap | transforma | Nucleus |
| Zc3h7a | zinc finger | Extracellul |
| Mthfr | methylene | Cytoplasm |
| Atp6v1a | ATPase H+ | Plasma Me |
| Stxbp2 | syntaxin bi | Plasma Me |
| Josd2 | Josephin c | Cytoplasm |
| Lgals2 | galectin 2 | Cytoplasm |
| Ift46 | intraflagell | Cytoplasm |
| Nadsyn1 | NAD synth | Cytoplasm |
| Hook2 | hook micro | Cytoplasm |
| Eef2kmt | eukaryotic | Cytoplasm |
| Fuca2 | alpha-L-fu | Extracellul |
| Ctsz | cathepsin | Cytoplasm |
| Renbp | renin bind | Cytoplasm |
| Kmt2d | lysine metl | Nucleus |
| Tbc1d10a | TBC1 dom | Plasma Me |
| Txndc11 | thioredoxi | Cytoplasm |
| Cmtm4 | CKLF like N | Extracellul |
| Muc13 | mucin 13, | Extracellul |
| Slc25a22 | solute carr | Cytoplasm |
| Hnf4a | hepatocyt | Nucleus |
| Xpo6 | exportin 6 | Cytoplasm |
| Lpar1 | lysophosp | Plasma Me |
| Atp6v0d1 | ATPase H+ | Cytoplasm |
| Wash1 | - | - |
| Ctsl | cathepsin | Cytoplasm |
| Pmm2 | phosphom | Cytoplasm |
| Pick1 | protein int | Cytoplasm |
| Dtx3 | deltex E3 u | Cytoplasm |
| Cltc | clathrin he | Plasma Me |
| Hps5 | HPS5, biog | Cytoplasm |
| Atp6v1h | ATPase H+ | Cytoplasm |

| | | |
|---|---|---|
| Commd4 | COMM dc | Cytoplasm |
| Tuba4a | tubulin alp | Cytoplasm |
| Nfx1 | nuclear tra | Nucleus |
| Narfl | nuclear pr | Cytoplasm |
| Ninj1 | ninjurin 1 | Plasma Me |
| Cgnl1 | cingulin lik | Plasma Me |
| Tlcd1 | TLC doma | Other |
| Ubqln2 | ubiquilin 2 | Nucleus |
| Rpusd3 | RNA pseu | Other |
| Uvrag | UV radiatio | Nucleus |
| Rogdi | rogdi hom | Nucleus |
| Camsap3 | calmodulir | Cytoplasm |
| Immp2l | inner mito | Cytoplasm |
| Prrc2a | proline ricl | Cytoplasm |
| Hmgxb3 | HMG-box | Nucleus |
| Plekhg3 | pleckstrin | Other |
| Atp1b1 | ATPase Na | Plasma Me |
| Cib1 | calcium ar | Nucleus |
| Smcr8 | Smith-Ma | Nucleus |
| Glis1 | GLIS family | Nucleus |
| Tcirg1 | T cell imm | Plasma Me |
| Pbld2 | phenazine | Other |
| Ralgps1 | Ral GEF wi | Cytoplasm |
| Rbm25 | RNA bindi | Nucleus |
| Grtp1 | growth ho | Other |
| Zdhhc12 | zinc finger | Other |
| 2510039O18Rik | KIAA2013 | Other |
| Ghdc | GH3 doma | Cytoplasm |
| Golgb1 | golgin B1 | Cytoplasm |
| B4galt7 | beta-1,4-g | Cytoplasm |
| Sigirr | single Ig a | Plasma Me |
| Klc4 | kinesin ligl | Other |
| Dgat2 | diacylglyce | Cytoplasm |
| Lrp10 | LDL recept | Plasma Me |
| Trafd1 | TRAF-type | Other |
| Trip12 | thyroid ho | Cytoplasm |
| Pafah2 | platelet ac | Cytoplasm |
| Zfp948 | zinc finger | Other |
| Rabggta | Rab geran | Extracellul |
| Polr1b | RNA polyr | Nucleus |
| Psph | phosphose | Cytoplasm |
| Preb | prolactin r | Nucleus |
| Ecm1 | extracellul | Extracellul |
| Fpgs | folylpolygl | Cytoplasm |
| Syvn1 | synoviolin | Cytoplasm |
| Gpatch3 | G-patch d | Other |
| Slc12a9 | solute carr | Plasma Me |
| Gjb2 | gap junctio | Plasma Me |
| Phyh | phytanoyl- | Cytoplasm |
| Hrsp12 | - | - |
| Man2c1 | mannosida | Cytoplasm |
| Cep192 | centrosom | Cytoplasm |
| Lama5 | laminin su | Extracellul |
| Hpn | hepsin | Plasma Me |
| Mttp | microsoma | Cytoplasm |
| Tpra1 | transmeml | Plasma Me |
| Ip6k1 | inositol he | Cytoplasm |
| 1600002H07Rik | - | - |

| | | |
|---|---|---|
| Selt | - | - |
| Dnmbp | dynamin b | Cytoplasm |
| Agfg1 | ArfGAP wi | Nucleus |
| Slc7a9 | solute carr | Plasma Me |
| Arpc1b | actin relate | Cytoplasm |
| Fam120a | family with | Cytoplasm |
| Dlg5 | discs large | Plasma Me |
| Socs5 | suppresso | Extracellula |
| Dram1 | DNA dama | Cytoplasm |
| Mgrn1 | mahoguni | Cytoplasm |
| Zfp871 | zinc finger | Other |
| Pstk | phosphose | Cytoplasm |
| Hexa | hexosamin | Cytoplasm |
| Slc27a1 | solute carr | Plasma Me |
| Nle1 | notchless l | Nucleus |
| Tom1l1 | target of n | Cytoplasm |
| Plekha7 | pleckstrin | Cytoplasm |
| Gmeb1 | glucocortic | Nucleus |
| Syne4 | spectrin re | Nucleus |
| Rbp4 | retinol bin | Extracellula |
| Fastk | Fas activat | Cytoplasm |
| Ptov1 | prostate tu | Nucleus |
| AI464131 | myogenes | Nucleus |
| Rnf111 | ring finger | Nucleus |
| Zfp69 | ZFP69 zinc | Other |
| Prdm6 | PR/SET do | Nucleus |
| Dnajc4 | DnaJ heat | Cytoplasm |
| Nxf7 | nuclear RN | Cytoplasm |
| Thnsl2 | threonine | Other |
| Gphn | gephyrin | Plasma Me |
| Pdzk1 | PDZ doma | Plasma Me |
| Nipa2 | non imprin | Cytoplasm |
| Rhou | ras homolo | Cytoplasm |
| Naaa | N-acyletha | Cytoplasm |
| Sec16a | SEC16 hor | Cytoplasm |
| Xlr4b | X-linked ly | Other |
| Lrrc19 | leucine ric | Other |
| Srrm1 | serine/arg | Nucleus |
| Zfp36 | ZFP36 ring | Nucleus |
| Tmem161b | transmeml | Other |
| Trpm7 | transient r | Plasma Me |
| Tgm1 | transglutai | Plasma Me |
| Mfsd5 | major facil | Other |
| Slc20a1 | solute carr | Plasma Me |
| Slc25a28 | solute carr | Cytoplasm |
| Dqx1 | DEAQ-box | Other |
| Fem1c | fem-1 hor | Nucleus |
| Galt | galactose- | Cytoplasm |
| Araf | A-Raf prot | Cytoplasm |
| Ly75 | lymphocyt | Plasma Me |
| Msx1 | msh home | Nucleus |
| Rap1gap2 | RAP1 GTPa | Cytoplasm |
| Vil1 | villin 1 | Cytoplasm |
| Mfsd1 | major facil | Other |
| Gns | glucosami | Cytoplasm |
| Slc3a2 | solute carr | Plasma Me |
| Msx2 | msh home | Nucleus |
| Fbxl3 | F-box and | Nucleus |

| | | |
|---|---|---|
| Pkp3 | plakophilir | Plasma Me |
| Aqp11 | aquaporin | Plasma Me |
| Usp7 | ubiquitin s | Nucleus |
| Ndor1 | NADPH de | Cytoplasm |
| Mtmr12 | myotubula | Plasma Me |
| Usp36 | ubiquitin s | Nucleus |
| Parvb | parvin bet | Cytoplasm |
| Eps8l2 | EPS8 like 2 | Cytoplasm |
| Inpp5f | inositol pc | Plasma Me |
| Lars2 | leucyl-tRN | Cytoplasm |
| Irf6 | interferon | Nucleus |
| Mtg1 | mitochonc | Cytoplasm |
| Uso1 | USO1 vesi | Cytoplasm |
| Klf9 | Kruppel lik | Nucleus |
| Lzts2 | leucine zip | Cytoplasm |
| Zswim1 | zinc finger | Nucleus |
| Akr1b10 | aldo-keto | Cytoplasm |
| Edem1 | ER degrad | Cytoplasm |
| Cars2 | cysteinyl-t | Cytoplasm |
| Ehhadh | enoyl-CoA | Cytoplasm |
| Znf512b | - | - |
| Plbd1 | phospholi | Extracellula |
| Abcc5 | ATP bindir | Plasma Me |
| Abhd11 | abhydrolas | Cytoplasm |
| Ino80 | INO80 cor | Nucleus |
| Foxm1 | forkhead b | Nucleus |
| 1700019D03Rik | chromosor | Other |
| Nfkbiz | NFKB inhib | Nucleus |
| Utp20 | UTP20, sm | Nucleus |
| B4galt3 | beta-1,4-g | Cytoplasm |
| Epb4.1l3 | - | - |
| Acot8 | acyl-CoA t | Cytoplasm |
| Fastkd2 | FAST kinas | Cytoplasm |
| Rbm26 | RNA bindi | Other |
| Aplp1 | amyloid be | Extracellula |
| Ankrd13a | ankyrin rep | Plasma Me |
| Apom | apolipoprc | Plasma Me |
| Mri1 | methylthic | Cytoplasm |
| Ttr | transthyre | Extracellula |
| Hmg20b | high mobi | Nucleus |
| Lactb2 | lactamase | Cytoplasm |
| Susd6 | sushi dom | Other |
| Cldn2 | claudin 2 | Plasma Me |
| Atp6v0b | ATPase H+ | Cytoplasm |
| Ncoa3 | nuclear rec | Nucleus |
| Vwce | von Willeb | Cytoplasm |
| Tlk2 | tousled lik | Cytoplasm |
| Pisd | phosphatic | Cytoplasm |
| Bend7 | BEN doma | Cytoplasm |
| Slc22a23 | solute carr | Other |
| Mms19 | MMS19 hc | Nucleus |
| Alg13 | asparagine | Other |
| Senp5 | SUMO1/se | Nucleus |
| Slc16a9 | solute carr | Other |
| Ccdc120 | coiled-coi | Plasma Me |
| Oat | ornithine a | Cytoplasm |
| Srsf2 | serine and | Nucleus |
| Ppp2r3d | protein ph | Other |

| | | |
|---|---|---|
| Pus1 | pseudouri | Nucleus |
| Nus1 | NUS1 deh | Cytoplasm |
| Abhd17b | abhydrola | Plasma Me |
| Sypl | synaptoph | Plasma Me |
| Ankrd11 | ankyrin re | Nucleus |
| Fxr2 | FMR1 auto | Cytoplasm |
| Dnajc22 | DnaJ heat | Other |
| Manba | mannosida | Cytoplasm |
| Slco4c1 | solute carr | Plasma Me |
| Rab24 | RAB24, me | Cytoplasm |
| Abhd16a | abhydrola | Other |
| Ctsc | cathepsin | Cytoplasm |
| Rab6b | RAB6B, me | Cytoplasm |
| Wbp2 | WW doma | Cytoplasm |
| Ptpn6 | protein tyr | Cytoplasm |
| Stx4a | syntaxin 4 | Plasma Me |
| Pomt1 | protein O- | Cytoplasm |
| Ifi30 | IFI30, lysos | Cytoplasm |
| Ddt | D-dopach | Cytoplasm |
| Dlg1 | discs large | Plasma Me |
| Tcn2 | transcobal | Extracellula |
| Cndp2 | carnosine | Cytoplasm |
| Kdf1 | keratinocy | Plasma Me |
| Hsf2 | heat shock | Nucleus |
| Slc7a7 | solute carr | Plasma Me |
| Etv4 | ETS varian | Nucleus |
| Gcsh | glycine cle | Cytoplasm |
| Daam1 | dishevelle | Cytoplasm |
| Shf | Src homol | Other |
| Napa | NSF attach | Cytoplasm |
| Lnx2 | ligand of r | Other |
| Zfp956 | zinc finger | Other |
| Pkp4 | plakophilir | Plasma Me |
| Mcee | methylmal | Cytoplasm |
| Far2 | fatty acyl- | Cytoplasm |
| Vps29 | VPS29, ret | Cytoplasm |
| Xlr3b | X-linked ly | Other |
| Xlr3a | X-linked ly | Other |
| Mrps17 | mitochonc | Cytoplasm |
| Crtap | cartilage a | Extracellula |
| Slc25a15 | solute carr | Cytoplasm |
| Glrx | glutaredo | Cytoplasm |
| Stx3 | syntaxin 3 | Plasma Me |
| Kidins220 | kinase D ir | Nucleus |
| Tmem63b | transmeml | Plasma Me |
| Emilin1 | elastin mic | Extracellula |
| E4f1 | E4F transc | Nucleus |
| Tmem241 | transmeml | Other |
| Foxn3 | forkhead b | Other |
| Flcn | folliculin | Cytoplasm |
| Lgalsl | galectin lik | Other |
| Yod1 | YOD1 deu | Cytoplasm |
| Vps11 | VPS11, CC | Cytoplasm |
| Ccdc84 | coiled-coi | Other |
| Sephs2 | selenopho | Cytoplasm |
| Gpn2 | GPN-loop | Other |
| Snx2 | sorting ne | Cytoplasm |
| Xlr4a | X-linked ly | Other |

| | | |
|---|---|---|
| Inpp5b | inositol pc | Plasma Me |
| Lztr1 | leucine zip | Cytoplasm |
| Cdc14b | cell divisio | Nucleus |
| Dlk1 | delta like r | Extracellul |
| Glul | glutamate | Cytoplasm |
| Gsn | gelsolin | Extracellul |
| Efnb1 | ephrin B1 | Plasma Me |
| Ap3d1 | adaptor re | Cytoplasm |
| Plekha6 | pleckstrin | Other |
| Oxsr1 | oxidative s | Nucleus |
| Igsf5 | immunogl | Plasma Me |
| Slc39a4 | solute carr | Plasma Me |
| Pth1r | parathyroi | Plasma Me |
| Rab4a | RAB4A, me | Cytoplasm |
| 2510002D24Rik | RIKEN cDN | Other |
| Snx3 | sorting ne: | Cytoplasm |
| Haus7 | HAUS aug | Cytoplasm |
| Taf5l | TATA-box | Nucleus |
| Pgm3 | phosphog | Cytoplasm |
| Csnk1g2 | casein kina | Cytoplasm |
| Fxyd2 | FXYD dom | Plasma Me |
| Xlr4c | X-linked ly | Other |
| Twf1 | twinfilin ac | Cytoplasm |
| Rpe | ribulose-5 | Cytoplasm |
| Mrps26 | mitochonc | Cytoplasm |
| Plekha3 | pleckstrin | Cytoplasm |
| Map2k2 | mitogen-a | Cytoplasm |
| D15Ertd621e | - | - |
| Kif12 | kinesin fan | Cytoplasm |
| Cdk11b | cyclin dep | Nucleus |
| Scarb1 | scavenger | Plasma Me |
| Spry4 | sprouty RT | Plasma Me |
| Pcgf5 | polycomb | Cytoplasm |
| Gas2l3 | growth arr | Other |
| Sh3bp5 | SH3 doma | Cytoplasm |
| Amt | aminomet | Cytoplasm |
| Soat1 | sterol O-a | Cytoplasm |
| Tysnd1 | trypsin do | Cytoplasm |
| Hdac10 | histone de | Nucleus |
| Sbno1 | strawberry | Other |
| Rnf6 | ring finger | Nucleus |
| Cgn | cingulin | Plasma Me |
| Tm4sf5 | transmeml | Plasma Me |
| Rictor | RPTOR inc | Cytoplasm |
| Apoc1 | apolipopro | Extracellul |
| Strn3 | striatin 3 | Nucleus |
| Galm | galactose | Cytoplasm |
| Lamtor4 | late endos | Cytoplasm |
| Crebbp | CREB bind | Nucleus |
| Ihh | indian hed | Extracellul |
| Plekhf2 | pleckstrin | Cytoplasm |
| Setd8 | - | - |
| Rfng | RFNG O-fu | Cytoplasm |
| Ap2a2 | adaptor re | Cytoplasm |
| Aprt | adenine pl | Cytoplasm |
| Spink3 | - | - |
| Slc39a8 | solute carr | Extracellul |
| Palb2 | partner an | Nucleus |

| | | |
|---|---|---|
| Mvb12a | multivesicu | Cytoplasm |
| Stk24 | serine/thre | Cytoplasm |
| Trim44 | tripartite m | Cytoplasm |
| Nat10 | N-acetyltra | Nucleus |
| Dhps | deoxyhypu | Cytoplasm |
| Tjp3 | tight junct | Cytoplasm |
| Gm21962 | - | - |
| Hars2 | histidyl-tR | Cytoplasm |
| Ak3 | adenylate | Cytoplasm |
| Mogs | mannosyl- | Cytoplasm |
| Tmed5 | transmemb | Cytoplasm |
| Cgref1 | cell growth | Extracellula |
| Elovl1 | ELOVL fatt | Cytoplasm |
| Trmt1 | tRNA meth | Extracellula |
| Bod1 | biorientati | Nucleus |
| Mrps24 | mitochonc | Cytoplasm |
| Ric8 | - | - |
| Setd5 | SET domai | Nucleus |
| Ripk3 | receptor ir | Plasma Me |
| Ltbr | lymphotox | Plasma Me |
| Snx27 | sorting ne | Cytoplasm |
| Rock1 | Rho associ | Cytoplasm |
| Cbx7 | chromobo | Nucleus |
| Khsrp | KH-type s | Nucleus |
| Zfand2b | zinc finger | Other |
| Stx7 | syntaxin 7 | Plasma Me |
| Fam195a | - | - |
| Scamp5 | secretory c | Cytoplasm |
| Rhebl1 | RHEB like 1 | Cytoplasm |
| Ppan | peter pan | Nucleus |
| Slc38a7 | solute carr | Plasma Me |
| Bop1 | block of pr | Nucleus |
| Glmp | glycosylate | Cytoplasm |
| Bckdk | branched c | Cytoplasm |
| Eps8 | epidermal | Plasma Me |
| Ascc2 | activating | Nucleus |
| Stard10 | StAR relate | Cytoplasm |
| Sin3b | SIN3 trans | Nucleus |
| Cnot6 | CCR4-NOT | Nucleus |
| Clcn7 | chloride vo | Plasma Me |
| 2310061I04Rik | chromosor | Cytoplasm |
| Mtmr7 | myotubula | Cytoplasm |
| Grb7 | growth fac | Plasma Me |
| Lace1 | - | - |
| Ddhd2 | DDHD dor | Cytoplasm |
| Rhox5 | reproducti | Nucleus |
| Ell | elongatior | Nucleus |
| Styx | serine/thre | Cytoplasm |
| Smyd5 | SMYD fam | Other |
| Arglu1 | arginine ar | Other |
| Arhgap18 | Rho GTPas | Cytoplasm |
| 1700017B05Rik | chromosor | Cytoplasm |
| Arhgap10 | Rho GTPas | Cytoplasm |
| Rbm38 | RNA bindi | Nucleus |
| Slc37a1 | solute carr | Plasma Me |
| Slc33a1 | solute carr | Cytoplasm |
| Zfp654 | zinc finger | Other |
| Gcdh | glutaryl-C | Cytoplasm |

| | | |
|---|---|---|
| Vps16 | VPS16, CC | Cytoplasm |
| Rnf181 | ring finger | Cytoplasm |
| Arl2 | ADP ribosy | Cytoplasm |
| Abcg2 | ATP bindir | Plasma Me |
| Lxn | latexin | Cytoplasm |
| Aldh7a1 | aldehyde c | Cytoplasm |
| Acadm | acyl-CoA c | Cytoplasm |
| Agrn | agrin | Plasma Me |
| Snf8 | SNF8, ESC | Cytoplasm |
| Rbks | ribokinase | Cytoplasm |
| Cog3 | componer | Cytoplasm |
| Wee1 | WEE1 G2 c | Nucleus |
| Ube2e1 | ubiquitin c | Cytoplasm |
| Atxn2l | ataxin 2 lik | Nucleus |
| Lpcat4 | lysophosp | Cytoplasm |
| Slc2a3 | solute carr | Plasma Me |
| Ptdss2 | phosphati | Cytoplasm |
| Xlr3c | X-linked ly | Other |
| Dhrs1 | dehydroge | Cytoplasm |
| Cyba | cytochrom | Cytoplasm |
| Tmprss2 | transmeml | Plasma Me |
| Itsn2 | intersectin | Cytoplasm |
| Aifm3 | apoptosis | Cytoplasm |
| Abcc4 | ATP bindir | Plasma Me |
| Stard4 | StAR relate | Cytoplasm |
| Crlf1 | cytokine re | Extracellul |
| Lpcat3 | lysophosp | Plasma Me |
| Tsr1 | TSR1, ribos | Nucleus |
| Ank | ANKH inor | Plasma Me |
| Ankzf1 | ankyrin rep | Nucleus |
| Klf6 | Kruppel lik | Nucleus |
| Arid3c | AT-rich int | Nucleus |
| Pop5 | POP5 hom | Nucleus |
| Ezr | ezrin | Plasma Me |
| BC029214 | - | - |
| Cc2d1b | coiled-coi | Nucleus |
| Phgdh | phosphog | Cytoplasm |
| Gata4 | GATA binc | Nucleus |
| Podxl | podocalyx | Plasma Me |
| Gm21833 | - | - |
| Cd2ap | CD2 assoc | Cytoplasm |
| Gpr137b | G protein- | Plasma Me |
| Clic6 | chloride in | Plasma Me |
| Ifrd2 | interferon | Nucleus |
| Mb21d2 | Mab-21 d | Other |
| Mon1a | MON1 hor | Cytoplasm |
| Timm44 | translocas | Cytoplasm |
| Gps1 | G protein | | Nucleus |
| Zfp655 | zinc finger | Nucleus |
| Fuk | fucokinase | Cytoplasm |
| S100g | S100 calciu | Cytoplasm |
| Pla2g12b | phospholi | Extracellul |
| Tiparp | TCDD indu | Nucleus |
| Tbc1d17 | TBC1 dom | Cytoplasm |
| Igf2r | insulin like | Plasma Me |
| Wibg | - | - |
| Sap130 | Sin3A assc | Nucleus |
| Srrm2 | serine/arg | Nucleus |

| | | |
|---|---|---|
| Ehd1 | EH domair | Cytoplasm |
| Prr5 | proline ricl | Cytoplasm |
| Btaf1 | B-TFIID TA | Nucleus |
| Bag2 | BCL2 asso | Cytoplasm |
| Rabggtb | Rab geran | Cytoplasm |
| Gm5884 | target of n | Other |
| Tdh | L-threonir | Cytoplasm |
| Ftsj2 | - | - |
| Cobll1 | cordon-bl | Extracellul |
| Tpd52l1 | tumor pro | Cytoplasm |
| Mau2 | MAU2 siste | Nucleus |
| Ndufs7 | NADH:ubi | Cytoplasm |
| Man2b1 | mannosida | Cytoplasm |
| Yipf2 | Yip1 doma | Cytoplasm |
| Apba3 | amyloid be | Cytoplasm |
| Tead4 | TEA doma | Nucleus |
| Slc9a3r1 | SLC9A3 re | Plasma Me |
| Gng12 | G protein s | Plasma Me |
| Ppap2c | - | - |
| Tagap1 | T cell activ | Other |
| Cenpt | centromer | Cytoplasm |
| Sat1 | spermidine | Cytoplasm |
| Srfbp1 | serum resp | Nucleus |
| Vps35 | VPS35, ret | Cytoplasm |
| Abhd17c | abhydrolas | Plasma Me |
| Wdfy3 | WD repeat | Cytoplasm |
| Pnpla6 | patatin like | Cytoplasm |
| Bri3 | brain prote | Cytoplasm |
| Gm17555 | - | - |
| Mcts1 | MCTS1, re | Cytoplasm |
| Smg5 | SMG5, nor | Nucleus |
| C230052I12Rik | - | - |
| Exoc6 | exocyst co | Plasma Me |
| Pabpc1 | poly(A) bir | Cytoplasm |
| Phlda2 | pleckstrin | Cytoplasm |
| Arfgef2 | ADP ribosy | Cytoplasm |
| Kat2a | lysine acet | Cytoplasm |
| Pepd | peptidase | Cytoplasm |
| N4bp3 | NEDD4 bir | Cytoplasm |
| Limk2 | LIM doma | Cytoplasm |
| Pls1 | plastin 1 | Plasma Me |
| Rnf14 | ring finger | Cytoplasm |
| Rbpms2 | RNA bindi | Cytoplasm |
| Smad2 | SMAD fam | Nucleus |
| Lrpprc | leucine ric | Cytoplasm |
| Plin3 | perilipin 3 | Cytoplasm |
| Vamp2 | vesicle ass | Plasma Me |
| Acsl3 | acyl-CoA s | Cytoplasm |
| Idh3g | isocitrate c | Cytoplasm |
| Rras2 | RAS relate | Plasma Me |
| Stub1 | STIP1 hom | Cytoplasm |
| Fga | fibrinogen | Extracellul |
| Qtrt1 | queuine tR | Cytoplasm |
| Atp1a1 | ATPase Na | Plasma Me |
| Gmpr2 | guanosine | Cytoplasm |
| Cotl1 | coactosin | Cytoplasm |
| Tom1 | target of n | Cytoplasm |
| S100a1 | S100 calciu | Cytoplasm |

| | | |
|---|---|---|
| Fcgrt | Fc fragmei | Plasma Me |
| Rab20 | RAB20, me | Cytoplasm |
| Cdca2 | cell divisio | Nucleus |
| Nckap1 | NCK assoc | Plasma Me |
| Idh3b | isocitrate c | Cytoplasm |
| Isca2 | iron-sulfur | Cytoplasm |
| Vmp1 | vacuole m | Plasma Me |
| Grhpr | glyoxylate | Cytoplasm |
| Rrp9 | ribosomal | Nucleus |
| Taf1d | TATA-box | Nucleus |
| Ndufaf4 | NADH:ubi | Cytoplasm |
| Fam134a | - | - |
| E130012A19Rik | - | - |
| Katnb1 | katanin re | Cytoplasm |
| Mapkap1 | mitogen-a | Cytoplasm |
| Sqstm1 | sequestos | Cytoplasm |
| Zfand5 | zinc finger | Nucleus |
| Zcchc6 | zinc finger | Cytoplasm |
| E430025E21Rik | - | - |
| Mrpl2 | mitochonc | Extracellul |
| Gm21814 | predicted | Other |
| Nup85 | nucleopor | Cytoplasm |
| Tmem51 | transmem | Other |
| Sord | sorbitol de | Cytoplasm |
| Xpot | exportin fc | Nucleus |
| Akr7a5 | aldo-keto | Cytoplasm |
| Rere | arginine-g | Nucleus |
| Fus | fused in sa | Nucleus |
| Dus1l | dihydrouri | Other |
| Ubiad1 | UbiA pren | Nucleus |
| Atp11b | ATPase ph | Plasma Me |
| Atp6v1c1 | ATPase H+ | Cytoplasm |
| Agpat1 | 1-acylglyc | Cytoplasm |
| Emb | embigin | Plasma Me |
| Rhpn1 | rhophilin F | Cytoplasm |
| Cabin1 | calcineurir | Nucleus |
| Ppp1r14b | protein ph | Cytoplasm |
| Pip5k1c | phosphati | Plasma Me |
| Txndc17 | thioredoxi | Cytoplasm |
| Ifrd1 | interferon | Nucleus |
| Gmppa | GDP-mani | Cytoplasm |
| Sertad1 | SERTA dor | Nucleus |
| Ep400 | E1A bindir | Nucleus |
| Abcf3 | ATP bindir | Other |
| Depdc7 | DEP doma | Cytoplasm |
| Id1 | inhibitor o | Nucleus |
| Bet1l | Bet1 golgi | Cytoplasm |
| Pcbp2 | poly(rC) bi | Nucleus |
| Nomo1 | NODAL m | Plasma Me |
| Mtif2 | mitochonc | Cytoplasm |
| Stat6 | signal tran | Nucleus |
| Map1s | microtubu | Cytoplasm |
| Flnb | filamin B | Cytoplasm |
| Odc1 | ornithine c | Cytoplasm |
| Ap1g2 | adaptor re | Cytoplasm |
| Tdrp | testis deve | Cytoplasm |
| Ubxn4 | UBX doma | Extracellul |
| Mllt10 | MLLT10, h | Nucleus |

| | | |
|---|---|---|
| Afp | alpha feto | Extracellul |
| Safb2 | scaffold at | Cytoplasm |
| Rap1b | RAP1B, me | Cytoplasm |
| Hdgfrp2 | - | - |
| Gstm1 | glutathion | Cytoplasm |
| Plscr1 | phospholi | Plasma Me |
| 0610011F06Rik | - | - |
| Lamtor5 | late endos | Cytoplasm |
| Mad2l1bp | MAD2L1 b | Nucleus |
| Rragc | Ras relatec | Cytoplasm |
| Tpp1 | tripeptidyl | Cytoplasm |
| Cmpk1 | cytidine/u | Nucleus |
| Tmem55b | phosphati | Cytoplasm |
| Ftl1 | ferritin ligl | Cytoplasm |
| Pdzd3 | PDZ doma | Plasma Me |
| Commd9 | COMM dc | Cytoplasm |
| Rps6kb2 | ribosomal | Cytoplasm |
| Atp6v1d | ATPase H+ | Cytoplasm |
| Gab1 | GRB2 asso | Cytoplasm |
| Med29 | mediator c | Nucleus |
| Snx29 | sorting ne: | Cytoplasm |
| Pecr | peroxisom | Cytoplasm |
| Chd4 | chromodo | Nucleus |
| Gm3550 | ribosomal | Other |
| Uqcrq | ubiquinol- | Cytoplasm |
| Gm10116 | - | - |
| Rpl10a | ribosomal | Nucleus |
| Snx9 | sorting ne: | Cytoplasm |
| Slc7a6 | solute carr | Plasma Me |
| Serpinf2 | serpin fam | Extracellul |
| Scpep1 | serine carb | Cytoplasm |
| Tctn3 | tectonic fa | Cytoplasm |
| Cdc25c | cell divisio | Nucleus |
| Shmt2 | serine hyd | Cytoplasm |
| Fem1b | fem-1 hor | Nucleus |
| Jtb | jumping tr | Plasma Me |
| Cdkn3 | cyclin dep | Cytoplasm |
| Rnf128 | ring finger | Cytoplasm |
| Id2 | inhibitor o | Nucleus |
| Tmem9b | TMEM9 dc | Other |
| Tor1aip2 | torsin 1A i | Cytoplasm |
| Acaa2 | acetyl-Co/ | Cytoplasm |
| Gm21750 | - | - |
| Maz | MYC assoc | Nucleus |
| Nhp2 | NHP2 ribo | Nucleus |
| Rhog | ras homol | Cytoplasm |
| Zfp280b | zinc finger | Nucleus |
| Slc16a13 | solute carr | Cytoplasm |
| Nrde2 | NRDE-2, n | Other |
| Setmar | SET domai | Nucleus |
| Grina | glutamate | Other |
| Eif4g1 | eukaryotic | Cytoplasm |
| Cd63 | CD63 mol | Plasma Me |
| 9430015G10Rik | chromoso | Other |
| Asah1 | N-acylsph | Cytoplasm |
| Igf2bp2 | insulin like | Cytoplasm |
| Ppfibp2 | PPFIA binc | Nucleus |
| Tnfrsf12a | TNF recep | Plasma Me |

| | | |
|---|---|---|
| Otub1 | OTU doma | Cytoplasm |
| Trappc6b | trafficking | Cytoplasm |
| Atad3a | ATPase far | Cytoplasm |
| 2210011C24Rik | RIKEN cDN | Other |
| Phlda1 | pleckstrin | Cytoplasm |
| Fam21 | - | - |
| Rbck1 | RANBP2-t | Cytoplasm |
| Fth1 | ferritin hea | Cytoplasm |
| Cdk20 | cyclin dep | Nucleus |
| Phf13 | PHD finge | Nucleus |
| Ugdh | UDP-gluc | Nucleus |
| Slc35f5 | solute carr | Other |
| Mapk6 | mitogen-a | Cytoplasm |
| Tmem120a | transmeml | Other |
| Pip4k2c | phosphati | Cytoplasm |
| Nt5dc2 | 5'-nucleot | Cytoplasm |
| Prss8 | protease, s | Extracellul |
| Emg1 | EMG1, N1 | Nucleus |
| Cyp26a1 | cytochrom | Cytoplasm |
| Pxdc1 | PX domair | Other |
| Usp15 | ubiquitin s | Cytoplasm |
| Ccdc61 | coiled-coi | Cytoplasm |
| Arhgap1 | Rho GTPas | Cytoplasm |
| Dusp4 | dual speci | Nucleus |
| Slc35b1 | solute carr | Cytoplasm |
| Nop56 | NOP56 rib | Nucleus |
| Atp5d | ATP synth | Cytoplasm |
| Gyltl1b | - | - |
| Sdc4 | syndecan 4 | Plasma Me |
| Fmr1nb | fragile X m | Nucleus |
| Noc2l | NOC2 like | Nucleus |
| Scyl1 | SCY1 like p | Cytoplasm |
| Snrnp48 | small nucl | Nucleus |
| Pigyl | phosphati | Plasma Me |
| Tomm40 | translocas | Cytoplasm |
| Yipf3 | Yip1 doma | Cytoplasm |
| Fgg | fibrinogen | Extracellul |
| Clasrp | CLK4-assc | Nucleus |
| Tmem180 | - | - |
| Pacsin2 | protein kir | Cytoplasm |
| Dpp4 | dipeptidyl | Plasma Me |
| Mogat2 | monoacylg | Cytoplasm |
| Lamp2 | lysosomal | Plasma Me |
| Nubp2 | nucleotide | Nucleus |
| Cldn6 | claudin 6 | Plasma Me |
| Scaf4 | SR-relatec | Nucleus |
| Tusc2 | tumor sup | Nucleus |
| Tob1 | transducer | Nucleus |
| Ybx3 | Y-box bin | Nucleus |
| Chchd3 | coiled-coi | Cytoplasm |
| Deb1 | - | - |
| Cyb5a | cytochrom | Cytoplasm |
| Snapin | SNAP assc | Plasma Me |
| Sec31a | SEC31 hor | Cytoplasm |
| Snrnp35 | small nucl | Nucleus |
| Gnpda1 | glucosami | Cytoplasm |
| Terf1 | telomeric | Nucleus |

e that the gene is more highly expressed in the given cluster.

| Family |
|---|
| transcription regulator |
| transmembrane receptor |
| enzyme |
| other |
| other |
| other |
| peptidase |
| transcription regulator |
| transcription regulator |
| other |
| other |
| transmembrane receptor |
| other |
| growth factor |
| growth factor |
| other |
| transmembrane receptor |
| transmembrane receptor |
| other |
| other |
| other |
| peptidase |
| other |
| enzyme |
| - |
| other |
| other |
| other |
| other |
| other |
| - |
| other |
| other |
| peptidase |
| peptidase |
| other |
| other |
| other |
| other |
| other |
| other |
| kinase |
| G-protein coupled receptor |
| transporter |
| other |

growth factor
other
other
peptidase
other
phosphatase
other
other
growth factor
enzyme
other
enzyme
other
other
other
enzyme
other
other
other
-
-
other
other
other
transporter
other
cytokine
other
other
ion channel
enzyme
kinase
G-protein coupled receptor
transporter
other
other
transporter
other
cytokine
other
other
other
other
transcription regulator
other
other
other
other
peptidase
other
other
other
enzyme
enzyme
other
other
other
other

other
other
other
enzyme
transcription regulator
enzyme
other
peptidase
growth factor
enzyme
ion channel
other
transmembrane receptor
other
other
transcription regulator
enzyme
phosphatase
other
other
peptidase
other
transporter
enzyme
transporter
other
other
other
other
kinase
other
transcription regulator
peptidase
other
phosphatase
other
other
other
other
other
other
transporter
peptidase
other
enzyme
kinase
growth factor
transcription regulator
enzyme
other
other
other
enzyme
other
other
other
-
transmembrane receptor

enzyme
transporter
ion channel
enzyme
other
cytokine
other
G-protein coupled receptor
transporter
transporter
ion channel
enzyme
other
other
peptidase
enzyme
other
other
transporter
transporter
transcription regulator
other
transporter
peptidase
other
other
other
other
transporter
other
enzyme
other
enzyme
enzyme
other
enzyme
enzyme
transporter
enzyme
transcription regulator
enzyme
other
other
other
other
enzyme
growth factor
enzyme
other
peptidase
transcription regulator
other
transcription regulator
other
other
-
transporter
enzyme

enzyme
peptidase
transporter
other
other
transporter
transporter
other
transcription regulator
transporter
other
enzyme
transporter
other
transporter
transcription regulator
transcription regulator
other
other
other
other
enzyme
transmembrane receptor
phosphatase
transcription regulator
other
other
enzyme
transmembrane receptor
other
other
transporter
other
kinase
transporter
other
other
other
transporter
transporter
other
other
enzyme
enzyme
other
other
other
enzyme
enzyme
transcription regulator
other
transcription regulator
enzyme
translation regulator
transporter
transporter
transporter
enzyme

enzyme
other
enzyme
other
other
other
enzyme
transporter
other
other
other
enzyme
other
ion channel
other
enzyme
other
other
other
peptidase
transcription regulator
other
other
other
other
enzyme
transporter
other
transporter
enzyme
peptidase
other
enzyme
other
other
transcription regulator
other
enzyme
transcription regulator
other
transporter
other
transporter
kinase
other
other
other
cytokine
enzyme
enzyme
transcription regulator
other
other
other
other
other
other
transporter

peptidase
other
G-protein coupled receptor
enzyme
other
other
other
other
enzyme
transporter
other
other
transporter
other
enzyme
other
peptidase
transcription regulator
other
transcription regulator
other
G-protein coupled receptor
other
peptidase
enzyme
enzyme
peptidase
enzyme
enzyme
other
other
other
other
enzyme
enzyme
enzyme
other
other
other
other
enzyme
other
other
other
transcription regulator
enzyme
other
enzyme
other
transmembrane receptor
enzyme
other
other
other
other
other
enzyme
other

transcription regulator
transcription regulator
transcription regulator
-
other
enzyme
transmembrane receptor
other
other
enzyme
other
enzyme
peptidase
enzyme
other
transcription regulator
other
other
enzyme
other
other
other
kinase
enzyme
other
other
transporter
ligand-dependent nuclear receptor
transporter
other
transcription regulator
other
other
transcription regulator
enzyme
enzyme
other
other
transporter
enzyme
enzyme
enzyme
other
other
other
other
other
transcription regulator
enzyme
other
enzyme
transcription regulator
transporter
other
transporter
enzyme
transporter
other

other
enzyme
other
enzyme
other
enzyme
peptidase
enzyme
other
peptidase
other
other
other
growth factor
other
growth factor
other
-
-
enzyme
other
transporter
other
enzyme
transcription regulator
other
phosphatase
enzyme
other
kinase
translation regulator
enzyme
enzyme
other
other
other
transporter
other
other
other
transcription regulator
other
peptidase
transcription regulator
transporter
enzyme
other
other
enzyme
transcription regulator
enzyme
other
other
other
other
other
enzyme
other

-
enzyme
transcription regulator
other
transporter
kinase
other
other
other
other
other
growth factor
other
transcription regulator
other
kinase
other
transcription regulator
transcription regulator
transcription regulator
transcription regulator
transcription regulator
other
other
other
enzyme
transporter
transporter
other
other
other
enzyme
other
transmembrane receptor
other
transcription regulator
transcription regulator
transporter
other
G-protein coupled receptor
kinase
ligand-dependent nuclear receptor
other
enzyme
other
other
other
other
other
transmembrane receptor
growth factor
kinase
other
other
enzyme
enzyme
transcription regulator
other

other
kinase
transmembrane receptor
other
enzyme
transcription regulator
other
other
other
transcription regulator
other
transcription regulator
kinase
transcription regulator
transcription regulator
other
transcription regulator
transcription regulator
enzyme
enzyme
other
other
transmembrane receptor
transcription regulator
transcription regulator
transcription regulator
other
other
transcription regulator
enzyme
peptidase
other
other
other
other
transcription regulator
enzyme
phosphatase
other
other
other
other
transporter
other
other
other
transcription regulator
other
other
enzyme
enzyme
G-protein coupled receptor
G-protein coupled receptor
other
other
enzyme
ion channel
ligand-dependent nuclear receptor

other
other
other
other
enzyme
other
other
other
other
other
enzyme
enzyme
other
-
other
phosphatase
other
other
enzyme
other
other
enzyme
kinase
transcription regulator
other
other
other
other
enzyme
other
growth factor
enzyme
enzyme
transporter
other
transcription regulator
-
other
other
transcription regulator
other
other
other
transcription regulator
kinase
enzyme
transcription regulator
enzyme
other
enzyme
-
transcription regulator
transcription regulator
kinase
transporter
other
enzyme
other

transcription regulator
other
other
other
kinase
translation regulator
kinase
transporter
other
enzyme
other
enzyme
other
enzyme
other
transcription regulator
transcription regulator
-
ion channel
other
kinase
transporter
other
other
other
kinase
phosphatase
other
other
other
other
other
other
growth factor
transcription regulator
other
other
other
other
transcription regulator
transcription regulator
kinase
other
enzyme
other
kinase
other
enzyme
other
translation regulator
other
enzyme
enzyme
other
enzyme
other
enzyme
enzyme

other
enzyme
transporter
transcription regulator
other
other
transcription regulator
transcription regulator
other
other
other
other
enzyme
translation regulator
kinase
other
other
phosphatase
phosphatase
other
other
other
enzyme
transporter
other
other
enzyme
other
other
other
other
other
transcription regulator
transporter
other
transcription regulator
other
other
-
transcription regulator
transporter
other
enzyme
other
other
other
kinase
other
enzyme
other
enzyme
other
other
kinase
enzyme
transmembrane receptor
other
enzyme

enzyme
enzyme
other
other
transporter
other
enzyme
enzyme
enzyme
transcription regulator
enzyme
other
transcription regulator
translation regulator
enzyme
other
enzyme
other
enzyme
transcription regulator
other
enzyme
transcription regulator
enzyme
enzyme
other
other
transporter
enzyme
other
other
other
transcription regulator
other
other
enzyme
enzyme
enzyme
kinase
transcription regulator
enzyme
other
other
cytokine
growth factor
other
growth factor
transcription regulator
transcription regulator
other
growth factor
kinase
transcription regulator
other
other
transcription regulator
other
-

other
other
other
other
other
other
other
other
other
growth factor
transmembrane receptor
other
other
other
other
enzyme
other
other
other
other
other
other
other
other
other
other
other
other
other
enzyme
other
other
enzyme
translation regulator
other
other
transcription regulator
other
other
other
enzyme
enzyme
other
transporter
other
other
other
other
other
other
other
other
other
other
other
translation regulator
peptidase
other

other
other
other
other
other
other
other
other
other
other
enzyme
other
transcription regulator
translation regulator
transcription regulator
other
other
other
other
other
other
kinase
other
other
transmembrane receptor
other
other
other
enzyme
other
other
other
other
translation regulator
other
other
enzyme
other
-
other
enzyme
translation regulator
phosphatase
other
other
other
other
other
translation regulator
other
other
other
transporter
other
enzyme
transporter
other
other

other
other
enzyme
other
enzyme
other
transcription regulator
transcription regulator
other
translation regulator
enzyme
enzyme
other
-
enzyme
other
enzyme
cytokine
other
transporter
enzyme
translation regulator
other
enzyme
enzyme
other
other
peptidase
other
other
other
growth factor
other
other
other
-
other
enzyme
transporter
peptidase
enzyme
enzyme
other
other
translation regulator
other
other
enzyme
other
other
-
enzyme
other
enzyme
other
enzyme
peptidase
transporter

enzyme
enzyme
other
other
enzyme
transporter
other
transcription regulator
other
other
transporter
transcription regulator
other
other
enzyme
other
enzyme
enzyme
other
other
transcription regulator
other
other
transporter
transcription regulator
-
enzyme
other
transcription regulator
enzyme
other
other
other
other
-
other
other
phosphatase
transcription regulator
other
other
other
other
other
other
transporter
other
translation regulator
other
peptidase
other
other
enzyme
peptidase
other
enzyme
other
peptidase

transcription regulator
peptidase
growth factor
other
other
other
kinase
other
enzyme
translation regulator
other
other
enzyme
enzyme
other
enzyme
other
enzyme
transporter
other
other
enzyme
other
transporter
enzyme
other
transporter
other
peptidase
transporter
enzyme
other
peptidase
enzyme
translation regulator
enzyme
enzyme
enzyme
other
transcription regulator
kinase
growth factor
enzyme
other
-
transcription regulator
peptidase
peptidase
transcription regulator
other
enzyme
other
enzyme
other
enzyme
transporter
other
other

other
other
other
other
enzyme
enzyme
transporter
other
enzyme
other
transcription regulator
other
phosphatase
enzyme
other
peptidase
enzyme
other
other
other
transporter
other
enzyme
other
transcription regulator
other
other
other
growth factor
transporter
transcription regulator
enzyme
transcription regulator
transcription regulator
other
enzyme
other
enzyme
-
other
transcription regulator
transporter
enzyme
peptidase
other
other
transporter
other
enzyme
other
other
other
other
other
other
translation regulator
other
enzyme

kinase
enzyme
enzyme
enzyme
phosphatase
other
enzyme
transporter
kinase
other
other
transcription regulator
other
other
translation regulator
other
transcription regulator
other
other
kinase
transporter
other
transmembrane receptor
kinase
transporter
transcription regulator
enzyme
transporter
ion channel
transporter
other
other
enzyme
other
enzyme
kinase
transcription regulator
other
other
phosphatase
transporter
transcription regulator
transporter
other
enzyme
other
enzyme
ion channel
peptidase
transporter
transporter
other
other
other
peptidase
other
phosphatase
other

enzyme
other
other
transporter
other
other
enzyme
enzyme
enzyme
transporter
other
enzyme
enzyme
transporter
transporter
other
enzyme
enzyme
transporter
enzyme
transporter
transcription regulator
other
kinase
other
transporter
phosphatase
other
transcription regulator
transporter
transcription regulator
enzyme
transcription regulator
transporter
kinase
transporter
transcription regulator
enzyme
other
transporter
other
enzyme
peptidase
peptidase
other
transcription regulator
transporter
peptidase
transporter
enzyme
other
transporter
other
enzyme
phosphatase
other
other
transporter

kinase
transcription regulator
-
transporter
enzyme
other
other
ion channel
transcription regulator
peptidase
other
ion channel
other
transcription regulator
transporter
translation regulator
enzyme
transporter
transcription regulator
other
other
other
peptidase
other
other
other
other
other
other
other
other
kinase
other
other
transporter
enzyme
other
other
peptidase
enzyme
other
enzyme
enzyme
ion channel
other
phosphatase
other
other
other
other
kinase
transporter
transporter
enzyme
enzyme
other
other
other

enzyme
enzyme
enzyme
transcription regulator
other
other
other
G-protein coupled receptor
other
ion channel
ion channel
enzyme
enzyme
other
transporter
other
transporter
transporter
transcription regulator
other
other
transcription regulator
other
enzyme
transporter
transcription regulator
transcription regulator
other
enzyme
transporter
transporter
enzyme
other
other
enzyme
other
enzyme
enzyme
peptidase
enzyme
transcription regulator
other
enzyme
cytokine
other
transporter
transcription regulator
transporter
G-protein coupled receptor
transporter
-
peptidase
enzyme
enzyme
other
other
other
transporter

other
other
transcription regulator
other
other
other
other
other
other
other
other
other
peptidase
other
other
other
transporter
other
other
transcription regulator
enzyme
other
other
other
other
enzyme
other
other
other
enzyme
transmembrane receptor
other
enzyme
transmembrane receptor
other
enzyme
enzyme
other
enzyme
enzyme
phosphatase
transcription regulator
transporter
enzyme
transporter
other
transporter
transporter
enzyme
-
enzyme
other
other
peptidase
transporter
G-protein coupled receptor
kinase
-

-
other
other
transporter
other
other
other
cytokine
other
enzyme
other
kinase
enzyme
transporter
enzyme
other
other
transcription regulator
other
other
kinase
other
other
enzyme
other
other
other
other
other
enzyme
transporter
other
enzyme
enzyme
phosphatase
other
other
other
transcription regulator
other
kinase
enzyme
transporter
transporter
transporter
enzyme
transcription regulator
enzyme
kinase
transmembrane receptor
transcription regulator
other
other
transporter
enzyme
transporter
transcription regulator
enzyme

other
transporter
peptidase
enzyme
other
peptidase
other
other
phosphatase
enzyme
transcription regulator
enzyme
transporter
transcription regulator
other
other
enzyme
enzyme
enzyme
enzyme
-
enzyme
transporter
enzyme
enzyme
transcription regulator
other
transcription regulator
other
enzyme
-
enzyme
other
other
other
other
transporter
translation regulator
transporter
transcription regulator
enzyme
other
other
transporter
transcription regulator
other
kinase
enzyme
other
transporter
transcription regulator
enzyme
peptidase
other
other
enzyme
transcription regulator
phosphatase

enzyme
enzyme
peptidase
transporter
other
other
other
enzyme
transporter
other
other
peptidase
enzyme
transcription regulator
phosphatase
transporter
enzyme
enzyme
enzyme
kinase
transporter
peptidase
other
transcription regulator
transporter
transcription regulator
enzyme
other
other
transporter
other
other
other
enzyme
enzyme
transporter
other
other
other
other
transporter
enzyme
transporter
transcription regulator
other
other
transcription regulator
other
other
other
other
enzyme
transporter
other
enzyme
other
transporter
other

phosphatase
transcription regulator
phosphatase
other
enzyme
other
other
transporter
other
kinase
other
transporter
G-protein coupled receptor
enzyme
other
transporter
other
transcription regulator
enzyme
kinase
other
other
kinase
enzyme
other
other
kinase
-
other
kinase
transporter
other
other
other
other
enzyme
enzyme
peptidase
transcription regulator
enzyme
transcription regulator
other
other
other
other
transcription regulator
enzyme
other
transcription regulator
enzyme
other
-
enzyme
transporter
enzyme
-
transporter
other

other
kinase
other
enzyme
enzyme
other
-
enzyme
kinase
enzyme
other
other
enzyme
enzyme
other
other
-
other
kinase
transmembrane receptor
other
kinase
other
enzyme
other
transporter
-
other
enzyme
other
transporter
other
transcription regulator
kinase
peptidase
other
other
transcription regulator
enzyme
ion channel
other
phosphatase
other
-
enzyme
transcription regulator
transcription regulator
phosphatase
other
other
other
other
other
other
transporter
transporter
other
enzyme

transporter
enzyme
enzyme
transporter
other
enzyme
enzyme
other
enzyme
kinase
transporter
kinase
enzyme
other
enzyme
transporter
enzyme
other
enzyme
enzyme
peptidase
other
enzyme
transporter
transporter
other
enzyme
other
transporter
transcription regulator
transcription regulator
transcription regulator
enzyme
other
-
transcription regulator
enzyme
transcription regulator
other
-
other
other
ion channel
other
other
other
transporter
other
other
kinase
other
enzyme
enzyme
other
transmembrane receptor
-
transcription regulator
other

other
other
transcription regulator
other
enzyme
other
enzyme
-
other
other
other
enzyme
enzyme
other
transporter
transcription regulator
other
enzyme
-
other
other
enzyme
other
transporter
enzyme
enzyme
enzyme
other
-
other
other
-
transporter
translation regulator
other
other
enzyme
peptidase
other
kinase
other
transcription regulator
other
transcription regulator
other
other
other
enzyme
enzyme
enzyme
enzyme
other
enzyme
transporter
enzyme
other
transporter
other

transmembrane receptor
enzyme
other
other
enzyme
other
other
enzyme
other
other
other
-
-
enzyme
other
transcription regulator
other
enzyme
-
other
other
other
other
enzyme
other
enzyme
transcription regulator
transcription regulator
other
enzyme
transporter
transporter
enzyme
transporter
other
other
phosphatase
kinase
enzyme
other
enzyme
transcription regulator
other
transporter
other
transcription regulator
transporter
other
other
translation regulator
transcription regulator
enzyme
other
enzyme
transporter
other
other
transcription regulator

transporter
other
enzyme
-
enzyme
enzyme
-
other
other
enzyme
peptidase
kinase
phosphatase
enzyme
transporter
other
kinase
transporter
kinase
other
other
enzyme
enzyme
other
enzyme
-
other
transporter
transporter
other
peptidase
other
phosphatase
enzyme
transcription regulator
other
phosphatase
enzyme
transcription regulator
other
other
enzyme
-
transcription regulator
other
enzyme
other
other
other
enzyme
ion channel
translation regulator
other
other
enzyme
translation regulator
phosphatase
transmembrane receptor

enzyme
other
other
other
other
-
transcription regulator
enzyme
kinase
other
enzyme
other
kinase
other
kinase
other
peptidase
enzyme
enzyme
other
peptidase
other
other
phosphatase
transporter
other
transporter
-
other
other
transcription regulator
kinase
other
other
ion channel
other
other
other
-
transporter
peptidase
enzyme
enzyme
other
other
other
other
transcription regulator
transcription regulator
transcription regulator
-
enzyme
other
other
other
enzyme
other

### Differentially expressed genes (DEGs) between FG-1

- **lb**:lower bound, **mle**:maximum likelihood estimate, **ub**: upper bo
- **ce:** conservative estimate of expression-fold change (equals to th
- **Z:** uncorrected Z-score of expression difference
- **cZ:** expression difference Z-score corrected for multiple hypothe
- **Genes highly expressed in the FG-1 presented positive value**

| | lb | mle | ub | ce | Z | cZ |
|---|---|---|---|---|---|---|
| ENSMUSG00000000805 | -5.21917 | -3.20267 | -2.31304 | -2.31304 | -7.16081 | -5.77894 |
| ENSMUSG00000061808 | -11.0611 | -8.09564 | -5.66398 | -5.66398 | -7.16085 | -5.77894 |
| ENSMUSG00000037379 | -7.8584 | -6.76119 | -5.48606 | -5.48606 | -6.03963 | -4.42483 |
| ENSMUSG00000018822 | -10.6163 | -9.37078 | -6.55361 | -6.55361 | -5.96338 | -4.41371 |
| ENSMUSG00000033831 | -9.16319 | -8.06598 | -4.00334 | -4.00334 | -5.94825 | -4.41371 |
| ENSMUSG00000019900 | -8.30322 | -7.32462 | -5.96052 | -5.96052 | -5.83864 | -4.30841 |
| ENSMUSG00000025075 | -8.86665 | -7.65082 | -5.75294 | -5.75294 | -5.66187 | -4.13766 |
| ENSMUSG00000037370 | -9.04458 | -8.15495 | -4.47781 | -4.47781 | -5.67347 | -4.13766 |
| ENSMUSG00000028583 | -6.1088 | -4.50746 | -2.84682 | -2.84682 | -5.50895 | -3.95924 |
| ENSMUSG00000026380 | -8.03633 | -7.05774 | -5.1302 | -5.1302 | -5.46799 | -3.92893 |
| ENSMUSG00000054932 | -10.3197 | -7.50255 | -4.06265 | -4.06265 | -5.42975 | -3.90005 |
| ENSMUSG00000024621 | -7.71013 | -6.79085 | -5.21917 | -5.21917 | -5.37605 | -3.86491 |
| ENSMUSG00000061751 | -5.96052 | -3.29163 | -2.10546 | -2.10546 | -5.37386 | -3.86491 |
| ENSMUSG00000028024 | -8.27356 | -7.35428 | -5.57502 | -5.57502 | -5.30131 | -3.78461 |
| ENSMUSG00000015627 | -7.50255 | -6.61292 | -4.29988 | -4.29988 | -5.12015 | -3.55253 |
| ENSMUSG00000024140 | -7.82875 | -6.64257 | -4.29988 | -4.29988 | -5.08881 | -3.52583 |
| ENSMUSG00000026295 | -8.00667 | -5.48606 | -3.85507 | -3.85507 | -5.02629 | -3.46983 |
| ENSMUSG00000039385 | -8.98527 | -8.00667 | -4.06265 | -4.06265 | -5.02805 | -3.46983 |
| ENSMUSG00000074457 | -7.17635 | -6.25707 | -3.94403 | -3.94403 | -4.9947 | -3.44014 |
| ENSMUSG00000026175 | -7.62117 | -6.55361 | -3.88472 | -3.88472 | -4.92279 | -3.36638 |
| ENSMUSG00000049154 | -6.46465 | -5.51571 | -3.58818 | -3.58818 | -4.92764 | -3.36638 |
| ENSMUSG00000021944 | -7.62117 | -6.79085 | -4.50746 | -4.50746 | -4.89582 | -3.35094 |
| ENSMUSG00000059481 | -8.59976 | -6.79085 | -4.32954 | -4.32954 | -4.89399 | -3.35094 |
| ENSMUSG00000024386 | -7.94737 | -6.37569 | -3.7661 | -3.7661 | -4.80853 | -3.25347 |
| ENSMUSG00000033577 | -8.06598 | -7.02808 | -4.47781 | -4.47781 | -4.81297 | -3.25347 |
| ENSMUSG00000020609 | -9.34112 | -5.30813 | -4.18126 | -4.18126 | -4.73357 | -3.17457 |
| ENSMUSG00000074217 | -6.67223 | -5.7826 | -3.08405 | -3.08405 | -4.73271 | -3.17457 |
| ENSMUSG00000024503 | -5.93087 | -4.21092 | -2.25373 | -2.25373 | -4.71362 | -3.16086 |
| ENSMUSG00000022790 | -6.76119 | -5.81225 | -3.23232 | -3.23232 | -4.6917 | -3.15824 |
| ENSMUSG00000036832 | -9.19285 | -7.71013 | -3.7661 | -3.7661 | -4.69431 | -3.15824 |
| ENSMUSG00000043013 | -7.4729 | -6.55361 | -3.88472 | -3.88472 | -4.70124 | -3.15824 |
| ENSMUSG00000021876 | -8.45149 | -7.41359 | -3.97368 | -3.97368 | -4.66863 | -3.14308 |
| ENSMUSG00000031621 | -7.20601 | -6.31638 | -3.23232 | -3.23232 | -4.5731 | -3.0499 |
| ENSMUSG00000030208 | -6.73154 | -5.60467 | -3.52887 | -3.52887 | -4.46465 | -2.92036 |
| ENSMUSG00000020785 | -7.26532 | -5.96052 | -2.34269 | -2.34269 | -4.441 | -2.90013 |
| ENSMUSG00000024391 | -8.89631 | -6.34603 | -4.03299 | -4.03299 | -4.42732 | -2.88709 |
| ENSMUSG00000059456 | -7.02808 | -5.99018 | -2.93578 | -2.93578 | -4.40178 | -2.85657 |
| ENSMUSG00000061947 | -6.67223 | -5.72329 | -3.17302 | -3.17302 | -4.38586 | -2.84654 |
| ENSMUSG00000001943 | -6.07914 | -3.82541 | -2.28339 | -2.28339 | -4.37309 | -2.83429 |
| ENSMUSG00000045064 | -6.37569 | -5.4564 | -1.95719 | -1.95719 | -4.29447 | -2.73107 |
| ENSMUSG00000029103 | -3.73645 | -2.63924 | -1.45306 | -1.45306 | -4.25927 | -2.68495 |
| ENSMUSG00000002847 | -5.93087 | -4.89297 | -2.402 | -2.402 | -4.23759 | -2.65875 |
| ENSMUSG00000024990 | -10.2308 | -9.37078 | -3.41025 | -3.41025 | -4.16357 | -2.56545 |
| ENSMUSG00000035407 | -6.4943 | -4.7447 | -1.92753 | -1.92753 | -4.16745 | -2.56545 |
| ENSMUSG00000091705 | -6.55361 | -5.07089 | -2.84682 | -2.84682 | -4.10847 | -2.49334 |
| ENSMUSG00000028076 | -6.52396 | -5.4564 | -2.87647 | -2.87647 | -4.10418 | -2.49271 |
| ENSMUSG00000066366 | -7.76944 | -5.54536 | -2.0165 | -2.0165 | -4.09569 | -2.49133 |

| | | | | | | |
|---|---|---|---|---|---|---|
| ENSMUSG00000011256 | -6.70188 | -5.18951 | -3.20267 | -3.20267 | -4.0023 | -2.37813 |
| ENSMUSG00000022150 | -5.51571 | -3.79576 | -1.80892 | -1.80892 | -3.94311 | -2.31936 |
| ENSMUSG00000034574 | -7.08739 | -6.25707 | -2.84682 | -2.84682 | -3.91732 | -2.28386 |
| ENSMUSG00000037440 | -8.21426 | -7.50255 | -2.0758 | -2.0758 | -3.87782 | -2.26027 |
| ENSMUSG00000027249 | -6.67223 | -5.66398 | -2.43166 | -2.43166 | -3.87073 | -2.2537 |
| ENSMUSG00000017493 | -5.10055 | -3.35094 | -1.48272 | -1.48272 | -3.83782 | -2.21068 |
| ENSMUSG00000026399 | -8.33287 | -7.26532 | -2.55027 | -2.55027 | -3.80038 | -2.17324 |
| ENSMUSG00000049858 | -6.90946 | -2.99509 | -1.54203 | -1.54203 | -3.74991 | -2.10516 |
| ENSMUSG00000069833 | -7.56186 | -6.73154 | -1.92753 | -1.92753 | -3.71093 | -2.05514 |
| ENSMUSG00000035376 | -4.47781 | -2.87647 | -1.3641 | -1.3641 | -3.69747 | -2.04535 |
| ENSMUSG00000041075 | -7.62117 | -3.94403 | -1.63099 | -1.63099 | -3.67601 | -2.0185 |
| ENSMUSG00000068874 | -5.93087 | -3.91437 | -1.0379 | -1.0379 | -3.66118 | -2.00631 |
| ENSMUSG00000021806 | -4.0923 | -2.25373 | -1.00825 | -1.00825 | -3.62421 | -1.95475 |
| ENSMUSG00000029735 | -6.99843 | -5.72329 | -3.02474 | -3.02474 | -3.62266 | -1.95475 |
| ENSMUSG00000024781 | -3.3806 | -2.22408 | -1.0379 | -1.0379 | -3.60567 | -1.93443 |
| ENSMUSG00000029634 | -7.32462 | -6.64257 | -1.54203 | -1.54203 | -3.58729 | -1.91162 |
| ENSMUSG00000028158 | -6.34603 | -4.00334 | -1.83857 | -1.83857 | -3.5293 | -1.84836 |
| ENSMUSG00000026726 | -6.58327 | -5.15986 | -2.43166 | -2.43166 | -3.50918 | -1.82762 |
| ENSMUSG00000019998 | -4.68539 | -2.25373 | -1.00825 | -1.00825 | -3.49266 | -1.8081 |
| ENSMUSG00000021850 | -7.97702 | -7.08739 | -2.22408 | -2.22408 | -3.45908 | -1.76851 |
| ENSMUSG00000043683 | -5.81225 | -4.35919 | -1.48272 | -1.48272 | -3.44909 | -1.75488 |
| ENSMUSG00000006538 | -6.61292 | -5.60467 | -2.28339 | -2.28339 | -3.44032 | -1.75018 |
| ENSMUSG00000017781 | -6.46465 | -5.30813 | -1.80892 | -1.80892 | -3.44168 | -1.75018 |
| ENSMUSG00000009563 | -4.83366 | -2.75785 | -1.15652 | -1.15652 | -3.42926 | -1.73798 |
| ENSMUSG00000030849 | -7.35428 | -6.4943 | -1.57168 | -1.57168 | -3.41565 | -1.72915 |
| ENSMUSG00000027082 | -8.27356 | -4.89297 | -1.92753 | -1.92753 | -3.3901 | -1.69694 |
| ENSMUSG00000021583 | -6.58327 | -5.39709 | -1.21583 | -1.21583 | -3.3525 | -1.64119 |
| ENSMUSG00000020679 | -7.73979 | -6.93912 | -1.60133 | -1.60133 | -3.34424 | -1.63067 |
| ENSMUSG00000023022 | -5.39709 | -2.96544 | -1.24548 | -1.24548 | -3.27855 | -1.53971 |
| ENSMUSG00000031216 | -7.65082 | -6.43499 | -1.92753 | -1.92753 | -3.27926 | -1.53971 |
| ENSMUSG00000038668 | -5.72329 | -2.90613 | -1.18617 | -1.18617 | -3.27715 | -1.53971 |
| ENSMUSG00000026880 | -7.26532 | -6.22741 | -2.04615 | -2.04615 | -3.27186 | -1.53339 |
| ENSMUSG00000026866 | -6.40534 | -4.53712 | -1.15652 | -1.15652 | -3.25218 | -1.51053 |
| ENSMUSG00000024486 | -7.26532 | -6.28672 | -1.95719 | -1.95719 | -3.24474 | -1.50022 |
| ENSMUSG00000063354 | -5.24882 | -2.93578 | -1.09721 | -1.09721 | -3.22726 | -1.47782 |
| ENSMUSG00000026235 | -6.85016 | -3.3806 | -1.54203 | -1.54203 | -3.22383 | -1.47699 |
| ENSMUSG00000041058 | -5.90122 | -4.0923 | -1.42341 | -1.42341 | -3.221 | -1.47563 |
| ENSMUSG00000048706 | -7.94737 | -7.08739 | -2.81716 | -2.81716 | -3.20926 | -1.46334 |
| ENSMUSG00000013418 | -7.62117 | -6.79085 | -1.77926 | -1.77926 | -3.20126 | -1.45175 |
| ENSMUSG00000026401 | -6.01983 | -4.59642 | -2.0165 | -2.0165 | -3.19546 | -1.45125 |
| ENSMUSG00000040152 | -6.43499 | -5.27847 | -1.30479 | -1.30479 | -3.17516 | -1.41879 |
| ENSMUSG00000028497 | -7.17635 | -6.1088 | -2.19442 | -2.19442 | -3.08556 | -1.27422 |
| ENSMUSG00000030847 | -5.84191 | -3.4399 | -1.18617 | -1.18617 | -3.08073 | -1.27099 |
| ENSMUSG00000061731 | -6.31638 | -3.02474 | -1.0379 | -1.0379 | -3.08156 | -1.27099 |
| ENSMUSG00000030559 | -5.99018 | -3.79576 | -1.66064 | -1.66064 | -3.06452 | -1.24269 |
| ENSMUSG00000021953 | -7.73979 | -6.52396 | -2.43166 | -2.43166 | -3.05286 | -1.22608 |
| ENSMUSG00000043154 | -5.60467 | -4.50746 | -1.60133 | -1.60133 | -3.03993 | -1.20686 |
| ENSMUSG00000027777 | -6.73154 | -5.42675 | -2.13511 | -2.13511 | -3.02912 | -1.19167 |
| ENSMUSG00000041757 | -7.38393 | -6.43499 | -1.30479 | -1.30479 | -3.02368 | -1.18396 |
| ENSMUSG00000007034 | -7.32462 | -6.43499 | -2.37235 | -2.37235 | -2.99384 | -1.1457 |
| ENSMUSG00000049922 | -6.76119 | -2.81716 | -1.0379 | -1.0379 | -2.97584 | -1.12735 |
| ENSMUSG00000017344 | -8.00667 | -7.26532 | -1.0379 | -1.0379 | -2.9533 | -1.09652 |
| ENSMUSG00000027978 | -7.59151 | -4.77435 | -1.74961 | -1.74961 | -2.93648 | -1.07466 |
| ENSMUSG00000000385 | -8.837 | -4.62608 | -1.15652 | -1.15652 | -2.89159 | -0.99872 |
| ENSMUSG00000035504 | -8.06598 | -7.26532 | -1.00825 | -1.00825 | -2.87817 | -0.99265 |
| ENSMUSG00000038224 | -8.24391 | -7.38393 | -1.95719 | -1.95719 | -2.883 | -0.99265 |
| ENSMUSG00000031714 | -7.32462 | -3.7661 | -1.30479 | -1.30479 | -2.86485 | -0.97412 |
| ENSMUSG00000031681 | -6.96877 | -5.90122 | -1.83857 | -1.83857 | -2.8445 | -0.94238 |

| | | | | | | |
|---|---|---|---|---|---|---|
| ENSMUSG00000020080 | -8.12529 | -3.70679 | -1.30479 | -1.30479 | -2.83622 | -0.93039 |
| ENSMUSG00000032083 | -8.74803 | -4.98193 | -1.74961 | -1.74961 | -2.8016 | -0.88385 |
| ENSMUSG00000034258 | -8.03633 | -7.11704 | -1.66064 | -1.66064 | -2.72794 | -0.78016 |
| ENSMUSG00000096054 | -6.37569 | -4.83366 | -1.33445 | -1.33445 | -2.72203 | -0.77246 |
| ENSMUSG00000034854 | -6.37569 | -5.30813 | -1.51237 | -1.51237 | -2.70934 | -0.76397 |
| ENSMUSG00000039878 | -7.35428 | -4.32954 | -1.92753 | -1.92753 | -2.71084 | -0.76397 |
| ENSMUSG00000022747 | -6.79085 | -5.93087 | -1.15652 | -1.15652 | -2.70329 | -0.75539 |
| ENSMUSG00000063652 | -5.30813 | -3.41025 | -1.15652 | -1.15652 | -2.69502 | -0.73929 |
| ENSMUSG00000001604 | -7.41359 | -6.55361 | -1.6903 | -1.6903 | -2.69105 | -0.7329 |
| ENSMUSG00000038267 | -6.4943 | -5.48606 | -1.51237 | -1.51237 | -2.67473 | -0.70077 |
| ENSMUSG00000050212 | -6.46465 | -5.51571 | -1.09721 | -1.09721 | -2.67126 | -0.69542 |
| ENSMUSG00000029130 | -6.58327 | -4.29988 | -1.09721 | -1.09721 | -2.66526 | -0.68684 |
| ENSMUSG00000049690 | -5.69364 | -4.21092 | -1.18617 | -1.18617 | -2.66097 | -0.68218 |
| ENSMUSG00000023067 | -7.44324 | -6.58327 | -1.71995 | -1.71995 | -2.63711 | -0.64852 |
| ENSMUSG00000027669 | -7.32462 | -6.31638 | -1.3641 | -1.3641 | -2.62884 | -0.63848 |
| ENSMUSG00000000253 | -6.52396 | -5.27847 | -1.21583 | -1.21583 | -2.59359 | -0.58298 |
| ENSMUSG00000054737 | -6.61292 | -5.39709 | -2.0165 | -2.0165 | -2.58349 | -0.56631 |
| ENSMUSG00000025790 | -7.02808 | -5.93087 | -1.42341 | -1.42341 | -2.57384 | -0.56353 |
| ENSMUSG00000037949 | -6.96877 | -5.96052 | -1.15652 | -1.15652 | -2.56883 | -0.55387 |
| ENSMUSG00000028357 | -7.82875 | -6.52396 | -1.3641 | -1.3641 | -2.55375 | -0.53341 |
| ENSMUSG00000019872 | -6.87981 | -5.96052 | -1.00825 | -1.00825 | -2.54488 | -0.52215 |
| ENSMUSG00000039670 | -5.93087 | -4.77435 | -1.15652 | -1.15652 | -2.54264 | -0.52215 |
| ENSMUSG00000031362 | -6.37569 | -5.30813 | -1.09721 | -1.09721 | -2.51682 | -0.491 |
| ENSMUSG00000029484 | -6.55361 | -5.54536 | -1.12687 | -1.12687 | -2.47994 | -0.44664 |
| ENSMUSG00000031788 | -6.99843 | -5.81225 | -1.09721 | -1.09721 | -2.48034 | -0.44664 |
| ENSMUSG00000021678 | -6.4943 | -5.42675 | -1.24548 | -1.24548 | -2.4622 | -0.4145 |
| ENSMUSG00000031398 | 1.275137 | 5.278475 | 6.138451 | 1.275137 | 2.426826 | 0.341695 |
| ENSMUSG00000047141 | 1.186174 | 4.922622 | 5.841907 | 1.186174 | 2.537085 | 0.513833 |
| ENSMUSG00000027238 | 1.275137 | 4.833659 | 5.841907 | 1.275137 | 2.563398 | 0.545709 |
| ENSMUSG00000075334 | 1.334446 | 6.523957 | 7.354279 | 1.334446 | 2.563666 | 0.545709 |
| ENSMUSG00000050973 | 1.008248 | 4.804005 | 5.634327 | 1.008248 | 2.577169 | 0.563533 |
| ENSMUSG00000001151 | 1.215828 | 4.952277 | 6.049488 | 1.215828 | 2.586436 | 0.570871 |
| ENSMUSG00000096528 | 1.15652 | 4.181264 | 5.456401 | 1.15652 | 2.616722 | 0.622806 |
| ENSMUSG00000020027 | 1.067557 | 5.93087 | 6.731538 | 1.067557 | 2.627732 | 0.638482 |
| ENSMUSG00000025507 | 1.215828 | 5.841907 | 6.553612 | 1.215828 | 2.637922 | 0.64852 |
| ENSMUSG00000027387 | 1.334446 | 5.456401 | 6.40534 | 1.334446 | 2.649066 | 0.665461 |
| ENSMUSG00000030284 | 1.097211 | 5.72329 | 6.494303 | 1.097211 | 2.678545 | 0.706897 |
| ENSMUSG00000051306 | 1.660644 | 5.515709 | 6.434994 | 1.660644 | 2.709945 | 0.763966 |
| ENSMUSG00000057228 | 1.393755 | 5.841907 | 6.553612 | 1.393755 | 2.721464 | 0.772458 |
| ENSMUSG00000025352 | 1.482718 | 4.804005 | 5.93087 | 1.482718 | 2.727288 | 0.780157 |
| ENSMUSG00000038593 | 1.215828 | 4.092301 | 5.070894 | 1.215828 | 2.733004 | 0.784863 |
| ENSMUSG00000049680 | 1.512372 | 4.952277 | 5.841907 | 1.512372 | 2.735194 | 0.785882 |
| ENSMUSG00000055884 | 1.275137 | 5.960525 | 6.790847 | 1.275137 | 2.746643 | 0.804435 |
| ENSMUSG00000018001 | 1.3641 | 4.774351 | 6.049488 | 1.3641 | 2.750266 | 0.809761 |
| ENSMUSG00000033762 | 1.304791 | 5.070894 | 5.901216 | 1.304791 | 2.756939 | 0.821818 |
| ENSMUSG00000030309 | 1.423409 | 4.744696 | 5.752944 | 1.423409 | 2.765408 | 0.832858 |
| ENSMUSG00000047342 | 1.719952 | 5.634327 | 6.346031 | 1.719952 | 2.795395 | 0.873173 |
| ENSMUSG00000041975 | 1.660644 | 4.181264 | 5.72329 | 1.660644 | 2.806342 | 0.883851 |
| ENSMUSG00000025408 | 1.215828 | 4.062646 | 5.693635 | 1.215828 | 2.83368 | 0.927771 |
| ENSMUSG00000001497 | 1.690298 | 6.079142 | 6.939118 | 1.690298 | 2.855158 | 0.956519 |
| ENSMUSG00000005718 | 2.224076 | 4.655733 | 5.871562 | 2.224076 | 2.883428 | 0.992645 |
| ENSMUSG00000039081 | 1.660644 | 4.062646 | 5.486055 | 1.660644 | 2.87773 | 0.992645 |
| ENSMUSG00000029641 | 1.097211 | 3.706794 | 6.523957 | 1.097211 | 2.899913 | 1.013148 |
| ENSMUSG00000055629 | 1.986842 | 5.159857 | 5.990179 | 1.986842 | 2.920192 | 1.049247 |
| ENSMUSG00000039910 | 1.037902 | 2.7282 | 7.117044 | 1.037902 | 2.965032 | 1.117258 |
| ENSMUSG00000019960 | 1.601335 | 6.553612 | 7.947366 | 1.601335 | 2.96986 | 1.118623 |
| ENSMUSG00000022367 | 2.016496 | 6.316377 | 7.354279 | 2.016496 | 2.983026 | 1.132965 |
| ENSMUSG00000030499 | 2.194422 | 5.219166 | 6.079142 | 2.194422 | 2.983954 | 1.132965 |

| | | | | | | |
|---|---|---|---|---|---|---|
| ENSMUSG00000044674 | 1.690298 | 5.04124 | 6.434994 | 1.690298 | 2.987909 | 1.136821 |
| ENSMUSG00000031393 | 2.105459 | 5.130203 | 6.049488 | 2.105459 | 3.011692 | 1.172253 |
| ENSMUSG00000051367 | 2.313039 | 6.108796 | 6.998427 | 2.313039 | 3.037665 | 1.205448 |
| ENSMUSG00000020642 | 1.423409 | 5.93087 | 6.761192 | 1.423409 | 3.060723 | 1.238362 |
| ENSMUSG00000048878 | 1.512372 | 4.655733 | 5.871562 | 1.512372 | 3.092468 | 1.281387 |
| ENSMUSG00000059921 | 2.550274 | 5.219166 | 6.61292 | 2.550274 | 3.151585 | 1.381066 |
| ENSMUSG00000040312 | 2.04615 | 4.507461 | 5.515709 | 2.04615 | 3.253425 | 1.510532 |
| ENSMUSG00000024427 | 2.402002 | 5.604672 | 6.553612 | 2.402002 | 3.285793 | 1.542332 |
| ENSMUSG00000017724 | 2.757855 | 4.952277 | 5.990179 | 2.757855 | 3.298361 | 1.561605 |
| ENSMUSG00000010051 | 1.304791 | 3.321287 | 5.159857 | 1.304791 | 3.324696 | 1.605194 |
| ENSMUSG00000013089 | 1.186174 | 3.054398 | 5.604672 | 1.186174 | 3.336951 | 1.623592 |
| ENSMUSG00000024420 | 2.342694 | 4.981931 | 5.990179 | 2.342694 | 3.342809 | 1.630669 |
| ENSMUSG00000021846 | 1.512372 | 3.439905 | 5.337783 | 1.512372 | 3.372222 | 1.672403 |
| ENSMUSG00000075271 | 2.52062 | 4.56677 | 5.93087 | 2.52062 | 3.397966 | 1.707288 |
| ENSMUSG00000018648 | 1.512372 | 4.596424 | 5.486055 | 1.512372 | 3.414445 | 1.729147 |
| ENSMUSG00000050447 | 2.372348 | 4.833659 | 5.901216 | 2.372348 | 3.418526 | 1.72957 |
| ENSMUSG00000050668 | 2.490966 | 5.278475 | 6.227414 | 2.490966 | 3.426035 | 1.735803 |
| ENSMUSG00000029211 | 1.393755 | 4.092301 | 5.960525 | 1.393755 | 3.446411 | 1.753754 |
| ENSMUSG00000040723 | 2.639237 | 5.397092 | 6.583266 | 2.639237 | 3.4802 | 1.794393 |
| ENSMUSG00000091337 | 1.275137 | 3.855066 | 5.871562 | 1.275137 | 3.490342 | 1.807724 |
| ENSMUSG00000020021 | 1.571681 | 4.418498 | 5.486055 | 1.571681 | 3.509058 | 1.827619 |
| ENSMUSG00000021118 | 1.571681 | 4.715042 | 5.693635 | 1.571681 | 3.508402 | 1.827619 |
| ENSMUSG00000020486 | 1.393755 | 3.973683 | 5.663981 | 1.393755 | 3.539714 | 1.858486 |
| ENSMUSG00000038550 | 1.630989 | 5.070894 | 6.019833 | 1.630989 | 3.545417 | 1.863895 |
| ENSMUSG00000042258 | 1.542026 | 3.469559 | 8.154947 | 1.542026 | 3.566824 | 1.885093 |
| ENSMUSG00000030347 | 2.253731 | 5.011585 | 6.346031 | 2.253731 | 3.573226 | 1.891963 |
| ENSMUSG00000036902 | 2.402002 | 5.219166 | 6.019833 | 2.402002 | 3.649326 | 1.990838 |
| ENSMUSG00000033174 | 1.690298 | 4.329535 | 5.782599 | 1.690298 | 3.664326 | 2.007464 |
| ENSMUSG00000002107 | 2.846818 | 5.04124 | 6.079142 | 2.846818 | 3.708602 | 2.055142 |
| ENSMUSG00000031478 | 2.668892 | 5.337783 | 6.257068 | 2.668892 | 3.709881 | 2.055142 |
| ENSMUSG00000028179 | 2.757855 | 4.863314 | 5.93087 | 2.757855 | 3.724372 | 2.068034 |
| ENSMUSG00000022022 | 3.173016 | 5.663981 | 6.494303 | 3.173016 | 3.770538 | 2.132857 |
| ENSMUSG00000054046 | 1.927533 | 4.892968 | 5.871562 | 1.927533 | 3.769767 | 2.132857 |
| ENSMUSG00000016262 | 2.639237 | 5.04124 | 6.138451 | 2.639237 | 3.802613 | 2.173243 |
| ENSMUSG00000031665 | 1.245483 | 2.639237 | 5.367438 | 1.245483 | 3.802216 | 2.173243 |
| ENSMUSG00000096847 | 3.143361 | 4.922622 | 6.672229 | 3.143361 | 3.811697 | 2.178073 |
| ENSMUSG00000054727 | 2.372348 | 5.337783 | 6.375686 | 2.372348 | 3.831654 | 2.205398 |
| ENSMUSG00000004113 | 2.846818 | 5.100548 | 6.079142 | 2.846818 | 3.855631 | 2.23439 |
| ENSMUSG00000039201 | 2.639237 | 5.456401 | 6.494303 | 2.639237 | 3.88124 | 2.260272 |
| ENSMUSG00000040289 | 3.469559 | 6.227414 | 7.057736 | 3.469559 | 3.877961 | 2.260272 |
| ENSMUSG00000071679 | 2.194422 | 4.981931 | 5.901216 | 2.194422 | 3.880067 | 2.260272 |
| ENSMUSG00000026482 | 2.579929 | 5.575018 | 6.375686 | 2.579929 | 3.900564 | 2.277229 |
| ENSMUSG00000040040 | 3.084053 | 5.130203 | 6.138451 | 3.084053 | 3.908894 | 2.280585 |
| ENSMUSG00000048216 | 2.372348 | 5.219166 | 6.257068 | 2.372348 | 3.954589 | 2.332251 |
| ENSMUSG00000020277 | 1.067557 | 2.194422 | 3.232324 | 1.067557 | 3.961364 | 2.3377 |
| ENSMUSG00000026360 | 2.876472 | 5.189512 | 6.019833 | 2.876472 | 3.978983 | 2.35097 |
| ENSMUSG00000038930 | 2.461311 | 4.685388 | 5.960525 | 2.461311 | 3.976558 | 2.35097 |
| ENSMUSG00000066392 | 2.609583 | 4.685388 | 6.583266 | 2.609583 | 4.00083 | 2.378126 |
| ENSMUSG00000038010 | 2.817163 | 5.011585 | 5.93087 | 2.817163 | 4.012982 | 2.386072 |
| ENSMUSG00000001986 | 3.291633 | 5.337783 | 6.257068 | 3.291633 | 4.027081 | 2.40242 |
| ENSMUSG00000070822 | 2.99509 | 5.426746 | 6.553612 | 2.99509 | 4.057809 | 2.438717 |
| ENSMUSG00000003955 | 1.3641 | 2.609583 | 4.151609 | 1.3641 | 4.099605 | 2.491563 |
| ENSMUSG00000041396 | 3.291633 | 5.337783 | 6.138451 | 3.291633 | 4.127828 | 2.516996 |
| ENSMUSG00000025161 | 1.097211 | 2.105459 | 3.084053 | 1.097211 | 4.169463 | 2.56545 |
| ENSMUSG00000002486 | 3.499213 | 5.24882 | 6.346031 | 3.499213 | 4.322241 | 2.765866 |
| ENSMUSG00000020363 | 1.897878 | 3.588177 | 7.56186 | 1.897878 | 4.388869 | 2.846542 |
| ENSMUSG00000044220 | 3.973683 | 8.303218 | 9.192849 | 3.973683 | 4.450574 | 2.907038 |
| ENSMUSG00000019789 | 3.855066 | 6.168105 | 6.909464 | 3.855066 | 4.48985 | 2.949667 |

| | | | | | | |
|---|---|---|---|---|---|---|
| ENSMUSG00000015579 | 3.825411 | 7.443242 | 8.125292 | 3.825411 | 4.569684 | 3.049901 |
| ENSMUSG00000032380 | 3.944029 | 6.257068 | 7.29497 | 3.944029 | 4.583049 | 3.053671 |
| ENSMUSG00000000127 | 4.151609 | 5.634327 | 6.40534 | 4.151609 | 4.596334 | 3.064748 |
| ENSMUSG00000023092 | 1.3641 | 2.372348 | 3.439905 | 1.3641 | 4.625654 | 3.098584 |
| ENSMUSG00000021848 | 4.329535 | 6.701883 | 7.472897 | 4.329535 | 4.661151 | 3.140922 |
| ENSMUSG00000039239 | 1.808915 | 3.439905 | 5.426746 | 1.808915 | 4.669721 | 3.143083 |
| ENSMUSG00000026208 | 4.35919 | 6.40534 | 7.176353 | 4.35919 | 4.730344 | 3.174565 |

#### and FG-2

ound of the 95
ne min(abs(c(lb, ub))), or 0 if the CI crosses the 0

sis testing using Holm procedure
**of ce, Z and cZ.**

| GeneName | Description | Location | Family |
|---|---|---|---|
| Car4 | carbonic anhydrase 4 | Plasma Membrane | enzyme |
| Ttr | transthyretin | Extracellular Space | transporter |
| Spon2 | spondin 2 | Extracellular Space | other |
| Sfrp5 | secreted frizzled relatec | Plasma Membrane | transmembrane receptor |
| Fgb | fibrinogen beta chain | Extracellular Space | other |
| Rfx6 | regulatory factor X6 | Nucleus | enzyme |
| Habp2 | hyaluronan binding prc | Extracellular Space | peptidase |
| Enpp1 | ectonucleotide pyropho | Plasma Membrane | enzyme |
| Pdpn | podoplanin | Plasma Membrane | other |
| Tfcp2l1 | transcription factor CP2 | Nucleus | transcription regulator |
| Afp | alpha fetoprotein | Extracellular Space | transporter |
| Csf1r | colony stimulating facto | Plasma Membrane | kinase |
| Kalrn | kalirin RhoGEF kinase | Cytoplasm | kinase |
| Enpep | glutamyl aminopeptida | Plasma Membrane | peptidase |
| Gata5 | GATA binding protein 5 | Nucleus | transcription regulator |
| Epas1 | endothelial PAS domair | Nucleus | transcription regulator |
| Spp2 | secreted phosphoprote | Extracellular Space | other |
| Cdh6 | cadherin 6 | Plasma Membrane | other |
| S100a16 | S100 calcium binding p | Nucleus | other |
| Vil1 | villin 1 | Cytoplasm | other |
| Fam183b | family with sequence si | Extracellular Space | other |
| Gata4 | GATA binding protein 4 | Nucleus | transcription regulator |
| Plg | plasminogen | Extracellular Space | peptidase |
| Proc | protein C, inactivator of | Extracellular Space | peptidase |
| Myo6 | myosin VI | Cytoplasm | other |
| Apob | apolipoprotein B | Extracellular Space | transporter |
| 2210011C24Rik | RIKEN cDNA 2210011C | Other | other |
| Spink3 | serine peptidase inhibit | Extracellular Space | other |
| Igsf11 | immunoglobulin superf | Plasma Membrane | other |
| Lpar3 | lysophosphatidic acid r | Plasma Membrane | G-protein coupled receptor |
| Onecut1 | one cut homeobox 1 | Nucleus | transcription regulator |
| Rnase4 | ribonuclease A family m | Extracellular Space | enzyme |
| Isx | intestine specific home | Nucleus | transcription regulator |
| Emp1 | epithelial membrane pr | Plasma Membrane | other |
| Camkk1 | calcium/calmodulin dep | Cytoplasm | kinase |
| Apom | apolipoprotein M | Plasma Membrane | transporter |
| Ptk2b | protein tyrosine kinase | Cytoplasm | kinase |
| Serpina10 | serpin family A membe | Extracellular Space | other |
| Vsig2 | V-set and immunoglob | Plasma Membrane | other |
| Zc2hc1c | zinc finger C2HC-type ( | Other | other |
| Lrpap1 | LDL receptor related pr | Plasma Membrane | other |
| Pla1a | phospholipase A1 mem | Extracellular Space | enzyme |
| Rbp4 | retinol binding protein | Extracellular Space | other |
| Kank4 | KN motif and ankyrin re | Cytoplasm | other |
| H2-Q2 | major histocompatibilit | Plasma Membrane | other |
| Cd1d1 | CD1d molecule | Plasma Membrane | other |
| Serpina1a | serpin family A membe | Extracellular Space | other |

| | | | |
|---|---|---|---|
| Adam19 | ADAM metallopeptidas | Plasma Membrane | peptidase |
| Dab2 | DAB2, clathrin adaptor | Plasma Membrane | other |
| Daam1 | dishevelled associated a | Cytoplasm | other |
| Vnn1 | vanin 1 | Plasma Membrane | enzyme |
| F2 | coagulation factor II, th | Extracellular Space | peptidase |
| Igfbp4 | insulin like growth facto | Extracellular Space | other |
| Cd55 | CD55 molecule (Crome | Plasma Membrane | other |
| Suox | sulfite oxidase | Cytoplasm | enzyme |
| Ahnak | AHNAK nucleoprotein | Nucleus | other |
| Hacd2 | 3-hydroxyacyl-CoA del | Cytoplasm | phosphatase |
| Fzd7 | frizzled class receptor 7 | Plasma Membrane | G-protein coupled receptor |
| Selenbp1 | selenium binding prote | Cytoplasm | other |
| Nid2 | nidogen 2 | Extracellular Space | other |
| Tpk1 | thiamin pyrophosphoki | Cytoplasm | kinase |
| Lipa | lipase A, lysosomal acid | Cytoplasm | enzyme |
| Rnf6 | ring finger protein 6 | Nucleus | transcription regulator |
| Mttp | microsomal triglyceride | Cytoplasm | transporter |
| Cubn | cubilin | Plasma Membrane | transmembrane receptor |
| Stx7 | syntaxin 7 | Plasma Membrane | transporter |
| 1700011H14Rik | coiled-coil domain con | Other | other |
| Fem1a | fem-1 homolog A | Nucleus | transcription regulator |
| Ihh | indian hedgehog | Extracellular Space | enzyme |
| Pitpna | phosphatidylinositol tra | Cytoplasm | transporter |
| Tor2a | torsin family 2 member | Extracellular Space | other |
| Fgfr2 | fibroblast growth factor | Plasma Membrane | kinase |
| Tfpi | tissue factor pathway in | Extracellular Space | other |
| Erap1 | endoplasmic reticulum | Extracellular Space | peptidase |
| Hnf1b | HNF1 homeobox B | Nucleus | transcription regulator |
| Lima1 | LIM domain and actin b | Cytoplasm | other |
| Stard8 | StAR related lipid transt | Cytoplasm | other |
| Lpar1 | lysophosphatidic acid r | Plasma Membrane | G-protein coupled receptor |
| Stom | stomatin | Plasma Membrane | other |
| Kynu | kynureninase | Cytoplasm | enzyme |
| Hbegf | heparin binding EGF lik | Extracellular Space | growth factor |
| Slc39a4 | solute carrier family 39 | Plasma Membrane | transporter |
| Epha4 | EPH receptor A4 | Plasma Membrane | kinase |
| Wwp1 | WW domain containing | Cytoplasm | enzyme |
| Lurap1l | leucine rich adaptor pro | Other | other |
| B4galnt2 | beta-1,4-N-acetyl-gala | Cytoplasm | enzyme |
| Daf2 | CD55 molecule (Crome | Plasma Membrane | other |
| Thbs1 | thrombospondin 1 | Extracellular Space | other |
| Hacd4 | 3-hydroxyacyl-CoA del | Cytoplasm | enzyme |
| Bag3 | BCL2 associated athano | Cytoplasm | other |
| Ext1 | exostosin glycosyltranst | Cytoplasm | enzyme |
| Rab38 | RAB38, member RAS or | Cytoplasm | enzyme |
| Tdh | L-threonine dehydroge | Cytoplasm | enzyme |
| Ppp2r3a | protein phosphatase 2 | Nucleus | phosphatase |
| Schip1 | schwannomin interactin | Cytoplasm | other |
| Plekha6 | pleckstrin homology do | Other | other |
| Slc44a4 | solute carrier family 44 | Plasma Membrane | transporter |
| Slc35c1 | solute carrier family 35 | Cytoplasm | transporter |
| Vtn | vitronectin | Extracellular Space | other |
| Prss12 | serine protease 12 | Extracellular Space | peptidase |
| Tmprss2 | transmembrane serine | Plasma Membrane | peptidase |
| Reep6 | receptor accessory prot | Plasma Membrane | other |
| Serpinf2 | serpin family F member | Extracellular Space | other |
| Gab1 | GRB2 associated bindin | Cytoplasm | kinase |
| Smad1 | SMAD family member 1 | Nucleus | transcription regulator |

| | | | |
|---|---|---|---|
| Hkdc1 | hexokinase domain cor | Cytoplasm | kinase |
| Apoa1 | apolipoprotein A1 | Extracellular Space | transporter |
| Mfsd7c | feline leukemia virus su | Plasma Membrane | transporter |
| Syne1 | spectrin repeat containi | Nucleus | other |
| Mfsd12 | major facilitator superfa | Cytoplasm | other |
| Slc39a5 | solute carrier family 39 | Plasma Membrane | transporter |
| St3gal6 | ST3 beta-galactoside al | Cytoplasm | enzyme |
| Slc22a21 | solute carrier family 22 | Cytoplasm | transporter |
| Tcea3 | transcription elongatior | Nucleus | transcription regulator |
| Slc22a23 | solute carrier family 22 | Other | transporter |
| Eva1b | eva-1 homolog B | Other | other |
| Rnf32 | ring finger protein 32 | Cytoplasm | other |
| Nckap5 | NCK associated protein | Other | other |
| Cdkn1a | cyclin dependent kinase | Nucleus | kinase |
| Gnb4 | G protein subunit beta | Plasma Membrane | enzyme |
| Gmpr | guanosine monophosp | Cytoplasm | enzyme |
| Zfp182 | zinc finger protein 182 | Nucleus | transcription regulator |
| Slco3a1 | solute carrier organic ar | Plasma Membrane | transporter |
| Ano10 | anoctamin 10 | Plasma Membrane | ion channel |
| Kif12 | kinesin family member | Cytoplasm | other |
| Smpdl3a | sphingomyelin phosph | Extracellular Space | enzyme |
| Oxld1 | oxidoreductase like dor | Cytoplasm | other |
| Xlr4c | X-linked lymphocyte-re | Other | other |
| Anxa3 | annexin A3 | Cytoplasm | enzyme |
| Kifc3 | kinesin family member | Cytoplasm | enzyme |
| F2rl1 | F2R like trypsin recepto | Plasma Membrane | G-protein coupled receptor |
| Plxna3 | plexin A3 | Plasma Membrane | transmembrane receptor |
| Zfp654 | zinc finger protein 654 | Other | other |
| Frmd5 | FERM domain containir | Plasma Membrane | other |
| Rprm | reprimo, TP53 depende | Cytoplasm | other |
| Gdpgp1 | GDP-D-glucose phospl | Cytoplasm | enzyme |
| Pcnt | pericentrin (kendrin) | Cytoplasm | other |
| G430049J08Rik | - | Other | other |
| Socs2 | suppressor of cytokine | Cytoplasm | other |
| Pidd1 | p53-induced death dor | Cytoplasm | other |
| Zc3h8 | zinc finger CCCH-type | Nucleus | transcription regulator |
| Creld1 | cysteine rich with EGF li | Other | other |
| Usp42 | ubiquitin specific peptic | Nucleus | peptidase |
| Aadat | aminoadipate aminotra | Cytoplasm | enzyme |
| Gdf11 | growth differentiation f | Extracellular Space | growth factor |
| Tctn1 | tectonic family member | Extracellular Space | other |
| Urgcp | upregulator of cell prol | Cytoplasm | other |
| Fancm | FA complementation gr | Nucleus | enzyme |
| Cyth3 | cytohesin 3 | Cytoplasm | other |
| Recql4 | RecQ like helicase 4 | Nucleus | enzyme |
| Caprin2 | caprin family member 2 | Cytoplasm | other |
| Zfp286 | zinc finger protein 286/ | Nucleus | transcription regulator |
| Mettl8 | methyltransferase like 8 | Other | enzyme |
| Ddit3 | DNA damage inducible | Nucleus | transcription regulator |
| Pax9 | paired box 9 | Nucleus | transcription regulator |
| Tfap4 | transcription factor AP- | Nucleus | transcription regulator |
| Zfp503 | zinc finger protein 503 | Nucleus | other |
| Rasl11a | RAS like family 11 mem | Nucleus | other |
| B4galnt4 | beta-1,4-N-acetyl-gala | Extracellular Space | enzyme |
| Cited2 | Cbp/p300 interacting tr | Nucleus | transcription regulator |
| Dusp6 | dual specificity phospha | Cytoplasm | phosphatase |
| Has2 | hyaluronan synthase 2 | Plasma Membrane | enzyme |
| Kctd15 | potassium channel tetra | Other | ion channel |

| | | | |
|---|---|---|---|
| Fzd1 | frizzled class receptor 1 | Plasma Membrane | G-protein coupled receptor |
| Mecp2 | methyl-CpG binding pr | Nucleus | transcription regulator |
| Six1 | SIX homeobox 1 | Nucleus | transcription regulator |
| Rnf144a | ring finger protein 144/ | Nucleus | enzyme |
| Hexim1 | hexamethylene bisaceta | Nucleus | transcription regulator |
| Unc5c | unc-5 netrin receptor C | Plasma Membrane | transmembrane receptor |
| Cchcr1 | coiled-coil alpha-helica | Cytoplasm | other |
| Spry4 | sprouty RTK signaling a | Plasma Membrane | other |
| Etv4 | ETS variant 4 | Nucleus | transcription regulator |
| Hyal1 | hyaluronoglucosaminid | Cytoplasm | enzyme |
| Etv5 | ETS variant 5 | Nucleus | transcription regulator |
| Zfp521 | zinc finger protein 521 | Nucleus | transcription regulator |
| Peli2 | pellino E3 ubiquitin pro | Cytoplasm | kinase |
| Ttc30a1 | tetratricopeptide repea | Extracellular Space | other |
| Dusp14 | dual specificity phospha | Other | phosphatase |
| Lypd6 | LY6/PLAUR domain cor | Extracellular Space | other |
| Gpatch11 | G-patch domain contai | Nucleus | other |
| Gabra4 | gamma-aminobutyric a | Plasma Membrane | ion channel |
| Rcsd1 | RCSD domain containir | Other | other |
| Eid1 | EP300 interacting inhib | Nucleus | transcription regulator |
| Fgd6 | FYVE, RhoGEF and PH c | Cytoplasm | other |
| Plek2 | pleckstrin 2 | Plasma Membrane | other |
| 4-Sep | septin 4 | Cytoplasm | enzyme |
| Ciart | circadian associated rep | Nucleus | other |
| Isl1 | ISL LIM homeobox 1 | Nucleus | transcription regulator |
| D6Wsu163e | chromosome 12 open r | Cytoplasm | other |
| Neto2 | neuropilin and tolloid li | Plasma Membrane | other |
| Mgll | monoglyceride lipase | Plasma Membrane | enzyme |
| Celf2 | CUGBP Elav-like family | Nucleus | other |
| Nek3 | NIMA related kinase 3 | Nucleus | kinase |
| Cth | cystathionine gamma-l | Cytoplasm | enzyme |
| Mtrf1 | mitochondrial translatic | Cytoplasm | translation regulator |
| Klk13 | kallikrein related peptid | Extracellular Space | peptidase |
| Sertad4 | SERTA domain containi | Other | other |
| Sall1 | spalt like transcription f | Nucleus | transcription regulator |
| Tmem151b | transmembrane proteir | Other | other |
| 1700013H16Rik | RIKEN cDNA 1700013H | Nucleus | other |
| Cacna1b | calcium voltage-gated | Plasma Membrane | ion channel |
| Tbc1d25 | TBC1 domain family me | Other | other |
| Hey1 | hes related family bHLH | Nucleus | transcription regulator |
| Zcchc16 | retrotransposon Gag lik | Other | other |
| Rgl1 | ral guanine nucleotide | Cytoplasm | other |
| Ift88 | intraflagellar transport 8 | Cytoplasm | other |
| Gpr85 | G protein-coupled rece | Plasma Membrane | G-protein coupled receptor |
| Pfkl | phosphofructokinase, li | Cytoplasm | kinase |
| Rgs2 | regulator of G protein s | Nucleus | other |
| Rccd1 | RCC1 domain containir | Other | other |
| Nrxn3 | neurexin III | Plasma Membrane | other |
| Ccdc138 | coiled-coil domain con | Other | other |
| Gria3 | glutamate ionotropic re | Plasma Membrane | ion channel |
| Zscan18 | zinc finger and SCAN d | Other | other |
| Fam162a | family with sequence si | Cytoplasm | other |
| Mettl18 | methyltransferase like 1 | Other | other |
| Slc16a3 | solute carrier family 16 | Plasma Membrane | transporter |
| Tchp | trichoplein keratin filam | Cytoplasm | other |
| Gfpt2 | glutamine-fructose-6-p | Cytoplasm | enzyme |
| Nkx2-3 | NK2 homeobox 3 | Nucleus | transcription regulator |
| Hey2 | hes related family bHLH | Nucleus | transcription regulator |

| | | | |
|---|---|---|---|
| Nkx2-5 | NK2 homeobox 5 | Nucleus | transcription regulator |
| Dapk2 | death associated protei | Cytoplasm | kinase |
| Fer | FER tyrosine kinase | Cytoplasm | kinase |
| Fhl1 | four and a half LIM dor | Cytoplasm | other |
| Otx2 | orthodenticle homeobc | Nucleus | transcription regulator |
| Tgfb2 | transforming growth fa | Extracellular Space | growth factor |
| Des | desmin | Cytoplasm | other |

### Differentially expressed genes (DEGs) between FG-1 and PS

- **lb**:lower bound, **mle**:maximum likelihood estimate, **ub**: upper bound of the
- **ce:** conservative estimate of expression-fold change (equals to the min(abs(
- **Z:** uncorrected Z-score of expression difference
- **cZ:** expression difference Z-score corrected for multiple hypothesis testing
- **Genes highly expressed in the FG-1 presented positive value of ce, Z an**

| | lb | mle | ub | ce | Z | cZ | GeneNam |
|---|---|---|---|---|---|---|---|
| ENSMUSG00000001435 | 2.746306 | 4.016104 | 5.551673 | 2.746306 | 7.160728 | 6.051966 | Col18a1 |
| ENSMUSG00000003153 | 1.476509 | 2.303354 | 3.514091 | 1.476509 | 7.160813 | 6.051966 | Slc2a3 |
| ENSMUSG00000020176 | 2.185233 | 2.982548 | 3.750332 | 2.185233 | 7.160813 | 6.051966 | Grb10 |
| ENSMUSG00000020644 | 3.071138 | 4.459056 | 6.260397 | 3.071138 | 7.134886 | 6.051966 | Id2 |
| ENSMUSG00000023092 | 4.163755 | 5.876505 | 8.150328 | 4.163755 | 7.160813 | 6.051966 | Fhl1 |
| ENSMUSG00000026185 | 8.120798 | 9.685897 | 10.36509 | 8.120798 | 7.160813 | 6.051966 | Igfbp5 |
| ENSMUSG00000042258 | 6.76241 | 7.825496 | 8.622811 | 6.76241 | 7.160735 | 6.051966 | Isl1 |
| ENSMUSG00000042712 | 2.008052 | 2.716776 | 3.39597 | 2.008052 | 7.160813 | 6.051966 | Wbp5 |
| ENSMUSG00000046432 | 1.71275 | 2.332884 | 2.864427 | 1.71275 | 7.160625 | 6.051966 | Ngfrap1 |
| ENSMUSG00000048583 | 3.868453 | 5.256371 | 7.618785 | 3.868453 | 7.141622 | 6.051966 | Igf2 |
| ENSMUSG00000055653 | 2.332884 | 2.893957 | 3.750332 | 2.332884 | 7.160813 | 6.051966 | Gpc3 |
| ENSMUSG00000078566 | 2.480535 | 3.573151 | 5.404022 | 2.480535 | 7.160377 | 6.051966 | Bnip3 |
| ENSMUSG00000047501 | 6.083216 | 7.441604 | 8.38657 | 6.083216 | 7.113951 | 6.051334 | Cldn4 |
| ENSMUSG00000022816 | 2.185233 | 3.159729 | 4.783888 | 2.185233 | 6.859967 | 5.779219 | Fstl1 |
| ENSMUSG00000089901 | 2.451004 | 4.193285 | 5.817444 | 2.451004 | 6.857738 | 5.779219 | Gm8113 |
| ENSMUSG00000018604 | 5.906035 | 7.707376 | 8.50469 | 5.906035 | 6.764238 | 5.693918 | Tbx3 |
| ENSMUSG00000025608 | 1.65369 | 2.598655 | 4.163755 | 1.65369 | 6.378078 | 5.264493 | Podxl |
| ENSMUSG00000048376 | 1.77181 | 2.628186 | 4.222815 | 1.77181 | 6.361926 | 5.252065 | F2r |
| ENSMUSG00000026193 | 2.155703 | 3.39597 | 4.577177 | 2.155703 | 6.344627 | 5.244294 | Fn1 |
| ENSMUSG00000017311 | 5.758384 | 7.530194 | 8.859052 | 5.758384 | 6.264013 | 5.157265 | Pyy |
| ENSMUSG00000029086 | 2.923487 | 4.075164 | 6.880531 | 2.923487 | 6.261569 | 5.157265 | Prom1 |
| ENSMUSG00000036030 | 1.594629 | 2.539595 | 3.36644 | 1.594629 | 6.157264 | 5.043485 | Prtg |
| ENSMUSG00000017491 | 4.902009 | 6.230867 | 7.116772 | 4.902009 | 6.15141 | 5.042132 | Rarb |
| ENSMUSG00000068876 | 2.214763 | 3.248319 | 4.872479 | 2.214763 | 6.059006 | 4.935952 | Cgn |
| ENSMUSG00000027560 | 4.990599 | 6.230867 | 7.087242 | 4.990599 | 6.031955 | 4.908636 | Dok5 |
| ENSMUSG00000032320 | 1.476509 | 2.332884 | 3.484561 | 1.476509 | 5.961308 | 4.828101 | Rcn2 |
| ENSMUSG00000027239 | 1.151677 | 1.77181 | 2.391944 | 1.151677 | 5.931026 | 4.80169 | Mdk |
| ENSMUSG00000018593 | 1.948992 | 3.189259 | 4.488587 | 1.948992 | 5.803398 | 4.660554 | Sparc |
| ENSMUSG00000021994 | 4.399996 | 7.057712 | 8.209388 | 4.399996 | 5.761545 | 4.61396 | Wnt5a |
| ENSMUSG00000031351 | 4.518117 | 6.024156 | 6.73288 | 4.518117 | 5.749331 | 4.603706 | Zfp185 |
| ENSMUSG00000026276 | 1.446979 | 2.273823 | 3.159729 | 1.446979 | 5.71955 | 4.571709 | Sept2 |
| ENSMUSG00000039239 | 2.834897 | 4.902009 | 7.648315 | 2.834897 | 5.70102 | 4.553469 | Tgfb2 |
| ENSMUSG00000015579 | 4.075164 | 7.323483 | 8.091268 | 4.075164 | 5.68263 | 4.539747 | Nkx2-5 |
| ENSMUSG00000057193 | 2.332884 | 4.104694 | 7.116772 | 2.332884 | 5.649673 | 4.511916 | Slc44a2 |
| ENSMUSG00000028645 | 1.387918 | 2.185233 | 2.923487 | 1.387918 | 5.624729 | 4.493296 | Slc2a1 |
| ENSMUSG00000044186 | 4.045634 | 5.581203 | 6.585229 | 4.045634 | 5.595946 | 4.465554 | Nkx2-6 |
| ENSMUSG00000032366 | 1.358388 | 2.480535 | 3.33691 | 1.358388 | 5.592528 | 4.465127 | Tpm1 |
| ENSMUSG00000022887 | 4.075164 | 5.522143 | 6.614759 | 4.075164 | 5.546808 | 4.416085 | Masp1 |
| ENSMUSG00000060572 | 1.978522 | 3.218789 | 5.433552 | 1.978522 | 5.53026 | 4.402845 | Mfap2 |
| ENSMUSG00000019789 | 3.809393 | 6.112746 | 6.939591 | 3.809393 | 5.485955 | 4.365213 | Hey2 |
| ENSMUSG00000044220 | 4.370466 | 8.297979 | 9.242945 | 4.370466 | 5.453982 | 4.32893 | Nkx2-3 |
| ENSMUSG00000017466 | 3.720802 | 5.610733 | 6.319457 | 3.720802 | 5.450479 | 4.327853 | Timp2 |
| ENSMUSG00000034112 | 3.809393 | 6.201337 | 7.087242 | 3.809393 | 5.417212 | 4.296163 | Atp2c2 |
| ENSMUSG00000038463 | 3.602681 | 5.787914 | 6.555699 | 3.602681 | 5.370635 | 4.244518 | Olfml2b |
| ENSMUSG00000027954 | 2.391944 | 3.779862 | 7.914087 | 2.391944 | 5.319501 | 4.183865 | Efna1 |
| ENSMUSG00000057615 | 3.661742 | 5.935565 | 6.73288 | 3.661742 | 5.303011 | 4.169339 | Ldoc1 |

| | | | | | | | |
|---|---|---|---|---|---|---|---|
| ENSMUSG00000036275 | 1.446979 | 2.303354 | 3.277849 | 1.446979 | 5.294292 | 4.164376 | 9530068E( |
| ENSMUSG00000017057 | 3.33691 | 5.787914 | 6.70335 | 3.33691 | 5.254177 | 4.12567 | Il13ra1 |
| ENSMUSG00000028943 | 4.075164 | 5.640263 | 6.378518 | 4.075164 | 5.256478 | 4.12567 | Espn |
| ENSMUSG00000029669 | 2.510065 | 3.897983 | 5.610733 | 2.510065 | 5.24975 | 4.122974 | Tspan12 |
| ENSMUSG00000025810 | 4.075164 | 7.618785 | 8.740932 | 4.075164 | 5.221278 | 4.090116 | Nrp1 |
| ENSMUSG00000020183 | 2.628186 | 4.695298 | 6.230867 | 2.628186 | 5.21529 | 4.085361 | Cpm |
| ENSMUSG00000020032 | 3.957043 | 5.522143 | 6.70335 | 3.957043 | 5.201412 | 4.070818 | Nuak1 |
| ENSMUSG00000055629 | 3.130198 | 5.404022 | 6.260397 | 3.130198 | 5.199365 | 4.070818 | B4galnt4 |
| ENSMUSG00000057530 | 2.214763 | 3.750332 | 5.817444 | 2.214763 | 5.188831 | 4.060281 | Ece1 |
| ENSMUSG00000031075 | 3.130198 | 5.463082 | 6.526169 | 3.130198 | 5.149033 | 4.012957 | Ano1 |
| ENSMUSG00000018906 | 2.569125 | 6.348988 | 7.175832 | 2.569125 | 5.121428 | 3.988462 | P4ha2 |
| ENSMUSG00000020911 | 1.978522 | 4.252345 | 6.496638 | 1.978522 | 5.121163 | 3.988462 | Krt19 |
| ENSMUSG00000022048 | 1.240267 | 2.126173 | 3.071138 | 1.240267 | 5.122551 | 3.988462 | Dpysl2 |
| ENSMUSG00000022426 | 3.36644 | 5.463082 | 6.408048 | 3.36644 | 5.119279 | 3.988462 | Josd1 |
| ENSMUSG00000039601 | 3.36644 | 5.758384 | 6.555699 | 3.36644 | 5.126554 | 3.988462 | Rcan2 |
| ENSMUSG00000028456 | 2.953017 | 5.374492 | 6.171806 | 2.953017 | 5.106524 | 3.977036 | Unc13b |
| ENSMUSG00000020160 | 3.927513 | 5.285901 | 6.644289 | 3.927513 | 5.098481 | 3.969762 | Meis1 |
| ENSMUSG00000038319 | 2.953017 | 5.256371 | 6.378518 | 2.953017 | 5.096159 | 3.969285 | Kcnh2 |
| ENSMUSG00000035967 | 3.543621 | 4.547647 | 6.76241 | 3.543621 | 5.080641 | 3.952175 | Ddx26b |
| ENSMUSG00000063632 | 1.299328 | 2.155703 | 2.923487 | 1.299328 | 5.07304 | 3.947391 | Sox11 |
| ENSMUSG00000001552 | 1.71275 | 2.982548 | 4.990599 | 1.71275 | 5.06741 | 3.942663 | Jup |
| ENSMUSG00000027820 | 3.720802 | 5.344962 | 6.348988 | 3.720802 | 5.042516 | 3.915965 | Mme |
| ENSMUSG00000070473 | 3.36644 | 5.167781 | 6.171806 | 3.36644 | 5.035834 | 3.912137 | Cldn3 |
| ENSMUSG00000030654 | 1.004026 | 1.74228 | 2.539595 | 1.004026 | 5.031436 | 3.908861 | Arl6ip1 |
| ENSMUSG00000026305 | 1.68322 | 2.893957 | 4.547647 | 1.68322 | 5.019379 | 3.895909 | Lrrfip1 |
| ENSMUSG00000036902 | 2.628186 | 5.404022 | 6.171806 | 2.628186 | 5.010485 | 3.886922 | Neto2 |
| ENSMUSG00000002980 | 1.74228 | 3.39597 | 4.724828 | 1.74228 | 4.998739 | 3.878702 | Bcam |
| ENSMUSG00000052516 | 2.657716 | 5.581203 | 6.408048 | 2.657716 | 4.986511 | 3.865435 | Robo2 |
| ENSMUSG00000022797 | 1.358388 | 2.391944 | 3.36644 | 1.358388 | 4.932656 | 3.803734 | Tfrc |
| ENSMUSG00000040652 | 1.387918 | 2.362414 | 3.33691 | 1.387918 | 4.928789 | 3.800957 | Oaz2 |
| ENSMUSG00000046822 | 2.923487 | 5.02013 | 5.876505 | 2.923487 | 4.857419 | 3.720683 | Slc39a3 |
| ENSMUSG00000036853 | 3.159729 | 5.04966 | 6.142276 | 3.159729 | 4.854394 | 3.71889 | Mcoln3 |
| ENSMUSG00000028527 | 1.535569 | 2.657716 | 4.547647 | 1.535569 | 4.849846 | 3.715138 | Ak4 |
| ENSMUSG00000031616 | 2.539595 | 5.226841 | 6.260397 | 2.539595 | 4.838243 | 3.706399 | Ednra |
| ENSMUSG00000034059 | 2.303354 | 4.488587 | 5.758384 | 2.303354 | 4.823766 | 3.689928 | Ypel4 |
| ENSMUSG00000002265 | 1.063086 | 1.919461 | 2.657716 | 1.063086 | 4.81866 | 3.687359 | Peg3 |
| ENSMUSG00000047986 | 3.838923 | 5.610733 | 6.348988 | 3.838923 | 4.812515 | 3.68147 | Palm3 |
| ENSMUSG00000048728 | 2.008052 | 4.961069 | 5.787914 | 2.008052 | 4.762988 | 3.625072 | Zfp454 |
| ENSMUSG00000032060 | 3.100668 | 5.404022 | 6.319457 | 3.100668 | 4.759664 | 3.623524 | Cryab |
| ENSMUSG00000032172 | 2.362414 | 4.931539 | 6.053686 | 2.362414 | 4.74974 | 3.614589 | Olfm2 |
| ENSMUSG00000039405 | 2.893957 | 4.783888 | 5.994625 | 2.893957 | 4.7413 | 3.605646 | Prss23 |
| ENSMUSG00000035775 | 1.889931 | 4.399996 | 5.787914 | 1.889931 | 4.733956 | 3.599944 | Krt20 |
| ENSMUSG00000018476 | 1.151677 | 2.008052 | 2.953017 | 1.151677 | 4.728848 | 3.595235 | Kdm6b |
| ENSMUSG00000041782 | 4.547647 | 6.70335 | 7.648315 | 4.547647 | 4.717187 | 3.582108 | Lad1 |
| ENSMUSG00000033860 | 2.864427 | 5.226841 | 6.024156 | 2.864427 | 4.708406 | 3.57447 | Fgg |
| ENSMUSG00000066392 | 2.746306 | 4.488587 | 6.348988 | 2.746306 | 4.70219 | 3.568291 | Nrxn3 |
| ENSMUSG00000015619 | 2.539595 | 6.998651 | 8.032207 | 2.539595 | 4.695054 | 3.560917 | Gata3 |
| ENSMUSG00000035835 | 2.037582 | 4.045634 | 6.76241 | 2.037582 | 4.678341 | 3.541219 | Lppr3 |
| ENSMUSG00000070436 | 1.063086 | 1.860401 | 2.628186 | 1.063086 | 4.676738 | 3.540934 | Serpinh1 |
| ENSMUSG00000032374 | 1.417448 | 2.539595 | 4.311405 | 1.417448 | 4.672713 | 3.537523 | Plod2 |
| ENSMUSG00000034687 | 3.36644 | 5.02013 | 5.994625 | 3.36644 | 4.657649 | 3.519893 | Fras1 |
| ENSMUSG00000066687 | 2.864427 | 5.07919 | 6.083216 | 2.864427 | 4.61355 | 3.469964 | Zbtb16 |
| ENSMUSG00000074925 | 2.569125 | 4.783888 | 5.640263 | 2.569125 | 4.602889 | 3.45854 | Ptar1 |
| ENSMUSG00000033487 | 1.71275 | 3.218789 | 5.640263 | 1.71275 | 4.595362 | 3.4533 | Fndc3a |
| ENSMUSG00000026343 | 1.71275 | 3.927513 | 5.876505 | 1.71275 | 4.576262 | 3.433631 | Gpr39 |
| ENSMUSG00000028082 | 3.33691 | 5.315431 | 6.348988 | 3.33691 | 4.56809 | 3.426375 | Sh3d19 |
| ENSMUSG00000010080 | 2.539595 | 4.724828 | 5.906035 | 2.539595 | 4.513232 | 3.363353 | Epn3 |
| ENSMUSG00000026600 | 1.181207 | 2.214763 | 3.33691 | 1.181207 | 4.509598 | 3.36025 | Soat1 |

| | | | | | | | |
|---|---|---|---|---|---|---|---|
| ENSMUSG00000039234 | 3.957043 | 6.939591 | 7.707376 | 3.957043 | 4.49527 | 3.346428 | Sec24d |
| ENSMUSG00000036968 | 1.210737 | 2.244293 | 4.104694 | 1.210737 | 4.488713 | 3.339482 | Cnpy4 |
| ENSMUSG00000071679 | 2.273823 | 4.990599 | 5.876505 | 2.273823 | 4.483011 | 3.335224 | Zcchc16 |
| ENSMUSG00000004885 | 2.008052 | 3.720802 | 6.024156 | 2.008052 | 4.449531 | 3.294659 | Crabp2 |
| ENSMUSG00000021413 | 1.092616 | 2.037582 | 3.33691 | 1.092616 | 4.447014 | 3.294506 | Prpf4b |
| ENSMUSG00000027315 | 1.328858 | 2.657716 | 4.281875 | 1.328858 | 4.442156 | 3.291261 | Spint1 |
| ENSMUSG00000023150 | 1.004026 | 1.74228 | 2.451004 | 1.004026 | 4.433752 | 3.283347 | Ivns1abp |
| ENSMUSG00000039952 | 1.122147 | 2.067112 | 3.012078 | 1.122147 | 4.430864 | 3.281096 | Dag1 |
| ENSMUSG00000042359 | 2.687246 | 4.636237 | 6.526169 | 2.687246 | 4.42499 | 3.276447 | Osbpl6 |
| ENSMUSG00000033174 | 1.299328 | 4.016104 | 5.846975 | 1.299328 | 4.423281 | 3.275719 | Mgll |
| ENSMUSG00000040998 | 3.45503 | 6.79194 | 7.677845 | 3.45503 | 4.405942 | 3.26193 | Npnt |
| ENSMUSG00000027210 | 1.889931 | 7.205363 | 7.648315 | 1.889931 | 4.396835 | 3.255809 | Meis2 |
| ENSMUSG00000007613 | 1.181207 | 2.185233 | 3.543621 | 1.181207 | 4.391736 | 3.250572 | Tgfbr1 |
| ENSMUSG00000019943 | 1.358388 | 2.510065 | 3.720802 | 1.358388 | 4.389643 | 3.249261 | Atp2b1 |
| ENSMUSG00000030800 | 1.358388 | 2.569125 | 5.787914 | 1.358388 | 4.379416 | 3.238987 | Prss8 |
| ENSMUSG00000045201 | 2.451004 | 5.07919 | 6.171806 | 2.451004 | 4.378549 | 3.238987 | Lrrc3b |
| ENSMUSG00000038366 | 1.210737 | 2.303354 | 3.661742 | 1.210737 | 4.370743 | 3.230173 | Lasp1 |
| ENSMUSG00000042515 | 1.446979 | 2.687246 | 4.636237 | 1.446979 | 4.365374 | 3.224541 | Mum1l1 |
| ENSMUSG00000059921 | 2.716776 | 4.399996 | 6.319457 | 2.716776 | 4.358095 | 3.216397 | Unc5c |
| ENSMUSG00000038903 | 2.273823 | 4.931539 | 5.846975 | 2.273823 | 4.35236 | 3.211654 | Ccdc68 |
| ENSMUSG00000031242 | 2.480535 | 5.256371 | 6.024156 | 2.480535 | 4.344269 | 3.202423 | 2610002M |
| ENSMUSG00000060681 | 2.834897 | 6.083216 | 6.79194 | 2.834897 | 4.337387 | 3.194767 | Slc9a6 |
| ENSMUSG00000022037 | 3.159729 | 6.644289 | 7.677845 | 3.159729 | 4.331077 | 3.187854 | Clu |
| ENSMUSG00000032363 | 1.240267 | 3.691272 | 5.492612 | 1.240267 | 4.326776 | 3.183573 | Adamts7 |
| ENSMUSG00000042372 | 1.033556 | 3.573151 | 5.404022 | 1.033556 | 4.309385 | 3.163431 | Dmrt3 |
| ENSMUSG00000067889 | 3.838923 | 6.053686 | 6.910061 | 3.838923 | 4.305797 | 3.160888 | Sptbn2 |
| ENSMUSG00000071771 | 1.092616 | 2.628186 | 4.724828 | 1.092616 | 4.305394 | 3.160888 | Rhox4b |
| ENSMUSG00000037062 | 1.122147 | 2.008052 | 2.864427 | 1.122147 | 4.29046 | 3.142567 | Sh3glb1 |
| ENSMUSG00000094945 | 1.033556 | 2.864427 | 4.931539 | 1.033556 | 4.261971 | 3.107642 | Rhox4a2 |
| ENSMUSG00000021665 | 3.041608 | 5.935565 | 7.057712 | 3.041608 | 4.255414 | 3.101154 | Hexb |
| ENSMUSG00000020674 | 1.063086 | 2.067112 | 3.36644 | 1.063086 | 4.250646 | 3.096682 | Pxdn |
| ENSMUSG00000023830 | 1.092616 | 2.096642 | 3.130198 | 1.092616 | 4.238839 | 3.085708 | Igf2r |
| ENSMUSG00000019916 | 1.092616 | 2.155703 | 3.248319 | 1.092616 | 4.225295 | 3.072337 | P4ha1 |
| ENSMUSG00000025351 | 1.122147 | 2.185233 | 3.248319 | 1.122147 | 4.207888 | 3.050544 | Cd63 |
| ENSMUSG00000087153 | 1.092616 | 2.096642 | 3.720802 | 1.092616 | 4.197536 | 3.040684 | Gm6483 |
| ENSMUSG00000043252 | 2.155703 | 5.02013 | 6.083216 | 2.155703 | 4.192783 | 3.03694 | Tmem64 |
| ENSMUSG00000059878 | 1.063086 | 2.067112 | 3.100668 | 1.063086 | 4.187303 | 3.032213 | Zfp422 |
| ENSMUSG00000037465 | 1.446979 | 2.953017 | 5.07919 | 1.446979 | 4.170135 | 3.013197 | Klf10 |
| ENSMUSG00000040249 | 1.299328 | 2.569125 | 5.02013 | 1.299328 | 4.169203 | 3.013197 | Lrp1 |
| ENSMUSG00000054814 | 1.476509 | 2.569125 | 4.872479 | 1.476509 | 4.167062 | 3.011605 | Usp46 |
| ENSMUSG00000027303 | 3.33691 | 5.640263 | 6.467108 | 3.33691 | 4.155313 | 3.000944 | Ptpra |
| ENSMUSG00000035863 | 1.240267 | 2.451004 | 4.370466 | 1.240267 | 4.155394 | 3.000944 | Palm |
| ENSMUSG00000028032 | 1.151677 | 2.244293 | 3.632211 | 1.151677 | 4.136318 | 2.979323 | Papss1 |
| ENSMUSG00000020695 | 2.510065 | 4.104694 | 5.935565 | 2.510065 | 4.122804 | 2.963731 | Mrc2 |
| ENSMUSG00000030659 | 1.919461 | 3.750332 | 7.057712 | 1.919461 | 4.121361 | 2.963027 | Nucb2 |
| ENSMUSG00000026944 | 1.830871 | 4.252345 | 5.551673 | 1.830871 | 4.110223 | 2.951785 | Abca2 |
| ENSMUSG00000058881 | 2.746306 | 5.13825 | 6.171806 | 2.746306 | 4.108482 | 2.950665 | Zfp516 |
| ENSMUSG00000014177 | 1.151677 | 2.362414 | 4.429526 | 1.151677 | 4.100614 | 2.942548 | Tvp23b |
| ENSMUSG00000049791 | 2.805367 | 6.230867 | 6.939591 | 2.805367 | 4.098776 | 2.941287 | Fzd4 |
| ENSMUSG00000028759 | 1.004026 | 1.889931 | 2.805367 | 1.004026 | 4.095567 | 2.938186 | Hp1bp3 |
| ENSMUSG00000022475 | 1.506039 | 4.459056 | 5.581203 | 1.506039 | 4.089792 | 2.936368 | Hdac7 |
| ENSMUSG00000036661 | 1.919461 | 4.252345 | 5.404022 | 1.919461 | 4.091164 | 2.936368 | Dennd3 |
| ENSMUSG00000060260 | 1.801341 | 4.518117 | 5.640263 | 1.801341 | 4.090539 | 2.936368 | Pwwp2b |
| ENSMUSG00000095403 | 1.004026 | 1.919461 | 2.953017 | 1.004026 | 4.090911 | 2.936368 | Gm21092 |
| ENSMUSG00000021366 | 3.159729 | 5.492612 | 6.437578 | 3.159729 | 4.081981 | 2.928254 | Hivep1 |
| ENSMUSG00000025776 | 2.628186 | 4.813418 | 6.053686 | 2.628186 | 4.079664 | 2.926316 | Crispld1 |
| ENSMUSG00000069378 | 2.391944 | 5.13825 | 5.994625 | 2.391944 | 4.069558 | 2.913945 | Prdm6 |
| ENSMUSG00000017723 | 1.151677 | 2.391944 | 3.809393 | 1.151677 | 4.06028 | 2.904977 | Wfdc2 |

| | | | | | | | |
|---|---|---|---|---|---|---|---|
| ENSMUSG00000028664 | 2.332884 | 5.581203 | 6.467108 | 2.332884 | 4.053332 | 2.896809 | Ephb2 |
| ENSMUSG00000033149 | 1.092616 | 2.155703 | 3.4255 | 1.092616 | 4.034312 | 2.874702 | Phldb2 |
| ENSMUSG00000036580 | 1.122147 | 2.185233 | 3.33691 | 1.122147 | 4.026596 | 2.86758 | Spg20 |
| ENSMUSG00000040111 | 2.953017 | 5.02013 | 6.644289 | 2.953017 | 4.025416 | 2.867288 | Gramd1b |
| ENSMUSG00000028776 | 2.421474 | 5.285901 | 6.230867 | 2.421474 | 4.012828 | 2.854848 | Tinagl1 |
| ENSMUSG00000062591 | 1.151677 | 2.332884 | 4.075164 | 1.151677 | 4.007643 | 2.851209 | Tubb4a |
| ENSMUSG00000007039 | 1.063086 | 2.214763 | 3.897983 | 1.063086 | 3.998038 | 2.841589 | Ddah2 |
| ENSMUSG00000032733 | 2.923487 | 5.374492 | 6.289927 | 2.923487 | 3.983971 | 2.823725 | Snx33 |
| ENSMUSG00000061048 | 1.889931 | 3.750332 | 8.032207 | 1.889931 | 3.978728 | 2.818845 | Cdh3 |
| ENSMUSG00000037995 | 1.269798 | 2.657716 | 4.842949 | 1.269798 | 3.972549 | 2.812688 | Igsf9 |
| ENSMUSG00000037989 | 1.62416 | 4.134224 | 5.463082 | 1.62416 | 3.948919 | 2.785079 | Wnk2 |
| ENSMUSG00000050732 | 1.181207 | 2.362414 | 3.661742 | 1.181207 | 3.941858 | 2.776596 | Vamp8 |
| ENSMUSG00000039154 | 2.746306 | 4.754358 | 5.787914 | 2.746306 | 3.924639 | 2.759643 | Shd |
| ENSMUSG00000041653 | 2.244293 | 4.193285 | 6.348988 | 2.244293 | 3.91972 | 2.754083 | Pnpla3 |
| ENSMUSG00000041215 | 1.033556 | 2.155703 | 3.691272 | 1.033556 | 3.894053 | 2.726597 | Yeats2 |
| ENSMUSG00000030747 | 1.240267 | 2.362414 | 4.399996 | 1.240267 | 3.888346 | 2.720865 | Dgat2 |
| ENSMUSG00000071855 | 1.387918 | 3.36644 | 4.872479 | 1.387918 | 3.856322 | 2.683482 | Ccdc112 |
| ENSMUSG00000020312 | 2.155703 | 4.990599 | 5.846975 | 2.155703 | 3.85081 | 2.679009 | Shc2 |
| ENSMUSG00000021831 | 1.506039 | 2.923487 | 4.754358 | 1.506039 | 3.849679 | 2.679009 | Ero1l |
| ENSMUSG00000017756 | 1.417448 | 3.218789 | 6.408048 | 1.417448 | 3.846977 | 2.677906 | Slc12a7 |
| ENSMUSG00000031995 | 1.387918 | 2.864427 | 5.522143 | 1.387918 | 3.842527 | 2.673856 | St14 |
| ENSMUSG00000023043 | 1.063086 | 2.362414 | 4.370466 | 1.063086 | 3.817697 | 2.644914 | Krt18 |
| ENSMUSG00000022837 | 1.240267 | 2.834897 | 4.370466 | 1.240267 | 3.805692 | 2.632455 | Iqcb1 |
| ENSMUSG00000015189 | 1.889931 | 5.846975 | 6.614759 | 1.889931 | 3.797369 | 2.624995 | Casd1 |
| ENSMUSG00000025161 | 1.033556 | 2.126173 | 2.923487 | 1.033556 | 3.787651 | 2.615601 | Slc16a3 |
| ENSMUSG00000028173 | 1.033556 | 2.214763 | 3.4255 | 1.033556 | 3.765651 | 2.589245 | Wls |
| ENSMUSG00000022483 | 2.096642 | 7.559725 | 8.179858 | 2.096642 | 3.745723 | 2.565648 | Col2a1 |
| ENSMUSG00000022367 | 2.332884 | 6.260397 | 7.264423 | 2.332884 | 3.703102 | 2.521993 | Has2 |
| ENSMUSG00000028413 | 1.210737 | 2.687246 | 4.222815 | 1.210737 | 3.700995 | 2.520014 | B4galt1 |
| ENSMUSG00000040272 | 1.476509 | 4.606707 | 5.551673 | 1.476509 | 3.695769 | 2.514646 | Accs |
| ENSMUSG00000035877 | 2.893957 | 5.344962 | 6.526169 | 2.893957 | 3.67795 | 2.495506 | Zhx3 |
| ENSMUSG00000010830 | 1.74228 | 4.340936 | 5.846975 | 1.74228 | 3.672068 | 2.488253 | Kdelr3 |
| ENSMUSG00000034055 | 1.948992 | 4.518117 | 5.965095 | 1.948992 | 3.669195 | 2.485184 | Phka1 |
| ENSMUSG00000039960 | 1.68322 | 5.551673 | 6.526169 | 1.68322 | 3.645055 | 2.456153 | Rhou |
| ENSMUSG00000059316 | 2.421474 | 4.577177 | 5.965095 | 2.421474 | 3.635947 | 2.445269 | Slc27a4 |
| ENSMUSG00000025352 | 2.480535 | 4.665768 | 5.758384 | 2.480535 | 3.622433 | 2.429112 | Gdf11 |
| ENSMUSG00000032353 | 1.446979 | 6.171806 | 6.70335 | 1.446979 | 3.608971 | 2.41482 | Tmed3 |
| ENSMUSG00000029781 | 1.033556 | 2.214763 | 3.720802 | 1.033556 | 3.600478 | 2.410154 | Fkbp9 |
| ENSMUSG00000067276 | 2.864427 | 7.146302 | 8.977173 | 2.864427 | 3.601621 | 2.410154 | Capn6 |
| ENSMUSG00000024665 | 1.063086 | 2.657716 | 4.311405 | 1.063086 | 3.594104 | 2.403001 | Fads2 |
| ENSMUSG00000031274 | 2.628186 | 5.699324 | 6.467108 | 2.628186 | 3.588839 | 2.3965 | Col4a5 |
| ENSMUSG00000035678 | 2.037582 | 4.104694 | 5.610733 | 2.037582 | 3.568261 | 2.373785 | Tnfsf9 |
| ENSMUSG00000038280 | 2.834897 | 5.10872 | 5.994625 | 2.834897 | 3.56426 | 2.369033 | Ostm1 |
| ENSMUSG00000055320 | 1.151677 | 2.480535 | 4.754358 | 1.151677 | 3.535973 | 2.335257 | Tead1 |
| ENSMUSG00000032231 | 2.805367 | 6.112746 | 7.175832 | 2.805367 | 3.53199 | 2.330497 | Anxa2 |
| ENSMUSG00000035403 | 2.185233 | 6.053686 | 6.880531 | 2.185233 | 3.522744 | 2.319636 | Crb2 |
| ENSMUSG00000024598 | 1.210737 | 2.893957 | 4.606707 | 1.210737 | 3.520691 | 2.31887 | Fbn2 |
| ENSMUSG00000020099 | 1.594629 | 5.256371 | 6.260397 | 1.594629 | 3.518126 | 2.316105 | Unc5b |
| ENSMUSG00000022329 | 1.151677 | 2.923487 | 6.526169 | 1.151677 | 3.517498 | 2.316087 | Stk3 |
| ENSMUSG00000023232 | 1.328858 | 6.083216 | 6.79194 | 1.328858 | 3.511643 | 2.312116 | Serinc2 |
| ENSMUSG00000053093 | 2.037582 | 4.134224 | 6.112746 | 2.037582 | 3.499468 | 2.299133 | Myh7 |
| ENSMUSG00000073295 | 1.269798 | 3.012078 | 6.555699 | 1.269798 | 3.492299 | 2.292364 | Nudt11 |
| ENSMUSG00000027217 | 1.446979 | 4.193285 | 6.319457 | 1.446979 | 3.480272 | 2.277032 | Tspan18 |
| ENSMUSG00000025082 | 2.657716 | 5.02013 | 5.906035 | 2.657716 | 3.450119 | 2.242387 | Vwa2 |
| ENSMUSG00000038732 | 2.067112 | 5.846975 | 6.998651 | 2.067112 | 3.445039 | 2.237642 | Mboat1 |
| ENSMUSG00000019975 | 1.269798 | 3.277849 | 5.07919 | 1.269798 | 3.434029 | 2.225225 | Ikbip |
| ENSMUSG00000057315 | 1.476509 | 6.201337 | 6.880531 | 1.476509 | 3.433298 | 2.225011 | Arhgap24 |
| ENSMUSG00000021196 | 2.569125 | 5.817444 | 6.880531 | 2.569125 | 3.429184 | 2.219948 | Pfkp |

| | | | | | | | |
|---|---|---|---|---|---|---|---|
| ENSMUSG00000060512 | 2.155703 | 6.260397 | 6.939591 | 2.155703 | 3.409336 | 2.197257 | 0610040J0 |
| ENSMUSG00000031740 | 1.358388 | 3.543621 | 7.736906 | 1.358388 | 3.407847 | 2.196537 | Mmp2 |
| ENSMUSG00000044365 | 1.68322 | 5.02013 | 5.994625 | 1.68322 | 3.396049 | 2.184178 | Cxxc4 |
| ENSMUSG00000028194 | 1.033556 | 2.510065 | 4.134224 | 1.033556 | 3.376408 | 2.160477 | Ddah1 |
| ENSMUSG00000070576 | 1.358388 | 3.45503 | 6.408048 | 1.358388 | 3.362029 | 2.149371 | Mn1 |
| ENSMUSG00000021597 | 1.299328 | 3.041608 | 5.07919 | 1.299328 | 3.356509 | 2.14381 | Ankrd32 |
| ENSMUSG00000009876 | 1.62416 | 5.965095 | 7.057712 | 1.62416 | 3.354705 | 2.142464 | Cox4i2 |
| ENSMUSG00000033233 | 2.421474 | 4.665768 | 6.024156 | 2.421474 | 3.349371 | 2.136663 | Trim45 |
| ENSMUSG00000039623 | 1.62416 | 3.809393 | 5.256371 | 1.62416 | 3.346758 | 2.13403 | Ap5z1 |
| ENSMUSG00000047557 | 2.096642 | 5.758384 | 6.555699 | 2.096642 | 3.338392 | 2.123927 | Lxn |
| ENSMUSG00000034613 | 1.978522 | 5.640263 | 6.467108 | 1.978522 | 3.334766 | 2.119461 | Ppm1h |
| ENSMUSG00000040856 | 1.68322 | 4.163755 | 5.669793 | 1.68322 | 3.31653 | 2.094539 | Dlk1 |
| ENSMUSG00000021065 | 1.063086 | 2.716776 | 6.142276 | 1.063086 | 3.304534 | 2.080219 | Fut8 |
| ENSMUSG00000029838 | 2.746306 | 5.10872 | 6.201337 | 2.746306 | 3.298757 | 2.074943 | Ptn |
| ENSMUSG00000025855 | 2.332884 | 5.610733 | 6.408048 | 2.332884 | 3.294014 | 2.070371 | Prkar1b |
| ENSMUSG00000031965 | 2.451004 | 6.201337 | 6.969121 | 2.451004 | 3.284198 | 2.06379 | Tbx20 |
| ENSMUSG00000021690 | 2.037582 | 4.783888 | 5.817444 | 2.037582 | 3.276954 | 2.056298 | Jmy |
| ENSMUSG00000001507 | 1.181207 | 2.805367 | 7.648315 | 1.181207 | 3.263572 | 2.039781 | Itga3 |
| ENSMUSG00000002668 | 1.092616 | 5.463082 | 6.083216 | 1.092616 | 3.228529 | 2.000466 | Dennd1c |
| ENSMUSG00000036687 | 2.155703 | 5.610733 | 6.496638 | 2.155703 | 3.221869 | 1.992913 | Tmem184a |
| ENSMUSG00000021371 | 1.62416 | 5.522143 | 6.378518 | 1.62416 | 3.220628 | 1.991837 | Mcur1 |
| ENSMUSG00000021709 | 1.299328 | 5.492612 | 6.171806 | 1.299328 | 3.206183 | 1.978737 | Erbb2ip |
| ENSMUSG00000041734 | 1.978522 | 4.518117 | 5.610733 | 1.978522 | 3.197134 | 1.969805 | Kirrel |
| ENSMUSG00000018166 | 1.446979 | 6.260397 | 7.205363 | 1.446979 | 3.185186 | 1.955075 | Erbb3 |
| ENSMUSG00000040322 | 1.299328 | 3.248319 | 6.79194 | 1.299328 | 3.17662 | 1.945316 | Slc25a24 |
| ENSMUSG00000010025 | 1.889931 | 5.433552 | 6.319457 | 1.889931 | 3.175572 | 1.944494 | Aldh3a2 |
| ENSMUSG00000040297 | 1.122147 | 3.484561 | 5.13825 | 1.122147 | 3.166074 | 1.936227 | Suco |
| ENSMUSG00000026208 | 2.598655 | 6.348988 | 7.205363 | 2.598655 | 3.146023 | 1.910756 | Des |
| ENSMUSG00000031209 | 1.033556 | 2.746306 | 6.851 | 1.033556 | 3.143793 | 1.908156 | Heph |
| ENSMUSG00000020642 | 1.210737 | 5.699324 | 6.467108 | 1.210737 | 3.116163 | 1.881924 | Rnf144a |
| ENSMUSG00000029999 | 1.476509 | 5.463082 | 6.201337 | 1.476509 | 3.116664 | 1.881924 | Tgfa |
| ENSMUSG00000041594 | 1.033556 | 3.012078 | 6.289927 | 1.033556 | 3.109445 | 1.873289 | Tmtc4 |
| ENSMUSG00000079042 | 1.033556 | 3.012078 | 4.990599 | 1.033556 | 3.101165 | 1.863003 | Apela |
| ENSMUSG00000032020 | 1.889931 | 5.522143 | 6.289927 | 1.889931 | 3.097526 | 1.858955 | Ubash3b |
| ENSMUSG00000029032 | 1.65369 | 5.669793 | 6.526169 | 1.65369 | 3.090997 | 1.851267 | Arhgef16 |
| ENSMUSG00000016494 | 1.62416 | 4.045634 | 5.285901 | 1.62416 | 3.074025 | 1.83135 | Cd34 |
| ENSMUSG00000048502 | 1.74228 | 4.518117 | 5.610733 | 1.74228 | 3.066118 | 1.823792 | Duxbl1 |
| ENSMUSG00000027583 | 1.387918 | 5.728854 | 6.467108 | 1.387918 | 3.055195 | 1.816706 | Zbtb46 |
| ENSMUSG00000028766 | 1.801341 | 5.935565 | 6.70335 | 1.801341 | 3.051764 | 1.814352 | Alpl |
| ENSMUSG00000041889 | 1.181207 | 5.699324 | 6.496638 | 1.181207 | 3.047149 | 1.810782 | Shisa4 |
| ENSMUSG00000035629 | 1.269798 | 3.750332 | 5.433552 | 1.269798 | 3.042101 | 1.805193 | 1700021K1 |
| ENSMUSG00000029729 | 1.801341 | 5.07919 | 6.083216 | 1.801341 | 3.033856 | 1.793412 | Zkscan1 |
| ENSMUSG00000046761 | 2.155703 | 5.404022 | 6.201337 | 2.155703 | 3.030797 | 1.790135 | Fam83h |
| ENSMUSG00000068130 | 1.269798 | 4.488587 | 5.522143 | 1.269798 | 3.024215 | 1.782193 | Zfp442 |
| ENSMUSG00000035498 | 1.004026 | 6.142276 | 6.79194 | 1.004026 | 3.015189 | 1.774538 | Cdcp1 |
| ENSMUSG00000038679 | 2.037582 | 5.551673 | 6.496638 | 2.037582 | 3.012197 | 1.772012 | Trps1 |
| ENSMUSG00000027520 | 1.387918 | 5.02013 | 6.024156 | 1.387918 | 3.005907 | 1.764024 | Zdbf2 |
| ENSMUSG00000072623 | 1.151677 | 3.543621 | 5.256371 | 1.151677 | 2.997955 | 1.755038 | Zfp9 |
| ENSMUSG00000022305 | 1.328858 | 5.965095 | 6.644289 | 1.328858 | 2.986571 | 1.743148 | Lrp12 |
| ENSMUSG00000090919 | 1.122147 | 6.053686 | 6.585229 | 1.122147 | 2.975289 | 1.730402 | Pabpc4l |
| ENSMUSG00000024451 | 1.565099 | 4.281875 | 5.551673 | 1.565099 | 2.969056 | 1.723505 | Arap3 |
| ENSMUSG00000030551 | 1.65369 | 5.463082 | 6.614759 | 1.65369 | 2.965753 | 1.720419 | Nr2f2 |
| ENSMUSG00000028786 | 2.303354 | 6.112746 | 7.028182 | 2.303354 | 2.963021 | 1.717571 | Tmem54 |
| ENSMUSG00000078350 | 1.919461 | 4.961069 | 5.846975 | 1.919461 | 2.962612 | 1.717571 | Smim1 |
| ENSMUSG00000074282 | 1.328858 | 4.636237 | 5.640263 | 1.328858 | 2.958198 | 1.712771 | Zfp94 |
| ENSMUSG00000028545 | 1.033556 | 5.610733 | 6.319457 | 1.033556 | 2.954529 | 1.709315 | Bend5 |
| ENSMUSG00000030064 | 1.063086 | 4.045634 | 6.437578 | 1.063086 | 2.953596 | 1.709291 | Frmd4b |
| ENSMUSG00000034463 | 1.889931 | 5.581203 | 6.73288 | 1.889931 | 2.949505 | 1.708243 | Scara3 |

| | | | | | | | |
|---|---|---|---|---|---|---|---|
| ENSMUSG00000015312 | 1.830871 | 4.813418 | 5.699324 | 1.830871 | 2.940657 | 1.697494 | Gadd45b |
| ENSMUSG00000034614 | 1.535569 | 4.961069 | 6.171806 | 1.535569 | 2.938329 | 1.694787 | Pik3ip1 |
| ENSMUSG00000044921 | 1.62416 | 4.311405 | 5.463082 | 1.62416 | 2.938059 | 1.694787 | Rassf9 |
| ENSMUSG00000048970 | 1.004026 | 2.893957 | 6.260397 | 1.004026 | 2.934482 | 1.689941 | C1galt1c1 |
| ENSMUSG00000001665 | 1.594629 | 5.463082 | 6.260397 | 1.594629 | 2.911484 | 1.662567 | Gstt3 |
| ENSMUSG00000031712 | 2.096642 | 6.024156 | 6.79194 | 2.096642 | 2.909753 | 1.661495 | Il15 |
| ENSMUSG00000021057 | 1.033556 | 3.39597 | 6.614759 | 1.033556 | 2.900452 | 1.65096 | Akap5 |
| ENSMUSG00000006586 | 1.65369 | 5.10872 | 6.437578 | 1.65369 | 2.89328 | 1.645734 | Runx1t1 |
| ENSMUSG00000031434 | 1.004026 | 4.577177 | 5.817444 | 1.004026 | 2.891861 | 1.644503 | Morc4 |
| ENSMUSG00000035992 | 1.65369 | 4.222815 | 5.581203 | 1.65369 | 2.889696 | 1.643025 | Fnip1 |
| ENSMUSG00000041605 | 1.033556 | 5.699324 | 6.496638 | 1.033556 | 2.889158 | 1.642813 | 5730559C1 |
| ENSMUSG00000018474 | 1.830871 | 4.577177 | 5.581203 | 1.830871 | 2.87468 | 1.625998 | Chd3 |
| ENSMUSG00000031557 | 1.181207 | 6.910061 | 7.707376 | 1.181207 | 2.86586 | 1.616029 | Plekha2 |
| ENSMUSG00000052915 | 1.063086 | 3.45503 | 5.226841 | 1.063086 | 2.860478 | 1.61036 | Msl1 |
| ENSMUSG00000053113 | 1.240267 | 5.581203 | 6.348988 | 1.240267 | 2.84196 | 1.590509 | Socs3 |
| ENSMUSG00000038205 | 1.004026 | 5.846975 | 6.585229 | 1.004026 | 2.830141 | 1.577847 | Prkab2 |
| ENSMUSG00000056383 | 1.210737 | 3.779862 | 5.344962 | 1.210737 | 2.829133 | 1.577178 | AI987944 |
| ENSMUSG00000001870 | 1.358388 | 3.986574 | 5.404022 | 1.358388 | 2.824911 | 1.572398 | Ltbp1 |
| ENSMUSG00000035476 | 1.299328 | 5.344962 | 6.348988 | 1.299328 | 2.819315 | 1.565463 | Tab3 |
| ENSMUSG00000039086 | 1.594629 | 5.433552 | 6.171806 | 1.594629 | 2.814119 | 1.559742 | Ss18l1 |
| ENSMUSG00000041117 | 1.004026 | 4.340936 | 5.374492 | 1.004026 | 2.814242 | 1.559742 | Ccdc8 |
| ENSMUSG00000031273 | 1.417448 | 5.551673 | 6.614759 | 1.417448 | 2.810924 | 1.555344 | Col4a6 |
| ENSMUSG00000039395 | 1.151677 | 4.577177 | 5.728854 | 1.151677 | 2.810367 | 1.555071 | Mreg |
| ENSMUSG00000021638 | 1.151677 | 5.935565 | 6.673819 | 1.151677 | 2.797786 | 1.540736 | Ocln |
| ENSMUSG00000089876 | 1.535569 | 5.256371 | 5.994625 | 1.535569 | 2.762978 | 1.494822 | Tmem102 |
| ENSMUSG00000028414 | 1.151677 | 4.045634 | 5.492612 | 1.151677 | 2.762017 | 1.493898 | Fktn |
| ENSMUSG00000015452 | 1.594629 | 4.754358 | 5.699324 | 1.594629 | 2.75749 | 1.489108 | Ager |
| ENSMUSG00000020282 | 1.299328 | 5.02013 | 6.467108 | 1.299328 | 2.756456 | 1.488065 | Rhbdf1 |
| ENSMUSG00000021706 | 1.269798 | 4.754358 | 6.053686 | 1.269798 | 2.752099 | 1.483668 | Zfyve16 |
| ENSMUSG00000049307 | 1.387918 | 5.787914 | 6.526169 | 1.387918 | 2.75181 | 1.483668 | Fut4 |
| ENSMUSG00000030917 | 1.240267 | 5.315431 | 6.289927 | 1.240267 | 2.746933 | 1.480811 | Tmem159 |
| ENSMUSG00000072675 | 1.476509 | 3.927513 | 5.581203 | 1.476509 | 2.740087 | 1.472732 | Duxbl2 |
| ENSMUSG00000047821 | 1.71275 | 5.758384 | 6.644289 | 1.71275 | 2.73683 | 1.471093 | Trim16 |
| ENSMUSG00000030691 | 1.476509 | 5.787914 | 6.673819 | 1.476509 | 2.727329 | 1.461791 | Fchsd2 |
| ENSMUSG00000072672 | 1.68322 | 4.783888 | 5.758384 | 1.68322 | 2.724067 | 1.45834 | Duxbl3 |
| ENSMUSG00000094595 | 1.299328 | 4.340936 | 5.315431 | 1.299328 | 2.716904 | 1.450403 | Fsbp |
| ENSMUSG00000029059 | 1.387918 | 4.783888 | 5.699324 | 1.387918 | 2.712269 | 1.447046 | Fam213b |
| ENSMUSG00000020246 | 1.033556 | 5.167781 | 6.201337 | 1.033556 | 2.700587 | 1.437247 | Hcfc2 |
| ENSMUSG00000039131 | 1.151677 | 4.281875 | 5.669793 | 1.151677 | 2.699161 | 1.436758 | Gipc2 |
| ENSMUSG00000047242 | 1.387918 | 4.961069 | 5.994625 | 1.387918 | 2.696078 | 1.43374 | Taf9b |
| ENSMUSG00000078452 | 1.328858 | 4.961069 | 5.935565 | 1.328858 | 2.696142 | 1.43374 | Raet1d |
| ENSMUSG00000039007 | 1.417448 | 5.669793 | 6.408048 | 1.417448 | 2.695549 | 1.433467 | Cpq |
| ENSMUSG00000017390 | 1.299328 | 5.197311 | 6.142276 | 1.299328 | 2.690531 | 1.430017 | Aldoc |
| ENSMUSG00000026042 | 1.387918 | 5.167781 | 6.230867 | 1.387918 | 2.691556 | 1.430017 | Col5a2 |
| ENSMUSG00000030606 | 1.68322 | 4.872479 | 5.994625 | 1.68322 | 2.67652 | 1.410349 | Hapln3 |
| ENSMUSG00000003812 | 1.033556 | 3.573151 | 5.581203 | 1.033556 | 2.675045 | 1.409107 | Dnase2a |
| ENSMUSG00000042156 | 1.506039 | 5.02013 | 5.965095 | 1.506039 | 2.664596 | 1.396777 | Dzip1 |
| ENSMUSG00000096847 | 1.063086 | 4.370466 | 6.024156 | 1.063086 | 2.655243 | 1.385689 | Tmem151l |
| ENSMUSG00000050503 | 1.417448 | 4.665768 | 5.581203 | 1.417448 | 2.638389 | 1.363515 | Fbxl22 |
| ENSMUSG00000021236 | 1.240267 | 5.02013 | 6.142276 | 1.240267 | 2.636425 | 1.360895 | Entpd5 |
| ENSMUSG00000056124 | 1.004026 | 4.222815 | 6.348988 | 1.004026 | 2.630127 | 1.354789 | B4galt6 |
| ENSMUSG00000038225 | 1.151677 | 4.665768 | 5.787914 | 1.151677 | 2.62082 | 1.348901 | Primpol |
| ENSMUSG00000068923 | 1.004026 | 4.222815 | 5.728854 | 1.004026 | 2.600324 | 1.328711 | Syt11 |
| ENSMUSG00000028337 | 1.269798 | 4.606707 | 5.581203 | 1.269798 | 2.598872 | 1.327432 | Coro2a |
| ENSMUSG00000031227 | 1.092616 | 4.931539 | 5.994625 | 1.092616 | 2.589777 | 1.318817 | Magee1 |
| ENSMUSG00000025076 | 1.122147 | 4.606707 | 5.551673 | 1.122147 | 2.583383 | 1.312966 | Casp7 |
| ENSMUSG00000043153 | 1.122147 | 4.429526 | 5.610733 | 1.122147 | 2.56873 | 1.298105 | Ispd |
| ENSMUSG00000025529 | 1.210737 | 5.492612 | 6.289927 | 1.210737 | 2.539927 | 1.264763 | Zfp711 |

| | | | | | | | |
|---|---|---|---|---|---|---|---|
| ENSMUSG00000027584 | 1.358388 | 5.374492 | 6.260397 | 1.358388 | 2.52935 | 1.255585 | Oprl1 |
| ENSMUSG00000027222 | 1.210737 | 4.636237 | 5.551673 | 1.210737 | 2.521678 | 1.248808 | Pex16 |
| ENSMUSG00000040511 | 1.122147 | 4.193285 | 5.433552 | 1.122147 | 2.511868 | 1.239172 | Pvr |
| ENSMUSG00000021453 | 1.092616 | 4.990599 | 5.787914 | 1.092616 | 2.499601 | 1.226583 | Gadd45g |
| ENSMUSG00000000753 | 1.151677 | 4.606707 | 6.142276 | 1.151677 | 2.479287 | 1.208111 | Serpinf1 |
| ENSMUSG00000026678 | 1.417448 | 6.526169 | 7.648315 | 1.417448 | 2.466022 | 1.197651 | Rgs5 |
| ENSMUSG00000073968 | 1.063086 | 5.463082 | 6.378518 | 1.063086 | 2.443497 | 1.173871 | Trim68 |
| ENSMUSG00000037625 | 1.004026 | 4.665768 | 7.530194 | 1.004026 | 2.358482 | 1.095274 | Cldn11 |
| ENSMUSG00000040726 | -6.79194 | -5.72885 | -1.06309 | -1.06309 | -2.40171 | -1.13706 | Hesx1 |
| ENSMUSG00000029648 | -6.76241 | -5.78791 | -1.00403 | -1.00403 | -2.42908 | -1.16136 | Flt1 |
| ENSMUSG00000026497 | -7.05771 | -6.31946 | -1.2698 | -1.2698 | -2.49242 | -1.21813 | Mixl1 |
| ENSMUSG00000052133 | -5.90603 | -5.19731 | -1.15168 | -1.15168 | -2.52839 | -1.25452 | Sema5b |
| ENSMUSG00000030844 | -5.34496 | -4.13422 | -1.18121 | -1.18121 | -2.55878 | -1.28555 | Rgs10 |
| ENSMUSG00000027500 | -5.43355 | -4.63624 | -1.09262 | -1.09262 | -2.58024 | -1.30944 | Stmn2 |
| ENSMUSG00000003436 | -7.41207 | -6.2604 | -1.53557 | -1.53557 | -2.58776 | -1.31635 | Dll3 |
| ENSMUSG00000001542 | -5.31543 | -3.89798 | -1.29933 | -1.29933 | -2.59706 | -1.32556 | Ell2 |
| ENSMUSG00000042675 | -5.19731 | -4.04563 | -1.15168 | -1.15168 | -2.63152 | -1.35574 | Ypel3 |
| ENSMUSG00000051427 | -5.04966 | -3.4255 | -1.09262 | -1.09262 | -2.66457 | -1.39678 | Ccdc157 |
| ENSMUSG00000022763 | -5.40402 | -4.0161 | -1.03356 | -1.03356 | -2.71339 | -1.44716 | Aifm3 |
| ENSMUSG00000046598 | -6.23087 | -5.34496 | -1.44698 | -1.44698 | -2.71465 | -1.44765 | Bdh1 |
| ENSMUSG00000017453 | -4.87248 | -3.57315 | -1.03356 | -1.03356 | -2.73029 | -1.46418 | Pipox |
| ENSMUSG00000006958 | -7.4416 | -6.46711 | -1.8604 | -1.8604 | -2.74802 | -1.48081 | Chrd |
| ENSMUSG00000045573 | -7.64832 | -6.52617 | -1.94899 | -1.94899 | -2.7487 | -1.4811 | Penk |
| ENSMUSG00000010136 | -6.76241 | -4.31141 | -1.77181 | -1.77181 | -2.76069 | -1.49267 | Pifo |
| ENSMUSG00000026833 | -5.46308 | -4.04563 | -1.38792 | -1.38792 | -2.7824 | -1.52072 | Olfm1 |
| ENSMUSG00000024678 | -6.14228 | -5.37449 | -1.77181 | -1.77181 | -2.81664 | -1.56188 | Ms4a4d |
| ENSMUSG00000011256 | -6.58523 | -5.9651 | -1.32886 | -1.32886 | -2.85654 | -1.60708 | Adam19 |
| ENSMUSG00000066440 | -5.40402 | -4.60671 | -1.80134 | -1.80134 | -2.85742 | -1.60785 | Zfyve26 |
| ENSMUSG00000038192 | -11.4282 | -8.7114 | -2.77584 | -2.77584 | -2.88591 | -1.63965 | Cer1 |
| ENSMUSG00000024987 | -8.38657 | -7.05771 | -1.15168 | -1.15168 | -2.88767 | -1.64175 | Cyp26a1 |
| ENSMUSG00000035305 | -6.49664 | -5.55167 | -1.50604 | -1.50604 | -2.89306 | -1.64573 | Ror1 |
| ENSMUSG00000032184 | -5.90603 | -5.07919 | -1.88993 | -1.88993 | -2.90011 | -1.65096 | Lysmd2 |
| ENSMUSG00000025386 | -4.57718 | -3.63221 | -1.24027 | -1.24027 | -2.91995 | -1.67271 | Pde6g |
| ENSMUSG00000031523 | -6.02416 | -5.2859 | -1.12215 | -1.12215 | -2.9884 | -1.74447 | Dlc1 |
| ENSMUSG00000072944 | -5.52214 | -4.6953 | -1.8604 | -1.8604 | -3.00522 | -1.76402 | Nup62cl |
| ENSMUSG00000041134 | -5.16778 | -4.4 | -1.88993 | -1.88993 | -3.05016 | -1.81392 | Cyyr1 |
| ENSMUSG00000046791 | -5.72885 | -4.9906 | -2.09664 | -2.09664 | -3.06083 | -1.81873 | 2410016O |
| ENSMUSG00000095098 | -5.02013 | -4.10469 | -1.80134 | -1.80134 | -3.08461 | -1.84465 | Ccdc85b |
| ENSMUSG00000015659 | -5.43355 | -4.63624 | -1.97852 | -1.97852 | -3.13189 | -1.8976 | Serac1 |
| ENSMUSG00000051851 | -5.34496 | -4.31141 | -2.24429 | -2.24429 | -3.14044 | -1.90532 | Cxx1c |
| ENSMUSG00000021765 | -6.43758 | -4.84295 | -2.09664 | -2.09664 | -3.14161 | -1.90635 | Fst |
| ENSMUSG00000101972 | -4.6953 | -3.80939 | -1.53557 | -1.53557 | -3.16323 | -1.93345 | Hist1h3i |
| ENSMUSG00000026872 | -4.9906 | -3.86845 | -1.8604 | -1.8604 | -3.19519 | -1.96767 | Zeb2 |
| ENSMUSG00000026124 | -6.93959 | -5.90603 | -1.38792 | -1.38792 | -3.19773 | -1.96981 | Cfc1 |
| ENSMUSG00000046634 | -5.72885 | -4.75436 | -2.36241 | -2.36241 | -3.23917 | -2.01385 | Pkd1l1 |
| ENSMUSG00000020902 | -6.52617 | -5.81744 | -1.80134 | -1.80134 | -3.24664 | -2.02105 | Ntn1 |
| ENSMUSG00000027358 | -7.29395 | -6.5557 | -2.42147 | -2.42147 | -3.31135 | -2.08858 | Bmp2 |
| ENSMUSG00000050010 | -4.84295 | -3.57315 | -1.18121 | -1.18121 | -3.36484 | -2.15025 | Shisa3 |
| ENSMUSG00000071719 | -5.22684 | -3.86845 | -1.32886 | -1.32886 | -3.36704 | -2.15181 | Tmem28 |
| ENSMUSG00000027349 | -5.34496 | -3.69127 | -1.24027 | -1.24027 | -3.39104 | -2.17966 | Fam98b |
| ENSMUSG00000027676 | -5.37449 | -4.13422 | -1.53557 | -1.53557 | -3.40154 | -2.19099 | Ccdc39 |
| ENSMUSG00000024565 | -5.5812 | -4.42953 | -2.12617 | -2.12617 | -3.40256 | -2.19163 | Sall3 |
| ENSMUSG00000031119 | -7.47113 | -2.1557 | -1.00403 | -1.00403 | -3.40711 | -2.19654 | Gpc4 |
| ENSMUSG00000037683 | -5.2859 | -4.31141 | -1.29933 | -1.29933 | -3.40712 | -2.19654 | Armc3 |
| ENSMUSG00000028519 | -6.67382 | -6.17181 | -1.24027 | -1.24027 | -3.41181 | -2.19916 | Dab1 |
| ENSMUSG00000060487 | -5.34496 | -4.28188 | -2.21476 | -2.21476 | -3.53821 | -2.33754 | Samd5 |
| ENSMUSG00000050555 | -5.07919 | -3.89798 | -1.50604 | -1.50604 | -3.55593 | -2.35883 | Hyls1 |
| ENSMUSG00000055254 | -5.61073 | -4.4 | -2.451 | -2.451 | -3.55577 | -2.35883 | Ntrk2 |

| | | | | | | | |
|---|---|---|---|---|---|---|---|
| ENSMUSG00000020140 | -5.55167 | -4.16375 | -2.30335 | -2.30335 | -3.56832 | -2.37378 | Lgr5 |
| ENSMUSG00000036885 | -6.34899 | -5.66979 | -1.8604 | -1.8604 | -3.57137 | -2.37583 | Arhgef26 |
| ENSMUSG00000075304 | -7.91409 | -3.07114 | -1.38792 | -1.38792 | -3.57098 | -2.37583 | Sp5 |
| ENSMUSG00000044338 | -7.38254 | -4.90201 | -2.27382 | -2.27382 | -3.59688 | -2.406 | Aplnr |
| ENSMUSG00000071398 | -5.84697 | -4.4 | -2.51006 | -2.51006 | -3.60592 | -2.41257 | 2410004P( |
| ENSMUSG00000025020 | -5.22684 | -4.07516 | -1.80134 | -1.80134 | -3.6134 | -2.42012 | Slit1 |
| ENSMUSG00000035849 | -4.93154 | -3.95704 | -1.53557 | -1.53557 | -3.62328 | -2.42937 | Krt222 |
| ENSMUSG00000026672 | -4.93154 | -3.89798 | -1.09262 | -1.09262 | -3.68398 | -2.50202 | Optn |
| ENSMUSG00000033308 | -5.49261 | -4.42953 | -1.5651 | -1.5651 | -3.70386 | -2.52199 | Dpyd |
| ENSMUSG00000022389 | -5.61073 | -4.6953 | -2.451 | -2.451 | -3.73516 | -2.5559 | Tef |
| ENSMUSG00000024304 | -6.99865 | -6.23087 | -2.24429 | -2.24429 | -3.74102 | -2.56205 | Cdh2 |
| ENSMUSG00000037868 | -5.72885 | -3.77986 | -2.18523 | -2.18523 | -3.74217 | -2.56269 | Egr2 |
| ENSMUSG00000027562 | -5.13825 | -2.71678 | -1.24027 | -1.24027 | -3.78167 | -2.60872 | Car2 |
| ENSMUSG00000039419 | -5.55167 | -4.0161 | -2.21476 | -2.21476 | -3.81092 | -2.63863 | Cntnap2 |
| ENSMUSG00000037716 | -5.84697 | -4.90201 | -2.27382 | -2.27382 | -3.81614 | -2.64378 | Ccdc33 |
| ENSMUSG00000034227 | -5.31543 | -4.22281 | -1.83087 | -1.83087 | -3.82662 | -2.65613 | Foxj1 |
| ENSMUSG00000047446 | -5.64026 | -4.0161 | -1.53557 | -1.53557 | -3.83503 | -2.66461 | Arl4a |
| ENSMUSG00000053684 | -4.63624 | -3.60268 | -1.15168 | -1.15168 | -3.85005 | -2.67901 | BC048403 |
| ENSMUSG00000041498 | -4.90201 | -4.0161 | -1.83087 | -1.83087 | -3.92942 | -2.76188 | Kif14 |
| ENSMUSG00000021944 | -5.37449 | -4.28188 | -2.68725 | -2.68725 | -4.01002 | -2.8533 | Gata4 |
| ENSMUSG00000024530 | -5.34496 | -3.80939 | -2.1557 | -2.1557 | -4.01426 | -2.8554 | Slmo1 |
| ENSMUSG00000062327 | -7.8255 | -6.02416 | -2.56913 | -2.56913 | -4.01595 | -2.85568 | T |
| ENSMUSG00000051184 | -5.16778 | -4.13422 | -2.27382 | -2.27382 | -4.17522 | -3.01741 | Zfp524 |
| ENSMUSG00000024039 | -5.93557 | -4.57718 | -2.65772 | -2.65772 | -4.1882 | -3.03221 | Cbs |
| ENSMUSG00000039634 | -5.69932 | -4.51812 | -2.56913 | -2.56913 | -4.19826 | -3.04068 | Zfp189 |
| ENSMUSG00000062939 | -5.2859 | -4.42953 | -1.47651 | -1.47651 | -4.20142 | -3.04325 | Stat4 |
| ENSMUSG00000039126 | -4.87248 | -3.95704 | -1.65369 | -1.65369 | -4.23351 | -3.08192 | Prune2 |
| ENSMUSG00000031847 | -5.43355 | -4.42953 | -1.68322 | -1.68322 | -4.23614 | -3.08409 | 1700030J2 |
| ENSMUSG00000000861 | -5.52214 | -4.42953 | -2.71678 | -2.71678 | -4.31694 | -3.17201 | Bcl11a |
| ENSMUSG00000031342 | -6.02416 | -5.19731 | -3.21879 | -3.21879 | -4.38828 | -3.24889 | Gpm6b |
| ENSMUSG00000037279 | -5.69932 | -4.54765 | -2.21476 | -2.21476 | -4.40365 | -3.2604 | Ovol2 |
| ENSMUSG00000024409 | -5.84697 | -4.07516 | -1.8604 | -1.8604 | -4.40621 | -3.26193 | Psors1c2 |
| ENSMUSG00000031160 | -6.93959 | -6.05369 | -3.54362 | -3.54362 | -4.42202 | -3.27557 | Eras |
| ENSMUSG00000026824 | -5.07919 | -4.10469 | -2.00805 | -2.00805 | -4.42939 | -3.28069 | Kcnj3 |
| ENSMUSG00000076441 | -5.31543 | -4.51812 | -2.48053 | -2.48053 | -4.44352 | -3.29149 | Ass1 |
| ENSMUSG00000028680 | -5.52214 | -4.63624 | -2.80537 | -2.80537 | -4.44745 | -3.29451 | Plk3 |
| ENSMUSG00000029162 | -5.25637 | -4.42953 | -2.74631 | -2.74631 | -4.48335 | -3.33522 | Khk |
| ENSMUSG00000022429 | -5.61073 | -4.75436 | -2.18523 | -2.18523 | -4.4973 | -3.34747 | Dmc1 |
| ENSMUSG00000015843 | -6.49664 | -4.90201 | -3.21879 | -3.21879 | -4.54924 | -3.40687 | Rxrg |
| ENSMUSG00000003657 | -5.66979 | -4.78389 | -2.451 | -2.451 | -4.57889 | -3.43536 | Calb2 |
| ENSMUSG00000055401 | -5.40402 | -4.48859 | -2.36241 | -2.36241 | -4.57927 | -3.43536 | Fbxo6 |
| ENSMUSG00000056367 | -5.81744 | -4.90201 | -2.74631 | -2.74631 | -4.6251 | -3.48316 | Actr3b |
| ENSMUSG00000061802 | -5.72885 | -4.84295 | -2.59866 | -2.59866 | -4.62511 | -3.48316 | Armc4 |
| ENSMUSG00000019880 | -5.66979 | -4.42953 | -2.74631 | -2.74631 | -4.73442 | -3.59994 | Rspo3 |
| ENSMUSG00000038065 | -5.75838 | -5.02013 | -3.39597 | -3.39597 | -4.76236 | -3.62507 | Mturn |
| ENSMUSG00000024697 | -5.61073 | -4.66577 | -2.89396 | -2.89396 | -4.8092 | -3.6811 | Gna14 |
| ENSMUSG00000032577 | -5.75838 | -4.60671 | -2.86443 | -2.86443 | -4.81898 | -3.68736 | Mapkapk3 |
| ENSMUSG00000030851 | -5.66979 | -4.93154 | -2.92349 | -2.92349 | -4.83959 | -3.7064 | Ldhc |
| ENSMUSG00000037171 | -6.91006 | -4.25235 | -3.07114 | -3.07114 | -4.84628 | -3.71262 | Nodal |
| ENSMUSG00000006442 | -3.24832 | -2.18523 | -1.24027 | -1.24027 | -4.87159 | -3.73667 | Srm |
| ENSMUSG00000058806 | -5.93557 | -4.78389 | -3.07114 | -3.07114 | -4.87459 | -3.7384 | Col13a1 |
| ENSMUSG00000032415 | -5.90603 | -5.07919 | -3.27785 | -3.27785 | -4.90636 | -3.77671 | Ube2cbp |
| ENSMUSG00000068302 | -6.37852 | -4.54765 | -3.48456 | -3.48456 | -4.92327 | -3.79607 | Noto |
| ENSMUSG00000033837 | -7.79597 | -2.59866 | -1.62416 | -1.62416 | -4.93899 | -3.80963 | Foxh1 |
| ENSMUSG00000030630 | -5.72885 | -4.93154 | -3.18926 | -3.18926 | -4.95751 | -3.83094 | Fah |
| ENSMUSG00000039001 | -2.42147 | -1.65369 | -1.00403 | -1.00403 | -5.00531 | -3.88261 | Rps21 |
| ENSMUSG00000032446 | -7.17583 | -6.52617 | -3.7208 | -3.7208 | -5.03852 | -3.91323 | Eomes |
| ENSMUSG00000031162 | -6.851 | -5.22684 | -3.51409 | -3.51409 | -5.07614 | -3.94891 | Gata1 |

| | | | | | | | |
|---|---|---|---|---|---|---|---|
| ENSMUSG00000060019 | -2.48053 | -1.71275 | -1.06309 | -1.06309 | -5.10622 | -3.97704 | Gm10073 |
| ENSMUSG00000018698 | -10.0698 | -9.50872 | -4.31141 | -4.31141 | -5.25414 | -4.12567 | Lhx1 |
| ENSMUSG00000023800 | -6.31946 | -5.46308 | -3.98657 | -3.98657 | -5.29969 | -4.16817 | Tiam2 |
| ENSMUSG00000021913 | -6.37852 | -5.46308 | -4.34094 | -4.34094 | -5.4079 | -4.28772 | Ogdhl |
| ENSMUSG00000034336 | -7.26442 | -6.46711 | -4.25235 | -4.25235 | -5.41913 | -4.29616 | Ina |
| ENSMUSG00000021018 | -3.21879 | -2.18523 | -1.35839 | -1.35839 | -5.43881 | -4.31662 | Polr2h |
| ENSMUSG00000020407 | -5.46308 | -3.45503 | -1.8604 | -1.8604 | -5.50077 | -4.37682 | Upp1 |
| ENSMUSG00000029337 | -8.97717 | -8.47516 | -2.33288 | -2.33288 | -5.50759 | -4.38182 | Fgf5 |
| ENSMUSG00000061524 | -6.11275 | -5.43355 | -4.51812 | -4.51812 | -5.53753 | -4.40825 | Zic2 |
| ENSMUSG00000034892 | -2.33288 | -1.68322 | -1.09262 | -1.09262 | -5.61336 | -4.48319 | Rps29 |
| ENSMUSG00000067860 | -7.70738 | -7.05771 | -5.84697 | -5.84697 | -5.66103 | -4.52173 | Zic3 |
| ENSMUSG00000025219 | -7.94362 | -7.29395 | -5.93557 | -5.93557 | -5.88176 | -4.75162 | Fgf8 |
| ENSMUSG00000019970 | -6.99865 | -6.23087 | -4.84295 | -4.84295 | -5.91513 | -4.78738 | Sgk1 |
| ENSMUSG00000041703 | -6.851 | -6.11275 | -5.10872 | -5.10872 | -5.93468 | -4.80169 | Zic5 |
| ENSMUSG00000051354 | -7.53019 | -6.70335 | -5.43355 | -5.43355 | -6.17415 | -5.05799 | Samd3 |
| ENSMUSG00000012848 | -2.39194 | -1.80134 | -1.24027 | -1.24027 | -6.34445 | -5.24429 | Rps5 |
| ENSMUSG00000071235 | -8.26845 | -7.76644 | -4.07516 | -4.07516 | -6.7231 | -5.65302 | Vrtn |
| ENSMUSG00000024406 | -7.97315 | -6.99865 | -5.9651 | -5.9651 | -6.76748 | -5.69392 | Pou5f1 |
| ENSMUSG00000032494 | -9.95167 | -9.0067 | -7.11677 | -7.11677 | -7.15986 | -6.05197 | Tdgf1 |
| ENSMUSG00000035202 | -2.98255 | -2.27382 | -1.65369 | -1.65369 | -7.1261 | -6.05197 | Lars2 |

s

e 95

(c(lb, ub))), or 0 if the CI crosses t

using Holm procedure

**d cZ.**

| Descriptio | Location | Family |
|---|---|---|
| collagen ty | Extracellula | other |
| solute carr | Plasma Me | transporter |
| growth fac | Cytoplasm | other |
| inhibitor o | Nucleus | transcription regulator |
| four and a | Cytoplasm | other |
| insulin like | Extracellula | other |
| ISL LIM ho | Nucleus | transcription regulator |
| transcriptio | Other | other |
| brain expr | Cytoplasm | other |
| insulin like | Extracellula | growth factor |
| glypican 3 | Plasma Me | other |
| BCL2 inter | Cytoplasm | other |
| claudin 4 | Plasma Me | transmembrane receptor |
| follistatin l | Extracellula | other |
| | Other | other |
| T-box 3 | Nucleus | transcription regulator |
| podocalyx | Plasma Me | other |
| coagulatio | Plasma Me | G-protein coupled receptor |
| fibronectin | Extracellula | enzyme |
| peptide YY | Extracellula | other |
| prominin 1 | Plasma Me | other |
| protogenin | Extracellula | other |
| retinoic ac | Nucleus | ligand-dependent nuclear receptor |
| cingulin | Plasma Me | other |
| docking pr | Plasma Me | other |
| reticulocal | Cytoplasm | other |
| midkine | Extracellula | growth factor |
| secreted p | Extracellula | other |
| Wnt family | Extracellula | cytokine |
| zinc finger | Nucleus | other |
| septin 2 | Cytoplasm | enzyme |
| transformi | Extracellula | growth factor |
| NK2 home | Nucleus | transcription regulator |
| solute carr | Extracellula | transporter |
| solute carr | Plasma Me | transporter |
| NK2 home | Nucleus | transcription regulator |
| tropomyos | Plasma Me | other |
| mannan-b | Extracellula | peptidase |
| microfibril | Extracellula | other |
| hes relatec | Nucleus | transcription regulator |
| NK2 home | Nucleus | transcription regulator |
| TIMP meta | Extracellula | other |
| ATPase se | Cytoplasm | enzyme |
| olfactome | Extracellula | other |
| ephrin A1 | Plasma Me | other |
| LDOC1, re | Nucleus | other |

| | | |
|---|---|---|
| chromosoı | Other | other |
| interleukin | Plasma Me | transmembrane receptor |
| espin | Cytoplasm | other |
| tetraspanii | Plasma Me | other |
| neuropilin | Plasma Me | transmembrane receptor |
| carboxype | Plasma Me | peptidase |
| NUAK fam | Nucleus | kinase |
| beta-1,4-N | Extracellula | enzyme |
| endothelir | Plasma Me | peptidase |
| anoctamin | Plasma Me | ion channel |
| prolyl 4-hy | Cytoplasm | transporter |
| keratin 19 | Cytoplasm | other |
| dihydropy | Cytoplasm | enzyme |
| Josephin c | Other | other |
| regulator c | Other | other |
| unc-13 ho | Cytoplasm | other |
| Meis home | Nucleus | transcription regulator |
| potassium | Plasma Me | ion channel |
| integrator | Nucleus | other |
| SRY-box 1 | Nucleus | transcription regulator |
| junction pl | Plasma Me | other |
| membrane | Plasma Me | peptidase |
| claudin 3 | Plasma Me | transmembrane receptor |
| ADP ribos | Cytoplasm | other |
| LRR bindir | Cytoplasm | transcription regulator |
| neuropilin | Plasma Me | other |
| basal cell a | Plasma Me | transmembrane receptor |
| roundabou | Plasma Me | transmembrane receptor |
| transferrin | Plasma Me | transporter |
| ornithine c | Cytoplasm | other |
| solute carr | Plasma Me | transporter |
| mucolipin | Plasma Me | ion channel |
| adenylate | Cytoplasm | kinase |
| endothelir | Plasma Me | transmembrane receptor |
| yippee like | Nucleus | other |
| paternally | Nucleus | kinase |
| paralemmi | Other | other |
| zinc finger | Nucleus | transcription regulator |
| crystallin a | Nucleus | other |
| olfactome | Cytoplasm | other |
| serine pro | Extracellula | peptidase |
| keratin 20 | Cytoplasm | other |
| lysine dem | Extracellula | enzyme |
| ladinin 1 | Extracellula | other |
| fibrinogen | Extracellula | other |
| neurexin II | Plasma Me | other |
| GATA binc | Nucleus | transcription regulator |
| phospholip | Other | phosphatase |
| serpin fam | Extracellula | other |
| procollage | Cytoplasm | enzyme |
| Fraser extr | Extracellula | other |
| zinc finger | Nucleus | transcription regulator |
| protein pr | Other | other |
| fibronectir | Cytoplasm | other |
| G protein- | Plasma Me | G-protein coupled receptor |
| SH3 doma | Plasma Me | other |
| epsin 3 | Cytoplasm | other |
| sterol O-a | Cytoplasm | enzyme |

| | | |
|---|---|---|
| SEC24 hor | Cytoplasm | transporter |
| canopy FG | Other | other |
| retrotransp | Other | other |
| cellular ret | Cytoplasm | transporter |
| pre-mRNA | Nucleus | kinase |
| serine pep | Extracellula | other |
| influenza v | Nucleus | other |
| dystroglyc | Plasma Me | transmembrane receptor |
| oxysterol b | Cytoplasm | transporter |
| monoglyce | Plasma Me | enzyme |
| nephronec | Plasma Me | other |
| Meis home | Nucleus | transcription regulator |
| transformi | Plasma Me | kinase |
| ATPase pla | Plasma Me | transporter |
| serine prot | Extracellula | peptidase |
| leucine ric | Other | other |
| LIM and SH | Cytoplasm | transporter |
| MUM1 like | Cytoplasm | other |
| unc-5 netr | Plasma Me | transmembrane receptor |
| coiled-coi | Other | other |
| RIKEN cDN | Cytoplasm | other |
| solute carr | Plasma Me | transporter |
| clusterin | Cytoplasm | other |
| ADAM me | Extracellula | peptidase |
| doublesex | Nucleus | transcription regulator |
| spectrin be | Cytoplasm | other |
| reproducti | Nucleus | transcription regulator |
| SH3 doma | Cytoplasm | enzyme |
| hexosamin | Cytoplasm | enzyme |
| peroxidasi | Extracellula | enzyme |
| insulin like | Plasma Me | transmembrane receptor |
| prolyl 4-hy | Cytoplasm | enzyme |
| CD63 mole | Plasma Me | other |
| cadherin 1 | Other | other |
| transmemb | Cytoplasm | other |
| zinc finger | Nucleus | transcription regulator |
| Kruppel lik | Nucleus | transcription regulator |
| LDL recept | Plasma Me | transmembrane receptor |
| ubiquitin s | Plasma Me | peptidase |
| protein tyr | Plasma Me | phosphatase |
| paralemmi | Plasma Me | other |
| 3'-phosph | Cytoplasm | enzyme |
| mannose r | Plasma Me | transmembrane receptor |
| nucleobind | Nucleus | other |
| ATP bindin | Plasma Me | transporter |
| zinc finger | Nucleus | transcription regulator |
| trans-golg | Other | other |
| frizzled cla | Plasma Me | G-protein coupled receptor |
| heterochro | Nucleus | other |
| histone de | Nucleus | transcription regulator |
| DENN dor | Cytoplasm | other |
| PWWP do | Nucleus | other |
| cadherin 1 | Other | other |
| human im | Nucleus | transcription regulator |
| cysteine ric | Cytoplasm | other |
| PR/SET do | Nucleus | transcription regulator |
| WAP four- | Extracellula | other |

| | | |
|---|---|---|
| EPH recep | Plasma Me | kinase |
| pleckstrin | Cytoplasm | other |
| spastic par | Plasma Me | other |
| GRAM dor | Other | other |
| tubulointe | Extracellula | transporter |
| tubulin be | Cytoplasm | other |
| dimethylar | Cytoplasm | enzyme |
| sorting ne: | Cytoplasm | other |
| cadherin 3 | Plasma Me | other |
| immunogl | Plasma Me | other |
| WNK lysin | Cytoplasm | kinase |
| vesicle ass | Plasma Me | transporter |
| Src homol | Cytoplasm | other |
| patatin like | Cytoplasm | enzyme |
| YEATS dor | Nucleus | transcription regulator |
| diacylglyce | Cytoplasm | enzyme |
| coiled-coi | Extracellula | other |
| SHC adapt | Cytoplasm | other |
| endoplasn | Cytoplasm | enzyme |
| solute carr | Plasma Me | transporter |
| suppressic | Plasma Me | peptidase |
| keratin 18 | Cytoplasm | other |
| IQ motif c | Extracellula | other |
| CAS1 dom | Cytoplasm | enzyme |
| solute carr | Plasma Me | transporter |
| wntless W | Cytoplasm | other |
| collagen ty | Extracellula | other |
| hyaluronar | Plasma Me | enzyme |
| beta-1,4-g | Cytoplasm | enzyme |
| 1-aminocy | Other | enzyme |
| zinc finger | Nucleus | transcription regulator |
| KDEL endo | Cytoplasm | transporter |
| phosphory | Cytoplasm | kinase |
| ras homol | Cytoplasm | enzyme |
| solute carr | Plasma Me | transporter |
| growth dif | Extracellula | growth factor |
| transmem | Cytoplasm | transporter |
| FK506 binc | Cytoplasm | enzyme |
| calpain 6 | Cytoplasm | peptidase |
| fatty acid c | Plasma Me | enzyme |
| collagen ty | Extracellula | other |
| tumor nec | Plasma Me | other |
| osteopetro | Cytoplasm | other |
| TEA doma | Nucleus | transcription regulator |
| annexin A | Plasma Me | other |
| crumbs 2, | Extracellula | other |
| fibrillin 2 | Extracellula | other |
| unc-5 netr | Plasma Me | transmembrane receptor |
| serine/thre | Cytoplasm | kinase |
| serine inco | Plasma Me | transporter |
| myosin he | Cytoplasm | enzyme |
| nudix hydr | Cytoplasm | phosphatase |
| tetraspani | Other | other |
| von Willeb | Extracellula | other |
| membrane | Cytoplasm | enzyme |
| IKBKB inte | Cytoplasm | other |
| Rho GTPas | Cytoplasm | other |
| phosphofr | Cytoplasm | kinase |

| | | |
|---|---|---|
| chromoso | Nucleus | other |
| matrix met | Extracellul | peptidase |
| CXXC fing | Cytoplasm | other |
| dimethylar | Cytoplasm | enzyme |
| MN1 proto | Nucleus | other |
| SMC5-SM | Nucleus | transcription regulator |
| cytochrom | Cytoplasm | enzyme |
| tripartite n | Cytoplasm | other |
| adaptor re | Nucleus | other |
| latexin | Cytoplasm | other |
| protein ph | Nucleus | phosphatase |
| delta like r | Extracellul | other |
| fucosyltrar | Cytoplasm | enzyme |
| pleiotroph | Extracellul | growth factor |
| protein kir | Cytoplasm | kinase |
| T-box 20 | Nucleus | transcription regulator |
| junction m | Nucleus | transcription regulator |
| integrin su | Plasma Me | other |
| DENN dor | Cytoplasm | other |
| transmeml | Cytoplasm | other |
| mitochonc | Cytoplasm | other |
| erbb2 inte | Cytoplasm | other |
| kirre like n | Plasma Me | other |
| erb-b2 rec | Plasma Me | kinase |
| solute carr | Cytoplasm | transporter |
| aldehyde ( | Cytoplasm | enzyme |
| SUN doma | Other | other |
| desmin | Cytoplasm | other |
| hephaestir | Plasma Me | transporter |
| ring finger | Nucleus | enzyme |
| transformi | Extracellul | growth factor |
| transmeml | Other | other |
| apelin rece | Extracellul | other |
| ubiquitin a | Other | enzyme |
| Rho guani | Cytoplasm | other |
| CD34 mol | Plasma Me | other |
| double ho | Nucleus | other |
| zinc finger | Nucleus | transcription regulator |
| alkaline ph | Plasma Me | phosphatase |
| shisa famil | Other | other |
| RUN and ( | Cytoplasm | other |
| zinc finger | Nucleus | transcription regulator |
| family with | Other | other |
| zinc finger | Other | other |
| CUB doma | Plasma Me | other |
| transcriptic | Nucleus | transcription regulator |
| zinc finger | Other | other |
| zinc finger | Nucleus | transcription regulator |
| LDL recept | Plasma Me | transmembrane receptor |
| poly(A) bir | Other | other |
| ArfGAP wi | Cytoplasm | other |
| nuclear re | Nucleus | ligand-dependent nuclear receptor |
| transmeml | Other | other |
| small inte | Cytoplasm | other |
| zinc finger | Other | other |
| BEN doma | Cytoplasm | other |
| FERM dom | Cytoplasm | other |
| scavenger | Plasma Me | transmembrane receptor |

| | | |
|---|---|---|
| growth arr | Cytoplasm | other |
| phosphoir | Cytoplasm | other |
| Ras associ | Cytoplasm | transporter |
| C1GALT1 s | Cytoplasm | other |
| glutathion | Other | enzyme |
| interleukin | Extracellula | cytokine |
| A-kinase a | Plasma Me | other |
| RUNX1 tra | Nucleus | transcription regulator |
| MORC fam | Nucleus | other |
| folliculin ir | Cytoplasm | other |
| innate imn | Other | other |
| chromodo | Nucleus | enzyme |
| pleckstrin | Cytoplasm | other |
| male speci | Nucleus | other |
| suppresso | Cytoplasm | phosphatase |
| protein kir | Cytoplasm | kinase |
| expressed | Other | other |
| latent tran | Extracellula | other |
| TGF-beta . | Cytoplasm | other |
| SS18L1, nE | Nucleus | transcription regulator |
| coiled-coi | Plasma Me | other |
| collagen ty | Extracellula | other |
| melanoreg | Cytoplasm | other |
| occludin | Plasma Me | other |
| fukutin | Extracellula | other |
| advanced | Plasma Me | transmembrane receptor |
| rhomboid | Cytoplasm | other |
| zinc finger | Nucleus | transporter |
| fucosyltrar | Cytoplasm | enzyme |
| transmeml | Other | other |
| double ho | Nucleus | other |
| tripartite n | Cytoplasm | transcription regulator |
| FCH and d | Other | other |
| double ho | Nucleus | other |
| fibrinogen | Nucleus | other |
| family with | Cytoplasm | enzyme |
| host cell fa | Nucleus | transcription regulator |
| GIPC PDZ | Cytoplasm | other |
| TATA-box | Nucleus | transcription regulator |
| retinoic ac | Plasma Me | other |
| carboxype | Extracellula | peptidase |
| aldolase, fi | Cytoplasm | enzyme |
| collagen ty | Extracellula | other |
| hyaluronai | Extracellula | other |
| deoxyribo | Cytoplasm | enzyme |
| DAZ intera | Cytoplasm | other |
| transmeml | Other | other |
| F-box and | Other | enzyme |
| ectonuclec | Cytoplasm | enzyme |
| beta-1,4-g | Cytoplasm | enzyme |
| primase ar | Extracellula | enzyme |
| synaptotag | Cytoplasm | transporter |
| coronin 2A | Cytoplasm | other |
| MAGE fam | Plasma Me | other |
| caspase 7 | Cytoplasm | peptidase |
| isoprenoic | Cytoplasm | enzyme |
| zinc finger | Nucleus | transcription regulator |

| | | |
|---|---|---|
| opioid rela | Plasma Me | G-protein coupled receptor |
| peroxisom | Cytoplasm | other |
| poliovirus | Plasma Me | other |
| growth arr | Nucleus | other |
| serpin fam | Extracellul | other |
| regulator ( | Plasma Me | other |
| tripartite n | Cytoplasm | enzyme |
| claudin 11 | Plasma Me | other |
| HESX hom | Nucleus | transcription regulator |
| fms relate | Plasma Me | kinase |
| Mix paired | Nucleus | transcription regulator |
| semaphori | Plasma Me | other |
| regulator ( | Cytoplasm | other |
| stathmin 2 | Plasma Me | other |
| delta like ( | Extracellul | other |
| elongation | Nucleus | transcription regulator |
| yippee like | Other | other |
| coiled-coi | Other | other |
| apoptosis | Cytoplasm | enzyme |
| 3-hydroxy | Cytoplasm | enzyme |
| pipecolic a | Cytoplasm | enzyme |
| chordin | Extracellul | other |
| proenkeph | Extracellul | other |
| primary cil | Cytoplasm | other |
| olfactome | Cytoplasm | other |
| membrane | Plasma Me | other |
| ADAM me | Plasma Me | peptidase |
| zinc finger | Cytoplasm | other |
| cerberus 1 | Extracellul | cytokine |
| cytochrom | Cytoplasm | enzyme |
| receptor ty | Plasma Me | kinase |
| LysM dom | Other | other |
| phosphod | Cytoplasm | enzyme |
| DLC1 Rho | Cytoplasm | other |
| nucleopor | Other | other |
| cysteine ar | Other | other |
| ribosomal | Nucleus | enzyme |
| coiled-coi | Cytoplasm | other |
| serine acti | Extracellul | other |
| retrotransp | Plasma Me | other |
| follistatin | Extracellul | other |
| histone clu | Nucleus | other |
| zinc finger | Nucleus | transcription regulator |
| cripto, FRL | Extracellul | other |
| polycystin | Plasma Me | ion channel |
| netrin 1 | Extracellul | other |
| bone morp | Extracellul | growth factor |
| shisa famil | Other | other |
| family with | Plasma Me | transmembrane receptor |
| family with | Nucleus | enzyme |
| coiled-coi | Cytoplasm | other |
| spalt like t | Nucleus | transcription regulator |
| glypican 4 | Plasma Me | transmembrane receptor |
| armadillo r | Cytoplasm | other |
| DAB1, reel | Cytoplasm | other |
| sterile alph | Other | other |
| HYLS1, cen | Nucleus | other |
| neurotrop | Plasma Me | kinase |

| | | |
|---|---|---|
| leucine ric | Plasma Me | transmembrane receptor |
| Rho guani | Plasma Me | other |
| Sp5 transc | Nucleus | transcription regulator |
| apelin rece | Plasma Me | G-protein coupled receptor |
| chromoso | Other | other |
| slit guidan | Extracellul | other |
| keratin 22 | Other | other |
| optineurin | Cytoplasm | other |
| dihydropy | Cytoplasm | enzyme |
| TEF, PAR b | Nucleus | transcription regulator |
| cadherin 2 | Plasma Me | other |
| early grow | Nucleus | transcription regulator |
| carbonic a | Cytoplasm | enzyme |
| contactin a | Plasma Me | other |
| coiled-coi | Cytoplasm | other |
| forkhead b | Nucleus | transcription regulator |
| ADP ribos | Nucleus | enzyme |
| chromoso | Plasma Me | other |
| kinesin fan | Cytoplasm | enzyme |
| GATA binc | Nucleus | transcription regulator |
| PRELI dom | Cytoplasm | transporter |
| T-box trar | Nucleus | transcription regulator |
| zinc finger | Other | other |
| cystathion | Cytoplasm | enzyme |
| zinc finger | Nucleus | transcription regulator |
| signal tran | Nucleus | transcription regulator |
| prune hon | Cytoplasm | other |
| chromoso | Nucleus | other |
| B cell CLL/ | Nucleus | transcription regulator |
| glycoprote | Plasma Me | other |
| ovo like zi | Nucleus | transcription regulator |
| psoriasis s | Extracellul | other |
| ES cell exp | Plasma Me | enzyme |
| potassium | Plasma Me | ion channel |
| argininosu | Cytoplasm | enzyme |
| polo like k | Nucleus | kinase |
| ketohexok | Cytoplasm | kinase |
| DNA meio | Nucleus | enzyme |
| retinoid X | Nucleus | ligand-dependent nuclear receptor |
| calbindin 2 | Cytoplasm | other |
| F-box pro | Cytoplasm | enzyme |
| ARP3 actir | Cytoplasm | other |
| armadillo r | Extracellul | other |
| R-spondin | Extracellul | kinase |
| maturin, n | Other | other |
| G protein s | Plasma Me | enzyme |
| mitogen-a | Nucleus | kinase |
| lactate del | Cytoplasm | enzyme |
| nodal grov | Extracellul | growth factor |
| spermidine | Cytoplasm | enzyme |
| collagen ty | Plasma Me | other |
| ubiquitin p | Other | other |
| notochord | Other | other |
| forkhead b | Nucleus | transcription regulator |
| fumarylace | Cytoplasm | enzyme |
| ribosomal | Cytoplasm | other |
| eomesode | Nucleus | transcription regulator |
| GATA binc | Nucleus | transcription regulator |

| | | |
|---|---|---|
| ribosomal | Other | other |
| LIM home | Nucleus | transcription regulator |
| T cell lymp | Cytoplasm | enzyme |
| oxoglutara | Other | enzyme |
| internexin | Cytoplasm | other |
| RNA polyr | Nucleus | enzyme |
| uridine ph | Cytoplasm | enzyme |
| fibroblast | Extracellula | growth factor |
| Zic family | Nucleus | transcription regulator |
| ribosomal | Cytoplasm | other |
| Zic family | Nucleus | transcription regulator |
| fibroblast | Extracellula | growth factor |
| serum/glu | Cytoplasm | kinase |
| Zic family | Nucleus | transcription regulator |
| sterile alpl | Other | other |
| ribosomal | Cytoplasm | other |
| vertebrae | Other | other |
| POU class | Nucleus | transcription regulator |
| teratocarci | Extracellula | growth factor |
| leucyl-tRN | Cytoplasm | enzyme |

### Differentially expressed genes (DEGs) of cells

**Instructions:**

- **p_val :** p-value (unadjusted)
- **avg_logFC :** log fold-chage of the average expression between the more highly expressed in the given cluster.
- **pct.1 :** The ratio of expression in the given cell cluster
- **pct.2 :** The ratio of expression in the left cell clusters from E9.5-E10.
- **p_val_adj :** Adjusted p-value, based on bonferroni correction using

| | p_val | avg_logFC | pct.1 | pct.2 | p_val_adj |
|---|---|---|---|---|---|
| Adgrv1 | 2.75E-86 | 0.332504264 | 1 | 0.8 | 3.54E-82 |
| Tubb3 | 7.11E-79 | 1.06497316 | 1 | 0.65 | 9.16E-75 |
| Lhfp | 3.31E-71 | 0.97818572 | 1 | 0.893 | 4.27E-67 |
| Ngfr | 8.54E-62 | 2.161460607 | 1 | 0.386 | 1.10E-57 |
| Adcyap1r1 | 2.00E-56 | 1.357677732 | 1 | 0.343 | 2.58E-52 |
| Cdh11 | 2.43E-55 | 0.536006304 | 1 | 0.979 | 3.13E-51 |
| Nes | 1.08E-52 | 1.415210191 | 1 | 0.407 | 1.40E-48 |
| Scube1 | 3.94E-49 | 0.557286168 | 1 | 0.743 | 5.08E-45 |
| Celsr2 | 1.15E-48 | 0.796150435 | 1 | 0.607 | 1.48E-44 |
| Zeb2 | 8.04E-48 | 0.296209354 | 1 | 0.636 | 1.04E-43 |
| Ncam1 | 5.11E-46 | 0.686803536 | 1 | 0.636 | 6.59E-42 |
| Plxna2 | 1.46E-42 | 0.470972222 | 1 | 0.943 | 1.88E-38 |
| Lrrn1 | 3.53E-42 | 2.349897864 | 1 | 0.45 | 4.55E-38 |
| Ednrb | 7.60E-42 | 1.642842862 | 1 | 0.393 | 9.79E-38 |
| Slit1 | 1.46E-41 | 1.972613156 | 1 | 0.293 | 1.88E-37 |
| Jam2 | 2.76E-41 | 0.498693043 | 0.75 | 0.364 | 3.55E-37 |
| Sema5b | 2.99E-40 | 1.072213591 | 1 | 0.321 | 3.86E-36 |
| Phox2b | 9.21E-39 | 1.352425399 | 1 | 0.221 | 1.19E-34 |
| Fgfr3 | 2.82E-38 | 0.89253273 | 1 | 0.943 | 3.64E-34 |
| Rfx4 | 1.14E-36 | 1.214980803 | 1 | 0.186 | 1.47E-32 |
| Srgap3 | 3.03E-36 | 0.56563609 | 1 | 0.357 | 3.91E-32 |
| Notch1 | 3.24E-33 | 0.343717021 | 1 | 0.7 | 4.18E-29 |
| Rgma | 5.48E-33 | 0.615092946 | 1 | 0.307 | 7.06E-29 |
| Entpd7 | 1.43E-32 | 0.304954739 | 1 | 0.736 | 1.85E-28 |
| Nrcam | 1.81E-32 | 0.354712394 | 1 | 0.55 | 2.34E-28 |
| Elavl4 | 2.06E-32 | 0.976993169 | 0.75 | 0.336 | 2.65E-28 |
| Dll3 | 5.22E-32 | 1.404919654 | 1 | 0.207 | 6.72E-28 |
| Osbpl7 | 9.71E-31 | 0.427878263 | 0.5 | 0.65 | 1.25E-26 |
| Olfml3 | 1.20E-30 | 0.66237169 | 1 | 0.486 | 1.54E-26 |
| Adgrl2 | 3.16E-30 | 0.669991738 | 1 | 0.879 | 4.08E-26 |
| Igsf9b | 1.70E-29 | 1.087748239 | 1 | 0.336 | 2.19E-25 |
| Sulf2 | 6.01E-29 | 1.02238366 | 1 | 0.814 | 7.74E-25 |
| Ptpru | 1.00E-28 | 0.443451289 | 0.75 | 0.443 | 1.29E-24 |
| Tulp4 | 1.23E-28 | 0.274581706 | 1 | 0.986 | 1.59E-24 |
| Tox2 | 2.16E-28 | 0.641539642 | 1 | 0.379 | 2.78E-24 |
| Tppp3 | 5.74E-28 | 1.237862252 | 1 | 0.514 | 7.40E-24 |
| Gm4707 | 1.14E-27 | 0.39570931 | 1 | 0.993 | 1.47E-23 |
| Mn1 | 1.56E-27 | 0.868793361 | 1 | 0.771 | 2.01E-23 |
| Spon1 | 1.94E-27 | 1.118815071 | 1 | 0.557 | 2.50E-23 |
| Ncor2 | 3.64E-27 | 0.322506426 | 1 | 0.864 | 4.69E-23 |
| Vangl2 | 6.71E-27 | 1.103737456 | 1 | 0.95 | 8.65E-23 |
| Ntn1 | 9.12E-27 | 1.262794958 | 1 | 0.3 | 1.18E-22 |
| Cdc42bpb | 2.05E-26 | 0.391070532 | 1 | 0.714 | 2.64E-22 |
| Sulf1 | 3.52E-26 | 0.625163192 | 1 | 0.464 | 4.54E-22 |

| | | | | | |
|---|---|---|---|---|---|
| Nell2 | 4.23E-26 | 1.298843863 | 1 | 0.2 | 5.45E-22 |
| Ckb | 1.31E-25 | 3.394114532 | 1 | 0.979 | 1.68E-21 |
| Lfng | 2.08E-25 | 0.808683733 | 1 | 0.129 | 2.68E-21 |
| 2310067B1 | 2.53E-25 | 0.642866974 | 1 | 0.764 | 3.27E-21 |
| Syt11 | 4.26E-25 | 1.101723775 | 1 | 0.8 | 5.49E-21 |
| Map1b | 5.12E-25 | 1.457849398 | 1 | 0.836 | 6.60E-21 |
| 5031439G | 8.45E-25 | 0.335826363 | 1 | 0.686 | 1.09E-20 |
| Slit3 | 9.85E-25 | 0.669614187 | 1 | 0.364 | 1.27E-20 |
| Sfrp1 | 1.23E-24 | 1.169132983 | 1 | 0.664 | 1.59E-20 |
| Zfp11 | 1.31E-24 | 0.308491338 | 1 | 0.964 | 1.69E-20 |
| Zc3h12c | 1.44E-24 | 0.54693957 | 1 | 0.429 | 1.86E-20 |
| Stxbp1 | 2.41E-24 | 0.557928373 | 0.75 | 0.407 | 3.10E-20 |
| Prtg | 4.23E-24 | 1.149314772 | 1 | 0.979 | 5.45E-20 |
| Atp8a1 | 4.82E-24 | 0.6117297 | 0.75 | 0.464 | 6.21E-20 |
| Map4k4 | 4.96E-24 | 0.438997724 | 1 | 0.979 | 6.39E-20 |
| Itsn1 | 2.09E-23 | 0.255470345 | 1 | 1 | 2.70E-19 |
| Crb2 | 2.75E-23 | 0.728372369 | 1 | 0.557 | 3.55E-19 |
| Dtx4 | 3.02E-23 | 0.786140264 | 1 | 0.564 | 3.89E-19 |
| Reep1 | 4.26E-23 | 1.043218888 | 1 | 0.686 | 5.50E-19 |
| Adgrg1 | 5.92E-23 | 1.407430375 | 1 | 0.764 | 7.63E-19 |
| Cacna1g | 9.33E-23 | 0.290246682 | 0.75 | 0.2 | 1.20E-18 |
| Ddr1 | 1.09E-22 | 1.188121814 | 1 | 0.664 | 1.41E-18 |
| Pcdhgc3 | 1.51E-22 | 0.31145538 | 0.75 | 0.414 | 1.95E-18 |
| Eml1 | 1.95E-22 | 0.56091682 | 1 | 0.864 | 2.52E-18 |
| Ube4a | 2.40E-22 | 0.467974121 | 1 | 0.85 | 3.09E-18 |
| Igdcc3 | 2.83E-22 | 1.878983438 | 1 | 0.686 | 3.64E-18 |
| Uggt1 | 4.54E-22 | 0.785242677 | 1 | 0.993 | 5.85E-18 |
| Nr2f1 | 9.15E-22 | 0.286818448 | 1 | 0.221 | 1.18E-17 |
| Fabp7 | 1.23E-21 | 1.288585778 | 1 | 0.086 | 1.58E-17 |
| Pou2f1 | 1.50E-21 | 0.261503449 | 1 | 0.907 | 1.93E-17 |
| Prkar2a | 1.52E-21 | 0.448712294 | 0.75 | 0.864 | 1.96E-17 |
| Trappc8 | 3.30E-21 | 0.367020629 | 0.5 | 0.821 | 4.25E-17 |
| Sorcs2 | 4.37E-21 | 0.420834711 | 1 | 0.443 | 5.64E-17 |
| Maml3 | 4.87E-21 | 0.258270146 | 1 | 0.957 | 6.28E-17 |
| Col22a1 | 6.30E-21 | 0.26659184 | 1 | 0.814 | 8.12E-17 |
| Brsk2 | 9.79E-21 | 0.258534055 | 0.75 | 0.2 | 1.26E-16 |
| Sept3 | 2.86E-20 | 0.78049733 | 1 | 0.629 | 3.68E-16 |
| Nuak2 | 4.65E-20 | 0.722495817 | 1 | 0.414 | 5.99E-16 |
| Cdh2 | 5.05E-20 | 1.16374382 | 1 | 0.993 | 6.51E-16 |
| Vim | 6.55E-20 | 1.70750641 | 1 | 0.893 | 8.44E-16 |
| Pknox2 | 7.19E-20 | 0.521076854 | 1 | 0.379 | 9.27E-16 |
| Nkx6-1 | 7.61E-20 | 1.011659188 | 1 | 0.171 | 9.81E-16 |
| Slc45a3 | 9.95E-20 | 0.546604189 | 0.75 | 0.343 | 1.28E-15 |
| Tcf12 | 1.06E-19 | 0.311976702 | 1 | 0.993 | 1.37E-15 |
| Tmem219 | 1.08E-19 | 0.676761841 | 1 | 0.671 | 1.39E-15 |
| Ptch1 | 1.37E-19 | 0.518448495 | 1 | 0.85 | 1.76E-15 |
| Arhgap35 | 1.60E-19 | 0.358345661 | 1 | 0.986 | 2.06E-15 |
| Prom1 | 1.90E-19 | 1.316587149 | 1 | 1 | 2.45E-15 |
| Rsg1 | 1.98E-19 | 0.269737731 | 0.75 | 0.707 | 2.56E-15 |
| Jam3 | 2.07E-19 | 1.555313521 | 1 | 0.979 | 2.67E-15 |
| Ptpn21 | 6.74E-19 | 0.26853097 | 0.5 | 0.6 | 8.69E-15 |
| Mfng | 6.94E-19 | 0.833268358 | 1 | 0.236 | 8.95E-15 |
| Rnf123 | 9.04E-19 | 0.318837789 | 1 | 0.714 | 1.17E-14 |
| Zswim4 | 1.01E-18 | 0.612078478 | 1 | 0.5 | 1.30E-14 |
| Scrn1 | 1.32E-18 | 0.364158387 | 1 | 0.229 | 1.70E-14 |
| Rcor1 | 1.67E-18 | 0.303003245 | 0.25 | 0.793 | 2.16E-14 |
| Ptprs | 2.36E-18 | 1.032995468 | 1 | 0.986 | 3.05E-14 |
| Fez1 | 2.68E-18 | 0.751420945 | 1 | 0.214 | 3.46E-14 |

| | | | | | |
|---|---|---|---|---|---|
| Myo19 | 3.43E-18 | 0.332248604 | 0.75 | 0.707 | 4.43E-14 |
| Sall1 | 4.95E-18 | 0.464752757 | 1 | 0.6 | 6.38E-14 |
| Hoxd3 | 8.09E-18 | 0.476585489 | 1 | 0.079 | 1.04E-13 |
| Tmem47 | 8.43E-18 | 0.814536801 | 1 | 0.7 | 1.09E-13 |
| Tfdp2 | 9.85E-18 | 0.365083741 | 1 | 0.971 | 1.27E-13 |
| Itgb8 | 9.92E-18 | 0.522002386 | 1 | 0.436 | 1.28E-13 |
| Pea15a | 1.10E-17 | 1.264458673 | 1 | 0.929 | 1.42E-13 |
| Ttyh2 | 1.21E-17 | 0.311450649 | 0.5 | 0.636 | 1.55E-13 |
| Nsun3 | 1.35E-17 | 0.316307219 | 0.5 | 0.693 | 1.74E-13 |
| Slc35e2 | 2.82E-17 | 0.364886964 | 1 | 0.993 | 3.63E-13 |
| Hes5 | 3.36E-17 | 1.042221998 | 0.75 | 0.071 | 4.33E-13 |
| E130308A1 | 4.00E-17 | 0.335117103 | 1 | 0.95 | 5.15E-13 |
| Pik3c2b | 5.34E-17 | 0.328886414 | 0.75 | 0.85 | 6.89E-13 |
| Cdh22 | 5.49E-17 | 0.416566736 | 0.75 | 0.121 | 7.07E-13 |
| Gpc1 | 7.33E-17 | 0.514939426 | 1 | 0.779 | 9.44E-13 |
| 1700106J1 | 7.98E-17 | 1.765443778 | 0.75 | 0.164 | 1.03E-12 |
| Cercam | 1.31E-16 | 0.623381846 | 1 | 0.864 | 1.69E-12 |
| Prmt10 | 1.36E-16 | 0.46793255 | 1 | 0.814 | 1.75E-12 |
| Ankrd11 | 1.73E-16 | 0.4199166 | 1 | 1 | 2.22E-12 |
| Msn | 1.77E-16 | 0.71404767 | 1 | 0.779 | 2.28E-12 |
| Lrp1 | 1.83E-16 | 0.422177499 | 1 | 0.943 | 2.35E-12 |
| Dgcr2 | 2.01E-16 | 0.697733002 | 1 | 0.993 | 2.59E-12 |
| Tnfaip6 | 2.20E-16 | 0.549012979 | 0.75 | 0.214 | 2.84E-12 |
| Ly6h | 2.93E-16 | 0.644639025 | 1 | 0.579 | 3.78E-12 |
| Tyw1 | 4.36E-16 | 0.413211191 | 1 | 0.95 | 5.62E-12 |
| Fbln2 | 4.63E-16 | 0.343537481 | 0.75 | 0.414 | 5.97E-12 |
| Rassf4 | 5.30E-16 | 0.719460462 | 1 | 0.3 | 6.83E-12 |
| Akna | 5.69E-16 | 0.408468118 | 0.75 | 0.343 | 7.33E-12 |
| Nr6a1 | 6.18E-16 | 0.33771041 | 1 | 0.879 | 7.96E-12 |
| Ephb1 | 7.81E-16 | 0.271745126 | 0.75 | 0.214 | 1.01E-11 |
| Slc38a6 | 8.10E-16 | 0.260560103 | 1 | 0.993 | 1.04E-11 |
| Zgrf1 | 9.29E-16 | 0.250361453 | 0.75 | 0.714 | 1.20E-11 |
| Ndufb2 | 1.44E-15 | 0.264212249 | 1 | 1 | 1.86E-11 |
| Ep400 | 1.55E-15 | 0.636688109 | 1 | 1 | 2.00E-11 |
| Jade1 | 2.28E-15 | 0.293738695 | 1 | 0.864 | 2.94E-11 |
| Qsox2 | 2.58E-15 | 0.636307781 | 1 | 0.85 | 3.33E-11 |
| Slc27a4 | 3.14E-15 | 0.538530023 | 1 | 0.679 | 4.05E-11 |
| Rps6kc1 | 3.82E-15 | 0.251448895 | 1 | 0.629 | 4.93E-11 |
| Ascl1 | 3.84E-15 | 1.17113543 | 1 | 0.164 | 4.95E-11 |
| Strada | 4.04E-15 | 0.437066594 | 1 | 0.75 | 5.20E-11 |
| Tmx4 | 5.43E-15 | 0.255234502 | 1 | 0.95 | 7.01E-11 |
| Klhdc8b | 5.48E-15 | 0.293124944 | 1 | 0.95 | 7.06E-11 |
| Acvr2b | 1.30E-14 | 1.004361784 | 1 | 0.771 | 1.67E-10 |
| Cblb | 1.47E-14 | 0.489849396 | 0.75 | 0.764 | 1.90E-10 |
| Usp42 | 1.92E-14 | 0.439473769 | 0.75 | 0.871 | 2.48E-10 |
| Tgfbr1 | 1.96E-14 | 0.701166323 | 1 | 0.986 | 2.52E-10 |
| Tagln3 | 1.99E-14 | 0.910636248 | 1 | 0.314 | 2.57E-10 |
| Uros | 2.15E-14 | 0.659796855 | 1 | 0.829 | 2.77E-10 |
| Zfp521 | 2.33E-14 | 0.571386974 | 0.75 | 0.671 | 3.00E-10 |
| Samd14 | 2.72E-14 | 0.86707529 | 1 | 0.579 | 3.51E-10 |
| Tmem2 | 3.39E-14 | 0.69743232 | 1 | 0.871 | 4.37E-10 |
| Plbd2 | 4.92E-14 | 0.54358342 | 1 | 0.964 | 6.34E-10 |
| Cdc25b | 5.03E-14 | 1.052082799 | 1 | 0.564 | 6.49E-10 |
| Xndc1 | 5.92E-14 | 0.304132071 | 1 | 0.743 | 7.63E-10 |
| Npl | 7.26E-14 | 0.315193969 | 1 | 0.736 | 9.36E-10 |
| Rpusd2 | 7.32E-14 | 0.447069637 | 0.5 | 0.693 | 9.43E-10 |
| Efnb3 | 8.64E-14 | 0.793286377 | 1 | 0.5 | 1.11E-09 |
| Rnf31 | 1.03E-13 | 0.334399929 | 0.75 | 0.736 | 1.32E-09 |

| | | | | | |
|---|---|---|---|---|---|
| Cog5 | 1.05E-13 | 0.66931802 | 1 | 0.864 | 1.36E-09 |
| Crmp1 | 1.06E-13 | 0.580495478 | 0.75 | 0.2 | 1.37E-09 |
| Rgmb | 1.28E-13 | 0.760939685 | 1 | 0.393 | 1.65E-09 |
| Epha7 | 1.73E-13 | 0.564905366 | 1 | 0.714 | 2.23E-09 |
| Etnk1 | 1.73E-13 | 0.27772558 | 0.5 | 0.85 | 2.24E-09 |
| Rhobtb3 | 1.74E-13 | 0.424424323 | 1 | 0.85 | 2.24E-09 |
| Dhx32 | 2.34E-13 | 0.686907204 | 0.75 | 0.679 | 3.01E-09 |
| Pcnxl3 | 2.58E-13 | 0.297809514 | 1 | 0.821 | 3.32E-09 |
| Dtl | 2.74E-13 | 0.434290089 | 1 | 1 | 3.53E-09 |
| Nup153 | 2.74E-13 | 0.362686194 | 1 | 0.964 | 3.53E-09 |
| Btbd17 | 3.27E-13 | 0.650876267 | 0.75 | 0.136 | 4.22E-09 |
| Zfp329 | 3.37E-13 | 0.382536214 | 1 | 0.9 | 4.34E-09 |
| Tcof1 | 3.60E-13 | 0.899884575 | 1 | 0.986 | 4.64E-09 |
| Camsap1 | 3.85E-13 | 0.366988466 | 1 | 0.9 | 4.96E-09 |
| Myh10 | 4.32E-13 | 0.77072076 | 1 | 1 | 5.57E-09 |
| Fgfbp3 | 5.36E-13 | 1.330761302 | 1 | 0.436 | 6.92E-09 |
| Rtn1 | 5.43E-13 | 1.030747939 | 1 | 0.3 | 7.00E-09 |
| Plxna1 | 5.96E-13 | 0.430257496 | 1 | 0.643 | 7.69E-09 |
| Cc2d1a | 6.42E-13 | 0.289261924 | 0.75 | 0.664 | 8.28E-09 |
| Ubn1 | 6.55E-13 | 0.328140595 | 1 | 0.814 | 8.44E-09 |
| Cbl | 6.87E-13 | 0.467930656 | 1 | 0.829 | 8.86E-09 |
| Pigm | 7.22E-13 | 0.33136646 | 1 | 0.957 | 9.31E-09 |
| Morc4 | 7.93E-13 | 0.355018705 | 1 | 0.971 | 1.02E-08 |
| Ddit4l | 8.19E-13 | 0.708575247 | 0.75 | 0.279 | 1.06E-08 |
| Zfp516 | 8.37E-13 | 0.254600337 | 1 | 0.821 | 1.08E-08 |
| Oxct1 | 9.73E-13 | 0.482011127 | 1 | 0.771 | 1.25E-08 |
| Msi1 | 1.29E-12 | 0.428402027 | 1 | 0.993 | 1.66E-08 |
| Meis2 | 1.52E-12 | 0.295755044 | 1 | 0.793 | 1.96E-08 |
| Ermp1 | 1.71E-12 | 0.253788842 | 1 | 0.986 | 2.21E-08 |
| Dpysl4 | 2.37E-12 | 0.585806362 | 1 | 0.479 | 3.06E-08 |
| Extl3 | 2.48E-12 | 0.51037229 | 1 | 0.9 | 3.19E-08 |
| Osgin2 | 2.58E-12 | 0.39645211 | 0.25 | 0.75 | 3.32E-08 |
| Brca1 | 3.96E-12 | 0.338302072 | 1 | 0.886 | 5.10E-08 |
| Sik3 | 5.15E-12 | 0.282729069 | 0.75 | 0.686 | 6.64E-08 |
| 2510009E( | 1.01E-11 | 0.273875953 | 0.75 | 0.157 | 1.30E-07 |
| Stxbp6 | 1.03E-11 | 0.430663063 | 1 | 0.936 | 1.32E-07 |
| Grik4 | 1.09E-11 | 0.260372916 | 0.5 | 0.714 | 1.40E-07 |
| Ankrd13b | 1.09E-11 | 0.419679278 | 0.75 | 0.364 | 1.41E-07 |
| Wdr76 | 1.30E-11 | 0.347572576 | 1 | 0.964 | 1.68E-07 |
| Brf2 | 1.52E-11 | 0.55767335 | 0.5 | 0.6 | 1.96E-07 |
| Ext1 | 1.55E-11 | 0.355378114 | 1 | 0.65 | 2.00E-07 |
| Spsb4 | 1.81E-11 | 0.463923312 | 1 | 0.721 | 2.33E-07 |
| Mxra7 | 1.97E-11 | 0.368620368 | 0.75 | 0.357 | 2.54E-07 |
| Manea | 2.00E-11 | 0.423536116 | 1 | 0.893 | 2.58E-07 |
| Stat5b | 2.04E-11 | 0.323039172 | 0.75 | 0.921 | 2.63E-07 |
| Ube4b | 2.05E-11 | 0.575675048 | 1 | 1 | 2.64E-07 |
| Ttll4 | 2.43E-11 | 0.316243966 | 1 | 0.929 | 3.13E-07 |
| Tmtc2 | 2.57E-11 | 0.367441663 | 1 | 0.314 | 3.31E-07 |
| Tmed7 | 3.50E-11 | 0.266050872 | 1 | 1 | 4.51E-07 |
| Bscl2 | 4.11E-11 | 0.398686693 | 1 | 0.986 | 5.30E-07 |
| Cep135 | 4.39E-11 | 0.326290506 | 1 | 0.814 | 5.66E-07 |
| Recql4 | 4.79E-11 | 0.522321922 | 1 | 0.536 | 6.18E-07 |
| Prpf8 | 5.52E-11 | 0.906359134 | 1 | 1 | 7.11E-07 |
| Vac14 | 5.53E-11 | 0.529218926 | 0.75 | 0.579 | 7.13E-07 |
| Cabin1 | 5.99E-11 | 0.440403899 | 1 | 0.907 | 7.72E-07 |
| Map6 | 6.46E-11 | 0.311193087 | 1 | 0.136 | 8.33E-07 |
| Slc43a1 | 7.67E-11 | 0.588973476 | 1 | 0.929 | 9.89E-07 |
| Vps54 | 7.90E-11 | 0.311729089 | 0.75 | 0.736 | 1.02E-06 |

| | | | | | |
|---|---|---|---|---|---|
| 1190002N | 8.23E-11 | 0.533055031 | 1 | 0.629 | 1.06E-06 |
| Fanca | 8.86E-11 | 0.309901539 | 1 | 0.879 | 1.14E-06 |
| Lancl2 | 8.91E-11 | 0.748079272 | 1 | 0.779 | 1.15E-06 |
| Tmem33 | 9.16E-11 | 0.529058691 | 1 | 1 | 1.18E-06 |
| Kdm4a | 9.24E-11 | 0.491596756 | 0.75 | 0.629 | 1.19E-06 |
| Espl1 | 9.80E-11 | 0.29113279 | 1 | 0.864 | 1.26E-06 |
| Ccdc57 | 1.08E-10 | 0.403699134 | 0.5 | 0.586 | 1.39E-06 |
| Timp2 | 1.11E-10 | 0.296221184 | 0.5 | 0.679 | 1.43E-06 |
| Zfp518b | 1.19E-10 | 0.395328817 | 0.75 | 0.757 | 1.54E-06 |
| Dll1 | 1.20E-10 | 1.166975719 | 1 | 0.443 | 1.54E-06 |
| Cpe | 1.22E-10 | 1.320794587 | 1 | 0.671 | 1.58E-06 |
| Gm10330 | 1.27E-10 | 0.394337341 | 0.75 | 0.807 | 1.64E-06 |
| Elovl4 | 1.28E-10 | 0.491326201 | 1 | 0.436 | 1.64E-06 |
| Tubb2b | 1.34E-10 | 0.817777536 | 1 | 0.979 | 1.72E-06 |
| Man2a2 | 1.39E-10 | 0.356600731 | 1 | 0.836 | 1.80E-06 |
| Itgb3bp | 1.47E-10 | 0.313185431 | 1 | 0.964 | 1.89E-06 |
| Rbmx | 1.61E-10 | 1.019802604 | 1 | 0.993 | 2.08E-06 |
| Zfp146 | 1.82E-10 | 0.353985665 | 0.75 | 0.743 | 2.34E-06 |
| Tmcc2 | 1.92E-10 | 0.397790464 | 1 | 0.286 | 2.48E-06 |
| Cmip | 1.93E-10 | 0.403364807 | 0.75 | 0.879 | 2.49E-06 |
| Taok2 | 2.07E-10 | 0.401849741 | 1 | 0.929 | 2.67E-06 |
| Rai14 | 2.09E-10 | 0.753849718 | 1 | 0.879 | 2.69E-06 |
| Atf6 | 2.21E-10 | 0.33158953 | 1 | 0.936 | 2.85E-06 |
| Ccdc15 | 2.27E-10 | 0.314640594 | 0.5 | 0.543 | 2.93E-06 |
| Cptp | 2.38E-10 | 0.560598503 | 1 | 0.629 | 3.07E-06 |
| Ext2 | 2.78E-10 | 0.680419387 | 1 | 0.993 | 3.59E-06 |
| Galnt16 | 2.90E-10 | 0.752897471 | 1 | 0.371 | 3.74E-06 |
| Pcgf1 | 2.95E-10 | 0.436616957 | 0.75 | 0.686 | 3.81E-06 |
| Dvl2 | 3.29E-10 | 0.29222424 | 1 | 0.957 | 4.24E-06 |
| Pnpla2 | 3.37E-10 | 0.568087943 | 0.75 | 0.721 | 4.34E-06 |
| Gadd45g | 3.81E-10 | 1.746347667 | 0.75 | 0.643 | 4.92E-06 |
| Slc39a7 | 4.24E-10 | 0.366804404 | 1 | 0.971 | 5.47E-06 |
| Glce | 4.65E-10 | 0.36595185 | 1 | 0.786 | 5.99E-06 |
| Zcchc14 | 4.82E-10 | 0.391992965 | 1 | 0.771 | 6.21E-06 |
| Tmem35 | 4.82E-10 | 0.720144698 | 0.75 | 0.114 | 6.21E-06 |
| Zfp664 | 5.36E-10 | 0.457240203 | 1 | 0.943 | 6.90E-06 |
| Ece1 | 5.76E-10 | 0.736643543 | 1 | 0.993 | 7.42E-06 |
| Rnf220 | 5.80E-10 | 0.49144475 | 1 | 0.836 | 7.48E-06 |
| Dync1h1 | 5.81E-10 | 0.414853241 | 1 | 0.979 | 7.49E-06 |
| Ints5 | 6.33E-10 | 0.652176499 | 1 | 0.893 | 8.16E-06 |
| Zgpat | 6.41E-10 | 0.328475829 | 0.75 | 0.814 | 8.26E-06 |
| Mcm10 | 6.77E-10 | 0.506024916 | 1 | 0.957 | 8.73E-06 |
| Fyn | 7.26E-10 | 0.317842383 | 0.75 | 0.6 | 9.36E-06 |
| Dmtf1 | 9.43E-10 | 0.394371181 | 1 | 0.971 | 1.22E-05 |
| Spata13 | 9.62E-10 | 0.304609045 | 1 | 0.879 | 1.24E-05 |
| Emc1 | 1.02E-09 | 0.35107157 | 1 | 0.85 | 1.32E-05 |
| Gnai1 | 1.03E-09 | 0.314224937 | 1 | 0.321 | 1.32E-05 |
| Cad | 1.06E-09 | 0.657862247 | 1 | 0.964 | 1.36E-05 |
| Clasp1 | 1.09E-09 | 0.31072138 | 1 | 0.979 | 1.40E-05 |
| March6 | 1.33E-09 | 0.610083712 | 1 | 0.979 | 1.72E-05 |
| Ndst1 | 1.37E-09 | 0.37461566 | 1 | 0.829 | 1.76E-05 |
| Hus1 | 1.50E-09 | 0.430072282 | 1 | 0.943 | 1.94E-05 |
| Neo1 | 1.51E-09 | 0.348980445 | 1 | 0.807 | 1.95E-05 |
| Tmem201 | 1.67E-09 | 0.26202788 | 1 | 0.979 | 2.15E-05 |
| Fance | 1.75E-09 | 0.608115551 | 1 | 0.95 | 2.26E-05 |
| Ktn1 | 1.91E-09 | 0.499754846 | 1 | 1 | 2.46E-05 |
| Sh3kbp1 | 1.93E-09 | 0.291918753 | 1 | 0.821 | 2.48E-05 |
| Ehmt1 | 2.01E-09 | 0.62352148 | 1 | 0.993 | 2.59E-05 |

| | | | | | |
|---|---|---|---|---|---|
| Pomt2 | 2.04E-09 | 0.27258606 | 0.75 | 0.786 | 2.63E-05 |
| Rcc2 | 2.13E-09 | 0.568679575 | 1 | 0.979 | 2.75E-05 |
| Trrap | 2.31E-09 | 0.259888165 | 1 | 1 | 2.98E-05 |
| Abi2 | 2.38E-09 | 0.304268943 | 1 | 0.986 | 3.07E-05 |
| Pcnxl4 | 2.74E-09 | 0.394491012 | 1 | 0.757 | 3.53E-05 |
| B3galnt2 | 2.83E-09 | 0.345581383 | 0.5 | 0.886 | 3.65E-05 |
| Peg12 | 3.11E-09 | 0.734305776 | 1 | 0.971 | 4.01E-05 |
| Fam188b | 3.37E-09 | 0.382288241 | 0.75 | 0.564 | 4.35E-05 |
| Rcor2 | 3.41E-09 | 0.375332444 | 1 | 0.671 | 4.39E-05 |
| Smarcd1 | 3.48E-09 | 0.632101727 | 1 | 0.95 | 4.49E-05 |
| Slc52a2 | 3.67E-09 | 0.431986128 | 1 | 0.7 | 4.73E-05 |
| Ezh2 | 3.90E-09 | 0.480843275 | 1 | 0.957 | 5.03E-05 |
| Rnd2 | 3.95E-09 | 0.677625368 | 0.75 | 0.514 | 5.09E-05 |
| Emilin2 | 4.19E-09 | 0.619827874 | 0.75 | 0.45 | 5.40E-05 |
| Prpf3 | 4.23E-09 | 0.654898934 | 1 | 0.993 | 5.46E-05 |
| Mafg | 4.25E-09 | 0.293078971 | 0.75 | 0.936 | 5.48E-05 |
| Scrib | 4.34E-09 | 0.406242087 | 1 | 0.9 | 5.60E-05 |
| Zfp687 | 4.93E-09 | 0.33628543 | 1 | 0.9 | 6.35E-05 |
| Dnajb2 | 5.19E-09 | 0.495097884 | 1 | 0.779 | 6.69E-05 |
| Clasp2 | 5.43E-09 | 0.392629994 | 1 | 0.986 | 7.00E-05 |
| Plch1 | 5.65E-09 | 0.305229214 | 0.75 | 0.436 | 7.28E-05 |
| Cstf2 | 5.65E-09 | 0.31463936 | 1 | 0.964 | 7.29E-05 |
| Fzr1 | 5.85E-09 | 0.553072065 | 1 | 0.95 | 7.54E-05 |
| Abcb8 | 5.96E-09 | 0.695629855 | 1 | 0.964 | 7.68E-05 |
| Zfp41 | 7.15E-09 | 0.810772376 | 1 | 0.714 | 9.21E-05 |
| Pomc | 7.37E-09 | 0.421365938 | 0.75 | 0.257 | 9.50E-05 |
| F2r | 7.55E-09 | 1.307925548 | 1 | 0.95 | 9.73E-05 |
| Zfp59 | 7.64E-09 | 0.317765623 | 0.75 | 0.786 | 9.85E-05 |
| Zfp445 | 8.43E-09 | 0.480371374 | 1 | 0.879 | 0.000108723 |
| Irs2 | 8.70E-09 | 0.393360049 | 0.75 | 0.743 | 0.000112181 |
| Rragd | 9.05E-09 | 0.320012512 | 0.75 | 0.536 | 0.000116672 |
| Ints1 | 1.19E-08 | 0.278923693 | 1 | 0.829 | 0.000153108 |
| Map4 | 1.21E-08 | 0.264522633 | 1 | 1 | 0.000156486 |
| Pltp | 1.25E-08 | 0.603754517 | 1 | 0.364 | 0.000160766 |
| Adpgk | 1.34E-08 | 0.818360391 | 0.75 | 0.964 | 0.000173384 |
| Kif5c | 1.40E-08 | 0.405712231 | 1 | 0.443 | 0.000180964 |
| Zfp568 | 1.47E-08 | 0.349889156 | 1 | 0.971 | 0.000189079 |
| Sptlc2 | 1.56E-08 | 0.299048866 | 1 | 0.964 | 0.000200977 |
| Raver1 | 1.61E-08 | 0.640861655 | 1 | 0.993 | 0.000207139 |
| Xpo5 | 1.61E-08 | 0.818233529 | 1 | 0.993 | 0.000207718 |
| Rfc1 | 1.70E-08 | 0.408078759 | 1 | 1 | 0.000219186 |
| Add1 | 1.71E-08 | 0.393869642 | 0.75 | 0.986 | 0.000221012 |
| Pds5a | 1.83E-08 | 0.481955841 | 1 | 0.9 | 0.00023623 |
| Gnl3l | 1.84E-08 | 0.435199306 | 1 | 0.993 | 0.000237418 |
| Tmco3 | 2.27E-08 | 0.361000385 | 1 | 0.979 | 0.000292112 |
| Llgl1 | 2.41E-08 | 0.735234004 | 1 | 0.914 | 0.000311 |
| Champ1 | 2.44E-08 | 0.707870979 | 0.75 | 0.929 | 0.000315067 |
| Scarb2 | 2.52E-08 | 0.75433157 | 1 | 0.636 | 0.000325208 |
| Fat1 | 2.58E-08 | 0.284121239 | 1 | 0.864 | 0.000332563 |
| Arnt2 | 2.61E-08 | 0.339505793 | 1 | 0.564 | 0.000335954 |
| Kat6a | 2.63E-08 | 0.272042922 | 1 | 0.8 | 0.000339553 |
| Lmnb2 | 2.68E-08 | 0.758919833 | 1 | 0.979 | 0.000345065 |
| Scube3 | 2.72E-08 | 0.303368475 | 0.75 | 0.929 | 0.000350059 |
| Zfp689 | 2.87E-08 | 0.53773706 | 0.75 | 0.557 | 0.000369452 |
| Ttll12 | 2.87E-08 | 0.632234644 | 1 | 0.979 | 0.000370626 |
| Odf2 | 2.90E-08 | 0.291722927 | 1 | 0.986 | 0.000373846 |
| Tmem191c | 3.08E-08 | 0.473617021 | 0.5 | 0.579 | 0.000397485 |
| Rbl1 | 3.09E-08 | 0.275316603 | 1 | 0.871 | 0.000398019 |

| | | | | | |
|---|---|---|---|---|---|
| Foxn3 | 3.09E-08 | 0.422710175 | 1 | 0.957 | 0.000398978 |
| Nup155 | 3.22E-08 | 0.436429654 | 1 | 0.979 | 0.000414576 |
| Ipo9 | 3.25E-08 | 0.541095167 | 1 | 1 | 0.000418369 |
| Tmem167l | 3.36E-08 | 0.478053814 | 1 | 0.814 | 0.000432664 |
| Ino80e | 3.49E-08 | 0.35436722 | 1 | 0.986 | 0.000450566 |
| Prim2 | 3.72E-08 | 0.384007992 | 1 | 0.971 | 0.000479818 |
| Slc35e1 | 3.88E-08 | 0.253368985 | 1 | 0.85 | 0.000500666 |
| Cadm1 | 3.94E-08 | 0.672740104 | 1 | 1 | 0.000507658 |
| Adck2 | 4.33E-08 | 0.388964776 | 1 | 0.55 | 0.000558111 |
| Rad17 | 4.40E-08 | 0.566853397 | 1 | 0.921 | 0.000566685 |
| Hes6 | 4.91E-08 | 0.956892136 | 1 | 0.936 | 0.000632358 |
| Akr1b3 | 4.98E-08 | 0.474130174 | 1 | 0.979 | 0.000642333 |
| Sfrp2 | 5.10E-08 | 0.685223074 | 1 | 0.464 | 0.000656917 |
| Zmiz1 | 5.25E-08 | 0.355768924 | 1 | 0.971 | 0.000676572 |
| Osbpl11 | 5.26E-08 | 0.453348847 | 1 | 0.886 | 0.000678131 |
| Zik1 | 5.33E-08 | 0.437586022 | 0.75 | 0.714 | 0.00068658 |
| Rpgr | 5.64E-08 | 0.271539787 | 0.75 | 0.793 | 0.000727477 |
| Skiv2l | 5.68E-08 | 0.483930881 | 0.75 | 0.921 | 0.000732785 |
| Igf2bp1 | 5.72E-08 | 0.577485673 | 1 | 1 | 0.000736786 |
| Fstl1 | 6.05E-08 | 1.324028537 | 1 | 0.936 | 0.000780125 |
| Golm1 | 6.85E-08 | 1.225897675 | 1 | 0.993 | 0.000883399 |
| Zbtb17 | 7.01E-08 | 0.443727834 | 0.75 | 0.886 | 0.000904346 |
| Npepps | 7.12E-08 | 0.362113239 | 1 | 0.986 | 0.000917995 |
| Ttf1 | 8.14E-08 | 0.299303555 | 1 | 1 | 0.001049567 |
| Epb4.1 | 8.34E-08 | 0.443503082 | 1 | 0.986 | 0.001074565 |
| Mdc1 | 8.60E-08 | 0.270017883 | 1 | 0.921 | 0.001108868 |
| Gng2 | 8.91E-08 | 0.443093517 | 0.75 | 0.464 | 0.001148263 |
| Kif21a | 8.92E-08 | 0.38604362 | 1 | 0.9 | 0.001150503 |
| Dleu7 | 9.22E-08 | 0.591991812 | 0.75 | 0.157 | 0.00118866 |
| Tnfrsf21 | 9.28E-08 | 0.695015685 | 1 | 0.714 | 0.001196599 |
| Slc7a6 | 9.31E-08 | 0.952819847 | 1 | 0.993 | 0.001200253 |
| Mycl | 9.59E-08 | 0.555121288 | 1 | 0.607 | 0.001236416 |
| Snrnp200 | 1.01E-07 | 0.627844156 | 1 | 1 | 0.001296824 |
| Haus8 | 1.03E-07 | 0.388046858 | 1 | 1 | 0.001326867 |
| Fkbp5 | 1.04E-07 | 0.291648114 | 1 | 0.786 | 0.001341214 |
| Ptms | 1.09E-07 | 0.648434816 | 1 | 0.964 | 0.00140273 |
| Fam171a1 | 1.10E-07 | 0.654005734 | 1 | 0.764 | 0.001414167 |
| Lman2l | 1.22E-07 | 0.358956621 | 1 | 1 | 0.001567294 |
| Ubqln4 | 1.33E-07 | 0.458788475 | 1 | 0.964 | 0.001720594 |
| Gpm6b | 1.34E-07 | 0.415214183 | 0.75 | 0.343 | 0.001724994 |
| Pom121 | 1.50E-07 | 0.377725188 | 1 | 0.836 | 0.001929783 |
| Wdr33 | 1.54E-07 | 0.584677346 | 1 | 0.993 | 0.001983792 |
| E130309D( | 1.57E-07 | 0.326353067 | 1 | 0.986 | 0.002021117 |
| Taf1c | 1.66E-07 | 0.26114127 | 0.75 | 0.593 | 0.002133971 |
| Map2k7 | 1.78E-07 | 0.283248818 | 1 | 0.943 | 0.002296515 |
| Slc12a7 | 1.82E-07 | 0.560551843 | 1 | 0.971 | 0.002350122 |
| Naa60 | 1.91E-07 | 0.389913001 | 0.75 | 0.943 | 0.002461694 |
| Ncln | 2.06E-07 | 0.805357871 | 1 | 0.993 | 0.002660388 |
| Man2b2 | 2.12E-07 | 0.48486457 | 1 | 0.657 | 0.002727046 |
| Vwa9 | 2.20E-07 | 0.799782504 | 1 | 0.979 | 0.002833747 |
| Slc9a6 | 2.20E-07 | 0.259310003 | 0.75 | 0.586 | 0.002839535 |
| Clip3 | 2.24E-07 | 0.715776944 | 1 | 0.571 | 0.002890847 |
| Tmem101 | 2.29E-07 | 0.745186115 | 1 | 0.907 | 0.002958615 |
| Cdc42se2 | 2.30E-07 | 0.897646338 | 1 | 0.971 | 0.002969175 |
| Zfp113 | 2.31E-07 | 0.310710301 | 1 | 1 | 0.002981436 |
| Crabp1 | 2.41E-07 | 2.081741802 | 1 | 0.6 | 0.003102912 |
| Six5 | 2.44E-07 | 0.283726667 | 0.25 | 0.514 | 0.003141969 |
| Zfp944 | 2.63E-07 | 0.321850895 | 1 | 0.9 | 0.003394445 |

| | | | | | |
|---|---|---|---|---|---|
| Loxl2 | 2.89E-07 | 0.302473495 | 1 | 1 | 0.003727085 |
| Pole2 | 3.36E-07 | 0.869736391 | 1 | 0.921 | 0.004327514 |
| Ppm1l | 3.40E-07 | 0.286031125 | 1 | 0.75 | 0.004379747 |
| Elovl6 | 3.50E-07 | 0.45111285 | 1 | 0.971 | 0.004518277 |
| Carkd | 3.74E-07 | 0.335831159 | 1 | 0.943 | 0.004823778 |
| Pprc1 | 3.74E-07 | 0.446332822 | 1 | 0.971 | 0.004824163 |
| Ctu2 | 3.75E-07 | 0.354656059 | 1 | 0.936 | 0.004831385 |
| Fzd3 | 4.03E-07 | 0.588328735 | 1 | 0.8 | 0.005195659 |
| Bptf | 4.19E-07 | 0.251337211 | 1 | 1 | 0.005398428 |
| E2f2 | 4.25E-07 | 0.310787302 | 0.5 | 0.586 | 0.005478701 |
| Pcyox1 | 4.35E-07 | 0.669178558 | 1 | 0.986 | 0.005610571 |
| Gpaa1 | 4.41E-07 | 0.792743134 | 1 | 0.986 | 0.005691816 |
| Zdhhc16 | 4.44E-07 | 0.916299089 | 1 | 0.936 | 0.005721532 |
| Gpd2 | 4.56E-07 | 0.262214565 | 0.75 | 0.793 | 0.005880773 |
| Dpysl5 | 4.72E-07 | 0.715134168 | 1 | 0.757 | 0.006078868 |
| Shroom2 | 5.07E-07 | 0.25680877 | 0.75 | 0.521 | 0.006534503 |
| Npdc1 | 5.24E-07 | 0.360299403 | 0.75 | 0.571 | 0.006752659 |
| Hap1 | 5.52E-07 | 0.308161686 | 1 | 0.286 | 0.007117531 |
| Mettl17 | 5.54E-07 | 0.508842274 | 1 | 0.964 | 0.007140566 |
| Rbm4b | 5.55E-07 | 0.286092593 | 1 | 1 | 0.007156855 |
| Zfp207 | 5.75E-07 | 0.716120103 | 1 | 1 | 0.007411167 |
| Myo1e | 5.96E-07 | 0.441536581 | 1 | 0.85 | 0.007684472 |
| Pak2 | 5.98E-07 | 0.476008837 | 1 | 1 | 0.00770725 |
| Casp8ap2 | 6.05E-07 | 0.441543796 | 1 | 0.957 | 0.007801057 |
| Smad2 | 6.29E-07 | 0.328682736 | 0.75 | 0.95 | 0.008106777 |
| Heatr3 | 6.40E-07 | 0.586823923 | 1 | 0.964 | 0.00824685 |
| Sufu | 6.71E-07 | 0.384554199 | 1 | 0.829 | 0.008644921 |
| Hipk2 | 6.75E-07 | 0.343232394 | 1 | 0.986 | 0.008707889 |
| Hmgxb4 | 7.08E-07 | 0.410137864 | 1 | 0.936 | 0.00912397 |
| Phf6 | 7.08E-07 | 0.358442546 | 1 | 0.9 | 0.009133874 |
| Aacs | 7.44E-07 | 0.547963822 | 1 | 0.864 | 0.009590391 |
| Pcbp3 | 7.45E-07 | 0.415803944 | 0.5 | 0.779 | 0.009608068 |
| Zfp511 | 7.67E-07 | 0.894700045 | 1 | 0.843 | 0.009883654 |
| Zswim5 | 8.11E-07 | 0.353203099 | 1 | 0.214 | 0.010454496 |
| Itfg2 | 8.39E-07 | 0.275824624 | 1 | 0.836 | 0.010819675 |
| Trib2 | 8.44E-07 | 0.521442478 | 1 | 0.386 | 0.010874755 |
| Nxf1 | 9.26E-07 | 0.634604996 | 1 | 1 | 0.011941975 |
| Prkrir | 9.84E-07 | 0.296576069 | 1 | 0.979 | 0.012681456 |
| Plagl2 | 9.90E-07 | 0.315675541 | 0.75 | 0.964 | 0.012763037 |
| Acin1 | 1.04E-06 | 0.403184266 | 1 | 0.993 | 0.013464848 |
| Snapc3 | 1.05E-06 | 0.269394861 | 1 | 0.943 | 0.013543934 |
| Bysl | 1.05E-06 | 0.555860104 | 1 | 0.993 | 0.013600077 |
| Slc46a1 | 1.06E-06 | 0.386069905 | 0.5 | 0.671 | 0.013638224 |
| Casp2 | 1.11E-06 | 0.46299413 | 1 | 0.957 | 0.014265135 |
| Zfp367 | 1.13E-06 | 0.388993828 | 1 | 0.957 | 0.014511824 |
| Ptbp2 | 1.14E-06 | 0.452706616 | 1 | 0.993 | 0.014662509 |
| Trp53inp2 | 1.14E-06 | 0.516627488 | 0.75 | 0.686 | 0.014684713 |
| Pkmyt1 | 1.17E-06 | 0.77397488 | 1 | 0.986 | 0.015101007 |
| Clstn1 | 1.20E-06 | 0.473974868 | 1 | 0.814 | 0.015528031 |
| Plk3 | 1.23E-06 | 0.81084759 | 0.75 | 0.629 | 0.015918461 |
| Thumpd2 | 1.28E-06 | 0.514496707 | 0.75 | 0.564 | 0.016474978 |
| Mllt1 | 1.37E-06 | 0.30234392 | 0.75 | 0.564 | 0.017668721 |
| Dbn1 | 1.38E-06 | 1.211505504 | 1 | 1 | 0.017820833 |
| Alg10b | 1.39E-06 | 0.883925083 | 1 | 0.871 | 0.017977438 |
| Zfp36l1 | 1.41E-06 | 0.762393709 | 1 | 1 | 0.01820511 |
| Fam172a | 1.42E-06 | 0.332969642 | 0.75 | 0.707 | 0.01836513 |
| Nkx2-2 | 1.48E-06 | 0.397420267 | 1 | 0.079 | 0.019144381 |
| Lgmn | 1.52E-06 | 0.828116816 | 1 | 0.95 | 0.019575061 |

| | | | | | |
|---|---|---|---|---|---|
| Eif5b | 1.54E-06 | 0.374021712 | 1 | 1 | 0.019840138 |
| Sppl2b | 1.60E-06 | 0.37488333 | 0.75 | 0.707 | 0.020671681 |
| Mkl1 | 1.67E-06 | 0.308257091 | 1 | 0.921 | 0.02150926 |
| Panx1 | 1.72E-06 | 0.562098733 | 1 | 0.836 | 0.022128736 |
| Poc1b | 1.80E-06 | 0.356385142 | 0.75 | 0.721 | 0.023148643 |
| Bclaf1 | 1.81E-06 | 0.567721009 | 1 | 1 | 0.023345686 |
| Epha4 | 1.85E-06 | 0.513042261 | 1 | 0.686 | 0.02381784 |
| Efna5 | 1.91E-06 | 0.387433273 | 0.5 | 0.65 | 0.024615904 |
| Zbtb22 | 1.91E-06 | 0.284075824 | 0.75 | 0.521 | 0.024671963 |
| Cfap43 | 2.00E-06 | 0.26453743 | 1 | 0.55 | 0.025807698 |
| Cdh24 | 2.28E-06 | 0.51510497 | 1 | 0.407 | 0.029345686 |
| Col18a1 | 2.36E-06 | 1.206832137 | 1 | 1 | 0.03037299 |
| Msh2 | 2.51E-06 | 0.710756146 | 1 | 0.993 | 0.032421168 |
| Glg1 | 2.59E-06 | 0.541987744 | 1 | 1 | 0.033441648 |
| Wdr91 | 2.90E-06 | 0.327978496 | 1 | 0.479 | 0.037400321 |
| Slc4a7 | 3.19E-06 | 0.343240579 | 1 | 0.879 | 0.041084969 |
| Mipep | 3.23E-06 | 0.437942031 | 1 | 0.779 | 0.041592871 |
| Ift140 | 3.24E-06 | 0.261237335 | 0.75 | 0.836 | 0.041819983 |
| Nedd4 | 3.25E-06 | 0.65269043 | 1 | 1 | 0.041930002 |
| U2af2 | 3.26E-06 | 0.539068379 | 1 | 0.993 | 0.042070456 |
| Tmem109 | 3.28E-06 | 0.397284971 | 0.75 | 0.95 | 0.042305198 |
| Nup188 | 3.29E-06 | 0.441236032 | 1 | 0.979 | 0.042438737 |
| Rcbtb1 | 3.29E-06 | 0.419175349 | 1 | 0.971 | 0.042456675 |
| Map4k1 | 3.38E-06 | 0.325891848 | 0.5 | 0.529 | 0.043562954 |
| Hyou1 | 3.38E-06 | 0.529561701 | 1 | 1 | 0.043623373 |
| Secisbp2 | 3.42E-06 | 0.289628374 | 1 | 0.879 | 0.044066664 |
| Eri3 | 3.74E-06 | 0.590424726 | 1 | 0.986 | 0.048212215 |
| Strip1 | 3.75E-06 | 0.456619334 | 1 | 0.907 | 0.048366446 |
| Med12 | 3.87E-06 | 0.250788209 | 1 | 0.679 | 0.04993928 |
| Rara | 3.91E-06 | 0.418512176 | 1 | 0.907 | 0.050467981 |
| Esyt1 | 4.71E-06 | 0.309937035 | 0.75 | 0.5 | 0.060729837 |
| Txlna | 5.00E-06 | 0.575058717 | 1 | 0.993 | 0.064445314 |
| Ppap2b | 5.42E-06 | 0.379620139 | 0.75 | 0.707 | 0.069927206 |
| Fuca1 | 6.40E-06 | 0.456549508 | 1 | 0.893 | 0.082482485 |
| Srprb | 6.48E-06 | 0.351173453 | 1 | 1 | 0.083568281 |
| Smc1a | 6.54E-06 | 0.578038398 | 1 | 1 | 0.084286576 |
| Khsrp | 6.56E-06 | 0.315582955 | 1 | 1 | 0.084596636 |
| Asphd2 | 6.60E-06 | 0.38630245 | 0.75 | 0.157 | 0.085062628 |
| Hmgcr | 6.62E-06 | 0.691927566 | 1 | 1 | 0.085402378 |
| Tmem231 | 6.72E-06 | 0.629910192 | 1 | 0.629 | 0.086581995 |
| Slc39a6 | 6.80E-06 | 0.528141421 | 1 | 0.907 | 0.087673402 |
| Cbfb | 6.81E-06 | 0.369927371 | 1 | 0.843 | 0.08773603 |
| Atp13a2 | 6.99E-06 | 0.300956819 | 1 | 0.95 | 0.090153118 |
| B4gat1 | 7.01E-06 | 0.534973483 | 1 | 0.793 | 0.090402628 |
| Chaf1b | 7.16E-06 | 0.664963277 | 1 | 0.986 | 0.092273967 |
| Sart3 | 7.17E-06 | 0.539362241 | 1 | 1 | 0.092495187 |
| Srd5a1 | 7.39E-06 | 0.300891873 | 0.5 | 0.7 | 0.095314474 |
| Rprd1b | 7.77E-06 | 0.285871391 | 1 | 1 | 0.1001458 |
| Prosc | 7.77E-06 | 0.374446285 | 1 | 0.921 | 0.100192201 |
| Fgfrl1 | 7.85E-06 | 0.288620495 | 0.5 | 0.743 | 0.101208558 |
| Arih2 | 8.08E-06 | 0.269689885 | 1 | 0.986 | 0.104150473 |
| Wscd1 | 8.74E-06 | 0.300967247 | 1 | 0.279 | 0.112664741 |
| Crabp2 | 8.94E-06 | 1.453099043 | 1 | 0.929 | 0.115281637 |
| Slc6a6 | 9.09E-06 | 0.544715113 | 1 | 0.986 | 0.117155135 |
| Tmem18 | 9.10E-06 | 0.343680659 | 0.5 | 0.593 | 0.117376287 |
| Mettl7a3 | 9.55E-06 | 0.266854152 | 1 | 1 | 0.123147511 |
| Alkbh3 | 9.86E-06 | 0.276526258 | 0.75 | 0.871 | 0.127169495 |
| Fxr1 | 1.01E-05 | 0.310759817 | 1 | 1 | 0.130834245 |

| | | | | | |
|---|---|---|---|---|---|
| Sapcd2 | 1.05E-05 | 0.322289468 | 0.75 | 0.721 | 0.134848075 |
| Tspan3 | 1.06E-05 | 1.04332315 | 1 | 1 | 0.137176065 |
| Paxbp1 | 1.10E-05 | 0.373037916 | 1 | 0.857 | 0.141311659 |
| Mns1 | 1.14E-05 | 0.325975815 | 1 | 0.929 | 0.146484411 |
| Atp1a1 | 1.16E-05 | 0.620472515 | 1 | 1 | 0.149837651 |
| Sall2 | 1.21E-05 | 0.290614443 | 1 | 0.714 | 0.156243872 |
| Jade3 | 1.21E-05 | 0.331011534 | 1 | 0.979 | 0.156634518 |
| Nova1 | 1.27E-05 | 0.261633365 | 0.75 | 0.279 | 0.163567966 |
| Slc10a3 | 1.27E-05 | 0.331289699 | 0.25 | 0.679 | 0.164274782 |
| Abhd17c | 1.28E-05 | 0.289174085 | 0.75 | 0.957 | 0.164899679 |
| Pcdh8 | 1.39E-05 | 0.30355412 | 0.75 | 0.129 | 0.178893154 |
| Utp6 | 1.39E-05 | 0.449622427 | 1 | 0.986 | 0.179761444 |
| Adamts7 | 1.43E-05 | 0.276271278 | 1 | 0.343 | 0.184902827 |
| Tril | 1.45E-05 | 0.2805182 | 1 | 0.336 | 0.187141488 |
| Ddx51 | 1.49E-05 | 0.295126585 | 1 | 0.886 | 0.191833771 |
| Fnbp4 | 1.52E-05 | 0.361947728 | 1 | 1 | 0.195336266 |
| Cx3cl1 | 1.52E-05 | 0.430722419 | 0.75 | 0.229 | 0.195463286 |
| Snx27 | 1.52E-05 | 0.264377685 | 1 | 0.95 | 0.19555238 |
| Ctnnb1 | 1.55E-05 | 0.921508418 | 1 | 1 | 0.200093848 |
| Arvcf | 1.55E-05 | 0.566613025 | 1 | 0.829 | 0.200364723 |
| Slc1a5 | 1.61E-05 | 0.439687373 | 1 | 0.957 | 0.207664542 |
| Eif4g1 | 1.63E-05 | 0.570844234 | 1 | 1 | 0.210078921 |
| Qk | 1.74E-05 | 0.285222735 | 1 | 1 | 0.223924819 |
| Nrm | 1.78E-05 | 0.911506405 | 1 | 0.95 | 0.228916321 |
| Rhpn1 | 1.87E-05 | 0.816295475 | 1 | 0.643 | 0.24066773 |
| Dnajc4 | 1.89E-05 | 0.505516134 | 0.5 | 0.607 | 0.243108675 |
| Fam3a | 1.90E-05 | 0.335871505 | 0.5 | 0.786 | 0.244703862 |
| Cyth2 | 1.97E-05 | 0.78564184 | 1 | 0.993 | 0.253900352 |
| Terf2 | 1.99E-05 | 0.26582291 | 0.75 | 0.679 | 0.256356485 |
| Rbak | 2.02E-05 | 0.380414456 | 1 | 0.779 | 0.261028905 |
| Ykt6 | 2.06E-05 | 0.304656167 | 1 | 0.871 | 0.265264045 |
| Nmral1 | 2.10E-05 | 0.615942474 | 1 | 1 | 0.270828327 |
| Dgcr8 | 2.16E-05 | 0.260648436 | 1 | 1 | 0.277929359 |
| Rnf44 | 2.20E-05 | 0.432361122 | 1 | 0.979 | 0.283568501 |
| Ska2 | 2.21E-05 | 0.647310329 | 1 | 0.986 | 0.285257382 |
| Cenpm | 2.25E-05 | 0.53093232 | 1 | 0.857 | 0.289630701 |
| Vamp2 | 2.26E-05 | 0.26909695 | 0.75 | 0.807 | 0.290863056 |
| Nedd9 | 2.34E-05 | 0.469172591 | 1 | 0.679 | 0.301811162 |
| Lias | 2.39E-05 | 0.40068959 | 1 | 0.957 | 0.307998753 |
| Bcs1l | 2.46E-05 | 0.383009433 | 1 | 0.821 | 0.316726207 |
| Naglu | 2.53E-05 | 0.266140247 | 0.25 | 0.664 | 0.325767499 |
| Fuk | 2.57E-05 | 0.268864922 | 1 | 0.643 | 0.331495642 |
| Gpc6 | 2.74E-05 | 0.392169198 | 1 | 0.914 | 0.353045015 |
| Crip2 | 2.87E-05 | 1.201912685 | 1 | 0.707 | 0.370143599 |
| Zfp202 | 2.88E-05 | 0.264711232 | 1 | 0.936 | 0.370694739 |
| Trip6 | 3.15E-05 | 0.682612601 | 1 | 0.864 | 0.405462983 |
| Mest | 3.41E-05 | 1.439702484 | 1 | 0.871 | 0.439320704 |
| Minpp1 | 3.46E-05 | 0.752576832 | 1 | 0.986 | 0.446527551 |
| Hnrnph1 | 3.54E-05 | 0.824271023 | 1 | 1 | 0.456405263 |
| Insr | 3.59E-05 | 0.284757922 | 1 | 0.857 | 0.462637659 |
| Ubp1 | 3.64E-05 | 0.443895773 | 1 | 0.886 | 0.469668793 |
| Top2a | 3.82E-05 | 0.854451862 | 1 | 1 | 0.492407494 |
| Mcm4 | 3.86E-05 | 1.073642408 | 1 | 1 | 0.497428245 |
| Fhl1 | 3.92E-05 | 0.558011825 | 1 | 0.771 | 0.505319731 |
| Dlst | 3.92E-05 | 0.848014501 | 1 | 1 | 0.505919581 |
| Scd1 | 3.97E-05 | 0.418932819 | 1 | 0.993 | 0.512430353 |
| Bicd2 | 3.99E-05 | 0.266018714 | 0.75 | 0.743 | 0.514320203 |
| Gid4 | 4.00E-05 | 0.335042792 | 1 | 0.971 | 0.516223192 |

| | | | | | |
|---|---|---|---|---|---|
| Lig1 | 4.01E-05 | 0.742040516 | 1 | 1 | 0.516576902 |
| Zfp710 | 4.12E-05 | 0.350563167 | 0.75 | 0.821 | 0.531222936 |
| Orc1 | 4.25E-05 | 0.31998358 | 1 | 0.757 | 0.548369709 |
| Ssbp3 | 4.42E-05 | 0.292992021 | 1 | 0.971 | 0.569453885 |
| Psap | 4.61E-05 | 1.114083694 | 1 | 0.993 | 0.594742434 |
| Pacsin3 | 4.66E-05 | 0.274061151 | 1 | 0.864 | 0.60052389 |
| Sort1 | 4.75E-05 | 0.281490362 | 1 | 0.893 | 0.611809242 |
| Cpsf1 | 4.82E-05 | 0.643050127 | 1 | 0.979 | 0.620947475 |
| Faf2 | 4.86E-05 | 0.494329491 | 1 | 1 | 0.626010959 |
| Ubap2 | 4.98E-05 | 0.634662189 | 1 | 0.993 | 0.641523356 |
| Zkscan3 | 5.15E-05 | 0.33786345 | 1 | 1 | 0.663302285 |
| Foxm1 | 5.21E-05 | 0.270437403 | 1 | 0.829 | 0.672043038 |
| Tmem168 | 5.28E-05 | 0.315624532 | 1 | 0.843 | 0.681234675 |
| Chek1 | 5.43E-05 | 0.400333006 | 1 | 0.986 | 0.699640618 |
| Porcn | 5.52E-05 | 0.676835024 | 1 | 0.657 | 0.711603423 |
| Pygo2 | 5.73E-05 | 0.371111693 | 1 | 0.886 | 0.738370192 |
| Ganab | 5.97E-05 | 0.754397934 | 1 | 1 | 0.769871706 |
| Rabepk | 6.15E-05 | 0.326637352 | 1 | 0.95 | 0.79263607 |
| Dynlt1c | 6.22E-05 | 0.406217078 | 1 | 1 | 0.801427444 |
| Nol8 | 6.31E-05 | 0.302902851 | 1 | 0.936 | 0.813194751 |
| Rfng | 6.42E-05 | 0.464810689 | 1 | 0.793 | 0.827519471 |
| Epn2 | 6.48E-05 | 0.313320171 | 1 | 0.8 | 0.835739791 |
| Vopp1 | 6.52E-05 | 0.299346151 | 1 | 0.807 | 0.840935112 |
| Prpf4b | 6.75E-05 | 0.524525591 | 1 | 1 | 0.869918429 |
| Elovl5 | 6.84E-05 | 0.62615229 | 1 | 1 | 0.881507542 |
| Tpcn2 | 6.90E-05 | 0.304909861 | 0.75 | 0.486 | 0.889587767 |
| Rmdn3 | 6.94E-05 | 0.283563511 | 0.75 | 0.957 | 0.894226904 |
| Zfp266 | 7.04E-05 | 0.492693979 | 1 | 0.986 | 0.908239632 |
| Mta2 | 7.09E-05 | 0.750247756 | 1 | 1 | 0.914667472 |
| Mki67 | 7.20E-05 | 0.342109265 | 1 | 0.971 | 0.92803407 |
| Fads1 | 7.22E-05 | 0.854790327 | 1 | 1 | 0.930783264 |
| Apbb1 | 7.40E-05 | 0.470166816 | 1 | 0.943 | 0.953555204 |
| Nelfa | 7.40E-05 | 0.44658202 | 1 | 0.979 | 0.954323096 |
| Gabpb1 | 7.72E-05 | 0.273814594 | 1 | 0.957 | 0.995383546 |
| Gns | 7.74E-05 | 0.53763164 | 1 | 0.979 | 0.997429042 |
| Arhgef1 | 7.79E-05 | 0.282717941 | 1 | 0.943 | 1 |
| Spop | 8.07E-05 | 0.712967039 | 1 | 0.993 | 1 |
| Fam13b | 8.17E-05 | 0.27319495 | 0.75 | 0.829 | 1 |
| Herc4 | 8.49E-05 | 0.409466506 | 1 | 0.929 | 1 |
| Ints9 | 8.50E-05 | 0.566934544 | 1 | 0.943 | 1 |
| Timm22 | 8.56E-05 | 0.282035477 | 1 | 1 | 1 |
| Zfand3 | 9.01E-05 | 0.401525158 | 0.75 | 0.893 | 1 |
| Carm1 | 9.09E-05 | 0.272273965 | 0.75 | 0.836 | 1 |
| Ppp2r4 | 9.19E-05 | 0.568523336 | 1 | 1 | 1 |
| Tm9sf4 | 9.58E-05 | 0.370633134 | 1 | 1 | 1 |
| Zmat5 | 9.77E-05 | 0.423290338 | 1 | 0.843 | 1 |
| Nptn | 9.85E-05 | 0.278400694 | 1 | 0.993 | 1 |
| Zfp821 | 0.000101772 | 0.424746594 | 1 | 0.557 | 1 |
| Ptk7 | 0.000103067 | 0.581805157 | 1 | 0.914 | 1 |
| Shd | 0.000108435 | 0.57599604 | 1 | 0.414 | 1 |
| Pigl | 0.000110029 | 0.40592031 | 1 | 0.993 | 1 |
| 2810408M | 0.000111729 | 0.379948102 | 1 | 0.95 | 1 |
| Prep | 0.000113394 | 0.718640833 | 1 | 1 | 1 |
| Nufip1 | 0.000113453 | 0.516854473 | 1 | 0.993 | 1 |
| Dhx38 | 0.000114093 | 0.504261771 | 1 | 0.943 | 1 |
| Zmynd11 | 0.000114537 | 0.352408566 | 1 | 0.986 | 1 |
| Cant1 | 0.000115507 | 0.382858598 | 1 | 0.893 | 1 |
| Ap1b1 | 0.00011689 | 0.529023861 | 1 | 0.986 | 1 |

| | | | | | |
|---|---|---|---|---|---|
| Plekha5 | 0.000117224 | 0.327935065 | 1 | 0.957 | 1 |
| Ddx17 | 0.000124395 | 0.432800541 | 1 | 0.993 | 1 |
| R3hdm4 | 0.000125994 | 0.81650056 | 1 | 0.986 | 1 |
| Cbx5 | 0.000131074 | 0.369720655 | 1 | 1 | 1 |
| Rere | 0.000131901 | 0.250564542 | 1 | 0.964 | 1 |
| Ptpn13 | 0.000132363 | 0.288034384 | 1 | 0.757 | 1 |
| Prps1 | 0.000137095 | 0.282314555 | 0.75 | 0.95 | 1 |
| Pald1 | 0.00013893 | 0.456330201 | 1 | 0.914 | 1 |
| Smad5 | 0.000143551 | 0.292794964 | 1 | 0.964 | 1 |
| Mau2 | 0.000149315 | 0.35992227 | 1 | 0.971 | 1 |
| Rbm12b2 | 0.000151543 | 0.251615266 | 0.75 | 0.793 | 1 |
| Ing4 | 0.000157093 | 0.450858557 | 1 | 1 | 1 |
| Kit | 0.000158812 | 0.528879351 | 1 | 0.75 | 1 |
| Ighmbp2 | 0.000161959 | 0.250078098 | 0.5 | 0.593 | 1 |
| 1110057K( | 0.000164769 | 0.36650319 | 1 | 0.993 | 1 |
| Cdc37l1 | 0.000167088 | 0.348387852 | 1 | 0.886 | 1 |
| Zkscan17 | 0.000169659 | 0.284883316 | 0.75 | 0.879 | 1 |
| Troap | 0.000172866 | 0.713343054 | 0.75 | 0.721 | 1 |
| Rad54l | 0.000174028 | 0.252084508 | 1 | 0.964 | 1 |
| Spns1 | 0.000174548 | 0.614446925 | 1 | 0.979 | 1 |
| Cxxc1 | 0.000175255 | 0.488090126 | 0.75 | 0.943 | 1 |
| Mpdu1 | 0.000175687 | 0.631367342 | 1 | 0.971 | 1 |
| Srsf1 | 0.000178585 | 0.46939145 | 1 | 1 | 1 |
| D930015E( | 0.000181448 | 0.317935373 | 1 | 0.929 | 1 |
| Ror2 | 0.000196327 | 0.568426835 | 1 | 0.75 | 1 |
| Ddx6 | 0.000196638 | 0.46844189 | 1 | 1 | 1 |
| Rgs12 | 0.000197051 | 0.285087693 | 1 | 0.8 | 1 |
| Bms1 | 0.000199446 | 0.624083482 | 1 | 0.979 | 1 |
| Ncstn | 0.000203832 | 0.765738081 | 1 | 0.986 | 1 |
| Tmem127 | 0.000206464 | 0.319164257 | 1 | 0.857 | 1 |
| Itfg1 | 0.000211167 | 0.546491056 | 1 | 0.979 | 1 |
| Sun1 | 0.00021207 | 0.329847006 | 1 | 0.993 | 1 |
| Hells | 0.000213878 | 0.401459388 | 1 | 1 | 1 |
| C730034F( | 0.000217417 | 0.365086877 | 1 | 0.986 | 1 |
| Trp53i11 | 0.000224768 | 0.683214374 | 1 | 0.643 | 1 |
| Wdr6 | 0.000225209 | 0.849297702 | 1 | 0.95 | 1 |
| Xrcc1 | 0.000226122 | 0.851392475 | 1 | 0.993 | 1 |
| Rad51 | 0.000228318 | 0.348197774 | 1 | 0.971 | 1 |
| Zfp503 | 0.000230738 | 0.294485613 | 1 | 0.379 | 1 |
| Uck1 | 0.000232039 | 0.595740079 | 1 | 0.95 | 1 |
| Ankrd54 | 0.000239451 | 0.362480326 | 1 | 0.957 | 1 |
| Ncapg2 | 0.000243859 | 0.314275684 | 1 | 0.886 | 1 |
| Mapk7 | 0.000248324 | 0.418669896 | 1 | 0.864 | 1 |
| Atf6b | 0.000253391 | 0.493643057 | 1 | 0.964 | 1 |
| Rtn3 | 0.000253399 | 0.606984762 | 1 | 1 | 1 |
| Wdr77 | 0.000260465 | 0.350149422 | 1 | 0.993 | 1 |
| Inppl1 | 0.000260554 | 0.374121032 | 1 | 0.814 | 1 |
| Ikbkap | 0.000267877 | 0.345736855 | 1 | 1 | 1 |
| Atxn2l | 0.000272304 | 0.487472762 | 1 | 0.986 | 1 |
| Vars | 0.000273983 | 0.811504338 | 1 | 1 | 1 |
| Pard6g | 0.000274333 | 0.308383407 | 1 | 0.736 | 1 |
| Kif4 | 0.000281135 | 0.262044682 | 0.75 | 0.9 | 1 |
| B3gnt5 | 0.000285694 | 0.262541634 | 1 | 0.614 | 1 |
| Hsph1 | 0.000285768 | 0.676164337 | 1 | 1 | 1 |
| Ptcd1 | 0.00030143 | 0.282068773 | 0.75 | 0.764 | 1 |
| Trappc6b | 0.000312621 | 0.688288245 | 1 | 0.95 | 1 |
| Ccnj | 0.00031412 | 0.294157885 | 1 | 0.879 | 1 |
| Ddx49 | 0.00031665 | 0.250763481 | 0.75 | 0.936 | 1 |

| | | | | | |
|---|---|---|---|---|---|
| Msh6 | 0.000319649 | 0.505567401 | 1 | 0.993 | 1 |
| Cyp51 | 0.000325519 | 0.769575151 | 1 | 1 | 1 |
| Blm | 0.000326459 | 0.396567431 | 1 | 0.943 | 1 |
| Pik3r3 | 0.0003271 | 0.34098209 | 1 | 0.75 | 1 |
| Ist1 | 0.000329959 | 0.873047915 | 1 | 1 | 1 |
| Slc38a10 | 0.000334404 | 0.542292643 | 1 | 0.971 | 1 |
| Nt5dc2 | 0.000335563 | 0.6229096 | 1 | 0.986 | 1 |
| Gtf2i | 0.000335945 | 0.47166262 | 1 | 0.993 | 1 |
| Uba1 | 0.00033668 | 0.813145326 | 1 | 1 | 1 |
| Phldb1 | 0.00033677 | 0.268352834 | 1 | 0.736 | 1 |
| Hcfc1 | 0.000336971 | 0.455639651 | 1 | 1 | 1 |
| Nhlrc2 | 0.000348034 | 0.30742399 | 1 | 0.964 | 1 |
| Zfp866 | 0.000352252 | 0.286134648 | 1 | 0.871 | 1 |
| Prss23 | 0.000360348 | 0.365337263 | 1 | 0.643 | 1 |
| Psip1 | 0.000361259 | 0.347081959 | 1 | 1 | 1 |
| Armc5 | 0.000363952 | 0.265343207 | 0.75 | 0.707 | 1 |
| Zfp719 | 0.000365132 | 0.281700126 | 0.5 | 0.657 | 1 |
| Ccdc91 | 0.000367674 | 0.317648269 | 1 | 0.729 | 1 |
| Sh3bgrl | 0.000380777 | 0.687949539 | 1 | 0.986 | 1 |
| 2610318N | 0.000390722 | 0.685843069 | 0.75 | 0.95 | 1 |
| Atp1b3 | 0.00040084 | 0.792368288 | 1 | 0.993 | 1 |
| Nup210 | 0.00040698 | 0.275246348 | 1 | 0.921 | 1 |
| Sdad1 | 0.000422752 | 0.284187416 | 1 | 0.979 | 1 |
| Pef1 | 0.000428713 | 0.53524109 | 1 | 0.943 | 1 |
| Tmem39b | 0.000441493 | 0.378287698 | 0.75 | 0.7 | 1 |
| Drosha | 0.000461389 | 0.271927312 | 1 | 0.993 | 1 |
| Pigq | 0.000465594 | 0.256969845 | 1 | 0.957 | 1 |
| Pcsk7 | 0.000469048 | 0.359442997 | 0.75 | 0.743 | 1 |
| Wsb1 | 0.00049097 | 0.808352042 | 1 | 0.986 | 1 |
| Luzp1 | 0.000494462 | 0.29802495 | 1 | 0.986 | 1 |
| Dtnbp1 | 0.000495978 | 0.366224865 | 0.25 | 0.55 | 1 |
| Tmub2 | 0.000501009 | 0.571569579 | 1 | 0.921 | 1 |
| Phactr4 | 0.000503673 | 0.317499746 | 1 | 0.979 | 1 |
| Nkiras2 | 0.000509741 | 0.347398137 | 0.75 | 0.879 | 1 |
| Nedd1 | 0.000517231 | 0.48798446 | 1 | 0.971 | 1 |
| Uhrf1 | 0.000520814 | 0.873614034 | 1 | 1 | 1 |
| Aagab | 0.000526796 | 0.678117752 | 1 | 0.943 | 1 |
| Sgta | 0.000534241 | 0.552133508 | 1 | 0.993 | 1 |
| Uqcc1 | 0.000536079 | 0.371419052 | 1 | 0.986 | 1 |
| Chmp6 | 0.000538184 | 0.48877121 | 1 | 0.971 | 1 |
| Sel1l | 0.000562835 | 0.273935582 | 1 | 0.986 | 1 |
| Dalrd3 | 0.00056601 | 0.363507892 | 1 | 0.843 | 1 |
| Cherp | 0.000568923 | 0.517337172 | 1 | 0.986 | 1 |
| Adam17 | 0.000575875 | 0.257683039 | 1 | 1 | 1 |
| Tfip11 | 0.000594609 | 0.307957266 | 1 | 0.893 | 1 |
| Abhd12 | 0.000600306 | 0.421448403 | 1 | 0.964 | 1 |
| Aldh18a1 | 0.000609566 | 0.390329701 | 1 | 0.964 | 1 |
| Anks3 | 0.000614315 | 0.379509352 | 0.75 | 0.9 | 1 |
| Ctr9 | 0.000616732 | 0.396919935 | 1 | 0.993 | 1 |
| Dck | 0.000619928 | 0.582420438 | 1 | 0.986 | 1 |
| Ctns | 0.000628868 | 0.25397426 | 1 | 0.714 | 1 |
| Bbs2 | 0.000653543 | 0.252855278 | 1 | 0.586 | 1 |
| Srrm2 | 0.000656544 | 0.45564765 | 1 | 1 | 1 |
| Emc10 | 0.000667667 | 0.677223428 | 1 | 1 | 1 |
| Man2b1 | 0.000671696 | 0.422830949 | 1 | 0.957 | 1 |
| Pls3 | 0.000672592 | 0.527267628 | 0.75 | 0.864 | 1 |
| Hmg20a | 0.000673737 | 0.505697529 | 1 | 0.964 | 1 |
| Dhx29 | 0.000673782 | 0.301948535 | 0.75 | 0.893 | 1 |

| | | | | | |
|---|---|---|---|---|---|
| 1110059E2 | 0.000679595 | 0.443700281 | 1 | 0.936 | 1 |
| Rce1 | 0.00069276 | 0.687943024 | 1 | 0.936 | 1 |
| Wars | 0.000701672 | 0.652824164 | 1 | 1 | 1 |
| Yipf3 | 0.000708098 | 0.553671369 | 1 | 0.964 | 1 |
| Sdc1 | 0.000723598 | 0.729180227 | 1 | 1 | 1 |
| Tcf19 | 0.000731521 | 0.650692611 | 1 | 0.757 | 1 |
| Hirip3 | 0.000737971 | 0.414571087 | 0.75 | 0.879 | 1 |
| Eif2s3x | 0.000742548 | 0.61223139 | 1 | 0.993 | 1 |
| Ddx20 | 0.000763551 | 0.282786812 | 1 | 0.986 | 1 |
| Pcdh18 | 0.000766587 | 0.30539309 | 0.75 | 0.407 | 1 |
| Srsf9 | 0.000789923 | 0.478562566 | 1 | 0.986 | 1 |
| Wee1 | 0.000792447 | 0.389719543 | 0.75 | 0.914 | 1 |
| B4galnt4 | 0.000792928 | 0.289839166 | 1 | 0.621 | 1 |
| Smpdl3a | 0.000796615 | 0.444081509 | 0.75 | 0.421 | 1 |
| 2610301B2 | 0.000809837 | 0.253138221 | 0.75 | 0.786 | 1 |
| Wbp1 | 0.000826816 | 0.719333624 | 1 | 0.929 | 1 |
| Zfp811 | 0.000848952 | 0.65150732 | 1 | 0.386 | 1 |
| Ciao1 | 0.000885454 | 0.351907046 | 1 | 0.971 | 1 |
| Get4 | 0.000888971 | 0.409360306 | 1 | 1 | 1 |
| Rad51ap1 | 0.000892899 | 0.351647166 | 1 | 0.971 | 1 |
| Ackr3 | 0.000893589 | 0.658178833 | 1 | 0.614 | 1 |
| Adprh | 0.000900091 | 0.882035929 | 1 | 0.993 | 1 |
| Trim32 | 0.000910418 | 0.466873434 | 1 | 0.9 | 1 |
| Mpv17l2 | 0.00091483 | 0.415747916 | 0.5 | 0.693 | 1 |
| Tmem234 | 0.000932764 | 0.25890494 | 1 | 0.993 | 1 |
| Zfp830 | 0.000950572 | 0.562739311 | 1 | 0.886 | 1 |
| Trafd1 | 0.000964871 | 0.289609937 | 0.75 | 0.821 | 1 |
| 4930452B0 | 0.00096962 | 0.362607138 | 0.75 | 0.621 | 1 |
| Hspa4l | 0.000991086 | 0.279112029 | 0.5 | 0.579 | 1 |
| Calu | 0.001015684 | 0.544973729 | 1 | 1 | 1 |
| Tcea1 | 0.001047104 | 0.284634343 | 1 | 1 | 1 |
| Zmynd8 | 0.001050631 | 0.277736533 | 1 | 1 | 1 |
| Vamp3 | 0.001068547 | 0.404686017 | 1 | 1 | 1 |
| Tmem43 | 0.00107195 | 0.386079897 | 1 | 0.879 | 1 |
| Tbpl1 | 0.001094971 | 0.461669128 | 1 | 0.95 | 1 |
| Ddx24 | 0.001125899 | 0.469384163 | 1 | 0.993 | 1 |
| Taf5l | 0.001127328 | 0.373685109 | 0.75 | 0.921 | 1 |
| Azin1 | 0.001131356 | 0.289376235 | 1 | 1 | 1 |
| Srebf2 | 0.001141669 | 0.628882673 | 1 | 0.986 | 1 |
| Lppr3 | 0.001152452 | 0.650099681 | 1 | 0.814 | 1 |
| Smarca4 | 0.001159348 | 0.477790197 | 1 | 0.993 | 1 |
| Wls | 0.001200932 | 0.438693775 | 1 | 0.829 | 1 |
| Rbm14 | 0.001201201 | 0.443336676 | 1 | 1 | 1 |
| Nfe2l1 | 0.001208885 | 0.307404207 | 1 | 0.979 | 1 |
| Sh3yl1 | 0.001259419 | 0.296573402 | 0.75 | 0.536 | 1 |
| Add3 | 0.00128651 | 0.268437397 | 1 | 0.986 | 1 |
| Glt8d1 | 0.001299292 | 0.268780047 | 1 | 0.907 | 1 |
| Alg8 | 0.001308612 | 0.448371203 | 1 | 1 | 1 |
| Slc25a30 | 0.001347153 | 0.266273869 | 0.75 | 0.343 | 1 |
| Cyb5r3 | 0.001397265 | 0.626345839 | 1 | 1 | 1 |
| Foxa2 | 0.001398426 | 0.509232443 | 1 | 1 | 1 |
| Fam83d | 0.001485633 | 0.295686449 | 1 | 0.714 | 1 |
| Lrrc8a | 0.001516011 | 0.393879452 | 1 | 0.979 | 1 |
| Tmem222 | 0.001525659 | 0.317332642 | 1 | 0.979 | 1 |
| G3bp1 | 0.00154748 | 0.476367164 | 1 | 1 | 1 |
| Entpd6 | 0.001582127 | 0.335137894 | 1 | 0.986 | 1 |
| Ube3b | 0.001586579 | 0.26883196 | 1 | 0.979 | 1 |
| Ppil2 | 0.001589944 | 0.411749071 | 1 | 0.993 | 1 |

| | | | | | |
|---|---|---|---|---|---|
| Uchl1 | 0.001607472 | 1.170260811 | 1 | 0.75 | 1 |
| P3h4 | 0.00161105 | 0.30185172 | 1 | 0.714 | 1 |
| Ogfr | 0.001629353 | 0.475176734 | 1 | 1 | 1 |
| Mfsd5 | 0.001661217 | 0.279022602 | 1 | 0.707 | 1 |
| Nol6 | 0.001669381 | 0.315200812 | 1 | 0.929 | 1 |
| Polr2b | 0.001687408 | 0.482581893 | 1 | 1 | 1 |
| Cklf | 0.001728943 | 0.292627723 | 0.75 | 0.721 | 1 |
| Mapk14 | 0.00179563 | 0.281759286 | 0.75 | 0.829 | 1 |
| Pdhx | 0.001804423 | 0.284262151 | 1 | 0.95 | 1 |
| Aco2 | 0.001825133 | 0.57801935 | 1 | 1 | 1 |
| Vgll4 | 0.001834457 | 0.393019021 | 1 | 0.95 | 1 |
| Hdhd2 | 0.001910287 | 0.283286914 | 1 | 0.964 | 1 |
| Usp39 | 0.001911414 | 0.708441781 | 1 | 0.986 | 1 |
| Cast | 0.001955365 | 0.274179929 | 1 | 0.721 | 1 |
| Cog4 | 0.001965417 | 0.388103421 | 1 | 0.964 | 1 |
| Rcn2 | 0.001979679 | 0.53867879 | 1 | 1 | 1 |
| Farsa | 0.001995518 | 0.509565257 | 1 | 1 | 1 |
| Igf1r | 0.002022865 | 0.404525651 | 1 | 0.729 | 1 |
| E2f5 | 0.002042729 | 0.466908507 | 1 | 0.893 | 1 |
| Tox4 | 0.002083766 | 0.291899309 | 1 | 0.986 | 1 |
| Fars2 | 0.002086069 | 0.254401932 | 0.75 | 0.879 | 1 |
| Sirt3 | 0.002086175 | 0.419263796 | 0.75 | 0.757 | 1 |
| Snf8 | 0.002109974 | 0.312954908 | 1 | 1 | 1 |
| Agpat6 | 0.00213818 | 0.589032379 | 1 | 1 | 1 |
| Armcx1 | 0.002181816 | 0.422066094 | 1 | 0.75 | 1 |
| Mtmr4 | 0.002191167 | 0.257499743 | 1 | 0.964 | 1 |
| Btd | 0.002229909 | 0.300636985 | 1 | 0.629 | 1 |
| Tmem132a | 0.002239454 | 0.497711762 | 1 | 0.95 | 1 |
| Sf1 | 0.002301003 | 0.360901435 | 1 | 0.993 | 1 |
| Ipp | 0.002317908 | 0.458010332 | 1 | 0.736 | 1 |
| Hadha | 0.002405341 | 0.292747721 | 1 | 1 | 1 |
| Zfp212 | 0.002437141 | 0.358459514 | 1 | 0.879 | 1 |
| Psme3 | 0.002464882 | 0.411309254 | 1 | 0.964 | 1 |
| Slc37a4 | 0.002469674 | 0.539752171 | 0.75 | 0.821 | 1 |
| Kiz | 0.002556057 | 0.342700637 | 1 | 0.643 | 1 |
| Ogt | 0.002559248 | 0.563070257 | 1 | 1 | 1 |
| Ash2l | 0.002587751 | 0.418568069 | 1 | 0.993 | 1 |
| Trip10 | 0.002631892 | 0.473437556 | 1 | 0.943 | 1 |
| Usp21 | 0.002635986 | 0.315725333 | 1 | 0.957 | 1 |
| Pxdn | 0.002664443 | 0.502832425 | 1 | 0.964 | 1 |
| Ivns1abp | 0.002731818 | 0.601518399 | 1 | 1 | 1 |
| Rbm38 | 0.002735071 | 0.375643487 | 1 | 0.986 | 1 |
| 9530068E( | 0.002756372 | 0.61358971 | 1 | 1 | 1 |
| Bag5 | 0.002773205 | 0.304796889 | 1 | 0.886 | 1 |
| Fubp1 | 0.002823584 | 0.314317523 | 1 | 1 | 1 |
| Dpysl2 | 0.002826612 | 0.976132358 | 1 | 0.993 | 1 |
| Mia3 | 0.002839689 | 0.254275193 | 0.75 | 0.971 | 1 |
| Supt5 | 0.002859987 | 0.56327703 | 1 | 1 | 1 |
| Dmap1 | 0.002906055 | 0.450974014 | 1 | 0.95 | 1 |
| Ptprf | 0.002989928 | 0.429995752 | 1 | 1 | 1 |
| Exo5 | 0.003019337 | 0.473988932 | 1 | 0.736 | 1 |
| Ppwd1 | 0.003022065 | 0.33842098 | 1 | 0.986 | 1 |
| Hip1 | 0.003031238 | 0.290238967 | 1 | 0.814 | 1 |
| Slc29a4 | 0.003076807 | 0.394079854 | 0.75 | 0.45 | 1 |
| Ccdc117 | 0.003089345 | 0.46585105 | 1 | 0.936 | 1 |
| Isca2 | 0.003094848 | 0.558743498 | 0.75 | 0.943 | 1 |
| Mcm2 | 0.003152023 | 0.843911028 | 1 | 1 | 1 |
| Cpsf4 | 0.003166914 | 0.335058815 | 1 | 0.943 | 1 |

| | | | | | |
|---|---|---|---|---|---|
| Slc44a2 | 0.003178122 | 0.532447488 | 1 | 0.743 | 1 |
| Cs | 0.003186969 | 0.525999775 | 1 | 1 | 1 |
| Dcun1d4 | 0.00325028 | 0.345500407 | 1 | 0.921 | 1 |
| Myg1 | 0.00332596 | 0.400159305 | 1 | 0.971 | 1 |
| Atp6ap1 | 0.003382423 | 0.614409586 | 1 | 1 | 1 |
| Prdx4 | 0.003437875 | 0.741733928 | 1 | 1 | 1 |
| Ddx56 | 0.003452091 | 0.722797127 | 1 | 1 | 1 |
| Med22 | 0.003497501 | 0.343531538 | 1 | 0.957 | 1 |
| Sgpl1 | 0.003511213 | 0.373891699 | 1 | 0.993 | 1 |
| Serpinh1 | 0.003520389 | 0.763735366 | 1 | 1 | 1 |
| Prkab1 | 0.003705587 | 0.474267422 | 1 | 0.957 | 1 |
| Csnk1a1 | 0.003753901 | 0.361567348 | 1 | 1 | 1 |
| Sra1 | 0.003910921 | 0.297570413 | 1 | 1 | 1 |
| Ctcf | 0.004016001 | 0.2620927 | 1 | 0.957 | 1 |
| C1galt1c1 | 0.004086717 | 0.479522968 | 1 | 0.686 | 1 |
| Tubgcp2 | 0.004099131 | 0.47590483 | 1 | 0.986 | 1 |
| Gtf2a1 | 0.004124896 | 0.351896949 | 1 | 0.993 | 1 |
| Hjurp | 0.00413883 | 0.438045055 | 1 | 1 | 1 |
| Thrap3 | 0.004146803 | 0.280424741 | 1 | 1 | 1 |
| Mcrs1 | 0.004214036 | 0.601412851 | 1 | 0.979 | 1 |
| Ethe1 | 0.004256292 | 0.307904762 | 0.75 | 0.686 | 1 |
| Morc2a | 0.004317574 | 0.333338253 | 1 | 0.971 | 1 |
| St5 | 0.004400247 | 0.268308928 | 1 | 0.921 | 1 |
| Cpsf3l | 0.004428038 | 0.382666204 | 1 | 0.971 | 1 |
| Tor1a | 0.004450871 | 0.582280924 | 1 | 0.979 | 1 |
| Cyb5b | 0.004451776 | 0.493269831 | 1 | 0.993 | 1 |
| Sil1 | 0.004468414 | 0.385289074 | 0.75 | 0.957 | 1 |
| Ogdhl | 0.004502614 | 0.296651152 | 1 | 0.579 | 1 |
| Ampd2 | 0.004604249 | 0.29772633 | 1 | 0.936 | 1 |
| Fam109a | 0.004628397 | 0.43798608 | 1 | 0.836 | 1 |
| Rbbp8 | 0.004797995 | 0.306270104 | 1 | 0.907 | 1 |
| Arhgap28 | 0.004831317 | 0.343999873 | 1 | 0.657 | 1 |
| Zfp397 | 0.004837905 | 0.258215819 | 1 | 0.936 | 1 |
| Mgme1 | 0.004865359 | 0.29523678 | 0.75 | 0.929 | 1 |
| Plcg1 | 0.004873799 | 0.300552342 | 1 | 0.943 | 1 |
| Flna | 0.005138997 | 0.515799066 | 1 | 0.964 | 1 |
| Tmem159 | 0.005163587 | 0.363904732 | 0.75 | 0.45 | 1 |
| Nucb2 | 0.005217959 | 0.311844286 | 0.75 | 0.557 | 1 |
| Angptl2 | 0.005322405 | 0.306990661 | 1 | 0.721 | 1 |
| Dnmt1 | 0.005368971 | 0.35986258 | 1 | 0.993 | 1 |
| Unc119b | 0.00537174 | 0.437135293 | 1 | 0.993 | 1 |
| Mlec | 0.005383354 | 0.470219306 | 1 | 1 | 1 |
| Capn10 | 0.005424334 | 0.336574118 | 1 | 0.85 | 1 |
| Sptlc1 | 0.005488637 | 0.422486099 | 1 | 0.957 | 1 |
| Gpr180 | 0.005516233 | 0.277684878 | 1 | 0.836 | 1 |
| Cdk16 | 0.0056217 | 0.493956104 | 1 | 0.993 | 1 |
| Tusc2 | 0.005887245 | 0.482183063 | 1 | 0.943 | 1 |
| Tcn2 | 0.005935467 | 0.482912971 | 0.75 | 0.793 | 1 |
| Ldlrad3 | 0.005956287 | 0.254010913 | 1 | 0.957 | 1 |
| Fads2 | 0.00598754 | 0.830703537 | 1 | 0.986 | 1 |
| Psmd8 | 0.005991314 | 0.702548718 | 1 | 1 | 1 |
| Zfp131 | 0.005994495 | 0.405998902 | 1 | 0.971 | 1 |
| Nnat | 0.006169314 | 1.239916744 | 1 | 0.914 | 1 |
| Slc29a3 | 0.006181133 | 0.263022234 | 0.75 | 0.629 | 1 |
| Cuta | 0.006243151 | 0.628808106 | 0.75 | 0.993 | 1 |
| Syngr2 | 0.006336552 | 0.621652521 | 1 | 0.993 | 1 |
| Cdk20 | 0.006425311 | 0.399782371 | 0.5 | 0.557 | 1 |
| Rhno1 | 0.006428096 | 0.386235867 | 1 | 0.993 | 1 |

| | | | | | |
|---|---|---|---|---|---|
| Nek4 | 0.006524967 | 0.306146769 | 0.75 | 0.771 | 1 |
| Chmp7 | 0.006617719 | 0.486045913 | 1 | 0.95 | 1 |
| Sec61a1 | 0.006817716 | 0.488917503 | 1 | 1 | 1 |
| Tpp1 | 0.006901119 | 0.445604521 | 1 | 0.993 | 1 |
| Derl1 | 0.006999532 | 0.576643463 | 1 | 0.993 | 1 |
| Chmp1a | 0.007015243 | 0.332353146 | 1 | 0.986 | 1 |
| Dph7 | 0.007017409 | 0.331399387 | 1 | 0.857 | 1 |
| Cdc16 | 0.007204361 | 0.514821039 | 1 | 0.993 | 1 |
| Tsr2 | 0.007273826 | 0.333938594 | 1 | 0.979 | 1 |
| Actn4 | 0.007334928 | 0.581742646 | 1 | 1 | 1 |
| Arhgdia | 0.007371912 | 0.42512861 | 1 | 1 | 1 |
| Kctd15 | 0.007380905 | 0.333513257 | 0.75 | 0.657 | 1 |
| Rrn3 | 0.007390159 | 0.441229391 | 1 | 1 | 1 |
| Tusc3 | 0.007411386 | 0.595312776 | 1 | 0.979 | 1 |
| Ap1s1 | 0.007446189 | 0.546541472 | 1 | 0.993 | 1 |
| Plxnb2 | 0.007452644 | 0.267017025 | 1 | 0.971 | 1 |
| EGFP | 0.007505786 | 0.613518526 | 1 | 1 | 1 |
| Laptm4b | 0.007523205 | 0.865025583 | 1 | 1 | 1 |
| Vmp1 | 0.007547697 | 0.333613817 | 1 | 0.993 | 1 |
| Srpr | 0.007634738 | 0.579813955 | 1 | 1 | 1 |
| Arhgef39 | 0.00769981 | 0.293780303 | 0.75 | 0.636 | 1 |
| Rbbp4 | 0.0077013 | 0.297831664 | 1 | 1 | 1 |
| Pcmt1 | 0.007761189 | 0.3357069 | 1 | 0.993 | 1 |
| Nap1l4 | 0.007853055 | 0.515435599 | 1 | 1 | 1 |
| Samm50 | 0.007855235 | 0.276694824 | 0.75 | 1 | 1 |
| Tmem214 | 0.007855242 | 0.544925379 | 1 | 0.993 | 1 |
| Chaf1a | 0.007999897 | 0.444367262 | 1 | 0.95 | 1 |
| Alkbh2 | 0.008072508 | 0.285134344 | 0.75 | 0.636 | 1 |
| Sugp1 | 0.008260472 | 0.435590517 | 1 | 0.957 | 1 |
| Lpcat1 | 0.00839433 | 0.331093106 | 1 | 0.979 | 1 |
| Dnaaf2 | 0.008404326 | 0.316988826 | 1 | 0.964 | 1 |
| Cfap20 | 0.00845503 | 0.494046852 | 1 | 0.979 | 1 |
| Gtf3c5 | 0.008513778 | 0.32667193 | 1 | 0.957 | 1 |
| Usp5 | 0.008517238 | 0.349776891 | 1 | 0.993 | 1 |
| Cenph | 0.008533648 | 0.412030393 | 1 | 0.993 | 1 |
| Gdpd5 | 0.008580572 | 0.302715121 | 1 | 0.507 | 1 |
| Stk3 | 0.008601524 | 0.288741828 | 1 | 0.886 | 1 |
| Agpat1 | 0.008660945 | 0.347103388 | 1 | 0.993 | 1 |
| Luc7l | 0.008816645 | 0.273583222 | 1 | 1 | 1 |
| Pinx1 | 0.008971433 | 0.569517922 | 1 | 0.843 | 1 |
| Nme6 | 0.009005302 | 0.474408153 | 1 | 0.986 | 1 |
| Aurkb | 0.009037409 | 0.617035319 | 1 | 0.957 | 1 |
| Gramd1a | 0.009242235 | 0.359338864 | 1 | 0.964 | 1 |
| Palm | 0.009325181 | 0.341110945 | 0.75 | 0.907 | 1 |
| Pigx | 0.009344395 | 0.411908669 | 1 | 1 | 1 |
| Nemf | 0.009367655 | 0.400785749 | 1 | 0.993 | 1 |
| Zfp410 | 0.009459959 | 0.321657554 | 1 | 0.986 | 1 |
| Ddx18 | 0.009460868 | 0.440144442 | 1 | 1 | 1 |
| Map2k5 | 0.009484448 | 0.469023481 | 1 | 0.936 | 1 |
| Pigt | 0.009557882 | 0.317265088 | 1 | 0.786 | 1 |
| Tjap1 | 0.009575521 | 0.515265747 | 1 | 0.929 | 1 |
| Id3 | 0.009613259 | 0.935319288 | 1 | 0.993 | 1 |
| Ddb1 | 0.009662357 | 0.613401206 | 1 | 1 | 1 |
| Klhl28 | 0.009729533 | 0.257237225 | 1 | 0.65 | 1 |
| Puf60 | 0.009753032 | 0.272170884 | 1 | 1 | 1 |
| Gnai2 | 0.009809786 | 0.284109708 | 1 | 0.986 | 1 |
| Cdca2 | 0.009870085 | 0.472931739 | 1 | 0.986 | 1 |
| Tarbp2 | 0.009928777 | 0.258345741 | 1 | 1 | 1 |

| | | | | | |
|---|---|---|---|---|---|
| Actn2 | 2.91E-106 | 0.527711624 | 0.966 | 0.696 | 3.75E-102 |
| Osr1 | 1.05E-83 | 0.91820985 | 0.931 | 0.461 | 1.35E-79 |
| Mybpc3 | 2.72E-80 | 0.630340219 | 0.966 | 0.522 | 3.51E-76 |
| Capn6 | 4.91E-80 | 0.956473881 | 1 | 0.713 | 6.34E-76 |
| Myh7 | 6.22E-77 | 0.582281486 | 1 | 0.53 | 8.02E-73 |
| Pitx2 | 1.71E-55 | 0.431959371 | 0.793 | 0.391 | 2.21E-51 |
| Krt7 | 2.34E-55 | 0.603167073 | 1 | 0.887 | 3.01E-51 |
| Myl3 | 1.76E-46 | 0.396309815 | 0.862 | 0.4 | 2.27E-42 |
| Hoxc4 | 3.24E-46 | 0.304928137 | 0.931 | 0.243 | 4.18E-42 |
| Myl2 | 1.29E-43 | 0.305028642 | 0.862 | 0.33 | 1.67E-39 |
| Myh6 | 5.10E-43 | 1.142685197 | 1 | 0.539 | 6.58E-39 |
| Bcam | 8.14E-43 | 0.757113315 | 1 | 0.852 | 1.05E-38 |
| Isx | 1.91E-40 | 0.313598459 | 0.655 | 0.174 | 2.47E-36 |
| Gprc5a | 1.65E-39 | 0.368468964 | 0.655 | 0.27 | 2.13E-35 |
| Tpm1 | 1.20E-38 | 1.074996051 | 1 | 1 | 1.55E-34 |
| Slc2a3 | 3.99E-38 | 1.436237729 | 1 | 1 | 5.14E-34 |
| Raet1e | 2.57E-37 | 0.454434451 | 0.862 | 0.409 | 3.32E-33 |
| Smoc2 | 2.12E-32 | 0.262255065 | 0.552 | 0.357 | 2.73E-28 |
| Ankrd1 | 1.91E-31 | 0.906811686 | 0.931 | 0.443 | 2.46E-27 |
| Tmprss2 | 6.23E-31 | 1.068023435 | 1 | 0.765 | 8.04E-27 |
| Nkx2-6 | 2.17E-30 | 0.390680437 | 0.69 | 0.183 | 2.80E-26 |
| Rfx6 | 5.47E-30 | 0.498648748 | 1 | 0.391 | 7.05E-26 |
| Ifitm1 | 7.74E-30 | 1.09191484 | 1 | 0.661 | 9.98E-26 |
| Myl4 | 2.66E-29 | 0.292384575 | 0.897 | 0.643 | 3.43E-25 |
| Nepn | 1.59E-28 | 2.207330345 | 1 | 0.626 | 2.06E-24 |
| Peg3 | 3.27E-28 | 0.999733561 | 1 | 1 | 4.22E-24 |
| Rdh10 | 8.23E-28 | 0.268625525 | 0.931 | 0.557 | 1.06E-23 |
| Vnn1 | 2.33E-27 | 0.549105953 | 0.897 | 0.426 | 3.01E-23 |
| Klf5 | 2.38E-27 | 0.449301296 | 0.897 | 0.278 | 3.06E-23 |
| Slc5a5 | 9.53E-27 | 0.320517178 | 0.724 | 0.452 | 1.23E-22 |
| Arg1 | 1.65E-26 | 1.794486298 | 1 | 0.861 | 2.13E-22 |
| Gata4 | 4.48E-26 | 0.584250866 | 1 | 0.47 | 5.77E-22 |
| Cyp2s1 | 4.00E-25 | 0.29250467 | 0.862 | 0.391 | 5.16E-21 |
| Car2 | 9.39E-25 | 0.346081988 | 0.69 | 0.287 | 1.21E-20 |
| Fam183b | 1.38E-24 | 1.285318101 | 1 | 0.435 | 1.78E-20 |
| Agrn | 6.79E-24 | 0.405445651 | 1 | 0.965 | 8.75E-20 |
| Gmpr | 8.77E-24 | 0.27102156 | 0.759 | 0.287 | 1.13E-19 |
| Cdh1 | 1.51E-22 | 0.564769909 | 0.966 | 0.774 | 1.94E-18 |
| Ociad2 | 2.13E-22 | 0.600808874 | 1 | 0.957 | 2.75E-18 |
| App | 2.74E-22 | 0.906556601 | 1 | 0.73 | 3.53E-18 |
| Vcan | 6.03E-22 | 0.350458427 | 1 | 0.852 | 7.78E-18 |
| Epb4.1l3 | 7.26E-22 | 0.729239617 | 1 | 0.922 | 9.36E-18 |
| Csrp3 | 8.57E-22 | 0.312405427 | 0.724 | 0.33 | 1.10E-17 |
| Igfbp5 | 1.32E-21 | 0.615881964 | 1 | 0.722 | 1.70E-17 |
| Tnnc1 | 1.48E-21 | 0.596616818 | 0.931 | 0.722 | 1.91E-17 |
| Dpp4 | 2.40E-21 | 0.31682841 | 0.966 | 0.843 | 3.10E-17 |
| Cd24a | 4.31E-21 | 1.220773411 | 1 | 1 | 5.55E-17 |
| Gpx3 | 6.53E-21 | 1.407381165 | 1 | 0.843 | 8.42E-17 |
| Entpd3 | 9.41E-21 | 0.279783554 | 0.793 | 0.165 | 1.21E-16 |
| Itga3 | 1.06E-20 | 0.333993321 | 0.966 | 0.574 | 1.37E-16 |
| Lrpap1 | 2.33E-20 | 0.745720933 | 1 | 1 | 3.01E-16 |
| Rarb | 3.71E-20 | 0.516279233 | 1 | 0.73 | 4.78E-16 |
| Sema6a | 3.94E-20 | 0.297982483 | 1 | 0.643 | 5.08E-16 |
| St14 | 5.47E-20 | 0.571630579 | 1 | 0.565 | 7.05E-16 |
| Tspan8 | 9.34E-20 | 0.389615236 | 0.552 | 0.235 | 1.20E-15 |
| Tdh | 1.20E-19 | 0.427913177 | 0.724 | 0.304 | 1.55E-15 |
| Prss8 | 1.36E-19 | 0.75942102 | 1 | 0.8 | 1.75E-15 |
| Podxl | 1.56E-19 | 0.569776867 | 1 | 0.774 | 2.01E-15 |

| | | | | | |
|---|---|---|---|---|---|
| Gja1 | 1.60E-19 | 0.880778808 | 1 | 0.687 | 2.06E-15 |
| Cldn8 | 2.01E-19 | 0.327136256 | 0.724 | 0.2 | 2.59E-15 |
| Mreg | 3.21E-19 | 0.501714709 | 0.966 | 0.948 | 4.14E-15 |
| Shh | 1.42E-18 | 0.442292456 | 0.966 | 0.33 | 1.83E-14 |
| Cryab | 1.52E-18 | 0.250263406 | 0.621 | 0.287 | 1.96E-14 |
| Igsf9 | 1.56E-18 | 0.551744355 | 1 | 0.765 | 2.01E-14 |
| Tmem88 | 1.59E-18 | 0.269218617 | 0.724 | 0.391 | 2.05E-14 |
| Slc39a10 | 2.20E-18 | 0.345666492 | 1 | 0.991 | 2.83E-14 |
| Hoxa1 | 2.36E-18 | 0.375390487 | 0.862 | 0.235 | 3.05E-14 |
| Flna | 3.35E-18 | 0.687467514 | 1 | 0.957 | 4.32E-14 |
| Palm3 | 3.67E-18 | 0.262093611 | 0.931 | 0.504 | 4.73E-14 |
| Efna1 | 4.71E-18 | 0.485858084 | 1 | 0.452 | 6.07E-14 |
| Tff3 | 7.17E-18 | 0.886231634 | 0.862 | 0.626 | 9.24E-14 |
| Obsl1 | 9.90E-18 | 0.285307544 | 1 | 0.983 | 1.28E-13 |
| Alpl | 2.21E-17 | 0.273720178 | 0.862 | 0.504 | 2.85E-13 |
| Sema3c | 2.23E-17 | 0.309519444 | 1 | 0.548 | 2.87E-13 |
| Gprc5c | 2.52E-17 | 0.476455993 | 1 | 0.965 | 3.25E-13 |
| Cldn4 | 2.57E-17 | 0.864615579 | 1 | 0.565 | 3.32E-13 |
| Folr1 | 6.25E-17 | 0.253965532 | 0.517 | 0.217 | 8.06E-13 |
| Cldn6 | 7.52E-17 | 1.405351858 | 1 | 0.983 | 9.69E-13 |
| Jup | 9.78E-17 | 0.256827865 | 0.966 | 0.965 | 1.26E-12 |
| Col4a1 | 1.03E-16 | 0.690527279 | 1 | 0.922 | 1.32E-12 |
| Acta2 | 1.10E-16 | 0.669683129 | 1 | 1 | 1.42E-12 |
| Cthrc1 | 2.55E-16 | 0.300008718 | 0.828 | 0.365 | 3.29E-12 |
| Foxa1 | 3.38E-16 | 0.386241326 | 0.966 | 0.791 | 4.36E-12 |
| Wfdc2 | 5.35E-16 | 0.916614791 | 1 | 0.887 | 6.90E-12 |
| Smtnl2 | 7.29E-16 | 0.360519797 | 1 | 0.939 | 9.40E-12 |
| Lad1 | 7.45E-16 | 0.592969758 | 1 | 0.687 | 9.60E-12 |
| Hmgn3 | 8.67E-16 | 0.504247025 | 0.966 | 0.626 | 1.12E-11 |
| Uchl1 | 2.08E-15 | 0.839495934 | 1 | 0.696 | 2.69E-11 |
| Ifi30 | 2.59E-15 | 0.740070707 | 0.966 | 0.583 | 3.33E-11 |
| Wls | 2.77E-15 | 0.339895235 | 1 | 0.791 | 3.57E-11 |
| Anxa6 | 3.27E-15 | 0.551376754 | 1 | 0.922 | 4.22E-11 |
| Lamb1 | 3.68E-15 | 0.402436788 | 1 | 0.835 | 4.75E-11 |
| Mapre2 | 7.52E-15 | 0.276022583 | 1 | 0.983 | 9.70E-11 |
| Actc1 | 9.71E-15 | 1.377100757 | 1 | 0.878 | 1.25E-10 |
| Dag1 | 1.13E-14 | 0.418681923 | 1 | 0.991 | 1.45E-10 |
| Ap1m2 | 1.71E-14 | 0.752313516 | 1 | 0.583 | 2.21E-10 |
| Prps2 | 1.87E-14 | 0.292858237 | 0.931 | 0.817 | 2.41E-10 |
| Epcam | 2.55E-14 | 1.130438099 | 1 | 0.974 | 3.28E-10 |
| Myl7 | 3.38E-14 | 0.832116984 | 1 | 0.557 | 4.35E-10 |
| Rdm1 | 7.80E-14 | 0.318994655 | 0.759 | 0.391 | 1.01E-09 |
| Gm21814 | 1.26E-13 | 0.519749748 | 0.759 | 0.557 | 1.62E-09 |
| Nrep | 1.30E-13 | 0.825581089 | 1 | 0.991 | 1.67E-09 |
| Ramp3 | 1.62E-13 | 0.33746776 | 0.69 | 0.217 | 2.09E-09 |
| Spint1 | 2.91E-13 | 0.690734272 | 1 | 0.783 | 3.75E-09 |
| Cdh6 | 4.69E-13 | 0.921708405 | 0.966 | 0.591 | 6.05E-09 |
| Dhrs3 | 4.95E-13 | 0.297622369 | 0.966 | 0.817 | 6.38E-09 |
| Col18a1 | 5.12E-13 | 0.68608712 | 1 | 1 | 6.60E-09 |
| Hsd11b2 | 5.18E-13 | 0.551288273 | 1 | 0.513 | 6.68E-09 |
| Lars2 | 5.26E-13 | 0.437703387 | 1 | 1 | 6.78E-09 |
| Fxyd6 | 5.69E-13 | 1.093410248 | 0.966 | 0.739 | 7.34E-09 |
| Efs | 6.83E-13 | 0.254648136 | 0.966 | 0.435 | 8.81E-09 |
| Kcnk1 | 9.20E-13 | 0.446674028 | 1 | 0.504 | 1.19E-08 |
| Spint2 | 1.10E-12 | 0.924105112 | 1 | 1 | 1.41E-08 |
| Dab2 | 1.10E-12 | 0.27575087 | 1 | 0.635 | 1.42E-08 |
| Atp2c2 | 1.77E-12 | 0.276810381 | 0.966 | 0.452 | 2.28E-08 |
| Krt19 | 2.44E-12 | 0.917006636 | 0.966 | 0.678 | 3.14E-08 |

| | | | | | |
|---|---|---|---|---|---|
| Sfrp5 | 2.75E-12 | 1.396521127 | 0.862 | 0.513 | 3.55E-08 |
| Cdh3 | 2.84E-12 | 0.456822515 | 0.966 | 0.713 | 3.67E-08 |
| Fam129b | 3.15E-12 | 0.293492682 | 0.966 | 0.6 | 4.06E-08 |
| Plac8 | 3.22E-12 | 0.299400386 | 0.759 | 0.435 | 4.15E-08 |
| Fam174b | 3.23E-12 | 0.363405093 | 0.931 | 0.574 | 4.16E-08 |
| Lpar3 | 6.12E-12 | 0.366356578 | 0.897 | 0.348 | 7.89E-08 |
| Tmsb4x | 6.29E-12 | 0.631596647 | 1 | 0.922 | 8.10E-08 |
| S1pr2 | 8.20E-12 | 0.336733567 | 0.931 | 0.609 | 1.06E-07 |
| Dbn1 | 8.67E-12 | 0.579510421 | 1 | 1 | 1.12E-07 |
| Calcoco1 | 8.69E-12 | 0.293210782 | 0.966 | 0.626 | 1.12E-07 |
| Nudt10 | 9.28E-12 | 0.253862117 | 1 | 0.557 | 1.20E-07 |
| Pvrl2 | 1.01E-11 | 0.519073336 | 1 | 0.991 | 1.30E-07 |
| 3632451O | 1.22E-11 | 0.438644232 | 0.966 | 0.591 | 1.58E-07 |
| Cnn3 | 1.29E-11 | 0.624205171 | 1 | 1 | 1.66E-07 |
| Amotl2 | 1.50E-11 | 0.345593594 | 1 | 0.983 | 1.93E-07 |
| Pttg1ip | 1.74E-11 | 0.505362154 | 1 | 1 | 2.24E-07 |
| Gm8113 | 1.83E-11 | 0.817000857 | 1 | 1 | 2.36E-07 |
| F11r | 1.84E-11 | 0.750718102 | 1 | 1 | 2.38E-07 |
| Dsc2 | 2.14E-11 | 0.250233849 | 0.966 | 0.783 | 2.75E-07 |
| Raet1d | 2.22E-11 | 0.366025063 | 0.862 | 0.417 | 2.86E-07 |
| Tpm4 | 2.23E-11 | 0.766006575 | 1 | 0.991 | 2.88E-07 |
| Ces1d | 2.58E-11 | 0.289249293 | 0.793 | 0.339 | 3.32E-07 |
| Mmp2 | 3.73E-11 | 0.276709547 | 0.862 | 0.357 | 4.81E-07 |
| Zfp956 | 4.69E-11 | 0.266270738 | 0.931 | 0.635 | 6.04E-07 |
| Tmem123 | 4.91E-11 | 0.576715103 | 1 | 0.965 | 6.33E-07 |
| Cldn9 | 4.98E-11 | 0.442403617 | 0.966 | 0.365 | 6.42E-07 |
| Fbln1 | 5.74E-11 | 0.427603458 | 1 | 1 | 7.40E-07 |
| Rnf167 | 6.26E-11 | 0.375244355 | 1 | 0.957 | 8.07E-07 |
| Nr2f2 | 7.34E-11 | 0.450755443 | 1 | 0.983 | 9.47E-07 |
| Nudt11 | 8.35E-11 | 0.272651017 | 1 | 0.67 | 1.08E-06 |
| Car4 | 8.35E-11 | 1.04999644 | 1 | 1 | 1.08E-06 |
| Plod2 | 8.42E-11 | 0.30256177 | 0.966 | 0.843 | 1.09E-06 |
| Fscn1 | 1.02E-10 | 0.478345149 | 1 | 1 | 1.31E-06 |
| Cnn2 | 1.05E-10 | 0.583400012 | 0.931 | 0.843 | 1.36E-06 |
| Ctsd | 1.19E-10 | 0.67461431 | 1 | 1 | 1.53E-06 |
| Ypel1 | 1.29E-10 | 0.258847707 | 1 | 0.643 | 1.66E-06 |
| Cldn3 | 1.44E-10 | 0.486262324 | 0.931 | 0.443 | 1.86E-06 |
| Shisa4 | 1.59E-10 | 0.319963464 | 0.966 | 0.704 | 2.06E-06 |
| Slc27a3 | 1.66E-10 | 0.466938897 | 1 | 0.843 | 2.14E-06 |
| Mmp11 | 1.92E-10 | 0.550773641 | 1 | 0.713 | 2.47E-06 |
| Marcksl1 | 2.00E-10 | 0.70859638 | 1 | 1 | 2.58E-06 |
| Ddb1 | 2.82E-10 | 0.45463273 | 1 | 1 | 3.64E-06 |
| Gyg | 2.87E-10 | 0.415929322 | 0.966 | 0.965 | 3.69E-06 |
| Trp53i11 | 2.97E-10 | 0.361739568 | 1 | 0.565 | 3.83E-06 |
| Fstl1 | 3.24E-10 | 0.602107662 | 1 | 0.922 | 4.18E-06 |
| Crb3 | 3.41E-10 | 0.540126529 | 1 | 0.765 | 4.39E-06 |
| S100a10 | 3.83E-10 | 0.863724261 | 1 | 0.617 | 4.94E-06 |
| Col4a2 | 4.24E-10 | 0.436328617 | 1 | 0.791 | 5.47E-06 |
| Slc2a1 | 4.90E-10 | 0.859215815 | 1 | 1 | 6.32E-06 |
| Pxdn | 5.15E-10 | 0.355100231 | 0.966 | 0.965 | 6.64E-06 |
| Kcnk16 | 5.28E-10 | 0.428882662 | 0.828 | 0.217 | 6.81E-06 |
| Krt8 | 5.33E-10 | 0.932285024 | 1 | 1 | 6.87E-06 |
| Fblim1 | 6.07E-10 | 0.30205276 | 1 | 0.73 | 7.82E-06 |
| Ezr | 7.14E-10 | 0.546437242 | 1 | 0.896 | 9.21E-06 |
| H2afy2 | 8.20E-10 | 0.619546613 | 1 | 0.991 | 1.06E-05 |
| H3f3a | 8.73E-10 | 0.398054893 | 1 | 1 | 1.13E-05 |
| Armc10 | 9.01E-10 | 0.389784199 | 1 | 0.957 | 1.16E-05 |
| S100a11 | 9.06E-10 | 0.933035372 | 1 | 0.817 | 1.17E-05 |

| | | | | | |
|---|---|---|---|---|---|
| Tspan3 | 9.19E-10 | 0.445159769 | 1 | 1 | 1.18E-05 |
| Pla2g7 | 9.37E-10 | 0.271896527 | 0.828 | 0.478 | 1.21E-05 |
| Nnat | 1.14E-09 | 0.836755563 | 1 | 0.896 | 1.47E-05 |
| Clic1 | 1.27E-09 | 0.589792772 | 1 | 0.87 | 1.64E-05 |
| Fhl2 | 1.36E-09 | 0.437418057 | 0.931 | 0.609 | 1.76E-05 |
| Cldn7 | 1.46E-09 | 0.632926082 | 1 | 0.765 | 1.88E-05 |
| Rcan2 | 1.50E-09 | 0.325586519 | 0.966 | 0.939 | 1.94E-05 |
| Igfbp2 | 1.57E-09 | 0.963298567 | 1 | 0.922 | 2.03E-05 |
| Lasp1 | 2.15E-09 | 0.291766884 | 1 | 0.948 | 2.78E-05 |
| Wdr6 | 2.41E-09 | 0.496527365 | 1 | 0.939 | 3.11E-05 |
| F2r | 2.58E-09 | 0.496835887 | 1 | 0.939 | 3.33E-05 |
| Mapk13 | 2.72E-09 | 0.563222801 | 1 | 0.835 | 3.50E-05 |
| Ddah2 | 2.73E-09 | 0.729114247 | 1 | 0.965 | 3.52E-05 |
| Cd320 | 2.75E-09 | 0.362796058 | 1 | 0.991 | 3.54E-05 |
| Ccnd2 | 3.08E-09 | 0.528130154 | 1 | 1 | 3.97E-05 |
| Rnf145 | 3.11E-09 | 0.339579921 | 1 | 0.965 | 4.01E-05 |
| Stmn1 | 3.14E-09 | 0.589848111 | 1 | 1 | 4.04E-05 |
| Upp1 | 3.25E-09 | 0.345894521 | 0.793 | 0.452 | 4.20E-05 |
| Rasl11b | 3.68E-09 | 0.386441379 | 0.897 | 0.643 | 4.75E-05 |
| Maged1 | 3.89E-09 | 0.480110736 | 1 | 1 | 5.01E-05 |
| Vps72 | 4.24E-09 | 0.357401834 | 1 | 1 | 5.46E-05 |
| Vash2 | 4.53E-09 | 0.321814039 | 1 | 0.774 | 5.83E-05 |
| Pak1 | 4.71E-09 | 0.407301323 | 0.966 | 0.678 | 6.08E-05 |
| Ptp4a1 | 4.72E-09 | 0.343297215 | 1 | 1 | 6.08E-05 |
| Limk2 | 4.76E-09 | 0.260540838 | 1 | 1 | 6.14E-05 |
| Stap2 | 5.37E-09 | 0.304158669 | 0.931 | 0.626 | 6.92E-05 |
| Vps36 | 5.62E-09 | 0.358995197 | 1 | 0.991 | 7.24E-05 |
| Arrdc1 | 5.85E-09 | 0.351562168 | 0.966 | 0.609 | 7.54E-05 |
| Tmem98 | 5.92E-09 | 0.475518919 | 1 | 0.835 | 7.64E-05 |
| Crip2 | 6.48E-09 | 0.608531661 | 0.966 | 0.652 | 8.35E-05 |
| Sh3bp4 | 6.70E-09 | 0.293286778 | 1 | 0.643 | 8.64E-05 |
| Sh3bp1 | 7.57E-09 | 0.270042629 | 0.966 | 0.965 | 9.76E-05 |
| Rcn1 | 7.57E-09 | 0.456903791 | 1 | 0.983 | 9.76E-05 |
| Ajuba | 7.58E-09 | 0.316532758 | 1 | 0.904 | 9.77E-05 |
| Laptm4b | 7.88E-09 | 0.474856939 | 1 | 1 | 0.00010161 |
| Fzd7 | 9.15E-09 | 0.268570057 | 1 | 0.843 | 0.000117971 |
| Slc16a13 | 1.13E-08 | 0.324124543 | 0.793 | 0.548 | 0.000145617 |
| Dpysl5 | 1.28E-08 | 0.30303445 | 0.966 | 0.713 | 0.000165143 |
| Pkm | 1.31E-08 | 0.520721365 | 1 | 1 | 0.000169206 |
| Syt5 | 1.40E-08 | 0.525767063 | 0.828 | 0.496 | 0.000179891 |
| Psca | 1.61E-08 | 0.424752551 | 0.517 | 0.165 | 0.000207407 |
| Ctnnb1 | 1.69E-08 | 0.434042415 | 1 | 1 | 0.000217442 |
| Hnrnph1 | 1.98E-08 | 0.425012671 | 1 | 1 | 0.000255129 |
| Krt18 | 2.16E-08 | 0.813684112 | 1 | 1 | 0.000278202 |
| Tmsb10 | 2.17E-08 | 0.818366202 | 1 | 1 | 0.000279325 |
| Etv5 | 2.30E-08 | 0.343580008 | 0.966 | 0.957 | 0.000296846 |
| Ctnna1 | 2.40E-08 | 0.520609019 | 1 | 0.991 | 0.000309379 |
| Mdk | 2.75E-08 | 0.73404765 | 1 | 1 | 0.000353932 |
| Slc38a1 | 2.99E-08 | 0.294771237 | 1 | 1 | 0.000385072 |
| Fam32a | 3.17E-08 | 0.501542972 | 1 | 0.957 | 0.000408942 |
| Alg10b | 3.35E-08 | 0.383097291 | 1 | 0.843 | 0.000432108 |
| Mfap2 | 3.55E-08 | 0.79550847 | 1 | 0.826 | 0.000457351 |
| Lxn | 3.67E-08 | 0.355040202 | 0.966 | 0.487 | 0.000473573 |
| Ocln | 3.75E-08 | 0.311153699 | 1 | 0.922 | 0.000483766 |
| Efna4 | 5.54E-08 | 0.408312912 | 0.966 | 0.661 | 0.000714284 |
| Rprm | 5.59E-08 | 0.269273121 | 0.621 | 0.157 | 0.000721069 |
| AA986860 | 5.94E-08 | 0.271823258 | 0.759 | 0.313 | 0.000765456 |
| Cd276 | 5.97E-08 | 0.347794284 | 1 | 0.878 | 0.000770218 |

| | | | | | |
|---|---|---|---|---|---|
| Zfp219 | 8.11E-08 | 0.285880022 | 0.897 | 0.809 | 0.001046048 |
| Mycn | 8.14E-08 | 0.401703035 | 1 | 0.939 | 0.001049576 |
| Myl6 | 1.14E-07 | 0.467662959 | 1 | 1 | 0.001466626 |
| Myl12b | 1.24E-07 | 0.44023647 | 1 | 1 | 0.001603131 |
| Wsb1 | 1.48E-07 | 0.430846713 | 1 | 0.983 | 0.001910264 |
| Gabarapl1 | 1.69E-07 | 0.509720644 | 0.966 | 0.948 | 0.00218065 |
| Actn4 | 1.76E-07 | 0.383006291 | 1 | 1 | 0.002262846 |
| Map7d1 | 2.08E-07 | 0.252757527 | 1 | 0.904 | 0.002675119 |
| Rgn | 2.12E-07 | 0.295819428 | 0.862 | 0.687 | 0.002726863 |
| Pls3 | 2.12E-07 | 0.315122622 | 1 | 0.826 | 0.002731532 |
| Tmem132a | 2.13E-07 | 0.29275982 | 1 | 0.939 | 0.00274846 |
| Zfp36l1 | 2.47E-07 | 0.32358025 | 1 | 1 | 0.003182079 |
| Nono | 2.48E-07 | 0.31158488 | 1 | 1 | 0.00320131 |
| Tuba1a | 2.64E-07 | 0.404955241 | 1 | 1 | 0.003400569 |
| Srebf2 | 2.96E-07 | 0.379793026 | 1 | 0.983 | 0.003814367 |
| Eif4ebp2 | 3.17E-07 | 0.380323886 | 0.931 | 0.948 | 0.004080479 |
| Hnrnpm | 3.37E-07 | 0.34413049 | 1 | 1 | 0.004342425 |
| Rem2 | 3.50E-07 | 0.343223292 | 0.931 | 0.548 | 0.004517318 |
| Slc16a1 | 3.66E-07 | 0.51706652 | 1 | 1 | 0.004721509 |
| Lppr3 | 4.15E-07 | 0.27111553 | 1 | 0.774 | 0.005353429 |
| Rasgrp3 | 4.30E-07 | 0.263257699 | 1 | 0.739 | 0.005543976 |
| Ndn | 4.64E-07 | 0.500203472 | 0.931 | 0.817 | 0.005988093 |
| Lgals2 | 4.81E-07 | 0.645156185 | 0.69 | 0.609 | 0.006199584 |
| Klk8 | 5.27E-07 | 0.330134622 | 0.655 | 0.304 | 0.006793141 |
| Pdpn | 5.60E-07 | 0.488337616 | 1 | 0.678 | 0.007222138 |
| Mapk6 | 6.41E-07 | 0.253370903 | 1 | 0.991 | 0.008258648 |
| Grn | 6.42E-07 | 0.348108139 | 1 | 0.948 | 0.008273172 |
| Ube2e3 | 6.47E-07 | 0.271970654 | 1 | 1 | 0.008337059 |
| Samd10 | 7.07E-07 | 0.346538817 | 0.931 | 0.617 | 0.009112975 |
| Tubb4a | 8.85E-07 | 0.562504383 | 1 | 0.835 | 0.011404925 |
| Rac3 | 8.89E-07 | 0.31031796 | 0.931 | 0.53 | 0.011455864 |
| Calu | 9.08E-07 | 0.341912843 | 1 | 1 | 0.011700059 |
| Cdkn1a | 9.56E-07 | 0.41937178 | 0.966 | 0.922 | 0.012320914 |
| Atp1a1 | 9.63E-07 | 0.290136421 | 1 | 1 | 0.012415428 |
| Cpe | 1.09E-06 | 0.288412774 | 1 | 0.6 | 0.014010491 |
| Cpm | 1.09E-06 | 0.467571467 | 1 | 1 | 0.014102403 |
| Snx17 | 1.25E-06 | 0.273286466 | 1 | 0.896 | 0.016064704 |
| Skap2 | 1.39E-06 | 0.342446387 | 0.828 | 0.557 | 0.017864785 |
| Aif1l | 1.50E-06 | 0.251772065 | 0.966 | 0.87 | 0.019344926 |
| 4930427A0 | 1.51E-06 | 0.253993857 | 0.828 | 0.774 | 0.019505062 |
| Prdx2 | 1.59E-06 | 0.37759588 | 1 | 1 | 0.020505739 |
| Tes | 1.61E-06 | 0.432374998 | 1 | 0.948 | 0.020812368 |
| Ogt | 1.64E-06 | 0.312685782 | 1 | 1 | 0.021124704 |
| Serpinh1 | 1.74E-06 | 0.442482765 | 1 | 1 | 0.022406119 |
| H2afv | 2.09E-06 | 0.546839543 | 1 | 1 | 0.026975817 |
| Cbx2 | 2.11E-06 | 0.278797441 | 0.966 | 0.948 | 0.027141686 |
| Dpp3 | 2.21E-06 | 0.360969446 | 1 | 1 | 0.028484879 |
| Tusc3 | 2.25E-06 | 0.392367602 | 1 | 0.974 | 0.028966019 |
| Ffar4 | 2.57E-06 | 0.420270784 | 0.828 | 0.713 | 0.0330935 |
| Cgn | 2.58E-06 | 0.250259221 | 1 | 1 | 0.033317648 |
| Flywch2 | 2.81E-06 | 0.333176655 | 0.966 | 0.687 | 0.036277857 |
| Igf2bp3 | 2.85E-06 | 0.298971845 | 1 | 1 | 0.036715865 |
| Gm1673 | 3.30E-06 | 0.367249023 | 0.966 | 0.791 | 0.042522297 |
| Dstn | 3.52E-06 | 0.407728104 | 1 | 1 | 0.045378779 |
| Nap1l1 | 3.69E-06 | 0.307101505 | 1 | 1 | 0.047605608 |
| Plat | 4.35E-06 | 0.267918898 | 0.793 | 0.374 | 0.056113523 |
| Me2 | 4.52E-06 | 0.370176247 | 0.966 | 0.974 | 0.058282368 |
| Slc7a6 | 5.44E-06 | 0.321512799 | 1 | 0.991 | 0.070082611 |

| | | | | | |
|---|---|---|---|---|---|
| Crabp2 | 5.69E-06 | 0.625335588 | 1 | 0.913 | 0.073405093 |
| Aplp2 | 6.53E-06 | 0.439787361 | 1 | 1 | 0.084147738 |
| Zfp422 | 7.39E-06 | 0.338325846 | 1 | 0.991 | 0.09525753 |
| Ctxn1 | 7.48E-06 | 0.529549894 | 1 | 0.896 | 0.096451671 |
| Usp22 | 7.63E-06 | 0.309052265 | 1 | 0.991 | 0.098407523 |
| Trappc6a | 8.41E-06 | 0.34473082 | 1 | 0.974 | 0.108402714 |
| Itm2c | 1.05E-05 | 0.410288145 | 1 | 1 | 0.134793453 |
| Tomm34 | 1.06E-05 | 0.31430939 | 1 | 0.983 | 0.136781405 |
| Ppp2r1a | 1.45E-05 | 0.354035402 | 1 | 1 | 0.187243511 |
| Usf1 | 1.46E-05 | 0.260373804 | 1 | 0.974 | 0.188463647 |
| Cdca7 | 1.47E-05 | 0.33780676 | 0.966 | 1 | 0.189705868 |
| Gsta4 | 1.51E-05 | 0.533520099 | 0.966 | 0.939 | 0.194797892 |
| Ash2l | 1.65E-05 | 0.257314707 | 1 | 0.991 | 0.212942679 |
| Ganab | 1.73E-05 | 0.298541084 | 1 | 1 | 0.222560991 |
| Rab25 | 1.94E-05 | 0.250701736 | 0.793 | 0.383 | 0.250616021 |
| Dctn2 | 2.34E-05 | 0.286119056 | 0.966 | 1 | 0.301255237 |
| Ccar2 | 2.40E-05 | 0.305408391 | 1 | 0.974 | 0.309866777 |
| Tra2a | 2.46E-05 | 0.390414578 | 1 | 0.991 | 0.317254192 |
| Lrrc42 | 2.48E-05 | 0.354977368 | 1 | 0.983 | 0.319957449 |
| Nradd | 2.61E-05 | 0.250691246 | 0.931 | 0.67 | 0.337124455 |
| Eef1a1 | 2.81E-05 | 0.323700897 | 1 | 1 | 0.362272005 |
| Xrcc1 | 2.90E-05 | 0.353350162 | 0.966 | 1 | 0.373492355 |
| Nedd4 | 2.98E-05 | 0.272274585 | 1 | 1 | 0.384265274 |
| Naa40 | 3.18E-05 | 0.278755917 | 0.966 | 0.957 | 0.410168733 |
| Zfp125 | 3.21E-05 | 0.373738079 | 1 | 0.948 | 0.414092027 |
| Tcf3 | 3.22E-05 | 0.308238122 | 1 | 1 | 0.41448232 |
| Ctbp1 | 3.35E-05 | 0.274650267 | 1 | 0.939 | 0.431957373 |
| Csrp1 | 3.46E-05 | 0.437057756 | 1 | 0.939 | 0.445948523 |
| Adcy8 | 3.49E-05 | 0.273815631 | 0.793 | 0.487 | 0.449526512 |
| Morn4 | 3.63E-05 | 0.269767978 | 0.897 | 0.67 | 0.468440303 |
| Serinc2 | 4.12E-05 | 0.313439494 | 0.966 | 0.609 | 0.531369288 |
| Ptk7 | 4.33E-05 | 0.256339917 | 1 | 0.896 | 0.558642943 |
| Rcn2 | 4.52E-05 | 0.255284895 | 1 | 1 | 0.582128671 |
| Rbpms | 4.70E-05 | 0.267902971 | 1 | 1 | 0.605414542 |
| Akr7a5 | 4.89E-05 | 0.306103546 | 1 | 0.991 | 0.630067689 |
| Laptm4a | 4.92E-05 | 0.387025348 | 1 | 1 | 0.63411923 |
| Myl9 | 5.69E-05 | 0.585323279 | 0.966 | 0.809 | 0.733474873 |
| Wbp1 | 6.14E-05 | 0.314433421 | 0.966 | 0.922 | 0.791612235 |
| Spns1 | 7.08E-05 | 0.297306561 | 1 | 0.974 | 0.913278334 |
| 5730507C( | 7.42E-05 | 0.346333864 | 1 | 0.991 | 0.956624668 |
| Cdk16 | 7.94E-05 | 0.259976746 | 1 | 0.991 | 1 |
| Tspan14 | 8.41E-05 | 0.333676073 | 1 | 0.948 | 1 |
| Col2a1 | 8.56E-05 | 0.365634649 | 0.966 | 0.678 | 1 |
| Klc4 | 8.73E-05 | 0.253601947 | 0.897 | 0.6 | 1 |
| Casp8 | 9.83E-05 | 0.260643115 | 0.897 | 0.704 | 1 |
| Chmp2b | 0.000103878 | 0.29955664 | 0.966 | 0.904 | 1 |
| Gpx8 | 0.000107673 | 0.38897555 | 0.897 | 0.557 | 1 |
| Gipc1 | 0.000112897 | 0.414514438 | 1 | 0.965 | 1 |
| Gins4 | 0.000115533 | 0.398012563 | 1 | 0.965 | 1 |
| Suv420h2 | 0.000133132 | 0.25031333 | 1 | 0.948 | 1 |
| Hnrnpc | 0.00014704 | 0.293584592 | 1 | 1 | 1 |
| Ivns1abp | 0.000158737 | 0.300466685 | 1 | 1 | 1 |
| Ift43 | 0.000172686 | 0.328009201 | 1 | 0.904 | 1 |
| R3hdm4 | 0.000199016 | 0.286028049 | 0.966 | 0.991 | 1 |
| Nudt14 | 0.000212629 | 0.378363978 | 0.931 | 0.904 | 1 |
| Glrx | 0.000213895 | 0.426190934 | 0.966 | 0.73 | 1 |
| Pigs | 0.000239925 | 0.311265319 | 1 | 0.991 | 1 |
| Golm1 | 0.000251357 | 0.356580314 | 1 | 0.991 | 1 |

| | | | | | |
|---|---|---|---|---|---|
| Mfge8 | 0.000272206 | 0.453482827 | 1 | 1 | 1 |
| Sap30 | 0.000345191 | 0.296114746 | 0.931 | 0.87 | 1 |
| Abracl | 0.000394284 | 0.331762876 | 1 | 0.913 | 1 |
| Adprh | 0.000619436 | 0.345281705 | 1 | 0.991 | 1 |
| Rcan1 | 0.000637387 | 0.294538092 | 0.897 | 0.704 | 1 |
| Mest | 0.0006444 | 0.445970047 | 1 | 0.843 | 1 |
| Psap | 0.000645016 | 0.408680389 | 1 | 0.991 | 1 |
| Cdk4 | 0.000714684 | 0.261346677 | 1 | 1 | 1 |
| Eno1b | 0.000773568 | 0.277191372 | 1 | 1 | 1 |
| Ncstn | 0.000791514 | 0.277107114 | 1 | 0.983 | 1 |
| Cst3 | 0.00081686 | 0.496128028 | 1 | 1 | 1 |
| Sf3b2 | 0.000945834 | 0.295608139 | 1 | 1 | 1 |
| Myl12a | 0.001087012 | 0.267826774 | 1 | 1 | 1 |
| Hmgn1 | 0.001093891 | 0.253021078 | 1 | 1 | 1 |
| Lypla2 | 0.001323047 | 0.255018116 | 1 | 0.974 | 1 |
| Ptma | 0.001372037 | 0.287658797 | 1 | 1 | 1 |
| Emb | 0.001411115 | 0.479876664 | 1 | 1 | 1 |
| Cdc42ep1 | 0.001521612 | 0.269346821 | 1 | 0.93 | 1 |
| Gabarap | 0.001860661 | 0.312456341 | 1 | 1 | 1 |
| Eno1 | 0.001869702 | 0.273650938 | 1 | 1 | 1 |
| Ttc5 | 0.001880706 | 0.251499084 | 1 | 0.913 | 1 |
| Ogfr | 0.002322993 | 0.281974395 | 1 | 1 | 1 |
| Zfand6 | 0.00285781 | 0.27329426 | 1 | 0.983 | 1 |
| Bsg | 0.002866974 | 0.251351623 | 1 | 1 | 1 |
| Xrcc5 | 0.002940594 | 0.292819378 | 1 | 0.991 | 1 |
| Smim22 | 0.003969828 | 0.28165097 | 0.724 | 0.67 | 1 |
| Ccdc23 | 0.004364059 | 0.253574029 | 1 | 0.948 | 1 |
| Ctsz | 0.004658858 | 0.305867899 | 1 | 0.991 | 1 |
| Rcn3 | 0.004848873 | 0.421387352 | 1 | 1 | 1 |
| Pgrmc1 | 0.005469058 | 0.303012132 | 1 | 1 | 1 |
| Csrp2 | 0.005907637 | 0.435607452 | 1 | 1 | 1 |
| Uba1 | 0.005968251 | 0.26711085 | 1 | 1 | 1 |
| Txnrd1 | 0.006817412 | 0.250337035 | 1 | 1 | 1 |
| Nap1l4 | 0.008029577 | 0.251353485 | 1 | 1 | 1 |
| Mcm2 | 0.008411533 | 0.296830997 | 1 | 1 | 1 |
| Ppdpf | 0.008608975 | 0.286179653 | 1 | 1 | 1 |
| Tmbim6 | 0.009831646 | 0.269758571 | 1 | 1 | 1 |
| Sox17 | 4.27E-45 | 0.958331536 | 1 | 0.5 | 5.51E-41 |
| Gjb2 | 4.96E-26 | 0.362543932 | 0.667 | 0.31 | 6.40E-22 |
| Lgr4 | 2.81E-25 | 0.425517866 | 1 | 0.984 | 3.62E-21 |
| Pgpep1 | 4.17E-21 | 0.467300863 | 1 | 0.944 | 5.38E-17 |
| Dctn4 | 7.93E-21 | 0.597720589 | 1 | 1 | 1.02E-16 |
| Flrt3 | 6.49E-20 | 1.121244018 | 0.944 | 0.794 | 8.37E-16 |
| 1700011H | 6.86E-20 | 1.534806838 | 1 | 0.563 | 8.84E-16 |
| Sox11 | 3.98E-19 | 1.046008713 | 1 | 1 | 5.13E-15 |
| Flrt2 | 5.75E-18 | 0.341537056 | 0.944 | 0.619 | 7.42E-14 |
| 2610008E1 | 5.22E-17 | 0.3514476 | 1 | 0.722 | 6.73E-13 |
| Prr15l | 1.10E-16 | 0.316550955 | 0.833 | 0.484 | 1.41E-12 |
| Itpripl1 | 3.78E-16 | 0.297012178 | 0.944 | 0.889 | 4.87E-12 |
| Ift22 | 7.45E-16 | 0.306992695 | 1 | 0.857 | 9.60E-12 |
| Grpel2 | 1.15E-15 | 0.278720911 | 0.833 | 0.889 | 1.49E-11 |
| Gsk3b | 3.20E-15 | 0.446240376 | 1 | 1 | 4.12E-11 |
| Cbfa2t2 | 3.86E-15 | 0.26663789 | 1 | 0.984 | 4.98E-11 |
| Onecut1 | 4.41E-15 | 0.584727277 | 1 | 0.937 | 5.69E-11 |
| Pdcd4 | 5.75E-15 | 0.522181316 | 1 | 1 | 7.41E-11 |
| Rab38 | 6.88E-15 | 0.707990823 | 0.833 | 0.476 | 8.87E-11 |
| Slc2a2 | 7.63E-15 | 0.282418921 | 1 | 0.833 | 9.83E-11 |
| Tcf7l2 | 1.51E-14 | 0.421004457 | 1 | 0.992 | 1.95E-10 |

| | | | | | |
|---|---|---|---|---|---|
| Igsf3 | 2.20E-14 | 0.385318911 | 1 | 0.762 | 2.84E-10 |
| Tjp2 | 2.68E-14 | 0.432658236 | 1 | 0.921 | 3.45E-10 |
| Dpysl2 | 4.63E-14 | 0.866308783 | 1 | 0.992 | 5.97E-10 |
| Gstm2 | 9.10E-14 | 0.34113373 | 0.944 | 0.952 | 1.17E-09 |
| Arl4c | 9.84E-14 | 0.269905105 | 0.944 | 0.595 | 1.27E-09 |
| Stk24 | 1.14E-13 | 0.320023041 | 0.778 | 0.897 | 1.47E-09 |
| Rab13 | 3.90E-13 | 0.31513291 | 0.778 | 0.881 | 5.02E-09 |
| Zfp691 | 5.43E-13 | 0.251590685 | 0.889 | 0.857 | 7.00E-09 |
| Sri | 5.76E-13 | 0.334078082 | 1 | 0.976 | 7.43E-09 |
| Rxrg | 6.39E-13 | 0.265861841 | 0.667 | 0.333 | 8.24E-09 |
| Klf13 | 6.59E-13 | 0.295392981 | 1 | 0.992 | 8.49E-09 |
| Sparc | 7.79E-13 | 0.933693962 | 1 | 0.984 | 1.00E-08 |
| Kif3a | 7.85E-13 | 0.368312794 | 1 | 0.96 | 1.01E-08 |
| Cenpe | 8.42E-13 | 0.321658628 | 0.944 | 0.96 | 1.09E-08 |
| Zfp37 | 1.13E-12 | 0.289426969 | 0.722 | 0.683 | 1.46E-08 |
| Sox9 | 1.21E-12 | 1.025963947 | 1 | 0.817 | 1.56E-08 |
| Tspyl1 | 1.23E-12 | 0.353611113 | 1 | 0.905 | 1.58E-08 |
| Ccdc92 | 1.46E-12 | 0.307224469 | 0.833 | 0.603 | 1.88E-08 |
| Ifitm3 | 1.72E-12 | 1.040948602 | 1 | 0.984 | 2.22E-08 |
| Id2 | 2.06E-12 | 1.258555251 | 1 | 1 | 2.66E-08 |
| Plk2 | 2.30E-12 | 0.830740436 | 1 | 0.944 | 2.97E-08 |
| Prr13 | 2.53E-12 | 0.872353101 | 1 | 0.96 | 3.26E-08 |
| Crabp1 | 2.59E-12 | 1.653541812 | 0.944 | 0.563 | 3.34E-08 |
| Camk1g | 3.79E-12 | 0.585617429 | 0.889 | 0.5 | 4.89E-08 |
| Sesn1 | 4.83E-12 | 0.279882121 | 0.667 | 0.603 | 6.23E-08 |
| Ppox | 8.70E-12 | 0.269731688 | 1 | 0.905 | 1.12E-07 |
| Gm10479 | 1.10E-11 | 0.333602533 | 0.944 | 0.913 | 1.42E-07 |
| Mgst1 | 1.19E-11 | 0.439147673 | 1 | 0.627 | 1.53E-07 |
| Usp11 | 1.21E-11 | 0.451358749 | 1 | 0.921 | 1.56E-07 |
| Fbxo44 | 1.42E-11 | 0.316559652 | 0.722 | 0.365 | 1.83E-07 |
| Fbxo18 | 1.76E-11 | 0.329656115 | 1 | 0.889 | 2.27E-07 |
| Sall4 | 3.52E-11 | 0.305172928 | 1 | 1 | 4.53E-07 |
| Ndufa3 | 4.57E-11 | 0.634095343 | 1 | 1 | 5.90E-07 |
| Zscan26 | 5.56E-11 | 0.360037551 | 1 | 1 | 7.17E-07 |
| Dcaf7 | 6.68E-11 | 0.308042491 | 1 | 0.968 | 8.61E-07 |
| Csde1 | 1.12E-10 | 0.349031168 | 1 | 1 | 1.44E-06 |
| Dynlt1f | 1.16E-10 | 0.525383135 | 1 | 1 | 1.50E-06 |
| Fabp3 | 1.17E-10 | 0.356062802 | 0.611 | 0.373 | 1.50E-06 |
| Gm10031 | 1.23E-10 | 0.284366785 | 1 | 0.984 | 1.59E-06 |
| Gpr108 | 1.58E-10 | 0.405800899 | 0.944 | 0.825 | 2.03E-06 |
| Stxbp2 | 2.50E-10 | 0.459285361 | 1 | 1 | 3.23E-06 |
| Med18 | 2.80E-10 | 0.282434649 | 0.944 | 0.881 | 3.61E-06 |
| Opa3 | 2.98E-10 | 0.275911907 | 1 | 0.968 | 3.84E-06 |
| 9030624G | 3.18E-10 | 0.288481336 | 1 | 0.944 | 4.10E-06 |
| Ppp1r1a | 3.65E-10 | 0.535683097 | 0.944 | 0.746 | 4.71E-06 |
| Krtcap3 | 3.89E-10 | 0.714124098 | 1 | 0.73 | 5.01E-06 |
| Dynlt1b | 4.53E-10 | 0.483835556 | 1 | 1 | 5.85E-06 |
| Tes | 4.73E-10 | 0.75295321 | 1 | 0.952 | 6.10E-06 |
| Dynlt1a | 5.38E-10 | 0.429437308 | 1 | 1 | 6.93E-06 |
| Tgif1 | 5.90E-10 | 0.35854795 | 1 | 0.96 | 7.61E-06 |
| Col2a1 | 6.43E-10 | 0.657293637 | 1 | 0.698 | 8.28E-06 |
| Cystm1 | 6.96E-10 | 0.282603458 | 1 | 0.77 | 8.98E-06 |
| Alas1 | 7.86E-10 | 0.260367692 | 0.889 | 0.817 | 1.01E-05 |
| Gstm7 | 8.04E-10 | 0.437813209 | 0.889 | 0.802 | 1.04E-05 |
| St3gal5 | 8.61E-10 | 0.331237119 | 0.889 | 0.762 | 1.11E-05 |
| Trim35 | 1.33E-09 | 0.512982591 | 1 | 0.992 | 1.71E-05 |
| Eci2 | 1.52E-09 | 0.331400105 | 0.944 | 0.921 | 1.96E-05 |
| Narf | 2.00E-09 | 0.416776207 | 0.944 | 0.913 | 2.58E-05 |

| | | | | | |
|---|---|---|---|---|---|
| Atat1 | 2.14E-09 | 0.293277131 | 1 | 0.865 | 2.77E-05 |
| B4galt1 | 2.37E-09 | 0.38847826 | 1 | 0.992 | 3.06E-05 |
| Serinc2 | 2.70E-09 | 0.653263142 | 1 | 0.635 | 3.48E-05 |
| Hbp1 | 3.28E-09 | 0.337631548 | 1 | 0.992 | 4.22E-05 |
| Clic6 | 3.59E-09 | 0.448322434 | 1 | 0.913 | 4.63E-05 |
| Fbxo36 | 5.19E-09 | 0.262035935 | 0.722 | 0.532 | 6.69E-05 |
| Anxa5 | 5.42E-09 | 0.873662813 | 1 | 0.944 | 6.99E-05 |
| Msantd4 | 5.97E-09 | 0.500556521 | 1 | 0.929 | 7.70E-05 |
| Zcchc18 | 6.03E-09 | 0.365514696 | 1 | 0.603 | 7.77E-05 |
| Bcl2l1 | 6.18E-09 | 0.36660885 | 1 | 0.968 | 7.97E-05 |
| Rhoc | 1.02E-08 | 0.435528297 | 0.889 | 0.833 | 0.000130931 |
| Arglu1 | 1.05E-08 | 0.938645048 | 1 | 1 | 0.000135221 |
| Ddah2 | 1.15E-08 | 0.697328799 | 1 | 0.968 | 0.000147755 |
| Bet1l | 1.21E-08 | 0.331910839 | 0.889 | 0.77 | 0.000155994 |
| Dynlt1c | 1.53E-08 | 0.256334162 | 1 | 1 | 0.000196618 |
| Rnase4 | 1.59E-08 | 0.591661616 | 0.944 | 0.786 | 0.000204828 |
| S100a16 | 1.76E-08 | 0.367035724 | 0.778 | 0.611 | 0.000227445 |
| Glrx | 2.02E-08 | 0.77072637 | 1 | 0.746 | 0.00026082 |
| Gstt3 | 2.16E-08 | 0.251797821 | 0.833 | 0.738 | 0.0002783 |
| Cacnb3 | 2.64E-08 | 0.283033367 | 0.944 | 0.619 | 0.00034001 |
| Cdca2 | 2.81E-08 | 0.316070085 | 1 | 0.984 | 0.000362361 |
| Ncapd2 | 2.85E-08 | 0.369480578 | 1 | 1 | 0.00036733 |
| Tmc4 | 3.09E-08 | 0.271344958 | 1 | 0.897 | 0.000398278 |
| Nmt2 | 3.56E-08 | 0.359601088 | 1 | 1 | 0.000459552 |
| Hsd17b10 | 4.05E-08 | 0.604895651 | 1 | 1 | 0.00052216 |
| Vamp8 | 4.17E-08 | 0.47795677 | 1 | 0.984 | 0.000538025 |
| Ccdc23 | 4.59E-08 | 0.565327041 | 1 | 0.952 | 0.000591116 |
| Aktip | 4.79E-08 | 0.352177412 | 1 | 0.944 | 0.000617042 |
| Ntan1 | 4.96E-08 | 0.289787436 | 1 | 0.952 | 0.000639741 |
| Nrp1 | 5.40E-08 | 0.339006861 | 1 | 0.897 | 0.000696618 |
| Hkdc1 | 6.08E-08 | 0.319065068 | 1 | 0.944 | 0.000784026 |
| 2700062C( | 7.88E-08 | 0.515560078 | 0.833 | 0.849 | 0.001015297 |
| Mboat2 | 8.86E-08 | 0.363090178 | 1 | 0.897 | 0.001141881 |
| Cmtm8 | 9.64E-08 | 0.603646129 | 1 | 0.976 | 0.001242577 |
| Meis3 | 1.13E-07 | 0.26720262 | 0.667 | 0.571 | 0.001451507 |
| Clic1 | 1.21E-07 | 0.892402945 | 1 | 0.881 | 0.00156631 |
| Ndufa12 | 1.23E-07 | 0.424451256 | 1 | 1 | 0.001580216 |
| Nusap1 | 1.27E-07 | 0.291427383 | 0.944 | 0.96 | 0.001631275 |
| Zfp61 | 1.28E-07 | 0.257314437 | 0.833 | 0.73 | 0.001648888 |
| Tmem206 | 1.41E-07 | 0.349858164 | 0.833 | 0.675 | 0.001817679 |
| Aip | 1.51E-07 | 0.277480693 | 1 | 0.897 | 0.00194469 |
| Rpusd1 | 1.64E-07 | 0.355162638 | 0.722 | 0.675 | 0.002109548 |
| Atp6v1g1 | 1.71E-07 | 0.565204228 | 1 | 0.992 | 0.002204594 |
| Grina | 1.71E-07 | 0.552425357 | 0.944 | 0.825 | 0.002205843 |
| Lgals1 | 1.74E-07 | 0.720627844 | 0.944 | 0.881 | 0.002248608 |
| Ptprk | 1.85E-07 | 0.326145627 | 1 | 0.96 | 0.002380387 |
| Zfp125 | 2.07E-07 | 0.629258042 | 0.944 | 0.96 | 0.002664231 |
| Cox7a1 | 2.14E-07 | 0.397026459 | 0.833 | 0.365 | 0.002762052 |
| Kdm6b | 2.26E-07 | 0.25984273 | 1 | 1 | 0.002909641 |
| Cd81 | 2.61E-07 | 0.569988955 | 1 | 1 | 0.003366338 |
| Styx | 2.81E-07 | 0.349794642 | 1 | 0.849 | 0.003626189 |
| Notch2 | 2.88E-07 | 0.292246295 | 1 | 0.992 | 0.003718678 |
| Hn1 | 3.46E-07 | 0.611625873 | 1 | 1 | 0.004455527 |
| Maged2 | 3.77E-07 | 0.40716589 | 1 | 0.952 | 0.004863048 |
| Emb | 3.84E-07 | 0.882629446 | 1 | 1 | 0.004949507 |
| Fgfr2 | 4.76E-07 | 0.257373594 | 1 | 0.873 | 0.006143025 |
| Psme2 | 5.58E-07 | 0.449550572 | 1 | 0.992 | 0.00719667 |
| Calm2 | 5.80E-07 | 0.704502894 | 1 | 1 | 0.007471525 |

| | | | | | |
|---|---|---|---|---|---|
| Chrnb1 | 6.39E-07 | 0.256800928 | 0.778 | 0.317 | 0.008241862 |
| Foxa2 | 6.96E-07 | 0.506950089 | 1 | 1 | 0.008977477 |
| Gltp | 7.22E-07 | 0.458654068 | 0.944 | 0.857 | 0.009304913 |
| Gng11 | 7.58E-07 | 0.458958035 | 0.944 | 0.587 | 0.009774509 |
| Pdrg1 | 7.93E-07 | 0.341243647 | 0.944 | 0.937 | 0.010228296 |
| Igfbp2 | 8.09E-07 | 0.992832208 | 1 | 0.929 | 0.010433317 |
| Tmem59 | 8.12E-07 | 0.529622184 | 1 | 1 | 0.010473679 |
| Tmem106( | 8.43E-07 | 0.417004494 | 0.889 | 0.73 | 0.010870717 |
| Rcan1 | 8.81E-07 | 0.462365592 | 0.944 | 0.714 | 0.011359041 |
| Cmtm3 | 9.53E-07 | 0.548226174 | 0.889 | 0.659 | 0.012290034 |
| Sdr39u1 | 1.07E-06 | 0.331010266 | 1 | 0.881 | 0.013830888 |
| Nras | 1.11E-06 | 0.356622709 | 1 | 1 | 0.014329799 |
| Tmem218 | 1.12E-06 | 0.610019291 | 0.944 | 0.722 | 0.014437085 |
| 1110007C( | 1.13E-06 | 0.505729293 | 0.889 | 0.762 | 0.014552334 |
| Cirbp | 1.14E-06 | 0.307127556 | 1 | 0.968 | 0.014634709 |
| Tfpt | 1.17E-06 | 0.373774155 | 0.833 | 0.937 | 0.015067581 |
| Pde6d | 1.21E-06 | 0.267440134 | 1 | 0.921 | 0.01562031 |
| BC005624 | 1.43E-06 | 0.295884939 | 0.944 | 0.976 | 0.018483676 |
| Ndfip1 | 1.48E-06 | 0.721937567 | 1 | 0.992 | 0.019061163 |
| Tead2 | 1.52E-06 | 0.32632872 | 1 | 0.992 | 0.019559591 |
| Pum1 | 1.67E-06 | 0.255314185 | 1 | 1 | 0.021543938 |
| Gc | 1.74E-06 | 0.28157467 | 0.722 | 0.627 | 0.022456269 |
| Prmt2 | 1.81E-06 | 0.321892409 | 1 | 0.794 | 0.023323136 |
| Tm4sf4 | 1.85E-06 | 0.418319469 | 0.944 | 0.754 | 0.023870262 |
| Ptp4a2 | 1.87E-06 | 0.437900614 | 1 | 1 | 0.024098695 |
| Kif23 | 2.05E-06 | 0.410419952 | 0.944 | 0.889 | 0.026455536 |
| Serping1 | 2.19E-06 | 0.417517793 | 1 | 0.849 | 0.028230172 |
| Nr2f2 | 2.29E-06 | 0.430303445 | 1 | 0.984 | 0.029549258 |
| Gbas | 2.48E-06 | 0.488155641 | 1 | 1 | 0.032022552 |
| Hnrnpa0 | 2.89E-06 | 0.289508719 | 1 | 1 | 0.03731785 |
| Zfp825 | 3.02E-06 | 0.345425748 | 0.944 | 0.865 | 0.038903429 |
| Pfn2 | 3.30E-06 | 0.310544 | 1 | 0.968 | 0.042590885 |
| Dolk | 3.47E-06 | 0.256646804 | 0.722 | 0.794 | 0.044755465 |
| Rgs19 | 3.51E-06 | 0.322647404 | 0.944 | 0.929 | 0.045275298 |
| Spp1 | 4.03E-06 | 0.297948637 | 0.667 | 0.429 | 0.051995217 |
| Cited2 | 4.35E-06 | 0.26270611 | 0.944 | 0.849 | 0.056088391 |
| Commd10 | 4.35E-06 | 0.437100022 | 1 | 0.96 | 0.056090748 |
| Polr3k | 4.43E-06 | 0.285834199 | 1 | 1 | 0.057103294 |
| Carhsp1 | 4.48E-06 | 0.456876183 | 1 | 0.984 | 0.05774308 |
| Nelfb | 4.59E-06 | 0.333342322 | 1 | 0.992 | 0.05919222 |
| Pdpn | 4.60E-06 | 0.655271551 | 1 | 0.706 | 0.059273258 |
| Psme1 | 4.66E-06 | 0.44374425 | 1 | 1 | 0.060030651 |
| Kdm3a | 4.69E-06 | 0.296555945 | 1 | 0.921 | 0.060470461 |
| Cox4i1 | 5.31E-06 | 0.429208956 | 1 | 1 | 0.068480016 |
| G3bp2 | 5.36E-06 | 0.266506519 | 1 | 1 | 0.069087625 |
| Cep57 | 5.49E-06 | 0.269968455 | 1 | 0.992 | 0.070786787 |
| 4833420G | 5.92E-06 | 0.295816435 | 1 | 0.944 | 0.076349957 |
| Car14 | 6.74E-06 | 0.436436827 | 1 | 0.992 | 0.086850374 |
| Eif2b2 | 6.77E-06 | 0.345338592 | 1 | 0.944 | 0.087263185 |
| Pgm1 | 6.82E-06 | 0.252876712 | 1 | 0.802 | 0.087964405 |
| Mapk3 | 6.89E-06 | 0.286551501 | 1 | 0.921 | 0.088844249 |
| Tax1bp3 | 6.94E-06 | 0.295078272 | 1 | 1 | 0.089419241 |
| Fblim1 | 7.39E-06 | 0.285927643 | 1 | 0.754 | 0.095277141 |
| Ift52 | 7.48E-06 | 0.262944715 | 0.833 | 0.905 | 0.096373873 |
| Npc2 | 9.83E-06 | 0.424822182 | 1 | 1 | 0.126694147 |
| Deaf1 | 1.01E-05 | 0.291134744 | 0.944 | 0.968 | 0.12985424 |
| Slc50a1 | 1.03E-05 | 0.352818086 | 0.833 | 0.738 | 0.133070419 |
| Fopnl | 1.12E-05 | 0.407903463 | 1 | 0.984 | 0.145029849 |

| | | | | | |
|---|---|---|---|---|---|
| Clybl | 1.16E-05 | 0.363040565 | 0.889 | 0.825 | 0.150127035 |
| Ttc5 | 1.23E-05 | 0.42638686 | 1 | 0.921 | 0.158564249 |
| Epcam | 1.35E-05 | 0.806551823 | 1 | 0.976 | 0.174426408 |
| Pigyl | 1.49E-05 | 0.383981127 | 0.944 | 0.913 | 0.192009622 |
| Cnot8 | 1.55E-05 | 0.327187102 | 1 | 0.976 | 0.199374331 |
| Map1lc3b | 1.69E-05 | 0.255260047 | 1 | 1 | 0.217596392 |
| Sf3b2 | 1.70E-05 | 0.417121995 | 1 | 1 | 0.218712862 |
| Myadm | 1.94E-05 | 0.279881206 | 0.889 | 0.754 | 0.249723066 |
| Spg21 | 2.03E-05 | 0.334253793 | 1 | 0.952 | 0.261422697 |
| Lrrc58 | 2.06E-05 | 0.303628782 | 1 | 1 | 0.266192325 |
| Zkscan6 | 2.39E-05 | 0.2563095 | 0.833 | 0.54 | 0.307662402 |
| Spry2 | 2.65E-05 | 0.336744245 | 1 | 0.944 | 0.341220029 |
| Ackr3 | 2.70E-05 | 0.375527296 | 0.889 | 0.587 | 0.348236497 |
| Hhex | 2.75E-05 | 0.743070896 | 1 | 1 | 0.353967973 |
| 1110059E2 | 2.95E-05 | 0.253728886 | 1 | 0.929 | 0.380645226 |
| Rab3ip | 3.07E-05 | 0.391841099 | 1 | 0.944 | 0.395443505 |
| Lxn | 3.14E-05 | 0.406205681 | 0.889 | 0.54 | 0.404173418 |
| Spon2 | 3.15E-05 | 0.321219852 | 0.833 | 0.5 | 0.406362926 |
| Zfr | 3.24E-05 | 0.253900853 | 1 | 1 | 0.417104728 |
| Gnb1 | 3.29E-05 | 0.471318717 | 1 | 1 | 0.424563111 |
| Morn2 | 3.30E-05 | 0.376793999 | 0.944 | 0.651 | 0.424884938 |
| Marcksl1 | 3.42E-05 | 0.455145586 | 1 | 1 | 0.440365382 |
| Tmem51 | 3.51E-05 | 0.351435485 | 0.889 | 0.96 | 0.452968574 |
| S100a11 | 3.92E-05 | 0.74706411 | 1 | 0.833 | 0.50567922 |
| Tmem242 | 3.96E-05 | 0.296392017 | 1 | 0.976 | 0.510504051 |
| Itm2c | 4.12E-05 | 0.501040629 | 1 | 1 | 0.531368577 |
| Coro1b | 4.13E-05 | 0.3616651 | 1 | 1 | 0.532503876 |
| Abracl | 4.37E-05 | 0.45248137 | 0.944 | 0.929 | 0.563739526 |
| Snx17 | 4.46E-05 | 0.444715602 | 1 | 0.905 | 0.575007404 |
| Paip1 | 4.63E-05 | 0.370144839 | 1 | 1 | 0.596953849 |
| Ddx42 | 4.75E-05 | 0.31038545 | 1 | 1 | 0.612746353 |
| Zfp105 | 5.03E-05 | 0.297520552 | 0.889 | 0.897 | 0.648482689 |
| 1500009L1 | 5.14E-05 | 0.336817486 | 0.944 | 0.651 | 0.662363368 |
| Dazap2 | 5.52E-05 | 0.517906636 | 1 | 1 | 0.711290937 |
| Smim22 | 5.88E-05 | 0.452581362 | 0.833 | 0.659 | 0.75749333 |
| Rp9 | 6.09E-05 | 0.309140111 | 1 | 0.944 | 0.784772178 |
| 2610524H | 6.33E-05 | 0.371234351 | 1 | 0.944 | 0.815803067 |
| Snupn | 6.40E-05 | 0.373871038 | 0.944 | 0.786 | 0.825105577 |
| Twf1 | 7.25E-05 | 0.383277254 | 1 | 0.984 | 0.934924024 |
| EGFP | 7.31E-05 | 0.56357223 | 1 | 1 | 0.942467023 |
| BC023829 | 7.47E-05 | 0.304511023 | 1 | 0.897 | 0.96262136 |
| Leo1 | 7.70E-05 | 0.363856101 | 1 | 1 | 0.992818009 |
| Fam173a | 8.09E-05 | 0.338037837 | 0.944 | 0.889 | 1 |
| Sh3bp4 | 9.00E-05 | 0.252653049 | 1 | 0.675 | 1 |
| Trappc2l | 9.32E-05 | 0.391914321 | 1 | 0.921 | 1 |
| 5730507C0 | 9.55E-05 | 0.424349908 | 1 | 0.992 | 1 |
| Mcoln3 | 0.000102293 | 0.273606188 | 0.722 | 0.452 | 1 |
| H3f3b | 0.000102306 | 0.548115471 | 1 | 1 | 1 |
| Pcbd2 | 0.000102559 | 0.358895093 | 0.833 | 0.619 | 1 |
| H2afv | 0.000103016 | 0.500596201 | 1 | 1 | 1 |
| Atp6v1d | 0.000104731 | 0.303852309 | 1 | 0.968 | 1 |
| Cmtm7 | 0.000111364 | 0.289331288 | 1 | 0.825 | 1 |
| Fam96a | 0.00011626 | 0.476944061 | 1 | 0.992 | 1 |
| Cdc123 | 0.000131866 | 0.302819452 | 1 | 1 | 1 |
| Idnk | 0.000133887 | 0.253735539 | 0.944 | 0.929 | 1 |
| Prdx2 | 0.00013999 | 0.3894462 | 1 | 1 | 1 |
| Chchd6 | 0.000142568 | 0.305694747 | 0.944 | 0.897 | 1 |
| Arpc5 | 0.000145256 | 0.316351695 | 1 | 0.992 | 1 |

| | | | | | |
|---|---|---|---|---|---|
| Adcy8 | 0.000148441 | 0.303926188 | 0.889 | 0.5 | 1 |
| Pafah1b3 | 0.000149541 | 0.344514059 | 1 | 1 | 1 |
| Gdi1 | 0.000153067 | 0.32795102 | 1 | 0.992 | 1 |
| Map1lc3a | 0.000154774 | 0.462638633 | 1 | 0.976 | 1 |
| Ankrd46 | 0.000164429 | 0.252808836 | 0.944 | 0.937 | 1 |
| Fam60a | 0.000168909 | 0.396295224 | 1 | 1 | 1 |
| Sfr1 | 0.000171743 | 0.53594885 | 1 | 1 | 1 |
| Fbxo25 | 0.000182717 | 0.257798962 | 0.889 | 0.675 | 1 |
| Stoml1 | 0.000184694 | 0.253377784 | 0.778 | 0.46 | 1 |
| Pak1 | 0.000190003 | 0.308598734 | 1 | 0.698 | 1 |
| Rab11a | 0.000194682 | 0.292606305 | 1 | 0.992 | 1 |
| Kif12 | 0.000194832 | 0.297630639 | 1 | 0.929 | 1 |
| Mdk | 0.000199902 | 0.555044065 | 1 | 1 | 1 |
| AW20949 | 0.000203088 | 0.323538965 | 1 | 0.825 | 1 |
| Lyrm2 | 0.000211917 | 0.309765897 | 1 | 0.857 | 1 |
| Gpx8 | 0.000221775 | 0.489869137 | 0.889 | 0.587 | 1 |
| Sdhc | 0.000225496 | 0.338634092 | 1 | 1 | 1 |
| Klhl9 | 0.000229177 | 0.259777997 | 1 | 1 | 1 |
| Cldn7 | 0.000256931 | 0.564177582 | 1 | 0.786 | 1 |
| Tmem254l | 0.000287034 | 0.297606885 | 1 | 0.992 | 1 |
| Ift43 | 0.000322686 | 0.377009327 | 1 | 0.913 | 1 |
| Haus1 | 0.000332462 | 0.318938848 | 0.944 | 0.992 | 1 |
| Cox4i2 | 0.000342431 | 0.278361412 | 0.722 | 0.373 | 1 |
| Hes1 | 0.000348028 | 0.404820513 | 1 | 0.976 | 1 |
| Anapc2 | 0.000357294 | 0.326160921 | 1 | 0.984 | 1 |
| Hnf1b | 0.000361858 | 0.288261542 | 1 | 0.881 | 1 |
| Gnb5 | 0.000373866 | 0.265917166 | 0.667 | 0.484 | 1 |
| Mapk13 | 0.000376198 | 0.497574782 | 1 | 0.849 | 1 |
| Pdk3 | 0.000378615 | 0.271588309 | 0.944 | 0.897 | 1 |
| Incenp | 0.000389026 | 0.372329679 | 0.944 | 0.976 | 1 |
| Mea1 | 0.000389511 | 0.47180413 | 1 | 0.992 | 1 |
| Casp8 | 0.000393388 | 0.257614883 | 0.944 | 0.714 | 1 |
| Spint1 | 0.000399531 | 0.425121169 | 1 | 0.802 | 1 |
| Tmem254a | 0.00040216 | 0.294154737 | 1 | 0.992 | 1 |
| Gas8 | 0.000427694 | 0.26111418 | 1 | 0.841 | 1 |
| Kif22 | 0.000428068 | 0.378545649 | 1 | 0.984 | 1 |
| Pdcd2l | 0.000440926 | 0.25875923 | 0.944 | 0.897 | 1 |
| Mageh1 | 0.000440969 | 0.271986407 | 0.611 | 0.587 | 1 |
| Uqcc3 | 0.00050902 | 0.386260005 | 1 | 0.96 | 1 |
| Stmn1 | 0.000541056 | 0.602901845 | 1 | 1 | 1 |
| Rassf7 | 0.000554523 | 0.368549351 | 1 | 0.976 | 1 |
| Rcn2 | 0.000569279 | 0.299888881 | 1 | 1 | 1 |
| Lppr3 | 0.000578214 | 0.293821889 | 1 | 0.794 | 1 |
| Tubb4a | 0.000641847 | 0.515925698 | 1 | 0.849 | 1 |
| Nrep | 0.000648746 | 0.592525712 | 1 | 0.992 | 1 |
| Mgst3 | 0.000648846 | 0.251962312 | 1 | 0.802 | 1 |
| Isca2 | 0.0006529 | 0.327842729 | 1 | 0.929 | 1 |
| Akap8l | 0.000673135 | 0.358782409 | 1 | 0.976 | 1 |
| Lipa | 0.00067541 | 0.293701719 | 1 | 0.968 | 1 |
| Snx6 | 0.000680621 | 0.369915319 | 1 | 1 | 1 |
| Kdf1 | 0.000684224 | 0.320966719 | 0.778 | 0.548 | 1 |
| Syt5 | 0.000699363 | 0.283505736 | 0.944 | 0.508 | 1 |
| Tubb5 | 0.000704016 | 0.430516772 | 1 | 1 | 1 |
| Pcyox1l | 0.000743937 | 0.274284189 | 1 | 0.976 | 1 |
| Lrrc42 | 0.000760143 | 0.366653808 | 1 | 0.984 | 1 |
| Cldn6 | 0.00076663 | 0.861989124 | 1 | 0.984 | 1 |
| Cdkn1a | 0.000770413 | 0.40408065 | 1 | 0.921 | 1 |
| Wbp5 | 0.000805096 | 0.543974427 | 1 | 1 | 1 |

| | | | | | |
|---|---|---|---|---|---|
| Ccnf | 0.000806315 | 0.252114117 | 1 | 0.921 | 1 |
| Cbx3 | 0.000830583 | 0.475872476 | 1 | 1 | 1 |
| Ctxn1 | 0.000831721 | 0.537402113 | 1 | 0.905 | 1 |
| 0610040J0 | 0.000860936 | 0.282998251 | 1 | 0.96 | 1 |
| Polb | 0.000867715 | 0.297226565 | 0.944 | 0.976 | 1 |
| Atp6v1f | 0.000876513 | 0.446012255 | 1 | 0.992 | 1 |
| Tmem254 | 0.000894329 | 0.286775996 | 1 | 0.992 | 1 |
| Armc10 | 0.000946528 | 0.296520258 | 1 | 0.96 | 1 |
| Zfand6 | 0.000975863 | 0.391317021 | 1 | 0.984 | 1 |
| Prelid2 | 0.000998911 | 0.470394842 | 0.944 | 0.754 | 1 |
| Sept2 | 0.001051333 | 0.274059495 | 1 | 1 | 1 |
| Mfap2 | 0.001077977 | 0.660506577 | 1 | 0.841 | 1 |
| Fxyd6 | 0.001088854 | 0.468645056 | 1 | 0.754 | 1 |
| Timm21 | 0.001099101 | 0.369890654 | 1 | 0.952 | 1 |
| Fam103a1 | 0.001135088 | 0.327218869 | 1 | 0.984 | 1 |
| 1810037I1 | 0.001137324 | 0.271724119 | 1 | 1 | 1 |
| Ubl7 | 0.001151276 | 0.343302054 | 0.944 | 0.984 | 1 |
| Tubb2a | 0.001172593 | 0.260230177 | 1 | 0.984 | 1 |
| Fibp | 0.001274746 | 0.263222307 | 1 | 0.802 | 1 |
| Cst3 | 0.001287981 | 0.433618881 | 1 | 1 | 1 |
| Erdr1 | 0.001378353 | 0.435931101 | 1 | 1 | 1 |
| Gm8797 | 0.001504235 | 0.381559688 | 1 | 1 | 1 |
| Coprs | 0.001508993 | 0.274313508 | 1 | 0.889 | 1 |
| Smim12 | 0.001538169 | 0.266072197 | 0.889 | 0.96 | 1 |
| Mrpl27 | 0.001539109 | 0.393577117 | 1 | 1 | 1 |
| Arf5 | 0.001546479 | 0.306180567 | 1 | 1 | 1 |
| Ppic | 0.001637575 | 0.464686854 | 0.944 | 0.952 | 1 |
| Prkar1a | 0.001672632 | 0.257711543 | 1 | 1 | 1 |
| Tma7 | 0.001745532 | 0.265797256 | 1 | 1 | 1 |
| Gm10282 | 0.001779316 | 0.417483888 | 1 | 1 | 1 |
| Lsm10 | 0.001795851 | 0.297682245 | 1 | 0.865 | 1 |
| Rrp36 | 0.001820785 | 0.298120166 | 1 | 0.952 | 1 |
| Leprot | 0.001842287 | 0.299567983 | 0.889 | 0.841 | 1 |
| S100a10 | 0.001853322 | 0.621258119 | 0.833 | 0.675 | 1 |
| Nono | 0.001953736 | 0.321754251 | 1 | 1 | 1 |
| Plat | 0.002003471 | 0.265013509 | 0.667 | 0.429 | 1 |
| Ogt | 0.002006799 | 0.290894504 | 1 | 1 | 1 |
| Gsta4 | 0.002049443 | 0.455088854 | 1 | 0.937 | 1 |
| Ttc1 | 0.002066341 | 0.265452382 | 1 | 1 | 1 |
| Col4a2 | 0.002087693 | 0.285838114 | 1 | 0.81 | 1 |
| Gm1673 | 0.002094696 | 0.375494784 | 0.944 | 0.81 | 1 |
| Tmem261 | 0.00214324 | 0.402292162 | 1 | 1 | 1 |
| Rtfdc1 | 0.002291679 | 0.289576816 | 1 | 0.968 | 1 |
| Adi1 | 0.002538499 | 0.323635294 | 1 | 0.976 | 1 |
| Tmem86a | 0.002573917 | 0.367183517 | 1 | 0.921 | 1 |
| Ubb | 0.002605452 | 0.387051707 | 1 | 1 | 1 |
| Dhrs4 | 0.002618348 | 0.395136977 | 1 | 1 | 1 |
| Cd24a | 0.002681266 | 0.543051829 | 1 | 1 | 1 |
| Dera | 0.002722189 | 0.277846646 | 1 | 0.952 | 1 |
| Ifitm2 | 0.002743704 | 0.641734717 | 1 | 1 | 1 |
| Slc25a17 | 0.00274542 | 0.352877191 | 1 | 0.96 | 1 |
| Fam32a | 0.002850807 | 0.433591222 | 1 | 0.96 | 1 |
| H2afy2 | 0.002865067 | 0.393767104 | 1 | 0.992 | 1 |
| Dctn3 | 0.002946295 | 0.290310523 | 1 | 0.976 | 1 |
| Gm21887 | 0.003076498 | 0.317650056 | 1 | 1 | 1 |
| Manbal | 0.003148913 | 0.318208368 | 1 | 0.937 | 1 |
| Dync2li1 | 0.003267677 | 0.277081804 | 0.889 | 0.841 | 1 |
| Tmsb10 | 0.003316891 | 0.763600942 | 1 | 1 | 1 |

| | | | | | |
|---|---|---|---|---|---|
| Rabac1 | 0.003438203 | 0.312116077 | 0.944 | 0.96 | 1 |
| Pts | 0.003445532 | 0.31818449 | 0.889 | 0.897 | 1 |
| Mzt1 | 0.003469098 | 0.270277539 | 1 | 0.968 | 1 |
| Higd2a | 0.003475337 | 0.447096112 | 1 | 1 | 1 |
| Tmbim6 | 0.003528051 | 0.337426614 | 1 | 1 | 1 |
| Romo1 | 0.003546269 | 0.359534082 | 1 | 1 | 1 |
| Rsrp1 | 0.00356091 | 0.360128716 | 1 | 1 | 1 |
| Lamtor5 | 0.003576216 | 0.348770065 | 1 | 0.976 | 1 |
| Psmb2 | 0.003792674 | 0.264403132 | 1 | 1 | 1 |
| Hmgb3 | 0.003818625 | 0.397203822 | 1 | 1 | 1 |
| 1700123O | 0.00388685 | 0.253209414 | 1 | 0.968 | 1 |
| Hsd11b2 | 0.003918395 | 0.332815333 | 0.889 | 0.571 | 1 |
| Nudt16l1 | 0.003988117 | 0.321414939 | 1 | 0.984 | 1 |
| Ndrg2 | 0.004040425 | 0.289698472 | 0.833 | 0.889 | 1 |
| Dctn5 | 0.004066257 | 0.251167033 | 1 | 0.984 | 1 |
| Nsmce1 | 0.004454843 | 0.25763842 | 1 | 1 | 1 |
| Gja1 | 0.004530014 | 0.287346597 | 1 | 0.714 | 1 |
| Pea15a | 0.004546961 | 0.254751353 | 1 | 0.921 | 1 |
| Apoa1bp | 0.004800714 | 0.404594607 | 1 | 0.992 | 1 |
| Cdh6 | 0.004847239 | 0.295928038 | 0.944 | 0.627 | 1 |
| Hsbp1 | 0.004937307 | 0.389437499 | 1 | 1 | 1 |
| Habp2 | 0.005282287 | 0.424652753 | 1 | 1 | 1 |
| Serpinh1 | 0.005602237 | 0.374318196 | 1 | 1 | 1 |
| Coa3 | 0.005624113 | 0.325876395 | 1 | 0.984 | 1 |
| Samd10 | 0.005634896 | 0.265024382 | 0.889 | 0.651 | 1 |
| Col4a1 | 0.005767732 | 0.361700554 | 1 | 0.929 | 1 |
| Fis1 | 0.006034029 | 0.317888494 | 1 | 1 | 1 |
| Flywch2 | 0.006103354 | 0.281800569 | 0.889 | 0.722 | 1 |
| Atxn10 | 0.006117297 | 0.380898016 | 1 | 1 | 1 |
| Hmgn2 | 0.00632874 | 0.372717004 | 1 | 1 | 1 |
| Polr2k | 0.00678093 | 0.337048265 | 1 | 0.984 | 1 |
| Cbr1 | 0.00696607 | 0.322307537 | 1 | 0.96 | 1 |
| Paip2 | 0.007219208 | 0.253754392 | 1 | 1 | 1 |
| Tfb1m | 0.007482378 | 0.27890446 | 0.833 | 0.706 | 1 |
| Cldn4 | 0.007941277 | 0.320456069 | 0.944 | 0.611 | 1 |
| Rpp30 | 0.007981904 | 0.307098275 | 1 | 0.952 | 1 |
| Krt19 | 0.008462582 | 0.513581666 | 0.944 | 0.706 | 1 |
| Akr7a5 | 0.008835372 | 0.262749319 | 1 | 0.992 | 1 |
| Praf2 | 0.009253417 | 0.319699722 | 1 | 0.944 | 1 |
| Efna4 | 0.009487679 | 0.299712967 | 0.833 | 0.706 | 1 |
| Cited1 | 0.009772533 | 0.757702936 | 1 | 0.992 | 1 |
| Sh3bgrl3 | 0.009944178 | 0.368063923 | 1 | 0.976 | 1 |
| Myc | 3.11E-07 | 0.52927475 | 0.957 | 0.992 | 0.004009661 |
| Tgm2 | 4.47E-07 | 0.460023862 | 1 | 1 | 0.005767832 |
| Gm17511 | 2.77E-06 | 0.299942234 | 1 | 1 | 0.03574862 |
| Cdkn1c | 3.09E-06 | 0.621223502 | 1 | 1 | 0.039876272 |
| Rps12-ps3 | 1.36E-05 | 0.295499051 | 1 | 0.992 | 0.175284612 |
| Lars2 | 3.37E-05 | 1.194201682 | 1 | 1 | 0.435074128 |
| Id3 | 6.39E-05 | 0.619828357 | 1 | 0.992 | 0.824177541 |
| Scd2 | 6.97E-05 | 0.447789211 | 1 | 1 | 0.898315059 |
| Id1 | 8.60E-05 | 0.609294928 | 1 | 0.959 | 1 |
| Hsdl1 | 8.73E-05 | 0.25852394 | 0.913 | 0.81 | 1 |
| Igf2 | 0.00034117 | 0.411825762 | 1 | 1 | 1 |
| Pcyt2 | 0.000504973 | 0.257258457 | 1 | 1 | 1 |
| Rps12 | 0.000564613 | 0.473537341 | 1 | 1 | 1 |
| Cald1 | 0.000612268 | 0.268596408 | 1 | 1 | 1 |
| Phlda2 | 0.000729509 | 0.447830914 | 0.739 | 0.537 | 1 |
| Gm10073 | 0.000742595 | 0.356455099 | 1 | 1 | 1 |

| | | | | | |
|---|---|---|---|---|---|
| Mvd | 0.001080893 | 0.306323863 | 1 | 0.942 | 1 |
| Dusp9 | 0.001492511 | 0.281923241 | 1 | 1 | 1 |
| Gm11808 | 0.00172562 | 0.266396872 | 1 | 1 | 1 |
| Gm21887 | 0.001918325 | 0.3367245 | 1 | 1 | 1 |
| Sqle | 0.001968773 | 0.293528153 | 1 | 1 | 1 |
| Pcsk9 | 0.002607624 | 0.422596712 | 1 | 1 | 1 |
| Dlk1 | 0.002722626 | 0.291716417 | 1 | 1 | 1 |
| Rps15 | 0.003818374 | 0.276109954 | 1 | 1 | 1 |
| Hhex | 0.005064808 | 0.420406878 | 1 | 1 | 1 |
| Crabp2 | 0.005866474 | 0.386436305 | 1 | 0.917 | 1 |
| Gap43 | 0.009741653 | 0.284036287 | 0.913 | 0.818 | 1 |
| Apoa2 | 1.20E-62 | 2.078294629 | 1 | 1 | 1.55E-58 |
| Apom | 5.86E-57 | 2.088766413 | 1 | 0.986 | 7.56E-53 |
| Apoe | 1.86E-54 | 1.329963736 | 1 | 1 | 2.40E-50 |
| Serpina6 | 9.69E-54 | 2.399311968 | 1 | 0.946 | 1.25E-49 |
| Rbp4 | 5.90E-52 | 1.290850359 | 1 | 1 | 7.61E-48 |
| Trf | 3.09E-51 | 0.878269255 | 1 | 1 | 3.98E-47 |
| Apoc2 | 3.79E-47 | 0.718911225 | 0.929 | 0.541 | 4.88E-43 |
| Serpinf2 | 6.76E-47 | 1.407818038 | 1 | 1 | 8.72E-43 |
| Apoa1 | 1.63E-46 | 1.62966391 | 1 | 0.986 | 2.10E-42 |
| Itih2 | 3.27E-46 | 1.126546476 | 1 | 1 | 4.22E-42 |
| Sepp1 | 1.64E-45 | 1.250656918 | 1 | 0.986 | 2.11E-41 |
| Afp | 1.87E-45 | 2.234208599 | 1 | 1 | 2.41E-41 |
| Kng1 | 3.20E-45 | 0.286025136 | 0.757 | 0.608 | 4.12E-41 |
| Gulo | 2.20E-44 | 1.368714805 | 1 | 0.986 | 2.84E-40 |
| Qprt | 3.02E-42 | 0.716894374 | 1 | 0.919 | 3.90E-38 |
| Serpind1 | 5.35E-42 | 1.017533433 | 1 | 0.878 | 6.90E-38 |
| Fga | 1.22E-40 | 1.076479504 | 1 | 0.851 | 1.58E-36 |
| Ambp | 1.70E-40 | 0.959744551 | 1 | 1 | 2.19E-36 |
| F12 | 3.85E-39 | 0.848882947 | 1 | 0.743 | 4.97E-35 |
| Fgb | 4.10E-39 | 1.229924361 | 1 | 1 | 5.29E-35 |
| Plg | 7.01E-39 | 1.089292331 | 0.986 | 0.878 | 9.03E-35 |
| Glud1 | 1.03E-37 | 0.842373559 | 1 | 1 | 1.33E-33 |
| Phgdh | 3.17E-37 | 1.029639963 | 1 | 1 | 4.09E-33 |
| Lect2 | 1.93E-36 | 1.065792546 | 1 | 0.797 | 2.48E-32 |
| Serpinc1 | 1.94E-36 | 0.43981766 | 1 | 0.986 | 2.50E-32 |
| Fgg | 1.87E-35 | 1.277228048 | 1 | 1 | 2.41E-31 |
| Srm | 8.12E-35 | 1.004542818 | 1 | 1 | 1.05E-30 |
| Lpl | 3.19E-34 | 0.655961867 | 1 | 1 | 4.11E-30 |
| Shbg | 4.08E-34 | 0.827889306 | 0.957 | 0.689 | 5.26E-30 |
| Psat1 | 9.24E-34 | 0.889954435 | 1 | 1 | 1.19E-29 |
| Soat2 | 1.60E-32 | 0.596815241 | 1 | 1 | 2.06E-28 |
| B2m | 5.17E-32 | 0.908714599 | 1 | 1 | 6.67E-28 |
| Serpina1b | 7.03E-32 | 0.928807333 | 1 | 1 | 9.07E-28 |
| Igfbp1 | 1.91E-31 | 0.89243119 | 0.957 | 0.743 | 2.47E-27 |
| Fabp5 | 3.04E-31 | 0.92725775 | 1 | 1 | 3.92E-27 |
| Fkbp11 | 3.68E-31 | 0.998852829 | 1 | 0.919 | 4.74E-27 |
| Serpinf1 | 2.47E-30 | 0.846359936 | 1 | 1 | 3.19E-26 |
| C1qbp | 6.09E-30 | 0.658449817 | 1 | 1 | 7.85E-26 |
| Lcat | 6.62E-30 | 0.86342869 | 0.986 | 0.716 | 8.53E-26 |
| Ttr | 6.92E-30 | 0.639257979 | 1 | 1 | 8.92E-26 |
| F10 | 2.07E-29 | 0.616281959 | 1 | 0.946 | 2.67E-25 |
| Herpud1 | 2.62E-29 | 1.223446359 | 1 | 1 | 3.38E-25 |
| Alb | 4.73E-28 | 1.538146576 | 1 | 0.973 | 6.10E-24 |
| Cth | 1.06E-27 | 0.493853119 | 1 | 0.892 | 1.37E-23 |
| Tmem97 | 1.32E-27 | 0.791437778 | 1 | 1 | 1.71E-23 |
| Gm5549 | 2.24E-27 | 0.710811497 | 0.914 | 0.649 | 2.89E-23 |
| Rpl36al | 6.42E-27 | 0.610449834 | 1 | 1 | 8.28E-23 |

| | | | | | |
|---|---|---|---|---|---|
| Ddt | 8.78E-27 | 0.743124762 | 1 | 1 | 1.13E-22 |
| Nipsnap1 | 1.02E-26 | 0.644559766 | 1 | 1 | 1.31E-22 |
| Serpina1a | 1.31E-26 | 0.621624568 | 1 | 1 | 1.69E-22 |
| Psph | 2.13E-26 | 0.564458401 | 1 | 0.986 | 2.75E-22 |
| Tst | 2.93E-26 | 0.913295469 | 1 | 0.919 | 3.78E-22 |
| Pla2g12b | 6.41E-26 | 0.737290091 | 1 | 0.689 | 8.27E-22 |
| Osgin1 | 7.21E-26 | 0.800830725 | 0.957 | 0.77 | 9.29E-22 |
| Hspe1 | 2.41E-25 | 0.647418428 | 1 | 1 | 3.10E-21 |
| Gm10053 | 2.81E-25 | 0.60596265 | 1 | 1 | 3.62E-21 |
| F2 | 3.13E-25 | 0.590144741 | 1 | 0.811 | 4.04E-21 |
| Itm2b | 3.46E-25 | 0.641859544 | 1 | 1 | 4.46E-21 |
| Hspd1 | 8.78E-25 | 0.593917055 | 1 | 1 | 1.13E-20 |
| Gm6484 | 9.24E-25 | 0.570028473 | 0.971 | 0.486 | 1.19E-20 |
| Cpb2 | 1.47E-24 | 0.66421393 | 0.986 | 0.797 | 1.90E-20 |
| Stard10 | 2.25E-24 | 0.571653235 | 1 | 1 | 2.90E-20 |
| Ddit3 | 2.75E-24 | 0.653610876 | 0.986 | 0.635 | 3.55E-20 |
| Hspa9 | 3.10E-24 | 0.473528386 | 1 | 1 | 4.00E-20 |
| Gcat | 5.14E-24 | 0.46949329 | 1 | 0.986 | 6.63E-20 |
| Gchfr | 5.97E-24 | 0.688897864 | 0.986 | 0.649 | 7.69E-20 |
| Cited1 | 8.18E-24 | 0.509714902 | 1 | 0.986 | 1.05E-19 |
| Cox8a | 9.36E-24 | 0.292930259 | 1 | 1 | 1.21E-19 |
| Pcbd1 | 1.57E-23 | 0.731004643 | 1 | 1 | 2.03E-19 |
| Serpina1d | 2.45E-23 | 0.657067621 | 1 | 1 | 3.16E-19 |
| Cyc1 | 3.61E-23 | 0.511146237 | 1 | 1 | 4.65E-19 |
| Gstm1 | 4.71E-23 | 0.880199301 | 1 | 1 | 6.07E-19 |
| Dgat2 | 5.07E-23 | 0.733806699 | 1 | 0.986 | 6.53E-19 |
| Igsf1 | 7.01E-23 | 0.312319722 | 1 | 0.878 | 9.04E-19 |
| Hpx | 7.54E-23 | 0.848824881 | 0.986 | 0.919 | 9.72E-19 |
| Phb2 | 8.73E-23 | 0.43756421 | 1 | 1 | 1.13E-18 |
| Rps27l | 1.33E-22 | 0.530063409 | 1 | 1 | 1.71E-18 |
| Spp2 | 1.91E-22 | 0.698143246 | 1 | 0.77 | 2.46E-18 |
| Ddah1 | 2.14E-22 | 0.528232987 | 1 | 1 | 2.76E-18 |
| Mogat2 | 2.69E-22 | 0.707836165 | 0.986 | 0.986 | 3.47E-18 |
| Bco1 | 2.76E-22 | 0.321015245 | 0.9 | 0.446 | 3.56E-18 |
| Cycs | 2.82E-22 | 0.476662304 | 1 | 1 | 3.64E-18 |
| Slc25a5 | 7.61E-22 | 0.648369835 | 1 | 1 | 9.81E-18 |
| Sdhd | 1.98E-21 | 0.678001707 | 1 | 1 | 2.55E-17 |
| A1cf | 4.41E-21 | 0.317718274 | 1 | 0.77 | 5.68E-17 |
| C3 | 6.04E-21 | 0.313845972 | 0.986 | 0.662 | 7.79E-17 |
| Eif5a | 6.43E-21 | 0.504458655 | 1 | 1 | 8.29E-17 |
| Prdx1 | 1.16E-20 | 0.704622898 | 1 | 1 | 1.50E-16 |
| Cyp2c68 | 1.96E-20 | 0.433254406 | 0.957 | 0.757 | 2.53E-16 |
| Acaa2 | 2.45E-20 | 0.715583292 | 1 | 1 | 3.16E-16 |
| Ncl | 2.61E-20 | 0.380926436 | 1 | 1 | 3.36E-16 |
| Gstp1 | 2.98E-20 | 0.702467165 | 1 | 1 | 3.84E-16 |
| Atp5c1 | 3.20E-20 | 0.351717009 | 1 | 1 | 4.13E-16 |
| Gstp2 | 3.62E-20 | 0.598954114 | 1 | 1 | 4.67E-16 |
| Asns | 4.14E-20 | 0.641192028 | 1 | 0.973 | 5.33E-16 |
| Got2 | 7.99E-20 | 0.450048331 | 1 | 1 | 1.03E-15 |
| Reep6 | 9.88E-20 | 0.628936618 | 1 | 1 | 1.27E-15 |
| Serpina1c | 1.55E-19 | 0.399755463 | 1 | 0.986 | 2.00E-15 |
| Cacybp | 1.79E-19 | 0.554903416 | 1 | 1 | 2.31E-15 |
| Gdf15 | 1.92E-19 | 0.282012059 | 0.857 | 0.541 | 2.48E-15 |
| Apoc1 | 3.06E-19 | 1.036729831 | 1 | 0.838 | 3.94E-15 |
| Eif2s2 | 3.18E-19 | 0.384863163 | 1 | 1 | 4.10E-15 |
| Uqcrq | 4.54E-19 | 0.519910473 | 1 | 1 | 5.86E-15 |
| Gstz1 | 9.57E-19 | 0.600186153 | 1 | 0.973 | 1.23E-14 |
| Junb | 1.10E-18 | 0.652781818 | 0.886 | 0.689 | 1.42E-14 |

| | | | | | |
|---|---|---|---|---|---|
| Mttp | 1.38E-18 | 0.416941502 | 1 | 0.986 | 1.78E-14 |
| Ndufs2 | 1.50E-18 | 0.443484064 | 1 | 1 | 1.94E-14 |
| Ndufv2 | 3.01E-18 | 0.25914686 | 1 | 1 | 3.88E-14 |
| Emc6 | 3.15E-18 | 0.473528942 | 1 | 0.986 | 4.06E-14 |
| Ppib | 3.40E-18 | 0.45720307 | 1 | 1 | 4.38E-14 |
| Ostc | 5.58E-18 | 0.44126718 | 1 | 1 | 7.19E-14 |
| Agmat | 6.72E-18 | 0.410010379 | 0.929 | 0.5 | 8.66E-14 |
| Ssr2 | 8.04E-18 | 0.551130568 | 1 | 1 | 1.04E-13 |
| Ssr4 | 8.66E-18 | 0.304845693 | 1 | 1 | 1.12E-13 |
| Vtn | 1.21E-17 | 0.382185565 | 1 | 1 | 1.56E-13 |
| Sec61b | 1.28E-17 | 0.562672227 | 1 | 1 | 1.65E-13 |
| Mrpl20 | 1.31E-17 | 0.353587353 | 1 | 1 | 1.69E-13 |
| Shmt1 | 1.32E-17 | 0.451626091 | 1 | 1 | 1.70E-13 |
| Serpina10 | 2.15E-17 | 0.262256437 | 0.957 | 0.784 | 2.77E-13 |
| Sgk1 | 2.19E-17 | 0.256563121 | 0.986 | 0.892 | 2.82E-13 |
| Fth1 | 2.62E-17 | 0.709206163 | 1 | 1 | 3.38E-13 |
| Gnpnat1 | 2.71E-17 | 0.401038616 | 1 | 0.973 | 3.49E-13 |
| Rsl1d1 | 5.05E-17 | 0.418575182 | 1 | 1 | 6.51E-13 |
| Nme1 | 5.18E-17 | 0.333242104 | 1 | 1 | 6.68E-13 |
| Ran | 6.56E-17 | 0.334542696 | 1 | 1 | 8.45E-13 |
| Serp1 | 6.58E-17 | 0.535203048 | 1 | 1 | 8.48E-13 |
| Bex2 | 8.03E-17 | 0.494557355 | 1 | 1 | 1.04E-12 |
| Cnbp | 8.43E-17 | 0.410189261 | 1 | 1 | 1.09E-12 |
| Tmed2 | 1.09E-16 | 0.391119363 | 1 | 1 | 1.40E-12 |
| Tomm20 | 1.12E-16 | 0.31856231 | 1 | 1 | 1.45E-12 |
| Tomm5 | 1.19E-16 | 0.549924959 | 1 | 0.986 | 1.53E-12 |
| Lsm5 | 1.69E-16 | 0.38601201 | 1 | 1 | 2.17E-12 |
| Tmem258 | 1.77E-16 | 0.314763702 | 1 | 1 | 2.28E-12 |
| Ndufa6 | 2.05E-16 | 0.528482318 | 1 | 1 | 2.64E-12 |
| Aes | 3.14E-16 | 0.327581215 | 1 | 1 | 4.04E-12 |
| Uqcr10 | 3.21E-16 | 0.413465366 | 1 | 1 | 4.13E-12 |
| Nostrin | 3.58E-16 | 0.296548154 | 0.829 | 0.541 | 4.61E-12 |
| Nhp2 | 5.54E-16 | 0.496854663 | 1 | 1 | 7.14E-12 |
| Uqcrfs1 | 6.13E-16 | 0.368996424 | 1 | 0.986 | 7.90E-12 |
| Dnajc3 | 6.41E-16 | 0.37290726 | 1 | 0.973 | 8.26E-12 |
| Mtfp1 | 8.38E-16 | 0.432597176 | 0.986 | 0.811 | 1.08E-11 |
| Hint1 | 1.21E-15 | 0.395445071 | 1 | 1 | 1.57E-11 |
| Manf | 1.22E-15 | 0.678444126 | 1 | 1 | 1.57E-11 |
| Idh3a | 1.31E-15 | 0.566438186 | 1 | 0.973 | 1.69E-11 |
| Gpc3 | 1.79E-15 | 0.303850139 | 1 | 1 | 2.31E-11 |
| Npm1 | 2.99E-15 | 0.339149602 | 1 | 1 | 3.85E-11 |
| Sec61g | 3.19E-15 | 0.307320144 | 1 | 1 | 4.12E-11 |
| Slc19a2 | 3.89E-15 | 0.30081242 | 0.971 | 0.892 | 5.02E-11 |
| Galk1 | 4.01E-15 | 0.498534642 | 1 | 1 | 5.17E-11 |
| Fam195a | 4.89E-15 | 0.409939985 | 0.957 | 0.622 | 6.31E-11 |
| Fst | 5.06E-15 | 0.366678917 | 0.857 | 0.622 | 6.52E-11 |
| Mrpl17 | 5.13E-15 | 0.287805513 | 1 | 1 | 6.61E-11 |
| Slc25a15 | 5.71E-15 | 0.29991072 | 0.971 | 0.892 | 7.37E-11 |
| Naa50 | 5.97E-15 | 0.352391827 | 1 | 1 | 7.70E-11 |
| Timm13 | 6.79E-15 | 0.470857045 | 1 | 1 | 8.75E-11 |
| Nhp2l1 | 7.37E-15 | 0.361197579 | 1 | 1 | 9.50E-11 |
| Gm9797 | 8.48E-15 | 0.364099667 | 1 | 1 | 1.09E-10 |
| Mydgf | 9.06E-15 | 0.367604081 | 1 | 1 | 1.17E-10 |
| Ass1 | 1.38E-14 | 0.427823145 | 0.957 | 0.824 | 1.78E-10 |
| Nadk | 1.60E-14 | 0.368476438 | 1 | 0.973 | 2.06E-10 |
| Mrps28 | 1.88E-14 | 0.417530207 | 1 | 0.986 | 2.42E-10 |
| Gstt1 | 1.97E-14 | 0.520448707 | 1 | 0.905 | 2.54E-10 |
| Adk | 2.26E-14 | 0.4194204 | 1 | 1 | 2.91E-10 |

| | | | | | |
|---|---|---|---|---|---|
| Cdk2ap2 | 2.28E-14 | 0.50670783 | 1 | 0.973 | 2.94E-10 |
| Nop10 | 2.31E-14 | 0.562829021 | 1 | 1 | 2.98E-10 |
| Mrpl50 | 2.51E-14 | 0.396774249 | 1 | 1 | 3.23E-10 |
| Vapa | 2.58E-14 | 0.306952411 | 1 | 1 | 3.33E-10 |
| Gjb1 | 2.83E-14 | 0.497245324 | 0.943 | 0.797 | 3.64E-10 |
| Itih3 | 3.58E-14 | 0.311976429 | 0.9 | 0.568 | 4.62E-10 |
| Timm8a1 | 3.88E-14 | 0.372940731 | 1 | 0.973 | 5.01E-10 |
| Mrpl22 | 4.02E-14 | 0.513345752 | 1 | 0.986 | 5.18E-10 |
| Cct8 | 4.06E-14 | 0.333109294 | 1 | 1 | 5.23E-10 |
| Rps11 | 4.53E-14 | 0.300666921 | 1 | 1 | 5.84E-10 |
| Fxyd1 | 5.46E-14 | 0.488560859 | 0.857 | 0.473 | 7.04E-10 |
| Akr1c13 | 5.71E-14 | 0.525826492 | 0.986 | 0.892 | 7.36E-10 |
| Psmd7 | 7.05E-14 | 0.351321847 | 1 | 1 | 9.08E-10 |
| Gstm3 | 8.51E-14 | 0.292242365 | 1 | 0.878 | 1.10E-09 |
| Mrpl16 | 9.64E-14 | 0.461804358 | 1 | 0.892 | 1.24E-09 |
| Aadac | 9.65E-14 | 0.380669205 | 0.743 | 0.432 | 1.24E-09 |
| Uqcrb | 1.13E-13 | 0.40430953 | 1 | 1 | 1.46E-09 |
| Ptges3 | 1.19E-13 | 0.321198875 | 1 | 1 | 1.54E-09 |
| Gm5611 | 1.24E-13 | 0.330087988 | 1 | 1 | 1.60E-09 |
| Grhpr | 1.29E-13 | 0.406707803 | 1 | 0.986 | 1.66E-09 |
| Etfa | 1.42E-13 | 0.396673145 | 1 | 1 | 1.83E-09 |
| Rpn1 | 1.46E-13 | 0.379747902 | 1 | 1 | 1.88E-09 |
| Uchl5 | 1.46E-13 | 0.315343527 | 1 | 1 | 1.89E-09 |
| Timm23 | 1.99E-13 | 0.358933131 | 1 | 1 | 2.57E-09 |
| H2-Q7 | 2.07E-13 | 0.254040013 | 1 | 1 | 2.67E-09 |
| Emg1 | 2.26E-13 | 0.4328256 | 1 | 1 | 2.92E-09 |
| Cfi | 2.67E-13 | 0.555255893 | 0.957 | 0.77 | 3.44E-09 |
| Sephs2 | 3.48E-13 | 0.483127973 | 1 | 1 | 4.49E-09 |
| Gm10273 | 3.62E-13 | 0.308409049 | 1 | 0.986 | 4.66E-09 |
| Pla2g12a | 3.75E-13 | 0.309450392 | 1 | 1 | 4.84E-09 |
| Cox7b | 4.91E-13 | 0.374173044 | 1 | 1 | 6.33E-09 |
| Ndufa2 | 4.93E-13 | 0.313865261 | 1 | 1 | 6.35E-09 |
| Edf1 | 5.56E-13 | 0.408508464 | 1 | 1 | 7.16E-09 |
| Mmd2 | 5.84E-13 | 0.289345892 | 1 | 0.824 | 7.53E-09 |
| Pdia6 | 7.94E-13 | 0.391114248 | 1 | 1 | 1.02E-08 |
| Etfb | 8.57E-13 | 0.394713289 | 1 | 1 | 1.10E-08 |
| Ndufb4 | 9.13E-13 | 0.392789912 | 1 | 1 | 1.18E-08 |
| Cct6a | 9.90E-13 | 0.317254254 | 1 | 1 | 1.28E-08 |
| Psmb10 | 1.06E-12 | 0.451643876 | 1 | 0.932 | 1.37E-08 |
| Mthfd2 | 1.22E-12 | 0.428666813 | 1 | 0.973 | 1.58E-08 |
| Cd302 | 1.39E-12 | 0.321311673 | 0.929 | 0.689 | 1.79E-08 |
| Brix1 | 1.39E-12 | 0.433848055 | 1 | 0.986 | 1.79E-08 |
| Gm5616 | 1.47E-12 | 0.272880132 | 1 | 0.973 | 1.89E-08 |
| Vimp | 1.90E-12 | 0.647305438 | 1 | 0.986 | 2.45E-08 |
| Rpf2 | 2.22E-12 | 0.396714661 | 1 | 0.986 | 2.86E-08 |
| Ccne1 | 2.35E-12 | 0.493743424 | 1 | 1 | 3.03E-08 |
| Psma5 | 2.43E-12 | 0.276065633 | 1 | 1 | 3.13E-08 |
| Qdpr | 2.62E-12 | 0.266656242 | 0.986 | 0.959 | 3.38E-08 |
| Chchd4 | 2.74E-12 | 0.453295376 | 0.986 | 0.973 | 3.53E-08 |
| Snx2 | 2.91E-12 | 0.338436648 | 1 | 1 | 3.76E-08 |
| Mpzl1 | 3.49E-12 | 0.255536537 | 1 | 1 | 4.50E-08 |
| Gamt | 3.69E-12 | 0.333092887 | 1 | 0.919 | 4.76E-08 |
| Sdhb | 4.31E-12 | 0.348454921 | 1 | 1 | 5.55E-08 |
| Tex30 | 5.59E-12 | 0.395529675 | 1 | 0.959 | 7.20E-08 |
| Ldhb | 6.10E-12 | 0.500217041 | 1 | 1 | 7.87E-08 |
| Tmem37 | 6.40E-12 | 0.440683626 | 0.986 | 0.865 | 8.25E-08 |
| Mdh2 | 7.01E-12 | 0.280851108 | 1 | 1 | 9.04E-08 |
| Igfbp4 | 8.88E-12 | 0.544847183 | 1 | 1 | 1.14E-07 |

| | | | | | |
|---|---|---|---|---|---|
| Stt3a | 1.21E-11 | 0.260389059 | 1 | 1 | 1.56E-07 |
| Nars | 1.35E-11 | 0.453400103 | 1 | 1 | 1.75E-07 |
| Scp2 | 1.36E-11 | 0.309905251 | 1 | 1 | 1.75E-07 |
| Pbk | 1.60E-11 | 0.409180388 | 1 | 0.919 | 2.06E-07 |
| Ndufb6 | 1.79E-11 | 0.292717919 | 1 | 1 | 2.31E-07 |
| Gcsh | 1.82E-11 | 0.28969229 | 1 | 1 | 2.34E-07 |
| Eci1 | 1.96E-11 | 0.379867748 | 1 | 0.959 | 2.52E-07 |
| Sdf2l1 | 1.97E-11 | 0.675612356 | 1 | 0.986 | 2.54E-07 |
| Cks2 | 2.00E-11 | 0.425913118 | 1 | 1 | 2.58E-07 |
| Sar1b | 2.20E-11 | 0.471898164 | 1 | 1 | 2.83E-07 |
| Anp32b | 2.35E-11 | 0.275509149 | 1 | 1 | 3.03E-07 |
| Btg2 | 2.89E-11 | 0.646018653 | 0.971 | 0.932 | 3.73E-07 |
| Psmg1 | 2.89E-11 | 0.340695413 | 1 | 1 | 3.73E-07 |
| Pold2 | 3.06E-11 | 0.339764644 | 1 | 1 | 3.94E-07 |
| Ndufs8 | 3.07E-11 | 0.280372217 | 1 | 0.986 | 3.95E-07 |
| Fkbp1a | 3.09E-11 | 0.303964548 | 1 | 1 | 3.99E-07 |
| Eif4ebp1 | 3.52E-11 | 0.559451382 | 1 | 1 | 4.54E-07 |
| Surf4 | 3.73E-11 | 0.372390999 | 1 | 1 | 4.81E-07 |
| Mrpl12 | 3.82E-11 | 0.293774708 | 1 | 1 | 4.93E-07 |
| Mrps7 | 4.05E-11 | 0.29846032 | 1 | 1 | 5.23E-07 |
| Hspa5 | 4.20E-11 | 0.542584376 | 1 | 1 | 5.42E-07 |
| Akr1c12 | 4.47E-11 | 0.450914319 | 0.986 | 0.851 | 5.76E-07 |
| Timm50 | 5.16E-11 | 0.30396702 | 1 | 1 | 6.66E-07 |
| Rcl1 | 5.49E-11 | 0.384965196 | 1 | 1 | 7.08E-07 |
| Apof | 5.94E-11 | 0.299180473 | 1 | 0.811 | 7.66E-07 |
| H2-Q6 | 6.04E-11 | 0.330399423 | 1 | 1 | 7.79E-07 |
| Xbp1 | 9.50E-11 | 0.439966385 | 1 | 1 | 1.22E-06 |
| Degs1 | 9.84E-11 | 0.356921881 | 1 | 1 | 1.27E-06 |
| Tspan12 | 9.91E-11 | 0.53231556 | 1 | 1 | 1.28E-06 |
| Mesdc2 | 1.06E-10 | 0.256869922 | 1 | 1 | 1.37E-06 |
| Creld2 | 1.10E-10 | 0.731188443 | 1 | 1 | 1.42E-06 |
| Eif2s1 | 1.24E-10 | 0.281288591 | 1 | 1 | 1.60E-06 |
| Aass | 1.49E-10 | 0.34391502 | 0.943 | 0.851 | 1.92E-06 |
| U2af1 | 1.62E-10 | 0.340922439 | 1 | 1 | 2.09E-06 |
| Akr1c19 | 1.75E-10 | 0.39075547 | 1 | 1 | 2.25E-06 |
| Dnajc12 | 1.80E-10 | 0.278290259 | 0.986 | 0.784 | 2.33E-06 |
| Hprt | 1.83E-10 | 0.449009964 | 1 | 0.986 | 2.36E-06 |
| Hmbs | 1.88E-10 | 0.346447585 | 1 | 1 | 2.42E-06 |
| Calr | 1.94E-10 | 0.38420695 | 1 | 1 | 2.50E-06 |
| Rpn2 | 1.97E-10 | 0.270220996 | 1 | 1 | 2.53E-06 |
| Egr1 | 2.00E-10 | 0.620655387 | 0.986 | 0.973 | 2.58E-06 |
| Bccip | 2.23E-10 | 0.340843338 | 1 | 0.986 | 2.87E-06 |
| Prdx3 | 2.41E-10 | 0.427126546 | 1 | 0.973 | 3.11E-06 |
| Sigmar1 | 2.41E-10 | 0.253214463 | 1 | 0.986 | 3.11E-06 |
| Dctpp1 | 2.46E-10 | 0.289287704 | 1 | 1 | 3.17E-06 |
| Ppp1r15a | 2.49E-10 | 0.428413191 | 0.986 | 0.919 | 3.22E-06 |
| Tceb1 | 3.42E-10 | 0.266170839 | 1 | 1 | 4.41E-06 |
| Rnf128 | 3.73E-10 | 0.272827727 | 0.957 | 0.932 | 4.81E-06 |
| Pno1 | 4.48E-10 | 0.263213025 | 1 | 1 | 5.77E-06 |
| Fos | 4.67E-10 | 1.057371339 | 0.986 | 0.946 | 6.02E-06 |
| Dnajc22 | 5.92E-10 | 0.381477675 | 1 | 0.865 | 7.63E-06 |
| Eif1a | 6.04E-10 | 0.346256719 | 1 | 0.986 | 7.78E-06 |
| Spcs1 | 6.96E-10 | 0.332495479 | 1 | 1 | 8.98E-06 |
| Mrpl10 | 7.18E-10 | 0.307213795 | 1 | 0.986 | 9.26E-06 |
| Gm10094 | 8.29E-10 | 0.268572239 | 1 | 1 | 1.07E-05 |
| Hsp90b1 | 9.27E-10 | 0.472665175 | 1 | 1 | 1.20E-05 |
| Nifk | 1.02E-09 | 0.322959761 | 1 | 1 | 1.31E-05 |
| Exosc7 | 1.28E-09 | 0.372897375 | 1 | 1 | 1.65E-05 |

| | | | | | |
|---|---|---|---|---|---|
| Spc24 | 1.42E-09 | 0.294773388 | 1 | 1 | 1.83E-05 |
| Lpcat3 | 1.47E-09 | 0.308923277 | 1 | 0.946 | 1.90E-05 |
| Pa2g4 | 1.48E-09 | 0.342869862 | 1 | 1 | 1.91E-05 |
| Slc25a39 | 1.54E-09 | 0.280462173 | 1 | 1 | 1.98E-05 |
| Atf3 | 1.55E-09 | 0.545662979 | 0.957 | 0.689 | 1.99E-05 |
| Tuba4a | 1.77E-09 | 0.294195366 | 1 | 0.986 | 2.28E-05 |
| Rexo2 | 2.06E-09 | 0.373946468 | 1 | 0.946 | 2.65E-05 |
| Mrpl42 | 2.10E-09 | 0.427888583 | 1 | 1 | 2.70E-05 |
| Ephx2 | 2.20E-09 | 0.281521362 | 0.9 | 0.703 | 2.84E-05 |
| Atp6v0c | 2.73E-09 | 0.272481256 | 1 | 1 | 3.52E-05 |
| Acadm | 2.89E-09 | 0.255618569 | 1 | 0.986 | 3.73E-05 |
| Farsb | 3.62E-09 | 0.337554964 | 1 | 1 | 4.66E-05 |
| Cks1b | 3.84E-09 | 0.344177146 | 1 | 1 | 4.95E-05 |
| Pdia3 | 4.10E-09 | 0.259555475 | 1 | 1 | 5.28E-05 |
| Lyar | 4.74E-09 | 0.34793276 | 1 | 0.986 | 6.11E-05 |
| Cops6 | 5.08E-09 | 0.328185997 | 1 | 1 | 6.55E-05 |
| Atp5a1 | 5.47E-09 | 0.384711881 | 1 | 1 | 7.05E-05 |
| Timm10 | 5.66E-09 | 0.33520752 | 0.986 | 0.838 | 7.29E-05 |
| Gsn | 6.45E-09 | 0.336748247 | 0.943 | 0.811 | 8.31E-05 |
| Erp44 | 8.20E-09 | 0.261439298 | 1 | 0.986 | 0.000105663 |
| Atp5g1 | 8.88E-09 | 0.26606985 | 1 | 1 | 0.00011445 |
| Slc35b1 | 9.62E-09 | 0.335114248 | 1 | 1 | 0.000123983 |
| Nudcd2 | 9.95E-09 | 0.280050008 | 1 | 1 | 0.000128311 |
| Sc5d | 1.05E-08 | 0.353925991 | 1 | 0.973 | 0.00013532 |
| Ppa1 | 1.25E-08 | 0.308574475 | 1 | 1 | 0.000160644 |
| Cinp | 1.26E-08 | 0.297209793 | 1 | 0.986 | 0.000162359 |
| Jagn1 | 1.59E-08 | 0.310358288 | 1 | 0.986 | 0.00020507 |
| Nop16 | 1.63E-08 | 0.383490645 | 1 | 1 | 0.000210358 |
| Nfkbia | 1.83E-08 | 0.493110605 | 0.943 | 0.892 | 0.000235634 |
| Tmem176a | 1.99E-08 | 0.382451466 | 0.943 | 0.811 | 0.000257103 |
| Lap3 | 2.03E-08 | 0.282925019 | 1 | 1 | 0.000261278 |
| Pcsk9 | 2.11E-08 | 0.286622235 | 1 | 1 | 0.000272097 |
| Cops7a | 2.27E-08 | 0.254419282 | 1 | 1 | 0.000292282 |
| Mak16 | 2.41E-08 | 0.267214605 | 1 | 1 | 0.000310769 |
| Dap | 2.85E-08 | 0.390363955 | 1 | 1 | 0.000367874 |
| Clpp | 2.93E-08 | 0.332767363 | 1 | 0.986 | 0.000378221 |
| Ccdc58 | 3.27E-08 | 0.34859948 | 1 | 1 | 0.000420926 |
| Cyb5a | 4.13E-08 | 0.314666792 | 1 | 1 | 0.000531879 |
| Tmem70 | 5.27E-08 | 0.30530854 | 1 | 1 | 0.000679274 |
| Jun | 5.50E-08 | 0.858074881 | 1 | 1 | 0.000709677 |
| Slmo2 | 5.51E-08 | 0.269298374 | 1 | 1 | 0.000710764 |
| Shmt2 | 5.87E-08 | 0.328353866 | 1 | 1 | 0.000756619 |
| Tomm22 | 6.38E-08 | 0.290573153 | 1 | 1 | 0.000821955 |
| Tmem147 | 8.29E-08 | 0.379263736 | 1 | 1 | 0.001068583 |
| Mt1 | 9.76E-08 | 1.045894733 | 0.986 | 1 | 0.001258335 |
| Gm10146 | 1.04E-07 | 0.304068118 | 1 | 0.986 | 0.00133937 |
| Ndufaf2 | 1.05E-07 | 0.26219524 | 1 | 0.973 | 0.001356457 |
| Gstm4 | 1.11E-07 | 0.294453074 | 0.914 | 0.77 | 0.001431755 |
| Pdia5 | 1.22E-07 | 0.385067242 | 0.986 | 0.959 | 0.001567039 |
| Tagln2 | 1.30E-07 | 0.304077843 | 1 | 1 | 0.00167841 |
| Mrpl39 | 1.34E-07 | 0.290777224 | 1 | 0.986 | 0.001728952 |
| Set | 1.50E-07 | 0.331007901 | 1 | 1 | 0.001939351 |
| Tmem41a | 1.57E-07 | 0.36816442 | 0.986 | 0.973 | 0.002021012 |
| Tars | 1.59E-07 | 0.333467526 | 1 | 0.986 | 0.002050745 |
| Sub1 | 1.70E-07 | 0.301810766 | 1 | 1 | 0.002194683 |
| Cnih1 | 1.93E-07 | 0.354213878 | 1 | 0.986 | 0.002492166 |
| Mrps31 | 2.03E-07 | 0.271019548 | 0.986 | 0.932 | 0.002622048 |
| Lsm7 | 2.21E-07 | 0.315563814 | 1 | 1 | 0.002851896 |

| | | | | | |
|---|---|---|---|---|---|
| Proc | 2.51E-07 | 0.28183302 | 0.814 | 0.554 | 0.00322969 |
| Psmd6 | 2.69E-07 | 0.251261721 | 1 | 1 | 0.003470202 |
| Tubb6 | 2.96E-07 | 0.35713479 | 1 | 0.986 | 0.003821803 |
| Uchl3 | 3.33E-07 | 0.316277841 | 1 | 0.973 | 0.004297038 |
| Eef1e1 | 3.51E-07 | 0.377291556 | 1 | 0.986 | 0.004529873 |
| Smdt1 | 3.82E-07 | 0.325905418 | 1 | 1 | 0.00492752 |
| Ndufa5 | 4.11E-07 | 0.268297783 | 1 | 1 | 0.005297355 |
| Mrps16 | 4.19E-07 | 0.337644962 | 1 | 0.986 | 0.005405336 |
| Fabp1 | 4.23E-07 | 0.318125818 | 0.629 | 0.405 | 0.005449625 |
| Spink3 | 4.26E-07 | 0.31872164 | 0.957 | 0.77 | 0.005492872 |
| Uqcr11 | 4.36E-07 | 0.255980031 | 1 | 1 | 0.005621476 |
| Ranbp1 | 4.51E-07 | 0.271210238 | 1 | 1 | 0.005814307 |
| Fkbp2 | 4.83E-07 | 0.346563369 | 1 | 0.959 | 0.006227478 |
| Tmem150a | 6.80E-07 | 0.347209478 | 0.971 | 0.851 | 0.008767696 |
| Mrpl36 | 9.00E-07 | 0.275136286 | 1 | 1 | 0.011604907 |
| Dnajb11 | 9.04E-07 | 0.284680776 | 1 | 0.986 | 0.011652869 |
| A430005L1 | 9.51E-07 | 0.251244178 | 1 | 0.946 | 0.012259527 |
| Mrpl32 | 1.23E-06 | 0.303660852 | 1 | 0.973 | 0.015837195 |
| Dkc1 | 1.32E-06 | 0.255810383 | 1 | 1 | 0.016964969 |
| Tmem176l | 1.39E-06 | 0.284621328 | 1 | 0.973 | 0.017918744 |
| Bckdhb | 1.65E-06 | 0.26241254 | 1 | 1 | 0.02125032 |
| Dbi | 1.79E-06 | 0.295301898 | 1 | 1 | 0.023057907 |
| Slc35b2 | 1.83E-06 | 0.267816726 | 1 | 0.973 | 0.023619825 |
| Fbl | 2.07E-06 | 0.405141339 | 1 | 1 | 0.026710736 |
| Tnfrsf12a | 2.09E-06 | 0.468276286 | 0.929 | 0.743 | 0.026981285 |
| Ndufb10 | 2.73E-06 | 0.333410633 | 1 | 1 | 0.035200371 |
| Gemin6 | 3.06E-06 | 0.268077483 | 1 | 0.865 | 0.039451757 |
| Spcs2 | 3.31E-06 | 0.275577045 | 1 | 1 | 0.042682011 |
| Deb1 | 3.79E-06 | 0.397375939 | 0.986 | 0.892 | 0.048815708 |
| Vdac1 | 3.79E-06 | 0.287658048 | 1 | 1 | 0.048892227 |
| Cdk1 | 3.83E-06 | 0.283381768 | 0.986 | 0.986 | 0.049394446 |
| Rogdi | 3.84E-06 | 0.328289521 | 1 | 0.959 | 0.049456374 |
| H2afx | 4.07E-06 | 0.456057138 | 1 | 1 | 0.052415749 |
| Atp5b | 4.34E-06 | 0.292963363 | 1 | 1 | 0.055934947 |
| Dnajb9 | 4.61E-06 | 0.369864842 | 0.986 | 0.959 | 0.059456668 |
| Gm5160 | 5.53E-06 | 0.261394257 | 1 | 1 | 0.071236624 |
| Ccnb1 | 8.49E-06 | 0.286634348 | 1 | 1 | 0.109449526 |
| Lyrm4 | 8.73E-06 | 0.284845705 | 1 | 0.946 | 0.11255623 |
| Cirh1a | 9.56E-06 | 0.254755416 | 1 | 0.986 | 0.123282944 |
| Cmss1 | 1.20E-05 | 0.259223476 | 0.971 | 0.892 | 0.155162202 |
| Nup37 | 1.82E-05 | 0.294379432 | 1 | 0.932 | 0.234110417 |
| Ubc | 1.84E-05 | 0.379215029 | 1 | 1 | 0.236788051 |
| Kdelr2 | 2.32E-05 | 0.261338989 | 1 | 1 | 0.299296398 |
| Mrpl33 | 3.49E-05 | 0.309105425 | 1 | 0.986 | 0.449489679 |
| Atf4 | 3.50E-05 | 0.451929064 | 1 | 1 | 0.451320071 |
| Dnaja2 | 3.57E-05 | 0.281905525 | 1 | 1 | 0.4606232 |
| Usmg5 | 3.70E-05 | 0.255720357 | 1 | 1 | 0.477134033 |
| Pnliprp1 | 4.46E-05 | 0.251591064 | 0.786 | 0.595 | 0.57450987 |
| Afm | 4.56E-05 | 0.261026992 | 0.943 | 0.905 | 0.588222512 |
| Ube2c | 5.83E-05 | 0.447012365 | 1 | 0.973 | 0.751784994 |
| Klf6 | 7.62E-05 | 0.339477805 | 1 | 0.959 | 0.98198356 |
| Fam107b | 0.00016101 | 0.258476998 | 0.971 | 0.959 | 1 |
| Aurka | 0.00016131 | 0.28128762 | 0.986 | 0.959 | 1 |
| Bub3 | 0.000175963 | 0.305571093 | 1 | 0.986 | 1 |
| Gm8186 | 0.000186476 | 0.280079965 | 1 | 1 | 1 |
| Snrpg | 0.000468895 | 0.320972221 | 1 | 1 | 1 |
| Gm10184 | 0.000665914 | 0.270125216 | 1 | 0.986 | 1 |
| Zfp36 | 0.000754639 | 0.256418471 | 0.943 | 0.865 | 1 |

| | | | | | |
|---|---|---|---|---|---|
| Kpna2 | 0.000783495 | 0.345870648 | 1 | 1 | 1 |
| Sqstm1 | 0.001129384 | 0.315982129 | 1 | 1 | 1 |
| Ier2 | 0.001320651 | 0.493286093 | 1 | 0.986 | 1 |
| Mt2 | 0.005072101 | 0.41931125 | 0.943 | 0.824 | 1 |

### s from E9.5-E10.5

given cell cluster and the left cell clusters. Positive values indicate that the gene is

5
all genes in the dataset.

| cluster | gene | Descriptio | Location | Family |
|---|---|---|---|---|
| Neural Tube | Adgrv1 | adhesion ( | Plasma Me | G-protein coupled re |
| Neural Tube | Tubb3 | tubulin be | Cytoplasm | other |
| Neural Tube | Lhfp | LHFPL tetr | Other | other |
| Neural Tube | Ngfr | nerve grov | Plasma Me | transmembrane recep |
| Neural Tube | Adcyap1r1 | ADCYAP r | Plasma Me | G-protein coupled re |
| Neural Tube | Cdh11 | cadherin 1 | Plasma Me | other |
| Neural Tube | Nes | nestin | Cytoplasm | other |
| Neural Tube | Scube1 | signal pep | Plasma Me | transmembrane recep |
| Neural Tube | Celsr2 | cadherin E | Plasma Me | G-protein coupled re |
| Neural Tube | Zeb2 | zinc finger | Nucleus | transcription regulato |
| Neural Tube | Ncam1 | neural cell | Plasma Me | other |
| Neural Tube | Plxna2 | plexin A2 | Plasma Me | transmembrane recep |
| Neural Tube | Lrrn1 | leucine ric | Other | other |
| Neural Tube | Ednrb | endothelir | Plasma Me | G-protein coupled re |
| Neural Tube | Slit1 | slit guidan | Extracellul | other |
| Neural Tube | Jam2 | junctional | Plasma Me | other |
| Neural Tube | Sema5b | semaphori | Plasma Me | other |
| Neural Tube | Phox2b | paired like | Nucleus | transcription regulato |
| Neural Tube | Fgfr3 | fibroblast | Plasma Me | kinase |
| Neural Tube | Rfx4 | regulatory | Nucleus | transcription regulato |
| Neural Tube | Srgap3 | SLIT-ROB( | Cytoplasm | other |
| Neural Tube | Notch1 | notch 1 | Plasma Me | transcription regulato |
| Neural Tube | Rgma | repulsive g | Plasma Me | other |
| Neural Tube | Entpd7 | ectonuclec | Cytoplasm | enzyme |
| Neural Tube | Nrcam | neuronal c | Plasma Me | other |
| Neural Tube | Elavl4 | ELAV like I | Cytoplasm | other |
| Neural Tube | Dll3 | delta like c | Extracellul | other |
| Neural Tube | Osbpl7 | oxysterol k | Cytoplasm | transporter |
| Neural Tube | Olfml3 | olfactome | Extracellul | other |
| Neural Tube | Adgrl2 | adhesion ( | Plasma Me | G-protein coupled re |
| Neural Tube | Igsf9b | immunogl | Plasma Me | other |
| Neural Tube | Sulf2 | sulfatase 2 | Plasma Me | enzyme |
| Neural Tube | Ptpru | protein tyr | Plasma Me | phosphatase |
| Neural Tube | Tulp4 | tubby like | Cytoplasm | transcription regulato |
| Neural Tube | Tox2 | TOX high | Nucleus | transcription regulato |
| Neural Tube | Tppp3 | tubulin po | Cytoplasm | other |
| Neural Tube | Gm4707 | predicted | Other | other |
| Neural Tube | Mn1 | MN1 prot | Nucleus | other |
| Neural Tube | Spon1 | spondin 1 | Extracellul | other |
| Neural Tube | Ncor2 | nuclear re | Nucleus | transcription regulato |
| Neural Tube | Vangl2 | VANGL pla | Plasma Me | other |
| Neural Tube | Ntn1 | netrin 1 | Extracellul | other |
| Neural Tube | Cdc42bpb | CDC42 bir | Cytoplasm | kinase |
| Neural Tube | Sulf1 | sulfatase 1 | Cytoplasm | enzyme |

| | | | | |
|---|---|---|---|---|
| Neural Tube | Nell2 | neural EGF | Extracellul | other |
| Neural Tube | Ckb | creatine ki | Cytoplasm | kinase |
| Neural Tube | Lfng | LFNG O-fu | Cytoplasm | enzyme |
| Neural Tube | 2310067B1 | - | - | - |
| Neural Tube | Syt11 | synaptotag | Cytoplasm | transporter |
| Neural Tube | Map1b | microtubu | Cytoplasm | other |
| Neural Tube | 5031439G( | KIAA0930 | Other | other |
| Neural Tube | Slit3 | slit guidan | Extracellul | other |
| Neural Tube | Sfrp1 | secreted fr | Plasma Me | transmembrane recep |
| Neural Tube | Zfp11 | zinc finger | Nucleus | other |
| Neural Tube | Zc3h12c | zinc finger | Other | other |
| Neural Tube | Stxbp1 | syntaxin bi | Cytoplasm | transporter |
| Neural Tube | Prtg | protogenir | Extracellul | other |
| Neural Tube | Atp8a1 | ATPase ph | Cytoplasm | transporter |
| Neural Tube | Map4k4 | mitogen-a | Cytoplasm | kinase |
| Neural Tube | Itsn1 | intersectin | Cytoplasm | other |
| Neural Tube | Crb2 | crumbs 2, | Extracellul | other |
| Neural Tube | Dtx4 | deltex E3 u | Cytoplasm | enzyme |
| Neural Tube | Reep1 | receptor a | Cytoplasm | other |
| Neural Tube | Adgrg1 | adhesion ( | Plasma Me | G-protein coupled re |
| Neural Tube | Cacna1g | calcium vo | Plasma Me | ion channel |
| Neural Tube | Ddr1 | discoidin c | Plasma Me | kinase |
| Neural Tube | Pcdhgc3 | protocadh | Plasma Me | other |
| Neural Tube | Eml1 | echinoderi | Cytoplasm | other |
| Neural Tube | Ube4a | ubiquitinat | Cytoplasm | enzyme |
| Neural Tube | Igdcc3 | immunogl | Other | other |
| Neural Tube | Uggt1 | UDP-gluco | Cytoplasm | enzyme |
| Neural Tube | Nr2f1 | nuclear re | Nucleus | ligand-dependent nu |
| Neural Tube | Fabp7 | fatty acid b | Cytoplasm | transporter |
| Neural Tube | Pou2f1 | POU class | Nucleus | transcription regulato |
| Neural Tube | Prkar2a | protein kir | Cytoplasm | kinase |
| Neural Tube | Trappc8 | trafficking | Cytoplasm | transporter |
| Neural Tube | Sorcs2 | sortilin rela | Plasma Me | transporter |
| Neural Tube | Maml3 | mastermin | Nucleus | transcription regulato |
| Neural Tube | Col22a1 | collagen ty | Extracellul | other |
| Neural Tube | Brsk2 | BR serine/t | Cytoplasm | kinase |
| Neural Tube | Sept3 | septin 3 | Cytoplasm | enzyme |
| Neural Tube | Nuak2 | NUAK fam | Other | kinase |
| Neural Tube | Cdh2 | cadherin 2 | Plasma Me | other |
| Neural Tube | Vim | vimentin | Cytoplasm | other |
| Neural Tube | Pknox2 | PBX/knotte | Nucleus | other |
| Neural Tube | Nkx6-1 | NK6 home | Nucleus | transcription regulato |
| Neural Tube | Slc45a3 | solute carr | Cytoplasm | transporter |
| Neural Tube | Tcf12 | transcriptio | Nucleus | transcription regulato |
| Neural Tube | Tmem219 | transmeml | Plasma Me | other |
| Neural Tube | Ptch1 | patched 1 | Plasma Me | transmembrane recep |
| Neural Tube | Arhgap35 | Rho GTPas | Nucleus | transcription regulato |
| Neural Tube | Prom1 | prominin 1 | Plasma Me | other |
| Neural Tube | Rsg1 | REM2 and | Cytoplasm | other |
| Neural Tube | Jam3 | junctional | Plasma Me | other |
| Neural Tube | Ptpn21 | protein tyr | Cytoplasm | phosphatase |
| Neural Tube | Mfng | MFNG O-f | Cytoplasm | enzyme |
| Neural Tube | Rnf123 | ring finger | Cytoplasm | other |
| Neural Tube | Zswim4 | zinc finger | Other | other |
| Neural Tube | Scrn1 | secernin 1 | Cytoplasm | other |
| Neural Tube | Rcor1 | REST corep | Nucleus | transcription regulato |
| Neural Tube | Ptprs | protein tyr | Plasma Me | phosphatase |
| Neural Tube | Fez1 | fasciculatio | Cytoplasm | other |

| | | | | |
|---|---|---|---|---|
| Neural Tube | Myo19 | myosin XIX | Cytoplasm | enzyme |
| Neural Tube | Sall1 | spalt like t | Nucleus | transcription regulato |
| Neural Tube | Hoxd3 | homeobox | Nucleus | transcription regulato |
| Neural Tube | Tmem47 | transmeml | Plasma Me | other |
| Neural Tube | Tfdp2 | transcriptio | Nucleus | transcription regulato |
| Neural Tube | Itgb8 | integrin su | Plasma Me | other |
| Neural Tube | Pea15a | proliferatio | Cytoplasm | transporter |
| Neural Tube | Ttyh2 | tweety fan | Other | ion channel |
| Neural Tube | Nsun3 | NOP2/Sur | Cytoplasm | enzyme |
| Neural Tube | Slc35e2 | solute carr | Other | other |
| Neural Tube | Hes5 | hes family | Nucleus | transcription regulato |
| Neural Tube | E130308A1 | RIKEN cDN | Other | other |
| Neural Tube | Pik3c2b | phosphatio | Cytoplasm | kinase |
| Neural Tube | Cdh22 | cadherin 2 | Plasma Me | other |
| Neural Tube | Gpc1 | glypican 1 | Plasma Me | transmembrane recep |
| Neural Tube | 1700106J1 | - | - | - |
| Neural Tube | Cercam | cerebral er | Other | other |
| Neural Tube | Prmt10 | - | - | - |
| Neural Tube | Ankrd11 | ankyrin rep | Nucleus | other |
| Neural Tube | Msn | moesin | Plasma Me | other |
| Neural Tube | Lrp1 | LDL recept | Plasma Me | transmembrane recep |
| Neural Tube | Dgcr2 | DiGeorge s | Plasma Me | other |
| Neural Tube | Tnfaip6 | TNF alpha | Extracellul | other |
| Neural Tube | Ly6h | lymphocyt | Plasma Me | other |
| Neural Tube | Tyw1 | tRNA-yW | Other | other |
| Neural Tube | Fbln2 | fibulin 2 | Extracellul | other |
| Neural Tube | Rassf4 | Ras associ | Other | other |
| Neural Tube | Akna | AT-hook t | Nucleus | transcription regulato |
| Neural Tube | Nr6a1 | nuclear rec | Nucleus | ligand-dependent nu |
| Neural Tube | Ephb1 | EPH recep | Plasma Me | kinase |
| Neural Tube | Slc38a6 | solute carr | Plasma Me | transporter |
| Neural Tube | Zgrf1 | zinc finger | Other | other |
| Neural Tube | Ndufb2 | NADH:ubi | Cytoplasm | enzyme |
| Neural Tube | Ep400 | E1A bindir | Nucleus | other |
| Neural Tube | Jade1 | jade family | Nucleus | transcription regulato |
| Neural Tube | Qsox2 | quiescin su | Other | enzyme |
| Neural Tube | Slc27a4 | solute carr | Plasma Me | transporter |
| Neural Tube | Rps6kc1 | ribosomal | Cytoplasm | kinase |
| Neural Tube | Ascl1 | achaete-sc | Nucleus | transcription regulato |
| Neural Tube | Strada | STE20-rela | Nucleus | kinase |
| Neural Tube | Tmx4 | thioredoxi | Cytoplasm | enzyme |
| Neural Tube | Klhdc8b | kelch dom | Cytoplasm | other |
| Neural Tube | Acvr2b | activin A re | Plasma Me | kinase |
| Neural Tube | Cblb | Cbl proto- | Nucleus | enzyme |
| Neural Tube | Usp42 | ubiquitin s | Nucleus | peptidase |
| Neural Tube | Tgfbr1 | transformi | Plasma Me | kinase |
| Neural Tube | Tagln3 | transgelin | Extracellul | other |
| Neural Tube | Uros | uroporphy | Cytoplasm | enzyme |
| Neural Tube | Zfp521 | zinc finger | Nucleus | other |
| Neural Tube | Samd14 | sterile alph | Other | other |
| Neural Tube | Tmem2 | transmeml | Cytoplasm | enzyme |
| Neural Tube | Plbd2 | phospholip | Extracellul | other |
| Neural Tube | Cdc25b | cell divisio | Nucleus | phosphatase |
| Neural Tube | Xndc1 | Xrcc1 N-te | Other | other |
| Neural Tube | Npl | N-acetylne | Cytoplasm | enzyme |
| Neural Tube | Rpusd2 | RNA pseud | Other | enzyme |
| Neural Tube | Efnb3 | ephrin B3 | Plasma Me | kinase |
| Neural Tube | Rnf31 | ring finger | Cytoplasm | enzyme |

| | | | | |
|---|---|---|---|---|
| Neural Tube | Cog5 | componer | Cytoplasm | transporter |
| Neural Tube | Crmp1 | collapsin r | Cytoplasm | enzyme |
| Neural Tube | Rgmb | repulsive g | Plasma Me | other |
| Neural Tube | Epha7 | EPH recep | Plasma Me | kinase |
| Neural Tube | Etnk1 | ethanolam | Cytoplasm | kinase |
| Neural Tube | Rhobtb3 | Rho relate | Cytoplasm | enzyme |
| Neural Tube | Dhx32 | DEAH-box | Other | enzyme |
| Neural Tube | Pcnxl3 | - | - | - |
| Neural Tube | Dtl | denticleles | Nucleus | other |
| Neural Tube | Nup153 | nucleopor | Nucleus | other |
| Neural Tube | Btbd17 | BTB doma | Plasma Me | other |
| Neural Tube | Zfp329 | zinc finger | Other | other |
| Neural Tube | Tcof1 | treacle ribo | Nucleus | transporter |
| Neural Tube | Camsap1 | calmodulir | Cytoplasm | other |
| Neural Tube | Myh10 | myosin he | Cytoplasm | enzyme |
| Neural Tube | Fgfbp3 | fibroblast g | Extracellula | other |
| Neural Tube | Rtn1 | reticulon 1 | Cytoplasm | other |
| Neural Tube | Plxna1 | plexin A1 | Plasma Me | transmembrane recep |
| Neural Tube | Cc2d1a | coiled-coi | Cytoplasm | transcription regulato |
| Neural Tube | Ubn1 | ubinuclein | Nucleus | transcription regulato |
| Neural Tube | Cbl | Cbl proto- | Nucleus | transcription regulato |
| Neural Tube | Pigm | phosphatic | Cytoplasm | enzyme |
| Neural Tube | Morc4 | MORC fam | Nucleus | other |
| Neural Tube | Ddit4l | DNA dama | Cytoplasm | other |
| Neural Tube | Zfp516 | zinc finger | Nucleus | other |
| Neural Tube | Oxct1 | 3-oxoacid | Cytoplasm | enzyme |
| Neural Tube | Msi1 | musashi R | Cytoplasm | other |
| Neural Tube | Meis2 | Meis home | Nucleus | transcription regulato |
| Neural Tube | Ermp1 | endoplasm | Cytoplasm | other |
| Neural Tube | Dpysl4 | dihydropy | Cytoplasm | enzyme |
| Neural Tube | Extl3 | exostosin l | Cytoplasm | enzyme |
| Neural Tube | Osgin2 | oxidative s | Other | other |
| Neural Tube | Brca1 | BRCA1, DN | Nucleus | transcription regulato |
| Neural Tube | Sik3 | SIK family | Cytoplasm | kinase |
| Neural Tube | 2510009E0 | chromosoi | Other | other |
| Neural Tube | Stxbp6 | syntaxin bi | Cytoplasm | other |
| Neural Tube | Grik4 | glutamate | Plasma Me | ion channel |
| Neural Tube | Ankrd13b | ankyrin rep | Plasma Me | other |
| Neural Tube | Wdr76 | WD repeat | Nucleus | other |
| Neural Tube | Brf2 | BRF2, RNA | Nucleus | transcription regulato |
| Neural Tube | Ext1 | exostosin g | Cytoplasm | enzyme |
| Neural Tube | Spsb4 | splA/ryanc | Cytoplasm | other |
| Neural Tube | Mxra7 | matrix-ren | Extracellula | other |
| Neural Tube | Manea | mannosida | Cytoplasm | enzyme |
| Neural Tube | Stat5b | signal tran | Nucleus | transcription regulato |
| Neural Tube | Ube4b | ubiquitinat | Cytoplasm | enzyme |
| Neural Tube | Ttll4 | tubulin tyr | Cytoplasm | enzyme |
| Neural Tube | Tmtc2 | transmeml | Cytoplasm | other |
| Neural Tube | Tmed7 | transmeml | Cytoplasm | transporter |
| Neural Tube | Bscl2 | BSCL2, sei | Cytoplasm | other |
| Neural Tube | Cep135 | centrosom | Cytoplasm | other |
| Neural Tube | Recql4 | RecQ like l | Nucleus | enzyme |
| Neural Tube | Prpf8 | pre-mRNA | Nucleus | other |
| Neural Tube | Vac14 | Vac14, PIK | Cytoplasm | other |
| Neural Tube | Cabin1 | calcineurir | Nucleus | other |
| Neural Tube | Map6 | microtubu | Cytoplasm | other |
| Neural Tube | Slc43a1 | solute carr | Plasma Me | transporter |
| Neural Tube | Vps54 | VPS54, GA | Extracellula | other |

| | | | | |
|---|---|---|---|---|
| Neural Tube | 1190002N | chromosoı | Cytoplasm | other |
| Neural Tube | Fanca | Fanconi ar | Nucleus | other |
| Neural Tube | Lancl2 | LanC like 2 | Plasma Me | other |
| Neural Tube | Tmem33 | transmeml | Cytoplasm | other |
| Neural Tube | Kdm4a | lysine dem | Nucleus | transcription regulato |
| Neural Tube | Espl1 | extra spinc | Nucleus | peptidase |
| Neural Tube | Ccdc57 | coiled-coi | Other | other |
| Neural Tube | Timp2 | TIMP meta | Extracellula | other |
| Neural Tube | Zfp518b | zinc finger | Other | other |
| Neural Tube | Dll1 | delta like c | Plasma Me | enzyme |
| Neural Tube | Cpe | carboxype | Cytoplasm | peptidase |
| Neural Tube | Gm10330 | - | - | - |
| Neural Tube | Elovl4 | ELOVL fatt | Cytoplasm | enzyme |
| Neural Tube | Tubb2b | tubulin be | Cytoplasm | other |
| Neural Tube | Man2a2 | mannosida | Cytoplasm | enzyme |
| Neural Tube | Itgb3bp | integrin su | Nucleus | other |
| Neural Tube | Rbmx | RNA bindi | Nucleus | other |
| Neural Tube | Zfp146 | zinc finger | Nucleus | other |
| Neural Tube | Tmcc2 | transmeml | Extracellula | other |
| Neural Tube | Cmip | c-Maf indu | Cytoplasm | other |
| Neural Tube | Taok2 | TAO kinas | Cytoplasm | kinase |
| Neural Tube | Rai14 | retinoic ac | Nucleus | transcription regulato |
| Neural Tube | Atf6 | activating | Cytoplasm | transcription regulato |
| Neural Tube | Ccdc15 | coiled-coi | Cytoplasm | other |
| Neural Tube | Cptp | ceramide- | Cytoplasm | transporter |
| Neural Tube | Ext2 | exostosin g | Cytoplasm | enzyme |
| Neural Tube | Galnt16 | polypeptic | Cytoplasm | enzyme |
| Neural Tube | Pcgf1 | polycomb | Nucleus | other |
| Neural Tube | Dvl2 | dishevelle | Cytoplasm | other |
| Neural Tube | Pnpla2 | patatin like | Cytoplasm | enzyme |
| Neural Tube | Gadd45g | growth arr | Nucleus | other |
| Neural Tube | Slc39a7 | solute carr | Plasma Me | transporter |
| Neural Tube | Glce | glucuronic | Cytoplasm | enzyme |
| Neural Tube | Zcchc14 | zinc finger | Other | other |
| Neural Tube | Tmem35 | - | - | - |
| Neural Tube | Zfp664 | zinc finger | Nucleus | other |
| Neural Tube | Ece1 | endothelir | Plasma Me | peptidase |
| Neural Tube | Rnf220 | ring finger | Cytoplasm | enzyme |
| Neural Tube | Dync1h1 | dynein cyt | Cytoplasm | peptidase |
| Neural Tube | Ints5 | integrator | Nucleus | other |
| Neural Tube | Zgpat | zinc finger | Nucleus | transcription regulato |
| Neural Tube | Mcm10 | minichrom | Nucleus | other |
| Neural Tube | Fyn | FYN proto | Plasma Me | kinase |
| Neural Tube | Dmtf1 | cyclin D bi | Nucleus | transcription regulato |
| Neural Tube | Spata13 | spermatog | Plasma Me | other |
| Neural Tube | Emc1 | ER membr | Plasma Me | other |
| Neural Tube | Gnai1 | G protein s | Plasma Me | enzyme |
| Neural Tube | Cad | carbamoyl | Cytoplasm | enzyme |
| Neural Tube | Clasp1 | cytoplasm | Cytoplasm | other |
| Neural Tube | March6 | membrane | Cytoplasm | enzyme |
| Neural Tube | Ndst1 | N-deacety | Cytoplasm | enzyme |
| Neural Tube | Hus1 | HUS1 chec | Nucleus | kinase |
| Neural Tube | Neo1 | neogenin | Plasma Me | transcription regulato |
| Neural Tube | Tmem201 | transmeml | Nucleus | other |
| Neural Tube | Fance | Fanconi ar | Nucleus | other |
| Neural Tube | Ktn1 | kinectin 1 | Cytoplasm | other |
| Neural Tube | Sh3kbp1 | SH3 doma | Cytoplasm | other |
| Neural Tube | Ehmt1 | euchroma | Nucleus | transcription regulato |

| | | | | |
|---|---|---|---|---|
| Neural Tube | Pomt2 | protein O- | Cytoplasm | enzyme |
| Neural Tube | Rcc2 | regulator c | Nucleus | other |
| Neural Tube | Trrap | transforma | Nucleus | transcription regulato |
| Neural Tube | Abi2 | abl interac | Cytoplasm | other |
| Neural Tube | Pcnxl4 | - | - | - |
| Neural Tube | B3galnt2 | beta-1,3-N | Cytoplasm | enzyme |
| Neural Tube | Peg12 | paternally | Other | other |
| Neural Tube | Fam188b | - | - | - |
| Neural Tube | Rcor2 | REST core | Nucleus | transcription regulato |
| Neural Tube | Smarcd1 | SWI/SNF r | Nucleus | transcription regulato |
| Neural Tube | Slc52a2 | solute carr | Plasma Me | transporter |
| Neural Tube | Ezh2 | enhancer c | Nucleus | transcription regulato |
| Neural Tube | Rnd2 | Rho family | Cytoplasm | enzyme |
| Neural Tube | Emilin2 | elastin mic | Extracellul | other |
| Neural Tube | Prpf3 | pre-mRNA | Nucleus | other |
| Neural Tube | Mafg | MAF bZIP | Nucleus | transcription regulato |
| Neural Tube | Scrib | scribbled p | Cytoplasm | other |
| Neural Tube | Zfp687 | zinc finger | Nucleus | other |
| Neural Tube | Dnajb2 | DnaJ heat | Nucleus | other |
| Neural Tube | Clasp2 | cytoplasm | Cytoplasm | other |
| Neural Tube | Plch1 | phospholi | Cytoplasm | enzyme |
| Neural Tube | Cstf2 | cleavage s | Nucleus | other |
| Neural Tube | Fzr1 | fizzy and c | Nucleus | kinase |
| Neural Tube | Abcb8 | ATP bindir | Cytoplasm | transporter |
| Neural Tube | Zfp41 | ZFP41 zinc | Nucleus | other |
| Neural Tube | Pomc | proopiome | Extracellul | other |
| Neural Tube | F2r | coagulatio | Plasma Me | G-protein coupled re |
| Neural Tube | Zfp59 | zinc finger | Nucleus | other |
| Neural Tube | Zfp445 | zinc finger | Nucleus | transcription regulato |
| Neural Tube | Irs2 | insulin rec | Cytoplasm | enzyme |
| Neural Tube | Rragd | Ras relatec | Cytoplasm | enzyme |
| Neural Tube | Ints1 | integrator | Nucleus | other |
| Neural Tube | Map4 | microtubu | Cytoplasm | other |
| Neural Tube | Pltp | phospholi | Extracellul | enzyme |
| Neural Tube | Adpgk | ADP deper | Plasma Me | kinase |
| Neural Tube | Kif5c | kinesin fan | Cytoplasm | enzyme |
| Neural Tube | Zfp568 | zinc finger | Nucleus | other |
| Neural Tube | Sptlc2 | serine palr | Cytoplasm | enzyme |
| Neural Tube | Raver1 | ribonucleo | Nucleus | other |
| Neural Tube | Xpo5 | exportin 5 | Nucleus | transporter |
| Neural Tube | Rfc1 | replication | Nucleus | transcription regulato |
| Neural Tube | Add1 | adducin 1 | Cytoplasm | other |
| Neural Tube | Pds5a | PDS5 cohe | Nucleus | other |
| Neural Tube | Gnl3l | G protein | Nucleus | other |
| Neural Tube | Tmco3 | transmeml | Other | transporter |
| Neural Tube | Llgl1 | LLGL1, scri | Cytoplasm | other |
| Neural Tube | Champ1 | chromosor | Nucleus | other |
| Neural Tube | Scarb2 | scavenger | Plasma Me | other |
| Neural Tube | Fat1 | FAT atypic | Plasma Me | other |
| Neural Tube | Arnt2 | aryl hydro | Nucleus | transcription regulato |
| Neural Tube | Kat6a | lysine acet | Nucleus | enzyme |
| Neural Tube | Lmnb2 | lamin B2 | Nucleus | other |
| Neural Tube | Scube3 | signal pep | Plasma Me | other |
| Neural Tube | Zfp689 | zinc finger | Nucleus | transcription regulato |
| Neural Tube | Ttll12 | tubulin tyr | Other | other |
| Neural Tube | Odf2 | outer dens | Cytoplasm | other |
| Neural Tube | Tmem191c | transmeml | Other | other |
| Neural Tube | Rbl1 | RB transcri | Nucleus | transcription regulato |

| | | | | |
|---|---|---|---|---|
| Neural Tube | Foxn3 | forkhead b | Other | other |
| Neural Tube | Nup155 | nucleopor | Nucleus | transporter |
| Neural Tube | Ipo9 | importin 9 | Nucleus | transporter |
| Neural Tube | Tmem167l | transmeml | Other | other |
| Neural Tube | Ino80e | INO80 cor | Nucleus | other |
| Neural Tube | Prim2 | DNA prima | Nucleus | enzyme |
| Neural Tube | Slc35e1 | solute carr | Cytoplasm | other |
| Neural Tube | Cadm1 | cell adhesi | Plasma Me | other |
| Neural Tube | Adck2 | aarF doma | Cytoplasm | kinase |
| Neural Tube | Rad17 | RAD17 che | Nucleus | enzyme |
| Neural Tube | Hes6 | hes family | Nucleus | transcription regulato |
| Neural Tube | Akr1b3 | aldo-keto | Cytoplasm | enzyme |
| Neural Tube | Sfrp2 | secreted fr | Plasma Me | transmembrane recep |
| Neural Tube | Zmiz1 | zinc finger | Nucleus | other |
| Neural Tube | Osbpl11 | oxysterol b | Cytoplasm | other |
| Neural Tube | Zik1 | zinc finger | Other | other |
| Neural Tube | Rpgr | retinitis pig | Cytoplasm | other |
| Neural Tube | Skiv2l | Ski2 like Rl | Nucleus | enzyme |
| Neural Tube | Igf2bp1 | insulin like | Cytoplasm | translation regulator |
| Neural Tube | Fstl1 | follistatin l | Extracellul | other |
| Neural Tube | Golm1 | golgi mem | Cytoplasm | other |
| Neural Tube | Zbtb17 | zinc finger | Nucleus | transcription regulato |
| Neural Tube | Npepps | aminopep | Cytoplasm | peptidase |
| Neural Tube | Ttf1 | transcriptio | Nucleus | other |
| Neural Tube | Epb4.1 | - | - | - |
| Neural Tube | Mdc1 | mediator c | Nucleus | other |
| Neural Tube | Gng2 | G protein s | Plasma Me | enzyme |
| Neural Tube | Kif21a | kinesin fan | Cytoplasm | other |
| Neural Tube | Dleu7 | deleted in | Other | other |
| Neural Tube | Tnfrsf21 | TNF recep | Plasma Me | transmembrane recep |
| Neural Tube | Slc7a6 | solute carr | Plasma Me | transporter |
| Neural Tube | Mycl | MYCL prot | Nucleus | transcription regulato |
| Neural Tube | Snrnp200 | small nucle | Nucleus | enzyme |
| Neural Tube | Haus8 | HAUS aug | Cytoplasm | other |
| Neural Tube | Fkbp5 | FK506 bin | Nucleus | enzyme |
| Neural Tube | Ptms | parathymo | Cytoplasm | other |
| Neural Tube | Fam171a1 | family with | Cytoplasm | other |
| Neural Tube | Lman2l | lectin, mar | Cytoplasm | other |
| Neural Tube | Ubqln4 | ubiquilin 4 | Cytoplasm | other |
| Neural Tube | Gpm6b | glycoprote | Plasma Me | other |
| Neural Tube | Pom121 | POM121 t | Nucleus | other |
| Neural Tube | Wdr33 | WD repeat | Nucleus | other |
| Neural Tube | E130309D0 | chromosor | Nucleus | other |
| Neural Tube | Taf1c | TATA-box | Nucleus | transcription regulato |
| Neural Tube | Map2k7 | mitogen-a | Cytoplasm | kinase |
| Neural Tube | Slc12a7 | solute carr | Plasma Me | transporter |
| Neural Tube | Naa60 | N(alpha)-a | Cytoplasm | enzyme |
| Neural Tube | Ncln | nicalin | Cytoplasm | peptidase |
| Neural Tube | Man2b2 | mannosida | Cytoplasm | enzyme |
| Neural Tube | Vwa9 | - | - | - |
| Neural Tube | Slc9a6 | solute carr | Plasma Me | transporter |
| Neural Tube | Clip3 | CAP-Gly d | Plasma Me | other |
| Neural Tube | Tmem101 | transmeml | Extracellul | other |
| Neural Tube | Cdc42se2 | CDC42 sm | Plasma Me | other |
| Neural Tube | Zfp113 | zinc finger | Nucleus | transcription regulato |
| Neural Tube | Crabp1 | cellular ret | Cytoplasm | transporter |
| Neural Tube | Six5 | SIX homeo | Nucleus | transcription regulato |
| Neural Tube | Zfp944 | zinc finger | Other | other |

| | | | | |
|---|---|---|---|---|
| Neural Tube | Loxl2 | lysyl oxida | Extracellul | enzyme |
| Neural Tube | Pole2 | DNA polyr | Nucleus | enzyme |
| Neural Tube | Ppm1l | protein ph | Cytoplasm | phosphatase |
| Neural Tube | Elovl6 | ELOVL fatt | Cytoplasm | enzyme |
| Neural Tube | Carkd | - | - | - |
| Neural Tube | Pprc1 | peroxisom | Nucleus | transcription regulato |
| Neural Tube | Ctu2 | cytosolic tl | Cytoplasm | other |
| Neural Tube | Fzd3 | frizzled cla | Plasma Me | G-protein coupled re |
| Neural Tube | Bptf | bromodor | Nucleus | transcription regulato |
| Neural Tube | E2f2 | E2F transc | Nucleus | transcription regulato |
| Neural Tube | Pcyox1 | prenylcyst | Cytoplasm | enzyme |
| Neural Tube | Gpaa1 | glycosylph | Cytoplasm | enzyme |
| Neural Tube | Zdhhc16 | zinc finger | Cytoplasm | enzyme |
| Neural Tube | Gpd2 | glycerol-3 | Cytoplasm | enzyme |
| Neural Tube | Dpysl5 | dihydropy | Cytoplasm | enzyme |
| Neural Tube | Shroom2 | shroom fa | Plasma Me | ion channel |
| Neural Tube | Npdc1 | neural pro | Extracellul | other |
| Neural Tube | Hap1 | huntingtin | Cytoplasm | other |
| Neural Tube | Mettl17 | methyltrar | Nucleus | other |
| Neural Tube | Rbm4b | RNA bindi | Nucleus | other |
| Neural Tube | Zfp207 | zinc finger | Nucleus | transcription regulato |
| Neural Tube | Myo1e | myosin IE | Cytoplasm | enzyme |
| Neural Tube | Pak2 | p21 (RAC1 | Cytoplasm | kinase |
| Neural Tube | Casp8ap2 | caspase 8 | Nucleus | transcription regulato |
| Neural Tube | Smad2 | SMAD fam | Nucleus | transcription regulato |
| Neural Tube | Heatr3 | HEAT repe | Other | other |
| Neural Tube | Sufu | SUFU nega | Nucleus | transcription regulato |
| Neural Tube | Hipk2 | homeodor | Nucleus | kinase |
| Neural Tube | Hmgxb4 | HMG-box | Nucleus | other |
| Neural Tube | Phf6 | PHD finge | Nucleus | transcription regulato |
| Neural Tube | Aacs | acetoacety | Cytoplasm | enzyme |
| Neural Tube | Pcbp3 | poly(rC) bi | Nucleus | transcription regulato |
| Neural Tube | Zfp511 | zinc finger | Nucleus | other |
| Neural Tube | Zswim5 | zinc finger | Extracellul | other |
| Neural Tube | Itfg2 | integrin al | Cytoplasm | other |
| Neural Tube | Trib2 | tribbles ps | Plasma Me | kinase |
| Neural Tube | Nxf1 | nuclear RN | Nucleus | other |
| Neural Tube | Prkrir | - | - | - |
| Neural Tube | Plagl2 | PLAG1 like | Nucleus | transcription regulato |
| Neural Tube | Acin1 | apoptotic | Nucleus | enzyme |
| Neural Tube | Snapc3 | small nucle | Nucleus | other |
| Neural Tube | Bysl | bystin like | Nucleus | other |
| Neural Tube | Slc46a1 | solute carr | Plasma Me | transporter |
| Neural Tube | Casp2 | caspase 2 | Cytoplasm | peptidase |
| Neural Tube | Zfp367 | zinc finger | Nucleus | transcription regulato |
| Neural Tube | Ptbp2 | polypyrimi | Nucleus | other |
| Neural Tube | Trp53inp2 | tumor pro | Nucleus | other |
| Neural Tube | Pkmyt1 | protein kir | Cytoplasm | kinase |
| Neural Tube | Clstn1 | calsynteni | Plasma Me | other |
| Neural Tube | Plk3 | polo like k | Nucleus | kinase |
| Neural Tube | Thumpd2 | THUMP dc | Other | enzyme |
| Neural Tube | Mllt1 | MLLT1, su | Nucleus | transcription regulato |
| Neural Tube | Dbn1 | drebrin 1 | Cytoplasm | other |
| Neural Tube | Alg10b | ALG10, alp | Cytoplasm | enzyme |
| Neural Tube | Zfp36l1 | ZFP36 ring | Nucleus | transcription regulato |
| Neural Tube | Fam172a | family with | Cytoplasm | transcription regulato |
| Neural Tube | Nkx2-2 | NK2 home | Nucleus | transcription regulato |
| Neural Tube | Lgmn | legumain | Cytoplasm | peptidase |

| | | | | |
|---|---|---|---|---|
| Neural Tube | Eif5b | eukaryotic | Cytoplasm | translation regulator |
| Neural Tube | Sppl2b | signal pep | Plasma Me | peptidase |
| Neural Tube | Mkl1 | megakaryo | Nucleus | transcription regulato |
| Neural Tube | Panx1 | pannexin 1 | Plasma Me | transporter |
| Neural Tube | Poc1b | POC1 cent | Cytoplasm | other |
| Neural Tube | Bclaf1 | BCL2 asso | Nucleus | transcription regulato |
| Neural Tube | Epha4 | EPH recep | Plasma Me | kinase |
| Neural Tube | Efna5 | ephrin A5 | Plasma Me | kinase |
| Neural Tube | Zbtb22 | zinc finger | Nucleus | other |
| Neural Tube | Cfap43 | cilia and fl | Other | other |
| Neural Tube | Cdh24 | cadherin 2 | Plasma Me | other |
| Neural Tube | Col18a1 | collagen ty | Extracellul | other |
| Neural Tube | Msh2 | mutS hom | Nucleus | enzyme |
| Neural Tube | Glg1 | golgi glyco | Cytoplasm | other |
| Neural Tube | Wdr91 | WD repeat | Cytoplasm | other |
| Neural Tube | Slc4a7 | solute carr | Plasma Me | transporter |
| Neural Tube | Mipep | mitochonc | Cytoplasm | peptidase |
| Neural Tube | Ift140 | intraflagell | Extracellul | other |
| Neural Tube | Nedd4 | neural pre | Cytoplasm | enzyme |
| Neural Tube | U2af2 | U2 small n | Nucleus | other |
| Neural Tube | Tmem109 | transmeml | Cytoplasm | other |
| Neural Tube | Nup188 | nucleopor | Nucleus | other |
| Neural Tube | Rcbtb1 | RCC1 and | Nucleus | other |
| Neural Tube | Map4k1 | mitogen-a | Cytoplasm | kinase |
| Neural Tube | Hyou1 | hypoxia up | Cytoplasm | other |
| Neural Tube | Secisbp2 | SECIS binc | Cytoplasm | translation regulator |
| Neural Tube | Eri3 | ERI1 exorib | Plasma Me | other |
| Neural Tube | Strip1 | striatin inte | Nucleus | other |
| Neural Tube | Med12 | mediator c | Nucleus | transcription regulato |
| Neural Tube | Rara | retinoic ac | Nucleus | ligand-dependent nu |
| Neural Tube | Esyt1 | extended s | Cytoplasm | other |
| Neural Tube | Txlna | taxilin alph | Extracellul | cytokine |
| Neural Tube | Ppap2b | - | - | - |
| Neural Tube | Fuca1 | alpha-L-fu | Cytoplasm | enzyme |
| Neural Tube | Srprb | SRP recept | Cytoplasm | other |
| Neural Tube | Smc1a | structural | Nucleus | transporter |
| Neural Tube | Khsrp | KH-type sp | Nucleus | enzyme |
| Neural Tube | Asphd2 | aspartate b | Other | enzyme |
| Neural Tube | Hmgcr | 3-hydroxy | Cytoplasm | enzyme |
| Neural Tube | Tmem231 | transmeml | Extracellul | other |
| Neural Tube | Slc39a6 | solute carr | Plasma Me | transporter |
| Neural Tube | Cbfb | core-bindi | Nucleus | transcription regulato |
| Neural Tube | Atp13a2 | ATPase 13 | Cytoplasm | transporter |
| Neural Tube | B4gat1 | beta-1,4-g | Cytoplasm | enzyme |
| Neural Tube | Chaf1b | chromatin | Nucleus | other |
| Neural Tube | Sart3 | squamous | Nucleus | other |
| Neural Tube | Srd5a1 | steroid 5 a | Cytoplasm | enzyme |
| Neural Tube | Rprd1b | regulation | Nucleus | other |
| Neural Tube | Prosc | - | - | - |
| Neural Tube | Fgfrl1 | fibroblast g | Plasma Me | transmembrane recep |
| Neural Tube | Arih2 | ariadne RB | Nucleus | enzyme |
| Neural Tube | Wscd1 | WSC doma | Other | other |
| Neural Tube | Crabp2 | cellular ret | Cytoplasm | transporter |
| Neural Tube | Slc6a6 | solute carr | Plasma Me | transporter |
| Neural Tube | Tmem18 | transmeml | Nucleus | other |
| Neural Tube | Mettl7a3 | methyltrar | Other | other |
| Neural Tube | Alkbh3 | alkB homo | Nucleus | enzyme |
| Neural Tube | Fxr1 | FMR1 auto | Cytoplasm | other |

| | | | | |
|---|---|---|---|---|
| Neural Tube | Sapcd2 | suppresso | Nucleus | other |
| Neural Tube | Tspan3 | tetraspanii | Plasma Me | other |
| Neural Tube | Paxbp1 | PAX3 and | Nucleus | other |
| Neural Tube | Mns1 | meiosis sp | Cytoplasm | other |
| Neural Tube | Atp1a1 | ATPase Na | Plasma Me | transporter |
| Neural Tube | Sall2 | spalt like ti | Nucleus | transcription regulato |
| Neural Tube | Jade3 | jade family | Nucleus | other |
| Neural Tube | Nova1 | NOVA alte | Nucleus | other |
| Neural Tube | Slc10a3 | solute carr | Plasma Me | transporter |
| Neural Tube | Abhd17c | abhydrolas | Plasma Me | enzyme |
| Neural Tube | Pcdh8 | protocadh | Plasma Me | other |
| Neural Tube | Utp6 | UTP6, sma | Nucleus | other |
| Neural Tube | Adamts7 | ADAM me | Extracellul | peptidase |
| Neural Tube | Tril | TLR4 intera | Other | other |
| Neural Tube | Ddx51 | DEAD-box | Other | enzyme |
| Neural Tube | Fnbp4 | formin bin | Nucleus | other |
| Neural Tube | Cx3cl1 | C-X3-C m | Extracellul | cytokine |
| Neural Tube | Snx27 | sorting ne | Cytoplasm | other |
| Neural Tube | Ctnnb1 | catenin be | Nucleus | transcription regulato |
| Neural Tube | Arvcf | ARVCF, de | Plasma Me | other |
| Neural Tube | Slc1a5 | solute carr | Plasma Me | transporter |
| Neural Tube | Eif4g1 | eukaryotic | Cytoplasm | translation regulator |
| Neural Tube | Qk | QKI, KH do | Nucleus | other |
| Neural Tube | Nrm | nurim | Nucleus | other |
| Neural Tube | Rhpn1 | rhophilin F | Cytoplasm | other |
| Neural Tube | Dnajc4 | DnaJ heat | Cytoplasm | other |
| Neural Tube | Fam3a | family with | Extracellul | other |
| Neural Tube | Cyth2 | cytohesin : | Cytoplasm | other |
| Neural Tube | Terf2 | telomeric i | Nucleus | other |
| Neural Tube | Rbak | RB associa | Nucleus | transcription regulato |
| Neural Tube | Ykt6 | YKT6 v-SN | Cytoplasm | enzyme |
| Neural Tube | Nmral1 | NmrA like | Nucleus | other |
| Neural Tube | Dgcr8 | DGCR8, m | Nucleus | enzyme |
| Neural Tube | Rnf44 | ring finger | Other | other |
| Neural Tube | Ska2 | spindle an | Nucleus | other |
| Neural Tube | Cenpm | centromer | Cytoplasm | other |
| Neural Tube | Vamp2 | vesicle ass | Plasma Me | other |
| Neural Tube | Nedd9 | neural pre | Nucleus | other |
| Neural Tube | Lias | lipoic acid | Cytoplasm | enzyme |
| Neural Tube | Bcs1l | BCS1 hom | Cytoplasm | other |
| Neural Tube | Naglu | N-acetyl-a | Cytoplasm | enzyme |
| Neural Tube | Fuk | fucokinase | Cytoplasm | kinase |
| Neural Tube | Gpc6 | glypican 6 | Plasma Me | transmembrane recep |
| Neural Tube | Crip2 | cysteine ri | Plasma Me | other |
| Neural Tube | Zfp202 | zinc finger | Nucleus | transcription regulato |
| Neural Tube | Trip6 | thyroid ho | Extracellul | cytokine |
| Neural Tube | Mest | mesoderm | Cytoplasm | peptidase |
| Neural Tube | Minpp1 | multiple in | Cytoplasm | phosphatase |
| Neural Tube | Hnrnph1 | heterogen | Nucleus | other |
| Neural Tube | Insr | insulin rec | Plasma Me | kinase |
| Neural Tube | Ubp1 | upstream I | Cytoplasm | transcription regulato |
| Neural Tube | Top2a | DNA topo | Nucleus | enzyme |
| Neural Tube | Mcm4 | minichrom | Nucleus | enzyme |
| Neural Tube | Fhl1 | four and a | Cytoplasm | other |
| Neural Tube | Dlst | dihydrolip | Cytoplasm | enzyme |
| Neural Tube | Scd1 | stearoyl-C | Cytoplasm | enzyme |
| Neural Tube | Bicd2 | BICD carg | Cytoplasm | other |
| Neural Tube | Gid4 | GID compl | Nucleus | other |

| | | | | |
|---|---|---|---|---|
| Neural Tube | Lig1 | DNA ligase | Nucleus | enzyme |
| Neural Tube | Zfp710 | zinc finger | Nucleus | other |
| Neural Tube | Orc1 | origin reco | Nucleus | other |
| Neural Tube | Ssbp3 | single stra | Nucleus | transcription regulato |
| Neural Tube | Psap | prosaposin | Extracellul | enzyme |
| Neural Tube | Pacsin3 | protein kin | Cytoplasm | other |
| Neural Tube | Sort1 | sortilin 1 | Plasma Me | G-protein coupled re |
| Neural Tube | Cpsf1 | cleavage a | Nucleus | other |
| Neural Tube | Faf2 | Fas associa | Cytoplasm | other |
| Neural Tube | Ubap2 | ubiquitin a | Cytoplasm | other |
| Neural Tube | Zkscan3 | zinc finger | Nucleus | transcription regulato |
| Neural Tube | Foxm1 | forkhead b | Nucleus | transcription regulato |
| Neural Tube | Tmem168 | transmemb | Cytoplasm | other |
| Neural Tube | Chek1 | checkpoint | Nucleus | kinase |
| Neural Tube | Porcn | porcupine | Cytoplasm | enzyme |
| Neural Tube | Pygo2 | pygopus fa | Nucleus | other |
| Neural Tube | Ganab | glucosidas | Cytoplasm | enzyme |
| Neural Tube | Rabepk | Rab9 effec | Cytoplasm | other |
| Neural Tube | Dynlt1c | dynein ligh | Cytoplasm | other |
| Neural Tube | Nol8 | nucleolar p | Nucleus | other |
| Neural Tube | Rfng | RFNG O-fu | Cytoplasm | enzyme |
| Neural Tube | Epn2 | epsin 2 | Cytoplasm | other |
| Neural Tube | Vopp1 | VOPP1, W | Nucleus | other |
| Neural Tube | Prpf4b | pre-mRNA | Nucleus | kinase |
| Neural Tube | Elovl5 | ELOVL fatt | Cytoplasm | enzyme |
| Neural Tube | Tpcn2 | two pore s | Cytoplasm | ion channel |
| Neural Tube | Rmdn3 | regulator o | Cytoplasm | other |
| Neural Tube | Zfp266 | zinc finger | Nucleus | other |
| Neural Tube | Mta2 | metastasis | Nucleus | transcription regulato |
| Neural Tube | Mki67 | marker of | Nucleus | other |
| Neural Tube | Fads1 | fatty acid d | Plasma Me | enzyme |
| Neural Tube | Apbb1 | amyloid be | Cytoplasm | transcription regulato |
| Neural Tube | Nelfa | negative e | Nucleus | other |
| Neural Tube | Gabpb1 | GA binding | Nucleus | transcription regulato |
| Neural Tube | Gns | glucosami | Cytoplasm | enzyme |
| Neural Tube | Arhgef1 | Rho guani | Cytoplasm | other |
| Neural Tube | Spop | speckle typ | Extracellul | other |
| Neural Tube | Fam13b | family with | Cytoplasm | other |
| Neural Tube | Herc4 | HECT and | Cytoplasm | enzyme |
| Neural Tube | Ints9 | integrator | Nucleus | other |
| Neural Tube | Timm22 | translocase | Cytoplasm | transporter |
| Neural Tube | Zfand3 | zinc finger | Other | other |
| Neural Tube | Carm1 | coactivato | Nucleus | transcription regulato |
| Neural Tube | Ppp2r4 | - | - | - |
| Neural Tube | Tm9sf4 | transmemb | Cytoplasm | transporter |
| Neural Tube | Zmat5 | zinc finger | Nucleus | other |
| Neural Tube | Nptn | neuroplast | Plasma Me | other |
| Neural Tube | Zfp821 | zinc finger | Other | other |
| Neural Tube | Ptk7 | protein tyr | Plasma Me | kinase |
| Neural Tube | Shd | Src homolo | Cytoplasm | other |
| Neural Tube | Pigl | phosphatid | Cytoplasm | enzyme |
| Neural Tube | 2810408M | - | - | - |
| Neural Tube | Prep | prolyl endo | Cytoplasm | peptidase |
| Neural Tube | Nufip1 | NUFIP1, FM | Nucleus | other |
| Neural Tube | Dhx38 | DEAH-box | Nucleus | enzyme |
| Neural Tube | Zmynd11 | zinc finger | Nucleus | transcription regulato |
| Neural Tube | Cant1 | calcium ac | Extracellul | enzyme |
| Neural Tube | Ap1b1 | adaptor re | Cytoplasm | transporter |

| | | | | |
|---|---|---|---|---|
| Neural Tube | Plekha5 | pleckstrin | Cytoplasm | other |
| Neural Tube | Ddx17 | DEAD-box | Nucleus | enzyme |
| Neural Tube | R3hdm4 | R3H doma | Other | other |
| Neural Tube | Cbx5 | chromobo | Nucleus | transcription regulato |
| Neural Tube | Rere | arginine-g | Nucleus | transcription regulato |
| Neural Tube | Ptpn13 | protein tyr | Cytoplasm | phosphatase |
| Neural Tube | Prps1 | phosphori | Cytoplasm | kinase |
| Neural Tube | Pald1 | phosphata | Cytoplasm | phosphatase |
| Neural Tube | Smad5 | SMAD fam | Nucleus | transcription regulato |
| Neural Tube | Mau2 | MAU2 sist | Nucleus | other |
| Neural Tube | Rbm12b2 | RNA bindi | Other | other |
| Neural Tube | Ing4 | inhibitor o | Nucleus | transcription regulato |
| Neural Tube | Kit | KIT proto- | Plasma Me | transmembrane recep |
| Neural Tube | Ighmbp2 | immunogl | Nucleus | enzyme |
| Neural Tube | 1110057K( | - | - | - |
| Neural Tube | Cdc37l1 | cell divisio | Cytoplasm | other |
| Neural Tube | Zkscan17 | zinc finger | Nucleus | transcription regulato |
| Neural Tube | Troap | trophinin a | Cytoplasm | peptidase |
| Neural Tube | Rad54l | RAD54 like | Nucleus | enzyme |
| Neural Tube | Spns1 | sphingolip | Cytoplasm | transporter |
| Neural Tube | Cxxc1 | CXXC fing | Nucleus | transcription regulato |
| Neural Tube | Mpdu1 | mannose- | Cytoplasm | other |
| Neural Tube | Srsf1 | serine and | Nucleus | other |
| Neural Tube | D930015E( | - | - | - |
| Neural Tube | Ror2 | receptor ty | Plasma Me | kinase |
| Neural Tube | Ddx6 | DEAD-box | Nucleus | enzyme |
| Neural Tube | Rgs12 | regulator ( | Nucleus | other |
| Neural Tube | Bms1 | BMS1, ribo | Nucleus | other |
| Neural Tube | Ncstn | nicastrin | Plasma Me | peptidase |
| Neural Tube | Tmem127 | transmeml | Other | other |
| Neural Tube | Itfg1 | integrin al | Plasma Me | other |
| Neural Tube | Sun1 | Sad1 and U | Nucleus | other |
| Neural Tube | Hells | helicase, ly | Nucleus | enzyme |
| Neural Tube | C730034F( | RIKEN cDN | Other | other |
| Neural Tube | Trp53i11 | tumor pro | Other | other |
| Neural Tube | Wdr6 | WD repeat | Cytoplasm | other |
| Neural Tube | Xrcc1 | X-ray repa | Nucleus | enzyme |
| Neural Tube | Rad51 | RAD51 rec | Nucleus | enzyme |
| Neural Tube | Zfp503 | zinc finger | Nucleus | other |
| Neural Tube | Uck1 | uridine-cy | Cytoplasm | kinase |
| Neural Tube | Ankrd54 | ankyrin rep | Nucleus | transcription regulato |
| Neural Tube | Ncapg2 | non-SMC | Nucleus | other |
| Neural Tube | Mapk7 | mitogen-a | Cytoplasm | kinase |
| Neural Tube | Atf6b | activating | Nucleus | transcription regulato |
| Neural Tube | Rtn3 | reticulon 3 | Cytoplasm | other |
| Neural Tube | Wdr77 | WD repeat | Nucleus | transcription regulato |
| Neural Tube | Inppl1 | inositol po | Cytoplasm | phosphatase |
| Neural Tube | Ikbkap | elongator | Cytoplasm | other |
| Neural Tube | Atxn2l | ataxin 2 lik | Nucleus | other |
| Neural Tube | Vars | valyl-tRNA | Cytoplasm | enzyme |
| Neural Tube | Pard6g | par-6 fam | Plasma Me | other |
| Neural Tube | Kif4 | kinesin fan | Nucleus | other |
| Neural Tube | B3gnt5 | UDP-GlcN | Cytoplasm | enzyme |
| Neural Tube | Hsph1 | heat shock | Cytoplasm | other |
| Neural Tube | Ptcd1 | ATP5MF-F | Cytoplasm | other |
| Neural Tube | Trappc6b | trafficking | Cytoplasm | other |
| Neural Tube | Ccnj | cyclin J | Nucleus | other |
| Neural Tube | Ddx49 | DEAD-box | Nucleus | enzyme |

| | | | | |
|---|---|---|---|---|
| Neural Tube | Msh6 | mutS hom | Nucleus | enzyme |
| Neural Tube | Cyp51 | cytochrom | Cytoplasm | enzyme |
| Neural Tube | Blm | Bloom syn | Nucleus | enzyme |
| Neural Tube | Pik3r3 | phosphoir | Cytoplasm | kinase |
| Neural Tube | Ist1 | IST1, ESCR | Cytoplasm | other |
| Neural Tube | Slc38a10 | solute carr | Cytoplasm | other |
| Neural Tube | Nt5dc2 | 5'-nucleot | Cytoplasm | other |
| Neural Tube | Gtf2i | general tra | Nucleus | transcription regulato |
| Neural Tube | Uba1 | ubiquitin li | Cytoplasm | enzyme |
| Neural Tube | Phldb1 | pleckstrin | Cytoplasm | other |
| Neural Tube | Hcfc1 | host cell fa | Nucleus | transcription regulato |
| Neural Tube | Nhlrc2 | NHL repea | Other | enzyme |
| Neural Tube | Zfp866 | zinc finger | Other | other |
| Neural Tube | Prss23 | protease, s | Extracellula | peptidase |
| Neural Tube | Psip1 | PC4 and S | Nucleus | transcription regulato |
| Neural Tube | Armc5 | armadillo r | Cytoplasm | other |
| Neural Tube | Zfp719 | zinc finger | Other | other |
| Neural Tube | Ccdc91 | coiled-coi | Cytoplasm | other |
| Neural Tube | Sh3bgrl | SH3 doma | Cytoplasm | other |
| Neural Tube | 2610318N | RIKEN cDN | Other | other |
| Neural Tube | Atp1b3 | ATPase Na | Plasma Me | transporter |
| Neural Tube | Nup210 | nucleopor | Nucleus | transporter |
| Neural Tube | Sdad1 | SDA1 dom | Nucleus | other |
| Neural Tube | Pef1 | penta-EF- | Cytoplasm | other |
| Neural Tube | Tmem39b | transmemb | Cytoplasm | other |
| Neural Tube | Drosha | drosha rib | Nucleus | enzyme |
| Neural Tube | Pigq | phosphatid | Cytoplasm | enzyme |
| Neural Tube | Pcsk7 | proprotein | Cytoplasm | peptidase |
| Neural Tube | Wsb1 | WD repeat | Cytoplasm | enzyme |
| Neural Tube | Luzp1 | leucine zip | Nucleus | other |
| Neural Tube | Dtnbp1 | dystrobrev | Plasma Me | other |
| Neural Tube | Tmub2 | transmemb | Other | other |
| Neural Tube | Phactr4 | phosphata | Plasma Me | other |
| Neural Tube | Nkiras2 | NFKB inhib | Cytoplasm | enzyme |
| Neural Tube | Nedd1 | neural pre | Cytoplasm | other |
| Neural Tube | Uhrf1 | ubiquitin li | Nucleus | transcription regulato |
| Neural Tube | Aagab | alpha and | Cytoplasm | other |
| Neural Tube | Sgta | small gluta | Cytoplasm | other |
| Neural Tube | Uqcc1 | ubiquinol- | Cytoplasm | other |
| Neural Tube | Chmp6 | charged m | Cytoplasm | other |
| Neural Tube | Sel1l | SEL1L ERA | Cytoplasm | other |
| Neural Tube | Dalrd3 | DALR antic | Other | other |
| Neural Tube | Cherp | calcium hc | Cytoplasm | other |
| Neural Tube | Adam17 | ADAM me | Plasma Me | peptidase |
| Neural Tube | Tfip11 | tuftelin int | Extracellula | other |
| Neural Tube | Abhd12 | abhydrolas | Plasma Me | enzyme |
| Neural Tube | Aldh18a1 | aldehyde c | Cytoplasm | kinase |
| Neural Tube | Anks3 | ankyrin rep | Nucleus | transcription regulato |
| Neural Tube | Ctr9 | CTR9 hom | Nucleus | other |
| Neural Tube | Dck | deoxycytic | Nucleus | kinase |
| Neural Tube | Ctns | cystinosin, | Cytoplasm | transporter |
| Neural Tube | Bbs2 | Bardet-Bie | Cytoplasm | other |
| Neural Tube | Srrm2 | serine/arg | Nucleus | other |
| Neural Tube | Emc10 | ER membr | Cytoplasm | other |
| Neural Tube | Man2b1 | mannosida | Cytoplasm | enzyme |
| Neural Tube | Pls3 | plastin 3 | Cytoplasm | other |
| Neural Tube | Hmg20a | high mobi | Nucleus | transcription regulato |
| Neural Tube | Dhx29 | DExH-box | Cytoplasm | enzyme |

| | | | | |
|---|---|---|---|---|
| Neural Tube | 1110059E2 | chromosoı | Other | other |
| Neural Tube | Rce1 | Ras conveı | Cytoplasm | peptidase |
| Neural Tube | Wars | tryptophaı | Cytoplasm | enzyme |
| Neural Tube | Yipf3 | Yip1 doma | Cytoplasm | other |
| Neural Tube | Sdc1 | syndecan : | Plasma Me | enzyme |
| Neural Tube | Tcf19 | transcripti | Nucleus | transcription regulato |
| Neural Tube | Hirip3 | HIRA inter | Nucleus | other |
| Neural Tube | Eif2s3x | eukaryotic | Cytoplasm | translation regulator |
| Neural Tube | Ddx20 | DEAD-box | Nucleus | transcription regulato |
| Neural Tube | Pcdh18 | protocadh | Extracellul | other |
| Neural Tube | Srsf9 | serine and | Nucleus | enzyme |
| Neural Tube | Wee1 | WEE1 G2 ( | Nucleus | kinase |
| Neural Tube | B4galnt4 | beta-1,4-N | Extracellul | enzyme |
| Neural Tube | Smpdl3a | sphingomy | Extracellul | enzyme |
| Neural Tube | 2610301B2 | chromosoı | Cytoplasm | other |
| Neural Tube | Wbp1 | WW doma | Cytoplasm | other |
| Neural Tube | Zfp811 | zinc finger | Other | other |
| Neural Tube | Ciao1 | cytosolic iı | Nucleus | transcription regulato |
| Neural Tube | Get4 | golgi to EF | Cytoplasm | other |
| Neural Tube | Rad51ap1 | RAD51 ass | Nucleus | other |
| Neural Tube | Ackr3 | atypical ch | Plasma Me | G-protein coupled re |
| Neural Tube | Adprh | ADP-ribos | Other | enzyme |
| Neural Tube | Trim32 | tripartite n | Nucleus | transcription regulato |
| Neural Tube | Mpv17l2 | MPV17 mi | Cytoplasm | other |
| Neural Tube | Tmem234 | transmeml | Other | other |
| Neural Tube | Zfp830 | zinc finger | Nucleus | other |
| Neural Tube | Trafd1 | TRAF-type | Other | other |
| Neural Tube | 4930452B0 | chromosoı | Other | other |
| Neural Tube | Hspa4l | heat shock | Cytoplasm | other |
| Neural Tube | Calu | calumenin | Cytoplasm | other |
| Neural Tube | Tcea1 | transcripti | Nucleus | transcription regulato |
| Neural Tube | Zmynd8 | zinc finger | Nucleus | transcription regulato |
| Neural Tube | Vamp3 | vesicle ass | Plasma Me | other |
| Neural Tube | Tmem43 | transmeml | Nucleus | other |
| Neural Tube | Tbpl1 | TATA-box | Nucleus | transcription regulato |
| Neural Tube | Ddx24 | DEAD-box | Nucleus | enzyme |
| Neural Tube | Taf5l | TATA-box | Nucleus | transcription regulato |
| Neural Tube | Azin1 | antizyme i | Cytoplasm | enzyme |
| Neural Tube | Srebf2 | sterol regu | Nucleus | transcription regulato |
| Neural Tube | Lppr3 | - | - | - |
| Neural Tube | Smarca4 | SWI/SNF r | Nucleus | transcription regulato |
| Neural Tube | Wls | wntless Wı | Cytoplasm | other |
| Neural Tube | Rbm14 | RNA bindi | Nucleus | transcription regulato |
| Neural Tube | Nfe2l1 | nuclear fa | Nucleus | transcription regulato |
| Neural Tube | Sh3yl1 | SH3 and S | Plasma Me | other |
| Neural Tube | Add3 | adducin 3 | Cytoplasm | other |
| Neural Tube | Glt8d1 | glycosyltra | Cytoplasm | enzyme |
| Neural Tube | Alg8 | ALG8, alph | Cytoplasm | enzyme |
| Neural Tube | Slc25a30 | solute carr | Cytoplasm | other |
| Neural Tube | Cyb5r3 | cytochrom | Cytoplasm | enzyme |
| Neural Tube | Foxa2 | forkhead b | Nucleus | transcription regulato |
| Neural Tube | Fam83d | family with | Cytoplasm | other |
| Neural Tube | Lrrc8a | leucine ric | Plasma Me | ion channel |
| Neural Tube | Tmem222 | transmeml | Other | other |
| Neural Tube | G3bp1 | G3BP stres | Nucleus | enzyme |
| Neural Tube | Entpd6 | ectonucle | Cytoplasm | enzyme |
| Neural Tube | Ube3b | ubiquitin p | Extracellul | enzyme |
| Neural Tube | Ppil2 | peptidylpr | Nucleus | enzyme |

| | | | | |
|---|---|---|---|---|
| Neural Tube | Uchl1 | ubiquitin C | Cytoplasm | peptidase |
| Neural Tube | P3h4 | prolyl 3-hy | Nucleus | other |
| Neural Tube | Ogfr | opioid gro | Plasma Me | other |
| Neural Tube | Mfsd5 | major facil | Other | transporter |
| Neural Tube | Nol6 | nucleolar p | Nucleus | other |
| Neural Tube | Polr2b | RNA polyr | Nucleus | enzyme |
| Neural Tube | Cklf | chemokine | Extracellul | cytokine |
| Neural Tube | Mapk14 | mitogen-a | Cytoplasm | kinase |
| Neural Tube | Pdhx | pyruvate c | Cytoplasm | enzyme |
| Neural Tube | Aco2 | aconitase 2 | Cytoplasm | enzyme |
| Neural Tube | Vgll4 | vestigial lik | Nucleus | other |
| Neural Tube | Hdhd2 | haloacid d | Cytoplasm | other |
| Neural Tube | Usp39 | ubiquitin s | Nucleus | peptidase |
| Neural Tube | Cast | calpastatin | Cytoplasm | peptidase |
| Neural Tube | Cog4 | componen | Cytoplasm | transporter |
| Neural Tube | Rcn2 | reticulocal | Cytoplasm | other |
| Neural Tube | Farsa | phenylalar | Cytoplasm | enzyme |
| Neural Tube | Igf1r | insulin like | Plasma Me | transmembrane recep |
| Neural Tube | E2f5 | E2F transc | Nucleus | transcription regulato |
| Neural Tube | Tox4 | TOX high m | Nucleus | other |
| Neural Tube | Fars2 | phenylalar | Cytoplasm | enzyme |
| Neural Tube | Sirt3 | sirtuin 3 | Cytoplasm | enzyme |
| Neural Tube | Snf8 | SNF8, ESC | Cytoplasm | enzyme |
| Neural Tube | Agpat6 | - | - | - |
| Neural Tube | Armcx1 | armadillo r | Other | other |
| Neural Tube | Mtmr4 | myotubula | Cytoplasm | phosphatase |
| Neural Tube | Btd | biotinidase | Extracellul | enzyme |
| Neural Tube | Tmem132a | transmeml | Cytoplasm | other |
| Neural Tube | Sf1 | splicing fa | Nucleus | other |
| Neural Tube | Ipp | intracistern | Cytoplasm | other |
| Neural Tube | Hadha | hydroxyac | Cytoplasm | enzyme |
| Neural Tube | Zfp212 | zinc finger | Nucleus | other |
| Neural Tube | Psme3 | proteasom | Cytoplasm | peptidase |
| Neural Tube | Slc37a4 | solute carr | Cytoplasm | transporter |
| Neural Tube | Kiz | kizuna cen | Cytoplasm | other |
| Neural Tube | Ogt | O-linked N | Cytoplasm | enzyme |
| Neural Tube | Ash2l | ASH2 like | Nucleus | transcription regulato |
| Neural Tube | Trip10 | thyroid ho | Cytoplasm | other |
| Neural Tube | Usp21 | ubiquitin s | Cytoplasm | peptidase |
| Neural Tube | Pxdn | peroxidasi | Extracellul | enzyme |
| Neural Tube | Ivns1abp | influenza v | Nucleus | other |
| Neural Tube | Rbm38 | RNA bindi | Nucleus | other |
| Neural Tube | 9530068E0 | chromosoı | Other | other |
| Neural Tube | Bag5 | BCL2 asso | Cytoplasm | other |
| Neural Tube | Fubp1 | far upstrea | Nucleus | transcription regulato |
| Neural Tube | Dpysl2 | dihydropy | Cytoplasm | enzyme |
| Neural Tube | Mia3 | MIA family | Cytoplasm | transporter |
| Neural Tube | Supt5 | SPT5 hom | Nucleus | transcription regulato |
| Neural Tube | Dmap1 | DNA meth | Nucleus | transcription regulato |
| Neural Tube | Ptprf | protein tyr | Plasma Me | phosphatase |
| Neural Tube | Exo5 | exonuclea: | Nucleus | enzyme |
| Neural Tube | Ppwd1 | peptidylpr | Nucleus | enzyme |
| Neural Tube | Hip1 | huntingtin | Cytoplasm | other |
| Neural Tube | Slc29a4 | solute carr | Plasma Me | transporter |
| Neural Tube | Ccdc117 | coiled-coi | Other | other |
| Neural Tube | Isca2 | iron-sulfur | Cytoplasm | other |
| Neural Tube | Mcm2 | minichrom | Nucleus | enzyme |
| Neural Tube | Cpsf4 | cleavage a | Nucleus | other |

| | | | | |
|---|---|---|---|---|
| Neural Tube | Slc44a2 | solute carr | Extracellula | transporter |
| Neural Tube | Cs | citrate syn | Cytoplasm | enzyme |
| Neural Tube | Dcun1d4 | defective i | Nucleus | other |
| Neural Tube | Myg1 | chromosor | Nucleus | other |
| Neural Tube | Atp6ap1 | ATPase H+ | Cytoplasm | transporter |
| Neural Tube | Prdx4 | peroxiredc | Cytoplasm | enzyme |
| Neural Tube | Ddx56 | DEAD-box | Nucleus | enzyme |
| Neural Tube | Med22 | mediator c | Cytoplasm | other |
| Neural Tube | Sgpl1 | sphingosir | Cytoplasm | enzyme |
| Neural Tube | Serpinh1 | serpin fam | Extracellula | other |
| Neural Tube | Prkab1 | protein kir | Nucleus | kinase |
| Neural Tube | Csnk1a1 | casein kina | Cytoplasm | kinase |
| Neural Tube | Sra1 | steroid rec | Nucleus | transcription regulato |
| Neural Tube | Ctcf | CCCTC-bir | Nucleus | transcription regulato |
| Neural Tube | C1galt1c1 | C1GALT1 s | Cytoplasm | other |
| Neural Tube | Tubgcp2 | tubulin ga | Cytoplasm | peptidase |
| Neural Tube | Gtf2a1 | general tra | Cytoplasm | transcription regulato |
| Neural Tube | Hjurp | Holliday ju | Nucleus | other |
| Neural Tube | Thrap3 | thyroid ho | Nucleus | transcription regulato |
| Neural Tube | Mcrs1 | microsphe | Nucleus | other |
| Neural Tube | Ethe1 | ETHE1, pe | Cytoplasm | enzyme |
| Neural Tube | Morc2a | MORC fam | Nucleus | other |
| Neural Tube | St5 | suppressic | Cytoplasm | enzyme |
| Neural Tube | Cpsf3l | - | - | - |
| Neural Tube | Tor1a | torsin fami | Cytoplasm | enzyme |
| Neural Tube | Cyb5b | cytochrom | Cytoplasm | enzyme |
| Neural Tube | Sil1 | SIL1 nucle | Cytoplasm | transporter |
| Neural Tube | Ogdhl | oxoglutara | Other | enzyme |
| Neural Tube | Ampd2 | adenosine | Cytoplasm | enzyme |
| Neural Tube | Fam109a | family with | Cytoplasm | other |
| Neural Tube | Rbbp8 | RB binding | Nucleus | enzyme |
| Neural Tube | Arhgap28 | Rho GTPas | Cytoplasm | other |
| Neural Tube | Zfp397 | zinc finger | Nucleus | transcription regulato |
| Neural Tube | Mgme1 | mitochonc | Cytoplasm | enzyme |
| Neural Tube | Plcg1 | phospholip | Cytoplasm | enzyme |
| Neural Tube | Flna | filamin A | Cytoplasm | other |
| Neural Tube | Tmem159 | transmeml | Other | other |
| Neural Tube | Nucb2 | nucleobinc | Nucleus | other |
| Neural Tube | Angptl2 | angiopoie | Extracellula | other |
| Neural Tube | Dnmt1 | DNA meth | Nucleus | enzyme |
| Neural Tube | Unc119b | unc-119 li | Cytoplasm | other |
| Neural Tube | Mlec | malectin | Plasma Me | other |
| Neural Tube | Capn10 | calpain 10 | Cytoplasm | peptidase |
| Neural Tube | Sptlc1 | serine palr | Cytoplasm | enzyme |
| Neural Tube | Gpr180 | G protein- | Cytoplasm | other |
| Neural Tube | Cdk16 | cyclin dep | Cytoplasm | kinase |
| Neural Tube | Tusc2 | tumor sup | Nucleus | other |
| Neural Tube | Tcn2 | transcobal | Extracellula | transporter |
| Neural Tube | Ldlrad3 | low densit | Plasma Me | other |
| Neural Tube | Fads2 | fatty acid c | Plasma Me | enzyme |
| Neural Tube | Psmd8 | proteasom | Cytoplasm | other |
| Neural Tube | Zfp131 | zinc finger | Nucleus | transcription regulato |
| Neural Tube | Nnat | neuronatir | Plasma Me | transporter |
| Neural Tube | Slc29a3 | solute carr | Plasma Me | transporter |
| Neural Tube | Cuta | cutA divale | Cytoplasm | other |
| Neural Tube | Syngr2 | synaptogy | Cytoplasm | other |
| Neural Tube | Cdk20 | cyclin dep | Nucleus | kinase |
| Neural Tube | Rhno1 | RAD9-HUS | Nucleus | other |

| | | | | |
|---|---|---|---|---|
| Neural Tube | Nek4 | NIMA rela | Nucleus | kinase |
| Neural Tube | Chmp7 | charged m | Cytoplasm | transporter |
| Neural Tube | Sec61a1 | Sec61 tran | Cytoplasm | transporter |
| Neural Tube | Tpp1 | tripeptidyl | Cytoplasm | peptidase |
| Neural Tube | Derl1 | derlin 1 | Cytoplasm | other |
| Neural Tube | Chmp1a | charged m | Extracellula | peptidase |
| Neural Tube | Dph7 | diphthami | Cytoplasm | other |
| Neural Tube | Cdc16 | cell divisio | Nucleus | other |
| Neural Tube | Tsr2 | TSR2, ribo: | Other | other |
| Neural Tube | Actn4 | actinin alp | Cytoplasm | transcription regulato |
| Neural Tube | Arhgdia | Rho GDP c | Cytoplasm | other |
| Neural Tube | Kctd15 | potassium | Other | ion channel |
| Neural Tube | Rrn3 | RRN3 hom | Nucleus | transcription regulato |
| Neural Tube | Tusc3 | tumor sup | Plasma Me | transporter |
| Neural Tube | Ap1s1 | adaptor re | Cytoplasm | transporter |
| Neural Tube | Plxnb2 | plexin B2 | Plasma Me | transmembrane recep |
| Neural Tube | EGFP | - | - | - |
| Neural Tube | Laptm4b | lysosomal | Cytoplasm | other |
| Neural Tube | Vmp1 | vacuole m | Plasma Me | other |
| Neural Tube | Srpr | SRP recept | Cytoplasm | other |
| Neural Tube | Arhgef39 | Rho guani | Plasma Me | other |
| Neural Tube | Rbbp4 | RB binding | Nucleus | enzyme |
| Neural Tube | Pcmt1 | protein-L- | Cytoplasm | enzyme |
| Neural Tube | Nap1l4 | nucleosom | Cytoplasm | other |
| Neural Tube | Samm50 | SAMM50 s | Cytoplasm | other |
| Neural Tube | Tmem214 | transmeml | Extracellula | other |
| Neural Tube | Chaf1a | chromatin | Nucleus | other |
| Neural Tube | Alkbh2 | alkB homc | Nucleus | enzyme |
| Neural Tube | Sugp1 | SURP and | Nucleus | other |
| Neural Tube | Lpcat1 | lysophosp | Cytoplasm | enzyme |
| Neural Tube | Dnaaf2 | dynein axc | Cytoplasm | other |
| Neural Tube | Cfap20 | cilia and fla | Cytoplasm | other |
| Neural Tube | Gtf3c5 | general tra | Nucleus | transcription regulato |
| Neural Tube | Usp5 | ubiquitin s | Cytoplasm | peptidase |
| Neural Tube | Cenph | centromer | Nucleus | other |
| Neural Tube | Gdpd5 | glyceropho | Plasma Me | enzyme |
| Neural Tube | Stk3 | serine/thre | Cytoplasm | kinase |
| Neural Tube | Agpat1 | 1-acylglyc | Cytoplasm | enzyme |
| Neural Tube | Luc7l | LUC7 like | Nucleus | other |
| Neural Tube | Pinx1 | PIN2/TERF | Nucleus | other |
| Neural Tube | Nme6 | NME/NM2 | Cytoplasm | kinase |
| Neural Tube | Aurkb | aurora kina | Nucleus | kinase |
| Neural Tube | Gramd1a | GRAM dor | Extracellula | other |
| Neural Tube | Palm | paralemmi | Plasma Me | other |
| Neural Tube | Pigx | phosphatio | Cytoplasm | other |
| Neural Tube | Nemf | nuclear ex | Nucleus | other |
| Neural Tube | Zfp410 | zinc finger | Nucleus | other |
| Neural Tube | Ddx18 | DEAD-box | Nucleus | enzyme |
| Neural Tube | Map2k5 | mitogen-a | Cytoplasm | kinase |
| Neural Tube | Pigt | phosphatio | Cytoplasm | enzyme |
| Neural Tube | Tjap1 | tight junct | Plasma Me | other |
| Neural Tube | Id3 | inhibitor o | Nucleus | transcription regulato |
| Neural Tube | Ddb1 | damage sp | Nucleus | other |
| Neural Tube | Klhl28 | kelch like f | Other | other |
| Neural Tube | Puf60 | poly(U) bir | Nucleus | other |
| Neural Tube | Gnai2 | G protein s | Plasma Me | enzyme |
| Neural Tube | Cdca2 | cell divisio | Nucleus | other |
| Neural Tube | Tarbp2 | TARBP2, R | Nucleus | other |

| | | | | |
|---|---|---|---|---|
| Gut Tube | Actn2 | actinin alp | Nucleus | transcription regulato |
| Gut Tube | Osr1 | odd-skipp | Nucleus | kinase |
| Gut Tube | Mybpc3 | myosin bir | Cytoplasm | other |
| Gut Tube | Capn6 | calpain 6 | Cytoplasm | peptidase |
| Gut Tube | Myh7 | myosin he | Cytoplasm | enzyme |
| Gut Tube | Pitx2 | paired like | Nucleus | transcription regulato |
| Gut Tube | Krt7 | keratin 7 | Cytoplasm | other |
| Gut Tube | Myl3 | myosin lig | Cytoplasm | other |
| Gut Tube | Hoxc4 | homeobox | Nucleus | transcription regulato |
| Gut Tube | Myl2 | myosin lig | Cytoplasm | other |
| Gut Tube | Myh6 | myosin he | Cytoplasm | enzyme |
| Gut Tube | Bcam | basal cell a | Plasma Me | transmembrane recep |
| Gut Tube | Isx | intestine s | Nucleus | transcription regulato |
| Gut Tube | Gprc5a | G protein- | Plasma Me | G-protein coupled re |
| Gut Tube | Tpm1 | tropomyos | Plasma Me | other |
| Gut Tube | Slc2a3 | solute carr | Plasma Me | transporter |
| Gut Tube | Raet1e | retinoic ac | Plasma Me | other |
| Gut Tube | Smoc2 | SPARC rela | Extracellul | other |
| Gut Tube | Ankrd1 | ankyrin rep | Cytoplasm | transcription regulato |
| Gut Tube | Tmprss2 | transmeml | Plasma Me | peptidase |
| Gut Tube | Nkx2-6 | NK2 home | Nucleus | transcription regulato |
| Gut Tube | Rfx6 | regulatory | Nucleus | enzyme |
| Gut Tube | Ifitm1 | interferon | Other | other |
| Gut Tube | Myl4 | myosin lig | Cytoplasm | other |
| Gut Tube | Nepn | nephrocar | Extracellul | other |
| Gut Tube | Peg3 | paternally | Nucleus | kinase |
| Gut Tube | Rdh10 | retinol del | Nucleus | enzyme |
| Gut Tube | Vnn1 | vanin 1 | Plasma Me | enzyme |
| Gut Tube | Klf5 | Kruppel lik | Nucleus | transcription regulato |
| Gut Tube | Slc5a5 | solute carr | Plasma Me | transporter |
| Gut Tube | Arg1 | arginase 1 | Cytoplasm | enzyme |
| Gut Tube | Gata4 | GATA binc | Nucleus | transcription regulato |
| Gut Tube | Cyp2s1 | cytochrom | Cytoplasm | enzyme |
| Gut Tube | Car2 | carbonic a | Cytoplasm | enzyme |
| Gut Tube | Fam183b | family with | Extracellul | other |
| Gut Tube | Agrn | agrin | Plasma Me | other |
| Gut Tube | Gmpr | guanosine | Cytoplasm | enzyme |
| Gut Tube | Cdh1 | cadherin 1 | Plasma Me | other |
| Gut Tube | Ociad2 | OCIA dom | Cytoplasm | other |
| Gut Tube | App | amyloid be | Plasma Me | other |
| Gut Tube | Vcan | versican | Extracellul | other |
| Gut Tube | Epb4.1l3 | - | - | - |
| Gut Tube | Csrp3 | cysteine ar | Nucleus | other |
| Gut Tube | Igfbp5 | insulin like | Extracellul | other |
| Gut Tube | Tnnc1 | troponin C | Cytoplasm | other |
| Gut Tube | Dpp4 | dipeptidyl | Plasma Me | peptidase |
| Gut Tube | Cd24a | CD24a ant | Plasma Me | other |
| Gut Tube | Gpx3 | glutathion | Extracellul | enzyme |
| Gut Tube | Entpd3 | ectonucleo | Plasma Me | enzyme |
| Gut Tube | Itga3 | integrin su | Plasma Me | other |
| Gut Tube | Lrpap1 | LDL recept | Plasma Me | other |
| Gut Tube | Rarb | retinoic ac | Nucleus | ligand-dependent nu |
| Gut Tube | Sema6a | semaphori | Plasma Me | transmembrane recep |
| Gut Tube | St14 | suppressio | Plasma Me | peptidase |
| Gut Tube | Tspan8 | tetraspanin | Plasma Me | other |
| Gut Tube | Tdh | L-threonin | Cytoplasm | enzyme |
| Gut Tube | Prss8 | protease, s | Extracellul | peptidase |
| Gut Tube | Podxl | podocalyx | Plasma Me | other |

| | | | | |
|---|---|---|---|---|
| Gut Tube | Gja1 | gap junctio | Plasma Me | transporter |
| Gut Tube | Cldn8 | claudin 8 | Plasma Me | other |
| Gut Tube | Mreg | melanoreg | Cytoplasm | other |
| Gut Tube | Shh | sonic hedg | Extracellula | peptidase |
| Gut Tube | Cryab | crystallin a | Nucleus | other |
| Gut Tube | Igsf9 | immunogl | Plasma Me | other |
| Gut Tube | Tmem88 | transmeml | Plasma Me | other |
| Gut Tube | Slc39a10 | solute carr | Extracellula | transporter |
| Gut Tube | Hoxa1 | homeobox | Nucleus | transcription regulato |
| Gut Tube | Flna | filamin A | Cytoplasm | other |
| Gut Tube | Palm3 | paralemmi | Other | other |
| Gut Tube | Efna1 | ephrin A1 | Plasma Me | other |
| Gut Tube | Tff3 | trefoil facto | Extracellula | other |
| Gut Tube | Obsl1 | obscurin-l | Other | other |
| Gut Tube | Alpl | alkaline ph | Plasma Me | phosphatase |
| Gut Tube | Sema3c | semaphori | Extracellula | other |
| Gut Tube | Gprc5c | G protein- | Plasma Me | G-protein coupled re |
| Gut Tube | Cldn4 | claudin 4 | Plasma Me | transmembrane recep |
| Gut Tube | Folr1 | folate rece | Plasma Me | transporter |
| Gut Tube | Cldn6 | claudin 6 | Plasma Me | other |
| Gut Tube | Jup | junction pl | Plasma Me | other |
| Gut Tube | Col4a1 | collagen ty | Extracellula | other |
| Gut Tube | Acta2 | actin, alph | Cytoplasm | other |
| Gut Tube | Cthrc1 | collagen tr | Extracellula | other |
| Gut Tube | Foxa1 | forkhead b | Nucleus | transcription regulato |
| Gut Tube | Wfdc2 | WAP four- | Extracellula | other |
| Gut Tube | Smtnl2 | smoothelin | Other | other |
| Gut Tube | Lad1 | ladinin 1 | Extracellula | other |
| Gut Tube | Hmgn3 | high mobi | Nucleus | other |
| Gut Tube | Uchl1 | ubiquitin C | Cytoplasm | peptidase |
| Gut Tube | Ifi30 | IFI30, lysos | Cytoplasm | enzyme |
| Gut Tube | Wls | wntless W | Cytoplasm | other |
| Gut Tube | Anxa6 | annexin A6 | Plasma Me | ion channel |
| Gut Tube | Lamb1 | laminin su | Extracellula | other |
| Gut Tube | Mapre2 | microtubu | Cytoplasm | other |
| Gut Tube | Actc1 | actin, alph | Cytoplasm | enzyme |
| Gut Tube | Dag1 | dystroglyc | Plasma Me | transmembrane recep |
| Gut Tube | Ap1m2 | adaptor re | Cytoplasm | transporter |
| Gut Tube | Prps2 | phosphori | Cytoplasm | kinase |
| Gut Tube | Epcam | epithelial c | Plasma Me | other |
| Gut Tube | Myl7 | myosin lig | Cytoplasm | enzyme |
| Gut Tube | Rdm1 | RAD52 mo | Nucleus | other |
| Gut Tube | Gm21814 | predicted | Other | other |
| Gut Tube | Nrep | neuronal r | Cytoplasm | other |
| Gut Tube | Ramp3 | receptor a | Plasma Me | transporter |
| Gut Tube | Spint1 | serine pep | Extracellula | other |
| Gut Tube | Cdh6 | cadherin 6 | Plasma Me | other |
| Gut Tube | Dhrs3 | dehydroge | Cytoplasm | enzyme |
| Gut Tube | Col18a1 | collagen ty | Extracellula | other |
| Gut Tube | Hsd11b2 | hydroxyste | Cytoplasm | enzyme |
| Gut Tube | Lars2 | leucyl-tRN | Cytoplasm | enzyme |
| Gut Tube | Fxyd6 | FXYD dom | Plasma Me | ion channel |
| Gut Tube | Efs | embryona | Cytoplasm | other |
| Gut Tube | Kcnk1 | potassium | Plasma Me | ion channel |
| Gut Tube | Spint2 | serine pep | Extracellula | other |
| Gut Tube | Dab2 | DAB2, clat | Plasma Me | other |
| Gut Tube | Atp2c2 | ATPase sec | Cytoplasm | enzyme |
| Gut Tube | Krt19 | keratin 19 | Cytoplasm | other |

| | | | | |
|---|---|---|---|---|
| Gut Tube | Sfrp5 | secreted fr | Plasma Me | transmembrane recep |
| Gut Tube | Cdh3 | cadherin 3 | Plasma Me | other |
| Gut Tube | Fam129b | family with | Cytoplasm | transcription regulato |
| Gut Tube | Plac8 | placenta s | Nucleus | other |
| Gut Tube | Fam174b | family with | Other | other |
| Gut Tube | Lpar3 | lysophosp | Plasma Me | G-protein coupled re |
| Gut Tube | Tmsb4x | thymosin, | Cytoplasm | other |
| Gut Tube | S1pr2 | sphingosir | Plasma Me | G-protein coupled re |
| Gut Tube | Dbn1 | drebrin 1 | Cytoplasm | other |
| Gut Tube | Calcoco1 | calcium bi | Nucleus | transcription regulato |
| Gut Tube | Nudt10 | nudix hydr | Cytoplasm | phosphatase |
| Gut Tube | Pvrl2 | - | - | - |
| Gut Tube | 3632451O | chromosor | Extracellul | other |
| Gut Tube | Cnn3 | calponin 3 | Cytoplasm | other |
| Gut Tube | Amotl2 | angiomoti | Plasma Me | other |
| Gut Tube | Pttg1ip | PTTG1 inte | Nucleus | other |
| Gut Tube | Gm8113 | - | - | - |
| Gut Tube | F11r | F11 recept | Plasma Me | other |
| Gut Tube | Dsc2 | desmocoll | Plasma Me | other |
| Gut Tube | Raet1d | retinoic ac | Plasma Me | other |
| Gut Tube | Tpm4 | tropomyos | Cytoplasm | other |
| Gut Tube | Ces1d | carboxyles | Cytoplasm | enzyme |
| Gut Tube | Mmp2 | matrix met | Extracellul | peptidase |
| Gut Tube | Zfp956 | zinc finger | Other | other |
| Gut Tube | Tmem123 | transmeml | Plasma Me | other |
| Gut Tube | Cldn9 | claudin 9 | Plasma Me | other |
| Gut Tube | Fbln1 | fibulin 1 | Extracellul | other |
| Gut Tube | Rnf167 | ring finger | Cytoplasm | enzyme |
| Gut Tube | Nr2f2 | nuclear rec | Nucleus | ligand-dependent nu |
| Gut Tube | Nudt11 | nudix hydr | Cytoplasm | phosphatase |
| Gut Tube | Car4 | carbonic a | Plasma Me | enzyme |
| Gut Tube | Plod2 | procollage | Cytoplasm | enzyme |
| Gut Tube | Fscn1 | fascin actir | Cytoplasm | other |
| Gut Tube | Cnn2 | calponin 2 | Cytoplasm | other |
| Gut Tube | Ctsd | cathepsin | Cytoplasm | peptidase |
| Gut Tube | Ypel1 | yippee like | Nucleus | enzyme |
| Gut Tube | Cldn3 | claudin 3 | Plasma Me | transmembrane recep |
| Gut Tube | Shisa4 | shisa famil | Other | other |
| Gut Tube | Slc27a3 | solute carr | Cytoplasm | transporter |
| Gut Tube | Mmp11 | matrix met | Extracellul | peptidase |
| Gut Tube | Marcksl1 | MARCKS li | Cytoplasm | other |
| Gut Tube | Ddb1 | damage sp | Nucleus | other |
| Gut Tube | Gyg | glycogenir | Cytoplasm | enzyme |
| Gut Tube | Trp53i11 | tumor pro | Other | other |
| Gut Tube | Fstl1 | follistatin l | Extracellul | other |
| Gut Tube | Crb3 | crumbs fa | Plasma Me | other |
| Gut Tube | S100a10 | S100 calciu | Cytoplasm | other |
| Gut Tube | Col4a2 | collagen ty | Extracellul | other |
| Gut Tube | Slc2a1 | solute carr | Plasma Me | transporter |
| Gut Tube | Pxdn | peroxidasi | Extracellul | enzyme |
| Gut Tube | Kcnk16 | potassium | Plasma Me | ion channel |
| Gut Tube | Krt8 | keratin 8 | Cytoplasm | other |
| Gut Tube | Fblim1 | filamin bin | Plasma Me | other |
| Gut Tube | Ezr | ezrin | Plasma Me | other |
| Gut Tube | H2afy2 | H2A histor | Nucleus | other |
| Gut Tube | H3f3a | H3 histone | Nucleus | other |
| Gut Tube | Armc10 | armadillo r | Cytoplasm | other |
| Gut Tube | S100a11 | S100 calciu | Cytoplasm | other |

| | | | | |
|---|---|---|---|---|
| Gut Tube | Tspan3 | tetraspanir | Plasma Me | other |
| Gut Tube | Pla2g7 | phospholi | Extracellul | enzyme |
| Gut Tube | Nnat | neuronatir | Plasma Me | transporter |
| Gut Tube | Clic1 | chloride in | Nucleus | ion channel |
| Gut Tube | Fhl2 | four and a | Nucleus | transcription regulato |
| Gut Tube | Cldn7 | claudin 7 | Plasma Me | other |
| Gut Tube | Rcan2 | regulator c | Other | other |
| Gut Tube | Igfbp2 | insulin like | Extracellul | other |
| Gut Tube | Lasp1 | LIM and SI | Cytoplasm | transporter |
| Gut Tube | Wdr6 | WD repeat | Cytoplasm | other |
| Gut Tube | F2r | coagulatio | Plasma Me | G-protein coupled re |
| Gut Tube | Mapk13 | mitogen-a | Cytoplasm | kinase |
| Gut Tube | Ddah2 | dimethylar | Cytoplasm | enzyme |
| Gut Tube | Cd320 | CD320 mc | Plasma Me | transmembrane recep |
| Gut Tube | Ccnd2 | cyclin D2 | Nucleus | other |
| Gut Tube | Rnf145 | ring finger | Other | other |
| Gut Tube | Stmn1 | stathmin 1 | Cytoplasm | other |
| Gut Tube | Upp1 | uridine ph | Cytoplasm | enzyme |
| Gut Tube | Rasl11b | RAS like fa | Plasma Me | enzyme |
| Gut Tube | Maged1 | MAGE fam | Plasma Me | transcription regulato |
| Gut Tube | Vps72 | vacuolar p | Nucleus | transcription regulato |
| Gut Tube | Vash2 | vasohibin | Cytoplasm | other |
| Gut Tube | Pak1 | p21 (RAC1 | Cytoplasm | kinase |
| Gut Tube | Ptp4a1 | protein tyr | Cytoplasm | phosphatase |
| Gut Tube | Limk2 | LIM doma | Cytoplasm | kinase |
| Gut Tube | Stap2 | signal tran | Cytoplasm | other |
| Gut Tube | Vps36 | vacuolar p | Cytoplasm | other |
| Gut Tube | Arrdc1 | arrestin dc | Cytoplasm | other |
| Gut Tube | Tmem98 | transmeml | Cytoplasm | other |
| Gut Tube | Crip2 | cysteine ri | Plasma Me | other |
| Gut Tube | Sh3bp4 | SH3 doma | Cytoplasm | other |
| Gut Tube | Sh3bp1 | SH3 doma | Plasma Me | other |
| Gut Tube | Rcn1 | reticulocal | Cytoplasm | other |
| Gut Tube | Ajuba | ajuba LIM | Nucleus | transcription regulato |
| Gut Tube | Laptm4b | lysosomal | Cytoplasm | other |
| Gut Tube | Fzd7 | frizzled cla | Plasma Me | G-protein coupled re |
| Gut Tube | Slc16a13 | solute carr | Cytoplasm | other |
| Gut Tube | Dpysl5 | dihydropy | Cytoplasm | enzyme |
| Gut Tube | Pkm | pyruvate k | Cytoplasm | kinase |
| Gut Tube | Syt5 | synaptota | Cytoplasm | transporter |
| Gut Tube | Psca | prostate st | Plasma Me | other |
| Gut Tube | Ctnnb1 | catenin be | Nucleus | transcription regulato |
| Gut Tube | Hnrnph1 | heterogen | Nucleus | other |
| Gut Tube | Krt18 | keratin 18 | Cytoplasm | other |
| Gut Tube | Tmsb10 | thymosin, | Cytoplasm | other |
| Gut Tube | Etv5 | ETS varian | Nucleus | transcription regulato |
| Gut Tube | Ctnna1 | catenin alp | Plasma Me | other |
| Gut Tube | Mdk | midkine | Extracellul | growth factor |
| Gut Tube | Slc38a1 | solute carr | Plasma Me | transporter |
| Gut Tube | Fam32a | family with | Nucleus | other |
| Gut Tube | Alg10b | ALG10, alp | Cytoplasm | enzyme |
| Gut Tube | Mfap2 | microfibril | Extracellul | other |
| Gut Tube | Lxn | latexin | Cytoplasm | other |
| Gut Tube | Ocln | occludin | Plasma Me | other |
| Gut Tube | Efna4 | ephrin A4 | Plasma Me | kinase |
| Gut Tube | Rprm | reprimo, T | Cytoplasm | other |
| Gut Tube | AA986860 | chromosoi | Cytoplasm | other |
| Gut Tube | Cd276 | CD276 mc | Plasma Me | other |

| | | | | |
|---|---|---|---|---|
| Gut Tube | Zfp219 | zinc finger | Nucleus | transcription regulato |
| Gut Tube | Mycn | MYCN pro | Nucleus | transcription regulato |
| Gut Tube | Myl6 | myosin lig | Cytoplasm | enzyme |
| Gut Tube | Myl12b | myosin lig | Cytoplasm | other |
| Gut Tube | Wsb1 | WD repeat | Cytoplasm | enzyme |
| Gut Tube | Gabarapl1 | GABA type | Cytoplasm | other |
| Gut Tube | Actn4 | actinin alp | Cytoplasm | transcription regulato |
| Gut Tube | Map7d1 | MAP7 don | Cytoplasm | other |
| Gut Tube | Rgn | regucalcin | Nucleus | enzyme |
| Gut Tube | Pls3 | plastin 3 | Cytoplasm | other |
| Gut Tube | Tmem132a | transmeml | Cytoplasm | other |
| Gut Tube | Zfp36l1 | ZFP36 ring | Nucleus | transcription regulato |
| Gut Tube | Nono | non-POU | Nucleus | other |
| Gut Tube | Tuba1a | tubulin alp | Cytoplasm | other |
| Gut Tube | Srebf2 | sterol regu | Nucleus | transcription regulato |
| Gut Tube | Eif4ebp2 | eukaryotic | Cytoplasm | translation regulator |
| Gut Tube | Hnrnpm | heterogen | Nucleus | other |
| Gut Tube | Rem2 | RRAD and | Plasma Me | enzyme |
| Gut Tube | Slc16a1 | solute carr | Plasma Me | transporter |
| Gut Tube | Lppr3 | - | - | - |
| Gut Tube | Rasgrp3 | RAS guany | Cytoplasm | other |
| Gut Tube | Ndn | necdin, MA | Nucleus | transcription regulato |
| Gut Tube | Lgals2 | galectin 2 | Cytoplasm | other |
| Gut Tube | Klk8 | kallikrein r | Extracellul | peptidase |
| Gut Tube | Pdpn | podoplani | Plasma Me | other |
| Gut Tube | Mapk6 | mitogen-a | Cytoplasm | kinase |
| Gut Tube | Grn | granulin p | Extracellul | growth factor |
| Gut Tube | Ube2e3 | ubiquitin c | Nucleus | enzyme |
| Gut Tube | Samd10 | sterile alph | Other | other |
| Gut Tube | Tubb4a | tubulin be | Cytoplasm | other |
| Gut Tube | Rac3 | Rac family | Cytoplasm | enzyme |
| Gut Tube | Calu | calumenin | Cytoplasm | other |
| Gut Tube | Cdkn1a | cyclin dep | Nucleus | kinase |
| Gut Tube | Atp1a1 | ATPase Na | Plasma Me | transporter |
| Gut Tube | Cpe | carboxype | Cytoplasm | peptidase |
| Gut Tube | Cpm | carboxype | Plasma Me | peptidase |
| Gut Tube | Snx17 | sorting ne | Cytoplasm | transporter |
| Gut Tube | Skap2 | src kinase | Cytoplasm | other |
| Gut Tube | Aif1l | allograft ir | Plasma Me | other |
| Gut Tube | 4930427A( | - | - | - |
| Gut Tube | Prdx2 | peroxiredo | Cytoplasm | enzyme |
| Gut Tube | Tes | testin LIM | Plasma Me | other |
| Gut Tube | Ogt | O-linked N | Cytoplasm | enzyme |
| Gut Tube | Serpinh1 | serpin fam | Extracellul | other |
| Gut Tube | H2afv | H2A histor | Nucleus | other |
| Gut Tube | Cbx2 | chromobo | Nucleus | transcription regulato |
| Gut Tube | Dpp3 | dipeptidyl | Cytoplasm | peptidase |
| Gut Tube | Tusc3 | tumor sup | Plasma Me | transporter |
| Gut Tube | Ffar4 | free fatty a | Plasma Me | G-protein coupled re |
| Gut Tube | Cgn | cingulin | Plasma Me | other |
| Gut Tube | Flywch2 | FLYWCH fa | Other | other |
| Gut Tube | Igf2bp3 | insulin like | Cytoplasm | translation regulator |
| Gut Tube | Gm1673 | predicted | Other | other |
| Gut Tube | Dstn | destrin, ac | Cytoplasm | other |
| Gut Tube | Nap1l1 | nucleosom | Nucleus | other |
| Gut Tube | Plat | plasminog | Extracellul | peptidase |
| Gut Tube | Me2 | malic enzy | Cytoplasm | enzyme |
| Gut Tube | Slc7a6 | solute carr | Plasma Me | transporter |

| | | | | |
|---|---|---|---|---|
| Gut Tube | Crabp2 | cellular ret | Cytoplasm | transporter |
| Gut Tube | Aplp2 | amyloid be | Cytoplasm | other |
| Gut Tube | Zfp422 | zinc finger | Nucleus | other |
| Gut Tube | Ctxn1 | cortexin 1 | Other | other |
| Gut Tube | Usp22 | ubiquitin s | Nucleus | peptidase |
| Gut Tube | Trappc6a | trafficking | Cytoplasm | other |
| Gut Tube | Itm2c | integral m | Cytoplasm | other |
| Gut Tube | Tomm34 | translocase | Cytoplasm | other |
| Gut Tube | Ppp2r1a | protein ph | Cytoplasm | phosphatase |
| Gut Tube | Usf1 | upstream t | Nucleus | transcription regulato |
| Gut Tube | Cdca7 | cell divisio | Nucleus | other |
| Gut Tube | Gsta4 | glutathion | Other | enzyme |
| Gut Tube | Ash2l | ASH2 like | Nucleus | transcription regulato |
| Gut Tube | Ganab | glucosidas | Cytoplasm | enzyme |
| Gut Tube | Rab25 | RAB25, me | Cytoplasm | enzyme |
| Gut Tube | Dctn2 | dynactin s | Cytoplasm | other |
| Gut Tube | Ccar2 | cell cycle a | Cytoplasm | peptidase |
| Gut Tube | Tra2a | transforme | Nucleus | other |
| Gut Tube | Lrrc42 | leucine ric | Other | other |
| Gut Tube | Nradd | neurotrop | Extracellul | cytokine |
| Gut Tube | Eef1a1 | eukaryotic | Cytoplasm | translation regulator |
| Gut Tube | Xrcc1 | X-ray repa | Nucleus | enzyme |
| Gut Tube | Nedd4 | neural pre | Cytoplasm | enzyme |
| Gut Tube | Naa40 | N(alpha)-a | Other | other |
| Gut Tube | Zfp125 | zinc finger | Nucleus | transcription regulato |
| Gut Tube | Tcf3 | transcripti | Nucleus | transcription regulato |
| Gut Tube | Ctbp1 | C-termina | Nucleus | enzyme |
| Gut Tube | Csrp1 | cysteine ar | Nucleus | other |
| Gut Tube | Adcy8 | adenylate | Plasma Me | enzyme |
| Gut Tube | Morn4 | MORN rep | Extracellul | other |
| Gut Tube | Serinc2 | serine inco | Plasma Me | transporter |
| Gut Tube | Ptk7 | protein tyr | Plasma Me | kinase |
| Gut Tube | Rcn2 | reticulocal | Cytoplasm | other |
| Gut Tube | Rbpms | RNA bindi | Cytoplasm | transcription regulato |
| Gut Tube | Akr7a5 | aldo-keto | Cytoplasm | enzyme |
| Gut Tube | Laptm4a | lysosomal | Cytoplasm | other |
| Gut Tube | Myl9 | myosin lig | Cytoplasm | other |
| Gut Tube | Wbp1 | WW doma | Cytoplasm | other |
| Gut Tube | Spns1 | sphingolip | Cytoplasm | transporter |
| Gut Tube | 5730507C0 | RIKEN cDN | Other | other |
| Gut Tube | Cdk16 | cyclin dep | Cytoplasm | kinase |
| Gut Tube | Tspan14 | tetraspani | Plasma Me | other |
| Gut Tube | Col2a1 | collagen ty | Extracellul | other |
| Gut Tube | Klc4 | kinesin ligl | Other | other |
| Gut Tube | Casp8 | caspase 8 | Nucleus | peptidase |
| Gut Tube | Chmp2b | charged m | Cytoplasm | other |
| Gut Tube | Gpx8 | glutathion | Cytoplasm | enzyme |
| Gut Tube | Gipc1 | GIPC PDZ | Cytoplasm | other |
| Gut Tube | Gins4 | GINS comp | Nucleus | other |
| Gut Tube | Suv420h2 | - | - | - |
| Gut Tube | Hnrnpc | heterogen | Nucleus | other |
| Gut Tube | Ivns1abp | influenza v | Nucleus | other |
| Gut Tube | Ift43 | intraflagell | Cytoplasm | other |
| Gut Tube | R3hdm4 | R3H doma | Other | other |
| Gut Tube | Nudt14 | nudix hydr | Cytoplasm | phosphatase |
| Gut Tube | Glrx | glutaredox | Cytoplasm | enzyme |
| Gut Tube | Pigs | phosphatid | Cytoplasm | enzyme |
| Gut Tube | Golm1 | golgi mem | Cytoplasm | other |

| | | | | |
|---|---|---|---|---|
| Gut Tube | Mfge8 | milk fat glo | Extracellula | other |
| Gut Tube | Sap30 | Sin3A asso | Nucleus | transcription regulato |
| Gut Tube | Abracl | ABRA C-te | Other | other |
| Gut Tube | Adprh | ADP-ribos | Other | enzyme |
| Gut Tube | Rcan1 | regulator c | Nucleus | transcription regulato |
| Gut Tube | Mest | mesoderm | Cytoplasm | peptidase |
| Gut Tube | Psap | prosaposir | Extracellula | enzyme |
| Gut Tube | Cdk4 | cyclin depe | Nucleus | kinase |
| Gut Tube | Eno1b | enolase 1 | Cytoplasm | enzyme |
| Gut Tube | Ncstn | nicastrin | Plasma Me | peptidase |
| Gut Tube | Cst3 | cystatin C | Extracellula | other |
| Gut Tube | Sf3b2 | splicing fa | Nucleus | other |
| Gut Tube | Myl12a | myosin lig | Cytoplasm | other |
| Gut Tube | Hmgn1 | high mobi | Nucleus | transcription regulato |
| Gut Tube | Lypla2 | lysophosp | Cytoplasm | enzyme |
| Gut Tube | Ptma | prothymos | Nucleus | other |
| Gut Tube | Emb | embigin | Plasma Me | transporter |
| Gut Tube | Cdc42ep1 | CDC42 effe | Extracellula | other |
| Gut Tube | Gabarap | GABA type | Cytoplasm | transporter |
| Gut Tube | Eno1 | enolase 1 | Cytoplasm | enzyme |
| Gut Tube | Ttc5 | tetratricop | Nucleus | other |
| Gut Tube | Ogfr | opioid gro | Plasma Me | other |
| Gut Tube | Zfand6 | zinc finger | Other | other |
| Gut Tube | Bsg | basigin (O | Plasma Me | transporter |
| Gut Tube | Xrcc5 | X-ray repa | Nucleus | enzyme |
| Gut Tube | Smim22 | small integ | Other | other |
| Gut Tube | Ccdc23 | - | - | - |
| Gut Tube | Ctsz | cathepsin | Cytoplasm | peptidase |
| Gut Tube | Rcn3 | reticulocal | Cytoplasm | other |
| Gut Tube | Pgrmc1 | progestero | Plasma Me | transmembrane recep |
| Gut Tube | Csrp2 | cysteine ar | Nucleus | other |
| Gut Tube | Uba1 | ubiquitin li | Cytoplasm | enzyme |
| Gut Tube | Txnrd1 | thioredoxi | Cytoplasm | enzyme |
| Gut Tube | Nap1l4 | nucleosom | Cytoplasm | other |
| Gut Tube | Mcm2 | minichrom | Nucleus | enzyme |
| Gut Tube | Ppdpf | pancreatic | Other | other |
| Gut Tube | Tmbim6 | transmeml | Nucleus | other |
| Gallbladder Primordium | Sox17 | SRY-box 1 | Nucleus | transcription regulato |
| Gallbladder Primordium | Gjb2 | gap juncti | Plasma Me | transporter |
| Gallbladder Primordium | Lgr4 | leucine ric | Plasma Me | transmembrane recep |
| Gallbladder Primordium | Pgpep1 | pyroglutar | Cytoplasm | peptidase |
| Gallbladder Primordium | Dctn4 | dynactin s | Nucleus | other |
| Gallbladder Primordium | Flrt3 | fibronectir | Plasma Me | other |
| Gallbladder Primordium | 1700011H | coiled-coi | Other | other |
| Gallbladder Primordium | Sox11 | SRY-box 1 | Nucleus | transcription regulato |
| Gallbladder Primordium | Flrt2 | fibronectir | Plasma Me | other |
| Gallbladder Primordium | 2610008E1 | RIKEN cDN | Other | other |
| Gallbladder Primordium | Prr15l | proline ricl | Other | other |
| Gallbladder Primordium | Itpripl1 | ITPRIP like | Other | other |
| Gallbladder Primordium | Ift22 | intraflagell | Extracellula | other |
| Gallbladder Primordium | Grpel2 | GrpE like 2 | Cytoplasm | other |
| Gallbladder Primordium | Gsk3b | glycogen s | Nucleus | kinase |
| Gallbladder Primordium | Cbfa2t2 | CBFA2/RU | Nucleus | transcription regulato |
| Gallbladder Primordium | Onecut1 | one cut hc | Nucleus | transcription regulato |
| Gallbladder Primordium | Pdcd4 | programm | Nucleus | other |
| Gallbladder Primordium | Rab38 | RAB38, me | Cytoplasm | enzyme |
| Gallbladder Primordium | Slc2a2 | solute carr | Plasma Me | transporter |
| Gallbladder Primordium | Tcf7l2 | transcriptio | Nucleus | transcription regulato |

| | | | | |
|---|---|---|---|---|
| Gallbladder Primordium | Igsf3 | immunogl | Plasma Me | other |
| Gallbladder Primordium | Tjp2 | tight junct | Plasma Me | kinase |
| Gallbladder Primordium | Dpysl2 | dihydropy | Cytoplasm | enzyme |
| Gallbladder Primordium | Gstm2 | glutathion | Cytoplasm | enzyme |
| Gallbladder Primordium | Arl4c | ADP ribosy | Nucleus | enzyme |
| Gallbladder Primordium | Stk24 | serine/thre | Cytoplasm | kinase |
| Gallbladder Primordium | Rab13 | RAB13, me | Plasma Me | enzyme |
| Gallbladder Primordium | Zfp691 | zinc finger | Nucleus | other |
| Gallbladder Primordium | Sri | sorcin | Cytoplasm | transporter |
| Gallbladder Primordium | Rxrg | retinoid X | Nucleus | ligand-dependent nu |
| Gallbladder Primordium | Klf13 | Kruppel lik | Nucleus | transcription regulato |
| Gallbladder Primordium | Sparc | secreted p | Extracellula | other |
| Gallbladder Primordium | Kif3a | kinesin fan | Cytoplasm | enzyme |
| Gallbladder Primordium | Cenpe | centromer | Nucleus | other |
| Gallbladder Primordium | Zfp37 | ZFP37 zinc | Nucleus | transcription regulato |
| Gallbladder Primordium | Sox9 | SRY-box 9 | Nucleus | transcription regulato |
| Gallbladder Primordium | Tspyl1 | TSPY like 1 | Nucleus | other |
| Gallbladder Primordium | Ccdc92 | coiled-coi | Cytoplasm | other |
| Gallbladder Primordium | Ifitm3 | interferon | Plasma Me | other |
| Gallbladder Primordium | Id2 | inhibitor o | Nucleus | transcription regulato |
| Gallbladder Primordium | Plk2 | polo like k | Nucleus | kinase |
| Gallbladder Primordium | Prr13 | proline ricl | Other | other |
| Gallbladder Primordium | Crabp1 | cellular ret | Cytoplasm | transporter |
| Gallbladder Primordium | Camk1g | calcium/ca | Cytoplasm | kinase |
| Gallbladder Primordium | Sesn1 | sestrin 1 | Nucleus | other |
| Gallbladder Primordium | Ppox | protoporp | Cytoplasm | enzyme |
| Gallbladder Primordium | Gm10479 | predicted | Other | other |
| Gallbladder Primordium | Mgst1 | microsoma | Cytoplasm | enzyme |
| Gallbladder Primordium | Usp11 | ubiquitin s | Nucleus | peptidase |
| Gallbladder Primordium | Fbxo44 | F-box pro | Cytoplasm | enzyme |
| Gallbladder Primordium | Fbxo18 | F-box pro | Nucleus | enzyme |
| Gallbladder Primordium | Sall4 | spalt like t | Nucleus | transcription regulato |
| Gallbladder Primordium | Ndufa3 | NADH:ubi | Cytoplasm | enzyme |
| Gallbladder Primordium | Zscan26 | zinc finger | Nucleus | transcription regulato |
| Gallbladder Primordium | Dcaf7 | DDB1 and | Cytoplasm | other |
| Gallbladder Primordium | Csde1 | cold shock | Cytoplasm | enzyme |
| Gallbladder Primordium | Dynlt1f | dynein ligh | Cytoplasm | other |
| Gallbladder Primordium | Fabp3 | fatty acid b | Cytoplasm | transporter |
| Gallbladder Primordium | Gm10031 | predicted | Other | other |
| Gallbladder Primordium | Gpr108 | G protein- | Plasma Me | G-protein coupled re |
| Gallbladder Primordium | Stxbp2 | syntaxin bi | Plasma Me | transporter |
| Gallbladder Primordium | Med18 | mediator c | Nucleus | other |
| Gallbladder Primordium | Opa3 | OPA3, out | Cytoplasm | other |
| Gallbladder Primordium | 9030624G | predicted | Cytoplasm | other |
| Gallbladder Primordium | Ppp1r1a | protein ph | Cytoplasm | phosphatase |
| Gallbladder Primordium | Krtcap3 | keratinocy | Other | other |
| Gallbladder Primordium | Dynlt1b | dynein ligh | Cytoplasm | other |
| Gallbladder Primordium | Tes | testin LIM | Plasma Me | other |
| Gallbladder Primordium | Dynlt1a | dynein ligh | Cytoplasm | other |
| Gallbladder Primordium | Tgif1 | TGFB indu | Nucleus | transcription regulato |
| Gallbladder Primordium | Col2a1 | collagen ty | Extracellula | other |
| Gallbladder Primordium | Cystm1 | cysteine-ri | Cytoplasm | other |
| Gallbladder Primordium | Alas1 | 5'-aminole | Cytoplasm | enzyme |
| Gallbladder Primordium | Gstm7 | glutathion | Cytoplasm | enzyme |
| Gallbladder Primordium | St3gal5 | ST3 beta-g | Cytoplasm | enzyme |
| Gallbladder Primordium | Trim35 | tripartite n | Cytoplasm | other |
| Gallbladder Primordium | Eci2 | enoyl-CoA | Cytoplasm | enzyme |
| Gallbladder Primordium | Narf | nuclear pr | Nucleus | enzyme |

| | | | | |
|---|---|---|---|---|
| Gallbladder Primordium | Atat1 | alpha tubu | Cytoplasm | enzyme |
| Gallbladder Primordium | B4galt1 | beta-1,4-g | Cytoplasm | enzyme |
| Gallbladder Primordium | Serinc2 | serine inco | Plasma Me | transporter |
| Gallbladder Primordium | Hbp1 | HMG-box | Nucleus | transcription regulato |
| Gallbladder Primordium | Clic6 | chloride in | Plasma Me | ion channel |
| Gallbladder Primordium | Fbxo36 | F-box pro | Other | other |
| Gallbladder Primordium | Anxa5 | annexin A5 | Plasma Me | transporter |
| Gallbladder Primordium | Msantd4 | Myb/SANT | Nucleus | other |
| Gallbladder Primordium | Zcchc18 | zinc finger | Other | other |
| Gallbladder Primordium | Bcl2l1 | BCL2 like 1 | Cytoplasm | other |
| Gallbladder Primordium | Rhoc | ras homolo | Plasma Me | enzyme |
| Gallbladder Primordium | Arglu1 | arginine ar | Other | other |
| Gallbladder Primordium | Ddah2 | dimethylar | Cytoplasm | enzyme |
| Gallbladder Primordium | Bet1l | Bet1 golgi | Cytoplasm | transporter |
| Gallbladder Primordium | Dynlt1c | dynein ligh | Cytoplasm | other |
| Gallbladder Primordium | Rnase4 | ribonuclea | Extracellula | enzyme |
| Gallbladder Primordium | S100a16 | S100 calciu | Nucleus | other |
| Gallbladder Primordium | Glrx | glutaredox | Cytoplasm | enzyme |
| Gallbladder Primordium | Gstt3 | glutathion | Other | enzyme |
| Gallbladder Primordium | Cacnb3 | calcium vo | Plasma Me | ion channel |
| Gallbladder Primordium | Cdca2 | cell divisio | Nucleus | other |
| Gallbladder Primordium | Ncapd2 | non-SMC | Nucleus | other |
| Gallbladder Primordium | Tmc4 | transmemb | Cytoplasm | other |
| Gallbladder Primordium | Nmt2 | N-myristo | Cytoplasm | enzyme |
| Gallbladder Primordium | Hsd17b10 | hydroxyste | Cytoplasm | enzyme |
| Gallbladder Primordium | Vamp8 | vesicle ass | Plasma Me | transporter |
| Gallbladder Primordium | Ccdc23 | - | - | - |
| Gallbladder Primordium | Aktip | AKT intera | Cytoplasm | other |
| Gallbladder Primordium | Ntan1 | N-termina | Nucleus | enzyme |
| Gallbladder Primordium | Nrp1 | neuropilin | Plasma Me | transmembrane recep |
| Gallbladder Primordium | Hkdc1 | hexokinase | Cytoplasm | kinase |
| Gallbladder Primordium | 2700062C0 | chromosom | Other | other |
| Gallbladder Primordium | Mboat2 | membrane | Cytoplasm | enzyme |
| Gallbladder Primordium | Cmtm8 | CKLF like M | Extracellula | cytokine |
| Gallbladder Primordium | Meis3 | Meis home | Nucleus | transcription regulato |
| Gallbladder Primordium | Clic1 | chloride in | Nucleus | ion channel |
| Gallbladder Primordium | Ndufa12 | NADH:ubi | Cytoplasm | enzyme |
| Gallbladder Primordium | Nusap1 | nucleolar a | Nucleus | other |
| Gallbladder Primordium | Zfp61 | zinc finger | Nucleus | peptidase |
| Gallbladder Primordium | Tmem206 | transmemb | Plasma Me | other |
| Gallbladder Primordium | Aip | aryl hydro | Nucleus | transcription regulato |
| Gallbladder Primordium | Rpusd1 | RNA pseud | Other | enzyme |
| Gallbladder Primordium | Atp6v1g1 | ATPase H+ | Cytoplasm | transporter |
| Gallbladder Primordium | Grina | glutamate | Other | ion channel |
| Gallbladder Primordium | Lgals1 | galectin 1 | Extracellula | other |
| Gallbladder Primordium | Ptprk | protein tyr | Plasma Me | phosphatase |
| Gallbladder Primordium | Zfp125 | zinc finger | Nucleus | transcription regulato |
| Gallbladder Primordium | Cox7a1 | cytochrom | Cytoplasm | enzyme |
| Gallbladder Primordium | Kdm6b | lysine dem | Extracellula | enzyme |
| Gallbladder Primordium | Cd81 | CD81 mole | Plasma Me | other |
| Gallbladder Primordium | Styx | serine/thre | Cytoplasm | phosphatase |
| Gallbladder Primordium | Notch2 | notch 2 | Plasma Me | transcription regulato |
| Gallbladder Primordium | Hn1 | - | - | - |
| Gallbladder Primordium | Maged2 | MAGE fam | Plasma Me | other |
| Gallbladder Primordium | Emb | embigin | Plasma Me | transporter |
| Gallbladder Primordium | Fgfr2 | fibroblast g | Plasma Me | kinase |
| Gallbladder Primordium | Psme2 | proteasom | Cytoplasm | peptidase |
| Gallbladder Primordium | Calm2 | calmodulin | Nucleus | other |

| | | | | |
|---|---|---|---|---|
| Gallbladder Primordium | Chrnb1 | cholinergi | Plasma Me | transmembrane recep |
| Gallbladder Primordium | Foxa2 | forkhead b | Nucleus | transcription regulato |
| Gallbladder Primordium | Gltp | glycolipid | Cytoplasm | transporter |
| Gallbladder Primordium | Gng11 | G protein s | Plasma Me | enzyme |
| Gallbladder Primordium | Pdrg1 | p53 and D | Cytoplasm | other |
| Gallbladder Primordium | Igfbp2 | insulin like | Extracellula | other |
| Gallbladder Primordium | Tmem59 | transmeml | Plasma Me | peptidase |
| Gallbladder Primordium | Tmem106c | transmeml | Other | other |
| Gallbladder Primordium | Rcan1 | regulator c | Nucleus | transcription regulato |
| Gallbladder Primordium | Cmtm3 | CKLF like M | Extracellula | cytokine |
| Gallbladder Primordium | Sdr39u1 | short chain | Nucleus | other |
| Gallbladder Primordium | Nras | NRAS prot | Plasma Me | enzyme |
| Gallbladder Primordium | Tmem218 | transmeml | Extracellula | other |
| Gallbladder Primordium | 1110007C( | - | - | - |
| Gallbladder Primordium | Cirbp | cold induc | Nucleus | translation regulator |
| Gallbladder Primordium | Tfpt | TCF3 fusio | Nucleus | other |
| Gallbladder Primordium | Pde6d | phosphod | Cytoplasm | enzyme |
| Gallbladder Primordium | BC005624 | chromosoi | Other | other |
| Gallbladder Primordium | Ndfip1 | Nedd4 fan | Cytoplasm | other |
| Gallbladder Primordium | Tead2 | TEA doma | Nucleus | transcription regulato |
| Gallbladder Primordium | Pum1 | pumilio RN | Cytoplasm | other |
| Gallbladder Primordium | Gc | GC, vitami | Extracellula | transporter |
| Gallbladder Primordium | Prmt2 | protein arg | Nucleus | enzyme |
| Gallbladder Primordium | Tm4sf4 | transmeml | Plasma Me | other |
| Gallbladder Primordium | Ptp4a2 | protein tyr | Cytoplasm | phosphatase |
| Gallbladder Primordium | Kif23 | kinesin fan | Cytoplasm | other |
| Gallbladder Primordium | Serping1 | serpin fam | Extracellula | other |
| Gallbladder Primordium | Nr2f2 | nuclear rec | Nucleus | ligand-dependent nu |
| Gallbladder Primordium | Gbas | - | - | - |
| Gallbladder Primordium | Hnrnpa0 | heterogen | Nucleus | other |
| Gallbladder Primordium | Zfp825 | expressed | Other | other |
| Gallbladder Primordium | Pfn2 | profilin 2 | Cytoplasm | enzyme |
| Gallbladder Primordium | Dolk | dolichol ki | Cytoplasm | kinase |
| Gallbladder Primordium | Rgs19 | regulator c | Cytoplasm | other |
| Gallbladder Primordium | Spp1 | secreted p | Extracellula | cytokine |
| Gallbladder Primordium | Cited2 | Cbp/p300 | Nucleus | transcription regulato |
| Gallbladder Primordium | Commd10 | COMM dc | Other | other |
| Gallbladder Primordium | Polr3k | RNA polyr | Nucleus | transcription regulato |
| Gallbladder Primordium | Carhsp1 | calcium re | Cytoplasm | other |
| Gallbladder Primordium | Nelfb | negative e | Nucleus | other |
| Gallbladder Primordium | Pdpn | podoplani | Plasma Me | other |
| Gallbladder Primordium | Psme1 | proteasom | Cytoplasm | other |
| Gallbladder Primordium | Kdm3a | lysine dem | Nucleus | transcription regulato |
| Gallbladder Primordium | Cox4i1 | cytochrom | Cytoplasm | enzyme |
| Gallbladder Primordium | G3bp2 | G3BP stres | Cytoplasm | enzyme |
| Gallbladder Primordium | Cep57 | centrosom | Cytoplasm | other |
| Gallbladder Primordium | 4833420G | chromosoi | Other | other |
| Gallbladder Primordium | Car14 | carbonic a | Plasma Me | enzyme |
| Gallbladder Primordium | Eif2b2 | eukaryotic | Cytoplasm | other |
| Gallbladder Primordium | Pgm1 | phosphog | Cytoplasm | enzyme |
| Gallbladder Primordium | Mapk3 | mitogen-a | Cytoplasm | kinase |
| Gallbladder Primordium | Tax1bp3 | Tax1 bindi | Cytoplasm | transcription regulato |
| Gallbladder Primordium | Fblim1 | filamin bin | Plasma Me | other |
| Gallbladder Primordium | Ift52 | intraflagell | Cytoplasm | other |
| Gallbladder Primordium | Npc2 | NPC intrac | Extracellula | transporter |
| Gallbladder Primordium | Deaf1 | DEAF1, tra | Nucleus | transcription regulato |
| Gallbladder Primordium | Slc50a1 | solute carr | Plasma Me | transporter |
| Gallbladder Primordium | Fopnl | FGFR1OP I | Cytoplasm | other |

| | | | | |
|---|---|---|---|---|
| Gallbladder Primordium | Clybl | citrate lyas | Cytoplasm | enzyme |
| Gallbladder Primordium | Ttc5 | tetratricop | Nucleus | other |
| Gallbladder Primordium | Epcam | epithelial c | Plasma Me | other |
| Gallbladder Primordium | Pigyl | phosphatic | Plasma Me | other |
| Gallbladder Primordium | Cnot8 | CCR4-NOT | Nucleus | transcription regulato |
| Gallbladder Primordium | Map1lc3b | microtubu | Cytoplasm | other |
| Gallbladder Primordium | Sf3b2 | splicing fa | Nucleus | other |
| Gallbladder Primordium | Myadm | myeloid as | Nucleus | other |
| Gallbladder Primordium | Spg21 | SPG21, ma | Plasma Me | enzyme |
| Gallbladder Primordium | Lrrc58 | leucine ric | Other | other |
| Gallbladder Primordium | Zkscan6 | zinc finger | Nucleus | transcription regulato |
| Gallbladder Primordium | Spry2 | sprouty RT | Plasma Me | other |
| Gallbladder Primordium | Ackr3 | atypical ch | Plasma Me | G-protein coupled re |
| Gallbladder Primordium | Hhex | hematopo | Nucleus | transcription regulato |
| Gallbladder Primordium | 1110059E2 | chromoso | Other | other |
| Gallbladder Primordium | Rab3ip | RAB3A inte | Cytoplasm | other |
| Gallbladder Primordium | Lxn | latexin | Cytoplasm | other |
| Gallbladder Primordium | Spon2 | spondin 2 | Extracellul | other |
| Gallbladder Primordium | Zfr | zinc finger | Nucleus | other |
| Gallbladder Primordium | Gnb1 | G protein s | Plasma Me | enzyme |
| Gallbladder Primordium | Morn2 | MORN rep | Other | other |
| Gallbladder Primordium | Marcksl1 | MARCKS li | Cytoplasm | other |
| Gallbladder Primordium | Tmem51 | transmeml | Other | other |
| Gallbladder Primordium | S100a11 | S100 calciu | Cytoplasm | other |
| Gallbladder Primordium | Tmem242 | transmeml | Other | other |
| Gallbladder Primordium | Itm2c | integral m | Cytoplasm | other |
| Gallbladder Primordium | Coro1b | coronin 1E | Cytoplasm | other |
| Gallbladder Primordium | Abracl | ABRA C-te | Other | other |
| Gallbladder Primordium | Snx17 | sorting ne | Cytoplasm | transporter |
| Gallbladder Primordium | Paip1 | poly(A) bir | Cytoplasm | translation regulator |
| Gallbladder Primordium | Ddx42 | DEAD-box | Cytoplasm | enzyme |
| Gallbladder Primordium | Zfp105 | zinc finger | Nucleus | transcription regulato |
| Gallbladder Primordium | 1500009L1 | RIKEN cDN | Other | other |
| Gallbladder Primordium | Dazap2 | DAZ assoc | Nucleus | other |
| Gallbladder Primordium | Smim22 | small integ | Other | other |
| Gallbladder Primordium | Rp9 | RP9, pre-r | Nucleus | other |
| Gallbladder Primordium | 2610524H | RIKEN cDN | Other | other |
| Gallbladder Primordium | Snupn | snurportin | Nucleus | other |
| Gallbladder Primordium | Twf1 | twinfilin ac | Cytoplasm | kinase |
| Gallbladder Primordium | EGFP | - | - | - |
| Gallbladder Primordium | BC023829 | - | - | - |
| Gallbladder Primordium | Leo1 | LEO1 hom | Nucleus | other |
| Gallbladder Primordium | Fam173a | family with | Other | other |
| Gallbladder Primordium | Sh3bp4 | SH3 doma | Cytoplasm | other |
| Gallbladder Primordium | Trappc2l | trafficking | Cytoplasm | other |
| Gallbladder Primordium | 5730507C | RIKEN cDN | Other | other |
| Gallbladder Primordium | Mcoln3 | mucolipin | Plasma Me | ion channel |
| Gallbladder Primordium | H3f3b | H3 histone | Nucleus | other |
| Gallbladder Primordium | Pcbd2 | pterin-4 al | Other | enzyme |
| Gallbladder Primordium | H2afv | H2A histor | Nucleus | other |
| Gallbladder Primordium | Atp6v1d | ATPase H+ | Cytoplasm | transporter |
| Gallbladder Primordium | Cmtm7 | CKLF like M | Extracellul | cytokine |
| Gallbladder Primordium | Fam96a | family with | Extracellul | other |
| Gallbladder Primordium | Cdc123 | cell divisio | Cytoplasm | other |
| Gallbladder Primordium | Idnk | IDNK, gluc | Cytoplasm | other |
| Gallbladder Primordium | Prdx2 | peroxiredo | Cytoplasm | enzyme |
| Gallbladder Primordium | Chchd6 | coiled-coi | Cytoplasm | other |
| Gallbladder Primordium | Arpc5 | actin relate | Cytoplasm | other |

| | | | | |
|---|---|---|---|---|
| Gallbladder Primordium | Adcy8 | adenylate | Plasma Me | enzyme |
| Gallbladder Primordium | Pafah1b3 | platelet ac | Cytoplasm | enzyme |
| Gallbladder Primordium | Gdi1 | GDP disso | Cytoplasm | other |
| Gallbladder Primordium | Map1lc3a | microtubu | Cytoplasm | other |
| Gallbladder Primordium | Ankrd46 | ankyrin rep | Nucleus | transcription regulato |
| Gallbladder Primordium | Fam60a | SIN3-HDA | Extracellul | other |
| Gallbladder Primordium | Sfr1 | SWI5 depe | Nucleus | transcription regulato |
| Gallbladder Primordium | Fbxo25 | F-box pro | Nucleus | enzyme |
| Gallbladder Primordium | Stoml1 | stomatin li | Other | other |
| Gallbladder Primordium | Pak1 | p21 (RAC1 | Cytoplasm | kinase |
| Gallbladder Primordium | Rab11a | RAB11A, n | Cytoplasm | enzyme |
| Gallbladder Primordium | Kif12 | kinesin fan | Cytoplasm | other |
| Gallbladder Primordium | Mdk | midkine | Extracellul | growth factor |
| Gallbladder Primordium | AW209491 | chromoso | Other | other |
| Gallbladder Primordium | Lyrm2 | LYR motif | Cytoplasm | other |
| Gallbladder Primordium | Gpx8 | glutathion | Cytoplasm | enzyme |
| Gallbladder Primordium | Sdhc | succinate | Cytoplasm | enzyme |
| Gallbladder Primordium | Klhl9 | kelch like f | Cytoplasm | other |
| Gallbladder Primordium | Cldn7 | claudin 7 | Plasma Me | other |
| Gallbladder Primordium | Tmem254b | transmeml | Other | other |
| Gallbladder Primordium | Ift43 | intraflagell | Cytoplasm | other |
| Gallbladder Primordium | Haus1 | HAUS aug | Nucleus | other |
| Gallbladder Primordium | Cox4i2 | cytochrom | Cytoplasm | enzyme |
| Gallbladder Primordium | Hes1 | hes family | Nucleus | transcription regulato |
| Gallbladder Primordium | Anapc2 | anaphase | Nucleus | other |
| Gallbladder Primordium | Hnf1b | HNF1 hom | Nucleus | transcription regulato |
| Gallbladder Primordium | Gnb5 | G protein s | Plasma Me | enzyme |
| Gallbladder Primordium | Mapk13 | mitogen-a | Cytoplasm | kinase |
| Gallbladder Primordium | Pdk3 | pyruvate d | Cytoplasm | kinase |
| Gallbladder Primordium | Incenp | inner centr | Nucleus | other |
| Gallbladder Primordium | Mea1 | male-enha | Cytoplasm | other |
| Gallbladder Primordium | Casp8 | caspase 8 | Nucleus | peptidase |
| Gallbladder Primordium | Spint1 | serine pep | Extracellul | other |
| Gallbladder Primordium | Tmem254a | transmeml | Other | other |
| Gallbladder Primordium | Gas8 | growth arr | Cytoplasm | other |
| Gallbladder Primordium | Kif22 | kinesin fan | Nucleus | other |
| Gallbladder Primordium | Pdcd2l | programm | Other | other |
| Gallbladder Primordium | Mageh1 | MAGE fam | Cytoplasm | other |
| Gallbladder Primordium | Uqcc3 | ubiquinol- | Extracellul | other |
| Gallbladder Primordium | Stmn1 | stathmin 1 | Cytoplasm | other |
| Gallbladder Primordium | Rassf7 | Ras associ | Cytoplasm | other |
| Gallbladder Primordium | Rcn2 | reticulocal | Cytoplasm | other |
| Gallbladder Primordium | Lppr3 | - | - | - |
| Gallbladder Primordium | Tubb4a | tubulin be | Cytoplasm | other |
| Gallbladder Primordium | Nrep | neuronal r | Cytoplasm | other |
| Gallbladder Primordium | Mgst3 | microsoma | Cytoplasm | enzyme |
| Gallbladder Primordium | Isca2 | iron-sulfur | Cytoplasm | other |
| Gallbladder Primordium | Akap8l | A-kinase a | Nucleus | other |
| Gallbladder Primordium | Lipa | lipase A, ly | Cytoplasm | enzyme |
| Gallbladder Primordium | Snx6 | sorting ne | Cytoplasm | transporter |
| Gallbladder Primordium | Kdf1 | keratinocy | Plasma Me | other |
| Gallbladder Primordium | Syt5 | synaptotag | Cytoplasm | transporter |
| Gallbladder Primordium | Tubb5 | tubulin be | Cytoplasm | other |
| Gallbladder Primordium | Pcyox1l | prenylcyst | Other | other |
| Gallbladder Primordium | Lrrc42 | leucine ric | Other | other |
| Gallbladder Primordium | Cldn6 | claudin 6 | Plasma Me | other |
| Gallbladder Primordium | Cdkn1a | cyclin dep | Nucleus | kinase |
| Gallbladder Primordium | Wbp5 | - | - | - |

| | | | | |
|---|---|---|---|---|
| Gallbladder Primordium | Ccnf | cyclin F | Nucleus | enzyme |
| Gallbladder Primordium | Cbx3 | chromobo | Nucleus | transcription regulato |
| Gallbladder Primordium | Ctxn1 | cortexin 1 | Other | other |
| Gallbladder Primordium | 0610040J0 | chromosoı | Nucleus | other |
| Gallbladder Primordium | Polb | DNA polyr | Nucleus | enzyme |
| Gallbladder Primordium | Atp6v1f | ATPase H+ | Cytoplasm | enzyme |
| Gallbladder Primordium | Tmem254c | transmeml | Other | other |
| Gallbladder Primordium | Armc10 | armadillo ı | Cytoplasm | other |
| Gallbladder Primordium | Zfand6 | zinc finger | Other | other |
| Gallbladder Primordium | Prelid2 | PRELI dom | Cytoplasm | other |
| Gallbladder Primordium | Sept2 | septin 2 | Cytoplasm | enzyme |
| Gallbladder Primordium | Mfap2 | microfibril | Extracellul | other |
| Gallbladder Primordium | Fxyd6 | FXYD dom | Plasma Me | ion channel |
| Gallbladder Primordium | Timm21 | translocase | Cytoplasm | other |
| Gallbladder Primordium | Fam103a1 | family with | Nucleus | other |
| Gallbladder Primordium | 1810037I1 | predicted | Other | other |
| Gallbladder Primordium | Ubl7 | ubiquitin li | Other | other |
| Gallbladder Primordium | Tubb2a | tubulin be | Cytoplasm | other |
| Gallbladder Primordium | Fibp | FGF1 intra | Nucleus | other |
| Gallbladder Primordium | Cst3 | cystatin C | Extracellul | other |
| Gallbladder Primordium | Erdr1 | erythroid c | Other | other |
| Gallbladder Primordium | Gm8797 | ubiquitin E | Other | other |
| Gallbladder Primordium | Coprs | coordinatc | Nucleus | other |
| Gallbladder Primordium | Smim12 | small integ | Other | other |
| Gallbladder Primordium | Mrpl27 | mitochonc | Cytoplasm | other |
| Gallbladder Primordium | Arf5 | ADP ribosy | Cytoplasm | enzyme |
| Gallbladder Primordium | Ppic | peptidylpr | Cytoplasm | enzyme |
| Gallbladder Primordium | Prkar1a | protein kir | Cytoplasm | kinase |
| Gallbladder Primordium | Tma7 | translation | Other | other |
| Gallbladder Primordium | Gm10282 | high mobi | Other | other |
| Gallbladder Primordium | Lsm10 | LSM10, U7 | Nucleus | other |
| Gallbladder Primordium | Rrp36 | ribosomal | Nucleus | other |
| Gallbladder Primordium | Leprot | leptin rece | Plasma Me | other |
| Gallbladder Primordium | S100a10 | S100 calciu | Cytoplasm | other |
| Gallbladder Primordium | Nono | non-POU | Nucleus | other |
| Gallbladder Primordium | Plat | plasminog | Extracellul | peptidase |
| Gallbladder Primordium | Ogt | O-linked N | Cytoplasm | enzyme |
| Gallbladder Primordium | Gsta4 | glutathion | Other | enzyme |
| Gallbladder Primordium | Ttc1 | tetratricop | Cytoplasm | other |
| Gallbladder Primordium | Col4a2 | collagen ty | Extracellul | other |
| Gallbladder Primordium | Gm1673 | predicted | Other | other |
| Gallbladder Primordium | Tmem261 | - | - | - |
| Gallbladder Primordium | Rtfdc1 | - | - | - |
| Gallbladder Primordium | Adi1 | acireducto | Nucleus | enzyme |
| Gallbladder Primordium | Tmem86a | transmeml | Other | other |
| Gallbladder Primordium | Ubb | ubiquitin E | Cytoplasm | other |
| Gallbladder Primordium | Dhrs4 | dehydroge | Cytoplasm | enzyme |
| Gallbladder Primordium | Cd24a | CD24a ant | Plasma Me | other |
| Gallbladder Primordium | Dera | deoxyribos | Cytoplasm | enzyme |
| Gallbladder Primordium | Ifitm2 | interferon | Cytoplasm | other |
| Gallbladder Primordium | Slc25a17 | solute carr | Cytoplasm | transporter |
| Gallbladder Primordium | Fam32a | family with | Nucleus | other |
| Gallbladder Primordium | H2afy2 | H2A histor | Nucleus | other |
| Gallbladder Primordium | Dctn3 | dynactin sı | Nucleus | other |
| Gallbladder Primordium | Gm21887 | - | - | - |
| Gallbladder Primordium | Manbal | mannosida | Other | other |
| Gallbladder Primordium | Dync2li1 | dynein cyt | Cytoplasm | other |
| Gallbladder Primordium | Tmsb10 | thymosin, | Cytoplasm | other |

| | | | | |
|---|---|---|---|---|
| Gallbladder Primordium | Rabac1 | Rab accep | Cytoplasm | other |
| Gallbladder Primordium | Pts | 6-pyruvoy | Cytoplasm | enzyme |
| Gallbladder Primordium | Mzt1 | mitotic spi | Cytoplasm | other |
| Gallbladder Primordium | Higd2a | HIG1 hypc | Cytoplasm | other |
| Gallbladder Primordium | Tmbim6 | transmeml | Nucleus | other |
| Gallbladder Primordium | Romo1 | reactive ox | Cytoplasm | other |
| Gallbladder Primordium | Rsrp1 | arginine ar | Other | other |
| Gallbladder Primordium | Lamtor5 | late endos | Cytoplasm | other |
| Gallbladder Primordium | Psmb2 | proteasom | Cytoplasm | peptidase |
| Gallbladder Primordium | Hmgb3 | high mobi | Nucleus | other |
| Gallbladder Primordium | 1700123O | chromosoı | Other | other |
| Gallbladder Primordium | Hsd11b2 | hydroxyste | Cytoplasm | enzyme |
| Gallbladder Primordium | Nudt16l1 | nudix hydı | Cytoplasm | other |
| Gallbladder Primordium | Ndrg2 | NDRG fam | Cytoplasm | other |
| Gallbladder Primordium | Dctn5 | dynactin sı | Cytoplasm | other |
| Gallbladder Primordium | Nsmce1 | NSE1 hom | Nucleus | transporter |
| Gallbladder Primordium | Gja1 | gap juncti | Plasma Me | transporter |
| Gallbladder Primordium | Pea15a | proliferatic | Cytoplasm | transporter |
| Gallbladder Primordium | Apoa1bp | - | - | - |
| Gallbladder Primordium | Cdh6 | cadherin 6 | Plasma Me | other |
| Gallbladder Primordium | Hsbp1 | heat shock | Nucleus | other |
| Gallbladder Primordium | Habp2 | hyaluronaı | Extracellul | peptidase |
| Gallbladder Primordium | Serpinh1 | serpin fam | Extracellul | other |
| Gallbladder Primordium | Coa3 | cytochrom | Cytoplasm | other |
| Gallbladder Primordium | Samd10 | sterile alpł | Other | other |
| Gallbladder Primordium | Col4a1 | collagen ty | Extracellul | other |
| Gallbladder Primordium | Fis1 | fission, mit | Cytoplasm | other |
| Gallbladder Primordium | Flywch2 | FLYWCH fa | Other | other |
| Gallbladder Primordium | Atxn10 | ataxin 10 | Cytoplasm | other |
| Gallbladder Primordium | Hmgn2 | high mobi | Nucleus | other |
| Gallbladder Primordium | Polr2k | RNA polyr | Nucleus | enzyme |
| Gallbladder Primordium | Cbr1 | carbonyl r | Cytoplasm | enzyme |
| Gallbladder Primordium | Paip2 | poly(A) bir | Cytoplasm | translation regulator |
| Gallbladder Primordium | Tfb1m | transcripti | Cytoplasm | transcription regulato |
| Gallbladder Primordium | Cldn4 | claudin 4 | Plasma Me | transmembrane recep |
| Gallbladder Primordium | Rpp30 | ribonuclea | Nucleus | enzyme |
| Gallbladder Primordium | Krt19 | keratin 19 | Cytoplasm | other |
| Gallbladder Primordium | Akr7a5 | aldo-keto | Cytoplasm | enzyme |
| Gallbladder Primordium | Praf2 | PRA1 dom | Other | other |
| Gallbladder Primordium | Efna4 | ephrin A4 | Plasma Me | kinase |
| Gallbladder Primordium | Cited1 | Cbp/p300 | Nucleus | transcription regulato |
| Gallbladder Primordium | Sh3bgrl3 | SH3 doma | Nucleus | other |
| Liver Primordium | Myc | MYC proto | Nucleus | transcription regulato |
| Liver Primordium | Tgm2 | transglutaı | Cytoplasm | enzyme |
| Liver Primordium | Gm17511 | ribosomal | Other | other |
| Liver Primordium | Cdkn1c | cyclin-dep | Nucleus | other |
| Liver Primordium | Rps12-ps3 | ribosomal | Other | other |
| Liver Primordium | Lars2 | leucyl-tRN | Cytoplasm | enzyme |
| Liver Primordium | Id3 | inhibitor o | Nucleus | transcription regulato |
| Liver Primordium | Scd2 | stearoyl-C | Cytoplasm | enzyme |
| Liver Primordium | Id1 | inhibitor o | Nucleus | transcription regulato |
| Liver Primordium | Hsdl1 | hydroxyste | Cytoplasm | enzyme |
| Liver Primordium | Igf2 | insulin like | Extracellul | growth factor |
| Liver Primordium | Pcyt2 | phosphate | Cytoplasm | enzyme |
| Liver Primordium | Rps12 | ribosomal | Cytoplasm | other |
| Liver Primordium | Cald1 | caldesmor | Plasma Me | other |
| Liver Primordium | Phlda2 | pleckstrin | Cytoplasm | other |
| Liver Primordium | Gm10073 | ribosomal | Other | other |

| | | | | |
|---|---|---|---|---|
| Liver Primordium | Mvd | mevalonat | Cytoplasm | enzyme |
| Liver Primordium | Dusp9 | dual speci | Nucleus | phosphatase |
| Liver Primordium | Gm11808 | ubiquitin A | Other | other |
| Liver Primordium | Gm21887 | - | - | - |
| Liver Primordium | Sqle | squalene e | Cytoplasm | enzyme |
| Liver Primordium | Pcsk9 | proproteir | Extracellul | peptidase |
| Liver Primordium | Dlk1 | delta like r | Extracellul | other |
| Liver Primordium | Rps15 | ribosomal | Cytoplasm | other |
| Liver Primordium | Hhex | hematopo | Nucleus | transcription regulato |
| Liver Primordium | Crabp2 | cellular ret | Cytoplasm | transporter |
| Liver Primordium | Gap43 | growth ass | Plasma Me | other |
| Liver Bud | Apoa2 | apolipopro | Extracellul | transporter |
| Liver Bud | Apom | apolipopro | Plasma Me | transporter |
| Liver Bud | Apoe | apolipopro | Extracellul | transporter |
| Liver Bud | Serpina6 | serpin fam | Extracellul | other |
| Liver Bud | Rbp4 | retinol bin | Extracellul | other |
| Liver Bud | Trf | transferrin | Extracellul | transporter |
| Liver Bud | Apoc2 | apolipopro | Extracellul | transporter |
| Liver Bud | Serpinf2 | serpin fam | Extracellul | other |
| Liver Bud | Apoa1 | apolipopro | Extracellul | transporter |
| Liver Bud | Itih2 | inter-alpha | Extracellul | other |
| Liver Bud | Sepp1 | - | - | - |
| Liver Bud | Afp | alpha feto | Extracellul | transporter |
| Liver Bud | Kng1 | kininogen | Extracellul | other |
| Liver Bud | Gulo | gulonolact | Cytoplasm | enzyme |
| Liver Bud | Qprt | quinolinate | Cytoplasm | enzyme |
| Liver Bud | Serpind1 | serpin fam | Extracellul | other |
| Liver Bud | Fga | fibrinogen | Extracellul | other |
| Liver Bud | Ambp | alpha-1-m | Extracellul | transporter |
| Liver Bud | F12 | coagulatio | Extracellul | peptidase |
| Liver Bud | Fgb | fibrinogen | Extracellul | other |
| Liver Bud | Plg | plasminog | Extracellul | peptidase |
| Liver Bud | Glud1 | glutamate | Cytoplasm | enzyme |
| Liver Bud | Phgdh | phosphog | Cytoplasm | enzyme |
| Liver Bud | Lect2 | leukocyte | Extracellul | other |
| Liver Bud | Serpinc1 | serpin fam | Extracellul | enzyme |
| Liver Bud | Fgg | fibrinogen | Extracellul | other |
| Liver Bud | Srm | spermidine | Cytoplasm | enzyme |
| Liver Bud | Lpl | lipoproteir | Cytoplasm | enzyme |
| Liver Bud | Shbg | sex hormo | Extracellul | other |
| Liver Bud | Psat1 | phosphose | Cytoplasm | enzyme |
| Liver Bud | Soat2 | sterol O-a | Cytoplasm | enzyme |
| Liver Bud | B2m | beta-2-mi | Plasma Me | transmembrane recep |
| Liver Bud | Serpina1b | serpin fam | Extracellul | other |
| Liver Bud | Igfbp1 | insulin like | Extracellul | other |
| Liver Bud | Fabp5 | fatty acid b | Cytoplasm | transporter |
| Liver Bud | Fkbp11 | FK506 binc | Cytoplasm | enzyme |
| Liver Bud | Serpinf1 | serpin fam | Extracellul | other |
| Liver Bud | C1qbp | compleme | Cytoplasm | transcription regulato |
| Liver Bud | Lcat | lecithin-ch | Extracellul | enzyme |
| Liver Bud | Ttr | transthyret | Extracellul | transporter |
| Liver Bud | F10 | coagulatio | Extracellul | peptidase |
| Liver Bud | Herpud1 | homocyste | Cytoplasm | other |
| Liver Bud | Alb | albumin | Extracellul | transporter |
| Liver Bud | Cth | cystathion | Cytoplasm | enzyme |
| Liver Bud | Tmem97 | transmeml | Extracellul | other |
| Liver Bud | Gm5549 | - | - | - |
| Liver Bud | Rpl36al | ribosomal | Cytoplasm | other |

| | | | | |
|---|---|---|---|---|
| Liver Bud | Ddt | D-dopach | Cytoplasm | enzyme |
| Liver Bud | Nipsnap1 | nipsnap h | Cytoplasm | enzyme |
| Liver Bud | Serpina1a | serpin fam | Extracellul | other |
| Liver Bud | Psph | phosphos | Cytoplasm | phosphatase |
| Liver Bud | Tst | thiosulfate | Cytoplasm | enzyme |
| Liver Bud | Pla2g12b | phospholi | Extracellul | enzyme |
| Liver Bud | Osgin1 | oxidative s | Other | growth factor |
| Liver Bud | Hspe1 | heat shock | Cytoplasm | enzyme |
| Liver Bud | Gm10053 | cytochrom | Other | other |
| Liver Bud | F2 | coagulatio | Extracellul | peptidase |
| Liver Bud | Itm2b | integral m | Plasma Me | other |
| Liver Bud | Hspd1 | heat shock | Cytoplasm | enzyme |
| Liver Bud | Gm6484 | - | - | - |
| Liver Bud | Cpb2 | carboxype | Extracellul | peptidase |
| Liver Bud | Stard10 | StAR relat | Cytoplasm | other |
| Liver Bud | Ddit3 | DNA dama | Nucleus | transcription regulato |
| Liver Bud | Hspa9 | heat shock | Cytoplasm | other |
| Liver Bud | Gcat | glycine C- | Cytoplasm | enzyme |
| Liver Bud | Gchfr | GTP cycloh | Cytoplasm | other |
| Liver Bud | Cited1 | Cbp/p300 | Nucleus | transcription regulato |
| Liver Bud | Cox8a | cytochrom | Cytoplasm | enzyme |
| Liver Bud | Pcbd1 | pterin-4 al | Nucleus | transcription regulato |
| Liver Bud | Serpina1d | serpin fam | Extracellul | other |
| Liver Bud | Cyc1 | cytochrom | Cytoplasm | enzyme |
| Liver Bud | Gstm1 | glutathion | Cytoplasm | enzyme |
| Liver Bud | Dgat2 | diacylglyce | Cytoplasm | enzyme |
| Liver Bud | Igsf1 | immunogl | Plasma Me | other |
| Liver Bud | Hpx | hemopexi | Extracellul | transporter |
| Liver Bud | Phb2 | prohibitin | Cytoplasm | transcription regulato |
| Liver Bud | Rps27l | ribosomal | Cytoplasm | translation regulator |
| Liver Bud | Spp2 | secreted p | Extracellul | other |
| Liver Bud | Ddah1 | dimethylar | Cytoplasm | enzyme |
| Liver Bud | Mogat2 | monoacylg | Cytoplasm | enzyme |
| Liver Bud | Bco1 | beta-carot | Cytoplasm | enzyme |
| Liver Bud | Cycs | cytochrom | Cytoplasm | transporter |
| Liver Bud | Slc25a5 | solute carr | Cytoplasm | transporter |
| Liver Bud | Sdhd | succinate | Cytoplasm | enzyme |
| Liver Bud | A1cf | APOBEC1 | Nucleus | other |
| Liver Bud | C3 | compleme | Extracellul | peptidase |
| Liver Bud | Eif5a | eukaryotic | Cytoplasm | translation regulator |
| Liver Bud | Prdx1 | peroxired | Cytoplasm | enzyme |
| Liver Bud | Cyp2c68 | cytochrom | Cytoplasm | enzyme |
| Liver Bud | Acaa2 | acetyl-Co | Cytoplasm | enzyme |
| Liver Bud | Ncl | nucleolin | Nucleus | other |
| Liver Bud | Gstp1 | glutathion | Cytoplasm | enzyme |
| Liver Bud | Atp5c1 | ATP synth | Cytoplasm | transporter |
| Liver Bud | Gstp2 | glutathion | Cytoplasm | enzyme |
| Liver Bud | Asns | asparagin | Cytoplasm | enzyme |
| Liver Bud | Got2 | glutamic- | Cytoplasm | enzyme |
| Liver Bud | Reep6 | receptor a | Plasma Me | other |
| Liver Bud | Serpina1c | serpin fam | Extracellul | other |
| Liver Bud | Cacybp | calcyclin b | Nucleus | other |
| Liver Bud | Gdf15 | growth dif | Extracellul | growth factor |
| Liver Bud | Apoc1 | apolipopr | Extracellul | other |
| Liver Bud | Eif2s2 | eukaryotic | Cytoplasm | translation regulator |
| Liver Bud | Uqcrq | ubiquinol- | Cytoplasm | enzyme |
| Liver Bud | Gstz1 | glutathion | Cytoplasm | enzyme |
| Liver Bud | Junb | JunB proto | Nucleus | transcription regulato |

| | | | | |
|---|---|---|---|---|
| Liver Bud | Mttp | microsoma | Cytoplasm | transporter |
| Liver Bud | Ndufs2 | NADH:ubi | Cytoplasm | enzyme |
| Liver Bud | Ndufv2 | NADH:ubi | Cytoplasm | enzyme |
| Liver Bud | Emc6 | ER membr | Cytoplasm | other |
| Liver Bud | Ppib | peptidylpr | Cytoplasm | enzyme |
| Liver Bud | Ostc | oligosacch | Cytoplasm | enzyme |
| Liver Bud | Agmat | agmatinas | Cytoplasm | enzyme |
| Liver Bud | Ssr2 | signal seq | Cytoplasm | other |
| Liver Bud | Ssr4 | signal seq | Cytoplasm | other |
| Liver Bud | Vtn | vitronectin | Extracellul | other |
| Liver Bud | Sec61b | Sec61 tran | Cytoplasm | transporter |
| Liver Bud | Mrpl20 | mitochonc | Cytoplasm | other |
| Liver Bud | Shmt1 | serine hyd | Cytoplasm | enzyme |
| Liver Bud | Serpina10 | serpin fam | Extracellul | other |
| Liver Bud | Sgk1 | serum/glu | Cytoplasm | kinase |
| Liver Bud | Fth1 | ferritin hea | Cytoplasm | enzyme |
| Liver Bud | Gnpnat1 | glucosami | Cytoplasm | enzyme |
| Liver Bud | Rsl1d1 | ribosomal | Nucleus | other |
| Liver Bud | Nme1 | NME/NM2 | Cytoplasm | kinase |
| Liver Bud | Ran | RAN, mem | Nucleus | enzyme |
| Liver Bud | Serp1 | stress asso | Cytoplasm | other |
| Liver Bud | Bex2 | brain expr | Other | other |
| Liver Bud | Cnbp | CCHC-typ | Nucleus | transcription regulato |
| Liver Bud | Tmed2 | transmeml | Cytoplasm | transporter |
| Liver Bud | Tomm20 | translocase | Cytoplasm | transporter |
| Liver Bud | Tomm5 | translocase | Cytoplasm | other |
| Liver Bud | Lsm5 | LSM5 hom | Cytoplasm | other |
| Liver Bud | Tmem258 | transmeml | Other | other |
| Liver Bud | Ndufa6 | NADH:ubi | Cytoplasm | enzyme |
| Liver Bud | Aes | amino-ter | Nucleus | transcription regulato |
| Liver Bud | Uqcr10 | ubiquinol- | Cytoplasm | enzyme |
| Liver Bud | Nostrin | nitric oxide | Cytoplasm | transcription regulato |
| Liver Bud | Nhp2 | NHP2 ribo | Nucleus | other |
| Liver Bud | Uqcrfs1 | ubiquinol- | Cytoplasm | enzyme |
| Liver Bud | Dnajc3 | DnaJ heat | Cytoplasm | other |
| Liver Bud | Mtfp1 | mitochonc | Cytoplasm | other |
| Liver Bud | Hint1 | histidine tr | Nucleus | enzyme |
| Liver Bud | Manf | mesencep | Extracellul | other |
| Liver Bud | Idh3a | isocitrate c | Cytoplasm | enzyme |
| Liver Bud | Gpc3 | glypican 3 | Plasma Me | other |
| Liver Bud | Npm1 | nucleophc | Nucleus | transcription regulato |
| Liver Bud | Sec61g | Sec61 tran | Plasma Me | transporter |
| Liver Bud | Slc19a2 | solute carr | Plasma Me | transporter |
| Liver Bud | Galk1 | galactokin | Cytoplasm | kinase |
| Liver Bud | Fam195a | - | - | - |
| Liver Bud | Fst | follistatin | Extracellul | other |
| Liver Bud | Mrpl17 | mitochonc | Cytoplasm | other |
| Liver Bud | Slc25a15 | solute carr | Cytoplasm | transporter |
| Liver Bud | Naa50 | N(alpha)-a | Cytoplasm | enzyme |
| Liver Bud | Timm13 | translocase | Cytoplasm | transporter |
| Liver Bud | Nhp2l1 | - | - | - |
| Liver Bud | Gm9797 | translocase | Other | other |
| Liver Bud | Mydgf | myeloid de | Extracellul | cytokine |
| Liver Bud | Ass1 | argininosu | Cytoplasm | enzyme |
| Liver Bud | Nadk | NAD kinas | Cytoplasm | kinase |
| Liver Bud | Mrps28 | mitochonc | Cytoplasm | other |
| Liver Bud | Gstt1 | glutathion | Cytoplasm | enzyme |
| Liver Bud | Adk | adenosine | Nucleus | kinase |

| | | | | |
|---|---|---|---|---|
| Liver Bud | Cdk2ap2 | cyclin dep | Nucleus | other |
| Liver Bud | Nop10 | NOP10 rib | Nucleus | other |
| Liver Bud | Mrpl50 | mitochonc | Cytoplasm | other |
| Liver Bud | Vapa | VAMP ass | Plasma Me | other |
| Liver Bud | Gjb1 | gap juncti | Plasma Me | transporter |
| Liver Bud | Itih3 | inter-alph | Extracellul | other |
| Liver Bud | Timm8a1 | translocas | Cytoplasm | transporter |
| Liver Bud | Mrpl22 | mitochonc | Cytoplasm | other |
| Liver Bud | Cct8 | chaperoni | Cytoplasm | enzyme |
| Liver Bud | Rps11 | ribosomal | Cytoplasm | other |
| Liver Bud | Fxyd1 | FXYD dom | Plasma Me | ion channel |
| Liver Bud | Akr1c13 | aldo-keto | Other | enzyme |
| Liver Bud | Psmd7 | proteasom | Cytoplasm | other |
| Liver Bud | Gstm3 | glutathion | Cytoplasm | enzyme |
| Liver Bud | Mrpl16 | mitochonc | Cytoplasm | other |
| Liver Bud | Aadac | arylacetam | Cytoplasm | enzyme |
| Liver Bud | Uqcrb | ubiquinol- | Cytoplasm | enzyme |
| Liver Bud | Ptges3 | prostaglar | Cytoplasm | enzyme |
| Liver Bud | Gm5611 | nucleopho | Other | other |
| Liver Bud | Grhpr | glyoxylate | Cytoplasm | enzyme |
| Liver Bud | Etfa | electron tr | Cytoplasm | transporter |
| Liver Bud | Rpn1 | ribophorin | Cytoplasm | enzyme |
| Liver Bud | Uchl5 | ubiquitin C | Cytoplasm | peptidase |
| Liver Bud | Timm23 | translocas | Cytoplasm | transporter |
| Liver Bud | H2-Q7 | major hist | Plasma Me | other |
| Liver Bud | Emg1 | EMG1, N1 | Nucleus | enzyme |
| Liver Bud | Cfi | compleme | Extracellul | peptidase |
| Liver Bud | Sephs2 | selenopho | Cytoplasm | enzyme |
| Liver Bud | Gm10273 | translocas | Other | other |
| Liver Bud | Pla2g12a | phospholi | Other | enzyme |
| Liver Bud | Cox7b | cytochrom | Cytoplasm | enzyme |
| Liver Bud | Ndufa2 | NADH:ubi | Cytoplasm | enzyme |
| Liver Bud | Edf1 | endothelia | Nucleus | transcription regulato |
| Liver Bud | Mmd2 | monocyte | Other | kinase |
| Liver Bud | Pdia6 | protein dis | Cytoplasm | enzyme |
| Liver Bud | Etfb | electron tr | Cytoplasm | transporter |
| Liver Bud | Ndufb4 | NADH:ubi | Cytoplasm | transporter |
| Liver Bud | Cct6a | chaperoni | Cytoplasm | other |
| Liver Bud | Psmb10 | proteasom | Cytoplasm | peptidase |
| Liver Bud | Mthfd2 | methylene | Cytoplasm | enzyme |
| Liver Bud | Cd302 | CD302 mc | Plasma Me | transmembrane recep |
| Liver Bud | Brix1 | BRX1, biog | Nucleus | other |
| Liver Bud | Gm5616 | NHP2 non | Nucleus | other |
| Liver Bud | Vimp | - | - | - |
| Liver Bud | Rpf2 | ribosome | Nucleus | other |
| Liver Bud | Ccne1 | cyclin E1 | Nucleus | transcription regulato |
| Liver Bud | Psma5 | proteasom | Cytoplasm | peptidase |
| Liver Bud | Qdpr | quinoid di | Cytoplasm | enzyme |
| Liver Bud | Chchd4 | coiled-coi | Cytoplasm | enzyme |
| Liver Bud | Snx2 | sorting ne | Cytoplasm | transporter |
| Liver Bud | Mpzl1 | myelin prc | Plasma Me | other |
| Liver Bud | Gamt | guanidino | Cytoplasm | enzyme |
| Liver Bud | Sdhb | succinate | Cytoplasm | enzyme |
| Liver Bud | Tex30 | testis expr | Other | other |
| Liver Bud | Ldhb | lactate del | Cytoplasm | enzyme |
| Liver Bud | Tmem37 | transmeml | Plasma Me | ion channel |
| Liver Bud | Mdh2 | malate del | Cytoplasm | enzyme |
| Liver Bud | Igfbp4 | insulin like | Extracellul | other |

| | | | | |
|---|---|---|---|---|
| Liver Bud | Stt3a | STT3A, cat | Plasma Me | enzyme |
| Liver Bud | Nars | asparaginy | Cytoplasm | enzyme |
| Liver Bud | Scp2 | sterol carri | Cytoplasm | transporter |
| Liver Bud | Pbk | PDZ bindi | Cytoplasm | kinase |
| Liver Bud | Ndufb6 | NADH:ubi | Cytoplasm | enzyme |
| Liver Bud | Gcsh | glycine cle | Cytoplasm | enzyme |
| Liver Bud | Eci1 | enoyl-CoA | Cytoplasm | enzyme |
| Liver Bud | Sdf2l1 | stromal ce | Cytoplasm | other |
| Liver Bud | Cks2 | CDC28 pro | Other | kinase |
| Liver Bud | Sar1b | secretion a | Cytoplasm | enzyme |
| Liver Bud | Anp32b | acidic (leu | Nucleus | other |
| Liver Bud | Btg2 | BTG anti-p | Nucleus | transcription regulato |
| Liver Bud | Psmg1 | proteasom | Plasma Me | other |
| Liver Bud | Pold2 | DNA polyr | Nucleus | enzyme |
| Liver Bud | Ndufs8 | NADH:ubi | Cytoplasm | enzyme |
| Liver Bud | Fkbp1a | FK506 bin | Cytoplasm | enzyme |
| Liver Bud | Eif4ebp1 | eukaryotic | Cytoplasm | translation regulator |
| Liver Bud | Surf4 | surfeit 4 | Cytoplasm | other |
| Liver Bud | Mrpl12 | mitochond | Cytoplasm | other |
| Liver Bud | Mrps7 | mitochond | Cytoplasm | other |
| Liver Bud | Hspa5 | heat shock | Cytoplasm | enzyme |
| Liver Bud | Akr1c12 | aldo-keto | Other | enzyme |
| Liver Bud | Timm50 | translocas | Cytoplasm | phosphatase |
| Liver Bud | Rcl1 | RNA termi | Nucleus | enzyme |
| Liver Bud | Apof | apolipopro | Extracellul | transporter |
| Liver Bud | H2-Q6 | major hist | Plasma Me | other |
| Liver Bud | Xbp1 | X-box bin | Nucleus | transcription regulato |
| Liver Bud | Degs1 | delta 4-de | Plasma Me | enzyme |
| Liver Bud | Tspan12 | tetraspani | Plasma Me | other |
| Liver Bud | Mesdc2 | - | - | - |
| Liver Bud | Creld2 | cysteine ri | Other | other |
| Liver Bud | Eif2s1 | eukaryotic | Cytoplasm | translation regulator |
| Liver Bud | Aass | aminoadip | Cytoplasm | enzyme |
| Liver Bud | U2af1 | U2 small n | Nucleus | other |
| Liver Bud | Akr1c19 | aldo-keto | Other | other |
| Liver Bud | Dnajc12 | DnaJ heat | Cytoplasm | other |
| Liver Bud | Hprt | hypoxanth | Cytoplasm | enzyme |
| Liver Bud | Hmbs | hydroxym | Cytoplasm | enzyme |
| Liver Bud | Calr | calreticulir | Cytoplasm | transcription regulato |
| Liver Bud | Rpn2 | ribophorin | Cytoplasm | enzyme |
| Liver Bud | Egr1 | early grow | Nucleus | transcription regulato |
| Liver Bud | Bccip | BRCA2 an | Nucleus | other |
| Liver Bud | Prdx3 | peroxired | Cytoplasm | enzyme |
| Liver Bud | Sigmar1 | sigma non | Plasma Me | transmembrane recep |
| Liver Bud | Dctpp1 | dCTP pyro | Cytoplasm | enzyme |
| Liver Bud | Ppp1r15a | protein ph | Cytoplasm | other |
| Liver Bud | Tceb1 | - | - | - |
| Liver Bud | Rnf128 | ring finger | Cytoplasm | enzyme |
| Liver Bud | Pno1 | partner of | Nucleus | other |
| Liver Bud | Fos | Fos proto- | Nucleus | transcription regulato |
| Liver Bud | Dnajc22 | DnaJ heat | Other | other |
| Liver Bud | Eif1a | eukaryotic | Cytoplasm | translation regulator |
| Liver Bud | Spcs1 | signal pep | Cytoplasm | peptidase |
| Liver Bud | Mrpl10 | mitochond | Cytoplasm | other |
| Liver Bud | Gm10094 | - | - | - |
| Liver Bud | Hsp90b1 | heat shock | Cytoplasm | other |
| Liver Bud | Nifk | nucleolar p | Nucleus | other |
| Liver Bud | Exosc7 | exosome c | Nucleus | enzyme |

| | | | | | |
|---|---|---|---|---|---|
| Liver Bud | | Spc24 | SPC24, NE | Cytoplasm | other |
| Liver Bud | | Lpcat3 | lysophosp | Plasma Me | enzyme |
| Liver Bud | | Pa2g4 | proliferatic | Nucleus | transcription regulato |
| Liver Bud | | Slc25a39 | solute carr | Cytoplasm | other |
| Liver Bud | | Atf3 | activating | Nucleus | transcription regulato |
| Liver Bud | | Tuba4a | tubulin alp | Cytoplasm | other |
| Liver Bud | | Rexo2 | RNA exonu | Cytoplasm | enzyme |
| Liver Bud | | Mrpl42 | mitochonc | Cytoplasm | other |
| Liver Bud | | Ephx2 | epoxide hy | Cytoplasm | enzyme |
| Liver Bud | | Atp6v0c | ATPase H+ | Cytoplasm | transporter |
| Liver Bud | | Acadm | acyl-CoA c | Cytoplasm | enzyme |
| Liver Bud | | Farsb | phenylalar | Cytoplasm | enzyme |
| Liver Bud | | Cks1b | CDC28 prc | Other | kinase |
| Liver Bud | | Pdia3 | protein dis | Cytoplasm | peptidase |
| Liver Bud | | Lyar | Ly1 antibo | Plasma Me | other |
| Liver Bud | | Cops6 | COP9 sign | Nucleus | other |
| Liver Bud | | Atp5a1 | ATP synth | Cytoplasm | transporter |
| Liver Bud | | Timm10 | translocas | Cytoplasm | transporter |
| Liver Bud | | Gsn | gelsolin | Extracellul | other |
| Liver Bud | | Erp44 | endoplasm | Cytoplasm | enzyme |
| Liver Bud | | Atp5g1 | ATP synth | Cytoplasm | transporter |
| Liver Bud | | Slc35b1 | solute carr | Cytoplasm | transporter |
| Liver Bud | | Nudcd2 | NudC dom | Cytoplasm | other |
| Liver Bud | | Sc5d | sterol-C5- | Cytoplasm | enzyme |
| Liver Bud | | Ppa1 | pyrophosp | Cytoplasm | enzyme |
| Liver Bud | | Cinp | cyclin dep | Nucleus | other |
| Liver Bud | | Jagn1 | jagunal hc | Cytoplasm | other |
| Liver Bud | | Nop16 | NOP16 nu | Nucleus | other |
| Liver Bud | | Nfkbia | NFKB inhib | Cytoplasm | transcription regulato |
| Liver Bud | | Tmem176a | transmeml | Other | other |
| Liver Bud | | Lap3 | leucine am | Cytoplasm | peptidase |
| Liver Bud | | Pcsk9 | proprotein | Extracellul | peptidase |
| Liver Bud | | Cops7a | COP9 sign | Cytoplasm | other |
| Liver Bud | | Mak16 | MAK16 ho | Nucleus | other |
| Liver Bud | | Dap | death asso | Cytoplasm | transcription regulato |
| Liver Bud | | Clpp | caseinolyti | Cytoplasm | peptidase |
| Liver Bud | | Ccdc58 | coiled-coi | Cytoplasm | other |
| Liver Bud | | Cyb5a | cytochrom | Cytoplasm | enzyme |
| Liver Bud | | Tmem70 | transmeml | Cytoplasm | other |
| Liver Bud | | Jun | Jun proto- | Nucleus | transcription regulato |
| Liver Bud | | Slmo2 | - | - | - |
| Liver Bud | | Shmt2 | serine hyd | Cytoplasm | enzyme |
| Liver Bud | | Tomm22 | translocas | Cytoplasm | transporter |
| Liver Bud | | Tmem147 | transmeml | Cytoplasm | other |
| Liver Bud | | Mt1 | metallothi | Cytoplasm | other |
| Liver Bud | | Gm10146 | LSM7 hom | Other | other |
| Liver Bud | | Ndufaf2 | NADH:ubi | Cytoplasm | enzyme |
| Liver Bud | | Gstm4 | glutathion | Cytoplasm | enzyme |
| Liver Bud | | Pdia5 | protein dis | Cytoplasm | enzyme |
| Liver Bud | | Tagln2 | transgelin | Cytoplasm | other |
| Liver Bud | | Mrpl39 | mitochonc | Cytoplasm | other |
| Liver Bud | | Set | SET nucle | Nucleus | phosphatase |
| Liver Bud | | Tmem41a | transmeml | Other | other |
| Liver Bud | | Tars | threonyl-t | Nucleus | enzyme |
| Liver Bud | | Sub1 | SUB1 hom | Nucleus | transcription regulato |
| Liver Bud | | Cnih1 | cornichon | Plasma Me | other |
| Liver Bud | | Mrps31 | mitochonc | Cytoplasm | other |
| Liver Bud | | Lsm7 | LSM7 hom | Nucleus | other |

| | | | | |
|---|---|---|---|---|
| Liver Bud | Proc | protein C, | Extracellul | peptidase |
| Liver Bud | Psmd6 | proteasom | Cytoplasm | enzyme |
| Liver Bud | Tubb6 | tubulin be | Cytoplasm | other |
| Liver Bud | Uchl3 | ubiquitin C | Cytoplasm | peptidase |
| Liver Bud | Eef1e1 | eukaryotic | Cytoplasm | translation regulator |
| Liver Bud | Smdt1 | single-pas | Extracellul | other |
| Liver Bud | Ndufa5 | NADH:ubi | Cytoplasm | enzyme |
| Liver Bud | Mrps16 | mitochonc | Cytoplasm | other |
| Liver Bud | Fabp1 | fatty acid b | Cytoplasm | transporter |
| Liver Bud | Spink3 | - | - | - |
| Liver Bud | Uqcr11 | ubiquinol- | Cytoplasm | enzyme |
| Liver Bud | Ranbp1 | RAN bindi | Nucleus | other |
| Liver Bud | Fkbp2 | FK506 binc | Cytoplasm | enzyme |
| Liver Bud | Tmem150a | transmeml | Plasma Me | other |
| Liver Bud | Mrpl36 | mitochonc | Cytoplasm | other |
| Liver Bud | Dnajb11 | DnaJ heat | Cytoplasm | other |
| Liver Bud | A430005L1 | chromosol | Nucleus | other |
| Liver Bud | Mrpl32 | mitochonc | Cytoplasm | translation regulator |
| Liver Bud | Dkc1 | dyskerin p | Nucleus | enzyme |
| Liver Bud | Tmem176b | transmeml | Other | other |
| Liver Bud | Bckdhb | branched | Cytoplasm | enzyme |
| Liver Bud | Dbi | diazepam | Cytoplasm | other |
| Liver Bud | Slc35b2 | solute carr | Cytoplasm | transporter |
| Liver Bud | Fbl | fibrillarin | Nucleus | enzyme |
| Liver Bud | Tnfrsf12a | TNF recep | Plasma Me | transmembrane recep |
| Liver Bud | Ndufb10 | NADH:ubi | Cytoplasm | enzyme |
| Liver Bud | Gemin6 | gem nucle | Nucleus | other |
| Liver Bud | Spcs2 | signal pep | Cytoplasm | other |
| Liver Bud | Deb1 | - | - | - |
| Liver Bud | Vdac1 | voltage de | Cytoplasm | ion channel |
| Liver Bud | Cdk1 | cyclin dep | Nucleus | kinase |
| Liver Bud | Rogdi | rogdi hom | Nucleus | other |
| Liver Bud | H2afx | H2A histor | Nucleus | transcription regulato |
| Liver Bud | Atp5b | ATP synth | Cytoplasm | transporter |
| Liver Bud | Dnajb9 | DnaJ heat | Nucleus | other |
| Liver Bud | Gm5160 | peptidylpr | Other | other |
| Liver Bud | Ccnb1 | cyclin B1 | Cytoplasm | kinase |
| Liver Bud | Lyrm4 | LYR motif | Cytoplasm | other |
| Liver Bud | Cirh1a | - | - | - |
| Liver Bud | Cmss1 | cms1 ribos | Other | other |
| Liver Bud | Nup37 | nucleopor | Nucleus | transporter |
| Liver Bud | Ubc | ubiquitin C | Cytoplasm | enzyme |
| Liver Bud | Kdelr2 | KDEL endc | Cytoplasm | other |
| Liver Bud | Mrpl33 | mitochonc | Cytoplasm | other |
| Liver Bud | Atf4 | activating | Nucleus | transcription regulato |
| Liver Bud | Dnaja2 | DnaJ heat | Nucleus | enzyme |
| Liver Bud | Usmg5 | ATP synth | Cytoplasm | other |
| Liver Bud | Pnliprp1 | pancreatic | Extracellul | enzyme |
| Liver Bud | Afm | afamin | Extracellul | transporter |
| Liver Bud | Ube2c | ubiquitin c | Cytoplasm | enzyme |
| Liver Bud | Klf6 | Kruppel lik | Nucleus | transcription regulato |
| Liver Bud | Fam107b | family with | Other | other |
| Liver Bud | Aurka | aurora kin | Nucleus | kinase |
| Liver Bud | Bub3 | BUB3, mitc | Nucleus | other |
| Liver Bud | Gm8186 | small nucl | Other | other |
| Liver Bud | Snrpg | small nucl | Nucleus | other |
| Liver Bud | Gm10184 | karyopheri | Other | other |
| Liver Bud | Zfp36 | ZFP36 ring | Nucleus | transcription regulato |

| | | | | | |
|---|---|---|---|---|---|
| Liver Bud | Kpna2 | karyopheri | Nucleus | transporter | |
| Liver Bud | Sqstm1 | sequestos | Cytoplasm | transcription | regulato |
| Liver Bud | Ier2 | immediate | Cytoplasm | transcription | regulato |
| Liver Bud | Mt2 | metallothi | Other | other | |

ıclear receptor

ıclear receptor

ıclear receptor

ıclear receptor

ıclear receptor

ıclear receptor

ıclear receptor

### Differentially expressed genes (DEGs) between gallbladde

- lb:lower bound, mle:maximum likelihood estimate, ub: upper bound of the
- ce: conservative estimate of expression-fold change (equals to the min(abs(
- Z: uncorrected Z-score of expression difference
- cZ: expression difference Z-score corrected for multiple hypothesis testing u
- **Genes highly expressed in the GBP presented positive value of ce, Z anc**

| | lb | mle | ub | ce | Z | cZ | GeneNam |
|---|---|---|---|---|---|---|---|
| ENSMUSG00000067276 | -7.61901 | -4.5714 | -3.55554 | -3.55554 | -7.16081 | -5.92167 | Capn6 |
| ENSMUSG00000035451 | -7.43974 | -6.69277 | -5.37812 | -5.37812 | -7.05797 | -5.82564 | Foxa1 |
| ENSMUSG00000032366 | -3.58542 | -2.6293 | -1.76283 | -1.76283 | -6.50024 | -5.24147 | Tpm1 |
| ENSMUSG00000048387 | -7.7684 | -6.84217 | -5.25861 | -5.25861 | -6.1691 | -4.87437 | Osr1 |
| ENSMUSG00000075394 | -6.69277 | -5.8263 | -4.87019 | -4.87019 | -6.17257 | -4.87437 | Hoxc4 |
| ENSMUSG00000039911 | -5.19885 | -4.36225 | -3.37627 | -3.37627 | -6.00054 | -4.68808 | Spsb1 |
| ENSMUSG00000005220 | -6.09521 | -5.01958 | -3.94396 | -3.94396 | -5.9722 | -4.66408 | Corin |
| ENSMUSG00000043924 | -5.07934 | -4.18298 | -3.16712 | -3.16712 | -5.89984 | -4.59454 | Ncmap |
| ENSMUSG00000048450 | -6.12508 | -4.54153 | -3.31651 | -3.31651 | -5.88822 | -4.59004 | Msx1 |
| ENSMUSG00000006538 | -5.8263 | -5.04946 | -4.15311 | -4.15311 | -5.85026 | -4.56116 | Ihh |
| ENSMUSG00000026639 | -6.60314 | -5.8263 | -4.60128 | -4.60128 | -5.85629 | -4.56116 | Lamb3 |
| ENSMUSG00000042258 | -6.60314 | -5.49764 | -4.48177 | -4.48177 | -5.82812 | -4.54249 | Isl1 |
| ENSMUSG00000079445 | -6.09521 | -5.16897 | -4.09335 | -4.09335 | -5.79271 | -4.52237 | B3gnt7 |
| ENSMUSG00000001496 | -6.2446 | -4.51165 | -3.79456 | -3.79456 | -5.71638 | -4.46077 | Nkx2-1 |
| ENSMUSG00000025921 | -7.23059 | -6.2446 | -4.90007 | -4.90007 | -5.71469 | -4.46077 | Rdh10 |
| ENSMUSG00000037621 | -5.19885 | -4.39213 | -3.34639 | -3.34639 | -5.71249 | -4.46077 | Atoh8 |
| ENSMUSG00000046157 | -4.90007 | -3.97383 | -2.74882 | -2.74882 | -5.71278 | -4.46077 | Tmem229l |
| ENSMUSG00000057069 | -6.78241 | -5.52751 | -4.48177 | -4.48177 | -5.70989 | -4.46077 | Ero1lb |
| ENSMUSG00000066026 | -6.60314 | -5.85618 | -4.39213 | -4.39213 | -5.72447 | -4.46077 | Dhrs3 |
| ENSMUSG00000075707 | -5.46776 | -4.63116 | -3.55554 | -3.55554 | -5.69016 | -4.44233 | Dio3 |
| ENSMUSG00000026765 | -6.03545 | -4.87019 | -3.82444 | -3.82444 | -5.67717 | -4.43221 | Lypd6b |
| ENSMUSG00000028023 | -7.11107 | -5.43788 | -4.54153 | -4.54153 | -5.65901 | -4.42104 | Pitx2 |
| ENSMUSG00000090439 | -4.90007 | -3.97383 | -2.7787 | -2.7787 | -5.65877 | -4.42104 | Gm17455 |
| ENSMUSG00000019909 | -5.37812 | -4.27262 | -3.22687 | -3.22687 | -5.65078 | -4.41674 | Fam162b |
| ENSMUSG00000030200 | -4.9897 | -4.09335 | -3.25675 | -3.25675 | -5.62225 | -4.38645 | Bcl2l14 |
| ENSMUSG00000022485 | -5.07934 | -4.18298 | -3.0476 | -3.0476 | -5.58482 | -4.35492 | Hoxc5 |
| ENSMUSG00000024485 | -4.9897 | -4.06347 | -3.197 | -3.197 | -5.58433 | -4.35492 | Slc4a9 |
| ENSMUSG00000050052 | -5.79642 | -5.01958 | -3.67505 | -3.67505 | -5.58135 | -4.35492 | Tdrp |
| ENSMUSG00000031621 | -5.73666 | -4.7208 | -3.79456 | -3.79456 | -5.53672 | -4.30502 | Isx |
| ENSMUSG00000084174 | -5.55739 | -4.45189 | -3.197 | -3.197 | -5.51826 | -4.28667 | Sycn |
| ENSMUSG00000031028 | -5.37812 | -4.48177 | -3.55554 | -3.55554 | -5.50506 | -4.27623 | Tub |
| ENSMUSG00000031952 | -4.92995 | -3.97383 | -2.89821 | -2.89821 | -5.50235 | -4.27623 | Chst5 |
| ENSMUSG00000026399 | -5.8263 | -4.66104 | -3.55554 | -3.55554 | -5.48523 | -4.25928 | Cd55 |
| ENSMUSG00000027004 | -4.81043 | -3.97383 | -2.83845 | -2.83845 | -5.47989 | -4.25715 | Frzb |
| ENSMUSG00000024897 | -5.07934 | -3.85432 | -2.7787 | -2.7787 | -5.43836 | -4.21358 | Apba1 |
| ENSMUSG00000007021 | -5.13909 | -4.12323 | -3.16712 | -3.16712 | -5.42982 | -4.20715 | Syngr3 |
| ENSMUSG00000001288 | -5.49764 | -4.69092 | -3.55554 | -3.55554 | -5.42317 | -4.20303 | Rarg |
| ENSMUSG00000029646 | -5.34824 | -4.42201 | -3.52566 | -3.52566 | -5.40621 | -4.18995 | Cdx2 |
| ENSMUSG00000060260 | -5.76654 | -4.87019 | -3.76469 | -3.76469 | -5.40741 | -4.18995 | Pwwp2b |
| ENSMUSG00000033278 | -5.34824 | -4.33238 | -3.16712 | -3.16712 | -5.40086 | -4.18727 | Ptprm |
| ENSMUSG00000025491 | -6.03545 | -2.68906 | -1.67319 | -1.67319 | -5.3902 | -4.1823 | Ifitm1 |
| ENSMUSG00000026986 | -4.5714 | -3.67505 | -2.27076 | -2.27076 | -5.38622 | -4.1823 | Hnmt |
| ENSMUSG00000039114 | -4.9897 | -4.27262 | -3.28663 | -3.28663 | -5.38453 | -4.1823 | Nrn1 |
| ENSMUSG00000040624 | -5.25861 | -4.15311 | -3.01772 | -3.01772 | -5.38821 | -4.1823 | Plekhg1 |
| ENSMUSG00000049929 | -5.19885 | -4.39213 | -3.25675 | -3.25675 | -5.3768 | -4.1774 | Lpar4 |
| ENSMUSG00000054667 | -5.61715 | -4.90007 | -3.64517 | -3.64517 | -5.37056 | -4.17575 | Irs4 |
| ENSMUSG00000102742 | -5.16897 | -3.8842 | -2.92809 | -2.92809 | -5.36594 | -4.17349 | Pcdhga11 |

| | | | | | | | |
|---|---|---|---|---|---|---|---|
| ENSMUSG00000036568 | -5.22873 | -4.39213 | -3.40614 | -3.40614 | -5.35413 | -4.16213 | Gltscr1l |
| ENSMUSG00000024713 | -5.79642 | -4.81043 | -3.55554 | -3.55554 | -5.34649 | -4.15941 | Pcsk5 |
| ENSMUSG00000029484 | -5.37812 | -4.60128 | -3.52566 | -3.52566 | -5.34664 | -4.15941 | Anxa3 |
| ENSMUSG00000028517 | -5.91593 | -5.04946 | -3.94396 | -3.94396 | -5.34238 | -4.15761 | Ppap2b |
| ENSMUSG00000019851 | -5.97569 | -4.92995 | -3.97383 | -3.97383 | -5.33174 | -4.14753 | Perp |
| ENSMUSG00000019987 | -5.43788 | -4.33238 | -2.33052 | -2.33052 | -5.30552 | -4.11772 | Arg1 |
| ENSMUSG00000038175 | -5.8263 | -4.78055 | -3.85432 | -3.85432 | -5.2981 | -4.11163 | Mylip |
| ENSMUSG00000034127 | -5.34824 | -4.5714 | -3.55554 | -3.55554 | -5.28327 | -4.0961 | Tspan8 |
| ENSMUSG00000049686 | -4.78055 | -3.91408 | -2.92809 | -2.92809 | -5.2783 | -4.09301 | Orai1 |
| ENSMUSG00000030353 | -5.52751 | -4.48177 | -3.197 | -3.197 | -5.27018 | -4.0859 | Tead4 |
| ENSMUSG00000005148 | -6.48363 | -5.43788 | -4.27262 | -4.27262 | -5.25301 | -4.0673 | Klf5 |
| ENSMUSG00000019989 | -5.34824 | -4.33238 | -3.43602 | -3.43602 | -5.24459 | -4.05972 | Enpp3 |
| ENSMUSG00000050737 | -4.95982 | -3.94396 | -2.3604 | -2.3604 | -5.23466 | -4.0502 | Ptges |
| ENSMUSG00000039904 | -5.04946 | -4.06347 | -2.83845 | -2.83845 | -5.21218 | -4.03069 | Gpr37 |
| ENSMUSG00000048612 | -5.64703 | -4.69092 | -3.67505 | -3.67505 | -5.20939 | -4.03007 | Myof |
| ENSMUSG00000009772 | -5.67691 | -4.78055 | -3.58542 | -3.58542 | -5.20169 | -4.02887 | Nuak2 |
| ENSMUSG00000038151 | -6.2446 | -5.13909 | -4.03359 | -4.03359 | -5.20327 | -4.02887 | Prdm1 |
| ENSMUSG00000050994 | -4.87019 | -3.94396 | -2.03174 | -2.03174 | -5.18958 | -4.01633 | Adgb |
| ENSMUSG00000022061 | -4.95982 | -3.97383 | -2.7787 | -2.7787 | -5.18352 | -4.01415 | Nkx3-1 |
| ENSMUSG00000022240 | -4.69092 | -3.82444 | -2.45003 | -2.45003 | -5.18518 | -4.01415 | Ctnnd2 |
| ENSMUSG00000016262 | -4.60128 | -3.64517 | -1.67319 | -1.67319 | -5.17797 | -4.00982 | Sertad4 |
| ENSMUSG00000028128 | -5.34824 | -4.39213 | -3.197 | -3.197 | -5.17571 | -4.00962 | F3 |
| ENSMUSG00000003153 | -3.97383 | -2.59943 | -1.49392 | -1.49392 | -5.1489 | -3.98096 | Slc2a3 |
| ENSMUSG00000036941 | -5.22873 | -4.33238 | -3.22687 | -3.22687 | -5.15048 | -3.98096 | Elac1 |
| ENSMUSG00000023039 | -3.64517 | -2.30064 | -1.40429 | -1.40429 | -5.14454 | -3.97802 | Krt7 |
| ENSMUSG00000021506 | -4.45189 | -3.67505 | -2.24088 | -2.24088 | -5.1384 | -3.97282 | Pitx1 |
| ENSMUSG00000103677 | -4.60128 | -3.52566 | -1.79271 | -1.79271 | -5.12456 | -3.96036 | Pcdhga4 |
| ENSMUSG00000044313 | -4.81043 | -3.94396 | -2.45003 | -2.45003 | -5.10526 | -3.93838 | Mab21l3 |
| ENSMUSG00000024206 | -5.16897 | -4.24274 | -3.13724 | -3.13724 | -5.09788 | -3.93151 | Rfx2 |
| ENSMUSG00000000792 | -5.70679 | -4.7208 | -3.76469 | -3.76469 | -5.0721 | -3.90611 | Slc5a5 |
| ENSMUSG00000057614 | -4.81043 | -4.00371 | -2.71894 | -2.71894 | -5.07267 | -3.90611 | Gnai1 |
| ENSMUSG00000046733 | -6.6629 | -5.8263 | -4.06347 | -4.06347 | -5.03331 | -3.86393 | Gprc5a |
| ENSMUSG00000032179 | -4.84031 | -4.06347 | -2.53967 | -2.53967 | -5.02572 | -3.86357 | Bmp5 |
| ENSMUSG00000038094 | -4.9897 | -4.00371 | -2.71894 | -2.71894 | -5.02763 | -3.86357 | Atp13a4 |
| ENSMUSG00000046223 | -4.54153 | -3.70493 | -2.53967 | -2.53967 | -5.02652 | -3.86357 | Plaur |
| ENSMUSG00000042190 | -4.45189 | -3.49578 | -2.27076 | -2.27076 | -4.99405 | -3.82777 | Cmklr1 |
| ENSMUSG00000020017 | -5.28849 | -4.03359 | -2.56955 | -2.56955 | -4.99031 | -3.82524 | Hal |
| ENSMUSG00000023886 | -5.8263 | -4.39213 | -3.31651 | -3.31651 | -4.98429 | -3.82079 | Smoc2 |
| ENSMUSG00000057615 | -4.92995 | -4.00371 | -2.98785 | -2.98785 | -4.98333 | -3.82079 | Ldoc1 |
| ENSMUSG00000026124 | -3.73481 | -2.59943 | -1.31465 | -1.31465 | -4.91674 | -3.74001 | Cfc1 |
| ENSMUSG00000047497 | -4.63116 | -3.64517 | -2.47991 | -2.47991 | -4.91323 | -3.7377 | Adamts12 |
| ENSMUSG00000059674 | -6.18484 | -5.19885 | -3.79456 | -3.79456 | -4.88594 | -3.70482 | Cdh24 |
| ENSMUSG00000031093 | -5.34824 | -4.3025 | -3.16712 | -3.16712 | -4.8842 | -3.70476 | Dock11 |
| ENSMUSG00000030256 | -4.9897 | -3.8842 | -2.6293 | -2.6293 | -4.85235 | -3.67019 | Bhlhe41 |
| ENSMUSG00000074505 | -5.408 | -4.54153 | -2.98785 | -2.98785 | -4.84156 | -3.66211 | Fat3 |
| ENSMUSG00000074637 | -4.66104 | -3.67505 | -1.91222 | -1.91222 | -4.83958 | -3.66211 | Sox2 |
| ENSMUSG00000075297 | -4.45189 | -2.98785 | -1.34453 | -1.34453 | -4.8407 | -3.66211 | H60b |
| ENSMUSG00000020427 | -5.61715 | -4.27262 | -3.25675 | -3.25675 | -4.81877 | -3.63733 | Igfbp3 |
| ENSMUSG00000025608 | -3.0476 | -2.00186 | -1.1055 | -1.1055 | -4.79918 | -3.61407 | Podxl |
| ENSMUSG00000039239 | -6.36411 | -4.78055 | -3.70493 | -3.70493 | -4.74838 | -3.55637 | Tgfb2 |
| ENSMUSG00000042428 | -5.408 | -4.3025 | -3.13724 | -3.13724 | -4.75053 | -3.55637 | Mgat3 |
| ENSMUSG00000055963 | -4.81043 | -3.8842 | -2.12137 | -2.12137 | -4.72653 | -3.52996 | Triqk |
| ENSMUSG00000006731 | -5.46776 | -4.09335 | -3.13724 | -3.13724 | -4.67855 | -3.47135 | B4galnt1 |
| ENSMUSG00000032537 | -4.36225 | -3.43602 | -1.91222 | -1.91222 | -4.67669 | -3.47115 | Ephb1 |
| ENSMUSG00000041801 | -5.01958 | -4.06347 | -2.80858 | -2.80858 | -4.67534 | -3.47115 | Phlda3 |
| ENSMUSG00000029121 | -4.84031 | -4.00371 | -2.53967 | -2.53967 | -4.52165 | -3.28217 | Crmp1 |
| ENSMUSG00000079560 | -4.21286 | -2.95797 | -1.67319 | -1.67319 | -4.51851 | -3.27996 | Hoxa3 |
| ENSMUSG00000019558 | -5.01958 | -4.21286 | -2.71894 | -2.71894 | -4.47538 | -3.23022 | Slc6a8 |

| | | | | | | | |
|---|---|---|---|---|---|---|---|
| ENSMUSG00000058153 | -4.78055 | -4.03359 | -1.34453 | -1.34453 | -4.43144 | -3.17318 | Sez6l |
| ENSMUSG00000024172 | -5.22873 | -4.15311 | -2.80858 | -2.80858 | -4.40037 | -3.13518 | St6gal2 |
| ENSMUSG00000025666 | -6.9318 | -6.42387 | -1.55368 | -1.55368 | -4.38888 | -3.12354 | Tmem47 |
| ENSMUSG00000029563 | -5.67691 | -4.69092 | -3.13724 | -3.13724 | -4.3856 | -3.12103 | Foxp2 |
| ENSMUSG00000031665 | -5.28849 | -4.42201 | -2.56955 | -2.56955 | -4.36996 | -3.10177 | Sall1 |
| ENSMUSG00000042680 | -4.95982 | -4.12323 | -1.85246 | -1.85246 | -4.36216 | -3.09311 | Garem |
| ENSMUSG00000001435 | -3.13724 | -2.09149 | -1.1055 | -1.1055 | -4.3512 | -3.08205 | Col18a1 |
| ENSMUSG00000002688 | -5.04946 | -4.09335 | -2.09149 | -2.09149 | -4.34352 | -3.0735 | Prkd1 |
| ENSMUSG00000044186 | -6.27448 | -5.13909 | -3.34639 | -3.34639 | -4.32902 | -3.05756 | Nkx2-6 |
| ENSMUSG00000019647 | -7.70864 | -2.98785 | -1.70307 | -1.70307 | -4.32647 | -3.05595 | Sema6a |
| ENSMUSG00000019900 | -7.46961 | -6.6629 | -1.67319 | -1.67319 | -4.32266 | -3.05262 | Rfx6 |
| ENSMUSG00000022892 | -3.8842 | -2.27076 | -1.16526 | -1.16526 | -4.30403 | -3.03094 | App |
| ENSMUSG00000024691 | -5.07934 | -4.36225 | -2.56955 | -2.56955 | -4.3016 | -3.02945 | Fam111a |
| ENSMUSG00000067219 | -4.87019 | -4.00371 | -1.49392 | -1.49392 | -4.29259 | -3.019 | Nipal1 |
| ENSMUSG00000029223 | -3.10736 | -1.91222 | -1.01587 | -1.01587 | -4.28223 | -3.00848 | Uchl1 |
| ENSMUSG00000033276 | -5.67691 | -4.78055 | -3.31651 | -3.31651 | -4.27247 | -2.99696 | Stk36 |
| ENSMUSG00000021678 | -4.36225 | -3.55554 | -1.9421 | -1.9421 | -4.26482 | -2.98828 | F2rl1 |
| ENSMUSG00000073402 | -5.37812 | -4.48177 | -2.56955 | -2.56955 | -4.22798 | -2.94312 | Gm8909 |
| ENSMUSG00000019872 | -5.28849 | -4.33238 | -3.0476 | -3.0476 | -4.21986 | -2.93727 | Smpdl3a |
| ENSMUSG00000078185 | -4.9897 | -3.94396 | -2.65918 | -2.65918 | -4.22099 | -2.93727 | Chml |
| ENSMUSG00000032816 | -5.13909 | -4.12323 | -2.47991 | -2.47991 | -4.20974 | -2.92721 | Igdcc4 |
| ENSMUSG00000034382 | -4.48177 | -3.34639 | -2.09149 | -2.09149 | -4.20873 | -2.92721 | AI661453 |
| ENSMUSG00000030340 | -5.55739 | -4.60128 | -2.95797 | -2.95797 | -4.1639 | -2.87397 | Scnn1a |
| ENSMUSG00000044674 | -5.52751 | -4.63116 | -3.0476 | -3.0476 | -4.15985 | -2.87009 | Fzd1 |
| ENSMUSG00000036390 | -5.13909 | -4.18298 | -2.59943 | -2.59943 | -4.15576 | -2.86613 | Gadd45a |
| ENSMUSG00000058145 | -4.75067 | -3.91408 | -1.9421 | -1.9421 | -4.12834 | -2.82977 | Adamts17 |
| ENSMUSG00000031937 | -4.95982 | -4.18298 | -1.04575 | -1.04575 | -4.11384 | -2.81129 | Vstm5 |
| ENSMUSG00000027007 | -4.9897 | -4.03359 | -2.68906 | -2.68906 | -4.11105 | -2.80911 | Ssfa2 |
| ENSMUSG00000021614 | -3.13724 | -2.03174 | -1.04575 | -1.04575 | -4.09868 | -2.79354 | Vcan |
| ENSMUSG00000019852 | -4.95982 | -4.15311 | -2.71894 | -2.71894 | -4.09569 | -2.79106 | Arfgef3 |
| ENSMUSG00000074873 | -4.63116 | -3.73481 | -2.03174 | -2.03174 | -4.08848 | -2.78268 | AI606181 |
| ENSMUSG00000002633 | -7.3501 | -6.72265 | -1.55368 | -1.55368 | -4.07809 | -2.77319 | Shh |
| ENSMUSG00000064247 | -4.69092 | -3.82444 | -2.06161 | -2.06161 | -4.02262 | -2.70029 | Plcxd1 |
| ENSMUSG00000020609 | -6.2446 | -4.18298 | -2.50979 | -2.50979 | -4.01596 | -2.69415 | Apob |
| ENSMUSG00000026932 | -4.9897 | -3.82444 | -2.47991 | -2.47991 | -4.01418 | -2.69329 | Nacc2 |
| ENSMUSG00000001504 | -4.84031 | -4.00371 | -1.04575 | -1.04575 | -3.97977 | -2.6495 | Irx2 |
| ENSMUSG00000022598 | -5.22873 | -4.60128 | -1.01587 | -1.01587 | -3.96398 | -2.62868 | Psca |
| ENSMUSG00000028889 | -5.76654 | -4.81043 | -3.01772 | -3.01772 | -3.95871 | -2.62283 | Yrdc |
| ENSMUSG00000022206 | -4.69092 | -3.64517 | -1.70307 | -1.70307 | -3.92662 | -2.58354 | Npr3 |
| ENSMUSG00000044968 | -5.13909 | -4.15311 | -2.71894 | -2.71894 | -3.92233 | -2.57902 | Napepld |
| ENSMUSG00000053693 | -4.51165 | -3.34639 | -1.61344 | -1.61344 | -3.91036 | -2.56582 | Mast1 |
| ENSMUSG00000053930 | -4.9897 | -4.12323 | -1.46404 | -1.46404 | -3.90771 | -2.56455 | Shisa6 |
| ENSMUSG00000001123 | -4.81043 | -4.12323 | -1.91222 | -1.91222 | -3.90424 | -2.56116 | Lgals9 |
| ENSMUSG00000030287 | -4.87019 | -3.91408 | -2.45003 | -2.45003 | -3.89845 | -2.55446 | Itpr2 |
| ENSMUSG00000047878 | -5.64703 | -4.66104 | -2.83845 | -2.83845 | -3.88504 | -2.53996 | A4galt |
| ENSMUSG00000037166 | -4.66104 | -3.49578 | -2.09149 | -2.09149 | -3.87532 | -2.53364 | Ppp1r14a |
| ENSMUSG00000039620 | -6.42387 | -5.43788 | -3.49578 | -3.49578 | -3.87519 | -2.53364 | 6430573F1 |
| ENSMUSG00000059741 | -6.33423 | -4.78055 | -2.56955 | -2.56955 | -3.87757 | -2.53364 | Myl3 |
| ENSMUSG00000020886 | -4.60128 | -3.43602 | -1.76283 | -1.76283 | -3.87095 | -2.53065 | Dlg4 |
| ENSMUSG00000031112 | -4.78055 | -3.73481 | -2.15125 | -2.15125 | -3.83844 | -2.48524 | Stk26 |
| ENSMUSG00000027684 | -7.23059 | -6.30435 | -1.97198 | -1.97198 | -3.82344 | -2.46659 | Mecom |
| ENSMUSG00000029093 | -5.8263 | -5.07934 | -2.6293 | -2.6293 | -3.80888 | -2.45007 | Sorcs2 |
| ENSMUSG00000008136 | -7.05132 | -6.2446 | -1.97198 | -1.97198 | -3.80752 | -2.44964 | Fhl2 |
| ENSMUSG00000048330 | -5.10922 | -4.03359 | -2.33052 | -2.33052 | -3.78543 | -2.41895 | Ric3 |
| ENSMUSG00000030431 | -4.92995 | -3.73481 | -2.21101 | -2.21101 | -3.78394 | -2.4183 | Tmem238 |
| ENSMUSG00000057286 | -6.33423 | -5.46776 | -3.01772 | -3.01772 | -3.76286 | -2.39204 | St6galnac2 |
| ENSMUSG00000069045 | -6.63302 | -5.58727 | -2.45003 | -2.45003 | -3.74811 | -2.37458 | Ddx3y |
| ENSMUSG00000043336 | -5.43788 | -4.63116 | -3.31651 | -3.31651 | -3.73606 | -2.36198 | Filip1l |

| | | | | | | | |
|---|---|---|---|---|---|---|---|
| ENSMUSG00000051499 | -4.5714 | -3.64517 | -1.46404 | -1.46404 | -3.71243 | -2.3329 | Zfp786 |
| ENSMUSG00000025789 | -5.52751 | -4.5714 | -2.74882 | -2.74882 | -3.70592 | -2.32627 | St8sia2 |
| ENSMUSG00000004665 | -7.49949 | -2.65918 | -1.49392 | -1.49392 | -3.70113 | -2.32216 | Cnn2 |
| ENSMUSG00000063415 | -4.5714 | -3.55554 | -1.01587 | -1.01587 | -3.69982 | -2.3217 | Cyp26b1 |
| ENSMUSG00000030731 | -4.7208 | -3.85432 | -2.09149 | -2.09149 | -3.66589 | -2.27572 | Syt3 |
| ENSMUSG00000020593 | -5.28849 | -4.39213 | -2.42016 | -2.42016 | -3.64418 | -2.24473 | Lpin1 |
| ENSMUSG00000075588 | -5.49764 | -4.7208 | -2.6293 | -2.6293 | -3.61267 | -2.20328 | Hoxb2 |
| ENSMUSG00000038119 | -4.90007 | -3.82444 | -1.9421 | -1.9421 | -3.59695 | -2.18694 | Cdon |
| ENSMUSG00000082766 | -3.8842 | -2.3604 | -1.19514 | -1.19514 | -3.59874 | -2.18694 | 1700064H |
| ENSMUSG00000030895 | -4.92995 | -3.8842 | -2.09149 | -2.09149 | -3.58109 | -2.16585 | Hpx |
| ENSMUSG00000027452 | -4.81043 | -4.09335 | -2.09149 | -2.09149 | -3.57027 | -2.15235 | Acss1 |
| ENSMUSG00000031833 | -5.76654 | -4.69092 | -2.12137 | -2.12137 | -3.56911 | -2.15205 | Mast3 |
| ENSMUSG00000046994 | -5.31837 | -4.39213 | -2.33052 | -2.33052 | -3.54038 | -2.11986 | Mars2 |
| ENSMUSG00000026235 | -5.55739 | -2.56955 | -1.16526 | -1.16526 | -3.53258 | -2.10937 | Epha4 |
| ENSMUSG00000015619 | -6.21472 | -5.408 | -1.82259 | -1.82259 | -3.52764 | -2.10324 | Gata3 |
| ENSMUSG00000049690 | -4.92995 | -4.03359 | -2.06161 | -2.06161 | -3.50488 | -2.07402 | Nckap5 |
| ENSMUSG00000025964 | -4.5714 | -3.52566 | -1.73295 | -1.73295 | -3.50036 | -2.06992 | Adam23 |
| ENSMUSG00000062184 | -6.57326 | -5.8263 | -1.85246 | -1.85246 | -3.48682 | -2.05189 | Hs6st2 |
| ENSMUSG00000066152 | -5.85618 | -4.54153 | -2.33052 | -2.33052 | -3.4781 | -2.04124 | Slc31a2 |
| ENSMUSG00000003746 | -5.13909 | -4.06347 | -2.00186 | -2.00186 | -3.46905 | -2.03006 | Man1a |
| ENSMUSG00000027562 | -6.6629 | -5.67691 | -2.15125 | -2.15125 | -3.46677 | -2.02833 | Car2 |
| ENSMUSG00000029844 | -6.72265 | -5.88606 | -1.82259 | -1.82259 | -3.45183 | -2.01731 | Hoxa1 |
| ENSMUSG00000031075 | -5.79642 | -4.60128 | -2.50979 | -2.50979 | -3.44913 | -2.01478 | Ano1 |
| ENSMUSG00000000253 | -6.15496 | -5.16897 | -2.21101 | -2.21101 | -3.43354 | -1.993 | Gmpr |
| ENSMUSG00000046782 | -6.15496 | -5.34824 | -2.6293 | -2.6293 | -3.43227 | -1.9924 | 4921506M |
| ENSMUSG00000069833 | -5.70679 | -4.78055 | -2.68906 | -2.68906 | -3.42828 | -1.98867 | Ahnak |
| ENSMUSG00000040734 | -4.51165 | -3.34639 | -1.5238 | -1.5238 | -3.38833 | -1.93365 | Ppp1r13l |
| ENSMUSG00000038738 | -5.01958 | -3.70493 | -1.97198 | -1.97198 | -3.38259 | -1.92684 | Shank1 |
| ENSMUSG00000020140 | -4.5714 | -3.58542 | -1.70307 | -1.70307 | -3.37683 | -1.92048 | Lgr5 |
| ENSMUSG00000024029 | -5.28849 | -4.5714 | -2.24088 | -2.24088 | -3.3445 | -1.87898 | Tff3 |
| ENSMUSG00000041907 | -4.45189 | -3.25675 | -1.64332 | -1.64332 | -3.33087 | -1.86244 | Gpr45 |
| ENSMUSG00000050520 | -6.30435 | -5.31837 | -1.97198 | -1.97198 | -3.31328 | -1.8397 | Cldn8 |
| ENSMUSG00000021469 | -6.00557 | -5.31837 | -1.61344 | -1.61344 | -3.2989 | -1.81795 | Msx2 |
| ENSMUSG00000004891 | -5.408 | -4.54153 | -1.01587 | -1.01587 | -3.27983 | -1.79658 | Nes |
| ENSMUSG00000041046 | -5.79642 | -4.63116 | -2.15125 | -2.15125 | -3.27333 | -1.79001 | Ramp3 |
| ENSMUSG00000043501 | -5.52751 | -4.7208 | -1.97198 | -1.97198 | -3.27451 | -1.79001 | Lgals2 |
| ENSMUSG00000010080 | -5.85618 | -5.04946 | -2.12137 | -2.12137 | -3.24697 | -1.75864 | Epn3 |
| ENSMUSG00000021062 | -6.5135 | -5.55739 | -1.91222 | -1.91222 | -3.24715 | -1.75864 | Rab15 |
| ENSMUSG00000032186 | -5.13909 | -4.06347 | -1.9421 | -1.9421 | -3.23733 | -1.74551 | Tmod2 |
| ENSMUSG00000056973 | -6.42387 | -5.37812 | -2.27076 | -2.27076 | -3.23013 | -1.73956 | Ces1d |
| ENSMUSG00000036022 | -5.04946 | -4.27262 | -1.91222 | -1.91222 | -3.20916 | -1.71043 | Fam122b |
| ENSMUSG00000030124 | -4.45189 | -3.40614 | -1.67319 | -1.67319 | -3.19986 | -1.69775 | Lag3 |
| ENSMUSG00000027357 | -5.10922 | -3.73481 | -1.61344 | -1.61344 | -3.16785 | -1.65776 | Crls1 |
| ENSMUSG00000068699 | -6.15496 | -4.39213 | -1.91222 | -1.91222 | -3.16406 | -1.65492 | Flnc |
| ENSMUSG00000062373 | -4.87019 | -4.09335 | -1.79271 | -1.79271 | -3.1602 | -1.65145 | Tmem65 |
| ENSMUSG00000000282 | -5.49764 | -4.33238 | -2.03174 | -2.03174 | -3.15373 | -1.64577 | Mnt |
| ENSMUSG00000053897 | -6.72265 | -6.09521 | -1.13538 | -1.13538 | -3.14961 | -1.64268 | Slc39a8 |
| ENSMUSG00000017802 | -6.5135 | -5.67691 | -1.70307 | -1.70307 | -3.14587 | -1.63771 | Fam134c |
| ENSMUSG00000022297 | -6.03545 | -4.87019 | -2.27076 | -2.27076 | -3.12838 | -1.61356 | Fzd6 |
| ENSMUSG00000021476 | -4.92995 | -4.09335 | -1.61344 | -1.61344 | -3.11591 | -1.59893 | Habp4 |
| ENSMUSG00000027340 | -5.49764 | -4.66104 | -1.76283 | -1.76283 | -3.11326 | -1.59572 | Slc23a2 |
| ENSMUSG00000032289 | -4.63116 | -3.4659 | -1.46404 | -1.46404 | -3.09185 | -1.56713 | Thsd4 |
| ENSMUSG00000078485 | -4.66104 | -3.82444 | -1.76283 | -1.76283 | -3.09009 | -1.56574 | Plekhn1 |
| ENSMUSG00000041112 | -6.48363 | -5.28849 | -1.40429 | -1.40429 | -3.06169 | -1.52587 | Elmo1 |
| ENSMUSG00000041351 | -5.07934 | -4.21286 | -1.88234 | -1.88234 | -3.06092 | -1.52581 | Rap1gap |
| ENSMUSG00000066687 | -5.31837 | -4.42201 | -2.06161 | -2.06161 | -3.04215 | -1.49865 | Zbtb16 |
| ENSMUSG00000026090 | -5.73666 | -4.5714 | -1.85246 | -1.85246 | -3.03687 | -1.4933 | 2010300C( |
| ENSMUSG00000021481 | -6.12508 | -5.34824 | -1.49392 | -1.49392 | -3.0304 | -1.48708 | Zfp346 |

| | | | | | | | |
|---|---|---|---|---|---|---|---|
| ENSMUSG00000042808 | -5.70679 | -4.60128 | -1.82259 | -1.82259 | -3.01919 | -1.46901 | Gpx2 |
| ENSMUSG00000030772 | -5.76654 | -4.66104 | -2.09149 | -2.09149 | -3.01035 | -1.45864 | Dkk3 |
| ENSMUSG00000032327 | -5.28849 | -4.36225 | -1.91222 | -1.91222 | -2.99819 | -1.44365 | Stra6 |
| ENSMUSG00000041608 | -6.48363 | -5.43788 | -1.58356 | -1.58356 | -2.99545 | -1.44011 | Entpd3 |
| ENSMUSG00000031517 | -5.22873 | -4.27262 | -1.67319 | -1.67319 | -2.98021 | -1.42206 | Gpm6a |
| ENSMUSG00000027954 | -7.26047 | -6.81229 | -1.13538 | -1.13538 | -2.96021 | -1.39309 | Efna1 |
| ENSMUSG00000043773 | -4.78055 | -3.67505 | -1.31465 | -1.31465 | -2.95552 | -1.38607 | 1700048O |
| ENSMUSG00000027524 | -6.15496 | -4.84031 | -1.9421 | -1.9421 | -2.94508 | -1.37134 | Edn3 |
| ENSMUSG00000032085 | -5.25861 | -4.36225 | -1.70307 | -1.70307 | -2.94443 | -1.37134 | Tagln |
| ENSMUSG00000057531 | -5.52751 | -4.39213 | -1.79271 | -1.79271 | -2.94263 | -1.36938 | Dtnbp1 |
| ENSMUSG00000039345 | -5.52751 | -4.48177 | -1.43417 | -1.43417 | -2.93937 | -1.36685 | Mettl22 |
| ENSMUSG00000035783 | -5.43788 | -3.82444 | -1.28477 | -1.28477 | -2.93009 | -1.35559 | Acta2 |
| ENSMUSG00000021211 | -4.54153 | -3.4659 | -1.22502 | -1.22502 | -2.92605 | -1.35438 | Akr1c12 |
| ENSMUSG00000036106 | -4.90007 | -3.97383 | -1.49392 | -1.49392 | -2.9163 | -1.34298 | Prr5 |
| ENSMUSG00000019817 | -6.87205 | -5.04946 | -1.43417 | -1.43417 | -2.90687 | -1.33326 | Plagl1 |
| ENSMUSG00000029868 | -5.46776 | -4.75067 | -1.64332 | -1.64332 | -2.88483 | -1.30821 | Trpv6 |
| ENSMUSG00000069793 | -4.27262 | -3.13724 | -1.1055 | -1.1055 | -2.85483 | -1.27116 | Slfn9 |
| ENSMUSG00000058900 | -4.48177 | -3.4659 | -1.19514 | -1.19514 | -2.84005 | -1.25464 | Rsl1 |
| ENSMUSG00000028532 | -6.00557 | -5.10922 | -1.43417 | -1.43417 | -2.83539 | -1.24958 | Cachd1 |
| ENSMUSG00000038582 | -5.10922 | -4.21286 | -1.61344 | -1.61344 | -2.82704 | -1.23769 | Pptc7 |
| ENSMUSG00000029592 | -5.16897 | -4.27262 | -1.70307 | -1.70307 | -2.82274 | -1.2325 | Usp30 |
| ENSMUSG00000031732 | -5.408 | -4.24274 | -1.37441 | -1.37441 | -2.81983 | -1.23027 | Phlpp2 |
| ENSMUSG00000057176 | -5.07934 | -4.12323 | -1.64332 | -1.64332 | -2.81322 | -1.22436 | Gm166 |
| ENSMUSG00000057914 | -4.75067 | -3.49578 | -1.34453 | -1.34453 | -2.81293 | -1.22436 | Cacnb2 |
| ENSMUSG00000038564 | -5.88606 | -4.81043 | -1.34453 | -1.34453 | -2.79553 | -1.20126 | Ift172 |
| ENSMUSG00000050989 | -5.408 | -4.60128 | -1.58356 | -1.58356 | -2.79545 | -1.20126 | Sepn1 |
| ENSMUSG00000037860 | -4.39213 | -3.13724 | -1.07562 | -1.07562 | -2.79239 | -1.19785 | Aim2 |
| ENSMUSG00000030257 | -4.7208 | -3.82444 | -1.37441 | -1.37441 | -2.78921 | -1.19421 | Srgap3 |
| ENSMUSG00000031504 | -4.81043 | -3.82444 | -1.5238 | -1.5238 | -2.77982 | -1.18352 | Rab20 |
| ENSMUSG00000039481 | -4.7208 | -3.70493 | -1.46404 | -1.46404 | -2.71612 | -1.10921 | Nrtn |
| ENSMUSG00000034936 | -5.10922 | -4.15311 | -1.28477 | -1.28477 | -2.71201 | -1.10362 | Arl4d |
| ENSMUSG00000020357 | -5.31837 | -4.45189 | -1.16526 | -1.16526 | -2.70734 | -1.09925 | Flt4 |
| ENSMUSG00000003545 | -5.43788 | -4.42201 | -1.28477 | -1.28477 | -2.6928 | -1.08017 | Fosb |
| ENSMUSG00000046634 | -5.52751 | -4.60128 | -1.46404 | -1.46404 | -2.69231 | -1.08017 | Pkd1l1 |
| ENSMUSG00000020469 | -6.09521 | -3.76469 | -1.01587 | -1.01587 | -2.69067 | -1.07917 | Myl7 |
| ENSMUSG00000021277 | -5.52751 | -4.66104 | -1.01587 | -1.01587 | -2.68616 | -1.07588 | Traf3 |
| ENSMUSG00000027030 | -4.63116 | -3.52566 | -1.04575 | -1.04575 | -2.68147 | -1.07211 | Stk39 |
| ENSMUSG00000038893 | -4.51165 | -3.55554 | -1.01587 | -1.01587 | -2.67809 | -1.06881 | Fam117a |
| ENSMUSG00000038388 | -5.408 | -4.42201 | -1.19514 | -1.19514 | -2.66417 | -1.05192 | Mpp6 |
| ENSMUSG00000074500 | -5.52751 | -4.12323 | -1.19514 | -1.19514 | -2.64307 | -1.02762 | Zfp558 |
| ENSMUSG00000020319 | -4.90007 | -3.91408 | -1.19514 | -1.19514 | -2.6314 | -1.01431 | Wdpcp |
| ENSMUSG00000006463 | -5.46776 | -4.5714 | -1.19514 | -1.19514 | -2.62772 | -1.00964 | Zdhhc24 |
| ENSMUSG00000059824 | -5.19885 | -4.09335 | -1.07562 | -1.07562 | -2.61294 | -0.99471 | Dbp |
| ENSMUSG00000037679 | -4.92995 | -3.8842 | -1.19514 | -1.19514 | -2.60614 | -0.98652 | Inf2 |
| ENSMUSG00000000627 | -5.04946 | -4.18298 | -1.37441 | -1.37441 | -2.60395 | -0.98462 | Sema4f |
| ENSMUSG00000038777 | -5.01958 | -4.03359 | -1.13538 | -1.13538 | -2.58445 | -0.96112 | Sema6c |
| ENSMUSG00000030029 | -4.87019 | -3.85432 | -1.16526 | -1.16526 | -2.58266 | -0.95985 | Lrig1 |
| ENSMUSG00000064267 | -4.84031 | -3.8842 | -1.13538 | -1.13538 | -2.55315 | -0.91932 | Hvcn1 |
| ENSMUSG00000048911 | -5.55739 | -4.63116 | -1.07562 | -1.07562 | -2.53848 | -0.90668 | Rnf24 |
| ENSMUSG00000040877 | -5.46776 | -4.36225 | -1.19514 | -1.19514 | -2.46773 | -0.81227 | Wdr25 |
| ENSMUSG00000040502 | 1.045746 | 4.212863 | 5.557393 | 1.045746 | 2.563639 | 0.933138 | March9 |
| ENSMUSG00000047904 | 1.225017 | 4.093349 | 5.736664 | 1.225017 | 2.615925 | 0.997506 | Sstr2 |
| ENSMUSG00000016942 | 1.344531 | 4.422012 | 5.587272 | 1.344531 | 2.638797 | 1.022474 | Tmprss6 |
| ENSMUSG00000027577 | 1.135381 | 4.810432 | 6.423869 | 1.135381 | 2.66597 | 1.051985 | Chrna4 |
| ENSMUSG00000046834 | 1.374409 | 4.362255 | 5.736664 | 1.374409 | 2.681906 | 1.07211 | Krt1 |
| ENSMUSG00000017724 | 1.075625 | 4.511647 | 6.513504 | 1.075625 | 2.696534 | 1.085009 | Etv4 |
| ENSMUSG00000059412 | 1.493923 | 4.959824 | 6.035449 | 1.493923 | 2.708191 | 1.099586 | Fxyd2 |
| ENSMUSG00000069206 | 1.135381 | 3.764686 | 5.527515 | 1.135381 | 2.775713 | 1.180172 | Zfp874a |

| | | | | | | | |
|---|---|---|---|---|---|---|---|
| ENSMUSG00000061877 | 1.55368 | 5.258609 | 6.154962 | 1.55368 | 2.835459 | 1.249578 | BC048679 |
| ENSMUSG00000052676 | 1.135381 | 4.033592 | 5.318366 | 1.135381 | 2.837147 | 1.250511 | Zmat1 |
| ENSMUSG00000033389 | 1.314652 | 5.706786 | 6.722653 | 1.314652 | 2.871472 | 1.295174 | Arhgap44 |
| ENSMUSG00000038037 | 1.493923 | 4.571404 | 5.647029 | 1.493923 | 2.881413 | 1.306343 | Socs1 |
| ENSMUSG00000046352 | 2.001857 | 5.915935 | 6.752532 | 2.001857 | 2.897665 | 1.323843 | Gjb2 |
| ENSMUSG00000069305 | 1.314652 | 3.645172 | 4.959824 | 1.314652 | 2.907059 | 1.333258 | Hist1h4n |
| ENSMUSG00000035397 | 1.284774 | 5.139095 | 6.184841 | 1.284774 | 2.913335 | 1.341236 | Klf16 |
| ENSMUSG00000068762 | 2.031735 | 5.318366 | 6.453747 | 2.031735 | 2.977098 | 1.417842 | Gstm6 |
| ENSMUSG00000048251 | 1.852464 | 4.870189 | 6.214719 | 1.852464 | 3.035376 | 1.493181 | Bcl11b |
| ENSMUSG00000061482 | 1.344531 | 3.49578 | 4.959824 | 1.344531 | 3.072745 | 1.540975 | Hist1h4d |
| ENSMUSG00000062444 | 2.121371 | 5.109216 | 6.304355 | 2.121371 | 3.123812 | 1.609666 | Ap3b2 |
| ENSMUSG00000037681 | 1.404288 | 3.943957 | 5.796421 | 1.404288 | 3.150294 | 1.642677 | Esyt3 |
| ENSMUSG00000028445 | 1.9421 | 4.780553 | 5.945813 | 1.9421 | 3.156694 | 1.649421 | Enho |
| ENSMUSG00000030145 | 1.16526 | 2.778697 | 5.587272 | 1.16526 | 3.157359 | 1.649421 | Zfp248 |
| ENSMUSG00000004035 | 1.16526 | 2.569547 | 4.481769 | 1.16526 | 3.246664 | 1.758641 | Gstm7 |
| ENSMUSG00000021684 | 2.121371 | 5.378123 | 6.453747 | 2.121371 | 3.246642 | 1.758641 | Pde8b |
| ENSMUSG00000052942 | 2.659183 | 4.959824 | 6.00557 | 2.659183 | 3.284692 | 1.803097 | Glis3 |
| ENSMUSG00000008575 | 1.045746 | 2.569547 | 5.826299 | 1.045746 | 3.292581 | 1.811805 | Nfib |
| ENSMUSG00000038233 | 1.523801 | 6.035449 | 6.961681 | 1.523801 | 3.320774 | 1.849002 | Fam198a |
| ENSMUSG00000026163 | 2.061614 | 4.481769 | 6.364112 | 2.061614 | 3.342945 | 1.878977 | Sphkap |
| ENSMUSG00000032120 | 2.420155 | 4.332376 | 5.886056 | 2.420155 | 3.352907 | 1.889402 | C2cd2l |
| ENSMUSG00000047040 | 1.045746 | 2.390277 | 3.704929 | 1.045746 | 3.39425 | 1.939726 | Prr15l |
| ENSMUSG00000027544 | 2.001857 | 4.690918 | 5.766542 | 2.001857 | 3.454617 | 2.018875 | Nfatc2 |
| ENSMUSG00000040938 | 1.075625 | 4.123227 | 5.258609 | 1.075625 | 3.459364 | 2.022606 | Slc16a11 |
| ENSMUSG00000027801 | 1.16526 | 2.450034 | 6.304355 | 1.16526 | 3.463593 | 2.025825 | Tm4sf4 |
| ENSMUSG00000038521 | 2.240884 | 4.392133 | 5.975692 | 2.240884 | 3.466112 | 2.028325 | C1s1 |
| ENSMUSG00000029648 | 1.374409 | 6.065327 | 7.140952 | 1.374409 | 3.472775 | 2.034413 | Flt1 |
| ENSMUSG00000018427 | 1.045746 | 2.151249 | 3.8842 | 1.045746 | 3.542871 | 2.122228 | Ypel2 |
| ENSMUSG00000026398 | 1.195138 | 2.509791 | 5.706786 | 1.195138 | 3.563838 | 2.148403 | Nr5a2 |
| ENSMUSG00000069306 | 2.001857 | 4.123227 | 5.258609 | 2.001857 | 3.576795 | 2.160797 | Hist1h4m |
| ENSMUSG00000036306 | 1.135381 | 2.569547 | 5.826299 | 1.135381 | 3.602887 | 2.191475 | Lzts1 |
| ENSMUSG00000019790 | 2.509791 | 4.870189 | 5.945813 | 2.509791 | 3.613366 | 2.203283 | Stxbp5 |
| ENSMUSG00000004633 | 2.838454 | 5.915935 | 6.633018 | 2.838454 | 3.617549 | 2.20765 | Chn2 |
| ENSMUSG00000003031 | 1.374409 | 3.226874 | 4.631161 | 1.374409 | 3.674544 | 2.287128 | Cdkn1b |
| ENSMUSG00000036256 | 2.748818 | 5.198852 | 6.274476 | 2.748818 | 3.703928 | 2.32481 | Igfbp7 |
| ENSMUSG00000035540 | 2.569547 | 5.766542 | 6.78241 | 2.569547 | 3.708876 | 2.329145 | Gc |
| ENSMUSG00000022490 | 1.015868 | 2.061614 | 3.585415 | 1.015868 | 3.782743 | 2.418097 | Ppp1r1a |
| ENSMUSG00000070287 | 2.659183 | 4.959824 | 6.065327 | 2.659183 | 3.819129 | 2.463436 | Slc35g2 |
| ENSMUSG00000042589 | 1.16526 | 2.390277 | 3.824443 | 1.16526 | 3.878119 | 2.533642 | Cux2 |
| ENSMUSG00000020467 | 2.928089 | 5.019581 | 6.39399 | 2.928089 | 3.896614 | 2.55342 | Efemp1 |
| ENSMUSG00000030559 | 1.254895 | 2.479912 | 4.27262 | 1.254895 | 3.909726 | 2.565822 | Rab38 |
| ENSMUSG00000027217 | 1.314652 | 2.509791 | 4.27262 | 1.314652 | 3.918393 | 2.575008 | Tspan18 |
| ENSMUSG00000010307 | 1.344531 | 2.629304 | 6.901924 | 1.344531 | 3.930002 | 2.586735 | Tmem86a |
| ENSMUSG00000050199 | 1.254895 | 2.509791 | 3.764686 | 1.254895 | 3.935342 | 2.592733 | Lgr4 |
| ENSMUSG00000079015 | 1.254895 | 2.509791 | 3.49578 | 1.254895 | 3.948833 | 2.610379 | Serpina1c |
| ENSMUSG00000023961 | 3.525658 | 5.408001 | 6.453747 | 3.525658 | 4.035767 | 2.71709 | Enpp4 |
| ENSMUSG00000037437 | 2.270763 | 4.989703 | 6.184841 | 2.270763 | 4.086169 | 2.781134 | Adam32 |
| ENSMUSG00000041202 | 2.509791 | 4.332376 | 6.095206 | 2.509791 | 4.174464 | 2.883411 | Pla2g2d |
| ENSMUSG00000020051 | 3.137238 | 4.959824 | 6.364112 | 3.137238 | 4.194381 | 2.909182 | Pah |
| ENSMUSG00000023224 | 1.493923 | 2.957967 | 4.153106 | 1.493923 | 4.256722 | 2.978992 | Serping1 |
| ENSMUSG00000000567 | 1.254895 | 2.868332 | 4.392133 | 1.254895 | 4.391555 | 3.125213 | Sox9 |
| ENSMUSG00000051379 | 1.225017 | 2.509791 | 3.615293 | 1.225017 | 4.424918 | 3.16636 | Flrt3 |
| ENSMUSG00000045725 | 2.240884 | 5.139095 | 6.154962 | 2.240884 | 4.526169 | 3.286201 | Prr15 |
| ENSMUSG00000021850 | 1.493923 | 2.778697 | 3.764686 | 1.493923 | 4.626222 | 3.412669 | 1700011H |
| ENSMUSG00000026643 | 1.045746 | 1.882343 | 2.748818 | 1.045746 | 4.645709 | 3.434309 | Nmt2 |
| ENSMUSG00000039533 | 2.659183 | 6.125084 | 7.200709 | 2.659183 | 4.749213 | 3.556371 | Mmd2 |
| ENSMUSG00000025075 | 1.284774 | 2.390277 | 4.392133 | 1.284774 | 4.77419 | 3.583758 | Habp2 |
| ENSMUSG00000064225 | 1.822586 | 2.987846 | 5.288487 | 1.822586 | 4.846078 | 3.664219 | Paqr9 |

| | | | | | | | |
|---|---|---|---|---|---|---|---|
| ENSMUSG00000032053 | 2.957967 | 4.810432 | 6.244598 | 2.957967 | 4.880733 | 3.702453 | Pou2af1 |
| ENSMUSG00000018593 | 1.225017 | 2.240884 | 3.376266 | 1.225017 | 4.939976 | 3.76759 | Sparc |
| ENSMUSG00000040459 | 1.045746 | 1.792708 | 2.450034 | 1.045746 | 5.021354 | 3.860287 | Arglu1 |
| ENSMUSG00000028194 | 1.314652 | 2.240884 | 3.316509 | 1.314652 | 5.028894 | 3.863567 | Ddah1 |
| ENSMUSG00000072849 | 1.9421 | 3.137238 | 4.332376 | 1.9421 | 5.127613 | 3.9617 | Serpina1e |
| ENSMUSG00000032291 | 2.987846 | 6.035449 | 8.037305 | 2.987846 | 5.227352 | 4.046952 | Crabp1 |
| ENSMUSG00000037254 | 1.792708 | 3.555537 | 4.929946 | 1.792708 | 5.580296 | 4.35492 | Itih2 |
| ENSMUSG00000027690 | 2.957967 | 5.826299 | 7.379979 | 2.957967 | 5.801313 | 4.525233 | Slc2a2 |
| ENSMUSG00000029761 | 1.792708 | 2.808575 | 3.824443 | 1.792708 | 5.94058 | 4.635341 | Cald1 |
| ENSMUSG00000045518 | 4.033592 | 5.467758 | 6.543382 | 4.033592 | 6.016135 | 4.695351 | Onecut3 |
| ENSMUSG00000030854 | 5.378123 | 6.931802 | 7.708642 | 5.378123 | 6.29449 | 5.002622 | Ptpn5 |
| ENSMUSG00000055254 | 4.481769 | 6.125084 | 6.901924 | 4.481769 | 6.614367 | 5.365183 | Ntrk2 |
| ENSMUSG00000040856 | 1.912221 | 2.957967 | 3.914078 | 1.912221 | 6.769431 | 5.536705 | Dlk1 |
| ENSMUSG00000000753 | 2.240884 | 3.256752 | 4.690918 | 2.240884 | 6.862176 | 5.630547 | Serpinf1 |
| ENSMUSG00000051159 | 2.629304 | 3.8842 | 5.109216 | 2.629304 | 7.040956 | 5.825642 | Cited1 |
| ENSMUSG00000020644 | 1.912221 | 2.629304 | 3.376266 | 1.912221 | 7.160795 | 5.921669 | Id2 |
| ENSMUSG00000024986 | 2.659183 | 3.943957 | 8.216576 | 2.659183 | 7.160806 | 5.921669 | Hhex |
| ENSMUSG00000025902 | 3.734807 | 8.485482 | 9.262322 | 3.734807 | 7.16081 | 5.921669 | Sox17 |
| ENSMUSG00000028356 | 2.360398 | 3.256752 | 5.019581 | 2.360398 | 7.160813 | 5.921669 | Ambp |

#### r primordium and gut tube

95
c(lb, ub))), or 0 if the CI crosses the 0

ısing Holm procedure
**d cZ.**

| Descriptio | Location | Family |
|---|---|---|
| calpain 6 | Cytoplasm | peptidase |
| forkhead b | Nucleus | transcription regulator |
| tropomyos | Plasma Membrane | other |
| odd-skipp | Nucleus | kinase |
| homeobox | Nucleus | transcription regulator |
| splA/ryanc | Cytoplasm | enzyme |
| corin, serin | Plasma Membrane | peptidase |
| non-comp | Plasma Membrane | other |
| msh home | Nucleus | transcription regulator |
| indian hed | Extracellular Space | enzyme |
| laminin su | Extracellular Space | transporter |
| ISL LIM ho | Nucleus | transcription regulator |
| UDP-GlcN | Cytoplasm | enzyme |
| NK2 home | Nucleus | transcription regulator |
| retinol deh | Nucleus | enzyme |
| atonal bHI | Nucleus | transcription regulator |
| transmeml | Other | other |
| endoplasn | Cytoplasm | enzyme |
| dehydroge | Cytoplasm | enzyme |
| iodothyror | Plasma Membrane | enzyme |
| LY6/PLAUI | Other | other |
| paired like | Nucleus | transcription regulator |
| chromoso | Other | other |
| family with | Other | other |
| BCL2 like 1 | Cytoplasm | other |
| homeobox | Nucleus | transcription regulator |
| solute carr | Plasma Membrane | transporter |
| testis deve | Cytoplasm | other |
| intestine s | Nucleus | transcription regulator |
| syncollin | Extracellular Space | other |
| tubby bipa | Cytoplasm | transcription regulator |
| carbohydr | Cytoplasm | enzyme |
| CD55 mole | Plasma Membrane | other |
| frizzled rel | Extracellular Space | other |
| amyloid be | Cytoplasm | transporter |
| synaptogy | Plasma Membrane | other |
| retinoic ac | Nucleus | ligand-dependent nuclear receptor |
| caudal typ | Nucleus | transcription regulator |
| PWWP do | Nucleus | other |
| protein tyr | Plasma Membrane | phosphatase |
| interferon | Other | other |
| histamine | Cytoplasm | enzyme |
| neuritin 1 | Cytoplasm | other |
| pleckstrin | Extracellular Space | other |
| lysophosp | Plasma Membrane | transmembrane receptor |
| insulin rec | Other | other |
| protocadh | Plasma Membrane | other |

| | | |
|---|---|---|
| BRD4 inter | Other | other |
| proprotei | Extracellular Space | peptidase |
| annexin A | Cytoplasm | enzyme |
| phosphol | Plasma Membrane | phosphatase |
| PERP, TP5 | Plasma Membrane | other |
| arginase 1 | Cytoplasm | enzyme |
| myosin re | Cytoplasm | enzyme |
| tetraspani | Plasma Membrane | other |
| ORAI calci | Plasma Membrane | ion channel |
| TEA doma | Nucleus | transcription regulator |
| Kruppel lik | Nucleus | transcription regulator |
| ectonucle | Plasma Membrane | enzyme |
| prostaglar | Cytoplasm | enzyme |
| G protein- | Plasma Membrane | G-protein coupled receptor |
| myoferlin | Nucleus | other |
| NUAK fam | Other | kinase |
| PR/SET do | Nucleus | transcription regulator |
| androglob | Other | other |
| NK3 home | Nucleus | transcription regulator |
| catenin de | Plasma Membrane | other |
| SERTA do | Other | other |
| coagulatio | Plasma Membrane | transmembrane receptor |
| solute carr | Plasma Membrane | transporter |
| elaC ribon | Nucleus | enzyme |
| keratin 7 | Cytoplasm | other |
| paired like | Nucleus | transcription regulator |
| protocadh | Other | other |
| mab-21 lik | Other | other |
| regulatory | Nucleus | transcription regulator |
| solute carr | Plasma Membrane | transporter |
| G protein | Plasma Membrane | enzyme |
| G protein- | Plasma Membrane | G-protein coupled receptor |
| bone mor | Extracellular Space | growth factor |
| ATPase 13 | Cytoplasm | transporter |
| plasminog | Plasma Membrane | transmembrane receptor |
| chemerin | Plasma Membrane | G-protein coupled receptor |
| histidine a | Cytoplasm | enzyme |
| SPARC rel | Extracellular Space | other |
| LDOC1, re | Nucleus | other |
| cripto, FRL | Extracellular Space | other |
| ADAM me | Extracellular Space | peptidase |
| cadherin 2 | Plasma Membrane | other |
| dedicator | Cytoplasm | other |
| basic helix | Nucleus | transcription regulator |
| FAT atypic | Plasma Membrane | other |
| SRY-box 2 | Nucleus | transcription regulator |
| histocomp | Plasma Membrane | other |
| insulin like | Extracellular Space | other |
| podocalyx | Plasma Membrane | other |
| transformi | Extracellular Space | growth factor |
| mannosyl | Cytoplasm | enzyme |
| triple Qxx | Cytoplasm | other |
| beta-1,4- | Cytoplasm | enzyme |
| EPH recep | Plasma Membrane | kinase |
| pleckstrin | Plasma Membrane | other |
| collapsin | Cytoplasm | enzyme |
| homeobo | Nucleus | transcription regulator |
| solute carr | Cytoplasm | transporter |

| | | |
|---|---|---|
| seizure rel | Plasma Membrane | other |
| ST6 beta-g | Cytoplasm | enzyme |
| transmeml | Plasma Membrane | other |
| forkhead k | Nucleus | transcription regulator |
| spalt like t | Nucleus | transcription regulator |
| GRB2 asso | Nucleus | other |
| collagen ty | Extracellular Space | other |
| protein kir | Cytoplasm | kinase |
| NK2 home | Nucleus | transcription regulator |
| semaphori | Plasma Membrane | transmembrane receptor |
| regulatory | Nucleus | enzyme |
| amyloid be | Plasma Membrane | other |
| family with | Nucleus | other |
| NIPA like c | Other | other |
| ubiquitin C | Cytoplasm | peptidase |
| serine/thre | Cytoplasm | kinase |
| F2R like try | Plasma Membrane | G-protein coupled receptor |
| major hist | Plasma Membrane | other |
| sphingomy | Extracellular Space | enzyme |
| CHM like, | Cytoplasm | enzyme |
| immunogl | Plasma Membrane | other |
| expressed | Other | other |
| sodium ch | Plasma Membrane | ion channel |
| frizzled cla | Plasma Membrane | G-protein coupled receptor |
| growth arr | Nucleus | other |
| ADAM me | Extracellular Space | other |
| V-set and | Plasma Membrane | other |
| ITPR intera | Plasma Membrane | other |
| versican | Extracellular Space | other |
| ARFGEF fa | Other | other |
| | Other | other |
| sonic hedg | Extracellular Space | peptidase |
| phosphatio | Other | other |
| apolipopro | Extracellular Space | transporter |
| NACC fam | Nucleus | transcription regulator |
| iroquois ho | Nucleus | transcription regulator |
| prostate st | Plasma Membrane | other |
| yrdC N6-t | Other | other |
| natriuretic | Plasma Membrane | G-protein coupled receptor |
| N-acyl pho | Cytoplasm | enzyme |
| microtubu | Cytoplasm | kinase |
| shisa famil | Plasma Membrane | other |
| galectin 9B | Cytoplasm | other |
| inositol 1,4 | Cytoplasm | ion channel |
| alpha 1,4- | Cytoplasm | enzyme |
| protein ph | Cytoplasm | phosphatase |
| tRNA meth | Cytoplasm | enzyme |
| myosin lig | Cytoplasm | other |
| discs large | Plasma Membrane | kinase |
| serine/thre | Nucleus | kinase |
| MDS1 and | Nucleus | transcription regulator |
| sortilin rela | Plasma Membrane | transporter |
| four and a | Nucleus | transcription regulator |
| RIC3 acety | Cytoplasm | other |
| transmeml | Other | other |
| ST6 N-ace | Cytoplasm | enzyme |
| DEAD-box | Cytoplasm | enzyme |
| filamin A ir | Nucleus | other |

| | | |
|---|---|---|
| zinc finger | Nucleus | transcription regulator |
| ST8 alpha- | Cytoplasm | enzyme |
| calponin 2 | Cytoplasm | other |
| cytochrom | Cytoplasm | enzyme |
| synaptotag | Cytoplasm | transporter |
| lipin 1 | Nucleus | phosphatase |
| homeobox | Nucleus | transcription regulator |
| cell adhesi | Plasma Membrane | other |
| RIKEN cDN | Other | other |
| hemopexin | Extracellular Space | transporter |
| acyl-CoA s | Cytoplasm | enzyme |
| microtubu | Other | kinase |
| methionyl- | Cytoplasm | enzyme |
| EPH recep | Plasma Membrane | kinase |
| GATA binc | Nucleus | transcription regulator |
| NCK assoc | Other | other |
| ADAM me | Plasma Membrane | peptidase |
| heparan su | Plasma Membrane | enzyme |
| solute carr | Plasma Membrane | transporter |
| mannosida | Cytoplasm | enzyme |
| carbonic a | Cytoplasm | enzyme |
| homeobox | Nucleus | transcription regulator |
| anoctamin | Plasma Membrane | ion channel |
| guanosine | Cytoplasm | enzyme |
| tetratricop | Other | other |
| AHNAK nu | Nucleus | other |
| protein ph | Nucleus | transcription regulator |
| SH3 and m | Cytoplasm | other |
| leucine ric | Plasma Membrane | transmembrane receptor |
| trefoil fact | Extracellular Space | other |
| G protein- | Plasma Membrane | G-protein coupled receptor |
| claudin 8 | Plasma Membrane | other |
| msh home | Nucleus | transcription regulator |
| nestin | Cytoplasm | other |
| receptor a | Plasma Membrane | transporter |
| galectin 2 | Cytoplasm | other |
| epsin 3 | Cytoplasm | other |
| RAB15, me | Cytoplasm | enzyme |
| tropomod | Cytoplasm | other |
| carboxyles | Cytoplasm | enzyme |
| family with | Other | other |
| lymphocyt | Plasma Membrane | transmembrane receptor |
| cardiolipin | Cytoplasm | enzyme |
| filamin C | Cytoplasm | other |
| transmeml | Cytoplasm | other |
| MAX netw | Nucleus | transcription regulator |
| solute carr | Extracellular Space | transporter |
| reticuloph | Other | other |
| frizzled cla | Plasma Membrane | G-protein coupled receptor |
| hyaluronan | Cytoplasm | other |
| solute carr | Plasma Membrane | transporter |
| thrombosp | Cytoplasm | other |
| pleckstrin | Other | other |
| engulfmer | Cytoplasm | other |
| RAP1 GTPa | Cytoplasm | other |
| zinc finger | Nucleus | transcription regulator |
| KIAA1211 | Other | other |
| zinc finger | Nucleus | other |

| | | |
|---|---|---|
| glutathion | Cytoplasm | enzyme |
| dickkopf W | Extracellular Space | cytokine |
| stimulated | Plasma Membrane | transporter |
| ectonucleo | Plasma Membrane | enzyme |
| glycoprote | Plasma Membrane | ion channel |
| ephrin A1 | Plasma Membrane | other |
| RIKEN cDN | Other | other |
| endothelin | Extracellular Space | other |
| transgelin | Cytoplasm | other |
| dystrobrev | Plasma Membrane | other |
| methyltran | Nucleus | enzyme |
| actin, alph | Cytoplasm | other |
| aldo-keto | Other | enzyme |
| proline ricl | Cytoplasm | other |
| PLAG1 like | Nucleus | transcription regulator |
| transient r | Plasma Membrane | ion channel |
| schlafen fa | Nucleus | enzyme |
| regulator c | Other | other |
| cache dom | Other | other |
| PTC7 prot | Cytoplasm | phosphatase |
| ubiquitin s | Cytoplasm | peptidase |
| PH domain | Cytoplasm | enzyme |
| coiled-coi | Other | other |
| calcium vo | Plasma Membrane | ion channel |
| intraflagell | Extracellular Space | other |
| selenoprot | Cytoplasm | other |
| absent in r | Cytoplasm | other |
| SLIT-ROBO | Cytoplasm | other |
| RAB20, me | Cytoplasm | enzyme |
| neurturin | Extracellular Space | growth factor |
| ADP ribosy | Nucleus | enzyme |
| fms relatec | Plasma Membrane | transmembrane receptor |
| FosB proto | Nucleus | transcription regulator |
| polycystin | Plasma Membrane | ion channel |
| myosin lig | Cytoplasm | enzyme |
| TNF recep | Cytoplasm | enzyme |
| serine/thre | Nucleus | kinase |
| family with | Other | transporter |
| membrane | Plasma Membrane | kinase |
| zinc finger | Nucleus | transcription regulator |
| WD repeat | Plasma Membrane | other |
| zinc finger | Plasma Membrane | other |
| D-box bin | Nucleus | transcription regulator |
| inverted fo | Cytoplasm | other |
| ssemapho | Plasma Membrane | other |
| semaphori | Plasma Membrane | other |
| leucine ric | Extracellular Space | other |
| hydrogen | Plasma Membrane | ion channel |
| ring finger | Cytoplasm | other |
| WD repeat | Other | other |
| membrane | Cytoplasm | other |
| somatosta | Plasma Membrane | G-protein coupled receptor |
| transmeml | Plasma Membrane | peptidase |
| cholinergic | Plasma Membrane | transmembrane receptor |
| keratin 1 | Cytoplasm | other |
| ETS varian | Nucleus | transcription regulator |
| FXYD dom | Plasma Membrane | other |
| zinc finger | Other | other |

| | | |
|---|---|---|
| cDNA sequ | Other | other |
| zinc finger | Nucleus | other |
| Rho GTPas | Cytoplasm | other |
| suppresso | Cytoplasm | other |
| gap juncti | Plasma Membrane | transporter |
| histone clu | Nucleus | other |
| Kruppel-lil | Nucleus | transcription regulator |
| glutathion | Cytoplasm | enzyme |
| B cell CLL/ | Nucleus | transcription regulator |
| histone clu | Nucleus | other |
| adaptor re | Cytoplasm | transporter |
| extended s | Plasma Membrane | other |
| energy ho | Plasma Membrane | other |
| zinc finger | Nucleus | transcription regulator |
| glutathion | Cytoplasm | enzyme |
| phosphod | Cytoplasm | enzyme |
| GLIS family | Nucleus | transcription regulator |
| nuclear fac | Nucleus | transcription regulator |
| family with | Other | other |
| SPHK1 inte | Cytoplasm | other |
| C2CD2 like | Cytoplasm | transporter |
| proline ricl | Other | other |
| nuclear fac | Nucleus | transcription regulator |
| solute carr | Cytoplasm | transporter |
| transmeml | Plasma Membrane | other |
| compleme | Extracellular Space | peptidase |
| fms relatec | Plasma Membrane | kinase |
| yippee like | Nucleus | other |
| nuclear rec | Nucleus | ligand-dependent nuclear receptor |
| histone clu | Nucleus | other |
| leucine zip | Nucleus | transcription regulator |
| syntaxin bi | Plasma Membrane | other |
| chimerin 2 | Cytoplasm | other |
| cyclin dep | Nucleus | kinase |
| insulin like | Extracellular Space | transporter |
| GC, vitami | Extracellular Space | transporter |
| protein ph | Cytoplasm | phosphatase |
| solute carr | Cytoplasm | other |
| cut like ho | Nucleus | transcription regulator |
| EGF contai | Extracellular Space | enzyme |
| RAB38, me | Cytoplasm | enzyme |
| tetraspanii | Other | other |
| transmeml | Other | other |
| leucine ric | Plasma Membrane | transmembrane receptor |
| serpin fam | Extracellular Space | other |
| ectonucle | Cytoplasm | enzyme |
| ADAM me | Other | peptidase |
| phospholi | Extracellular Space | enzyme |
| phenylalar | Cytoplasm | enzyme |
| serpin fam | Extracellular Space | other |
| SRY-box 9 | Nucleus | transcription regulator |
| fibronectir | Plasma Membrane | other |
| proline ricl | Other | other |
| coiled-coi | Other | other |
| N-myristo | Cytoplasm | enzyme |
| monocyte | Other | kinase |
| hyaluronai | Extracellular Space | peptidase |
| progestin | Plasma Membrane | other |

| | | |
|---|---|---|
| POU class | Nucleus | transcription regulator |
| secreted p | Extracellular Space | other |
| arginine ai | Other | other |
| dimethylar | Cytoplasm | enzyme |
| serpin fam | Extracellular Space | other |
| cellular ret | Cytoplasm | transporter |
| inter-alpha | Extracellular Space | other |
| solute carr | Plasma Membrane | transporter |
| caldesmor | Plasma Membrane | other |
| one cut dc | Other | transcription regulator |
| protein tyr | Plasma Membrane | phosphatase |
| neurotrop | Plasma Membrane | kinase |
| delta like r | Extracellular Space | other |
| serpin fam | Extracellular Space | other |
| Cbp/p300 | Nucleus | transcription regulator |
| inhibitor o | Nucleus | transcription regulator |
| hematopo | Nucleus | transcription regulator |
| SRY-box 1 | Nucleus | transcription regulator |
| alpha-1-m | Extracellular Space | transporter |

### Gene groups of the epithelial-hepatic transition

| GeneName | Gene group | Description |
|---|---|---|
| Prox1 | L1 | prospero homeobox 1 |
| Dlk1 | L1 | delta like non-canonical Notch ligand 1 |
| Hhex | L1 | hematopoietically expressed homeobox |
| Iah1 | L1 | isoamyl acetate hydrolyzing esterase 1 (putative) |
| Dnajc22 | L1 | DnaJ heat shock protein family (Hsp40) member C22 |
| Mrpl16 | L1 | mitochondrial ribosomal protein L16 |
| Pecr | L1 | peroxisomal trans-2-enoyl-CoA reductase |
| Maob | L1 | monoamine oxidase B |
| Tuba4a | L1 | tubulin alpha 4a |
| Ppif | L1 | peptidylprolyl isomerase F |
| Mt2 | L1 | metallothionein 2 |
| Junb | L1 | JunB proto-oncogene, AP-1 transcription factor subunit |
| Osgin1 | L1 | oxidative stress induced growth inhibitor 1 |
| Fam195a | L1 | - |
| Fxyd1 | L1 | FXYD domain containing ion transport regulator 1 |
| Proc | L1 | protein C, inactivator of coagulation factors Va and VIIIa |
| Cd302 | L1 | CD40 molecule |
| Cyp2c68 | L1 | cytochrome P450, family 2, subfamily c, polypeptide 40 |
| C3 | L1 | calpain 6 |
| Agmat | L1 | agmatinase |
| Alb | L1 | albumin |
| Serpina6 | L1 | serpin family A member 6 |
| Gulo | L1 | gulonolactone (L-) oxidase |
| Lect2 | L1 | leukocyte cell derived chemotaxin 2 |
| Spp2 | L1 | secreted phosphoprotein 2 |
| Pla2g12b | L1 | phospholipase A2 group XIIB |
| Serpind1 | L1 | serpin family D member 1 |
| F10 | L1 | coagulation factor X |
| Qprt | L1 | quinolinate phosphoribosyltransferase |
| Apoc2 | L1 | apolipoprotein C2 |
| Gchfr | L1 | GTP cyclohydrolase I feedback regulator |
| F12 | L1 | coagulation factor XII |
| Shbg | L1 | sex hormone binding globulin |
| Plg | L1 | plasminogen |
| Igfbp1 | L1 | insulin like growth factor binding protein 1 |
| Lcat | L1 | lecithin-cholesterol acyltransferase |
| Fga | L1 | fibrinogen alpha chain |
| Fabp1 | L2 | fatty acid binding protein 1 |
| Aadac | L2 | arylacetamide deacetylase |
| Ahsg | L2 | alpha 2-HS glycoprotein |
| Chst13 | L2 | carbohydrate sulfotransferase 13 |
| Kng1 | L2 | kininogen 1 |
| Sord | L2 | sorbitol dehydrogenase |
| Hsd17b2 | L2 | hydroxysteroid 17-beta dehydrogenase 2 |
| Creb3l3 | L2 | cAMP responsive element binding protein 3 like 3 |
| Cxcl10 | L2 | C-X-C motif chemokine ligand 10 |
| Icam1 | L2 | intercellular adhesion molecule 1 |
| Proz | L2 | protein Z, vitamin K dependent plasma glycoprotein |
| Hgd | L2 | homogentisate 1,2-dioxygenase |
| Agt | L2 | angiotensinogen |
| Blvrb | L2 | complement C1r |
| Hmgcs2 | L2 | 3-hydroxy-3-methylglutaryl-CoA synthase 2 |
| Lgals3bp | L2 | galectin 3 binding protein |
| Nostrin | L2 | nitric oxide synthase trafficking |
| G0s2 | L2 | G0/G1 switch 2 |
| Gltpd2 | L2 | glycolipid transfer protein domain containing 2 |

| | | |
|---|---|---|
| Slc6a13 | L2 | solute carrier family 6 member 13 |
| Apob | L2 | apolipoprotein B |
| Acss1 | L2 | acyl-CoA synthetase short chain family member 1 |
| Bdh1 | L2 | basic helix-loop-helix family member e40 |
| Ube2l6 | L2 | ubiquitin conjugating enzyme E2 L6 |
| Apoc4 | L2 | apolipoprotein C4 |
| Arhgap36 | L2 | Rho GTPase activating protein 36 |
| Leap2 | L2 | liver enriched antimicrobial peptide 2 |
| Naprt | L2 | nicotinate phosphoribosyltransferase |
| Aldob | L2 | aldolase, fructose-bisphosphate B |
| BC021614 | L2 | 3-hydroxybutyrate dehydrogenase 1 |
| Slc27a2 | L2 | solute carrier family 27 member 2 |
| Mbl1 | L2 | mannose-binding lectin (protein A) 1 |
| Saa4 | L2 | SAA2-SAA4 readthrough |
| Otc | L2 | ornithine carbamoyltransferase |
| Sardh | L2 | sarcosine dehydrogenase |
| Kng2 | L2 | kininogen 1 |
| Fabp2 | L2 | fatty acid binding protein 2 |
| Slc22a18 | L2 | solute carrier family 22 member 18 |
| Hebp1 | L2 | heme binding protein 1 |
| Bhlhe40 | L2 | biliverdin reductase B |
| Marc1 | L2 | mitochondrial amidoxime reducing component 1 |
| Slc36a2 | L2 | solute carrier family 36 member 2 |
| Cyp2c44 | L2 | - |
| Thnsl2 | L2 | threonine synthase like 2 |
| Ttc36 | L2 | tetratricopeptide repeat domain 36 |
| Hoga1 | L2 | 4-hydroxy-2-oxoglutarate aldolase 1 |
| Haao | L2 | 3-hydroxyanthranilate 3,4-dioxygenase |
| Mst1 | L2 | macrophage stimulating 1 |
| Hsd3b7 | L2 | hydroxy-delta-5-steroid dehydrogenase, 3 beta- and sterc |
| Apoh | L2 | apolipoprotein H |
| Tfr2 | L2 | transferrin receptor 2 |
| Itih4 | L2 | inter-alpha-trypsin inhibitor heavy chain family member 4 |
| Cyp2d10 | L2 | cytochrome P450, family 2, subfamily d, polypeptide 9 |
| Hba−a1 | L3 | hemoglobin subunit alpha 2 |
| Hba−a2 | L3 | hemoglobin subunit alpha 2 |
| Hbb−bs | L3 | hemoglobin subunit epsilon 1 |
| Hbb−bt | L3 | hemoglobin subunit gamma 2 |
| Hbb−y | L3 | hemoglobin subunit zeta |
| Hbb−bh1 | L3 | hemoglobin subunit beta |
| Hba−x | L3 | hemoglobin subunit beta |
| Miox | L3 | myo-inositol oxygenase |
| Ces2a | L3 | carboxylesterase 2A |
| Slc27a5 | L3 | solute carrier family 27 member 5 |
| Nlrp6 | L3 | NLR family pyrin domain containing 6 |
| Mocos | L3 | molybdenum cofactor sulfurase |
| Adra1b | L3 | adrenoceptor alpha 1B |
| Bbox1 | L3 | gamma-butyrobetaine hydroxylase 1 |
| Ropn1l | L3 | rhophilin associated tail protein 1 like |
| Crp | L3 | C-reactive protein |
| Sgk2 | L3 | SGK2, serine/threonine kinase 2 |
| Ngef | L3 | neuronal guanine nucleotide exchange factor |
| Cpt1a | L3 | carnitine palmitoyltransferase 1A |
| Serpina11 | L3 | serpin family A member 11 |
| Pdzk1 | L3 | PDZ domain containing 1 |
| C1ra | L3 | complement C3 |
| Pdk4 | L3 | pyruvate dehydrogenase kinase 4 |
| Slc17a9 | L3 | solute carrier family 17 member 9 |
| Rasgrp2 | L3 | RAS guanyl releasing protein 2 |

| | | |
|---|---|---|
| Cd40 | L3 | cell migration inducing hyaluronidase 2 |
| Kcnk2 | L3 | potassium two pore domain channel subfamily K member [illegible] |
| Prdm1 | G1 | PR/SET domain 1 |
| Nsg1 | G1 | neuronal vesicle trafficking associated 1 |
| Klk8 | G1 | kallikrein related peptidase 8 |
| Plat | G1 | plasminogen activator, tissue type |
| Hoxa1 | G1 | homeobox A1 |
| Grhl2 | G1 | grainyhead like transcription factor 2 |
| B4galt4 | G2 | beta-1,4-galactosyltransferase 4 |
| Sipa1l2 | G2 | signal induced proliferation associated 1 like 2 |
| Sox17 | G2 | SRY-box 17 |
| Adcy8 | G2 | adenylate cyclase 8 |
| Akap12 | G2 | A-kinase anchoring protein 12 |
| Flrt2 | G2 | fibronectin leucine rich transmembrane protein 2 |
| Igdcc3 | G2 | immunoglobulin superfamily DCC subclass member 3 |
| Clstn1 | G2 | calsyntenin 1 |
| Syt11 | G2 | synaptotagmin 11 |
| Map1b | G2 | microtubule associated protein 1B |
| Ddr1 | G2 | discoidin domain receptor tyrosine kinase 1 |
| Tmem2 | G2 | - |
| Kit | G2 | KIT proto-oncogene receptor tyrosine kinase |
| Hip1 | G2 | huntingtin interacting protein 1 |
| Igf1r | G2 | insulin like growth factor 1 receptor |
| Firre | G2 | functional intergenic repeating RNA element |
| Fgfr1 | G2 | fibroblast growth factor receptor 1 |
| Paxbp1 | G2 | PAX3 and PAX7 binding protein 1 |
| Dll1 | G2 | delta like canonical Notch ligand 1 |
| Epha4 | G2 | EPH receptor A4 |
| Lama1 | G2 | laminin subunit alpha 1 |
| Sema6a | G2 | semaphorin 6A |
| Eml1 | G2 | echinoderm microtubule associated protein like 1 |
| Dpysl4 | G2 | dihydropyrimidinase like 4 |
| Sh3pxd2b | G2 | SH3 and PX domains 2B |
| Sh3rf1 | G2 | SH3 domain containing ring finger 1 |
| Tctn2 | G2 | tectonic family member 2 |
| Meis2 | G2 | Meis homeobox 2 |
| Crb2 | G2 | crumbs 2, cell polarity complex component |
| Epha7 | G2 | EPH receptor A7 |
| Mpdz | G2 | multiple PDZ domain crumbs cell polarity complex compor |
| Ptpn13 | G2 | protein tyrosine phosphatase, non-receptor type 13 |
| Megf8 | G2 | multiple EGF like domains 8 |
| Hspg2 | G2 | heparan sulfate proteoglycan 2 |
| Itpr1 | G2 | inositol 1,4,5-trisphosphate receptor type 1 |
| Tead1 | G2 | TEA domain transcription factor 1 |
| Syne2 | G2 | spectrin repeat containing nuclear envelope protein 2 |
| Macf1 | G2 | microtubule-actin crosslinking factor 1 |
| Sfrp5 | G2 | secreted frizzled related protein 5 |
| Nepn | G2 | nephrocan |
| Capn6 | G2 | CD302 molecule |
| Ifitm1 | G2 | interferon induced transmembrane protein 1 |
| Fam183b | G2 | family with sequence similarity 183, member B |
| Cldn4 | G2 | claudin 4 |
| Gpx8 | G2 | glutathione peroxidase 8 (putative) |
| Igfbp5 | G2 | insulin like growth factor binding protein 5 |
| Hmgn3 | G2 | high mobility group nucleosomal binding domain 3 |
| Rem2 | G2 | RRAD and GEM like GTPase 2 |
| Rab25 | G2 | RAB25, member RAS oncogene family |
| Cldn3 | G2 | claudin 3 |
| Lxn | G2 | latexin |

| | | |
|---|---|---|
| Hsd11b2 | G2 | hydroxysteroid 11-beta dehydrogenase 2 |
| Zcchc18 | G2 | zinc finger CCHC-type containing 18 |
| Epb4.1l4a | G2 | - |
| Fzd2 | G2 | frizzled class receptor 2 |
| Pdpn | G2 | podoplanin |
| S100a10 | G2 | S100 calcium binding protein A10 |
| Spint1 | G2 | serine peptidase inhibitor, Kunitz type 1 |
| Lad1 | G2 | ladinin 1 |
| Ap1m2 | G2 | adaptor related protein complex 1 subunit mu 2 |
| Krt19 | G2 | keratin 19 |
| Crip2 | G2 | cysteine rich protein 2 |
| Serinc2 | G2 | serine incorporator 2 |
| Flrt3 | G2 | fibronectin leucine rich transmembrane protein 3 |
| Col4a2 | G2 | collagen type IV alpha 2 chain |
| Col2a1 | G2 | collagen type II alpha 1 chain |
| Igsf3 | G2 | immunoglobulin superfamily member 3 |
| Notch2 | G3 | notch 2 |
| Fstl1 | G3 | follistatin like 1 |
| Pxdn | G3 | peroxidasin |
| Col4a1 | G3 | collagen type IV alpha 1 chain |
| Epcam | G3 | epithelial cell adhesion molecule |
| Cldn6 | G3 | claudin 6 |
| Mapk13 | G3 | mitogen-activated protein kinase 13 |
| Wfdc2 | G3 | WAP four-disulfide core domain 2 |
| Nnat | G3 | neuronatin |
| Mfap2 | G3 | microfibril associated protein 2 |
| Sox11 | G3 | SRY-box 11 |
| H2afy2 | G3 | H2A histone family member Y2 |

#### (EHT)

| Location | Family |
|---|---|
| Nucleus | transcription regulator |
| Extracellular Space | other |
| Nucleus | transcription regulator |
| Cytoplasm | other |
| Other | other |
| Cytoplasm | other |
| Cytoplasm | enzyme |
| Cytoplasm | enzyme |
| Cytoplasm | other |
| Cytoplasm | enzyme |
| Other | other |
| Nucleus | transcription regulator |
| Other | growth factor |
| - | - |
| Plasma Membrane | ion channel |
| Extracellular Space | peptidase |
| Plasma Membrane | transmembrane receptor |
| Cytoplasm | enzyme |
| Cytoplasm | peptidase |
| Cytoplasm | enzyme |
| Extracellular Space | transporter |
| Extracellular Space | other |
| Cytoplasm | enzyme |
| Extracellular Space | other |
| Extracellular Space | other |
| Extracellular Space | enzyme |
| Extracellular Space | other |
| Extracellular Space | peptidase |
| Cytoplasm | enzyme |
| Extracellular Space | transporter |
| Cytoplasm | other |
| Extracellular Space | peptidase |
| Extracellular Space | other |
| Extracellular Space | peptidase |
| Extracellular Space | other |
| Extracellular Space | enzyme |
| Extracellular Space | other |
| Cytoplasm | transporter |
| Cytoplasm | enzyme |
| Extracellular Space | other |
| Cytoplasm | enzyme |
| Extracellular Space | other |
| Cytoplasm | enzyme |
| Cytoplasm | enzyme |
| Cytoplasm | transcription regulator |
| Extracellular Space | cytokine |
| Plasma Membrane | transmembrane receptor |
| Extracellular Space | peptidase |
| Cytoplasm | enzyme |
| Extracellular Space | growth factor |
| Extracellular Space | peptidase |
| Cytoplasm | enzyme |
| Plasma Membrane | transmembrane receptor |
| Cytoplasm | transcription regulator |
| Cytoplasm | other |
| Other | other |

| | |
|---|---|
| Plasma Membrane | transporter |
| Extracellular Space | transporter |
| Cytoplasm | enzyme |
| Nucleus | transcription regulator |
| Cytoplasm | enzyme |
| Extracellular Space | transporter |
| Cytoplasm | other |
| Extracellular Space | other |
| Cytoplasm | enzyme |
| Cytoplasm | enzyme |
| Cytoplasm | enzyme |
| Cytoplasm | transporter |
| Cytoplasm | other |
| Extracellular Space | transporter |
| Cytoplasm | enzyme |
| Cytoplasm | enzyme |
| Extracellular Space | other |
| Cytoplasm | transporter |
| Plasma Membrane | transporter |
| Cytoplasm | other |
| Cytoplasm | enzyme |
| Cytoplasm | enzyme |
| Plasma Membrane | transporter |
| - | - |
| Other | other |
| Other | other |
| Cytoplasm | enzyme |
| Cytoplasm | enzyme |
| Extracellular Space | growth factor |
| Cytoplasm | enzyme |
| Extracellular Space | transporter |
| Plasma Membrane | transporter |
| Extracellular Space | other |
| Cytoplasm | other |
| Extracellular Space | transporter |
| Extracellular Space | transporter |
| Cytoplasm | transporter |
| Cytoplasm | other |
| Cytoplasm | transporter |
| Cytoplasm | transporter |
| Cytoplasm | transporter |
| Cytoplasm | enzyme |
| Cytoplasm | enzyme |
| Cytoplasm | transporter |
| Plasma Membrane | G-protein coupled receptor |
| Cytoplasm | enzyme |
| Plasma Membrane | G-protein coupled receptor |
| Cytoplasm | enzyme |
| Cytoplasm | kinase |
| Extracellular Space | other |
| Cytoplasm | kinase |
| Cytoplasm | other |
| Cytoplasm | enzyme |
| Extracellular Space | other |
| Plasma Membrane | transporter |
| Extracellular Space | peptidase |
| Cytoplasm | kinase |
| Cytoplasm | other |
| Cytoplasm | other |

| | |
|---|---|
| Cytoplasm | enzyme |
| Plasma Membrane | ion channel |
| Nucleus | transcription regulator |
| Cytoplasm | other |
| Extracellular Space | peptidase |
| Extracellular Space | peptidase |
| Nucleus | transcription regulator |
| Nucleus | transcription regulator |
| Cytoplasm | enzyme |
| Other | other |
| Nucleus | transcription regulator |
| Plasma Membrane | enzyme |
| Cytoplasm | transporter |
| Plasma Membrane | other |
| Other | other |
| Plasma Membrane | other |
| Cytoplasm | transporter |
| Cytoplasm | other |
| Plasma Membrane | kinase |
| - | - |
| Plasma Membrane | transmembrane receptor |
| Cytoplasm | other |
| Plasma Membrane | transmembrane receptor |
| Other | other |
| Plasma Membrane | kinase |
| Nucleus | other |
| Plasma Membrane | enzyme |
| Plasma Membrane | kinase |
| Extracellular Space | other |
| Plasma Membrane | transmembrane receptor |
| Cytoplasm | other |
| Cytoplasm | enzyme |
| Cytoplasm | other |
| Cytoplasm | other |
| Extracellular Space | other |
| Nucleus | transcription regulator |
| Extracellular Space | other |
| Plasma Membrane | kinase |
| Plasma Membrane | other |
| Cytoplasm | phosphatase |
| Extracellular Space | other |
| Extracellular Space | enzyme |
| Cytoplasm | ion channel |
| Nucleus | transcription regulator |
| Nucleus | other |
| Cytoplasm | enzyme |
| Plasma Membrane | transmembrane receptor |
| Extracellular Space | other |
| Plasma Membrane | transmembrane receptor |
| Other | other |
| Extracellular Space | other |
| Plasma Membrane | transmembrane receptor |
| Cytoplasm | enzyme |
| Extracellular Space | other |
| Nucleus | other |
| Plasma Membrane | enzyme |
| Cytoplasm | enzyme |
| Plasma Membrane | transmembrane receptor |
| Cytoplasm | other |

| | |
|---|---|
| Cytoplasm | enzyme |
| Other | other |
| - | - |
| Plasma Membrane | G-protein coupled receptor |
| Plasma Membrane | other |
| Cytoplasm | other |
| Extracellular Space | other |
| Extracellular Space | other |
| Cytoplasm | transporter |
| Cytoplasm | other |
| Plasma Membrane | other |
| Plasma Membrane | transporter |
| Plasma Membrane | other |
| Extracellular Space | other |
| Extracellular Space | other |
| Plasma Membrane | other |
| Plasma Membrane | transcription regulator |
| Extracellular Space | other |
| Extracellular Space | enzyme |
| Extracellular Space | other |
| Plasma Membrane | other |
| Plasma Membrane | other |
| Cytoplasm | kinase |
| Extracellular Space | other |
| Plasma Membrane | transporter |
| Extracellular Space | other |
| Nucleus | transcription regulator |
| Nucleus | other |

### differentially expressed genes(DEGs) between liver pri

- **lb**:lower bound, **mle**:maximum likelihood estimate, **ub**: upper boun
- **ce:** conservative estimate of expression-fold change (equals to the r
- **Z:** uncorrected Z-score of expression difference
- **cZ:** expression difference Z-score corrected for multiple hypothesis
- **Genes highly expressed in the LP presented positive value of ce,**

| | lb | mle | ub | ce |
|---|---|---|---|---|
| ENSMUSG00000003153 | -3.263719739 | -2.575044932 | -1.94625489 | -1.94625489 |
| ENSMUSG00000032366 | -3.114007824 | -2.425333017 | -1.766600593 | -1.766600593 |
| ENSMUSG00000049154 | -7.335883817 | -6.766978541 | -5.988476585 | -5.988476585 |
| ENSMUSG00000005681 | 3.233777356 | 4.371587907 | 7.3658262 | 3.233777356 |
| ENSMUSG00000037254 | 3.143950207 | 4.40153029 | 5.748937522 | 3.143950207 |
| ENSMUSG00000033831 | 2.96429591 | 4.132048844 | 5.778879905 | 2.96429591 |
| ENSMUSG00000042453 | 2.934353527 | 3.89250978 | 5.060262715 | 2.934353527 |
| ENSMUSG00000002985 | 2.575044932 | 3.563143568 | 4.970435566 | 2.575044932 |
| ENSMUSG00000054932 | 1.616888678 | 3.922452163 | 4.281760758 | 1.616888678 |
| ENSMUSG00000040856 | 2.485217783 | 3.533201185 | 4.40153029 | 2.485217783 |
| ENSMUSG00000038224 | 2.545102549 | 3.952394546 | 5.539340841 | 2.545102549 |
| ENSMUSG00000024986 | 2.575044932 | 3.652970717 | 7.844904327 | 2.575044932 |
| ENSMUSG00000022346 | 2.425333017 | 3.353546888 | 5.030320332 | 2.425333017 |
| ENSMUSG00000067276 | -7.695192412 | -5.41957131 | -4.40153029 | -4.40153029 |
| ENSMUSG00000028780 | -7.814961944 | -7.186171902 | -4.072164078 | -4.072164078 |
| ENSMUSG00000038624 | -8.80306058 | -7.305941434 | -5.000377949 | -5.000377949 |
| ENSMUSG00000050953 | -5.030320332 | -3.563143568 | -2.724756846 | -2.724756846 |
| ENSMUSG00000023039 | -5.389628927 | -3.89250978 | -2.994238293 | -2.994238293 |
| ENSMUSG00000022892 | -5.000377949 | -3.6829131 | -2.634929698 | -2.634929698 |
| ENSMUSG00000000385 | -5.030320332 | -3.862567397 | -2.784641612 | -2.784641612 |
| ENSMUSG00000015568 | 2.215736337 | 2.96429591 | 3.922452163 | 2.215736337 |
| ENSMUSG00000027533 | 1.467176763 | 2.066024422 | 2.724756846 | 1.467176763 |
| ENSMUSG00000002633 | -7.335883817 | -6.527439478 | -5.090205097 | -5.090205097 |
| ENSMUSG00000029761 | 2.155851571 | 3.114007824 | 4.221875993 | 2.155851571 |
| ENSMUSG00000010175 | 2.275621102 | 3.233777356 | 4.431472673 | 2.275621102 |
| ENSMUSG00000044337 | -6.976575222 | -6.1381885 | -5.299801778 | -5.299801778 |
| ENSMUSG00000025491 | -7.335883817 | -3.6829131 | -2.784641612 | -2.784641612 |
| ENSMUSG00000026235 | -7.515538114 | -6.737036158 | -5.150089863 | -5.150089863 |
| ENSMUSG00000029844 | -6.737036158 | -5.838764671 | -5.030320332 | -5.030320332 |
| ENSMUSG00000027684 | -7.395768583 | -6.317842797 | -5.000377949 | -5.000377949 |
| ENSMUSG00000002265 | -2.934353527 | -2.275621102 | -1.676773444 | -1.676773444 |
| ENSMUSG00000048387 | -7.814961944 | -6.617266627 | -5.030320332 | -5.030320332 |
| ENSMUSG00000047139 | -2.814583995 | -2.095966805 | -1.497119146 | -1.497119146 |
| ENSMUSG00000019900 | -7.695192412 | -7.006517605 | -3.353546888 | -3.353546888 |
| ENSMUSG00000031538 | -6.946632839 | -6.108246117 | -5.000377949 | -5.000377949 |
| ENSMUSG00000075394 | -6.587324244 | -5.718995139 | -4.880608417 | -4.880608417 |
| ENSMUSG00000021850 | -6.557381861 | -5.748937522 | -4.940493183 | -4.940493183 |
| ENSMUSG00000039385 | -4.730896502 | -3.772740249 | -2.365448251 | -2.365448251 |
| ENSMUSG00000029153 | -6.826863307 | -5.718995139 | -3.802682632 | -3.802682632 |
| ENSMUSG00000022286 | -7.096344754 | -6.228015649 | -5.000377949 | -5.000377949 |
| ENSMUSG00000050520 | -6.437612329 | -5.41957131 | -4.581184588 | -4.581184588 |
| ENSMUSG00000000303 | -4.641069354 | -3.383489271 | -2.275621102 | -2.275621102 |
| ENSMUSG00000019987 | -3.114007824 | -2.275621102 | -1.586946295 | -1.586946295 |
| ENSMUSG00000031740 | -6.617266627 | -5.539340841 | -4.700954119 | -4.700954119 |
| ENSMUSG00000044186 | -6.34778518 | -5.509398458 | -4.461415056 | -4.461415056 |
| ENSMUSG00000046733 | -6.916690456 | -6.018418968 | -4.461415056 | -4.461415056 |
| ENSMUSG00000022330 | -5.060262715 | -4.19193361 | -3.084065441 | -3.084065441 |
| ENSMUSG00000053219 | -7.036459988 | -6.257958032 | -4.341645524 | -4.341645524 |

| | | | | |
|---|---|---|---|---|
| ENSMUSG00000075334 | -6.257958032 | -5.030320332 | -4.102106461 | -4.102106461 |
| ENSMUSG00000026639 | -6.85680569 | -5.838764671 | -4.581184588 | -4.581184588 |
| ENSMUSG00000005148 | -6.437612329 | -5.359686544 | -4.371587907 | -4.371587907 |
| ENSMUSG00000014773 | -6.108246117 | -5.239917012 | -4.341645524 | -4.341645524 |
| ENSMUSG00000039323 | -3.742797866 | -2.634929698 | -1.766600593 | -1.766600593 |
| ENSMUSG00000022577 | -6.078303734 | -5.389628927 | -4.551242205 | -4.551242205 |
| ENSMUSG00000031502 | -3.413431654 | -2.245678719 | -1.527061529 | -1.527061529 |
| ENSMUSG00000030551 | -4.161991227 | -3.084065441 | -2.066024422 | -2.066024422 |
| ENSMUSG00000031503 | -4.341645524 | -3.17389259 | -2.066024422 | -2.066024422 |
| ENSMUSG00000023886 | -5.988476585 | -4.641069354 | -3.832625015 | -3.832625015 |
| ENSMUSG00000019909 | -5.329744161 | -4.251818376 | -3.233777356 | -3.233777356 |
| ENSMUSG00000028173 | -4.700954119 | -3.383489271 | -2.335505868 | -2.335505868 |
| ENSMUSG00000031517 | -5.62916799 | -4.431472673 | -3.413431654 | -3.413431654 |
| ENSMUSG00000026765 | -6.1381885 | -4.9105508 | -3.862567397 | -3.862567397 |
| ENSMUSG00000037995 | -3.802682632 | -2.545102549 | -1.676773444 | -1.676773444 |
| ENSMUSG00000005220 | -6.048361351 | -4.820723651 | -3.862567397 | -3.862567397 |
| ENSMUSG00000031028 | -5.389628927 | -4.491357439 | -3.623028334 | -3.623028334 |
| ENSMUSG00000038151 | -5.988476585 | -5.030320332 | -4.102106461 | -4.102106461 |
| ENSMUSG00000028356 | 1.856427741 | 2.934353527 | 4.671011736 | 1.856427741 |
| ENSMUSG00000038094 | -4.940493183 | -4.132048844 | -3.114007824 | -3.114007824 |
| ENSMUSG00000002980 | -3.802682632 | -2.545102549 | -1.676773444 | -1.676773444 |
| ENSMUSG00000032334 | -5.958534202 | -4.940493183 | -3.802682632 | -3.802682632 |
| ENSMUSG00000024713 | -5.718995139 | -4.730896502 | -3.742797866 | -3.742797866 |
| ENSMUSG00000018604 | 1.886370124 | 2.754699229 | 4.341645524 | 1.886370124 |
| ENSMUSG00000001943 | -5.299801778 | -4.281760758 | -3.293662122 | -3.293662122 |
| ENSMUSG00000043924 | -5.090205097 | -4.161991227 | -3.143950207 | -3.143950207 |
| ENSMUSG00000025969 | -7.485595732 | -4.790781268 | -3.89250978 | -3.89250978 |
| ENSMUSG00000034463 | -4.9105508 | -4.072164078 | -2.934353527 | -2.934353527 |
| ENSMUSG00000019823 | -5.539340841 | -4.611126971 | -3.712855483 | -3.712855483 |
| ENSMUSG00000047501 | -4.850666034 | -2.994238293 | -2.006139656 | -2.006139656 |
| ENSMUSG00000041309 | -5.509398458 | -4.581184588 | -3.6829131 | -3.6829131 |
| ENSMUSG00000061808 | 1.766600593 | 2.874468761 | 4.341645524 | 1.766600593 |
| ENSMUSG00000042808 | -5.898649436 | -4.940493183 | -3.712855483 | -3.712855483 |
| ENSMUSG00000021953 | -6.34778518 | -4.790781268 | -3.982336929 | -3.982336929 |
| ENSMUSG00000029644 | -5.808822288 | -4.700954119 | -3.6829131 | -3.6829131 |
| ENSMUSG00000028023 | -6.946632839 | -4.790781268 | -3.922452163 | -3.922452163 |
| ENSMUSG00000020297 | -5.659110373 | -4.431472673 | -3.353546888 | -3.353546888 |
| ENSMUSG00000046546 | -6.467554712 | -5.329744161 | -4.311703141 | -4.311703141 |
| ENSMUSG00000090439 | -5.000377949 | -3.952394546 | -3.024180676 | -3.024180676 |
| ENSMUSG00000048502 | -4.940493183 | -3.652970717 | -2.515160166 | -2.515160166 |
| ENSMUSG00000022061 | -4.970435566 | -4.072164078 | -2.874468761 | -2.874468761 |
| ENSMUSG00000019647 | -7.425710966 | -3.503258802 | -2.515160166 | -2.515160166 |
| ENSMUSG00000020357 | -5.509398458 | -4.611126971 | -3.593085951 | -3.593085951 |
| ENSMUSG00000000753 | 2.036082039 | 3.203834973 | 5.239917012 | 2.036082039 |
| ENSMUSG00000024897 | -4.970435566 | -3.862567397 | -2.784641612 | -2.784641612 |
| ENSMUSG00000057615 | -5.030320332 | -4.042221695 | -3.143950207 | -3.143950207 |
| ENSMUSG00000074505 | -5.41957131 | -4.641069354 | -3.263719739 | -3.263719739 |
| ENSMUSG00000031621 | -5.479456076 | -4.551242205 | -3.593085951 | -3.593085951 |
| ENSMUSG00000060487 | -5.239917012 | -4.19193361 | -3.054123058 | -3.054123058 |
| ENSMUSG00000022270 | -5.389628927 | -4.521299822 | -3.503258802 | -3.503258802 |
| ENSMUSG00000024036 | -5.090205097 | -4.251818376 | -2.904411144 | -2.904411144 |
| ENSMUSG00000020173 | -5.41957131 | -3.802682632 | -2.724756846 | -2.724756846 |
| ENSMUSG00000072672 | -5.12014748 | -3.862567397 | -2.724756846 | -2.724756846 |
| ENSMUSG00000031891 | -6.377727563 | -4.491357439 | -2.754699229 | -2.754699229 |
| ENSMUSG00000025921 | -7.126287136 | -6.078303734 | -3.982336929 | -3.982336929 |
| ENSMUSG00000026866 | -5.12014748 | -4.311703141 | -3.203834973 | -3.203834973 |
| ENSMUSG00000022485 | -5.090205097 | -4.161991227 | -3.084065441 | -3.084065441 |
| ENSMUSG00000038175 | -5.868707054 | -4.940493183 | -3.772740249 | -3.772740249 |
| ENSMUSG00000048281 | -4.491357439 | -3.323604505 | -2.155851571 | -2.155851571 |

| | | | | |
|---|---|---|---|---|
| ENSMUSG00000041608 | -6.527439478 | -5.359686544 | -4.161991227 | -4.161991227 |
| ENSMUSG00000043496 | -5.748937522 | -4.970435566 | -3.772740249 | -3.772740249 |
| ENSMUSG00000025473 | -4.9105508 | -4.132048844 | -2.784641612 | -2.784641612 |
| ENSMUSG00000028289 | -6.946632839 | -3.233777356 | -2.155851571 | -2.155851571 |
| ENSMUSG00000024206 | -5.299801778 | -4.311703141 | -3.233777356 | -3.233777356 |
| ENSMUSG00000041605 | -6.796920924 | -6.078303734 | -3.233777356 | -3.233777356 |
| ENSMUSG00000028716 | -5.269859395 | -4.371587907 | -3.233777356 | -3.233777356 |
| ENSMUSG00000037016 | -4.012279312 | -2.994238293 | -2.036082039 | -2.036082039 |
| ENSMUSG00000039081 | -5.239917012 | -4.341645524 | -3.024180676 | -3.024180676 |
| ENSMUSG00000072675 | -5.030320332 | -3.383489271 | -2.395390634 | -2.395390634 |
| ENSMUSG00000042607 | 1.94625489 | 3.17389259 | 4.521299822 | 1.94625489 |
| ENSMUSG00000044254 | 1.616888678 | 2.604987315 | 3.952394546 | 1.616888678 |
| ENSMUSG00000026796 | -6.826863307 | -2.96429591 | -1.826485359 | -1.826485359 |
| ENSMUSG00000026748 | -5.180032246 | -4.221875993 | -3.024180676 | -3.024180676 |
| ENSMUSG00000053398 | 1.018041019 | 1.646831061 | 2.335505868 | 1.018041019 |
| ENSMUSG00000075588 | -5.659110373 | -4.940493183 | -3.233777356 | -3.233777356 |
| ENSMUSG00000036832 | -7.006517605 | -6.377727563 | -3.383489271 | -3.383489271 |
| ENSMUSG00000013418 | -5.209974629 | -4.251818376 | -3.024180676 | -3.024180676 |
| ENSMUSG00000021087 | -5.359686544 | -4.611126971 | -3.503258802 | -3.503258802 |
| ENSMUSG00000035580 | -5.389628927 | -4.461415056 | -2.96429591 | -2.96429591 |
| ENSMUSG00000033805 | -4.880608417 | -3.982336929 | -2.335505868 | -2.335505868 |
| ENSMUSG00000022816 | -3.533201185 | -2.365448251 | -1.43723438 | -1.43723438 |
| ENSMUSG00000022203 | -6.707093775 | -6.168130883 | -2.4552754 | -2.4552754 |
| ENSMUSG00000039904 | -5.150089863 | -4.19193361 | -2.934353527 | -2.934353527 |
| ENSMUSG00000050994 | -4.880608417 | -3.862567397 | -2.575044932 | -2.575044932 |
| ENSMUSG00000070867 | -5.659110373 | -4.611126971 | -3.533201185 | -3.533201185 |
| ENSMUSG00000054162 | -4.970435566 | -3.982336929 | -2.724756846 | -2.724756846 |
| ENSMUSG00000052430 | -5.180032246 | -4.311703141 | -2.96429591 | -2.96429591 |
| ENSMUSG00000038602 | -4.970435566 | -4.042221695 | -2.814583995 | -2.814583995 |
| ENSMUSG00000037664 | 1.287522466 | 2.185793954 | 3.143950207 | 1.287522466 |
| ENSMUSG00000002900 | -3.203834973 | -2.006139656 | -1.197695317 | -1.197695317 |
| ENSMUSG00000066720 | -6.1381885 | -5.389628927 | -3.293662122 | -3.293662122 |
| ENSMUSG00000026494 | -5.838764671 | -4.790781268 | -3.593085951 | -3.593085951 |
| ENSMUSG00000026014 | 1.257580083 | 2.036082039 | 2.814583995 | 1.257580083 |
| ENSMUSG00000055320 | -6.34778518 | -5.599225607 | -3.323604505 | -3.323604505 |
| ENSMUSG00000042834 | -3.263719739 | -2.365448251 | -1.527061529 | -1.527061529 |
| ENSMUSG00000024140 | -4.671011736 | -3.832625015 | -2.545102549 | -2.545102549 |
| ENSMUSG00000070348 | 1.018041019 | 1.73665821 | 2.694814463 | 1.018041019 |
| ENSMUSG00000034127 | -5.12014748 | -4.341645524 | -3.233777356 | -3.233777356 |
| ENSMUSG00000037820 | 1.2276377 | 2.365448251 | 3.952394546 | 1.2276377 |
| ENSMUSG00000096001 | -4.880608417 | -4.072164078 | -2.844526378 | -2.844526378 |
| ENSMUSG00000007021 | -5.030320332 | -4.042221695 | -2.844526378 | -2.844526378 |
| ENSMUSG00000073207 | -4.671011736 | -3.89250978 | -2.634929698 | -2.634929698 |
| ENSMUSG00000033032 | -6.946632839 | -6.228015649 | -3.17389259 | -3.17389259 |
| ENSMUSG00000025608 | -2.994238293 | -2.036082039 | -1.197695317 | -1.197695317 |
| ENSMUSG00000020911 | -3.533201185 | -2.305563485 | -1.317464849 | -1.317464849 |
| ENSMUSG00000029223 | -2.994238293 | -2.006139656 | -1.2276377 | -1.2276377 |
| ENSMUSG00000074500 | -5.569283224 | -4.311703141 | -3.114007824 | -3.114007824 |
| ENSMUSG00000028167 | -6.34778518 | -5.329744161 | -3.652970717 | -3.652970717 |
| ENSMUSG00000028039 | -4.641069354 | -3.623028334 | -2.365448251 | -2.365448251 |
| ENSMUSG00000032179 | -4.940493183 | -4.102106461 | -2.275621102 | -2.275621102 |
| ENSMUSG00000032515 | -5.539340841 | -4.491357439 | -3.323604505 | -3.323604505 |
| ENSMUSG00000054667 | -5.509398458 | -4.671011736 | -3.413431654 | -3.413431654 |
| ENSMUSG00000001663 | 1.73665821 | 2.694814463 | 6.228015649 | 1.73665821 |
| ENSMUSG00000027562 | -6.737036158 | -5.718995139 | -3.862567397 | -3.862567397 |
| ENSMUSG00000024501 | -5.868707054 | -4.760838885 | -3.563143568 | -3.563143568 |
| ENSMUSG00000026773 | 1.167752934 | 1.94625489 | 2.724756846 | 1.167752934 |
| ENSMUSG00000069833 | -5.509398458 | -4.521299822 | -3.203834973 | -3.203834973 |
| ENSMUSG00000024044 | -3.6829131 | -2.365448251 | -1.407291998 | -1.407291998 |

| | | | | |
|---|---|---|---|---|
| ENSMUSG00000064036 | -4.491357439 | -3.712855483 | -2.305563485 | -2.305563485 |
| ENSMUSG00000006538 | -5.748937522 | -4.9105508 | -3.623028334 | -3.623028334 |
| ENSMUSG00000049313 | -6.677151393 | -4.012279312 | -2.245678719 | -2.245678719 |
| ENSMUSG00000033860 | 1.796542976 | 3.203834973 | 7.156229519 | 1.796542976 |
| ENSMUSG00000041936 | -2.814583995 | -1.94625489 | -1.167752934 | -1.167752934 |
| ENSMUSG00000036306 | 1.766600593 | 2.694814463 | 6.317842797 | 1.766600593 |
| ENSMUSG00000078452 | -6.497497095 | -5.808822288 | -2.874468761 | -2.874468761 |
| ENSMUSG00000049946 | -4.730896502 | -3.623028334 | -1.766600593 | -1.766600593 |
| ENSMUSG00000075297 | -4.641069354 | -3.084065441 | -1.616888678 | -1.616888678 |
| ENSMUSG00000026380 | -5.12014748 | -4.461415056 | -3.263719739 | -3.263719739 |
| ENSMUSG00000049690 | -5.299801778 | -4.551242205 | -3.203834973 | -3.203834973 |
| ENSMUSG00000028532 | -5.958534202 | -5.090205097 | -3.623028334 | -3.623028334 |
| ENSMUSG00000063838 | -4.940493183 | -4.072164078 | -2.904411144 | -2.904411144 |
| ENSMUSG00000071178 | 1.676773444 | 3.233777356 | 4.221875993 | 1.676773444 |
| ENSMUSG00000025932 | -5.359686544 | -4.281760758 | -2.934353527 | -2.934353527 |
| ENSMUSG00000045394 | -2.754699229 | -1.766600593 | -1.018041019 | -1.018041019 |
| ENSMUSG00000050737 | -4.9105508 | -3.89250978 | -2.275621102 | -2.275621102 |
| ENSMUSG00000032363 | -5.599225607 | -4.700954119 | -3.263719739 | -3.263719739 |
| ENSMUSG00000049929 | -5.060262715 | -4.281760758 | -3.084065441 | -3.084065441 |
| ENSMUSG00000022537 | -6.257958032 | -5.479456076 | -2.036082039 | -2.036082039 |
| ENSMUSG00000032079 | 1.856427741 | 3.6829131 | 4.700954119 | 1.856427741 |
| ENSMUSG00000022382 | -5.838764671 | -4.551242205 | -3.533201185 | -3.533201185 |
| ENSMUSG00000007908 | -5.928591819 | -4.820723651 | -3.383489271 | -3.383489271 |
| ENSMUSG00000020806 | -5.000377949 | -4.132048844 | -2.844526378 | -2.844526378 |
| ENSMUSG00000036158 | -6.228015649 | -5.239917012 | -3.054123058 | -3.054123058 |
| ENSMUSG00000079445 | -6.108246117 | -5.239917012 | -3.413431654 | -3.413431654 |
| ENSMUSG00000047485 | -4.40153029 | -3.323604505 | -1.856427741 | -1.856427741 |
| ENSMUSG00000029167 | -5.090205097 | -4.40153029 | -2.604987315 | -2.604987315 |
| ENSMUSG00000058135 | 1.107868168 | 1.94625489 | 2.934353527 | 1.107868168 |
| ENSMUSG00000048376 | -3.413431654 | -1.976197273 | -1.107868168 | -1.107868168 |
| ENSMUSG00000042359 | -6.677151393 | -6.168130883 | -2.036082039 | -2.036082039 |
| ENSMUSG00000031351 | -5.569283224 | -4.671011736 | -3.413431654 | -3.413431654 |
| ENSMUSG00000022099 | -5.030320332 | -3.802682632 | -2.395390634 | -2.395390634 |
| ENSMUSG00000063428 | -4.9105508 | -3.952394546 | -2.245678719 | -2.245678719 |
| ENSMUSG00000070473 | -6.677151393 | -2.515160166 | -1.616888678 | -1.616888678 |
| ENSMUSG00000028737 | 1.167752934 | 1.976197273 | 3.024180676 | 1.167752934 |
| ENSMUSG00000103677 | -4.491357439 | -3.473316419 | -1.2276377 | -1.2276377 |
| ENSMUSG00000022483 | -6.976575222 | -5.030320332 | -2.365448251 | -2.365448251 |
| ENSMUSG00000001946 | -4.880608417 | -3.593085951 | -2.066024422 | -2.066024422 |
| ENSMUSG00000047443 | -4.880608417 | -4.012279312 | -2.155851571 | -2.155851571 |
| ENSMUSG00000037440 | -7.216114285 | -6.497497095 | -2.575044932 | -2.575044932 |
| ENSMUSG00000064225 | 1.377349615 | 2.335505868 | 5.299801778 | 1.377349615 |
| ENSMUSG00000072849 | 1.73665821 | 2.994238293 | 4.042221695 | 1.73665821 |
| ENSMUSG00000058498 | -5.12014748 | -4.281760758 | -2.96429591 | -2.96429591 |
| ENSMUSG00000052397 | -3.922452163 | -2.754699229 | -1.557003912 | -1.557003912 |
| ENSMUSG00000079606 | 2.694814463 | 3.832625015 | 5.269859395 | 2.694814463 |
| ENSMUSG00000041907 | -4.700954119 | -3.593085951 | -2.006139656 | -2.006139656 |
| ENSMUSG00000031952 | -4.940493183 | -3.922452163 | -2.694814463 | -2.694814463 |
| ENSMUSG00000029446 | 1.167752934 | 2.095966805 | 2.994238293 | 1.167752934 |
| ENSMUSG00000031995 | -3.593085951 | -2.185793954 | -1.2276377 | -1.2276377 |
| ENSMUSG00000047986 | -6.1381885 | -3.503258802 | -2.006139656 | -2.006139656 |
| ENSMUSG00000042258 | -6.617266627 | -5.389628927 | -3.233777356 | -3.233777356 |
| ENSMUSG00000042190 | -4.461415056 | -3.593085951 | -1.467176763 | -1.467176763 |
| ENSMUSG00000032232 | -4.521299822 | -2.634929698 | -1.616888678 | -1.616888678 |
| ENSMUSG00000058145 | -4.730896502 | -3.89250978 | -2.515160166 | -2.515160166 |
| ENSMUSG00000022665 | -6.497497095 | -5.659110373 | -2.335505868 | -2.335505868 |
| ENSMUSG00000001870 | -4.730896502 | -2.575044932 | -1.407291998 | -1.407291998 |
| ENSMUSG00000018822 | -7.156229519 | -4.251818376 | -2.185793954 | -2.185793954 |
| ENSMUSG00000010660 | -6.557381861 | -5.599225607 | -2.485217783 | -2.485217783 |

| | | | | |
|---|---|---|---|---|
| ENSMUSG00000057137 | -4.431472673 | -3.17389259 | -1.047983402 | -1.047983402 |
| ENSMUSG00000022240 | -4.671011736 | -3.6829131 | -2.155851571 | -2.155851571 |
| ENSMUSG00000025885 | -5.180032246 | -4.461415056 | -2.874468761 | -2.874468761 |
| ENSMUSG00000066705 | -4.102106461 | -2.814583995 | -1.646831061 | -1.646831061 |
| ENSMUSG00000029369 | 1.916312507 | 5.778879905 | 6.766978541 | 1.916312507 |
| ENSMUSG00000038587 | -6.198073266 | -5.329744161 | -2.4552754 | -2.4552754 |
| ENSMUSG00000057751 | -5.359686544 | -4.341645524 | -2.814583995 | -2.814583995 |
| ENSMUSG00000018398 | -5.41957131 | -4.491357439 | -3.024180676 | -3.024180676 |
| ENSMUSG00000057092 | -4.341645524 | -3.413431654 | -1.257580083 | -1.257580083 |
| ENSMUSG00000034382 | -4.641069354 | -3.413431654 | -2.095966805 | -2.095966805 |
| ENSMUSG00000051159 | 1.107868168 | 2.095966805 | 3.17389259 | 1.107868168 |
| ENSMUSG00000046204 | -5.060262715 | -4.132048844 | -2.994238293 | -2.994238293 |
| ENSMUSG00000029368 | 1.557003912 | 2.934353527 | 6.976575222 | 1.557003912 |
| ENSMUSG00000022150 | -5.958534202 | -2.874468761 | -1.527061529 | -1.527061529 |
| ENSMUSG00000047261 | 1.257580083 | 2.245678719 | 4.581184588 | 1.257580083 |
| ENSMUSG00000000253 | -6.078303734 | -4.9105508 | -2.934353527 | -2.934353527 |
| ENSMUSG00000008136 | -7.006517605 | -6.34778518 | -2.125909188 | -2.125909188 |
| ENSMUSG00000035517 | -5.449513693 | -4.281760758 | -2.784641612 | -2.784641612 |
| ENSMUSG00000024500 | -6.228015649 | -5.12014748 | -2.425333017 | -2.425333017 |
| ENSMUSG00000020695 | -5.000377949 | -4.221875993 | -2.515160166 | -2.515160166 |
| ENSMUSG00000066366 | 1.407291998 | 2.814583995 | 3.832625015 | 1.407291998 |
| ENSMUSG00000004961 | -6.34778518 | -5.509398458 | -2.066024422 | -2.066024422 |
| ENSMUSG00000034687 | -3.652970717 | -2.604987315 | -1.43723438 | -1.43723438 |
| ENSMUSG00000031444 | 2.634929698 | 5.239917012 | 6.407669946 | 2.634929698 |
| ENSMUSG00000051652 | 1.347407232 | 2.485217783 | 5.62916799 | 1.347407232 |
| ENSMUSG00000020902 | -5.479456076 | -4.281760758 | -2.515160166 | -2.515160166 |
| ENSMUSG00000010080 | -5.778879905 | -5.000377949 | -2.634929698 | -2.634929698 |
| ENSMUSG00000041782 | -3.6829131 | -2.545102549 | -1.257580083 | -1.257580083 |
| ENSMUSG00000060314 | -4.820723651 | -3.89250978 | -2.275621102 | -2.275621102 |
| ENSMUSG00000039239 | -6.467554712 | -4.730896502 | -2.485217783 | -2.485217783 |
| ENSMUSG00000061288 | 1.257580083 | 2.215736337 | 3.383489271 | 1.257580083 |
| ENSMUSG00000040152 | -6.018418968 | -5.000377949 | -2.305563485 | -2.305563485 |
| ENSMUSG00000041362 | -6.078303734 | -5.449513693 | -2.006139656 | -2.006139656 |
| ENSMUSG00000040728 | -3.652970717 | -2.066024422 | -1.047983402 | -1.047983402 |
| ENSMUSG00000027954 | -4.281760758 | -2.515160166 | -1.317464849 | -1.317464849 |
| ENSMUSG00000060206 | -5.988476585 | -2.4552754 | -1.377349615 | -1.377349615 |
| ENSMUSG00000021703 | -5.329744161 | -4.40153029 | -2.814583995 | -2.814583995 |
| ENSMUSG00000019810 | -4.102106461 | -2.275621102 | -1.167752934 | -1.167752934 |
| ENSMUSG00000021057 | -5.868707054 | -3.533201185 | -1.796542976 | -1.796542976 |
| ENSMUSG00000056900 | -5.449513693 | -4.371587907 | -2.245678719 | -2.245678719 |
| ENSMUSG00000055675 | -4.760838885 | -3.952394546 | -1.2276377 | -1.2276377 |
| ENSMUSG00000047557 | -4.611126971 | -2.96429591 | -1.407291998 | -1.407291998 |
| ENSMUSG00000029228 | -6.1381885 | -5.269859395 | -2.395390634 | -2.395390634 |
| ENSMUSG00000066687 | -5.329744161 | -4.341645524 | -2.694814463 | -2.694814463 |
| ENSMUSG00000027993 | -5.988476585 | -4.042221695 | -2.036082039 | -2.036082039 |
| ENSMUSG00000028019 | -4.790781268 | -2.604987315 | -1.317464849 | -1.317464849 |
| ENSMUSG00000026715 | 1.197695317 | 2.425333017 | 6.078303734 | 1.197695317 |
| ENSMUSG00000057969 | -6.1381885 | -5.12014748 | -2.365448251 | -2.365448251 |
| ENSMUSG00000029304 | -5.090205097 | -4.042221695 | -2.575044932 | -2.575044932 |
| ENSMUSG00000039620 | -6.34778518 | -5.479456076 | -2.994238293 | -2.994238293 |
| ENSMUSG00000032796 | -6.497497095 | -2.215736337 | -1.2276377 | -1.2276377 |
| ENSMUSG00000001507 | -3.802682632 | -2.634929698 | -1.317464849 | -1.317464849 |
| ENSMUSG00000032083 | 1.137810551 | 2.634929698 | 3.563143568 | 1.137810551 |
| ENSMUSG00000059602 | -4.431472673 | -3.443374037 | -1.676773444 | -1.676773444 |
| ENSMUSG00000032014 | 1.137810551 | 2.095966805 | 3.233777356 | 1.137810551 |
| ENSMUSG00000006356 | -4.730896502 | -2.994238293 | -1.467176763 | -1.467176763 |
| ENSMUSG00000032394 | -4.042221695 | -2.425333017 | -1.257580083 | -1.257580083 |
| ENSMUSG00000020150 | 1.018041019 | 1.916312507 | 3.772740249 | 1.018041019 |
| ENSMUSG00000031290 | -4.40153029 | -3.473316419 | -2.066024422 | -2.066024422 |

| | | | | |
|---|---|---|---|---|
| ENSMUSG00000032718 | -4.581184588 | -3.623028334 | -2.215736337 | -2.215736337 |
| ENSMUSG00000001300 | -5.090205097 | -2.814583995 | -1.407291998 | -1.407291998 |
| ENSMUSG00000001827 | -5.060262715 | -4.19193361 | -2.305563485 | -2.305563485 |
| ENSMUSG00000020601 | -6.287900415 | -5.389628927 | -1.886370124 | -1.886370124 |
| ENSMUSG00000031770 | 1.018041019 | 1.976197273 | 2.934353527 | 1.018041019 |
| ENSMUSG00000062184 | -6.437612329 | -5.748937522 | -1.706715827 | -1.706715827 |
| ENSMUSG00000068699 | -5.898649436 | -4.461415056 | -2.694814463 | -2.694814463 |
| ENSMUSG00000022766 | 1.706715827 | 5.389628927 | 6.587324244 | 1.706715827 |
| ENSMUSG00000029108 | -5.12014748 | -4.311703141 | -2.66487208 | -2.66487208 |
| ENSMUSG00000028100 | -5.748937522 | -4.880608417 | -2.904411144 | -2.904411144 |
| ENSMUSG00000023055 | -3.89250978 | -2.365448251 | -1.2276377 | -1.2276377 |
| ENSMUSG00000039542 | -5.599225607 | -4.671011736 | -2.365448251 | -2.365448251 |
| ENSMUSG00000034917 | -6.377727563 | -5.689052756 | -1.527061529 | -1.527061529 |
| ENSMUSG00000084174 | -5.449513693 | -4.19193361 | -2.4552754 | -2.4552754 |
| ENSMUSG00000041046 | -5.898649436 | -4.760838885 | -2.4552754 | -2.4552754 |
| ENSMUSG00000021318 | -5.778879905 | -2.814583995 | -1.467176763 | -1.467176763 |
| ENSMUSG00000070883 | -4.700954119 | -3.712855483 | -1.976197273 | -1.976197273 |
| ENSMUSG00000066798 | -5.000377949 | -3.832625015 | -2.125909188 | -2.125909188 |
| ENSMUSG00000022376 | -6.826863307 | -5.479456076 | -1.467176763 | -1.467176763 |
| ENSMUSG00000075254 | 1.2276377 | 2.275621102 | 3.503258802 | 1.2276377 |
| ENSMUSG00000042035 | -3.263719739 | -2.155851571 | -1.018041019 | -1.018041019 |
| ENSMUSG00000055980 | -5.389628927 | -4.281760758 | -2.215736337 | -2.215736337 |
| ENSMUSG00000024485 | -4.970435566 | -4.042221695 | -2.66487208 | -2.66487208 |
| ENSMUSG00000068962 | -4.551242205 | -3.084065441 | -1.796542976 | -1.796542976 |
| ENSMUSG00000079343 | 1.018041019 | 2.724756846 | 4.132048844 | 1.018041019 |
| ENSMUSG00000022951 | -5.449513693 | -2.425333017 | -1.257580083 | -1.257580083 |
| ENSMUSG00000079481 | -5.958534202 | -5.000377949 | -2.125909188 | -2.125909188 |
| ENSMUSG00000033060 | -6.228015649 | -5.060262715 | -2.095966805 | -2.095966805 |
| ENSMUSG00000036242 | -5.389628927 | -3.233777356 | -1.407291998 | -1.407291998 |
| ENSMUSG00000015133 | -5.988476585 | -5.060262715 | -2.095966805 | -2.095966805 |
| ENSMUSG00000047631 | 1.377349615 | 5.41957131 | 6.108246117 | 1.377349615 |
| ENSMUSG00000029826 | 2.185793954 | 4.431472673 | 5.599225607 | 2.185793954 |
| ENSMUSG00000017057 | -6.108246117 | -5.030320332 | -2.006139656 | -2.006139656 |
| ENSMUSG00000057286 | -6.407669946 | -5.359686544 | -1.676773444 | -1.676773444 |
| ENSMUSG00000025188 | -6.257958032 | -5.389628927 | -1.766600593 | -1.766600593 |
| ENSMUSG00000031434 | -5.569283224 | -2.754699229 | -1.197695317 | -1.197695317 |
| ENSMUSG00000079560 | -4.221875993 | -2.844526378 | -1.107868168 | -1.107868168 |
| ENSMUSG00000054196 | -6.108246117 | -5.150089863 | -1.646831061 | -1.646831061 |
| ENSMUSG00000026090 | -5.659110373 | -4.611126971 | -1.646831061 | -1.646831061 |
| ENSMUSG00000058254 | -4.461415056 | -2.4552754 | -1.2276377 | -1.2276377 |
| ENSMUSG00000023387 | -6.257958032 | -5.599225607 | -1.557003912 | -1.557003912 |
| ENSMUSG00000025789 | -5.748937522 | -4.521299822 | -1.886370124 | -1.886370124 |
| ENSMUSG00000028158 | 1.2276377 | 2.395390634 | 3.952394546 | 1.2276377 |
| ENSMUSG00000045377 | -5.928591819 | -5.000377949 | -2.036082039 | -2.036082039 |
| ENSMUSG00000026399 | -5.808822288 | -4.700954119 | -2.395390634 | -2.395390634 |
| ENSMUSG00000069324 | 1.527061529 | 2.994238293 | 4.880608417 | 1.527061529 |
| ENSMUSG00000028024 | -6.317842797 | -3.383489271 | -1.467176763 | -1.467176763 |
| ENSMUSG00000027983 | 2.036082039 | 3.89250978 | 4.940493183 | 2.036082039 |
| ENSMUSG00000030905 | 1.467176763 | 3.323604505 | 4.311703141 | 1.467176763 |
| ENSMUSG00000034957 | 3.203834973 | 4.9105508 | 5.928591819 | 3.203834973 |
| ENSMUSG00000074768 | 1.347407232 | 2.904411144 | 5.000377949 | 1.347407232 |
| ENSMUSG00000023913 | -6.527439478 | -2.425333017 | -1.2276377 | -1.2276377 |
| ENSMUSG00000079015 | 1.077925785 | 2.335505868 | 3.353546888 | 1.077925785 |
| ENSMUSG00000025742 | -3.862567397 | -2.395390634 | -1.137810551 | -1.137810551 |
| ENSMUSG00000041361 | -4.521299822 | -2.96429591 | -1.497119146 | -1.497119146 |
| ENSMUSG00000045382 | -4.850666034 | -3.263719739 | -1.856427741 | -1.856427741 |
| ENSMUSG00000002769 | 2.125909188 | 3.862567397 | 5.12014748 | 2.125909188 |
| ENSMUSG00000040706 | 2.305563485 | 3.862567397 | 4.880608417 | 2.305563485 |
| ENSMUSG00000021752 | -5.299801778 | -4.042221695 | -2.4552754 | -2.4552754 |

| | | | | |
|---|---|---|---|---|
| ENSMUSG00000003541 | -5.030320332 | -4.012279312 | -2.215736337 | -2.215736337 |
| ENSMUSG00000024043 | -4.161991227 | -2.575044932 | -1.2276377 | -1.2276377 |
| ENSMUSG00000030340 | -5.389628927 | -4.221875993 | -2.006139656 | -2.006139656 |
| ENSMUSG00000022686 | -6.198073266 | -2.575044932 | -1.257580083 | -1.257580083 |
| ENSMUSG00000001348 | -5.090205097 | -4.161991227 | -2.515160166 | -2.515160166 |
| ENSMUSG00000028845 | -5.509398458 | -4.611126971 | -2.365448251 | -2.365448251 |
| ENSMUSG00000024302 | -5.868707054 | -5.209974629 | -1.347407232 | -1.347407232 |
| ENSMUSG00000064373 | 1.107868168 | 2.275621102 | 3.323604505 | 1.107868168 |
| ENSMUSG00000016382 | -3.413431654 | -2.245678719 | -1.018041019 | -1.018041019 |
| ENSMUSG00000071177 | 1.018041019 | 2.305563485 | 3.293662122 | 1.018041019 |
| ENSMUSG00000032231 | -5.748937522 | -4.700954119 | -1.73665821 | -1.73665821 |
| ENSMUSG00000019478 | -6.078303734 | -5.41957131 | -1.497119146 | -1.497119146 |
| ENSMUSG00000020940 | -4.730896502 | -3.413431654 | -1.856427741 | -1.856427741 |
| ENSMUSG00000040717 | -5.239917012 | -4.581184588 | -2.125909188 | -2.125909188 |
| ENSMUSG00000029126 | -5.838764671 | -5.030320332 | -1.527061529 | -1.527061529 |
| ENSMUSG00000022425 | -5.569283224 | -4.611126971 | -1.976197273 | -1.976197273 |
| ENSMUSG00000031762 | -5.659110373 | -3.054123058 | -1.2276377 | -1.2276377 |
| ENSMUSG00000022941 | -4.760838885 | -3.623028334 | -1.616888678 | -1.616888678 |
| ENSMUSG00000078670 | -5.12014748 | -2.694814463 | -1.287522466 | -1.287522466 |
| ENSMUSG00000073293 | -4.671011736 | -3.054123058 | -1.137810551 | -1.137810551 |
| ENSMUSG00000044948 | -6.317842797 | -2.335505868 | -1.047983402 | -1.047983402 |
| ENSMUSG00000024391 | 1.047983402 | 2.365448251 | 4.072164078 | 1.047983402 |
| ENSMUSG00000037169 | -4.521299822 | -2.844526378 | -1.317464849 | -1.317464849 |
| ENSMUSG00000028681 | -5.62916799 | -4.700954119 | -1.497119146 | -1.497119146 |
| ENSMUSG00000034435 | -5.299801778 | -4.341645524 | -2.185793954 | -2.185793954 |
| ENSMUSG00000051910 | -5.299801778 | -4.491357439 | -2.305563485 | -2.305563485 |
| ENSMUSG00000061751 | -4.19193361 | -2.575044932 | -1.047983402 | -1.047983402 |
| ENSMUSG00000035934 | -5.389628927 | -4.431472673 | -2.125909188 | -2.125909188 |
| ENSMUSG00000026042 | -5.180032246 | -4.461415056 | -2.066024422 | -2.066024422 |
| ENSMUSG00000048612 | -5.449513693 | -4.760838885 | -2.006139656 | -2.006139656 |
| ENSMUSG00000064267 | -4.850666034 | -3.832625015 | -1.886370124 | -1.886370124 |
| ENSMUSG00000031274 | -5.958534202 | -2.575044932 | -1.137810551 | -1.137810551 |
| ENSMUSG00000019817 | -5.718995139 | -3.473316419 | -1.467176763 | -1.467176763 |
| ENSMUSG00000009772 | -5.689052756 | -4.760838885 | -2.275621102 | -2.275621102 |
| ENSMUSG00000022867 | -5.718995139 | -4.641069354 | -1.407291998 | -1.407291998 |
| ENSMUSG00000067771 | -4.850666034 | -3.862567397 | -2.066024422 | -2.066024422 |
| ENSMUSG00000075304 | -5.449513693 | -4.491357439 | -2.305563485 | -2.305563485 |
| ENSMUSG00000031838 | -3.952394546 | -2.515160166 | -1.077925785 | -1.077925785 |
| ENSMUSG00000007888 | -4.611126971 | -3.473316419 | -1.646831061 | -1.646831061 |
| ENSMUSG00000078954 | -5.778879905 | -4.970435566 | -1.796542976 | -1.796542976 |
| ENSMUSG00000037206 | -5.62916799 | -4.671011736 | -1.347407232 | -1.347407232 |
| ENSMUSG00000071745 | -4.611126971 | -3.623028334 | -1.886370124 | -1.886370124 |
| ENSMUSG00000014837 | -5.958534202 | -2.245678719 | -1.077925785 | -1.077925785 |
| ENSMUSG00000037624 | -5.449513693 | -4.641069354 | -1.557003912 | -1.557003912 |
| ENSMUSG00000090919 | -5.329744161 | -4.341645524 | -2.185793954 | -2.185793954 |
| ENSMUSG00000025813 | -5.180032246 | -4.19193361 | -2.066024422 | -2.066024422 |
| ENSMUSG00000001942 | -5.62916799 | -4.760838885 | -1.2276377 | -1.2276377 |
| ENSMUSG00000021062 | -6.317842797 | -2.515160166 | -1.197695317 | -1.197695317 |
| ENSMUSG00000039164 | 2.784641612 | 4.102106461 | 5.12014748 | 2.784641612 |
| ENSMUSG00000032714 | -5.689052756 | -4.940493183 | -1.73665821 | -1.73665821 |
| ENSMUSG00000023961 | 2.724756846 | 4.730896502 | 5.808822288 | 2.724756846 |
| ENSMUSG00000073988 | 1.047983402 | 3.143950207 | 4.40153029 | 1.047983402 |
| ENSMUSG00000015627 | -6.287900415 | -2.215736337 | -1.018041019 | -1.018041019 |
| ENSMUSG00000031075 | -5.689052756 | -4.730896502 | -1.586946295 | -1.586946295 |
| ENSMUSG00000025355 | 2.185793954 | 3.89250978 | 5.389628927 | 2.185793954 |
| ENSMUSG00000044034 | 1.407291998 | 3.143950207 | 4.371587907 | 1.407291998 |
| ENSMUSG00000046814 | 2.335505868 | 4.19193361 | 5.479456076 | 2.335505868 |
| ENSMUSG00000030882 | -5.030320332 | -3.742797866 | -1.916312507 | -1.916312507 |
| ENSMUSG00000031376 | 1.73665821 | 3.6829131 | 5.359686544 | 1.73665821 |

| | | | | |
|---|---|---|---|---|
| ENSMUSG00000036533 | -5.000377949 | -4.19193361 | -2.036082039 | -2.036082039 |
| ENSMUSG00000021469 | -6.108246117 | -5.479456076 | -1.167752934 | -1.167752934 |
| ENSMUSG00000039405 | -4.820723651 | -2.814583995 | -1.137810551 | -1.137810551 |
| ENSMUSG00000060212 | 2.425333017 | 3.832625015 | 5.569283224 | 2.425333017 |
| ENSMUSG00000027966 | -4.760838885 | -3.742797866 | -1.796542976 | -1.796542976 |
| ENSMUSG00000020467 | 1.676773444 | 3.473316419 | 4.581184588 | 1.676773444 |
| ENSMUSG00000035274 | -4.281760758 | -2.66487208 | -1.018041019 | -1.018041019 |
| ENSMUSG00000038521 | 2.095966805 | 3.862567397 | 5.060262715 | 2.095966805 |
| ENSMUSG00000103567 | -4.850666034 | -4.012279312 | -1.856427741 | -1.856427741 |
| ENSMUSG00000028807 | -4.880608417 | -3.802682632 | -1.886370124 | -1.886370124 |
| ENSMUSG00000027792 | 1.2276377 | 2.395390634 | 4.161991227 | 1.2276377 |
| ENSMUSG00000020427 | -5.898649436 | -4.790781268 | -1.856427741 | -1.856427741 |
| ENSMUSG00000034303 | -5.868707054 | -5.239917012 | -1.347407232 | -1.347407232 |
| ENSMUSG00000033985 | 1.916312507 | 3.353546888 | 4.880608417 | 1.916312507 |
| ENSMUSG00000059493 | -4.641069354 | -3.623028334 | -1.706715827 | -1.706715827 |
| ENSMUSG00000020000 | -5.060262715 | -4.102106461 | -1.676773444 | -1.676773444 |
| ENSMUSG00000041351 | -5.030320332 | -4.072164078 | -2.155851571 | -2.155851571 |
| ENSMUSG00000021271 | -4.491357439 | -3.623028334 | -1.676773444 | -1.676773444 |
| ENSMUSG00000036885 | -5.090205097 | -4.221875993 | -1.826485359 | -1.826485359 |
| ENSMUSG00000033542 | -5.12014748 | -2.844526378 | -1.107868168 | -1.107868168 |
| ENSMUSG00000020799 | -5.479456076 | -4.491357439 | -1.856427741 | -1.856427741 |
| ENSMUSG00000058656 | -4.641069354 | -3.712855483 | -1.886370124 | -1.886370124 |
| ENSMUSG00000031112 | -4.880608417 | -3.89250978 | -1.616888678 | -1.616888678 |
| ENSMUSG00000030587 | -4.700954119 | -3.024180676 | -1.197695317 | -1.197695317 |
| ENSMUSG00000047878 | -5.599225607 | -4.671011736 | -1.407291998 | -1.407291998 |
| ENSMUSG00000027330 | -5.898649436 | -5.150089863 | -1.077925785 | -1.077925785 |
| ENSMUSG00000060519 | -6.198073266 | -5.239917012 | -1.2276377 | -1.2276377 |
| ENSMUSG00000041245 | -5.778879905 | -4.671011736 | -1.2276377 | -1.2276377 |
| ENSMUSG00000031342 | -5.509398458 | -4.341645524 | -1.73665821 | -1.73665821 |
| ENSMUSG00000021280 | 2.365448251 | 3.742797866 | 5.329744161 | 2.365448251 |
| ENSMUSG00000027030 | -4.970435566 | -4.012279312 | -1.497119146 | -1.497119146 |
| ENSMUSG00000042066 | -5.359686544 | -4.581184588 | -1.94625489 | -1.94625489 |
| ENSMUSG00000062312 | -4.102106461 | -2.425333017 | -1.018041019 | -1.018041019 |
| ENSMUSG00000029130 | -5.778879905 | -5.090205097 | -1.197695317 | -1.197695317 |
| ENSMUSG00000026479 | -5.209974629 | -4.461415056 | -1.73665821 | -1.73665821 |
| ENSMUSG00000064023 | -6.048361351 | -5.329744161 | -1.137810551 | -1.137810551 |
| ENSMUSG00000030020 | -5.030320332 | -4.19193361 | -1.916312507 | -1.916312507 |
| ENSMUSG00000020102 | 2.365448251 | 4.671011736 | 5.868707054 | 2.365448251 |
| ENSMUSG00000019851 | -5.958534202 | -5.239917012 | -1.047983402 | -1.047983402 |
| ENSMUSG00000087408 | -4.820723651 | -3.89250978 | -1.826485359 | -1.826485359 |
| ENSMUSG00000029127 | -5.239917012 | -4.221875993 | -1.287522466 | -1.287522466 |
| ENSMUSG00000030782 | -4.790781268 | -3.89250978 | -1.616888678 | -1.616888678 |
| ENSMUSG00000068299 | -5.150089863 | -4.311703141 | -1.916312507 | -1.916312507 |
| ENSMUSG00000055202 | -5.62916799 | -4.521299822 | -1.2276377 | -1.2276377 |
| ENSMUSG00000075592 | -4.521299822 | -2.934353527 | -1.018041019 | -1.018041019 |
| ENSMUSG00000002032 | -5.329744161 | -4.251818376 | -1.347407232 | -1.347407232 |
| ENSMUSG00000039607 | -5.209974629 | -4.431472673 | -1.796542976 | -1.796542976 |
| ENSMUSG00000016179 | -5.778879905 | -5.000377949 | -1.137810551 | -1.137810551 |
| ENSMUSG00000016918 | -5.539340841 | -4.790781268 | -1.467176763 | -1.467176763 |
| ENSMUSG00000029646 | -5.090205097 | -4.341645524 | -1.377349615 | -1.377349615 |
| ENSMUSG00000042249 | -5.239917012 | -4.281760758 | -1.706715827 | -1.706715827 |
| ENSMUSG00000036002 | -4.9105508 | -3.952394546 | -1.347407232 | -1.347407232 |
| ENSMUSG00000000275 | 1.2276377 | 2.4552754 | 3.6829131 | 1.2276377 |
| ENSMUSG00000020658 | -4.700954119 | -3.383489271 | -1.347407232 | -1.347407232 |
| ENSMUSG00000032776 | -5.868707054 | -4.820723651 | -1.167752934 | -1.167752934 |
| ENSMUSG00000042743 | -5.180032246 | -4.40153029 | -1.886370124 | -1.886370124 |
| ENSMUSG00000103472 | -5.748937522 | -4.9105508 | -1.018041019 | -1.018041019 |
| ENSMUSG00000022297 | -5.659110373 | -4.730896502 | -1.018041019 | -1.018041019 |
| ENSMUSG00000096847 | -5.928591819 | -5.060262715 | -1.257580083 | -1.257580083 |

| | | | | |
|---|---|---|---|---|
| ENSMUSG00000000732 | 1.467176763 | 2.96429591 | 4.551242205 | 1.467176763 |
| ENSMUSG00000034993 | -5.000377949 | -2.844526378 | -1.018041019 | -1.018041019 |
| ENSMUSG00000043556 | -4.221875993 | -3.203834973 | -1.077925785 | -1.077925785 |
| ENSMUSG00000035948 | -4.850666034 | -3.89250978 | -1.646831061 | -1.646831061 |
| ENSMUSG00000020309 | -4.641069354 | -3.802682632 | -1.586946295 | -1.586946295 |
| ENSMUSG00000042846 | 1.766600593 | 3.533201185 | 5.209974629 | 1.766600593 |
| ENSMUSG00000031217 | -5.389628927 | -3.233777356 | -1.077925785 | -1.077925785 |
| ENSMUSG00000021539 | 1.796542976 | 4.40153029 | 5.62916799 | 1.796542976 |
| ENSMUSG00000028776 | -5.180032246 | -4.371587907 | -1.856427741 | -1.856427741 |
| ENSMUSG00000015401 | -5.479456076 | -4.371587907 | -1.467176763 | -1.467176763 |
| ENSMUSG00000014158 | 1.916312507 | 3.802682632 | 5.389628927 | 1.916312507 |
| ENSMUSG00000056972 | -4.850666034 | -3.862567397 | -1.497119146 | -1.497119146 |
| ENSMUSG00000069270 | 1.287522466 | 2.994238293 | 3.982336929 | 1.287522466 |
| ENSMUSG00000022474 | -5.479456076 | -4.790781268 | -1.018041019 | -1.018041019 |
| ENSMUSG00000032849 | -5.090205097 | -4.221875993 | -1.766600593 | -1.766600593 |
| ENSMUSG00000028399 | -5.090205097 | -4.19193361 | -1.616888678 | -1.616888678 |
| ENSMUSG00000045215 | -5.299801778 | -4.461415056 | -1.317464849 | -1.317464849 |
| ENSMUSG00000052142 | 2.245678719 | 3.89250978 | 5.239917012 | 2.245678719 |
| ENSMUSG00000028445 | 2.215736337 | 4.072164078 | 5.389628927 | 2.215736337 |
| ENSMUSG00000017697 | 1.706715827 | 3.143950207 | 4.521299822 | 1.706715827 |
| ENSMUSG00000027329 | -4.461415056 | -3.503258802 | -1.047983402 | -1.047983402 |
| ENSMUSG00000063446 | -6.108246117 | -5.000377949 | -1.167752934 | -1.167752934 |
| ENSMUSG00000061482 | 1.287522466 | 3.323604505 | 4.461415056 | 1.287522466 |
| ENSMUSG00000047281 | -4.611126971 | -3.802682632 | -1.527061529 | -1.527061529 |
| ENSMUSG00000019872 | -5.269859395 | -4.311703141 | -1.287522466 | -1.287522466 |
| ENSMUSG00000031778 | -5.269859395 | -4.371587907 | -1.407291998 | -1.407291998 |
| ENSMUSG00000016942 | 2.305563485 | 4.730896502 | 5.928591819 | 2.305563485 |
| ENSMUSG00000074063 | 1.706715827 | 4.611126971 | 6.048361351 | 1.706715827 |
| ENSMUSG00000032358 | -4.760838885 | -3.772740249 | -1.43723438 | -1.43723438 |
| ENSMUSG00000032120 | 1.796542976 | 3.263719739 | 4.611126971 | 1.796542976 |
| ENSMUSG00000046634 | -5.299801778 | -4.371587907 | -1.167752934 | -1.167752934 |
| ENSMUSG00000037166 | -4.611126971 | -3.413431654 | -1.43723438 | -1.43723438 |
| ENSMUSG00000052595 | 1.257580083 | 4.581184588 | 5.958534202 | 1.257580083 |
| ENSMUSG00000003949 | 1.826485359 | 3.982336929 | 5.509398458 | 1.826485359 |
| ENSMUSG00000034881 | 1.347407232 | 3.143950207 | 4.311703141 | 1.347407232 |
| ENSMUSG00000025931 | -4.880608417 | -4.102106461 | -1.377349615 | -1.377349615 |
| ENSMUSG00000051022 | -5.479456076 | -4.251818376 | -1.377349615 | -1.377349615 |
| ENSMUSG00000028919 | -5.150089863 | -4.132048844 | -1.646831061 | -1.646831061 |
| ENSMUSG00000018474 | -5.868707054 | -2.844526378 | -1.197695317 | -1.197695317 |
| ENSMUSG00000031970 | -4.9105508 | -3.89250978 | -1.167752934 | -1.167752934 |
| ENSMUSG00000048332 | -4.730896502 | -3.6829131 | -1.197695317 | -1.197695317 |
| ENSMUSG00000020926 | -5.389628927 | -4.341645524 | -1.047983402 | -1.047983402 |
| ENSMUSG00000094595 | -4.161991227 | -2.96429591 | -1.107868168 | -1.107868168 |
| ENSMUSG00000054065 | -5.180032246 | -4.251818376 | -1.287522466 | -1.287522466 |
| ENSMUSG00000022512 | 2.125909188 | 4.491357439 | 6.168130883 | 2.125909188 |
| ENSMUSG00000048440 | -5.569283224 | -4.790781268 | -1.137810551 | -1.137810551 |
| ENSMUSG00000071478 | 1.047983402 | 2.934353527 | 3.922452163 | 1.047983402 |
| ENSMUSG00000027007 | -4.9105508 | -3.982336929 | -1.2276377 | -1.2276377 |
| ENSMUSG00000043085 | 1.047983402 | 3.143950207 | 4.431472673 | 1.047983402 |
| ENSMUSG00000024558 | -4.730896502 | -3.772740249 | -1.467176763 | -1.467176763 |
| ENSMUSG00000019989 | -5.299801778 | -4.40153029 | -1.047983402 | -1.047983402 |
| ENSMUSG00000082229 | -5.329744161 | -4.371587907 | -1.018041019 | -1.018041019 |
| ENSMUSG00000038119 | -4.820723651 | -3.652970717 | -1.167752934 | -1.167752934 |
| ENSMUSG00000058297 | -4.940493183 | -3.982336929 | -1.287522466 | -1.287522466 |
| ENSMUSG00000006522 | 1.2276377 | 3.6829131 | 4.9105508 | 1.2276377 |
| ENSMUSG00000036834 | -5.12014748 | -4.341645524 | -1.347407232 | -1.347407232 |
| ENSMUSG00000020032 | -4.880608417 | -4.042221695 | -1.167752934 | -1.167752934 |
| ENSMUSG00000031443 | 1.706715827 | 3.233777356 | 4.521299822 | 1.706715827 |
| ENSMUSG00000032589 | -4.850666034 | -3.862567397 | -1.347407232 | -1.347407232 |

| | | | | |
|---|---|---|---|---|
| ENSMUSG00000069237 | -4.491357439 | -3.263719739 | -1.107868168 | -1.107868168 |
| ENSMUSG00000062309 | -4.9105508 | -3.982336929 | -1.557003912 | -1.557003912 |
| ENSMUSG00000025427 | -5.060262715 | -4.251818376 | -1.197695317 | -1.197695317 |
| ENSMUSG00000002190 | -4.760838885 | -3.922452163 | -1.077925785 | -1.077925785 |
| ENSMUSG00000079557 | -4.9105508 | -3.862567397 | -1.107868168 | -1.107868168 |
| ENSMUSG00000059741 | -5.988476585 | -4.251818376 | -1.137810551 | -1.137810551 |
| ENSMUSG00000045180 | -5.329744161 | -4.431472673 | -1.137810551 | -1.137810551 |
| ENSMUSG00000022598 | -5.150089863 | -4.581184588 | -1.257580083 | -1.257580083 |
| ENSMUSG00000029102 | 1.976197273 | 4.281760758 | 5.838764671 | 1.976197273 |
| ENSMUSG00000031465 | -4.730896502 | -3.89250978 | -1.047983402 | -1.047983402 |
| ENSMUSG00000032085 | -4.820723651 | -3.832625015 | -1.137810551 | -1.137810551 |
| ENSMUSG00000008734 | -4.820723651 | -3.982336929 | -1.047983402 | -1.047983402 |
| ENSMUSG00000040808 | -4.671011736 | -3.772740249 | -1.077925785 | -1.077925785 |
| ENSMUSG00000068015 | -5.060262715 | -4.251818376 | -1.107868168 | -1.107868168 |
| ENSMUSG00000028621 | -4.9105508 | -3.982336929 | -1.077925785 | -1.077925785 |
| ENSMUSG00000068762 | 1.167752934 | 4.40153029 | 5.479456076 | 1.167752934 |
| ENSMUSG00000024451 | -4.491357439 | -3.6829131 | -1.018041019 | -1.018041019 |
| ENSMUSG00000026208 | -5.928591819 | -4.521299822 | -1.018041019 | -1.018041019 |
| ENSMUSG00000031273 | -5.299801778 | -4.431472673 | -1.347407232 | -1.347407232 |
| ENSMUSG00000036459 | -4.700954119 | -3.652970717 | -1.018041019 | -1.018041019 |
| ENSMUSG00000034156 | -4.730896502 | -3.862567397 | -1.077925785 | -1.077925785 |
| ENSMUSG00000043760 | -4.880608417 | -3.89250978 | -1.107868168 | -1.107868168 |
| ENSMUSG00000075420 | -4.671011736 | -3.6829131 | -1.047983402 | -1.047983402 |
| ENSMUSG00000028337 | -4.641069354 | -3.623028334 | -1.287522466 | -1.287522466 |
| ENSMUSG00000006529 | 1.047983402 | 3.563143568 | 4.700954119 | 1.047983402 |
| ENSMUSG00000035493 | 1.43723438 | 3.89250978 | 4.820723651 | 1.43723438 |
| ENSMUSG00000003752 | -4.940493183 | -3.89250978 | -1.107868168 | -1.107868168 |
| ENSMUSG00000027276 | -4.970435566 | -4.221875993 | -1.107868168 | -1.107868168 |

#### mordium and gut tube

d of the 95
nin(abs(c(lb, ub))), or 0 if the CI crosses the 0

testing using Holm procedure

**Z and cZ.**

| Z | cZ | GeneName | Description | Location | Family |
|---|---|---|---|---|---|
| -7.160846644 | -6.190643472 | Slc2a3 | solute carrier | Plasma Mem | transporter |
| -7.160846644 | -6.190643472 | Tpm1 | tropomyosin | Plasma Mem | other |
| -7.160846644 | -6.190643472 | Fam183b | family with se | Other | other |
| 7.160812982 | 6.190643472 | Apoa2 | apolipoprote | Extracellular : | transporter |
| 7.160812982 | 6.190643472 | Itih2 | inter-alpha-t | Extracellular : | other |
| 7.160812982 | 6.190643472 | Fgb | fibrinogen be | Extracellular : | other |
| 7.160812982 | 6.190643472 | Reln | reelin | Extracellular : | peptidase |
| 7.160812982 | 6.190643472 | Apoe | apolipoprote | Extracellular : | transporter |
| 7.160812982 | 6.190643472 | Afp | alpha fetopro | Extracellular : | transporter |
| 7.160812982 | 6.190643472 | Dlk1 | delta like nor | Extracellular : | other |
| 7.160812982 | 6.190643472 | Serpinf2 | serpin family | Extracellular : | other |
| 7.160812982 | 6.190643472 | Hhex | hematopoiet | Nucleus | transcription regulator |
| 7.160808732 | 6.190643472 | Myc | MYC proto-c | Nucleus | transcription regulator |
| -7.16080869 | -6.190643472 | Capn6 | calpain 6 | Cytoplasm | peptidase |
| -7.16080869 | -6.190643472 | Sema3c | semaphorin : | Extracellular : | other |
| -7.16080869 | -6.190643472 | Nepn | nephrocan | Extracellular : | other |
| -7.16080869 | -6.190643472 | Gja1 | gap junction | Plasma Mem | transporter |
| -7.16080869 | -6.190643472 | Krt7 | keratin 7 | Cytoplasm | other |
| -7.16080869 | -6.190643472 | App | amyloid beta | Plasma Mem | other |
| -7.16080869 | -6.190643472 | Tmprss2 | transmembra | Plasma Mem | peptidase |
| 7.160786749 | 6.190643472 | Lpl | lipoprotein li | Cytoplasm | enzyme |
| 7.160753117 | 6.190643472 | Fabp5 | fatty acid bin | Cytoplasm | transporter |
| -7.160732813 | -6.190643472 | Shh | sonic hedgeh | Extracellular : | peptidase |
| 7.160729239 | 6.190643472 | Cald1 | caldesmon 1 | Plasma Mem | other |
| 7.160718103 | 6.190643472 | Prox1 | prospero hor | Nucleus | transcription regulator |
| -7.16069489 | -6.190643472 | Ackr3 | atypical chen | Plasma Mem | G-protein coupled recepto |
| -7.160581182 | -6.190643472 | Ifitm1 | interferon inc | Other | other |
| -7.160165045 | -6.190643472 | Epha4 | EPH receptor | Plasma Mem | kinase |
| -7.159712487 | -6.190643472 | Hoxa1 | homeobox A | Nucleus | transcription regulator |
| -7.159336472 | -6.190643472 | Mecom | MDS1 and E\ | Nucleus | transcription regulator |
| -7.158251718 | -6.190643472 | Peg3 | paternally ex | Nucleus | kinase |
| -7.158102756 | -6.190643472 | Osr1 | odd-skipped | Nucleus | kinase |
| -7.157805307 | -6.190643472 | Cd24a | CD24a antige | Plasma Mem | other |
| -7.147325682 | -6.185934903 | Rfx6 | regulatory fa | Nucleus | enzyme |
| -7.142249947 | -6.185934903 | Plat | plasminogen | Extracellular : | peptidase |
| -7.138479476 | -6.185934903 | Hoxc4 | homeobox C | Nucleus | transcription regulator |
| -7.137995075 | -6.185934903 | 1700011H14I | chromosome | Other | other |
| -7.128371703 | -6.179105258 | Cdh6 | cadherin 6 | Plasma Mem | other |
| -7.119366331 | -6.172881883 | Ociad2 | OCIA domair | Cytoplasm | other |
| -7.107049862 | -6.163097567 | Grhl2 | grainyhead li | Nucleus | transcription regulator |
| -7.103929695 | -6.163097567 | Cldn8 | claudin 8 | Plasma Mem | other |
| -7.073210095 | -6.131706991 | Cdh1 | cadherin 1 | Plasma Mem | other |
| -7.026334113 | -6.081673923 | Arg1 | arginase 1 | Cytoplasm | enzyme |
| -6.995296055 | -6.049713795 | Mmp2 | matrix metall | Extracellular : | peptidase |
| -6.948872718 | -5.999961696 | Nkx2-6 | NK2 homeob | Nucleus | transcription regulator |
| -6.874458476 | -5.917798411 | Gprc5a | G protein-co | Plasma Mem | G-protein coupled recepto |
| -6.813590023 | -5.851035799 | Osr2 | odd-skipped | Nucleus | transcription regulator |
| -6.775993188 | -5.811039432 | Raet1e | retinoic acid | Plasma Mem | other |

| | | | | | |
|---|---|---|---|---|---|
| -6.756525338 | -5.794268572 | Rprm | reprimo, TP5 | Cytoplasm | other |
| -6.755599126 | -5.794268572 | Lamb3 | laminin subu | Extracellular ! | transporter |
| -6.727716256 | -5.768756115 | Klf5 | Kruppel like f | Nucleus | transcription regulator |
| -6.72509542 | -5.768756115 | Dll1 | delta like can | Plasma Mem | enzyme |
| -6.719563489 | -5.765504719 | Igfbp2 | insulin like gr | Extracellular ! | other |
| -6.715812222 | -5.764258897 | Ly6h | lymphocyte a | Plasma Mem | other |
| -6.684740024 | -5.731341775 | Col4a1 | collagen type | Extracellular ! | other |
| -6.675492719 | -5.723637131 | Nr2f2 | nuclear recep | Nucleus | ligand-dependent nuclear |
| -6.647284279 | -5.693919483 | Col4a2 | collagen type | Extracellular ! | other |
| -6.598954786 | -5.643678521 | Smoc2 | SPARC relate | Extracellular ! | other |
| -6.583164052 | -5.628197759 | Fam162b | family with se | Other | other |
| -6.548954687 | -5.591523142 | Wls | wntless Wnt l | Cytoplasm | other |
| -6.546779176 | -5.591523142 | Gpm6a | glycoprotein | Plasma Mem | ion channel |
| -6.526768488 | -5.573391799 | Lypd6b | LY6/PLAUR d | Other | other |
| -6.526498897 | -5.573391799 | Igsf9 | immunoglob | Plasma Mem | other |
| -6.507508269 | -5.553981621 | Corin | corin, serine | Plasma Mem | peptidase |
| -6.44919497 | -5.491359312 | Tub | tubby biparti | Cytoplasm | transcription regulator |
| -6.436648271 | -5.479321153 | Prdm1 | PR/SET doma | Nucleus | transcription regulator |
| 6.432867341 | 5.477462392 | Ambp | alpha-1-mic | Extracellular ! | transporter |
| -6.384288636 | -5.425830532 | Atp13a4 | ATPase 13A4 | Cytoplasm | transporter |
| -6.380121861 | -5.423432593 | Bcam | basal cell adh | Plasma Mem | transmembrane receptor |
| -6.355077979 | -5.396637556 | Loxl1 | lysyl oxidase | Extracellular ! | enzyme |
| -6.337154091 | -5.378114839 | Pcsk5 | proprotein co | Extracellular ! | peptidase |
| 6.323679131 | 5.364756773 | Tbx3 | T-box 3 | Nucleus | transcription regulator |
| -6.31033053 | -5.351515893 | Vsig2 | V-set and im | Plasma Mem | other |
| -6.30016588 | -5.341968289 | Ncmap | non-compac | Plasma Mem | other |
| -6.295217171 | -5.340377635 | Nrp2 | neuropilin 2 | Plasma Mem | kinase |
| -6.29487936 | -5.340377635 | Scara3 | scavenger re | Plasma Mem | transmembrane receptor |
| -6.289731894 | -5.336615193 | Mical1 | microtubule a | Cytoplasm | enzyme |
| -6.277523055 | -5.324575501 | Cldn4 | claudin 4 | Plasma Mem | transmembrane receptor |
| -6.274016296 | -5.322681039 | Nkx6-2 | NK6 homeob | Nucleus | transcription regulator |
| 6.235270153 | 5.279550523 | Ttr | transthyretin | Extracellular ! | transporter |
| -6.227648928 | -5.274105327 | Gpx2 | glutathione p | Cytoplasm | enzyme |
| -6.226932049 | -5.274105327 | Tdh | L-threonine c | Cytoplasm | enzyme |
| -6.18945642 | -5.234452882 | Pdx1 | pancreatic ar | Nucleus | transcription regulator |
| -6.187607864 | -5.234374354 | Pitx2 | paired like hc | Nucleus | transcription regulator |
| -6.178762908 | -5.226077693 | Nsg2 | HMP19 prote | Cytoplasm | other |
| -6.173069801 | -5.221452826 | Fam43a | family with se | Other | other |
| -6.153882435 | -5.200993162 | Gm17455 | chromosome | Other | other |
| -6.144901676 | -5.192474097 | Duxbl1 | double home | Nucleus | other |
| -6.118366562 | -5.165317945 | Nkx3-1 | NK3 homeob | Nucleus | transcription regulator |
| -6.108590951 | -5.155805547 | Sema6a | semaphorin 6 | Plasma Mem | transmembrane receptor |
| -6.078491722 | -5.122400872 | Flt4 | fms related ty | Plasma Mem | transmembrane receptor |
| 6.059777469 | 5.102334268 | Serpinf1 | serpin family | Extracellular ! | other |
| -6.035095486 | -5.077123732 | Apba1 | amyloid beta | Cytoplasm | transporter |
| -6.027636593 | -5.07023616 | Ldoc1 | LDOC1, regu | Nucleus | other |
| -6.024335346 | -5.068225669 | Fat3 | FAT atypical c | Plasma Mem | other |
| -6.014549843 | -5.059400979 | Isx | intestine spec | Nucleus | transcription regulator |
| -6.013693961 | -5.059400979 | Samd5 | sterile alpha r | Other | other |
| -6.010050182 | -5.056937992 | Fam134b | reticulophagy | Cytoplasm | other |
| -6.006790984 | -5.054911559 | Slc37a1 | solute carrier | Plasma Mem | transporter |
| -6.001093385 | -5.050000452 | Cobl | cordon-bleu | Plasma Mem | other |
| -5.998895189 | -5.049191366 | Duxbl3 | double home | Nucleus | other |
| -5.976784297 | -5.024932135 | Hsd11b2 | hydroxystero | Cytoplasm | enzyme |
| -5.967448937 | -5.015692083 | Rdh10 | retinol dehyc | Nucleus | enzyme |
| -5.957132903 | -5.005281208 | Kynu | kynureninase | Cytoplasm | enzyme |
| -5.955188929 | -5.004718715 | Hoxc5 | homeobox C | Nucleus | transcription regulator |
| -5.90487301 | -4.947090597 | Mylip | myosin regul | Cytoplasm | enzyme |
| -5.903012662 | -4.946609217 | Dleu7 | deleted in lyr | Other | other |

| | | | | | |
|---|---|---|---|---|---|
| -5.901527297 | -4.946556191 | Entpd3 | ectonucleosi( | Plasma Mem | enzyme |
| -5.886462886 | -4.930453753 | Tril | TLR4 interact | Other | other |
| -5.856156976 | -4.896305561 | Adam8 | ADAM metal | Plasma Mem | peptidase |
| -5.847045338 | -4.887197838 | Epha7 | EPH receptor | Plasma Mem | kinase |
| -5.844922837 | -4.886348203 | Rfx2 | regulatory fa | Nucleus | transcription regulator |
| -5.833297884 | -4.874239753 | 5730559C18I | chromosome | Other | other |
| -5.800498227 | -4.837021463 | Pdzk1ip1 | PDZK1 intera | Extracellular S | other |
| -5.779392402 | -4.813619201 | Frem2 | FRAS1 relate | Extracellular S | other |
| -5.777283968 | -4.812747113 | Zfp503 | zinc finger pr | Nucleus | other |
| -5.768079426 | -4.80343945 | Duxbl2 | double home | Nucleus | other |
| 5.754566121 | 4.789000653 | Asb4 | ankyrin repea | Nucleus | transcription regulator |
| 5.735248953 | 4.770177047 | Pcsk9 | proprotein co | Extracellular S | peptidase |
| -5.734526104 | -4.770177047 | Fam129b | family with se | Cytoplasm | transcription regulator |
| -5.733391195 | -4.770177047 | Plxdc2 | plexin domai | Extracellular S | other |
| 5.71859971 | 4.754154627 | Phgdh | phosphoglyc | Cytoplasm | enzyme |
| -5.714799819 | -4.751185694 | Hoxb2 | homeobox B | Nucleus | transcription regulator |
| -5.712324045 | -4.750141474 | Lpar3 | lysophosphat | Plasma Mem | G-protein coupled recepto |
| -5.711343944 | -4.750141474 | B4galnt2 | beta-1,4-N-a | Cytoplasm | enzyme |
| -5.700826651 | -4.739156132 | Rtn1 | reticulon 1 | Cytoplasm | other |
| -5.684537015 | -4.721290775 | Kcnh8 | potassium vo | Plasma Mem | ion channel |
| -5.681591884 | -4.719287781 | Ephx4 | epoxide hydr | Other | enzyme |
| -5.660819383 | -4.69605407 | Fstl1 | follistatin like | Extracellular S | other |
| -5.640076395 | -4.672824041 | Efs | embryonal Fy | Cytoplasm | other |
| -5.619421579 | -4.649667574 | Gpr37 | G protein-co | Plasma Mem | G-protein coupled recepto |
| -5.606745915 | -4.636012486 | Adgb | androglobin | Other | other |
| -5.603023279 | -4.633038629 | Trabd2b | TraB domain | Plasma Mem | peptidase |
| -5.601138187 | -4.632249894 | Spock3 | SPARC/ostec | Extracellular S | other |
| -5.574618495 | -4.603463467 | Bmpr1b | bone morpho | Plasma Mem | kinase |
| -5.559113469 | -4.586359676 | Slc35f1 | solute carrier | Other | other |
| 5.539626703 | 4.564466558 | Cdkn1c | cyclin-depen | Nucleus | other |
| -5.530966345 | -4.555523342 | Lamb1 | laminin subu | Extracellular S | other |
| -5.514824757 | -4.53759881 | Cldn9 | claudin 9 | Plasma Mem | other |
| -5.50359654 | -4.525550184 | Kif26b | kinesin family | Other | other |
| 5.498153137 | 4.520432696 | Raph1 | Ras associati | Plasma Mem | other |
| -5.47166541 | -4.491433052 | Tead1 | TEA domain | Nucleus | transcription regulator |
| -5.46293947 | -4.482341785 | Nrep | neuronal reg | Cytoplasm | other |
| -5.459920775 | -4.480102818 | Epas1 | endothelial P | Nucleus | transcription regulator |
| 5.440674344 | 4.46108436 | Ccnd1 | cyclin D1 | Nucleus | transcription regulator |
| -5.423184086 | -4.441398104 | Tspan8 | tetraspanin 8 | Plasma Mem | other |
| 5.397071917 | 4.411290981 | Tgm2 | transglutamii | Cytoplasm | enzyme |
| -5.391953288 | -4.406474746 | 2610528A11I | RIKEN cDNA | Extracellular S | other |
| -5.352281982 | -4.36394815 | Syngr3 | synaptogyrin | Plasma Mem | other |
| -5.345233576 | -4.357351007 | Ccdc160 | coiled-coil do | Other | other |
| -5.34462951 | -4.357351007 | Afap1l1 | actin filamen | Other | other |
| -5.33077332 | -4.341913653 | Podxl | podocalyxin- | Plasma Mem | other |
| -5.322817583 | -4.33360079 | Krt19 | keratin 19 | Cytoplasm | other |
| -5.314439173 | -4.324767397 | Uchl1 | ubiquitin C-t | Cytoplasm | peptidase |
| -5.303000591 | -4.312218422 | Zfp558 | zinc finger pr | Nucleus | other |
| -5.300092829 | -4.309993019 | Bdh2 | 3-hydroxybu | Cytoplasm | enzyme |
| -5.286203093 | -4.294911635 | Efna3 | ephrin A3 | Plasma Mem | kinase |
| -5.285519957 | -4.294911635 | Bmp5 | bone morpho | Extracellular S | growth factor |
| -5.278550716 | -4.287743329 | Csrnp1 | cysteine and | Nucleus | transcription regulator |
| -5.27570835 | -4.28557105 | Irs4 | insulin recept | Other | other |
| 5.27371533 | 4.284421833 | Gstt1 | glutathione S | Cytoplasm | enzyme |
| -5.26713681 | -4.277965577 | Car2 | carbonic anh | Cytoplasm | enzyme |
| -5.266315236 | -4.277965577 | Dpysl3 | dihydropyrim | Cytoplasm | enzyme |
| 5.255629051 | 4.266256437 | Pfkfb3 | 6-phosphofr | Cytoplasm | kinase |
| -5.251990702 | -4.263087147 | Ahnak | AHNAK nucle | Nucleus | other |
| -5.229351224 | -4.23684491 | Epb4.1l3 | erythrocyte n | Plasma Mem | other |

| | | | | | |
|---|---|---|---|---|---|
| -5.221759305 | -4.229901078 | Mro | maestro | Nucleus | other |
| -5.221602822 | -4.229901078 | Ihh | indian hedge | Extracellular S | enzyme |
| -5.186700556 | -4.188669339 | Sorl1 | sortilin relate | Cytoplasm | transporter |
| 5.184987851 | 4.187813319 | Fgg | fibrinogen ga | Extracellular S | other |
| -5.165087297 | -4.164782513 | Agrn | agrin | Plasma Mem | other |
| 5.157049323 | 4.156200207 | Lzts1 | leucine zippe | Nucleus | transcription regulator |
| -5.153556289 | -4.153157068 | Raet1d | retinoic acid | Plasma Mem | other |
| -5.142634053 | -4.141038713 | BC030500 | cDNA sequer | Other | other |
| -5.136904161 | -4.135251797 | H60b | histocompati | Plasma Mem | other |
| -5.116748412 | -4.111823077 | Tfcp2l1 | transcription | Nucleus | transcription regulator |
| -5.115395928 | -4.11137418 | Nckap5 | NCK associat | Other | other |
| -5.104826075 | -4.099643051 | Cachd1 | cache domai | Other | other |
| -5.087589968 | -4.079733875 | 1700027J19R | CDC42 effect | Cytoplasm | other |
| 5.072962104 | 4.062997104 | Serpina1b | serpin family | Extracellular S | other |
| -5.031930323 | -4.013778924 | Eya1 | EYA transcrip | Nucleus | phosphatase |
| -5.024930128 | -4.006895275 | Epcam | epithelial cell | Plasma Mem | other |
| -5.003030424 | -3.981770298 | Ptges | prostaglandi | Cytoplasm | enzyme |
| -4.99678777 | -3.975262901 | Adamts7 | ADAM metal | Extracellular S | peptidase |
| -4.993751236 | -3.972704452 | Lpar4 | lysophosphat | Plasma Mem | transmembrane receptor |
| -4.987786734 | -3.966528366 | Tmem44 | transmembra | Other | other |
| 4.968228718 | 3.943557102 | Apoa5 | apolipoprote | Extracellular S | transporter |
| -4.964901211 | -3.941795762 | Wnt7b | Wnt family m | Extracellular S | other |
| -4.963334538 | -3.941027137 | Hmgcll1 | 3-hydroxyme | Cytoplasm | enzyme |
| -4.951915499 | -3.92807709 | Rhbdf2 | rhomboid 5 h | Cytoplasm | other |
| -4.946888085 | -3.923507765 | Prickle1 | prickle plana | Nucleus | other |
| -4.946348269 | -3.923507765 | B3gnt7 | UDP-GlcNAc | Cytoplasm | enzyme |
| -4.923982018 | -3.898131001 | Klhl34 | kelch like fam | Extracellular S | other |
| -4.915624969 | -3.888924935 | Ppargc1a | PPARG coact | Nucleus | transcription regulator |
| 4.913430336 | 3.887348073 | Gstm1 | glutathione S | Cytoplasm | enzyme |
| -4.908436068 | -3.882297234 | F2r | coagulation f | Plasma Mem | G-protein coupled recepto |
| -4.905811479 | -3.880178152 | Osbpl6 | oxysterol bin | Cytoplasm | transporter |
| -4.898286354 | -3.8727518 | Zfp185 | zinc finger pr | Nucleus | other |
| -4.898001349 | -3.8727518 | Dmtn | dematin actir | Plasma Mem | other |
| -4.888484842 | -3.862072703 | Ddo | D-aspartate c | Cytoplasm | enzyme |
| -4.883412661 | -3.856898734 | Cldn3 | claudin 3 | Plasma Mem | transmembrane receptor |
| 4.855871334 | 3.823803767 | Aldh4a1 | aldehyde deh | Cytoplasm | enzyme |
| -4.852706192 | -3.82114416 | Pcdhga4 | protocadheri | Other | other |
| -4.851939173 | -3.82114416 | Col2a1 | collagen type | Extracellular S | other |
| -4.842655884 | -3.810704939 | Esam | endothelial c | Plasma Mem | other |
| -4.799935688 | -3.758545802 | Fam132b | erythroferror | Extracellular S | other |
| -4.783273666 | -3.738827667 | Vnn1 | vanin 1 | Plasma Mem | enzyme |
| 4.776037421 | 3.730886086 | Paqr9 | progestin and | Plasma Mem | other |
| 4.754922281 | 3.705539347 | Serpina1e | serpin family | Extracellular S | other |
| -4.744456579 | -3.693520484 | Rnf207 | ring finger pr | Other | other |
| -4.740598103 | -3.68978853 | Ezr | ezrin | Plasma Mem | other |
| 4.72769283 | 3.67468457 | Gm595 | predicted ge | Other | other |
| -4.709121616 | -3.652428455 | Gpr45 | G protein-co | Plasma Mem | G-protein coupled recepto |
| -4.703359981 | -3.646281063 | Chst5 | carbohydrate | Cytoplasm | enzyme |
| 4.686258027 | 3.625827346 | Psph | phosphoserir | Cytoplasm | phosphatase |
| -4.67853518 | -3.617187293 | St14 | suppression c | Plasma Mem | peptidase |
| -4.661041096 | -3.597253449 | Palm3 | paralemmin 3 | Other | other |
| -4.661009721 | -3.597253449 | Isl1 | ISL LIM home | Nucleus | transcription regulator |
| -4.659355737 | -3.596262629 | Cmklr1 | chemerin che | Plasma Mem | G-protein coupled recepto |
| -4.648156885 | -3.58319487 | Cgnl1 | cingulin like 1 | Plasma Mem | other |
| -4.606012655 | -3.53194462 | Adamts17 | ADAM metal | Extracellular S | other |
| -4.593790306 | -3.517502655 | Ccdc80 | coiled-coil d | Nucleus | other |
| -4.588893099 | -3.51236985 | Ltbp1 | latent transfo | Extracellular S | other |
| -4.582293727 | -3.506157742 | Sfrp5 | secreted frizz | Plasma Mem | transmembrane receptor |
| -4.567134367 | -3.49023894 | Plcd1 | phospholipas | Cytoplasm | enzyme |

| | | | | | |
|---|---|---|---|---|---|
| -4.567087577 | -3.49023894 | Tmem140 | transmembra | Other | other |
| -4.566905708 | -3.49023894 | Ctnnd2 | catenin delta | Plasma Mem | other |
| -4.566390442 | -3.49023894 | Myo5b | myosin VB | Cytoplasm | enzyme |
| -4.55444 | -3.47608056 | Fxyd6 | FXYD domair | Plasma Mem | ion channel |
| 4.550294643 | 3.471867412 | Afm | afamin | Extracellular S | transporter |
| -4.529838349 | -3.446921896 | Akap12 | A-kinase anc | Cytoplasm | transporter |
| -4.529083416 | -3.446921896 | Megf6 | multiple EGF | Cytoplasm | other |
| -4.508810253 | -3.422120443 | Sept8 | septin 8 | Extracellular S | other |
| -4.508030567 | -3.422120443 | Fxyd3 | FXYD domair | Plasma Mem | other |
| -4.495013399 | -3.40650294 | AI661453 | expressed se | Other | other |
| 4.480220911 | 3.388582129 | Cited1 | Cbp/p300 int | Nucleus | transcription regulator |
| -4.462584159 | -3.368041573 | Pnma2 | paraneoplast | Nucleus | transporter |
| 4.441498203 | 3.34300935 | Alb | albumin | Extracellular S | transporter |
| -4.437221314 | -3.338557955 | Dab2 | DAB2, clathri | Plasma Mem | other |
| 4.436109417 | 3.338185947 | Gap43 | growth assoc | Plasma Mem | other |
| -4.425795609 | -3.326991635 | Gmpr | guanosine m | Cytoplasm | enzyme |
| -4.415178146 | -3.314331684 | Fhl2 | four and a ha | Nucleus | transcription regulator |
| -4.412484813 | -3.311903855 | Tdrd7 | tudor domair | Cytoplasm | other |
| -4.409905222 | -3.309619501 | Ppp2r2b | protein phos | Cytoplasm | phosphatase |
| -4.404897272 | -3.304191873 | Mrc2 | mannose rec | Plasma Mem | transmembrane receptor |
| 4.40377469 | 3.303784407 | Serpina1a | serpin family | Extracellular S | other |
| -4.390390996 | -3.288549633 | Syt5 | synaptotagm | Cytoplasm | transporter |
| -4.34485623 | -3.230472814 | Fras1 | Fraser extrac | Extracellular S | other |
| 4.32275254 | 3.204803066 | F10 | coagulation f | Extracellular S | peptidase |
| 4.314712101 | 3.196394395 | Lrrc3 | leucine rich r | Other | other |
| -4.287242438 | -3.161502058 | Ntn1 | netrin 1 | Extracellular S | other |
| -4.26996504 | -3.140929799 | Epn3 | epsin 3 | Cytoplasm | other |
| -4.232393641 | -3.09394543 | Lad1 | ladinin 1 | Extracellular S | other |
| -4.23188297 | -3.09394543 | Zfp941 | zinc finger pr | Other | other |
| -4.218479908 | -3.077295028 | Tgfb2 | transforming | Extracellular S | growth factor |
| 4.215204578 | 3.075058561 | Taok3 | TAO kinase 3 | Cytoplasm | kinase |
| -4.206633849 | -3.064768485 | Thbs1 | thrombospor | Extracellular S | other |
| -4.192619131 | -3.047254017 | 4930506M07 | shootin 1 | Plasma Mem | other |
| -4.185544917 | -3.038917406 | Esrp1 | epithelial spli | Nucleus | other |
| -4.170981931 | -3.020625025 | Efna1 | ephrin A1 | Plasma Mem | other |
| -4.16926731 | -3.020022485 | Zfp462 | zinc finger pr | Nucleus | other |
| -4.168962179 | -3.020022485 | Serinc5 | serine incorp | Plasma Mem | transporter |
| -4.165126848 | -3.016994787 | Fuca2 | fucosidase, a | Extracellular S | enzyme |
| -4.163789633 | -3.016247512 | Akap5 | A-kinase anc | Plasma Mem | other |
| -4.161453383 | -3.014169802 | Usp13 | ubiquitin spe | Cytoplasm | peptidase |
| -4.160496658 | -3.013921909 | Kbtbd11 | kelch repeat | Other | other |
| -4.15050853 | -3.001665637 | Lxn | latexin | Cytoplasm | other |
| -4.138596193 | -2.987849747 | Lnx1 | ligand of nur | Cytoplasm | enzyme |
| -4.124936928 | -2.971241922 | Zbtb16 | zinc finger ar | Nucleus | transcription regulator |
| -4.120418617 | -2.96727546 | Trim2 | tripartite mot | Cytoplasm | enzyme |
| -4.120117829 | -2.96727546 | Pdgfc | platelet deriv | Extracellular S | growth factor |
| 4.111477273 | 2.956750726 | Serpinc1 | serpin family | Extracellular S | enzyme |
| -4.106772993 | -2.952482733 | Sema3b | semaphorin 3 | Extracellular S | other |
| -4.095445202 | -2.940348179 | Spp1 | secreted pho | Extracellular S | cytokine |
| -4.090828234 | -2.935170643 | 6430573F11F | KIAA1456 | Cytoplasm | enzyme |
| -4.087623799 | -2.931879137 | Lama1 | laminin subu | Extracellular S | other |
| -4.085090114 | -2.929482206 | Itga3 | integrin subu | Plasma Mem | other |
| 4.081509195 | 2.927655193 | Apoa1 | apolipoprote | Extracellular S | transporter |
| -4.074048197 | -2.918650401 | Syn3 | synapsin III | Plasma Mem | other |
| 4.061107134 | 2.904248656 | Oaf | out at first hc | Cytoplasm | other |
| -4.056771912 | -2.90065659 | Crip2 | cysteine rich | Plasma Mem | other |
| -4.056249272 | -2.90065659 | Igdcc3 | immunoglob | Other | other |
| 4.053850581 | 2.898408645 | Gamt | guanidinoace | Cytoplasm | enzyme |
| -4.049641172 | -2.89372872 | Lrch2 | leucine rich r | Other | other |

| | | | | | |
|---|---|---|---|---|---|
| -4.04571831 | -2.890392245 | Mansc1 | MANSC dom | Other | other |
| -4.040471387 | -2.884309809 | Efnb2 | ephrin B2 | Plasma Mem | kinase |
| -4.035990198 | -2.879647558 | Folr1 | folate recept | Plasma Mem | transporter |
| -4.032530659 | -2.876515156 | Trib2 | tribbles pseu | Plasma Mem | kinase |
| 4.030192169 | 2.874325811 | Herpud1 | homocystein | Cytoplasm | other |
| -4.024838203 | -2.868080915 | Hs6st2 | heparan sulfa | Plasma Mem | enzyme |
| -4.019656944 | -2.862062719 | Flnc | filamin C | Cytoplasm | other |
| 4.013695139 | 2.855740313 | Serpind1 | serpin family | Extracellular S | other |
| -4.01355049 | -2.855740313 | Pcdh7 | protocadheri | Plasma Mem | other |
| -4.003767182 | -2.844453769 | Nudt17 | nudix hydrola | Cytoplasm | other |
| -3.986972724 | -2.826499491 | Calcoco1 | calcium bind | Nucleus | transcription regulator |
| -3.971760895 | -2.807818828 | Ncam1 | neural cell ac | Plasma Mem | other |
| -3.962253885 | -2.795888295 | Tjp3 | tight junction | Cytoplasm | other |
| -3.957280287 | -2.790085162 | Sycn | syncollin | Extracellular S | other |
| -3.951469035 | -2.7831412 | Ramp3 | receptor acti | Plasma Mem | transporter |
| -3.946996483 | -2.77815322 | Gli3 | GLI family zin | Nucleus | transcription regulator |
| -3.946419013 | -2.77815322 | Ccdc173 | coiled-coil d | Other | other |
| -3.944417781 | -2.776365939 | Zbtb6 | zinc finger ar | Nucleus | other |
| -3.942590091 | -2.774811721 | Adcy8 | adenylate cyc | Plasma Mem | enzyme |
| 3.941677486 | 2.77449651 | Heg1 | heart develop | Plasma Mem | other |
| -3.935390383 | -2.767805804 | Igsf3 | immunoglob | Plasma Mem | other |
| -3.934093685 | -2.766962927 | Irs1 | insulin recept | Cytoplasm | enzyme |
| -3.932821031 | -2.766150412 | Slc4a9 | solute carrier | Plasma Mem | transporter |
| -3.930662531 | -2.764133465 | Zfp114 | zinc finger pr | Other | other |
| 3.919144296 | 2.750310513 | C1s2 | complement | Extracellular S | peptidase |
| -3.916753229 | -2.749742333 | Rcan1 | regulator of c | Nucleus | transcription regulator |
| -3.916725264 | -2.749742333 | Nhsl2 | NHS like 2 | Other | other |
| -3.903092177 | -2.733958583 | Lmo7 | LIM domain 7 | Cytoplasm | enzyme |
| -3.887857703 | -2.714948839 | 3632451O06 | chromosome | Extracellular S | other |
| -3.850621232 | -2.666709001 | Lrrk1 | leucine rich r | Cytoplasm | kinase |
| 3.843915437 | 2.659311217 | Apof | apolipoprote | Extracellular S | transporter |
| 3.841156849 | 2.656424255 | Zc3hav1 | zinc finger CC | Plasma Mem | other |
| -3.835310496 | -2.650190211 | Il13ra1 | interleukin 13 | Plasma Mem | transmembrane receptor |
| -3.830899016 | -2.645022505 | St6galnac2 | ST6 N-acetyl | Cytoplasm | enzyme |
| -3.824724458 | -2.639213018 | Hps1 | HPS1, bioger | Cytoplasm | other |
| -3.819351073 | -2.632708887 | Morc4 | MORC family | Nucleus | other |
| -3.816359217 | -2.629703749 | Hoxa3 | homeobox A | Nucleus | transcription regulator |
| -3.815876615 | -2.629703749 | Cthrc1 | collagen tripl | Extracellular S | other |
| -3.795842924 | -2.604998615 | 2010300C02I | KIAA1211 like | Other | other |
| -3.794159002 | -2.604998615 | Tspan7 | tetraspanin 7 | Plasma Mem | other |
| -3.793486277 | -2.604998615 | Kcnk16 | potassium tw | Plasma Mem | ion channel |
| -3.788344368 | -2.598791323 | St8sia2 | ST8 alpha-N- | Cytoplasm | enzyme |
| 3.779956716 | 2.588074065 | Mttp | microsomal t | Cytoplasm | transporter |
| -3.777185137 | -2.585993317 | Tmem88 | transmembra | Plasma Mem | other |
| -3.774006116 | -2.583341142 | Cd55 | CD55 molecu | Plasma Mem | other |
| 3.766187086 | 2.573387599 | Gm5096 | betaine--hor | Cytoplasm | enzyme |
| -3.746395507 | -2.548541587 | Enpep | glutamyl ami | Plasma Mem | peptidase |
| 3.740744404 | 2.543295059 | Cyp2u1 | cytochrome P | Cytoplasm | enzyme |
| 3.738256441 | 2.540702105 | Crym | crystallin mu | Cytoplasm | enzyme |
| 3.737313301 | 2.540255201 | Cebpa | CCAAT/enha | Nucleus | transcription regulator |
| 3.7327174 | 2.535300155 | Bhmt | betaine--hor | Cytoplasm | enzyme |
| -3.732510635 | -2.535300155 | Pla2g7 | phospholipas | Extracellular S | enzyme |
| 3.728543972 | 2.530704228 | Serpina1c | serpin family | Extracellular S | other |
| -3.727972253 | -2.530704228 | Prps2 | phosphoribo | Cytoplasm | kinase |
| -3.719282735 | -2.521008812 | Myzap | myocardial zo | Plasma Mem | other |
| -3.715426699 | -2.516654946 | Cxcr4 | C-X-C motif | Plasma Mem | G-protein coupled recepto |
| 3.71478863 | 2.516617968 | Gnmt | glycine N-me | Cytoplasm | enzyme |
| 3.707492424 | 2.508994784 | Agmat | agmatinase | Cytoplasm | enzyme |
| -3.705624915 | -2.507236838 | Kctd6 | potassium ch | Other | ion channel |

| | | | | | |
|---|---|---|---|---|---|
| -3.690541131 | -2.487017857 | Ier3 | immediate ea | Cytoplasm | other |
| -3.686262117 | -2.481879869 | Arhgap28 | Rho GTPase a | Cytoplasm | other |
| -3.674099992 | -2.467117569 | Scnn1a | sodium chan | Plasma Mem | ion channel |
| -3.66686528 | -2.458082687 | B3gnt5 | UDP-GlcNAc | Cytoplasm | enzyme |
| -3.641454231 | -2.426567929 | Acp5 | acid phospha | Cytoplasm | phosphatase |
| -3.626103395 | -2.409047105 | Tekt2 | tektin 2 | Cytoplasm | other |
| -3.625454919 | -2.409047105 | Dtna | dystrobrevin | Plasma Mem | other |
| 3.622040341 | 2.405061982 | Sepp1 | selenoprotein | Extracellular S | other |
| -3.620590686 | -2.403850912 | Pls3 | plastin 3 | Cytoplasm | other |
| 3.613221473 | 2.395103636 | Serpina1d | serpin family | Extracellular S | other |
| -3.606794754 | -2.388509128 | Anxa2 | annexin A2 | Plasma Mem | other |
| -3.605078321 | -2.386909984 | Rab4a | RAB4A, mem | Cytoplasm | enzyme |
| -3.603834311 | -2.385977972 | 1700023F06R | RIKEN cDNA | Other | other |
| -3.597764648 | -2.379036541 | Il17rd | interleukin 17 | Cytoplasm | other |
| -3.5931454 | -2.37331562 | Nsg1 | neuron speci | Cytoplasm | other |
| -3.592022513 | -2.37254885 | Enpp2 | ectonucleotic | Plasma Mem | enzyme |
| -3.578404296 | -2.355949242 | Mt2 | metallothione | Other | other |
| -3.57800786 | -2.355949242 | Ripply3 | ripply transcr | Nucleus | other |
| -3.571375422 | -2.348982152 | Fam174b | family with se | Other | other |
| -3.560941356 | -2.338224848 | Nudt10 | nudix hydrola | Cytoplasm | phosphatase |
| -3.560651061 | -2.338224848 | Cfap43 | NA | NA | NA |
| 3.556444766 | 2.333790062 | Apom | apolipoprote | Plasma Mem | transporter |
| -3.556125435 | -2.333790062 | Mycn | MYCN proto- | Nucleus | transcription regulator |
| -3.551072683 | -2.327400929 | Ptch2 | patched 2 | Plasma Mem | transmembrane receptor |
| -3.548283156 | -2.324232686 | Tmem30b | transmembra | Cytoplasm | other |
| -3.535702816 | -2.307891733 | Sox6 | SRY-box 6 | Nucleus | transcription regulator |
| -3.523219738 | -2.290837899 | Kalrn | kalirin RhoGE | Cytoplasm | kinase |
| -3.52173105 | -2.28951534 | Pknox2 | PBX/knotted | Nucleus | other |
| -3.514816434 | -2.282027571 | Col5a2 | collagen type | Extracellular S | other |
| -3.512659085 | -2.279739405 | Myof | myoferlin | Nucleus | other |
| -3.507963998 | -2.273806631 | Hvcn1 | hydrogen vo | Plasma Mem | ion channel |
| -3.506243679 | -2.272141555 | Col4a5 | collagen type | Extracellular S | other |
| -3.501952819 | -2.266781693 | Plagl1 | PLAG1 like zi | Nucleus | transcription regulator |
| -3.499952675 | -2.264710696 | Nuak2 | NUAK family | Other | kinase |
| -3.496598519 | -2.260690857 | Usp25 | ubiquitin spe | Cytoplasm | peptidase |
| -3.494853197 | -2.258982583 | Gm14685 | predicted ge | Other | other |
| -3.493756555 | -2.258205919 | Sp5 | Sp5 transcrip | Nucleus | other |
| -3.47410793 | -2.23308983 | Ifi30 | IFI30, lysosor | Cytoplasm | enzyme |
| -3.466933136 | -2.223707538 | Crlf1 | cytokine rece | Extracellular S | other |
| -3.466506878 | -2.223707538 | Arhgap8 | Rho GTPase a | Cytoplasm | other |
| -3.452528209 | -2.205868793 | Islr | immunoglob | Extracellular S | other |
| -3.438063947 | -2.188068605 | DXBay18 | predicted ge | Other | other |
| -3.43715872 | -2.187550494 | 4931428F04R | KIAA0895 like | Other | other |
| -3.429742415 | -2.178357274 | Kcnk2 | potassium tw | Plasma Mem | ion channel |
| -3.429029274 | -2.178114887 | Pabpc4l | poly(A) bindi | Other | other |
| -3.425816616 | -2.174232814 | Homer2 | homer scaffo | Plasma Mem | other |
| -3.423221243 | -2.172039408 | Siae | sialic acid ace | Cytoplasm | enzyme |
| -3.417553449 | -2.165215266 | Rab15 | RAB15, mem | Cytoplasm | enzyme |
| 3.416909632 | 2.165215266 | Naif1 | nuclear apop | Nucleus | other |
| -3.41635007 | -2.165187854 | Syde1 | synapse defe | Cytoplasm | other |
| 3.413055865 | 2.161173183 | Enpp4 | ectonucleotic | Cytoplasm | enzyme |
| 3.412271798 | 2.16081611 | Ttpa | alpha tocoph | Cytoplasm | transporter |
| -3.40884644 | -2.156605778 | Gata5 | GATA bindin | Nucleus | transcription regulator |
| -3.406283432 | -2.153650646 | Ano1 | anoctamin 1 | Plasma Mem | ion channel |
| 3.403852002 | 2.151642634 | Mmp19 | matrix metall | Extracellular S | peptidase |
| 3.394248893 | 2.138760268 | Npb | neuropeptide | Extracellular S | other |
| 3.393946868 | 2.138760268 | Gchfr | GTP cyclohyd | Cytoplasm | other |
| -3.387211908 | -2.129685383 | Dnhd1 | dynein heavy | Cytoplasm | other |
| 3.386095224 | 2.129613506 | Atp2b3 | ATPase plasn | Plasma Mem | transporter |

| | | | | | |
|---|---|---|---|---|---|
| -3.384998227 | -2.128786714 | Cdc42ep3 | CDC42 effect | Cytoplasm | other |
| -3.384453417 | -2.128767201 | Msx2 | msh homeob | Nucleus | transcription regulator |
| -3.376227794 | -2.118268172 | Prss23 | protease, ser | Extracellular S | peptidase |
| 3.36831816 | 2.108213235 | Pcnxl2 | pecanex hom | Other | other |
| -3.355879621 | -2.092250257 | Col11a1 | collagen type | Extracellular S | other |
| 3.350819228 | 2.086348036 | Efemp1 | EGF containi | Extracellular S | enzyme |
| -3.349282651 | -2.084970013 | Tpbg | trophoblast g | Plasma Mem | other |
| 3.344390558 | 2.079192712 | C1s1 | complement | Extracellular S | peptidase |
| -3.34381031 | -2.079109626 | Pcdhga5 | protocadheri | Other | other |
| -3.339228639 | -2.073120094 | Zbtb8a | zinc finger ar | Nucleus | other |
| 3.335411576 | 2.070563593 | Bche | butyrylcholin | Plasma Mem | enzyme |
| -3.33374116 | -2.068863899 | Igfbp3 | insulin like gr | Extracellular S | other |
| -3.331528542 | -2.066361889 | Ccdc15 | coiled-coil de | Cytoplasm | other |
| 3.329945112 | 2.066078368 | Tesk2 | testis-specifi | Nucleus | kinase |
| -3.329785641 | -2.066078368 | Nhs | NHS actin rei | Nucleus | other |
| -3.326014146 | -2.061268186 | Moxd1 | monooxygen | Cytoplasm | enzyme |
| -3.317590806 | -2.049568914 | Rap1gap | RAP1 GTPase | Cytoplasm | other |
| -3.31076993 | -2.040986713 | Zfp839 | zinc finger pr | Other | other |
| -3.308561738 | -2.039235349 | Arhgef26 | Rho guanine | Plasma Mem | other |
| -3.306102553 | -2.036347687 | Arhgef5 | Rho guanine | Cytoplasm | other |
| -3.3014319 | -2.03017432 | Tekt1 | tektin 1 | Cytoplasm | other |
| -3.300456585 | -2.029485466 | Samd12 | sterile alpha | Other | other |
| -3.299244563 | -2.02844384 | Stk26 | serine/threor | Nucleus | kinase |
| -3.295750655 | -2.025417646 | 2200002D01I | RIKEN cDNA | Nucleus | other |
| -3.295679781 | -2.025417646 | A4galt | alpha 1,4-ga | Cytoplasm | enzyme |
| -3.291930479 | -2.021354119 | Cdc25b | cell division c | Nucleus | phosphatase |
| -3.290087718 | -2.019367499 | Tor3a | torsin family | Cytoplasm | enzyme |
| -3.289328395 | -2.018990381 | Wnk3 | WNK lysine c | Plasma Mem | kinase |
| -3.284580274 | -2.015186369 | Gpm6b | glycoprotein | Plasma Mem | other |
| 3.282217985 | 2.012905379 | Exoc3l4 | exocyst comp | Other | other |
| -3.280953879 | -2.011769588 | Stk39 | serine/threor | Nucleus | kinase |
| -3.280370971 | -2.011645823 | Tmcc2 | transmembra | Extracellular S | other |
| -3.262023331 | -1.987274288 | Erbb2 | erb-b2 recep | Plasma Mem | kinase |
| -3.251952599 | -1.975191936 | Rnf32 | ring finger pr | Cytoplasm | other |
| -3.240833448 | -1.961502242 | Lamc2 | laminin subu | Extracellular S | other |
| -3.238828126 | -1.959235159 | Klk8 | kallikrein rela | Extracellular S | peptidase |
| -3.237233065 | -1.95831184 | Prickle2 | prickle plana | Nucleus | other |
| 3.226709928 | 1.945469141 | Slc16a7 | solute carrier | Plasma Mem | transporter |
| -3.22491155 | -1.943501871 | Perp | PERP, TP53 a | Plasma Mem | other |
| -3.221954647 | -1.940525455 | Cers1 | growth differ | Extracellular S | growth factor |
| -3.22139901 | -1.940423337 | Zbtb49 | zinc finger ar | Nucleus | other |
| -3.216017744 | -1.933952194 | Tgfb1i1 | transforming | Nucleus | transcription regulator |
| -3.215880268 | -1.933952194 | 1700019G17I | N-acetyltrans | Other | enzyme |
| -3.204279566 | -1.919026472 | Zfp811 | zinc finger pr | Other | other |
| -3.197083641 | -1.90962074 | Nynrin | NYN domain | Other | other |
| -3.191326981 | -1.903835561 | Tmem25 | transmembra | Extracellular S | other |
| -3.190734475 | -1.903667089 | Rbms3 | RNA binding | Other | other |
| -3.187036543 | -1.898798667 | Camk1g | calcium/calm | Cytoplasm | kinase |
| -3.185366917 | -1.897723689 | Sulf1 | sulfatase 1 | Cytoplasm | enzyme |
| -3.183734535 | -1.895977174 | Cdx2 | caudal type h | Nucleus | transcription regulator |
| -3.175197891 | -1.884487586 | Adrbk2 | G protein-co | Cytoplasm | kinase |
| -3.155154571 | -1.859080719 | Fam214b | family with se | Nucleus | other |
| 3.151048987 | 1.85354458 | Trim25 | tripartite mot | Cytoplasm | transcription regulator |
| -3.148475235 | -1.850810405 | Efr3b | EFR3 homolo | Plasma Mem | other |
| -3.143410405 | -1.844776599 | Mctp2 | multiple C2 a | Other | other |
| -3.133874932 | -1.831636596 | Sgtb | small glutam | Other | other |
| -3.129531113 | -1.82714952 | Pcdhga7 | protocadheri | Other | other |
| -3.124085542 | -1.820962291 | Fzd6 | frizzled class | Plasma Mem | G-protein coupled recepto |
| -3.118038065 | -1.814699379 | Tmem151b | transmembra | Other | other |

| | | | | | |
|---|---|---|---|---|---|
| 3.117665013 | 1.814699379 | Icosl | inducible T-c | Plasma Mem | other |
| -3.110619884 | -1.806023075 | Vat1 | vesicle amine | Plasma Mem | transporter |
| -3.10996324 | -1.805728087 | Fbxl7 | F-box and le | Cytoplasm | enzyme |
| -3.104663381 | -1.799932801 | Acss3 | acyl-CoA syr | Cytoplasm | enzyme |
| -3.104341385 | -1.799932801 | Chac2 | ChaC cation | Cytoplasm | enzyme |
| 3.10233723 | 1.797559913 | Lrrtm3 | leucine rich r | Plasma Mem | other |
| -3.101826527 | -1.797484119 | Efnb1 | ephrin B1 | Plasma Mem | other |
| 3.096777598 | 1.791129062 | Lect2 | leukocyte cel | Extracellular S | other |
| -3.095448849 | -1.790287667 | Tinagl1 | tubulointerst | Extracellular S | transporter |
| -3.090807757 | -1.786170147 | Tmem27 | transmembra | Plasma Mem | other |
| 3.089753833 | 1.78524952 | Trpv4 | transient rece | Plasma Mem | ion channel |
| -3.087615222 | -1.783358017 | Magel2 | MAGE family | Cytoplasm | enzyme |
| 3.084595798 | 1.781502748 | Hist1h2ac | histone clust | Nucleus | other |
| -3.071470324 | -1.76681508 | Pmm1 | phosphomar | Cytoplasm | enzyme |
| -3.06632356 | -1.760939177 | Abcc4 | ATP binding | Plasma Mem | transporter |
| -3.049039317 | -1.736771443 | Ptprd | protein tyros | Plasma Mem | phosphatase |
| -3.045094475 | -1.731508529 | Asxl3 | additional se | Other | other |
| 3.043857058 | 1.731508529 | Rasal3 | RAS protein a | Cytoplasm | other |
| 3.033473297 | 1.71962384 | Enho | energy home | Plasma Mem | other |
| 3.024244489 | 1.708674103 | Ada | adenosine de | Cytoplasm | enzyme |
| -3.021551557 | -1.705625616 | Spef1 | sperm flagell | Cytoplasm | other |
| -3.017893223 | -1.702185124 | Lppr1 | phospholipid | Nucleus | other |
| 3.016312842 | 1.701082889 | Hist1h4d | histone clust | Nucleus | other |
| -3.015580318 | -1.700774444 | Sfn | stratifin | Cytoplasm | other |
| -3.001724722 | -1.687763483 | Smpdl3a | sphingomyel | Extracellular S | enzyme |
| -2.999589684 | -1.686206049 | Cx3cl1 | C-X3-C moti | Extracellular S | cytokine |
| 2.988815708 | 1.670899486 | Tmprss6 | transmembra | Plasma Mem | peptidase |
| 2.985244121 | 1.667954512 | Osgin1 | oxidative stre | Other | growth factor |
| -2.98148314 | -1.662740528 | Fam83b | family with se | Cytoplasm | other |
| 2.976097184 | 1.656612749 | C2cd2l | C2CD2 like | Cytoplasm | transporter |
| -2.974561429 | -1.6558681 | Pkd1l1 | polycystin 1 l | Plasma Mem | ion channel |
| -2.965033405 | -1.644197535 | Ppp1r14a | protein phos | Cytoplasm | phosphatase |
| 2.964301434 | 1.644197535 | A1cf | APOBEC1 co | Nucleus | other |
| 2.959188157 | 1.639865493 | Hlf | HLF, PAR bZI | Nucleus | transcription regulator |
| 2.958091607 | 1.639865493 | Tbxa2r | thromboxane | Plasma Mem | G-protein coupled recepto |
| -2.957059067 | -1.639515198 | Paqr8 | progestin an | Plasma Mem | other |
| -2.952779146 | -1.634769568 | Hs3st1 | heparan sulfa | Cytoplasm | enzyme |
| -2.95139231 | -1.633236363 | Arhgef19 | Rho guanine | Cytoplasm | other |
| -2.943986959 | -1.62309698 | Chd3 | chromodoma | Nucleus | enzyme |
| -2.94099761 | -1.619676531 | Dbndd1 | dysbindin do | Other | other |
| -2.937295275 | -1.616186907 | Lhfp | lipoma HMG | Other | other |
| -2.922653556 | -1.598833695 | Adam11 | ADAM metal | Plasma Mem | peptidase |
| -2.919612849 | -1.59715039 | Fsbp | fibrinogen sil | Nucleus | other |
| -2.914151553 | -1.591829086 | Pkp3 | plakophilin 3 | Plasma Mem | other |
| 2.913577937 | 1.591829086 | Cldn1 | claudin 1 | Plasma Mem | other |
| -2.90926474 | -1.586617735 | Cyp4f16 | cytochrome l | Cytoplasm | enzyme |
| 2.901075064 | 1.577397439 | Hist1h2ad | histone clust | Nucleus | other |
| -2.891110266 | -1.564097943 | Ssfa2 | sperm specifi | Plasma Mem | other |
| 2.890011541 | 1.563585809 | Tmem82 | transmembra | Other | other |
| -2.884710859 | -1.558299278 | Mapk4 | mitogen-acti | Cytoplasm | kinase |
| -2.861559846 | -1.528177103 | Enpp3 | ectonucleoti | Plasma Mem | enzyme |
| -2.855634767 | -1.521179548 | Nap1l2 | nucleosome | Nucleus | other |
| -2.851240555 | -1.517261197 | Cdon | cell adhesion | Plasma Mem | other |
| -2.848624035 | -1.514305783 | Spock2 | SPARC/ostec | Extracellular S | other |
| 2.83169464 | 1.492627877 | Itih3 | inter-alpha-t | Extracellular S | other |
| -2.830191138 | -1.49082229 | Plch1 | phospholipas | Cytoplasm | enzyme |
| -2.829087046 | -1.489662337 | Nuak1 | NUAK family | Nucleus | kinase |
| 2.823531888 | 1.483794817 | F7 | coagulation f | Plasma Mem | peptidase |
| -2.81504286 | -1.473769712 | Bsn | bassoon pres | Plasma Mem | other |

| | | | | | |
|---|---|---|---|---|---|
| -2.812736061 | -1.471891499 | C78339 | family with se | Other | other |
| -2.803532713 | -1.460317438 | Rpp25 | ribonuclease | Nucleus | other |
| -2.795315947 | -1.450366595 | Rnf165 | ring finger pr | Nucleus | enzyme |
| -2.792790068 | -1.449325079 | Clgn | calmegin | Cytoplasm | peptidase |
| -2.783972867 | -1.439251439 | March2 | membrane as | Cytoplasm | enzyme |
| -2.777605084 | -1.431907387 | Myl3 | myosin light | Cytoplasm | other |
| -2.776720276 | -1.431072474 | Shroom2 | shroom famil | Plasma Mem | ion channel |
| -2.765869403 | -1.418183618 | Psca | prostate sten | Plasma Mem | other |
| 2.761782535 | 1.413305333 | Hgfac | HGF activato | Extracellular S | peptidase |
| -2.749871816 | -1.396441821 | Angpt2 | angiopoietin | Extracellular S | growth factor |
| -2.74577314 | -1.39426178 | Tagln | transgelin | Cytoplasm | other |
| -2.738868622 | -1.387842014 | Gprc5b | G protein-co | Plasma Mem | G-protein coupled recepto |
| -2.738582175 | -1.387842014 | S100g | S100 calcium | Cytoplasm | other |
| -2.710593538 | -1.360022062 | Lrch1 | leucine rich r | Other | other |
| -2.705198249 | -1.353612258 | Cyb5rl | cytochrome b | Nucleus | enzyme |
| 2.704661853 | 1.353612258 | Gstm6 | glutathione S | Cytoplasm | enzyme |
| -2.704583668 | -1.353612258 | Arap3 | ArfGAP with | Cytoplasm | other |
| -2.70269771 | -1.353414608 | Des | desmin | Cytoplasm | other |
| -2.679370827 | -1.328593515 | Col4a6 | collagen type | Extracellular S | other |
| -2.66905487 | -1.317724333 | Wtip | WT1 interact | Nucleus | transcription regulator |
| -2.653928467 | -1.296443518 | Bzrap1 | TSPO associa | Cytoplasm | other |
| -2.634702663 | -1.271575414 | Pkhd1 | PKHD1, fibro | Plasma Mem | other |
| -2.614152964 | -1.251093864 | Smim6 | small integra | Other | other |
| -2.589755099 | -1.218686705 | Coro2a | coronin 2A | Cytoplasm | other |
| 2.571187712 | 1.200798726 | Itih1 | inter-alpha-t | Extracellular S | other |
| 2.554654369 | 1.181835313 | Tgfbi | transforming | Extracellular S | other |
| -2.554504219 | -1.181835313 | Itpkc | inositol-trisp | Cytoplasm | kinase |
| -2.534937417 | -1.162999287 | Jag1 | jagged 1 | Extracellular S | growth factor |

ɔr

ɔr

receptor

or

or

or

or

or

or

ɔr

or

or

### RXRA motif genes

| ID | Entrez Gene Name | Location | Family |
|---|---|---|---|
| Agt | angiotensinogen | Extracellular Space | growth factor |
| Ahsg | alpha 2-HS glycoprotein | Extracellular Space | other |
| Alb | albumin | Extracellular Space | transporter |
| Aldob | aldolase, fructose-bisphosphate B | Cytoplasm | enzyme |
| Apob | apolipoprotein B | Extracellular Space | transporter |
| Apoc2 | apolipoprotein C2 | Extracellular Space | transporter |
| Arhgap36 | Rho GTPase activating protein 36 | Cytoplasm | other |
| Bbox1 | gamma-butyrobetaine hydroxylase 1 | Cytoplasm | enzyme |
| C1ra | complement C1r | Extracellular Space | peptidase |
| C3 | complement C3 | Extracellular Space | peptidase |
| Ces2a | carboxylesterase 2A | Cytoplasm | enzyme |
| Cxcl10 | C-X-C motif chemokine ligand 10 | Extracellular Space | cytokine |
| Cyp2d10 | cytochrome P450, family 2, subfamily d, poly | Cytoplasm | other |
| Dnajc22 | DnaJ heat shock protein family (Hsp40) mem | Other | other |
| F12 | coagulation factor XII | Extracellular Space | peptidase |
| Fabp1 | fatty acid binding protein 1 | Cytoplasm | transporter |
| Fga | fibrinogen alpha chain | Extracellular Space | other |
| Gchfr | GTP cyclohydrolase I feedback regulator | Cytoplasm | other |
| Gltpd2 | glycolipid transfer protein domain containin | Other | other |
| Gulo | gulonolactone (L-) oxidase | Cytoplasm | enzyme |
| Hba-x | hemoglobin subunit zeta | Cytoplasm | transporter |
| Hebp1 | heme binding protein 1 | Cytoplasm | other |
| Hmgcs2 | 3-hydroxy-3-methylglutaryl-CoA synthase 2 | Cytoplasm | enzyme |
| Icam1 | intercellular adhesion molecule 1 | Plasma Membrane | transmembra |
| Itih4 | inter-alpha-trypsin inhibitor heavy chain fan | Extracellular Space | other |
| Kng1 | kininogen 1 | Extracellular Space | other |
| Kng2 | kininogen 1 | Extracellular Space | other |
| Lcat | lecithin-cholesterol acyltransferase | Extracellular Space | enzyme |
| Leap2 | liver enriched antimicrobial peptide 2 | Extracellular Space | other |
| Lgals3bp | galectin 3 binding protein | Plasma Membrane | transmembra |
| Mocos | molybdenum cofactor sulfurase | Cytoplasm | enzyme |
| Naprt | nicotinate phosphoribosyltransferase | Cytoplasm | enzyme |
| Ngef | neuronal guanine nucleotide exchange facto | Cytoplasm | other |
| Nlrp6 | NLR family pyrin domain containing 6 | Plasma Membrane | G-protein co |
| Osgin1 | oxidative stress induced growth inhibitor 1 | Other | growth factor |
| Otc | ornithine carbamoyltransferase | Cytoplasm | enzyme |
| Pdk4 | pyruvate dehydrogenase kinase 4 | Cytoplasm | kinase |
| Pdzk1 | PDZ domain containing 1 | Plasma Membrane | transporter |
| Pla2g12b | phospholipase A2 group XIIB | Extracellular Space | enzyme |
| Proc | protein C, inactivator of coagulation factors | Extracellular Space | peptidase |
| Sardh | sarcosine dehydrogenase | Cytoplasm | enzyme |
| Serpina11 | serpin family A member 11 | Extracellular Space | other |
| Serpina6 | serpin family A member 6 | Extracellular Space | other |
| Sgk2 | SGK2, serine/threonine kinase 2 | Cytoplasm | kinase |
| Slc27a5 | solute carrier family 27 member 5 | Cytoplasm | transporter |
| Slc36a2 | solute carrier family 36 member 2 | Plasma Membrane | transporter |
| Slc6a13 | solute carrier family 6 member 13 | Plasma Membrane | transporter |
| Spp2 | secreted phosphoprotein 2 | Extracellular Space | other |
| Fam195a | - | - | - |

upled receptor

### Dynamic gene expression during hepatoblast developmen

**Instructions:**

- **p_value :** p-value (unadjusted)
- **q_value :** Adjusted p-value, based on bonferroni correction using all genes

| ID | p-value | q-value | gene_short_name | Description | Location |
|---|---|---|---|---|---|
| ENSMUSG00000022868 | 0 | 0 | Ahsg | alpha 2-HS g | Extracellular S |
| ENSMUSG00000029368 | 0 | 0 | Alb | albumin | Extracellular S |
| ENSMUSG00000028307 | 0 | 0 | Aldob | aldolase, fruc | Cytoplasm |
| ENSMUSG00000032083 | 0 | 0 | Apoa1 | apolipoprote | Extracellular S |
| ENSMUSG00000000049 | 0 | 0 | Apoh | apolipoprote | Extracellular S |
| ENSMUSG00000021728 | 0 | 0 | Emb | embigin | Plasma Mem |
| ENSMUSG00000054422 | 0 | 0 | Fabp1 | fatty acid bin | Cytoplasm |
| ENSMUSG00000035540 | 0 | 0 | Gc | GC, vitamin D | Extracellular S |
| ENSMUSG00000069919 | 0 | 0 | Hba-a1 | hemoglobin | Extracellular S |
| ENSMUSG00000069917 | 0 | 0 | Hba-a2 | hemoglobin | Extracellular S |
| ENSMUSG00000073940 | 0 | 0 | Hbb-bt | hemoglobin | Cytoplasm |
| ENSMUSG00000052305 | 0 | 0 | Hbb-bs | hemoglobin | Cytoplasm |
| ENSMUSG00000020429 | 0 | 0 | Igfbp1 | insulin like gr | Extracellular S |
| ENSMUSG00000022875 | 0 | 0 | Kng1 | kininogen 1 | Extracellular S |
| ENSMUSG00000027239 | 0 | 0 | Mdk | midkine | Extracellular S |
| ENSMUSG00000032294 | 0 | 0 | Pkm | pyruvate kina | Cytoplasm |
| ENSMUSG00000001525 | 1.16E-307 | 4.80E-305 | Tubb5 | tubulin beta | Cytoplasm |
| ENSMUSG00000031762 | 1.16E-302 | 4.59E-300 | Mt2 | metallothion | Other |
| ENSMUSG00000031765 | 3.91E-300 | 1.47E-297 | Mt1 | metallothion | Cytoplasm |
| ENSMUSG00000046434 | 3.95E-298 | 1.43E-295 | Hnrnpa1 | heterogeneo | Nucleus |
| ENSMUSG00000039438 | 2.57E-295 | 8.90E-293 | Ttc36 | tetratricopep | Other |
| ENSMUSG00000047945 | 1.72E-293 | 5.70E-291 | Marcksl1 | MARCKS like | Cytoplasm |
| ENSMUSG00000030246 | 3.07E-293 | 9.80E-291 | Ldhb | lactate dehyc | Cytoplasm |
| ENSMUSG00000072849 | 2.03E-277 | 6.25E-275 | Serpina1e | serpin family | Extracellular S |
| ENSMUSG00000027559 | 1.25E-274 | 3.71E-272 | Car3 | carbonic anh | Cytoplasm |
| ENSMUSG00000061808 | 5.98E-274 | 1.71E-271 | Ttr | transthyretin | Extracellular S |
| ENSMUSG00000030359 | 3.37E-267 | 9.32E-265 | Pzp | PZP, alpha-2 | Extracellular S |
| ENSMUSG00000074336 | 4.42E-263 | 1.18E-260 | Apoc4 | apolipoprote | Extracellular S |
| ENSMUSG00000028001 | 1.70E-259 | 4.41E-257 | Fga | fibrinogen al | Extracellular S |
| ENSMUSG00000005732 | 2.51E-253 | 6.32E-251 | Ranbp1 | RAN binding | Nucleus |
| ENSMUSG00000057113 | 1.01E-250 | 2.46E-248 | Npm1 | nucleophosm | Nucleus |
| ENSMUSG00000026295 | 2.51E-246 | 5.95E-244 | Spp2 | secreted pho | Extracellular S |
| ENSMUSG00000000805 | 3.06E-246 | 7.05E-244 | Car4 | carbonic anh | Plasma Mem |
| ENSMUSG00000058216 | 1.29E-245 | 2.90E-243 | BC021614 | glutathione S | Other |
| ENSMUSG00000023906 | 3.29E-244 | 7.19E-242 | Cldn6 | claudin 6 | Plasma Mem |
| ENSMUSG00000071177 | 1.46E-243 | 3.10E-241 | Serpina1d | serpin family | Extracellular S |
| ENSMUSG00000022445 | 1.42E-238 | 2.94E-236 | Cyp2d26 | cytochrome P | Cytoplasm |
| ENSMUSG00000079015 | 9.90E-238 | 2.00E-235 | Serpina1c | serpin family | Extracellular S |
| ENSMUSG00000060459 | 1.48E-228 | 2.93E-226 | Kng2 | kininogen 1 | Extracellular S |
| ENSMUSG00000031980 | 2.60E-226 | 5.01E-224 | Agt | angiotensino | Extracellular S |
| ENSMUSG00000021922 | 5.00E-225 | 9.44E-223 | Itih4 | inter-alpha-t | Extracellular S |
| ENSMUSG00000066366 | 2.12E-214 | 3.91E-212 | Serpina1a | serpin family | Extracellular S |
| ENSMUSG00000026238 | 8.07E-212 | 1.46E-209 | Ptma | prothymosin | Nucleus |
| ENSMUSG00000030895 | 1.83E-206 | 3.24E-204 | Hpx | hemopexin | Extracellular S |
| ENSMUSG00000033860 | 7.04E-206 | 1.22E-203 | Fgg | fibrinogen ga | Extracellular S |
| ENSMUSG00000032554 | 2.17E-205 | 3.68E-203 | Trf | transferrin | Extracellular S |
| ENSMUSG00000057278 | 2.04E-196 | 3.39E-194 | Snrpg | small nuclear | Nucleus |
| ENSMUSG00000003038 | 2.45E-194 | 3.98E-192 | Hmgn2 | high mobility | Nucleus |
| ENSMUSG00000037894 | 4.67E-194 | 7.46E-192 | H2afz | H2A histone | Nucleus |
| ENSMUSG00000023004 | 1.29E-192 | 2.01E-190 | Tuba1b | tubulin alpha | Cytoplasm |

| | | | | | |
|---|---|---|---|---|---|
| ENSMUSG00000027875 | 1.31E-186 | 2.01E-184 | Hmgcs2 | 3-hydroxy-3 | Cytoplasm |
| ENSMUSG00000076441 | 2.24E-185 | 3.38E-183 | Ass1 | argininosucci | Cytoplasm |
| ENSMUSG00000025991 | 6.52E-182 | 9.66E-180 | Cps1 | carbamoyl-p | Cytoplasm |
| ENSMUSG00000029445 | 1.27E-181 | 1.85E-179 | Hpd | 4-hydroxyph | Cytoplasm |
| ENSMUSG00000071178 | 2.26E-181 | 3.23E-179 | Serpina1b | serpin family | Extracellular S |
| ENSMUSG00000026542 | 5.56E-181 | 7.83E-179 | Apcs | amyloid P co | Extracellular S |
| ENSMUSG00000008540 | 1.75E-180 | 2.43E-178 | Mgst1 | microsomal g | Cytoplasm |
| ENSMUSG00000031722 | 3.26E-180 | 4.43E-178 | Hp | haptoglobin | Extracellular S |
| ENSMUSG00000028044 | 3.64E-180 | 4.87E-178 | Cks1b | CDC28 prote | Other |
| ENSMUSG00000021270 | 1.36E-179 | 1.79E-177 | Hsp90aa1 | heat shock p | Cytoplasm |
| ENSMUSG00000027761 | 1.70E-177 | 2.20E-175 | Aadac | arylacetamid | Cytoplasm |
| ENSMUSG00000023070 | 8.27E-176 | 1.06E-173 | Rgn | regucalcin | Nucleus |
| ENSMUSG00000020649 | 7.81E-175 | 9.82E-173 | Rrm2 | ribonucleotic | Nucleus |
| ENSMUSG00000024164 | 1.03E-174 | 1.28E-172 | C3 | complement | Extracellular S |
| ENSMUSG00000054717 | 1.60E-171 | 1.93E-169 | Hmgb2 | high mobility | Nucleus |
| ENSMUSG00000030695 | 1.53E-169 | 1.82E-167 | Aldoa | aldolase, fruc | Cytoplasm |
| ENSMUSG00000091898 | 5.26E-169 | 6.15E-167 | Tnnc1 | troponin C1, | Cytoplasm |
| ENSMUSG00000024386 | 2.91E-168 | 3.35E-166 | Proc | protein C, ina | Extracellular S |
| ENSMUSG00000030711 | 3.96E-167 | 4.51E-165 | Sult1a1 | sulfotransfera | Cytoplasm |
| ENSMUSG00000001123 | 4.38E-166 | 4.91E-164 | Lgals9 | galectin 9B | Cytoplasm |
| ENSMUSG00000037798 | 3.51E-165 | 3.88E-163 | Mat1a | methionine a | Cytoplasm |
| ENSMUSG00000064373 | 1.12E-164 | 1.22E-162 | Sepp1 | selenoprotei | Extracellular S |
| ENSMUSG00000054932 | 1.38E-164 | 1.49E-162 | Afp | alpha fetopro | Extracellular S |
| ENSMUSG00000015217 | 1.54E-163 | 1.64E-161 | Hmgb3 | high mobility | Nucleus |
| ENSMUSG00000032324 | 6.79E-161 | 7.13E-159 | Tspan3 | tetraspanin 3 | Plasma Mem |
| ENSMUSG00000031844 | 5.24E-158 | 5.44E-156 | Hsd17b2 | hydroxystero | Cytoplasm |
| ENSMUSG00000035561 | 7.39E-158 | 7.57E-156 | Aldh1b1 | aldehyde deh | Cytoplasm |
| ENSMUSG00000024503 | 2.72E-157 | 2.75E-155 | Spink3 | serine peptid | Extracellular S |
| ENSMUSG00000021131 | 1.47E-156 | 1.47E-154 | Erh | ERH, mRNA s | Nucleus |
| ENSMUSG00000060807 | 3.98E-156 | 3.93E-154 | Serpina6 | serpin family | Extracellular S |
| ENSMUSG00000054766 | 6.40E-155 | 6.24E-153 | Set | SET nuclear p | Nucleus |
| ENSMUSG00000040564 | 3.41E-153 | 3.29E-151 | Apoc1 | apolipoprote | Extracellular S |
| ENSMUSG00000025161 | 1.27E-152 | 1.21E-150 | Slc16a3 | solute carrier | Plasma Mem |
| ENSMUSG00000039114 | 3.89E-151 | 3.67E-149 | Nrn1 | neuritin 1 | Cytoplasm |
| ENSMUSG00000014226 | 1.70E-149 | 1.58E-147 | Cacybp | calcyclin binc | Nucleus |
| ENSMUSG00000024097 | 2.40E-146 | 2.21E-144 | Srsf7 | serine and ar | Nucleus |
| ENSMUSG00000072235 | 4.17E-146 | 3.81E-144 | Tuba1a | tubulin alpha | Cytoplasm |
| ENSMUSG00000028851 | 6.45E-146 | 5.82E-144 | Nudc | nuclear distri | Cytoplasm |
| ENSMUSG00000028832 | 9.16E-146 | 8.18E-144 | Stmn1 | stathmin 1 | Cytoplasm |
| ENSMUSG00000090553 | 9.46E-146 | 8.35E-144 | Snrpe | small nuclear | Nucleus |
| ENSMUSG00000015656 | 3.03E-144 | 2.64E-142 | Hspa8 | heat shock p | Cytoplasm |
| ENSMUSG00000029102 | 3.64E-142 | 3.14E-140 | Hgfac | HGF activato | Extracellular S |
| ENSMUSG00000034192 | 6.43E-142 | 5.50E-140 | Lsm3 | LSM3 homolo | Nucleus |
| ENSMUSG00000046711 | 1.86E-140 | 1.56E-138 | Hmga1 | high mobility | Nucleus |
| ENSMUSG00000049775 | 2.25E-140 | 1.86E-138 | Tmsb4x | thymosin, be | Cytoplasm |
| ENSMUSG00000059040 | 1.11E-139 | 9.09E-138 | Eno1b | enolase 1 | Cytoplasm |
| ENSMUSG00000029430 | 2.86E-139 | 2.33E-137 | Ran | RAN, membe | Nucleus |
| ENSMUSG00000000673 | 1.20E-138 | 9.64E-137 | Haao | 3-hydroxyan | Cytoplasm |
| ENSMUSG00000020186 | 1.78E-137 | 1.42E-135 | Csrp2 | cysteine and | Nucleus |
| ENSMUSG00000056201 | 5.77E-137 | 4.56E-135 | Cfl1 | cofilin 1 | Nucleus |
| ENSMUSG00000029656 | 2.48E-135 | 1.94E-133 | C8b | complement | Extracellular S |
| ENSMUSG00000063524 | 3.46E-135 | 2.68E-133 | Eno1 | enolase 1 | Cytoplasm |
| ENSMUSG00000058952 | 6.60E-135 | 5.07E-133 | Cfi | complement | Extracellular S |
| ENSMUSG00000028333 | 4.54E-132 | 3.46E-130 | Anp32b | acidic (leucin | Nucleus |
| ENSMUSG00000022769 | 9.39E-132 | 7.09E-130 | Sdf2l1 | stromal cell c | Cytoplasm |
| ENSMUSG00000002769 | 7.13E-131 | 5.33E-129 | Gnmt | glycine N-me | Cytoplasm |
| ENSMUSG00000028553 | 2.36E-130 | 1.75E-128 | Angptl3 | angiopoietin | Extracellular S |
| ENSMUSG00000078249 | 2.45E-130 | 1.80E-128 | Hmga1-rs1 | high mobility | Nucleus |

| | | | | | |
|---|---|---|---|---|---|
| ENSMUSG00000049932 | 1.22E-129 | 8.87E-128 | H2afx | H2A histone | Nucleus |
| ENSMUSG00000020949 | 1.59E-127 | 1.15E-125 | Fkbp3 | FK506 bindin | Nucleus |
| ENSMUSG00000016559 | 1.67E-126 | 1.20E-124 | H3f3b | H3 histone fa | Nucleus |
| ENSMUSG00000003429 | 3.22E-126 | 2.29E-124 | Rps11 | ribosomal pr | Cytoplasm |
| ENSMUSG00000039985 | 2.00E-124 | 1.41E-122 | Fam60a | SIN3-HDAC c | Extracellular S |
| ENSMUSG00000040681 | 3.10E-124 | 2.16E-122 | Hmgn1 | high mobility | Nucleus |
| ENSMUSG00000001056 | 1.37E-123 | 9.50E-122 | Nhp2 | NHP2 ribonu | Nucleus |
| ENSMUSG00000004885 | 1.67E-123 | 1.15E-121 | Crabp2 | cellular retinc | Cytoplasm |
| ENSMUSG00000067377 | 3.85E-120 | 2.60E-118 | Tspan6 | tetraspanin 6 | Plasma Mem |
| ENSMUSG00000025934 | 1.77E-119 | 1.19E-117 | Gsta3 | glutathione S | Cytoplasm |
| ENSMUSG00000025439 | 6.47E-119 | 4.29E-117 | Clns1a | chloride nucl | Plasma Mem |
| ENSMUSG00000023944 | 7.49E-119 | 4.94E-117 | Hsp90ab1 | heat shock p | Cytoplasm |
| ENSMUSG00000026879 | 2.67E-118 | 1.75E-116 | Gsn | gelsolin | Extracellular S |
| ENSMUSG00000024966 | 2.89E-117 | 1.87E-115 | Stip1 | stress induce | Cytoplasm |
| ENSMUSG00000006728 | 4.92E-117 | 3.17E-115 | Cdk4 | cyclin depend | Nucleus |
| ENSMUSG00000033307 | 1.78E-115 | 1.13E-113 | Mif | macrophage | Extracellular S |
| ENSMUSG00000040459 | 2.37E-115 | 1.50E-113 | Arglu1 | arginine and | Other |
| ENSMUSG00000009927 | 1.22E-113 | 7.59E-112 | Rps25 | ribosomal pr | Cytoplasm |
| ENSMUSG00000002847 | 1.93E-113 | 1.20E-111 | Pla1a | phospholipas | Extracellular S |
| ENSMUSG00000000184 | 8.60E-113 | 5.29E-111 | Ccnd2 | cyclin D2 | Nucleus |
| ENSMUSG00000040883 | 1.17E-112 | 7.14E-111 | Tmem205 | transmembra | Cytoplasm |
| ENSMUSG00000059481 | 2.46E-112 | 1.49E-110 | Plg | plasminogen | Extracellular S |
| ENSMUSG00000078812 | 1.30E-110 | 7.79E-109 | Eif5a | eukaryotic tra | Cytoplasm |
| ENSMUSG00000005846 | 1.32E-110 | 7.91E-109 | Rsl1d1 | ribosomal L1 | Nucleus |
| ENSMUSG00000035215 | 1.84E-110 | 1.09E-108 | Lsm7 | LSM7 homolo | Nucleus |
| ENSMUSG00000026664 | 1.31E-109 | 7.72E-108 | Phyh | phytanoyl-Co | Cytoplasm |
| ENSMUSG00000067235 | 3.66E-109 | 2.14E-107 | H2-Q10 | major histocc | Plasma Mem |
| ENSMUSG00000044533 | 1.33E-108 | 7.72E-107 | Rps2 | ribosomal pr | Cytoplasm |
| ENSMUSG00000029447 | 9.91E-108 | 5.71E-106 | Cct6a | chaperonin c | Cytoplasm |
| ENSMUSG00000020547 | 2.26E-107 | 1.29E-105 | Bzw2 | basic leucine | Cytoplasm |
| ENSMUSG00000090733 | 2.30E-107 | 1.31E-105 | Rps27 | ribosomal pr | Other |
| ENSMUSG00000030111 | 7.97E-107 | 4.50E-105 | A2m | alpha-2-mac | Extracellular S |
| ENSMUSG00000046865 | 9.13E-107 | 5.12E-105 | Fbl | fibrillarin | Nucleus |
| ENSMUSG00000003955 | 2.09E-106 | 1.16E-104 | Fam162a | family with se | Cytoplasm |
| ENSMUSG00000043091 | 2.50E-106 | 1.38E-104 | Tuba1c | tubulin alpha | Cytoplasm |
| ENSMUSG00000009633 | 2.15E-105 | 1.18E-103 | G0s2 | G0/G1 switch | Cytoplasm |
| ENSMUSG00000025362 | 2.50E-105 | 1.36E-103 | Rps26 | ribosomal pr | Cytoplasm |
| ENSMUSG00000066551 | 8.67E-105 | 4.67E-103 | Hmgb1 | high mobility | Nucleus |
| ENSMUSG00000026021 | 3.47E-103 | 1.86E-101 | Sumo1 | small ubiquit | Nucleus |
| ENSMUSG00000029730 | 3.56E-103 | 1.89E-101 | Mcm7 | minichromos | Nucleus |
| ENSMUSG00000006442 | 6.51E-103 | 3.44E-101 | Srm | spermidine sy | Cytoplasm |
| ENSMUSG00000002477 | 1.45E-102 | 7.60E-101 | Snrpd1 | small nuclear | Nucleus |
| ENSMUSG00000008450 | 1.65E-102 | 8.60E-101 | Nutf2 | nuclear trans | Nucleus |
| ENSMUSG00000019539 | 1.83E-102 | 9.47E-101 | Rcn3 | reticulocalbir | Cytoplasm |
| ENSMUSG00000067367 | 3.04E-102 | 1.57E-100 | Lyar | Ly1 antibody | Plasma Mem |
| ENSMUSG00000003868 | 3.84E-102 | 1.97E-100 | Ruvbl2 | RuvB like AA | Nucleus |
| ENSMUSG00000032397 | 7.52E-102 | 3.83E-100 | Tipin | TIMELESS int | Nucleus |
| ENSMUSG00000050621 | 5.01E-101 | 2.53E-99 | Rps27rt | ribosomal pr | Other |
| ENSMUSG00000029836 | 9.29E-101 | 4.67E-99 | Cbx3 | chromobox 3 | Nucleus |
| ENSMUSG00000040824 | 9.56E-101 | 4.78E-99 | Snrpd2 | small nuclear | Nucleus |
| ENSMUSG00000022766 | 1.03E-99 | 5.11E-98 | Serpind1 | serpin family | Extracellular S |
| ENSMUSG00000071497 | 2.75E-99 | 1.36E-97 | Nutf2-ps1 | nuclear trans | Other |
| ENSMUSG00000020738 | 1.23E-98 | 6.02E-97 | Sumo2 | small ubiquit | Nucleus |
| ENSMUSG00000045394 | 9.40E-98 | 4.59E-96 | Epcam | epithelial cell | Plasma Mem |
| ENSMUSG00000003873 | 3.01E-97 | 1.46E-95 | Bax | BCL2 associa | Cytoplasm |
| ENSMUSG00000021999 | 3.51E-97 | 1.69E-95 | Cpb2 | carboxypepti | Extracellular S |
| ENSMUSG00000047631 | 3.56E-97 | 1.71E-95 | Apof | apolipoprote | Extracellular S |
| ENSMUSG00000027133 | 4.69E-97 | 2.24E-95 | Nop10 | NOP10 ribon | Nucleus |

| | | | | | |
|---|---|---|---|---|---|
| ENSMUSG00000063480 | 5.59E-97 | 2.65E-95 | Nhp2l1 | small nuclear | Nucleus |
| ENSMUSG00000025364 | 9.07E-97 | 4.28E-95 | Pa2g4 | proliferation- | Nucleus |
| ENSMUSG00000071644 | 1.56E-96 | 7.31E-95 | Eef1g | eukaryotic tra | Cytoplasm |
| ENSMUSG00000060373 | 2.41E-96 | 1.12E-94 | Hnrnpc | heterogeneo | Nucleus |
| ENSMUSG00000042462 | 4.64E-96 | 2.15E-94 | Dctpp1 | dCTP pyroph | Cytoplasm |
| ENSMUSG00000039878 | 6.13E-96 | 2.83E-94 | Slc39a5 | solute carrier | Plasma Mem |
| ENSMUSG00000001270 | 8.62E-96 | 3.95E-94 | Ckb | creatine kina: | Cytoplasm |
| ENSMUSG00000026234 | 1.40E-94 | 6.38E-93 | Ncl | nucleolin | Nucleus |
| ENSMUSG00000035202 | 2.33E-94 | 1.05E-92 | Lars2 | leucyl-tRNA : | Cytoplasm |
| ENSMUSG00000049517 | 2.34E-94 | 1.05E-92 | Rps23 | ribosomal pr | Cytoplasm |
| ENSMUSG00000053565 | 1.21E-93 | 5.42E-92 | Eif3k | eukaryotic tra | Cytoplasm |
| ENSMUSG00000019987 | 3.23E-93 | 1.44E-91 | Arg1 | arginase 1 | Cytoplasm |
| ENSMUSG00000036880 | 6.58E-93 | 2.92E-91 | Acaa2 | acetyl-CoA a | Cytoplasm |
| ENSMUSG00000025613 | 1.51E-91 | 6.67E-90 | Cct8 | chaperonin c | Cytoplasm |
| ENSMUSG00000031697 | 3.04E-91 | 1.34E-89 | Orc6 | origin recogr | Nucleus |
| ENSMUSG00000025176 | 3.10E-91 | 1.35E-89 | Hoga1 | 4-hydroxy-2 | Cytoplasm |
| ENSMUSG00000021751 | 5.20E-91 | 2.26E-89 | Acox2 | acyl-CoA oxi | Cytoplasm |
| ENSMUSG00000002068 | 7.22E-91 | 3.12E-89 | Ccne1 | cyclin E1 | Nucleus |
| ENSMUSG00000021811 | 1.82E-90 | 7.85E-89 | Dnajc9 | DnaJ heat sh | Nucleus |
| ENSMUSG00000027404 | 2.64E-90 | 1.13E-88 | Snrpb | small nuclear | Nucleus |
| ENSMUSG00000024844 | 4.54E-90 | 1.93E-88 | Banf1 | barrier to aut | Nucleus |
| ENSMUSG00000042607 | 6.15E-90 | 2.60E-88 | Asb4 | ankyrin repea | Nucleus |
| ENSMUSG00000062825 | 7.70E-90 | 3.24E-88 | Actg1 | actin gamma | Cytoplasm |
| ENSMUSG00000005681 | 2.02E-89 | 8.47E-88 | Apoa2 | apolipoprote | Extracellular S |
| ENSMUSG00000061360 | 2.12E-89 | 8.83E-88 | Phf5a | PHD finger p | Nucleus |
| ENSMUSG00000027405 | 2.34E-89 | 9.72E-88 | Nop56 | NOP56 ribon | Nucleus |
| ENSMUSG00000026621 | 9.24E-89 | 3.82E-87 | Marc1 | mitochondria | Cytoplasm |
| ENSMUSG00000026368 | 1.03E-88 | 4.24E-87 | F13b | coagulation f | Extracellular S |
| ENSMUSG00000060572 | 1.40E-88 | 5.73E-87 | Mfap2 | microfibril as | Extracellular S |
| ENSMUSG00000059796 | 1.60E-88 | 6.52E-87 | Eif4a1 | eukaryotic tra | Cytoplasm |
| ENSMUSG00000067274 | 1.63E-88 | 6.61E-87 | Rplp0 | ribosomal pr | Cytoplasm |
| ENSMUSG00000030630 | 2.17E-88 | 8.75E-87 | Fah | fumarylaceto | Cytoplasm |
| ENSMUSG00000019139 | 2.46E-88 | 9.86E-87 | Isyna1 | inositol-3-ph | Cytoplasm |
| ENSMUSG00000005410 | 4.75E-88 | 1.89E-86 | Mcm5 | minichromos | Nucleus |
| ENSMUSG00000015176 | 1.94E-87 | 7.72E-86 | Nolc1 | nucleolar anc | Nucleus |
| ENSMUSG00000028609 | 2.58E-87 | 1.02E-85 | Magoh | mago homol | Nucleus |
| ENSMUSG00000017344 | 3.81E-87 | 1.50E-85 | Vtn | vitronectin | Extracellular S |
| ENSMUSG00000052833 | 1.00E-86 | 3.92E-85 | Sae1 | SUMO1 activ | Cytoplasm |
| ENSMUSG00000064326 | 1.77E-86 | 6.91E-85 | Siva1 | SIVA1 apopto | Cytoplasm |
| ENSMUSG00000032047 | 5.82E-86 | 2.26E-84 | Acat1 | acetyl-CoA a | Cytoplasm |
| ENSMUSG00000019432 | 6.95E-86 | 2.68E-84 | Ddx39b | DExD-box he | Nucleus |
| ENSMUSG00000024608 | 6.99E-86 | 2.69E-84 | Rps14 | ribosomal pr | Cytoplasm |
| ENSMUSG00000026003 | 2.95E-85 | 1.13E-83 | Acadl | acyl-CoA deh | Cytoplasm |
| ENSMUSG00000023272 | 3.07E-85 | 1.17E-83 | Creld2 | cysteine rich | Other |
| ENSMUSG00000028081 | 3.93E-85 | 1.49E-83 | Rps3a1 | ribosomal pr | Cytoplasm |
| ENSMUSG00000029260 | 4.76E-85 | 1.80E-83 | Ugt2b34 | UDP glucuro | Cytoplasm |
| ENSMUSG00000022512 | 6.10E-85 | 2.29E-83 | Cldn1 | claudin 1 | Plasma Mem |
| ENSMUSG00000078713 | 8.17E-85 | 3.05E-83 | Tomm5 | translocase o | Cytoplasm |
| ENSMUSG00000036216 | 1.14E-84 | 4.26E-83 | Leap2 | liver enrichec | Extracellular S |
| ENSMUSG00000022234 | 3.39E-84 | 1.26E-82 | Cct5 | chaperonin c | Cytoplasm |
| ENSMUSG00000037780 | 4.13E-84 | 1.52E-82 | Mbl1 | mannose-bir | Cytoplasm |
| ENSMUSG00000020018 | 4.74E-84 | 1.74E-82 | Snrpf | small nuclear | Nucleus |
| ENSMUSG00000027359 | 4.91E-84 | 1.79E-82 | Slc27a2 | solute carrier | Cytoplasm |
| ENSMUSG00000027203 | 7.85E-84 | 2.86E-82 | Dut | deoxyuridine | Nucleus |
| ENSMUSG00000055730 | 8.20E-84 | 2.97E-82 | Ces2a | carboxylester | Cytoplasm |
| ENSMUSG00000063316 | 1.40E-83 | 5.05E-82 | Rpl27 | ribosomal pr | Cytoplasm |
| ENSMUSG00000019122 | 3.55E-83 | 1.27E-81 | Ccl9 | chemokine (C | Extracellular S |
| ENSMUSG00000006333 | 5.29E-83 | 1.89E-81 | Rps9 | ribosomal pr | Cytoplasm |

| | | | | | |
|---|---|---|---|---|---|
| ENSMUSG00000015083 | 5.84E-83 | 2.08E-81 | C8g | complement | Extracellular S |
| ENSMUSG00000015749 | 5.98E-83 | 2.12E-81 | Anp32e | acidic (leucin | Cytoplasm |
| ENSMUSG00000027249 | 8.36E-83 | 2.95E-81 | F2 | coagulation f | Extracellular S |
| ENSMUSG00000009281 | 2.03E-82 | 7.14E-81 | Rarres2 | retinoic acid | Plasma Mem |
| ENSMUSG00000011752 | 2.11E-82 | 7.40E-81 | Pgam1 | phosphoglyc | Cytoplasm |
| ENSMUSG00000031848 | 2.61E-82 | 9.11E-81 | Lsm4 | LSM4 homol | Nucleus |
| ENSMUSG00000034681 | 8.43E-82 | 2.93E-80 | Rnps1 | RNA binding | Nucleus |
| ENSMUSG00000047139 | 1.64E-81 | 5.64E-80 | Cd24a | CD24a antige | Plasma Mem |
| ENSMUSG00000021037 | 1.67E-81 | 5.71E-80 | Ahsa1 | activator of H | Cytoplasm |
| ENSMUSG00000061613 | 1.78E-81 | 6.09E-80 | U2af1 | U2 small nuc | Nucleus |
| ENSMUSG00000025980 | 4.81E-81 | 1.64E-79 | Hspd1 | heat shock p | Cytoplasm |
| ENSMUSG00000030057 | 7.17E-81 | 2.43E-79 | Cnbp | CCHC-type z | Nucleus |
| ENSMUSG00000022111 | 9.04E-81 | 3.05E-79 | Uchl3 | ubiquitin C-t | Cytoplasm |
| ENSMUSG00000028639 | 1.52E-80 | 5.11E-79 | Ybx1 | Y-box bindin | Nucleus |
| ENSMUSG00000047261 | 4.06E-80 | 1.36E-78 | Gap43 | growth assoc | Plasma Mem |
| ENSMUSG00000068877 | 5.44E-80 | 1.81E-78 | Selenbp2 | selenium bin | Cytoplasm |
| ENSMUSG00000062070 | 1.36E-79 | 4.49E-78 | Pgk1 | phosphoglyc | Cytoplasm |
| ENSMUSG00000057841 | 2.78E-79 | 9.17E-78 | Rpl32 | ribosomal pr | Cytoplasm |
| ENSMUSG00000076432 | 1.01E-78 | 3.31E-77 | Ywhaq | tyrosine 3-m | Cytoplasm |
| ENSMUSG00000056973 | 1.74E-78 | 5.70E-77 | Ces1d | carboxylester | Cytoplasm |
| ENSMUSG00000031173 | 2.06E-78 | 6.69E-77 | Otc | ornithine carl | Cytoplasm |
| ENSMUSG00000030357 | 2.14E-78 | 6.95E-77 | Fkbp4 | FK506 bindin | Nucleus |
| ENSMUSG00000058799 | 3.30E-78 | 1.07E-76 | Nap1l1 | nucleosome | Nucleus |
| ENSMUSG00000031939 | 3.53E-78 | 1.14E-76 | Taf1d | TATA-box bi | Nucleus |
| ENSMUSG00000020739 | 3.86E-78 | 1.24E-76 | Nup85 | nucleoporin | Cytoplasm |
| ENSMUSG00000027342 | 4.33E-78 | 1.38E-76 | Pcna | proliferating | Nucleus |
| ENSMUSG00000025794 | 4.45E-78 | 1.41E-76 | Rpl14 | ribosomal pr | Cytoplasm |
| ENSMUSG00000022336 | 5.44E-78 | 1.72E-76 | Eif3e | eukaryotic tra | Cytoplasm |
| ENSMUSG00000053907 | 9.50E-78 | 3.00E-76 | Mat2a | methionine a | Cytoplasm |
| ENSMUSG00000034024 | 1.29E-77 | 4.07E-76 | Cct2 | chaperonin c | Cytoplasm |
| ENSMUSG00000028693 | 1.62E-77 | 5.06E-76 | Nasp | nuclear auto | Nucleus |
| ENSMUSG00000030744 | 1.73E-77 | 5.41E-76 | Rps3 | ribosomal pr | Cytoplasm |
| ENSMUSG00000091625 | 1.82E-77 | 5.67E-76 | Lsm5 | LSM5 homol | Cytoplasm |
| ENSMUSG00000024833 | 2.20E-77 | 6.82E-76 | Pola2 | DNA polyme | Nucleus |
| ENSMUSG00000030867 | 2.61E-77 | 8.03E-76 | Plk1 | polo like kina | Nucleus |
| ENSMUSG00000027067 | 2.60E-77 | 8.03E-76 | Ssrp1 | structure spe | Nucleus |
| ENSMUSG00000039067 | 4.99E-77 | 1.53E-75 | Psmd7 | proteasome | Cytoplasm |
| ENSMUSG00000041841 | 5.65E-77 | 1.72E-75 | Rpl37 | ribosomal pr | Cytoplasm |
| ENSMUSG00000018446 | 8.90E-77 | 2.71E-75 | C1qbp | complement | Cytoplasm |
| ENSMUSG00000078427 | 9.89E-77 | 3.00E-75 | Sarnp | SAP domain | Nucleus |
| ENSMUSG00000007892 | 1.33E-76 | 4.02E-75 | Rplp1 | ribosomal pr | Nucleus |
| ENSMUSG00000003153 | 2.13E-76 | 6.41E-75 | Slc2a3 | solute carrier | Plasma Mem |
| ENSMUSG00000060703 | 2.66E-76 | 7.93E-75 | Cd302 | CD302 molec | Plasma Mem |
| ENSMUSG00000048856 | 7.33E-76 | 2.18E-74 | Slc25a47 | solute carrier | Cytoplasm |
| ENSMUSG00000030978 | 9.44E-76 | 2.80E-74 | Rrm1 | ribonucleotic | Nucleus |
| ENSMUSG00000029649 | 1.38E-75 | 4.07E-74 | Pomp | proteasome | Nucleus |
| ENSMUSG00000025508 | 3.65E-75 | 1.07E-73 | Rplp2 | ribosomal pr | Cytoplasm |
| ENSMUSG00000040658 | 1.83E-74 | 5.36E-73 | Dnph1 | 2'-deoxynucl | Nucleus |
| ENSMUSG00000037563 | 2.21E-74 | 6.45E-73 | Rps16 | ribosomal pr | Cytoplasm |
| ENSMUSG00000024258 | 3.20E-74 | 9.31E-73 | Polr2d | RNA polymer | Nucleus |
| ENSMUSG00000020277 | 3.80E-74 | 1.10E-72 | Pfkl | phosphofruc | Cytoplasm |
| ENSMUSG00000060938 | 5.56E-74 | 1.61E-72 | Rpl26 | ribosomal pr | Cytoplasm |
| ENSMUSG00000031645 | 1.42E-73 | 4.09E-72 | F11 | coagulation f | Extracellular S |
| ENSMUSG00000030762 | 2.04E-73 | 5.85E-72 | Aqp8 | aquaporin 8 | Plasma Mem |
| ENSMUSG00000033831 | 2.44E-73 | 6.99E-72 | Fgb | fibrinogen be | Extracellular S |
| ENSMUSG00000068749 | 1.78E-72 | 5.07E-71 | Psma5 | proteasome | Cytoplasm |
| ENSMUSG00000070704 | 2.41E-72 | 6.85E-71 | Ugt2b36 | UDP glucuro | Cytoplasm |
| ENSMUSG00000034855 | 3.94E-72 | 1.12E-70 | Cxcl10 | C-X-C motif | Extracellular S |

| | | | | | |
|---|---|---|---|---|---|
| ENSMUSG00000068882 | 4.61E-72 | 1.30E-70 | Ssb | Sjogren synd | Nucleus |
| ENSMUSG00000067608 | 9.29E-72 | 2.61E-70 | Pcna-ps2 | proliferating | Other |
| ENSMUSG00000005667 | 1.61E-71 | 4.53E-70 | Mthfd2 | methylenetet | Cytoplasm |
| ENSMUSG00000026715 | 1.77E-71 | 4.96E-70 | Serpinc1 | serpin family | Extracellular S |
| ENSMUSG00000033732 | 2.04E-71 | 5.67E-70 | Sf3b3 | splicing facto | Nucleus |
| ENSMUSG00000071072 | 2.07E-71 | 5.73E-70 | Ptges3 | prostaglandi | Cytoplasm |
| ENSMUSG00000079435 | 2.46E-71 | 6.82E-70 | Rpl36a | ribosomal pr | Cytoplasm |
| ENSMUSG00000079641 | 3.76E-71 | 1.04E-69 | Rpl39 | ribosomal pr | Cytoplasm |
| ENSMUSG00000003970 | 5.00E-71 | 1.37E-69 | Rpl8 | ribosomal pr | Cytoplasm |
| ENSMUSG00000071172 | 8.22E-71 | 2.25E-69 | Srsf3 | serine and ar | Nucleus |
| ENSMUSG00000029147 | 1.13E-70 | 3.09E-69 | Ppm1g | protein phos | Nucleus |
| ENSMUSG00000020849 | 1.52E-70 | 4.14E-69 | Ywhae | tyrosine 3-m | Cytoplasm |
| ENSMUSG00000004268 | 3.57E-70 | 9.67E-69 | Emg1 | EMG1, N1-sp | Nucleus |
| ENSMUSG00000006095 | 4.03E-70 | 1.09E-68 | Tbcb | tubulin foldir | Cytoplasm |
| ENSMUSG00000015671 | 5.29E-70 | 1.43E-68 | Psma2 | proteasome s | Cytoplasm |
| ENSMUSG00000066979 | 6.45E-70 | 1.73E-68 | Bub3 | BUB3, mitotic | Nucleus |
| ENSMUSG00000037070 | 7.33E-70 | 1.96E-68 | Rbmxl1 | RNA binding | Nucleus |
| ENSMUSG00000059775 | 1.45E-69 | 3.86E-68 | Rps26-ps1 | ribosomal pr | Other |
| ENSMUSG00000030007 | 2.17E-69 | 5.77E-68 | Cct7 | chaperonin c | Cytoplasm |
| ENSMUSG00000030697 | 4.16E-69 | 1.10E-67 | Ppp4c | protein phos | Cytoplasm |
| ENSMUSG00000032399 | 4.77E-69 | 1.26E-67 | Rpl4 | ribosomal pr | Cytoplasm |
| ENSMUSG00000067786 | 7.13E-69 | 1.87E-67 | Nnat | neuronatin | Plasma Mem |
| ENSMUSG00000073640 | 1.05E-68 | 2.74E-67 | Rpl27-ps3 | ribosomal pr | Other |
| ENSMUSG00000038279 | 1.26E-68 | 3.30E-67 | Nop2 | NOP2 nucleo | Nucleus |
| ENSMUSG00000041859 | 1.29E-68 | 3.37E-67 | Mcm3 | minichromos | Nucleus |
| ENSMUSG00000019773 | 1.77E-68 | 4.58E-67 | Fbxo5 | F-box protei | Nucleus |
| ENSMUSG00000022673 | 2.64E-68 | 6.83E-67 | Mcm4 | minichromos | Nucleus |
| ENSMUSG00000023495 | 4.48E-68 | 1.16E-66 | Pcbp4 | poly(rC) bind | Nucleus |
| ENSMUSG00000028953 | 5.10E-68 | 1.31E-66 | Abcf2 | ATP binding | Cytoplasm |
| ENSMUSG00000032199 | 5.92E-68 | 1.52E-66 | Polr2m | RNA polymer | Nucleus |
| ENSMUSG00000090862 | 6.08E-68 | 1.55E-66 | Rps13 | ribosomal pr | Cytoplasm |
| ENSMUSG00000021178 | 7.79E-68 | 1.98E-66 | Psmc1 | proteasome : | Nucleus |
| ENSMUSG00000025580 | 9.62E-68 | 2.44E-66 | Eif4a3 | eukaryotic tra | Nucleus |
| ENSMUSG00000028902 | 1.55E-67 | 3.93E-66 | Sf3a3 | splicing facto | Nucleus |
| ENSMUSG00000028186 | 2.14E-67 | 5.40E-66 | Uox | urate oxidase | Cytoplasm |
| ENSMUSG00000031403 | 3.33E-67 | 8.37E-66 | Dkc1 | dyskerin pseu | Nucleus |
| ENSMUSG00000093674 | 3.53E-67 | 8.86E-66 | Rpl41 | ribosomal pr | Cytoplasm |
| ENSMUSG00000028896 | 3.93E-67 | 9.82E-66 | Rcc1 | regulator of c | Cytoplasm |
| ENSMUSG00000048644 | 4.89E-67 | 1.22E-65 | Ctxn1 | cortexin 1 | Other |
| ENSMUSG00000039001 | 6.92E-67 | 1.71E-65 | Rps21 | ribosomal pr | Cytoplasm |
| ENSMUSG00000038374 | 1.22E-66 | 3.02E-65 | Rbm8a | RNA binding | Nucleus |
| ENSMUSG00000048758 | 1.40E-66 | 3.46E-65 | Rpl29 | ribosomal pr | Cytoplasm |
| ENSMUSG00000020708 | 1.41E-66 | 3.47E-65 | Psmc5 | proteasome : | Nucleus |
| ENSMUSG00000074476 | 1.55E-66 | 3.79E-65 | Spc24 | SPC24, NDC8 | Cytoplasm |
| ENSMUSG00000037197 | 2.50E-66 | 6.09E-65 | Rbm17 | RNA binding | Nucleus |
| ENSMUSG00000061477 | 3.21E-66 | 7.81E-65 | Rps7 | ribosomal pr | Cytoplasm |
| ENSMUSG00000017404 | 3.91E-66 | 9.49E-65 | Rpl19 | ribosomal pr | Cytoplasm |
| ENSMUSG00000051234 | 3.93E-66 | 9.50E-65 | Rnf7 | ring finger pr | Nucleus |
| ENSMUSG00000027801 | 5.30E-66 | 1.28E-64 | Tm4sf4 | transmembra | Plasma Mem |
| ENSMUSG00000066632 | 5.58E-66 | 1.34E-64 | Pgk1-rs7 | phosphoglyc | Other |
| ENSMUSG00000027078 | 6.25E-66 | 1.50E-64 | Ube2l6 | ubiquitin cor | Cytoplasm |
| ENSMUSG00000055612 | 1.05E-65 | 2.51E-64 | Cdca7 | cell division c | Nucleus |
| ENSMUSG00000022193 | 1.17E-65 | 2.80E-64 | Psmb5 | proteasome s | Cytoplasm |
| ENSMUSG00000012848 | 3.11E-65 | 7.40E-64 | Rps5 | ribosomal pr | Cytoplasm |
| ENSMUSG00000025290 | 7.68E-65 | 1.82E-63 | Rps24 | ribosomal pr | Cytoplasm |
| ENSMUSG00000036892 | 8.02E-65 | 1.90E-63 | Prodh2 | proline dehy | Cytoplasm |
| ENSMUSG00000053398 | 8.92E-65 | 2.10E-63 | Phgdh | phosphoglyc | Cytoplasm |
| ENSMUSG00000073676 | 1.21E-64 | 2.85E-63 | Hspe1 | heat shock p | Cytoplasm |

| | | | | | |
|---|---|---|---|---|---|
| ENSMUSG00000023367 | 1.86E-64 | 4.37E-63 | Tmem176a | transmembra | Other |
| ENSMUSG00000071662 | 3.61E-64 | 8.44E-63 | Polr2g | RNA polymei | Nucleus |
| ENSMUSG00000021069 | 5.11E-64 | 1.19E-62 | Pygl | glycogen pho | Cytoplasm |
| ENSMUSG00000016921 | 7.83E-64 | 1.82E-62 | Srsf6 | serine and ar | Nucleus |
| ENSMUSG00000008668 | 7.88E-64 | 1.83E-62 | Rps18 | ribosomal pr | Cytoplasm |
| ENSMUSG00000062997 | 8.44E-64 | 1.95E-62 | Rpl35 | ribosomal pr | Cytoplasm |
| ENSMUSG00000028837 | 1.51E-63 | 3.49E-62 | Psmb2 | proteasome s | Cytoplasm |
| ENSMUSG00000020644 | 1.55E-63 | 3.56E-62 | Id2 | inhibitor of D | Nucleus |
| ENSMUSG00000061479 | 2.56E-63 | 5.86E-62 | Snrpa | small nuclear | Nucleus |
| ENSMUSG00000027018 | 5.29E-63 | 1.21E-61 | Hat1 | histone acety | Nucleus |
| ENSMUSG00000021546 | 5.91E-63 | 1.35E-61 | Hnrnpk | heterogeneo | Nucleus |
| ENSMUSG00000034892 | 6.20E-63 | 1.41E-61 | Rps29 | ribosomal pr | Cytoplasm |
| ENSMUSG00000029223 | 7.04E-63 | 1.60E-61 | Uchl1 | ubiquitin C-t | Cytoplasm |
| ENSMUSG00000055172 | 7.34E-63 | 1.66E-61 | C1ra | complement | Extracellular S |
| ENSMUSG00000002985 | 1.15E-62 | 2.59E-61 | Apoe | apolipoprote | Extracellular S |
| ENSMUSG00000047675 | 1.80E-62 | 4.06E-61 | Rps8 | ribosomal pr | Cytoplasm |
| ENSMUSG00000030603 | 1.82E-62 | 4.09E-61 | Psmc4 | proteasome 2 | Nucleus |
| ENSMUSG00000046330 | 2.60E-62 | 5.82E-61 | Rpl37a | ribosomal pr | Cytoplasm |
| ENSMUSG00000029810 | 3.53E-62 | 7.87E-61 | Tmem176b | transmembra | Other |
| ENSMUSG00000042354 | 3.71E-62 | 8.26E-61 | Gnl3 | G protein nu | Nucleus |
| ENSMUSG00000037053 | 3.72E-62 | 8.26E-61 | Azgp1 | alpha-2-glyc | Extracellular S |
| ENSMUSG00000061787 | 8.32E-62 | 1.84E-60 | Rps17 | ribosomal pr | Cytoplasm |
| ENSMUSG00000032959 | 9.99E-62 | 2.21E-60 | Pebp1 | phosphatidyl | Cytoplasm |
| ENSMUSG00000035875 | 1.06E-61 | 2.34E-60 | AI182371 | expressed se | Other |
| ENSMUSG00000071415 | 1.07E-61 | 2.35E-60 | Rpl23 | ribosomal pr | Cytoplasm |
| ENSMUSG00000022853 | 1.45E-61 | 3.18E-60 | Ehhadh | enoyl-CoA h | Cytoplasm |
| ENSMUSG00000012405 | 1.50E-61 | 3.28E-60 | Rpl15 | ribosomal pr | Cytoplasm |
| ENSMUSG00000033020 | 1.71E-61 | 3.73E-60 | Polr2f | RNA polymei | Nucleus |
| ENSMUSG00000063787 | 1.72E-61 | 3.73E-60 | Chchd1 | coiled-coil-h | Nucleus |
| ENSMUSG00000027566 | 1.83E-61 | 3.97E-60 | Psma7 | proteasome s | Cytoplasm |
| ENSMUSG00000023104 | 2.01E-61 | 4.34E-60 | Rfc2 | replication fa | Nucleus |
| ENSMUSG00000041126 | 2.47E-61 | 5.32E-60 | H2afv | H2A histone | Nucleus |
| ENSMUSG00000028495 | 2.66E-61 | 5.71E-60 | Rps6 | ribosomal pr | Cytoplasm |
| ENSMUSG00000029657 | 3.17E-61 | 6.76E-60 | Hsph1 | heat shock p | Cytoplasm |
| ENSMUSG00000036752 | 3.19E-61 | 6.79E-60 | Tubb4b | tubulin beta | Cytoplasm |
| ENSMUSG00000073702 | 4.14E-61 | 8.77E-60 | Rpl31 | ribosomal pr | Cytoplasm |
| ENSMUSG00000021377 | 5.98E-61 | 1.26E-59 | Dek | DEK proto-o | Nucleus |
| ENSMUSG00000032591 | 7.38E-61 | 1.56E-59 | Mst1 | macrophage | Extracellular S |
| ENSMUSG00000007739 | 1.14E-60 | 2.40E-59 | Cct4 | chaperonin c | Cytoplasm |
| ENSMUSG00000034528 | 1.29E-60 | 2.71E-59 | Hsd17b13 | hydroxystero | Extracellular S |
| ENSMUSG00000020264 | 1.36E-60 | 2.85E-59 | Slc36a2 | solute carrier | Plasma Mem |
| ENSMUSG00000052798 | 2.08E-60 | 4.35E-59 | Nup107 | nucleoporin 1 | Nucleus |
| ENSMUSG00000034675 | 2.76E-60 | 5.75E-59 | Dbn1 | drebrin 1 | Cytoplasm |
| ENSMUSG00000020086 | 3.11E-60 | 6.45E-59 | H2afy2 | H2A histone | Nucleus |
| ENSMUSG00000004980 | 3.55E-60 | 7.35E-59 | Hnrnpa2b1 | heterogeneo | Nucleus |
| ENSMUSG00000002870 | 4.68E-60 | 9.66E-59 | Mcm2 | minichromos | Nucleus |
| ENSMUSG00000022391 | 4.75E-60 | 9.78E-59 | Rangap1 | Ran GTPase a | Nucleus |
| ENSMUSG00000010376 | 5.70E-60 | 1.17E-58 | Nedd8 | neural precu | Nucleus |
| ENSMUSG00000068874 | 6.57E-60 | 1.35E-58 | Selenbp1 | selenium bin | Cytoplasm |
| ENSMUSG00000020496 | 6.79E-60 | 1.39E-58 | Rnf187 | ring finger pr | Nucleus |
| ENSMUSG00000022033 | 8.15E-60 | 1.66E-58 | Pbk | PDZ binding | Cytoplasm |
| ENSMUSG00000062981 | 1.06E-59 | 2.16E-58 | Mrpl42 | mitochondria | Cytoplasm |
| ENSMUSG00000022205 | 1.11E-59 | 2.26E-58 | Sub1 | SUB1 homolo | Nucleus |
| ENSMUSG00000063457 | 2.01E-59 | 4.07E-58 | Rps15 | ribosomal pr | Cytoplasm |
| ENSMUSG00000093930 | 2.28E-59 | 4.60E-58 | Hmgcs1 | 3-hydroxy-3 | Cytoplasm |
| ENSMUSG00000030662 | 2.76E-59 | 5.55E-58 | Ipo5 | importin 5 | Nucleus |
| ENSMUSG00000036438 | 3.57E-59 | 7.16E-58 | Calm2 | calmodulin 1 | Nucleus |
| ENSMUSG00000017723 | 4.14E-59 | 8.27E-58 | Wfdc2 | WAP four-di | Extracellular S |

| | | | | | |
|---|---|---|---|---|---|
| ENSMUSG00000074207 | 5.21E-59 | 1.04E-57 | Adh1 | alcohol dehy | Cytoplasm |
| ENSMUSG00000042118 | 5.23E-59 | 1.04E-57 | Bhmt2 | betaine--hor | Cytoplasm |
| ENSMUSG00000022858 | 6.57E-59 | 1.30E-57 | Tra2b | transformer 2 | Nucleus |
| ENSMUSG00000054405 | 1.25E-58 | 2.46E-57 | Dnajc8 | DnaJ heat sh | Nucleus |
| ENSMUSG00000027261 | 1.79E-58 | 3.53E-57 | Hao1 | hydroxyacid | Cytoplasm |
| ENSMUSG00000024067 | 2.16E-58 | 4.24E-57 | Dpy30 | dpy-30, histo | Nucleus |
| ENSMUSG00000057657 | 2.16E-58 | 4.24E-57 | Rps18-ps3 | ribosomal pr | Other |
| ENSMUSG00000010064 | 2.61E-58 | 5.11E-57 | Slc38a3 | solute carrier | Plasma Mem |
| ENSMUSG00000062006 | 3.30E-58 | 6.45E-57 | Rpl34 | ribosomal pr | Cytoplasm |
| ENSMUSG00000021024 | 3.91E-58 | 7.62E-57 | Psma6 | proteasome s | Cytoplasm |
| ENSMUSG00000006529 | 3.96E-58 | 7.70E-57 | Itih1 | inter-alpha-t | Extracellular S |
| ENSMUSG00000009079 | 4.82E-58 | 9.35E-57 | Ewsr1 | Ewing sarcon | Nucleus |
| ENSMUSG00000029614 | 5.90E-58 | 1.14E-56 | Rpl6 | ribosomal pr | Nucleus |
| ENSMUSG00000067288 | 6.64E-58 | 1.28E-56 | Rps28 | ribosomal pr | Cytoplasm |
| ENSMUSG00000022400 | 6.76E-58 | 1.30E-56 | Rbx1 | ring-box 1 | Cytoplasm |
| ENSMUSG00000018583 | 7.33E-58 | 1.41E-56 | G3bp1 | G3BP stress g | Nucleus |
| ENSMUSG00000060600 | 9.62E-58 | 1.84E-56 | Eno3 | enolase 3 | Cytoplasm |
| ENSMUSG00000019505 | 1.57E-57 | 3.01E-56 | Ubb | ubiquitin B | Cytoplasm |
| ENSMUSG00000067219 | 1.59E-57 | 3.04E-56 | Nipal1 | NIPA like dor | Other |
| ENSMUSG00000025967 | 2.83E-57 | 5.38E-56 | Eef1b2 | eukaryotic tra | Cytoplasm |
| ENSMUSG00000025197 | 6.11E-57 | 1.16E-55 | Cyp2c44 | cytochrome I | Cytoplasm |
| ENSMUSG00000038482 | 8.89E-57 | 1.68E-55 | Tfdp1 | transcription | Nucleus |
| ENSMUSG00000032939 | 9.00E-57 | 1.70E-55 | Nup93 | nucleoporin 9 | Nucleus |
| ENSMUSG00000026874 | 1.21E-56 | 2.29E-55 | Hc | complement | Extracellular S |
| ENSMUSG00000021660 | 1.92E-56 | 3.61E-55 | Btf3 | basic transcri | Nucleus |
| ENSMUSG00000020051 | 2.05E-56 | 3.85E-55 | Pah | phenylalanin | Cytoplasm |
| ENSMUSG00000040731 | 2.33E-56 | 4.36E-55 | Eif4h | eukaryotic tra | Cytoplasm |
| ENSMUSG00000028633 | 2.40E-56 | 4.48E-55 | Ctps | CTP synthase | Nucleus |
| ENSMUSG00000007872 | 2.88E-56 | 5.37E-55 | Id3 | inhibitor of D | Nucleus |
| ENSMUSG00000038510 | 4.08E-56 | 7.59E-55 | Rpf2 | ribosome pro | Nucleus |
| ENSMUSG00000043716 | 4.35E-56 | 8.07E-55 | Rpl7 | ribosomal pr | Nucleus |
| ENSMUSG00000025066 | 5.61E-56 | 1.04E-54 | Sfr1 | SWI5 depenc | Nucleus |
| ENSMUSG00000021537 | 8.04E-56 | 1.49E-54 | Cetn3 | centrin 3 | Cytoplasm |
| ENSMUSG00000001403 | 8.22E-56 | 1.52E-54 | Ube2c | ubiquitin con | Cytoplasm |
| ENSMUSG00000022285 | 1.08E-55 | 1.98E-54 | Ywhaz | tyrosine 3-m | Cytoplasm |
| ENSMUSG00000006398 | 2.53E-55 | 4.64E-54 | Cdc20 | cell division c | Nucleus |
| ENSMUSG00000040413 | 2.78E-55 | 5.10E-54 | Timd2 | T cell immun | Plasma Mem |
| ENSMUSG00000022992 | 3.25E-55 | 5.95E-54 | Kansl2 | KAT8 regulat | Nucleus |
| ENSMUSG00000022247 | 5.59E-55 | 1.02E-53 | Brix1 | BRX1, bioger | Nucleus |
| ENSMUSG00000094806 | 8.06E-55 | 1.47E-53 | Cyp2d10 | cytochrome I | Cytoplasm |
| ENSMUSG00000009013 | 9.04E-55 | 1.64E-53 | Dynll1 | dynein light c | Cytoplasm |
| ENSMUSG00000026377 | 1.74E-54 | 3.15E-53 | Nifk | nucleolar pro | Nucleus |
| ENSMUSG00000057322 | 2.07E-54 | 3.73E-53 | Rpl38 | ribosomal pr | Cytoplasm |
| ENSMUSG00000005481 | 2.12E-54 | 3.81E-53 | Ddx39 | DExD-box he | Nucleus |
| ENSMUSG00000016024 | 2.52E-54 | 4.53E-53 | Lbp | lipopolysacch | Plasma Mem |
| ENSMUSG00000032207 | 2.87E-54 | 5.15E-53 | Lipc | lipase C, hep | Extracellular S |
| ENSMUSG00000027496 | 3.03E-54 | 5.41E-53 | Aurka | aurora kinase | Nucleus |
| ENSMUSG00000024007 | 3.09E-54 | 5.51E-53 | Ppil1 | peptidylproly | Plasma Mem |
| ENSMUSG00000029711 | 3.64E-54 | 6.47E-53 | Epo | erythropoieti | Extracellular S |
| ENSMUSG00000024029 | 5.06E-54 | 8.97E-53 | Tff3 | trefoil factor | Extracellular S |
| ENSMUSG00000001416 | 5.30E-54 | 9.38E-53 | Cct3 | chaperonin c | Cytoplasm |
| ENSMUSG00000060036 | 6.23E-54 | 1.10E-52 | Rpl3 | ribosomal pr | Nucleus |
| ENSMUSG00000045128 | 6.64E-54 | 1.17E-52 | Rpl18a | ribosomal pr | Cytoplasm |
| ENSMUSG00000027804 | 1.40E-53 | 2.46E-52 | Ppid | peptidylproly | Cytoplasm |
| ENSMUSG00000032279 | 1.61E-53 | 2.82E-52 | Idh3a | isocitrate deh | Cytoplasm |
| ENSMUSG00000042770 | 3.45E-53 | 6.04E-52 | Hebp1 | heme binding | Cytoplasm |
| ENSMUSG00000029580 | 4.31E-53 | 7.52E-52 | Actb | actin beta | Cytoplasm |
| ENSMUSG00000053801 | 5.59E-53 | 9.74E-52 | Grwd1 | glutamate ric | Nucleus |

| | | | | | |
|---|---|---|---|---|---|
| ENSMUSG00000022881 | 5.84E-53 | 1.02E-51 | Rfc4 | replication fa | Nucleus |
| ENSMUSG00000000628 | 6.92E-53 | 1.20E-51 | Hk2 | hexokinase 2 | Cytoplasm |
| ENSMUSG00000037542 | 8.81E-53 | 1.53E-51 | Aldh8a1 | aldehyde deh | Cytoplasm |
| ENSMUSG00000031311 | 1.04E-52 | 1.80E-51 | Nono | non-POU do | Nucleus |
| ENSMUSG00000042729 | 1.15E-52 | 1.99E-51 | Wdr74 | WD repeat d | Nucleus |
| ENSMUSG00000038274 | 1.43E-52 | 2.47E-51 | Fau | FAU, ubiquiti | Cytoplasm |
| ENSMUSG00000033898 | 1.80E-52 | 3.09E-51 | Cfhr2 | complement | Other |
| ENSMUSG00000030245 | 2.66E-52 | 4.55E-51 | Golt1b | golgi transpo | Cytoplasm |
| ENSMUSG00000037361 | 2.71E-52 | 4.62E-51 | Sf3b6 | splicing facto | Nucleus |
| ENSMUSG00000028330 | 2.84E-52 | 4.84E-51 | Ncbp1 | nuclear cap b | Nucleus |
| ENSMUSG00000042079 | 4.24E-52 | 7.20E-51 | Hnrnpf | heterogeneo | Nucleus |
| ENSMUSG00000015120 | 4.42E-52 | 7.49E-51 | Ube2i | ubiquitin con | Nucleus |
| ENSMUSG00000001155 | 4.44E-52 | 7.50E-51 | Ftcd | formimidoylt | Cytoplasm |
| ENSMUSG00000058558 | 5.35E-52 | 9.02E-51 | Rpl5 | ribosomal pro | Cytoplasm |
| ENSMUSG00000005683 | 6.01E-52 | 1.01E-50 | Cs | citrate syntha | Cytoplasm |
| ENSMUSG00000021116 | 6.59E-52 | 1.11E-50 | Eif2s1 | eukaryotic tra | Cytoplasm |
| ENSMUSG00000021832 | 7.67E-52 | 1.29E-50 | Psmc6 | proteasome ( | Nucleus |
| ENSMUSG00000062867 | 9.95E-52 | 1.67E-50 | Impdh2 | inosine mono | Cytoplasm |
| ENSMUSG00000026750 | 1.21E-51 | 2.02E-50 | Psmb7 | proteasome s | Cytoplasm |
| ENSMUSG00000031904 | 1.35E-51 | 2.25E-50 | Slc7a6 | solute carrier | Plasma Mem |
| ENSMUSG00000021282 | 1.48E-51 | 2.46E-50 | Eif5 | eukaryotic tra | Cytoplasm |
| ENSMUSG00000067148 | 2.75E-51 | 4.56E-50 | Polr1c | RNA polymer | Nucleus |
| ENSMUSG00000000028 | 2.79E-51 | 4.61E-50 | Cdc45 | cell division c | Nucleus |
| ENSMUSG00000036371 | 3.31E-51 | 5.45E-50 | Serbp1 | SERPINE1 mR | Cytoplasm |
| ENSMUSG00000018293 | 3.98E-51 | 6.54E-50 | Pfn1 | profilin 1 | Cytoplasm |
| ENSMUSG00000005233 | 5.44E-51 | 8.93E-50 | Spc25 | SPC25, NDC8 | Cytoplasm |
| ENSMUSG00000074088 | 5.67E-51 | 9.29E-50 | Snrnp40 | small nuclear | Nucleus |
| ENSMUSG00000032575 | 6.25E-51 | 1.02E-49 | Manf | mesencephal | Extracellular S |
| ENSMUSG00000023110 | 8.36E-51 | 1.36E-49 | Prmt5 | protein argin | Cytoplasm |
| ENSMUSG00000041057 | 8.74E-51 | 1.42E-49 | Wdr43 | WD repeat d | Nucleus |
| ENSMUSG00000022109 | 1.03E-50 | 1.67E-49 | Med4 | mediator con | Nucleus |
| ENSMUSG00000046364 | 1.09E-50 | 1.77E-49 | Rpl27a | ribosomal pro | Cytoplasm |
| ENSMUSG00000024556 | 1.14E-50 | 1.84E-49 | Me2 | malic enzyme | Cytoplasm |
| ENSMUSG00000007039 | 1.40E-50 | 2.25E-49 | Ddah2 | dimethylargi | Cytoplasm |
| ENSMUSG00000041840 | 1.46E-50 | 2.34E-49 | Haus1 | HAUS augmi | Nucleus |
| ENSMUSG00000020720 | 1.71E-50 | 2.75E-49 | Psmd12 | proteasome ( | Cytoplasm |
| ENSMUSG00000036570 | 1.82E-50 | 2.91E-49 | Fxyd1 | FXYD domain | Plasma Mem |
| ENSMUSG00000027170 | 2.00E-50 | 3.20E-49 | Eif3m | eukaryotic tra | Cytoplasm |
| ENSMUSG00000053898 | 2.27E-50 | 3.62E-49 | Ech1 | enoyl-CoA h | Cytoplasm |
| ENSMUSG00000055302 | 2.31E-50 | 3.68E-49 | Mrfap1 | Morf4 family | Nucleus |
| ENSMUSG00000047797 | 2.36E-50 | 3.75E-49 | Gjb1 | gap junction | Plasma Mem |
| ENSMUSG00000040026 | 2.75E-50 | 4.37E-49 | Saa3 | serum amylo | Extracellular S |
| ENSMUSG00000033047 | 3.41E-50 | 5.39E-49 | Eif3l | eukaryotic tra | Cytoplasm |
| ENSMUSG00000020116 | 3.43E-50 | 5.42E-49 | Pno1 | partner of NO | Nucleus |
| ENSMUSG00000041431 | 6.32E-50 | 9.96E-49 | Ccnb1 | cyclin B1 | Cytoplasm |
| ENSMUSG00000029380 | 6.62E-50 | 1.04E-48 | Cxcl1 | C-X-C motif | Extracellular S |
| ENSMUSG00000031445 | 1.01E-49 | 1.58E-48 | Proz | protein Z, vita | Extracellular S |
| ENSMUSG00000062248 | 1.15E-49 | 1.80E-48 | Cks2 | CDC28 prote | Other |
| ENSMUSG00000034203 | 2.02E-49 | 3.16E-48 | Chchd4 | coiled-coil-h | Cytoplasm |
| ENSMUSG00000074656 | 2.09E-49 | 3.25E-48 | Eif2s2 | eukaryotic tra | Cytoplasm |
| ENSMUSG00000000740 | 4.10E-49 | 6.36E-48 | Rpl13 | ribosomal pro | Nucleus |
| ENSMUSG00000030983 | 4.77E-49 | 7.39E-48 | Bccip | BRCA2 and C | Nucleus |
| ENSMUSG00000036781 | 4.83E-49 | 7.46E-48 | Rps27l | ribosomal pro | Cytoplasm |
| ENSMUSG00000060288 | 5.02E-49 | 7.75E-48 | Ppih | peptidylproly | Nucleus |
| ENSMUSG00000062647 | 5.14E-49 | 7.92E-48 | Rpl7a | ribosomal pro | Cytoplasm |
| ENSMUSG00000018362 | 5.54E-49 | 8.52E-48 | Kpna2 | karyopherin s | Nucleus |
| ENSMUSG00000028010 | 5.59E-49 | 8.58E-48 | Gar1 | GAR1 ribonu | Nucleus |
| ENSMUSG00000022304 | 6.65E-49 | 1.02E-47 | Dpys | dihydropyrim | Cytoplasm |

| | | | | | |
|---|---|---|---|---|---|
| ENSMUSG00000029777 | 7.78E-49 | 1.19E-47 | Gars | glycyl-tRNA : | Cytoplasm |
| ENSMUSG00000032412 | 8.64E-49 | 1.32E-47 | Atp1b3 | ATPase Na+/ | Plasma Mem |
| ENSMUSG00000044986 | 1.06E-48 | 1.61E-47 | Tst | thiosulfate su | Cytoplasm |
| ENSMUSG00000023169 | 1.43E-48 | 2.18E-47 | Slc38a1 | solute carrier | Plasma Mem |
| ENSMUSG00000038393 | 1.45E-48 | 2.21E-47 | Txnip | thioredoxin i | Cytoplasm |
| ENSMUSG00000060407 | 1.74E-48 | 2.63E-47 | Cyp2a12 | cytochrome I | Cytoplasm |
| ENSMUSG00000095427 | 2.00E-48 | 3.02E-47 | Rps2-ps6 | ribosomal pr | Other |
| ENSMUSG00000035960 | 2.54E-48 | 3.84E-47 | Apex1 | apurinic/apyı | Nucleus |
| ENSMUSG00000037942 | 2.59E-48 | 3.91E-47 | Crp | C-reactive pr | Extracellular S |
| ENSMUSG00000030677 | 2.67E-48 | 4.01E-47 | Kif22 | kinesin family | Nucleus |
| ENSMUSG00000026281 | 3.19E-48 | 4.79E-47 | Dtymk | deoxythymid | Cytoplasm |
| ENSMUSG00000028322 | 3.22E-48 | 4.82E-47 | Exosc3 | exosome cor | Nucleus |
| ENSMUSG00000058655 | 3.40E-48 | 5.09E-47 | Eif4b | eukaryotic tra | Cytoplasm |
| ENSMUSG00000010095 | 4.04E-48 | 6.03E-47 | Slc3a2 | solute carrier | Plasma Mem |
| ENSMUSG00000047963 | 4.74E-48 | 7.07E-47 | Stbd1 | starch bindin | Cytoplasm |
| ENSMUSG00000021018 | 9.79E-48 | 1.45E-46 | Polr2h | RNA polymeı | Nucleus |
| ENSMUSG00000055760 | 1.09E-47 | 1.61E-46 | Gemin6 | gem nuclear | Nucleus |
| ENSMUSG00000059208 | 1.11E-47 | 1.64E-46 | Hnrnpm | heterogeneo | Nucleus |
| ENSMUSG00000001707 | 1.14E-47 | 1.69E-46 | Eef1e1 | eukaryotic tra | Cytoplasm |
| ENSMUSG00000074781 | 1.33E-47 | 1.96E-46 | Ube2n | ubiquitin cor | Cytoplasm |
| ENSMUSG00000031320 | 1.40E-47 | 2.06E-46 | Rps4x | ribosomal pr | Cytoplasm |
| ENSMUSG00000074768 | 1.83E-47 | 2.68E-46 | Bhmt | betaine--hoı | Cytoplasm |
| ENSMUSG00000070697 | 2.14E-47 | 3.13E-46 | Utp3 | UTP3, small s | Nucleus |
| ENSMUSG00000020808 | 2.24E-47 | 3.28E-46 | Fam64a | PICALM inter | Nucleus |
| ENSMUSG00000052146 | 2.77E-47 | 4.04E-46 | Rps10 | ribosomal pr | Cytoplasm |
| ENSMUSG00000017716 | 3.39E-47 | 4.94E-46 | Birc5 | baculoviral IA | Cytoplasm |
| ENSMUSG00000004642 | 3.40E-47 | 4.94E-46 | Slbp | stem-loop bi | Nucleus |
| ENSMUSG00000031578 | 4.57E-47 | 6.63E-46 | Mak16 | MAK16 homo | Nucleus |
| ENSMUSG00000024177 | 5.20E-47 | 7.54E-46 | Nme4 | NME/NM23 ı | Cytoplasm |
| ENSMUSG00000064264 | 8.27E-47 | 1.20E-45 | Zfp428 | zinc finger pı | Other |
| ENSMUSG00000024165 | 9.23E-47 | 1.33E-45 | Hn1l | Jupiter micro | Cytoplasm |
| ENSMUSG00000021569 | 9.25E-47 | 1.33E-45 | Trip13 | thyroid horm | Cytoplasm |
| ENSMUSG00000020180 | 1.11E-46 | 1.59E-45 | Snrpd3 | small nuclear | Nucleus |
| ENSMUSG00000009614 | 1.18E-46 | 1.69E-45 | Sardh | sarcosine del | Cytoplasm |
| ENSMUSG00000059070 | 1.46E-46 | 2.10E-45 | Rpl18 | ribosomal pr | Cytoplasm |
| ENSMUSG00000006998 | 1.71E-46 | 2.44E-45 | Psmd2 | proteasome : | Cytoplasm |
| ENSMUSG00000070436 | 2.00E-46 | 2.85E-45 | Serpinh1 | serpin family | Extracellular S |
| ENSMUSG00000024791 | 2.12E-46 | 3.02E-45 | Cdca5 | cell division c | Cytoplasm |
| ENSMUSG00000021340 | 2.26E-46 | 3.21E-45 | Gpld1 | glycosylphos | Cytoplasm |
| ENSMUSG00000025869 | 2.34E-46 | 3.33E-45 | Nop16 | NOP16 nucle | Nucleus |
| ENSMUSG00000041571 | 2.73E-46 | 3.88E-45 | Sepw1 | selenoproteiı | Cytoplasm |
| ENSMUSG00000063511 | 2.84E-46 | 4.02E-45 | Snrnp70 | small nuclear | Nucleus |
| ENSMUSG00000031458 | 2.99E-46 | 4.23E-45 | Coprs | coordinator c | Nucleus |
| ENSMUSG00000045273 | 3.37E-46 | 4.76E-45 | Cenph | centromere p | Nucleus |
| ENSMUSG00000022312 | 3.52E-46 | 4.95E-45 | Eif3h | eukaryotic tra | Cytoplasm |
| ENSMUSG00000029064 | 3.53E-46 | 4.96E-45 | Gnb1 | G protein sul | Plasma Mem |
| ENSMUSG00000030224 | 3.88E-46 | 5.46E-45 | Strap | serine/threor | Plasma Mem |
| ENSMUSG00000068039 | 4.26E-46 | 5.97E-45 | Tcp1 | t-complex 1 | Cytoplasm |
| ENSMUSG00000030088 | 4.32E-46 | 6.05E-45 | Aldh1l1 | aldehyde del | Cytoplasm |
| ENSMUSG00000021591 | 4.80E-46 | 6.71E-45 | Glrx | glutaredoxin | Cytoplasm |
| ENSMUSG00000002102 | 5.86E-46 | 8.17E-45 | Psmc3 | proteasome : | Nucleus |
| ENSMUSG00000034932 | 7.14E-46 | 9.95E-45 | Mrpl54 | mitochondria | Cytoplasm |
| ENSMUSG00000054630 | 8.62E-46 | 1.20E-44 | Ugt2b5 | UDP glucuro | Cytoplasm |
| ENSMUSG00000021737 | 8.86E-46 | 1.23E-44 | Psmd6 | proteasome : | Cytoplasm |
| ENSMUSG00000010122 | 9.96E-46 | 1.38E-44 | Slc47a1 | solute carrier | Plasma Mem |
| ENSMUSG00000029432 | 1.06E-45 | 1.46E-44 | Gbas | nipsnap hom | Plasma Mem |
| ENSMUSG00000051989 | 1.26E-45 | 1.74E-44 | Smim11 | small integra | Other |
| ENSMUSG00000026049 | 1.60E-45 | 2.20E-44 | Tex30 | testis express | Other |

| | | | | | |
|---|---|---|---|---|---|
| ENSMUSG00000002718 | 2.03E-45 | 2.79E-44 | Cse1l | chromosome | Nucleus |
| ENSMUSG00000038521 | 3.14E-45 | 4.31E-44 | C1s1 | complement | Extracellular S |
| ENSMUSG00000057863 | 5.66E-45 | 7.77E-44 | Rpl36 | ribosomal pr | Nucleus |
| ENSMUSG00000001674 | 7.15E-45 | 9.80E-44 | Ddx18 | DEAD-box h | Nucleus |
| ENSMUSG00000022322 | 9.03E-45 | 1.23E-43 | Shcbp1 | SHC binding | Other |
| ENSMUSG00000028603 | 9.79E-45 | 1.34E-43 | Scp2 | sterol carrier | Cytoplasm |
| ENSMUSG00000022037 | 1.03E-44 | 1.40E-43 | Clu | clusterin | Cytoplasm |
| ENSMUSG00000024853 | 1.05E-44 | 1.43E-43 | Sf3b2 | splicing facto | Nucleus |
| ENSMUSG00000038622 | 1.21E-44 | 1.64E-43 | Med30 | mediator cor | Nucleus |
| ENSMUSG00000070348 | 1.52E-44 | 2.06E-43 | Ccnd1 | cyclin D1 | Nucleus |
| ENSMUSG00000030512 | 1.73E-44 | 2.34E-43 | Snrpa1 | small nuclear | Nucleus |
| ENSMUSG00000028156 | 2.07E-44 | 2.80E-43 | Eif4e | eukaryotic tra | Cytoplasm |
| ENSMUSG00000030382 | 2.44E-44 | 3.29E-43 | Slc27a5 | solute carrier | Cytoplasm |
| ENSMUSG00000032966 | 3.36E-44 | 4.53E-43 | Fkbp1a | FK506 bindin | Cytoplasm |
| ENSMUSG00000061032 | 3.73E-44 | 5.02E-43 | Rrp1 | ribosomal RN | Nucleus |
| ENSMUSG00000031954 | 3.87E-44 | 5.20E-43 | Cfdp1 | craniofacial c | Extracellular S |
| ENSMUSG00000063632 | 3.99E-44 | 5.36E-43 | Sox11 | SRY-box 11 | Nucleus |
| ENSMUSG00000063888 | 4.48E-44 | 6.00E-43 | Rpl7l1 | ribosomal pr | Cytoplasm |
| ENSMUSG00000013155 | 6.30E-44 | 8.41E-43 | Enkd1 | enkurin dom | Cytoplasm |
| ENSMUSG00000019471 | 7.02E-44 | 9.37E-43 | Cdc37 | cell division c | Cytoplasm |
| ENSMUSG00000039356 | 8.06E-44 | 1.07E-42 | Exosc2 | exosome cor | Nucleus |
| ENSMUSG00000044786 | 8.19E-44 | 1.09E-42 | Zfp36 | ZFP36 ring fi | Nucleus |
| ENSMUSG00000041438 | 8.27E-44 | 1.10E-42 | Cirh1a | UTP4, small s | Nucleus |
| ENSMUSG00000038412 | 8.94E-44 | 1.18E-42 | Higd1a | HIG1 hypoxia | Cytoplasm |
| ENSMUSG00000002833 | 1.09E-43 | 1.44E-42 | Hdgfrp2 | HDGF like 2 | Nucleus |
| ENSMUSG00000074882 | 1.10E-43 | 1.46E-42 | Cyp2c68 | cytochrome I | Cytoplasm |
| ENSMUSG00000005413 | 1.18E-43 | 1.55E-42 | Hmox1 | heme oxyger | Cytoplasm |
| ENSMUSG00000027715 | 1.22E-43 | 1.61E-42 | Ccna2 | cyclin A2 | Nucleus |
| ENSMUSG00000075229 | 1.31E-43 | 1.72E-42 | Ccdc58 | coiled-coil d | Cytoplasm |
| ENSMUSG00000030079 | 1.56E-43 | 2.05E-42 | Ruvbl1 | RuvB like AA | Nucleus |
| ENSMUSG00000062397 | 1.71E-43 | 2.24E-42 | Zfp706 | zinc finger pr | Other |
| ENSMUSG00000021135 | 1.74E-43 | 2.27E-42 | Slc10a1 | solute carrier | Plasma Mem |
| ENSMUSG00000036309 | 1.89E-43 | 2.47E-42 | Skp1a | S-phase kina | Nucleus |
| ENSMUSG00000067818 | 2.03E-43 | 2.65E-42 | Myl9 | myosin light | Cytoplasm |
| ENSMUSG00000006369 | 2.04E-43 | 2.65E-42 | Fbln1 | fibulin 1 | Extracellular S |
| ENSMUSG00000025872 | 2.14E-43 | 2.78E-42 | Thoc3 | THO comple | Nucleus |
| ENSMUSG00000033294 | 2.25E-43 | 2.92E-42 | Noc4l | nucleolar cor | Nucleus |
| ENSMUSG00000093904 | 2.82E-43 | 3.65E-42 | Tomm20 | translocase o | Cytoplasm |
| ENSMUSG00000021607 | 3.84E-43 | 4.97E-42 | Mrpl36 | mitochondria | Cytoplasm |
| ENSMUSG00000021094 | 4.15E-43 | 5.36E-42 | Dhrs7 | dehydrogena | Other |
| ENSMUSG00000024990 | 4.57E-43 | 5.89E-42 | Rbp4 | retinol bindir | Extracellular S |
| ENSMUSG00000028884 | 4.64E-43 | 5.97E-42 | Rpa2 | replication pr | Nucleus |
| ENSMUSG00000020075 | 4.87E-43 | 6.26E-42 | Ddx21 | DExD-box he | Nucleus |
| ENSMUSG00000024039 | 5.20E-43 | 6.67E-42 | Cbs | cystathionine | Cytoplasm |
| ENSMUSG00000091537 | 5.39E-43 | 6.91E-42 | Tma7 | translation m | Other |
| ENSMUSG00000053560 | 5.51E-43 | 7.05E-42 | Ier2 | immediate ea | Cytoplasm |
| ENSMUSG00000031444 | 5.76E-43 | 7.35E-42 | F10 | coagulation f | Extracellular S |
| ENSMUSG00000028367 | 5.86E-43 | 7.47E-42 | Txn1 | thioredoxin | Cytoplasm |
| ENSMUSG00000032454 | 6.54E-43 | 8.32E-42 | Rbp2 | retinol bindir | Cytoplasm |
| ENSMUSG00000025358 | 7.14E-43 | 9.08E-42 | Cdk2 | cyclin depen | Nucleus |
| ENSMUSG00000031917 | 9.15E-43 | 1.16E-41 | Nip7 | NIP7, nucleol | Nucleus |
| ENSMUSG00000024391 | 9.85E-43 | 1.25E-41 | Apom | apolipoprote | Plasma Mem |
| ENSMUSG00000007050 | 1.28E-42 | 1.62E-41 | Lsm2 | LSM2 homolo | Nucleus |
| ENSMUSG00000032359 | 1.34E-42 | 1.69E-41 | Ctsh | cathepsin H | Cytoplasm |
| ENSMUSG00000061024 | 1.39E-42 | 1.75E-41 | Rrs1 | ribosome bio | Nucleus |
| ENSMUSG00000068566 | 1.40E-42 | 1.77E-41 | Myadm | myeloid asso | Nucleus |
| ENSMUSG00000071533 | 1.65E-42 | 2.08E-41 | Pcnp | PEST proteol | Nucleus |
| ENSMUSG00000001348 | 1.82E-42 | 2.29E-41 | Acp5 | acid phospha | Cytoplasm |

| | | | | | |
|---|---|---|---|---|---|
| ENSMUSG00000031843 | 1.86E-42 | 2.34E-41 | Mphosph6 | M-phase pho | Nucleus |
| ENSMUSG00000020460 | 2.10E-42 | 2.63E-41 | Rps27a | ribosomal pr | Cytoplasm |
| ENSMUSG00000032518 | 2.15E-42 | 2.69E-41 | Rpsa | ribosomal pr | Cytoplasm |
| ENSMUSG00000022427 | 2.21E-42 | 2.76E-41 | Tomm22 | translocase o | Cytoplasm |
| ENSMUSG00000001228 | 3.02E-42 | 3.76E-41 | Uhrf1 | ubiquitin like | Nucleus |
| ENSMUSG00000001642 | 3.19E-42 | 3.97E-41 | Akr1b3 | aldo-keto rec | Cytoplasm |
| ENSMUSG00000022962 | 3.69E-42 | 4.58E-41 | Gart | phosphoribo | Cytoplasm |
| ENSMUSG00000034321 | 3.73E-42 | 4.62E-41 | Exosc1 | exosome cor | Nucleus |
| ENSMUSG00000059835 | 3.85E-42 | 4.77E-41 | Rpl13-ps3 | ribosomal pr | Other |
| ENSMUSG00000029910 | 4.00E-42 | 4.94E-41 | Mad2l1 | mitotic arrest | Nucleus |
| ENSMUSG00000028741 | 4.31E-42 | 5.32E-41 | Mrto4 | MRT4 homol | Cytoplasm |
| ENSMUSG00000036678 | 4.50E-42 | 5.54E-41 | Aaas | aladin WD re | Nucleus |
| ENSMUSG00000060073 | 4.66E-42 | 5.73E-41 | Psma3 | proteasome s | Cytoplasm |
| ENSMUSG00000068184 | 4.81E-42 | 5.91E-41 | Ndufaf2 | NADH:ubiqu | Cytoplasm |
| ENSMUSG00000024986 | 5.36E-42 | 6.57E-41 | Hhex | hematopoiet | Nucleus |
| ENSMUSG00000031068 | 6.61E-42 | 8.09E-41 | Glrx3 | glutaredoxin | Cytoplasm |
| ENSMUSG00000025630 | 8.40E-42 | 1.03E-40 | Hprt | hypoxanthine | Cytoplasm |
| ENSMUSG00000040463 | 9.15E-42 | 1.12E-40 | Mybbp1a | MYB binding | Nucleus |
| ENSMUSG00000040017 | 9.44E-42 | 1.15E-40 | Saa4 | SAA2-SAA4 r | Extracellular S |
| ENSMUSG00000050440 | 1.06E-41 | 1.29E-40 | Hamp | hepcidin anti | Extracellular S |
| ENSMUSG00000078126 | 1.28E-41 | 1.56E-40 | Rpl23a-ps3 | ribosomal pr | Other |
| ENSMUSG00000022580 | 1.39E-41 | 1.68E-40 | Rhpn1 | rhophilin Rho | Cytoplasm |
| ENSMUSG00000052997 | 1.52E-41 | 1.84E-40 | Uba2 | ubiquitin like | Cytoplasm |
| ENSMUSG00000016946 | 1.67E-41 | 2.02E-40 | Kctd5 | potassium ch | Cytoplasm |
| ENSMUSG00000069805 | 1.70E-41 | 2.05E-40 | Fbp1 | fructose-bisp | Cytoplasm |
| ENSMUSG00000040767 | 1.85E-41 | 2.23E-40 | Snrnp25 | small nuclear | Nucleus |
| ENSMUSG00000040952 | 1.91E-41 | 2.29E-40 | Rps19 | ribosomal pr | Cytoplasm |
| ENSMUSG00000040084 | 2.11E-41 | 2.54E-40 | Bub1b | BUB1 mitotic | Nucleus |
| ENSMUSG00000029642 | 2.33E-41 | 2.80E-40 | Polr1d | RNA polymer | Nucleus |
| ENSMUSG00000058135 | 2.46E-41 | 2.95E-40 | Gstm1 | glutathione S | Cytoplasm |
| ENSMUSG00000023224 | 2.55E-41 | 3.05E-40 | Serping1 | serpin family | Extracellular S |
| ENSMUSG00000029552 | 2.62E-41 | 3.13E-40 | Tes | testin LIM do | Plasma Mem |
| ENSMUSG00000028645 | 2.72E-41 | 3.24E-40 | Slc2a1 | solute carrier | Plasma Mem |
| ENSMUSG00000024590 | 2.98E-41 | 3.56E-40 | Lmnb1 | lamin B1 | Nucleus |
| ENSMUSG00000028356 | 3.03E-41 | 3.61E-40 | Ambp | alpha-1-micr | Extracellular S |
| ENSMUSG00000018189 | 3.03E-41 | 3.61E-40 | Uchl5 | ubiquitin C-t | Cytoplasm |
| ENSMUSG00000019814 | 3.20E-41 | 3.80E-40 | Ltv1 | LTV1 ribosom | Other |
| ENSMUSG00000058355 | 3.21E-41 | 3.81E-40 | Abce1 | ATP binding | Cytoplasm |
| ENSMUSG00000000154 | 3.32E-41 | 3.93E-40 | Slc22a18 | solute carrier | Plasma Mem |
| ENSMUSG00000013822 | 3.40E-41 | 4.02E-40 | Elof1 | elongation fa | Nucleus |
| ENSMUSG00000037601 | 3.74E-41 | 4.41E-40 | Nme1 | NME/NM23 r | Cytoplasm |
| ENSMUSG00000019857 | 4.30E-41 | 5.06E-40 | Asf1a | anti-silencing | Nucleus |
| ENSMUSG00000026688 | 4.84E-41 | 5.69E-40 | Mgst3 | microsomal g | Cytoplasm |
| ENSMUSG00000005779 | 4.94E-41 | 5.81E-40 | Psmb4 | proteasome s | Cytoplasm |
| ENSMUSG00000021276 | 6.29E-41 | 7.39E-40 | Cinp | cyclin depend | Nucleus |
| ENSMUSG00000024800 | 7.81E-41 | 9.16E-40 | Rpp30 | ribonuclease | Nucleus |
| ENSMUSG00000047832 | 9.93E-41 | 1.16E-39 | Cdca4 | cell division c | Nucleus |
| ENSMUSG00000028932 | 1.32E-40 | 1.55E-39 | Psmc2 | proteasome 2 | Nucleus |
| ENSMUSG00000040771 | 1.33E-40 | 1.55E-39 | Oard1 | O-acyl-ADP- | Other |
| ENSMUSG00000025503 | 1.40E-40 | 1.64E-39 | Taldo1 | transaldolase | Cytoplasm |
| ENSMUSG00000020485 | 1.84E-40 | 2.14E-39 | Supt4a | SPT4 homolo | Nucleus |
| ENSMUSG00000035351 | 1.93E-40 | 2.24E-39 | Nup37 | nucleoporin 3 | Nucleus |
| ENSMUSG00000037570 | 2.36E-40 | 2.74E-39 | Mcrs1 | microspherul | Nucleus |
| ENSMUSG00000047215 | 2.39E-40 | 2.77E-39 | Rpl9 | ribosomal pr | Nucleus |
| ENSMUSG00000020646 | 2.71E-40 | 3.14E-39 | Mboat2 | membrane b | Cytoplasm |
| ENSMUSG00000028729 | 2.77E-40 | 3.20E-39 | Ebna1bp2 | EBNA1 bindi | Nucleus |
| ENSMUSG00000028800 | 3.39E-40 | 3.91E-39 | Hdac1 | histone deac | Nucleus |
| ENSMUSG00000029014 | 3.71E-40 | 4.28E-39 | Dnajc2 | DnaJ heat sh | Nucleus |

| | | | | | |
|---|---|---|---|---|---|
| ENSMUSG00000039055 | 4.27E-40 | 4.91E-39 | Eme1 | essential mei | Nucleus |
| ENSMUSG00000042289 | 4.67E-40 | 5.37E-39 | Hsd3b7 | hydroxy-delt | Cytoplasm |
| ENSMUSG00000002660 | 4.82E-40 | 5.53E-39 | Clpp | caseinolytic r | Cytoplasm |
| ENSMUSG00000033940 | 5.92E-40 | 6.79E-39 | Brk1 | BRICK1, SCAI | Cytoplasm |
| ENSMUSG00000024498 | 6.06E-40 | 6.93E-39 | Tcerg1 | transcription | Nucleus |
| ENSMUSG00000034994 | 6.25E-40 | 7.14E-39 | Eef2 | eukaryotic tra | Cytoplasm |
| ENSMUSG00000024841 | 6.49E-40 | 7.41E-39 | Eif1ad | eukaryotic tra | Nucleus |
| ENSMUSG00000030342 | 7.50E-40 | 8.55E-39 | Cd9 | CD9 molecul | Plasma Mem |
| ENSMUSG00000020706 | 9.65E-40 | 1.10E-38 | Ftsj3 | FtsJ RNA met | Nucleus |
| ENSMUSG00000032171 | 1.04E-39 | 1.18E-38 | Pin1 | peptidylproly | Nucleus |
| ENSMUSG00000020737 | 1.19E-39 | 1.35E-38 | Hn1 | Jupiter micro | Nucleus |
| ENSMUSG00000020224 | 1.21E-39 | 1.37E-38 | Llph | LLP homolog | Extracellular S |
| ENSMUSG00000005470 | 1.40E-39 | 1.59E-38 | Asf1b | anti-silencing | Nucleus |
| ENSMUSG00000029328 | 1.42E-39 | 1.61E-38 | Hnrnpdl | heterogeneo | Nucleus |
| ENSMUSG00000029247 | 1.56E-39 | 1.76E-38 | Paics | phosphoribo | Cytoplasm |
| ENSMUSG00000029911 | 1.59E-39 | 1.79E-38 | Ssbp1 | single strand | Cytoplasm |
| ENSMUSG00000064037 | 2.07E-39 | 2.33E-38 | Gpn1 | GPN-loop GT | Nucleus |
| ENSMUSG00000074129 | 2.17E-39 | 2.43E-38 | Rpl13a | ribosomal pr | Cytoplasm |
| ENSMUSG00000035198 | 2.30E-39 | 2.57E-38 | Tubg1 | tubulin gamr | Cytoplasm |
| ENSMUSG00000056035 | 3.16E-39 | 3.53E-38 | Cyp3a11 | cytochrome I | Cytoplasm |
| ENSMUSG00000023456 | 3.61E-39 | 4.03E-38 | Tpi1 | triosephosph | Cytoplasm |
| ENSMUSG00000020897 | 3.95E-39 | 4.40E-38 | Aurkb | aurora kinase | Nucleus |
| ENSMUSG00000060636 | 4.09E-39 | 4.56E-38 | Rpl35a | ribosomal pr | Cytoplasm |
| ENSMUSG00000009092 | 4.40E-39 | 4.90E-38 | Derl3 | derlin 3 | Cytoplasm |
| ENSMUSG00000008333 | 4.77E-39 | 5.30E-38 | Snrpb2 | small nuclear | Nucleus |
| ENSMUSG00000056305 | 6.22E-39 | 6.90E-38 | Usp39 | ubiquitin spe | Nucleus |
| ENSMUSG00000027227 | 6.66E-39 | 7.38E-38 | Sord | sorbitol dehy | Cytoplasm |
| ENSMUSG00000014769 | 7.07E-39 | 7.83E-38 | Psmb1 | proteasome s | Cytoplasm |
| ENSMUSG00000022774 | 9.26E-39 | 1.02E-37 | Ncbp2 | nuclear cap b | Nucleus |
| ENSMUSG00000026558 | 9.85E-39 | 1.09E-37 | Uck2 | uridine-cytid | Cytoplasm |
| ENSMUSG00000020471 | 1.08E-38 | 1.19E-37 | Pold2 | DNA polyme | Nucleus |
| ENSMUSG00000069744 | 1.11E-38 | 1.22E-37 | Psmb3 | proteasome s | Cytoplasm |
| ENSMUSG00000031168 | 1.14E-38 | 1.25E-37 | Ebp | EBP, choleste | Cytoplasm |
| ENSMUSG00000013701 | 1.18E-38 | 1.30E-37 | Timm23 | translocase o | Cytoplasm |
| ENSMUSG00000026355 | 1.26E-38 | 1.38E-37 | Mcm6 | minichromos | Nucleus |
| ENSMUSG00000024537 | 1.28E-38 | 1.40E-37 | Psmg2 | proteasome a | Nucleus |
| ENSMUSG00000024309 | 1.30E-38 | 1.42E-37 | Pfdn6 | prefoldin sub | Cytoplasm |
| ENSMUSG00000032215 | 1.34E-38 | 1.47E-37 | Rsl24d1 | ribosomal L2 | Nucleus |
| ENSMUSG00000057666 | 1.49E-38 | 1.62E-37 | Gapdh | glyceraldehy | Cytoplasm |
| ENSMUSG00000030796 | 1.55E-38 | 1.69E-37 | Tead2 | TEA domain | Nucleus |
| ENSMUSG00000029716 | 1.60E-38 | 1.74E-37 | Tfr2 | transferrin re | Plasma Mem |
| ENSMUSG00000039756 | 1.76E-38 | 1.91E-37 | Dnttip2 | deoxynucleo | Nucleus |
| ENSMUSG00000026914 | 1.82E-38 | 1.98E-37 | Psmd14 | proteasome 2 | Cytoplasm |
| ENSMUSG00000028676 | 1.92E-38 | 2.08E-37 | Srsf10 | serine and ar | Nucleus |
| ENSMUSG00000068856 | 1.95E-38 | 2.11E-37 | Sf3b4 | splicing facto | Nucleus |
| ENSMUSG00000024660 | 2.19E-38 | 2.36E-37 | Incenp | inner centror | Nucleus |
| ENSMUSG00000019782 | 2.19E-38 | 2.36E-37 | Rwdd1 | RWD domain | Cytoplasm |
| ENSMUSG00000031783 | 2.67E-38 | 2.87E-37 | Polr2c | RNA polymer | Nucleus |
| ENSMUSG00000031585 | 2.69E-38 | 2.90E-37 | Gtf2e2 | general trans | Nucleus |
| ENSMUSG00000006412 | 2.91E-38 | 3.12E-37 | Pfdn2 | prefoldin sub | Cytoplasm |
| ENSMUSG00000031948 | 3.23E-38 | 3.46E-37 | Kars | lysyl-tRNA sy | Cytoplasm |
| ENSMUSG00000030751 | 3.31E-38 | 3.54E-37 | Psma1 | proteasome s | Cytoplasm |
| ENSMUSG00000007721 | 3.34E-38 | 3.56E-37 | Ccdc124 | coiled-coil do | Other |
| ENSMUSG00000029066 | 3.59E-38 | 3.82E-37 | Mrpl20 | mitochondria | Cytoplasm |
| ENSMUSG00000035237 | 3.60E-38 | 3.83E-37 | Lcat | lecithin-chol | Extracellular S |
| ENSMUSG00000031799 | 3.71E-38 | 3.94E-37 | Tpm4 | tropomyosin | Cytoplasm |
| ENSMUSG00000020328 | 3.93E-38 | 4.18E-37 | Nudcd2 | NudC domai | Cytoplasm |
| ENSMUSG00000061286 | 4.43E-38 | 4.70E-37 | Exosc5 | exosome cor | Nucleus |

| | | | | | |
|---|---|---|---|---|---|
| ENSMUSG00000027671 | 4.61E-38 | 4.88E-37 | Actl6a | actin like 6A | Nucleus |
| ENSMUSG00000039771 | 4.89E-38 | 5.17E-37 | Polr2j | RNA polymeı | Nucleus |
| ENSMUSG00000031562 | 5.22E-38 | 5.51E-37 | Dctd | dCMP deami | Cytoplasm |
| ENSMUSG00000023150 | 5.72E-38 | 6.03E-37 | Ivns1abp | influenza viru | Nucleus |
| ENSMUSG00000025785 | 6.09E-38 | 6.41E-37 | Exosc7 | exosome cor | Nucleus |
| ENSMUSG00000022574 | 6.35E-38 | 6.68E-37 | Naprt | nicotinate ph | Cytoplasm |
| ENSMUSG00000042406 | 6.45E-38 | 6.78E-37 | Atf4 | activating tra | Nucleus |
| ENSMUSG00000026272 | 6.57E-38 | 6.89E-37 | Agxt | alanine--gly | Cytoplasm |
| ENSMUSG00000021492 | 8.24E-38 | 8.62E-37 | F12 | coagulation f | Extracellular S |
| ENSMUSG00000031434 | 9.84E-38 | 1.03E-36 | Morc4 | MORC family | Nucleus |
| ENSMUSG00000023913 | 1.05E-37 | 1.10E-36 | Pla2g7 | phospholipas | Extracellular S |
| ENSMUSG00000041506 | 1.11E-37 | 1.16E-36 | Rrp9 | ribosomal RN | Nucleus |
| ENSMUSG00000058546 | 1.15E-37 | 1.20E-36 | Rpl23a | ribosomal pr | Nucleus |
| ENSMUSG00000035455 | 1.28E-37 | 1.33E-36 | Fignl1 | fidgetin like 1 | Nucleus |
| ENSMUSG00000022052 | 1.80E-37 | 1.86E-36 | Ppp2r2a | protein phos | Cytoplasm |
| ENSMUSG00000031756 | 1.85E-37 | 1.92E-36 | Cenpn | centromere p | Nucleus |
| ENSMUSG00000001305 | 1.99E-37 | 2.06E-36 | Rrp15 | ribosomal RN | Nucleus |
| ENSMUSG00000055762 | 2.42E-37 | 2.50E-36 | Eef1d | eukaryotic tra | Cytoplasm |
| ENSMUSG00000007850 | 2.65E-37 | 2.73E-36 | Hnrnph1 | heterogeneo | Nucleus |
| ENSMUSG00000079658 | 2.77E-37 | 2.86E-36 | Tceb1 | elongin C | Nucleus |
| ENSMUSG00000003131 | 2.80E-37 | 2.89E-36 | Pafah1b2 | platelet activ | Cytoplasm |
| ENSMUSG00000003299 | 2.96E-37 | 3.04E-36 | Mrpl4 | mitochondria | Cytoplasm |
| ENSMUSG00000038252 | 3.05E-37 | 3.13E-36 | Ncapd2 | non-SMC co | Nucleus |
| ENSMUSG00000059291 | 3.10E-37 | 3.18E-36 | Rpl11 | ribosomal pr | Cytoplasm |
| ENSMUSG00000001663 | 3.37E-37 | 3.46E-36 | Gstt1 | glutathione S | Cytoplasm |
| ENSMUSG00000027076 | 8.01E-37 | 8.20E-36 | Timm10 | translocase o | Cytoplasm |
| ENSMUSG00000021595 | 8.46E-37 | 8.65E-36 | Nsun2 | NOP2/Sun R | Nucleus |
| ENSMUSG00000028936 | 9.15E-37 | 9.34E-36 | Rpl22 | ribosomal pr | Nucleus |
| ENSMUSG00000024271 | 9.63E-37 | 9.82E-36 | Elp2 | elongator ac | Cytoplasm |
| ENSMUSG00000020914 | 1.09E-36 | 1.11E-35 | Top2a | DNA topoiso | Nucleus |
| ENSMUSG00000022706 | 1.12E-36 | 1.14E-35 | Mrpl40 | mitochondria | Cytoplasm |
| ENSMUSG00000037805 | 1.44E-36 | 1.46E-35 | Rpl10a | ribosomal pr | Nucleus |
| ENSMUSG00000055839 | 1.95E-36 | 1.98E-35 | Tceb2 | elongin B | Nucleus |
| ENSMUSG00000022241 | 2.13E-36 | 2.16E-35 | Tars | threonyl-tRN | Nucleus |
| ENSMUSG00000017499 | 2.44E-36 | 2.47E-35 | Cdc6 | cell division c | Nucleus |
| ENSMUSG00000029246 | 2.46E-36 | 2.49E-35 | Ppat | phosphoribo | Cytoplasm |
| ENSMUSG00000006731 | 2.59E-36 | 2.62E-35 | B4galnt1 | beta-1,4-N-a | Cytoplasm |
| ENSMUSG00000027509 | 2.93E-36 | 2.95E-35 | Rae1 | ribonucleic a | Nucleus |
| ENSMUSG00000022821 | 2.96E-36 | 2.98E-35 | Hgd | homogentisa | Cytoplasm |
| ENSMUSG00000020372 | 2.99E-36 | 3.00E-35 | Gnb2l1 | receptor for a | Cytoplasm |
| ENSMUSG00000019916 | 3.26E-36 | 3.28E-35 | P4ha1 | prolyl 4-hydr | Cytoplasm |
| ENSMUSG00000034450 | 3.33E-36 | 3.34E-35 | Gulo | gulonolactor | Cytoplasm |
| ENSMUSG00000062591 | 3.87E-36 | 3.88E-35 | Tubb4a | tubulin beta | Cytoplasm |
| ENSMUSG00000020719 | 5.58E-36 | 5.58E-35 | Ddx5 | DEAD-box h | Nucleus |
| ENSMUSG00000016319 | 7.27E-36 | 7.26E-35 | Slc25a5 | solute carrier | Cytoplasm |
| ENSMUSG00000027752 | 7.89E-36 | 7.87E-35 | Exosc8 | exosome cor | Nucleus |
| ENSMUSG00000028409 | 8.38E-36 | 8.35E-35 | Smu1 | SMU1, DNA r | Nucleus |
| ENSMUSG00000031990 | 8.94E-36 | 8.90E-35 | Jam3 | junctional ad | Plasma Mem |
| ENSMUSG00000005161 | 1.05E-35 | 1.05E-34 | Prdx2 | peroxiredoxi | Cytoplasm |
| ENSMUSG00000023057 | 1.07E-35 | 1.06E-34 | Fabp2 | fatty acid bin | Cytoplasm |
| ENSMUSG00000028998 | 1.10E-35 | 1.09E-34 | Tomm7 | translocase o | Cytoplasm |
| ENSMUSG00000026864 | 1.21E-35 | 1.20E-34 | Hspa5 | heat shock p | Cytoplasm |
| ENSMUSG00000028873 | 1.33E-35 | 1.31E-34 | Cdca8 | cell division c | Nucleus |
| ENSMUSG00000031490 | 1.47E-35 | 1.46E-34 | Eif4ebp1 | eukaryotic tra | Cytoplasm |
| ENSMUSG00000028394 | 1.48E-35 | 1.46E-34 | Pole3 | DNA polyme | Nucleus |
| ENSMUSG00000027454 | 1.59E-35 | 1.57E-34 | Gins1 | GINS comple | Nucleus |
| ENSMUSG00000031701 | 1.61E-35 | 1.58E-34 | Dnaja2 | DnaJ heat sh | Nucleus |
| ENSMUSG00000020183 | 1.70E-35 | 1.67E-34 | Cpm | carboxypepti | Plasma Mem |

| | | | | | |
|---|---|---|---|---|---|
| ENSMUSG00000026174 | 1.88E-35 | 1.84E-34 | Rqcd1 | CCR4-NOT tı | Cytoplasm |
| ENSMUSG00000003779 | 2.01E-35 | 1.97E-34 | Kif20a | kinesin family | Cytoplasm |
| ENSMUSG00000004460 | 2.17E-35 | 2.13E-34 | Dnajb11 | DnaJ heat sh | Cytoplasm |
| ENSMUSG00000020777 | 2.19E-35 | 2.14E-34 | Acox1 | acyl-CoA oxi | Cytoplasm |
| ENSMUSG00000069755 | 2.23E-35 | 2.18E-34 | Zfp125 | | Other |
| ENSMUSG00000030056 | 2.35E-35 | 2.30E-34 | Isy1 | ISY1-RAB43 ı | Nucleus |
| ENSMUSG00000029388 | 2.61E-35 | 2.54E-34 | Eif2b1 | eukaryotic tra | Cytoplasm |
| ENSMUSG00000019923 | 2.88E-35 | 2.80E-34 | Zwint | ZW10 interac | Nucleus |
| ENSMUSG00000016554 | 3.00E-35 | 2.92E-34 | Eif3d | eukaryotic tra | Cytoplasm |
| ENSMUSG00000019942 | 3.11E-35 | 3.02E-34 | Cdk1 | cyclin depen | Nucleus |
| ENSMUSG00000001383 | 3.12E-35 | 3.03E-34 | Zmat2 | zinc finger m | Nucleus |
| ENSMUSG00000016257 | 3.30E-35 | 3.20E-34 | Slmo2 | PRELI domair | Cytoplasm |
| ENSMUSG00000024646 | 3.86E-35 | 3.74E-34 | Cyb5a | cytochrome I | Cytoplasm |
| ENSMUSG00000004665 | 3.89E-35 | 3.77E-34 | Cnn2 | calponin 2 | Cytoplasm |
| ENSMUSG00000062456 | 4.28E-35 | 4.14E-34 | Rpl9-ps6 | ribosomal pr | Other |
| ENSMUSG00000022300 | 4.31E-35 | 4.16E-34 | Dcaf13 | DDB1 and CU | Nucleus |
| ENSMUSG00000071866 | 4.91E-35 | 4.74E-34 | Ppia | peptidylproly | Cytoplasm |
| ENSMUSG00000046352 | 5.15E-35 | 4.96E-34 | Gjb2 | gap junction | Plasma Mem |
| ENSMUSG00000001440 | 5.32E-35 | 5.12E-34 | Kpnb1 | karyopherin s | Nucleus |
| ENSMUSG00000031546 | 5.44E-35 | 5.23E-34 | Gins4 | GINS comple | Nucleus |
| ENSMUSG00000061315 | 5.55E-35 | 5.32E-34 | Naca | nascent poly | Cytoplasm |
| ENSMUSG00000004864 | 5.89E-35 | 5.64E-34 | Mapk13 | mitogen-acti | Cytoplasm |
| ENSMUSG00000056394 | 6.86E-35 | 6.56E-34 | Lig1 | DNA ligase 1 | Nucleus |
| ENSMUSG00000020457 | 7.01E-35 | 6.70E-34 | Drg1 | development | Cytoplasm |
| ENSMUSG00000020022 | 8.33E-35 | 7.94E-34 | Ndufa12 | NADH:ubiqu | Cytoplasm |
| ENSMUSG00000027193 | 9.03E-35 | 8.60E-34 | Api5 | apoptosis inł | Cytoplasm |
| ENSMUSG00000041453 | 9.26E-35 | 8.81E-34 | Rpl21 | ribosomal pr | Cytoplasm |
| ENSMUSG00000028651 | 9.31E-35 | 8.84E-34 | Ppie | peptidylproly | Nucleus |
| ENSMUSG00000022856 | 9.32E-35 | 8.84E-34 | Tmem41a | transmembra | Other |
| ENSMUSG00000033272 | 9.51E-35 | 9.01E-34 | Slc35a4 | solute carrier | Cytoplasm |
| ENSMUSG00000002825 | 9.67E-35 | 9.16E-34 | Qtrt1 | queuine tRN | Cytoplasm |
| ENSMUSG00000022945 | 1.00E-34 | 9.45E-34 | Chaf1b | chromatin as | Nucleus |
| ENSMUSG00000032026 | 1.10E-34 | 1.04E-33 | Rexo2 | RNA exonucl | Cytoplasm |
| ENSMUSG00000061947 | 1.11E-34 | 1.05E-33 | Serpina10 | serpin family | Extracellular S |
| ENSMUSG00000035041 | 1.13E-34 | 1.07E-33 | Creb3l3 | cAMP respor | Cytoplasm |
| ENSMUSG00000041360 | 1.19E-34 | 1.12E-33 | D19Bwg1357e | pumilio RNA | Nucleus |
| ENSMUSG00000036992 | 1.39E-34 | 1.31E-33 | Nxt1 | nuclear trans | Nucleus |
| ENSMUSG00000040385 | 1.42E-34 | 1.33E-33 | Ppp1ca | protein phos | Cytoplasm |
| ENSMUSG00000059119 | 1.65E-34 | 1.54E-33 | Nap1l4 | nucleosome | Cytoplasm |
| ENSMUSG00000020171 | 1.85E-34 | 1.73E-33 | Yeats4 | YEATS doma | Nucleus |
| ENSMUSG00000020361 | 1.93E-34 | 1.80E-33 | Hspa4 | heat shock p | Cytoplasm |
| ENSMUSG00000039007 | 2.28E-34 | 2.13E-33 | Cpq | carboxypepti | Extracellular S |
| ENSMUSG00000057388 | 2.61E-34 | 2.44E-33 | Mrpl18 | mitochondria | Cytoplasm |
| ENSMUSG00000024661 | 2.93E-34 | 2.73E-33 | Fth1 | ferritin heavy | Cytoplasm |
| ENSMUSG00000030591 | 3.28E-34 | 3.05E-33 | Psmd8 | proteasome 2 | Cytoplasm |
| ENSMUSG00000030432 | 3.55E-34 | 3.30E-33 | Rpl28 | ribosomal pr | Cytoplasm |
| ENSMUSG00000026385 | 3.96E-34 | 3.68E-33 | Dbi | diazepam bir | Cytoplasm |
| ENSMUSG00000098470 | 3.97E-34 | 3.68E-33 | C1rb | complement | Extracellular S |
| ENSMUSG00000050777 | 4.16E-34 | 3.86E-33 | Tmem37 | transmembra | Plasma Mem |
| ENSMUSG00000059552 | 5.69E-34 | 5.26E-33 | Trp53 | tumor protei | Nucleus |
| ENSMUSG00000006498 | 5.87E-34 | 5.43E-33 | Ptbp1 | polypyrimidir | Nucleus |
| ENSMUSG00000032114 | 6.16E-34 | 5.69E-33 | Slc37a4 | solute carrier | Cytoplasm |
| ENSMUSG00000028964 | 6.47E-34 | 5.97E-33 | Park7 | Parkinsonism | Nucleus |
| ENSMUSG00000021243 | 8.65E-34 | 7.97E-33 | Fcf1 | FCF1, rRNA-p | Nucleus |
| ENSMUSG00000079563 | 9.08E-34 | 8.36E-33 | Pglyrp2 | peptidoglyca | Plasma Mem |
| ENSMUSG00000027479 | 1.10E-33 | 1.01E-32 | Mapre1 | microtubule | Cytoplasm |
| ENSMUSG00000046432 | 1.75E-33 | 1.61E-32 | Ngfrap1 | brain express | Cytoplasm |
| ENSMUSG00000021175 | 2.03E-33 | 1.85E-32 | Cdca7l | cell division c | Nucleus |

| | | | | | |
|---|---|---|---|---|---|
| ENSMUSG00000049115 | 2.16E-33 | 1.97E-32 | Agtr1a | angiotensin I | Plasma Mem |
| ENSMUSG00000025451 | 2.32E-33 | 2.11E-32 | Paip1 | poly(A) bindi | Cytoplasm |
| ENSMUSG00000025156 | 2.42E-33 | 2.21E-32 | Gps1 | G protein pat | Nucleus |
| ENSMUSG00000021645 | 2.48E-33 | 2.26E-32 | Smn1 | survival of m | Nucleus |
| ENSMUSG00000030423 | 2.60E-33 | 2.37E-32 | Pop4 | POP4 homol | Nucleus |
| ENSMUSG00000036023 | 2.73E-33 | 2.49E-32 | Parp2 | poly(ADP-rib | Nucleus |
| ENSMUSG00000025127 | 3.43E-33 | 3.11E-32 | Gcgr | glucagon rec | Plasma Mem |
| ENSMUSG00000025130 | 3.45E-33 | 3.13E-32 | P4hb | prolyl 4-hydr | Cytoplasm |
| ENSMUSG00000040749 | 3.65E-33 | 3.31E-32 | Siah1b | siah E3 ubiqu | Nucleus |
| ENSMUSG00000027313 | 3.70E-33 | 3.35E-32 | Chac1 | ChaC glutath | Cytoplasm |
| ENSMUSG00000035726 | 3.79E-33 | 3.42E-32 | Supt16 | SPT16 homo | Nucleus |
| ENSMUSG00000045427 | 3.87E-33 | 3.49E-32 | Hnrnph2 | heterogeneo | Nucleus |
| ENSMUSG00000024767 | 4.81E-33 | 4.34E-32 | Otub1 | OTU domain | Cytoplasm |
| ENSMUSG00000038550 | 4.90E-33 | 4.42E-32 | Ciart | circadian ass | Nucleus |
| ENSMUSG00000027469 | 5.34E-33 | 4.81E-32 | Tpx2 | TPX2, microt | Nucleus |
| ENSMUSG00000026189 | 5.97E-33 | 5.36E-32 | Pecr | peroxisomal | Cytoplasm |
| ENSMUSG00000020514 | 6.13E-33 | 5.50E-32 | Mrpl22 | mitochondria | Cytoplasm |
| ENSMUSG00000033715 | 8.02E-33 | 7.19E-32 | Akr1c14 | aldo-keto re | Cytoplasm |
| ENSMUSG00000021714 | 8.71E-33 | 7.79E-32 | Cenpk | centromere p | Nucleus |
| ENSMUSG00000062328 | 9.96E-33 | 8.90E-32 | Rpl17 | ribosomal pr | Cytoplasm |
| ENSMUSG00000057130 | 1.16E-32 | 1.03E-31 | Txnl4a | thioredoxin li | Nucleus |
| ENSMUSG00000056758 | 1.69E-32 | 1.50E-31 | Hmga2 | high mobility | Nucleus |
| ENSMUSG00000002524 | 1.71E-32 | 1.53E-31 | Puf60 | poly(U) bindi | Nucleus |
| ENSMUSG00000042043 | 1.76E-32 | 1.57E-31 | Tbca | tubulin foldir | Cytoplasm |
| ENSMUSG00000074797 | 1.81E-32 | 1.61E-31 | Itpa | inosine triph | Cytoplasm |
| ENSMUSG00000031383 | 1.88E-32 | 1.67E-31 | Dusp9 | dual specifici | Nucleus |
| ENSMUSG00000028195 | 2.23E-32 | 1.97E-31 | Cyr61 | cysteine rich | Extracellular |
| ENSMUSG00000031683 | 2.23E-32 | 1.98E-31 | Lsm6 | LSM6 homol | Nucleus |
| ENSMUSG00000041645 | 2.34E-32 | 2.07E-31 | Ddx24 | DEAD-box h | Nucleus |
| ENSMUSG00000029463 | 2.54E-32 | 2.24E-31 | Fam216a | family with se | Other |
| ENSMUSG00000030942 | 2.59E-32 | 2.29E-31 | Thumpd1 | THUMP dom | Nucleus |
| ENSMUSG00000031107 | 2.63E-32 | 2.32E-31 | Rbmx2 | RNA binding | Other |
| ENSMUSG00000024528 | 2.72E-32 | 2.40E-31 | Srfbp1 | serum respor | Nucleus |
| ENSMUSG00000012483 | 2.73E-32 | 2.40E-31 | Rpa3 | replication pr | Nucleus |
| ENSMUSG00000020526 | 3.09E-32 | 2.72E-31 | Znhit3 | zinc finger H | Nucleus |
| ENSMUSG00000051159 | 3.13E-32 | 2.75E-31 | Cited1 | Cbp/p300 int | Nucleus |
| ENSMUSG00000030738 | 3.20E-32 | 2.81E-31 | Eif3c | eukaryotic tra | Cytoplasm |
| ENSMUSG00000024084 | 3.21E-32 | 2.82E-31 | Qpct | glutaminyl-p | Cytoplasm |
| ENSMUSG00000019303 | 3.38E-32 | 2.96E-31 | Psmc3ip | PSMC3 intera | Nucleus |
| ENSMUSG00000028212 | 3.40E-32 | 2.97E-31 | Ccne2 | cyclin E2 | Nucleus |
| ENSMUSG00000079614 | 3.50E-32 | 3.06E-31 | Seh1l | SEH1 like nuc | Cytoplasm |
| ENSMUSG00000051223 | 4.55E-32 | 3.97E-31 | Bzw1 | basic leucine | Cytoplasm |
| ENSMUSG00000037313 | 4.59E-32 | 4.00E-31 | Tacc3 | transforming | Nucleus |
| ENSMUSG00000002372 | 6.15E-32 | 5.36E-31 | Ranbp3 | RAN binding | Nucleus |
| ENSMUSG00000007564 | 7.29E-32 | 6.34E-31 | Ppp2r1a | protein phos | Cytoplasm |
| ENSMUSG00000003099 | 7.77E-32 | 6.76E-31 | Ppp5c | protein phos | Nucleus |
| ENSMUSG00000025050 | 7.90E-32 | 6.85E-31 | Pcgf6 | polycomb gr | Nucleus |
| ENSMUSG00000033318 | 8.51E-32 | 7.38E-31 | Gstt2 | glutathione S | Cytoplasm |
| ENSMUSG00000063884 | 8.72E-32 | 7.55E-31 | Ptcd3 | pentatricope | Cytoplasm |
| ENSMUSG00000032178 | 8.82E-32 | 7.62E-31 | Ilf3 | interleukin er | Nucleus |
| ENSMUSG00000053453 | 1.21E-31 | 1.04E-30 | Thoc7 | THO comple: | Nucleus |
| ENSMUSG00000029405 | 1.21E-31 | 1.04E-30 | G3bp2 | G3BP stress g | Cytoplasm |
| ENSMUSG00000005687 | 1.21E-31 | 1.04E-30 | Bcas2 | BCAS2, pre-r | Nucleus |
| ENSMUSG00000030521 | 1.42E-31 | 1.22E-30 | Mphosph10 | M-phase pho | Nucleus |
| ENSMUSG00000023861 | 1.60E-31 | 1.37E-30 | Mpc1 | mitochondria | Cytoplasm |
| ENSMUSG00000035530 | 1.61E-31 | 1.38E-30 | Eif1 | eukaryotic tra | Other |
| ENSMUSG00000023019 | 1.76E-31 | 1.51E-30 | Gpd1 | glycerol-3-p | Cytoplasm |
| ENSMUSG00000037935 | 1.78E-31 | 1.53E-30 | Smarce1 | SWI/SNF rela | Nucleus |

| | | | | | |
|---|---|---|---|---|---|
| ENSMUSG00000058600 | 1.79E-31 | 1.53E-30 | Rpl30 | ribosomal pr | Cytoplasm |
| ENSMUSG00000031167 | 1.91E-31 | 1.63E-30 | Rbm3 | RNA binding | Cytoplasm |
| ENSMUSG00000021149 | 2.02E-31 | 1.72E-30 | Gtpbp4 | GTP binding | Nucleus |
| ENSMUSG00000005610 | 2.10E-31 | 1.80E-30 | Eif4g2 | eukaryotic tra | Cytoplasm |
| ENSMUSG00000038975 | 2.30E-31 | 1.96E-30 | Rabggtb | Rab geranylg | Cytoplasm |
| ENSMUSG00000022383 | 2.61E-31 | 2.22E-30 | Ppara | peroxisome p | Nucleus |
| ENSMUSG00000011179 | 2.65E-31 | 2.25E-30 | Odc1 | ornithine dec | Cytoplasm |
| ENSMUSG00000021115 | 2.68E-31 | 2.28E-30 | Vrk1 | vaccinia relat | Nucleus |
| ENSMUSG00000033970 | 2.87E-31 | 2.43E-30 | Rfc3 | replication fa | Nucleus |
| ENSMUSG00000020935 | 3.03E-31 | 2.57E-30 | Dcakd | dephospho-( | Cytoplasm |
| ENSMUSG00000040652 | 4.44E-31 | 3.76E-30 | Oaz2 | ornithine dec | Cytoplasm |
| ENSMUSG00000003438 | 4.69E-31 | 3.97E-30 | Timm50 | translocase o | Cytoplasm |
| ENSMUSG00000003623 | 6.55E-31 | 5.53E-30 | Crot | carnitine O-c | Cytoplasm |
| ENSMUSG00000047844 | 6.65E-31 | 5.61E-30 | Bex4 | brain express | Cytoplasm |
| ENSMUSG00000035683 | 8.08E-31 | 6.81E-30 | Melk | maternal emb | Cytoplasm |
| ENSMUSG00000025487 | 8.84E-31 | 7.44E-30 | Psmd13 | proteasome ( | Cytoplasm |
| ENSMUSG00000028234 | 1.01E-30 | 8.51E-30 | Rps20 | ribosomal pr | Cytoplasm |
| ENSMUSG00000032314 | 1.02E-30 | 8.57E-30 | Etfa | electron trans | Cytoplasm |
| ENSMUSG00000000561 | 1.03E-30 | 8.68E-30 | Wdr77 | WD repeat d | Nucleus |
| ENSMUSG00000016756 | 1.12E-30 | 9.43E-30 | Cmah | cytidine mon | Cytoplasm |
| ENSMUSG00000030417 | 1.21E-30 | 1.01E-29 | Pdcd5 | programmed | Nucleus |
| ENSMUSG00000035278 | 1.29E-30 | 1.08E-29 | Plekhj1 | pleckstrin ho | Other |
| ENSMUSG00000032058 | 1.46E-30 | 1.22E-29 | Ppp2r1b | protein phos | Plasma Mem |
| ENSMUSG00000045996 | 1.60E-30 | 1.34E-29 | Polr2k | RNA polymer | Nucleus |
| ENSMUSG00000001785 | 1.73E-30 | 1.44E-29 | Pwp1 | PWP1 homol | Nucleus |
| ENSMUSG00000029363 | 1.76E-30 | 1.47E-29 | Rfc5 | replication fa | Nucleus |
| ENSMUSG00000018326 | 1.78E-30 | 1.48E-29 | Ywhab | tyrosine 3-m | Cytoplasm |
| ENSMUSG00000054079 | 1.83E-30 | 1.52E-29 | Utp18 | UTP18, small | Nucleus |
| ENSMUSG00000024892 | 1.97E-30 | 1.63E-29 | Pcx | pyruvate carb | Cytoplasm |
| ENSMUSG00000006941 | 2.01E-30 | 1.66E-29 | Eif1b | eukaryotic tra | Cytoplasm |
| ENSMUSG00000029817 | 2.08E-30 | 1.72E-29 | Tra2a | transformer 2 | Nucleus |
| ENSMUSG00000105617 | 2.13E-30 | 1.76E-29 | Mrpl33 | | Other |
| ENSMUSG00000028345 | 2.20E-30 | 1.82E-29 | Tex10 | testis express | Nucleus |
| ENSMUSG00000024974 | 2.38E-30 | 1.96E-29 | Smc3 | structural ma | Nucleus |
| ENSMUSG00000024312 | 2.39E-30 | 1.97E-29 | Wdr46 | WD repeat d | Nucleus |
| ENSMUSG00000031149 | 2.41E-30 | 1.98E-29 | Praf2 | PRA1 domain | Other |
| ENSMUSG00000015889 | 2.74E-30 | 2.26E-29 | Lta4h | leukotriene A | Cytoplasm |
| ENSMUSG00000062937 | 2.75E-30 | 2.26E-29 | Mtap | methylthioad | Nucleus |
| ENSMUSG00000001017 | 2.78E-30 | 2.29E-29 | Chtop | chromatin ta | Nucleus |
| ENSMUSG00000034906 | 3.01E-30 | 2.47E-29 | Ncaph | non-SMC co | Nucleus |
| ENSMUSG00000028678 | 4.66E-30 | 3.83E-29 | Kif2c | kinesin family | Nucleus |
| ENSMUSG00000029695 | 4.77E-30 | 3.91E-29 | Aass | aminoadipate | Cytoplasm |
| ENSMUSG00000014402 | 4.94E-30 | 4.04E-29 | Tsg101 | tumor suscep | Cytoplasm |
| ENSMUSG00000022748 | 5.15E-30 | 4.21E-29 | Cmss1 | cms1 ribosom | Other |
| ENSMUSG00000031609 | 5.22E-30 | 4.27E-29 | Sap30 | Sin3A associa | Nucleus |
| ENSMUSG00000040181 | 5.49E-30 | 4.48E-29 | Fmo1 | flavin contain | Cytoplasm |
| ENSMUSG00000020918 | 5.57E-30 | 4.54E-29 | Kat2a | lysine acetylt | Cytoplasm |
| ENSMUSG00000060802 | 5.62E-30 | 4.58E-29 | B2m | beta-2-micro | Plasma Mem |
| ENSMUSG00000038628 | 5.83E-30 | 4.74E-29 | Polr3k | RNA polymer | Nucleus |
| ENSMUSG00000044080 | 6.03E-30 | 4.90E-29 | S100a1 | S100 calcium | Cytoplasm |
| ENSMUSG00000051768 | 6.66E-30 | 5.41E-29 | Xrcc1 | X-ray repair ( | Nucleus |
| ENSMUSG00000033585 | 7.33E-30 | 5.94E-29 | Ndn | necdin, MAG | Nucleus |
| ENSMUSG00000034349 | 8.82E-30 | 7.14E-29 | Smc4 | structural ma | Nucleus |
| ENSMUSG00000002679 | 9.07E-30 | 7.33E-29 | Med6 | mediator cor | Nucleus |
| ENSMUSG00000024056 | 9.99E-30 | 8.06E-29 | Ndc80 | NDC80, kinet | Nucleus |
| ENSMUSG00000030842 | 1.07E-29 | 8.59E-29 | Lamtor1 | late endosom | Plasma Mem |
| ENSMUSG00000014956 | 1.13E-29 | 9.08E-29 | Ppp1cb | protein phos | Cytoplasm |
| ENSMUSG00000063354 | 1.51E-29 | 1.21E-28 | Slc39a4 | solute carrier | Plasma Mem |

| | | | | | |
|---|---|---|---|---|---|
| ENSMUSG00000032288 | 1.76E-29 | 1.41E-28 | Imp3 | IMP3, U3 sma | Cytoplasm |
| ENSMUSG00000061740 | 1.82E-29 | 1.46E-28 | Cyp2d22 | cytochrome I | Cytoplasm |
| ENSMUSG00000051256 | 1.83E-29 | 1.47E-28 | Jagn1 | jagunal hom | Cytoplasm |
| ENSMUSG00000001473 | 1.86E-29 | 1.49E-28 | Tubb6 | tubulin beta | Cytoplasm |
| ENSMUSG00000001774 | 2.10E-29 | 1.68E-28 | Chordc1 | cysteine and | Other |
| ENSMUSG00000004285 | 2.21E-29 | 1.77E-28 | Atp6v1f | ATPase H+ tr | Cytoplasm |
| ENSMUSG00000021218 | 2.27E-29 | 1.81E-28 | Gdi2 | GDP dissocia | Cytoplasm |
| ENSMUSG00000000751 | 2.36E-29 | 1.88E-28 | Rpa1 | replication pr | Nucleus |
| ENSMUSG00000019851 | 2.57E-29 | 2.05E-28 | Perp | PERP, TP53 a | Plasma Mem |
| ENSMUSG00000003848 | 2.71E-29 | 2.15E-28 | Nob1 | NIN1 (RPN12 | Nucleus |
| ENSMUSG00000020150 | 2.74E-29 | 2.18E-28 | Gamt | guanidinoace | Cytoplasm |
| ENSMUSG00000005370 | 2.78E-29 | 2.21E-28 | Msh6 | mutS homolo | Nucleus |
| ENSMUSG00000030747 | 2.80E-29 | 2.22E-28 | Dgat2 | diacylglycero | Cytoplasm |
| ENSMUSG00000030089 | 2.80E-29 | 2.22E-28 | Slc41a3 | solute carrier | Plasma Mem |
| ENSMUSG00000021876 | 3.02E-29 | 2.39E-28 | Rnase4 | ribonuclease | Extracellular S |
| ENSMUSG00000025132 | 3.24E-29 | 2.56E-28 | Arhgdia | Rho GDP diss | Cytoplasm |
| ENSMUSG00000031879 | 3.49E-29 | 2.76E-28 | Fam96b | cytosolic iron | Cytoplasm |
| ENSMUSG00000025408 | 3.68E-29 | 2.91E-28 | Ddit3 | DNA damage | Nucleus |
| ENSMUSG00000028452 | 3.85E-29 | 3.04E-28 | Vcp | valosin conta | Cytoplasm |
| ENSMUSG00000021958 | 4.01E-29 | 3.16E-28 | Pinx1 | PIN2 (TERF1) | Nucleus |
| ENSMUSG00000026655 | 4.38E-29 | 3.45E-28 | Fam107b | family with se | Other |
| ENSMUSG00000029610 | 4.81E-29 | 3.79E-28 | Aimp2 | aminoacyl tR | Plasma Mem |
| ENSMUSG00000020307 | 5.04E-29 | 3.96E-28 | Cdc34 | cell division c | Nucleus |
| ENSMUSG00000036693 | 5.17E-29 | 4.06E-28 | Nop14 | NOP14 nucle | Nucleus |
| ENSMUSG00000034435 | 5.24E-29 | 4.11E-28 | Tmem30b | transmembra | Cytoplasm |
| ENSMUSG00000034826 | 5.50E-29 | 4.31E-28 | Nup54 | nucleoporin 5 | Nucleus |
| ENSMUSG00000031774 | 5.57E-29 | 4.36E-28 | Fam192a | family with se | Nucleus |
| ENSMUSG00000001016 | 5.69E-29 | 4.45E-28 | Ilf2 | interleukin en | Nucleus |
| ENSMUSG00000004100 | 6.16E-29 | 4.82E-28 | Ppan | peter pan ho | Nucleus |
| ENSMUSG00000019837 | 6.31E-29 | 4.93E-28 | Gtf3c6 | general trans | Nucleus |
| ENSMUSG00000016018 | 6.57E-29 | 5.12E-28 | Skiv2l2 | Mtr4 exosom | Nucleus |
| ENSMUSG00000031060 | 6.90E-29 | 5.38E-28 | Rbm10 | RNA binding | Nucleus |
| ENSMUSG00000070319 | 7.66E-29 | 5.97E-28 | Eif3g | eukaryotic tra | Cytoplasm |
| ENSMUSG00000033880 | 8.42E-29 | 6.55E-28 | Lgals3bp | galectin 3 bir | Plasma Mem |
| ENSMUSG00000028937 | 8.61E-29 | 6.68E-28 | Acot7 | acyl-CoA thio | Cytoplasm |
| ENSMUSG00000040034 | 8.60E-29 | 6.68E-28 | Nup43 | nucleoporin 4 | Nucleus |
| ENSMUSG00000062908 | 9.02E-29 | 7.00E-28 | Acadm | acyl-CoA deh | Cytoplasm |
| ENSMUSG00000057948 | 9.67E-29 | 7.49E-28 | Unc13d | unc-13 homo | Cytoplasm |
| ENSMUSG00000039783 | 1.01E-28 | 7.83E-28 | Kmo | kynurenine 3 | Cytoplasm |
| ENSMUSG00000002129 | 1.02E-28 | 7.87E-28 | Sf3a1 | splicing facto | Nucleus |
| ENSMUSG00000005625 | 1.05E-28 | 8.13E-28 | Psmd4 | proteasome 2 | Cytoplasm |
| ENSMUSG00000048327 | 1.10E-28 | 8.50E-28 | Ckap2l | cytoskeleton | Cytoplasm |
| ENSMUSG00000022557 | 1.19E-28 | 9.18E-28 | Bop1 | block of proli | Nucleus |
| ENSMUSG00000036376 | 1.22E-28 | 9.43E-28 | Abt1 | activator of b | Nucleus |
| ENSMUSG00000009549 | 1.23E-28 | 9.50E-28 | Srp14 | signal recogn | Cytoplasm |
| ENSMUSG00000035478 | 1.27E-28 | 9.74E-28 | Mbd3 | methyl-CpG | Nucleus |
| ENSMUSG00000040025 | 1.36E-28 | 1.04E-27 | Ythdf2 | YTH N6-met | Cytoplasm |
| ENSMUSG00000028675 | 1.37E-28 | 1.05E-27 | Pnrc2 | proline rich r | Nucleus |
| ENSMUSG00000034317 | 1.39E-28 | 1.07E-27 | Trim59 | tripartite mot | Cytoplasm |
| ENSMUSG00000007029 | 1.48E-28 | 1.14E-27 | Vars | valyl-tRNA sy | Cytoplasm |
| ENSMUSG00000024145 | 1.55E-28 | 1.19E-27 | Pigf | phosphatidyl | Cytoplasm |
| ENSMUSG00000035958 | 2.09E-28 | 1.60E-27 | Tdp2 | tyrosyl-DNA | Cytoplasm |
| ENSMUSG00000001435 | 2.30E-28 | 1.76E-27 | Col18a1 | collagen type | Extracellular S |
| ENSMUSG00000028614 | 2.41E-28 | 1.84E-27 | Ndc1 | NDC1 transm | Nucleus |
| ENSMUSG00000021079 | 2.47E-28 | 1.88E-27 | Timm9 | translocase o | Cytoplasm |
| ENSMUSG00000008301 | 2.53E-28 | 1.92E-27 | Phax | phosphorylat | Cytoplasm |
| ENSMUSG00000021965 | 2.63E-28 | 2.00E-27 | Ska3 | spindle and k | Nucleus |
| ENSMUSG00000028882 | 2.77E-28 | 2.10E-27 | Ppp1r8 | protein phos | Nucleus |

| | | | | | |
|---|---|---|---|---|---|
| ENSMUSG00000029283 | 3.09E-28 | 2.35E-27 | Cdc7 | cell division c | Nucleus |
| ENSMUSG00000028792 | 3.25E-28 | 2.47E-27 | Ak2 | adenylate kir | Cytoplasm |
| ENSMUSG00000018286 | 3.96E-28 | 3.00E-27 | Psmb6 | proteasome s | Nucleus |
| ENSMUSG00000027408 | 4.06E-28 | 3.08E-27 | Cpxm1 | carboxypepti | Extracellular S |
| ENSMUSG00000038224 | 4.56E-28 | 3.45E-27 | Serpinf2 | serpin family | Extracellular S |
| ENSMUSG00000030619 | 4.59E-28 | 3.47E-27 | Eed | embryonic ec | Nucleus |
| ENSMUSG00000027187 | 4.76E-28 | 3.59E-27 | Cat | catalase | Cytoplasm |
| ENSMUSG00000020430 | 4.76E-28 | 3.59E-27 | Pes1 | pescadillo rib | Nucleus |
| ENSMUSG00000020392 | 4.79E-28 | 3.61E-27 | Cdkn2aipnl | CDKN2A inte | Other |
| ENSMUSG00000021266 | 4.90E-28 | 3.69E-27 | Wars | tryptophanyl | Cytoplasm |
| ENSMUSG00000022466 | 5.28E-28 | 3.97E-27 | Rpap3 | RNA polymer | Cytoplasm |
| ENSMUSG00000020469 | 5.41E-28 | 4.07E-27 | Myl7 | myosin light | Cytoplasm |
| ENSMUSG00000030244 | 5.95E-28 | 4.47E-27 | Gys2 | glycogen syn | Cytoplasm |
| ENSMUSG00000042541 | 6.40E-28 | 4.80E-27 | Shfm1 | SEM1, 26S pr | Nucleus |
| ENSMUSG00000038503 | 6.73E-28 | 5.04E-27 | Mesdc2 | mesoderm de | Extracellular S |
| ENSMUSG00000028034 | 6.82E-28 | 5.11E-27 | Fubp1 | far upstream | Nucleus |
| ENSMUSG00000004264 | 7.16E-28 | 5.35E-27 | Phb2 | prohibitin 2 | Cytoplasm |
| ENSMUSG00000022698 | 7.88E-28 | 5.88E-27 | Naa50 | N(alpha)-ace | Cytoplasm |
| ENSMUSG00000039680 | 7.95E-28 | 5.93E-27 | Mrps6 | mitochondria | Cytoplasm |
| ENSMUSG00000039231 | 7.95E-28 | 5.93E-27 | Suv39h1 | suppressor o | Nucleus |
| ENSMUSG00000039748 | 8.05E-28 | 5.99E-27 | Exo1 | exonuclease | Nucleus |
| ENSMUSG00000020219 | 8.59E-28 | 6.39E-27 | Timm13 | translocase o | Cytoplasm |
| ENSMUSG00000019478 | 8.74E-28 | 6.50E-27 | Rab4a | RAB4A, mem | Cytoplasm |
| ENSMUSG00000022884 | 8.91E-28 | 6.61E-27 | Eif4a2 | eukaryotic tra | Cytoplasm |
| ENSMUSG00000004096 | 9.73E-28 | 7.22E-27 | Cwc15 | CWC15 splice | Nucleus |
| ENSMUSG00000002984 | 9.93E-28 | 7.36E-27 | Tomm40 | translocase o | Cytoplasm |
| ENSMUSG00000027870 | 1.03E-27 | 7.65E-27 | Hao2 | hydroxyacid | Cytoplasm |
| ENSMUSG00000003355 | 1.21E-27 | 8.94E-27 | Fkbp11 | FK506 bindin | Cytoplasm |
| ENSMUSG00000001783 | 1.23E-27 | 9.09E-27 | Rtcb | RNA 2',3'-cyc | Cytoplasm |
| ENSMUSG00000004032 | 1.23E-27 | 9.10E-27 | Gstm5 | glutathione S | Cytoplasm |
| ENSMUSG00000059343 | 1.32E-27 | 9.74E-27 | Aldoart1 | aldolase 1 A, | Extracellular S |
| ENSMUSG00000043284 | 1.38E-27 | 1.02E-26 | Tmem11 | transmembra | Plasma Mem |
| ENSMUSG00000056820 | 1.39E-27 | 1.02E-26 | Tsnax | translin assoc | Nucleus |
| ENSMUSG00000036968 | 1.39E-27 | 1.02E-26 | Cnpy4 | canopy FGF s | Other |
| ENSMUSG00000020899 | 1.43E-27 | 1.05E-26 | Pfas | phosphoribo | Cytoplasm |
| ENSMUSG00000018796 | 1.55E-27 | 1.14E-26 | Acsl1 | acyl-CoA syr | Cytoplasm |
| ENSMUSG00000025481 | 1.67E-27 | 1.23E-26 | Urah | urate (5-hydı | Cytoplasm |
| ENSMUSG00000028979 | 2.01E-27 | 1.47E-26 | Masp2 | mannan binc | Extracellular S |
| ENSMUSG00000035754 | 2.02E-27 | 1.48E-26 | Wdr18 | WD repeat d | Nucleus |
| ENSMUSG00000071451 | 2.23E-27 | 1.63E-26 | Psmg4 | proteasome a | Other |
| ENSMUSG00000004667 | 2.41E-27 | 1.76E-26 | Polr2e | RNA polymer | Nucleus |
| ENSMUSG00000024991 | 2.58E-27 | 1.89E-26 | Eif3a | eukaryotic tra | Cytoplasm |
| ENSMUSG00000037149 | 3.07E-27 | 2.24E-26 | Ddx1 | DEAD-box he | Nucleus |
| ENSMUSG00000022070 | 3.10E-27 | 2.26E-26 | Bora | bora, aurora | Cytoplasm |
| ENSMUSG00000034880 | 3.24E-27 | 2.36E-26 | Mrpl34 | mitochondria | Cytoplasm |
| ENSMUSG00000025995 | 3.34E-27 | 2.43E-26 | Wdr75 | WD repeat d | Nucleus |
| ENSMUSG00000038812 | 3.52E-27 | 2.56E-26 | Trmt112 | tRNA methyl | Cytoplasm |
| ENSMUSG00000079480 | 3.70E-27 | 2.69E-26 | Pin4 | peptidylproly | Nucleus |
| ENSMUSG00000038580 | 3.71E-27 | 2.70E-26 | Sct | secretin | Extracellular S |
| ENSMUSG00000024613 | 3.78E-27 | 2.74E-26 | Tcof1 | treacle ribosc | Nucleus |
| ENSMUSG00000024181 | 3.87E-27 | 2.81E-26 | Mrpl28 | mitochondria | Cytoplasm |
| ENSMUSG00000028398 | 4.46E-27 | 3.23E-26 | Tmem261 | distal membr | Cytoplasm |
| ENSMUSG00000057506 | 5.02E-27 | 3.63E-26 | Bloc1s2 | biogenesis of | Cytoplasm |
| ENSMUSG00000024248 | 5.19E-27 | 3.75E-26 | Cox7a2l | cytochrome c | Cytoplasm |
| ENSMUSG00000034883 | 5.56E-27 | 4.01E-26 | Lrr1 | leucine rich r | Cytoplasm |
| ENSMUSG00000026496 | 5.56E-27 | 4.01E-26 | Parp1 | poly(ADP-rib | Nucleus |
| ENSMUSG00000002835 | 5.96E-27 | 4.28E-26 | Chaf1a | chromatin as | Nucleus |
| ENSMUSG00000040028 | 6.47E-27 | 4.65E-26 | Elavl1 | ELAV like RN | Cytoplasm |

| | | | | | |
|---|---|---|---|---|---|
| ENSMUSG00000023908 | 6.53E-27 | 4.69E-26 | Pkmyt1 | protein kinas | Cytoplasm |
| ENSMUSG00000040134 | 8.23E-27 | 5.91E-26 | Rdh7 | retinol dehyc | Cytoplasm |
| ENSMUSG00000004929 | 8.78E-27 | 6.29E-26 | Thop1 | thimet oligop | Cytoplasm |
| ENSMUSG00000030654 | 9.07E-27 | 6.49E-26 | Arl6ip1 | ADP ribosyla | Cytoplasm |
| ENSMUSG00000044034 | 9.24E-27 | 6.61E-26 | Npb | neuropeptide | Extracellular S |
| ENSMUSG00000027315 | 9.37E-27 | 6.70E-26 | Spint1 | serine peptid | Extracellular S |
| ENSMUSG00000090145 | 9.90E-27 | 7.07E-26 | Ugt1a6b | UDP glucuro | Cytoplasm |
| ENSMUSG00000031271 | 1.07E-26 | 7.67E-26 | Serpina7 | serpin family | Extracellular S |
| ENSMUSG00000005566 | 1.17E-26 | 8.33E-26 | Trim28 | tripartite mot | Nucleus |
| ENSMUSG00000073705 | 1.24E-26 | 8.86E-26 | Apitd1 | centromere p | Nucleus |
| ENSMUSG00000036112 | 1.29E-26 | 9.16E-26 | Metap2 | methionyl an | Cytoplasm |
| ENSMUSG00000023044 | 1.36E-26 | 9.68E-26 | Csad | cysteine sulfi | Cytoplasm |
| ENSMUSG00000016637 | 1.37E-26 | 9.75E-26 | Ift27 | intraflagellar | Extracellular S |
| ENSMUSG00000033166 | 1.43E-26 | 1.02E-25 | Dis3 | DIS3 homolo | Nucleus |
| ENSMUSG00000020801 | 1.77E-26 | 1.25E-25 | Med31 | mediator cor | Nucleus |
| ENSMUSG00000012443 | 1.79E-26 | 1.27E-25 | Kif11 | kinesin family | Nucleus |
| ENSMUSG00000027968 | 2.16E-26 | 1.53E-25 | Larp7 | La ribonuclec | Nucleus |
| ENSMUSG00000060613 | 2.22E-26 | 1.57E-25 | Cyp2c70 | cytochrome I | Cytoplasm |
| ENSMUSG00000024587 | 2.45E-26 | 1.73E-25 | Nars | asparaginyl-t | Cytoplasm |
| ENSMUSG00000024989 | 2.57E-26 | 1.82E-25 | Cep55 | centrosomal | Cytoplasm |
| ENSMUSG00000027455 | 2.67E-26 | 1.89E-25 | Nsfl1c | NSFL1 cofact | Cytoplasm |
| ENSMUSG00000037204 | 2.69E-26 | 1.89E-25 | Atg101 | autophagy re | Cytoplasm |
| ENSMUSG00000079555 | 2.75E-26 | 1.93E-25 | Haus3 | HAUS augmi | Cytoplasm |
| ENSMUSG00000027184 | 2.96E-26 | 2.08E-25 | Caprin1 | cell cycle ass | Plasma Mem |
| ENSMUSG00000004937 | 3.05E-26 | 2.15E-25 | Sgta | small glutam | Cytoplasm |
| ENSMUSG00000030615 | 3.26E-26 | 2.29E-25 | Tmem126a | transmembra | Cytoplasm |
| ENSMUSG00000029250 | 3.27E-26 | 2.29E-25 | Polr2b | RNA polymer | Nucleus |
| ENSMUSG00000032231 | 3.46E-26 | 2.43E-25 | Anxa2 | annexin A2 | Plasma Mem |
| ENSMUSG00000028066 | 3.58E-26 | 2.51E-25 | Pmf1 | polyamine m | Nucleus |
| ENSMUSG00000020929 | 3.73E-26 | 2.61E-25 | Eftud2 | elongation fa | Nucleus |
| ENSMUSG00000025381 | 3.81E-26 | 2.66E-25 | Cnpy2 | canopy FGF s | Plasma Mem |
| ENSMUSG00000022009 | 3.81E-26 | 2.66E-25 | Nufip1 | NUFIP1, FMR | Nucleus |
| ENSMUSG00000029326 | 4.03E-26 | 2.81E-25 | Enoph1 | enolase-pho | Cytoplasm |
| ENSMUSG00000096768 | 4.12E-26 | 2.87E-25 | Erdr1 | | Other |
| ENSMUSG00000022219 | 4.35E-26 | 3.03E-25 | Cideb | cell death-in | Cytoplasm |
| ENSMUSG00000022556 | 4.47E-26 | 3.11E-25 | Hsf1 | heat shock tr | Nucleus |
| ENSMUSG00000041548 | 4.72E-26 | 3.28E-25 | Hspb8 | heat shock p | Cytoplasm |
| ENSMUSG00000002458 | 4.87E-26 | 3.38E-25 | Rgs19 | regulator of ( | Cytoplasm |
| ENSMUSG00000022814 | 5.47E-26 | 3.79E-25 | Umps | uridine monc | Cytoplasm |
| ENSMUSG00000015672 | 6.04E-26 | 4.19E-25 | Mrpl32 | mitochondria | Cytoplasm |
| ENSMUSG00000024799 | 6.10E-26 | 4.23E-25 | Tm7sf2 | transmembra | Cytoplasm |
| ENSMUSG00000031029 | 6.56E-26 | 4.53E-25 | Eif3f | eukaryotic tra | Cytoplasm |
| ENSMUSG00000029390 | 6.76E-26 | 4.67E-25 | Tmed2 | transmembra | Cytoplasm |
| ENSMUSG00000026245 | 7.99E-26 | 5.51E-25 | Farsb | phenylalanyl- | Cytoplasm |
| ENSMUSG00000001175 | 8.13E-26 | 5.60E-25 | Calm1 | calmodulin 1 | Nucleus |
| ENSMUSG00000027423 | 8.46E-26 | 5.83E-25 | Snx5 | sorting nexin | Cytoplasm |
| ENSMUSG00000026229 | 8.80E-26 | 6.06E-25 | Psmd1 | proteasome : | Cytoplasm |
| ENSMUSG00000024902 | 9.80E-26 | 6.74E-25 | Mrpl11 | mitochondria | Cytoplasm |
| ENSMUSG00000001082 | 1.00E-25 | 6.90E-25 | Mfsd10 | major facilita | Other |
| ENSMUSG00000003617 | 1.06E-25 | 7.30E-25 | Cp | ceruloplasmi | Extracellular S |
| ENSMUSG00000031157 | 1.06E-25 | 7.30E-25 | Pqbp1 | polyglutamin | Nucleus |
| ENSMUSG00000061983 | 1.09E-25 | 7.48E-25 | Rps12 | ribosomal pr | Cytoplasm |
| ENSMUSG00000018565 | 1.11E-25 | 7.60E-25 | Elp5 | elongator ac | Cytoplasm |
| ENSMUSG00000041012 | 1.12E-25 | 7.65E-25 | Cmtm8 | CKLF like MA | Extracellular S |
| ENSMUSG00000026566 | 1.13E-25 | 7.71E-25 | Mpzl1 | myelin protei | Plasma Mem |
| ENSMUSG00000020492 | 1.25E-25 | 8.55E-25 | Ska2 | spindle and k | Nucleus |
| ENSMUSG00000029387 | 1.50E-25 | 1.02E-24 | Gtf2h3 | general trans | Nucleus |
| ENSMUSG00000046562 | 1.59E-25 | 1.08E-24 | Unc119b | unc-119 lipic | Cytoplasm |

| | | | | | |
|---|---|---|---|---|---|
| ENSMUSG00000004558 | 1.63E-25 | 1.11E-24 | Ndrg2 | NDRG family | Cytoplasm |
| ENSMUSG00000049960 | 1.69E-25 | 1.15E-24 | Mrps16 | mitochondria | Cytoplasm |
| ENSMUSG00000011884 | 1.72E-25 | 1.17E-24 | Gltp | glycolipid tra | Cytoplasm |
| ENSMUSG00000032301 | 1.85E-25 | 1.26E-24 | Psma4 | proteasome s | Cytoplasm |
| ENSMUSG00000006585 | 1.95E-25 | 1.32E-24 | Cdt1 | chromatin lic | Nucleus |
| ENSMUSG00000057497 | 2.08E-25 | 1.41E-24 | Fam136a | family with se | Cytoplasm |
| ENSMUSG00000021606 | 2.21E-25 | 1.50E-24 | Ndufs6 | NADH:ubiqu | Cytoplasm |
| ENSMUSG00000040354 | 2.33E-25 | 1.58E-24 | Mars | methionyl-tR | Cytoplasm |
| ENSMUSG00000059409 | 2.34E-25 | 1.58E-24 | Ppp2r5d | protein phos | Nucleus |
| ENSMUSG00000033809 | 2.39E-25 | 1.62E-24 | Alg3 | ALG3, alpha- | Cytoplasm |
| ENSMUSG00000037058 | 2.40E-25 | 1.62E-24 | Paip2 | poly(A) bindi | Cytoplasm |
| ENSMUSG00000027206 | 2.68E-25 | 1.81E-24 | Cops2 | COP9 signalc | Nucleus |
| ENSMUSG00000071528 | 2.72E-25 | 1.83E-24 | Usmg5 | ATP synthase | Cytoplasm |
| ENSMUSG00000001767 | 3.06E-25 | 2.06E-24 | Crnkl1 | crooked neck | Nucleus |
| ENSMUSG00000035835 | 3.10E-25 | 2.09E-24 | Lppr3 | phospholipid | Other |
| ENSMUSG00000067150 | 3.39E-25 | 2.28E-24 | Xpo5 | exportin 5 | Nucleus |
| ENSMUSG00000034432 | 3.41E-25 | 2.29E-24 | Cops8 | COP9 signalc | Nucleus |
| ENSMUSG00000063052 | 3.43E-25 | 2.30E-24 | Lrrc40 | leucine rich r | Nucleus |
| ENSMUSG00000075266 | 3.59E-25 | 2.41E-24 | Cenpw | centromere p | Nucleus |
| ENSMUSG00000006494 | 3.89E-25 | 2.61E-24 | Pdk1 | pyruvate deh | Cytoplasm |
| ENSMUSG00000048007 | 3.96E-25 | 2.66E-24 | Timm8a1 | translocase o | Cytoplasm |
| ENSMUSG00000007589 | 4.29E-25 | 2.87E-24 | Tinf2 | TERF1 interac | Nucleus |
| ENSMUSG00000020167 | 4.39E-25 | 2.94E-24 | Tcf3 | transcription | Nucleus |
| ENSMUSG00000024091 | 4.49E-25 | 3.00E-24 | Vapa | VAMP associ | Plasma Mem |
| ENSMUSG00000056671 | 4.61E-25 | 3.08E-24 | Prelid2 | PRELI domair | Cytoplasm |
| ENSMUSG00000008683 | 4.78E-25 | 3.19E-24 | Rps15a | ribosomal pr | Cytoplasm |
| ENSMUSG00000070003 | 5.19E-25 | 3.46E-24 | Ssbp4 | single strand | Nucleus |
| ENSMUSG00000022283 | 6.12E-25 | 4.07E-24 | Pabpc1 | poly(A) bindi | Cytoplasm |
| ENSMUSG00000022149 | 6.35E-25 | 4.23E-24 | C9 | complement | Extracellular S |
| ENSMUSG00000031781 | 6.90E-25 | 4.59E-24 | Ciapin1 | cytokine indu | Cytoplasm |
| ENSMUSG00000028455 | 7.10E-25 | 4.72E-24 | Stoml2 | stomatin like | Plasma Mem |
| ENSMUSG00000034889 | 7.16E-25 | 4.75E-24 | Cactin | cactin, splice | Nucleus |
| ENSMUSG00000001249 | 7.20E-25 | 4.77E-24 | Hpn | hepsin | Plasma Mem |
| ENSMUSG00000041406 | 8.60E-25 | 5.70E-24 | BC055324 | chromosome | Other |
| ENSMUSG00000037296 | 8.72E-25 | 5.78E-24 | Lsm1 | LSM1 homol | Nucleus |
| ENSMUSG00000022913 | 9.10E-25 | 6.02E-24 | Psmg1 | proteasome a | Plasma Mem |
| ENSMUSG00000022346 | 1.05E-24 | 6.95E-24 | Myc | MYC proto-c | Nucleus |
| ENSMUSG00000022024 | 1.11E-24 | 7.33E-24 | Sugt1 | SGT1 homolc | Nucleus |
| ENSMUSG00000017421 | 1.17E-24 | 7.74E-24 | Zfp207 | zinc finger pr | Nucleus |
| ENSMUSG00000036223 | 1.19E-24 | 7.82E-24 | Ska1 | spindle and k | Nucleus |
| ENSMUSG00000018965 | 1.19E-24 | 7.82E-24 | Ywhah | tyrosine 3-m | Cytoplasm |
| ENSMUSG00000035914 | 1.23E-24 | 8.10E-24 | Cd276 | CD276 molec | Plasma Mem |
| ENSMUSG00000054115 | 1.27E-24 | 8.35E-24 | Skp2 | S-phase kina | Nucleus |
| ENSMUSG00000021692 | 1.28E-24 | 8.40E-24 | Dimt1 | DIM1 dimeth | Cytoplasm |
| ENSMUSG00000006932 | 1.35E-24 | 8.85E-24 | Ctnnb1 | catenin beta | Nucleus |
| ENSMUSG00000091931 | 1.41E-24 | 9.24E-24 | AK010878 | GON7, KEOP | Nucleus |
| ENSMUSG00000017999 | 1.48E-24 | 9.73E-24 | Ddx27 | DEAD-box h | Nucleus |
| ENSMUSG00000019373 | 1.52E-24 | 9.94E-24 | Cops3 | COP9 signalc | Cytoplasm |
| ENSMUSG00000017176 | 1.55E-24 | 1.02E-23 | Nt5c3b | 5'-nucleotida | Cytoplasm |
| ENSMUSG00000025747 | 1.59E-24 | 1.04E-23 | Tyms | thymidylate s | Nucleus |
| ENSMUSG00000007338 | 1.64E-24 | 1.07E-23 | Mrpl49 | mitochondria | Cytoplasm |
| ENSMUSG00000020994 | 1.64E-24 | 1.07E-23 | Pnn | pinin, desmo | Plasma Mem |
| ENSMUSG00000031508 | 1.65E-24 | 1.08E-23 | Ankrd10 | ankyrin repea | Nucleus |
| ENSMUSG00000018604 | 1.76E-24 | 1.15E-23 | Tbx3 | T-box 3 | Nucleus |
| ENSMUSG00000022682 | 1.86E-24 | 1.21E-23 | Rrn3 | RRN3 homol | Nucleus |
| ENSMUSG00000018666 | 1.90E-24 | 1.23E-23 | Cbx1 | chromobox 1 | Nucleus |
| ENSMUSG00000074272 | 2.10E-24 | 1.36E-23 | Ceacam1 | carcinoembry | Plasma Mem |
| ENSMUSG00000071646 | 2.09E-24 | 1.36E-23 | Mta2 | metastasis as | Nucleus |

| | | | | | |
|---|---|---|---|---|---|
| ENSMUSG00000036768 | 2.17E-24 | 1.41E-23 | Kif15 | kinesin family | Nucleus |
| ENSMUSG00000024493 | 2.20E-24 | 1.42E-23 | Lars | leucyl-tRNA : | Cytoplasm |
| ENSMUSG00000038416 | 2.38E-24 | 1.54E-23 | Cdc16 | cell division c | Nucleus |
| ENSMUSG00000020074 | 2.46E-24 | 1.59E-23 | Ccar1 | cell division c | Nucleus |
| ENSMUSG00000027134 | 2.51E-24 | 1.62E-23 | Lpcat4 | lysophosphat | Cytoplasm |
| ENSMUSG00000025007 | 2.59E-24 | 1.67E-23 | Aldh18a1 | aldehyde del | Cytoplasm |
| ENSMUSG00000016253 | 2.71E-24 | 1.75E-23 | Nelfcd | negative elor | Nucleus |
| ENSMUSG00000031400 | 2.76E-24 | 1.78E-23 | G6pdx | glucose-6-pl | Cytoplasm |
| ENSMUSG00000034456 | 3.17E-24 | 2.04E-23 | Uroc1 | urocanate hy | Cytoplasm |
| ENSMUSG00000061959 | 3.25E-24 | 2.10E-23 | Ces1e | carboxylester | Cytoplasm |
| ENSMUSG00000026999 | 3.32E-24 | 2.13E-23 | Nup35 | nucleoporin : | Nucleus |
| ENSMUSG00000021326 | 3.42E-24 | 2.20E-23 | Trim27 | tripartite mot | Nucleus |
| ENSMUSG00000052752 | 3.94E-24 | 2.53E-23 | Traf7 | TNF receptor | Cytoplasm |
| ENSMUSG00000019738 | 4.38E-24 | 2.81E-23 | Polr2i | RNA polymer | Nucleus |
| ENSMUSG00000026966 | 4.55E-24 | 2.92E-23 | Ssna1 | SS nuclear au | Cytoplasm |
| ENSMUSG00000028583 | 4.73E-24 | 3.03E-23 | Pdpn | podoplanin | Plasma Mem |
| ENSMUSG00000022360 | 5.38E-24 | 3.44E-23 | Atad2 | ATPase famil | Nucleus |
| ENSMUSG00000024666 | 5.40E-24 | 3.46E-23 | Tmem138 | transmembra | Extracellular S |
| ENSMUSG00000028069 | 5.41E-24 | 3.46E-23 | Gpatch4 | G-patch dom | Other |
| ENSMUSG00000032320 | 5.93E-24 | 3.79E-23 | Rcn2 | reticulocalbir | Cytoplasm |
| ENSMUSG00000027998 | 6.07E-24 | 3.87E-23 | Plrg1 | pleiotropic re | Nucleus |
| ENSMUSG00000047260 | 6.12E-24 | 3.90E-23 | Emc6 | ER membran | Cytoplasm |
| ENSMUSG00000049751 | 6.21E-24 | 3.96E-23 | Rpl36al | ribosomal pr | Cytoplasm |
| ENSMUSG00000022678 | 7.50E-24 | 4.78E-23 | Nde1 | nudE neurod | Nucleus |
| ENSMUSG00000038489 | 7.84E-24 | 4.99E-23 | Polr2l | RNA polymer | Nucleus |
| ENSMUSG00000052456 | 7.86E-24 | 5.00E-23 | Asna1 | arsA arsenite | Nucleus |
| ENSMUSG00000021039 | 7.97E-24 | 5.06E-23 | Snw1 | SNW domair | Nucleus |
| ENSMUSG00000004035 | 8.25E-24 | 5.23E-23 | Gstm7 | glutathione S | Cytoplasm |
| ENSMUSG00000033581 | 8.24E-24 | 5.23E-23 | Igf2bp2 | insulin like gr | Cytoplasm |
| ENSMUSG00000028187 | 8.93E-24 | 5.66E-23 | Rpf1 | ribosome prc | Nucleus |
| ENSMUSG00000003269 | 9.63E-24 | 6.10E-23 | Cyth2 | cytohesin 2 | Cytoplasm |
| ENSMUSG00000054312 | 1.06E-23 | 6.73E-23 | Mrps21 | mitochondria | Cytoplasm |
| ENSMUSG00000032076 | 1.09E-23 | 6.90E-23 | Cadm1 | cell adhesion | Plasma Mem |
| ENSMUSG00000061474 | 1.28E-23 | 8.05E-23 | Mrps36 | mitochondria | Cytoplasm |
| ENSMUSG00000024947 | 1.30E-23 | 8.18E-23 | Men1 | menin 1 | Nucleus |
| ENSMUSG00000020677 | 1.35E-23 | 8.53E-23 | Ddx52 | DExD-box he | Nucleus |
| ENSMUSG00000031897 | 1.38E-23 | 8.67E-23 | Psmb10 | proteasome s | Cytoplasm |
| ENSMUSG00000072980 | 1.39E-23 | 8.71E-23 | Oip5 | Opa interacti | Nucleus |
| ENSMUSG00000010760 | 1.41E-23 | 8.83E-23 | Phlda2 | pleckstrin ho | Cytoplasm |
| ENSMUSG00000029581 | 1.45E-23 | 9.11E-23 | Fscn1 | fascin actin-b | Cytoplasm |
| ENSMUSG00000075517 | 1.62E-23 | 1.02E-22 | Cyp2d37-ps | cytochrome I | Other |
| ENSMUSG00000018574 | 1.63E-23 | 1.02E-22 | Acadvl | acyl-CoA del | Cytoplasm |
| ENSMUSG00000020571 | 1.65E-23 | 1.03E-22 | Pdia6 | protein disulf | Cytoplasm |
| ENSMUSG00000022407 | 1.76E-23 | 1.10E-22 | Adsl | adenylosucci | Cytoplasm |
| ENSMUSG00000041660 | 1.83E-23 | 1.14E-22 | Bbox1 | gamma-buty | Cytoplasm |
| ENSMUSG00000057074 | 1.83E-23 | 1.15E-22 | Ces1g | carboxylester | Other |
| ENSMUSG00000025374 | 2.10E-23 | 1.31E-22 | Nabp2 | nucleic acid b | Nucleus |
| ENSMUSG00000028568 | 2.11E-23 | 1.32E-22 | Btf3l4 | basic transcri | Other |
| ENSMUSG00000022204 | 2.20E-23 | 1.37E-22 | Ngdn | neuroguidin | Nucleus |
| ENSMUSG00000027871 | 2.24E-23 | 1.40E-22 | Hsd3b1 | hydroxy-delt | Cytoplasm |
| ENSMUSG00000054428 | 2.29E-23 | 1.42E-22 | Atpif1 | ATP synthase | Cytoplasm |
| ENSMUSG00000074247 | 2.37E-23 | 1.48E-22 | Dda1 | DET1 and DD | Nucleus |
| ENSMUSG00000041491 | 2.47E-23 | 1.53E-22 | Cep78 | centrosomal | Cytoplasm |
| ENSMUSG00000078348 | 2.52E-23 | 1.56E-22 | Sf3b5 | splicing facto | Nucleus |
| ENSMUSG00000038722 | 2.59E-23 | 1.60E-22 | Bud31 | BUD31 homo | Nucleus |
| ENSMUSG00000042660 | 2.61E-23 | 1.61E-22 | Wdr55 | WD repeat d | Nucleus |
| ENSMUSG00000042797 | 2.65E-23 | 1.64E-22 | Aqp11 | aquaporin 11 | Plasma Mem |
| ENSMUSG00000022887 | 2.67E-23 | 1.65E-22 | Masp1 | mannan-binc | Extracellular S |

| | | | | | |
|---|---|---|---|---|---|
| ENSMUSG00000027323 | 2.76E-23 | 1.71E-22 | Rad51 | RAD51 recon | Nucleus |
| ENSMUSG00000042590 | 2.78E-23 | 1.72E-22 | Ipo11 | importin 11 | Nucleus |
| ENSMUSG00000034674 | 2.83E-23 | 1.75E-22 | Tdg | thymine DNA | Nucleus |
| ENSMUSG00000028798 | 2.88E-23 | 1.78E-22 | Eif3i | eukaryotic tra | Cytoplasm |
| ENSMUSG00000038286 | 2.89E-23 | 1.78E-22 | Bphl | biphenyl hyd | Cytoplasm |
| ENSMUSG00000070544 | 3.13E-23 | 1.93E-22 | Top1 | DNA topoiso | Nucleus |
| ENSMUSG00000004099 | 3.57E-23 | 2.20E-22 | Dnmt1 | DNA methylt | Nucleus |
| ENSMUSG00000030802 | 3.65E-23 | 2.24E-22 | Bckdk | branched cha | Cytoplasm |
| ENSMUSG00000031628 | 3.71E-23 | 2.28E-22 | Casp3 | caspase 3 | Cytoplasm |
| ENSMUSG00000029686 | 3.82E-23 | 2.34E-22 | Cul1 | cullin 1 | Nucleus |
| ENSMUSG00000036860 | 3.95E-23 | 2.42E-22 | Mrpl55 | mitochondria | Cytoplasm |
| ENSMUSG00000015312 | 4.17E-23 | 2.55E-22 | Gadd45b | growth arrest | Cytoplasm |
| ENSMUSG00000066705 | 4.65E-23 | 2.84E-22 | Fxyd6 | FXYD domair | Plasma Mem |
| ENSMUSG00000074264 | 5.44E-23 | 3.32E-22 | Amy1 | amylase, alpl | Extracellular S |
| ENSMUSG00000026037 | 5.86E-23 | 3.58E-22 | Orc2 | origin recogr | Nucleus |
| ENSMUSG00000032400 | 6.33E-23 | 3.86E-22 | Zwilch | zwilch kineto | Nucleus |
| ENSMUSG00000024829 | 6.46E-23 | 3.93E-22 | Mrpl21 | mitochondria | Cytoplasm |
| ENSMUSG00000035473 | 6.90E-23 | 4.20E-22 | Galm | galactose mu | Cytoplasm |
| ENSMUSG00000023021 | 7.27E-23 | 4.42E-22 | Cers5 | ceramide syn | Cytoplasm |
| ENSMUSG00000038900 | 7.64E-23 | 4.64E-22 | Rpl12 | ribosomal pr | Nucleus |
| ENSMUSG00000036285 | 8.00E-23 | 4.86E-22 | Noa1 | nitric oxide a | Cytoplasm |
| ENSMUSG00000028312 | 8.28E-23 | 5.02E-22 | Smc2 | structural ma | Nucleus |
| ENSMUSG00000038299 | 8.98E-23 | 5.45E-22 | Wdr36 | WD repeat d | Extracellular S |
| ENSMUSG00000038398 | 9.11E-23 | 5.52E-22 | Upf3a | UPF3A, regul | Nucleus |
| ENSMUSG00000072082 | 9.43E-23 | 5.71E-22 | Ccnf | cyclin F | Nucleus |
| ENSMUSG00000004788 | 9.57E-23 | 5.79E-22 | Eif2b2 | eukaryotic tra | Cytoplasm |
| ENSMUSG00000031328 | 9.68E-23 | 5.85E-22 | Flna | filamin A | Cytoplasm |
| ENSMUSG00000027805 | 9.80E-23 | 5.92E-22 | Pfn2 | profilin 2 | Cytoplasm |
| ENSMUSG00000024346 | 1.18E-22 | 7.13E-22 | Pfdn1 | prefoldin sub | Cytoplasm |
| ENSMUSG00000024045 | 1.21E-22 | 7.27E-22 | Akap8 | A-kinase anc | Nucleus |
| ENSMUSG00000054474 | 1.21E-22 | 7.27E-22 | Thnsl2 | threonine syr | Other |
| ENSMUSG00000000303 | 1.21E-22 | 7.27E-22 | Cdh1 | cadherin 1 | Plasma Mem |
| ENSMUSG00000013415 | 1.24E-22 | 7.44E-22 | Igf2bp1 | insulin like gr | Cytoplasm |
| ENSMUSG00000074227 | 1.27E-22 | 7.63E-22 | Spint2 | serine peptid | Extracellular S |
| ENSMUSG00000021608 | 1.40E-22 | 8.38E-22 | Lpcat1 | lysophosphat | Cytoplasm |
| ENSMUSG00000062075 | 1.41E-22 | 8.44E-22 | Lmnb2 | lamin B2 | Nucleus |
| ENSMUSG00000051695 | 1.42E-22 | 8.49E-22 | Pcbp1 | poly(rC) bind | Nucleus |
| ENSMUSG00000032113 | 1.50E-22 | 9.01E-22 | Chek1 | checkpoint ki | Nucleus |
| ENSMUSG00000022797 | 1.51E-22 | 9.02E-22 | Tfrc | transferrin re | Plasma Mem |
| ENSMUSG00000041459 | 1.57E-22 | 9.39E-22 | Tardbp | TAR DNA bir | Nucleus |
| ENSMUSG00000045763 | 1.59E-22 | 9.52E-22 | Basp1 | brain abunda | Nucleus |
| ENSMUSG00000079426 | 1.61E-22 | 9.59E-22 | Arpc4 | actin related | Cytoplasm |
| ENSMUSG00000041697 | 1.68E-22 | 1.00E-21 | Cox6a1 | cytochrome c | Cytoplasm |
| ENSMUSG00000034064 | 1.71E-22 | 1.02E-21 | Poglut1 | protein O-gl | Extracellular S |
| ENSMUSG00000030726 | 1.80E-22 | 1.07E-21 | Pold3 | DNA polyme | Nucleus |
| ENSMUSG00000028820 | 1.80E-22 | 1.07E-21 | Sfpq | splicing facto | Nucleus |
| ENSMUSG00000021033 | 1.83E-22 | 1.09E-21 | Gstz1 | glutathione S | Cytoplasm |
| ENSMUSG00000056962 | 1.89E-22 | 1.13E-21 | Jmjd6 | arginine dem | Plasma Mem |
| ENSMUSG00000078521 | 2.03E-22 | 1.20E-21 | Aunip | aurora kinase | Cytoplasm |
| ENSMUSG00000071708 | 2.08E-22 | 1.23E-21 | Sms | spermine syn | Cytoplasm |
| ENSMUSG00000035150 | 2.09E-22 | 1.24E-21 | Eif2s3x | eukaryotic tra | Cytoplasm |
| ENSMUSG00000015126 | 2.13E-22 | 1.26E-21 | Tsr3 | TSR3, acp tra | Cytoplasm |
| ENSMUSG00000028433 | 2.13E-22 | 1.26E-21 | Ubap2 | ubiquitin ass | Cytoplasm |
| ENSMUSG00000032180 | 2.15E-22 | 1.27E-21 | Tmed1 | transmembra | Extracellular S |
| ENSMUSG00000036442 | 2.16E-22 | 1.28E-21 | Thap11 | THAP domai | Nucleus |
| ENSMUSG00000063445 | 2.21E-22 | 1.31E-21 | Nmral1 | NmrA like re | Nucleus |
| ENSMUSG00000060860 | 2.23E-22 | 1.32E-21 | Ube2s | ubiquitin cor | Nucleus |
| ENSMUSG00000020973 | 2.49E-22 | 1.47E-21 | Dnaaf2 | dynein axone | Cytoplasm |

| | | | | | |
|---|---|---|---|---|---|
| ENSMUSG00000025492 | 2.57E-22 | 1.52E-21 | Ifitm3 | interferon inc | Plasma Mem |
| ENSMUSG00000029404 | 2.62E-22 | 1.55E-21 | Arl6ip4 | ADP ribosyla | Nucleus |
| ENSMUSG00000057895 | 2.68E-22 | 1.58E-21 | Zfp105 | zinc finger pr | Nucleus |
| ENSMUSG00000005973 | 2.70E-22 | 1.59E-21 | Rcn1 | reticulocalbir | Cytoplasm |
| ENSMUSG00000027248 | 2.73E-22 | 1.60E-21 | Pdia3 | protein disulf | Cytoplasm |
| ENSMUSG00000022177 | 2.98E-22 | 1.75E-21 | Haus4 | HAUS augmi | Cytoplasm |
| ENSMUSG00000021620 | 3.16E-22 | 1.86E-21 | Acot12 | acyl-CoA thic | Cytoplasm |
| ENSMUSG00000021391 | 3.18E-22 | 1.86E-21 | Cenpp | centromere p | Nucleus |
| ENSMUSG00000018848 | 3.35E-22 | 1.96E-21 | Rars | arginyl-tRNA | Cytoplasm |
| ENSMUSG00000037025 | 3.70E-22 | 2.17E-21 | Foxa2 | forkhead box | Nucleus |
| ENSMUSG00000059005 | 3.85E-22 | 2.26E-21 | Hnrnpa3 | heterogeneo | Nucleus |
| ENSMUSG00000022613 | 3.92E-22 | 2.30E-21 | Miox | myo-inositol | Cytoplasm |
| ENSMUSG00000052033 | 4.06E-22 | 2.38E-21 | Pfdn4 | prefoldin sub | Cytoplasm |
| ENSMUSG00000031197 | 4.07E-22 | 2.38E-21 | Vbp1 | VHL binding | Cytoplasm |
| ENSMUSG00000030612 | 4.09E-22 | 2.39E-21 | Mrpl46 | mitochondria | Cytoplasm |
| ENSMUSG00000032607 | 4.15E-22 | 2.42E-21 | Amt | aminomethyl | Cytoplasm |
| ENSMUSG00000047459 | 4.52E-22 | 2.64E-21 | Dynlrb1 | dynein light c | Cytoplasm |
| ENSMUSG00000063802 | 4.55E-22 | 2.65E-21 | Hspbp1 | HSPA (Hsp70 | Cytoplasm |
| ENSMUSG00000025591 | 4.56E-22 | 2.66E-21 | Tma16 | translation m | Nucleus |
| ENSMUSG00000028088 | 4.73E-22 | 2.75E-21 | Fmo5 | flavin contair | Cytoplasm |
| ENSMUSG00000060743 | 4.80E-22 | 2.79E-21 | H3f3a | H3 histone fa | Nucleus |
| ENSMUSG00000018239 | 4.81E-22 | 2.80E-21 | Zcchc10 | zinc finger C( | Other |
| ENSMUSG00000030876 | 5.11E-22 | 2.97E-21 | Mettl9 | methyltransfe | Other |
| ENSMUSG00000025264 | 5.15E-22 | 2.99E-21 | Tsr2 | TSR2, ribosor | Other |
| ENSMUSG00000029715 | 5.26E-22 | 3.05E-21 | Pop7 | POP7 homolo | Nucleus |
| ENSMUSG00000024188 | 5.31E-22 | 3.08E-21 | Luc7l | LUC7 like | Nucleus |
| ENSMUSG00000021477 | 5.44E-22 | 3.15E-21 | Ctsl | cathepsin V | Cytoplasm |
| ENSMUSG00000041231 | 5.52E-22 | 3.19E-21 | Ublcp1 | ubiquitin like | Nucleus |
| ENSMUSG00000019297 | 5.61E-22 | 3.24E-21 | Nop9 | NOP9 nucleo | Nucleus |
| ENSMUSG00000013089 | 5.68E-22 | 3.28E-21 | Etv5 | ETS variant 5 | Nucleus |
| ENSMUSG00000008682 | 5.87E-22 | 3.39E-21 | Rpl10 | ribosomal pr | Cytoplasm |
| ENSMUSG00000026960 | 6.02E-22 | 3.47E-21 | Arl6ip6 | ADP ribosyla | Other |
| ENSMUSG00000048170 | 6.03E-22 | 3.48E-21 | Mcmbp | minichromos | Nucleus |
| ENSMUSG00000096546 | 6.64E-22 | 3.83E-21 | Smlr1 | small leucine | Other |
| ENSMUSG00000060475 | 6.64E-22 | 3.83E-21 | Wtap | WT1 associat | Nucleus |
| ENSMUSG00000022949 | 6.77E-22 | 3.90E-21 | Clic6 | chloride intra | Plasma Mem |
| ENSMUSG00000045948 | 7.03E-22 | 4.04E-21 | Mrps12 | mitochondria | Cytoplasm |
| ENSMUSG00000030682 | 7.22E-22 | 4.15E-21 | Cdipt | CDP-diacylgl | Cytoplasm |
| ENSMUSG00000023932 | 7.45E-22 | 4.28E-21 | Cdc5l | cell division c | Nucleus |
| ENSMUSG00000024474 | 7.75E-22 | 4.45E-21 | Ik | IK cytokine | Extracellular S |
| ENSMUSG00000025911 | 7.80E-22 | 4.47E-21 | Adhfe1 | alcohol dehy | Cytoplasm |
| ENSMUSG00000025001 | 8.03E-22 | 4.60E-21 | Hells | helicase, lym | Nucleus |
| ENSMUSG00000048039 | 8.04E-22 | 4.61E-21 | Isg20l2 | interferon sti | Nucleus |
| ENSMUSG00000035443 | 8.21E-22 | 4.70E-21 | Thyn1 | thymocyte nu | Nucleus |
| ENSMUSG00000030800 | 8.66E-22 | 4.95E-21 | Prss8 | serine protea | Extracellular S |
| ENSMUSG00000001445 | 8.77E-22 | 5.01E-21 | Mrpl10 | mitochondria | Cytoplasm |
| ENSMUSG00000041028 | 9.05E-22 | 5.17E-21 | Ghitm | growth horm | Cytoplasm |
| ENSMUSG00000032112 | 9.08E-22 | 5.18E-21 | Trappc4 | trafficking pr | Cytoplasm |
| ENSMUSG00000001158 | 9.39E-22 | 5.36E-21 | Snrnp27 | small nuclear | Nucleus |
| ENSMUSG00000017428 | 9.71E-22 | 5.54E-21 | Psmd11 | proteasome 2 | Cytoplasm |
| ENSMUSG00000016942 | 1.07E-21 | 6.11E-21 | Tmprss6 | transmembra | Plasma Mem |
| ENSMUSG00000026072 | 1.19E-21 | 6.77E-21 | Il1r1 | interleukin 1 | Plasma Mem |
| ENSMUSG00000039865 | 1.24E-21 | 7.03E-21 | Slc44a3 | solute carrier | Plasma Mem |
| ENSMUSG00000056076 | 1.28E-21 | 7.25E-21 | Eif3b | eukaryotic tra | Cytoplasm |
| ENSMUSG00000037572 | 1.31E-21 | 7.46E-21 | Wdhd1 | WD repeat ar | Nucleus |
| ENSMUSG00000030750 | 1.42E-21 | 8.07E-21 | Nsmce1 | NSE1 homolo | Nucleus |
| ENSMUSG00000029169 | 1.43E-21 | 8.12E-21 | Dhx15 | DEAH-box h | Nucleus |
| ENSMUSG00000031821 | 1.54E-21 | 8.72E-21 | Gins2 | GINS comple | Nucleus |

| | | | | | |
|---|---|---|---|---|---|
| ENSMUSG00000001039 | 1.55E-21 | 8.79E-21 | B9d1 | B9 domain cc | Plasma Mem |
| ENSMUSG00000017453 | 1.61E-21 | 9.12E-21 | Pipox | pipecolic acid | Cytoplasm |
| ENSMUSG00000021701 | 1.71E-21 | 9.69E-21 | Plk2 | polo like kina | Nucleus |
| ENSMUSG00000000346 | 1.79E-21 | 1.01E-20 | Dazap2 | DAZ associat | Nucleus |
| ENSMUSG00000039183 | 1.79E-21 | 1.01E-20 | Nubp2 | nucleotide bi | Nucleus |
| ENSMUSG00000070526 | 1.83E-21 | 1.03E-20 | Peg12 | paternally ex | Other |
| ENSMUSG00000039221 | 1.95E-21 | 1.10E-20 | Rpl22l1 | ribosomal pr | Other |
| ENSMUSG00000025534 | 1.99E-21 | 1.12E-20 | Gusb | glucuronidas | Cytoplasm |
| ENSMUSG00000059791 | 1.99E-21 | 1.12E-20 | Nrm | nurim | Nucleus |
| ENSMUSG00000020609 | 2.00E-21 | 1.13E-20 | Apob | apolipoprote | Extracellular S |
| ENSMUSG00000060126 | 2.10E-21 | 1.18E-20 | Tpt1 | tumor protei | Cytoplasm |
| ENSMUSG00000019659 | 2.15E-21 | 1.21E-20 | Ccdc12 | coiled-coil d | Other |
| ENSMUSG00000030316 | 2.26E-21 | 1.27E-20 | Tamm41 | TAM41 mitoc | Cytoplasm |
| ENSMUSG00000031226 | 2.33E-21 | 1.31E-20 | Pbdc1 | polysaccharic | Other |
| ENSMUSG00000038641 | 2.36E-21 | 1.32E-20 | Akr1d1 | aldo-keto rec | Cytoplasm |
| ENSMUSG00000037742 | 2.41E-21 | 1.35E-20 | Eef1a1 | eukaryotic tra | Cytoplasm |
| ENSMUSG00000022043 | 2.43E-21 | 1.36E-20 | Trim35 | tripartite mot | Cytoplasm |
| ENSMUSG00000037568 | 2.55E-21 | 1.43E-20 | Vash2 | vasohibin 2 | Cytoplasm |
| ENSMUSG00000048355 | 2.69E-21 | 1.50E-20 | Arxes1 | adipocyte-re | Cytoplasm |
| ENSMUSG00000039041 | 3.02E-21 | 1.68E-20 | Adrm1 | adhesion reg | Plasma Mem |
| ENSMUSG00000022490 | 3.18E-21 | 1.77E-20 | Ppp1r1a | protein phos | Cytoplasm |
| ENSMUSG00000019966 | 3.32E-21 | 1.85E-20 | Kitl | KIT ligand | Extracellular S |
| ENSMUSG00000023723 | 3.41E-21 | 1.90E-20 | Mrps23 | mitochondria | Cytoplasm |
| ENSMUSG00000031402 | 3.41E-21 | 1.90E-20 | Mpp1 | membrane p | Plasma Mem |
| ENSMUSG00000021939 | 3.49E-21 | 1.94E-20 | Ctsb | cathepsin B | Cytoplasm |
| ENSMUSG00000041782 | 3.54E-21 | 1.97E-20 | Lad1 | ladinin 1 | Extracellular S |
| ENSMUSG00000005204 | 3.70E-21 | 2.05E-20 | Senp3 | SUMO specif | Nucleus |
| ENSMUSG00000021028 | 3.78E-21 | 2.09E-20 | Mbip | MAP3K12 bir | Nucleus |
| ENSMUSG00000021134 | 3.79E-21 | 2.10E-20 | Srsf5 | serine/argini | Nucleus |
| ENSMUSG00000028221 | 3.87E-21 | 2.14E-20 | Tmem55a | phosphatidyl | Cytoplasm |
| ENSMUSG00000051855 | 3.88E-21 | 2.15E-20 | Mest | mesoderm sp | Cytoplasm |
| ENSMUSG00000008136 | 4.01E-21 | 2.21E-20 | Fhl2 | four and a ha | Nucleus |
| ENSMUSG00000024645 | 4.06E-21 | 2.24E-20 | Timm21 | translocase o | Cytoplasm |
| ENSMUSG00000014980 | 4.15E-21 | 2.29E-20 | Tsen15 | tRNA splicing | Nucleus |
| ENSMUSG00000028790 | 4.31E-21 | 2.38E-20 | Khdrbs1 | KH RNA binc | Nucleus |
| ENSMUSG00000024151 | 4.43E-21 | 2.44E-20 | Msh2 | mutS homolo | Nucleus |
| ENSMUSG00000027379 | 4.54E-21 | 2.50E-20 | Bub1 | BUB1 mitotic | Nucleus |
| ENSMUSG00000023905 | 4.84E-21 | 2.66E-20 | Tnfrsf12a | TNF receptor | Plasma Mem |
| ENSMUSG00000029617 | 4.88E-21 | 2.68E-20 | Ccz1 | CCZ1 homolo | Cytoplasm |
| ENSMUSG00000024387 | 5.18E-21 | 2.85E-20 | Csnk2b | casein kinase | Cytoplasm |
| ENSMUSG00000017778 | 5.33E-21 | 2.93E-20 | Cox7c | cytochrome c | Cytoplasm |
| ENSMUSG00000027999 | 5.48E-21 | 3.01E-20 | Pla2g12a | phospholipas | Other |
| ENSMUSG00000075232 | 5.53E-21 | 3.03E-20 | Amd1 | adenosylmet | Cytoplasm |
| ENSMUSG00000047977 | 5.63E-21 | 3.09E-20 | Synb | endogenous | Other |
| ENSMUSG00000020537 | 5.67E-21 | 3.11E-20 | Drg2 | development | Cytoplasm |
| ENSMUSG00000033813 | 6.06E-21 | 3.31E-20 | Tcea1 | transcription | Nucleus |
| ENSMUSG00000015932 | 6.08E-21 | 3.33E-20 | Dstn | destrin, actin | Cytoplasm |
| ENSMUSG00000033712 | 6.12E-21 | 3.35E-20 | Ccar2 | cell cycle anc | Cytoplasm |
| ENSMUSG00000046598 | 6.51E-21 | 3.55E-20 | Bdh1 | 3-hydroxybu | Cytoplasm |
| ENSMUSG00000053841 | 7.17E-21 | 3.92E-20 | Txlna | taxilin alpha | Extracellular S |
| ENSMUSG00000048040 | 7.51E-21 | 4.10E-20 | Arxes2 | adipocyte-re | Cytoplasm |
| ENSMUSG00000040667 | 7.53E-21 | 4.10E-20 | Nup88 | nucleoporin 8 | Nucleus |
| ENSMUSG00000037236 | 8.03E-21 | 4.37E-20 | Matr3 | matrin 3 | Nucleus |
| ENSMUSG00000020250 | 8.44E-21 | 4.59E-20 | Txnrd1 | thioredoxin r | Cytoplasm |
| ENSMUSG00000026020 | 8.47E-21 | 4.61E-20 | Nop58 | NOP58 ribon | Nucleus |
| ENSMUSG00000008958 | 8.72E-21 | 4.74E-20 | Vps72 | vacuolar prot | Nucleus |
| ENSMUSG00000030042 | 8.93E-21 | 4.85E-20 | Pole4 | DNA polyme | Nucleus |
| ENSMUSG00000030647 | 8.98E-21 | 4.87E-20 | Ndufc2 | NADH:ubiqu | Cytoplasm |

| | | | | | |
|---|---|---|---|---|---|
| ENSMUSG00000023393 | 1.04E-20 | 5.65E-20 | Slc17a9 | solute carrier | Cytoplasm |
| ENSMUSG00000004865 | 1.07E-20 | 5.82E-20 | Srpk1 | SRSF protein | Nucleus |
| ENSMUSG00000058057 | 1.09E-20 | 5.90E-20 | Mettl7a3 | methyltransfe | Other |
| ENSMUSG00000051329 | 1.12E-20 | 6.06E-20 | Nup160 | nucleoporin 1 | Nucleus |
| ENSMUSG00000030346 | 1.14E-20 | 6.18E-20 | Rad51ap1 | RAD51 assoc | Nucleus |
| ENSMUSG00000022554 | 1.15E-20 | 6.24E-20 | Hgh1 | HGH1 homol | Other |
| ENSMUSG00000030980 | 1.16E-20 | 6.29E-20 | Knop1 | lysine rich nu | Other |
| ENSMUSG00000040945 | 1.17E-20 | 6.33E-20 | Rcc2 | regulator of c | Nucleus |
| ENSMUSG00000052488 | 1.24E-20 | 6.72E-20 | Cherp | calcium hom | Cytoplasm |
| ENSMUSG00000021929 | 1.29E-20 | 6.96E-20 | Kpna3 | karyopherin s | Nucleus |
| ENSMUSG00000095567 | 1.38E-20 | 7.42E-20 | Noc2l | NOC2 like nu | Nucleus |
| ENSMUSG00000043445 | 1.47E-20 | 7.92E-20 | Pgp | phosphoglyc | Cytoplasm |
| ENSMUSG00000074682 | 1.48E-20 | 7.98E-20 | Zcchc3 | zinc finger C | Other |
| ENSMUSG00000022634 | 1.51E-20 | 8.11E-20 | Yaf2 | YY1 associate | Nucleus |
| ENSMUSG00000033427 | 1.57E-20 | 8.43E-20 | Upb1 | beta-ureidop | Cytoplasm |
| ENSMUSG00000028788 | 1.58E-20 | 8.48E-20 | Ptp4a2 | protein tyros | Cytoplasm |
| ENSMUSG00000063694 | 1.62E-20 | 8.71E-20 | Cycs | cytochrome c | Cytoplasm |
| ENSMUSG00000033917 | 1.80E-20 | 9.67E-20 | Gde1 | glycerophosp | Plasma Mem |
| ENSMUSG00000026773 | 1.86E-20 | 9.99E-20 | Pfkfb3 | 6-phosphofr | Cytoplasm |
| ENSMUSG00000001436 | 1.98E-20 | 1.06E-19 | Slc19a1 | solute carrier | Plasma Mem |
| ENSMUSG00000022425 | 2.00E-20 | 1.07E-19 | Enpp2 | ectonucleotic | Plasma Mem |
| ENSMUSG00000058704 | 2.23E-20 | 1.20E-19 | Memo1 | mediator of c | Cytoplasm |
| ENSMUSG00000015092 | 2.26E-20 | 1.21E-19 | Edf1 | endothelial d | Nucleus |
| ENSMUSG00000021975 | 2.33E-20 | 1.24E-19 | Ints9 | integrator co | Nucleus |
| ENSMUSG00000062270 | 2.48E-20 | 1.32E-19 | Morf4l1 | mortality fact | Nucleus |
| ENSMUSG00000021494 | 2.57E-20 | 1.37E-19 | Ddx41 | DEAD-box he | Nucleus |
| ENSMUSG00000042814 | 2.63E-20 | 1.40E-19 | Mcts2 | malignant T c | Other |
| ENSMUSG00000032078 | 2.66E-20 | 1.42E-19 | Zpr1 | ZPR1 zinc fin | Nucleus |
| ENSMUSG00000024371 | 2.70E-20 | 1.44E-19 | C2 | complement | Extracellular S |
| ENSMUSG00000018899 | 2.77E-20 | 1.48E-19 | Irf1 | interferon reg | Nucleus |
| ENSMUSG00000022235 | 2.88E-20 | 1.53E-19 | Cmbl | carboxymeth | Cytoplasm |
| ENSMUSG00000023988 | 2.95E-20 | 1.57E-19 | Bysl | bystin like | Nucleus |
| ENSMUSG00000005656 | 3.00E-20 | 1.59E-19 | Snx6 | sorting nexin | Cytoplasm |
| ENSMUSG00000045752 | 3.03E-20 | 1.61E-19 | Tssc4 | tumor suppre | Other |
| ENSMUSG00000017677 | 3.08E-20 | 1.63E-19 | Wsb1 | WD repeat ar | Cytoplasm |
| ENSMUSG00000024667 | 3.23E-20 | 1.72E-19 | Tmem216 | transmembra | Extracellular S |
| ENSMUSG00000007656 | 3.29E-20 | 1.74E-19 | Arpp19 | cAMP regula | Cytoplasm |
| ENSMUSG00000028948 | 3.32E-20 | 1.76E-19 | Nol9 | nucleolar pro | Nucleus |
| ENSMUSG00000001891 | 3.42E-20 | 1.81E-19 | Ugp2 | UDP-glucose | Cytoplasm |
| ENSMUSG00000042487 | 3.51E-20 | 1.86E-19 | Leo1 | LEO1 homolo | Nucleus |
| ENSMUSG00000023827 | 3.54E-20 | 1.87E-19 | Agpat4 | 1-acylglycero | Cytoplasm |
| ENSMUSG00000054309 | 3.60E-20 | 1.90E-19 | Cpsf3 | cleavage and | Nucleus |
| ENSMUSG00000030934 | 3.70E-20 | 1.96E-19 | Oat | ornithine am | Cytoplasm |
| ENSMUSG00000023147 | 3.77E-20 | 1.99E-19 | Wrb | tryptophan ri | Extracellular S |
| ENSMUSG00000000088 | 3.87E-20 | 2.04E-19 | Cox5a | cytochrome c | Cytoplasm |
| ENSMUSG00000000339 | 4.21E-20 | 2.22E-19 | Rtca | RNA 3'-term | Nucleus |
| ENSMUSG00000003309 | 4.23E-20 | 2.23E-19 | Ap1m2 | adaptor relat | Cytoplasm |
| ENSMUSG00000024732 | 4.33E-20 | 2.28E-19 | Ccdc86 | coiled-coil do | Nucleus |
| ENSMUSG00000041278 | 4.41E-20 | 2.32E-19 | Ttc1 | tetratricopep | Cytoplasm |
| ENSMUSG00000094870 | 4.64E-20 | 2.44E-19 | Zfp131 | zinc finger pr | Nucleus |
| ENSMUSG00000040220 | 4.75E-20 | 2.50E-19 | Gas8 | growth arrest | Cytoplasm |
| ENSMUSG00000021288 | 4.78E-20 | 2.51E-19 | Klc1 | kinesin light c | Cytoplasm |
| ENSMUSG00000030204 | 4.86E-20 | 2.55E-19 | Ddx47 | DEAD-box he | Nucleus |
| ENSMUSG00000010097 | 4.91E-20 | 2.57E-19 | Nxf1 | nuclear RNA | Nucleus |
| ENSMUSG00000039220 | 4.97E-20 | 2.61E-19 | Ppp1r10 | protein phos | Nucleus |
| ENSMUSG00000028140 | 5.03E-20 | 2.63E-19 | Mrpl9 | mitochondria | Cytoplasm |
| ENSMUSG00000021840 | 5.08E-20 | 2.66E-19 | Mapk1ip1l | mitogen-acti | Nucleus |
| ENSMUSG00000001918 | 5.15E-20 | 2.69E-19 | Slc1a5 | solute carrier | Plasma Mem |

| | | | | | |
|---|---|---|---|---|---|
| ENSMUSG00000033965 | 5.83E-20 | 3.05E-19 | Slc16a2 | solute carrier | Plasma Mem |
| ENSMUSG00000063800 | 5.88E-20 | 3.07E-19 | Prpf38a | pre-mRNA p | Nucleus |
| ENSMUSG00000037169 | 6.12E-20 | 3.19E-19 | Mycn | MYCN proto- | Nucleus |
| ENSMUSG00000030103 | 6.23E-20 | 3.25E-19 | Bhlhe40 | basic helix-lo | Nucleus |
| ENSMUSG00000021639 | 6.29E-20 | 3.28E-19 | Gtf2h2 | general trans | Nucleus |
| ENSMUSG00000037254 | 6.45E-20 | 3.36E-19 | Itih2 | inter-alpha-t | Extracellular S |
| ENSMUSG00000017221 | 6.59E-20 | 3.43E-19 | Psmd3 | proteasome 2 | Cytoplasm |
| ENSMUSG00000032330 | 6.86E-20 | 3.57E-19 | Cox7a2 | cytochrome c | Cytoplasm |
| ENSMUSG00000042709 | 7.01E-20 | 3.64E-19 | Atpaf2 | ATP synthase | Cytoplasm |
| ENSMUSG00000027944 | 7.22E-20 | 3.75E-19 | Hax1 | HCLS1 associ | Cytoplasm |
| ENSMUSG00000057236 | 8.13E-20 | 4.22E-19 | Rbbp4 | RB binding p | Nucleus |
| ENSMUSG00000025035 | 8.25E-20 | 4.28E-19 | Arl3 | ADP ribosyla | Cytoplasm |
| ENSMUSG00000062300 | 9.26E-20 | 4.80E-19 | Pvrl2 | nectin cell ad | Plasma Mem |
| ENSMUSG00000022674 | 9.39E-20 | 4.87E-19 | Ube2v2 | ubiquitin con | Cytoplasm |
| ENSMUSG00000031985 | 1.01E-19 | 5.22E-19 | Gnpat | glyceroneph | Cytoplasm |
| ENSMUSG00000024736 | 1.02E-19 | 5.26E-19 | Tmem132a | transmembra | Cytoplasm |
| ENSMUSG00000029551 | 1.05E-19 | 5.43E-19 | Psmg3 | proteasome a | Other |
| ENSMUSG00000052298 | 1.06E-19 | 5.45E-19 | Cdc42se2 | CDC42 small | Plasma Mem |
| ENSMUSG00000048782 | 1.09E-19 | 5.61E-19 | Insc | INSC, spindle | Plasma Mem |
| ENSMUSG00000026917 | 1.10E-19 | 5.70E-19 | Wdr5 | WD repeat d | Nucleus |
| ENSMUSG00000019312 | 1.18E-19 | 6.08E-19 | Grb7 | growth facto | Plasma Mem |
| ENSMUSG00000022385 | 1.18E-19 | 6.10E-19 | Gtse1 | G2 and S-ph | Cytoplasm |
| ENSMUSG00000022816 | 1.20E-19 | 6.18E-19 | Fstl1 | follistatin like | Extracellular S |
| ENSMUSG00000031146 | 1.32E-19 | 6.78E-19 | Plp2 | proteolipid p | Cytoplasm |
| ENSMUSG00000039450 | 1.39E-19 | 7.18E-19 | Dcxr | dicarbonyl ar | Cytoplasm |
| ENSMUSG00000021951 | 1.45E-19 | 7.43E-19 | N6amt2 | EEF1A lysine | Cytoplasm |
| ENSMUSG00000038704 | 1.49E-19 | 7.64E-19 | Aspdh | aspartate deh | Other |
| ENSMUSG00000078941 | 1.53E-19 | 7.87E-19 | Ak6 | adenylate kir | Nucleus |
| ENSMUSG00000038335 | 1.63E-19 | 8.38E-19 | Tsr1 | TSR1, ribosor | Nucleus |
| ENSMUSG00000042426 | 1.64E-19 | 8.39E-19 | Dhx29 | DExH-box he | Cytoplasm |
| ENSMUSG00000067194 | 1.65E-19 | 8.47E-19 | Eif1ax | eukaryotic tra | Other |
| ENSMUSG00000025395 | 1.71E-19 | 8.75E-19 | Prim1 | DNA primase | Nucleus |
| ENSMUSG00000040147 | 1.71E-19 | 8.76E-19 | Maob | monoamine o | Cytoplasm |
| ENSMUSG00000003808 | 1.75E-19 | 8.96E-19 | Farsa | phenylalanyl- | Cytoplasm |
| ENSMUSG00000029273 | 1.76E-19 | 9.01E-19 | Sult1d1 | sulfotransfera | Cytoplasm |
| ENSMUSG00000068922 | 1.77E-19 | 9.06E-19 | Msto1 | misato 1, mit | Cytoplasm |
| ENSMUSG00000027108 | 1.92E-19 | 9.82E-19 | Ola1 | Obg like ATP | Cytoplasm |
| ENSMUSG00000026248 | 1.99E-19 | 1.01E-18 | Mrpl44 | mitochondria | Cytoplasm |
| ENSMUSG00000024454 | 2.01E-19 | 1.02E-18 | Hdac3 | histone deac | Nucleus |
| ENSMUSG00000058454 | 2.11E-19 | 1.07E-18 | Dhcr7 | 7-dehydroch | Cytoplasm |
| ENSMUSG00000054200 | 2.20E-19 | 1.12E-18 | Ffar4 | free fatty acid | Plasma Mem |
| ENSMUSG00000028869 | 2.23E-19 | 1.13E-18 | Gnl2 | G protein nu | Nucleus |
| ENSMUSG00000071547 | 2.68E-19 | 1.37E-18 | Nt5dc2 | 5'-nucleotida | Cytoplasm |
| ENSMUSG00000001847 | 2.75E-19 | 1.40E-18 | Rac1 | Rac family sm | Plasma Mem |
| ENSMUSG00000054693 | 2.76E-19 | 1.40E-18 | Adam10 | ADAM metal | Plasma Mem |
| ENSMUSG00000058443 | 2.78E-19 | 1.41E-18 | Rpl10-ps3 | ribosomal pr | Other |
| ENSMUSG00000051232 | 2.80E-19 | 1.42E-18 | Tmem199 | transmembra | Cytoplasm |
| ENSMUSG00000046573 | 2.87E-19 | 1.45E-18 | Lyrm4 | LYR motif co | Cytoplasm |
| ENSMUSG00000024317 | 2.87E-19 | 1.45E-18 | Rnf138 | ring finger pr | Nucleus |
| ENSMUSG00000004038 | 3.10E-19 | 1.57E-18 | Gstm3 | glutathione S | Cytoplasm |
| ENSMUSG00000063232 | 3.14E-19 | 1.59E-18 | Serpina11 | serpin family | Extracellular S |
| ENSMUSG00000038527 | 3.16E-19 | 1.60E-18 | C1rl | complement | Extracellular S |
| ENSMUSG00000035775 | 3.21E-19 | 1.62E-18 | Krt20 | keratin 20 | Cytoplasm |
| ENSMUSG00000067212 | 3.24E-19 | 1.64E-18 | H2-T23 | major histoco | Plasma Mem |
| ENSMUSG00000032018 | 3.33E-19 | 1.68E-18 | Sc5d | sterol-C5-de | Cytoplasm |
| ENSMUSG00000054648 | 3.64E-19 | 1.84E-18 | Zfp869 | zinc finger pr | Nucleus |
| ENSMUSG00000025270 | 3.79E-19 | 1.91E-18 | Alas2 | 5'-aminolevu | Cytoplasm |
| ENSMUSG00000027905 | 3.90E-19 | 1.97E-18 | Ddx20 | DEAD-box h | Nucleus |

| | | | | | |
|---|---|---|---|---|---|
| ENSMUSG00000019362 | 4.06E-19 | 2.04E-18 | D8Ertd738e | chromosome | Nucleus |
| ENSMUSG00000068085 | 4.31E-19 | 2.17E-18 | Cyp2d11 | cytochrome I | Cytoplasm |
| ENSMUSG00000032381 | 4.34E-19 | 2.18E-18 | Fam96a | cytosolic iron | Extracellular S |
| ENSMUSG00000023067 | 4.41E-19 | 2.22E-18 | Cdkn1a | cyclin depend | Nucleus |
| ENSMUSG00000091896 | 4.42E-19 | 2.22E-18 | Ube2d2a | ubiquitin cor | Cytoplasm |
| ENSMUSG00000021901 | 4.45E-19 | 2.24E-18 | Bap1 | BRCA1 assoc | Nucleus |
| ENSMUSG00000024683 | 4.57E-19 | 2.29E-18 | Mrpl16 | mitochondria | Cytoplasm |
| ENSMUSG00000030108 | 4.68E-19 | 2.35E-18 | Slc6a13 | solute carrier | Plasma Mem |
| ENSMUSG00000022671 | 4.74E-19 | 2.38E-18 | Mzt2 | mitotic spind | Cytoplasm |
| ENSMUSG00000010651 | 4.80E-19 | 2.41E-18 | Acaa1b | acetyl-Coenz | Cytoplasm |
| ENSMUSG00000040263 | 4.81E-19 | 2.41E-18 | Klhdc4 | kelch domair | Other |
| ENSMUSG00000038379 | 5.22E-19 | 2.61E-18 | Ttk | TTK protein k | Nucleus |
| ENSMUSG00000030421 | 5.23E-19 | 2.62E-18 | Uri1 | URI1, prefold | Nucleus |
| ENSMUSG00000051934 | 5.67E-19 | 2.83E-18 | Spats2 | spermatoger | Cytoplasm |
| ENSMUSG00000031078 | 5.68E-19 | 2.84E-18 | Cttn | cortactin | Plasma Mem |
| ENSMUSG00000026180 | 5.89E-19 | 2.94E-18 | Cxcr2 | C-X-C motif | Plasma Mem |
| ENSMUSG00000063273 | 5.96E-19 | 2.97E-18 | Naa15 | N(alpha)-ace | Nucleus |
| ENSMUSG00000014195 | 6.20E-19 | 3.09E-18 | Dnajc7 | DnaJ heat sh | Cytoplasm |
| ENSMUSG00000008036 | 6.42E-19 | 3.20E-18 | Ap2s1 | adaptor relat | Cytoplasm |
| ENSMUSG00000061374 | 6.46E-19 | 3.22E-18 | Fiz1 | FLT3 interact | Nucleus |
| ENSMUSG00000030605 | 6.62E-19 | 3.30E-18 | Mfge8 | milk fat glob | Extracellular S |
| ENSMUSG00000027215 | 6.69E-19 | 3.33E-18 | Cd82 | CD82 molecu | Plasma Mem |
| ENSMUSG00000022978 | 6.70E-19 | 3.33E-18 | Mis18a | MIS18 kineto | Nucleus |
| ENSMUSG00000015759 | 6.73E-19 | 3.34E-18 | Cnih1 | cornichon fa | Plasma Mem |
| ENSMUSG00000020476 | 6.73E-19 | 3.34E-18 | Dbnl | drebrin like | Cytoplasm |
| ENSMUSG00000022889 | 7.06E-19 | 3.50E-18 | Mrpl39 | mitochondria | Cytoplasm |
| ENSMUSG00000029669 | 7.39E-19 | 3.67E-18 | Tspan12 | tetraspanin 1 | Plasma Mem |
| ENSMUSG00000022744 | 7.41E-19 | 3.67E-18 | Cldnd1 | claudin doma | Plasma Mem |
| ENSMUSG00000003200 | 7.42E-19 | 3.67E-18 | Sh3gl1 | SH3 domain | Cytoplasm |
| ENSMUSG00000032715 | 7.98E-19 | 3.95E-18 | Trib3 | tribbles pseu | Nucleus |
| ENSMUSG00000020140 | 8.10E-19 | 4.01E-18 | Lgr5 | leucine rich r | Plasma Mem |
| ENSMUSG00000022031 | 8.16E-19 | 4.04E-18 | Elp3 | elongator ac | Nucleus |
| ENSMUSG00000066629 | 8.18E-19 | 4.05E-18 | Rpl36-ps3 | ribosomal pr | Other |
| ENSMUSG00000042102 | 8.26E-19 | 4.08E-18 | Dmgdh | dimethylglyc | Cytoplasm |
| ENSMUSG00000020911 | 8.37E-19 | 4.13E-18 | Krt19 | keratin 19 | Cytoplasm |
| ENSMUSG00000015804 | 8.41E-19 | 4.15E-18 | Med28 | mediator cor | Nucleus |
| ENSMUSG00000068823 | 8.44E-19 | 4.16E-18 | Csde1 | cold shock d | Cytoplasm |
| ENSMUSG00000041747 | 8.45E-19 | 4.17E-18 | Utp15 | UTP15, small | Nucleus |
| ENSMUSG00000024360 | 8.54E-19 | 4.21E-18 | Etf1 | eukaryotic tra | Cytoplasm |
| ENSMUSG00000071078 | 8.66E-19 | 4.26E-18 | Nr2c2ap | nuclear recep | Nucleus |
| ENSMUSG00000026249 | 8.91E-19 | 4.38E-18 | Serpine2 | serpin family | Extracellular S |
| ENSMUSG00000023940 | 9.16E-19 | 4.50E-18 | Sgol1 | shugoshin 1 | Nucleus |
| ENSMUSG00000030120 | 9.17E-19 | 4.51E-18 | Mlf2 | myeloid leuk | Nucleus |
| ENSMUSG00000016427 | 9.58E-19 | 4.71E-18 | Ndufa1 | NADH:ubiqu | Cytoplasm |
| ENSMUSG00000030138 | 9.69E-19 | 4.75E-18 | Bms1 | BMS1, riboso | Nucleus |
| ENSMUSG00000044201 | 9.76E-19 | 4.79E-18 | Cdc25c | cell division c | Nucleus |
| ENSMUSG00000028669 | 9.86E-19 | 4.83E-18 | Pithd1 | PITH domain | Nucleus |
| ENSMUSG00000039509 | 1.09E-18 | 5.33E-18 | Nup133 | nucleoporin 1 | Nucleus |
| ENSMUSG00000022722 | 1.10E-18 | 5.37E-18 | Arl6 | ADP ribosyla | Cytoplasm |
| ENSMUSG00000020124 | 1.10E-18 | 5.37E-18 | Usp15 | ubiquitin spe | Cytoplasm |
| ENSMUSG00000032002 | 1.11E-18 | 5.41E-18 | Dcun1d5 | defective in c | Other |
| ENSMUSG00000023277 | 1.11E-18 | 5.43E-18 | Twf2 | twinfilin actin | Cytoplasm |
| ENSMUSG00000027353 | 1.11E-18 | 5.44E-18 | Mcm8 | minichromos | Nucleus |
| ENSMUSG00000048583 | 1.12E-18 | 5.45E-18 | Igf2 | insulin like gr | Extracellular S |
| ENSMUSG00000063785 | 1.14E-18 | 5.56E-18 | Utp14a | UTP14A, sma | Nucleus |
| ENSMUSG00000034263 | 1.20E-18 | 5.83E-18 | Vwa9 | integrator co | Cytoplasm |
| ENSMUSG00000022051 | 1.23E-18 | 6.00E-18 | Bnip3l | BCL2 interact | Cytoplasm |
| ENSMUSG00000003500 | 1.23E-18 | 6.00E-18 | Impdh1 | inosine monc | Cytoplasm |

| | | | | | |
|---|---|---|---|---|---|
| ENSMUSG00000002728 | 1.25E-18 | 6.07E-18 | Naa20 | N(alpha)-ace | Cytoplasm |
| ENSMUSG00000002658 | 1.27E-18 | 6.20E-18 | Gtf2f1 | general trans | Nucleus |
| ENSMUSG00000066232 | 1.29E-18 | 6.26E-18 | Ipo7 | importin 7 | Nucleus |
| ENSMUSG00000031796 | 1.29E-18 | 6.27E-18 | Cfap20 | cilia and flag | Cytoplasm |
| ENSMUSG00000039063 | 1.36E-18 | 6.58E-18 | Echdc3 | enoyl-CoA h | Cytoplasm |
| ENSMUSG00000079478 | 1.36E-18 | 6.58E-18 | Sssca1 | Sjogren synd | Other |
| ENSMUSG00000037608 | 1.37E-18 | 6.67E-18 | Bclaf1 | BCL2 associa | Nucleus |
| ENSMUSG00000019845 | 1.39E-18 | 6.74E-18 | Tube1 | tubulin epsilo | Cytoplasm |
| ENSMUSG00000062797 | 1.40E-18 | 6.76E-18 | I7Rn6 | Hikeshi, heat | Cytoplasm |
| ENSMUSG00000042111 | 1.41E-18 | 6.81E-18 | Ccdc115 | coiled-coil d | Cytoplasm |
| ENSMUSG00000004897 | 1.42E-18 | 6.86E-18 | Hdgf | heparin bind | Extracellular S |
| ENSMUSG00000003437 | 1.58E-18 | 7.62E-18 | Paf1 | PAF1 homolo | Nucleus |
| ENSMUSG00000025982 | 1.58E-18 | 7.64E-18 | Sf3b1 | splicing facto | Nucleus |
| ENSMUSG00000037013 | 1.62E-18 | 7.82E-18 | Ss18 | SS18, nBAF c | Nucleus |
| ENSMUSG00000024037 | 1.71E-18 | 8.27E-18 | Wdr4 | WD repeat d | Nucleus |
| ENSMUSG00000022812 | 1.85E-18 | 8.90E-18 | Gsk3b | glycogen syn | Nucleus |
| ENSMUSG00000042712 | 1.89E-18 | 9.13E-18 | Wbp5 | transcription | Other |
| ENSMUSG00000047879 | 1.90E-18 | 9.16E-18 | Usp14 | ubiquitin spe | Cytoplasm |
| ENSMUSG00000036030 | 1.90E-18 | 9.17E-18 | Prtg | protogenin | Extracellular S |
| ENSMUSG00000029759 | 1.92E-18 | 9.24E-18 | Pon3 | paraoxonase | Extracellular S |
| ENSMUSG00000028980 | 1.92E-18 | 9.25E-18 | H6pd | hexose-6-ph | Cytoplasm |
| ENSMUSG00000005397 | 1.93E-18 | 9.26E-18 | Nid1 | nidogen 1 | Extracellular S |
| ENSMUSG00000044155 | 1.94E-18 | 9.32E-18 | Lsm8 | LSM8 homol | Nucleus |
| ENSMUSG00000029499 | 1.99E-18 | 9.54E-18 | Pxmp2 | peroxisomal | Cytoplasm |
| ENSMUSG00000022536 | 2.08E-18 | 9.99E-18 | Glyr1 | glyoxylate re | Nucleus |
| ENSMUSG00000050043 | 2.08E-18 | 9.99E-18 | Tmx2 | thioredoxin r | Other |
| ENSMUSG00000025208 | 2.11E-18 | 1.01E-17 | Mrpl43 | mitochondria | Cytoplasm |
| ENSMUSG00000026981 | 2.16E-18 | 1.03E-17 | Il1rn | interleukin 1 | Extracellular S |
| ENSMUSG00000056851 | 2.27E-18 | 1.09E-17 | Pcbp2 | poly(rC) bind | Nucleus |
| ENSMUSG00000026622 | 2.40E-18 | 1.15E-17 | Nek2 | NIMA relatec | Cytoplasm |
| ENSMUSG00000066026 | 2.40E-18 | 1.15E-17 | Dhrs3 | dehydrogena | Cytoplasm |
| ENSMUSG00000024764 | 2.45E-18 | 1.17E-17 | Naa40 | N(alpha)-ace | Other |
| ENSMUSG00000022160 | 2.50E-18 | 1.19E-17 | Mettl3 | methyltransfe | Nucleus |
| ENSMUSG00000025024 | 2.51E-18 | 1.20E-17 | Smndc1 | survival moto | Nucleus |
| ENSMUSG00000011254 | 2.58E-18 | 1.23E-17 | Thg1l | tRNA-histidir | Cytoplasm |
| ENSMUSG00000044573 | 2.62E-18 | 1.25E-17 | Acp1 | acid phospha | Cytoplasm |
| ENSMUSG00000059878 | 2.63E-18 | 1.25E-17 | Zfp422 | zinc finger pr | Nucleus |
| ENSMUSG00000041889 | 2.66E-18 | 1.27E-17 | Shisa4 | shisa family r | Other |
| ENSMUSG00000089678 | 2.70E-18 | 1.28E-17 | Agxt2 | alanine--gly | Cytoplasm |
| ENSMUSG00000034928 | 2.71E-18 | 1.29E-17 | Rnf44 | ring finger pr | Other |
| ENSMUSG00000022142 | 2.76E-18 | 1.31E-17 | Nup155 | nucleoporin 1 | Nucleus |
| ENSMUSG00000078652 | 2.86E-18 | 1.36E-17 | Psme3 | proteasome a | Cytoplasm |
| ENSMUSG00000024232 | 2.99E-18 | 1.42E-17 | Bambi | BMP and acti | Plasma Mem |
| ENSMUSG00000026520 | 3.08E-18 | 1.46E-17 | Pycr2 | pyrroline-5-c | Cytoplasm |
| ENSMUSG00000028410 | 3.16E-18 | 1.50E-17 | Dnaja1 | DnaJ heat sh | Nucleus |
| ENSMUSG00000090000 | 3.35E-18 | 1.59E-17 | Ier3ip1 | immediate ea | Cytoplasm |
| ENSMUSG00000080268 | 3.40E-18 | 1.61E-17 | Brms1 | BRMS1, trans | Nucleus |
| ENSMUSG00000071014 | 3.41E-18 | 1.62E-17 | Ndufb6 | NADH:ubiqu | Cytoplasm |
| ENSMUSG00000051518 | 3.42E-18 | 1.62E-17 | Rps19bp1 | ribosomal pr | Nucleus |
| ENSMUSG00000038759 | 3.57E-18 | 1.69E-17 | Nup205 | nucleoporin 2 | Nucleus |
| ENSMUSG00000022191 | 3.58E-18 | 1.69E-17 | Drosha | drosha ribon | Nucleus |
| ENSMUSG00000027185 | 3.97E-18 | 1.88E-17 | Nat10 | N-acetyltrans | Nucleus |
| ENSMUSG00000048922 | 4.16E-18 | 1.96E-17 | Cdca2 | cell division c | Nucleus |
| ENSMUSG00000024790 | 4.17E-18 | 1.97E-17 | Sac3d1 | SAC3 domair | Cytoplasm |
| ENSMUSG00000022575 | 4.30E-18 | 2.03E-17 | Gsdmd | gasdermin D | Extracellular S |
| ENSMUSG00000021238 | 4.37E-18 | 2.06E-17 | Aldh6a1 | aldehyde deh | Cytoplasm |
| ENSMUSG00000061028 | 4.80E-18 | 2.26E-17 | Clasrp | CLK4-associa | Nucleus |
| ENSMUSG00000031669 | 4.92E-18 | 2.31E-17 | Gins3 | GINS comple | Nucleus |

| | | | | | |
|---|---|---|---|---|---|
| ENSMUSG00000032586 | 4.92E-18 | 2.31E-17 | Traip | TRAF interact | Cytoplasm |
| ENSMUSG00000024414 | 5.06E-18 | 2.38E-17 | Mrpl27 | mitochondria | Cytoplasm |
| ENSMUSG00000025986 | 5.23E-18 | 2.46E-17 | Slc39a10 | solute carrier | Extracellular S |
| ENSMUSG00000071359 | 5.56E-18 | 2.61E-17 | Tbpl1 | TATA-box bi | Nucleus |
| ENSMUSG00000037474 | 5.57E-18 | 2.61E-17 | Dtl | denticleless E | Nucleus |
| ENSMUSG00000068220 | 5.57E-18 | 2.61E-17 | Lgals1 | galectin 1 | Extracellular S |
| ENSMUSG00000021891 | 5.58E-18 | 2.62E-17 | Mettl6 | methyltransfe | Other |
| ENSMUSG00000027620 | 5.63E-18 | 2.64E-17 | Rbm39 | RNA binding | Nucleus |
| ENSMUSG00000018340 | 5.73E-18 | 2.68E-17 | Anxa6 | annexin A6 | Plasma Mem |
| ENSMUSG00000066037 | 5.79E-18 | 2.71E-17 | Hnrnpr | heterogeneo | Nucleus |
| ENSMUSG00000018379 | 5.81E-18 | 2.72E-17 | Srsf1 | serine and ar | Nucleus |
| ENSMUSG00000021500 | 5.83E-18 | 2.72E-17 | Ddx46 | DEAD-box h | Nucleus |
| ENSMUSG00000009575 | 6.17E-18 | 2.88E-17 | Cbx5 | chromobox 5 | Nucleus |
| ENSMUSG00000042396 | 6.24E-18 | 2.91E-17 | Rbm7 | RNA binding | Nucleus |
| ENSMUSG00000026683 | 6.24E-18 | 2.91E-17 | Nuf2 | NUF2, NDC80 | Nucleus |
| ENSMUSG00000078768 | 6.28E-18 | 2.93E-17 | Zfp566 | zinc finger pr | Nucleus |
| ENSMUSG00000058126 | 6.62E-18 | 3.08E-17 | Tpm3-rs7 | tropomyosin | Other |
| ENSMUSG00000024863 | 6.68E-18 | 3.11E-17 | Mbl2 | mannose bin | Extracellular S |
| ENSMUSG00000033953 | 6.68E-18 | 3.11E-17 | Ppp3r1 | protein phos | Cytoplasm |
| ENSMUSG00000051396 | 6.83E-18 | 3.18E-17 | Hspa14 | heat shock p | Cytoplasm |
| ENSMUSG00000028318 | 6.99E-18 | 3.25E-17 | Polr1e | RNA polymer | Nucleus |
| ENSMUSG00000022403 | 7.11E-18 | 3.30E-17 | St13 | ST13, Hsp70 | Cytoplasm |
| ENSMUSG00000020925 | 7.20E-18 | 3.34E-17 | Ccdc43 | coiled-coil d | Cytoplasm |
| ENSMUSG00000030060 | 7.35E-18 | 3.41E-17 | Hmces | 5-hydroxyme | Other |
| ENSMUSG00000019762 | 7.37E-18 | 3.42E-17 | Iyd | iodotyrosine | Plasma Mem |
| ENSMUSG00000026851 | 7.49E-18 | 3.47E-17 | BC005624 | chromosome | Other |
| ENSMUSG00000035232 | 7.68E-18 | 3.56E-17 | Pdk3 | pyruvate deh | Cytoplasm |
| ENSMUSG00000032349 | 7.94E-18 | 3.67E-17 | Elovl5 | ELOVL fatty a | Cytoplasm |
| ENSMUSG00000033533 | 8.44E-18 | 3.91E-17 | Acsm1 | acyl-CoA syr | Cytoplasm |
| ENSMUSG00000031776 | 8.60E-18 | 3.98E-17 | Arl2bp | ADP ribosyla | Cytoplasm |
| ENSMUSG00000004880 | 8.89E-18 | 4.11E-17 | Lbr | lamin B recep | Nucleus |
| ENSMUSG00000070493 | 8.94E-18 | 4.13E-17 | Chchd2 | coiled-coil-h | Cytoplasm |
| ENSMUSG00000021957 | 9.05E-18 | 4.18E-17 | Tkt | transketolase | Cytoplasm |
| ENSMUSG00000040688 | 9.15E-18 | 4.22E-17 | Tbl3 | transducin be | Cytoplasm |
| ENSMUSG00000039640 | 9.30E-18 | 4.29E-17 | Mrpl12 | mitochondria | Cytoplasm |
| ENSMUSG00000038646 | 9.42E-18 | 4.34E-17 | Fam103a1 | RNA guanine | Nucleus |
| ENSMUSG00000027787 | 9.73E-18 | 4.48E-17 | Nmd3 | NMD3 riboso | Nucleus |
| ENSMUSG00000026669 | 1.06E-17 | 4.87E-17 | Mcm10 | minichromos | Nucleus |
| ENSMUSG00000030061 | 1.07E-17 | 4.92E-17 | Uba3 | ubiquitin like | Cytoplasm |
| ENSMUSG00000040808 | 1.09E-17 | 4.99E-17 | S100g | S100 calcium | Cytoplasm |
| ENSMUSG00000052533 | 1.13E-17 | 5.19E-17 | Nup188 | nucleoporin 1 | Nucleus |
| ENSMUSG00000026036 | 1.15E-17 | 5.26E-17 | Nif3l1 | NGG1 interac | Cytoplasm |
| ENSMUSG00000019189 | 1.16E-17 | 5.34E-17 | Rnf145 | ring finger pr | Other |
| ENSMUSG00000045007 | 1.19E-17 | 5.46E-17 | Tubg2 | tubulin gamr | Cytoplasm |
| ENSMUSG00000024217 | 1.22E-17 | 5.58E-17 | Snrpc | U1 small nuc | Nucleus |
| ENSMUSG00000027160 | 1.23E-17 | 5.65E-17 | Ccdc34 | coiled-coil d | Other |
| ENSMUSG00000063576 | 1.24E-17 | 5.68E-17 | Klhdc3 | kelch domair | Cytoplasm |
| ENSMUSG00000021040 | 1.24E-17 | 5.68E-17 | Slirp | SRA stem-loo | Cytoplasm |
| ENSMUSG00000024231 | 1.32E-17 | 6.05E-17 | Cul2 | cullin 2 | Nucleus |
| ENSMUSG00000061666 | 1.33E-17 | 6.07E-17 | Gdpd1 | glycerophosp | Other |
| ENSMUSG00000022013 | 1.38E-17 | 6.32E-17 | Dnajc15 | DnaJ heat sh | Cytoplasm |
| ENSMUSG00000049091 | 1.39E-17 | 6.36E-17 | Sephs2 | selenophosp | Cytoplasm |
| ENSMUSG00000059897 | 1.41E-17 | 6.46E-17 | Zfp930 | expressed se | Other |
| ENSMUSG00000051220 | 1.45E-17 | 6.62E-17 | Ercc6l | ERCC excisio | Nucleus |
| ENSMUSG00000031633 | 1.48E-17 | 6.74E-17 | Slc25a4 | solute carrier | Cytoplasm |
| ENSMUSG00000020451 | 1.48E-17 | 6.77E-17 | Limk2 | LIM domain I | Cytoplasm |
| ENSMUSG00000023911 | 1.53E-17 | 6.97E-17 | Flywch2 | FLYWCH fam | Other |
| ENSMUSG00000020692 | 1.56E-17 | 7.10E-17 | Nle1 | notchless hoi | Nucleus |

| | | | | | |
|---|---|---|---|---|---|
| ENSMUSG00000003992 | 1.57E-17 | 7.13E-17 | Ssbp2 | single strand | Nucleus |
| ENSMUSG00000034120 | 1.63E-17 | 7.41E-17 | Srsf2 | serine and ar | Nucleus |
| ENSMUSG00000020232 | 1.63E-17 | 7.42E-17 | Hmg20b | high mobility | Nucleus |
| ENSMUSG00000008373 | 1.66E-17 | 7.53E-17 | Prpf31 | pre-mRNA p | Nucleus |
| ENSMUSG00000020840 | 1.68E-17 | 7.62E-17 | Blmh | bleomycin hy | Cytoplasm |
| ENSMUSG00000021431 | 1.74E-17 | 7.87E-17 | Snrnp48 | small nuclear | Nucleus |
| ENSMUSG00000024429 | 1.80E-17 | 8.17E-17 | Gnl1 | G protein nu | Nucleus |
| ENSMUSG00000042502 | 1.83E-17 | 8.28E-17 | Cd2bp2 | CD2 cytoplas | Cytoplasm |
| ENSMUSG00000021022 | 1.86E-17 | 8.42E-17 | Ppp2r3c | protein phos | Cytoplasm |
| ENSMUSG00000028745 | 1.87E-17 | 8.45E-17 | Capzb | capping actir | Cytoplasm |
| ENSMUSG00000025236 | 1.89E-17 | 8.54E-17 | Adpgk | ADP depend | Plasma Mem |
| ENSMUSG00000028617 | 1.91E-17 | 8.62E-17 | Lrrc42 | leucine rich r | Other |
| ENSMUSG00000029394 | 1.94E-17 | 8.77E-17 | Cdk2ap1 | cyclin depen | Nucleus |
| ENSMUSG00000026374 | 1.94E-17 | 8.77E-17 | Tsn | translin | Nucleus |
| ENSMUSG00000057541 | 2.00E-17 | 9.02E-17 | Pus7 | pseudouridyl | Nucleus |
| ENSMUSG00000031864 | 2.00E-17 | 9.03E-17 | Ints10 | integrator co | Nucleus |
| ENSMUSG00000036977 | 2.04E-17 | 9.17E-17 | Anapc10 | anaphase pro | Nucleus |
| ENSMUSG00000024604 | 2.08E-17 | 9.36E-17 | Rbm22 | RNA binding | Nucleus |
| ENSMUSG00000024048 | 2.09E-17 | 9.39E-17 | Myl12a | myosin light | Cytoplasm |
| ENSMUSG00000022538 | 2.12E-17 | 9.54E-17 | Lsg1 | large 60S suk | Cytoplasm |
| ENSMUSG00000024270 | 2.21E-17 | 9.93E-17 | Slc39a6 | solute carrier | Plasma Mem |
| ENSMUSG00000037993 | 2.23E-17 | 1.00E-16 | Dhx38 | DEAH-box h | Nucleus |
| ENSMUSG00000063953 | 2.24E-17 | 1.00E-16 | Amd2 | adenosylmet | Cytoplasm |
| ENSMUSG00000027533 | 2.27E-17 | 1.02E-16 | Fabp5 | fatty acid bin | Cytoplasm |
| ENSMUSG00000071660 | 2.34E-17 | 1.05E-16 | Ttc9c | tetratricopep | Other |
| ENSMUSG00000033998 | 2.35E-17 | 1.05E-16 | Kcnk1 | potassium tw | Plasma Mem |
| ENSMUSG00000032423 | 2.40E-17 | 1.07E-16 | Syncrip | synaptotagm | Nucleus |
| ENSMUSG00000037740 | 2.41E-17 | 1.08E-16 | Mrps26 | mitochondria | Cytoplasm |
| ENSMUSG00000015165 | 2.54E-17 | 1.14E-16 | Hnrnpl | heterogeneo | Nucleus |
| ENSMUSG00000037379 | 2.73E-17 | 1.22E-16 | Spon2 | spondin 2 | Extracellular S |
| ENSMUSG00000048076 | 2.77E-17 | 1.24E-16 | Arf1 | ADP ribosyla | Cytoplasm |
| ENSMUSG00000025232 | 2.80E-17 | 1.25E-16 | Hexa | hexosaminida | Cytoplasm |
| ENSMUSG00000020900 | 2.85E-17 | 1.27E-16 | Myh10 | myosin heavy | Cytoplasm |
| ENSMUSG00000066319 | 3.06E-17 | 1.37E-16 | Rtp3 | receptor tran | Cytoplasm |
| ENSMUSG00000000567 | 3.08E-17 | 1.37E-16 | Sox9 | SRY-box 9 | Nucleus |
| ENSMUSG00000003444 | 3.11E-17 | 1.38E-16 | Med29 | mediator cor | Nucleus |
| ENSMUSG00000021794 | 3.15E-17 | 1.40E-16 | Glud1 | glutamate de | Cytoplasm |
| ENSMUSG00000020079 | 3.15E-17 | 1.40E-16 | Supv3l1 | Suv3 like RNA | Cytoplasm |
| ENSMUSG00000010609 | 3.18E-17 | 1.42E-16 | Psen2 | presenilin 2 | Cytoplasm |
| ENSMUSG00000066148 | 3.20E-17 | 1.42E-16 | Prpf4 | pre-mRNA p | Nucleus |
| ENSMUSG00000055436 | 3.39E-17 | 1.51E-16 | Srsf11 | serine and ar | Nucleus |
| ENSMUSG00000007950 | 3.39E-17 | 1.51E-16 | Abhd8 | abhydrolase | Cytoplasm |
| ENSMUSG00000019777 | 3.44E-17 | 1.53E-16 | Hdac2 | histone deac | Nucleus |
| ENSMUSG00000037257 | 3.55E-17 | 1.57E-16 | Aagab | alpha and ga | Cytoplasm |
| ENSMUSG00000096210 | 3.60E-17 | 1.60E-16 | H1f0 | H1 histone fa | Nucleus |
| ENSMUSG00000020088 | 3.64E-17 | 1.61E-16 | Sar1a | secretion ass | Cytoplasm |
| ENSMUSG00000021111 | 3.66E-17 | 1.62E-16 | Papola | poly(A) polyr | Nucleus |
| ENSMUSG00000050213 | 3.70E-17 | 1.64E-16 | Snip1 | Smad nuclea | Nucleus |
| ENSMUSG00000030528 | 3.78E-17 | 1.67E-16 | Blm | Bloom syndro | Nucleus |
| ENSMUSG00000006740 | 3.86E-17 | 1.71E-16 | Kif5b | kinesin family | Cytoplasm |
| ENSMUSG00000022849 | 3.87E-17 | 1.71E-16 | Hspbap1 | HSPB1 assoc | Other |
| ENSMUSG00000031584 | 3.93E-17 | 1.74E-16 | Gsr | glutathione-c | Cytoplasm |
| ENSMUSG00000032898 | 3.95E-17 | 1.74E-16 | Fbxo21 | F-box protei | Extracellular S |
| ENSMUSG00000039601 | 3.99E-17 | 1.76E-16 | Rcan2 | regulator of c | Other |
| ENSMUSG00000034164 | 4.01E-17 | 1.77E-16 | Emid1 | EMI domain c | Extracellular S |
| ENSMUSG00000025577 | 4.05E-17 | 1.78E-16 | Cbx2 | chromobox 2 | Nucleus |
| ENSMUSG00000028857 | 4.12E-17 | 1.82E-16 | Tmem222 | transmembra | Other |
| ENSMUSG00000029623 | 4.31E-17 | 1.90E-16 | Pdap1 | PDGFA assoc | Cytoplasm |

| | | | | | |
|---|---|---|---|---|---|
| ENSMUSG00000040097 | 4.40E-17 | 1.94E-16 | Flywch1 | FLYWCH-typ | Nucleus |
| ENSMUSG00000045969 | 4.40E-17 | 1.94E-16 | Ing1 | inhibitor of g | Nucleus |
| ENSMUSG00000024579 | 4.41E-17 | 1.94E-16 | Pcyox1l | prenylcystein | Other |
| ENSMUSG00000008859 | 4.46E-17 | 1.96E-16 | Rala | RAS like prot | Cytoplasm |
| ENSMUSG00000042472 | 4.53E-17 | 1.99E-16 | Zfp410 | zinc finger pr | Nucleus |
| ENSMUSG00000038542 | 4.63E-17 | 2.03E-16 | Pcid2 | PCI domain c | Nucleus |
| ENSMUSG00000041261 | 4.81E-17 | 2.11E-16 | Car8 | carbonic anh | Cytoplasm |
| ENSMUSG00000020156 | 4.99E-17 | 2.19E-16 | Mum1 | melanoma as | Nucleus |
| ENSMUSG00000033596 | 5.49E-17 | 2.40E-16 | Rfwd3 | ring finger ar | Nucleus |
| ENSMUSG00000021963 | 5.55E-17 | 2.43E-16 | Sap18 | Sin3A associa | Nucleus |
| ENSMUSG00000004610 | 5.60E-17 | 2.45E-16 | Etfb | electron trans | Cytoplasm |
| ENSMUSG00000002804 | 5.78E-17 | 2.53E-16 | Nudt14 | nudix hydrola | Cytoplasm |
| ENSMUSG00000032254 | 5.80E-17 | 2.53E-16 | Kif23 | kinesin family | Cytoplasm |
| ENSMUSG00000024287 | 5.85E-17 | 2.56E-16 | Thoc1 | THO complex | Nucleus |
| ENSMUSG00000079343 | 5.91E-17 | 2.58E-16 | C1s2 | complement | Extracellular S |
| ENSMUSG00000026755 | 5.95E-17 | 2.60E-16 | Arpc5l | actin related | Cytoplasm |
| ENSMUSG00000001248 | 6.02E-17 | 2.63E-16 | Gramd1a | GRAM doma | Extracellular S |
| ENSMUSG00000005469 | 6.08E-17 | 2.65E-16 | Prkaca | protein kinas | Cytoplasm |
| ENSMUSG00000000787 | 6.21E-17 | 2.71E-16 | Ddx3x | DEAD-box h | Cytoplasm |
| ENSMUSG00000026032 | 6.23E-17 | 2.71E-16 | Ndufb3 | NADH:ubiqu | Cytoplasm |
| ENSMUSG00000038298 | 6.35E-17 | 2.77E-16 | Pdzk1 | PDZ domain | Plasma Mem |
| ENSMUSG00000063358 | 6.47E-17 | 2.82E-16 | Mapk1 | mitogen-acti | Cytoplasm |
| ENSMUSG00000018459 | 6.56E-17 | 2.85E-16 | Slc13a3 | solute carrier | Plasma Mem |
| ENSMUSG00000026019 | 6.76E-17 | 2.94E-16 | Wdr12 | WD repeat d | Cytoplasm |
| ENSMUSG00000031837 | 6.78E-17 | 2.95E-16 | Necab2 | N-terminal E | Cytoplasm |
| ENSMUSG00000006418 | 6.89E-17 | 2.99E-16 | Rnf114 | ring finger pr | Extracellular S |
| ENSMUSG00000027433 | 6.99E-17 | 3.04E-16 | Xrn2 | 5'-3' exoribo | Nucleus |
| ENSMUSG00000022761 | 7.16E-17 | 3.11E-16 | Lztr1 | leucine zippe | Cytoplasm |
| ENSMUSG00000035337 | 7.19E-17 | 3.12E-16 | Uchl4 | ubiquitin carl | Cytoplasm |
| ENSMUSG00000016534 | 7.25E-17 | 3.14E-16 | Lamp2 | lysosomal ass | Plasma Mem |
| ENSMUSG00000027466 | 7.48E-17 | 3.24E-16 | Rbck1 | RANBP2-type | Cytoplasm |
| ENSMUSG00000002846 | 7.50E-17 | 3.25E-16 | Timmdc1 | translocase o | Cytoplasm |
| ENSMUSG00000000902 | 7.84E-17 | 3.39E-16 | Smarcb1 | SWI/SNF rela | Nucleus |
| ENSMUSG00000023505 | 7.92E-17 | 3.43E-16 | Cdca3 | cell division c | Cytoplasm |
| ENSMUSG00000003235 | 8.04E-17 | 3.47E-16 | Eif2b5 | eukaryotic tra | Cytoplasm |
| ENSMUSG00000042369 | 8.34E-17 | 3.60E-16 | Rbm45 | RNA binding | Cytoplasm |
| ENSMUSG00000004393 | 8.55E-17 | 3.69E-16 | Ddx56 | DEAD-box h | Nucleus |
| ENSMUSG00000029119 | 9.00E-17 | 3.88E-16 | Man2b2 | mannosidase | Cytoplasm |
| ENSMUSG00000006299 | 9.01E-17 | 3.88E-16 | Aamp | angio associa | Plasma Mem |
| ENSMUSG00000022428 | 9.12E-17 | 3.93E-16 | Cby1 | chibby family | Cytoplasm |
| ENSMUSG00000026579 | 9.32E-17 | 4.01E-16 | F5 | coagulation f | Extracellular S |
| ENSMUSG00000038697 | 9.53E-17 | 4.10E-16 | Taf5l | TATA-box bi | Nucleus |
| ENSMUSG00000028843 | 9.83E-17 | 4.23E-16 | Sh3bgrl3 | SH3 domain | Nucleus |
| ENSMUSG00000015748 | 9.95E-17 | 4.28E-16 | Prpf3 | pre-mRNA p | Nucleus |
| ENSMUSG00000005198 | 1.02E-16 | 4.36E-16 | Polr2a | RNA polymer | Nucleus |
| ENSMUSG00000102252 | 1.02E-16 | 4.40E-16 | Snrpn | small nuclear | Nucleus |
| ENSMUSG00000042380 | 1.03E-16 | 4.40E-16 | Smim12 | small integra | Other |
| ENSMUSG00000027091 | 1.04E-16 | 4.45E-16 | Zc3h15 | zinc finger C | Nucleus |
| ENSMUSG00000038552 | 1.05E-16 | 4.48E-16 | Fndc4 | fibronectin ty | Extracellular S |
| ENSMUSG00000022960 | 1.10E-16 | 4.70E-16 | Donson | downstream | Extracellular S |
| ENSMUSG00000058013 | 1.10E-16 | 4.73E-16 | Sept11 | septin 11 | Nucleus |
| ENSMUSG00000037725 | 1.11E-16 | 4.77E-16 | Ckap2 | cytoskeleton | Cytoplasm |
| ENSMUSG00000028576 | 1.12E-16 | 4.79E-16 | Ift74 | intraflagellar | Cytoplasm |
| ENSMUSG00000036427 | 1.12E-16 | 4.80E-16 | Gpi1 | glucose-6-pl | Extracellular S |
| ENSMUSG00000025403 | 1.23E-16 | 5.24E-16 | Shmt2 | serine hydrox | Cytoplasm |
| ENSMUSG00000098274 | 1.23E-16 | 5.24E-16 | Rpl24 | ribosomal pr | Cytoplasm |
| ENSMUSG00000024370 | 1.23E-16 | 5.24E-16 | Cdc23 | cell division c | Nucleus |
| ENSMUSG00000041560 | 1.26E-16 | 5.36E-16 | Gltscr2 | NOP53 ribos | Cytoplasm |

| | | | | | |
|---|---|---|---|---|---|
| ENSMUSG00000069662 | 1.26E-16 | 5.37E-16 | Marcks | myristoylatec | Plasma Mem |
| ENSMUSG00000020974 | 1.27E-16 | 5.40E-16 | Pole2 | DNA polyme | Nucleus |
| ENSMUSG00000020114 | 1.35E-16 | 5.75E-16 | Cand1 | cullin associa | Cytoplasm |
| ENSMUSG00000029521 | 1.37E-16 | 5.82E-16 | Chek2 | checkpoint ki | Nucleus |
| ENSMUSG00000006304 | 1.41E-16 | 6.00E-16 | Arpc2 | actin related | Cytoplasm |
| ENSMUSG00000029036 | 1.42E-16 | 6.06E-16 | Atad3a | ATPase famil | Cytoplasm |
| ENSMUSG00000029366 | 1.46E-16 | 6.23E-16 | Dck | deoxycytidin | Nucleus |
| ENSMUSG00000041881 | 1.54E-16 | 6.52E-16 | Ndufa7 | NADH:ubiqu | Cytoplasm |
| ENSMUSG00000030609 | 1.59E-16 | 6.75E-16 | Aen | apoptosis en | Nucleus |
| ENSMUSG00000030689 | 1.60E-16 | 6.79E-16 | Ino80e | INO80 comp | Nucleus |
| ENSMUSG00000020922 | 1.61E-16 | 6.84E-16 | Lsm12 | LSM12 homc | Other |
| ENSMUSG00000029063 | 1.62E-16 | 6.86E-16 | Nadk | NAD kinase | Cytoplasm |
| ENSMUSG00000035863 | 1.62E-16 | 6.86E-16 | Palm | paralemmin | Plasma Mem |
| ENSMUSG00000031590 | 1.64E-16 | 6.94E-16 | Frg1 | FSHD region | Nucleus |
| ENSMUSG00000027378 | 1.67E-16 | 7.08E-16 | Nphp1 | nephrocystin | Nucleus |
| ENSMUSG00000010175 | 1.72E-16 | 7.29E-16 | Prox1 | prospero hor | Nucleus |
| ENSMUSG00000046811 | 1.74E-16 | 7.37E-16 | Gltpd2 | glycolipid tra | Other |
| ENSMUSG00000024538 | 1.74E-16 | 7.37E-16 | Ppic | peptidylproly | Cytoplasm |
| ENSMUSG00000020936 | 1.78E-16 | 7.51E-16 | Nmt1 | N-myristoylt | Cytoplasm |
| ENSMUSG00000026341 | 1.81E-16 | 7.64E-16 | Actr3 | ARP3 actin re | Plasma Mem |
| ENSMUSG00000024781 | 1.86E-16 | 7.86E-16 | Lipa | lipase A, lyso | Cytoplasm |
| ENSMUSG00000038644 | 1.91E-16 | 8.05E-16 | Pold1 | DNA polyme | Nucleus |
| ENSMUSG00000046959 | 1.91E-16 | 8.05E-16 | Slc26a1 | solute carrier | Plasma Mem |
| ENSMUSG00000027954 | 1.94E-16 | 8.16E-16 | Efna1 | ephrin A1 | Plasma Mem |
| ENSMUSG00000019054 | 1.98E-16 | 8.33E-16 | Fis1 | fission, mitoc | Cytoplasm |
| ENSMUSG00000036167 | 1.99E-16 | 8.38E-16 | Pphln1 | periphilin 1 | Nucleus |
| ENSMUSG00000020069 | 2.20E-16 | 9.26E-16 | Hnrnph3 | heterogeneo | Nucleus |
| ENSMUSG00000025491 | 2.23E-16 | 9.38E-16 | Ifitm1 | interferon inc | Other |
| ENSMUSG00000040550 | 2.24E-16 | 9.41E-16 | Otud6b | OTU domain | Other |
| ENSMUSG00000027430 | 2.24E-16 | 9.42E-16 | Dtd1 | D-tyrosyl-tRI | Cytoplasm |
| ENSMUSG00000021368 | 2.27E-16 | 9.51E-16 | Tbc1d7 | TBC1 domair | Cytoplasm |
| ENSMUSG00000014633 | 2.28E-16 | 9.55E-16 | Cmc2 | C-X9-C moti | Cytoplasm |
| ENSMUSG00000079104 | 2.31E-16 | 9.70E-16 | Prps1l3 | phosphoribo | Other |
| ENSMUSG00000027395 | 2.38E-16 | 9.99E-16 | Polr1b | RNA polymei | Nucleus |
| ENSMUSG00000020873 | 2.39E-16 | 9.99E-16 | Slc35b1 | solute carrier | Cytoplasm |
| ENSMUSG00000033285 | 2.40E-16 | 1.00E-15 | Wdr3 | WD repeat d | Cytoplasm |
| ENSMUSG00000030435 | 2.41E-16 | 1.01E-15 | U2af2 | U2 small nuc | Nucleus |
| ENSMUSG00000030189 | 2.48E-16 | 1.04E-15 | Ybx3 | Y-box bindin | Nucleus |
| ENSMUSG00000033105 | 2.57E-16 | 1.07E-15 | Lss | lanosterol syi | Cytoplasm |
| ENSMUSG00000026578 | 2.68E-16 | 1.12E-15 | Ccdc181 | coiled-coil d | Nucleus |
| ENSMUSG00000037376 | 2.70E-16 | 1.13E-15 | Trmt6 | tRNA methyl | Nucleus |
| ENSMUSG00000034918 | 2.78E-16 | 1.16E-15 | Cdhr2 | cadherin rela | Plasma Mem |
| ENSMUSG00000030254 | 2.84E-16 | 1.18E-15 | Rad18 | RAD18, E3 ul | Nucleus |
| ENSMUSG00000031775 | 3.01E-16 | 1.25E-15 | Pllp | plasmolipin | Plasma Mem |
| ENSMUSG00000063439 | 3.12E-16 | 1.30E-15 | B9d2 | B9 domain c | Cytoplasm |
| ENSMUSG00000037461 | 3.12E-16 | 1.30E-15 | Ints7 | integrator co | Nucleus |
| ENSMUSG00000022721 | 3.15E-16 | 1.31E-15 | Trmt2a | tRNA methyl | Other |
| ENSMUSG00000015890 | 3.29E-16 | 1.37E-15 | Amdhd1 | amidohydrol | Cytoplasm |
| ENSMUSG00000037493 | 3.38E-16 | 1.41E-15 | Cib2 | calcium and i | Cytoplasm |
| ENSMUSG00000020048 | 3.48E-16 | 1.45E-15 | Hsp90b1 | heat shock p | Cytoplasm |
| ENSMUSG00000020661 | 3.55E-16 | 1.47E-15 | Dnmt3a | DNA methylt | Nucleus |
| ENSMUSG00000024856 | 3.62E-16 | 1.50E-15 | Cdk2ap2 | cyclin depen | Nucleus |
| ENSMUSG00000021774 | 3.65E-16 | 1.51E-15 | Ube2e1 | ubiquitin cor | Cytoplasm |
| ENSMUSG00000030045 | 3.65E-16 | 1.51E-15 | Mrpl19 | mitochondria | Cytoplasm |
| ENSMUSG00000002767 | 3.71E-16 | 1.54E-15 | Mrpl2 | mitochondria | Extracellular S |
| ENSMUSG00000044763 | 3.75E-16 | 1.55E-15 | Trmt10c | tRNA methyl | Cytoplasm |
| ENSMUSG00000027699 | 3.76E-16 | 1.56E-15 | Ect2 | epithelial cell | Cytoplasm |
| ENSMUSG00000063200 | 3.79E-16 | 1.57E-15 | Nol7 | nucleolar prc | Nucleus |

| | | | | | |
|---|---|---|---|---|---|
| ENSMUSG00000021302 | 3.99E-16 | 1.65E-15 | Ggps1 | geranylgeran | Cytoplasm |
| ENSMUSG00000051413 | 4.06E-16 | 1.68E-15 | Plagl2 | PLAG1 like zi | Nucleus |
| ENSMUSG00000047649 | 4.20E-16 | 1.74E-15 | Cd3eap | CD3e molecu | Nucleus |
| ENSMUSG00000006456 | 4.25E-16 | 1.75E-15 | Rbm14 | RNA binding | Nucleus |
| ENSMUSG00000061079 | 4.41E-16 | 1.82E-15 | Zfp143 | zinc finger pı | Nucleus |
| ENSMUSG00000007836 | 4.52E-16 | 1.87E-15 | Hnrnpa0 | heterogeneo | Nucleus |
| ENSMUSG00000018697 | 4.56E-16 | 1.88E-15 | Aatf | apoptosis an | Nucleus |
| ENSMUSG00000010554 | 4.70E-16 | 1.94E-15 | Mettl16 | methyltransfe | Nucleus |
| ENSMUSG00000058392 | 4.73E-16 | 1.95E-15 | Rrp1b | ribosomal RN | Nucleus |
| ENSMUSG00000028447 | 4.89E-16 | 2.01E-15 | Dctn3 | dynactin sub | Nucleus |
| ENSMUSG00000029535 | 4.96E-16 | 2.04E-15 | Triap1 | TP53 regulat | Cytoplasm |
| ENSMUSG00000008690 | 5.04E-16 | 2.07E-15 | Ncaph2 | non-SMC co | Nucleus |
| ENSMUSG00000031198 | 5.17E-16 | 2.13E-15 | Fundc2 | FUN14 doma | Cytoplasm |
| ENSMUSG00000020467 | 5.18E-16 | 2.13E-15 | Efemp1 | EGF containir | Extracellular S |
| ENSMUSG00000030505 | 5.19E-16 | 2.13E-15 | Prmt3 | protein argin | Nucleus |
| ENSMUSG00000030315 | 5.23E-16 | 2.15E-15 | Vgll4 | vestigial like | Nucleus |
| ENSMUSG00000002015 | 5.24E-16 | 2.15E-15 | Bcap31 | B cell recepto | Cytoplasm |
| ENSMUSG00000039693 | 5.62E-16 | 2.31E-15 | Msantd3 | Myb/SANT D | Other |
| ENSMUSG00000022010 | 5.88E-16 | 2.41E-15 | Tsc22d1 | TSC22 doma | Nucleus |
| ENSMUSG00000023919 | 5.94E-16 | 2.43E-15 | Cenpq | centromere p | Nucleus |
| ENSMUSG00000048109 | 5.96E-16 | 2.44E-15 | Rbm15 | RNA binding | Nucleus |
| ENSMUSG00000044783 | 6.06E-16 | 2.48E-15 | Hjurp | Holliday junc | Nucleus |
| ENSMUSG00000024234 | 6.31E-16 | 2.58E-15 | Mtpap | mitochondria | Cytoplasm |
| ENSMUSG00000027452 | 6.32E-16 | 2.58E-15 | Acss1 | acyl-CoA syr | Cytoplasm |
| ENSMUSG00000030888 | 6.45E-16 | 2.64E-15 | Rrp8 | ribosomal RN | Nucleus |
| ENSMUSG00000039347 | 6.50E-16 | 2.65E-15 | Atp6v0e2 | ATPase H+ tı | Cytoplasm |
| ENSMUSG00000045751 | 6.64E-16 | 2.71E-15 | Mms22l | MMS22 like, | Nucleus |
| ENSMUSG00000030216 | 6.70E-16 | 2.73E-15 | Wbp11 | WW domain | Nucleus |
| ENSMUSG00000005312 | 6.72E-16 | 2.74E-15 | Ubqln1 | ubiquilin 1 | Cytoplasm |
| ENSMUSG00000031657 | 6.87E-16 | 2.80E-15 | Heatr3 | HEAT repeat | Other |
| ENSMUSG00000028059 | 6.97E-16 | 2.84E-15 | Arhgef2 | Rho/Rac gua | Cytoplasm |
| ENSMUSG00000041429 | 7.06E-16 | 2.87E-15 | Nthl1 | nth like DNA | Nucleus |
| ENSMUSG00000018882 | 7.16E-16 | 2.91E-15 | Mrpl45 | mitochondria | Cytoplasm |
| ENSMUSG00000041632 | 7.37E-16 | 3.00E-15 | Mrps27 | mitochondria | Cytoplasm |
| ENSMUSG00000033918 | 7.55E-16 | 3.07E-15 | Parl | presenilin ass | Cytoplasm |
| ENSMUSG00000071054 | 7.65E-16 | 3.11E-15 | Safb | scaffold attac | Nucleus |
| ENSMUSG00000022677 | 7.71E-16 | 3.13E-15 | Fopnl | FGFR1OP N- | Cytoplasm |
| ENSMUSG00000022023 | 7.92E-16 | 3.21E-15 | Wbp4 | WW domain | Cytoplasm |
| ENSMUSG00000038619 | 7.96E-16 | 3.23E-15 | Ensa | endosulfine a | Cytoplasm |
| ENSMUSG00000079562 | 8.02E-16 | 3.25E-15 | Maea | macrophage | Plasma Mem |
| ENSMUSG00000016477 | 8.05E-16 | 3.26E-15 | E2f3 | E2F transcrip | Nucleus |
| ENSMUSG00000048668 | 8.28E-16 | 3.35E-15 | Rhno1 | RAD9-HUS1- | Nucleus |
| ENSMUSG00000038526 | 8.41E-16 | 3.40E-15 | Car14 | carbonic anh | Plasma Mem |
| ENSMUSG00000029467 | 8.43E-16 | 3.41E-15 | Atp2a2 | ATPase sarcc | Cytoplasm |
| ENSMUSG00000038845 | 8.66E-16 | 3.50E-15 | Phb | prohibitin | Nucleus |
| ENSMUSG00000073775 | 8.72E-16 | 3.53E-15 | Kti12 | KTI12 chroma | Other |
| ENSMUSG00000018572 | 8.73E-16 | 3.53E-15 | Phf23 | PHD finger p | Nucleus |
| ENSMUSG00000023106 | 8.77E-16 | 3.54E-15 | Denr | density regul | Other |
| ENSMUSG00000031536 | 8.86E-16 | 3.58E-15 | Polb | DNA polyme | Nucleus |
| ENSMUSG00000026568 | 9.06E-16 | 3.66E-15 | Mpc2 | mitochondria | Plasma Mem |
| ENSMUSG00000035575 | 9.30E-16 | 3.75E-15 | Utp6 | UTP6, small s | Nucleus |
| ENSMUSG00000007867 | 9.35E-16 | 3.77E-15 | Ift43 | intraflagellar | Cytoplasm |
| ENSMUSG00000003778 | 9.53E-16 | 3.84E-15 | Brd8 | bromodomai | Nucleus |
| ENSMUSG00000061518 | 9.74E-16 | 3.92E-15 | Cox5b | cytochrome ( | Cytoplasm |
| ENSMUSG00000091867 | 1.02E-15 | 4.12E-15 | Cyp2a22 | cytochrome I | Cytoplasm |
| ENSMUSG00000013662 | 1.03E-15 | 4.16E-15 | Atad1 | ATPase famil | Plasma Mem |
| ENSMUSG00000020705 | 1.05E-15 | 4.22E-15 | Ddx42 | DEAD-box h | Cytoplasm |
| ENSMUSG00000068101 | 1.07E-15 | 4.30E-15 | Cenpm | centromere p | Cytoplasm |

| | | | | | |
|---|---|---|---|---|---|
| ENSMUSG00000034371 | 1.07E-15 | 4.31E-15 | Dak | triokinase an | Cytoplasm |
| ENSMUSG00000039828 | 1.12E-15 | 4.52E-15 | Wdr70 | WD repeat d | Other |
| ENSMUSG00000062624 | 1.13E-15 | 4.52E-15 | Cyp2c67 | cytochrome I | Cytoplasm |
| ENSMUSG00000024668 | 1.13E-15 | 4.53E-15 | Sdhaf2 | succinate del | Extracellular S |
| ENSMUSG00000029177 | 1.15E-15 | 4.60E-15 | Cenpa | centromere p | Nucleus |
| ENSMUSG00000027649 | 1.15E-15 | 4.60E-15 | Ctnnbl1 | catenin beta | Nucleus |
| ENSMUSG00000008348 | 1.16E-15 | 4.66E-15 | Ubc | ubiquitin C | Cytoplasm |
| ENSMUSG00000024560 | 1.21E-15 | 4.84E-15 | Cxxc1 | CXXC finger | Nucleus |
| ENSMUSG00000063253 | 1.21E-15 | 4.84E-15 | Scoc | short coiled- | Cytoplasm |
| ENSMUSG00000032251 | 1.22E-15 | 4.89E-15 | Irak1bp1 | interleukin 1 | Other |
| ENSMUSG00000024539 | 1.22E-15 | 4.89E-15 | Ptpn2 | protein tyros | Cytoplasm |
| ENSMUSG00000037262 | 1.23E-15 | 4.92E-15 | Kin | Kin17 DNA a | Nucleus |
| ENSMUSG00000028494 | 1.23E-15 | 4.93E-15 | Plin2 | perilipin 2 | Plasma Mem |
| ENSMUSG00000046814 | 1.25E-15 | 4.99E-15 | Gchfr | GTP cyclohyc | Cytoplasm |
| ENSMUSG00000005378 | 1.25E-15 | 5.01E-15 | Wbscr22 | BUD23, rRNA | Nucleus |
| ENSMUSG00000025745 | 1.28E-15 | 5.11E-15 | Hadha | hydroxyacyl- | Cytoplasm |
| ENSMUSG00000042133 | 1.28E-15 | 5.11E-15 | Ppig | peptidylproly | Nucleus |
| ENSMUSG00000045598 | 1.30E-15 | 5.19E-15 | Zfp553 | zinc finger pr | Nucleus |
| ENSMUSG00000016619 | 1.30E-15 | 5.19E-15 | Nup50 | nucleoporin 5 | Nucleus |
| ENSMUSG00000050856 | 1.31E-15 | 5.20E-15 | Atp5k | ATP synthase | Cytoplasm |
| ENSMUSG00000058258 | 1.33E-15 | 5.30E-15 | Idi1 | isopentenyl-c | Cytoplasm |
| ENSMUSG00000029038 | 1.34E-15 | 5.32E-15 | Ssu72 | SSU72 homo | Other |
| ENSMUSG00000022621 | 1.35E-15 | 5.35E-15 | Rabl2 | RAB, membe | Cytoplasm |
| ENSMUSG00000029110 | 1.37E-15 | 5.43E-15 | Rnf4 | ring finger pr | Nucleus |
| ENSMUSG00000031826 | 1.38E-15 | 5.46E-15 | Usp10 | ubiquitin spe | Cytoplasm |
| ENSMUSG00000032366 | 1.38E-15 | 5.47E-15 | Tpm1 | tropomyosin | Plasma Mem |
| ENSMUSG00000059434 | 1.38E-15 | 5.48E-15 | Gckr | glucokinase r | Nucleus |
| ENSMUSG00000038671 | 1.39E-15 | 5.51E-15 | Arfrp1 | ADP ribosyla | Cytoplasm |
| ENSMUSG00000035637 | 1.45E-15 | 5.74E-15 | Grhpr | glyoxylate an | Cytoplasm |
| ENSMUSG00000025532 | 1.47E-15 | 5.83E-15 | Crcp | CGRP recept | Plasma Mem |
| ENSMUSG00000024949 | 1.49E-15 | 5.90E-15 | Sf1 | splicing facto | Nucleus |
| ENSMUSG00000037364 | 1.49E-15 | 5.90E-15 | Srrt | serrate, RNA | Nucleus |
| ENSMUSG00000066357 | 1.50E-15 | 5.92E-15 | Wdr6 | WD repeat d | Cytoplasm |
| ENSMUSG00000039671 | 1.50E-15 | 5.93E-15 | Zmynd8 | zinc finger M | Nucleus |
| ENSMUSG00000071645 | 1.58E-15 | 6.24E-15 | Tut1 | terminal urid | Nucleus |
| ENSMUSG00000025722 | 1.64E-15 | 6.48E-15 | Wdr73 | WD repeat d | Extracellular S |
| ENSMUSG00000019849 | 1.70E-15 | 6.71E-15 | Prep | prolyl endop | Cytoplasm |
| ENSMUSG00000046295 | 1.71E-15 | 6.76E-15 | Ankle1 | ankyrin repea | Cytoplasm |
| ENSMUSG00000042073 | 1.73E-15 | 6.81E-15 | Abhd14b | abhydrolase | Cytoplasm |
| ENSMUSG00000073481 | 1.73E-15 | 6.81E-15 | Marc2 | mitochondria | Cytoplasm |
| ENSMUSG00000029687 | 1.73E-15 | 6.84E-15 | Ezh2 | enhancer of z | Nucleus |
| ENSMUSG00000039737 | 1.84E-15 | 7.25E-15 | Prkrip1 | PRKR interac | Nucleus |
| ENSMUSG00000025857 | 1.86E-15 | 7.33E-15 | Dnaaf5 | dynein axone | Cytoplasm |
| ENSMUSG00000031371 | 1.90E-15 | 7.49E-15 | Haus7 | HAUS augmi | Cytoplasm |
| ENSMUSG00000004044 | 1.91E-15 | 7.50E-15 | Ptrf | caveolae asso | Nucleus |
| ENSMUSG00000029145 | 1.91E-15 | 7.50E-15 | Eif2b4 | eukaryotic tra | Cytoplasm |
| ENSMUSG00000024982 | 1.92E-15 | 7.56E-15 | Zdhhc6 | zinc finger DI | Nucleus |
| ENSMUSG00000031467 | 1.93E-15 | 7.60E-15 | Agpat5 | 1-acylglycero | Cytoplasm |
| ENSMUSG00000030127 | 1.94E-15 | 7.61E-15 | Cops7a | COP9 signalo | Cytoplasm |
| ENSMUSG00000063334 | 1.96E-15 | 7.70E-15 | Krr1 | KRR1, small s | Nucleus |
| ENSMUSG00000034793 | 1.99E-15 | 7.79E-15 | G6pc3 | glucose-6-pl | Cytoplasm |
| ENSMUSG00000034254 | 2.04E-15 | 8.00E-15 | Agpat1 | 1-acylglycero | Cytoplasm |
| ENSMUSG00000065990 | 2.04E-15 | 8.00E-15 | Aurkaip1 | aurora kinase | Nucleus |
| ENSMUSG00000026869 | 2.11E-15 | 8.25E-15 | Psmd5 | proteasome 2 | Other |
| ENSMUSG00000032621 | 2.11E-15 | 8.27E-15 | Srek1 | splicing regu | Nucleus |
| ENSMUSG00000028347 | 2.11E-15 | 8.27E-15 | Tmeff1 | transmembra | Plasma Mem |
| ENSMUSG00000049422 | 2.27E-15 | 8.89E-15 | Chchd10 | coiled-coil-h | Cytoplasm |
| ENSMUSG00000018569 | 2.30E-15 | 9.00E-15 | Cldn7 | claudin 7 | Plasma Mem |

| | | | | | |
|---|---|---|---|---|---|
| ENSMUSG00000038047 | 2.36E-15 | 9.21E-15 | Haus6 | HAUS augmi | Cytoplasm |
| ENSMUSG00000015757 | 2.40E-15 | 9.36E-15 | Ppil4 | peptidylproly | Nucleus |
| ENSMUSG00000018433 | 2.41E-15 | 9.41E-15 | Nol11 | nucleolar prc | Nucleus |
| ENSMUSG00000042719 | 2.42E-15 | 9.43E-15 | Naa25 | N(alpha)-ace | Cytoplasm |
| ENSMUSG00000049950 | 2.47E-15 | 9.62E-15 | Rpp38 | ribonuclease | Nucleus |
| ENSMUSG00000039660 | 2.48E-15 | 9.66E-15 | D2Wsu81e | SPOUT doma | Nucleus |
| ENSMUSG00000032126 | 2.62E-15 | 1.02E-14 | Hmbs | hydroxymeth | Cytoplasm |
| ENSMUSG00000027690 | 2.65E-15 | 1.03E-14 | Slc2a2 | solute carrier | Plasma Mem |
| ENSMUSG00000033706 | 2.70E-15 | 1.05E-14 | Smyd5 | SMYD family | Other |
| ENSMUSG00000040435 | 2.72E-15 | 1.06E-14 | Ppp1r15a | protein phos | Cytoplasm |
| ENSMUSG00000030086 | 2.74E-15 | 1.07E-14 | Chchd6 | coiled-coil-h | Cytoplasm |
| ENSMUSG00000024050 | 2.87E-15 | 1.11E-14 | Wiz | widely intersp | Nucleus |
| ENSMUSG00000005103 | 2.97E-15 | 1.15E-14 | Wdr1 | WD repeat d | Extracellular S |
| ENSMUSG00000026975 | 2.99E-15 | 1.16E-14 | Dph7 | diphthamide | Cytoplasm |
| ENSMUSG00000000295 | 3.08E-15 | 1.20E-14 | Hddc2 | HD domain c | Cytoplasm |
| ENSMUSG00000030611 | 3.13E-15 | 1.21E-14 | Mrps11 | mitochondria | Cytoplasm |
| ENSMUSG00000034118 | 3.17E-15 | 1.23E-14 | Tpst1 | tyrosylprotei | Cytoplasm |
| ENSMUSG00000031754 | 3.26E-15 | 1.27E-14 | Nudt21 | nudix hydrola | Nucleus |
| ENSMUSG00000028261 | 3.27E-15 | 1.27E-14 | Ndufaf4 | NADH:ubiqu | Cytoplasm |
| ENSMUSG00000021065 | 3.46E-15 | 1.34E-14 | Fut8 | fucosyltransfe | Cytoplasm |
| ENSMUSG00000032221 | 3.49E-15 | 1.35E-14 | Mns1 | meiosis speci | Cytoplasm |
| ENSMUSG00000032040 | 3.66E-15 | 1.41E-14 | Dcps | decapping er | Nucleus |
| ENSMUSG00000008475 | 3.77E-15 | 1.46E-14 | Arpc5 | actin related | Cytoplasm |
| ENSMUSG00000036822 | 3.77E-15 | 1.46E-14 | Topors | TOP1 binding | Nucleus |
| ENSMUSG00000038155 | 3.82E-15 | 1.47E-14 | Gstp2 | glutathione S | Cytoplasm |
| ENSMUSG00000007880 | 3.83E-15 | 1.48E-14 | Arid1a | AT-rich inter | Nucleus |
| ENSMUSG00000025474 | 3.91E-15 | 1.51E-14 | Tubgcp2 | tubulin gamr | Cytoplasm |
| ENSMUSG00000031711 | 3.98E-15 | 1.53E-14 | Zfp330 | zinc finger pr | Nucleus |
| ENSMUSG00000079259 | 4.31E-15 | 1.66E-14 | Trim71 | tripartite mot | Cytoplasm |
| ENSMUSG00000040250 | 4.42E-15 | 1.70E-14 | Asun | integrator co | Cytoplasm |
| ENSMUSG00000025608 | 4.43E-15 | 1.71E-14 | Podxl | podocalyxin- | Plasma Mem |
| ENSMUSG00000011837 | 4.72E-15 | 1.82E-14 | Snapc2 | small nuclear | Nucleus |
| ENSMUSG00000026918 | 4.77E-15 | 1.83E-14 | Brd3 | bromodomai | Nucleus |
| ENSMUSG00000020265 | 4.80E-15 | 1.85E-14 | Sumo3 | small ubiquit | Nucleus |
| ENSMUSG00000037214 | 4.83E-15 | 1.86E-14 | Thap1 | THAP domai | Nucleus |
| ENSMUSG00000021810 | 5.00E-15 | 1.92E-14 | Ecd | ecdysoneless | Nucleus |
| ENSMUSG00000030960 | 5.10E-15 | 1.96E-14 | Mettl10 | EEF1A lysine | Cytoplasm |
| ENSMUSG00000025925 | 5.25E-15 | 2.01E-14 | Terf1 | telomeric rep | Nucleus |
| ENSMUSG00000034403 | 5.46E-15 | 2.09E-14 | Pja1 | praja ring fin | Cytoplasm |
| ENSMUSG00000039128 | 5.58E-15 | 2.14E-14 | Cdc123 | cell division c | Cytoplasm |
| ENSMUSG00000015880 | 5.58E-15 | 2.14E-14 | Ncapg | non-SMC co | Nucleus |
| ENSMUSG00000059273 | 5.65E-15 | 2.16E-14 | Zc3h4 | zinc finger C( | Nucleus |
| ENSMUSG00000020709 | 5.66E-15 | 2.16E-14 | Adap2 | ArfGAP with | Cytoplasm |
| ENSMUSG00000078440 | 5.71E-15 | 2.18E-14 | Dohh | deoxyhypusi | Cytoplasm |
| ENSMUSG00000058587 | 5.80E-15 | 2.22E-14 | Tmod3 | tropomoduli | Cytoplasm |
| ENSMUSG00000027650 | 5.83E-15 | 2.23E-14 | Tti1 | TELO2 intera | Cytoplasm |
| ENSMUSG00000015354 | 6.03E-15 | 2.30E-14 | Pcolce2 | procollagen ( | Extracellular S |
| ENSMUSG00000038079 | 6.08E-15 | 2.32E-14 | Tmem237 | transmembra | Other |
| ENSMUSG00000037938 | 6.22E-15 | 2.37E-14 | Chchd5 | coiled-coil-h | Other |
| ENSMUSG00000035297 | 6.32E-15 | 2.41E-14 | Cops4 | COP9 signalc | Cytoplasm |
| ENSMUSG00000032469 | 6.33E-15 | 2.41E-14 | Dbr1 | debranching | Nucleus |
| ENSMUSG00000042506 | 6.43E-15 | 2.45E-14 | Usp22 | ubiquitin spe | Nucleus |
| ENSMUSG00000050071 | 6.63E-15 | 2.52E-14 | Bex1 | brain express | Other |
| ENSMUSG00000050244 | 6.73E-15 | 2.56E-14 | Heatr1 | HEAT repeat | Nucleus |
| ENSMUSG00000026111 | 6.80E-15 | 2.59E-14 | Unc50 | unc-50 inner | Cytoplasm |
| ENSMUSG00000038576 | 6.81E-15 | 2.59E-14 | Susd4 | sushi domair | Other |
| ENSMUSG00000026489 | 6.94E-15 | 2.64E-14 | Adck3 | coenzyme Q8 | Cytoplasm |
| ENSMUSG00000050428 | 6.99E-15 | 2.65E-14 | Fbxo46 | F-box protei | Other |

| | | | | | |
|---|---|---|---|---|---|
| ENSMUSG00000029062 | 7.05E-15 | 2.68E-14 | Cdk11b | cyclin depend | Nucleus |
| ENSMUSG00000030834 | 7.15E-15 | 2.71E-14 | Abcc6 | ATP binding | Plasma Mem |
| ENSMUSG00000035964 | 7.16E-15 | 2.72E-14 | Tmem59l | transmembra | Cytoplasm |
| ENSMUSG00000031931 | 7.18E-15 | 2.72E-14 | Ankrd49 | ankyrin repea | Nucleus |
| ENSMUSG00000029840 | 7.67E-15 | 2.91E-14 | Mtpn | myotrophin | Nucleus |
| ENSMUSG00000032264 | 7.85E-15 | 2.98E-14 | Zw10 | zw10 kinetoc | Nucleus |
| ENSMUSG00000036315 | 7.86E-15 | 2.98E-14 | Znrd1 | zinc ribbon d | Nucleus |
| ENSMUSG00000027714 | 7.93E-15 | 3.00E-14 | Exosc9 | exosome com | Nucleus |
| ENSMUSG00000026205 | 8.02E-15 | 3.04E-14 | Slc23a3 | solute carrier | Other |
| ENSMUSG00000025665 | 8.38E-15 | 3.17E-14 | Rps6ka6 | ribosomal pr | Cytoplasm |
| ENSMUSG00000026816 | 8.53E-15 | 3.22E-14 | Gtf3c5 | general trans | Nucleus |
| ENSMUSG00000070520 | 8.56E-15 | 3.24E-14 | Ndnl2 | NSE3 homolo | Nucleus |
| ENSMUSG00000004748 | 8.89E-15 | 3.36E-14 | Mtfp1 | mitochondria | Cytoplasm |
| ENSMUSG00000069135 | 8.95E-15 | 3.38E-14 | Fgfr1op | FGFR1 oncog | Cytoplasm |
| ENSMUSG00000035504 | 9.09E-15 | 3.43E-14 | Reep6 | receptor acce | Plasma Mem |
| ENSMUSG00000035227 | 9.15E-15 | 3.45E-14 | Spcs2 | signal peptid | Cytoplasm |
| ENSMUSG00000014294 | 9.16E-15 | 3.45E-14 | Ndufa2 | NADH:ubiqu | Cytoplasm |
| ENSMUSG00000024359 | 9.28E-15 | 3.50E-14 | Hspa9 | heat shock p | Cytoplasm |
| ENSMUSG00000078695 | 9.36E-15 | 3.52E-14 | Cisd3 | CDGSH iron s | Cytoplasm |
| ENSMUSG00000069910 | 9.68E-15 | 3.64E-14 | Spdl1 | spindle appa | Nucleus |
| ENSMUSG00000023175 | 9.89E-15 | 3.72E-14 | Bsg | basigin (Ok b | Plasma Mem |
| ENSMUSG00000002307 | 1.00E-14 | 3.77E-14 | Daxx | death domai | Nucleus |
| ENSMUSG00000020444 | 1.02E-14 | 3.85E-14 | Guk1 | guanylate kin | Cytoplasm |
| ENSMUSG00000018548 | 1.03E-14 | 3.86E-14 | Trim37 | tripartite mot | Cytoplasm |
| ENSMUSG00000021392 | 1.04E-14 | 3.90E-14 | Nol8 | nucleolar pro | Nucleus |
| ENSMUSG00000028821 | 1.05E-14 | 3.94E-14 | Syf2 | SYF2 pre-mR | Nucleus |
| ENSMUSG00000029203 | 1.05E-14 | 3.96E-14 | Ube2k | ubiquitin con | Cytoplasm |
| ENSMUSG00000025004 | 1.07E-14 | 4.01E-14 | Cyp2c40 | cytochrome I | Cytoplasm |
| ENSMUSG00000002635 | 1.07E-14 | 4.02E-14 | Pdcd2l | programmed | Other |
| ENSMUSG00000031862 | 1.08E-14 | 4.04E-14 | Atp13a1 | ATPase 13A1 | Extracellular S |
| ENSMUSG00000062661 | 1.11E-14 | 4.16E-14 | Ncs1 | neuronal calc | Plasma Mem |
| ENSMUSG00000006315 | 1.11E-14 | 4.17E-14 | Tmem147 | transmembra | Cytoplasm |
| ENSMUSG00000020358 | 1.12E-14 | 4.18E-14 | Hnrnpab | heterogeneo | Nucleus |
| ENSMUSG00000018287 | 1.13E-14 | 4.22E-14 | Spag7 | sperm associ | Nucleus |
| ENSMUSG00000016344 | 1.15E-14 | 4.30E-14 | Ppdpf | pancreatic pr | Other |
| ENSMUSG00000039515 | 1.18E-14 | 4.42E-14 | Ppp2r4 | protein phos | Cytoplasm |
| ENSMUSG00000020850 | 1.21E-14 | 4.51E-14 | Prpf8 | pre-mRNA p | Nucleus |
| ENSMUSG00000022792 | 1.21E-14 | 4.51E-14 | Yars2 | tyrosyl-tRNA | Cytoplasm |
| ENSMUSG00000031819 | 1.22E-14 | 4.53E-14 | Emc8 | ER membran | Cytoplasm |
| ENSMUSG00000020464 | 1.23E-14 | 4.59E-14 | Pnpt1 | polyribonucle | Cytoplasm |
| ENSMUSG00000003762 | 1.24E-14 | 4.62E-14 | Adck4 | coenzyme Q8 | Cytoplasm |
| ENSMUSG00000032216 | 1.26E-14 | 4.69E-14 | Nedd4 | neural precur | Cytoplasm |
| ENSMUSG00000036893 | 1.28E-14 | 4.76E-14 | Ehmt1 | euchromatic | Nucleus |
| ENSMUSG00000062421 | 1.28E-14 | 4.77E-14 | Arf2 | ADP-ribosyla | Cytoplasm |
| ENSMUSG00000041879 | 1.29E-14 | 4.81E-14 | Ipo9 | importin 9 | Nucleus |
| ENSMUSG00000027593 | 1.34E-14 | 4.99E-14 | Raly | RALY heterog | Nucleus |
| ENSMUSG00000017485 | 1.36E-14 | 5.05E-14 | Top2b | DNA topoiso | Nucleus |
| ENSMUSG00000042489 | 1.38E-14 | 5.14E-14 | Clspn | claspin | Nucleus |
| ENSMUSG00000033684 | 1.39E-14 | 5.14E-14 | Qsox1 | quiescin sulfh | Cytoplasm |
| ENSMUSG00000028291 | 1.39E-14 | 5.16E-14 | Akirin2 | akirin 2 | Nucleus |
| ENSMUSG00000029267 | 1.40E-14 | 5.21E-14 | Mtf2 | metal respon | Nucleus |
| ENSMUSG00000037017 | 1.45E-14 | 5.38E-14 | Zscan21 | zinc finger an | Nucleus |
| ENSMUSG00000020166 | 1.46E-14 | 5.41E-14 | Cnot2 | CCR4-NOT tr | Nucleus |
| ENSMUSG00000024826 | 1.48E-14 | 5.47E-14 | Dpf2 | double PHD | Nucleus |
| ENSMUSG00000021177 | 1.49E-14 | 5.52E-14 | Tdp1 | tyrosyl-DNA | Nucleus |
| ENSMUSG00000026134 | 1.51E-14 | 5.59E-14 | Prim2 | DNA primase | Nucleus |
| ENSMUSG00000029591 | 1.56E-14 | 5.76E-14 | Ung | uracil DNA g | Nucleus |
| ENSMUSG00000020330 | 1.58E-14 | 5.85E-14 | Hmmr | hyaluronan n | Plasma Mem |

| | | | | | |
|---|---|---|---|---|---|
| ENSMUSG00000015652 | 1.61E-14 | 5.97E-14 | Steap1 | STEAP family | Plasma Mem |
| ENSMUSG00000033323 | 1.63E-14 | 6.02E-14 | Ctdp1 | CTD phospha | Nucleus |
| ENSMUSG00000020674 | 1.73E-14 | 6.37E-14 | Pxdn | peroxidasin | Extracellular S |
| ENSMUSG00000028180 | 1.78E-14 | 6.55E-14 | Zranb2 | zinc finger, R | Nucleus |
| ENSMUSG00000028848 | 1.84E-14 | 6.78E-14 | Gpn2 | GPN-loop GT | Other |
| ENSMUSG00000028573 | 1.91E-14 | 7.04E-14 | Fggy | FGGY carboh | Other |
| ENSMUSG00000020755 | 1.96E-14 | 7.21E-14 | Sap30bp | SAP30 bindir | Nucleus |
| ENSMUSG00000006276 | 2.03E-14 | 7.46E-14 | Eps15l1 | epidermal gr | Plasma Mem |
| ENSMUSG00000040599 | 2.08E-14 | 7.67E-14 | Mis12 | MIS12, kineto | Nucleus |
| ENSMUSG00000024208 | 2.09E-14 | 7.69E-14 | Uqcc2 | ubiquinol-cy | Cytoplasm |
| ENSMUSG00000020462 | 2.11E-14 | 7.75E-14 | Cfap36 | cilia and flag | Extracellular S |
| ENSMUSG00000035585 | 2.14E-14 | 7.87E-14 | Tsen34 | tRNA splicing | Nucleus |
| ENSMUSG00000059248 | 2.15E-14 | 7.90E-14 | Sept9 | septin 9 | Cytoplasm |
| ENSMUSG00000049489 | 2.21E-14 | 8.11E-14 | Fam58b | cyclin Q | Other |
| ENSMUSG00000030718 | 2.24E-14 | 8.22E-14 | Ppme1 | protein phos | Nucleus |
| ENSMUSG00000021532 | 2.27E-14 | 8.32E-14 | Fastkd3 | FAST kinase d | Cytoplasm |
| ENSMUSG00000038943 | 2.30E-14 | 8.42E-14 | Prc1 | protein regul | Nucleus |
| ENSMUSG00000042066 | 2.32E-14 | 8.51E-14 | Tmcc2 | transmembra | Extracellular S |
| ENSMUSG00000041238 | 2.35E-14 | 8.60E-14 | Rbbp8 | RB binding p | Nucleus |
| ENSMUSG00000060739 | 2.38E-14 | 8.71E-14 | Nsa2 | NSA2, riboso | Nucleus |
| ENSMUSG00000024889 | 2.40E-14 | 8.79E-14 | Rce1 | Ras convertir | Cytoplasm |
| ENSMUSG00000020072 | 2.44E-14 | 8.93E-14 | Pbld2 | phenazine bi | Other |
| ENSMUSG00000033589 | 2.77E-14 | 1.01E-13 | Reep4 | receptor acce | Cytoplasm |
| ENSMUSG00000029701 | 2.79E-14 | 1.02E-13 | Rbm28 | RNA binding | Nucleus |
| ENSMUSG00000032348 | 2.86E-14 | 1.05E-13 | Gsta4 | glutathione S | Other |
| ENSMUSG00000038195 | 2.92E-14 | 1.07E-13 | Rilp | Rab interactir | Cytoplasm |
| ENSMUSG00000045193 | 3.03E-14 | 1.11E-13 | Cirbp | cold inducibl | Nucleus |
| ENSMUSG00000043424 | 3.05E-14 | 1.11E-13 | Eif3j2 | eukaryotic tra | Cytoplasm |
| ENSMUSG00000031729 | 3.29E-14 | 1.20E-13 | Ist1 | IST1, ESCRT- | Cytoplasm |
| ENSMUSG00000049764 | 3.34E-14 | 1.22E-13 | Zfp280b | zinc finger pr | Nucleus |
| ENSMUSG00000040383 | 3.35E-14 | 1.22E-13 | Aqr | aquarius intro | Nucleus |
| ENSMUSG00000028292 | 3.42E-14 | 1.24E-13 | Rars2 | arginyl-tRNA | Cytoplasm |
| ENSMUSG00000027367 | 3.48E-14 | 1.27E-13 | Stard7 | StAR related | Cytoplasm |
| ENSMUSG00000029918 | 3.54E-14 | 1.29E-13 | Mrps33 | mitochondria | Cytoplasm |
| ENSMUSG00000015341 | 3.56E-14 | 1.30E-13 | Golga7 | golgin A7 | Cytoplasm |
| ENSMUSG00000029500 | 3.60E-14 | 1.31E-13 | Pgam5 | PGAM family | Cytoplasm |
| ENSMUSG00000030188 | 3.72E-14 | 1.35E-13 | Magohb | mago homol | Nucleus |
| ENSMUSG00000055917 | 3.79E-14 | 1.38E-13 | Zfp277 | zinc finger pr | Nucleus |
| ENSMUSG00000011008 | 3.91E-14 | 1.42E-13 | Mcoln2 | mucolipin 2 | Plasma Mem |
| ENSMUSG00000014245 | 3.95E-14 | 1.43E-13 | Pigl | phosphatidyl | Cytoplasm |
| ENSMUSG00000030680 | 3.99E-14 | 1.45E-13 | Pagr1a | PAXIP1 assoc | Nucleus |
| ENSMUSG00000019590 | 4.00E-14 | 1.45E-13 | Cyb561 | cytochrome b | Cytoplasm |
| ENSMUSG00000025350 | 4.04E-14 | 1.47E-13 | Rdh5 | retinol dehyd | Cytoplasm |
| ENSMUSG00000007670 | 4.08E-14 | 1.48E-13 | Khsrp | KH-type splic | Nucleus |
| ENSMUSG00000034868 | 4.21E-14 | 1.53E-13 | Myl12b | myosin light | Cytoplasm |
| ENSMUSG00000026730 | 4.28E-14 | 1.55E-13 | Pter | phosphotries | Cytoplasm |
| ENSMUSG00000030180 | 4.28E-14 | 1.55E-13 | Kdm5a | lysine demetl | Nucleus |
| ENSMUSG00000031935 | 4.43E-14 | 1.60E-13 | Med17 | mediator cor | Nucleus |
| ENSMUSG00000026047 | 4.43E-14 | 1.60E-13 | Kdelc1 | KDEL motif c | Cytoplasm |
| ENSMUSG00000003208 | 4.49E-14 | 1.62E-13 | Ccdc94 | YJU2 splicing | Other |
| ENSMUSG00000022434 | 4.51E-14 | 1.63E-13 | Fam118a | family with se | Other |
| ENSMUSG00000022257 | 4.52E-14 | 1.63E-13 | Laptm4b | lysosomal pro | Cytoplasm |
| ENSMUSG00000038403 | 4.54E-14 | 1.64E-13 | Hfe2 | hemojuvelin | Plasma Mem |
| ENSMUSG00000039130 | 4.62E-14 | 1.67E-13 | Zc3hc1 | zinc finger C3 | Nucleus |
| ENSMUSG00000024146 | 4.68E-14 | 1.69E-13 | Cript | CXXC repeat | Cytoplasm |
| ENSMUSG00000026556 | 4.84E-14 | 1.74E-13 | Vangl2 | VANGL plana | Plasma Mem |
| ENSMUSG00000027940 | 4.93E-14 | 1.78E-13 | Tpm3 | tropomyosin | Cytoplasm |
| ENSMUSG00000028797 | 5.06E-14 | 1.82E-13 | Tmem234 | transmembra | Other |

| | | | | | |
|---|---|---|---|---|---|
| ENSMUSG00000031347 | 5.17E-14 | 1.86E-13 | Cetn2 | centrin 2 | Nucleus |
| ENSMUSG00000020089 | 5.30E-14 | 1.91E-13 | Ppa1 | pyrophospha | Cytoplasm |
| ENSMUSG00000017188 | 5.33E-14 | 1.92E-13 | Coa3 | cytochrome ( | Cytoplasm |
| ENSMUSG00000032449 | 5.36E-14 | 1.92E-13 | Slc25a36 | solute carrier | Cytoplasm |
| ENSMUSG00000034708 | 5.36E-14 | 1.93E-13 | Grn | granulin prec | Extracellular S |
| ENSMUSG00000032491 | 5.40E-14 | 1.94E-13 | Nradd | neurotrophin | Extracellular S |
| ENSMUSG00000024740 | 5.41E-14 | 1.94E-13 | Ddb1 | damage spec | Nucleus |
| ENSMUSG00000028478 | 5.47E-14 | 1.96E-13 | Clta | clathrin light | Plasma Mem |
| ENSMUSG00000028099 | 5.53E-14 | 1.98E-13 | Polr3c | RNA polymer | Nucleus |
| ENSMUSG00000022635 | 5.54E-14 | 1.99E-13 | Zcrb1 | zinc finger C( | Nucleus |
| ENSMUSG00000003824 | 5.66E-14 | 2.03E-13 | Syce2 | synaptonema | Nucleus |
| ENSMUSG00000028945 | 5.83E-14 | 2.09E-13 | Rheb | Ras homolog | Cytoplasm |
| ENSMUSG00000028911 | 5.85E-14 | 2.09E-13 | Srsf4 | serine and ar | Nucleus |
| ENSMUSG00000020775 | 6.02E-14 | 2.15E-13 | Mrpl38 | mitochondria | Cytoplasm |
| ENSMUSG00000026516 | 6.13E-14 | 2.19E-13 | Nvl | nuclear VCP- | Nucleus |
| ENSMUSG00000020549 | 6.24E-14 | 2.23E-13 | Elac2 | elaC ribonucl | Nucleus |
| ENSMUSG00000039449 | 6.24E-14 | 2.23E-13 | Prpf18 | pre-mRNA p | Nucleus |
| ENSMUSG00000054034 | 6.26E-14 | 2.23E-13 | Tceal5 | transcription | Other |
| ENSMUSG00000058427 | 6.30E-14 | 2.25E-13 | Cxcl2 | C-X-C motif | Extracellular S |
| ENSMUSG00000029162 | 6.31E-14 | 2.25E-13 | Khk | ketohexokina | Cytoplasm |
| ENSMUSG00000055720 | 6.46E-14 | 2.30E-13 | Ubl7 | ubiquitin like | Other |
| ENSMUSG00000047638 | 6.48E-14 | 2.31E-13 | Nr1h4 | nuclear recep | Nucleus |
| ENSMUSG00000079042 | 6.51E-14 | 2.32E-13 | Apela | apelin recept | Extracellular S |
| ENSMUSG00000021576 | 6.53E-14 | 2.33E-13 | Pdcd6 | programmed | Cytoplasm |
| ENSMUSG00000022436 | 6.58E-14 | 2.34E-13 | Sh3bp1 | SH3 domain | Plasma Mem |
| ENSMUSG00000040725 | 6.60E-14 | 2.35E-13 | Hnrnpul1 | heterogeneo | Nucleus |
| ENSMUSG00000031617 | 6.60E-14 | 2.35E-13 | Tmem184c | transmembra | Other |
| ENSMUSG00000060803 | 6.67E-14 | 2.37E-13 | Gstp1 | glutathione S | Cytoplasm |
| ENSMUSG00000002731 | 6.67E-14 | 2.37E-13 | Prkra | protein activa | Cytoplasm |
| ENSMUSG00000022553 | 6.74E-14 | 2.39E-13 | Maf1 | MAF1 homol | Nucleus |
| ENSMUSG00000028672 | 6.74E-14 | 2.39E-13 | Hmgcl | 3-hydroxy-3 | Cytoplasm |
| ENSMUSG00000023330 | 7.10E-14 | 2.52E-13 | Dtwd1 | DTW domain | Other |
| ENSMUSG00000022820 | 7.14E-14 | 2.54E-13 | Ndufb4 | NADH:ubiqu | Cytoplasm |
| ENSMUSG00000019961 | 7.17E-14 | 2.54E-13 | Tmpo | thymopoietin | Nucleus |
| ENSMUSG00000020528 | 7.21E-14 | 2.56E-13 | Prpsap2 | phosphoribo | Other |
| ENSMUSG00000030879 | 7.23E-14 | 2.56E-13 | Mrpl17 | mitochondria | Cytoplasm |
| ENSMUSG00000031883 | 7.26E-14 | 2.57E-13 | Car7 | carbonic anh | Cytoplasm |
| ENSMUSG00000022120 | 7.39E-14 | 2.62E-13 | Rnf219 | ring finger pr | Other |
| ENSMUSG00000055137 | 7.48E-14 | 2.65E-13 | Sugct | succinyl-CoA | Cytoplasm |
| ENSMUSG00000027424 | 7.55E-14 | 2.67E-13 | Mgme1 | mitochondria | Cytoplasm |
| ENSMUSG00000033543 | 7.61E-14 | 2.69E-13 | Gtf2a2 | general trans | Nucleus |
| ENSMUSG00000018921 | 7.77E-14 | 2.74E-13 | Pelp1 | proline, gluta | Nucleus |
| ENSMUSG00000031429 | 7.79E-14 | 2.75E-13 | Psmd10 | proteasome ( | Cytoplasm |
| ENSMUSG00000034543 | 7.87E-14 | 2.78E-13 | Morc2a | MORC family | Nucleus |
| ENSMUSG00000035783 | 7.94E-14 | 2.80E-13 | Acta2 | actin, alpha 2 | Cytoplasm |
| ENSMUSG00000029422 | 8.03E-14 | 2.83E-13 | Rsrc2 | arginine/serin | Other |
| ENSMUSG00000038525 | 8.07E-14 | 2.85E-13 | Armc10 | armadillo rep | Cytoplasm |
| ENSMUSG00000036777 | 8.14E-14 | 2.87E-13 | Anln | anillin actin b | Cytoplasm |
| ENSMUSG00000047721 | 8.16E-14 | 2.87E-13 | Bola2 | bolA-like 2 (E | Cytoplasm |
| ENSMUSG00000096145 | 8.49E-14 | 2.99E-13 | Vkorc1 | vitamin K epo | Cytoplasm |
| ENSMUSG00000047757 | 8.91E-14 | 3.14E-13 | Fancb | FA complemen | Nucleus |
| ENSMUSG00000023075 | 8.97E-14 | 3.16E-13 | Akirin1 | akirin 1 | Cytoplasm |
| ENSMUSG00000028683 | 8.98E-14 | 3.16E-13 | Eif2b3 | eukaryotic tra | Cytoplasm |
| ENSMUSG00000045282 | 9.11E-14 | 3.20E-13 | Tmem86b | transmembra | Cytoplasm |
| ENSMUSG00000022844 | 9.37E-14 | 3.29E-13 | Pdia5 | protein disulf | Cytoplasm |
| ENSMUSG00000030471 | 9.52E-14 | 3.34E-13 | Zdhhc13 | zinc finger DI | Nucleus |
| ENSMUSG00000003809 | 9.52E-14 | 3.34E-13 | Gcdh | glutaryl-CoA | Cytoplasm |
| ENSMUSG00000036875 | 9.87E-14 | 3.46E-13 | Dna2 | DNA replicat | Cytoplasm |

| | | | | | |
|---|---|---|---|---|---|
| ENSMUSG00000027562 | 9.92E-14 | 3.48E-13 | Car2 | carbonic anh | Cytoplasm |
| ENSMUSG00000011306 | 9.93E-14 | 3.48E-13 | Sugp1 | SURP and G- | Nucleus |
| ENSMUSG00000029782 | 1.00E-13 | 3.52E-13 | Tmem209 | transmembra | Other |
| ENSMUSG00000051675 | 1.02E-13 | 3.55E-13 | Trim32 | tripartite mot | Nucleus |
| ENSMUSG00000041740 | 1.05E-13 | 3.67E-13 | Rnf10 | ring finger pr | Cytoplasm |
| ENSMUSG00000031386 | 1.05E-13 | 3.68E-13 | Hcfc1 | host cell fact | Nucleus |
| ENSMUSG00000008892 | 1.07E-13 | 3.74E-13 | Vdac3 | voltage depe | Cytoplasm |
| ENSMUSG00000022035 | 1.11E-13 | 3.88E-13 | Ccdc25 | coiled-coil d | Cytoplasm |
| ENSMUSG00000025352 | 1.13E-13 | 3.94E-13 | Gdf11 | growth differ | Extracellular S |
| ENSMUSG00000058440 | 1.14E-13 | 3.96E-13 | Nrf1 | nuclear respi | Nucleus |
| ENSMUSG00000012535 | 1.16E-13 | 4.05E-13 | Tnpo3 | transportin 3 | Cytoplasm |
| ENSMUSG00000019470 | 1.17E-13 | 4.10E-13 | Xab2 | XPA binding | Nucleus |
| ENSMUSG00000033106 | 1.22E-13 | 4.26E-13 | Slc7a6os | solute carrier | Other |
| ENSMUSG00000068329 | 1.24E-13 | 4.31E-13 | Htra2 | HtrA serine p | Cytoplasm |
| ENSMUSG00000050912 | 1.25E-13 | 4.34E-13 | Tmem123 | transmembra | Plasma Mem |
| ENSMUSG00000014859 | 1.27E-13 | 4.42E-13 | E2f4 | E2F transcrip | Nucleus |
| ENSMUSG00000025504 | 1.29E-13 | 4.49E-13 | Eps8l2 | EPS8 like 2 | Cytoplasm |
| ENSMUSG00000030393 | 1.29E-13 | 4.50E-13 | Zik1 | zinc finger pr | Other |
| ENSMUSG00000026806 | 1.34E-13 | 4.66E-13 | Ddx31 | DEAD-box h | Nucleus |
| ENSMUSG00000031629 | 1.36E-13 | 4.72E-13 | Cenpu | centromere p | Nucleus |
| ENSMUSG00000020368 | 1.36E-13 | 4.74E-13 | Canx | calnexin | Cytoplasm |
| ENSMUSG00000028419 | 1.37E-13 | 4.75E-13 | Chmp5 | charged mult | Cytoplasm |
| ENSMUSG00000000901 | 1.37E-13 | 4.75E-13 | Mmp11 | matrix metall | Extracellular S |
| ENSMUSG00000058886 | 1.40E-13 | 4.86E-13 | Deaf1 | DEAF1, trans | Nucleus |
| ENSMUSG00000046179 | 1.44E-13 | 4.98E-13 | E2f8 | E2F transcrip | Nucleus |
| ENSMUSG00000012296 | 1.44E-13 | 4.98E-13 | Tjap1 | tight junction | Plasma Mem |
| ENSMUSG00000001380 | 1.44E-13 | 4.99E-13 | Hars | histidyl-tRNA | Cytoplasm |
| ENSMUSG00000010205 | 1.45E-13 | 5.01E-13 | Raver1 | ribonucleopr | Nucleus |
| ENSMUSG00000028580 | 1.45E-13 | 5.02E-13 | Pum1 | pumilio RNA | Cytoplasm |
| ENSMUSG00000021707 | 1.49E-13 | 5.15E-13 | Dhfr | dihydrofolate | Nucleus |
| ENSMUSG00000002455 | 1.49E-13 | 5.17E-13 | Prpf6 | pre-mRNA p | Nucleus |
| ENSMUSG00000060510 | 1.50E-13 | 5.19E-13 | Zfp266 | zinc finger pr | Nucleus |
| ENSMUSG00000027331 | 1.51E-13 | 5.22E-13 | Knstrn | kinetochore l | Cytoplasm |
| ENSMUSG00000021820 | 1.52E-13 | 5.25E-13 | Camk2g | calcium/calm | Cytoplasm |
| ENSMUSG00000025856 | 1.57E-13 | 5.41E-13 | Pdgfa | platelet deriv | Extracellular S |
| ENSMUSG00000028990 | 1.58E-13 | 5.44E-13 | Lzic | leucine zippe | Other |
| ENSMUSG00000049086 | 1.59E-13 | 5.47E-13 | Bmyc | brain express | Nucleus |
| ENSMUSG00000061904 | 1.61E-13 | 5.55E-13 | Slc25a3 | solute carrier | Cytoplasm |
| ENSMUSG00000021124 | 1.65E-13 | 5.68E-13 | Vti1b | vesicle transp | Plasma Mem |
| ENSMUSG00000031575 | 1.66E-13 | 5.73E-13 | Ash2l | ASH2 like, his | Nucleus |
| ENSMUSG00000020608 | 1.70E-13 | 5.86E-13 | Smc6 | structural ma | Nucleus |
| ENSMUSG00000025162 | 1.72E-13 | 5.93E-13 | Csnk1d | casein kinase | Cytoplasm |
| ENSMUSG00000036097 | 1.74E-13 | 6.00E-13 | Fam178a | SMC5-SMC6 | Extracellular S |
| ENSMUSG00000027498 | 1.78E-13 | 6.12E-13 | Cstf1 | cleavage stim | Nucleus |
| ENSMUSG00000020288 | 1.78E-13 | 6.13E-13 | Ahsa2 | AHA1, activa | Other |
| ENSMUSG00000024576 | 1.82E-13 | 6.24E-13 | Csnk1a1 | casein kinase | Cytoplasm |
| ENSMUSG00000018377 | 1.83E-13 | 6.29E-13 | Vezf1 | vascular endo | Nucleus |
| ENSMUSG00000000301 | 1.85E-13 | 6.37E-13 | Pemt | phosphatidyl | Cytoplasm |
| ENSMUSG00000032977 | 1.89E-13 | 6.49E-13 | Fam207a | family with se | Other |
| ENSMUSG00000037351 | 1.93E-13 | 6.62E-13 | Actr1b | ARP1 actin re | Cytoplasm |
| ENSMUSG00000026926 | 1.94E-13 | 6.65E-13 | Pmpca | peptidase, m | Cytoplasm |
| ENSMUSG00000039182 | 1.96E-13 | 6.72E-13 | AW209491 | chromosome | Other |
| ENSMUSG00000060098 | 1.98E-13 | 6.80E-13 | Prmt7 | protein argin | Cytoplasm |
| ENSMUSG00000037339 | 1.99E-13 | 6.80E-13 | Fam53a | family with se | Nucleus |
| ENSMUSG00000056612 | 2.02E-13 | 6.91E-13 | Ppp1r14b | protein phos | Cytoplasm |
| ENSMUSG00000026870 | 2.02E-13 | 6.92E-13 | Cutal | cutA divalent | Other |
| ENSMUSG00000038762 | 2.03E-13 | 6.93E-13 | Abcf1 | ATP binding | Cytoplasm |
| ENSMUSG00000003923 | 2.05E-13 | 7.01E-13 | Tfam | transcription | Cytoplasm |

| | | | | | |
|---|---|---|---|---|---|
| ENSMUSG00000037851 | 2.07E-13 | 7.08E-13 | Iars | isoleucyl-tRN | Cytoplasm |
| ENSMUSG00000027131 | 2.15E-13 | 7.36E-13 | Emc4 | ER membran | Cytoplasm |
| ENSMUSG00000035399 | 2.18E-13 | 7.46E-13 | Oser1 | oxidative stre | Other |
| ENSMUSG00000036398 | 2.29E-13 | 7.81E-13 | Ppp1r11 | protein phos | Cytoplasm |
| ENSMUSG00000092008 | 2.31E-13 | 7.87E-13 | Cyp2c69 | cytochrome I | Cytoplasm |
| ENSMUSG00000059586 | 2.31E-13 | 7.88E-13 | Nsmce2 | NSE2 (MMS2 | Nucleus |
| ENSMUSG00000023015 | 2.35E-13 | 8.00E-13 | Racgap1 | Rac GTPase a | Cytoplasm |
| ENSMUSG00000018040 | 2.37E-13 | 8.07E-13 | Rrp7a | ribosomal RN | Cytoplasm |
| ENSMUSG00000028907 | 2.39E-13 | 8.16E-13 | Utp11l | UTP11, small | Nucleus |
| ENSMUSG00000031934 | 2.52E-13 | 8.56E-13 | Panx1 | pannexin 1 | Plasma Mem |
| ENSMUSG00000051674 | 2.55E-13 | 8.67E-13 | Dcun1d4 | defective in c | Nucleus |
| ENSMUSG00000033099 | 2.56E-13 | 8.69E-13 | Nol12 | nucleolar pro | Nucleus |
| ENSMUSG00000028577 | 2.56E-13 | 8.70E-13 | Plaa | phospholipas | Cytoplasm |
| ENSMUSG00000041781 | 2.57E-13 | 8.72E-13 | Cpsf2 | cleavage and | Nucleus |
| ENSMUSG00000048277 | 2.60E-13 | 8.82E-13 | Syngr2 | synaptogyrin | Cytoplasm |
| ENSMUSG00000027247 | 2.61E-13 | 8.85E-13 | Arhgap1 | Rho GTPase a | Cytoplasm |
| ENSMUSG00000037852 | 2.62E-13 | 8.88E-13 | Cpe | carboxypepti | Cytoplasm |
| ENSMUSG00000020253 | 2.62E-13 | 8.89E-13 | Ppm1m | protein phos | Nucleus |
| ENSMUSG00000047714 | 2.71E-13 | 9.18E-13 | Ppp1r2 | protein phos | Cytoplasm |
| ENSMUSG00000024384 | 2.72E-13 | 9.22E-13 | Iws1 | IWS1, SUPT6 | Nucleus |
| ENSMUSG00000035674 | 2.72E-13 | 9.22E-13 | Ndufa3 | NADH:ubiqu | Cytoplasm |
| ENSMUSG00000032580 | 2.78E-13 | 9.40E-13 | Rbm5 | RNA binding | Nucleus |
| ENSMUSG00000026034 | 2.79E-13 | 9.45E-13 | Clk1 | CDC like kina | Nucleus |
| ENSMUSG00000027428 | 2.82E-13 | 9.54E-13 | Rbbp9 | RB binding p | Nucleus |
| ENSMUSG00000031527 | 2.92E-13 | 9.86E-13 | Eri1 | exoribonucle | Nucleus |
| ENSMUSG00000030319 | 2.99E-13 | 1.01E-12 | Cand2 | cullin associa | Nucleus |
| ENSMUSG00000061906 | 3.09E-13 | 1.04E-12 | Ugt2b38 | UDP glucuro | Cytoplasm |
| ENSMUSG00000025786 | 3.13E-13 | 1.06E-12 | Zdhhc3 | zinc finger D | Cytoplasm |
| ENSMUSG00000019214 | 3.15E-13 | 1.06E-12 | Chtf18 | chromosome | Nucleus |
| ENSMUSG00000070372 | 3.17E-13 | 1.07E-12 | Capza1 | capping actir | Cytoplasm |
| ENSMUSG00000028702 | 3.28E-13 | 1.11E-12 | Rad54l | RAD54 like | Nucleus |
| ENSMUSG00000029472 | 3.35E-13 | 1.13E-12 | Anapc5 | anaphase pro | Nucleus |
| ENSMUSG00000092607 | 3.36E-13 | 1.13E-12 | Scnm1 | sodium chan | Nucleus |
| ENSMUSG00000006392 | 3.39E-13 | 1.14E-12 | Med8 | mediator cor | Nucleus |
| ENSMUSG00000025133 | 3.49E-13 | 1.18E-12 | Ints4 | integrator co | Nucleus |
| ENSMUSG00000027889 | 3.51E-13 | 1.18E-12 | Ampd2 | adenosine m | Cytoplasm |
| ENSMUSG00000020432 | 3.53E-13 | 1.19E-12 | Tcn2 | transcobalam | Extracellular S |
| ENSMUSG00000010755 | 3.53E-13 | 1.19E-12 | Cars | cysteinyl-tRN | Cytoplasm |
| ENSMUSG00000030298 | 3.57E-13 | 1.20E-12 | Sec13 | SEC13 homo | Cytoplasm |
| ENSMUSG00000031586 | 3.77E-13 | 1.27E-12 | Rbpms | RNA binding | Cytoplasm |
| ENSMUSG00000021952 | 3.79E-13 | 1.27E-12 | Xpo4 | exportin 4 | Nucleus |
| ENSMUSG00000042453 | 4.14E-13 | 1.39E-12 | Reln | reelin | Extracellular S |
| ENSMUSG00000045294 | 4.35E-13 | 1.46E-12 | Insig1 | insulin induc | Cytoplasm |
| ENSMUSG00000029128 | 4.49E-13 | 1.51E-12 | Rab28 | RAB28, mem | Plasma Mem |
| ENSMUSG00000024283 | 4.56E-13 | 1.53E-12 | Wac | WW domain | Nucleus |
| ENSMUSG00000040785 | 4.68E-13 | 1.57E-12 | Ttc3 | tetratricopep | Cytoplasm |
| ENSMUSG00000074698 | 4.85E-13 | 1.62E-12 | Csnk2a1 | casein kinase | Nucleus |
| ENSMUSG00000020536 | 4.85E-13 | 1.62E-12 | Llgl1 | LLGL1, scribb | Cytoplasm |
| ENSMUSG00000027722 | 4.91E-13 | 1.64E-12 | Spata5 | spermatogen | Cytoplasm |
| ENSMUSG00000041133 | 4.94E-13 | 1.65E-12 | Smc1a | structural ma | Nucleus |
| ENSMUSG00000034773 | 4.95E-13 | 1.65E-12 | BC030867 | chromosome | Other |
| ENSMUSG00000020078 | 5.05E-13 | 1.69E-12 | Vps26a | VPS26, retror | Cytoplasm |
| ENSMUSG00000029111 | 5.08E-13 | 1.70E-12 | Nelfa | negative elor | Nucleus |
| ENSMUSG00000032212 | 5.09E-13 | 1.70E-12 | Sltm | SAFB like trar | Nucleus |
| ENSMUSG00000039768 | 5.12E-13 | 1.71E-12 | Dnajc11 | DnaJ heat sh | Cytoplasm |
| ENSMUSG00000022261 | 5.43E-13 | 1.81E-12 | Sdc2 | syndecan 2 | Plasma Mem |
| ENSMUSG00000020630 | 5.58E-13 | 1.86E-12 | Rnaseh1 | ribonuclease | Nucleus |
| ENSMUSG00000042305 | 5.61E-13 | 1.87E-12 | Tmem183a | transmembra | Other |

| | | | | | |
|---|---|---|---|---|---|
| ENSMUSG00000028165 | 5.67E-13 | 1.89E-12 | Cisd2 | CDGSH iron : | Cytoplasm |
| ENSMUSG00000036372 | 5.68E-13 | 1.89E-12 | Tmem258 | transmembra | Other |
| ENSMUSG00000062169 | 5.69E-13 | 1.89E-12 | Cnih4 | cornichon fai | Plasma Mem |
| ENSMUSG00000039994 | 5.74E-13 | 1.91E-12 | Timeless | timeless circa | Nucleus |
| ENSMUSG00000063145 | 5.88E-13 | 1.96E-12 | Bbs5 | Bardet-Biedl | Cytoplasm |
| ENSMUSG00000057561 | 5.89E-13 | 1.96E-12 | Eif1a | eukaryotic tra | Cytoplasm |
| ENSMUSG00000057134 | 5.95E-13 | 1.98E-12 | Ado | 2-aminoetha | Cytoplasm |
| ENSMUSG00000010911 | 5.98E-13 | 1.99E-12 | Apip | APAF1 intera | Cytoplasm |
| ENSMUSG00000054604 | 5.99E-13 | 1.99E-12 | Cggbp1 | CGG triplet re | Nucleus |
| ENSMUSG00000062203 | 6.21E-13 | 2.06E-12 | Gspt1 | G1 to S phase | Cytoplasm |
| ENSMUSG00000038429 | 6.39E-13 | 2.12E-12 | Usp5 | ubiquitin spe | Cytoplasm |
| ENSMUSG00000025413 | 6.66E-13 | 2.21E-12 | Ttc4 | tetratricopep | Other |
| ENSMUSG00000070923 | 6.78E-13 | 2.25E-12 | Klhl9 | kelch like fam | Cytoplasm |
| ENSMUSG00000054836 | 6.86E-13 | 2.27E-12 | Elp6 | elongator acc | Other |
| ENSMUSG00000026922 | 6.90E-13 | 2.29E-12 | Agpat2 | 1-acylglycerc | Cytoplasm |
| ENSMUSG00000029191 | 6.96E-13 | 2.31E-12 | Rfc1 | replication fa | Nucleus |
| ENSMUSG00000023971 | 7.26E-13 | 2.40E-12 | Rrp36 | ribosomal RN | Nucleus |
| ENSMUSG00000028233 | 7.33E-13 | 2.43E-12 | Tgs1 | trimethylguai | Nucleus |
| ENSMUSG00000028988 | 7.33E-13 | 2.43E-12 | Ctnnbip1 | catenin beta | Cytoplasm |
| ENSMUSG00000025040 | 7.56E-13 | 2.50E-12 | Fundc1 | FUN14 doma | Cytoplasm |
| ENSMUSG00000002289 | 7.71E-13 | 2.55E-12 | Angptl4 | angiopoietin | Extracellular S |
| ENSMUSG00000024423 | 7.73E-13 | 2.56E-12 | Impact | impact RWD | Cytoplasm |
| ENSMUSG00000047710 | 7.76E-13 | 2.56E-12 | Champ1 | chromosome | Nucleus |
| ENSMUSG00000020098 | 7.79E-13 | 2.57E-12 | Pcbd1 | pterin-4 alph | Nucleus |
| ENSMUSG00000018651 | 8.06E-13 | 2.66E-12 | Tada2a | transcriptiona | Nucleus |
| ENSMUSG00000027285 | 8.08E-13 | 2.67E-12 | Haus2 | HAUS augmi | Cytoplasm |
| ENSMUSG00000048100 | 8.11E-13 | 2.68E-12 | Taf13 | TATA-box bi | Nucleus |
| ENSMUSG00000036550 | 8.27E-13 | 2.73E-12 | Cnot1 | CCR4-NOT tr | Cytoplasm |
| ENSMUSG00000031592 | 8.89E-13 | 2.93E-12 | Pcm1 | pericentriolar | Cytoplasm |
| ENSMUSG00000016757 | 8.95E-13 | 2.95E-12 | Ttll12 | tubulin tyrosi | Other |
| ENSMUSG00000034158 | 8.98E-13 | 2.95E-12 | Lrrc58 | leucine rich r | Other |
| ENSMUSG00000026761 | 8.98E-13 | 2.95E-12 | Orc4 | origin recogr | Nucleus |
| ENSMUSG00000034108 | 9.11E-13 | 3.00E-12 | Ccs | copper chape | Cytoplasm |
| ENSMUSG00000031201 | 9.18E-13 | 3.02E-12 | Brcc3 | BRCA1/BRCA | Nucleus |
| ENSMUSG00000038656 | 9.28E-13 | 3.05E-12 | Cyp3a16 | cytochrome I | Cytoplasm |
| ENSMUSG00000019804 | 9.45E-13 | 3.10E-12 | Snx3 | sorting nexin | Cytoplasm |
| ENSMUSG00000026856 | 9.65E-13 | 3.17E-12 | Dolpp1 | dolichyldipho | Cytoplasm |
| ENSMUSG00000001833 | 9.71E-13 | 3.19E-12 | Sept7 | septin 7 | Cytoplasm |
| ENSMUSG00000024098 | 9.79E-13 | 3.21E-12 | Twsg1 | twisted gastr | Extracellular S |
| ENSMUSG00000054967 | 1.03E-12 | 3.38E-12 | Zfp647 | zinc finger pr | Nucleus |
| ENSMUSG00000020869 | 1.04E-12 | 3.40E-12 | Lrrc59 | leucine rich r | Cytoplasm |
| ENSMUSG00000021771 | 1.10E-12 | 3.60E-12 | Vdac2 | voltage depe | Cytoplasm |
| ENSMUSG00000021241 | 1.12E-12 | 3.66E-12 | Isca2 | iron-sulfur cl | Cytoplasm |
| ENSMUSG00000025993 | 1.12E-12 | 3.66E-12 | Slc40a1 | solute carrier | Plasma Mem |
| ENSMUSG00000011257 | 1.12E-12 | 3.67E-12 | Pabpc4 | poly(A) bindi | Cytoplasm |
| ENSMUSG00000026790 | 1.13E-12 | 3.70E-12 | Odf2 | outer dense f | Cytoplasm |
| ENSMUSG00000026429 | 1.13E-12 | 3.70E-12 | Ube2t | ubiquitin cor | Nucleus |
| ENSMUSG00000039630 | 1.16E-12 | 3.79E-12 | Hnrnpu | heterogeneo | Nucleus |
| ENSMUSG00000028861 | 1.18E-12 | 3.86E-12 | Mrps15 | mitochondria | Cytoplasm |
| ENSMUSG00000020390 | 1.20E-12 | 3.92E-12 | Ube2b | ubiquitin cor | Cytoplasm |
| ENSMUSG00000020650 | 1.20E-12 | 3.92E-12 | Bcap29 | B cell recepto | Cytoplasm |
| ENSMUSG00000021671 | 1.21E-12 | 3.94E-12 | Poc5 | POC5 centric | Cytoplasm |
| ENSMUSG00000020290 | 1.22E-12 | 3.97E-12 | Xpo1 | exportin 1 | Nucleus |
| ENSMUSG00000024870 | 1.23E-12 | 4.00E-12 | Rab1b | RAB1B, mem | Cytoplasm |
| ENSMUSG00000036980 | 1.24E-12 | 4.05E-12 | Taf6 | TATA-box bi | Nucleus |
| ENSMUSG00000017386 | 1.25E-12 | 4.09E-12 | Traf4 | TNF receptor | Cytoplasm |
| ENSMUSG00000022201 | 1.33E-12 | 4.34E-12 | Zfr | zinc finger RN | Nucleus |
| ENSMUSG00000034274 | 1.34E-12 | 4.35E-12 | Thoc5 | THO comple: | Cytoplasm |

| | | | | | |
|---|---|---|---|---|---|
| ENSMUSG00000007987 | 1.36E-12 | 4.43E-12 | Ift22 | intraflagellar | Extracellular S |
| ENSMUSG00000026662 | 1.37E-12 | 4.47E-12 | Sephs1 | selenophosp | Other |
| ENSMUSG00000006599 | 1.38E-12 | 4.50E-12 | Gtf2h1 | general trans | Nucleus |
| ENSMUSG00000022685 | 1.41E-12 | 4.59E-12 | Parn | poly(A)-spec | Nucleus |
| ENSMUSG00000026484 | 1.44E-12 | 4.68E-12 | Rnf2 | ring finger pr | Nucleus |
| ENSMUSG00000011096 | 1.45E-12 | 4.71E-12 | Akt1s1 | AKT1 substra | Cytoplasm |
| ENSMUSG00000027881 | 1.46E-12 | 4.74E-12 | Prpf38b | pre-mRNA p | Other |
| ENSMUSG00000049401 | 1.47E-12 | 4.77E-12 | Ogfr | opioid growt | Plasma Mem |
| ENSMUSG00000019794 | 1.48E-12 | 4.82E-12 | Katna1 | katanin catal | Cytoplasm |
| ENSMUSG00000005899 | 1.50E-12 | 4.85E-12 | Smpd4 | sphingomyel | Cytoplasm |
| ENSMUSG00000033931 | 1.50E-12 | 4.85E-12 | Rbm34 | RNA binding | Nucleus |
| ENSMUSG00000022855 | 1.52E-12 | 4.91E-12 | Senp2 | SUMO specif | Nucleus |
| ENSMUSG00000031095 | 1.52E-12 | 4.92E-12 | Cul4b | cullin 4B | Nucleus |
| ENSMUSG00000026211 | 1.53E-12 | 4.94E-12 | Obsl1 | obscurin-like | Other |
| ENSMUSG00000039323 | 1.57E-12 | 5.09E-12 | Igfbp2 | insulin like gr | Extracellular S |
| ENSMUSG00000066613 | 1.58E-12 | 5.10E-12 | Zfp932 | zinc finger pr | Nucleus |
| ENSMUSG00000025196 | 1.61E-12 | 5.22E-12 | Cpn1 | carboxypepti | Extracellular S |
| ENSMUSG00000026709 | 1.62E-12 | 5.24E-12 | Dars2 | aspartyl-tRN | Cytoplasm |
| ENSMUSG00000075054 | 1.64E-12 | 5.30E-12 | Yae1d1 | YAE1, ABCE1 | Other |
| ENSMUSG00000038611 | 1.65E-12 | 5.34E-12 | Phrf1 | PHD and ring | Nucleus |
| ENSMUSG00000020516 | 1.67E-12 | 5.38E-12 | Rps6kb1 | ribosomal pr | Cytoplasm |
| ENSMUSG00000045624 | 1.68E-12 | 5.43E-12 | Esf1 | ESF1 nucleol | Nucleus |
| ENSMUSG00000020612 | 1.70E-12 | 5.47E-12 | Prkar1a | protein kinas | Cytoplasm |
| ENSMUSG00000051557 | 1.70E-12 | 5.49E-12 | Pusl1 | pseudouridyl | Other |
| ENSMUSG00000030946 | 1.72E-12 | 5.55E-12 | Lhpp | phospholysir | Cytoplasm |
| ENSMUSG00000048930 | 1.73E-12 | 5.56E-12 | Tada3 | transcriptiona | Nucleus |
| ENSMUSG00000056812 | 1.78E-12 | 5.74E-12 | St8sia3 | ST8 alpha-N | Cytoplasm |
| ENSMUSG00000029034 | 1.80E-12 | 5.80E-12 | Cpsf3l | integrator co | Nucleus |
| ENSMUSG00000002733 | 1.84E-12 | 5.91E-12 | Plekha3 | pleckstrin ho | Cytoplasm |
| ENSMUSG00000030880 | 1.84E-12 | 5.91E-12 | Polr3e | RNA polymer | Nucleus |
| ENSMUSG00000021488 | 1.90E-12 | 6.10E-12 | Nsd1 | nuclear recep | Nucleus |
| ENSMUSG00000026927 | 1.91E-12 | 6.15E-12 | Sdccag3 | endosome as | Cytoplasm |
| ENSMUSG00000031922 | 1.92E-12 | 6.16E-12 | Cep57 | centrosomal | Cytoplasm |
| ENSMUSG00000031392 | 1.92E-12 | 6.17E-12 | Irak1 | interleukin 1 | Plasma Mem |
| ENSMUSG00000024078 | 1.94E-12 | 6.21E-12 | Ttc27 | tetratricopep | Other |
| ENSMUSG00000035104 | 2.04E-12 | 6.52E-12 | Eva1a | eva-1 homol | Plasma Mem |
| ENSMUSG00000062580 | 2.05E-12 | 6.58E-12 | Timm17a | translocase o | Cytoplasm |
| ENSMUSG00000028224 | 2.12E-12 | 6.79E-12 | Nbn | nibrin | Nucleus |
| ENSMUSG00000063904 | 2.13E-12 | 6.82E-12 | Dpp3 | dipeptidyl pe | Cytoplasm |
| ENSMUSG00000021713 | 2.13E-12 | 6.82E-12 | Ppwd1 | peptidylproly | Nucleus |
| ENSMUSG00000028643 | 2.20E-12 | 7.05E-12 | Ccdc23 | small vasohib | Other |
| ENSMUSG00000029522 | 2.21E-12 | 7.07E-12 | Pla2g1b | phospholipas | Extracellular S |
| ENSMUSG00000034748 | 2.24E-12 | 7.16E-12 | Sirt6 | sirtuin 6 | Nucleus |
| ENSMUSG00000040269 | 2.25E-12 | 7.19E-12 | Mrps28 | mitochondria | Cytoplasm |
| ENSMUSG00000054999 | 2.25E-12 | 7.20E-12 | Naaladl1 | N-acetylated | Plasma Mem |
| ENSMUSG00000018974 | 2.29E-12 | 7.31E-12 | Sart3 | squamous ce | Nucleus |
| ENSMUSG00000026112 | 2.30E-12 | 7.35E-12 | Coa5 | cytochrome c | Cytoplasm |
| ENSMUSG00000029198 | 2.31E-12 | 7.37E-12 | Grpel1 | GrpE like 1, n | Cytoplasm |
| ENSMUSG00000020402 | 2.34E-12 | 7.47E-12 | Vdac1 | voltage depe | Cytoplasm |
| ENSMUSG00000025337 | 2.35E-12 | 7.49E-12 | Sbds | SBDS, riboso | Nucleus |
| ENSMUSG00000022034 | 2.38E-12 | 7.58E-12 | Esco2 | establishmen | Nucleus |
| ENSMUSG00000006673 | 2.41E-12 | 7.68E-12 | Qrich1 | glutamine ric | Nucleus |
| ENSMUSG00000069089 | 2.42E-12 | 7.71E-12 | Cdk7 | cyclin depend | Nucleus |
| ENSMUSG00000032783 | 2.44E-12 | 7.76E-12 | Troap | trophinin ass | Cytoplasm |
| ENSMUSG00000021670 | 2.45E-12 | 7.81E-12 | Hmgcr | 3-hydroxy-3 | Cytoplasm |
| ENSMUSG00000021486 | 2.48E-12 | 7.88E-12 | Prelid1 | PRELI domair | Cytoplasm |
| ENSMUSG00000031661 | 2.49E-12 | 7.90E-12 | Nkd1 | naked cuticle | Other |
| ENSMUSG00000025228 | 2.55E-12 | 8.11E-12 | Actr1a | ARP1 actin re | Cytoplasm |

| | | | | | |
|---|---|---|---|---|---|
| ENSMUSG00000022462 | 2.56E-12 | 8.12E-12 | Slc38a2 | solute carrier | Plasma Mem |
| ENSMUSG00000025173 | 2.57E-12 | 8.15E-12 | Wdr45b | WD repeat d | Other |
| ENSMUSG00000009535 | 2.61E-12 | 8.29E-12 | Rnmt | RNA guanine | Nucleus |
| ENSMUSG00000006345 | 2.62E-12 | 8.33E-12 | Ggt1 | gamma-glut | Plasma Mem |
| ENSMUSG00000027933 | 2.65E-12 | 8.40E-12 | Ints3 | integrator co | Nucleus |
| ENSMUSG00000037531 | 2.71E-12 | 8.58E-12 | Mrpl47 | mitochondria | Cytoplasm |
| ENSMUSG00000031886 | 2.72E-12 | 8.64E-12 | Ces2e | carboxyleste | Other |
| ENSMUSG00000031787 | 2.74E-12 | 8.67E-12 | Katnb1 | katanin regul | Cytoplasm |
| ENSMUSG00000067847 | 2.77E-12 | 8.78E-12 | Romo1 | reactive oxyg | Cytoplasm |
| ENSMUSG00000037275 | 2.79E-12 | 8.83E-12 | Gemin5 | gem nuclear | Nucleus |
| ENSMUSG00000032192 | 2.79E-12 | 8.85E-12 | Gnb5 | G protein sul | Plasma Mem |
| ENSMUSG00000002748 | 2.95E-12 | 9.33E-12 | Baz1b | bromodomai | Nucleus |
| ENSMUSG00000032480 | 2.98E-12 | 9.41E-12 | Dhx30 | DExH-box he | Nucleus |
| ENSMUSG00000005682 | 2.99E-12 | 9.45E-12 | Pan2 | poly(A) speci | Cytoplasm |
| ENSMUSG00000015335 | 3.02E-12 | 9.53E-12 | Zdhhc12 | zinc finger D | Other |
| ENSMUSG00000027364 | 3.08E-12 | 9.72E-12 | Usp50 | ubiquitin spe | Other |
| ENSMUSG00000038806 | 3.19E-12 | 1.01E-11 | Sde2 | SDE2 telome | Other |
| ENSMUSG00000024081 | 3.21E-12 | 1.01E-11 | Cebpz | CCAAT enha | Nucleus |
| ENSMUSG00000026434 | 3.26E-12 | 1.03E-11 | Nucks1 | nuclear casei | Nucleus |
| ENSMUSG00000021193 | 3.26E-12 | 1.03E-11 | Pitrm1 | pitrilysin met | Cytoplasm |
| ENSMUSG00000038965 | 3.32E-12 | 1.05E-11 | Ube2l3 | ubiquitin cor | Nucleus |
| ENSMUSG00000045328 | 3.36E-12 | 1.06E-11 | Cenpe | centromere p | Nucleus |
| ENSMUSG00000027107 | 3.47E-12 | 1.09E-11 | Chrna1 | cholinergic re | Plasma Mem |
| ENSMUSG00000021484 | 3.54E-12 | 1.12E-11 | Lman2 | lectin, mannc | Cytoplasm |
| ENSMUSG00000030541 | 3.60E-12 | 1.13E-11 | Idh2 | isocitrate del | Cytoplasm |
| ENSMUSG00000034484 | 3.71E-12 | 1.17E-11 | Snx2 | sorting nexin | Cytoplasm |
| ENSMUSG00000053768 | 3.77E-12 | 1.19E-11 | Chchd3 | coiled-coil-h | Cytoplasm |
| ENSMUSG00000001761 | 3.81E-12 | 1.20E-11 | Smo | smoothened, | Plasma Mem |
| ENSMUSG00000018449 | 3.89E-12 | 1.22E-11 | Rpain | RPA interacti | Nucleus |
| ENSMUSG00000067995 | 3.90E-12 | 1.22E-11 | Gtf2f2 | general trans | Nucleus |
| ENSMUSG00000032316 | 3.94E-12 | 1.24E-11 | Clk3 | CDC like kina | Nucleus |
| ENSMUSG00000000876 | 4.03E-12 | 1.27E-11 | Pxmp4 | peroxisomal | Cytoplasm |
| ENSMUSG00000020863 | 4.03E-12 | 1.27E-11 | Luc7l3 | LUC7 like 3 p | Nucleus |
| ENSMUSG00000032436 | 4.05E-12 | 1.27E-11 | Cmtm7 | CKLF like MA | Extracellular S |
| ENSMUSG00000042535 | 4.14E-12 | 1.30E-11 | Gtpbp1 | GTP binding | Cytoplasm |
| ENSMUSG00000052131 | 4.17E-12 | 1.31E-11 | Akr1b7 | aldo-keto rec | Cytoplasm |
| ENSMUSG00000021650 | 4.18E-12 | 1.31E-11 | Ptcd2 | pentatricope | Cytoplasm |
| ENSMUSG00000029415 | 4.28E-12 | 1.34E-11 | Sdad1 | SDA1 domair | Nucleus |
| ENSMUSG00000061136 | 4.29E-12 | 1.34E-11 | Prpf40a | pre-mRNA p | Nucleus |
| ENSMUSG00000058503 | 4.32E-12 | 1.35E-11 | Fam133b | family with se | Other |
| ENSMUSG00000024213 | 4.41E-12 | 1.38E-11 | Nudt3 | nudix hydrol | Cytoplasm |
| ENSMUSG00000057649 | 4.44E-12 | 1.39E-11 | Brd9 | bromodomai | Other |
| ENSMUSG00000024137 | 4.49E-12 | 1.40E-11 | E4f1 | E4F transcrip | Nucleus |
| ENSMUSG00000079057 | 4.49E-12 | 1.40E-11 | Cyp4v3 | cytochrome I | Cytoplasm |
| ENSMUSG00000019989 | 4.55E-12 | 1.42E-11 | Enpp3 | ectonucleoti | Plasma Mem |
| ENSMUSG00000020076 | 4.62E-12 | 1.44E-11 | Ddx50 | DExD-box he | Nucleus |
| ENSMUSG00000035268 | 4.63E-12 | 1.45E-11 | Pkig | cAMP-deper | Other |
| ENSMUSG00000062510 | 4.70E-12 | 1.47E-11 | Nsl1 | NSL1, MIS12 | Nucleus |
| ENSMUSG00000006134 | 4.76E-12 | 1.48E-11 | Crkl | CRK like prot | Cytoplasm |
| ENSMUSG00000034160 | 4.76E-12 | 1.48E-11 | Ogt | O-linked N-a | Cytoplasm |
| ENSMUSG00000027932 | 4.95E-12 | 1.54E-11 | Slc27a3 | solute carrier | Cytoplasm |
| ENSMUSG00000039781 | 5.00E-12 | 1.56E-11 | Cep131 | centrosomal | Cytoplasm |
| ENSMUSG00000028587 | 5.02E-12 | 1.56E-11 | Orc1 | origin recogr | Nucleus |
| ENSMUSG00000031928 | 5.07E-12 | 1.58E-11 | Mre11a | MRE11 homc | Nucleus |
| ENSMUSG00000038909 | 5.07E-12 | 1.58E-11 | Kat7 | lysine acetylt | Nucleus |
| ENSMUSG00000074182 | 5.13E-12 | 1.60E-11 | Znhit6 | zinc finger H | Cytoplasm |
| ENSMUSG00000036083 | 5.17E-12 | 1.61E-11 | Slc17a3 | solute carrier | Plasma Mem |
| ENSMUSG00000092417 | 5.22E-12 | 1.62E-11 | Gpank1 | G-patch dom | Nucleus |

| | | | | | |
|---|---|---|---|---|---|
| ENSMUSG00000074802 | 5.25E-12 | 1.63E-11 | Gas2l3 | growth arrest | Other |
| ENSMUSG00000034532 | 5.26E-12 | 1.64E-11 | Fbxo16 | F-box protei | Other |
| ENSMUSG00000005125 | 5.27E-12 | 1.64E-11 | Ndrg1 | N-myc down | Nucleus |
| ENSMUSG00000025494 | 5.33E-12 | 1.65E-11 | Sigirr | single Ig and | Plasma Mem |
| ENSMUSG00000042606 | 5.41E-12 | 1.68E-11 | Hirip3 | HIRA interact | Nucleus |
| ENSMUSG00000025858 | 5.46E-12 | 1.69E-11 | Get4 | golgi to ER tr | Cytoplasm |
| ENSMUSG00000028431 | 5.53E-12 | 1.71E-11 | Ikbkap | elongator co | Cytoplasm |
| ENSMUSG00000027288 | 5.58E-12 | 1.73E-11 | Zfp106 | zinc finger pr | Cytoplasm |
| ENSMUSG00000025855 | 5.63E-12 | 1.74E-11 | Prkar1b | protein kinas | Cytoplasm |
| ENSMUSG00000042650 | 5.69E-12 | 1.76E-11 | Alkbh5 | alkB homolog | Nucleus |
| ENSMUSG00000006288 | 5.71E-12 | 1.77E-11 | Ttc5 | tetratricopep | Nucleus |
| ENSMUSG00000027080 | 6.14E-12 | 1.90E-11 | Med19 | mediator cor | Nucleus |
| ENSMUSG00000015961 | 6.22E-12 | 1.93E-11 | Adss | adenylosucci | Cytoplasm |
| ENSMUSG00000028416 | 6.31E-12 | 1.95E-11 | Bag1 | BCL2 associa | Cytoplasm |
| ENSMUSG00000087260 | 6.30E-12 | 1.95E-11 | Lamtor5 | late endoson | Cytoplasm |
| ENSMUSG00000037979 | 6.57E-12 | 2.03E-11 | Ccdc92 | coiled-coil d | Cytoplasm |
| ENSMUSG00000049882 | 6.97E-12 | 2.15E-11 | Vcpkmt | valosin conta | Cytoplasm |
| ENSMUSG00000027634 | 6.99E-12 | 2.16E-11 | Ndrg3 | NDRG family | Cytoplasm |
| ENSMUSG00000026411 | 7.17E-12 | 2.21E-11 | Tmem9 | transmembra | Cytoplasm |
| ENSMUSG00000029253 | 7.28E-12 | 2.25E-11 | Cenpc1 | centromere p | Nucleus |
| ENSMUSG00000030613 | 7.33E-12 | 2.26E-11 | Ccdc90b | coiled-coil d | Cytoplasm |
| ENSMUSG00000017264 | 7.33E-12 | 2.26E-11 | Exosc10 | exosome cor | Nucleus |
| ENSMUSG00000019179 | 7.44E-12 | 2.29E-11 | Mdh2 | malate dehyd | Cytoplasm |
| ENSMUSG00000068579 | 7.80E-12 | 2.40E-11 | Rpl7a-ps3 | ribosomal pr | Other |
| ENSMUSG00000046756 | 7.91E-12 | 2.43E-11 | Mrps7 | mitochondria | Cytoplasm |
| ENSMUSG00000013629 | 7.97E-12 | 2.45E-11 | Cad | carbamoyl-p | Cytoplasm |
| ENSMUSG00000032468 | 8.04E-12 | 2.47E-11 | Armc8 | armadillo rep | Nucleus |
| ENSMUSG00000042745 | 8.04E-12 | 2.47E-11 | Id1 | inhibitor of D | Nucleus |
| ENSMUSG00000003813 | 8.08E-12 | 2.48E-11 | Rad23a | RAD23 homo | Nucleus |
| ENSMUSG00000050966 | 8.10E-12 | 2.49E-11 | Lin28a | lin-28 homol | Cytoplasm |
| ENSMUSG00000028555 | 8.27E-12 | 2.54E-11 | Ttc39a | tetratricopep | Other |
| ENSMUSG00000063801 | 8.32E-12 | 2.55E-11 | Ap3s2 | adaptor relat | Cytoplasm |
| ENSMUSG00000090150 | 8.38E-12 | 2.57E-11 | Acad11 | acyl-CoA del | Cytoplasm |
| ENSMUSG00000029632 | 8.54E-12 | 2.62E-11 | Ndufa4 | NDUFA4, mit | Cytoplasm |
| ENSMUSG00000021629 | 8.63E-12 | 2.65E-11 | Slc30a5 | solute carrier | Plasma Mem |
| ENSMUSG00000032115 | 8.63E-12 | 2.65E-11 | Hyou1 | hypoxia up-r | Cytoplasm |
| ENSMUSG00000041846 | 8.71E-12 | 2.67E-11 | Smek1 | protein phos | Plasma Mem |
| ENSMUSG00000000916 | 8.78E-12 | 2.69E-11 | Nsun5 | NOP2/Sun R | Nucleus |
| ENSMUSG00000041815 | 8.84E-12 | 2.71E-11 | Poldip3 | DNA polyme | Nucleus |
| ENSMUSG00000028040 | 9.17E-12 | 2.81E-11 | Efna4 | ephrin A4 | Plasma Mem |
| ENSMUSG00000003070 | 9.18E-12 | 2.81E-11 | Efna2 | ephrin A2 | Plasma Mem |
| ENSMUSG00000064030 | 9.36E-12 | 2.86E-11 | Wibg | PYM homolo | Nucleus |
| ENSMUSG00000021597 | 9.41E-12 | 2.88E-11 | Ankrd32 | SMC5-SMC6 | Nucleus |
| ENSMUSG00000027201 | 9.50E-12 | 2.90E-11 | Myef2 | myelin expre | Nucleus |
| ENSMUSG00000025204 | 9.66E-12 | 2.95E-11 | Ndufb8 | NADH:ubiqu | Cytoplasm |
| ENSMUSG00000030499 | 1.02E-11 | 3.11E-11 | Kctd15 | potassium ch | Other |
| ENSMUSG00000010517 | 1.03E-11 | 3.15E-11 | Faf1 | Fas associate | Nucleus |
| ENSMUSG00000037991 | 1.03E-11 | 3.16E-11 | Rmi2 | RecQ mediat | Nucleus |
| ENSMUSG00000027651 | 1.04E-11 | 3.17E-11 | Rprd1b | regulation of | Nucleus |
| ENSMUSG00000026799 | 1.07E-11 | 3.25E-11 | Med27 | mediator cor | Nucleus |
| ENSMUSG00000001524 | 1.08E-11 | 3.29E-11 | Gtf2h4 | general trans | Nucleus |
| ENSMUSG00000030801 | 1.08E-11 | 3.30E-11 | Kat8 | lysine acetylt | Nucleus |
| ENSMUSG00000028134 | 1.10E-11 | 3.34E-11 | Ptbp2 | polypyrimidi | Nucleus |
| ENSMUSG00000025940 | 1.11E-11 | 3.37E-11 | Tmem70 | transmembra | Cytoplasm |
| ENSMUSG00000047728 | 1.13E-11 | 3.44E-11 | BC025446 | cDNA seque | Plasma Mem |
| ENSMUSG00000001924 | 1.16E-11 | 3.53E-11 | Uba1 | ubiquitin like | Cytoplasm |
| ENSMUSG00000031422 | 1.18E-11 | 3.59E-11 | Morf4l2 | mortality fact | Nucleus |
| ENSMUSG00000017615 | 1.19E-11 | 3.61E-11 | Tnfaip1 | TNF alpha in | Plasma Mem |

| | | | | | |
|---|---|---|---|---|---|
| ENSMUSG00000034258 | 1.22E-11 | 3.72E-11 | Mfsd7c | feline leukem | Plasma Mem |
| ENSMUSG00000031256 | 1.23E-11 | 3.73E-11 | Cstf2 | cleavage stim | Nucleus |
| ENSMUSG00000032077 | 1.25E-11 | 3.78E-11 | Bud13 | BUD13 homo | Nucleus |
| ENSMUSG00000022871 | 1.34E-11 | 4.08E-11 | Fetub | fetuin B | Extracellular S |
| ENSMUSG00000024975 | 1.36E-11 | 4.11E-11 | Pdcd4 | programmed | Nucleus |
| ENSMUSG00000024044 | 1.39E-11 | 4.21E-11 | Epb4.1l3 | erythrocyte n | Plasma Mem |
| ENSMUSG00000029920 | 1.40E-11 | 4.24E-11 | Smarcad1 | SWI/SNF-rela | Nucleus |
| ENSMUSG00000009292 | 1.43E-11 | 4.32E-11 | Trpm2 | transient rece | Plasma Mem |
| ENSMUSG00000023010 | 1.43E-11 | 4.32E-11 | Tmbim6 | transmembra | Nucleus |
| ENSMUSG00000018068 | 1.43E-11 | 4.34E-11 | Ints2 | integrator co | Nucleus |
| ENSMUSG00000036572 | 1.46E-11 | 4.43E-11 | Upf3b | UPF3B, regul | Nucleus |
| ENSMUSG00000031568 | 1.48E-11 | 4.48E-11 | Rwdd4a | RWD domain | Other |
| ENSMUSG00000000753 | 1.51E-11 | 4.55E-11 | Serpinf1 | serpin family | Extracellular S |
| ENSMUSG00000017561 | 1.51E-11 | 4.57E-11 | Crlf3 | cytokine rece | Cytoplasm |
| ENSMUSG00000013465 | 1.55E-11 | 4.68E-11 | Nelfb | negative elon | Nucleus |
| ENSMUSG00000044221 | 1.55E-11 | 4.69E-11 | Grsf1 | G-rich RNA s | Cytoplasm |
| ENSMUSG00000021760 | 1.56E-11 | 4.70E-11 | Gpx8 | glutathione p | Cytoplasm |
| ENSMUSG00000006005 | 1.57E-11 | 4.74E-11 | Tpr | translocated | Nucleus |
| ENSMUSG00000038059 | 1.58E-11 | 4.77E-11 | Smim3 | small integra | Other |
| ENSMUSG00000002981 | 1.61E-11 | 4.87E-11 | Clptm1 | CLPTM1, tran | Plasma Mem |
| ENSMUSG00000060152 | 1.61E-11 | 4.87E-11 | Pop5 | POP5 homolo | Nucleus |
| ENSMUSG00000023973 | 1.63E-11 | 4.92E-11 | Cnpy3 | canopy FGF s | Cytoplasm |
| ENSMUSG00000016503 | 1.64E-11 | 4.94E-11 | Gtf3a | general trans | Nucleus |
| ENSMUSG00000030062 | 1.66E-11 | 5.01E-11 | Rpn1 | ribophorin I | Cytoplasm |
| ENSMUSG00000012076 | 1.68E-11 | 5.06E-11 | Brms1l | BRMS1 like tr | Other |
| ENSMUSG00000027852 | 1.72E-11 | 5.17E-11 | Nras | NRAS proto- | Plasma Mem |
| ENSMUSG00000032265 | 1.77E-11 | 5.34E-11 | Fam46a | terminal nucl | Other |
| ENSMUSG00000036850 | 1.78E-11 | 5.35E-11 | Mrpl41 | mitochondria | Cytoplasm |
| ENSMUSG00000071337 | 1.80E-11 | 5.41E-11 | Tia1 | TIA1 cytotoxi | Nucleus |
| ENSMUSG00000020982 | 1.81E-11 | 5.43E-11 | Nemf | nuclear expo | Nucleus |
| ENSMUSG00000031438 | 1.84E-11 | 5.52E-11 | Rnf128 | ring finger pr | Cytoplasm |
| ENSMUSG00000020308 | 1.86E-11 | 5.60E-11 | Tpgs1 | tubulin polyg | Cytoplasm |
| ENSMUSG00000020515 | 1.87E-11 | 5.63E-11 | Cnot8 | CCR4-NOT tr | Nucleus |
| ENSMUSG00000003814 | 1.88E-11 | 5.63E-11 | Calr | calreticulin | Cytoplasm |
| ENSMUSG00000026942 | 1.89E-11 | 5.66E-11 | Traf2 | TNF receptor | Cytoplasm |
| ENSMUSG00000049904 | 1.90E-11 | 5.69E-11 | Tmem17 | transmembra | Extracellular S |
| ENSMUSG00000040464 | 1.92E-11 | 5.74E-11 | Gtpbp10 | GTP binding | Cytoplasm |
| ENSMUSG00000031820 | 1.92E-11 | 5.74E-11 | Babam1 | BRISC and BF | Nucleus |
| ENSMUSG00000032553 | 1.92E-11 | 5.74E-11 | Srprb | SRP receptor | Cytoplasm |
| ENSMUSG00000029703 | 1.93E-11 | 5.79E-11 | Lrwd1 | leucine rich r | Nucleus |
| ENSMUSG00000024941 | 1.94E-11 | 5.80E-11 | Scyl1 | SCY1 like pse | Cytoplasm |
| ENSMUSG00000031851 | 1.96E-11 | 5.86E-11 | Ntpcr | nucleoside-tr | Cytoplasm |
| ENSMUSG00000021748 | 1.96E-11 | 5.86E-11 | Pdhb | pyruvate deh | Cytoplasm |
| ENSMUSG00000027984 | 1.97E-11 | 5.89E-11 | Hadh | hydroxyacyl- | Cytoplasm |
| ENSMUSG00000053460 | 1.98E-11 | 5.92E-11 | Ggcx | gamma-gluta | Cytoplasm |
| ENSMUSG00000029641 | 2.00E-11 | 5.98E-11 | Rasl11a | RAS like fami | Nucleus |
| ENSMUSG00000051379 | 2.06E-11 | 6.16E-11 | Flrt3 | fibronectin le | Plasma Mem |
| ENSMUSG00000019969 | 2.07E-11 | 6.18E-11 | Psen1 | presenilin 1 | Plasma Mem |
| ENSMUSG00000036639 | 2.13E-11 | 6.36E-11 | Nudt1 | nudix hydrola | Extracellular S |
| ENSMUSG00000041346 | 2.18E-11 | 6.49E-11 | Wrap53 | WD repeat co | Nucleus |
| ENSMUSG00000021273 | 2.18E-11 | 6.51E-11 | Fdft1 | farnesyl-diph | Cytoplasm |
| ENSMUSG00000034758 | 2.20E-11 | 6.57E-11 | Tle6 | transducin lik | Nucleus |
| ENSMUSG00000030652 | 2.22E-11 | 6.60E-11 | Coq7 | coenzyme Q7 | Cytoplasm |
| ENSMUSG00000104445 | 2.22E-11 | 6.60E-11 | Rhbg | Rh family B g | Plasma Mem |
| ENSMUSG00000022964 | 2.22E-11 | 6.62E-11 | Tmem50b | transmembra | Plasma Mem |
| ENSMUSG00000029213 | 2.25E-11 | 6.69E-11 | Commd8 | COMM doma | Other |
| ENSMUSG00000027774 | 2.33E-11 | 6.91E-11 | Gfm1 | G elongation | Cytoplasm |
| ENSMUSG00000022471 | 2.33E-11 | 6.92E-11 | Xrcc6 | X-ray repair c | Nucleus |

| | | | | | |
|---|---|---|---|---|---|
| ENSMUSG00000024925 | 2.37E-11 | 7.04E-11 | Rnaseh2c | ribonuclease | Other |
| ENSMUSG00000037242 | 2.38E-11 | 7.08E-11 | Clic4 | chloride intra | Plasma Mem |
| ENSMUSG00000022370 | 2.43E-11 | 7.21E-11 | Mrpl13 | mitochondria | Cytoplasm |
| ENSMUSG00000022971 | 2.44E-11 | 7.26E-11 | Ifnar2 | interferon alp | Plasma Mem |
| ENSMUSG00000023980 | 2.45E-11 | 7.28E-11 | Taf8 | TATA-box bi | Nucleus |
| ENSMUSG00000021072 | 2.46E-11 | 7.30E-11 | Tmx1 | thioredoxin r | Cytoplasm |
| ENSMUSG00000050605 | 2.47E-11 | 7.32E-11 | Zfp61 | zinc finger pr | Nucleus |
| ENSMUSG00000037815 | 2.50E-11 | 7.40E-11 | Ctnna1 | catenin alpha | Plasma Mem |
| ENSMUSG00000036672 | 2.53E-11 | 7.49E-11 | Cenpt | centromere p | Cytoplasm |
| ENSMUSG00000019792 | 2.53E-11 | 7.49E-11 | Trmt11 | tRNA methyl | Other |
| ENSMUSG00000026643 | 2.56E-11 | 7.58E-11 | Nmt2 | N-myristoyltr | Cytoplasm |
| ENSMUSG00000028648 | 2.63E-11 | 7.78E-11 | Ndufs5 | NADH:ubiqu | Cytoplasm |
| ENSMUSG00000021731 | 2.64E-11 | 7.80E-11 | Mrps30 | mitochondria | Cytoplasm |
| ENSMUSG00000021665 | 2.73E-11 | 8.07E-11 | Hexb | hexosaminida | Cytoplasm |
| ENSMUSG00000011589 | 2.82E-11 | 8.34E-11 | Fsd1 | fibronectin ty | Cytoplasm |
| ENSMUSG00000019927 | 2.87E-11 | 8.48E-11 | Ube2d1 | ubiquitin con | Cytoplasm |
| ENSMUSG00000003549 | 2.88E-11 | 8.49E-11 | Ercc1 | ERCC excisio | Nucleus |
| ENSMUSG00000002319 | 2.88E-11 | 8.51E-11 | Ipo4 | importin 4 | Nucleus |
| ENSMUSG00000028955 | 2.89E-11 | 8.53E-11 | Vamp3 | vesicle associ | Plasma Mem |
| ENSMUSG00000042705 | 2.89E-11 | 8.53E-11 | Commd10 | COMM doma | Other |
| ENSMUSG00000022452 | 2.93E-11 | 8.64E-11 | Smdt1 | single-pass r | Extracellular S |
| ENSMUSG00000032902 | 2.98E-11 | 8.78E-11 | Slc16a1 | solute carrier | Plasma Mem |
| ENSMUSG00000041817 | 3.02E-11 | 8.89E-11 | Fam169a | family with se | Other |
| ENSMUSG00000038745 | 3.06E-11 | 9.01E-11 | Nlrp6 | NLR family py | Plasma Mem |
| ENSMUSG00000004455 | 3.14E-11 | 9.24E-11 | Ppp1cc | protein phos | Cytoplasm |
| ENSMUSG00000015966 | 3.18E-11 | 9.35E-11 | Il17rb | interleukin 17 | Plasma Mem |
| ENSMUSG00000019261 | 3.23E-11 | 9.50E-11 | Map1s | microtubule a | Cytoplasm |
| ENSMUSG00000026617 | 3.26E-11 | 9.58E-11 | Bpnt1 | 3'(2'), 5'-bisp | Nucleus |
| ENSMUSG00000024992 | 3.28E-11 | 9.66E-11 | Pde6c | phosphodies | Cytoplasm |
| ENSMUSG00000028029 | 3.30E-11 | 9.70E-11 | Aimp1 | aminoacyl tR | Extracellular S |
| ENSMUSG00000022420 | 3.33E-11 | 9.78E-11 | Dnal4 | dynein axone | Cytoplasm |
| ENSMUSG00000029012 | 3.36E-11 | 9.88E-11 | Orc5 | origin recogr | Nucleus |
| ENSMUSG00000020794 | 3.39E-11 | 9.97E-11 | Ube2g1 | ubiquitin con | Cytoplasm |
| ENSMUSG00000034022 | 3.40E-11 | 9.98E-11 | Cpsf1 | cleavage and | Nucleus |
| ENSMUSG00000090841 | 3.40E-11 | 9.98E-11 | Myl6 | myosin light | Cytoplasm |
| ENSMUSG00000026605 | 3.44E-11 | 1.01E-10 | Cenpf | centromere p | Nucleus |
| ENSMUSG00000096740 | 3.57E-11 | 1.05E-10 | Lbhd1 | LBH domain | Other |
| ENSMUSG00000022408 | 3.67E-11 | 1.08E-10 | Fam83f | family with se | Other |
| ENSMUSG00000078452 | 3.75E-11 | 1.10E-10 | Raet1d | retinoic acid | Plasma Mem |
| ENSMUSG00000068086 | 3.79E-11 | 1.11E-10 | Cyp2d9 | cytochrome P | Cytoplasm |
| ENSMUSG00000026083 | 3.92E-11 | 1.15E-10 | Eif5b | eukaryotic tra | Cytoplasm |
| ENSMUSG00000044005 | 3.96E-11 | 1.16E-10 | Gls2 | glutaminase | Cytoplasm |
| ENSMUSG00000028960 | 3.97E-11 | 1.16E-10 | Ube4b | ubiquitinatio | Cytoplasm |
| ENSMUSG00000032407 | 4.00E-11 | 1.17E-10 | U2surp | U2 snRNP as | Nucleus |
| ENSMUSG00000034544 | 4.03E-11 | 1.18E-10 | Rsrc1 | arginine and | Nucleus |
| ENSMUSG00000032745 | 4.06E-11 | 1.19E-10 | Gpbp1 | GC-rich pron | Nucleus |
| ENSMUSG00000067722 | 4.06E-11 | 1.19E-10 | BC003965 | chromosome | Other |
| ENSMUSG00000000776 | 4.13E-11 | 1.21E-10 | Polr3d | RNA polymer | Nucleus |
| ENSMUSG00000029131 | 4.19E-11 | 1.22E-10 | Dnajb6 | DnaJ heat sh | Nucleus |
| ENSMUSG00000029227 | 4.20E-11 | 1.23E-10 | Fip1l1 | FIP1 like 1 (S. | Nucleus |
| ENSMUSG00000022114 | 4.21E-11 | 1.23E-10 | Spry2 | sprouty RTK s | Plasma Mem |
| ENSMUSG00000015087 | 4.26E-11 | 1.24E-10 | Rabl6 | RAB, membe | Cytoplasm |
| ENSMUSG00000029433 | 4.34E-11 | 1.26E-10 | Diablo | diablo IAP-b | Cytoplasm |
| ENSMUSG00000027810 | 4.38E-11 | 1.28E-10 | Eif2a | eukaryotic tra | Cytoplasm |
| ENSMUSG00000018476 | 4.48E-11 | 1.30E-10 | Kdm6b | lysine demetl | Extracellular S |
| ENSMUSG00000018415 | 4.51E-11 | 1.31E-10 | Gid4 | GID complex | Nucleus |
| ENSMUSG00000001555 | 4.52E-11 | 1.32E-10 | Fkbp10 | FK506 bindin | Cytoplasm |
| ENSMUSG00000046010 | 4.60E-11 | 1.34E-10 | Zfp830 | zinc finger pr | Nucleus |

| | | | | | |
|---|---|---|---|---|---|
| ENSMUSG00000020184 | 4.61E-11 | 1.34E-10 | Mdm2 | MDM2 proto | Nucleus |
| ENSMUSG00000054364 | 4.63E-11 | 1.35E-10 | Rhob | ras homolog | Cytoplasm |
| ENSMUSG00000031015 | 4.63E-11 | 1.35E-10 | Swap70 | switching B c | Cytoplasm |
| ENSMUSG00000028114 | 4.70E-11 | 1.37E-10 | Mettl14 | methyltransfe | Nucleus |
| ENSMUSG00000031601 | 4.72E-11 | 1.37E-10 | Cnot7 | CCR4-NOT t | Nucleus |
| ENSMUSG00000021621 | 4.82E-11 | 1.40E-10 | Zcchc9 | zinc finger C | Other |
| ENSMUSG00000041035 | 4.88E-11 | 1.42E-10 | Dpcd | deleted in pr | Nucleus |
| ENSMUSG00000027361 | 4.90E-11 | 1.42E-10 | Gabpb1 | GA binding p | Nucleus |
| ENSMUSG00000039826 | 4.98E-11 | 1.45E-10 | Trub2 | TruB pseudo | Cytoplasm |
| ENSMUSG00000022787 | 5.06E-11 | 1.47E-10 | Wdr53 | WD repeat d | Other |
| ENSMUSG00000028690 | 5.08E-11 | 1.47E-10 | Mmachc | methylmalon | Cytoplasm |
| ENSMUSG00000052337 | 5.14E-11 | 1.49E-10 | Immt | inner membr | Cytoplasm |
| ENSMUSG00000005936 | 5.19E-11 | 1.50E-10 | Kctd20 | potassium ch | Cytoplasm |
| ENSMUSG00000029625 | 5.23E-11 | 1.51E-10 | Cpsf4 | cleavage and | Nucleus |
| ENSMUSG00000035704 | 5.26E-11 | 1.52E-10 | Alg8 | ALG8, alpha- | Cytoplasm |
| ENSMUSG00000024400 | 5.30E-11 | 1.53E-10 | Wdr33 | WD repeat d | Nucleus |
| ENSMUSG00000030067 | 5.32E-11 | 1.54E-10 | Foxp1 | forkhead box | Nucleus |
| ENSMUSG00000020152 | 5.33E-11 | 1.54E-10 | Actr2 | ARP2 actin re | Plasma Mem |
| ENSMUSG00000025154 | 5.36E-11 | 1.55E-10 | Arhgap19 | Rho GTPase a | Cytoplasm |
| ENSMUSG00000017210 | 5.42E-11 | 1.57E-10 | Med24 | mediator cor | Nucleus |
| ENSMUSG00000057037 | 5.44E-11 | 1.57E-10 | Cfhr1 | complement | Extracellular S |
| ENSMUSG00000031065 | 5.48E-11 | 1.58E-10 | Cdk16 | cyclin depen | Cytoplasm |
| ENSMUSG00000032046 | 5.51E-11 | 1.59E-10 | Abhd12 | abhydrolase | Plasma Mem |
| ENSMUSG00000027599 | 5.58E-11 | 1.61E-10 | Armc1 | armadillo rep | Cytoplasm |
| ENSMUSG00000020108 | 5.62E-11 | 1.62E-10 | Ddit4 | DNA damage | Cytoplasm |
| ENSMUSG00000031835 | 5.71E-11 | 1.65E-10 | Mbtps1 | membrane b | Cytoplasm |
| ENSMUSG00000030772 | 5.74E-11 | 1.65E-10 | Dkk3 | dickkopf WN | Extracellular S |
| ENSMUSG00000022837 | 5.86E-11 | 1.69E-10 | Iqcb1 | IQ motif cont | Extracellular S |
| ENSMUSG00000020803 | 5.88E-11 | 1.69E-10 | Txndc17 | thioredoxin d | Cytoplasm |
| ENSMUSG00000036202 | 5.92E-11 | 1.70E-10 | Rif1 | replication tir | Nucleus |
| ENSMUSG00000031245 | 5.95E-11 | 1.71E-10 | Hmgn5 | high-mobility | Nucleus |
| ENSMUSG00000037536 | 6.01E-11 | 1.73E-10 | Fbxo34 | F-box protei | Other |
| ENSMUSG00000000827 | 6.13E-11 | 1.76E-10 | Tpd52l2 | tumor protei | Cytoplasm |
| ENSMUSG00000019878 | 6.16E-11 | 1.77E-10 | Hsf2 | heat shock tr | Nucleus |
| ENSMUSG00000036180 | 6.20E-11 | 1.78E-10 | Gatad2a | GATA zinc fir | Nucleus |
| ENSMUSG00000022551 | 6.22E-11 | 1.79E-10 | Cyc1 | cytochrome c | Cytoplasm |
| ENSMUSG00000024644 | 6.30E-11 | 1.81E-10 | Cndp2 | carnosine dip | Cytoplasm |
| ENSMUSG00000031924 | 6.30E-11 | 1.81E-10 | Cyb5b | cytochrome b | Cytoplasm |
| ENSMUSG00000032803 | 6.32E-11 | 1.81E-10 | Cdv3 | CDV3 homol | Cytoplasm |
| ENSMUSG00000042694 | 6.34E-11 | 1.82E-10 | Obfc1 | STN1, CST cc | Nucleus |
| ENSMUSG00000046070 | 6.35E-11 | 1.82E-10 | Igfals | insulin like gr | Extracellular S |
| ENSMUSG00000022255 | 6.42E-11 | 1.84E-10 | Mtdh | metadherin | Cytoplasm |
| ENSMUSG00000051235 | 6.52E-11 | 1.87E-10 | Gen1 | GEN1, Hollid | Cytoplasm |
| ENSMUSG00000032299 | 6.59E-11 | 1.89E-10 | Commd4 | COMM doma | Cytoplasm |
| ENSMUSG00000029504 | 6.65E-11 | 1.90E-10 | Ddx51 | DEAD-box h | Other |
| ENSMUSG00000022359 | 6.75E-11 | 1.93E-10 | Wdyhv1 | WDYHV mot | Cytoplasm |
| ENSMUSG00000052593 | 6.77E-11 | 1.94E-10 | Adam17 | ADAM metal | Plasma Mem |
| ENSMUSG00000026641 | 6.95E-11 | 1.99E-10 | Usf1 | upstream tra | Nucleus |
| ENSMUSG00000061882 | 7.08E-11 | 2.02E-10 | Ccdc62 | coiled-coil d | Nucleus |
| ENSMUSG00000027387 | 7.11E-11 | 2.03E-10 | Zc3h8 | zinc finger C | Nucleus |
| ENSMUSG00000009470 | 7.14E-11 | 2.04E-10 | Tnpo1 | transportin 1 | Nucleus |
| ENSMUSG00000043131 | 7.17E-11 | 2.05E-10 | Mob1a | MOB kinase a | Plasma Mem |
| ENSMUSG00000024899 | 7.17E-11 | 2.05E-10 | Papss2 | 3'-phosphoa | Cytoplasm |
| ENSMUSG00000037300 | 7.37E-11 | 2.10E-10 | Ttc13 | tetratricopep | Other |
| ENSMUSG00000056999 | 7.40E-11 | 2.11E-10 | Ide | insulin degra | Extracellular S |
| ENSMUSG00000061559 | 7.40E-11 | 2.11E-10 | Wdr61 | WD repeat d | Nucleus |
| ENSMUSG00000020477 | 7.46E-11 | 2.13E-10 | Mrps24 | mitochondria | Cytoplasm |
| ENSMUSG00000029916 | 7.48E-11 | 2.13E-10 | Agk | acylglycerol k | Cytoplasm |

| | | | | | |
|---|---|---|---|---|---|
| ENSMUSG00000029670 | 7.48E-11 | 2.13E-10 | Ing3 | inhibitor of g | Nucleus |
| ENSMUSG00000003527 | 7.53E-11 | 2.14E-10 | Dgcr14 | ess-2 splicing | Nucleus |
| ENSMUSG00000070934 | 7.59E-11 | 2.16E-10 | Rraga | Ras related G | Cytoplasm |
| ENSMUSG00000022369 | 7.67E-11 | 2.18E-10 | Mtbp | MDM2 bindi | Cytoplasm |
| ENSMUSG00000039616 | 7.73E-11 | 2.20E-10 | Mocos | molybdenum | Cytoplasm |
| ENSMUSG00000030816 | 7.83E-11 | 2.23E-10 | Rnf40 | ring finger pr | Cytoplasm |
| ENSMUSG00000006058 | 7.85E-11 | 2.23E-10 | Snf8 | SNF8, ESCRT | Cytoplasm |
| ENSMUSG00000019158 | 7.89E-11 | 2.24E-10 | Tmem160 | transmembra | Cytoplasm |
| ENSMUSG00000026240 | 7.94E-11 | 2.25E-10 | Cops7b | COP9 signalo | Cytoplasm |
| ENSMUSG00000058267 | 8.08E-11 | 2.30E-10 | Mrps14 | mitochondria | Cytoplasm |
| ENSMUSG00000025817 | 8.45E-11 | 2.40E-10 | Nudt5 | nudix hydrola | Cytoplasm |
| ENSMUSG00000031960 | 8.49E-11 | 2.41E-10 | Aars | alanyl-tRNA | Cytoplasm |
| ENSMUSG00000024603 | 8.70E-11 | 2.47E-10 | Dctn4 | dynactin sub | Nucleus |
| ENSMUSG00000002308 | 8.97E-11 | 2.54E-10 | Cd320 | CD320 molec | Plasma Mem |
| ENSMUSG00000000384 | 9.07E-11 | 2.57E-10 | Tbrg4 | transforming | Cytoplasm |
| ENSMUSG00000062054 | 9.11E-11 | 2.58E-10 | Iah1 | isoamyl aceta | Cytoplasm |
| ENSMUSG00000035378 | 9.14E-11 | 2.59E-10 | Shq1 | SHQ1, H/ACA | Nucleus |
| ENSMUSG00000033186 | 9.18E-11 | 2.60E-10 | Mzt1 | mitotic spind | Cytoplasm |
| ENSMUSG00000013736 | 9.24E-11 | 2.62E-10 | Trnt1 | tRNA nucleot | Cytoplasm |
| ENSMUSG00000078974 | 9.50E-11 | 2.69E-10 | Sec61g | Sec61 translo | Plasma Mem |
| ENSMUSG00000020413 | 9.52E-11 | 2.69E-10 | Hus1 | HUS1 checkp | Nucleus |
| ENSMUSG00000043872 | 9.54E-11 | 2.70E-10 | Zmym1 | zinc finger M | Other |
| ENSMUSG00000058569 | 9.57E-11 | 2.71E-10 | Tmed9 | transmembra | Cytoplasm |
| ENSMUSG00000029149 | 9.62E-11 | 2.72E-10 | Krtcap3 | keratinocyte | Other |
| ENSMUSG00000021843 | 9.68E-11 | 2.73E-10 | Ktn1 | kinectin 1 | Cytoplasm |
| ENSMUSG00000053931 | 9.71E-11 | 2.74E-10 | Cnn3 | calponin 3 | Cytoplasm |
| ENSMUSG00000032306 | 9.83E-11 | 2.78E-10 | Mpi | mannose pho | Cytoplasm |
| ENSMUSG00000005447 | 9.84E-11 | 2.78E-10 | Pafah1b3 | platelet activ | Cytoplasm |
| ENSMUSG00000001506 | 9.85E-11 | 2.78E-10 | Col1a1 | collagen type | Extracellular S |
| ENSMUSG00000024446 | 9.85E-11 | 2.78E-10 | Rpp21 | ribonuclease | Nucleus |
| ENSMUSG00000060679 | 9.87E-11 | 2.78E-10 | Mrps9 | mitochondria | Cytoplasm |
| ENSMUSG00000030641 | 9.91E-11 | 2.79E-10 | Ddias | DNA damage | Cytoplasm |
| ENSMUSG00000037720 | 1.00E-10 | 2.83E-10 | Tmem33 | transmembra | Cytoplasm |
| ENSMUSG00000001415 | 1.01E-10 | 2.84E-10 | Smg5 | SMG5, nonse | Nucleus |
| ENSMUSG00000038383 | 1.02E-10 | 2.86E-10 | Pigu | phosphatidyl | Cytoplasm |
| ENSMUSG00000025017 | 1.04E-10 | 2.93E-10 | Pik3ap1 | phosphoinos | Cytoplasm |
| ENSMUSG00000035632 | 1.06E-10 | 2.98E-10 | Cnot3 | CCR4-NOT tr | Cytoplasm |
| ENSMUSG00000042688 | 1.08E-10 | 3.05E-10 | Mapk6 | mitogen-acti | Cytoplasm |
| ENSMUSG00000020781 | 1.09E-10 | 3.08E-10 | Tsen54 | tRNA splicing | Nucleus |
| ENSMUSG00000043384 | 1.11E-10 | 3.12E-10 | Gprasp1 | G protein-co | Cytoplasm |
| ENSMUSG00000036817 | 1.11E-10 | 3.12E-10 | Sun1 | Sad1 and UN | Nucleus |
| ENSMUSG00000036450 | 1.13E-10 | 3.18E-10 | Hif1an | hypoxia indu | Nucleus |
| ENSMUSG00000022961 | 1.14E-10 | 3.19E-10 | Son | SON DNA bin | Nucleus |
| ENSMUSG00000001120 | 1.14E-10 | 3.20E-10 | Pcbp3 | poly(rC) bind | Nucleus |
| ENSMUSG00000061111 | 1.14E-10 | 3.21E-10 | Fam195b | MAPK regula | Nucleus |
| ENSMUSG00000002059 | 1.15E-10 | 3.22E-10 | Rab34 | RAB34, mem | Cytoplasm |
| ENSMUSG00000041712 | 1.16E-10 | 3.24E-10 | Ubr7 | ubiquitin pro | Other |
| ENSMUSG00000040018 | 1.18E-10 | 3.31E-10 | Cox15 | cytochrome c | Cytoplasm |
| ENSMUSG00000020577 | 1.20E-10 | 3.36E-10 | Tspan13 | tetraspanin 1 | Plasma Mem |
| ENSMUSG00000022564 | 1.22E-10 | 3.41E-10 | Grina | glutamate ion | Other |
| ENSMUSG00000003660 | 1.24E-10 | 3.47E-10 | Snrnp200 | small nuclear | Nucleus |
| ENSMUSG00000040415 | 1.24E-10 | 3.47E-10 | Dtx3 | deltex E3 ubi | Cytoplasm |
| ENSMUSG00000056167 | 1.24E-10 | 3.48E-10 | Cnot10 | CCR4-NOT tr | Cytoplasm |
| ENSMUSG00000020691 | 1.27E-10 | 3.55E-10 | Mettl2 | methyltransfe | Other |
| ENSMUSG00000002996 | 1.27E-10 | 3.56E-10 | Hbp1 | HMG-box tra | Nucleus |
| ENSMUSG00000028540 | 1.28E-10 | 3.58E-10 | Dph2 | DPH2 homol | Cytoplasm |
| ENSMUSG00000038717 | 1.30E-10 | 3.63E-10 | Atp5l | ATP synthase | Cytoplasm |
| ENSMUSG00000027079 | 1.31E-10 | 3.65E-10 | Clp1 | cleavage and | Nucleus |

| | | | | | |
|---|---|---|---|---|---|
| ENSMUSG00000021868 | 1.31E-10 | 3.66E-10 | Ppif | peptidylproly | Cytoplasm |
| ENSMUSG00000072582 | 1.32E-10 | 3.67E-10 | Ptrh2 | peptidyl-tRN | Cytoplasm |
| ENSMUSG00000078853 | 1.32E-10 | 3.67E-10 | Igtp | interferon ga | Cytoplasm |
| ENSMUSG00000092416 | 1.33E-10 | 3.72E-10 | Zfp141 | zinc finger pr | Other |
| ENSMUSG00000022394 | 1.36E-10 | 3.79E-10 | L3mbtl2 | L3MBTL2, po | Nucleus |
| ENSMUSG00000024357 | 1.37E-10 | 3.81E-10 | Sil1 | SIL1 nucleoti | Cytoplasm |
| ENSMUSG00000040963 | 1.38E-10 | 3.84E-10 | Asgr2 | asialoglycopr | Plasma Mem |
| ENSMUSG00000020962 | 1.38E-10 | 3.84E-10 | Gtf2a1 | general trans | Cytoplasm |
| ENSMUSG00000008140 | 1.38E-10 | 3.84E-10 | Emc10 | ER membran | Cytoplasm |
| ENSMUSG00000021025 | 1.40E-10 | 3.89E-10 | Nfkbia | NFKB inhibito | Cytoplasm |
| ENSMUSG00000041303 | 1.43E-10 | 3.98E-10 | Gtf3c3 | general trans | Nucleus |
| ENSMUSG00000041168 | 1.44E-10 | 3.99E-10 | Lonp1 | lon peptidase | Cytoplasm |
| ENSMUSG00000036114 | 1.44E-10 | 4.01E-10 | Rpp25l | ribonuclease | Other |
| ENSMUSG00000022040 | 1.44E-10 | 4.01E-10 | Ephx2 | epoxide hydr | Cytoplasm |
| ENSMUSG00000009073 | 1.46E-10 | 4.04E-10 | Nf2 | neurofibromi | Plasma Mem |
| ENSMUSG00000039715 | 1.47E-10 | 4.07E-10 | Wdr34 | WD repeat d | Cytoplasm |
| ENSMUSG00000028833 | 1.47E-10 | 4.08E-10 | Ncdn | neurochondr | Cytoplasm |
| ENSMUSG00000024811 | 1.48E-10 | 4.11E-10 | Tnks2 | tankyrase 2 | Nucleus |
| ENSMUSG00000039178 | 1.49E-10 | 4.12E-10 | Tbc1d19 | TBC1 domair | Other |
| ENSMUSG00000025157 | 1.52E-10 | 4.21E-10 | Zdhhc16 | zinc finger DI | Cytoplasm |
| ENSMUSG00000063281 | 1.52E-10 | 4.22E-10 | Zfp35 | zinc finger pr | Nucleus |
| ENSMUSG00000005506 | 1.56E-10 | 4.33E-10 | Celf1 | CUGBP, Elav- | Nucleus |
| ENSMUSG00000029538 | 1.57E-10 | 4.34E-10 | Srsf9 | serine and ar | Nucleus |
| ENSMUSG00000013787 | 1.59E-10 | 4.40E-10 | Ehmt2 | euchromatic | Nucleus |
| ENSMUSG00000053483 | 1.61E-10 | 4.45E-10 | Usp21 | ubiquitin spe | Cytoplasm |
| ENSMUSG00000011960 | 1.62E-10 | 4.49E-10 | Ccnt1 | cyclin T1 | Nucleus |
| ENSMUSG00000020525 | 1.64E-10 | 4.55E-10 | Ppm1d | protein phos | Cytoplasm |
| ENSMUSG00000063895 | 1.65E-10 | 4.57E-10 | Nupl1 | nucleoporin ! | Nucleus |
| ENSMUSG00000050103 | 1.67E-10 | 4.62E-10 | Agmo | alkylglycerol | Cytoplasm |
| ENSMUSG00000021905 | 1.73E-10 | 4.77E-10 | Dph3 | diphthamide | Cytoplasm |
| ENSMUSG00000027306 | 1.73E-10 | 4.77E-10 | Nusap1 | nucleolar anc | Nucleus |
| ENSMUSG00000026603 | 1.73E-10 | 4.78E-10 | Smyd2 | SET and MYN | Cytoplasm |
| ENSMUSG00000031596 | 1.75E-10 | 4.83E-10 | Slc7a2 | solute carrier | Plasma Mem |
| ENSMUSG00000022718 | 1.76E-10 | 4.85E-10 | Dgcr8 | DGCR8, micr | Nucleus |
| ENSMUSG00000029474 | 1.76E-10 | 4.85E-10 | Rnf34 | ring finger pr | Cytoplasm |
| ENSMUSG00000045983 | 1.77E-10 | 4.87E-10 | Eif4g1 | eukaryotic tra | Cytoplasm |
| ENSMUSG00000014551 | 1.77E-10 | 4.87E-10 | Mrps25 | mitochondria | Cytoplasm |
| ENSMUSG00000032463 | 1.78E-10 | 4.91E-10 | Faim | Fas apoptotic | Plasma Mem |
| ENSMUSG00000023852 | 1.80E-10 | 4.96E-10 | Chd1 | chromodoma | Nucleus |
| ENSMUSG00000025889 | 1.81E-10 | 4.99E-10 | Snca | synuclein alp | Cytoplasm |
| ENSMUSG00000014873 | 1.82E-10 | 5.01E-10 | Surf2 | surfeit 2 | Nucleus |
| ENSMUSG00000047045 | 1.82E-10 | 5.01E-10 | Tmem164 | transmembra | Other |
| ENSMUSG00000003573 | 1.84E-10 | 5.06E-10 | Homer3 | homer scaffo | Plasma Mem |
| ENSMUSG00000071379 | 1.84E-10 | 5.06E-10 | Hpcal1 | hippocalcin li | Cytoplasm |
| ENSMUSG00000096199 | 1.85E-10 | 5.08E-10 | Ptrhd1 | peptidyl-tRN | Cytoplasm |
| ENSMUSG00000022877 | 1.85E-10 | 5.10E-10 | Hrg | histidine-rich | Cytoplasm |
| ENSMUSG00000027236 | 1.87E-10 | 5.15E-10 | Eif3j1 | eukaryotic tra | Cytoplasm |
| ENSMUSG00000016541 | 1.89E-10 | 5.21E-10 | Atxn10 | ataxin 10 | Cytoplasm |
| ENSMUSG00000021681 | 1.92E-10 | 5.27E-10 | Aggf1 | angiogenic fa | Cytoplasm |
| ENSMUSG00000003316 | 1.94E-10 | 5.32E-10 | Glg1 | golgi glycopr | Cytoplasm |
| ENSMUSG00000009640 | 1.95E-10 | 5.35E-10 | Dmap1 | DNA methylt | Nucleus |
| ENSMUSG00000022476 | 2.00E-10 | 5.48E-10 | Polr3h | RNA polymer | Nucleus |
| ENSMUSG00000043962 | 2.04E-10 | 5.59E-10 | Thrap3 | thyroid horm | Nucleus |
| ENSMUSG00000039615 | 2.04E-10 | 5.60E-10 | Stub1 | STIP1 homolo | Cytoplasm |
| ENSMUSG00000037405 | 2.05E-10 | 5.61E-10 | Icam1 | intercellular a | Plasma Mem |
| ENSMUSG00000040044 | 2.05E-10 | 5.62E-10 | Orc3 | origin recogr | Nucleus |
| ENSMUSG00000017843 | 2.06E-10 | 5.64E-10 | Ppp2r5c | protein phos | Nucleus |
| ENSMUSG00000019897 | 2.10E-10 | 5.74E-10 | Ccdc59 | coiled-coil d | Nucleus |

| | | | | | |
|---|---|---|---|---|---|
| ENSMUSG00000031982 | 2.12E-10 | 5.80E-10 | Arv1 | ARV1 homolo | Cytoplasm |
| ENSMUSG00000023048 | 2.13E-10 | 5.82E-10 | Prr13 | proline rich 1 | Other |
| ENSMUSG00000063856 | 2.17E-10 | 5.94E-10 | Gpx1 | glutathione p | Cytoplasm |
| ENSMUSG00000022800 | 2.19E-10 | 5.99E-10 | Fyttd1 | forty-two-th | Nucleus |
| ENSMUSG00000072258 | 2.20E-10 | 6.01E-10 | Taf1a | TATA-box bi | Nucleus |
| ENSMUSG00000030752 | 2.22E-10 | 6.07E-10 | Kdm8 | lysine demetl | Nucleus |
| ENSMUSG00000036599 | 2.24E-10 | 6.11E-10 | Chst12 | carbohydrate | Cytoplasm |
| ENSMUSG00000035885 | 2.26E-10 | 6.17E-10 | Cox8a | cytochrome c | Cytoplasm |
| ENSMUSG00000022789 | 2.32E-10 | 6.35E-10 | Dnm1l | dynamin 1 lik | Cytoplasm |
| ENSMUSG00000030602 | 2.33E-10 | 6.37E-10 | Pak4 | p21 (RAC1) a | Cytoplasm |
| ENSMUSG00000002055 | 2.37E-10 | 6.46E-10 | Spag5 | sperm associ | Nucleus |
| ENSMUSG00000031715 | 2.38E-10 | 6.48E-10 | Smarca5 | SWI/SNF rela | Nucleus |
| ENSMUSG00000019066 | 2.39E-10 | 6.51E-10 | Rab3d | RAB3D, mem | Cytoplasm |
| ENSMUSG00000026127 | 2.39E-10 | 6.51E-10 | Imp4 | IMP4, U3 sma | Nucleus |
| ENSMUSG00000027552 | 2.53E-10 | 6.90E-10 | E2f5 | E2F transcrip | Nucleus |
| ENSMUSG00000025193 | 2.60E-10 | 7.09E-10 | Cutc | cutC copper | Cytoplasm |
| ENSMUSG00000060261 | 2.62E-10 | 7.13E-10 | Gtf2i | general trans | Nucleus |
| ENSMUSG00000041718 | 2.63E-10 | 7.16E-10 | Alg13 | asparagine-li | Other |
| ENSMUSG00000066568 | 2.67E-10 | 7.26E-10 | Lsm14a | LSM14A, mR | Cytoplasm |
| ENSMUSG00000027176 | 2.74E-10 | 7.46E-10 | Cstf3 | cleavage stim | Nucleus |
| ENSMUSG00000015461 | 2.77E-10 | 7.53E-10 | Atf6b | activating tra | Nucleus |
| ENSMUSG00000028670 | 2.78E-10 | 7.56E-10 | Lypla2 | lysophospho | Cytoplasm |
| ENSMUSG00000063882 | 2.78E-10 | 7.56E-10 | Uqcrh | ubiquinol-cy | Cytoplasm |
| ENSMUSG00000022528 | 2.82E-10 | 7.67E-10 | Hes1 | hes family bH | Nucleus |
| ENSMUSG00000037958 | 2.84E-10 | 7.70E-10 | Ccdc55 | nuclear speck | Nucleus |
| ENSMUSG00000019433 | 2.89E-10 | 7.85E-10 | Gipc1 | GIPC PDZ do | Cytoplasm |
| ENSMUSG00000030725 | 2.90E-10 | 7.86E-10 | Lipt2 | lipoyl(octano | Cytoplasm |
| ENSMUSG00000003531 | 2.92E-10 | 7.93E-10 | Dgcr6 | DiGeorge syr | Other |
| ENSMUSG00000025862 | 2.93E-10 | 7.95E-10 | Stag2 | stromal antig | Nucleus |
| ENSMUSG00000032519 | 2.95E-10 | 8.01E-10 | Slc25a38 | solute carrier | Cytoplasm |
| ENSMUSG00000049404 | 2.98E-10 | 8.08E-10 | Rarres1 | retinoic acid | Plasma Mem |
| ENSMUSG00000035781 | 3.02E-10 | 8.17E-10 | R3hdm4 | R3H domain | Other |
| ENSMUSG00000031479 | 3.04E-10 | 8.22E-10 | Vps36 | vacuolar prot | Cytoplasm |
| ENSMUSG00000021938 | 3.09E-10 | 8.36E-10 | Pspc1 | paraspeckle c | Nucleus |
| ENSMUSG00000005699 | 3.10E-10 | 8.40E-10 | Pard6a | par-6 family | Plasma Mem |
| ENSMUSG00000068479 | 3.15E-10 | 8.51E-10 | Mfap1a | microfibril as | Extracellular S |
| ENSMUSG00000035834 | 3.18E-10 | 8.59E-10 | Polr3g | RNA polymer | Nucleus |
| ENSMUSG00000042787 | 3.22E-10 | 8.69E-10 | Exog | exo/endonuc | Cytoplasm |
| ENSMUSG00000001864 | 3.23E-10 | 8.74E-10 | Aif1l | allograft infla | Plasma Mem |
| ENSMUSG00000020482 | 3.25E-10 | 8.78E-10 | Ccdc117 | coiled-coil d | Other |
| ENSMUSG00000020397 | 3.27E-10 | 8.84E-10 | Med7 | mediator cor | Nucleus |
| ENSMUSG00000029599 | 3.34E-10 | 9.01E-10 | Ddx54 | DEAD-box he | Nucleus |
| ENSMUSG00000034820 | 3.38E-10 | 9.12E-10 | Cpsf7 | cleavage and | Nucleus |
| ENSMUSG00000031352 | 3.40E-10 | 9.18E-10 | Hccs | holocytochro | Cytoplasm |
| ENSMUSG00000035890 | 3.46E-10 | 9.34E-10 | Rnf126 | ring finger pr | Other |
| ENSMUSG00000062083 | 3.48E-10 | 9.38E-10 | Rpl13a-ps1 | ribosomal pr | Other |
| ENSMUSG00000039218 | 3.49E-10 | 9.40E-10 | Srrm2 | serine/argini | Nucleus |
| ENSMUSG00000020255 | 3.52E-10 | 9.48E-10 | D10Wsu102e | chromosome | Other |
| ENSMUSG00000049553 | 3.54E-10 | 9.54E-10 | Polr1a | RNA polymer | Nucleus |
| ENSMUSG00000023156 | 3.56E-10 | 9.59E-10 | Rpp14 | ribonuclease | Nucleus |
| ENSMUSG00000007613 | 3.58E-10 | 9.63E-10 | Tgfbr1 | transforming | Plasma Mem |
| ENSMUSG00000023387 | 3.58E-10 | 9.63E-10 | Kcnk16 | potassium tw | Plasma Mem |
| ENSMUSG00000029594 | 3.72E-10 | 1.00E-09 | Rbm19 | RNA binding | Nucleus |
| ENSMUSG00000035024 | 3.83E-10 | 1.03E-09 | Ncapd3 | non-SMC co | Nucleus |
| ENSMUSG00000068739 | 4.05E-10 | 1.09E-09 | Sars | seryl-tRNA sy | Cytoplasm |
| ENSMUSG00000032777 | 4.05E-10 | 1.09E-09 | Gtf3c1 | general trans | Nucleus |
| ENSMUSG00000030681 | 4.14E-10 | 1.11E-09 | Mvp | major vault p | Nucleus |
| ENSMUSG00000035062 | 4.20E-10 | 1.13E-09 | Zc4h2 | zinc finger C4 | Nucleus |

| | | | | | |
|---|---|---|---|---|---|
| ENSMUSG00000096472 | 4.23E-10 | 1.14E-09 | Cdkn2d | cyclin depen( | Cytoplasm |
| ENSMUSG00000046201 | 4.30E-10 | 1.15E-09 | Scaf8 | SR-related C | Nucleus |
| ENSMUSG00000016526 | 4.32E-10 | 1.16E-09 | Dyrk3 | dual specifici | Nucleus |
| ENSMUSG00000015937 | 4.39E-10 | 1.18E-09 | H2afy | H2A histone | Nucleus |
| ENSMUSG00000005718 | 4.40E-10 | 1.18E-09 | Tfap4 | transcription | Nucleus |
| ENSMUSG00000027547 | 4.45E-10 | 1.19E-09 | Sall4 | spalt like trar | Nucleus |
| ENSMUSG00000021287 | 4.54E-10 | 1.22E-09 | Xrcc3 | X-ray repair ( | Nucleus |
| ENSMUSG00000053838 | 4.56E-10 | 1.22E-09 | Nudcd3 | NudC domai | Cytoplasm |
| ENSMUSG00000026921 | 4.61E-10 | 1.23E-09 | Egfl7 | EGF like dom | Extracellular S |
| ENSMUSG00000059669 | 4.61E-10 | 1.23E-09 | Taf1b | TATA-box bi | Nucleus |
| ENSMUSG00000027829 | 4.61E-10 | 1.23E-09 | Ccnl1 | cyclin L1 | Nucleus |
| ENSMUSG00000010025 | 4.64E-10 | 1.24E-09 | Aldh3a2 | aldehyde del | Cytoplasm |
| ENSMUSG00000002052 | 4.68E-10 | 1.25E-09 | Supt6 | SPT6 homolc | Nucleus |
| ENSMUSG00000005483 | 4.80E-10 | 1.28E-09 | Dnajb1 | DnaJ heat sh | Nucleus |
| ENSMUSG00000028920 | 4.84E-10 | 1.29E-09 | Fbxo42 | F-box protei | Other |
| ENSMUSG00000055184 | 4.85E-10 | 1.29E-09 | Fam72a | family with se | Cytoplasm |
| ENSMUSG00000024795 | 4.89E-10 | 1.31E-09 | Kif20b | kinesin family | Nucleus |
| ENSMUSG00000028809 | 4.89E-10 | 1.31E-09 | Srrm1 | serine/argini | Nucleus |
| ENSMUSG00000075327 | 4.90E-10 | 1.31E-09 | Zbtb2 | zinc finger ar | Nucleus |
| ENSMUSG00000022314 | 4.90E-10 | 1.31E-09 | Rad21 | RAD21 cohes | Nucleus |
| ENSMUSG00000030795 | 4.97E-10 | 1.33E-09 | Fus | fused in sarc | Nucleus |
| ENSMUSG00000021932 | 4.98E-10 | 1.33E-09 | Rnaseh2b | ribonuclease | Nucleus |
| ENSMUSG00000025289 | 5.08E-10 | 1.35E-09 | Prdx4 | peroxiredoxi | Cytoplasm |
| ENSMUSG00000006335 | 5.08E-10 | 1.35E-09 | Tfpt | TCF3 fusion p | Nucleus |
| ENSMUSG00000028484 | 5.10E-10 | 1.36E-09 | Psip1 | PC4 and SFRS | Nucleus |
| ENSMUSG00000024816 | 5.19E-10 | 1.38E-09 | Frmd8 | FERM domai | Other |
| ENSMUSG00000037544 | 5.25E-10 | 1.40E-09 | Dlgap5 | DLG associat | Nucleus |
| ENSMUSG00000027531 | 5.25E-10 | 1.40E-09 | Impa1 | inositol mon | Cytoplasm |
| ENSMUSG00000002010 | 5.34E-10 | 1.42E-09 | Idh3g | isocitrate del | Cytoplasm |
| ENSMUSG00000018209 | 5.58E-10 | 1.48E-09 | Stk4 | serine/threor | Cytoplasm |
| ENSMUSG00000060950 | 5.59E-10 | 1.49E-09 | Trmt61a | tRNA methyl | Nucleus |
| ENSMUSG00000022438 | 5.59E-10 | 1.49E-09 | Parvb | parvin beta | Cytoplasm |
| ENSMUSG00000063019 | 5.60E-10 | 1.49E-09 | Manbal | mannosidase | Other |
| ENSMUSG00000030533 | 5.62E-10 | 1.49E-09 | Unc45a | unc-45 myos | Plasma Mem |
| ENSMUSG00000034343 | 5.62E-10 | 1.49E-09 | Ube2f | ubiquitin cor | Cytoplasm |
| ENSMUSG00000043501 | 5.75E-10 | 1.53E-09 | Lgals2 | galectin 2 | Cytoplasm |
| ENSMUSG00000042557 | 5.75E-10 | 1.53E-09 | Sin3a | SIN3 transcri | Nucleus |
| ENSMUSG00000020594 | 5.84E-10 | 1.55E-09 | Pum2 | pumilio RNA | Cytoplasm |
| ENSMUSG00000041124 | 5.84E-10 | 1.55E-09 | Msantd4 | Myb/SANT D | Nucleus |
| ENSMUSG00000075706 | 5.88E-10 | 1.56E-09 | Gpx4 | glutathione p | Cytoplasm |
| ENSMUSG00000021962 | 5.97E-10 | 1.58E-09 | Dcp1a | decapping m | Nucleus |
| ENSMUSG00000052926 | 5.97E-10 | 1.58E-09 | Rnaseh2a | ribonuclease | Nucleus |
| ENSMUSG00000098134 | 6.05E-10 | 1.60E-09 | Rnf113a2 | ring finger pr | Extracellular S |
| ENSMUSG00000034345 | 6.10E-10 | 1.62E-09 | Gtf2h5 | general trans | Nucleus |
| ENSMUSG00000014504 | 6.21E-10 | 1.64E-09 | Srp19 | signal recogr | Cytoplasm |
| ENSMUSG00000028149 | 6.25E-10 | 1.65E-09 | Rap1gds1 | Rap1 GTPase | Cytoplasm |
| ENSMUSG00000054237 | 6.29E-10 | 1.66E-09 | Fra10ac1 | FRA10A asso | Nucleus |
| ENSMUSG00000005804 | 6.32E-10 | 1.67E-09 | Bloc1s6 | biogenesis o | Cytoplasm |
| ENSMUSG00000028274 | 6.44E-10 | 1.70E-09 | Rngtt | RNA guanyly | Nucleus |
| ENSMUSG00000018593 | 6.60E-10 | 1.75E-09 | Sparc | secreted prot | Extracellular S |
| ENSMUSG00000000568 | 6.69E-10 | 1.77E-09 | Hnrnpd | heterogeneo | Nucleus |
| ENSMUSG00000018567 | 6.79E-10 | 1.79E-09 | Gabarap | GABA type A | Cytoplasm |
| ENSMUSG00000000001 | 6.81E-10 | 1.80E-09 | Gnai3 | G protein sul | Cytoplasm |
| ENSMUSG00000038685 | 6.90E-10 | 1.82E-09 | Rtel1 | regulator of t | Nucleus |
| ENSMUSG00000025470 | 6.96E-10 | 1.83E-09 | Zfp511 | zinc finger pr | Nucleus |
| ENSMUSG00000027680 | 7.15E-10 | 1.89E-09 | Fxr1 | FMR1 autosc | Cytoplasm |
| ENSMUSG00000060477 | 7.16E-10 | 1.89E-09 | Irak2 | interleukin 1 | Plasma Mem |
| ENSMUSG00000027613 | 7.17E-10 | 1.89E-09 | Eif6 | eukaryotic tra | Cytoplasm |

| | | | | | |
|---|---|---|---|---|---|
| ENSMUSG00000033808 | 7.24E-10 | 1.91E-09 | Tmem87a | transmembra | Cytoplasm |
| ENSMUSG00000027048 | 7.28E-10 | 1.92E-09 | Abcb11 | ATP binding | Plasma Mem |
| ENSMUSG00000029860 | 7.34E-10 | 1.93E-09 | Zyx | zyxin | Plasma Mem |
| ENSMUSG00000004561 | 7.48E-10 | 1.97E-09 | Mettl17 | methyltransfe | Nucleus |
| ENSMUSG00000032459 | 7.48E-10 | 1.97E-09 | Mrps22 | mitochondria | Cytoplasm |
| ENSMUSG00000066149 | 7.55E-10 | 1.99E-09 | Cdc26 | cell division c | Nucleus |
| ENSMUSG00000031951 | 7.65E-10 | 2.01E-09 | Tmem231 | transmembra | Extracellular S |
| ENSMUSG00000027011 | 7.66E-10 | 2.01E-09 | Ube2e3 | ubiquitin con | Nucleus |
| ENSMUSG00000034189 | 7.69E-10 | 2.02E-09 | Hsdl1 | hydroxystero | Cytoplasm |
| ENSMUSG00000024308 | 7.78E-10 | 2.04E-09 | Tapbp | TAP binding | Cytoplasm |
| ENSMUSG00000027422 | 7.79E-10 | 2.05E-09 | Rrbp1 | ribosome bin | Cytoplasm |
| ENSMUSG00000040225 | 7.84E-10 | 2.06E-09 | Prrc2c | proline rich c | Cytoplasm |
| ENSMUSG00000047554 | 7.92E-10 | 2.08E-09 | Tmem41b | transmembra | Other |
| ENSMUSG00000025135 | 7.93E-10 | 2.08E-09 | Anapc11 | anaphase pro | Cytoplasm |
| ENSMUSG00000001289 | 8.05E-10 | 2.11E-09 | Pfdn5 | prefoldin sub | Nucleus |
| ENSMUSG00000037458 | 8.06E-10 | 2.11E-09 | Azin1 | antizyme inh | Cytoplasm |
| ENSMUSG00000063605 | 8.21E-10 | 2.15E-09 | Ccdc102a | coiled-coil de | Other |
| ENSMUSG00000042625 | 8.23E-10 | 2.16E-09 | Safb2 | scaffold attac | Cytoplasm |
| ENSMUSG00000028766 | 8.35E-10 | 2.19E-09 | Alpl | alkaline phos | Plasma Mem |
| ENSMUSG00000022607 | 8.43E-10 | 2.21E-09 | Ptk2 | protein tyros | Cytoplasm |
| ENSMUSG00000055782 | 8.61E-10 | 2.25E-09 | Abcd2 | ATP binding | Cytoplasm |
| ENSMUSG00000034021 | 8.69E-10 | 2.27E-09 | Pds5b | PDS5 cohesin | Nucleus |
| ENSMUSG00000047228 | 8.77E-10 | 2.29E-09 | BC048546 | alpha-2-mac | Extracellular S |
| ENSMUSG00000030016 | 8.84E-10 | 2.31E-09 | Zfml | zinc finger pr | Nucleus |
| ENSMUSG00000005823 | 8.93E-10 | 2.33E-09 | Gpr108 | G protein-co | Plasma Mem |
| ENSMUSG00000014355 | 8.98E-10 | 2.35E-09 | Anapc1 | anaphase pro | Nucleus |
| ENSMUSG00000028863 | 8.99E-10 | 2.35E-09 | Meaf6 | MYST/Esa1 a | Nucleus |
| ENSMUSG00000038446 | 9.00E-10 | 2.35E-09 | Cdc40 | cell division c | Nucleus |
| ENSMUSG00000039483 | 9.24E-10 | 2.41E-09 | Asb6 | ankyrin repea | Nucleus |
| ENSMUSG00000020017 | 9.24E-10 | 2.41E-09 | Hal | histidine amm | Cytoplasm |
| ENSMUSG00000003420 | 9.30E-10 | 2.43E-09 | Fcgrt | Fc fragment o | Plasma Mem |
| ENSMUSG00000034621 | 9.49E-10 | 2.48E-09 | Gpatch8 | G-patch dom | Other |
| ENSMUSG00000041319 | 9.63E-10 | 2.51E-09 | Thoc6 | THO comple: | Nucleus |
| ENSMUSG00000033114 | 9.65E-10 | 2.52E-09 | Slc35d2 | solute carrier | Cytoplasm |
| ENSMUSG00000025371 | 9.66E-10 | 2.52E-09 | Chmp6 | charged mult | Cytoplasm |
| ENSMUSG00000022983 | 9.79E-10 | 2.55E-09 | Scaf4 | SR-related C | Nucleus |
| ENSMUSG00000054302 | 1.01E-09 | 2.62E-09 | Eapp | E2F associate | Cytoplasm |
| ENSMUSG00000001729 | 1.02E-09 | 2.66E-09 | Akt1 | AKT serine/th | Cytoplasm |
| ENSMUSG00000020271 | 1.03E-09 | 2.67E-09 | Fbxw11 | F-box and W | Cytoplasm |
| ENSMUSG00000042520 | 1.05E-09 | 2.74E-09 | Ubap2l | ubiquitin-ass | Nucleus |
| ENSMUSG00000026857 | 1.05E-09 | 2.74E-09 | Ntmt1 | N-terminal X | Nucleus |
| ENSMUSG00000023967 | 1.06E-09 | 2.76E-09 | Mrps18a | mitochondria | Cytoplasm |
| ENSMUSG00000037089 | 1.06E-09 | 2.77E-09 | Slc35b2 | solute carrier | Cytoplasm |
| ENSMUSG00000095098 | 1.07E-09 | 2.77E-09 | Ccdc85b | coiled-coil de | Cytoplasm |
| ENSMUSG00000027322 | 1.09E-09 | 2.82E-09 | Siglec1 | sialic acid bin | Plasma Mem |
| ENSMUSG00000041629 | 1.09E-09 | 2.84E-09 | Fam104a | family with se | Other |
| ENSMUSG00000033295 | 1.12E-09 | 2.89E-09 | Ptprf | protein tyros | Plasma Mem |
| ENSMUSG00000022108 | 1.12E-09 | 2.91E-09 | Itm2b | integral mem | Plasma Mem |
| ENSMUSG00000071052 | 1.13E-09 | 2.93E-09 | Rpl7a-ps5 | ribosomal pro | Other |
| ENSMUSG00000033760 | 1.13E-09 | 2.94E-09 | Rbm4b | RNA binding | Nucleus |
| ENSMUSG00000035413 | 1.15E-09 | 2.98E-09 | Tmem98 | transmembra | Cytoplasm |
| ENSMUSG00000030220 | 1.18E-09 | 3.05E-09 | Arhgdib | Rho GDP diss | Cytoplasm |
| ENSMUSG00000028560 | 1.18E-09 | 3.06E-09 | Usp1 | ubiquitin spe | Cytoplasm |
| ENSMUSG00000016308 | 1.18E-09 | 3.06E-09 | Ube2a | ubiquitin con | Cytoplasm |
| ENSMUSG00000024891 | 1.20E-09 | 3.10E-09 | Slc29a2 | solute carrier | Plasma Mem |
| ENSMUSG00000031730 | 1.20E-09 | 3.12E-09 | Dhodh | dihydroorota | Cytoplasm |
| ENSMUSG00000023034 | 1.22E-09 | 3.15E-09 | Nr4a1 | nuclear recep | Nucleus |
| ENSMUSG00000029507 | 1.22E-09 | 3.16E-09 | Pus1 | pseudouridyl | Nucleus |

| | | | | | |
|---|---|---|---|---|---|
| ENSMUSG00000013921 | 1.23E-09 | 3.18E-09 | Clip3 | CAP-Gly don | Plasma Mem |
| ENSMUSG00000020235 | 1.23E-09 | 3.18E-09 | Fzr1 | fizzy and cell | Nucleus |
| ENSMUSG00000028282 | 1.25E-09 | 3.23E-09 | Casp8ap2 | caspase 8 ass | Nucleus |
| ENSMUSG00000066306 | 1.25E-09 | 3.23E-09 | Numa1 | nuclear mitot | Nucleus |
| ENSMUSG00000022781 | 1.27E-09 | 3.29E-09 | Pak2 | p21 (RAC1) a | Cytoplasm |
| ENSMUSG00000040325 | 1.28E-09 | 3.30E-09 | Vprbp | DDB1 and CU | Nucleus |
| ENSMUSG00000022451 | 1.28E-09 | 3.30E-09 | Twf1 | twinfilin actin | Cytoplasm |
| ENSMUSG00000078765 | 1.30E-09 | 3.36E-09 | U2af1l4 | U2 small nuc | Other |
| ENSMUSG00000031297 | 1.31E-09 | 3.39E-09 | Slc7a3 | solute carrier | Plasma Mem |
| ENSMUSG00000032187 | 1.33E-09 | 3.43E-09 | Smarca4 | SWI/SNF rela | Nucleus |
| ENSMUSG00000030286 | 1.36E-09 | 3.50E-09 | Emc3 | ER membran | Nucleus |
| ENSMUSG00000024369 | 1.38E-09 | 3.56E-09 | Nelfe | negative elor | Nucleus |
| ENSMUSG00000078970 | 1.39E-09 | 3.58E-09 | Wdr92 | WD repeat d | Other |
| ENSMUSG00000026309 | 1.39E-09 | 3.58E-09 | Ilkap | ILK associate | Cytoplasm |
| ENSMUSG00000021109 | 1.40E-09 | 3.60E-09 | Hif1a | hypoxia indu | Nucleus |
| ENSMUSG00000024354 | 1.42E-09 | 3.66E-09 | Slc23a1 | solute carrier | Plasma Mem |
| ENSMUSG00000033701 | 1.44E-09 | 3.71E-09 | Acbd6 | acyl-CoA bin | Cytoplasm |
| ENSMUSG00000035597 | 1.45E-09 | 3.72E-09 | Prpf39 | pre-mRNA p | Nucleus |
| ENSMUSG00000036990 | 1.47E-09 | 3.78E-09 | Otud4 | OTU deubiqu | Other |
| ENSMUSG00000008200 | 1.47E-09 | 3.78E-09 | Fnbp4 | formin bindir | Nucleus |
| ENSMUSG00000022559 | 1.48E-09 | 3.79E-09 | Fbxl6 | F-box and le | Other |
| ENSMUSG00000024381 | 1.48E-09 | 3.80E-09 | Bin1 | bridging inte | Nucleus |
| ENSMUSG00000031134 | 1.53E-09 | 3.93E-09 | Rbmx | RNA binding | Nucleus |
| ENSMUSG00000035828 | 1.55E-09 | 3.97E-09 | Pim3 | Pim-3 proto- | Cytoplasm |
| ENSMUSG00000034518 | 1.57E-09 | 4.03E-09 | Hmgxb4 | HMG-box co | Nucleus |
| ENSMUSG00000030560 | 1.58E-09 | 4.06E-09 | Ctsc | cathepsin C | Cytoplasm |
| ENSMUSG00000019868 | 1.58E-09 | 4.06E-09 | Vta1 | vesicle traffic | Cytoplasm |
| ENSMUSG00000017861 | 1.59E-09 | 4.08E-09 | Mybl2 | MYB proto-c | Nucleus |
| ENSMUSG00000006699 | 1.61E-09 | 4.12E-09 | Cdc42 | cell division c | Plasma Mem |
| ENSMUSG00000073139 | 1.62E-09 | 4.15E-09 | BC023829 | transmembra | Nucleus |
| ENSMUSG00000031556 | 1.63E-09 | 4.16E-09 | Tm2d2 | TM2 domain | Extracellular S |
| ENSMUSG00000040549 | 1.67E-09 | 4.27E-09 | Ckap5 | cytoskeleton | Nucleus |
| ENSMUSG00000015697 | 1.68E-09 | 4.29E-09 | Setdb1 | SET domain I | Nucleus |
| ENSMUSG00000018322 | 1.70E-09 | 4.35E-09 | Tomm34 | translocase o | Cytoplasm |
| ENSMUSG00000017146 | 1.70E-09 | 4.36E-09 | Brca1 | BRCA1, DNA | Nucleus |
| ENSMUSG00000031148 | 1.70E-09 | 4.36E-09 | Gpkow | G-patch dom | Nucleus |
| ENSMUSG00000029536 | 1.71E-09 | 4.38E-09 | Gatc | glutamyl-tRN | Nucleus |
| ENSMUSG00000021102 | 1.74E-09 | 4.44E-09 | Glrx5 | glutaredoxin | Cytoplasm |
| ENSMUSG00000021779 | 1.76E-09 | 4.50E-09 | Thrb | thyroid horm | Nucleus |
| ENSMUSG00000047777 | 1.82E-09 | 4.64E-09 | Phf13 | PHD finger p | Nucleus |
| ENSMUSG00000028992 | 1.83E-09 | 4.68E-09 | Nmnat1 | nicotinamide | Nucleus |
| ENSMUSG00000019907 | 1.84E-09 | 4.69E-09 | Ppp1r12a | protein phos | Cytoplasm |
| ENSMUSG00000026627 | 1.85E-09 | 4.71E-09 | Tmem206 | transmembra | Plasma Mem |
| ENSMUSG00000041199 | 1.85E-09 | 4.72E-09 | Rpusd1 | RNA pseudo | Other |
| ENSMUSG00000042207 | 1.87E-09 | 4.77E-09 | Kdm5b | lysine demet | Nucleus |
| ENSMUSG00000015733 | 1.87E-09 | 4.77E-09 | Capza2 | capping actir | Cytoplasm |
| ENSMUSG00000024218 | 1.96E-09 | 5.00E-09 | Taf11 | TATA-box bi | Nucleus |
| ENSMUSG00000003166 | 1.97E-09 | 5.02E-09 | Dgcr2 | DiGeorge syr | Plasma Mem |
| ENSMUSG00000050248 | 1.97E-09 | 5.02E-09 | Evc2 | EvC ciliary co | Extracellular S |
| ENSMUSG00000020561 | 1.98E-09 | 5.03E-09 | Twistnb | TWIST neighl | Nucleus |
| ENSMUSG00000019577 | 2.00E-09 | 5.08E-09 | Pdk4 | pyruvate deh | Cytoplasm |
| ENSMUSG00000023206 | 2.02E-09 | 5.13E-09 | Il15ra | interleukin 15 | Plasma Mem |
| ENSMUSG00000022106 | 2.02E-09 | 5.14E-09 | Rcbtb2 | RCC1 and BT | Cytoplasm |
| ENSMUSG00000024583 | 2.03E-09 | 5.15E-09 | Txnl1 | thioredoxin li | Cytoplasm |
| ENSMUSG00000046794 | 2.05E-09 | 5.21E-09 | Ppp1r3b | protein phos | Cytoplasm |
| ENSMUSG00000022185 | 2.06E-09 | 5.23E-09 | Acin1 | apoptotic chr | Nucleus |
| ENSMUSG00000046691 | 2.07E-09 | 5.24E-09 | Chtf8 | chromosome | Cytoplasm |
| ENSMUSG00000067038 | 2.08E-09 | 5.28E-09 | Rps12-ps3 | ribosomal pr | Other |

| | | | | | |
|---|---|---|---|---|---|
| ENSMUSG00000032329 | 2.13E-09 | 5.40E-09 | Hmg20a | high mobility | Nucleus |
| ENSMUSG00000001964 | 2.15E-09 | 5.46E-09 | Emd | emerin | Nucleus |
| ENSMUSG00000034278 | 2.18E-09 | 5.54E-09 | Dnajc17 | DnaJ heat sh | Other |
| ENSMUSG00000005299 | 2.22E-09 | 5.62E-09 | Letm1 | leucine zippe | Cytoplasm |
| ENSMUSG00000001285 | 2.23E-09 | 5.64E-09 | Myg1 | chromosome | Nucleus |
| ENSMUSG00000024194 | 2.25E-09 | 5.71E-09 | Cuta | cutA divalent | Cytoplasm |
| ENSMUSG00000027132 | 2.26E-09 | 5.72E-09 | Katnbl1 | katanin regul | Nucleus |
| ENSMUSG00000031918 | 2.29E-09 | 5.79E-09 | Mtmr2 | myotubularin | Cytoplasm |
| ENSMUSG00000018160 | 2.30E-09 | 5.81E-09 | Med1 | mediator cor | Nucleus |
| ENSMUSG00000041264 | 2.31E-09 | 5.83E-09 | Uspl1 | ubiquitin spe | Cytoplasm |
| ENSMUSG00000025781 | 2.35E-09 | 5.94E-09 | Atp5c1 | ATP synthase | Cytoplasm |
| ENSMUSG00000036270 | 2.36E-09 | 5.97E-09 | Edc4 | enhancer of | Cytoplasm |
| ENSMUSG00000027997 | 2.38E-09 | 6.01E-09 | Casp6 | caspase 6 | Cytoplasm |
| ENSMUSG00000037787 | 2.40E-09 | 6.07E-09 | Apopt1 | apoptogenic | Cytoplasm |
| ENSMUSG00000023939 | 2.41E-09 | 6.08E-09 | Mrpl14 | mitochondria | Cytoplasm |
| ENSMUSG00000029863 | 2.55E-09 | 6.44E-09 | Casp2 | caspase 2 | Cytoplasm |
| ENSMUSG00000021711 | 2.59E-09 | 6.55E-09 | Trappc13 | trafficking pr | Cytoplasm |
| ENSMUSG00000079508 | 2.61E-09 | 6.59E-09 | Apoo | apolipoprote | Extracellular S |
| ENSMUSG00000025246 | 2.62E-09 | 6.61E-09 | Tbl1x | transducin be | Nucleus |
| ENSMUSG00000062761 | 2.63E-09 | 6.63E-09 | Zfp512 | zinc finger pr | Nucleus |
| ENSMUSG00000046997 | 2.65E-09 | 6.68E-09 | Spsb4 | splA/ryanodi | Cytoplasm |
| ENSMUSG00000019988 | 2.74E-09 | 6.91E-09 | Nedd1 | neural precu | Cytoplasm |
| ENSMUSG00000003435 | 2.78E-09 | 7.01E-09 | Supt5 | SPT5 homolo | Nucleus |
| ENSMUSG00000031231 | 2.85E-09 | 7.18E-09 | Cox7b | cytochrome c | Cytoplasm |
| ENSMUSG00000026283 | 2.88E-09 | 7.25E-09 | Ing5 | inhibitor of g | Nucleus |
| ENSMUSG00000001034 | 2.88E-09 | 7.25E-09 | Mapk7 | mitogen-acti | Cytoplasm |
| ENSMUSG00000027510 | 2.89E-09 | 7.27E-09 | Rbm38 | RNA binding | Nucleus |
| ENSMUSG00000037343 | 2.94E-09 | 7.38E-09 | Taf2 | TATA-box bi | Nucleus |
| ENSMUSG00000050445 | 2.94E-09 | 7.40E-09 | Cyp8b1 | cytochrome P | Cytoplasm |
| ENSMUSG00000010358 | 2.97E-09 | 7.47E-09 | Ifi35 | interferon inc | Nucleus |
| ENSMUSG00000046591 | 2.99E-09 | 7.52E-09 | Ticrr | TOPBP1 inter | Nucleus |
| ENSMUSG00000017715 | 3.05E-09 | 7.66E-09 | Pgs1 | phosphatidyl | Cytoplasm |
| ENSMUSG00000021428 | 3.06E-09 | 7.68E-09 | Riok1 | RIO kinase 1 | Cytoplasm |
| ENSMUSG00000029578 | 3.07E-09 | 7.70E-09 | Wipi2 | WD repeat d | Cytoplasm |
| ENSMUSG00000026280 | 3.15E-09 | 7.89E-09 | Atg4b | autophagy re | Cytoplasm |
| ENSMUSG00000030868 | 3.16E-09 | 7.93E-09 | Dctn5 | dynactin sub | Cytoplasm |
| ENSMUSG00000040446 | 3.17E-09 | 7.94E-09 | Rprd1a | regulation of | Extracellular S |
| ENSMUSG00000035367 | 3.19E-09 | 7.99E-09 | Rmi1 | RecQ mediat | Nucleus |
| ENSMUSG00000048799 | 3.22E-09 | 8.06E-09 | Cep120 | centrosomal | Cytoplasm |
| ENSMUSG00000042210 | 3.28E-09 | 8.22E-09 | Abhd14a | abhydrolase | Cytoplasm |
| ENSMUSG00000022793 | 3.30E-09 | 8.26E-09 | B4galt4 | beta-1,4-gal | Cytoplasm |
| ENSMUSG00000051579 | 3.34E-09 | 8.35E-09 | Tceal8 | transcription | Other |
| ENSMUSG00000027326 | 3.36E-09 | 8.41E-09 | Casc5 | kinetochore s | Nucleus |
| ENSMUSG00000038007 | 3.43E-09 | 8.57E-09 | Acer2 | alkaline cerar | Cytoplasm |
| ENSMUSG00000051839 | 3.45E-09 | 8.62E-09 | Gypa | glycophorin A | Plasma Mem |
| ENSMUSG00000002550 | 3.46E-09 | 8.65E-09 | Uck1 | uridine-cytid | Cytoplasm |
| ENSMUSG00000022378 | 3.56E-09 | 8.88E-09 | Fam49b | family with se | Extracellular S |
| ENSMUSG00000001666 | 3.56E-09 | 8.89E-09 | Ddt | D-dopachrom | Cytoplasm |
| ENSMUSG00000029152 | 3.62E-09 | 9.04E-09 | Ociad1 | OCIA domain | Cytoplasm |
| ENSMUSG00000029148 | 3.64E-09 | 9.08E-09 | Nrbp1 | nuclear recep | Nucleus |
| ENSMUSG00000033411 | 3.66E-09 | 9.12E-09 | Ctdspl2 | CTD small ph | Nucleus |
| ENSMUSG00000020196 | 3.68E-09 | 9.17E-09 | Cabin1 | calcineurin b | Nucleus |
| ENSMUSG00000047989 | 3.68E-09 | 9.17E-09 | Ino80c | INO80 comp | Nucleus |
| ENSMUSG00000005262 | 3.74E-09 | 9.33E-09 | Ufd1l | ubiquitin rec | Cytoplasm |
| ENSMUSG00000021395 | 3.80E-09 | 9.46E-09 | Spin1 | spindlin 1 | Nucleus |
| ENSMUSG00000032563 | 3.83E-09 | 9.55E-09 | Mrpl3 | mitochondria | Cytoplasm |
| ENSMUSG00000034205 | 3.88E-09 | 9.68E-09 | Loxl2 | lysyl oxidase | Extracellular S |
| ENSMUSG00000021928 | 3.90E-09 | 9.72E-09 | Ebpl | EBP like | Cytoplasm |

| | | | | | |
|---|---|---|---|---|---|
| ENSMUSG00000028089 | 3.94E-09 | 9.80E-09 | Chd1l | chromodoma | Nucleus |
| ENSMUSG00000033157 | 3.94E-09 | 9.81E-09 | Abhd10 | abhydrolase | Cytoplasm |
| ENSMUSG00000020380 | 3.99E-09 | 9.93E-09 | Rad50 | RAD50 doub | Nucleus |
| ENSMUSG00000021144 | 3.99E-09 | 9.93E-09 | Mta1 | metastasis as | Nucleus |
| ENSMUSG00000028811 | 4.00E-09 | 9.95E-09 | Yars | tyrosyl-tRNA | Cytoplasm |
| ENSMUSG00000027166 | 4.03E-09 | 1.00E-08 | Dnajc24 | DnaJ heat sh | Other |
| ENSMUSG00000032512 | 4.21E-09 | 1.05E-08 | Wdr48 | WD repeat d | Cytoplasm |
| ENSMUSG00000025145 | 4.21E-09 | 1.05E-08 | Lrrc45 | leucine rich r | Cytoplasm |
| ENSMUSG00000052763 | 4.24E-09 | 1.05E-08 | Zfp212 | zinc finger pr | Nucleus |
| ENSMUSG00000032239 | 4.38E-09 | 1.09E-08 | Rp9 | RP9, pre-mR | Nucleus |
| ENSMUSG00000028847 | 4.38E-09 | 1.09E-08 | Trappc3 | trafficking pr | Cytoplasm |
| ENSMUSG00000062683 | 4.41E-09 | 1.10E-08 | Atp5g2 | ATP synthase | Cytoplasm |
| ENSMUSG00000022710 | 4.44E-09 | 1.10E-08 | Usp7 | ubiquitin spe | Nucleus |
| ENSMUSG00000031875 | 4.46E-09 | 1.10E-08 | Cmtm3 | CKLF like MA | Extracellular S |
| ENSMUSG00000040462 | 4.54E-09 | 1.13E-08 | Os9 | OS9, endopla | Nucleus |
| ENSMUSG00000049299 | 4.58E-09 | 1.14E-08 | Trappc1 | trafficking pr | Cytoplasm |
| ENSMUSG00000002297 | 4.59E-09 | 1.14E-08 | Dbf4 | DBF4 zinc fin | Nucleus |
| ENSMUSG00000070811 | 4.62E-09 | 1.14E-08 | Sult2a2 | sulfotransfera | Cytoplasm |
| ENSMUSG00000020687 | 4.63E-09 | 1.15E-08 | Cdc27 | cell division c | Nucleus |
| ENSMUSG00000053094 | 4.65E-09 | 1.15E-08 | Tmem248 | transmembra | Other |
| ENSMUSG00000060032 | 4.70E-09 | 1.16E-08 | H2afj | H2A histone | Cytoplasm |
| ENSMUSG00000060427 | 4.85E-09 | 1.20E-08 | Zfp868 | zinc finger pr | Other |
| ENSMUSG00000028329 | 4.89E-09 | 1.21E-08 | Xpa | XPA, DNA da | Nucleus |
| ENSMUSG00000021715 | 4.92E-09 | 1.22E-08 | Cwc27 | CWC27 splic | Nucleus |
| ENSMUSG00000044627 | 4.94E-09 | 1.22E-08 | Swi5 | SWI5 homolo | Nucleus |
| ENSMUSG00000057522 | 4.97E-09 | 1.23E-08 | Spop | speckle type | Extracellular S |
| ENSMUSG00000022324 | 5.00E-09 | 1.23E-08 | Matn2 | matrilin 2 | Extracellular S |
| ENSMUSG00000050471 | 5.02E-09 | 1.24E-08 | Fam118b | family with se | Other |
| ENSMUSG00000047719 | 5.03E-09 | 1.24E-08 | Ubiad1 | UbiA prenyltr | Nucleus |
| ENSMUSG00000036905 | 5.15E-09 | 1.27E-08 | C1qb | complement | Extracellular S |
| ENSMUSG00000007415 | 5.19E-09 | 1.28E-08 | Gatad1 | GATA zinc fir | Nucleus |
| ENSMUSG00000055491 | 5.24E-09 | 1.29E-08 | Pprc1 | peroxisome p | Nucleus |
| ENSMUSG00000006763 | 5.25E-09 | 1.29E-08 | Saal1 | serum amylo | Other |
| ENSMUSG00000025917 | 5.26E-09 | 1.30E-08 | Cops5 | COP9 signalo | Nucleus |
| ENSMUSG00000004895 | 5.32E-09 | 1.31E-08 | Prcc | PRCC, proline | Nucleus |
| ENSMUSG00000096795 | 5.33E-09 | 1.31E-08 | Zfp433 | zinc finger pr | Nucleus |
| ENSMUSG00000020923 | 5.45E-09 | 1.34E-08 | Ubtf | upstream bin | Nucleus |
| ENSMUSG00000025278 | 5.46E-09 | 1.34E-08 | Flnb | filamin B | Cytoplasm |
| ENSMUSG00000027162 | 5.55E-09 | 1.36E-08 | Lin7c | lin-7 homolo | Cytoplasm |
| ENSMUSG00000024824 | 5.56E-09 | 1.37E-08 | Rad9a | RAD9 checkp | Nucleus |
| ENSMUSG00000068744 | 5.62E-09 | 1.38E-08 | Psrc1 | proline and s | Cytoplasm |
| ENSMUSG00000032599 | 5.64E-09 | 1.38E-08 | Ip6k2 | inositol hexal | Cytoplasm |
| ENSMUSG00000040482 | 5.65E-09 | 1.39E-08 | Dxo | decapping ex | Nucleus |
| ENSMUSG00000039016 | 5.66E-09 | 1.39E-08 | Timm8b | translocase o | Cytoplasm |
| ENSMUSG00000038776 | 5.71E-09 | 1.40E-08 | Ephx1 | epoxide hydr | Cytoplasm |
| ENSMUSG00000038069 | 5.75E-09 | 1.41E-08 | Cdkn2aip | CDKN2A inte | Nucleus |
| ENSMUSG00000052310 | 5.79E-09 | 1.42E-08 | Slc39a1 | solute carrier | Plasma Mem |
| ENSMUSG00000072964 | 5.86E-09 | 1.44E-08 | Bhlhb9 | basic helix-lo | Cytoplasm |
| ENSMUSG00000024014 | 5.89E-09 | 1.44E-08 | Pim1 | Pim-1 proto- | Cytoplasm |
| ENSMUSG00000063804 | 5.96E-09 | 1.46E-08 | Lin28b | lin-28 homol | Cytoplasm |
| ENSMUSG00000055531 | 6.03E-09 | 1.48E-08 | Cpsf6 | cleavage and | Nucleus |
| ENSMUSG00000050017 | 6.11E-09 | 1.50E-08 | Pitpnb | phosphatidyl | Cytoplasm |
| ENSMUSG00000024847 | 6.13E-09 | 1.50E-08 | Aip | aryl hydrocar | Nucleus |
| ENSMUSG00000055334 | 6.13E-09 | 1.50E-08 | Snupn | snurportin 1 | Nucleus |
| ENSMUSG00000067365 | 6.14E-09 | 1.50E-08 | Tmem128 | transmembra | Other |
| ENSMUSG00000041974 | 6.18E-09 | 1.51E-08 | Spidr | scaffold prot | Nucleus |
| ENSMUSG00000031370 | 6.19E-09 | 1.51E-08 | Zrsr2 | zinc finger C | Nucleus |
| ENSMUSG00000079477 | 6.20E-09 | 1.52E-08 | Rab7 | RAB7A, mem | Cytoplasm |

| | | | | | |
|---|---|---|---|---|---|
| ENSMUSG00000028772 | 6.20E-09 | 1.52E-08 | Zcchc17 | zinc finger C( | Nucleus |
| ENSMUSG00000025538 | 6.21E-09 | 1.52E-08 | Sumf2 | sulfatase mo( | Cytoplasm |
| ENSMUSG00000034032 | 6.31E-09 | 1.54E-08 | Rpap1 | RNA polymer | Other |
| ENSMUSG00000020898 | 6.39E-09 | 1.56E-08 | Ctc1 | CST telomere | Nucleus |
| ENSMUSG00000026867 | 6.58E-09 | 1.61E-08 | Gapvd1 | GTPase activa | Cytoplasm |
| ENSMUSG00000021156 | 6.61E-09 | 1.61E-08 | Zmynd11 | zinc finger M | Nucleus |
| ENSMUSG00000026519 | 6.62E-09 | 1.62E-08 | Tmem63a | transmembra | Cytoplasm |
| ENSMUSG00000044066 | 6.63E-09 | 1.62E-08 | Cep68 | centrosomal | Cytoplasm |
| ENSMUSG00000026822 | 6.65E-09 | 1.62E-08 | Lcn2 | lipocalin 2 | Extracellular S |
| ENSMUSG00000007610 | 6.67E-09 | 1.62E-08 | Gtpbp3 | GTP binding | Cytoplasm |
| ENSMUSG00000039568 | 6.69E-09 | 1.63E-08 | Ubald1 | UBA like dom | Other |
| ENSMUSG00000021113 | 6.71E-09 | 1.63E-08 | Snapc1 | small nuclear | Nucleus |
| ENSMUSG00000021948 | 6.80E-09 | 1.66E-08 | Prkcd | protein kinas | Cytoplasm |
| ENSMUSG00000075470 | 6.89E-09 | 1.68E-08 | Alg10b | ALG10, alpha | Cytoplasm |
| ENSMUSG00000059851 | 6.97E-09 | 1.70E-08 | Suv420h2 | lysine methyl | Nucleus |
| ENSMUSG00000026615 | 7.06E-09 | 1.72E-08 | Eprs | glutamyl-pro | Cytoplasm |
| ENSMUSG00000025507 | 7.10E-09 | 1.72E-08 | Pidd1 | p53-induced | Cytoplasm |
| ENSMUSG00000022752 | 7.10E-09 | 1.72E-08 | Tomm70a | translocase o | Cytoplasm |
| ENSMUSG00000038500 | 7.11E-09 | 1.73E-08 | Prr3 | proline rich 3 | Other |
| ENSMUSG00000059278 | 7.18E-09 | 1.74E-08 | Naa38 | N(alpha)-ace | Nucleus |
| ENSMUSG00000032244 | 7.20E-09 | 1.75E-08 | Fem1b | fem-1 homo | Nucleus |
| ENSMUSG00000023170 | 7.24E-09 | 1.76E-08 | Gps2 | G protein pat | Nucleus |
| ENSMUSG00000031988 | 7.26E-09 | 1.76E-08 | Vps26b | VPS26, retror | Cytoplasm |
| ENSMUSG00000040236 | 7.29E-09 | 1.77E-08 | Trappc5 | trafficking pr | Cytoplasm |
| ENSMUSG00000037344 | 7.32E-09 | 1.78E-08 | Slc12a9 | solute carrier | Plasma Mem |
| ENSMUSG00000022679 | 7.33E-09 | 1.78E-08 | Mpv17l | MPV17 mitoc | Cytoplasm |
| ENSMUSG00000035678 | 7.34E-09 | 1.78E-08 | Tnfsf9 | tumor necros | Plasma Mem |
| ENSMUSG00000044702 | 7.40E-09 | 1.79E-08 | Palb2 | partner and l | Nucleus |
| ENSMUSG00000020423 | 7.47E-09 | 1.81E-08 | Btg2 | BTG anti-pro | Nucleus |
| ENSMUSG00000025194 | 7.49E-09 | 1.81E-08 | Abcc2 | ATP binding | Plasma Mem |
| ENSMUSG00000000409 | 7.56E-09 | 1.83E-08 | Lck | LCK proto-o | Cytoplasm |
| ENSMUSG00000008435 | 7.62E-09 | 1.85E-08 | Rdh13 | retinol dehyc | Cytoplasm |
| ENSMUSG00000020248 | 7.85E-09 | 1.90E-08 | Nfyb | nuclear trans | Nucleus |
| ENSMUSG00000052915 | 7.90E-09 | 1.91E-08 | Msl1 | male specific | Nucleus |
| ENSMUSG00000022337 | 8.09E-09 | 1.96E-08 | Emc2 | ER membran | Nucleus |
| ENSMUSG00000036813 | 8.27E-09 | 2.00E-08 | Entpd8 | ectonucleosi | Plasma Mem |
| ENSMUSG00000032640 | 8.35E-09 | 2.02E-08 | Chsy1 | chondroitin s | Cytoplasm |
| ENSMUSG00000073542 | 8.49E-09 | 2.05E-08 | Cep76 | centrosomal | Cytoplasm |
| ENSMUSG00000020211 | 8.55E-09 | 2.07E-08 | Sf3a2 | splicing facto | Nucleus |
| ENSMUSG00000055447 | 8.56E-09 | 2.07E-08 | Cd47 | CD47 molecu | Plasma Mem |
| ENSMUSG00000068114 | 9.09E-09 | 2.19E-08 | Ccdc134 | coiled-coil d | Other |
| ENSMUSG00000064145 | 9.15E-09 | 2.21E-08 | Arih2 | ariadne RBR | Nucleus |
| ENSMUSG00000024997 | 9.18E-09 | 2.22E-08 | Prdx3 | peroxiredoxi | Cytoplasm |
| ENSMUSG00000021228 | 9.23E-09 | 2.23E-08 | Acot3 | acyl-CoA thi | Cytoplasm |
| ENSMUSG00000063406 | 9.28E-09 | 2.24E-08 | Tmed5 | transmembra | Cytoplasm |
| ENSMUSG00000095677 | 9.30E-09 | 2.24E-08 | Dynlt1f | dynein light | Cytoplasm |
| ENSMUSG00000003421 | 9.66E-09 | 2.33E-08 | Nosip | nitric oxide s | Cytoplasm |
| ENSMUSG00000029664 | 9.76E-09 | 2.35E-08 | Tfpi2 | tissue factor | Extracellular S |
| ENSMUSG00000023089 | 9.77E-09 | 2.35E-08 | Ndufa5 | NADH:ubiqu | Cytoplasm |
| ENSMUSG00000024570 | 9.89E-09 | 2.38E-08 | Rbfa | ribosome bir | Cytoplasm |
| ENSMUSG00000036613 | 1.00E-08 | 2.41E-08 | Tssc1 | EARP comple | Other |
| ENSMUSG00000001665 | 1.01E-08 | 2.42E-08 | Gstt3 | glutathione S | Other |
| ENSMUSG00000033022 | 1.01E-08 | 2.42E-08 | Cdo1 | cysteine diox | Cytoplasm |
| ENSMUSG00000040699 | 1.01E-08 | 2.43E-08 | Limd2 | LIM domain | Other |
| ENSMUSG00000000168 | 1.02E-08 | 2.45E-08 | Dlat | dihydrolipoa | Cytoplasm |
| ENSMUSG00000022338 | 1.03E-08 | 2.48E-08 | Eny2 | ENY2, transc | Nucleus |
| ENSMUSG00000004936 | 1.04E-08 | 2.49E-08 | Map2k1 | mitogen-acti | Cytoplasm |
| ENSMUSG00000040466 | 1.04E-08 | 2.49E-08 | Blvrb | biliverdin red | Cytoplasm |

| | | | | | |
|---|---|---|---|---|---|
| ENSMUSG00000033065 | 1.04E-08 | 2.49E-08 | Pfkm | phosphofruc | Cytoplasm |
| ENSMUSG00000033554 | 1.05E-08 | 2.51E-08 | Dph5 | diphthamide | Cytoplasm |
| ENSMUSG00000029171 | 1.05E-08 | 2.52E-08 | Pgm1 | phosphogluc | Cytoplasm |
| ENSMUSG00000032481 | 1.06E-08 | 2.54E-08 | Smarcc1 | SWI/SNF rela | Nucleus |
| ENSMUSG00000020163 | 1.09E-08 | 2.61E-08 | Uqcr11 | ubiquinol-cy | Cytoplasm |
| ENSMUSG00000025809 | 1.09E-08 | 2.61E-08 | Itgb1 | integrin subu | Plasma Mem |
| ENSMUSG00000025266 | 1.10E-08 | 2.62E-08 | Gnl3l | G protein nu | Nucleus |
| ENSMUSG00000021519 | 1.10E-08 | 2.63E-08 | Mterf3 | mitochondria | Cytoplasm |
| ENSMUSG00000021540 | 1.10E-08 | 2.63E-08 | Smad5 | SMAD family | Nucleus |
| ENSMUSG00000019370 | 1.11E-08 | 2.66E-08 | Calm3 | calmodulin 1 | Nucleus |
| ENSMUSG00000061979 | 1.13E-08 | 2.70E-08 | Wbscr16 | RCC1 like | Cytoplasm |
| ENSMUSG00000036835 | 1.13E-08 | 2.70E-08 | Psenen | presenilin en | Plasma Mem |
| ENSMUSG00000002329 | 1.15E-08 | 2.74E-08 | Mdp1 | magnesium d | Cytoplasm |
| ENSMUSG00000034617 | 1.15E-08 | 2.75E-08 | Mtrr | 5-methyltetra | Cytoplasm |
| ENSMUSG00000018765 | 1.16E-08 | 2.76E-08 | Fxr2 | FMR1 autosc | Cytoplasm |
| ENSMUSG00000051469 | 1.17E-08 | 2.80E-08 | Zfp191 | zinc finger pr | Nucleus |
| ENSMUSG00000029221 | 1.17E-08 | 2.80E-08 | Slc30a9 | solute carrier | Nucleus |
| ENSMUSG00000073639 | 1.18E-08 | 2.82E-08 | Rab18 | RAB18, mem | Cytoplasm |
| ENSMUSG00000020415 | 1.18E-08 | 2.82E-08 | Pttg1 | pituitary tum | Nucleus |
| ENSMUSG00000053289 | 1.20E-08 | 2.86E-08 | Ddx10 | DEAD-box h | Nucleus |
| ENSMUSG00000021556 | 1.21E-08 | 2.89E-08 | Golm1 | golgi membr | Cytoplasm |
| ENSMUSG00000007570 | 1.22E-08 | 2.90E-08 | Fance | FA complem | Nucleus |
| ENSMUSG00000008604 | 1.23E-08 | 2.93E-08 | Ubqln4 | ubiquilin 4 | Cytoplasm |
| ENSMUSG00000026179 | 1.25E-08 | 2.97E-08 | Pnkd | PNKD, MBL c | Nucleus |
| ENSMUSG00000024878 | 1.25E-08 | 2.99E-08 | Cbwd1 | COBW doma | Other |
| ENSMUSG00000040842 | 1.28E-08 | 3.05E-08 | Szrd1 | SUZ RNA bin | Other |
| ENSMUSG00000042032 | 1.28E-08 | 3.06E-08 | Mat2b | methionine a | Cytoplasm |
| ENSMUSG00000021548 | 1.31E-08 | 3.11E-08 | Ccnh | cyclin H | Nucleus |
| ENSMUSG00000050796 | 1.31E-08 | 3.12E-08 | B3galt6 | beta-1,3-gal | Cytoplasm |
| ENSMUSG00000057788 | 1.34E-08 | 3.19E-08 | Ddx49 | DEAD-box h | Nucleus |
| ENSMUSG00000034171 | 1.35E-08 | 3.22E-08 | Faah | fatty acid am | Plasma Mem |
| ENSMUSG00000034931 | 1.36E-08 | 3.22E-08 | Dhx8 | DEAH-box h | Nucleus |
| ENSMUSG00000015627 | 1.36E-08 | 3.22E-08 | Gata5 | GATA bindin | Nucleus |
| ENSMUSG00000032834 | 1.36E-08 | 3.22E-08 | Pwp2 | PWP2 period | Nucleus |
| ENSMUSG00000031154 | 1.38E-08 | 3.27E-08 | Otud5 | OTU domain | Other |
| ENSMUSG00000022325 | 1.38E-08 | 3.28E-08 | Pop1 | POP1 homol | Nucleus |
| ENSMUSG00000021013 | 1.39E-08 | 3.29E-08 | Ttc8 | tetratricopep | Cytoplasm |
| ENSMUSG00000018858 | 1.40E-08 | 3.33E-08 | Ict1 | mitochondria | Cytoplasm |
| ENSMUSG00000021900 | 1.41E-08 | 3.33E-08 | Btd | biotinidase | Extracellular S |
| ENSMUSG00000024778 | 1.41E-08 | 3.34E-08 | Fas | Fas cell surfa | Plasma Mem |
| ENSMUSG00000020463 | 1.42E-08 | 3.36E-08 | Smek2 | protein phos | Nucleus |
| ENSMUSG00000023991 | 1.43E-08 | 3.39E-08 | Foxp4 | forkhead box | Nucleus |
| ENSMUSG00000030714 | 1.47E-08 | 3.47E-08 | Ccdc101 | SAGA compl | Nucleus |
| ENSMUSG00000030031 | 1.47E-08 | 3.49E-08 | Kbtbd8 | kelch repeat | Cytoplasm |
| ENSMUSG00000040888 | 1.48E-08 | 3.51E-08 | Gfer | growth facto | Nucleus |
| ENSMUSG00000024436 | 1.48E-08 | 3.51E-08 | Mrps18b | mitochondria | Cytoplasm |
| ENSMUSG00000021578 | 1.49E-08 | 3.52E-08 | Ccdc127 | coiled-coil d | Other |
| ENSMUSG00000024885 | 1.49E-08 | 3.53E-08 | Aldh3b1 | aldehyde deh | Cytoplasm |
| ENSMUSG00000047109 | 1.52E-08 | 3.60E-08 | Cldn14 | claudin 14 | Plasma Mem |
| ENSMUSG00000006373 | 1.53E-08 | 3.62E-08 | Pgrmc1 | progesterone | Plasma Mem |
| ENSMUSG00000050608 | 1.53E-08 | 3.62E-08 | Minos1 | mitochondria | Cytoplasm |
| ENSMUSG00000071477 | 1.54E-08 | 3.65E-08 | Zfp777 | zinc finger pr | Other |
| ENSMUSG00000017491 | 1.56E-08 | 3.68E-08 | Rarb | retinoic acid | Nucleus |
| ENSMUSG00000066839 | 1.56E-08 | 3.68E-08 | Ecsit | ECSIT signalli | Nucleus |
| ENSMUSG00000029455 | 1.59E-08 | 3.75E-08 | Aldh2 | aldehyde deh | Cytoplasm |
| ENSMUSG00000026972 | 1.59E-08 | 3.76E-08 | Arrdc1 | arrestin dom | Cytoplasm |
| ENSMUSG00000020522 | 1.59E-08 | 3.76E-08 | Mfap3 | microfibril as | Extracellular S |
| ENSMUSG00000035451 | 1.61E-08 | 3.81E-08 | Foxa1 | forkhead box | Nucleus |

| | | | | | |
|---|---|---|---|---|---|
| ENSMUSG00000039219 | 1.62E-08 | 3.82E-08 | Arid4b | AT-rich inter | Nucleus |
| ENSMUSG00000057228 | 1.63E-08 | 3.83E-08 | Aadat | aminoadipat | Cytoplasm |
| ENSMUSG00000057858 | 1.66E-08 | 3.91E-08 | Fam204a | family with se | Other |
| ENSMUSG00000002588 | 1.66E-08 | 3.91E-08 | Pon1 | paraoxonase | Extracellular S |
| ENSMUSG00000003039 | 1.66E-08 | 3.92E-08 | Fam32a | family with se | Nucleus |
| ENSMUSG00000033405 | 1.67E-08 | 3.93E-08 | Nudt15 | nudix hydrola | Cytoplasm |
| ENSMUSG00000013236 | 1.69E-08 | 3.97E-08 | Ptprs | protein tyros | Plasma Mem |
| ENSMUSG00000020585 | 1.70E-08 | 3.99E-08 | Laptm4a | lysosomal pr | Cytoplasm |
| ENSMUSG00000029554 | 1.71E-08 | 4.01E-08 | Mad1l1 | mitotic arrest | Nucleus |
| ENSMUSG00000041961 | 1.71E-08 | 4.02E-08 | Znrf3 | zinc and ring | Plasma Mem |
| ENSMUSG00000027014 | 1.71E-08 | 4.02E-08 | Cwc22 | CWC22 splic | Nucleus |
| ENSMUSG00000031878 | 1.72E-08 | 4.05E-08 | Nae1 | NEDD8 activa | Cytoplasm |
| ENSMUSG00000032705 | 1.73E-08 | 4.06E-08 | Exd2 | exonuclease | Other |
| ENSMUSG00000036840 | 1.74E-08 | 4.10E-08 | Siah1a | siah E3 ubiqu | Nucleus |
| ENSMUSG00000006024 | 1.75E-08 | 4.10E-08 | Napa | NSF attachm | Cytoplasm |
| ENSMUSG00000028437 | 1.76E-08 | 4.14E-08 | Ubap1 | ubiquitin ass | Cytoplasm |
| ENSMUSG00000033364 | 1.77E-08 | 4.16E-08 | Usp37 | ubiquitin spe | Nucleus |
| ENSMUSG00000037628 | 1.78E-08 | 4.18E-08 | Cdkn3 | cyclin depen | Cytoplasm |
| ENSMUSG00000031138 | 1.79E-08 | 4.19E-08 | F9 | coagulation f | Extracellular S |
| ENSMUSG00000028480 | 1.81E-08 | 4.23E-08 | Glipr2 | GLI pathoger | Cytoplasm |
| ENSMUSG00000035439 | 1.82E-08 | 4.26E-08 | Haus8 | HAUS augmi | Cytoplasm |
| ENSMUSG00000030330 | 1.83E-08 | 4.28E-08 | Ing4 | inhibitor of g | Nucleus |
| ENSMUSG00000024253 | 1.83E-08 | 4.29E-08 | Dync2li1 | dynein cytop | Cytoplasm |
| ENSMUSG00000014074 | 1.85E-08 | 4.33E-08 | Rnf168 | ring finger pr | Nucleus |
| ENSMUSG00000079215 | 1.86E-08 | 4.35E-08 | Zfp664 | zinc finger pr | Nucleus |
| ENSMUSG00000016933 | 1.86E-08 | 4.36E-08 | Plcg1 | phospholipas | Cytoplasm |
| ENSMUSG00000056214 | 1.86E-08 | 4.37E-08 | Pard6g | par-6 family | Plasma Mem |
| ENSMUSG00000003062 | 1.88E-08 | 4.41E-08 | Stard3nl | STARD3 N-te | Cytoplasm |
| ENSMUSG00000041119 | 1.89E-08 | 4.43E-08 | Pde9a | phosphodies | Cytoplasm |
| ENSMUSG00000032582 | 1.90E-08 | 4.45E-08 | Rbm6 | RNA binding | Nucleus |
| ENSMUSG00000025899 | 1.91E-08 | 4.47E-08 | Alkbh8 | alkB homolo | Cytoplasm |
| ENSMUSG00000033054 | 1.93E-08 | 4.51E-08 | Npat | nuclear prote | Nucleus |
| ENSMUSG00000072772 | 1.96E-08 | 4.58E-08 | Grcc10 | chromosome | Cytoplasm |
| ENSMUSG00000022119 | 1.99E-08 | 4.64E-08 | Rbm26 | RNA binding | Other |
| ENSMUSG00000041763 | 1.99E-08 | 4.65E-08 | Tpp2 | tripeptidyl pe | Cytoplasm |
| ENSMUSG00000001750 | 2.01E-08 | 4.68E-08 | Tcirg1 | T cell immun | Plasma Mem |
| ENSMUSG00000021944 | 2.01E-08 | 4.70E-08 | Gata4 | GATA bindin | Nucleus |
| ENSMUSG00000015575 | 2.02E-08 | 4.73E-08 | Atp6v0e | ATPase H+ tr | Cytoplasm |
| ENSMUSG00000028898 | 2.03E-08 | 4.73E-08 | Trnau1ap | tRNA selenoc | Nucleus |
| ENSMUSG00000029505 | 2.07E-08 | 4.83E-08 | Ep400 | E1A binding | Nucleus |
| ENSMUSG00000022885 | 2.08E-08 | 4.86E-08 | St6gal1 | ST6 beta-gal | Cytoplasm |
| ENSMUSG00000040565 | 2.11E-08 | 4.91E-08 | Btaf1 | B-TFIID TATA | Nucleus |
| ENSMUSG00000030678 | 2.12E-08 | 4.95E-08 | Maz | MYC associat | Nucleus |
| ENSMUSG00000036858 | 2.13E-08 | 4.96E-08 | Ptcra | pre T cell ant | Plasma Mem |
| ENSMUSG00000042772 | 2.13E-08 | 4.96E-08 | Smg7 | SMG7, nonse | Nucleus |
| ENSMUSG00000058290 | 2.16E-08 | 5.03E-08 | Espl1 | extra spindle | Nucleus |
| ENSMUSG00000059363 | 2.19E-08 | 5.10E-08 | Fxn | frataxin | Cytoplasm |
| ENSMUSG00000029265 | 2.30E-08 | 5.35E-08 | Dr1 | down-regula | Nucleus |
| ENSMUSG00000031221 | 2.32E-08 | 5.40E-08 | Igbp1 | immunoglob | Cytoplasm |
| ENSMUSG00000058444 | 2.35E-08 | 5.46E-08 | Map2k5 | mitogen-acti | Cytoplasm |
| ENSMUSG00000043421 | 2.36E-08 | 5.49E-08 | Hilpda | hypoxia indu | Cytoplasm |
| ENSMUSG00000036327 | 2.36E-08 | 5.49E-08 | Qsox2 | quiescin sulfh | Other |
| ENSMUSG00000028923 | 2.39E-08 | 5.55E-08 | Necap2 | NECAP endo | Cytoplasm |
| ENSMUSG00000073411 | 2.40E-08 | 5.58E-08 | H2-D1 | major histoc | Plasma Mem |
| ENSMUSG00000042029 | 2.41E-08 | 5.59E-08 | Ncapg2 | non-SMC co | Nucleus |
| ENSMUSG00000031570 | 2.42E-08 | 5.63E-08 | Ppapdc1b | phospholipid | Other |
| ENSMUSG00000028175 | 2.63E-08 | 6.10E-08 | Depdc1a | DEP domain | Nucleus |
| ENSMUSG00000037032 | 2.65E-08 | 6.15E-08 | Apbb1 | amyloid beta | Cytoplasm |

| | | | | | |
|---|---|---|---|---|---|
| ENSMUSG00000037525 | 2.68E-08 | 6.23E-08 | Bcdin3d | BCDIN3 dom | Cytoplasm |
| ENSMUSG00000001909 | 2.70E-08 | 6.26E-08 | Trmt1 | tRNA methyl | Extracellular S |
| ENSMUSG00000047534 | 2.71E-08 | 6.30E-08 | Mis18bp1 | MIS18 bindin | Nucleus |
| ENSMUSG00000073616 | 2.74E-08 | 6.34E-08 | Myeov2 | COP9 signalo | Nucleus |
| ENSMUSG00000032869 | 2.75E-08 | 6.38E-08 | Psmf1 | proteasome i | Cytoplasm |
| ENSMUSG00000022828 | 2.79E-08 | 6.47E-08 | Gtf2e1 | general trans | Nucleus |
| ENSMUSG00000026492 | 2.80E-08 | 6.48E-08 | Tfb2m | transcription | Cytoplasm |
| ENSMUSG00000020453 | 2.82E-08 | 6.54E-08 | Patz1 | POZ/BTB and | Nucleus |
| ENSMUSG00000036103 | 2.83E-08 | 6.56E-08 | Colec12 | collectin subf | Plasma Mem |
| ENSMUSG00000029103 | 2.87E-08 | 6.64E-08 | Lrpap1 | LDL receptor | Plasma Mem |
| ENSMUSG00000040675 | 2.88E-08 | 6.67E-08 | Mthfd1l | methylenetet | Cytoplasm |
| ENSMUSG00000034659 | 2.91E-08 | 6.73E-08 | Tmem109 | transmembra | Cytoplasm |
| ENSMUSG00000025894 | 2.93E-08 | 6.77E-08 | Aasdhppt | aminoadipate | Cytoplasm |
| ENSMUSG00000022503 | 2.95E-08 | 6.82E-08 | Nubp1 | nucleotide bi | Cytoplasm |
| ENSMUSG00000038888 | 2.96E-08 | 6.85E-08 | Ctu1 | cytosolic thic | Cytoplasm |
| ENSMUSG00000003526 | 2.98E-08 | 6.89E-08 | Prodh | proline dehy | Cytoplasm |
| ENSMUSG00000025521 | 3.00E-08 | 6.93E-08 | Tmem192 | transmembra | Cytoplasm |
| ENSMUSG00000023755 | 3.01E-08 | 6.96E-08 | Rhebl1 | RHEB like 1 | Cytoplasm |
| ENSMUSG00000024240 | 3.05E-08 | 7.05E-08 | Epc1 | enhancer of p | Nucleus |
| ENSMUSG00000026427 | 3.06E-08 | 7.06E-08 | Eif2d | eukaryotic tra | Cytoplasm |
| ENSMUSG00000034112 | 3.08E-08 | 7.11E-08 | Atp2c2 | ATPase secre | Cytoplasm |
| ENSMUSG00000022244 | 3.09E-08 | 7.13E-08 | Amacr | alpha-methy | Cytoplasm |
| ENSMUSG00000055319 | 3.10E-08 | 7.15E-08 | Sec23ip | SEC23 intera | Cytoplasm |
| ENSMUSG00000020978 | 3.11E-08 | 7.18E-08 | Klhdc2 | kelch domain | Cytoplasm |
| ENSMUSG00000020495 | 3.12E-08 | 7.20E-08 | Smg8 | SMG8, nonse | Cytoplasm |
| ENSMUSG00000021884 | 3.12E-08 | 7.20E-08 | Hacl1 | 2-hydroxyacy | Cytoplasm |
| ENSMUSG00000026878 | 3.12E-08 | 7.20E-08 | Rab14 | RAB14, mem | Cytoplasm |
| ENSMUSG00000037337 | 3.13E-08 | 7.20E-08 | Map4k1 | mitogen-acti | Cytoplasm |
| ENSMUSG00000027628 | 3.16E-08 | 7.28E-08 | Aar2 | AAR2 splicing | Other |
| ENSMUSG00000029174 | 3.19E-08 | 7.33E-08 | Tbc1d1 | TBC1 domain | Nucleus |
| ENSMUSG00000024335 | 3.23E-08 | 7.43E-08 | Brd2 | bromodomai | Nucleus |
| ENSMUSG00000039456 | 3.23E-08 | 7.43E-08 | Morc3 | MORC family | Nucleus |
| ENSMUSG00000028411 | 3.26E-08 | 7.51E-08 | Aptx | aprataxin | Nucleus |
| ENSMUSG00000032374 | 3.30E-08 | 7.58E-08 | Plod2 | procollagen- | Cytoplasm |
| ENSMUSG00000036721 | 3.43E-08 | 7.89E-08 | Zscan12 | zinc finger ar | Nucleus |
| ENSMUSG00000033149 | 3.46E-08 | 7.94E-08 | Phldb2 | pleckstrin ho | Cytoplasm |
| ENSMUSG00000034152 | 3.47E-08 | 7.98E-08 | Exoc3 | exocyst comp | Plasma Mem |
| ENSMUSG00000020230 | 3.50E-08 | 8.03E-08 | Prmt2 | protein argin | Nucleus |
| ENSMUSG00000070953 | 3.51E-08 | 8.05E-08 | Rabepk | Rab9 effector | Cytoplasm |
| ENSMUSG00000021482 | 3.54E-08 | 8.12E-08 | Aaed1 | AhpC/TSA ar | Other |
| ENSMUSG00000024426 | 3.59E-08 | 8.24E-08 | Atat1 | alpha tubulin | Cytoplasm |
| ENSMUSG00000026005 | 3.59E-08 | 8.24E-08 | Rpe | ribulose-5-p | Cytoplasm |
| ENSMUSG00000015943 | 3.61E-08 | 8.27E-08 | Bola1 | bolA family n | Cytoplasm |
| ENSMUSG00000053286 | 3.65E-08 | 8.36E-08 | Trmt1l | tRNA methyl | Other |
| ENSMUSG00000046909 | 3.67E-08 | 8.41E-08 | Tefm | transcription | Cytoplasm |
| ENSMUSG00000043388 | 3.72E-08 | 8.52E-08 | Tmem130 | transmembra | Cytoplasm |
| ENSMUSG00000038215 | 3.72E-08 | 8.52E-08 | Cep44 | centrosomal | Cytoplasm |
| ENSMUSG00000074221 | 3.76E-08 | 8.60E-08 | Zfp568 | zinc finger pr | Nucleus |
| ENSMUSG00000018906 | 3.79E-08 | 8.69E-08 | P4ha2 | prolyl 4-hydr | Cytoplasm |
| ENSMUSG00000091705 | 3.84E-08 | 8.79E-08 | H2-Q2 | major histoco | Plasma Mem |
| ENSMUSG00000054715 | 3.88E-08 | 8.87E-08 | Zscan22 | zinc finger ar | Nucleus |
| ENSMUSG00000047003 | 3.88E-08 | 8.87E-08 | Zfp41 | ZFP41 zinc fi | Nucleus |
| ENSMUSG00000039682 | 3.89E-08 | 8.91E-08 | Lap3 | leucine amin | Cytoplasm |
| ENSMUSG00000028252 | 3.89E-08 | 8.91E-08 | Ccnc | cyclin C | Nucleus |
| ENSMUSG00000036775 | 3.90E-08 | 8.91E-08 | Decr2 | 2,4-dienoyl-( | Cytoplasm |
| ENSMUSG00000003464 | 3.90E-08 | 8.91E-08 | Pex19 | peroxisomal | Cytoplasm |
| ENSMUSG00000006720 | 3.90E-08 | 8.92E-08 | Zfp184 | zinc finger pr | Nucleus |
| ENSMUSG00000027667 | 3.91E-08 | 8.92E-08 | Zfp639 | zinc finger pr | Nucleus |

| | | | | | |
|---|---|---|---|---|---|
| ENSMUSG00000028243 | 4.06E-08 | 9.28E-08 | Ubxn2b | UBX domain | Cytoplasm |
| ENSMUSG00000071042 | 4.15E-08 | 9.48E-08 | Rasgrp3 | RAS guanyl r | Cytoplasm |
| ENSMUSG00000072664 | 4.20E-08 | 9.59E-08 | Ugt3a1 | UDP glycosyl | Cytoplasm |
| ENSMUSG00000020190 | 4.24E-08 | 9.67E-08 | Mknk2 | MAP kinase i | Cytoplasm |
| ENSMUSG00000036805 | 4.24E-08 | 9.67E-08 | Noxa1 | NADPH oxida | Plasma Mem |
| ENSMUSG00000041075 | 4.26E-08 | 9.72E-08 | Fzd7 | frizzled class | Plasma Mem |
| ENSMUSG00000026955 | 4.27E-08 | 9.73E-08 | Sapcd2 | suppressor A | Nucleus |
| ENSMUSG00000020366 | 4.29E-08 | 9.78E-08 | Mapk9 | mitogen-acti | Cytoplasm |
| ENSMUSG00000000826 | 4.33E-08 | 9.87E-08 | Dnajc5 | DnaJ heat sh | Plasma Mem |
| ENSMUSG00000026187 | 4.35E-08 | 9.91E-08 | Xrcc5 | X-ray repair | Nucleus |
| ENSMUSG00000031696 | 4.40E-08 | 1.00E-07 | Vps35 | VPS35, retror | Cytoplasm |
| ENSMUSG00000079184 | 4.43E-08 | 1.01E-07 | Mphosph8 | M-phase pho | Nucleus |
| ENSMUSG00000021572 | 4.46E-08 | 1.02E-07 | Cep72 | centrosomal | Cytoplasm |
| ENSMUSG00000032637 | 4.47E-08 | 1.02E-07 | Atxn2l | ataxin 2 like | Nucleus |
| ENSMUSG00000048222 | 4.47E-08 | 1.02E-07 | Mfap1b | microfibril as | Extracellular S |
| ENSMUSG00000037902 | 4.54E-08 | 1.03E-07 | Sirpa | signal regula | Plasma Mem |
| ENSMUSG00000054814 | 4.56E-08 | 1.04E-07 | Usp46 | ubiquitin spe | Other |
| ENSMUSG00000031388 | 4.60E-08 | 1.04E-07 | Naa10 | N(alpha)-ace | Nucleus |
| ENSMUSG00000022808 | 4.60E-08 | 1.05E-07 | Snx4 | sorting nexin | Cytoplasm |
| ENSMUSG00000026154 | 4.61E-08 | 1.05E-07 | Sdhaf4 | succinate de | Cytoplasm |
| ENSMUSG00000032604 | 4.61E-08 | 1.05E-07 | Qars | glutaminyl-tR | Cytoplasm |
| ENSMUSG00000026275 | 4.67E-08 | 1.06E-07 | Ppp1r7 | protein phos | Nucleus |
| ENSMUSG00000032498 | 4.82E-08 | 1.09E-07 | Mlh1 | mutL homolo | Nucleus |
| ENSMUSG00000062210 | 4.90E-08 | 1.11E-07 | Tnfaip8 | TNF alpha in | Cytoplasm |
| ENSMUSG00000048578 | 4.95E-08 | 1.12E-07 | Mlec | malectin | Plasma Mem |
| ENSMUSG00000025436 | 4.99E-08 | 1.13E-07 | Xrcc6bp1 | ATP23 metall | Plasma Mem |
| ENSMUSG00000035493 | 5.00E-08 | 1.13E-07 | Tgfbi | transforming | Extracellular S |
| ENSMUSG00000031972 | 5.02E-08 | 1.14E-07 | Acta1 | actin, alpha 1 | Cytoplasm |
| ENSMUSG00000025086 | 5.09E-08 | 1.15E-07 | Trub1 | TruB pseudo | Other |
| ENSMUSG00000031608 | 5.11E-08 | 1.16E-07 | Galnt7 | polypeptide | Cytoplasm |
| ENSMUSG00000024875 | 5.12E-08 | 1.16E-07 | Yif1a | Yip1 interacti | Cytoplasm |
| ENSMUSG00000041396 | 5.13E-08 | 1.16E-07 | Mettl18 | methyltransfe | Other |
| ENSMUSG00000037503 | 5.23E-08 | 1.18E-07 | Fam168b | family with se | Cytoplasm |
| ENSMUSG00000050410 | 5.24E-08 | 1.19E-07 | Tcf19 | transcription | Nucleus |
| ENSMUSG00000023791 | 5.28E-08 | 1.19E-07 | Pigx | phosphatidyl | Cytoplasm |
| ENSMUSG00000026843 | 5.35E-08 | 1.21E-07 | Fubp3 | far upstream | Nucleus |
| ENSMUSG00000029501 | 5.39E-08 | 1.22E-07 | Ankle2 | ankyrin repea | Nucleus |
| ENSMUSG00000042155 | 5.42E-08 | 1.23E-07 | Klhl23 | kelch like fan | Extracellular S |
| ENSMUSG00000075053 | 5.43E-08 | 1.23E-07 | Vdac3-ps1 | voltage-depe | Other |
| ENSMUSG00000028107 | 5.45E-08 | 1.23E-07 | Tars2 | threonyl-tRN | Cytoplasm |
| ENSMUSG00000037331 | 5.47E-08 | 1.24E-07 | Larp1 | La ribonucle | Cytoplasm |
| ENSMUSG00000052684 | 5.53E-08 | 1.25E-07 | Jun | Jun proto-or | Nucleus |
| ENSMUSG00000028718 | 5.57E-08 | 1.26E-07 | Stil | STIL, centriol | Nucleus |
| ENSMUSG00000038848 | 5.60E-08 | 1.26E-07 | Ythdf1 | YTH N6-met | Other |
| ENSMUSG00000061207 | 5.69E-08 | 1.28E-07 | Stk19 | serine/threor | Nucleus |
| ENSMUSG00000024787 | 5.71E-08 | 1.29E-07 | Snx15 | sorting nexin | Cytoplasm |
| ENSMUSG00000028522 | 5.75E-08 | 1.30E-07 | Mier1 | MIER1 transc | Nucleus |
| ENSMUSG00000026782 | 5.78E-08 | 1.30E-07 | Abi2 | abl interacto | Cytoplasm |
| ENSMUSG00000026078 | 5.79E-08 | 1.31E-07 | Pdcl3 | phosducin lik | Cytoplasm |
| ENSMUSG00000034881 | 5.96E-08 | 1.34E-07 | Tbxa2r | thromboxane | Plasma Mem |
| ENSMUSG00000009741 | 6.06E-08 | 1.36E-07 | Ubp1 | upstream bin | Cytoplasm |
| ENSMUSG00000029022 | 6.08E-08 | 1.37E-07 | Miip | migration an | Other |
| ENSMUSG00000034566 | 6.10E-08 | 1.37E-07 | Atp5h | ATP synthase | Cytoplasm |
| ENSMUSG00000071652 | 6.19E-08 | 1.39E-07 | Ints5 | integrator co | Nucleus |
| ENSMUSG00000073838 | 6.19E-08 | 1.39E-07 | Tufm | Tu translatior | Cytoplasm |
| ENSMUSG00000051483 | 6.25E-08 | 1.40E-07 | Cbr1 | carbonyl red | Cytoplasm |
| ENSMUSG00000028049 | 6.26E-08 | 1.41E-07 | Scamp3 | secretory car | Cytoplasm |
| ENSMUSG00000036160 | 6.27E-08 | 1.41E-07 | Surf6 | surfeit 6 | Nucleus |

| | | | | | |
|---|---|---|---|---|---|
| ENSMUSG00000061458 | 6.29E-08 | 1.41E-07 | Nol10 | nucleolar prc | Nucleus |
| ENSMUSG00000015023 | 6.34E-08 | 1.42E-07 | Ddx19a | DEAD-box h | Nucleus |
| ENSMUSG00000034259 | 6.36E-08 | 1.43E-07 | Exosc4 | exosome cor | Nucleus |
| ENSMUSG00000001247 | 6.39E-08 | 1.43E-07 | Lsr | lipolysis stim | Nucleus |
| ENSMUSG00000031903 | 6.41E-08 | 1.44E-07 | Pla2g15 | phospholipas | Cytoplasm |
| ENSMUSG00000037348 | 6.43E-08 | 1.44E-07 | Paqr7 | progestin and | Plasma Mem |
| ENSMUSG00000028016 | 6.48E-08 | 1.45E-07 | Ints12 | integrator co | Nucleus |
| ENSMUSG00000030264 | 6.53E-08 | 1.46E-07 | Thumpd3 | THUMP dom | Other |
| ENSMUSG00000038116 | 6.56E-08 | 1.47E-07 | Phf20 | PHD finger p | Nucleus |
| ENSMUSG00000021038 | 6.68E-08 | 1.50E-07 | Vipas39 | VPS33B inter | Plasma Mem |
| ENSMUSG00000009863 | 6.70E-08 | 1.50E-07 | Sdhb | succinate del | Cytoplasm |
| ENSMUSG00000033365 | 6.74E-08 | 1.51E-07 | Ipo13 | importin 13 | Nucleus |
| ENSMUSG00000003190 | 6.77E-08 | 1.52E-07 | Bcl2l12 | BCL2 like 12 | Nucleus |
| ENSMUSG00000052423 | 6.78E-08 | 1.52E-07 | B4galt3 | beta-1,4-gal | Cytoplasm |
| ENSMUSG00000014075 | 6.80E-08 | 1.52E-07 | Tctex1d2 | Tctex1 doma | Cytoplasm |
| ENSMUSG00000056310 | 6.81E-08 | 1.52E-07 | Tyw1 | tRNA-yW syr | Other |
| ENSMUSG00000024780 | 6.88E-08 | 1.54E-07 | Cdc37l1 | cell division c | Cytoplasm |
| ENSMUSG00000010290 | 6.90E-08 | 1.54E-07 | AI597479 | chromosome | Nucleus |
| ENSMUSG00000033228 | 6.99E-08 | 1.56E-07 | Scaf11 | SR-related C | Nucleus |
| ENSMUSG00000018585 | 7.01E-08 | 1.57E-07 | Atox1 | antioxidant 1 | Cytoplasm |
| ENSMUSG00000038160 | 7.06E-08 | 1.58E-07 | Atg5 | autophagy re | Cytoplasm |
| ENSMUSG00000026728 | 7.08E-08 | 1.58E-07 | Vim | vimentin | Cytoplasm |
| ENSMUSG00000028393 | 7.10E-08 | 1.58E-07 | Alad | aminolevulin | Cytoplasm |
| ENSMUSG00000009876 | 7.20E-08 | 1.61E-07 | Cox4i2 | cytochrome c | Cytoplasm |
| ENSMUSG00000041219 | 7.21E-08 | 1.61E-07 | Arhgap11a | Rho GTPase a | Cytoplasm |
| ENSMUSG00000053205 | 7.25E-08 | 1.62E-07 | Styx | serine/threor | Cytoplasm |
| ENSMUSG00000038803 | 7.27E-08 | 1.62E-07 | Ost4 | oligosacchar | Other |
| ENSMUSG00000036513 | 7.29E-08 | 1.63E-07 | Commd2 | COMM doma | Other |
| ENSMUSG00000037885 | 7.33E-08 | 1.64E-07 | Stk35 | serine/threor | Cytoplasm |
| ENSMUSG00000057400 | 7.49E-08 | 1.67E-07 | Ces1c | carboxylester | Cytoplasm |
| ENSMUSG00000032097 | 7.67E-08 | 1.71E-07 | Ddx6 | DEAD-box h | Nucleus |
| ENSMUSG00000023286 | 7.68E-08 | 1.71E-07 | Ube2j2 | ubiquitin cor | Cytoplasm |
| ENSMUSG00000041623 | 7.70E-08 | 1.72E-07 | D11Wsu47e | chromosome | Cytoplasm |
| ENSMUSG00000052299 | 7.72E-08 | 1.72E-07 | Ltn1 | listerin E3 ub | Nucleus |
| ENSMUSG00000022110 | 7.73E-08 | 1.72E-07 | Sucla2 | succinate-Co | Cytoplasm |
| ENSMUSG00000022151 | 7.74E-08 | 1.72E-07 | Ttc33 | tetratricopep | Other |
| ENSMUSG00000042148 | 7.76E-08 | 1.73E-07 | Cox10 | cytochrome c | Cytoplasm |
| ENSMUSG00000024142 | 7.82E-08 | 1.74E-07 | Mlst8 | MTOR associ | Cytoplasm |
| ENSMUSG00000034509 | 7.86E-08 | 1.75E-07 | Mad2l1bp | MAD2L1 bind | Nucleus |
| ENSMUSG00000042508 | 7.88E-08 | 1.75E-07 | Dmtf1 | cyclin D bind | Nucleus |
| ENSMUSG00000033933 | 7.92E-08 | 1.76E-07 | Vhl | von Hippel-L | Nucleus |
| ENSMUSG00000021190 | 8.05E-08 | 1.79E-07 | Lgmn | legumain | Cytoplasm |
| ENSMUSG00000052139 | 8.12E-08 | 1.80E-07 | Bre | BRISC and BI | Cytoplasm |
| ENSMUSG00000020363 | 8.12E-08 | 1.80E-07 | Gfpt2 | glutamine-fr | Cytoplasm |
| ENSMUSG00000029287 | 8.15E-08 | 1.81E-07 | Tgfbr3 | transforming | Plasma Mem |
| ENSMUSG00000041958 | 8.17E-08 | 1.81E-07 | Pigs | phosphatidyl | Cytoplasm |
| ENSMUSG00000020736 | 8.20E-08 | 1.82E-07 | Nt5c | 5', 3'-nucleot | Cytoplasm |
| ENSMUSG00000044709 | 8.28E-08 | 1.84E-07 | Gemin7 | gem nuclear | Nucleus |
| ENSMUSG00000006732 | 8.28E-08 | 1.84E-07 | Mettl1 | methyltransfe | Nucleus |
| ENSMUSG00000026889 | 8.30E-08 | 1.84E-07 | Rbm18 | RNA binding | Other |
| ENSMUSG00000035049 | 8.31E-08 | 1.84E-07 | Rrp12 | ribosomal RN | Nucleus |
| ENSMUSG00000055239 | 8.40E-08 | 1.86E-07 | Kcmf1 | potassium ch | Extracellular S |
| ENSMUSG00000038683 | 8.42E-08 | 1.87E-07 | Pak1ip1 | PAK1 interac | Nucleus |
| ENSMUSG00000025602 | 8.43E-08 | 1.87E-07 | Zfp202 | zinc finger pr | Nucleus |
| ENSMUSG00000031545 | 8.52E-08 | 1.89E-07 | Agpat6 | glycerol-3-p | Cytoplasm |
| ENSMUSG00000026497 | 8.54E-08 | 1.89E-07 | Mixl1 | Mix paired-li | Nucleus |
| ENSMUSG00000017776 | 8.57E-08 | 1.90E-07 | Crk | CRK proto-o | Cytoplasm |
| ENSMUSG00000024121 | 8.61E-08 | 1.91E-07 | Atp6v0c | ATPase H+ tr | Cytoplasm |

| | | | | | |
|---|---|---|---|---|---|
| ENSMUSG00000026014 | 8.64E-08 | 1.91E-07 | Raph1 | Ras associatio | Plasma Mem |
| ENSMUSG00000025010 | 8.71E-08 | 1.92E-07 | Ccnj | cyclin J | Nucleus |
| ENSMUSG00000033610 | 8.82E-08 | 1.95E-07 | Pank1 | pantothenate | Cytoplasm |
| ENSMUSG00000021413 | 8.83E-08 | 1.95E-07 | Prpf4b | pre-mRNA p | Nucleus |
| ENSMUSG00000074748 | 8.87E-08 | 1.96E-07 | Atxn7l3b | ataxin 7 like 3 | Cytoplasm |
| ENSMUSG00000049792 | 8.91E-08 | 1.97E-07 | Bag5 | BCL2 associa | Cytoplasm |
| ENSMUSG00000001098 | 8.93E-08 | 1.97E-07 | Kctd10 | potassium ch | Nucleus |
| ENSMUSG00000022704 | 8.95E-08 | 1.97E-07 | Qtrtd1 | queuine tRN/ | Cytoplasm |
| ENSMUSG00000019487 | 8.98E-08 | 1.98E-07 | Trip10 | thyroid horm | Cytoplasm |
| ENSMUSG00000039850 | 8.98E-08 | 1.98E-07 | Endov | endonuclease | Cytoplasm |
| ENSMUSG00000022905 | 9.02E-08 | 1.99E-07 | Kpna1 | karyopherin s | Nucleus |
| ENSMUSG00000035293 | 9.11E-08 | 2.01E-07 | G2e3 | G2/M-phase | Cytoplasm |
| ENSMUSG00000006676 | 9.18E-08 | 2.02E-07 | Usp19 | ubiquitin spe | Cytoplasm |
| ENSMUSG00000029681 | 9.22E-08 | 2.03E-07 | Bcl7b | BCL tumor su | Nucleus |
| ENSMUSG00000025068 | 9.25E-08 | 2.04E-07 | Gsto1 | glutathione S | Cytoplasm |
| ENSMUSG00000016386 | 9.32E-08 | 2.05E-07 | Mpped2 | metallophosp | Other |
| ENSMUSG00000044475 | 9.42E-08 | 2.08E-07 | Ascc1 | activating sig | Nucleus |
| ENSMUSG00000042700 | 9.47E-08 | 2.08E-07 | Sipa1l1 | signal induce | Cytoplasm |
| ENSMUSG00000035342 | 9.48E-08 | 2.09E-07 | Lzts2 | leucine zippe | Cytoplasm |
| ENSMUSG00000058407 | 9.48E-08 | 2.09E-07 | Txndc9 | thioredoxin c | Cytoplasm |
| ENSMUSG00000039069 | 9.51E-08 | 2.09E-07 | Mtg2 | mitochondria | Cytoplasm |
| ENSMUSG00000022323 | 9.63E-08 | 2.12E-07 | Hrsp12 | reactive inter | Cytoplasm |
| ENSMUSG00000021911 | 9.67E-08 | 2.13E-07 | Parg | poly(ADP-rib | Cytoplasm |
| ENSMUSG00000039275 | 9.70E-08 | 2.13E-07 | Foxk2 | forkhead box | Nucleus |
| ENSMUSG00000024319 | 9.77E-08 | 2.15E-07 | Vps52 | VPS52, GARP | Cytoplasm |
| ENSMUSG00000037400 | 9.86E-08 | 2.17E-07 | Atp11b | ATPase phos | Plasma Mem |
| ENSMUSG00000026389 | 1.02E-07 | 2.23E-07 | Steap3 | STEAP3 meta | Cytoplasm |
| ENSMUSG00000016831 | 1.02E-07 | 2.24E-07 | Tox4 | TOX high mc | Nucleus |
| ENSMUSG00000034300 | 1.02E-07 | 2.25E-07 | Fam53c | family with se | Other |
| ENSMUSG00000029466 | 1.03E-07 | 2.25E-07 | Anapc7 | anaphase pro | Nucleus |
| ENSMUSG00000064210 | 1.03E-07 | 2.26E-07 | Ano6 | anoctamin 6 | Plasma Mem |
| ENSMUSG00000042215 | 1.03E-07 | 2.26E-07 | Bag2 | BCL2 associa | Cytoplasm |
| ENSMUSG00000026356 | 1.05E-07 | 2.29E-07 | Dars | aspartyl-tRN/ | Cytoplasm |
| ENSMUSG00000078578 | 1.06E-07 | 2.31E-07 | Ube2d3 | ubiquitin con | Cytoplasm |
| ENSMUSG00000026259 | 1.07E-07 | 2.34E-07 | Ngef | neuronal gua | Cytoplasm |
| ENSMUSG00000030447 | 1.08E-07 | 2.36E-07 | Cyfip1 | cytoplasmic F | Cytoplasm |
| ENSMUSG00000021374 | 1.08E-07 | 2.38E-07 | Nup153 | nucleoporin 1 | Nucleus |
| ENSMUSG00000022807 | 1.09E-07 | 2.38E-07 | Osbpl11 | oxysterol bin | Cytoplasm |
| ENSMUSG00000025816 | 1.09E-07 | 2.39E-07 | Sec61a2 | Sec61 translo | Cytoplasm |
| ENSMUSG00000062078 | 1.10E-07 | 2.41E-07 | Qk | QKI, KH dom | Nucleus |
| ENSMUSG00000003865 | 1.12E-07 | 2.45E-07 | Gys1 | glycogen syn | Cytoplasm |
| ENSMUSG00000039648 | 1.14E-07 | 2.50E-07 | Ccbl1 | kynurenine a | Cytoplasm |
| ENSMUSG00000025626 | 1.14E-07 | 2.50E-07 | Phf6 | PHD finger p | Nucleus |
| ENSMUSG00000034190 | 1.15E-07 | 2.52E-07 | Chmp7 | charged mult | Cytoplasm |
| ENSMUSG00000028647 | 1.16E-07 | 2.54E-07 | Mycbp | MYC binding | Nucleus |
| ENSMUSG00000067873 | 1.17E-07 | 2.55E-07 | Htatsf1 | HIV-1 Tat spe | Nucleus |
| ENSMUSG00000031667 | 1.18E-07 | 2.58E-07 | Aktip | AKT interacti | Cytoplasm |
| ENSMUSG00000031785 | 1.19E-07 | 2.60E-07 | Adgrg1 | adhesion G p | Plasma Mem |
| ENSMUSG00000059540 | 1.20E-07 | 2.62E-07 | Tcea2 | transcription | Nucleus |
| ENSMUSG00000057594 | 1.20E-07 | 2.63E-07 | Arl16 | ADP ribosyla | Cytoplasm |
| ENSMUSG00000014243 | 1.21E-07 | 2.63E-07 | Zswim7 | zinc finger SV | Other |
| ENSMUSG00000032217 | 1.22E-07 | 2.67E-07 | Rnf111 | ring finger pr | Nucleus |
| ENSMUSG00000026784 | 1.23E-07 | 2.67E-07 | Pdss1 | decaprenyl d | Cytoplasm |
| ENSMUSG00000027835 | 1.24E-07 | 2.70E-07 | Pdcd10 | programmed | Cytoplasm |
| ENSMUSG00000038668 | 1.25E-07 | 2.71E-07 | Lpar1 | lysophosphat | Plasma Mem |
| ENSMUSG00000024985 | 1.25E-07 | 2.73E-07 | Tcf7l2 | transcription | Nucleus |
| ENSMUSG00000021493 | 1.28E-07 | 2.79E-07 | Pdlim7 | PDZ and LIM | Cytoplasm |
| ENSMUSG00000074358 | 1.28E-07 | 2.79E-07 | Ccdc61 | coiled-coil de | Cytoplasm |

| | | | | | |
|---|---|---|---|---|---|
| ENSMUSG00000028572 | 1.30E-07 | 2.83E-07 | Hook1 | hook microtu | Cytoplasm |
| ENSMUSG00000004945 | 1.31E-07 | 2.86E-07 | Tmem242 | transmembra | Other |
| ENSMUSG00000014301 | 1.31E-07 | 2.86E-07 | Pam16 | presequence | Cytoplasm |
| ENSMUSG00000038827 | 1.33E-07 | 2.88E-07 | Fam206a | family with se | Extracellular S |
| ENSMUSG00000008999 | 1.33E-07 | 2.89E-07 | Bmp7 | bone morpho | Extracellular S |
| ENSMUSG00000015994 | 1.35E-07 | 2.93E-07 | Fnta | farnesyltransf | Cytoplasm |
| ENSMUSG00000042569 | 1.38E-07 | 2.99E-07 | Dhrs7b | dehydrogena | Other |
| ENSMUSG00000057421 | 1.38E-07 | 3.00E-07 | Las1l | LAS1 like, rib | Nucleus |
| ENSMUSG00000033379 | 1.38E-07 | 3.01E-07 | Atp6v0b | ATPase H+ tr | Cytoplasm |
| ENSMUSG00000034667 | 1.41E-07 | 3.07E-07 | Xpot | exportin for t | Nucleus |
| ENSMUSG00000064181 | 1.42E-07 | 3.08E-07 | Rab3ip | RAB3A intera | Cytoplasm |
| ENSMUSG00000025823 | 1.43E-07 | 3.10E-07 | Pdia4 | protein disulf | Cytoplasm |
| ENSMUSG00000020946 | 1.43E-07 | 3.10E-07 | Gosr2 | golgi SNAP r | Cytoplasm |
| ENSMUSG00000030869 | 1.43E-07 | 3.11E-07 | Ndufab1 | NADH:ubiqu | Cytoplasm |
| ENSMUSG00000032555 | 1.43E-07 | 3.11E-07 | Topbp1 | DNA topoiso | Nucleus |
| ENSMUSG00000026634 | 1.44E-07 | 3.13E-07 | Angel2 | angel homolo | Nucleus |
| ENSMUSG00000027605 | 1.44E-07 | 3.13E-07 | Acss2 | acyl-CoA syn | Cytoplasm |
| ENSMUSG00000025373 | 1.47E-07 | 3.19E-07 | Rnf41 | ring finger pr | Cytoplasm |
| ENSMUSG00000029577 | 1.47E-07 | 3.19E-07 | Ube3b | ubiquitin pro | Extracellular S |
| ENSMUSG00000028309 | 1.49E-07 | 3.22E-07 | Rnf20 | ring finger pr | Nucleus |
| ENSMUSG00000018012 | 1.54E-07 | 3.33E-07 | Rac3 | Rac family sm | Cytoplasm |
| ENSMUSG00000029469 | 1.55E-07 | 3.36E-07 | Ift81 | intraflagellar | Extracellular S |
| ENSMUSG00000040532 | 1.55E-07 | 3.36E-07 | Abhd11 | abhydrolase | Cytoplasm |
| ENSMUSG00000025873 | 1.58E-07 | 3.41E-07 | Faf2 | Fas associate | Cytoplasm |
| ENSMUSG00000026342 | 1.60E-07 | 3.47E-07 | Slc35f5 | solute carrier | Other |
| ENSMUSG00000056666 | 1.61E-07 | 3.48E-07 | Retsat | retinol satura | Cytoplasm |
| ENSMUSG00000022000 | 1.62E-07 | 3.49E-07 | Zc3h13 | zinc finger CC | Extracellular S |
| ENSMUSG00000025134 | 1.62E-07 | 3.51E-07 | Alyref | Aly/REF expo | Nucleus |
| ENSMUSG00000038816 | 1.63E-07 | 3.51E-07 | Ctnnal1 | catenin alpha | Plasma Mem |
| ENSMUSG00000057572 | 1.63E-07 | 3.52E-07 | Zbtb8os | zinc finger ar | Cytoplasm |
| ENSMUSG00000021945 | 1.64E-07 | 3.54E-07 | Zmym2 | zinc finger M | Nucleus |
| ENSMUSG00000020409 | 1.65E-07 | 3.55E-07 | Slu7 | SLU7 homolo | Nucleus |
| ENSMUSG00000063808 | 1.66E-07 | 3.58E-07 | Gpatch1 | G-patch dom | Extracellular S |
| ENSMUSG00000014426 | 1.67E-07 | 3.61E-07 | Map3k4 | mitogen-acti | Cytoplasm |
| ENSMUSG00000074867 | 1.68E-07 | 3.61E-07 | Zfp808 | zinc finger pr | Other |
| ENSMUSG00000027935 | 1.68E-07 | 3.62E-07 | Rab13 | RAB13, mem | Plasma Mem |
| ENSMUSG00000027655 | 1.69E-07 | 3.64E-07 | Dhx35 | DEAH-box he | Other |
| ENSMUSG00000031827 | 1.69E-07 | 3.65E-07 | Cotl1 | coactosin like | Cytoplasm |
| ENSMUSG00000049670 | 1.73E-07 | 3.73E-07 | Morn4 | MORN repea | Extracellular S |
| ENSMUSG00000029276 | 1.73E-07 | 3.73E-07 | Glmn | glomulin, FKB | Cytoplasm |
| ENSMUSG00000028179 | 1.73E-07 | 3.73E-07 | Cth | cystathionine | Cytoplasm |
| ENSMUSG00000021555 | 1.74E-07 | 3.75E-07 | Naa35 | N(alpha)-ace | Cytoplasm |
| ENSMUSG00000024038 | 1.75E-07 | 3.77E-07 | Ndufv3 | NADH:ubiqu | Cytoplasm |
| ENSMUSG00000032897 | 1.76E-07 | 3.80E-07 | Nfyc | nuclear trans | Nucleus |
| ENSMUSG00000031016 | 1.77E-07 | 3.80E-07 | Wee1 | WEE1 G2 che | Nucleus |
| ENSMUSG00000020572 | 1.77E-07 | 3.82E-07 | Nampt | nicotinamide | Extracellular S |
| ENSMUSG00000053219 | 1.78E-07 | 3.82E-07 | Raet1e | retinoic acid | Plasma Mem |
| ENSMUSG00000028961 | 1.79E-07 | 3.84E-07 | Pgd | phosphogluc | Cytoplasm |
| ENSMUSG00000064128 | 1.82E-07 | 3.92E-07 | Cenpj | centromere p | Nucleus |
| ENSMUSG00000031613 | 1.82E-07 | 3.92E-07 | Hpgd | 15-hydroxyp | Cytoplasm |
| ENSMUSG00000042447 | 1.84E-07 | 3.94E-07 | Mios | meiosis regul | Cytoplasm |
| ENSMUSG00000038544 | 1.85E-07 | 3.97E-07 | Inip | INTS3 and N | Nucleus |
| ENSMUSG00000034329 | 1.86E-07 | 3.99E-07 | Brip1 | BRCA1 intera | Nucleus |
| ENSMUSG00000039740 | 1.87E-07 | 4.01E-07 | Alg2 | ALG2, alpha- | Cytoplasm |
| ENSMUSG00000069045 | 1.87E-07 | 4.02E-07 | Ddx3y | DEAD-box he | Cytoplasm |
| ENSMUSG00000033728 | 1.88E-07 | 4.03E-07 | Lrrc14 | leucine rich r | Cytoplasm |
| ENSMUSG00000028549 | 1.89E-07 | 4.05E-07 | Itgb3bp | integrin subu | Nucleus |
| ENSMUSG00000041483 | 1.91E-07 | 4.08E-07 | Zfp281 | zinc finger pr | Nucleus |

| | | | | | |
|---|---|---|---|---|---|
| ENSMUSG00000063015 | 1.93E-07 | 4.14E-07 | Ccni | cyclin I | Other |
| ENSMUSG00000038633 | 1.95E-07 | 4.18E-07 | Degs1 | delta 4-desat | Plasma Mem |
| ENSMUSG00000060227 | 1.96E-07 | 4.20E-07 | Casc4 | cancer susce | Cytoplasm |
| ENSMUSG00000046753 | 2.00E-07 | 4.27E-07 | Ccdc66 | coiled-coil d | Cytoplasm |
| ENSMUSG00000030304 | 2.01E-07 | 4.31E-07 | Ergic2 | ERGIC and g | Cytoplasm |
| ENSMUSG00000085793 | 2.03E-07 | 4.34E-07 | Lin52 | lin-52 DREAM | Nucleus |
| ENSMUSG00000016382 | 2.04E-07 | 4.37E-07 | Pls3 | plastin 3 | Cytoplasm |
| ENSMUSG00000053317 | 2.05E-07 | 4.39E-07 | Sec61b | Sec61 translo | Cytoplasm |
| ENSMUSG00000039396 | 2.05E-07 | 4.39E-07 | Neil3 | nei like DNA | Nucleus |
| ENSMUSG00000034311 | 2.06E-07 | 4.41E-07 | Kif4 | kinesin family | Nucleus |
| ENSMUSG00000025137 | 2.06E-07 | 4.41E-07 | Pcyt2 | phosphate cy | Cytoplasm |
| ENSMUSG00000021112 | 2.09E-07 | 4.46E-07 | Mpp5 | membrane p | Plasma Mem |
| ENSMUSG00000037315 | 2.10E-07 | 4.48E-07 | Jade3 | jade family P | Nucleus |
| ENSMUSG00000026977 | 2.12E-07 | 4.52E-07 | Marc7 | membrane as | Extracellular S |
| ENSMUSG00000019461 | 2.15E-07 | 4.59E-07 | Plscr3 | phospholipid | Plasma Mem |
| ENSMUSG00000063550 | 2.16E-07 | 4.61E-07 | Nup98 | nucleoporin 9 | Nucleus |
| ENSMUSG00000017868 | 2.17E-07 | 4.64E-07 | Sgk2 | SGK2, serine/ | Cytoplasm |
| ENSMUSG00000021501 | 2.20E-07 | 4.70E-07 | Caml | calcium mod | Cytoplasm |
| ENSMUSG00000042389 | 2.21E-07 | 4.71E-07 | Tsen2 | tRNA splicing | Cytoplasm |
| ENSMUSG00000035770 | 2.25E-07 | 4.80E-07 | Dync1li2 | dynein cytop | Cytoplasm |
| ENSMUSG00000001569 | 2.25E-07 | 4.80E-07 | Nom1 | nucleolar pro | Nucleus |
| ENSMUSG00000033356 | 2.27E-07 | 4.83E-07 | Pus7l | pseudouridyl | Other |
| ENSMUSG00000051716 | 2.27E-07 | 4.84E-07 | Apon | apolipoprote | Other |
| ENSMUSG00000038880 | 2.28E-07 | 4.85E-07 | Mrps34 | mitochondria | Cytoplasm |
| ENSMUSG00000036430 | 2.29E-07 | 4.88E-07 | Tbcc | tubulin foldin | Cytoplasm |
| ENSMUSG00000028799 | 2.31E-07 | 4.91E-07 | Zfp362 | zinc finger pr | Other |
| ENSMUSG00000020868 | 2.33E-07 | 4.96E-07 | Xylt2 | xylosyltransfe | Cytoplasm |
| ENSMUSG00000040843 | 2.34E-07 | 4.98E-07 | Tiprl | TOR signaling | Cytoplasm |
| ENSMUSG00000037772 | 2.35E-07 | 5.01E-07 | Mrpl23 | mitochondria | Cytoplasm |
| ENSMUSG00000024140 | 2.35E-07 | 5.01E-07 | Epas1 | endothelial P | Nucleus |
| ENSMUSG00000022020 | 2.36E-07 | 5.01E-07 | Naa16 | N(alpha)-ace | Nucleus |
| ENSMUSG00000028423 | 2.36E-07 | 5.01E-07 | Nfx1 | nuclear trans | Nucleus |
| ENSMUSG00000010607 | 2.37E-07 | 5.03E-07 | Pigyl | phosphatidyl | Plasma Mem |
| ENSMUSG00000021969 | 2.37E-07 | 5.04E-07 | Zdhhc20 | zinc finger D | Plasma Mem |
| ENSMUSG00000021594 | 2.38E-07 | 5.05E-07 | Srd5a1 | steroid 5 alph | Cytoplasm |
| ENSMUSG00000036940 | 2.40E-07 | 5.09E-07 | Kdm1a | lysine demeth | Nucleus |
| ENSMUSG00000023092 | 2.40E-07 | 5.10E-07 | Fhl1 | four and a ha | Cytoplasm |
| ENSMUSG00000025979 | 2.41E-07 | 5.12E-07 | Mob4 | MOB family r | Cytoplasm |
| ENSMUSG00000042195 | 2.43E-07 | 5.15E-07 | Slc35f2 | solute carrier | Other |
| ENSMUSG00000005621 | 2.43E-07 | 5.15E-07 | Zfp592 | zinc finger pr | Nucleus |
| ENSMUSG00000031066 | 2.43E-07 | 5.16E-07 | Usp11 | ubiquitin spe | Nucleus |
| ENSMUSG00000020592 | 2.44E-07 | 5.17E-07 | Sdc1 | syndecan 1 | Plasma Mem |
| ENSMUSG00000029613 | 2.45E-07 | 5.19E-07 | Eif2ak1 | eukaryotic tra | Cytoplasm |
| ENSMUSG00000020523 | 2.46E-07 | 5.21E-07 | Fam114a2 | family with se | Other |
| ENSMUSG00000025203 | 2.47E-07 | 5.24E-07 | Scd2 | stearoyl-Coe | Cytoplasm |
| ENSMUSG00000048285 | 2.48E-07 | 5.25E-07 | Frmd6 | FERM domain | Cytoplasm |
| ENSMUSG00000034998 | 2.48E-07 | 5.25E-07 | Foxn2 | forkhead box | Nucleus |
| ENSMUSG00000029070 | 2.50E-07 | 5.28E-07 | Mxra8 | matrix remod | Cytoplasm |
| ENSMUSG00000092035 | 2.51E-07 | 5.30E-07 | Peg10 | paternally ex | Nucleus |
| ENSMUSG00000018412 | 2.52E-07 | 5.32E-07 | Kansl1 | KAT8 regulat | Nucleus |
| ENSMUSG00000002625 | 2.52E-07 | 5.34E-07 | Akap8l | A-kinase anc | Nucleus |
| ENSMUSG00000037826 | 2.56E-07 | 5.41E-07 | Ppm1k | protein phos | Cytoplasm |
| ENSMUSG00000089824 | 2.56E-07 | 5.42E-07 | Rbm12 | RNA binding | Nucleus |
| ENSMUSG00000030559 | 2.64E-07 | 5.58E-07 | Rab38 | RAB38, mem | Cytoplasm |
| ENSMUSG00000020212 | 2.66E-07 | 5.62E-07 | Mdm1 | Mdm1 nuclea | Cytoplasm |
| ENSMUSG00000048217 | 2.66E-07 | 5.62E-07 | Nags | N-acetylgluta | Cytoplasm |
| ENSMUSG00000035086 | 2.66E-07 | 5.63E-07 | Becn1 | beclin 1 | Cytoplasm |
| ENSMUSG00000020718 | 2.67E-07 | 5.64E-07 | Polg2 | DNA polyme | Cytoplasm |

| | | | | | |
|---|---|---|---|---|---|
| ENSMUSG00000026094 | 2.71E-07 | 5.73E-07 | Stk17b | serine/threor | Nucleus |
| ENSMUSG00000071655 | 2.73E-07 | 5.75E-07 | Ubxn1 | UBX domain | Cytoplasm |
| ENSMUSG00000049672 | 2.75E-07 | 5.80E-07 | Zbtb14 | zinc finger ar | Nucleus |
| ENSMUSG00000029480 | 2.76E-07 | 5.81E-07 | Dhx37 | DEAH-box h | Nucleus |
| ENSMUSG00000071650 | 2.77E-07 | 5.84E-07 | Ganab | glucosidase I | Cytoplasm |
| ENSMUSG00000030704 | 2.79E-07 | 5.88E-07 | Rab6a | RAB6A, mem | Cytoplasm |
| ENSMUSG00000038784 | 2.80E-07 | 5.91E-07 | Cnot4 | CCR4-NOT t | Cytoplasm |
| ENSMUSG00000021792 | 2.81E-07 | 5.92E-07 | Fam213a | family with se | Extracellular S |
| ENSMUSG00000027962 | 2.87E-07 | 6.04E-07 | Vcam1 | vascular cell a | Plasma Mem |
| ENSMUSG00000006519 | 2.87E-07 | 6.04E-07 | Cyba | cytochrome b | Cytoplasm |
| ENSMUSG00000055633 | 2.92E-07 | 6.15E-07 | Zfp580 | zinc finger pr | Nucleus |
| ENSMUSG00000021890 | 2.93E-07 | 6.17E-07 | Eaf1 | ELL associate | Nucleus |
| ENSMUSG00000024926 | 2.93E-07 | 6.17E-07 | Kat5 | lysine acetylt | Nucleus |
| ENSMUSG00000028032 | 2.96E-07 | 6.23E-07 | Papss1 | 3'-phosphoa | Cytoplasm |
| ENSMUSG00000033423 | 3.02E-07 | 6.35E-07 | Eri3 | ERI1 exoribo | Plasma Mem |
| ENSMUSG00000044367 | 3.06E-07 | 6.43E-07 | Slc16a13 | solute carrier | Cytoplasm |
| ENSMUSG00000071648 | 3.07E-07 | 6.46E-07 | Rom1 | retinal outer | Plasma Mem |
| ENSMUSG00000026849 | 3.08E-07 | 6.47E-07 | Tor1a | torsin family | Cytoplasm |
| ENSMUSG00000067931 | 3.09E-07 | 6.49E-07 | Zfp948 | zinc finger pr | Other |
| ENSMUSG00000017858 | 3.11E-07 | 6.53E-07 | Ift52 | intraflagellar | Cytoplasm |
| ENSMUSG00000008393 | 3.11E-07 | 6.53E-07 | Carhsp1 | calcium regu | Cytoplasm |
| ENSMUSG00000029202 | 3.12E-07 | 6.55E-07 | Pds5a | PDS5 cohesir | Nucleus |
| ENSMUSG00000039048 | 3.13E-07 | 6.57E-07 | Foxred1 | FAD depende | Cytoplasm |
| ENSMUSG00000021806 | 3.17E-07 | 6.64E-07 | Nid2 | nidogen 2 | Extracellular S |
| ENSMUSG00000009291 | 3.17E-07 | 6.66E-07 | Pttg1ip | PTTG1 intera | Nucleus |
| ENSMUSG00000026276 | 3.18E-07 | 6.66E-07 | Sept2 | septin 2 | Cytoplasm |
| ENSMUSG00000037243 | 3.22E-07 | 6.74E-07 | Zfp692 | zinc finger pr | Nucleus |
| ENSMUSG00000044600 | 3.24E-07 | 6.78E-07 | Smim7 | small integra | Other |
| ENSMUSG00000015668 | 3.26E-07 | 6.82E-07 | Pdzd11 | PDZ domain | Extracellular S |
| ENSMUSG00000002900 | 3.27E-07 | 6.85E-07 | Lamb1 | laminin subu | Extracellular S |
| ENSMUSG00000009646 | 3.29E-07 | 6.88E-07 | Pla2g12b | phospholipas | Extracellular S |
| ENSMUSG00000044134 | 3.29E-07 | 6.89E-07 | Fam109a | PH domain c | Cytoplasm |
| ENSMUSG00000005698 | 3.31E-07 | 6.93E-07 | Ctcf | CCCTC-bindi | Nucleus |
| ENSMUSG00000030846 | 3.35E-07 | 7.00E-07 | Tial1 | TIA1 cytotoxi | Nucleus |
| ENSMUSG00000035031 | 3.36E-07 | 7.03E-07 | C8a | complement | Extracellular S |
| ENSMUSG00000028671 | 3.39E-07 | 7.10E-07 | Gale | UDP-galacto | Cytoplasm |
| ENSMUSG00000029621 | 3.40E-07 | 7.10E-07 | Arpc1a | actin related | Extracellular S |
| ENSMUSG00000039953 | 3.40E-07 | 7.11E-07 | Clstn1 | calsyntenin 1 | Plasma Mem |
| ENSMUSG00000003198 | 3.43E-07 | 7.18E-07 | Zfp959 | zinc finger pr | Other |
| ENSMUSG00000044934 | 3.47E-07 | 7.25E-07 | Zfp367 | zinc finger pr | Nucleus |
| ENSMUSG00000027598 | 3.47E-07 | 7.26E-07 | Itch | itchy E3 ubiq | Nucleus |
| ENSMUSG00000078676 | 3.48E-07 | 7.26E-07 | Casc3 | CASC3, exon | Nucleus |
| ENSMUSG00000026743 | 3.52E-07 | 7.34E-07 | Mllt10 | MLLT10, hist | Nucleus |
| ENSMUSG00000034729 | 3.54E-07 | 7.39E-07 | Mrps10 | mitochondria | Cytoplasm |
| ENSMUSG00000029402 | 3.54E-07 | 7.39E-07 | Snrnp35 | small nuclear | Nucleus |
| ENSMUSG00000020534 | 3.56E-07 | 7.43E-07 | Shmt1 | serine hydro | Cytoplasm |
| ENSMUSG00000028435 | 3.60E-07 | 7.52E-07 | Aqp3 | aquaporin 3 | Plasma Mem |
| ENSMUSG00000019917 | 3.63E-07 | 7.57E-07 | Spet10 | septin 10 | Cytoplasm |
| ENSMUSG00000037316 | 3.72E-07 | 7.76E-07 | Bag4 | BCL2 associa | Cytoplasm |
| ENSMUSG00000057363 | 3.74E-07 | 7.79E-07 | Uxs1 | UDP-glucurc | Cytoplasm |
| ENSMUSG00000061650 | 3.76E-07 | 7.84E-07 | Med9 | mediator cor | Nucleus |
| ENSMUSG00000023020 | 3.77E-07 | 7.85E-07 | Cox14 | cytochrome c | Cytoplasm |
| ENSMUSG00000029059 | 3.81E-07 | 7.93E-07 | Fam213b | family with se | Cytoplasm |
| ENSMUSG00000021051 | 3.81E-07 | 7.94E-07 | Ppp2r5e | protein phos | Cytoplasm |
| ENSMUSG00000033352 | 3.82E-07 | 7.95E-07 | Map2k4 | mitogen-acti | Cytoplasm |
| ENSMUSG00000028248 | 3.83E-07 | 7.96E-07 | Pnisr | PNN interact | Nucleus |
| ENSMUSG00000030335 | 3.83E-07 | 7.96E-07 | Mrpl51 | mitochondria | Cytoplasm |
| ENSMUSG00000019970 | 3.85E-07 | 8.00E-07 | Sgk1 | serum/gluco | Cytoplasm |

| | | | | | |
|---|---|---|---|---|---|
| ENSMUSG00000043140 | 3.87E-07 | 8.04E-07 | Tmem186 | transmembra | Cytoplasm |
| ENSMUSG00000037905 | 3.89E-07 | 8.08E-07 | Bri3bp | BRI3 binding | Extracellular S |
| ENSMUSG00000022472 | 3.95E-07 | 8.21E-07 | Desi1 | desumoylatir | Other |
| ENSMUSG00000007603 | 3.99E-07 | 8.28E-07 | Dus3l | dihydrouridir | Other |
| ENSMUSG00000015776 | 4.04E-07 | 8.39E-07 | Med22 | mediator cor | Cytoplasm |
| ENSMUSG00000014313 | 4.11E-07 | 8.52E-07 | Cox6c | cytochrome c | Cytoplasm |
| ENSMUSG00000025423 | 4.11E-07 | 8.52E-07 | Pias2 | protein inhib | Nucleus |
| ENSMUSG00000025151 | 4.14E-07 | 8.59E-07 | Maged1 | MAGE family | Plasma Mem |
| ENSMUSG00000090266 | 4.15E-07 | 8.60E-07 | Mettl23 | methyltransfe | Cytoplasm |
| ENSMUSG00000036504 | 4.18E-07 | 8.66E-07 | Phpt1 | phosphohisti | Cytoplasm |
| ENSMUSG00000026150 | 4.18E-07 | 8.67E-07 | Mff | mitochondria | Cytoplasm |
| ENSMUSG00000079677 | 4.19E-07 | 8.68E-07 | Fdx1l | ferredoxin 2 | Cytoplasm |
| ENSMUSG00000019808 | 4.20E-07 | 8.71E-07 | Adat2 | adenosine de | Nucleus |
| ENSMUSG00000019715 | 4.24E-07 | 8.78E-07 | Gle1 | GLE1, RNA e: | Nucleus |
| ENSMUSG00000079036 | 4.30E-07 | 8.90E-07 | Alkbh1 | alkB homolog | Cytoplasm |
| ENSMUSG00000066324 | 4.37E-07 | 9.06E-07 | Impad1 | inositol mon | Cytoplasm |
| ENSMUSG00000022757 | 4.38E-07 | 9.08E-07 | Tfg | TRK-fused ge | Cytoplasm |
| ENSMUSG00000025962 | 4.40E-07 | 9.10E-07 | Fastkd2 | FAST kinase c | Cytoplasm |
| ENSMUSG00000033417 | 4.40E-07 | 9.11E-07 | Cacul1 | CDK2 associa | Other |
| ENSMUSG00000020921 | 4.51E-07 | 9.33E-07 | Tmem101 | transmembra | Extracellular S |
| ENSMUSG00000070304 | 4.54E-07 | 9.39E-07 | Scn2b | sodium volta | Plasma Mem |
| ENSMUSG00000027243 | 4.54E-07 | 9.39E-07 | Harbi1 | harbinger tra | Cytoplasm |
| ENSMUSG00000026766 | 4.55E-07 | 9.40E-07 | Mmadhc | methylmalon | Cytoplasm |
| ENSMUSG00000026035 | 4.56E-07 | 9.42E-07 | Ppil3 | peptidylproly | Cytoplasm |
| ENSMUSG00000001054 | 4.62E-07 | 9.55E-07 | Rmnd5b | required for r | Other |
| ENSMUSG00000020954 | 4.70E-07 | 9.72E-07 | Strn3 | striatin 3 | Nucleus |
| ENSMUSG00000035614 | 4.73E-07 | 9.76E-07 | Fam179b | TOG array re | Extracellular S |
| ENSMUSG00000060224 | 4.79E-07 | 9.89E-07 | Pyroxd2 | pyridine nucl | Other |
| ENSMUSG00000030850 | 4.83E-07 | 9.97E-07 | Ate1 | arginyltransfe | Nucleus |
| ENSMUSG00000020260 | 4.84E-07 | 9.99E-07 | Pofut2 | protein O-fu | Cytoplasm |
| ENSMUSG00000025421 | 4.88E-07 | 1.01E-06 | Hdhd2 | haloacid deh | Cytoplasm |
| ENSMUSG00000030214 | 4.90E-07 | 1.01E-06 | Plbd1 | phospholipas | Extracellular S |
| ENSMUSG00000028430 | 4.96E-07 | 1.02E-06 | Nol6 | nucleolar pro | Nucleus |
| ENSMUSG00000021418 | 5.00E-07 | 1.03E-06 | Rpp40 | ribonuclease | Nucleus |
| ENSMUSG00000002416 | 5.05E-07 | 1.04E-06 | Ndufb2 | NADH:ubiqu | Cytoplasm |
| ENSMUSG00000024479 | 5.12E-07 | 1.06E-06 | Mal2 | mal, T cell dif | Plasma Mem |
| ENSMUSG00000019864 | 5.24E-07 | 1.08E-06 | Rtn4ip1 | reticulon 4 in | Cytoplasm |
| ENSMUSG00000041957 | 5.28E-07 | 1.09E-06 | Pkp2 | plakophilin 2 | Plasma Mem |
| ENSMUSG00000032309 | 5.29E-07 | 1.09E-06 | Fbxo22 | F-box protei | Cytoplasm |
| ENSMUSG00000061273 | 5.33E-07 | 1.10E-06 | Mmgt1 | membrane m | Cytoplasm |
| ENSMUSG00000032872 | 5.45E-07 | 1.12E-06 | Cyb5r4 | cytochrome b | Cytoplasm |
| ENSMUSG00000025144 | 5.50E-07 | 1.13E-06 | Stra13 | centromere p | Nucleus |
| ENSMUSG00000029710 | 5.51E-07 | 1.13E-06 | Ephb4 | EPH receptor | Plasma Mem |
| ENSMUSG00000035048 | 5.63E-07 | 1.16E-06 | Anapc13 | anaphase pro | Nucleus |
| ENSMUSG00000030870 | 5.65E-07 | 1.16E-06 | Ubfd1 | ubiquitin fam | Other |
| ENSMUSG00000035824 | 5.68E-07 | 1.17E-06 | Tk2 | thymidine kir | Cytoplasm |
| ENSMUSG00000021481 | 5.70E-07 | 1.17E-06 | Zfp346 | zinc finger pr | Nucleus |
| ENSMUSG00000024944 | 5.79E-07 | 1.19E-06 | Arl2 | ADP ribosyla | Cytoplasm |
| ENSMUSG00000023045 | 5.82E-07 | 1.20E-06 | Soat2 | sterol O-acyl | Cytoplasm |
| ENSMUSG00000026827 | 5.91E-07 | 1.21E-06 | Gpd2 | glycerol-3-p | Cytoplasm |
| ENSMUSG00000056978 | 5.95E-07 | 1.22E-06 | Hamp2 | hepcidin anti | Extracellular S |
| ENSMUSG00000044949 | 5.96E-07 | 1.22E-06 | Ubtd2 | ubiquitin dor | Other |
| ENSMUSG00000002343 | 5.96E-07 | 1.22E-06 | Armc6 | armadillo rep | Cytoplasm |
| ENSMUSG00000024855 | 6.02E-07 | 1.24E-06 | Pacs1 | phosphofurir | Cytoplasm |
| ENSMUSG00000037072 | 6.03E-07 | 1.24E-06 | Spe15 | selenoprotei | Cytoplasm |
| ENSMUSG00000030672 | 6.12E-07 | 1.26E-06 | Mylpf | myosin light | Cytoplasm |
| ENSMUSG00000046722 | 6.15E-07 | 1.26E-06 | Cdc42se1 | CDC42 small | Plasma Mem |
| ENSMUSG00000002949 | 6.20E-07 | 1.27E-06 | Timm44 | translocase o | Cytoplasm |

| | | | | | |
|---|---|---|---|---|---|
| ENSMUSG00000038736 | 6.21E-07 | 1.27E-06 | Nudcd1 | NudC domai | Nucleus |
| ENSMUSG00000031023 | 6.31E-07 | 1.29E-06 | Akip1 | A-kinase inte | Nucleus |
| ENSMUSG00000027495 | 6.34E-07 | 1.30E-06 | Fam210b | family with se | Extracellular S |
| ENSMUSG00000069208 | 6.37E-07 | 1.30E-06 | Zfp825 | expressed se | Other |
| ENSMUSG00000075486 | 6.46E-07 | 1.32E-06 | Commd6 | COMM doma | Other |
| ENSMUSG00000037820 | 6.48E-07 | 1.33E-06 | Tgm2 | transglutamir | Cytoplasm |
| ENSMUSG00000026171 | 6.49E-07 | 1.33E-06 | Rnf25 | ring finger pr | Nucleus |
| ENSMUSG00000021918 | 6.53E-07 | 1.34E-06 | Nek4 | NIMA relatec | Nucleus |
| ENSMUSG00000015971 | 6.58E-07 | 1.34E-06 | Actr8 | ARP8 actin re | Nucleus |
| ENSMUSG00000036306 | 6.62E-07 | 1.35E-06 | Lzts1 | leucine zippe | Nucleus |
| ENSMUSG00000046841 | 6.77E-07 | 1.38E-06 | Ckap4 | cytoskeleton | Cytoplasm |
| ENSMUSG00000022180 | 6.80E-07 | 1.39E-06 | Slc7a8 | solute carrier | Plasma Mem |
| ENSMUSG00000061353 | 6.82E-07 | 1.39E-06 | Cxcl12 | C-X-C motif | Extracellular S |
| ENSMUSG00000018678 | 6.89E-07 | 1.41E-06 | Sp2 | Sp2 transcrip | Nucleus |
| ENSMUSG00000018537 | 7.12E-07 | 1.45E-06 | Pcgf2 | polycomb gr | Nucleus |
| ENSMUSG00000019977 | 7.17E-07 | 1.46E-06 | Hbs1l | HBS1 like trai | Cytoplasm |
| ENSMUSG00000031849 | 7.17E-07 | 1.46E-06 | Comp | cartilage olig | Extracellular S |
| ENSMUSG00000017286 | 7.18E-07 | 1.46E-06 | Glod4 | glyoxalase dc | Cytoplasm |
| ENSMUSG00000032435 | 7.29E-07 | 1.49E-06 | Dync1li1 | dynein cytop | Cytoplasm |
| ENSMUSG00000024507 | 7.30E-07 | 1.49E-06 | Hsd17b4 | hydroxystero | Cytoplasm |
| ENSMUSG00000050846 | 7.37E-07 | 1.50E-06 | Zfp623 | zinc finger pr | Nucleus |
| ENSMUSG00000020744 | 7.41E-07 | 1.51E-06 | Slc25a19 | solute carrier | Cytoplasm |
| ENSMUSG00000049957 | 7.46E-07 | 1.52E-06 | Ccdc137 | coiled-coil d | Nucleus |
| ENSMUSG00000022206 | 7.47E-07 | 1.52E-06 | Npr3 | natriuretic pe | Plasma Mem |
| ENSMUSG00000045665 | 7.59E-07 | 1.55E-06 | Mfsd5 | major facilita | Other |
| ENSMUSG00000020257 | 7.61E-07 | 1.55E-06 | Wdr82 | WD repeat d | Nucleus |
| ENSMUSG00000063870 | 7.67E-07 | 1.56E-06 | Chd4 | chromodoma | Nucleus |
| ENSMUSG00000034121 | 7.81E-07 | 1.59E-06 | Mks1 | Meckel syndr | Cytoplasm |
| ENSMUSG00000074457 | 7.83E-07 | 1.59E-06 | S100a16 | S100 calcium | Nucleus |
| ENSMUSG00000029993 | 7.92E-07 | 1.61E-06 | Nfu1 | NFU1 iron-su | Cytoplasm |
| ENSMUSG00000027478 | 7.92E-07 | 1.61E-06 | Dnmt3b | DNA methylt | Nucleus |
| ENSMUSG00000050107 | 8.03E-07 | 1.63E-06 | Gsg2 | histone H3 as | Nucleus |
| ENSMUSG00000030685 | 8.08E-07 | 1.64E-06 | Kctd13 | potassium ch | Nucleus |
| ENSMUSG00000031631 | 8.08E-07 | 1.64E-06 | Cfap97 | cilia and flag | Other |
| ENSMUSG00000045502 | 8.26E-07 | 1.68E-06 | Hcar2 | hydroxycarbo | Plasma Mem |
| ENSMUSG00000048058 | 8.43E-07 | 1.71E-06 | Ldlrad3 | low density li | Plasma Mem |
| ENSMUSG00000026260 | 8.46E-07 | 1.72E-06 | Ndufa10 | NADH:ubiqu | Cytoplasm |
| ENSMUSG00000002741 | 8.52E-07 | 1.73E-06 | Ykt6 | YKT6 v-SNAF | Cytoplasm |
| ENSMUSG00000032702 | 8.55E-07 | 1.74E-06 | Kank1 | KN motif anc | Nucleus |
| ENSMUSG00000015291 | 8.57E-07 | 1.74E-06 | Gdi1 | GDP dissocia | Cytoplasm |
| ENSMUSG00000048351 | 8.65E-07 | 1.76E-06 | Coa7 | cytochrome c | Other |
| ENSMUSG00000059355 | 8.68E-07 | 1.76E-06 | Wdr83os | WD repeat d | Other |
| ENSMUSG00000024170 | 8.68E-07 | 1.76E-06 | Telo2 | telomere mai | Cytoplasm |
| ENSMUSG00000041801 | 8.70E-07 | 1.76E-06 | Phlda3 | pleckstrin ho | Plasma Mem |
| ENSMUSG00000017478 | 8.81E-07 | 1.78E-06 | Zc3h18 | zinc finger C | Nucleus |
| ENSMUSG00000037373 | 8.83E-07 | 1.79E-06 | Ctbp1 | C-terminal b | Nucleus |
| ENSMUSG00000063410 | 8.89E-07 | 1.80E-06 | Stk24 | serine/threor | Cytoplasm |
| ENSMUSG00000048728 | 8.90E-07 | 1.80E-06 | Zfp454 | zinc finger pr | Nucleus |
| ENSMUSG00000022906 | 8.91E-07 | 1.80E-06 | Parp9 | poly(ADP-rib | Nucleus |
| ENSMUSG00000009378 | 8.93E-07 | 1.81E-06 | Slc16a12 | solute carrier | Plasma Mem |
| ENSMUSG00000020956 | 8.97E-07 | 1.81E-06 | Dtd2 | D-tyrosyl-tRI | Other |
| ENSMUSG00000022753 | 8.99E-07 | 1.82E-06 | Tmem30c | transmembra | Other |
| ENSMUSG00000013593 | 9.09E-07 | 1.84E-06 | Ndufs2 | NADH:ubiqu | Cytoplasm |
| ENSMUSG00000006273 | 9.10E-07 | 1.84E-06 | Atp6v1b2 | ATPase H+ tr | Cytoplasm |
| ENSMUSG00000037152 | 9.12E-07 | 1.84E-06 | Ndufc1 | NADH:ubiqu | Cytoplasm |
| ENSMUSG00000022779 | 9.18E-07 | 1.85E-06 | Top3b | DNA topoiso | Nucleus |
| ENSMUSG00000028223 | 9.18E-07 | 1.86E-06 | Decr1 | 2,4-dienoyl-( | Cytoplasm |
| ENSMUSG00000063171 | 9.21E-07 | 1.86E-06 | Rps4l | ribosomal pr | Cytoplasm |

| | | | | | |
|---|---|---|---|---|---|
| ENSMUSG00000024150 | 9.34E-07 | 1.89E-06 | Mcfd2 | multiple coag | Cytoplasm |
| ENSMUSG00000034442 | 9.36E-07 | 1.89E-06 | Trmt5 | tRNA methyl | Cytoplasm |
| ENSMUSG00000026687 | 9.42E-07 | 1.90E-06 | Aldh9a1 | aldehyde deh | Cytoplasm |
| ENSMUSG00000029413 | 9.44E-07 | 1.90E-06 | Naaa | N-acylethano | Cytoplasm |
| ENSMUSG00000041997 | 9.53E-07 | 1.92E-06 | Tlk1 | tousled like k | Nucleus |
| ENSMUSG00000015289 | 9.67E-07 | 1.95E-06 | Lage3 | L antigen fan | Nucleus |
| ENSMUSG00000021114 | 9.70E-07 | 1.96E-06 | Atp6v1d | ATPase H+ tr | Cytoplasm |
| ENSMUSG00000026584 | 9.78E-07 | 1.97E-06 | Scyl3 | SCY1 like pse | Cytoplasm |
| ENSMUSG00000025574 | 9.82E-07 | 1.98E-06 | Tk1 | thymidine kir | Cytoplasm |
| ENSMUSG00000049038 | 9.91E-07 | 2.00E-06 | Mterf2 | mitochondria | Cytoplasm |
| ENSMUSG00000049482 | 9.99E-07 | 2.01E-06 | Ctu2 | cytosolic thic | Cytoplasm |
| ENSMUSG00000058192 | 1.01E-06 | 2.04E-06 | Zfp846 | zinc finger pr | Other |
| ENSMUSG00000046671 | 1.02E-06 | 2.05E-06 | Mtfr1l | mitochondria | Other |
| ENSMUSG00000025155 | 1.03E-06 | 2.08E-06 | Dus1l | dihydrouridir | Other |
| ENSMUSG00000057193 | 1.05E-06 | 2.11E-06 | Slc44a2 | solute carrier | Extracellular S |
| ENSMUSG00000049504 | 1.05E-06 | 2.12E-06 | Proser1 | proline and s | Other |
| ENSMUSG00000036916 | 1.07E-06 | 2.14E-06 | Zfp280c | zinc finger pr | Nucleus |
| ENSMUSG00000028933 | 1.07E-06 | 2.15E-06 | Xrcc2 | X-ray repair c | Nucleus |
| ENSMUSG00000040521 | 1.07E-06 | 2.16E-06 | Tsfm | Ts translatior | Cytoplasm |
| ENSMUSG00000053390 | 1.12E-06 | 2.25E-06 | Zfp952 | zinc finger pr | Other |
| ENSMUSG00000030551 | 1.13E-06 | 2.27E-06 | Nr2f2 | nuclear recep | Nucleus |
| ENSMUSG00000040722 | 1.13E-06 | 2.27E-06 | Scamp5 | secretory car | Cytoplasm |
| ENSMUSG00000073643 | 1.15E-06 | 2.30E-06 | Wdfy1 | WD repeat ar | Cytoplasm |
| ENSMUSG00000032766 | 1.15E-06 | 2.31E-06 | Gng11 | G protein sub | Plasma Mem |
| ENSMUSG00000048495 | 1.16E-06 | 2.33E-06 | Tyw5 | tRNA-yW syr | Cytoplasm |
| ENSMUSG00000050914 | 1.18E-06 | 2.37E-06 | Ankrd37 | ankyrin repea | Cytoplasm |
| ENSMUSG00000029056 | 1.20E-06 | 2.41E-06 | Pank4 | pantothenate | Cytoplasm |
| ENSMUSG00000032092 | 1.21E-06 | 2.42E-06 | Mpzl2 | myelin protei | Plasma Mem |
| ENSMUSG00000074886 | 1.21E-06 | 2.43E-06 | Grk6 | G protein-co | Plasma Mem |
| ENSMUSG00000026708 | 1.21E-06 | 2.43E-06 | Cenpl | centromere p | Cytoplasm |
| ENSMUSG00000041926 | 1.21E-06 | 2.43E-06 | Rnpep | arginyl aminc | Cytoplasm |
| ENSMUSG00000045854 | 1.22E-06 | 2.44E-06 | Lyrm2 | LYR motif co | Cytoplasm |
| ENSMUSG00000032816 | 1.23E-06 | 2.46E-06 | Igdcc4 | immunoglob | Plasma Mem |
| ENSMUSG00000057229 | 1.23E-06 | 2.46E-06 | Atp5sl | distal membr | Cytoplasm |
| ENSMUSG00000084957 | 1.23E-06 | 2.47E-06 | Bbip1 | BBSome inte | Cytoplasm |
| ENSMUSG00000009145 | 1.26E-06 | 2.52E-06 | Dqx1 | DEAQ-box R | Other |
| ENSMUSG00000027425 | 1.27E-06 | 2.53E-06 | Csrp2bp | lysine acetylt | Nucleus |
| ENSMUSG00000070427 | 1.27E-06 | 2.55E-06 | Il18bp | interleukin 18 | Extracellular S |
| ENSMUSG00000026088 | 1.27E-06 | 2.55E-06 | Mitd1 | microtubule | Cytoplasm |
| ENSMUSG00000042737 | 1.28E-06 | 2.56E-06 | Dpm3 | dolichyl-pho | Cytoplasm |
| ENSMUSG00000046862 | 1.29E-06 | 2.57E-06 | Pramef8 | PRAME famil | Other |
| ENSMUSG00000027381 | 1.29E-06 | 2.57E-06 | Bcl2l11 | BCL2 like 11 | Cytoplasm |
| ENSMUSG00000020770 | 1.32E-06 | 2.64E-06 | Unk | unkempt fam | Nucleus |
| ENSMUSG00000072115 | 1.32E-06 | 2.64E-06 | Ang | angiogenin | Extracellular S |
| ENSMUSG00000052214 | 1.32E-06 | 2.64E-06 | Opa3 | OPA3, outer | Cytoplasm |
| ENSMUSG00000040374 | 1.32E-06 | 2.64E-06 | Pex2 | peroxisomal | Cytoplasm |
| ENSMUSG00000081984 | 1.33E-06 | 2.65E-06 | Dnajb3 | DnaJ heat sh | Cytoplasm |
| ENSMUSG00000025317 | 1.34E-06 | 2.67E-06 | Car5a | carbonic anh | Cytoplasm |
| ENSMUSG00000029076 | 1.34E-06 | 2.68E-06 | Sdf4 | stromal cell c | Cytoplasm |
| ENSMUSG00000066042 | 1.35E-06 | 2.69E-06 | Med18 | mediator cor | Nucleus |
| ENSMUSG00000059772 | 1.35E-06 | 2.69E-06 | Slx1b | SLX1 homolc | Nucleus |
| ENSMUSG00000018774 | 1.35E-06 | 2.70E-06 | Cd68 | CD68 molecu | Plasma Mem |
| ENSMUSG00000026608 | 1.37E-06 | 2.72E-06 | Kctd3 | potassium ch | Plasma Mem |
| ENSMUSG00000038250 | 1.37E-06 | 2.73E-06 | Usp38 | ubiquitin spe | Other |
| ENSMUSG00000027569 | 1.39E-06 | 2.76E-06 | Mrgbp | MRG domain | Nucleus |
| ENSMUSG00000073988 | 1.40E-06 | 2.78E-06 | Ttpa | alpha tocoph | Cytoplasm |
| ENSMUSG00000016833 | 1.41E-06 | 2.80E-06 | Mrps18c | mitochondria | Cytoplasm |
| ENSMUSG00000059447 | 1.42E-06 | 2.83E-06 | Hadhb | hydroxyacyl- | Cytoplasm |

| | | | | | |
|---|---|---|---|---|---|
| ENSMUSG00000069808 | 1.44E-06 | 2.86E-06 | Fam57a | family with se | Plasma Mem |
| ENSMUSG00000072949 | 1.44E-06 | 2.87E-06 | Acot1 | acyl-CoA thio | Cytoplasm |
| ENSMUSG00000040006 | 1.44E-06 | 2.87E-06 | Ginm1 | glycoprotein | Cytoplasm |
| ENSMUSG00000021209 | 1.45E-06 | 2.87E-06 | Ppp4r4 | protein phos | Cytoplasm |
| ENSMUSG00000033752 | 1.45E-06 | 2.88E-06 | Mnd1 | meiotic nucle | Nucleus |
| ENSMUSG00000034157 | 1.45E-06 | 2.89E-06 | Cipc | CLOCK intera | Nucleus |
| ENSMUSG00000018425 | 1.46E-06 | 2.90E-06 | Dhx40 | DEAH-box h | Other |
| ENSMUSG00000038876 | 1.46E-06 | 2.91E-06 | Rnf146 | ring finger pr | Cytoplasm |
| ENSMUSG00000030322 | 1.47E-06 | 2.91E-06 | Mbd4 | methyl-CpG | Nucleus |
| ENSMUSG00000022561 | 1.48E-06 | 2.94E-06 | Gpaa1 | glycosylphos | Cytoplasm |
| ENSMUSG00000031262 | 1.48E-06 | 2.94E-06 | Cenpi | centromere p | Nucleus |
| ENSMUSG00000001134 | 1.49E-06 | 2.95E-06 | Uxt | ubiquitously | Cytoplasm |
| ENSMUSG00000036810 | 1.49E-06 | 2.95E-06 | Cnep1r1 | CTD nuclear | Nucleus |
| ENSMUSG00000052446 | 1.49E-06 | 2.95E-06 | Zfp961 | zinc finger pr | Other |
| ENSMUSG00000053536 | 1.50E-06 | 2.97E-06 | Cstf2t | cleavage stin | Nucleus |
| ENSMUSG00000023072 | 1.55E-06 | 3.08E-06 | Cep89 | centrosomal | Cytoplasm |
| ENSMUSG00000035933 | 1.57E-06 | 3.10E-06 | Cog5 | component c | Cytoplasm |
| ENSMUSG00000038650 | 1.57E-06 | 3.10E-06 | Rnh1 | ribonuclease, | Cytoplasm |
| ENSMUSG00000038127 | 1.58E-06 | 3.14E-06 | Ccdc50 | coiled-coil d | Cytoplasm |
| ENSMUSG00000006611 | 1.60E-06 | 3.16E-06 | Hfe | homeostatic | Plasma Mem |
| ENSMUSG00000028607 | 1.60E-06 | 3.17E-06 | Cpt2 | carnitine palr | Cytoplasm |
| ENSMUSG00000066278 | 1.60E-06 | 3.17E-06 | Vps37b | VPS37B, ESCI | Cytoplasm |
| ENSMUSG00000005354 | 1.61E-06 | 3.18E-06 | Txn2 | thioredoxin 2 | Cytoplasm |
| ENSMUSG00000029392 | 1.61E-06 | 3.18E-06 | Rilpl1 | Rab interactir | Cytoplasm |
| ENSMUSG00000064284 | 1.65E-06 | 3.26E-06 | Cdpf1 | cysteine rich | Extracellular S |
| ENSMUSG00000000959 | 1.65E-06 | 3.27E-06 | Oxa1l | OXA1L, mito | Cytoplasm |
| ENSMUSG00000001052 | 1.67E-06 | 3.30E-06 | Sec24b | SEC24 homo | Cytoplasm |
| ENSMUSG00000047492 | 1.68E-06 | 3.32E-06 | Inhbe | inhibin subur | Extracellular S |
| ENSMUSG00000010914 | 1.69E-06 | 3.34E-06 | Pdhx | pyruvate deh | Cytoplasm |
| ENSMUSG00000044700 | 1.72E-06 | 3.40E-06 | Tmem201 | transmembra | Nucleus |
| ENSMUSG00000066829 | 1.72E-06 | 3.40E-06 | Zfp810 | zinc finger pr | Other |
| ENSMUSG00000027635 | 1.73E-06 | 3.42E-06 | Dsn1 | DSN1 homol | Nucleus |
| ENSMUSG00000050270 | 1.74E-06 | 3.44E-06 | Tmem220 | transmembra | Other |
| ENSMUSG00000033319 | 1.75E-06 | 3.45E-06 | Fem1c | fem-1 homo | Nucleus |
| ENSMUSG00000032382 | 1.76E-06 | 3.48E-06 | Snx1 | sorting nexin | Cytoplasm |
| ENSMUSG00000042831 | 1.77E-06 | 3.50E-06 | Alkbh6 | alkB homolo | Nucleus |
| ENSMUSG00000019948 | 1.78E-06 | 3.51E-06 | Actr6 | ARP6 actin re | Nucleus |
| ENSMUSG00000024758 | 1.79E-06 | 3.53E-06 | Rtn3 | reticulon 3 | Cytoplasm |
| ENSMUSG00000038967 | 1.79E-06 | 3.54E-06 | Pdk2 | pyruvate deh | Cytoplasm |
| ENSMUSG00000032737 | 1.81E-06 | 3.56E-06 | Inppl1 | inositol polyp | Cytoplasm |
| ENSMUSG00000029185 | 1.81E-06 | 3.57E-06 | Fam114a1 | family with se | Extracellular S |
| ENSMUSG00000051306 | 1.83E-06 | 3.61E-06 | Usp42 | ubiquitin spe | Nucleus |
| ENSMUSG00000052566 | 1.85E-06 | 3.64E-06 | Hook2 | hook microtu | Cytoplasm |
| ENSMUSG00000023942 | 1.87E-06 | 3.68E-06 | Slc29a1 | solute carrier | Plasma Mem |
| ENSMUSG00000073565 | 1.88E-06 | 3.70E-06 | Prr16 | proline rich 1 | Other |
| ENSMUSG00000032477 | 1.90E-06 | 3.74E-06 | Cdc25a | cell division c | Nucleus |
| ENSMUSG00000028680 | 1.91E-06 | 3.77E-06 | Plk3 | polo like kina | Nucleus |
| ENSMUSG00000044674 | 1.93E-06 | 3.80E-06 | Fzd1 | frizzled class | Plasma Mem |
| ENSMUSG00000028710 | 1.94E-06 | 3.82E-06 | Atpaf1 | ATP synthase | Cytoplasm |
| ENSMUSG00000072825 | 2.00E-06 | 3.93E-06 | Cep170b | centrosomal | Other |
| ENSMUSG00000018995 | 2.01E-06 | 3.95E-06 | Nars2 | asparaginyl-t | Cytoplasm |
| ENSMUSG00000078798 | 2.02E-06 | 3.97E-06 | Sult2a1 | sulfotransfera | Cytoplasm |
| ENSMUSG00000022422 | 2.04E-06 | 4.00E-06 | Dscc1 | DNA replicat | Nucleus |
| ENSMUSG00000022442 | 2.06E-06 | 4.04E-06 | Ttll1 | tubulin tyrosi | Extracellular S |
| ENSMUSG00000040188 | 2.07E-06 | 4.07E-06 | Scamp2 | secretory car | Cytoplasm |
| ENSMUSG00000036748 | 2.08E-06 | 4.08E-06 | Cuedc2 | CUE domain | Extracellular S |
| ENSMUSG00000034187 | 2.11E-06 | 4.13E-06 | Nsf | N-ethylmalei | Cytoplasm |
| ENSMUSG00000029427 | 2.15E-06 | 4.22E-06 | Zcchc8 | zinc finger CC | Nucleus |

| | | | | | |
|---|---|---|---|---|---|
| ENSMUSG00000024182 | 2.17E-06 | 4.25E-06 | Axin1 | axin 1 | Cytoplasm |
| ENSMUSG00000027167 | 2.18E-06 | 4.27E-06 | Elp4 | elongator ac | Cytoplasm |
| ENSMUSG00000030793 | 2.18E-06 | 4.27E-06 | Pycard | PYD and CAF | Cytoplasm |
| ENSMUSG00000003345 | 2.20E-06 | 4.30E-06 | Csnk1g2 | casein kinase | Cytoplasm |
| ENSMUSG00000035944 | 2.21E-06 | 4.33E-06 | Ttc38 | tetratricopep | Cytoplasm |
| ENSMUSG00000029199 | 2.23E-06 | 4.36E-06 | Lias | lipoic acid sy | Cytoplasm |
| ENSMUSG00000030291 | 2.23E-06 | 4.36E-06 | Med21 | mediator cor | Nucleus |
| ENSMUSG00000019528 | 2.23E-06 | 4.36E-06 | Gyg | glycogenin 1 | Cytoplasm |
| ENSMUSG00000018339 | 2.24E-06 | 4.38E-06 | Gpx3 | glutathione p | Extracellular S |
| ENSMUSG00000025326 | 2.25E-06 | 4.40E-06 | Ube3a | ubiquitin pro | Nucleus |
| ENSMUSG00000033845 | 2.26E-06 | 4.42E-06 | Mrpl15 | mitochondria | Cytoplasm |
| ENSMUSG00000061244 | 2.26E-06 | 4.43E-06 | Exoc5 | exocyst comp | Cytoplasm |
| ENSMUSG00000063796 | 2.29E-06 | 4.47E-06 | Slc22a8 | solute carrier | Plasma Mem |
| ENSMUSG00000034175 | 2.29E-06 | 4.48E-06 | Rhbdd3 | rhomboid dc | Other |
| ENSMUSG00000024382 | 2.30E-06 | 4.49E-06 | Ercc3 | ERCC excisio | Nucleus |
| ENSMUSG00000029345 | 2.31E-06 | 4.52E-06 | Tfip11 | tuftelin intera | Extracellular S |
| ENSMUSG00000024491 | 2.32E-06 | 4.53E-06 | Rbm27 | RNA binding | Nucleus |
| ENSMUSG00000018672 | 2.32E-06 | 4.54E-06 | Copz2 | coatomer pro | Cytoplasm |
| ENSMUSG00000031604 | 2.32E-06 | 4.54E-06 | Msmo1 | methylsterol | Cytoplasm |
| ENSMUSG00000030822 | 2.33E-06 | 4.55E-06 | Prr14 | proline rich 1 | Other |
| ENSMUSG00000026170 | 2.33E-06 | 4.55E-06 | Cyp27a1 | cytochrome I | Cytoplasm |
| ENSMUSG00000031976 | 2.35E-06 | 4.58E-06 | Urb2 | URB2 ribosor | Nucleus |
| ENSMUSG00000022516 | 2.37E-06 | 4.63E-06 | Nudt16l1 | nudix hydrola | Cytoplasm |
| ENSMUSG00000041936 | 2.38E-06 | 4.64E-06 | Agrn | agrin | Plasma Mem |
| ENSMUSG00000020412 | 2.40E-06 | 4.68E-06 | Ascc2 | activating sig | Nucleus |
| ENSMUSG00000025353 | 2.41E-06 | 4.69E-06 | Ormdl2 | ORMDL sphi | Cytoplasm |
| ENSMUSG00000026803 | 2.49E-06 | 4.84E-06 | Ttf1 | transcription | Nucleus |
| ENSMUSG00000059436 | 2.50E-06 | 4.86E-06 | Max | MYC associat | Nucleus |
| ENSMUSG00000000782 | 2.50E-06 | 4.86E-06 | Tcf7 | transcription | Nucleus |
| ENSMUSG00000042312 | 2.51E-06 | 4.88E-06 | S100a13 | S100 calcium | Cytoplasm |
| ENSMUSG00000052144 | 2.53E-06 | 4.93E-06 | Ppp4r2 | protein phos | Cytoplasm |
| ENSMUSG00000028629 | 2.54E-06 | 4.93E-06 | Exo5 | exonuclease | Nucleus |
| ENSMUSG00000028899 | 2.56E-06 | 4.97E-06 | Taf12 | TATA-box bi | Nucleus |
| ENSMUSG00000032185 | 2.57E-06 | 4.99E-06 | Carm1 | coactivator a | Nucleus |
| ENSMUSG00000002814 | 2.58E-06 | 5.02E-06 | Top3a | DNA topoiso | Nucleus |
| ENSMUSG00000034908 | 2.59E-06 | 5.04E-06 | Sidt2 | SID1 transme | Cytoplasm |
| ENSMUSG00000045973 | 2.60E-06 | 5.05E-06 | Slc25a51 | solute carrier | Cytoplasm |
| ENSMUSG00000055835 | 2.61E-06 | 5.07E-06 | Zfp1 | ZFP1 zinc fin | Nucleus |
| ENSMUSG00000029407 | 2.62E-06 | 5.09E-06 | Uso1 | USO1 vesicle | Cytoplasm |
| ENSMUSG00000027217 | 2.64E-06 | 5.13E-06 | Tspan18 | tetraspanin 1 | Other |
| ENSMUSG00000001948 | 2.67E-06 | 5.18E-06 | Spa17 | sperm autoa | Plasma Mem |
| ENSMUSG00000041215 | 2.69E-06 | 5.22E-06 | Yeats2 | YEATS doma | Nucleus |
| ENSMUSG00000031885 | 2.70E-06 | 5.24E-06 | Cbfb | core-binding | Nucleus |
| ENSMUSG00000031671 | 2.71E-06 | 5.26E-06 | Setd6 | SET domain c | Nucleus |
| ENSMUSG00000031740 | 2.73E-06 | 5.30E-06 | Mmp2 | matrix metall | Extracellular S |
| ENSMUSG00000029290 | 2.74E-06 | 5.32E-06 | Zfp326 | zinc finger pr | Nucleus |
| ENSMUSG00000035027 | 2.76E-06 | 5.36E-06 | Map2k2 | mitogen-acti | Cytoplasm |
| ENSMUSG00000030269 | 2.78E-06 | 5.39E-06 | Mtmr14 | myotubularir | Cytoplasm |
| ENSMUSG00000039804 | 2.80E-06 | 5.43E-06 | Ncoa5 | nuclear recep | Nucleus |
| ENSMUSG00000025008 | 2.81E-06 | 5.44E-06 | Tctn3 | tectonic fami | Cytoplasm |
| ENSMUSG00000029439 | 2.81E-06 | 5.45E-06 | Sfswap | splicing facto | Nucleus |
| ENSMUSG00000020107 | 2.85E-06 | 5.52E-06 | Anapc16 | anaphase pro | Nucleus |
| ENSMUSG00000036078 | 2.86E-06 | 5.53E-06 | Sigmar1 | sigma non-o | Plasma Mem |
| ENSMUSG00000010608 | 2.88E-06 | 5.57E-06 | Rbm25 | RNA binding | Nucleus |
| ENSMUSG00000030177 | 2.89E-06 | 5.58E-06 | Ccdc77 | coiled-coil d | Cytoplasm |
| ENSMUSG00000060038 | 2.89E-06 | 5.59E-06 | Dhps | deoxyhypusi | Cytoplasm |
| ENSMUSG00000014846 | 2.89E-06 | 5.59E-06 | Tppp3 | tubulin polyn | Cytoplasm |
| ENSMUSG00000084786 | 2.91E-06 | 5.63E-06 | Ubl5 | ubiquitin like | Cytoplasm |

| | | | | | |
|---|---|---|---|---|---|
| ENSMUSG00000057068 | 2.92E-06 | 5.63E-06 | Fam47e | family with se | Cytoplasm |
| ENSMUSG00000047824 | 2.94E-06 | 5.67E-06 | Pygo2 | pygopus fam | Nucleus |
| ENSMUSG00000028687 | 2.94E-06 | 5.68E-06 | Mutyh | mutY DNA gl | Nucleus |
| ENSMUSG00000073791 | 2.95E-06 | 5.68E-06 | Efcab7 | EF-hand calc | Extracellular S |
| ENSMUSG00000037868 | 2.96E-06 | 5.71E-06 | Egr2 | early growth | Nucleus |
| ENSMUSG00000025724 | 2.99E-06 | 5.76E-06 | Sec11a | SEC11 homo | Cytoplasm |
| ENSMUSG00000033068 | 2.99E-06 | 5.77E-06 | Entpd6 | ectonucleosi | Cytoplasm |
| ENSMUSG00000027952 | 3.04E-06 | 5.86E-06 | Pmvk | phosphomev | Cytoplasm |
| ENSMUSG00000028943 | 3.06E-06 | 5.89E-06 | Espn | espin | Cytoplasm |
| ENSMUSG00000029913 | 3.08E-06 | 5.93E-06 | Prdm5 | PR/SET doma | Nucleus |
| ENSMUSG00000002043 | 3.08E-06 | 5.93E-06 | Trappc6a | trafficking pr | Cytoplasm |
| ENSMUSG00000028879 | 3.11E-06 | 5.99E-06 | Stx12 | syntaxin 12 | Cytoplasm |
| ENSMUSG00000032218 | 3.21E-06 | 6.18E-06 | Ccnb2 | cyclin B2 | Cytoplasm |
| ENSMUSG00000043013 | 3.21E-06 | 6.19E-06 | Onecut1 | one cut hom | Nucleus |
| ENSMUSG00000063229 | 3.24E-06 | 6.23E-06 | Ldha | lactate dehyc | Cytoplasm |
| ENSMUSG00000072714 | 3.27E-06 | 6.29E-06 | Rpl21-ps4 | ribosomal pr | Other |
| ENSMUSG00000014554 | 3.27E-06 | 6.30E-06 | Dguok | deoxyguanos | Cytoplasm |
| ENSMUSG00000006471 | 3.29E-06 | 6.32E-06 | Ndor1 | NADPH depe | Cytoplasm |
| ENSMUSG00000019734 | 3.29E-06 | 6.32E-06 | Tmc4 | transmembra | Cytoplasm |
| ENSMUSG00000026456 | 3.37E-06 | 6.49E-06 | Cyb5r1 | cytochrome b | Cytoplasm |
| ENSMUSG00000029369 | 3.38E-06 | 6.50E-06 | Afm | afamin | Extracellular S |
| ENSMUSG00000026201 | 3.44E-06 | 6.60E-06 | Stk16 | serine/threor | Cytoplasm |
| ENSMUSG00000022521 | 3.52E-06 | 6.77E-06 | Crebbp | CREB binding | Nucleus |
| ENSMUSG00000061461 | 3.54E-06 | 6.80E-06 | Smim20 | small integra | Cytoplasm |
| ENSMUSG00000022437 | 3.54E-06 | 6.81E-06 | Samm50 | SAMM50 sor | Cytoplasm |
| ENSMUSG00000000759 | 3.65E-06 | 7.01E-06 | Tubgcp3 | tubulin gamr | Cytoplasm |
| ENSMUSG00000062931 | 3.67E-06 | 7.05E-06 | Zfp938 | zinc finger pr | Other |
| ENSMUSG00000075304 | 3.67E-06 | 7.05E-06 | Sp5 | Sp5 transcrip | Nucleus |
| ENSMUSG00000004530 | 3.68E-06 | 7.07E-06 | Coro1c | coronin 1C | Cytoplasm |
| ENSMUSG00000024393 | 3.70E-06 | 7.10E-06 | Prrc2a | proline rich c | Cytoplasm |
| ENSMUSG00000053730 | 3.75E-06 | 7.19E-06 | Tmem39b | transmembra | Cytoplasm |
| ENSMUSG00000021520 | 3.75E-06 | 7.20E-06 | Uqcrb | ubiquinol-cy | Cytoplasm |
| ENSMUSG00000008090 | 3.77E-06 | 7.22E-06 | Fgfrl1 | fibroblast gro | Plasma Mem |
| ENSMUSG00000042548 | 3.77E-06 | 7.23E-06 | Asxl1 | ASXL transcri | Nucleus |
| ENSMUSG00000048772 | 3.78E-06 | 7.24E-06 | Tmem53 | transmembra | Nucleus |
| ENSMUSG00000031513 | 3.78E-06 | 7.25E-06 | Leprotl1 | leptin recept | Plasma Mem |
| ENSMUSG00000030595 | 3.82E-06 | 7.31E-06 | Nfkbib | NFKB inhibito | Nucleus |
| ENSMUSG00000024158 | 3.83E-06 | 7.34E-06 | Hagh | hydroxyacylg | Cytoplasm |
| ENSMUSG00000022863 | 3.85E-06 | 7.36E-06 | Btg3 | BTG anti-pro | Nucleus |
| ENSMUSG00000036446 | 3.87E-06 | 7.39E-06 | Lum | lumican | Extracellular S |
| ENSMUSG00000061950 | 3.87E-06 | 7.39E-06 | Ppp4r1 | protein phos | Other |
| ENSMUSG00000022842 | 3.88E-06 | 7.42E-06 | Ece2 | endothelin co | Plasma Mem |
| ENSMUSG00000033392 | 4.00E-06 | 7.64E-06 | Clasp2 | cytoplasmic l | Cytoplasm |
| ENSMUSG00000027893 | 4.01E-06 | 7.66E-06 | Ahcyl1 | adenosylhom | Cytoplasm |
| ENSMUSG00000014232 | 4.07E-06 | 7.78E-06 | Cluap1 | clusterin asso | Nucleus |
| ENSMUSG00000020267 | 4.08E-06 | 7.79E-06 | Hint1 | histidine triac | Nucleus |
| ENSMUSG00000021913 | 4.08E-06 | 7.79E-06 | Ogdhl | oxoglutarate | Other |
| ENSMUSG00000020653 | 4.10E-06 | 7.84E-06 | Klf11 | Kruppel like f | Nucleus |
| ENSMUSG00000021518 | 4.15E-06 | 7.92E-06 | Ptdss1 | phosphatidyl | Cytoplasm |
| ENSMUSG00000039089 | 4.17E-06 | 7.95E-06 | L3mbtl3 | L3MBTL3, his | Nucleus |
| ENSMUSG00000033773 | 4.18E-06 | 7.97E-06 | Rpap2 | RNA polymer | Nucleus |
| ENSMUSG00000042423 | 4.21E-06 | 8.03E-06 | Fbrs | fibrosin | Extracellular S |
| ENSMUSG00000027712 | 4.22E-06 | 8.04E-06 | Anxa5 | annexin A5 | Plasma Mem |
| ENSMUSG00000009907 | 4.23E-06 | 8.05E-06 | Vps4b | vacuolar prot | Cytoplasm |
| ENSMUSG00000025147 | 4.24E-06 | 8.07E-06 | Mob2 | MOB kinase a | Nucleus |
| ENSMUSG00000027122 | 4.24E-06 | 8.08E-06 | Arl14ep | ADP ribosyla | Other |
| ENSMUSG00000034210 | 4.28E-06 | 8.16E-06 | Efcab14 | EF-hand calc | Other |
| ENSMUSG00000020280 | 4.32E-06 | 8.23E-06 | Pus10 | pseudouridyl | Other |

| | | | | | |
|---|---|---|---|---|---|
| ENSMUSG00000031099 | 4.33E-06 | 8.24E-06 | Smarca1 | SWI/SNF rela | Nucleus |
| ENSMUSG00000028567 | 4.45E-06 | 8.47E-06 | Txndc12 | thioredoxin c | Cytoplasm |
| ENSMUSG00000029647 | 4.46E-06 | 8.48E-06 | Pan3 | poly(A) speci | Cytoplasm |
| ENSMUSG00000027006 | 4.53E-06 | 8.61E-06 | Dnajc10 | DnaJ heat sh | Cytoplasm |
| ENSMUSG00000027582 | 4.54E-06 | 8.63E-06 | Zgpat | zinc finger C | Nucleus |
| ENSMUSG00000025037 | 4.57E-06 | 8.70E-06 | Maoa | monoamine | Cytoplasm |
| ENSMUSG00000018481 | 4.61E-06 | 8.76E-06 | Appbp2 | amyloid beta | Cytoplasm |
| ENSMUSG00000023027 | 4.65E-06 | 8.84E-06 | Atf1 | activating tra | Nucleus |
| ENSMUSG00000092558 | 4.66E-06 | 8.85E-06 | Med20 | mediator cor | Nucleus |
| ENSMUSG00000007815 | 4.67E-06 | 8.87E-06 | Rhoa | ras homolog | Cytoplasm |
| ENSMUSG00000032643 | 4.69E-06 | 8.91E-06 | Fhl3 | four and a ha | Plasma Mem |
| ENSMUSG00000079003 | 4.69E-06 | 8.91E-06 | Samd1 | sterile alpha | Other |
| ENSMUSG00000028232 | 4.70E-06 | 8.91E-06 | Tmem68 | transmembra | Other |
| ENSMUSG00000043510 | 4.75E-06 | 9.01E-06 | Hscb | HscB mitoch | Cytoplasm |
| ENSMUSG00000029249 | 4.76E-06 | 9.03E-06 | Rest | RE1 silencing | Nucleus |
| ENSMUSG00000028962 | 4.76E-06 | 9.03E-06 | Slc4a2 | solute carrier | Plasma Mem |
| ENSMUSG00000039191 | 4.81E-06 | 9.12E-06 | Rbpj | recombinatic | Nucleus |
| ENSMUSG00000035505 | 4.92E-06 | 9.32E-06 | Cox18 | cytochrome c | Cytoplasm |
| ENSMUSG00000029029 | 4.93E-06 | 9.34E-06 | Wrap73 | WD repeat c | Nucleus |
| ENSMUSG00000026575 | 4.99E-06 | 9.45E-06 | Nme7 | NME/NM23 f | Cytoplasm |
| ENSMUSG00000060992 | 5.02E-06 | 9.51E-06 | Copz1 | coatomer pro | Cytoplasm |
| ENSMUSG00000026585 | 5.03E-06 | 9.52E-06 | Kifap3 | kinesin assoc | Cytoplasm |
| ENSMUSG00000031818 | 5.06E-06 | 9.58E-06 | Cox4i1 | cytochrome c | Cytoplasm |
| ENSMUSG00000030538 | 5.08E-06 | 9.62E-06 | Cib1 | calcium and i | Nucleus |
| ENSMUSG00000027472 | 5.09E-06 | 9.63E-06 | Pdrg1 | p53 and DNA | Cytoplasm |
| ENSMUSG00000028070 | 5.10E-06 | 9.65E-06 | Apoa1bp | NAD(P)HX ep | Extracellular S |
| ENSMUSG00000022313 | 5.10E-06 | 9.66E-06 | Utp23 | UTP23, small | Other |
| ENSMUSG00000090137 | 5.13E-06 | 9.70E-06 | Uba52 | ubiquitin A-5 | Cytoplasm |
| ENSMUSG00000017291 | 5.13E-06 | 9.71E-06 | Taok1 | TAO kinase 1 | Cytoplasm |
| ENSMUSG00000038506 | 5.14E-06 | 9.72E-06 | Dcun1d2 | defective in c | Other |
| ENSMUSG00000030763 | 5.16E-06 | 9.74E-06 | Lcmt1 | leucine carbo | Cytoplasm |
| ENSMUSG00000009733 | 5.22E-06 | 9.86E-06 | Tfcp2 | transcription | Nucleus |
| ENSMUSG00000073423 | 5.24E-06 | 9.89E-06 | Zfp414 | zinc finger pr | Other |
| ENSMUSG00000022019 | 5.25E-06 | 9.91E-06 | Tdrd3 | tudor domair | Nucleus |
| ENSMUSG00000050188 | 5.26E-06 | 9.94E-06 | Lsm10 | LSM10, U7 sr | Nucleus |
| ENSMUSG00000029373 | 5.31E-06 | 1.00E-05 | Pf4 | platelet facto | Extracellular S |
| ENSMUSG00000002083 | 5.32E-06 | 1.00E-05 | Bbc3 | BCL2 binding | Cytoplasm |
| ENSMUSG00000020659 | 5.36E-06 | 1.01E-05 | Cbll1 | Cbl proto-or | Nucleus |
| ENSMUSG00000018401 | 5.40E-06 | 1.02E-05 | Mtmr4 | myotubularir | Cytoplasm |
| ENSMUSG00000045252 | 5.40E-06 | 1.02E-05 | Zfp574 | zinc finger pr | Other |
| ENSMUSG00000038268 | 5.45E-06 | 1.03E-05 | Ovca2 | OVCA2, serir | Cytoplasm |
| ENSMUSG00000005882 | 5.49E-06 | 1.03E-05 | Uqcc1 | ubiquinol-cy | Cytoplasm |
| ENSMUSG00000048915 | 5.49E-06 | 1.03E-05 | Efna5 | ephrin A5 | Plasma Mem |
| ENSMUSG00000030107 | 5.52E-06 | 1.04E-05 | Usp18 | ubiquitin spe | Cytoplasm |
| ENSMUSG00000027282 | 5.53E-06 | 1.04E-05 | Mtch2 | mitochondria | Cytoplasm |
| ENSMUSG00000067071 | 5.53E-06 | 1.04E-05 | Hes6 | hes family bH | Nucleus |
| ENSMUSG00000074480 | 5.53E-06 | 1.04E-05 | Mex3a | mex-3 RNA b | Cytoplasm |
| ENSMUSG00000027878 | 5.54E-06 | 1.04E-05 | Notch2 | notch 2 | Plasma Mem |
| ENSMUSG00000058794 | 5.55E-06 | 1.04E-05 | Nfe2 | nuclear facto | Nucleus |
| ENSMUSG00000056941 | 5.56E-06 | 1.05E-05 | Commd7 | COMM doma | Cytoplasm |
| ENSMUSG00000030452 | 5.64E-06 | 1.06E-05 | Nipa2 | NIPA magnes | Cytoplasm |
| ENSMUSG00000056116 | 5.65E-06 | 1.06E-05 | H2-T22 | histocompati | Other |
| ENSMUSG00000030741 | 5.67E-06 | 1.07E-05 | Spns1 | sphingolipid | Cytoplasm |
| ENSMUSG00000023972 | 5.77E-06 | 1.08E-05 | Ptk7 | protein tyros | Plasma Mem |
| ENSMUSG00000025465 | 5.82E-06 | 1.09E-05 | Echs1 | enoyl-CoA h | Cytoplasm |
| ENSMUSG00000061436 | 5.82E-06 | 1.09E-05 | Hipk2 | homeodoma | Nucleus |
| ENSMUSG00000072944 | 5.85E-06 | 1.10E-05 | Nup62cl | nucleoporin | Other |
| ENSMUSG00000045991 | 5.85E-06 | 1.10E-05 | Onecut2 | one cut hom | Nucleus |

| | | | | | |
|---|---|---|---|---|---|
| ENSMUSG00000031834 | 5.85E-06 | 1.10E-05 | Pik3r2 | phosphoinos | Cytoplasm |
| ENSMUSG00000021242 | 5.87E-06 | 1.10E-05 | Npc2 | NPC intracell | Extracellular S |
| ENSMUSG00000041777 | 5.90E-06 | 1.11E-05 | Cir1 | corepressor i | Nucleus |
| ENSMUSG00000028293 | 5.94E-06 | 1.11E-05 | Slc35a1 | solute carrier | Cytoplasm |
| ENSMUSG00000027610 | 5.99E-06 | 1.12E-05 | Gss | glutathione s | Cytoplasm |
| ENSMUSG00000025200 | 6.01E-06 | 1.13E-05 | Cwf19l1 | CWF19 like 1 | Other |
| ENSMUSG00000026181 | 6.02E-06 | 1.13E-05 | Ppm1f | protein phos | Cytoplasm |
| ENSMUSG00000022404 | 6.03E-06 | 1.13E-05 | Slc25a17 | solute carrier | Cytoplasm |
| ENSMUSG00000020349 | 6.04E-06 | 1.13E-05 | Ppp2ca | protein phos | Cytoplasm |
| ENSMUSG00000024525 | 6.09E-06 | 1.14E-05 | Impa2 | inositol mon | Cytoplasm |
| ENSMUSG00000019952 | 6.09E-06 | 1.14E-05 | Poc1b | POC1 centric | Cytoplasm |
| ENSMUSG00000026202 | 6.16E-06 | 1.15E-05 | Tuba4a | tubulin alpha | Cytoplasm |
| ENSMUSG00000027937 | 6.19E-06 | 1.16E-05 | Jtb | jumping tran | Plasma Mem |
| ENSMUSG00000031691 | 6.21E-06 | 1.16E-05 | Tnpo2 | transportin 2 | Nucleus |
| ENSMUSG00000061665 | 6.23E-06 | 1.17E-05 | Cd2ap | CD2 associat | Cytoplasm |
| ENSMUSG00000020456 | 6.26E-06 | 1.17E-05 | Ogdh | oxoglutarate | Cytoplasm |
| ENSMUSG00000031516 | 6.35E-06 | 1.19E-05 | Dctn6 | dynactin sub | Cytoplasm |
| ENSMUSG00000059743 | 6.37E-06 | 1.19E-05 | Fdps | farnesyl diph | Cytoplasm |
| ENSMUSG00000021635 | 6.42E-06 | 1.20E-05 | Rad17 | RAD17 check | Nucleus |
| ENSMUSG00000028545 | 6.43E-06 | 1.20E-05 | Bend5 | BEN domain | Cytoplasm |
| ENSMUSG00000025280 | 6.44E-06 | 1.20E-05 | Polr3a | RNA polymer | Nucleus |
| ENSMUSG00000024099 | 6.46E-06 | 1.21E-05 | Ndufv2 | NADH:ubiqu | Cytoplasm |
| ENSMUSG00000044749 | 6.46E-06 | 1.21E-05 | Abca6 | ATP binding | Plasma Mem |
| ENSMUSG00000021221 | 6.46E-06 | 1.21E-05 | Dpf3 | double PHD | Nucleus |
| ENSMUSG00000032085 | 6.54E-06 | 1.22E-05 | Tagln | transgelin | Cytoplasm |
| ENSMUSG00000031757 | 6.56E-06 | 1.22E-05 | Mt4 | metallothion | Other |
| ENSMUSG00000037936 | 6.57E-06 | 1.22E-05 | Scarb1 | scavenger re | Plasma Mem |
| ENSMUSG00000000884 | 6.57E-06 | 1.23E-05 | Gnb1l | G protein sub | Cytoplasm |
| ENSMUSG00000057342 | 6.70E-06 | 1.25E-05 | Sphk2 | sphingosine | Cytoplasm |
| ENSMUSG00000000743 | 6.78E-06 | 1.26E-05 | Chmp1a | charged mult | Extracellular S |
| ENSMUSG00000024135 | 6.89E-06 | 1.28E-05 | Srbd1 | S1 RNA bind | Other |
| ENSMUSG00000032393 | 6.94E-06 | 1.29E-05 | Dpp8 | dipeptidyl pe | Cytoplasm |
| ENSMUSG00000051391 | 6.95E-06 | 1.29E-05 | Ywhag | tyrosine 3-m | Cytoplasm |
| ENSMUSG00000025782 | 7.04E-06 | 1.31E-05 | Taf3 | TATA-box bi | Nucleus |
| ENSMUSG00000029545 | 7.06E-06 | 1.31E-05 | Acads | acyl-CoA del | Cytoplasm |
| ENSMUSG00000090231 | 7.10E-06 | 1.32E-05 | Cfb | complement | Extracellular S |
| ENSMUSG00000073700 | 7.15E-06 | 1.33E-05 | Klhl21 | kelch like fan | Cytoplasm |
| ENSMUSG00000027641 | 7.15E-06 | 1.33E-05 | Rbl1 | RB transcripti | Nucleus |
| ENSMUSG00000007827 | 7.21E-06 | 1.34E-05 | Ankrd26 | ankyrin repea | Nucleus |
| ENSMUSG00000032336 | 7.25E-06 | 1.35E-05 | Nptn | neuroplastin | Plasma Mem |
| ENSMUSG00000035772 | 7.37E-06 | 1.37E-05 | Mrps2 | mitochondria | Cytoplasm |
| ENSMUSG00000041408 | 7.40E-06 | 1.37E-05 | Wapal | WAPL cohesi | Nucleus |
| ENSMUSG00000007041 | 7.48E-06 | 1.39E-05 | Clic1 | chloride intra | Nucleus |
| ENSMUSG00000047632 | 7.55E-06 | 1.40E-05 | Fgfbp3 | fibroblast gro | Extracellular S |
| ENSMUSG00000053470 | 7.60E-06 | 1.41E-05 | Kdm3a | lysine demet | Nucleus |
| ENSMUSG00000005102 | 7.61E-06 | 1.41E-05 | Eif2ak4 | eukaryotic tra | Cytoplasm |
| ENSMUSG00000031521 | 7.70E-06 | 1.43E-05 | Aga | aspartylgluco | Cytoplasm |
| ENSMUSG00000014668 | 7.73E-06 | 1.43E-05 | Chfr | checkpoint w | Nucleus |
| ENSMUSG00000078681 | 7.76E-06 | 1.44E-05 | Tm2d3 | TM2 domain | Other |
| ENSMUSG00000020843 | 7.83E-06 | 1.45E-05 | Timm22 | translocase o | Cytoplasm |
| ENSMUSG00000091512 | 7.86E-06 | 1.46E-05 | Lamtor3 | late endosom | Cytoplasm |
| ENSMUSG00000024588 | 7.91E-06 | 1.47E-05 | Fech | ferrochelatas | Cytoplasm |
| ENSMUSG00000025047 | 8.00E-06 | 1.48E-05 | Pdcd11 | programmed | Nucleus |
| ENSMUSG00000026192 | 8.06E-06 | 1.49E-05 | Atic | 5-aminoimid | Cytoplasm |
| ENSMUSG00000022464 | 8.12E-06 | 1.50E-05 | Slc38a4 | solute carrier | Plasma Mem |
| ENSMUSG00000033628 | 8.15E-06 | 1.51E-05 | Pik3c3 | phosphatidyl | Cytoplasm |
| ENSMUSG00000056917 | 8.20E-06 | 1.52E-05 | Sipa1 | signal-induc | Cytoplasm |
| ENSMUSG00000022064 | 8.21E-06 | 1.52E-05 | Pibf1 | progesterone | Nucleus |

| | | | | | |
|---|---|---|---|---|---|
| ENSMUSG00000045730 | 8.33E-06 | 1.54E-05 | Adrb2 | adrenocepto | Plasma Mem |
| ENSMUSG00000078789 | 8.40E-06 | 1.55E-05 | Dph1 | diphthamide | Cytoplasm |
| ENSMUSG00000066880 | 8.44E-06 | 1.56E-05 | Zfp617 | zinc finger pr | Other |
| ENSMUSG00000039234 | 8.59E-06 | 1.59E-05 | Sec24d | SEC24 homo | Cytoplasm |
| ENSMUSG00000016409 | 8.68E-06 | 1.60E-05 | Nkap | NFKB activati | Nucleus |
| ENSMUSG00000058325 | 8.70E-06 | 1.61E-05 | Dock1 | dedicator of | Cytoplasm |
| ENSMUSG00000046027 | 8.79E-06 | 1.62E-05 | Stard5 | StAR related | Cytoplasm |
| ENSMUSG00000061393 | 8.82E-06 | 1.63E-05 | Acvr2b | activin A rece | Plasma Mem |
| ENSMUSG00000033460 | 8.86E-06 | 1.64E-05 | Armcx1 | armadillo rep | Other |
| ENSMUSG00000028414 | 8.88E-06 | 1.64E-05 | Fktn | fukutin | Extracellular S |
| ENSMUSG00000005813 | 8.90E-06 | 1.64E-05 | Metap1 | methionyl am | Cytoplasm |
| ENSMUSG00000028357 | 8.98E-06 | 1.66E-05 | Kif12 | kinesin family | Cytoplasm |
| ENSMUSG00000025870 | 9.11E-06 | 1.68E-05 | Arl10 | ADP ribosyla | Other |
| ENSMUSG00000025742 | 9.13E-06 | 1.68E-05 | Prps2 | phosphoribo | Cytoplasm |
| ENSMUSG00000025825 | 9.14E-06 | 1.69E-05 | Iscu | iron-sulfur cl | Cytoplasm |
| ENSMUSG00000074218 | 9.18E-06 | 1.69E-05 | Cox7a1 | cytochrome c | Cytoplasm |
| ENSMUSG00000028247 | 9.23E-06 | 1.70E-05 | Coq3 | coenzyme Q3 | Cytoplasm |
| ENSMUSG00000029141 | 9.28E-06 | 1.71E-05 | Slc4a1ap | solute carrier | Nucleus |
| ENSMUSG00000037887 | 9.32E-06 | 1.72E-05 | Dusp8 | dual specifici | Nucleus |
| ENSMUSG00000036751 | 9.32E-06 | 1.72E-05 | Cox6b1 | cytochrome c | Cytoplasm |
| ENSMUSG00000057156 | 9.42E-06 | 1.73E-05 | Homez | homeobox an | Nucleus |
| ENSMUSG00000017550 | 9.53E-06 | 1.75E-05 | Atad5 | ATPase family | Extracellular S |
| ENSMUSG00000036845 | 9.57E-06 | 1.76E-05 | Lin37 | lin-37 DREAM | Nucleus |
| ENSMUSG00000014748 | 9.62E-06 | 1.77E-05 | Tex261 | testis express | Extracellular S |
| ENSMUSG00000077450 | 9.73E-06 | 1.79E-05 | Rab11b | RAB11B, mer | Cytoplasm |
| ENSMUSG00000035397 | 9.77E-06 | 1.80E-05 | Klf16 | Kruppel-like | Nucleus |
| ENSMUSG00000070699 | 9.80E-06 | 1.80E-05 | Sars2 | seryl-tRNA sy | Cytoplasm |
| ENSMUSG00000030203 | 9.82E-06 | 1.81E-05 | Dusp16 | dual specifici | Nucleus |
| ENSMUSG00000019831 | 9.84E-06 | 1.81E-05 | Wasf1 | WAS protein | Nucleus |
| ENSMUSG00000020472 | 9.85E-06 | 1.81E-05 | Zkscan17 | zinc finger pr | Nucleus |
| ENSMUSG00000017548 | 9.98E-06 | 1.83E-05 | Suz12 | SUZ12, polyc | Nucleus |
| ENSMUSG00000033739 | 1.00E-05 | 1.84E-05 | Fkbpl | FK506 bindin | Nucleus |
| ENSMUSG00000070644 | 1.00E-05 | 1.85E-05 | Etnk2 | ethanolamine | Cytoplasm |
| ENSMUSG00000023018 | 1.01E-05 | 1.85E-05 | Smarcd1 | SWI/SNF rela | Nucleus |
| ENSMUSG00000027222 | 1.02E-05 | 1.87E-05 | Pex16 | peroxisomal | Cytoplasm |
| ENSMUSG00000028906 | 1.02E-05 | 1.87E-05 | Epb4.1 | erythrocyte m | Plasma Mem |
| ENSMUSG00000027088 | 1.03E-05 | 1.89E-05 | Phospho2 | phosphatase | Other |
| ENSMUSG00000031432 | 1.03E-05 | 1.89E-05 | Prps1 | phosphoribo | Cytoplasm |
| ENSMUSG00000052837 | 1.05E-05 | 1.93E-05 | Junb | JunB proto-o | Nucleus |
| ENSMUSG00000029815 | 1.08E-05 | 1.98E-05 | Malsu1 | mitochondria | Extracellular S |
| ENSMUSG00000026087 | 1.08E-05 | 1.98E-05 | Mrpl30 | mitochondria | Cytoplasm |
| ENSMUSG00000007476 | 1.08E-05 | 1.98E-05 | Lrrc8a | leucine rich r | Plasma Mem |
| ENSMUSG00000095595 | 1.09E-05 | 1.99E-05 | Fam177a | family with se | Other |
| ENSMUSG00000001143 | 1.10E-05 | 2.01E-05 | Lman2l | lectin, manno | Cytoplasm |
| ENSMUSG00000031865 | 1.10E-05 | 2.01E-05 | Dctn1 | dynactin sub | Cytoplasm |
| ENSMUSG00000028295 | 1.12E-05 | 2.04E-05 | Smim8 | small integra | Other |
| ENSMUSG00000014550 | 1.12E-05 | 2.04E-05 | Rbsn | rabenosyn, R | Cytoplasm |
| ENSMUSG00000053799 | 1.12E-05 | 2.05E-05 | Exoc6 | exocyst comp | Plasma Mem |
| ENSMUSG00000038618 | 1.13E-05 | 2.07E-05 | Rassf7 | Ras associatio | Cytoplasm |
| ENSMUSG00000063065 | 1.13E-05 | 2.07E-05 | Mapk3 | mitogen-acti | Cytoplasm |
| ENSMUSG00000026199 | 1.14E-05 | 2.08E-05 | Ankzf1 | ankyrin repea | Nucleus |
| ENSMUSG00000031896 | 1.16E-05 | 2.12E-05 | Ctrl | chymotrypsin | Extracellular S |
| ENSMUSG00000036299 | 1.16E-05 | 2.12E-05 | BC031181 | chromosome | Other |
| ENSMUSG00000022339 | 1.16E-05 | 2.12E-05 | Ebag9 | estrogen rece | Cytoplasm |
| ENSMUSG00000060429 | 1.17E-05 | 2.13E-05 | Sntb1 | syntrophin be | Plasma Mem |
| ENSMUSG00000060548 | 1.19E-05 | 2.17E-05 | Tnfrsf19 | TNF receptor | Plasma Mem |
| ENSMUSG00000031004 | 1.19E-05 | 2.17E-05 | Mki67 | marker of pro | Nucleus |
| ENSMUSG00000047904 | 1.20E-05 | 2.19E-05 | Sstr2 | somatostatin | Plasma Mem |

| | | | | | |
|---|---|---|---|---|---|
| ENSMUSG00000027104 | 1.22E-05 | 2.22E-05 | Atf2 | activating tra | Nucleus |
| ENSMUSG00000007891 | 1.22E-05 | 2.22E-05 | Ctsd | cathepsin D | Cytoplasm |
| ENSMUSG00000019863 | 1.24E-05 | 2.25E-05 | Qrsl1 | glutaminyl-tF | Cytoplasm |
| ENSMUSG00000015189 | 1.25E-05 | 2.27E-05 | Casd1 | CAS1 domair | Cytoplasm |
| ENSMUSG00000020884 | 1.25E-05 | 2.28E-05 | Asgr1 | asialoglycopr | Plasma Mem |
| ENSMUSG00000029725 | 1.25E-05 | 2.28E-05 | Ppp1r35 | protein phos | Other |
| ENSMUSG00000029376 | 1.26E-05 | 2.30E-05 | Mthfd2l | methylenetet | Cytoplasm |
| ENSMUSG00000045316 | 1.26E-05 | 2.30E-05 | Fahd1 | fumarylaceto | Cytoplasm |
| ENSMUSG00000036323 | 1.27E-05 | 2.32E-05 | Srp72 | signal recogr | Nucleus |
| ENSMUSG00000001441 | 1.28E-05 | 2.34E-05 | Npepps | aminopeptid | Cytoplasm |
| ENSMUSG00000043702 | 1.28E-05 | 2.34E-05 | Pde12 | phosphodies | Cytoplasm |
| ENSMUSG00000026798 | 1.29E-05 | 2.35E-05 | Coq4 | coenzyme Q | Cytoplasm |
| ENSMUSG00000026097 | 1.30E-05 | 2.37E-05 | Ormdl1 | ORMDL sphi | Cytoplasm |
| ENSMUSG00000023830 | 1.32E-05 | 2.40E-05 | Igf2r | insulin like gr | Plasma Mem |
| ENSMUSG00000038696 | 1.34E-05 | 2.44E-05 | Mapkap1 | mitogen-acti | Cytoplasm |
| ENSMUSG00000000420 | 1.34E-05 | 2.45E-05 | Galnt1 | polypeptide | Cytoplasm |
| ENSMUSG00000034875 | 1.35E-05 | 2.46E-05 | Nudt19 | nudix hydrol | Cytoplasm |
| ENSMUSG00000026980 | 1.36E-05 | 2.47E-05 | Ly75 | lymphocyte a | Plasma Mem |
| ENSMUSG00000025026 | 1.36E-05 | 2.47E-05 | Add3 | adducin 3 | Cytoplasm |
| ENSMUSG00000025410 | 1.36E-05 | 2.47E-05 | Dctn2 | dynactin sub | Cytoplasm |
| ENSMUSG00000006215 | 1.38E-05 | 2.51E-05 | Zbtb17 | zinc finger ar | Nucleus |
| ENSMUSG00000026193 | 1.39E-05 | 2.52E-05 | Fn1 | fibronectin 1 | Extracellular S |
| ENSMUSG00000003644 | 1.39E-05 | 2.53E-05 | Rps6ka1 | ribosomal pr | Cytoplasm |
| ENSMUSG00000035790 | 1.39E-05 | 2.53E-05 | Cep19 | centrosomal | Cytoplasm |
| ENSMUSG00000022540 | 1.40E-05 | 2.53E-05 | Rogdi | rogdi homolo | Nucleus |
| ENSMUSG00000011427 | 1.40E-05 | 2.55E-05 | Zfp790 | zinc finger pr | Other |
| ENSMUSG00000038312 | 1.40E-05 | 2.55E-05 | Edem2 | ER degradati | Cytoplasm |
| ENSMUSG00000087687 | 1.41E-05 | 2.56E-05 | Pet100 | PET100 hom | Other |
| ENSMUSG00000028476 | 1.42E-05 | 2.57E-05 | Reck | reversion ind | Plasma Mem |
| ENSMUSG00000026860 | 1.42E-05 | 2.57E-05 | Sh3glb2 | SH3 domain | Cytoplasm |
| ENSMUSG00000029998 | 1.43E-05 | 2.59E-05 | Pcyox1 | prenylcystein | Cytoplasm |
| ENSMUSG00000020329 | 1.43E-05 | 2.59E-05 | Polrmt | RNA polymer | Cytoplasm |
| ENSMUSG00000034194 | 1.45E-05 | 2.63E-05 | R3hcc1 | R3H domain | Other |
| ENSMUSG00000025791 | 1.46E-05 | 2.64E-05 | Pgm2 | phosphogluc | Cytoplasm |
| ENSMUSG00000013646 | 1.46E-05 | 2.64E-05 | Sh3bp5l | SH3 binding | Other |
| ENSMUSG00000028426 | 1.50E-05 | 2.72E-05 | Rad23b | RAD23 homo | Nucleus |
| ENSMUSG00000094483 | 1.52E-05 | 2.75E-05 | Purb | purine rich el | Nucleus |
| ENSMUSG00000047514 | 1.52E-05 | 2.75E-05 | Tspyl1 | TSPY like 1 | Nucleus |
| ENSMUSG00000003541 | 1.53E-05 | 2.77E-05 | Ier3 | immediate ea | Cytoplasm |
| ENSMUSG00000020454 | 1.54E-05 | 2.78E-05 | Eif4enif1 | eukaryotic tra | Cytoplasm |
| ENSMUSG00000009630 | 1.54E-05 | 2.78E-05 | Ppp2cb | protein phos | Cytoplasm |
| ENSMUSG00000014601 | 1.54E-05 | 2.78E-05 | Strip1 | striatin intera | Nucleus |
| ENSMUSG00000022771 | 1.54E-05 | 2.79E-05 | Ppil2 | peptidylproly | Nucleus |
| ENSMUSG00000026175 | 1.54E-05 | 2.79E-05 | Vil1 | villin 1 | Cytoplasm |
| ENSMUSG00000030091 | 1.55E-05 | 2.80E-05 | Nup210 | nucleoporin | Nucleus |
| ENSMUSG00000002768 | 1.56E-05 | 2.82E-05 | Mea1 | male-enhanc | Cytoplasm |
| ENSMUSG00000031577 | 1.57E-05 | 2.83E-05 | Tti2 | TELO2 intera | Cytoplasm |
| ENSMUSG00000033055 | 1.57E-05 | 2.84E-05 | Ankrd54 | ankyrin repea | Nucleus |
| ENSMUSG00000021120 | 1.58E-05 | 2.86E-05 | Pigh | phosphatidyl | Cytoplasm |
| ENSMUSG00000028229 | 1.59E-05 | 2.86E-05 | Rmdn1 | regulator of | Cytoplasm |
| ENSMUSG00000039187 | 1.59E-05 | 2.87E-05 | Fanci | FA complem | Nucleus |
| ENSMUSG00000032754 | 1.59E-05 | 2.87E-05 | Slc8b1 | solute carrier | Plasma Mem |
| ENSMUSG00000051502 | 1.60E-05 | 2.88E-05 | Ufsp1 | UFM1 specifi | Cytoplasm |
| ENSMUSG00000024908 | 1.61E-05 | 2.90E-05 | Ppp6r3 | protein phos | Cytoplasm |
| ENSMUSG00000024792 | 1.61E-05 | 2.91E-05 | Zfpl1 | zinc finger pr | Cytoplasm |
| ENSMUSG00000067713 | 1.62E-05 | 2.92E-05 | Prkag1 | protein kinas | Nucleus |
| ENSMUSG00000048118 | 1.62E-05 | 2.92E-05 | Arid4a | AT-rich inter | Nucleus |
| ENSMUSG00000023345 | 1.62E-05 | 2.92E-05 | Poc1a | POC1 centric | Cytoplasm |

| | | | | | |
|---|---|---|---|---|---|
| ENSMUSG00000035069 | 1.62E-05 | 2.92E-05 | Oma1 | OMA1 zinc m | Cytoplasm |
| ENSMUSG00000031443 | 1.62E-05 | 2.92E-05 | F7 | coagulation f | Plasma Mem |
| ENSMUSG00000041975 | 1.62E-05 | 2.92E-05 | Mettl8 | methyltransfe | Other |
| ENSMUSG00000032067 | 1.63E-05 | 2.93E-05 | Pts | 6-pyruvoylte | Cytoplasm |
| ENSMUSG00000025036 | 1.63E-05 | 2.93E-05 | Sfxn2 | sideroflexin 2 | Cytoplasm |
| ENSMUSG00000029464 | 1.64E-05 | 2.94E-05 | Gpn3 | GPN-loop G | Cytoplasm |
| ENSMUSG00000034269 | 1.64E-05 | 2.95E-05 | Setd5 | SET domain c | Nucleus |
| ENSMUSG00000002732 | 1.64E-05 | 2.96E-05 | Fkbp7 | FK506 bindin | Cytoplasm |
| ENSMUSG00000026577 | 1.65E-05 | 2.97E-05 | Blzf1 | basic leucine | Cytoplasm |
| ENSMUSG00000025780 | 1.65E-05 | 2.97E-05 | Itih5 | inter-alpha-t | Plasma Mem |
| ENSMUSG00000079317 | 1.65E-05 | 2.97E-05 | Trappc2 | trafficking pr | Cytoplasm |
| ENSMUSG00000034837 | 1.65E-05 | 2.97E-05 | Gnat1 | G protein sub | Plasma Mem |
| ENSMUSG00000050530 | 1.68E-05 | 3.02E-05 | Fam171a1 | family with se | Cytoplasm |
| ENSMUSG00000030674 | 1.72E-05 | 3.08E-05 | Qprt | quinolinate p | Cytoplasm |
| ENSMUSG00000021076 | 1.72E-05 | 3.09E-05 | Actr10 | actin related | Cytoplasm |
| ENSMUSG00000049728 | 1.78E-05 | 3.19E-05 | Zfp668 | zinc finger pr | Cytoplasm |
| ENSMUSG00000026692 | 1.78E-05 | 3.19E-05 | Fmo4 | flavin contair | Cytoplasm |
| ENSMUSG00000029389 | 1.78E-05 | 3.20E-05 | Ddx55 | DEAD-box he | Nucleus |
| ENSMUSG00000028885 | 1.79E-05 | 3.20E-05 | Smpdl3b | sphingomyel | Extracellular S |
| ENSMUSG00000012429 | 1.80E-05 | 3.22E-05 | Mplkip | M-phase spe | Nucleus |
| ENSMUSG00000008976 | 1.81E-05 | 3.25E-05 | Gabpa | GA binding p | Nucleus |
| ENSMUSG00000017009 | 1.82E-05 | 3.26E-05 | Sdc4 | syndecan 4 | Plasma Mem |
| ENSMUSG00000034211 | 1.82E-05 | 3.26E-05 | Mrps17 | mitochondria | Cytoplasm |
| ENSMUSG00000026213 | 1.82E-05 | 3.27E-05 | Stk11ip | serine/threor | Cytoplasm |
| ENSMUSG00000030965 | 1.84E-05 | 3.30E-05 | Fam175b | abraxas 2, BR | Nucleus |
| ENSMUSG00000038539 | 1.85E-05 | 3.31E-05 | Atf5 | activating tra | Nucleus |
| ENSMUSG00000009112 | 1.86E-05 | 3.32E-05 | Bcl2l13 | BCL2 like 13 | Cytoplasm |
| ENSMUSG00000051355 | 1.86E-05 | 3.32E-05 | Commd1 | copper meta | Nucleus |
| ENSMUSG00000045954 | 1.86E-05 | 3.33E-05 | Sdpr | caveolae asso | Plasma Mem |
| ENSMUSG00000030313 | 1.88E-05 | 3.36E-05 | Dennd5b | DENN domai | Cytoplasm |
| ENSMUSG00000053950 | 1.88E-05 | 3.36E-05 | Adnp2 | ADNP homeo | Other |
| ENSMUSG00000050628 | 1.90E-05 | 3.39E-05 | Ubald2 | UBA like dom | Other |
| ENSMUSG00000028969 | 1.90E-05 | 3.40E-05 | Cdk5 | cyclin depend | Nucleus |
| ENSMUSG00000053600 | 1.90E-05 | 3.40E-05 | Zfp472 | zinc finger pr | Nucleus |
| ENSMUSG00000020745 | 1.90E-05 | 3.40E-05 | Pafah1b1 | platelet activ | Cytoplasm |
| ENSMUSG00000018923 | 1.91E-05 | 3.42E-05 | Med11 | mediator cor | Nucleus |
| ENSMUSG00000038014 | 1.93E-05 | 3.44E-05 | Fam120a | family with se | Cytoplasm |
| ENSMUSG00000010054 | 1.95E-05 | 3.49E-05 | Tusc2 | tumor suppre | Nucleus |
| ENSMUSG00000026418 | 1.96E-05 | 3.50E-05 | Tnni1 | troponin I1, s | Cytoplasm |
| ENSMUSG00000037440 | 1.96E-05 | 3.50E-05 | Vnn1 | vanin 1 | Plasma Mem |
| ENSMUSG00000045160 | 1.96E-05 | 3.50E-05 | Bola3 | bolA family n | Cytoplasm |
| ENSMUSG00000042211 | 1.98E-05 | 3.54E-05 | Fbxo38 | F-box protei | Other |
| ENSMUSG00000019087 | 1.99E-05 | 3.54E-05 | Atp6ap1 | ATPase H+ tr | Cytoplasm |
| ENSMUSG00000017679 | 1.99E-05 | 3.55E-05 | Ttpal | alpha tocoph | Other |
| ENSMUSG00000046062 | 1.99E-05 | 3.55E-05 | Ppp1r15b | protein phos | Cytoplasm |
| ENSMUSG00000090935 | 2.00E-05 | 3.57E-05 | Synj2bp | synaptojanin | Cytoplasm |
| ENSMUSG00000031502 | 2.00E-05 | 3.57E-05 | Col4a1 | collagen type | Extracellular S |
| ENSMUSG00000026048 | 2.02E-05 | 3.60E-05 | Ercc5 | ERCC excisio | Nucleus |
| ENSMUSG00000052392 | 2.02E-05 | 3.60E-05 | Acot4 | acyl-CoA thio | Cytoplasm |
| ENSMUSG00000027654 | 2.02E-05 | 3.60E-05 | Fam83d | family with se | Cytoplasm |
| ENSMUSG00000027708 | 2.03E-05 | 3.61E-05 | Dcun1d1 | defective in c | Nucleus |
| ENSMUSG00000043998 | 2.05E-05 | 3.66E-05 | Mgat2 | mannosyl (al | Cytoplasm |
| ENSMUSG00000029557 | 2.06E-05 | 3.68E-05 | Ftsj2 | mitochondria | Nucleus |
| ENSMUSG00000018599 | 2.07E-05 | 3.68E-05 | Mief2 | mitochondria | Cytoplasm |
| ENSMUSG00000027363 | 2.07E-05 | 3.69E-05 | Usp8 | ubiquitin spe | Cytoplasm |
| ENSMUSG00000016984 | 2.08E-05 | 3.70E-05 | Etaa1 | ETAA1, ATR k | Nucleus |
| ENSMUSG00000039461 | 2.09E-05 | 3.71E-05 | Tcta | T cell leukem | Other |
| ENSMUSG00000020818 | 2.09E-05 | 3.72E-05 | Mfsd11 | major facilita | Other |

| | | | | | |
|---|---|---|---|---|---|
| ENSMUSG00000040669 | 2.12E-05 | 3.76E-05 | Phc1 | polyhomeoti | Nucleus |
| ENSMUSG00000056228 | 2.12E-05 | 3.77E-05 | Cars2 | cysteinyl-tRN | Cytoplasm |
| ENSMUSG00000002395 | 2.12E-05 | 3.78E-05 | Use1 | unconventior | Cytoplasm |
| ENSMUSG00000027953 | 2.14E-05 | 3.81E-05 | Slc50a1 | solute carrier | Plasma Mem |
| ENSMUSG00000040112 | 2.15E-05 | 3.82E-05 | Mrps35 | mitochondria | Cytoplasm |
| ENSMUSG00000020778 | 2.16E-05 | 3.84E-05 | Ten1 | TEN1, CST cc | Nucleus |
| ENSMUSG00000027601 | 2.17E-05 | 3.86E-05 | Mtfr1 | mitochondria | Cytoplasm |
| ENSMUSG00000027646 | 2.18E-05 | 3.86E-05 | Src | SRC proto-oı | Cytoplasm |
| ENSMUSG00000020780 | 2.18E-05 | 3.88E-05 | Srp68 | signal recogr | Nucleus |
| ENSMUSG00000034893 | 2.21E-05 | 3.92E-05 | Cog3 | component c | Cytoplasm |
| ENSMUSG00000039703 | 2.26E-05 | 4.01E-05 | Nploc4 | NPL4 homolc | Nucleus |
| ENSMUSG00000028759 | 2.28E-05 | 4.04E-05 | Hp1bp3 | heterochrom | Nucleus |
| ENSMUSG00000018372 | 2.28E-05 | 4.04E-05 | Cep95 | centrosomal | Cytoplasm |
| ENSMUSG00000049233 | 2.29E-05 | 4.06E-05 | Apoo-ps | apolipoprote | Other |
| ENSMUSG00000025223 | 2.29E-05 | 4.06E-05 | Ldb1 | LIM domain I | Nucleus |
| ENSMUSG00000031561 | 2.30E-05 | 4.07E-05 | Tenm3 | teneurin tran | Plasma Mem |
| ENSMUSG00000028064 | 2.34E-05 | 4.14E-05 | Sema4a | semaphorin 4 | Plasma Mem |
| ENSMUSG00000007034 | 2.34E-05 | 4.15E-05 | Slc44a4 | solute carrier | Plasma Mem |
| ENSMUSG00000013997 | 2.36E-05 | 4.18E-05 | Nit1 | nitrilase 1 | Cytoplasm |
| ENSMUSG00000037995 | 2.36E-05 | 4.19E-05 | Igsf9 | immunoglob | Plasma Mem |
| ENSMUSG00000019960 | 2.38E-05 | 4.22E-05 | Dusp6 | dual specifici | Cytoplasm |
| ENSMUSG00000050323 | 2.39E-05 | 4.23E-05 | Ndufaf6 | NADH:ubiqu | Cytoplasm |
| ENSMUSG00000049439 | 2.40E-05 | 4.24E-05 | Cyp20a1 | cytochrome I | Other |
| ENSMUSG00000033088 | 2.40E-05 | 4.24E-05 | Triobp | TRIO and F-a | Nucleus |
| ENSMUSG00000024350 | 2.40E-05 | 4.25E-05 | Dnajc18 | DnaJ heat sh | Other |
| ENSMUSG00000002210 | 2.41E-05 | 4.26E-05 | Smg9 | SMG9, nonse | Cytoplasm |
| ENSMUSG00000038690 | 2.41E-05 | 4.27E-05 | Atp5j2 | ATP synthase | Cytoplasm |
| ENSMUSG00000034361 | 2.41E-05 | 4.27E-05 | Cpne2 | copine 2 | Cytoplasm |
| ENSMUSG00000015337 | 2.42E-05 | 4.27E-05 | Endog | endonucleas | Cytoplasm |
| ENSMUSG00000033444 | 2.42E-05 | 4.28E-05 | Specc1l | sperm antige | Extracellular S |
| ENSMUSG00000033373 | 2.43E-05 | 4.29E-05 | Fntb | farnesyltrans | Cytoplasm |
| ENSMUSG00000058351 | 2.46E-05 | 4.34E-05 | Smim4 | small integra | Cytoplasm |
| ENSMUSG00000022883 | 2.47E-05 | 4.37E-05 | Robo1 | roundabout g | Plasma Mem |
| ENSMUSG00000022295 | 2.48E-05 | 4.38E-05 | Atp6v1c1 | ATPase H+ tı | Cytoplasm |
| ENSMUSG00000028552 | 2.49E-05 | 4.40E-05 | Eps15 | epidermal gr | Cytoplasm |
| ENSMUSG00000046718 | 2.52E-05 | 4.45E-05 | Bst2 | bone marrow | Plasma Mem |
| ENSMUSG00000027012 | 2.53E-05 | 4.46E-05 | Dync1i2 | dynein cytop | Cytoplasm |
| ENSMUSG00000058396 | 2.53E-05 | 4.46E-05 | Gpr182 | G protein-co | Plasma Mem |
| ENSMUSG00000033790 | 2.54E-05 | 4.47E-05 | Tubgcp5 | tubulin gamr | Cytoplasm |
| ENSMUSG00000096910 | 2.55E-05 | 4.50E-05 | Zfp955b | zinc finger pı | Other |
| ENSMUSG00000022571 | 2.57E-05 | 4.53E-05 | Pycrl | pyrroline-5-c | Cytoplasm |
| ENSMUSG00000020513 | 2.58E-05 | 4.54E-05 | Tubd1 | tubulin delta | Cytoplasm |
| ENSMUSG00000021103 | 2.58E-05 | 4.55E-05 | Mnat1 | MNAT1, CDK | Nucleus |
| ENSMUSG00000024352 | 2.59E-05 | 4.57E-05 | Spata24 | spermatoger | Nucleus |
| ENSMUSG00000028563 | 2.61E-05 | 4.59E-05 | Tm2d1 | TM2 domain | Plasma Mem |
| ENSMUSG00000027506 | 2.64E-05 | 4.65E-05 | Tpd52 | tumor proteiı | Cytoplasm |
| ENSMUSG00000017764 | 2.67E-05 | 4.70E-05 | Zswim1 | zinc finger S\ | Nucleus |
| ENSMUSG00000022760 | 2.67E-05 | 4.71E-05 | Thap7 | THAP domaiı | Nucleus |
| ENSMUSG00000025078 | 2.70E-05 | 4.75E-05 | Nhlrc2 | NHL repeat c | Other |
| ENSMUSG00000045251 | 2.73E-05 | 4.79E-05 | Zfp688 | zinc finger pı | Other |
| ENSMUSG00000057103 | 2.73E-05 | 4.80E-05 | Cml1 | N-acetyltrans | Cytoplasm |
| ENSMUSG00000029826 | 2.76E-05 | 4.85E-05 | Zc3hav1 | zinc finger C( | Plasma Mem |
| ENSMUSG00000000326 | 2.77E-05 | 4.86E-05 | Comt | catechol-O-ı | Cytoplasm |
| ENSMUSG00000034173 | 2.77E-05 | 4.87E-05 | Zbed5 | zinc finger BI | Nucleus |
| ENSMUSG00000018395 | 2.78E-05 | 4.88E-05 | Kif3a | kinesin family | Cytoplasm |
| ENSMUSG00000053581 | 2.80E-05 | 4.92E-05 | Zfand2a | zinc finger Al | Other |
| ENSMUSG00000029661 | 2.80E-05 | 4.92E-05 | Col1a2 | collagen type | Extracellular S |
| ENSMUSG00000040447 | 2.85E-05 | 5.00E-05 | Spns2 | sphingolipid | Other |

| | | | | | |
|---|---|---|---|---|---|
| ENSMUSG00000056050 | 2.87E-05 | 5.04E-05 | Mia3 | MIA SH3 dor | Cytoplasm |
| ENSMUSG00000006574 | 2.88E-05 | 5.06E-05 | Slc4a1 | solute carrier | Plasma Mem |
| ENSMUSG00000040472 | 2.91E-05 | 5.10E-05 | Rabggta | Rab geranylg | Extracellular S |
| ENSMUSG00000089911 | 2.92E-05 | 5.12E-05 | Hiat1 | major facilita | Other |
| ENSMUSG00000058793 | 2.92E-05 | 5.13E-05 | Cds2 | CDP-diacylgl | Cytoplasm |
| ENSMUSG00000062929 | 2.93E-05 | 5.14E-05 | Cfl2 | cofilin 2 | Extracellular S |
| ENSMUSG00000020530 | 2.95E-05 | 5.17E-05 | Ggnbp2 | gametogene | Other |
| ENSMUSG00000023079 | 2.95E-05 | 5.18E-05 | Gtf2ird1 | GTF2I repeat | Nucleus |
| ENSMUSG00000030815 | 2.96E-05 | 5.19E-05 | Phkg2 | phosphorylas | Cytoplasm |
| ENSMUSG00000003423 | 2.99E-05 | 5.25E-05 | Pih1d1 | PIH1 domain | Nucleus |
| ENSMUSG00000052712 | 3.01E-05 | 5.27E-05 | BC004004 | chromosome | Cytoplasm |
| ENSMUSG00000047921 | 3.02E-05 | 5.28E-05 | Trappc9 | trafficking pr | Plasma Mem |
| ENSMUSG00000068394 | 3.02E-05 | 5.29E-05 | Cep152 | centrosomal | Cytoplasm |
| ENSMUSG00000038150 | 3.03E-05 | 5.30E-05 | Ormdl3 | ORMDL sphi | Cytoplasm |
| ENSMUSG00000019039 | 3.06E-05 | 5.35E-05 | Dalrd3 | DALR anticod | Other |
| ENSMUSG00000038732 | 3.06E-05 | 5.35E-05 | Mboat1 | membrane b | Cytoplasm |
| ENSMUSG00000028688 | 3.09E-05 | 5.40E-05 | Toe1 | target of EGF | Nucleus |
| ENSMUSG00000001150 | 3.09E-05 | 5.41E-05 | Mcm3ap | minichromos | Nucleus |
| ENSMUSG00000033436 | 3.09E-05 | 5.41E-05 | Armcx2 | armadillo rep | Other |
| ENSMUSG00000064289 | 3.13E-05 | 5.47E-05 | Tank | TRAF family r | Cytoplasm |
| ENSMUSG00000037949 | 3.16E-05 | 5.51E-05 | Ano10 | anoctamin 10 | Plasma Mem |
| ENSMUSG00000020955 | 3.16E-05 | 5.52E-05 | Ap4s1 | adaptor relat | Cytoplasm |
| ENSMUSG00000028483 | 3.17E-05 | 5.54E-05 | Snapc3 | small nuclear | Nucleus |
| ENSMUSG00000015869 | 3.24E-05 | 5.65E-05 | Prpsap1 | phosphoribo | Extracellular S |
| ENSMUSG00000046679 | 3.28E-05 | 5.72E-05 | C87436 | chromosome | Nucleus |
| ENSMUSG00000029003 | 3.29E-05 | 5.75E-05 | Mad2l2 | mitotic arrest | Nucleus |
| ENSMUSG00000035234 | 3.36E-05 | 5.86E-05 | Fam175a | abraxas 1, BR | Nucleus |
| ENSMUSG00000036199 | 3.36E-05 | 5.86E-05 | Ndufa13 | NADH:ubiqu | Cytoplasm |
| ENSMUSG00000030035 | 3.40E-05 | 5.93E-05 | Wbp1 | WW domain | Cytoplasm |
| ENSMUSG00000038084 | 3.41E-05 | 5.95E-05 | Opa1 | OPA1, mitocl | Cytoplasm |
| ENSMUSG00000003033 | 3.41E-05 | 5.95E-05 | Ap1m1 | adaptor relat | Cytoplasm |
| ENSMUSG00000075467 | 3.46E-05 | 6.03E-05 | Dnlz | DNL-type zir | Cytoplasm |
| ENSMUSG00000048439 | 3.46E-05 | 6.03E-05 | Nupl2 | nucleoporin l | Nucleus |
| ENSMUSG00000024220 | 3.48E-05 | 6.05E-05 | Zfp523 | zinc finger pr | Nucleus |
| ENSMUSG00000015377 | 3.52E-05 | 6.13E-05 | Dennd6b | DENN domai | Cytoplasm |
| ENSMUSG00000045107 | 3.54E-05 | 6.16E-05 | Saysd1 | SAYSVFN mc | Other |
| ENSMUSG00000030272 | 3.57E-05 | 6.21E-05 | Camk1 | calcium/calm | Cytoplasm |
| ENSMUSG00000031767 | 3.57E-05 | 6.21E-05 | Nudt7 | nudix hydrola | Cytoplasm |
| ENSMUSG00000029761 | 3.58E-05 | 6.23E-05 | Cald1 | caldesmon 1 | Plasma Mem |
| ENSMUSG00000019813 | 3.59E-05 | 6.24E-05 | Cep57l1 | centrosomal | Other |
| ENSMUSG00000021967 | 3.60E-05 | 6.27E-05 | Mrpl57 | mitochondria | Cytoplasm |
| ENSMUSG00000022048 | 3.64E-05 | 6.34E-05 | Dpysl2 | dihydropyrim | Cytoplasm |
| ENSMUSG00000073889 | 3.66E-05 | 6.36E-05 | Il11ra1 | interleukin 11 | Plasma Mem |
| ENSMUSG00000050732 | 3.74E-05 | 6.49E-05 | Vamp8 | vesicle associ | Plasma Mem |
| ENSMUSG00000028636 | 3.75E-05 | 6.52E-05 | Ppcs | phosphopan | Cytoplasm |
| ENSMUSG00000053192 | 3.78E-05 | 6.57E-05 | Mllt11 | MLLT11, tran | Cytoplasm |
| ENSMUSG00000029176 | 3.80E-05 | 6.61E-05 | Anapc4 | anaphase pro | Nucleus |
| ENSMUSG00000074102 | 3.81E-05 | 6.62E-05 | Rbm15b | RNA binding | Nucleus |
| ENSMUSG00000030881 | 3.82E-05 | 6.63E-05 | Arfip2 | ADP ribosyla | Cytoplasm |
| ENSMUSG00000024750 | 3.82E-05 | 6.64E-05 | Zfand5 | zinc finger AI | Nucleus |
| ENSMUSG00000039163 | 3.84E-05 | 6.67E-05 | Cmc1 | C-X9-C moti | Cytoplasm |
| ENSMUSG00000024854 | 3.86E-05 | 6.70E-05 | Pold4 | DNA polyme | Nucleus |
| ENSMUSG00000026153 | 3.87E-05 | 6.71E-05 | Fam135a | family with se | Other |
| ENSMUSG00000025165 | 3.90E-05 | 6.76E-05 | Sectm1a | secreted and | Extracellular S |
| ENSMUSG00000026239 | 3.95E-05 | 6.84E-05 | Pde6d | phosphodies | Cytoplasm |
| ENSMUSG00000046836 | 3.96E-05 | 6.86E-05 | Brox | BRO1 domai | Cytoplasm |
| ENSMUSG00000035601 | 3.96E-05 | 6.86E-05 | Trmt10b | tRNA methyl | Cytoplasm |
| ENSMUSG00000060601 | 3.96E-05 | 6.87E-05 | Nr1h2 | nuclear recep | Nucleus |

| | | | | | |
|---|---|---|---|---|---|
| ENSMUSG00000023963 | 4.00E-05 | 6.93E-05 | Cyp39a1 | cytochrome I | Cytoplasm |
| ENSMUSG00000022329 | 4.01E-05 | 6.94E-05 | Stk3 | serine/threor | Cytoplasm |
| ENSMUSG00000044502 | 4.04E-05 | 6.99E-05 | Bod1 | biorientation | Nucleus |
| ENSMUSG00000000581 | 4.06E-05 | 7.03E-05 | C1d | C1D nuclear | Nucleus |
| ENSMUSG00000020160 | 4.07E-05 | 7.04E-05 | Meis1 | Meis homeol | Nucleus |
| ENSMUSG00000039656 | 4.07E-05 | 7.05E-05 | Rxrb | retinoid X rec | Nucleus |
| ENSMUSG00000025077 | 4.08E-05 | 7.05E-05 | Dclre1a | DNA cross-li | Nucleus |
| ENSMUSG00000043411 | 4.09E-05 | 7.08E-05 | Usp48 | ubiquitin spe | Plasma Mem |
| ENSMUSG00000049354 | 4.17E-05 | 7.22E-05 | Dcaf7 | DDB1 and CU | Cytoplasm |
| ENSMUSG00000026547 | 4.20E-05 | 7.25E-05 | Tagln2 | transgelin 2 | Cytoplasm |
| ENSMUSG00000051730 | 4.20E-05 | 7.26E-05 | Mettl5 | methyltransfe | Other |
| ENSMUSG00000039068 | 4.22E-05 | 7.29E-05 | Zzz3 | zinc finger ZZ | Nucleus |
| ENSMUSG00000074865 | 4.23E-05 | 7.31E-05 | Zfp934 | expressed se | Other |
| ENSMUSG00000015013 | 4.26E-05 | 7.36E-05 | Trappc2l | trafficking pr | Cytoplasm |
| ENSMUSG00000027429 | 4.26E-05 | 7.36E-05 | Sec23b | Sec23 homol | Extracellular S |
| ENSMUSG00000020198 | 4.28E-05 | 7.39E-05 | Ap3d1 | adaptor relat | Cytoplasm |
| ENSMUSG00000029655 | 4.28E-05 | 7.39E-05 | N4bp2l2 | NEDD4 bindi | Nucleus |
| ENSMUSG00000031381 | 4.33E-05 | 7.47E-05 | Piga | phosphatidyl | Cytoplasm |
| ENSMUSG00000027349 | 4.37E-05 | 7.53E-05 | Fam98b | family with se | Nucleus |
| ENSMUSG00000027075 | 4.38E-05 | 7.55E-05 | Slc43a1 | solute carrier | Plasma Mem |
| ENSMUSG00000021765 | 4.38E-05 | 7.56E-05 | Fst | follistatin | Extracellular S |
| ENSMUSG00000020671 | 4.41E-05 | 7.61E-05 | Rab10 | RAB10, mem | Cytoplasm |
| ENSMUSG00000019467 | 4.42E-05 | 7.61E-05 | Arhgef25 | Rho guanine | Cytoplasm |
| ENSMUSG00000030410 | 4.42E-05 | 7.62E-05 | Dmwd | DM1 locus, W | Nucleus |
| ENSMUSG00000003545 | 4.44E-05 | 7.66E-05 | Fosb | FosB proto-c | Nucleus |
| ENSMUSG00000073125 | 4.45E-05 | 7.67E-05 | Xlr3b | X-linked lym | Other |
| ENSMUSG00000032570 | 4.47E-05 | 7.69E-05 | Atp2c1 | ATPase secre | Cytoplasm |
| ENSMUSG00000038324 | 4.52E-05 | 7.78E-05 | Trpc4ap | transient rece | Cytoplasm |
| ENSMUSG00000041995 | 4.55E-05 | 7.84E-05 | Zbed3 | zinc finger BE | Cytoplasm |
| ENSMUSG00000034290 | 4.57E-05 | 7.87E-05 | Nek9 | NIMA relatec | Nucleus |
| ENSMUSG00000002032 | 4.57E-05 | 7.87E-05 | Tmem25 | transmembra | Extracellular S |
| ENSMUSG00000024052 | 4.58E-05 | 7.88E-05 | Lpin2 | lipin 2 | Nucleus |
| ENSMUSG00000032688 | 4.61E-05 | 7.93E-05 | Malt1 | MALT1 parac | Cytoplasm |
| ENSMUSG00000028266 | 4.62E-05 | 7.94E-05 | Lmo4 | LIM domain c | Nucleus |
| ENSMUSG00000067787 | 4.68E-05 | 8.05E-05 | Blcap | BLCAP, apop | Other |
| ENSMUSG00000038508 | 4.68E-05 | 8.05E-05 | Gdf15 | growth differ | Extracellular S |
| ENSMUSG00000000439 | 4.69E-05 | 8.06E-05 | Mkrn2 | makorin ring | Other |
| ENSMUSG00000029814 | 4.70E-05 | 8.07E-05 | Igf2bp3 | insulin like gr | Cytoplasm |
| ENSMUSG00000036606 | 4.71E-05 | 8.10E-05 | Plxnb2 | plexin B2 | Plasma Mem |
| ENSMUSG00000001419 | 4.75E-05 | 8.16E-05 | Mef2d | myocyte enh | Nucleus |
| ENSMUSG00000022544 | 4.77E-05 | 8.20E-05 | Eef2kmt | eukaryotic el | Cytoplasm |
| ENSMUSG00000032184 | 4.79E-05 | 8.23E-05 | Lysmd2 | LysM domair | Other |
| ENSMUSG00000042229 | 4.80E-05 | 8.23E-05 | Rabif | RAB interacti | Other |
| ENSMUSG00000044279 | 4.80E-05 | 8.25E-05 | Crb3 | crumbs famil | Plasma Mem |
| ENSMUSG00000001576 | 4.81E-05 | 8.26E-05 | Ergic1 | endoplasmic | Cytoplasm |
| ENSMUSG00000015478 | 4.82E-05 | 8.26E-05 | Rnf5 | ring finger pr | Cytoplasm |
| ENSMUSG00000040940 | 4.82E-05 | 8.27E-05 | Arhgef1 | Rho guanine | Cytoplasm |
| ENSMUSG00000040998 | 4.85E-05 | 8.32E-05 | Npnt | nephronectir | Plasma Mem |
| ENSMUSG00000000811 | 4.90E-05 | 8.40E-05 | Txnrd3 | thioredoxin r | Cytoplasm |
| ENSMUSG00000057551 | 4.90E-05 | 8.40E-05 | Zfp317 | zinc finger pr | Nucleus |
| ENSMUSG00000020733 | 4.93E-05 | 8.45E-05 | Slc9a3r1 | SLC9A3 regu | Plasma Mem |
| ENSMUSG00000020895 | 4.95E-05 | 8.49E-05 | Tmem107 | transmembra | Extracellular S |
| ENSMUSG00000040043 | 4.97E-05 | 8.51E-05 | Rbms2 | RNA binding | Nucleus |
| ENSMUSG00000061048 | 4.97E-05 | 8.52E-05 | Cdh3 | cadherin 3 | Plasma Mem |
| ENSMUSG00000019494 | 4.98E-05 | 8.53E-05 | Cops6 | COP9 signalc | Nucleus |
| ENSMUSG00000024970 | 5.01E-05 | 8.58E-05 | AI846148 | spindlin inter | Other |
| ENSMUSG00000002608 | 5.01E-05 | 8.58E-05 | Ccdc97 | coiled-coil de | Other |
| ENSMUSG00000024580 | 5.01E-05 | 8.58E-05 | Grpel2 | GrpE like 2, n | Cytoplasm |

| | | | | | |
|---|---|---|---|---|---|
| ENSMUSG00000050541 | 5.03E-05 | 8.61E-05 | Adra1b | adrenocepto | Plasma Mem |
| ENSMUSG00000045078 | 5.05E-05 | 8.64E-05 | Rnf216 | ring finger pr | Cytoplasm |
| ENSMUSG00000035365 | 5.07E-05 | 8.67E-05 | Parpbp | PARP1 bindir | Nucleus |
| ENSMUSG00000022414 | 5.10E-05 | 8.72E-05 | Tab1 | TGF-beta act | Cytoplasm |
| ENSMUSG00000062794 | 5.13E-05 | 8.77E-05 | Zfp599 | zinc finger pr | Other |
| ENSMUSG00000025732 | 5.15E-05 | 8.80E-05 | Fam195a | MAPK regula | Cytoplasm |
| ENSMUSG00000017466 | 5.15E-05 | 8.81E-05 | Timp2 | TIMP metallo | Extracellular S |
| ENSMUSG00000003346 | 5.16E-05 | 8.81E-05 | Abhd17a | abhydrolase | Plasma Mem |
| ENSMUSG00000031950 | 5.16E-05 | 8.82E-05 | Gabarapl2 | GABA type A | Cytoplasm |
| ENSMUSG00000019791 | 5.17E-05 | 8.83E-05 | Hint3 | histidine triac | Cytoplasm |
| ENSMUSG00000022956 | 5.19E-05 | 8.86E-05 | Atp5o | ATP synthase | Cytoplasm |
| ENSMUSG00000028893 | 5.19E-05 | 8.87E-05 | Sesn2 | sestrin 2 | Cytoplasm |
| ENSMUSG00000001089 | 5.25E-05 | 8.95E-05 | Luzp1 | leucine zippe | Nucleus |
| ENSMUSG00000033735 | 5.28E-05 | 9.00E-05 | Spr | sepiapterin re | Cytoplasm |
| ENSMUSG00000032249 | 5.29E-05 | 9.02E-05 | Anp32a | acidic (leucin | Nucleus |
| ENSMUSG00000039842 | 5.32E-05 | 9.07E-05 | Mcph1 | microcephali | Nucleus |
| ENSMUSG00000024817 | 5.34E-05 | 9.10E-05 | Uhrf2 | ubiquitin like | Nucleus |
| ENSMUSG00000021589 | 5.34E-05 | 9.11E-05 | Rhobtb3 | Rho related E | Cytoplasm |
| ENSMUSG00000047648 | 5.52E-05 | 9.41E-05 | Fbxo30 | F-box protei | Extracellular S |
| ENSMUSG00000032417 | 5.58E-05 | 9.51E-05 | Rwdd2a | RWD domain | Other |
| ENSMUSG00000035472 | 5.62E-05 | 9.58E-05 | Slc25a21 | solute carrier | Cytoplasm |
| ENSMUSG00000001829 | 5.68E-05 | 9.67E-05 | Clpb | ClpB homolo | Nucleus |
| ENSMUSG00000023051 | 5.68E-05 | 9.67E-05 | Tarbp2 | TARBP2, RISC | Nucleus |
| ENSMUSG00000039458 | 5.69E-05 | 9.69E-05 | Mtmr12 | myotubularir | Plasma Mem |
| ENSMUSG00000014959 | 5.77E-05 | 9.81E-05 | Gorasp2 | golgi reassen | Cytoplasm |
| ENSMUSG00000033909 | 5.77E-05 | 9.83E-05 | Usp36 | ubiquitin spe | Nucleus |
| ENSMUSG00000070407 | 5.93E-05 | 0.00010082 | Hs3st3b1 | heparan sulfa | Cytoplasm |
| ENSMUSG00000031838 | 5.94E-05 | 0.00010102 | Ifi30 | IFI30, lysosor | Cytoplasm |
| ENSMUSG00000035949 | 5.96E-05 | 0.00010131 | Fbxw2 | F-box and W | Cytoplasm |
| ENSMUSG00000004934 | 5.96E-05 | 0.00010132 | Pias4 | protein inhib | Nucleus |
| ENSMUSG00000069844 | 5.98E-05 | 0.00010161 | Sco1 | SCO1, cytoch | Cytoplasm |
| ENSMUSG00000021236 | 6.10E-05 | 0.0001036 | Entpd5 | ectonucleosi | Cytoplasm |
| ENSMUSG00000020541 | 6.16E-05 | 0.0001047 | Tom1l1 | target of myb | Cytoplasm |
| ENSMUSG00000006777 | 6.20E-05 | 0.00010531 | Krt23 | keratin 23 | Cytoplasm |
| ENSMUSG00000007989 | 6.21E-05 | 0.00010544 | Fzd3 | frizzled class | Plasma Mem |
| ENSMUSG00000025607 | 6.25E-05 | 0.00010609 | Copg2 | coatomer pro | Cytoplasm |
| ENSMUSG00000039018 | 6.28E-05 | 0.00010659 | Mtg1 | mitochondria | Cytoplasm |
| ENSMUSG00000047084 | 6.30E-05 | 0.00010689 | Ngrn | neugrin, neu | Nucleus |
| ENSMUSG00000004947 | 6.36E-05 | 0.00010793 | Dtx2 | deltex E3 ubi | Nucleus |
| ENSMUSG00000028464 | 6.39E-05 | 0.00010841 | Tpm2 | tropomyosin | Cytoplasm |
| ENSMUSG00000057101 | 6.39E-05 | 0.00010841 | Zfp180 | zinc finger pr | Nucleus |
| ENSMUSG00000096215 | 6.42E-05 | 0.00010891 | Smim22 | small integra | Other |
| ENSMUSG00000024566 | 6.49E-05 | 0.00010998 | Atp9b | ATPase phos | Cytoplasm |
| ENSMUSG00000035266 | 6.50E-05 | 0.0001101 | Helq | helicase, POL | Nucleus |
| ENSMUSG00000062012 | 6.50E-05 | 0.0001101 | Zfp13 | zinc finger pr | Nucleus |
| ENSMUSG00000041733 | 6.50E-05 | 0.00011015 | Coq5 | coenzyme Q5 | Cytoplasm |
| ENSMUSG00000030532 | 6.52E-05 | 0.00011042 | Hddc3 | HD domain c | Other |
| ENSMUSG00000038417 | 6.53E-05 | 0.00011065 | Fig4 | FIG4 phosph | Cytoplasm |
| ENSMUSG00000039354 | 6.55E-05 | 0.00011088 | Smarcal1 | SWI/SNF rela | Nucleus |
| ENSMUSG00000030917 | 6.59E-05 | 0.00011164 | Tmem159 | transmembra | Other |
| ENSMUSG00000050627 | 6.60E-05 | 0.00011169 | Gpd1l | glycerol-3-p | Cytoplasm |
| ENSMUSG00000027016 | 6.60E-05 | 0.00011169 | Zfp385b | zinc finger pr | Nucleus |
| ENSMUSG00000019826 | 6.61E-05 | 0.0001119 | Zbtb24 | zinc finger ar | Nucleus |
| ENSMUSG00000030469 | 6.62E-05 | 0.00011208 | Zfp719 | zinc finger pr | Other |
| ENSMUSG00000002384 | 6.70E-05 | 0.00011326 | Bmp8b | bone morpho | Extracellular S |
| ENSMUSG00000040859 | 6.77E-05 | 0.00011449 | Bsdc1 | BSD domain | Other |
| ENSMUSG00000029629 | 6.81E-05 | 0.00011519 | Phf14 | PHD finger p | Nucleus |
| ENSMUSG00000022890 | 6.81E-05 | 0.0001152 | Atp5j | ATP synthase | Cytoplasm |

| | | | | | |
|---|---|---|---|---|---|
| ENSMUSG00000050199 | 6.94E-05 | 0.00011723 | Lgr4 | leucine rich r | Plasma Mem |
| ENSMUSG00000005078 | 6.95E-05 | 0.00011739 | Jkamp | JNK1/MAPK8 | Cytoplasm |
| ENSMUSG00000037251 | 7.01E-05 | 0.00011847 | Pomk | protein-O-m | Cytoplasm |
| ENSMUSG00000046897 | 7.02E-05 | 0.0001185 | Zfp740 | zinc finger pr | Other |
| ENSMUSG00000027242 | 7.05E-05 | 0.00011904 | Wdr76 | WD repeat d | Nucleus |
| ENSMUSG00000052584 | 7.05E-05 | 0.00011906 | Serp2 | stress associa | Other |
| ENSMUSG00000019795 | 7.06E-05 | 0.00011916 | Pcmt1 | protein-L-isc | Cytoplasm |
| ENSMUSG00000026779 | 7.07E-05 | 0.00011927 | Mastl | microtubule a | Cytoplasm |
| ENSMUSG00000045667 | 7.09E-05 | 0.0001197 | Smtnl2 | smoothelin li | Other |
| ENSMUSG00000026663 | 7.14E-05 | 0.00012038 | Atf6 | activating tra | Cytoplasm |
| ENSMUSG00000032679 | 7.22E-05 | 0.00012176 | Cd59a | CD59a antige | Plasma Mem |
| ENSMUSG00000073131 | 7.29E-05 | 0.00012297 | Vma21 | VMA21 vacuo | Cytoplasm |
| ENSMUSG00000037475 | 7.37E-05 | 0.00012421 | Thoc2 | THO complex | Nucleus |
| ENSMUSG00000037795 | 7.38E-05 | 0.0001243 | N4bp2 | NEDD4 bindi | Cytoplasm |
| ENSMUSG00000004462 | 7.38E-05 | 0.00012434 | Tbccd1 | TBCC domair | Cytoplasm |
| ENSMUSG00000026278 | 7.42E-05 | 0.00012495 | Bok | BOK, BCL2 fa | Cytoplasm |
| ENSMUSG00000027551 | 7.43E-05 | 0.00012516 | Zfp64 | ZFP64 zinc fi | Nucleus |
| ENSMUSG00000022570 | 7.46E-05 | 0.00012562 | Tsta3 | tissue specific | Plasma Mem |
| ENSMUSG00000046550 | 7.52E-05 | 0.00012655 | Spin2c | spindlin fami | Other |
| ENSMUSG00000022200 | 7.62E-05 | 0.00012831 | Golph3 | golgi phosph | Cytoplasm |
| ENSMUSG00000069874 | 7.71E-05 | 0.00012977 | Irgm2 | immunity rela | Cytoplasm |
| ENSMUSG00000021796 | 7.78E-05 | 0.00013089 | Bmpr1a | bone morpho | Plasma Mem |
| ENSMUSG00000033916 | 7.86E-05 | 0.00013217 | Chmp2a | charged mult | Cytoplasm |
| ENSMUSG00000021427 | 7.91E-05 | 0.00013304 | Ssr1 | signal sequer | Cytoplasm |
| ENSMUSG00000027706 | 7.92E-05 | 0.00013318 | Sec62 | SEC62 homo | Cytoplasm |
| ENSMUSG00000029319 | 7.93E-05 | 0.00013334 | Coq2 | coenzyme Q2 | Cytoplasm |
| ENSMUSG00000037253 | 7.96E-05 | 0.0001338 | Mex3c | mex-3 RNA b | Cytoplasm |
| ENSMUSG00000000378 | 8.03E-05 | 0.00013498 | Ccm2 | CCM2 scaffol | Cytoplasm |
| ENSMUSG00000071180 | 8.10E-05 | 0.00013614 | Smim15 | small integra | Other |
| ENSMUSG00000045248 | 8.11E-05 | 0.00013629 | Med26 | mediator cor | Nucleus |
| ENSMUSG00000073400 | 8.17E-05 | 0.00013712 | Trim10 | tripartite mot | Cytoplasm |
| ENSMUSG00000018347 | 8.28E-05 | 0.00013894 | Zkscan6 | zinc finger pr | Nucleus |
| ENSMUSG00000031153 | 8.33E-05 | 0.00013975 | Gripap1 | GRIP1 associa | Plasma Mem |
| ENSMUSG00000040525 | 8.48E-05 | 0.00014224 | Cblc | Cbl proto-or | Nucleus |
| ENSMUSG00000028743 | 8.48E-05 | 0.00014225 | Akr7a5 | aldo-keto rec | Cytoplasm |
| ENSMUSG00000042419 | 8.49E-05 | 0.00014236 | Nfkbil1 | NFKB inhibito | Nucleus |
| ENSMUSG00000024563 | 8.58E-05 | 0.00014388 | Smad2 | SMAD family | Nucleus |
| ENSMUSG00000049922 | 8.61E-05 | 0.00014439 | Slc35c1 | solute carrier | Cytoplasm |
| ENSMUSG00000004931 | 8.65E-05 | 0.00014503 | Apba3 | amyloid beta | Cytoplasm |
| ENSMUSG00000021044 | 8.66E-05 | 0.0001452 | Adck1 | aarF domain | Cytoplasm |
| ENSMUSG00000073421 | 8.67E-05 | 0.00014526 | H2-Ab1 | major histocc | Plasma Mem |
| ENSMUSG00000027678 | 8.69E-05 | 0.00014564 | Ncoa3 | nuclear recep | Nucleus |
| ENSMUSG00000042670 | 8.71E-05 | 0.00014595 | Immp1l | inner mitoch | Cytoplasm |
| ENSMUSG00000061689 | 8.72E-05 | 0.00014599 | Dlgap4 | DLG associat | Plasma Mem |
| ENSMUSG00000063179 | 8.79E-05 | 0.00014713 | Pstk | phosphosery | Cytoplasm |
| ENSMUSG00000029020 | 8.83E-05 | 0.00014775 | Mfn2 | mitofusin 2 | Cytoplasm |
| ENSMUSG00000039357 | 8.83E-05 | 0.00014787 | Fut11 | fucosyltransf | Cytoplasm |
| ENSMUSG00000037816 | 8.94E-05 | 0.00014961 | Fbxw17 | F-box and W | Other |
| ENSMUSG00000032044 | 8.95E-05 | 0.00014967 | Rpusd4 | RNA pseudo | Cytoplasm |
| ENSMUSG00000034620 | 8.97E-05 | 0.00014997 | Tmem5 | ribitol xylosyl | Plasma Mem |
| ENSMUSG00000015536 | 9.01E-05 | 0.00015066 | Mocs2 | molybdenum | Cytoplasm |
| ENSMUSG00000028982 | 9.05E-05 | 0.0001513 | Slc25a33 | solute carrier | Cytoplasm |
| ENSMUSG00000020029 | 9.06E-05 | 0.00015137 | Nudt4 | nudix hydrola | Cytoplasm |
| ENSMUSG00000039242 | 9.18E-05 | 0.00015341 | B3galnt2 | beta-1,3-N-a | Cytoplasm |
| ENSMUSG00000021981 | 9.21E-05 | 0.00015391 | Cab39l | calcium bind | Cytoplasm |
| ENSMUSG00000039952 | 9.27E-05 | 0.00015492 | Dag1 | dystroglycan | Plasma Mem |
| ENSMUSG00000022864 | 9.28E-05 | 0.00015505 | D16Ertd472e | chromosome | Other |
| ENSMUSG00000025505 | 9.29E-05 | 0.00015513 | Tmem80 | transmembra | Other |

| | | | | | |
|---|---|---|---|---|---|
| ENSMUSG00000028453 | 9.31E-05 | 0.00015546 | Fancg | FA complem | Nucleus |
| ENSMUSG00000032030 | 9.34E-05 | 0.00015593 | Cul5 | cullin 5 | Nucleus |
| ENSMUSG00000070730 | 9.41E-05 | 0.00015698 | Rmdn3 | regulator of | Cytoplasm |
| ENSMUSG00000051705 | 9.46E-05 | 0.00015781 | Senp8 | SUMO peptic | Cytoplasm |
| ENSMUSG00000002845 | 9.50E-05 | 0.00015847 | Tmem39a | transmembra | Other |
| ENSMUSG00000015027 | 9.51E-05 | 0.00015855 | Galns | galactosamin | Cytoplasm |
| ENSMUSG00000000441 | 9.64E-05 | 0.00016068 | Raf1 | Raf-1 proto- | Cytoplasm |
| ENSMUSG00000027602 | 9.67E-05 | 0.00016122 | Map1lc3a | microtubule | Cytoplasm |
| ENSMUSG00000023046 | 9.68E-05 | 0.00016141 | Igfbp6 | insulin like gr | Extracellular S |
| ENSMUSG00000024260 | 9.70E-05 | 0.00016165 | Sap130 | Sin3A associa | Nucleus |
| ENSMUSG00000004446 | 9.78E-05 | 0.00016286 | Bid | BH3 interacti | Cytoplasm |
| ENSMUSG00000035228 | 9.92E-05 | 0.00016517 | Ccdc106 | coiled-coil d | Extracellular S |
| ENSMUSG00000000934 | 9.99E-05 | 0.00016637 | Top1mt | DNA topoiso | Cytoplasm |
| ENSMUSG00000045211 | 1.00E-04 | 0.00016638 | Nudt18 | nudix hydrola | Cytoplasm |
| ENSMUSG00000035529 | 1.00E-04 | 0.00016639 | Prdm4 | PR/SET doma | Nucleus |
| ENSMUSG00000039745 | 0.00010026 | 0.00016684 | Htatip2 | HIV-1 Tat int | Nucleus |
| ENSMUSG00000031161 | 0.00010034 | 0.00016693 | Hdac6 | histone deac | Nucleus |
| ENSMUSG00000021759 | 0.00010111 | 0.00016818 | Ppap2a | phospholipid | Plasma Mem |
| ENSMUSG00000031660 | 0.00010117 | 0.00016821 | Brd7 | bromodomai | Nucleus |
| ENSMUSG00000047242 | 0.00010115 | 0.00016821 | Taf9b | TATA-box bi | Nucleus |
| ENSMUSG00000048970 | 0.00010192 | 0.00016943 | C1galt1c1 | C1GALT1 spe | Cytoplasm |
| ENSMUSG00000019763 | 0.00010225 | 0.00016994 | Rmnd1 | required for | Cytoplasm |
| ENSMUSG00000031365 | 0.00010258 | 0.00017045 | Zfp275 | zinc finger pr | Nucleus |
| ENSMUSG00000032997 | 0.00010323 | 0.0001715 | Chpf | chondroitin p | Cytoplasm |
| ENSMUSG00000002233 | 0.00010383 | 0.00017242 | Rhoc | ras homolog | Plasma Mem |
| ENSMUSG00000021468 | 0.00010412 | 0.00017287 | Sptlc1 | serine palmit | Cytoplasm |
| ENSMUSG00000050229 | 0.00010419 | 0.00017296 | Pigm | phosphatidyl | Cytoplasm |
| ENSMUSG00000029146 | 0.00010491 | 0.00017412 | Snx17 | sorting nexin | Cytoplasm |
| ENSMUSG00000020176 | 0.00010507 | 0.00017431 | Grb10 | growth facto | Cytoplasm |
| ENSMUSG00000031155 | 0.00010559 | 0.00017514 | Pim2 | Pim-2 proto- | Nucleus |
| ENSMUSG00000055900 | 0.00010585 | 0.00017554 | Tmem69 | transmembra | Other |
| ENSMUSG00000074925 | 0.00010593 | 0.00017564 | Ptar1 | protein preny | Other |
| ENSMUSG00000028622 | 0.00010616 | 0.00017598 | Mrpl37 | mitochondria | Cytoplasm |
| ENSMUSG00000061046 | 0.00010653 | 0.00017656 | Haghl | hydroxyacylg | Other |
| ENSMUSG00000047407 | 0.00010691 | 0.00017715 | Tgif1 | TGFB induce | Nucleus |
| ENSMUSG00000027710 | 0.00010742 | 0.00017796 | Acad9 | acyl-CoA del | Cytoplasm |
| ENSMUSG00000059820 | 0.00010766 | 0.00017833 | AU019823 | NKAP domai | Other |
| ENSMUSG00000031229 | 0.00010833 | 0.00017939 | Atrx | ATRX, chrom | Nucleus |
| ENSMUSG00000031447 | 0.00010893 | 0.00018036 | Lamp1 | lysosomal as | Plasma Mem |
| ENSMUSG00000027397 | 0.0001094 | 0.00018109 | Slc20a1 | solute carrier | Plasma Mem |
| ENSMUSG00000027540 | 0.0001096 | 0.00018139 | Ptpn1 | protein tyros | Cytoplasm |
| ENSMUSG00000019761 | 0.00010979 | 0.00018167 | Krt10 | keratin 10 | Cytoplasm |
| ENSMUSG00000029470 | 0.00011006 | 0.00018209 | P2rx4 | purinergic re | Plasma Mem |
| ENSMUSG00000005373 | 0.0001101 | 0.00018212 | Mlxipl | MLX interacti | Nucleus |
| ENSMUSG00000032534 | 0.00011014 | 0.00018214 | Cep63 | centrosomal | Cytoplasm |
| ENSMUSG00000037710 | 0.00011149 | 0.00018433 | Cisd1 | CDGSH iron | Cytoplasm |
| ENSMUSG00000026439 | 0.00011227 | 0.00018559 | Rbbp5 | RB binding p | Nucleus |
| ENSMUSG00000028696 | 0.00011239 | 0.00018575 | Ipp | intracisternal | Cytoplasm |
| ENSMUSG00000038141 | 0.0001128 | 0.00018639 | Tmem181a | transmembra | Other |
| ENSMUSG00000042699 | 0.00011322 | 0.00018706 | Dhx9 | DExH-box he | Nucleus |
| ENSMUSG00000024535 | 0.00011325 | 0.00018706 | Snx24 | sorting nexin | Other |
| ENSMUSG00000035495 | 0.00011337 | 0.00018721 | Tstd2 | thiosulfate su | Other |
| ENSMUSG00000034640 | 0.00011389 | 0.00018805 | Tiparp | TCDD inducil | Nucleus |
| ENSMUSG00000032905 | 0.00011416 | 0.00018846 | Atg12 | autophagy re | Cytoplasm |
| ENSMUSG00000016256 | 0.00011427 | 0.00018859 | Ctsz | cathepsin Z | Cytoplasm |
| ENSMUSG00000042834 | 0.00011446 | 0.00018888 | Nrep | neuronal reg | Cytoplasm |
| ENSMUSG00000044018 | 0.00011515 | 0.00018997 | Mrpl50 | mitochondria | Cytoplasm |
| ENSMUSG00000029234 | 0.00011517 | 0.00018997 | Tmem165 | transmembra | Plasma Mem |

| | | | | | |
|---|---|---|---|---|---|
| ENSMUSG00000038515 | 0.00011657 | 0.00019224 | Grtp1 | growth horm | Other |
| ENSMUSG00000043461 | 0.0001178 | 0.00019423 | Sptssb | serine palmit | Cytoplasm |
| ENSMUSG00000059816 | 0.00011913 | 0.00019638 | Fam159a | shisa like 2A | Other |
| ENSMUSG00000028405 | 0.00012006 | 0.00019788 | Aco1 | aconitase 1 | Cytoplasm |
| ENSMUSG00000024654 | 0.00012012 | 0.00019794 | Asrgl1 | asparaginase | Cytoplasm |
| ENSMUSG00000093803 | 0.00012111 | 0.00019953 | Ppp2r3d | protein phos | Other |
| ENSMUSG00000049076 | 0.00012127 | 0.00019974 | Acap2 | ArfGAP with | Nucleus |
| ENSMUSG00000030779 | 0.00012134 | 0.00019983 | Rbbp6 | RB binding p | Nucleus |
| ENSMUSG00000035142 | 0.00012195 | 0.00020078 | Nubpl | nucleotide bi | Cytoplasm |
| ENSMUSG00000031478 | 0.00012239 | 0.00020147 | Nek3 | NIMA relatec | Nucleus |
| ENSMUSG00000039852 | 0.00012293 | 0.00020233 | Rere | arginine-glut | Nucleus |
| ENSMUSG00000050854 | 0.0001242 | 0.00020438 | Tmem125 | transmembra | Other |
| ENSMUSG00000025354 | 0.00012456 | 0.00020493 | Dnajc14 | DnaJ heat sh | Cytoplasm |
| ENSMUSG00000034926 | 0.00012487 | 0.00020539 | Dhcr24 | 24-dehydroc | Cytoplasm |
| ENSMUSG00000052852 | 0.00012513 | 0.00020579 | Reep1 | receptor acce | Cytoplasm |
| ENSMUSG00000041915 | 0.00012589 | 0.00020699 | Ammecr1l | AMMECR1 lik | Other |
| ENSMUSG00000020385 | 0.00012628 | 0.00020759 | Clk4 | CDC like kina | Nucleus |
| ENSMUSG00000002812 | 0.00012638 | 0.00020772 | Flii | FLII, actin ren | Nucleus |
| ENSMUSG00000047213 | 0.00012694 | 0.0002086 | Ythdf3 | YTH N6-met | Cytoplasm |
| ENSMUSG00000051978 | 0.0001276 | 0.00020964 | Erich1 | glutamate ric | Other |
| ENSMUSG00000020362 | 0.00012803 | 0.00021026 | Cnot6 | CCR4-NOT t | Nucleus |
| ENSMUSG00000035620 | 0.00012803 | 0.00021026 | Ric8b | RIC8 guanine | Cytoplasm |
| ENSMUSG00000034453 | 0.0001281 | 0.00021034 | Polr3b | RNA polymer | Nucleus |
| ENSMUSG00000030281 | 0.00012979 | 0.00021306 | Il17rc | interleukin 17 | Plasma Mem |
| ENSMUSG00000025027 | 0.00012988 | 0.00021317 | Xpnpep1 | X-prolyl amir | Cytoplasm |
| ENSMUSG00000041935 | 0.0001302 | 0.00021365 | AW549877 | chromosome | Other |
| ENSMUSG00000050989 | 0.00013024 | 0.00021367 | Sepn1 | selenoprotein | Cytoplasm |
| ENSMUSG00000000078 | 0.00013157 | 0.00021582 | Klf6 | Kruppel like f | Nucleus |
| ENSMUSG00000061232 | 0.00013172 | 0.00021602 | H2-K1 | major histocc | Plasma Mem |
| ENSMUSG00000034744 | 0.00013221 | 0.00021678 | Nagk | N-acetylgluc | Cytoplasm |
| ENSMUSG00000059263 | 0.00013231 | 0.0002169 | Usp47 | ubiquitin spe | Cytoplasm |
| ENSMUSG00000003662 | 0.00013328 | 0.00021844 | Ciao1 | cytosolic iron | Nucleus |
| ENSMUSG00000006289 | 0.00013365 | 0.00021902 | Osgep | O-sialoglyco | Nucleus |
| ENSMUSG00000020038 | 0.00013437 | 0.00022016 | Cry1 | cryptochrom | Nucleus |
| ENSMUSG00000032673 | 0.00013502 | 0.00022116 | Prorsd1 | prolyl-tRNA s | Other |
| ENSMUSG00000035842 | 0.00013512 | 0.00022129 | Ddx11 | DEAD/H-box | Cytoplasm |
| ENSMUSG00000031930 | 0.00013539 | 0.00022168 | Wwp2 | WW domain | Cytoplasm |
| ENSMUSG00000024190 | 0.00013703 | 0.00022433 | Dusp1 | dual specifici | Nucleus |
| ENSMUSG00000026753 | 0.00013716 | 0.00022449 | Ppp6c | protein phos | Nucleus |
| ENSMUSG00000042515 | 0.00013746 | 0.00022495 | Mum1l1 | MUM1 like 1 | Cytoplasm |
| ENSMUSG00000010048 | 0.00013819 | 0.00022609 | Ifrd2 | interferon rel | Nucleus |
| ENSMUSG00000075000 | 0.00013878 | 0.00022702 | Nrbf2 | nuclear recep | Nucleus |
| ENSMUSG00000042323 | 0.00014055 | 0.00022987 | Pbrm1 | polybromo 1 | Nucleus |
| ENSMUSG00000021906 | 0.00014167 | 0.00023166 | Oxnad1 | oxidoreducta | Cytoplasm |
| ENSMUSG00000032010 | 0.0001421 | 0.00023231 | Usp2 | ubiquitin spe | Cytoplasm |
| ENSMUSG00000041236 | 0.00014222 | 0.00023245 | Vps41 | VPS41, HOPS | Cytoplasm |
| ENSMUSG00000027341 | 0.00014237 | 0.00023265 | Tmem230 | transmembra | Cytoplasm |
| ENSMUSG00000025103 | 0.00014285 | 0.00023339 | Btbd1 | BTB domain | Cytoplasm |
| ENSMUSG00000000976 | 0.00014323 | 0.00023398 | Heatr6 | HEAT repeat | Other |
| ENSMUSG00000054716 | 0.0001438 | 0.00023486 | Zfp771 | zinc finger pr | Nucleus |
| ENSMUSG00000006641 | 0.00014428 | 0.0002356 | Slc5a6 | solute carrier | Plasma Mem |
| ENSMUSG00000040048 | 0.00014449 | 0.00023589 | Ndufb10 | NADH:ubiqu | Cytoplasm |
| ENSMUSG00000036983 | 0.00014494 | 0.00023657 | Tfb1m | transcription | Cytoplasm |
| ENSMUSG00000012422 | 0.00014516 | 0.00023689 | Tmem167 | transmembra | Other |
| ENSMUSG00000029752 | 0.00014599 | 0.00023815 | Asns | asparagine sy | Cytoplasm |
| ENSMUSG00000071350 | 0.00014654 | 0.000239 | Setdb2 | SET domain b | Nucleus |
| ENSMUSG00000025235 | 0.00014697 | 0.00023966 | Bbs4 | Bardet-Biedl | Cytoplasm |
| ENSMUSG00000040280 | 0.00014823 | 0.00024162 | Ndufa4l2 | NDUFA4, mit | Other |

| | | | | | |
|---|---|---|---|---|---|
| ENSMUSG00000026775 | 0.00014821 | 0.00024162 | Yme1l1 | YME1 like 1 A | Cytoplasm |
| ENSMUSG00000054408 | 0.00014842 | 0.00024189 | Spcs3 | signal peptid | Cytoplasm |
| ENSMUSG00000035242 | 0.00014917 | 0.00024305 | Oaz1 | ornithine dec | Cytoplasm |
| ENSMUSG00000044030 | 0.00014932 | 0.00024324 | Irf2bp1 | interferon re | Nucleus |
| ENSMUSG00000075254 | 0.00014974 | 0.00024389 | Heg1 | heart develo | Plasma Mem |
| ENSMUSG00000025868 | 0.0001498 | 0.00024394 | Higd2a | HIG1 hypoxia | Cytoplasm |
| ENSMUSG00000028383 | 0.00015271 | 0.00024862 | Hsdl2 | hydroxystero | Cytoplasm |
| ENSMUSG00000059183 | 0.0001528 | 0.00024873 | Mtfmt | mitochondria | Cytoplasm |
| ENSMUSG00000070394 | 0.00015458 | 0.00025152 | Tmem256 | transmembra | Cytoplasm |
| ENSMUSG00000006360 | 0.00015525 | 0.00025256 | Crip1 | cysteine rich | Cytoplasm |
| ENSMUSG00000040713 | 0.00015617 | 0.00025401 | Creg1 | cellular repre | Nucleus |
| ENSMUSG00000032612 | 0.00015641 | 0.00025435 | Usp4 | ubiquitin spe | Nucleus |
| ENSMUSG00000027502 | 0.00015663 | 0.00025465 | Rtfdc1 | replication te | Other |
| ENSMUSG00000027828 | 0.00015757 | 0.00025613 | Ssr3 | signal seque | Cytoplasm |
| ENSMUSG00000051378 | 0.00015819 | 0.0002571 | Kif18b | kinesin family | Cytoplasm |
| ENSMUSG00000038425 | 0.00015828 | 0.00025719 | Poli | DNA polyme | Nucleus |
| ENSMUSG00000038451 | 0.00015879 | 0.00025797 | Spsb2 | splA/ryanodi | Cytoplasm |
| ENSMUSG00000021411 | 0.00015929 | 0.00025873 | Pxdc1 | PX domain c | Other |
| ENSMUSG00000057836 | 0.00015977 | 0.00025946 | Xlr3a | X-linked lym | Other |
| ENSMUSG00000030105 | 0.00016244 | 0.00026374 | Arl8b | ADP ribosyla | Plasma Mem |
| ENSMUSG00000021264 | 0.00016309 | 0.00026474 | Yy1 | YY1 transcrip | Nucleus |
| ENSMUSG00000032431 | 0.0001634 | 0.00026519 | Crtap | cartilage ass | Extracellular S |
| ENSMUSG00000022412 | 0.00016397 | 0.00026607 | Mief1 | mitochondria | Cytoplasm |
| ENSMUSG00000021385 | 0.00016454 | 0.00026694 | Ippk | inositol-pent | Cytoplasm |
| ENSMUSG00000025950 | 0.00016488 | 0.00026744 | Idh1 | isocitrate de | Cytoplasm |
| ENSMUSG00000030688 | 0.00016537 | 0.00026819 | Stard10 | StAR related | Cytoplasm |
| ENSMUSG00000027963 | 0.00016542 | 0.00026821 | Extl2 | exostosin like | Cytoplasm |
| ENSMUSG00000022216 | 0.00016546 | 0.00026823 | Psme1 | proteasome a | Cytoplasm |
| ENSMUSG00000045100 | 0.00016689 | 0.00027044 | Slc25a26 | solute carrier | Cytoplasm |
| ENSMUSG00000060279 | 0.00016707 | 0.00027068 | Ap2a1 | adaptor relat | Cytoplasm |
| ENSMUSG00000049421 | 0.00016783 | 0.00027186 | Zfp260 | zinc finger pr | Nucleus |
| ENSMUSG00000029416 | 0.0001687 | 0.0002732 | Slc15a4 | solute carrier | Cytoplasm |
| ENSMUSG00000031993 | 0.00016997 | 0.00027522 | Snx19 | sorting nexin | Cytoplasm |
| ENSMUSG00000058979 | 0.00017139 | 0.00027741 | Cecr5 | haloacid deh | Cytoplasm |
| ENSMUSG00000025142 | 0.00017343 | 0.00028065 | Aspscr1 | ASPSCR1, UB | Cytoplasm |
| ENSMUSG00000039199 | 0.00017354 | 0.00028077 | Zdhhc1 | zinc finger D | Nucleus |
| ENSMUSG00000021643 | 0.00017404 | 0.00028153 | Serf1 | small EDRK-r | Other |
| ENSMUSG00000018669 | 0.00017453 | 0.00028227 | Cdk5rap3 | CDK5 regulat | Nucleus |
| ENSMUSG00000064254 | 0.00017495 | 0.00028289 | Ethe1 | ETHE1, persu | Cytoplasm |
| ENSMUSG00000059486 | 0.00017513 | 0.00028313 | Kbtbd2 | kelch repeat | Other |
| ENSMUSG00000062190 | 0.00017657 | 0.0002854 | Lancl2 | LanC like 2 | Plasma Mem |
| ENSMUSG00000004846 | 0.00017683 | 0.00028576 | Plod3 | procollagen- | Cytoplasm |
| ENSMUSG00000052253 | 0.00017729 | 0.00028645 | Zfp622 | zinc finger pr | Nucleus |
| ENSMUSG00000050565 | 0.00017776 | 0.00028715 | Tor1aip2 | torsin 1A inte | Cytoplasm |
| ENSMUSG00000019872 | 0.00017906 | 0.00028915 | Smpdl3a | sphingomyel | Extracellular S |
| ENSMUSG00000028466 | 0.00017949 | 0.00028978 | Creb3 | cAMP respor | Nucleus |
| ENSMUSG00000031979 | 0.00018018 | 0.00029084 | Cog2 | component c | Cytoplasm |
| ENSMUSG00000028684 | 0.0001806 | 0.00029146 | Urod | uroporphyrin | Cytoplasm |
| ENSMUSG00000056938 | 0.00018124 | 0.00029244 | Acbd4 | acyl-CoA bin | Other |
| ENSMUSG00000001018 | 0.00018225 | 0.000294 | Snapin | SNAP associa | Plasma Mem |
| ENSMUSG00000054136 | 0.00018304 | 0.00029523 | Adm2 | adrenomedu | Extracellular S |
| ENSMUSG00000026887 | 0.00018316 | 0.00029531 | Mrrf | mitochondria | Cytoplasm |
| ENSMUSG00000029627 | 0.00018315 | 0.00029531 | Zkscan14 | zinc finger pr | Nucleus |
| ENSMUSG00000030720 | 0.00018575 | 0.00029943 | Cln3 | CLN3, batten | Cytoplasm |
| ENSMUSG00000038615 | 0.0001873 | 0.00030186 | Nfe2l1 | nuclear facto | Nucleus |
| ENSMUSG00000025220 | 0.00018773 | 0.00030249 | Mgea5 | O-GlcNAcase | Cytoplasm |
| ENSMUSG00000041625 | 0.00018794 | 0.00030277 | Ggact | gamma-gluta | Cytoplasm |
| ENSMUSG00000047221 | 0.00019094 | 0.00030755 | Fam185a | family with se | Cytoplasm |

| | | | | | |
|---|---|---|---|---|---|
| ENSMUSG00000043019 | 0.00019314 | 0.00031098 | Edem3 | ER degradati | Cytoplasm |
| ENSMUSG00000020305 | 0.00019404 | 0.00031236 | Asb3 | ankyrin repea | Cytoplasm |
| ENSMUSG00000047539 | 0.00019491 | 0.0003137 | Fbxo28 | F-box protei | Nucleus |
| ENSMUSG00000032624 | 0.00019502 | 0.00031381 | Eml4 | echinoderm | Cytoplasm |
| ENSMUSG00000024914 | 0.00019545 | 0.00031445 | Drap1 | DR1 associat | Nucleus |
| ENSMUSG00000033972 | 0.00019604 | 0.00031534 | Zfp944 | zinc finger pr | Other |
| ENSMUSG00000058147 | 0.00019781 | 0.00031812 | Xlr3c | X-linked lym | Other |
| ENSMUSG00000029156 | 0.00019972 | 0.00032106 | Sgcb | sarcoglycan | Plasma Mem |
| ENSMUSG00000037966 | 0.0002035 | 0.00032708 | Ninj1 | ninjurin 1 | Plasma Mem |
| ENSMUSG00000087370 | 0.00020376 | 0.00032743 | Tmem170b | transmembra | Plasma Mem |
| ENSMUSG00000020459 | 0.00020397 | 0.00032764 | Mtif2 | mitochondria | Cytoplasm |
| ENSMUSG00000020134 | 0.00020394 | 0.00032764 | Peli1 | pellino E3 ub | Cytoplasm |
| ENSMUSG00000021764 | 0.0002046 | 0.0003286 | Ndufs4 | NADH:ubiqu | Cytoplasm |
| ENSMUSG00000044442 | 0.00020583 | 0.00033051 | N6amt1 | N-6 adenine | Cytoplasm |
| ENSMUSG00000025616 | 0.00020834 | 0.00033446 | Usp16 | ubiquitin spe | Cytoplasm |
| ENSMUSG00000006589 | 0.00020848 | 0.00033464 | Aprt | adenine phos | Cytoplasm |
| ENSMUSG00000015095 | 0.00020854 | 0.00033466 | Fbxw5 | F-box and W | Cytoplasm |
| ENSMUSG00000094441 | 0.00021055 | 0.00033782 | Zfp955a | zinc finger pr | Other |
| ENSMUSG00000023025 | 0.00021153 | 0.00033932 | Larp4 | La ribonucle | Cytoplasm |
| ENSMUSG00000002808 | 0.00021248 | 0.00034079 | Epdr1 | ependymin r | Nucleus |
| ENSMUSG00000024943 | 0.00021279 | 0.00034123 | Smc5 | structural ma | Nucleus |
| ENSMUSG00000071454 | 0.00021367 | 0.00034257 | Dtnb | dystrobrevin | Plasma Mem |
| ENSMUSG00000055228 | 0.00021468 | 0.00034411 | Zfp935 | expressed se | Other |
| ENSMUSG00000018541 | 0.00021555 | 0.00034545 | Cwc25 | CWC25 splic | Nucleus |
| ENSMUSG00000053113 | 0.00021732 | 0.00034821 | Socs3 | suppressor o | Cytoplasm |
| ENSMUSG00000030579 | 0.00021861 | 0.00035022 | Tyrobp | TYRO protei | Plasma Mem |
| ENSMUSG00000038957 | 0.00021971 | 0.0003519 | Edc3 | enhancer of | Cytoplasm |
| ENSMUSG00000034789 | 0.00022008 | 0.00035243 | Rab24 | RAB24, mem | Cytoplasm |
| ENSMUSG00000034308 | 0.00022088 | 0.00035365 | Sdr42e1 | short chain d | Other |
| ENSMUSG00000025533 | 0.00022216 | 0.00035562 | Asl | argininosucci | Cytoplasm |
| ENSMUSG00000017670 | 0.00022371 | 0.00035805 | Elmo2 | engulfment a | Cytoplasm |
| ENSMUSG00000040891 | 0.00022376 | 0.00035806 | Foxa3 | forkhead box | Nucleus |
| ENSMUSG00000034781 | 0.00022479 | 0.00035962 | Gna11 | G protein sul | Plasma Mem |
| ENSMUSG00000031104 | 0.00022523 | 0.00036027 | Rab33a | RAB33A, mer | Cytoplasm |
| ENSMUSG00000028889 | 0.00022668 | 0.00036252 | Yrdc | yrdC N6-thre | Other |
| ENSMUSG00000031451 | 0.00022793 | 0.00036444 | Gas6 | growth arres | Extracellular S |
| ENSMUSG00000019518 | 0.00022875 | 0.00036568 | Ap4m1 | adaptor relat | Cytoplasm |
| ENSMUSG00000064068 | 0.00022999 | 0.00036758 | Mtx1 | metaxin 1 | Cytoplasm |
| ENSMUSG00000026466 | 0.00023012 | 0.00036766 | Tor1aip1 | torsin 1A inte | Nucleus |
| ENSMUSG00000092074 | 0.00023164 | 0.00036995 | Dynlt1a | dynein light | Cytoplasm |
| ENSMUSG00000029086 | 0.00023293 | 0.00037187 | Prom1 | prominin 1 | Plasma Mem |
| ENSMUSG00000027673 | 0.0002333 | 0.00037238 | Ndufb5 | NADH:ubiqu | Cytoplasm |
| ENSMUSG00000028277 | 0.00023429 | 0.0003739 | Ube2j1 | ubiquitin cor | Cytoplasm |
| ENSMUSG00000040795 | 0.00023565 | 0.00037599 | Iqcc | IQ motif cont | Other |
| ENSMUSG00000032526 | 0.00023774 | 0.00037925 | Deb1 | SS18 like 2 | Other |
| ENSMUSG00000031987 | 0.00023789 | 0.00037941 | Egln1 | egl-9 family | Cytoplasm |
| ENSMUSG00000037103 | 0.00023878 | 0.00038077 | Dcaf15 | DDB1 and CU | Other |
| ENSMUSG00000028582 | 0.00024067 | 0.00038371 | Cc2d1b | coiled-coil ar | Nucleus |
| ENSMUSG00000074916 | 0.00024113 | 0.00038437 | Chst14 | carbohydrate | Cytoplasm |
| ENSMUSG00000037210 | 0.00024164 | 0.00038511 | Fam193a | family with se | Other |
| ENSMUSG00000022141 | 0.00024185 | 0.00038537 | Nipbl | NIPBL, cohes | Nucleus |
| ENSMUSG00000003031 | 0.00024304 | 0.00038718 | Cdkn1b | cyclin depen | Nucleus |
| ENSMUSG00000029465 | 0.00024314 | 0.00038726 | Arpc3 | actin related | Cytoplasm |
| ENSMUSG00000025939 | 0.00024482 | 0.00038988 | Ube2w | ubiquitin cor | Nucleus |
| ENSMUSG00000002496 | 0.00024594 | 0.00039158 | Tsc2 | TSC complex | Cytoplasm |
| ENSMUSG00000003037 | 0.00024848 | 0.00039554 | Rab8a | RAB8A, mem | Plasma Mem |
| ENSMUSG00000042292 | 0.00025003 | 0.00039794 | Mkl1 | myocardin re | Nucleus |
| ENSMUSG00000027465 | 0.00025062 | 0.0003988 | Tbc1d20 | TBC1 domair | Nucleus |

| | | | | | |
|---|---|---|---|---|---|
| ENSMUSG00000049305 | 0.00025085 | 0.00039909 | Ccdc71 | coiled-coil do | Other |
| ENSMUSG00000048920 | 0.00025106 | 0.00039935 | Fkrp | fukutin relate | Cytoplasm |
| ENSMUSG00000058900 | 0.00025154 | 0.00040003 | Rsl1 | regulator of s | Other |
| ENSMUSG00000066838 | 0.00025172 | 0.00040024 | Zfp772 | zinc finger pr | Other |
| ENSMUSG00000036580 | 0.00025264 | 0.00040164 | Spg20 | spastic parap | Plasma Mem |
| ENSMUSG00000021910 | 0.00025446 | 0.00040445 | Nisch | nischarin | Cytoplasm |
| ENSMUSG00000028716 | 0.00025452 | 0.00040447 | Pdzk1ip1 | PDZK1 intera | Extracellular S |
| ENSMUSG00000027770 | 0.00025464 | 0.00040458 | Dhx36 | DEAH-box he | Cytoplasm |
| ENSMUSG00000025428 | 0.00025571 | 0.0004062 | Atp5a1 | ATP synthase | Cytoplasm |
| ENSMUSG00000055371 | 0.00025626 | 0.000407 | Stam2 | signal transdu | Cytoplasm |
| ENSMUSG00000060121 | 0.00025646 | 0.00040723 | Gemin2 | gem nuclear | Nucleus |
| ENSMUSG00000001472 | 0.00025656 | 0.00040731 | Tcf25 | transcription | Nucleus |
| ENSMUSG00000020123 | 0.0002583 | 0.00041 | Avpr1a | arginine vaso | Plasma Mem |
| ENSMUSG00000062234 | 0.00025902 | 0.00041097 | Gak | cyclin G asso | Nucleus |
| ENSMUSG00000029486 | 0.00025906 | 0.00041097 | Mrpl1 | mitochondria | Cytoplasm |
| ENSMUSG00000039849 | 0.00025903 | 0.00041097 | Pcif1 | PDX1 C-term | Nucleus |
| ENSMUSG00000041420 | 0.00025995 | 0.00041231 | Meis3 | Meis homeob | Nucleus |
| ENSMUSG00000030659 | 0.00026143 | 0.00041457 | Nucb2 | nucleobindin | Nucleus |
| ENSMUSG00000047264 | 0.00026175 | 0.000415 | Zfp358 | zinc finger pr | Nucleus |
| ENSMUSG00000041096 | 0.00026248 | 0.000416 | Tspyl2 | TSPY like 2 | Nucleus |
| ENSMUSG00000024614 | 0.00026266 | 0.00041621 | Tmx3 | thioredoxin r | Cytoplasm |
| ENSMUSG00000044881 | 0.00026341 | 0.00041731 | Coa4 | cytochrome c | Cytoplasm |
| ENSMUSG00000040124 | 0.00026438 | 0.00041878 | Gorab | golgin, RAB6 | Cytoplasm |
| ENSMUSG00000036291 | 0.00026554 | 0.00042053 | Ap5m1 | adaptor relat | Cytoplasm |
| ENSMUSG00000034412 | 0.00026584 | 0.00042092 | Tbc1d10a | TBC1 domain | Plasma Mem |
| ENSMUSG00000023232 | 0.0002684 | 0.00042481 | Serinc2 | serine incorp | Plasma Mem |
| ENSMUSG00000047866 | 0.00027285 | 0.00043177 | Lonp2 | lon peptidase | Cytoplasm |
| ENSMUSG00000018733 | 0.00027441 | 0.00043416 | Pex12 | peroxisomal | Cytoplasm |
| ENSMUSG00000030754 | 0.00027897 | 0.00044128 | Copb1 | coatomer pro | Cytoplasm |
| ENSMUSG00000025217 | 0.00027914 | 0.00044148 | Btrc | beta-transdu | Cytoplasm |
| ENSMUSG00000005505 | 0.00027953 | 0.00044201 | Kbtbd4 | kelch repeat | Other |
| ENSMUSG00000040434 | 0.0002796 | 0.00044203 | Gyltl1b | LARGE xylosy | Cytoplasm |
| ENSMUSG00000026554 | 0.00028418 | 0.00044919 | Dcaf8 | DDB1 and CU | Nucleus |
| ENSMUSG00000022194 | 0.00028463 | 0.00044981 | Pabpn1 | poly(A) bindi | Nucleus |
| ENSMUSG00000074063 | 0.00028937 | 0.00045721 | Osgin1 | oxidative stre | Other |
| ENSMUSG00000024304 | 0.00028983 | 0.00045786 | Cdh2 | cadherin 2 | Plasma Mem |
| ENSMUSG00000024900 | 0.0002904 | 0.00045866 | Cpt1a | carnitine palr | Cytoplasm |
| ENSMUSG00000005873 | 0.00029099 | 0.00045952 | Reep5 | receptor acce | Extracellular S |
| ENSMUSG00000024691 | 0.0002931 | 0.0004627 | Fam111a | family with se | Nucleus |
| ENSMUSG00000024712 | 0.00029312 | 0.0004627 | Rfk | riboflavin kin | Cytoplasm |
| ENSMUSG00000028849 | 0.00029353 | 0.00046326 | Map7d1 | MAP7 domai | Cytoplasm |
| ENSMUSG00000031207 | 0.00029697 | 0.00046861 | Msn | moesin | Plasma Mem |
| ENSMUSG00000007411 | 0.0002974 | 0.0004692 | Mark3 | microtubule a | Cytoplasm |
| ENSMUSG00000038010 | 0.00029792 | 0.00046992 | Ccdc138 | coiled-coil do | Other |
| ENSMUSG00000030979 | 0.00029808 | 0.00047008 | Uros | uroporphyrin | Cytoplasm |
| ENSMUSG00000068206 | 0.00030012 | 0.00047321 | Pick1 | protein intera | Cytoplasm |
| ENSMUSG00000060708 | 0.00030338 | 0.00047826 | Bloc1s4 | biogenesis of | Cytoplasm |
| ENSMUSG00000057554 | 0.00030372 | 0.00047871 | Lgals8 | galectin 8 | Extracellular S |
| ENSMUSG00000073409 | 0.00030449 | 0.00047984 | H2-Q6 | major histoco | Plasma Mem |
| ENSMUSG00000070803 | 0.00030612 | 0.0004823 | Cited4 | Cbp/p300 int | Nucleus |
| ENSMUSG00000083282 | 0.00030944 | 0.00048744 | Ctsf | cathepsin F | Cytoplasm |
| ENSMUSG00000069049 | 0.00031013 | 0.00048844 | Eif2s3y | eukaryotic tra | Other |
| ENSMUSG00000020623 | 0.00031118 | 0.00049 | Map2k6 | mitogen-acti | Cytoplasm |
| ENSMUSG00000032788 | 0.0003116 | 0.00049057 | Pdxk | pyridoxal kin | Cytoplasm |
| ENSMUSG00000028757 | 0.00031296 | 0.00049261 | Ddost | dolichyl-diph | Cytoplasm |
| ENSMUSG00000031446 | 0.00031329 | 0.00049304 | Cul4a | cullin 4A | Nucleus |
| ENSMUSG00000032305 | 0.00031348 | 0.00049325 | Fam219b | family with se | Other |
| ENSMUSG00000021127 | 0.00031548 | 0.0004963 | Zfp36l1 | ZFP36 ring fi | Nucleus |

| | | | | | |
|---|---|---|---|---|---|
| ENSMUSG00000051517 | 0.00031749 | 0.00049928 | Arhgef39 | Rho guanine | Plasma Mem |
| ENSMUSG00000027198 | 0.00031765 | 0.00049944 | Ext2 | exostosin gly | Cytoplasm |
| ENSMUSG00000026895 | 0.00031939 | 0.00050207 | Ndufa8 | NADH:ubiqu | Cytoplasm |
| ENSMUSG00000002661 | 0.00032249 | 0.00050684 | Alkbh7 | alkB homolo | Cytoplasm |
| ENSMUSG00000022601 | 0.0003259 | 0.00051211 | Zbtb11 | zinc finger ar | Nucleus |
| ENSMUSG00000034126 | 0.00032597 | 0.00051212 | Pomt2 | protein O-m | Cytoplasm |
| ENSMUSG00000002393 | 0.00032678 | 0.0005133 | Nr2f6 | nuclear recep | Nucleus |
| ENSMUSG00000054733 | 0.00032686 | 0.00051333 | Msra | methionine s | Cytoplasm |
| ENSMUSG00000022982 | 0.00032744 | 0.00051415 | Sod1 | superoxide d | Cytoplasm |
| ENSMUSG00000041354 | 0.00032794 | 0.00051482 | Rgl2 | ral guanine n | Cytoplasm |
| ENSMUSG00000025878 | 0.00032815 | 0.00051506 | Uimc1 | ubiquitin inte | Nucleus |
| ENSMUSG00000096255 | 0.00032991 | 0.00051772 | Dynlt1b | dynein light | Cytoplasm |
| ENSMUSG00000034610 | 0.00033031 | 0.00051826 | Zcchc11 | terminal urid | Nucleus |
| ENSMUSG00000028391 | 0.0003325 | 0.0005216 | Wdr31 | WD repeat d | Other |
| ENSMUSG00000030753 | 0.00033397 | 0.0005238 | Prkrir | THAP domai | Nucleus |
| ENSMUSG00000027751 | 0.00033415 | 0.00052398 | Supt20 | suppressor o | Nucleus |
| ENSMUSG00000022131 | 0.00033528 | 0.00052566 | Gpr180 | G protein-co | Cytoplasm |
| ENSMUSG00000002320 | 0.00033574 | 0.00052628 | Tm9sf1 | transmembra | Plasma Mem |
| ENSMUSG00000039285 | 0.0003362 | 0.0005269 | Azi2 | 5-azacytidine | Cytoplasm |
| ENSMUSG00000090523 | 0.00033859 | 0.00053044 | Gypc | glycophorin | Plasma Mem |
| ENSMUSG00000025728 | 0.00033854 | 0.00053044 | Pigq | phosphatidyl | Cytoplasm |
| ENSMUSG00000024976 | 0.00034264 | 0.00053668 | Shoc2 | SHOC2, leuci | Cytoplasm |
| ENSMUSG00000032293 | 0.00034357 | 0.00053793 | Ireb2 | iron responsi | Cytoplasm |
| ENSMUSG00000002064 | 0.00034354 | 0.00053793 | Sdf2 | stromal cell c | Extracellular S |
| ENSMUSG00000078773 | 0.00034479 | 0.00053974 | Rad54b | RAD54 homo | Nucleus |
| ENSMUSG00000040659 | 0.0003458 | 0.00054123 | Efhd2 | EF-hand don | Other |
| ENSMUSG00000019210 | 0.00034774 | 0.00054416 | Atp6v1e1 | ATPase H+ tr | Cytoplasm |
| ENSMUSG00000017801 | 0.00034835 | 0.00054501 | Mlx | MLX, MAX di | Nucleus |
| ENSMUSG00000038095 | 0.00034912 | 0.00054612 | Sbno1 | strawberry no | Other |
| ENSMUSG00000026718 | 0.00035019 | 0.00054758 | Stam | signal transd | Cytoplasm |
| ENSMUSG00000036402 | 0.00035453 | 0.00055427 | Gng12 | G protein sub | Plasma Mem |
| ENSMUSG00000024191 | 0.00035505 | 0.00055498 | Bnip1 | BCL2 interact | Cytoplasm |
| ENSMUSG00000031431 | 0.00035677 | 0.00055755 | Tsc22d3 | TSC22 doma | Nucleus |
| ENSMUSG00000024277 | 0.0003571 | 0.00055797 | Mapre2 | microtubule- | Cytoplasm |
| ENSMUSG00000046229 | 0.00036024 | 0.00056276 | Scand1 | SCAN domai | Nucleus |
| ENSMUSG00000039646 | 0.00036125 | 0.00056424 | Vasn | vasorin | Plasma Mem |
| ENSMUSG00000019774 | 0.00036251 | 0.0005661 | Mtrf1l | mitochondria | Cytoplasm |
| ENSMUSG00000047153 | 0.00036303 | 0.0005668 | Khnyn | KH and NYN | Other |
| ENSMUSG00000021661 | 0.00036678 | 0.00057256 | Ankra2 | ankyrin repea | Cytoplasm |
| ENSMUSG00000078994 | 0.00036725 | 0.00057318 | Zfp429 | regulator of s | Other |
| ENSMUSG00000066456 | 0.00036858 | 0.00057515 | Hmgn3 | high mobility | Nucleus |
| ENSMUSG00000054400 | 0.00037 | 0.00057725 | Cklf | chemokine lil | Extracellular S |
| ENSMUSG00000024069 | 0.00037069 | 0.00057822 | Slc30a6 | solute carrier | Cytoplasm |
| ENSMUSG00000039197 | 0.0003717 | 0.00057963 | Adk | adenosine ki | Nucleus |
| ENSMUSG00000070583 | 0.00037173 | 0.00057963 | Fv1 | Friend virus s | Cytoplasm |
| ENSMUSG00000024843 | 0.00037406 | 0.00058315 | Chka | choline kinas | Cytoplasm |
| ENSMUSG00000063698 | 0.00037471 | 0.00058406 | Sfxn4 | sideroflexin 4 | Cytoplasm |
| ENSMUSG00000022965 | 0.0003752 | 0.0005847 | Ifngr2 | interferon ga | Plasma Mem |
| ENSMUSG00000061619 | 0.00037756 | 0.00058827 | Vma21-ps | VMA21 vacu | Other |
| ENSMUSG00000021012 | 0.00037772 | 0.00058842 | Zc3h14 | zinc finger C | Nucleus |
| ENSMUSG00000057469 | 0.00037841 | 0.00058938 | E2f6 | E2F transcrip | Nucleus |
| ENSMUSG00000028886 | 0.00037926 | 0.00059059 | Eya3 | EYA transcrip | Nucleus |
| ENSMUSG00000058729 | 0.00037955 | 0.00059093 | Lin9 | lin-9 DREAM | Nucleus |
| ENSMUSG00000026701 | 0.00038052 | 0.00059233 | Prdx6 | peroxiredoxi | Cytoplasm |
| ENSMUSG00000046441 | 0.00038239 | 0.00059514 | Cmtr2 | cap methyltra | Other |
| ENSMUSG00000021280 | 0.00038342 | 0.00059663 | Exoc3l4 | exocyst comp | Other |
| ENSMUSG00000073563 | 0.00038821 | 0.00060396 | Csnk1g3 | casein kinase | Cytoplasm |
| ENSMUSG00000053510 | 0.0003901 | 0.00060679 | Nrd1 | nardilysin co | Cytoplasm |

| | | | | | | |
|---|---|---|---|---|---|---|
| ENSMUSG00000004069 | 0.00039853 | 0.00061967 | Dnaja3 | | DnaJ heat sh | Cytoplasm |
| ENSMUSG00000032312 | 0.00039906 | 0.00062038 | Csk | | C-terminal Si | Cytoplasm |
| ENSMUSG00000046798 | 0.00040074 | 0.00062287 | Cldn12 | | claudin 12 | Plasma Mem |
| ENSMUSG00000019173 | 0.00040099 | 0.00062314 | Rab5c | | RAB5C, mem | Cytoplasm |
| ENSMUSG00000004394 | 0.00040236 | 0.00062516 | Tmed4 | | transmembra | Cytoplasm |
| ENSMUSG00000049536 | 0.00040828 | 0.00063424 | Tceal1 | | transcription | Nucleus |
| ENSMUSG00000030835 | 0.00040891 | 0.00063509 | Nomo1 | | NODAL mod | Plasma Mem |
| ENSMUSG00000035547 | 0.00041056 | 0.00063754 | Capn5 | | calpain 5 | Cytoplasm |
| ENSMUSG00000074261 | 0.00041095 | 0.00063802 | Erich4 | | glutamate ric | Other |
| ENSMUSG00000001131 | 0.00041202 | 0.00063957 | Timp1 | | TIMP metallc | Extracellular S |
| ENSMUSG00000024002 | 0.0004126 | 0.00064035 | Brd4 | | bromodomai | Nucleus |
| ENSMUSG00000010453 | 0.00041407 | 0.0006425 | Kansl3 | | KAT8 regulat | Nucleus |
| ENSMUSG00000027618 | 0.00041549 | 0.0006446 | Nfs1 | | NFS1, cystein | Cytoplasm |
| ENSMUSG00000028334 | 0.00041606 | 0.00064535 | Nans | | N-acetylneur | Cytoplasm |
| ENSMUSG00000045503 | 0.00041621 | 0.00064546 | Sys1 | | SYS1, golgi tr | Cytoplasm |
| ENSMUSG00000024197 | 0.00041738 | 0.00064717 | Plin3 | | perilipin 3 | Cytoplasm |
| ENSMUSG00000054226 | 0.00041803 | 0.00064805 | Tprkb | | TP53RK bind | Cytoplasm |
| ENSMUSG00000061887 | 0.00042015 | 0.00065121 | Ssbp3 | | single strand | Nucleus |
| ENSMUSG00000056501 | 0.00042032 | 0.00065136 | Cebpb | | CCAAT enhai | Nucleus |
| ENSMUSG00000060733 | 0.00042523 | 0.00065884 | Ipmk | | inositol polyp | Nucleus |
| ENSMUSG00000037706 | 0.00042585 | 0.00065968 | Cd81 | | CD81 molecu | Plasma Mem |
| ENSMUSG00000005951 | 0.00042608 | 0.00065992 | Shpk | | sedoheptulol | Cytoplasm |
| ENSMUSG00000022008 | 0.00042902 | 0.00066435 | Gpalpp1 | | GPALPP mot | Other |
| ENSMUSG00000020326 | 0.00042935 | 0.00066472 | Ccng1 | | cyclin G1 | Nucleus |
| ENSMUSG00000047036 | 0.0004318 | 0.0006684 | Zfp445 | | zinc finger pr | Nucleus |
| ENSMUSG00000001151 | 0.00043293 | 0.00067001 | Pcnt | | pericentrin (k | Cytoplasm |
| ENSMUSG00000013833 | 0.00043529 | 0.00067355 | Med16 | | mediator cor | Nucleus |
| ENSMUSG00000043648 | 0.0004362 | 0.00067484 | Pld6 | | phospholipas | Cytoplasm |
| ENSMUSG00000032806 | 0.00043741 | 0.00067657 | Slc10a3 | | solute carrier | Plasma Mem |
| ENSMUSG00000027002 | 0.00044386 | 0.00068642 | Nckap1 | | NCK associat | Plasma Mem |
| ENSMUSG00000038102 | 0.00044405 | 0.0006866 | Trappc11 | | trafficking pr | Cytoplasm |
| ENSMUSG00000030109 | 0.00044974 | 0.00069513 | Slc6a12 | | solute carrier | Plasma Mem |
| ENSMUSG00000055312 | 0.00045021 | 0.00069573 | Them7 | | thioesterase s | Cytoplasm |
| ENSMUSG00000002058 | 0.00045034 | 0.00069579 | Unc119 | | unc-119 lipic | Cytoplasm |
| ENSMUSG00000034854 | 0.00045135 | 0.00069723 | Mfsd12 | | major facilita | Cytoplasm |
| ENSMUSG00000021895 | 0.00045263 | 0.00069907 | Arhgef3 | | Rho guanine | Cytoplasm |
| ENSMUSG00000024477 | 0.00045616 | 0.0007044 | Pggt1b | | protein gerar | Cytoplasm |
| ENSMUSG00000026880 | 0.0004589 | 0.00070849 | Stom | | stomatin | Plasma Mem |
| ENSMUSG00000022176 | 0.00046068 | 0.0007111 | Rem2 | | RRAD and GI | Plasma Mem |
| ENSMUSG00000021210 | 0.00046308 | 0.00071469 | Akr1c6 | | aldo-keto red | Cytoplasm |
| ENSMUSG00000040621 | 0.00046373 | 0.00071556 | Gemin8 | | gem nuclear | Cytoplasm |
| ENSMUSG00000017697 | 0.00046561 | 0.00071832 | Ada | | adenosine de | Cytoplasm |
| ENSMUSG00000038518 | 0.00046637 | 0.00071937 | Jarid2 | | jumonji and / | Nucleus |
| ENSMUSG00000068762 | 0.00046845 | 0.00072243 | Gstm6 | | glutathione S | Cytoplasm |
| ENSMUSG00000032014 | 0.00046892 | 0.00072302 | Oaf | | out at first hc | Cytoplasm |
| ENSMUSG00000032497 | 0.00046916 | 0.00072326 | Lrrfip2 | | leucine rich r | Other |
| ENSMUSG00000039236 | 0.00046977 | 0.00072407 | Isg20 | | interferon sti | Nucleus |
| ENSMUSG00000033216 | 0.00047175 | 0.00072699 | Eefsec | | eukaryotic el | Cytoplasm |
| ENSMUSG00000040331 | 0.00047227 | 0.00072765 | Nsmce4a | | NSE4 homolc | Nucleus |
| ENSMUSG00000036529 | 0.00047387 | 0.00072998 | Sbf1 | | SET binding f | Plasma Mem |
| ENSMUSG00000025283 | 0.00047499 | 0.00073157 | Sat1 | | spermidine/s | Cytoplasm |
| ENSMUSG00000025726 | 0.00047841 | 0.00073669 | Slc28a1 | | solute carrier | Plasma Mem |
| ENSMUSG00000032119 | 0.0004848 | 0.00074639 | Hinfp | | histone H4 tr | Nucleus |
| ENSMUSG00000005881 | 0.00048606 | 0.0007482 | Ergic3 | | ERGIC and gc | Cytoplasm |
| ENSMUSG00000001100 | 0.00048961 | 0.00075353 | Poldip2 | | DNA polyme | Cytoplasm |
| ENSMUSG00000024960 | 0.00048989 | 0.00075381 | Plcb3 | | phospholipas | Cytoplasm |
| ENSMUSG00000006782 | 0.00049406 | 0.00076009 | Cnp | | 2',3'-cyclic nu | Cytoplasm |
| ENSMUSG00000047368 | 0.00049491 | 0.00076125 | Abhd17b | | abhydrolase | Plasma Mem |

| | | | | | |
|---|---|---|---|---|---|
| ENSMUSG00000028126 | 0.00049555 | 0.00076211 | Pip5k1a | phosphatidyl | Cytoplasm |
| ENSMUSG00000039367 | 0.00049604 | 0.00076272 | Sec24c | SEC24 homo | Cytoplasm |
| ENSMUSG00000032727 | 0.00049864 | 0.00076657 | Mier3 | MIER family r | Nucleus |
| ENSMUSG00000046519 | 0.00049912 | 0.00076717 | Golph3l | golgi phosph | Cytoplasm |
| ENSMUSG00000019718 | 0.00050013 | 0.00076858 | L3hypdh | trans-L-3-hy | Other |
| ENSMUSG00000005142 | 0.00050105 | 0.00076985 | Man2b1 | mannosidase | Cytoplasm |
| ENSMUSG00000022529 | 0.00050157 | 0.0007705 | Zfp263 | zinc finger pr | Nucleus |
| ENSMUSG00000020340 | 0.00050561 | 0.00077657 | Cyfip2 | cytoplasmic F | Cytoplasm |
| ENSMUSG00000028194 | 0.00050597 | 0.00077698 | Ddah1 | dimethylargi | Cytoplasm |
| ENSMUSG00000026504 | 0.0005063 | 0.00077734 | Sdccag8 | serologically | Cytoplasm |
| ENSMUSG00000034714 | 0.00050805 | 0.00077989 | Ttyh2 | tweety family | Other |
| ENSMUSG00000047843 | 0.00050907 | 0.00078131 | Bri3 | brain protein | Cytoplasm |
| ENSMUSG00000019943 | 0.00051072 | 0.00078369 | Atp2b1 | ATPase plasn | Plasma Mem |
| ENSMUSG00000041684 | 0.00051762 | 0.00079413 | Bivm | basic, immun | Extracellular S |
| ENSMUSG00000025139 | 0.00052178 | 0.00080037 | Tollip | toll interactin | Cytoplasm |
| ENSMUSG00000040373 | 0.00052194 | 0.00080046 | Cacng5 | calcium volta | Plasma Mem |
| ENSMUSG00000004849 | 0.00052233 | 0.00080091 | Ap1s1 | adaptor relat | Cytoplasm |
| ENSMUSG00000031171 | 0.00052798 | 0.00080943 | Ftsj1 | FtsJ RNA met | Cytoplasm |
| ENSMUSG00000028825 | 0.00053005 | 0.00081245 | Rhd | Rh blood grc | Plasma Mem |
| ENSMUSG00000027286 | 0.00053082 | 0.00081348 | Lrrc57 | leucine rich r | Cytoplasm |
| ENSMUSG00000021203 | 0.00053498 | 0.0008197 | Otub2 | OTU deubiqu | Nucleus |
| ENSMUSG00000045817 | 0.00053737 | 0.00082322 | Zfp36l2 | ZFP36 ring fi | Nucleus |
| ENSMUSG00000017405 | 0.00053927 | 0.00082598 | Nek8 | NIMA relatec | Nucleus |
| ENSMUSG00000037916 | 0.00053993 | 0.00082682 | Ndufv1 | NADH:ubiqu | Cytoplasm |
| ENSMUSG00000035845 | 0.00054154 | 0.00082915 | Alg12 | ALG12, alpha | Cytoplasm |
| ENSMUSG00000020258 | 0.00054224 | 0.00083005 | Glyctk | glycerate kin | Cytoplasm |
| ENSMUSG00000020832 | 0.00054261 | 0.00083047 | Eral1 | Era like 12S r | Cytoplasm |
| ENSMUSG00000021716 | 0.00054296 | 0.00083086 | Srek1ip1 | splicing regu | Other |
| ENSMUSG00000042726 | 0.00054431 | 0.00083277 | Trafd1 | TRAF-type zi | Other |
| ENSMUSG00000024909 | 0.00055214 | 0.00084459 | Efemp2 | EGF containir | Extracellular S |
| ENSMUSG00000054611 | 0.00055416 | 0.00084753 | Kdm2a | lysine demetl | Nucleus |
| ENSMUSG00000000159 | 0.00055753 | 0.00085252 | Igsf5 | immunoglob | Plasma Mem |
| ENSMUSG00000062312 | 0.00056163 | 0.00085863 | Erbb2 | erb-b2 recep | Plasma Mem |
| ENSMUSG00000048921 | 0.00056217 | 0.0008593 | Zfp689 | zinc finger pr | Nucleus |
| ENSMUSG00000021610 | 0.00056249 | 0.00085964 | Clptm1l | CLPTM1 like | Other |
| ENSMUSG00000025959 | 0.00056428 | 0.0008622 | Klf7 | Kruppel like f | Nucleus |
| ENSMUSG00000036975 | 0.00056853 | 0.00086854 | Tmem177 | transmembra | Cytoplasm |
| ENSMUSG00000004040 | 0.00057405 | 0.00087682 | Stat3 | signal transd | Nucleus |
| ENSMUSG00000032235 | 0.00057421 | 0.0008769 | Ice2 | interactor of | Nucleus |
| ENSMUSG00000042178 | 0.00058124 | 0.00088743 | Armc5 | armadillo rep | Cytoplasm |
| ENSMUSG00000028952 | 0.00058132 | 0.00088743 | Zbtb48 | zinc finger ar | Nucleus |
| ENSMUSG00000055148 | 0.00058751 | 0.00089672 | Klf2 | Kruppel like f | Nucleus |
| ENSMUSG00000027900 | 0.00058766 | 0.00089677 | Dram2 | DNA damage | Cytoplasm |
| ENSMUSG00000029556 | 0.00058844 | 0.0008978 | Hnf1a | HNF1 homec | Nucleus |
| ENSMUSG00000035310 | 0.00059041 | 0.00090064 | Lin54 | lin-54 DREAN | Nucleus |
| ENSMUSG00000053964 | 0.00059451 | 0.00090673 | Lgals4 | galectin 4 | Extracellular S |
| ENSMUSG00000012117 | 0.00059633 | 0.00090917 | Dhdds | dehydrodolic | Cytoplasm |
| ENSMUSG00000059316 | 0.00059629 | 0.00090917 | Slc27a4 | solute carrier | Plasma Mem |
| ENSMUSG00000040820 | 0.00059719 | 0.00091032 | Hlcs | holocarboxyl | Cytoplasm |
| ENSMUSG00000049300 | 0.00059895 | 0.00091283 | Prmt6 | protein argin | Nucleus |
| ENSMUSG00000043789 | 0.00060682 | 0.00092466 | Vwce | von Willebrai | Cytoplasm |
| ENSMUSG00000037190 | 0.00060924 | 0.00092817 | Cyb561d2 | cytochrome l | Other |
| ENSMUSG00000059518 | 0.00061122 | 0.00093102 | Znhit1 | zinc finger H | Nucleus |
| ENSMUSG00000058620 | 0.00061313 | 0.00093375 | Adra2b | adrenocepto | Plasma Mem |
| ENSMUSG00000031634 | 0.00061443 | 0.00093557 | Ufsp2 | UFM1 specifi | Other |
| ENSMUSG00000025369 | 0.0006178 | 0.00094052 | Smarcc2 | SWI/SNF rela | Nucleus |
| ENSMUSG00000033720 | 0.00061882 | 0.0009419 | Sfxn5 | sideroflexin 5 | Cytoplasm |
| ENSMUSG00000024906 | 0.00062527 | 0.00095137 | Mus81 | MUS81 struc | Nucleus |

| | | | | | |
|---|---|---|---|---|---|
| ENSMUSG00000025153 | 0.00062891 | 0.00095673 | Fasn | fatty acid syn | Cytoplasm |
| ENSMUSG00000051495 | 0.00062914 | 0.00095691 | Irf2bp2 | interferon re( | Nucleus |
| ENSMUSG00000041870 | 0.00063065 | 0.00095903 | Ankrd13a | ankyrin repea | Plasma Mem |
| ENSMUSG00000034424 | 0.00063098 | 0.00095936 | Gcsh | glycine cleav | Cytoplasm |
| ENSMUSG00000047909 | 0.00063263 | 0.00096168 | Ankrd16 | ankyrin repea | Other |
| ENSMUSG00000056153 | 0.00063326 | 0.00096247 | Socs6 | suppressor o | Cytoplasm |
| ENSMUSG00000052406 | 0.0006336 | 0.00096281 | Rexo4 | REX4 homolo | Nucleus |
| ENSMUSG00000031266 | 0.00063412 | 0.00096343 | Gla | galactosidase | Cytoplasm |
| ENSMUSG00000026810 | 0.00063871 | 0.00097022 | Dpm2 | dolichyl-pho | Cytoplasm |
| ENSMUSG00000010045 | 0.00064375 | 0.0009777 | Tmem115 | transmembra | Other |
| ENSMUSG00000058239 | 0.00064676 | 0.0009821 | Usf2 | upstream tra | Nucleus |
| ENSMUSG00000010057 | 0.00064716 | 0.00098251 | Nprl2 | NPR2 like, GA | Cytoplasm |
| ENSMUSG00000049878 | 0.0006474 | 0.00098271 | Rlf | rearranged L | Nucleus |
| ENSMUSG00000022711 | 0.00064965 | 0.00098593 | Pmm2 | phosphomar | Cytoplasm |
| ENSMUSG00000032042 | 0.0006552 | 0.00099418 | Srpr | SRP receptor | Cytoplasm |
| ENSMUSG00000073424 | 0.00066113 | 0.001003 | Cyp4f15 | cytochrome I | Cytoplasm |
| ENSMUSG00000027739 | 0.00066235 | 0.00100466 | Rab33b | RAB33B, mer | Cytoplasm |
| ENSMUSG00000066043 | 0.00066477 | 0.00100814 | Phactr4 | phosphatase | Plasma Mem |
| ENSMUSG00000030301 | 0.00066867 | 0.00101388 | Ccdc91 | coiled-coil d | Cytoplasm |
| ENSMUSG00000030400 | 0.00066926 | 0.00101459 | Ercc2 | ERCC excisio | Nucleus |
| ENSMUSG00000036214 | 0.00067372 | 0.00102116 | Znrd1as | zinc ribbon d | Other |
| ENSMUSG00000022186 | 0.00067389 | 0.00102124 | Oxct1 | 3-oxoacid Co | Cytoplasm |
| ENSMUSG00000062859 | 0.00067818 | 0.00102754 | Tcp11 | t-complex 11 | Cytoplasm |
| ENSMUSG00000034639 | 0.00068168 | 0.00103266 | Setmar | SET domain v | Nucleus |
| ENSMUSG00000006931 | 0.00068558 | 0.00103838 | P3h4 | prolyl 3-hydr | Nucleus |
| ENSMUSG00000056459 | 0.00068969 | 0.00104442 | Zbtb25 | zinc finger ar | Nucleus |
| ENSMUSG00000020087 | 0.00069051 | 0.00104547 | Tysnd1 | trypsin doma | Cytoplasm |
| ENSMUSG00000027099 | 0.00069344 | 0.00104971 | Mtx2 | metaxin 2 | Cytoplasm |
| ENSMUSG00000027199 | 0.00069439 | 0.00105096 | Gatm | glycine amidi | Cytoplasm |
| ENSMUSG00000028057 | 0.00069579 | 0.00105278 | Rit1 | Ras like witho | Plasma Mem |
| ENSMUSG00000062116 | 0.00069585 | 0.00105278 | Zfp954 | zinc finger pr | Other |
| ENSMUSG00000035836 | 0.00070112 | 0.00106056 | Ugt2b1 | UDP glucuro | Cytoplasm |
| ENSMUSG00000050697 | 0.00070361 | 0.00106413 | Prkaa1 | protein kinas | Cytoplasm |
| ENSMUSG00000075700 | 0.00070714 | 0.00106927 | Selt | selenoprotei | Cytoplasm |
| ENSMUSG00000039759 | 0.00070739 | 0.00106946 | Thap3 | THAP domai | Nucleus |
| ENSMUSG00000020153 | 0.00070793 | 0.00107007 | Ndufs7 | NADH:ubiqu | Cytoplasm |
| ENSMUSG00000055660 | 0.00071174 | 0.00107564 | Mettl4 | methyltransfe | Other |
| ENSMUSG00000040648 | 0.00071202 | 0.00107587 | Ppip5k2 | diphosphoin | Cytoplasm |
| ENSMUSG00000025377 | 0.00071986 | 0.00108753 | Enthd2 | TEPSIN, adap | Nucleus |
| ENSMUSG00000017837 | 0.00072493 | 0.00109479 | Nkiras2 | NFKB inhibito | Cytoplasm |
| ENSMUSG00000047454 | 0.00072604 | 0.00109604 | Gphn | gephyrin | Plasma Mem |
| ENSMUSG00000028894 | 0.00072601 | 0.00109604 | Inpp5b | inositol polyp | Plasma Mem |
| ENSMUSG00000052738 | 0.00072616 | 0.00109604 | Suclg1 | succinate-Co | Cytoplasm |
| ENSMUSG00000028132 | 0.00072633 | 0.00109609 | Tmem56 | transmembra | Other |
| ENSMUSG00000055609 | 0.00073133 | 0.00110345 | Hba-x | hemoglobin | Cytoplasm |
| ENSMUSG00000042349 | 0.00073321 | 0.00110608 | Ikbke | inhibitor of n | Cytoplasm |
| ENSMUSG00000045268 | 0.00073527 | 0.00110878 | Zfp691 | zinc finger pr | Nucleus |
| ENSMUSG00000022377 | 0.00073866 | 0.00111369 | Asap1 | ArfGAP with | Plasma Mem |
| ENSMUSG00000030760 | 0.00074132 | 0.0011175 | Acer3 | alkaline ceran | Cytoplasm |
| ENSMUSG00000051007 | 0.00074549 | 0.00112359 | Pddc1 | glutamine an | Cytoplasm |
| ENSMUSG00000031072 | 0.00074799 | 0.00112715 | Oraov1 | LTO1, ABCE1 | Other |
| ENSMUSG00000025495 | 0.00075227 | 0.00113339 | Ptdss2 | phosphatidyl | Cytoplasm |
| ENSMUSG00000039967 | 0.00075544 | 0.00113796 | Zfp292 | zinc finger pr | Nucleus |
| ENSMUSG00000081534 | 0.00076181 | 0.00114735 | Slc48a1 | solute carrier | Cytoplasm |
| ENSMUSG00000041769 | 0.00076293 | 0.00114882 | Ppp2r2d | protein phos | Nucleus |
| ENSMUSG00000033430 | 0.00076614 | 0.00115345 | Terf2ip | TERF2 intera | Nucleus |
| ENSMUSG00000031007 | 0.0007663 | 0.00115348 | Atp6ap2 | ATPase H+ tr | Cytoplasm |
| ENSMUSG00000021400 | 0.00077718 | 0.00116964 | Wrnip1 | Werner helica | Nucleus |

| | | | | | |
|---|---|---|---|---|---|
| ENSMUSG00000010067 | 0.00077828 | 0.00117108 | Rassf1 | Ras associatio | Nucleus |
| ENSMUSG00000001467 | 0.00078043 | 0.00117411 | Cyp51 | cytochrome I | Cytoplasm |
| ENSMUSG00000027411 | 0.00078239 | 0.00117685 | Vps16 | VPS16, CORV | Cytoplasm |
| ENSMUSG00000038046 | 0.00078891 | 0.00118644 | Rnmtl1 | mitochondria | Cytoplasm |
| ENSMUSG00000037643 | 0.00079033 | 0.00118814 | Prkci | protein kinas | Cytoplasm |
| ENSMUSG00000048478 | 0.00079092 | 0.00118881 | Spata33 | spermatogen | Other |
| ENSMUSG00000026991 | 0.00079476 | 0.00119437 | Pkp4 | plakophilin 4 | Plasma Mem |
| ENSMUSG00000035112 | 0.00079743 | 0.00119816 | Wnk4 | WNK lysine c | Plasma Mem |
| ENSMUSG00000019889 | 0.00080157 | 0.00120417 | Ptprk | protein tyros | Plasma Mem |
| ENSMUSG00000025091 | 0.00080314 | 0.00120631 | Pnliprp2 | pancreatic lip | Extracellular S |
| ENSMUSG00000024678 | 0.00080599 | 0.00121036 | Ms4a4d | membrane-s | Plasma Mem |
| ENSMUSG00000029151 | 0.00080614 | 0.00121038 | Slc30a3 | solute carrier | Plasma Mem |
| ENSMUSG00000032515 | 0.00081009 | 0.00121609 | Csrnp1 | cysteine and | Nucleus |
| ENSMUSG00000038365 | 0.00081397 | 0.00122169 | Fbxo25 | F-box protei | Nucleus |
| ENSMUSG00000009035 | 0.00081569 | 0.00122406 | Tmem184b | transmembra | Other |
| ENSMUSG00000028063 | 0.00081817 | 0.00122755 | Lmna | lamin A/C | Nucleus |
| ENSMUSG00000024983 | 0.00082559 | 0.00123846 | Vti1a | vesicle transp | Plasma Mem |
| ENSMUSG00000026000 | 0.00082611 | 0.00123902 | Lancl1 | LanC like 1 | Plasma Mem |
| ENSMUSG00000017950 | 0.00082743 | 0.00124077 | Hnf4a | hepatocyte n | Nucleus |
| ENSMUSG00000073096 | 0.00083425 | 0.00125077 | Lrrc61 | leucine rich r | Other |
| ENSMUSG00000024055 | 0.00084123 | 0.00126101 | Cyp4f13 | cytochrome I | Cytoplasm |
| ENSMUSG00000006021 | 0.00084442 | 0.00126532 | Kptn | kaptin, actin | Plasma Mem |
| ENSMUSG00000029166 | 0.00084456 | 0.00126532 | Mapre3 | microtubule a | Cytoplasm |
| ENSMUSG00000073079 | 0.00084435 | 0.00126532 | Srp54a | signal recogn | Cytoplasm |
| ENSMUSG00000030766 | 0.00084725 | 0.00126911 | Arhgap17 | Rho GTPase a | Cytoplasm |
| ENSMUSG00000006736 | 0.00084872 | 0.00127109 | Tspan31 | tetraspanin 3 | Plasma Mem |
| ENSMUSG00000024276 | 0.00084998 | 0.00127275 | Zfp397 | zinc finger pr | Nucleus |
| ENSMUSG00000038240 | 0.00085072 | 0.00127362 | Pdss2 | decaprenyl d | Cytoplasm |
| ENSMUSG00000015002 | 0.00085154 | 0.00127462 | Efr3a | EFR3 homolo | Plasma Mem |
| ENSMUSG00000018428 | 0.0008531 | 0.00127672 | Akap1 | A-kinase anc | Cytoplasm |
| ENSMUSG00000027534 | 0.00085367 | 0.00127735 | Snx16 | sorting nexin | Cytoplasm |
| ENSMUSG00000026544 | 0.0008544 | 0.00127822 | Dusp23 | dual specifici | Cytoplasm |
| ENSMUSG00000040302 | 0.00085754 | 0.00128269 | Rbm48 | RNA binding | Nucleus |
| ENSMUSG00000028668 | 0.00085903 | 0.00128467 | Tceb3 | elongin A | Nucleus |
| ENSMUSG00000029426 | 0.0008601 | 0.00128604 | Scarb2 | scavenger re | Plasma Mem |
| ENSMUSG00000022401 | 0.00086422 | 0.00129197 | Xpnpep3 | X-prolyl amir | Cytoplasm |
| ENSMUSG00000026172 | 0.00086874 | 0.0012985 | Bcs1l | BCS1 homolo | Cytoplasm |
| ENSMUSG00000046351 | 0.00087654 | 0.00130991 | Zfp322a | zinc finger pr | Cytoplasm |
| ENSMUSG00000079108 | 0.00088614 | 0.00132402 | Srp54b | signal recogn | Other |
| ENSMUSG00000005374 | 0.00089164 | 0.00133201 | Tbl2 | transducin be | Plasma Mem |
| ENSMUSG00000025316 | 0.00089815 | 0.00134149 | Banp | BTG3 associa | Nucleus |
| ENSMUSG00000004054 | 0.00090033 | 0.00134451 | Map3k11 | mitogen-acti | Cytoplasm |
| ENSMUSG00000046985 | 0.0009048 | 0.00135094 | Tapt1 | transmembra | Plasma Mem |
| ENSMUSG00000036764 | 0.00090593 | 0.00135238 | Dnajc12 | DnaJ heat sh | Cytoplasm |
| ENSMUSG00000024622 | 0.00090977 | 0.00135786 | Hmgxb3 | HMG-box co | Nucleus |
| ENSMUSG00000021930 | 0.00092431 | 0.00137932 | Spryd7 | SPRY domain | Other |
| ENSMUSG00000068917 | 0.00092747 | 0.00138378 | Clk2 | CDC like kina | Nucleus |
| ENSMUSG00000023806 | 0.00092836 | 0.00138487 | Rsph3b | radial spoke | Extracellular S |
| ENSMUSG00000032415 | 0.00092893 | 0.00138547 | Ube2cbp | ubiquitin pro | Other |
| ENSMUSG00000027304 | 0.000932 | 0.00138979 | Rtf1 | RTF1 homolo | Nucleus |
| ENSMUSG00000027522 | 0.00093805 | 0.00139857 | Stx16 | syntaxin 16 | Cytoplasm |
| ENSMUSG00000032420 | 0.00094447 | 0.00140788 | Nt5e | 5'-nucleotida | Plasma Mem |
| ENSMUSG00000053128 | 0.00094886 | 0.00141417 | Rnf26 | ring finger pr | Cytoplasm |
| ENSMUSG00000074749 | 0.00095204 | 0.0014184 | Kiz | kizuna centro | Cytoplasm |
| ENSMUSG00000046962 | 0.00095198 | 0.0014184 | Zbtb21 | zinc finger ar | Nucleus |
| ENSMUSG00000014932 | 0.0009533 | 0.00142003 | Yes1 | YES proto-or | Cytoplasm |
| ENSMUSG00000028703 | 0.00095859 | 0.00142765 | Lrrc41 | leucine rich r | Cytoplasm |
| ENSMUSG00000069678 | 0.0009667 | 0.00143947 | Pcgf1 | polycomb gr | Nucleus |

| | | | | | |
|---|---|---|---|---|---|
| ENSMUSG00000035152 | 0.00096891 | 0.0014425 | Ap2b1 | adaptor-rela | Plasma Mem |
| ENSMUSG00000044433 | 0.0009754 | 0.00145191 | Camsap3 | calmodulin re | Cytoplasm |
| ENSMUSG00000022361 | 0.00098078 | 0.00145965 | Zhx1 | zinc fingers a | Nucleus |
| ENSMUSG00000050953 | 0.00098367 | 0.0014637 | Gja1 | gap junction | Plasma Mem |
| ENSMUSG00000039254 | 0.00098457 | 0.00146477 | Pomt1 | protein O-m | Cytoplasm |
| ENSMUSG00000019055 | 0.00098582 | 0.00146636 | Plod1 | procollagen- | Cytoplasm |
| ENSMUSG00000030002 | 0.00098684 | 0.00146762 | Dusp11 | dual specifici | Nucleus |
| ENSMUSG00000031349 | 0.00098718 | 0.00146786 | Nsdhl | NAD(P) depe | Cytoplasm |
| ENSMUSG00000006390 | 0.00099628 | 0.00148112 | Elovl1 | ELOVL fatty a | Cytoplasm |
| ENSMUSG00000020834 | 0.0009967 | 0.00148149 | Dhrs13 | dehydrogena | Other |
| ENSMUSG00000029512 | 0.00099848 | 0.00148386 | Ulk1 | unc-51 like a | Cytoplasm |
| ENSMUSG00000005575 | 0.00099983 | 0.0014856 | Ube2m | ubiquitin cor | Cytoplasm |
| ENSMUSG00000024212 | 0.00100429 | 0.00149197 | Mllt1 | MLLT1, super | Nucleus |
| ENSMUSG00000031823 | 0.00100705 | 0.0014958 | Zdhhc7 | zinc finger Dl | Cytoplasm |
| ENSMUSG00000057411 | 0.00100933 | 0.00149892 | Fam173a | family with se | Other |
| ENSMUSG00000020115 | 0.00101083 | 0.00150087 | Tbk1 | TANK binding | Cytoplasm |
| ENSMUSG00000067279 | 0.00101365 | 0.00150479 | Ppp1r3c | protein phos | Cytoplasm |
| ENSMUSG00000047547 | 0.00101538 | 0.00150709 | Cltb | clathrin light | Plasma Mem |
| ENSMUSG00000032376 | 0.00102002 | 0.00151371 | Usp3 | ubiquitin spe | Cytoplasm |
| ENSMUSG00000028954 | 0.00102297 | 0.00151782 | Nub1 | negative regu | Nucleus |
| ENSMUSG00000036061 | 0.00102445 | 0.00151974 | Smug1 | single-stranc | Nucleus |
| ENSMUSG00000019978 | 0.00104299 | 0.00154669 | Epb4.1l2 | erythrocyte n | Plasma Mem |
| ENSMUSG00000022498 | 0.00104283 | 0.00154669 | Txndc11 | thioredoxin c | Cytoplasm |
| ENSMUSG00000021215 | 0.00104405 | 0.00154798 | Net1 | neuroepithel | Nucleus |
| ENSMUSG00000024831 | 0.00104993 | 0.00155642 | Ighmbp2 | immunoglob | Nucleus |
| ENSMUSG00000018196 | 0.00105374 | 0.0015618 | Glrx2 | glutaredoxin | Cytoplasm |
| ENSMUSG00000002108 | 0.00105582 | 0.0015646 | Nr1h3 | nuclear recep | Nucleus |
| ENSMUSG00000032492 | 0.00105719 | 0.00156635 | Pth1r | parathyroid h | Plasma Mem |
| ENSMUSG00000070044 | 0.00106455 | 0.00157697 | Fam149a | family with se | Other |
| ENSMUSG00000020218 | 0.00106972 | 0.00158435 | Wif1 | WNT inhibito | Extracellular S |
| ENSMUSG00000038936 | 0.00107165 | 0.00158692 | Sccpdh | saccharopine | Cytoplasm |
| ENSMUSG00000007207 | 0.00107581 | 0.0015928 | Stx1a | syntaxin 1A | Cytoplasm |
| ENSMUSG00000058291 | 0.00108305 | 0.00160323 | Zfp68 | zinc finger pr | Nucleus |
| ENSMUSG00000046668 | 0.001092 | 0.00161619 | Cxxc5 | CXXC finger p | Nucleus |
| ENSMUSG00000029462 | 0.00109252 | 0.00161667 | Vps29 | VPS29, retror | Cytoplasm |
| ENSMUSG00000009076 | 0.00109366 | 0.00161808 | Zmat5 | zinc finger m | Nucleus |
| ENSMUSG00000022892 | 0.00109693 | 0.00162262 | App | amyloid beta | Plasma Mem |
| ENSMUSG00000037730 | 0.00110086 | 0.00162814 | Mynn | myoneurin | Nucleus |
| ENSMUSG00000024797 | 0.00111975 | 0.00165579 | Vps51 | VPS51, GARP | Cytoplasm |
| ENSMUSG00000004364 | 0.00112686 | 0.001666 | Cul3 | cullin 3 | Nucleus |
| ENSMUSG00000063172 | 0.00112797 | 0.00166735 | Hspb11 | heat shock p | Extracellular S |
| ENSMUSG00000051316 | 0.00114254 | 0.00168859 | Taf7 | TATA-box bi | Nucleus |
| ENSMUSG00000031969 | 0.00114399 | 0.00169043 | Acad8 | acyl-CoA deh | Cytoplasm |
| ENSMUSG00000004552 | 0.00115339 | 0.00170401 | Ctse | cathepsin E | Cytoplasm |
| ENSMUSG00000037583 | 0.00116718 | 0.00172407 | Nr0b2 | nuclear recep | Nucleus |
| ENSMUSG00000035762 | 0.0011757 | 0.00173604 | Tmem161b | transmembra | Other |
| ENSMUSG00000047030 | 0.00118379 | 0.00174762 | Spata2 | spermatogen | Extracellular S |
| ENSMUSG00000054519 | 0.00118396 | 0.00174762 | Zfp867 | zinc finger pr | Other |
| ENSMUSG00000040274 | 0.00119412 | 0.00176231 | Cdk6 | cyclin depen | Nucleus |
| ENSMUSG00000025340 | 0.00119452 | 0.00176259 | Rabgef1 | RAB guanine | Cytoplasm |
| ENSMUSG00000000355 | 0.00119486 | 0.00176277 | Mcts1 | MCTS1, re-in | Cytoplasm |
| ENSMUSG00000040850 | 0.00119543 | 0.0017633 | Psme4 | proteasome a | Cytoplasm |
| ENSMUSG00000025956 | 0.00120365 | 0.0017751 | Mettl21a | methyltransfe | Nucleus |
| ENSMUSG00000031901 | 0.00120825 | 0.00178158 | Dus2 | dihydrouridir | Cytoplasm |
| ENSMUSG00000031358 | 0.00121375 | 0.00178937 | Msl3 | MSL complex | Nucleus |
| ENSMUSG00000024999 | 0.00121613 | 0.00179255 | Noc3l | NOC3 like DN | Nucleus |
| ENSMUSG00000028737 | 0.00121929 | 0.00179689 | Aldh4a1 | aldehyde deh | Cytoplasm |
| ENSMUSG00000035898 | 0.00122285 | 0.00180182 | Uba6 | ubiquitin like | Cytoplasm |

| | | | | | |
|---|---|---|---|---|---|
| ENSMUSG00000009293 | 0.0012247 | 0.00180422 | Ube2g2 | ubiquitin con | Cytoplasm |
| ENSMUSG00000021179 | 0.00122584 | 0.00180559 | Nrde2 | NRDE-2, nec | Other |
| ENSMUSG00000066441 | 0.00122887 | 0.00180972 | Rdh11 | retinol dehyc | Cytoplasm |
| ENSMUSG00000029048 | 0.00124197 | 0.0018287 | Rer1 | retention in e | Cytoplasm |
| ENSMUSG00000052155 | 0.00124459 | 0.00183189 | Acvr2a | activin A rece | Plasma Mem |
| ENSMUSG00000068391 | 0.00124481 | 0.00183189 | Chrac1 | chromatin ac | Nucleus |
| ENSMUSG00000027305 | 0.00124477 | 0.00183189 | Ndufaf1 | NADH:ubiqu | Cytoplasm |
| ENSMUSG00000040242 | 0.00125737 | 0.00185006 | Fgfr1op2 | FGFR1 oncog | Cytoplasm |
| ENSMUSG00000030643 | 0.00126755 | 0.00186438 | Rab30 | RAB30, mem | Cytoplasm |
| ENSMUSG00000026349 | 0.00128435 | 0.00188875 | Ccnt2 | cyclin T2 | Nucleus |
| ENSMUSG00000048550 | 0.00129382 | 0.00190233 | Thnsl1 | threonine syr | Nucleus |
| ENSMUSG00000025898 | 0.00129476 | 0.00190339 | Cwf19l2 | CWF19 like 2 | Other |
| ENSMUSG00000031487 | 0.00130059 | 0.00191162 | Brf2 | BRF2, RNA p | Nucleus |
| ENSMUSG00000062762 | 0.00130411 | 0.00191646 | Ei24 | EI24, autopha | Other |
| ENSMUSG00000005609 | 0.00130875 | 0.00192293 | Ctr9 | CTR9 homolc | Nucleus |
| ENSMUSG00000018861 | 0.00131556 | 0.0019326 | Fdxr | ferredoxin re | Cytoplasm |
| ENSMUSG00000071037 | 0.00131873 | 0.00193691 | Camkmt | calmodulin-l | Nucleus |
| ENSMUSG00000020024 | 0.00132367 | 0.00194382 | Cep83 | centrosomal | Extracellular S |
| ENSMUSG00000002881 | 0.00132868 | 0.00195082 | Nab1 | NGFI-A bind | Nucleus |
| ENSMUSG00000015750 | 0.00132935 | 0.00195146 | Aph1a | aph-1 homo | Cytoplasm |
| ENSMUSG00000026749 | 0.00132967 | 0.00195159 | Nek6 | NIMA relatec | Nucleus |
| ENSMUSG00000029859 | 0.00133215 | 0.00195488 | Epha1 | EPH receptor | Plasma Mem |
| ENSMUSG00000031755 | 0.00133621 | 0.0019605 | Bbs2 | Bardet-Biedl | Cytoplasm |
| ENSMUSG00000029434 | 0.00134404 | 0.00197164 | Vps33a | VPS33A, COF | Cytoplasm |
| ENSMUSG00000026785 | 0.00135451 | 0.00198665 | Pkn3 | protein kinas | Nucleus |
| ENSMUSG00000037166 | 0.00135943 | 0.00199352 | Ppp1r14a | protein phos | Cytoplasm |
| ENSMUSG00000020697 | 0.00136137 | 0.00199601 | Lig3 | DNA ligase 3 | Nucleus |
| ENSMUSG00000059060 | 0.00136179 | 0.00199626 | Rad51b | RAD51 paralc | Nucleus |
| ENSMUSG00000032398 | 0.00137046 | 0.00200862 | Snapc5 | small nuclear | Nucleus |
| ENSMUSG00000020056 | 0.00137173 | 0.00201013 | Ccdc53 | WASH comp | Cytoplasm |
| ENSMUSG00000040158 | 0.001372 | 0.00201017 | Tax1bp3 | Tax1 binding | Cytoplasm |
| ENSMUSG00000026162 | 0.0013753 | 0.00201465 | Nhej1 | non-homolo | Nucleus |
| ENSMUSG00000024896 | 0.00138561 | 0.00202903 | Minpp1 | multiple inos | Cytoplasm |
| ENSMUSG00000023068 | 0.00138671 | 0.00203028 | Nus1 | NUS1, dehyd | Cytoplasm |
| ENSMUSG00000012114 | 0.00138749 | 0.00203107 | Med15 | mediator cor | Nucleus |
| ENSMUSG00000067430 | 0.00139496 | 0.00204165 | Zfp763 | zinc finger pr | Nucleus |
| ENSMUSG00000024404 | 0.00139833 | 0.00204622 | Riok3 | RIO kinase 3 | Cytoplasm |
| ENSMUSG00000021311 | 0.00139953 | 0.00204761 | Mtr | 5-methyltetra | Cytoplasm |
| ENSMUSG00000029344 | 0.00140313 | 0.00205251 | Tpst2 | tyrosylproteir | Cytoplasm |
| ENSMUSG00000058748 | 0.00140718 | 0.00205807 | Zfp958 | zinc finger pr | Other |
| ENSMUSG00000034501 | 0.00141053 | 0.00206262 | Pcnxl4 | pecanex 4 | Other |
| ENSMUSG00000049493 | 0.00141472 | 0.00206838 | Pls1 | plastin 1 | Plasma Mem |
| ENSMUSG00000030839 | 0.00143241 | 0.00209387 | Sergef | secretion reg | Nucleus |
| ENSMUSG00000006235 | 0.0014541 | 0.00212483 | Epor | erythropoieti | Plasma Mem |
| ENSMUSG00000030990 | 0.00145404 | 0.00212483 | Pgap2 | post-GPI atta | Cytoplasm |
| ENSMUSG00000037808 | 0.00145682 | 0.00212843 | Fam76b | family with se | Nucleus |
| ENSMUSG00000056209 | 0.00146097 | 0.00213411 | Npm3 | nucleophosm | Nucleus |
| ENSMUSG00000028138 | 0.00147123 | 0.00214873 | Adh5 | alcohol dehy | Cytoplasm |
| ENSMUSG00000022668 | 0.00147366 | 0.00215189 | Gtpbp8 | GTP binding | Cytoplasm |
| ENSMUSG00000022967 | 0.00147504 | 0.00215353 | Ifnar1 | interferon alp | Plasma Mem |
| ENSMUSG00000030279 | 0.00147743 | 0.00215664 | C2cd5 | C2 calcium d | Cytoplasm |
| ENSMUSG00000028013 | 0.00148591 | 0.00216865 | Ppa2 | pyrophospha | Cytoplasm |
| ENSMUSG00000062101 | 0.00148694 | 0.00216975 | Zfp119b | zinc finger pr | Other |
| ENSMUSG00000042251 | 0.00150421 | 0.00219458 | Pm20d1 | peptidase M2 | Cytoplasm |
| ENSMUSG00000025199 | 0.00150937 | 0.00220172 | Chuk | conserved he | Cytoplasm |
| ENSMUSG00000005362 | 0.00152813 | 0.00222869 | Crbn | cereblon | Cytoplasm |
| ENSMUSG00000024299 | 0.00153112 | 0.00223266 | Adamts10 | ADAM metal | Extracellular S |
| ENSMUSG00000031949 | 0.00154139 | 0.00224724 | Adat1 | adenosine de | Other |

| | | | | | |
|---|---|---|---|---|---|
| ENSMUSG00000032050 | 0.00155801 | 0.00227107 | Rdx | radixin | Cytoplasm |
| ENSMUSG00000025159 | 0.00155928 | 0.00227252 | Mms19 | MMS19 hom | Nucleus |
| ENSMUSG00000007458 | 0.0015621 | 0.00227623 | M6pr | mannose-6- | Cytoplasm |
| ENSMUSG00000078429 | 0.0015648 | 0.00227977 | Ctdsp2 | CTD small ph | Nucleus |
| ENSMUSG00000034487 | 0.00157309 | 0.00229144 | Kdelc2 | KDEL motif c | Other |
| ENSMUSG00000028218 | 0.00158509 | 0.00230852 | Fam92a | family with se | Cytoplasm |
| ENSMUSG00000044894 | 0.00159029 | 0.00231528 | Uqcrq | ubiquinol-cy | Cytoplasm |
| ENSMUSG00000021327 | 0.00159013 | 0.00231528 | Zkscan3 | zinc finger wi | Nucleus |
| ENSMUSG00000022089 | 0.00159165 | 0.00231685 | Bin3 | bridging inte | Cytoplasm |
| ENSMUSG00000024953 | 0.00160245 | 0.00233217 | Prdx5 | peroxiredoxir | Cytoplasm |
| ENSMUSG00000078861 | 0.00160753 | 0.00233914 | Zfp931 | ZNF559-ZNF | Other |
| ENSMUSG00000036698 | 0.00160925 | 0.00234124 | Ago2 | argonaute 2, | Cytoplasm |
| ENSMUSG00000037656 | 0.00161857 | 0.00235439 | Slc20a2 | solute carrier | Plasma Mem |
| ENSMUSG00000028101 | 0.00162507 | 0.00236343 | Pias3 | protein inhib | Nucleus |
| ENSMUSG00000023921 | 0.00162548 | 0.0023636 | Mut | methylmalon | Cytoplasm |
| ENSMUSG00000003226 | 0.0016283 | 0.0023673 | Ranbp2 | RANBP2-like | Nucleus |
| ENSMUSG00000041959 | 0.00162922 | 0.00236822 | S100a10 | S100 calcium | Cytoplasm |
| ENSMUSG00000084128 | 0.00163111 | 0.00237055 | Esrp2 | epithelial spli | Nucleus |
| ENSMUSG00000024286 | 0.00163681 | 0.00237841 | Ccny | cyclin Y | Nucleus |
| ENSMUSG00000054452 | 0.00164477 | 0.00238831 | Aes | amino-termi | Nucleus |
| ENSMUSG00000078735 | 0.00164433 | 0.00238831 | Il11ra2 | interleukin 1 | Plasma Mem |
| ENSMUSG00000003402 | 0.00164457 | 0.00238831 | Prkcsh | protein kinas | Cytoplasm |
| ENSMUSG00000042216 | 0.00164394 | 0.00238831 | Sgsm1 | small G prote | Nucleus |
| ENSMUSG00000022865 | 0.0016495 | 0.00239476 | Cxadr | CXADR, Ig-lil | Plasma Mem |
| ENSMUSG00000033161 | 0.00165336 | 0.00239994 | Atp1a1 | ATPase Na+/ | Plasma Mem |
| ENSMUSG00000017760 | 0.0016577 | 0.00240582 | Ctsa | cathepsin A | Cytoplasm |
| ENSMUSG00000031728 | 0.00166383 | 0.0024143 | Zfp821 | zinc finger pr | Other |
| ENSMUSG00000055912 | 0.00167771 | 0.00243402 | Tmem150a | transmembra | Plasma Mem |
| ENSMUSG00000074909 | 0.00168464 | 0.00244365 | Ranbp6 | RAN binding | Cytoplasm |
| ENSMUSG00000038057 | 0.00168566 | 0.0024447 | Dbil5 | diazepam bir | Cytoplasm |
| ENSMUSG00000009647 | 0.00169004 | 0.00245062 | Mcu | mitochondria | Cytoplasm |
| ENSMUSG00000039345 | 0.00169094 | 0.00245149 | Mettl22 | methyltransfe | Nucleus |
| ENSMUSG00000022217 | 0.00169679 | 0.00245954 | Emc9 | ER membran | Plasma Mem |
| ENSMUSG00000048755 | 0.00169722 | 0.00245974 | Mcat | malonyl-CoA | Cytoplasm |
| ENSMUSG00000031807 | 0.00170243 | 0.00246686 | Pgls | 6-phosphogl | Cytoplasm |
| ENSMUSG00000026965 | 0.00170649 | 0.00247231 | Anapc2 | anaphase prc | Nucleus |
| ENSMUSG00000073792 | 0.00170881 | 0.00247524 | Alg6 | ALG6, alpha- | Cytoplasm |
| ENSMUSG00000030327 | 0.00172135 | 0.00249254 | Necap1 | NECAP endo | Plasma Mem |
| ENSMUSG00000044339 | 0.00174394 | 0.00252481 | Alkbh2 | alkB homolog | Nucleus |
| ENSMUSG00000042558 | 0.00174518 | 0.00252616 | Adprhl2 | ADP-ribosylh | Cytoplasm |
| ENSMUSG00000017754 | 0.00174994 | 0.0025326 | Pltp | phospholipid | Extracellular S |
| ENSMUSG00000006850 | 0.00175295 | 0.00253653 | Tmco6 | transmembra | Other |
| ENSMUSG00000033446 | 0.00175682 | 0.00254168 | Lpar6 | lysophosphat | Plasma Mem |
| ENSMUSG00000020092 | 0.0017655 | 0.00255379 | Pald1 | phosphatase | Cytoplasm |
| ENSMUSG00000073968 | 0.00177227 | 0.00256314 | Trim68 | tripartite mot | Cytoplasm |
| ENSMUSG00000073471 | 0.00177483 | 0.0025664 | Rsph3a | radial spoke | Extracellular S |
| ENSMUSG00000024785 | 0.00177772 | 0.00257012 | Rcl1 | RNA termina | Nucleus |
| ENSMUSG00000032123 | 0.00179113 | 0.00258905 | Dpagt1 | dolichyl-pho | Cytoplasm |
| ENSMUSG00000037443 | 0.00179406 | 0.00259284 | Cep85 | centrosomal | Nucleus |
| ENSMUSG00000031791 | 0.00180492 | 0.00260809 | Tmem38a | transmembra | Cytoplasm |
| ENSMUSG00000036073 | 0.00181949 | 0.00262868 | Galt | galactose-1- | Cytoplasm |
| ENSMUSG00000044328 | 0.00182044 | 0.0026296 | Trp53i13 | tumor protei | Cytoplasm |
| ENSMUSG00000020664 | 0.00182615 | 0.00263738 | Dld | dihydrolipoa | Cytoplasm |
| ENSMUSG00000048445 | 0.00183312 | 0.00264698 | Ccdc57 | coiled-coil d | Other |
| ENSMUSG00000035765 | 0.00184041 | 0.0026566 | Dym | dymeclin | Cytoplasm |
| ENSMUSG00000045538 | 0.00185945 | 0.00268361 | Ddx28 | DEAD-box h | Nucleus |
| ENSMUSG00000048497 | 0.00187602 | 0.00270705 | Mmgt2 | membrane m | Cytoplasm |
| ENSMUSG00000026796 | 0.00187896 | 0.00271082 | Fam129b | family with se | Cytoplasm |

| | | | | | |
|---|---|---|---|---|---|
| ENSMUSG00000029161 | 0.00188078 | 0.00271298 | Cgref1 | cell growth re | Extracellular S |
| ENSMUSG00000058672 | 0.00188437 | 0.00271768 | Tubb2a | tubulin beta 2 | Cytoplasm |
| ENSMUSG00000001983 | 0.00188536 | 0.00271864 | Taco1 | translational | Cytoplasm |
| ENSMUSG00000018770 | 0.00188611 | 0.00271924 | Atp5g3 | ATP synthase | Cytoplasm |
| ENSMUSG00000006651 | 0.00188714 | 0.00272026 | Aplp1 | amyloid beta | Extracellular S |
| ENSMUSG00000034867 | 0.00189167 | 0.00272584 | Ankrd27 | ankyrin repea | Cytoplasm |
| ENSMUSG00000050148 | 0.00189167 | 0.00272584 | Ubqln2 | ubiquilin 2 | Nucleus |
| ENSMUSG00000014077 | 0.0019002 | 0.00273725 | Chp1 | calcineurin lik | Cytoplasm |
| ENSMUSG00000027324 | 0.00190025 | 0.00273725 | Rpusd2 | RNA pseudou | Other |
| ENSMUSG00000039917 | 0.00191154 | 0.00275304 | Rhbdd2 | rhomboid dc | Plasma Mem |
| ENSMUSG00000021248 | 0.00191696 | 0.00276038 | Tmed10 | transmembra | Cytoplasm |
| ENSMUSG00000028822 | 0.00192994 | 0.00277858 | Tmem50a | transmembra | Plasma Mem |
| ENSMUSG00000025737 | 0.00193057 | 0.002779 | Wdr24 | WD repeat d | Cytoplasm |
| ENSMUSG00000028124 | 0.00194082 | 0.00279327 | Gclm | glutamate-cy | Cytoplasm |
| ENSMUSG00000037470 | 0.00194201 | 0.0027945 | Uggt1 | UDP-glucose | Cytoplasm |
| ENSMUSG00000063275 | 0.0019593 | 0.00281841 | Hacd1 | 3-hydroxyacy | Cytoplasm |
| ENSMUSG00000005510 | 0.00196315 | 0.00282324 | Ndufs3 | NADH:ubiqu | Cytoplasm |
| ENSMUSG00000027642 | 0.00196334 | 0.00282324 | Rpn2 | ribophorin II | Cytoplasm |
| ENSMUSG00000066150 | 0.00196536 | 0.00282566 | Slc31a1 | solute carrier | Plasma Mem |
| ENSMUSG00000031916 | 0.00197388 | 0.00283741 | Cog8 | component o | Cytoplasm |
| ENSMUSG00000044813 | 0.00197497 | 0.00283848 | Shb | SH2 domain | Cytoplasm |
| ENSMUSG00000025579 | 0.00198645 | 0.00285449 | Gaa | glucosidase a | Cytoplasm |
| ENSMUSG00000025158 | 0.00198974 | 0.00285872 | Rfng | RFNG O-fuco | Cytoplasm |
| ENSMUSG00000029310 | 0.00199425 | 0.00286421 | Nudt9 | nudix hydrola | Cytoplasm |
| ENSMUSG00000036368 | 0.00199416 | 0.00286421 | Rmdn2 | regulator of r | Cytoplasm |
| ENSMUSG00000029598 | 0.00200025 | 0.00287233 | Plbd2 | phospholipas | Extracellular S |
| ENSMUSG00000021339 | 0.00201193 | 0.00288811 | Mrs2 | MRS2, magne | Cytoplasm |
| ENSMUSG00000028656 | 0.00201599 | 0.00289344 | Cap1 | cyclase assoc | Plasma Mem |
| ENSMUSG00000030872 | 0.00202247 | 0.00290211 | Gga2 | golgi associa | Cytoplasm |
| ENSMUSG00000023953 | 0.00202274 | 0.00290211 | Polh | DNA polyme | Nucleus |
| ENSMUSG00000014303 | 0.00202392 | 0.00290331 | Glis2 | GLIS family z | Nucleus |
| ENSMUSG00000019132 | 0.00203015 | 0.00291175 | BC005537 | chromosome | Other |
| ENSMUSG00000044950 | 0.00204032 | 0.00292582 | Pwwp2a | PWWP doma | Nucleus |
| ENSMUSG00000019295 | 0.0020579 | 0.00295052 | Tmem129 | transmembra | Cytoplasm |
| ENSMUSG00000057982 | 0.00206906 | 0.00296601 | Zfp809 | zinc finger pr | Nucleus |
| ENSMUSG00000034075 | 0.0020848 | 0.00298805 | Zdhhc5 | zinc finger DI | Nucleus |
| ENSMUSG00000029600 | 0.0020861 | 0.00298941 | Rita1 | RBPJ interact | Cytoplasm |
| ENSMUSG00000024201 | 0.00208818 | 0.00299187 | Kdm4b | lysine demetl | Nucleus |
| ENSMUSG00000057667 | 0.00209013 | 0.00299415 | Bloc1s3 | biogenesis of | Cytoplasm |
| ENSMUSG00000059475 | 0.00210165 | 0.00301012 | Zfp426 | zinc finger pr | Nucleus |
| ENSMUSG00000025810 | 0.00210557 | 0.00301522 | Nrp1 | neuropilin 1 | Plasma Mem |
| ENSMUSG00000024096 | 0.00211169 | 0.00302346 | Ralbp1 | ralA binding | Cytoplasm |
| ENSMUSG00000004771 | 0.00211863 | 0.00303288 | Rab11a | RAB11A, mer | Cytoplasm |
| ENSMUSG00000034807 | 0.00211978 | 0.003034 | Colgalt1 | collagen beta | Cytoplasm |
| ENSMUSG00000028436 | 0.00213202 | 0.003051 | Dcaf12 | DDB1 and CU | Cytoplasm |
| ENSMUSG00000009406 | 0.00213354 | 0.00305211 | Elk1 | ELK1, ETS tra | Nucleus |
| ENSMUSG00000020475 | 0.00213322 | 0.00305211 | Pgam2 | phosphoglyc | Cytoplasm |
| ENSMUSG00000021585 | 0.00214869 | 0.00307326 | Cast | calpastatin | Cytoplasm |
| ENSMUSG00000030095 | 0.00215483 | 0.0030815 | Tmem43 | transmembra | Nucleus |
| ENSMUSG00000062906 | 0.00216094 | 0.00308971 | Hdac10 | histone deac | Nucleus |
| ENSMUSG00000023243 | 0.00216136 | 0.00308978 | Kcnk5 | potassium tw | Plasma Mem |
| ENSMUSG00000015568 | 0.00217568 | 0.00310972 | Lpl | lipoprotein li | Cytoplasm |
| ENSMUSG00000005836 | 0.00218131 | 0.00311723 | Gata6 | GATA bindin | Nucleus |
| ENSMUSG00000027907 | 0.00219512 | 0.00313643 | S100a11 | S100 calcium | Cytoplasm |
| ENSMUSG00000015837 | 0.00219877 | 0.0031411 | Sqstm1 | sequestosom | Cytoplasm |
| ENSMUSG00000031299 | 0.00220615 | 0.0031511 | Pdha1 | pyruvate deh | Cytoplasm |
| ENSMUSG00000021611 | 0.00221223 | 0.00315925 | Tert | telomerase re | Nucleus |
| ENSMUSG00000019984 | 0.00221402 | 0.00316125 | Med23 | mediator cor | Nucleus |

| | | | | | |
|---|---|---|---|---|---|
| ENSMUSG00000049657 | 0.00222748 | 0.00317993 | Zbtb5 | zinc finger ar | Nucleus |
| ENSMUSG00000050144 | 0.00223336 | 0.00318777 | Slc25a44 | solute carrier | Cytoplasm |
| ENSMUSG00000022614 | 0.0022444 | 0.00320297 | Lmf2 | lipase matura | Other |
| ENSMUSG00000058997 | 0.0022784 | 0.00325094 | Vwa8 | von Willebrai | Cytoplasm |
| ENSMUSG00000049044 | 0.00227936 | 0.00325175 | Rapgef4 | Rap guanine | Cytoplasm |
| ENSMUSG00000027287 | 0.00232097 | 0.00331054 | Snap23 | synaptosome | Plasma Mem |
| ENSMUSG00000051853 | 0.00233883 | 0.00333545 | Arf3 | ADP ribosyla | Cytoplasm |
| ENSMUSG00000033938 | 0.00234082 | 0.00333772 | Ndufb7 | NADH:ubiqu | Cytoplasm |
| ENSMUSG00000039031 | 0.00235942 | 0.00336365 | Arhgap18 | Rho GTPase a | Cytoplasm |
| ENSMUSG00000022809 | 0.00236624 | 0.00337222 | Nr1i2 | nuclear recep | Nucleus |
| ENSMUSG00000042632 | 0.002366 | 0.00337222 | Pla2g6 | phospholipas | Cytoplasm |
| ENSMUSG00000054277 | 0.00236774 | 0.00337378 | Arfgap3 | ADP ribosyla | Cytoplasm |
| ENSMUSG00000000340 | 0.00237965 | 0.00339017 | Dbt | dihydrolipoai | Cytoplasm |
| ENSMUSG00000022724 | 0.00238415 | 0.00339599 | Mina | ribosomal ox | Nucleus |
| ENSMUSG00000024527 | 0.00238666 | 0.00339898 | Afg3l2 | AFG3 like ma | Cytoplasm |
| ENSMUSG00000030051 | 0.00238794 | 0.00340022 | Aplf | aprataxin anc | Cytoplasm |
| ENSMUSG00000026511 | 0.00239866 | 0.0034149 | Srp9 | signal recogr | Cytoplasm |
| ENSMUSG00000023074 | 0.00240029 | 0.00341664 | Mospd1 | motile sperm | Other |
| ENSMUSG00000013698 | 0.00240361 | 0.00342078 | Pea15a | proliferation | Cytoplasm |
| ENSMUSG00000035798 | 0.00242683 | 0.00345323 | Zdhhc17 | zinc finger DI | Cytoplasm |
| ENSMUSG00000008822 | 0.00243551 | 0.00346499 | Acyp1 | acylphosphat | Cytoplasm |
| ENSMUSG00000023007 | 0.00243824 | 0.00346828 | Prpf40b | pre-mRNA p | Nucleus |
| ENSMUSG00000024777 | 0.00247204 | 0.00351515 | Ppp2r5b | protein phos | Cytoplasm |
| ENSMUSG00000028826 | 0.00247656 | 0.00352097 | Tmem57 | macoilin 1 | Nucleus |
| ENSMUSG00000024981 | 0.00247879 | 0.00352354 | Acsl5 | acyl-CoA syr | Cytoplasm |
| ENSMUSG00000032392 | 0.00248338 | 0.00352946 | Parp16 | poly(ADP-rib | Cytoplasm |
| ENSMUSG00000029726 | 0.00249261 | 0.00354197 | Mepce | methylphosp | Other |
| ENSMUSG00000040616 | 0.00251984 | 0.00358006 | Tmem51 | transmembra | Other |
| ENSMUSG00000022663 | 0.00253028 | 0.00359427 | Atg3 | autophagy re | Cytoplasm |
| ENSMUSG00000037487 | 0.00256937 | 0.00364917 | Ubr5 | ubiquitin pro | Nucleus |
| ENSMUSG00000015747 | 0.00257577 | 0.00365764 | Vps45 | vacuolar prot | Cytoplasm |
| ENSMUSG00000078515 | 0.00259753 | 0.00368791 | Ddi2 | DNA damage | Plasma Mem |
| ENSMUSG00000034429 | 0.00260756 | 0.00370152 | Zfp707 | zinc finger pr | Other |
| ENSMUSG00000032571 | 0.00263709 | 0.00374279 | Pik3r4 | phosphoinos | Cytoplasm |
| ENSMUSG00000035206 | 0.00265043 | 0.00376108 | Sppl2b | signal peptid | Plasma Mem |
| ENSMUSG00000039013 | 0.0026618 | 0.00377657 | Siglecf | sialic acid bir | Plasma Mem |
| ENSMUSG00000032396 | 0.00266357 | 0.00377844 | Dis3l | DIS3 like exo | Cytoplasm |
| ENSMUSG00000078622 | 0.00267401 | 0.00379195 | Ccdc47 | coiled-coil d | Extracellular S |
| ENSMUSG00000071654 | 0.00267384 | 0.00379195 | Uqcc3 | ubiquinol-cy | Extracellular S |
| ENSMUSG00000050786 | 0.00269087 | 0.0038152 | Ccdc126 | coiled-coil d | Other |
| ENSMUSG00000025968 | 0.00269851 | 0.00382538 | Ndufs1 | NADH:ubiqu | Cytoplasm |
| ENSMUSG00000039233 | 0.00270877 | 0.00383927 | Tbce | tubulin foldir | Cytoplasm |
| ENSMUSG00000001844 | 0.00272111 | 0.00385611 | Zdhhc4 | zinc finger DI | Cytoplasm |
| ENSMUSG00000040841 | 0.00273369 | 0.00387327 | Six5 | SIX homeobc | Nucleus |
| ENSMUSG00000032440 | 0.00274158 | 0.00388379 | Tgfbr2 | transforming | Plasma Mem |
| ENSMUSG00000025257 | 0.0027432 | 0.00388542 | Ribc1 | RIB43A doma | Other |
| ENSMUSG00000048701 | 0.00275162 | 0.00389669 | Ccdc6 | coiled-coil d | Cytoplasm |
| ENSMUSG00000053119 | 0.00276892 | 0.00392051 | Chmp3 | charged mult | Cytoplasm |
| ENSMUSG00000028136 | 0.00277164 | 0.0039237 | Snx27 | sorting nexin | Cytoplasm |
| ENSMUSG00000030768 | 0.00278013 | 0.00393503 | Disp1 | dispatched R | Plasma Mem |
| ENSMUSG00000010047 | 0.00278453 | 0.0039406 | Hyal2 | hyaluronoglu | Cytoplasm |
| ENSMUSG00000039262 | 0.00278782 | 0.00394458 | Prrc2b | proline rich c | Other |
| ENSMUSG00000025405 | 0.00279549 | 0.00395476 | Inhbc | inhibin subur | Extracellular S |
| ENSMUSG00000027255 | 0.00280076 | 0.00396153 | Arfgap2 | ADP ribosyla | Nucleus |
| ENSMUSG00000040212 | 0.00281416 | 0.00397914 | Emp3 | epithelial me | Plasma Mem |
| ENSMUSG00000022453 | 0.00284134 | 0.00401688 | Naga | alpha-N-ace | Cytoplasm |
| ENSMUSG00000051285 | 0.00285779 | 0.00403945 | Pcmtd1 | protein-L-isc | Cytoplasm |
| ENSMUSG00000000823 | 0.00286473 | 0.00404858 | Znf512b | zinc finger pr | Nucleus |

| | | | | | |
|---|---|---|---|---|---|
| ENSMUSG00000034177 | 0.00288277 | 0.00407338 | Rnf43 | ring finger pr | Plasma Mem |
| ENSMUSG00000028653 | 0.00288691 | 0.00407853 | Trit1 | tRNA isopent | Cytoplasm |
| ENSMUSG00000030265 | 0.00288971 | 0.00408179 | Kras | KRAS proto-( | Cytoplasm |
| ENSMUSG00000024127 | 0.00290795 | 0.00410686 | Prepl | prolyl endop | Other |
| ENSMUSG00000056131 | 0.00291844 | 0.00412097 | Pgm3 | phosphogluc | Cytoplasm |
| ENSMUSG00000044795 | 0.00293095 | 0.00413793 | Cyb5d1 | cytochrome I | Other |
| ENSMUSG00000037325 | 0.00293368 | 0.00414107 | Bbs7 | Bardet-Biedl | Cytoplasm |
| ENSMUSG00000017288 | 0.00294639 | 0.00415831 | Vps53 | VPS53, GARP | Cytoplasm |
| ENSMUSG00000024259 | 0.00295073 | 0.00416372 | Slc25a46 | solute carrier | Cytoplasm |
| ENSMUSG00000001156 | 0.00295573 | 0.00417008 | Mxd1 | MAX dimeriz | Nucleus |
| ENSMUSG00000032096 | 0.00297353 | 0.00419448 | Arcn1 | archain 1 | Cytoplasm |
| ENSMUSG00000053644 | 0.00297628 | 0.00419764 | Aldh7a1 | aldehyde deh | Cytoplasm |
| ENSMUSG00000031600 | 0.00299346 | 0.00422116 | Vps37a | VPS37A, ESC | Cytoplasm |
| ENSMUSG00000063568 | 0.00299675 | 0.00422507 | Jazf1 | JAZF zinc fing | Nucleus |
| ENSMUSG00000044876 | 0.00300616 | 0.00423762 | Zfp444 | zinc finger pr | Nucleus |
| ENSMUSG00000003161 | 0.0030285 | 0.0042684 | Sri | sorcin | Cytoplasm |
| ENSMUSG00000044252 | 0.00303548 | 0.0042775 | Osbpl1a | oxysterol bin | Cytoplasm |
| ENSMUSG00000038462 | 0.00304615 | 0.00429181 | Uqcrfs1 | ubiquinol-cy | Cytoplasm |
| ENSMUSG00000023892 | 0.003059 | 0.00430919 | Zfp51 | zinc finger pr | Other |
| ENSMUSG00000066900 | 0.00306244 | 0.00431329 | Suds3 | SDS3 homolo | Nucleus |
| ENSMUSG00000035285 | 0.00307096 | 0.00432457 | Nat14 | N-acetyltrans | Extracellular S |
| ENSMUSG00000024072 | 0.00311238 | 0.00438215 | Yipf4 | Yip1 domain | Cytoplasm |
| ENSMUSG00000024959 | 0.00311859 | 0.00439015 | Bad | BCL2 associa | Cytoplasm |
| ENSMUSG00000028098 | 0.00312678 | 0.00440093 | Rnf115 | ring finger pr | Cytoplasm |
| ENSMUSG00000021250 | 0.00313458 | 0.00441116 | Fos | Fos proto-or | Nucleus |
| ENSMUSG00000028700 | 0.00316428 | 0.0044522 | Pomgnt1 | protein O-lin | Cytoplasm |
| ENSMUSG00000037896 | 0.00316621 | 0.00445415 | Rcor1 | REST corepre | Nucleus |
| ENSMUSG00000042284 | 0.00317194 | 0.00446146 | Itga1 | integrin subu | Plasma Mem |
| ENSMUSG00000028773 | 0.00320013 | 0.00450035 | Fabp3 | fatty acid bin | Cytoplasm |
| ENSMUSG00000048756 | 0.00323789 | 0.00455213 | Foxo3 | forkhead box | Nucleus |
| ENSMUSG00000026548 | 0.00323804 | 0.00455213 | Slamf9 | SLAM family | Extracellular S |
| ENSMUSG00000023143 | 0.00326074 | 0.00458326 | Nagpa | N-acetylgluc | Cytoplasm |
| ENSMUSG00000031792 | 0.00326225 | 0.00458461 | Usb1 | U6 snRNA bi | Nucleus |
| ENSMUSG00000037526 | 0.0032646 | 0.00458712 | Atg14 | autophagy re | Cytoplasm |
| ENSMUSG00000026811 | 0.00327043 | 0.00459454 | St6galnac6 | ST6 N-acetyl | Cytoplasm |
| ENSMUSG00000030826 | 0.00329175 | 0.00462358 | Bcat2 | branched cha | Cytoplasm |
| ENSMUSG00000051390 | 0.00329221 | 0.00462358 | Zbtb22 | zinc finger ar | Nucleus |
| ENSMUSG00000052428 | 0.00329632 | 0.00462857 | Tmco1 | transmembra | Cytoplasm |
| ENSMUSG00000026273 | 0.00329694 | 0.00462865 | Mterf4 | mitochondria | Cytoplasm |
| ENSMUSG00000046380 | 0.00332032 | 0.00466069 | Jrk | Jrk helix-turn | Nucleus |
| ENSMUSG00000014856 | 0.00333164 | 0.00467579 | Tmem208 | transmembra | Other |
| ENSMUSG00000068747 | 0.00333392 | 0.00467819 | Sort1 | sortilin 1 | Plasma Mem |
| ENSMUSG00000038733 | 0.00337122 | 0.00472973 | Wdr26 | WD repeat d | Cytoplasm |
| ENSMUSG00000029370 | 0.00341272 | 0.00478634 | Rassf6 | Ras associatio | Other |
| ENSMUSG00000017756 | 0.0034447 | 0.00483039 | Slc12a7 | solute carrier | Plasma Mem |
| ENSMUSG00000026788 | 0.00345412 | 0.00484277 | Zbtb43 | zinc finger ar | Nucleus |
| ENSMUSG00000058638 | 0.00346903 | 0.00486286 | Zfp110 | zinc finger pr | Nucleus |
| ENSMUSG00000039530 | 0.00347607 | 0.0048719 | Tusc3 | tumor suppre | Plasma Mem |
| ENSMUSG00000068580 | 0.00347791 | 0.00487365 | Zfyve19 | zinc finger FY | Cytoplasm |
| ENSMUSG00000015653 | 0.00349457 | 0.00489617 | Steap2 | STEAP2 meta | Plasma Mem |
| ENSMUSG00000037957 | 0.00355144 | 0.00497501 | Wdr20 | WD repeat d | Nucleus |
| ENSMUSG00000005150 | 0.00356025 | 0.00498651 | Wdr83 | WD repeat d | Nucleus |
| ENSMUSG00000038871 | 0.00361471 | 0.00506193 | Bpgm | bisphosphog | Extracellular S |
| ENSMUSG00000075702 | 0.0036179 | 0.00506555 | Selm | selenoprotein | Cytoplasm |
| ENSMUSG00000043241 | 0.00362121 | 0.00506932 | Upf2 | UPF2, regulat | Cytoplasm |
| ENSMUSG00000053916 | 0.00364316 | 0.0050992 | Nanp | N-acetylneur | Cytoplasm |
| ENSMUSG00000002844 | 0.00367177 | 0.00513837 | Adprh | ADP-ribosyla | Other |
| ENSMUSG00000050708 | 0.00367501 | 0.00514204 | Ftl1 | ferritin light c | Cytoplasm |

| | | | | | |
|---|---|---|---|---|---|
| ENSMUSG00000006057 | 0.00367855 | 0.00514612 | Atp5g1 | ATP synthase | Cytoplasm |
| ENSMUSG00000055866 | 0.00371553 | 0.00519698 | Per2 | period circad | Nucleus |
| ENSMUSG00000017765 | 0.00378388 | 0.00529169 | Slc12a4 | solute carrier | Plasma Mem |
| ENSMUSG00000034485 | 0.00380771 | 0.00532413 | Uaca | uveal autoan | Cytoplasm |
| ENSMUSG00000021361 | 0.00382682 | 0.00534995 | Tmem14c | transmembra | Plasma Mem |
| ENSMUSG00000006342 | 0.00382781 | 0.00535043 | Susd2 | sushi domair | Extracellular S |
| ENSMUSG00000004270 | 0.00383426 | 0.00535854 | Lpcat3 | lysophosphat | Plasma Mem |
| ENSMUSG00000026502 | 0.00384588 | 0.00537387 | Desi2 | desumoylatir | Cytoplasm |
| ENSMUSG00000028708 | 0.00387912 | 0.00541941 | Mknk1 | MAP kinase i | Cytoplasm |
| ENSMUSG00000029776 | 0.00388519 | 0.00542697 | Hibadh | 3-hydroxyisc | Cytoplasm |
| ENSMUSG00000061288 | 0.00388798 | 0.00542995 | Taok3 | TAO kinase 3 | Cytoplasm |
| ENSMUSG00000053134 | 0.00391268 | 0.00546353 | Supt7l | SPT7 like, ST/ | Nucleus |
| ENSMUSG00000061119 | 0.00392202 | 0.00547565 | Prcp | prolylcarboxy | Cytoplasm |
| ENSMUSG00000027374 | 0.00393479 | 0.00549256 | Mrps5 | mitochondria | Cytoplasm |
| ENSMUSG00000010392 | 0.00394521 | 0.00550619 | Gosr1 | golgi SNAP r | Cytoplasm |
| ENSMUSG00000039033 | 0.00394662 | 0.00550722 | Tasp1 | taspase 1 | Nucleus |
| ENSMUSG00000026219 | 0.00395237 | 0.00551431 | Trip12 | thyroid horm | Cytoplasm |
| ENSMUSG00000072501 | 0.00395891 | 0.00552251 | Phf20l1 | PHD finger p | Other |
| ENSMUSG00000037477 | 0.00396409 | 0.00552881 | Tbx10 | T-box 10 | Nucleus |
| ENSMUSG00000026404 | 0.00400241 | 0.00558132 | Ddx59 | DEAD-box h | Other |
| ENSMUSG00000061371 | 0.00402504 | 0.00561193 | Zfp873 | zinc finger pr | Nucleus |
| ENSMUSG00000001240 | 0.00406106 | 0.0056612 | Ramp2 | receptor acti | Plasma Mem |
| ENSMUSG00000031387 | 0.00406227 | 0.00566193 | Renbp | renin binding | Cytoplasm |
| ENSMUSG00000028454 | 0.00406948 | 0.00567103 | Pigo | phosphatidyl | Cytoplasm |
| ENSMUSG00000025485 | 0.00411002 | 0.00572657 | Ric8 | RIC8 guanine | Cytoplasm |
| ENSMUSG00000040213 | 0.00413575 | 0.00576145 | Ccbl2 | kynurenine a | Cytoplasm |
| ENSMUSG00000020439 | 0.00414068 | 0.00576735 | Smtn | smoothelin | Extracellular S |
| ENSMUSG00000055044 | 0.00414231 | 0.00576865 | Pdlim1 | PDZ and LIM | Cytoplasm |
| ENSMUSG00000028986 | 0.00416142 | 0.00579332 | Klhl7 | kelch like fam | Nucleus |
| ENSMUSG00000020544 | 0.00422237 | 0.00587718 | Cox11 | cytochrome c | Cytoplasm |
| ENSMUSG00000021993 | 0.00422307 | 0.00587718 | Mipep | mitochondria | Cytoplasm |
| ENSMUSG00000032782 | 0.00423269 | 0.00588958 | Cntrob | centrobin, ce | Cytoplasm |
| ENSMUSG00000042570 | 0.00423521 | 0.0058921 | Mier2 | MIER family r | Extracellular S |
| ENSMUSG00000014349 | 0.00428078 | 0.00595449 | Ube2z | ubiquitin cor | Nucleus |
| ENSMUSG00000059734 | 0.00428634 | 0.00596123 | Ndufs8 | NADH:ubiqu | Cytoplasm |
| ENSMUSG00000017144 | 0.00429338 | 0.00597002 | Rnd3 | Rho family G | Cytoplasm |
| ENSMUSG00000035545 | 0.00429417 | 0.00597013 | Leng8 | leukocyte rec | Other |
| ENSMUSG00000029050 | 0.00430153 | 0.00597935 | Ski | SKI proto-on | Nucleus |
| ENSMUSG00000038533 | 0.00431091 | 0.00599139 | Cbfa2t2 | CBFA2/RUNX | Nucleus |
| ENSMUSG00000042167 | 0.00431334 | 0.00599376 | Papd4 | terminal nucl | Nucleus |
| ENSMUSG00000027272 | 0.0043201 | 0.00600215 | Ubr1 | ubiquitin pro | Cytoplasm |
| ENSMUSG00000022827 | 0.00435313 | 0.00604702 | Rabl3 | RAB, membe | Other |
| ENSMUSG00000044147 | 0.00439411 | 0.00610293 | Arf6 | ADP ribosyla | Plasma Mem |
| ENSMUSG00000021235 | 0.00445484 | 0.00618624 | Coq6 | coenzyme Q( | Cytoplasm |
| ENSMUSG00000029068 | 0.00445867 | 0.00619053 | Ccnl2 | cyclin L2 | Nucleus |
| ENSMUSG00000030347 | 0.00446704 | 0.00620111 | D6Wsu163e | chromosome | Cytoplasm |
| ENSMUSG00000009248 | 0.00448237 | 0.00622135 | Ascl2 | achaete-scut | Nucleus |
| ENSMUSG00000025351 | 0.00450055 | 0.00624553 | Cd63 | CD63 molecu | Plasma Mem |
| ENSMUSG00000029672 | 0.00450322 | 0.00624819 | Fam3c | family with se | Extracellular S |
| ENSMUSG00000019838 | 0.00451898 | 0.00626902 | Slc16a10 | solute carrier | Plasma Mem |
| ENSMUSG00000050668 | 0.00455597 | 0.00631928 | Gpatch11 | G-patch dom | Nucleus |
| ENSMUSG00000045287 | 0.00457251 | 0.00634116 | Rtn4rl1 | reticulon 4 re | Plasma Mem |
| ENSMUSG00000042821 | 0.00459862 | 0.0063763 | Snai1 | snail family tr | Nucleus |
| ENSMUSG00000030431 | 0.00460171 | 0.00637952 | Tmem238 | transmembra | Other |
| ENSMUSG00000038695 | 0.00462935 | 0.00641677 | Josd2 | Josephin dor | Cytoplasm |
| ENSMUSG00000020806 | 0.00464241 | 0.00643272 | Rhbdf2 | rhomboid 5 h | Cytoplasm |
| ENSMUSG00000002205 | 0.00464188 | 0.00643272 | Vrk3 | vaccinia relat | Nucleus |
| ENSMUSG00000022022 | 0.00464587 | 0.00643644 | Mtrf1 | mitochondria | Cytoplasm |

| | | | | | | |
|---|---|---|---|---|---|---|
| ENSMUSG00000027459 | 0.00465639 | 0.00644994 | Fam110a | | family with se | Other |
| ENSMUSG00000034757 | 0.00466697 | 0.00646352 | Tmub2 | | transmembra | Other |
| ENSMUSG00000033416 | 0.00468159 | 0.00648269 | Gucd1 | | guanylyl cycl | Other |
| ENSMUSG00000033429 | 0.00469976 | 0.00650676 | Mcee | | methylmalon | Cytoplasm |
| ENSMUSG00000043923 | 0.00470712 | 0.00651587 | Ccdc84 | | coiled-coil d | Other |
| ENSMUSG00000023938 | 0.00471329 | 0.00652332 | Aars2 | | alanyl-tRNA | Cytoplasm |
| ENSMUSG00000021539 | 0.00474044 | 0.00655915 | Lect2 | | leukocyte cel | Extracellular S |
| ENSMUSG00000039100 | 0.00474077 | 0.00655915 | | 6-Mar | membrane a | Cytoplasm |
| ENSMUSG00000001211 | 0.00474496 | 0.00656386 | Agpat3 | | 1-acylglycerc | Cytoplasm |
| ENSMUSG00000035139 | 0.00475891 | 0.00658206 | Secisbp2 | | SECIS binding | Cytoplasm |
| ENSMUSG00000024747 | 0.00476148 | 0.00658452 | Aldh1a7 | | aldehyde deh | Cytoplasm |
| ENSMUSG00000075703 | 0.00477094 | 0.0065965 | Ept1 | | selenoprotei | Cytoplasm |
| ENSMUSG00000038418 | 0.004797 | 0.0066307 | Egr1 | | early growth | Nucleus |
| ENSMUSG00000052031 | 0.00479727 | 0.0066307 | Tagap1 | | T cell activati | Other |
| ENSMUSG00000004018 | 0.00480771 | 0.00664402 | Fancl | | FA complem | Nucleus |
| ENSMUSG00000031858 | 0.0048148 | 0.00665271 | Mau2 | | MAU2 sister | Nucleus |
| ENSMUSG00000031105 | 0.00482032 | 0.00665922 | Slc25a14 | | solute carrier | Cytoplasm |
| ENSMUSG00000006445 | 0.00483494 | 0.00667832 | Epha2 | | EPH receptor | Plasma Mem |
| ENSMUSG00000038775 | 0.00483627 | 0.00667904 | Vill | | villin like | Cytoplasm |
| ENSMUSG00000073758 | 0.00489676 | 0.00676146 | Sh3d21 | | SH3 domain | Cytoplasm |
| ENSMUSG00000053293 | 0.00489971 | 0.00676441 | Pom121 | | POM121 trar | Nucleus |
| ENSMUSG00000034902 | 0.004937 | 0.00681475 | Pip5k1c | | phosphatidyl | Plasma Mem |
| ENSMUSG00000048897 | 0.00495289 | 0.00683554 | Zfp710 | | zinc finger pr | Nucleus |
| ENSMUSG00000033355 | 0.00495932 | 0.00684328 | Rtp4 | | receptor tran | Plasma Mem |
| ENSMUSG00000024807 | 0.00497364 | 0.00686191 | Syvn1 | | synoviolin 1 | Cytoplasm |
| ENSMUSG00000003849 | 0.00497574 | 0.00686367 | Nqo1 | | NAD(P)H qui | Cytoplasm |
| ENSMUSG00000030536 | 0.00498715 | 0.00687826 | Iqgap1 | | IQ motif cont | Cytoplasm |
| ENSMUSG00000026576 | 0.00502028 | 0.0069228 | Atp1b1 | | ATPase Na+/ | Plasma Mem |
| ENSMUSG00000032602 | 0.00505571 | 0.00697049 | Slc25a20 | | solute carrier | Cytoplasm |
| ENSMUSG00000022130 | 0.00508505 | 0.00700978 | Tgds | | TDP-glucose | Extracellular S |
| ENSMUSG00000001786 | 0.00510418 | 0.00703499 | Fbxo7 | | F-box protei | Cytoplasm |
| ENSMUSG00000025140 | 0.00512335 | 0.00706023 | Pycr1 | | pyrroline-5-c | Cytoplasm |
| ENSMUSG00000042105 | 0.00512498 | 0.00706132 | Inpp5f | | inositol polyp | Plasma Mem |
| ENSMUSG00000004565 | 0.00516335 | 0.007113 | Pnpla6 | | patatin like p | Cytoplasm |
| ENSMUSG00000061313 | 0.00517454 | 0.00712723 | Ddhd2 | | DDHD doma | Cytoplasm |
| ENSMUSG00000044857 | 0.00518913 | 0.00714613 | Lemd2 | | LEM domain | Nucleus |
| ENSMUSG00000037750 | 0.00521458 | 0.0071787 | Fam222b | | family with se | Nucleus |
| ENSMUSG00000015605 | 0.00521392 | 0.0071787 | Srf | | serum respor | Nucleus |
| ENSMUSG00000039531 | 0.00521537 | 0.0071787 | Zufsp | | zinc finger cc | Other |
| ENSMUSG00000000605 | 0.0052421 | 0.0072143 | Clcn4-2 | | chloride volta | Plasma Mem |
| ENSMUSG00000017802 | 0.00526123 | 0.00723942 | Fam134c | | reticulophagy | Other |
| ENSMUSG00000002963 | 0.00526949 | 0.00724959 | Pnkp | | polynucleotic | Nucleus |
| ENSMUSG00000052040 | 0.00528024 | 0.00726317 | Klf13 | | Kruppel like f | Nucleus |
| ENSMUSG00000018900 | 0.00531819 | 0.00731416 | Slc22a5 | | solute carrier | Plasma Mem |
| ENSMUSG00000062753 | 0.005346 | 0.00735119 | AI413582 | | expressed se | Other |
| ENSMUSG00000020483 | 0.0053549 | 0.00736221 | Dynll2 | | dynein light c | Cytoplasm |
| ENSMUSG00000006356 | 0.00535676 | 0.00736355 | Crip2 | | cysteine rich | Plasma Mem |
| ENSMUSG00000002580 | 0.00537007 | 0.00738062 | Mien1 | | migration an | Cytoplasm |
| ENSMUSG00000027447 | 0.00537205 | 0.00738213 | Cst3 | | cystatin C | Extracellular S |
| ENSMUSG00000027189 | 0.00538 | 0.00739182 | Trim44 | | tripartite mot | Cytoplasm |
| ENSMUSG00000047417 | 0.00540133 | 0.0074199 | Rexo1 | | RNA exonucl | Nucleus |
| ENSMUSG00000030166 | 0.00548904 | 0.00753915 | Rad52 | | RAD52 homc | Nucleus |
| ENSMUSG00000031485 | 0.00549266 | 0.00754286 | Prosc | | pyridoxal phc | Cytoplasm |
| ENSMUSG00000030986 | 0.00552748 | 0.00758943 | Dhx32 | | DEAH-box h | Other |
| ENSMUSG00000045822 | 0.00553466 | 0.00759803 | Zswim3 | | zinc finger SV | Other |
| ENSMUSG00000015522 | 0.00555745 | 0.00762806 | Arnt | | aryl hydrocar | Nucleus |
| ENSMUSG00000041598 | 0.00556017 | 0.00763052 | Cdc42ep4 | | CDC42 effect | Cytoplasm |
| ENSMUSG00000019891 | 0.00562374 | 0.00771394 | Dcbld1 | | discoidin, CU | Extracellular S |

| | | | | | |
|---|---|---|---|---|---|
| ENSMUSG00000025764 | 0.00562338 | 0.00771394 | Jade1 | jade family P | Nucleus |
| ENSMUSG00000074211 | 0.00563762 | 0.00773171 | Sdhaf1 | succinate del | Cytoplasm |
| ENSMUSG00000003234 | 0.00564204 | 0.00773648 | Abcf3 | ATP binding | Other |
| ENSMUSG00000074825 | 0.00564628 | 0.00774102 | Itpripl1 | ITPRIP like 1 | Other |
| ENSMUSG00000019832 | 0.00565562 | 0.00775254 | Rab32 | RAB32, mem | Cytoplasm |
| ENSMUSG00000005982 | 0.00566224 | 0.00776034 | Naa60 | N(alpha)-ace | Cytoplasm |
| ENSMUSG00000021495 | 0.00572294 | 0.00784223 | Fam193b | family with se | Cytoplasm |
| ENSMUSG00000029869 | 0.00575714 | 0.00788779 | Ephb6 | EPH receptor | Plasma Mem |
| ENSMUSG00000036537 | 0.00579339 | 0.00793615 | Rnf113a1 | ring finger pr | Other |
| ENSMUSG00000026649 | 0.00581672 | 0.0079668 | Cfap126 | cilia and flag | Other |
| ENSMUSG00000059554 | 0.00582343 | 0.00797467 | Ccdc28a | coiled-coil d | Other |
| ENSMUSG00000032540 | 0.00585101 | 0.00801112 | Abhd5 | abhydrolase | Cytoplasm |
| ENSMUSG00000029616 | 0.00588511 | 0.00805648 | Erp29 | endoplasmic | Cytoplasm |
| ENSMUSG00000038742 | 0.00589225 | 0.00806492 | Angptl6 | angiopoietin | Extracellular S |
| ENSMUSG00000019143 | 0.0058939 | 0.00806584 | Hars2 | histidyl-tRNA | Cytoplasm |
| ENSMUSG00000041949 | 0.00590144 | 0.00807484 | Tango6 | transport anc | Other |
| ENSMUSG00000025575 | 0.0059756 | 0.00817496 | Cant1 | calcium activ | Extracellular S |
| ENSMUSG00000024422 | 0.00604526 | 0.00826889 | Dhx16 | DEAH-box h | Nucleus |
| ENSMUSG00000030101 | 0.00609423 | 0.0083345 | Sumf1 | sulfatase mo | Cytoplasm |
| ENSMUSG00000032342 | 0.00613803 | 0.00839302 | Mto1 | mitochondria | Cytoplasm |
| ENSMUSG00000019850 | 0.00619702 | 0.00847228 | Tnfaip3 | TNF alpha in | Nucleus |
| ENSMUSG00000021338 | 0.00621068 | 0.00848957 | Lrrc16a | capping prot | Cytoplasm |
| ENSMUSG00000019853 | 0.00621949 | 0.00850021 | Hebp2 | heme binding | Cytoplasm |
| ENSMUSG00000009030 | 0.00623074 | 0.00851418 | Pdcl | phosducin lik | Cytoplasm |
| ENSMUSG00000016528 | 0.00627341 | 0.00857108 | Mapkapk2 | mitogen-acti | Nucleus |
| ENSMUSG00000002012 | 0.00628885 | 0.00859076 | Pnck | pregnancy u | Other |
| ENSMUSG00000028868 | 0.00632927 | 0.00864455 | Wasf2 | WAS protein | Plasma Mem |
| ENSMUSG00000029017 | 0.00633782 | 0.0086548 | Pmpcb | peptidase, m | Cytoplasm |
| ENSMUSG00000032531 | 0.00634094 | 0.00865764 | Amotl2 | angiomotin l | Plasma Mem |
| ENSMUSG00000022552 | 0.00635909 | 0.00868099 | Sharpin | SHANK assoc | Plasma Mem |
| ENSMUSG00000021824 | 0.00640061 | 0.00873623 | Ap3m1 | adaptor relat | Cytoplasm |
| ENSMUSG00000051154 | 0.00641072 | 0.00874716 | Commd3 | COMM doma | Extracellular S |
| ENSMUSG00000066892 | 0.00641039 | 0.00874716 | Fbxl12 | F-box and le | Cytoplasm |
| ENSMUSG00000027180 | 0.00641704 | 0.00875435 | Fbxo3 | F-box protei | Cytoplasm |
| ENSMUSG00000039990 | 0.00647052 | 0.00882585 | Edrf1 | erythroid diff | Other |
| ENSMUSG00000027293 | 0.0064739 | 0.00882902 | Ehd4 | EH domain c | Plasma Mem |
| ENSMUSG00000026618 | 0.00648489 | 0.00884255 | Iars2 | isoleucyl-tRN | Cytoplasm |
| ENSMUSG00000033991 | 0.00648964 | 0.00884758 | Ttc37 | tetratricopep | Nucleus |
| ENSMUSG00000009566 | 0.00649595 | 0.00885472 | Fpgs | folylpolygluta | Cytoplasm |
| ENSMUSG00000061607 | 0.00656738 | 0.00895062 | Mdc1 | mediator of I | Nucleus |
| ENSMUSG00000003344 | 0.00657763 | 0.00896312 | Btbd2 | BTB domain | Cytoplasm |
| ENSMUSG00000037857 | 0.00658702 | 0.00897444 | Nufip2 | NUFIP2, FMR | Cytoplasm |
| ENSMUSG00000022750 | 0.00662498 | 0.00902467 | Klhl22 | kelch like fan | Cytoplasm |
| ENSMUSG00000050373 | 0.00665725 | 0.00906714 | Snx21 | sorting nexin | Cytoplasm |
| ENSMUSG00000022450 | 0.0066648 | 0.00907593 | Ndufa6 | NADH:ubiqu | Cytoplasm |
| ENSMUSG00000062480 | 0.0066733 | 0.00908602 | Acat3 | acetyl-CoA a | Cytoplasm |
| ENSMUSG00000029802 | 0.00671597 | 0.00914261 | Abcg2 | ATP binding | Plasma Mem |
| ENSMUSG00000024782 | 0.00674287 | 0.00917774 | Ak3 | adenylate kir | Cytoplasm |
| ENSMUSG00000055128 | 0.00679094 | 0.00924164 | Cgrrf1 | cell growth re | Nucleus |
| ENSMUSG00000040563 | 0.00679287 | 0.00924276 | Lppr2 | phospholipid | Other |
| ENSMUSG00000032649 | 0.00680679 | 0.00925866 | Colgalt2 | collagen beta | Cytoplasm |
| ENSMUSG00000000563 | 0.00681149 | 0.00926354 | Atp5f1 | ATP synthase | Cytoplasm |
| ENSMUSG00000020593 | 0.00681921 | 0.00927252 | Lpin1 | lipin 1 | Nucleus |
| ENSMUSG00000068876 | 0.00682506 | 0.00927895 | Cgn | cingulin | Plasma Mem |
| ENSMUSG00000029518 | 0.00682794 | 0.00928135 | Rab35 | RAB35, mem | Cytoplasm |
| ENSMUSG00000025893 | 0.00688709 | 0.00936022 | Kbtbd3 | kelch repeat | Extracellular S |
| ENSMUSG00000020064 | 0.00689077 | 0.00936369 | Herc4 | HECT and RL | Cytoplasm |
| ENSMUSG00000035376 | 0.00691392 | 0.00939361 | Hacd2 | 3-hydroxyac | Cytoplasm |

| | | | | | |
|---|---|---|---|---|---|
| ENSMUSG00000032422 | 0.00691521 | 0.00939381 | Snx14 | sorting nexin | Cytoplasm |
| ENSMUSG00000036820 | 0.00693946 | 0.00942523 | Amdhd2 | amidohydrol | Nucleus |
| ENSMUSG00000025240 | 0.00696189 | 0.00945413 | Sacm1l | SAC1 like ph | Cytoplasm |
| ENSMUSG00000045435 | 0.00704214 | 0.00956156 | Tmem60 | transmembra | Other |
| ENSMUSG00000021982 | 0.0070495 | 0.00956998 | Cdadc1 | cytidine and | Other |
| ENSMUSG00000012640 | 0.007069 | 0.00959488 | Zfp715 | zinc finger pr | Nucleus |
| ENSMUSG00000019883 | 0.00710042 | 0.00963596 | Echdc1 | ethylmalonyl | Cytoplasm |
| ENSMUSG00000026727 | 0.00719394 | 0.00976128 | Rsu1 | Ras suppress | Cytoplasm |
| ENSMUSG00000026360 | 0.007215 | 0.00978825 | Rgs2 | regulator of ( | Nucleus |
| ENSMUSG00000020629 | 0.00721852 | 0.00979142 | Adi1 | acireductone | Nucleus |
| ENSMUSG00000078619 | 0.00725706 | 0.00984209 | Smarcd2 | SWI/SNF rela | Nucleus |
| ENSMUSG00000039536 | 0.00727608 | 0.00986627 | Stau1 | staufen doub | Cytoplasm |
| ENSMUSG00000041845 | 0.00733276 | 0.00993988 | Rhod | ras homolog | Cytoplasm |
| ENSMUSG00000035517 | 0.00733178 | 0.00993988 | Tdrd7 | tudor domair | Cytoplasm |
| ENSMUSG00000022299 | 0.00736799 | 0.00998601 | Slc25a32 | solute carrier | Cytoplasm |
| ENSMUSG00000052707 | 0.00737194 | 0.00998973 | Tnrc6a | trinucleotide | Nucleus |
| ENSMUSG00000006522 | 5.385315539 | 2.629318554 | Itih3 | inter-alpha-t | Extracellular S |
| ENSMUSG00000079523 | 1.078322975 | 4.710568802 | Tmsb10 | thymosin, be | Cytoplasm |
| ENSMUSG00000002992 | 1.200579519 | 5.536005561 | Apoc2 | apolipoprote | Extracellular S |

**t**

s in the dataset.

| Family | expression pattern |
|---|---|
| other | Up-regulated |
| transporter | Up-regulated |
| enzyme | Up-regulated |
| transporter | Up-regulated |
| transporter | Up-regulated |
| transporter | Down-regulated |
| transporter | Up-regulated |
| transporter | Up-regulated |
| transporter | Up-regulated |
| transporter | Up-regulated |
| transporter | Up-regulated |
| transporter | Up-regulated |
| other | Up-regulated |
| other | Up-regulated |
| growth facto | Down-regulated |
| kinase | Down-regulated |
| other | Down-regulated |
| other | Up-regulated |
| other | Up-regulated |
| other | Down-regulated |
| other | Up-regulated |
| other | Down-regulated |
| enzyme | Down-regulated |
| other | Up-regulated |
| enzyme | Up-regulated |
| transporter | Up-regulated |
| other | Up-regulated |
| transporter | Up-regulated |
| other | Up-regulated |
| other | Down-regulated |
| transcription | Down-regulated |
| other | Up-regulated |
| enzyme | Down-regulated |
| other | Up-regulated |
| other | Down-regulated |
| other | Up-regulated |
| enzyme | Up-regulated |
| other | Up-regulated |
| other | Up-regulated |
| growth facto | Up-regulated |
| other | Up-regulated |
| other | Up-regulated |
| other | Down-regulated |
| transporter | Up-regulated |
| other | Up-regulated |
| transporter | Up-regulated |
| other | Down-regulated |
| other | Down-regulated |
| other | Down-regulated |
| other | Down-regulated |

| | |
|---|---|
| enzyme | Up-regulated |
| enzyme | Up-regulated |
| enzyme | Up-regulated |
| enzyme | Up-regulated |
| other | Up-regulated |
| other | Up-regulated |
| enzyme | Up-regulated |
| peptidase | Up-regulated |
| kinase | Down-regulated |
| enzyme | Down-regulated |
| enzyme | Up-regulated |
| enzyme | Up-regulated |
| enzyme | Down-regulated |
| peptidase | Up-regulated |
| transcription | Down-regulated |
| enzyme | Down-regulated |
| other | Up-regulated |
| peptidase | Up-regulated |
| enzyme | Up-regulated |
| other | Up-regulated |
| enzyme | Up-regulated |
| other | Up-regulated |
| transporter | Up-regulated |
| other | Down-regulated |
| other | Down-regulated |
| enzyme | Up-regulated |
| enzyme | Up-regulated |
| other | Up-regulated |
| other | Down-regulated |
| other | Up-regulated |
| phosphatase | Down-regulated |
| other | Up-regulated |
| transporter | Down-regulated |
| other | Up-regulated |
| other | Down-regulated |
| other | Down-regulated |
| other | Down-regulated |
| other | Down-regulated |
| other | Down-regulated |
| other | Down-regulated |
| enzyme | Down-regulated |
| peptidase | Up-regulated |
| other | Down-regulated |
| transcription | Down-regulated |
| other | Down-regulated |
| enzyme | Down-regulated |
| enzyme | Down-regulated |
| enzyme | Up-regulated |
| other | Down-regulated |
| other | Down-regulated |
| other | Up-regulated |
| enzyme | Down-regulated |
| peptidase | Up-regulated |
| other | Down-regulated |
| other | Down-regulated |
| enzyme | Up-regulated |
| growth facto | Up-regulated |
| transcription | Down-regulated |

| | |
|---|---|
| transcription | Down-regulated |
| enzyme | Down-regulated |
| other | Down-regulated |
| other | Down-regulated |
| other | Down-regulated |
| transcription | Down-regulated |
| other | Down-regulated |
| transporter | Down-regulated |
| other | Down-regulated |
| enzyme | Up-regulated |
| ion channel | Down-regulated |
| enzyme | Down-regulated |
| other | Up-regulated |
| other | Down-regulated |
| kinase | Down-regulated |
| cytokine | Down-regulated |
| other | Down-regulated |
| other | Down-regulated |
| enzyme | Up-regulated |
| other | Down-regulated |
| other | Up-regulated |
| peptidase | Up-regulated |
| translation re | Down-regulated |
| other | Down-regulated |
| other | Down-regulated |
| enzyme | Up-regulated |
| other | Up-regulated |
| other | Down-regulated |
| other | Down-regulated |
| translation re | Down-regulated |
| other | Down-regulated |
| transporter | Up-regulated |
| enzyme | Down-regulated |
| other | Down-regulated |
| other | Down-regulated |
| other | Up-regulated |
| other | Down-regulated |
| transcription | Down-regulated |
| enzyme | Down-regulated |
| enzyme | Down-regulated |
| enzyme | Down-regulated |
| other | Down-regulated |
| transporter | Down-regulated |
| other | Down-regulated |
| other | Down-regulated |
| transcription | Down-regulated |
| other | Down-regulated |
| other | Down-regulated |
| transcription | Down-regulated |
| other | Down-regulated |
| other | Up-regulated |
| other | Down-regulated |
| enzyme | Down-regulated |
| other | Down-regulated |
| transporter | Down-regulated |
| peptidase | Up-regulated |
| transporter | Up-regulated |
| other | Down-regulated |

| | |
|---|---|
| other | Down-regulated |
| transcription | Down-regulated |
| translation re | Down-regulated |
| other | Down-regulated |
| enzyme | Down-regulated |
| transporter | Up-regulated |
| kinase | Down-regulated |
| other | Down-regulated |
| enzyme | Down-regulated |
| translation re | Down-regulated |
| translation re | Down-regulated |
| enzyme | Up-regulated |
| enzyme | Up-regulated |
| enzyme | Down-regulated |
| other | Down-regulated |
| enzyme | Up-regulated |
| enzyme | Up-regulated |
| transcription | Down-regulated |
| other | Down-regulated |
| other | Down-regulated |
| other | Down-regulated |
| transcription | Down-regulated |
| other | Down-regulated |
| transporter | Up-regulated |
| transcription | Down-regulated |
| other | Down-regulated |
| enzyme | Up-regulated |
| enzyme | Up-regulated |
| other | Down-regulated |
| translation re | Down-regulated |
| other | Down-regulated |
| enzyme | Up-regulated |
| enzyme | Down-regulated |
| enzyme | Down-regulated |
| other | Down-regulated |
| other | Down-regulated |
| other | Up-regulated |
| enzyme | Down-regulated |
| other | Down-regulated |
| enzyme | Up-regulated |
| enzyme | Down-regulated |
| translation re | Down-regulated |
| enzyme | Up-regulated |
| other | Down-regulated |
| other | Down-regulated |
| enzyme | Up-regulated |
| other | Up-regulated |
| other | Down-regulated |
| other | Up-regulated |
| other | Down-regulated |
| other | Up-regulated |
| other | Down-regulated |
| transporter | Up-regulated |
| enzyme | Down-regulated |
| enzyme | Up-regulated |
| other | Down-regulated |
| cytokine | Up-regulated |
| translation re | Down-regulated |

| | |
|---|---|
| transporter | Up-regulated |
| other | Down-regulated |
| peptidase | Up-regulated |
| transmembra | Up-regulated |
| phosphatase | Down-regulated |
| other | Down-regulated |
| other | Down-regulated |
| other | Down-regulated |
| other | Down-regulated |
| other | Down-regulated |
| enzyme | Down-regulated |
| transcription | Down-regulated |
| peptidase | Down-regulated |
| transcription | Down-regulated |
| other | Down-regulated |
| other | Up-regulated |
| kinase | Down-regulated |
| other | Down-regulated |
| other | Down-regulated |
| enzyme | Up-regulated |
| enzyme | Up-regulated |
| enzyme | Down-regulated |
| other | Down-regulated |
| other | Down-regulated |
| other | Down-regulated |
| enzyme | Down-regulated |
| other | Down-regulated |
| other | Down-regulated |
| enzyme | Down-regulated |
| kinase | Down-regulated |
| other | Down-regulated |
| enzyme | Down-regulated |
| other | Down-regulated |
| enzyme | Down-regulated |
| kinase | Down-regulated |
| transcription | Down-regulated |
| other | Down-regulated |
| other | Down-regulated |
| transcription | Down-regulated |
| transcription | Down-regulated |
| other | Down-regulated |
| transporter | Down-regulated |
| transmembra | Up-regulated |
| other | Up-regulated |
| enzyme | Down-regulated |
| other | Down-regulated |
| other | Down-regulated |
| enzyme | Down-regulated |
| other | Down-regulated |
| enzyme | Down-regulated |
| kinase | Down-regulated |
| other | Down-regulated |
| peptidase | Up-regulated |
| transporter | Up-regulated |
| other | Up-regulated |
| peptidase | Down-regulated |
| enzyme | Up-regulated |
| cytokine | Up-regulated |

| | |
|---|---|
| enzyme | Down-regulated |
| other | Down-regulated |
| enzyme | Down-regulated |
| enzyme | Up-regulated |
| other | Down-regulated |
| enzyme | Down-regulated |
| other | Down-regulated |
| other | Down-regulated |
| other | Down-regulated |
| other | Down-regulated |
| phosphatase | Down-regulated |
| other | Down-regulated |
| enzyme | Down-regulated |
| other | Down-regulated |
| peptidase | Down-regulated |
| other | Down-regulated |
| other | Down-regulated |
| other | Down-regulated |
| other | Down-regulated |
| phosphatase | Down-regulated |
| enzyme | Down-regulated |
| transporter | Down-regulated |
| other | Down-regulated |
| other | Down-regulated |
| enzyme | Down-regulated |
| enzyme | Down-regulated |
| enzyme | Down-regulated |
| other | Down-regulated |
| transporter | Down-regulated |
| other | Down-regulated |
| other | Down-regulated |
| peptidase | Down-regulated |
| enzyme | Down-regulated |
| other | Down-regulated |
| enzyme | Up-regulated |
| enzyme | Down-regulated |
| other | Down-regulated |
| other | Down-regulated |
| other | Down-regulated |
| other | Down-regulated |
| other | Down-regulated |
| other | Down-regulated |
| transcription | Down-regulated |
| other | Down-regulated |
| other | Down-regulated |
| other | Down-regulated |
| other | Down-regulated |
| enzyme | Down-regulated |
| other | Up-regulated |
| other | Down-regulated |
| enzyme | Up-regulated |
| other | Down-regulated |
| peptidase | Down-regulated |
| other | Down-regulated |
| other | Down-regulated |
| enzyme | Up-regulated |
| enzyme | Down-regulated |
| enzyme | Down-regulated |

| | |
|---|---|
| other | Up-regulated |
| enzyme | Down-regulated |
| enzyme | Up-regulated |
| other | Down-regulated |
| other | Down-regulated |
| other | Down-regulated |
| peptidase | Down-regulated |
| transcription | Down-regulated |
| other | Down-regulated |
| enzyme | Down-regulated |
| transcription | Down-regulated |
| other | Down-regulated |
| peptidase | Down-regulated |
| peptidase | Up-regulated |
| transporter | Up-regulated |
| other | Down-regulated |
| peptidase | Down-regulated |
| other | Down-regulated |
| other | Up-regulated |
| other | Down-regulated |
| transporter | Up-regulated |
| other | Down-regulated |
| other | Down-regulated |
| other | Up-regulated |
| other | Down-regulated |
| enzyme | Up-regulated |
| other | Down-regulated |
| enzyme | Down-regulated |
| other | Down-regulated |
| peptidase | Down-regulated |
| other | Down-regulated |
| other | Down-regulated |
| other | Down-regulated |
| other | Down-regulated |
| other | Down-regulated |
| other | Down-regulated |
| transcription | Down-regulated |
| growth facto | Up-regulated |
| other | Down-regulated |
| enzyme | Up-regulated |
| transporter | Up-regulated |
| other | Down-regulated |
| other | Down-regulated |
| other | Down-regulated |
| other | Down-regulated |
| enzyme | Down-regulated |
| other | Down-regulated |
| enzyme | Down-regulated |
| other | Up-regulated |
| enzyme | Down-regulated |
| kinase | Down-regulated |
| other | Down-regulated |
| transcription | Down-regulated |
| other | Down-regulated |
| enzyme | Up-regulated |
| transporter | Down-regulated |
| other | Down-regulated |
| other | Down-regulated |

| | |
|---|---|
| enzyme | Up-regulated |
| enzyme | Up-regulated |
| other | Down-regulated |
| other | Down-regulated |
| enzyme | Up-regulated |
| enzyme | Down-regulated |
| other | Down-regulated |
| transporter | Up-regulated |
| other | Down-regulated |
| peptidase | Down-regulated |
| other | Up-regulated |
| other | Down-regulated |
| other | Down-regulated |
| other | Down-regulated |
| enzyme | Down-regulated |
| enzyme | Down-regulated |
| enzyme | Down-regulated |
| other | Down-regulated |
| other | Up-regulated |
| translation re | Down-regulated |
| enzyme | Up-regulated |
| transcription | Down-regulated |
| other | Down-regulated |
| cytokine | Up-regulated |
| transcription | Down-regulated |
| enzyme | Up-regulated |
| translation re | Down-regulated |
| enzyme | Down-regulated |
| transcription | Down-regulated |
| other | Down-regulated |
| transcription | Down-regulated |
| transcription | Down-regulated |
| other | Down-regulated |
| enzyme | Down-regulated |
| enzyme | Down-regulated |
| other | Down-regulated |
| transmembra | Up-regulated |
| other | Down-regulated |
| other | Down-regulated |
| other | Up-regulated |
| other | Down-regulated |
| other | Down-regulated |
| other | Down-regulated |
| enzyme | Down-regulated |
| transporter | Up-regulated |
| enzyme | Up-regulated |
| kinase | Down-regulated |
| enzyme | Down-regulated |
| cytokine | Down-regulated |
| other | Up-regulated |
| other | Down-regulated |
| other | Down-regulated |
| other | Down-regulated |
| enzyme | Down-regulated |
| enzyme | Down-regulated |
| other | Up-regulated |
| other | Down-regulated |
| other | Down-regulated |

| | |
|---|---|
| other | Down-regulated |
| kinase | Down-regulated |
| enzyme | Up-regulated |
| transcription | Down-regulated |
| other | Down-regulated |
| other | Down-regulated |
| other | Up-regulated |
| other | Down-regulated |
| other | Down-regulated |
| other | Down-regulated |
| other | Down-regulated |
| enzyme | Down-regulated |
| enzyme | Up-regulated |
| other | Down-regulated |
| enzyme | Down-regulated |
| translation re | Down-regulated |
| peptidase | Down-regulated |
| enzyme | Down-regulated |
| peptidase | Down-regulated |
| transporter | Down-regulated |
| translation re | Down-regulated |
| enzyme | Down-regulated |
| other | Down-regulated |
| other | Down-regulated |
| other | Down-regulated |
| other | Down-regulated |
| other | Down-regulated |
| other | Down-regulated |
| enzyme | Down-regulated |
| other | Down-regulated |
| transcription | Down-regulated |
| other | Down-regulated |
| enzyme | Down-regulated |
| enzyme | Down-regulated |
| other | Down-regulated |
| other | Down-regulated |
| ion channel | Up-regulated |
| other | Down-regulated |
| enzyme | Up-regulated |
| other | Down-regulated |
| transporter | Up-regulated |
| other | Up-regulated |
| other | Down-regulated |
| other | Down-regulated |
| kinase | Down-regulated |
| cytokine | Up-regulated |
| peptidase | Up-regulated |
| kinase | Down-regulated |
| enzyme | Down-regulated |
| translation re | Down-regulated |
| other | Down-regulated |
| other | Down-regulated |
| translation re | Down-regulated |
| enzyme | Down-regulated |
| other | Down-regulated |
| transporter | Down-regulated |
| other | Down-regulated |
| enzyme | Up-regulated |

| | |
|---|---|
| enzyme | Down-regulated |
| transporter | Down-regulated |
| enzyme | Up-regulated |
| transporter | Down-regulated |
| other | Up-regulated |
| enzyme | Up-regulated |
| other | Down-regulated |
| enzyme | Down-regulated |
| other | Up-regulated |
| other | Down-regulated |
| kinase | Down-regulated |
| enzyme | Down-regulated |
| translation re | Down-regulated |
| transporter | Down-regulated |
| other | Up-regulated |
| enzyme | Down-regulated |
| other | Down-regulated |
| other | Down-regulated |
| translation re | Down-regulated |
| enzyme | Down-regulated |
| other | Down-regulated |
| enzyme | Up-regulated |
| other | Down-regulated |
| other | Down-regulated |
| other | Down-regulated |
| other | Down-regulated |
| other | Down-regulated |
| other | Down-regulated |
| kinase | Down-regulated |
| other | Down-regulated |
| other | Down-regulated |
| transcription | Down-regulated |
| other | Down-regulated |
| enzyme | Up-regulated |
| other | Down-regulated |
| other | Down-regulated |
| other | Down-regulated |
| other | Down-regulated |
| enzyme | Up-regulated |
| other | Down-regulated |
| enzyme | Down-regulated |
| other | Down-regulated |
| other | Down-regulated |
| other | Down-regulated |
| other | Down-regulated |
| enzyme | Down-regulated |
| other | Down-regulated |
| other | Down-regulated |
| enzyme | Up-regulated |
| enzyme | Up-regulated |
| enzyme | Down-regulated |
| other | Down-regulated |
| enzyme | Up-regulated |
| enzyme | Down-regulated |
| transporter | Up-regulated |
| other | Down-regulated |
| other | Down-regulated |
| other | Down-regulated |

| | |
|---|---|
| transporter | Down-regulated |
| peptidase | Up-regulated |
| other | Down-regulated |
| enzyme | Down-regulated |
| other | Down-regulated |
| transporter | Up-regulated |
| other | Up-regulated |
| other | Down-regulated |
| transcription | Down-regulated |
| transcription | Down-regulated |
| other | Down-regulated |
| translation re | Down-regulated |
| transporter | Up-regulated |
| enzyme | Down-regulated |
| other | Down-regulated |
| other | Down-regulated |
| transcription | Down-regulated |
| transcription | Down-regulated |
| other | Down-regulated |
| kinase | Down-regulated |
| enzyme | Down-regulated |
| transcription | Up-regulated |
| other | Down-regulated |
| other | Down-regulated |
| other | Down-regulated |
| enzyme | Up-regulated |
| enzyme | Up-regulated |
| other | Down-regulated |
| other | Down-regulated |
| transcription | Down-regulated |
| other | Down-regulated |
| transporter | Up-regulated |
| transcription | Down-regulated |
| other | Down-regulated |
| other | Down-regulated |
| other | Down-regulated |
| transcription | Down-regulated |
| transporter | Down-regulated |
| other | Down-regulated |
| enzyme | Up-regulated |
| other | Up-regulated |
| other | Down-regulated |
| enzyme | Down-regulated |
| enzyme | Up-regulated |
| other | Down-regulated |
| transcription | Down-regulated |
| peptidase | Up-regulated |
| enzyme | Down-regulated |
| transporter | Up-regulated |
| kinase | Down-regulated |
| other | Down-regulated |
| transporter | Up-regulated |
| other | Down-regulated |
| peptidase | Up-regulated |
| other | Down-regulated |
| other | Up-regulated |
| other | Down-regulated |
| phosphatase | Up-regulated |

| | |
|---|---|
| other | Down-regulated |
| other | Down-regulated |
| translation re | Down-regulated |
| transporter | Down-regulated |
| transcription | Down-regulated |
| enzyme | Down-regulated |
| enzyme | Down-regulated |
| enzyme | Down-regulated |
| other | Down-regulated |
| other | Down-regulated |
| other | Down-regulated |
| other | Down-regulated |
| peptidase | Down-regulated |
| enzyme | Down-regulated |
| transcription | Down-regulated |
| enzyme | Down-regulated |
| enzyme | Down-regulated |
| transcription | Down-regulated |
| transporter | Up-regulated |
| other | Up-regulated |
| other | Down-regulated |
| other | Down-regulated |
| enzyme | Down-regulated |
| other | Down-regulated |
| phosphatase | Up-regulated |
| other | Down-regulated |
| other | Down-regulated |
| kinase | Down-regulated |
| enzyme | Down-regulated |
| enzyme | Up-regulated |
| other | Up-regulated |
| other | Down-regulated |
| transporter | Down-regulated |
| other | Down-regulated |
| transporter | Up-regulated |
| peptidase | Down-regulated |
| other | Down-regulated |
| transporter | Down-regulated |
| transporter | Up-regulated |
| other | Down-regulated |
| kinase | Down-regulated |
| other | Down-regulated |
| enzyme | Up-regulated |
| peptidase | Down-regulated |
| other | Down-regulated |
| enzyme | Down-regulated |
| other | Down-regulated |
| peptidase | Down-regulated |
| enzyme | Down-regulated |
| enzyme | Down-regulated |
| transcription | Down-regulated |
| transporter | Down-regulated |
| other | Down-regulated |
| other | Down-regulated |
| enzyme | Down-regulated |
| other | Down-regulated |
| transcription | Down-regulated |
| transcription | Down-regulated |

| | |
|---|---|
| enzyme | Down-regulated |
| enzyme | Up-regulated |
| peptidase | Down-regulated |
| other | Down-regulated |
| transcription | Down-regulated |
| translation re | Down-regulated |
| other | Down-regulated |
| other | Up-regulated |
| enzyme | Down-regulated |
| enzyme | Down-regulated |
| other | Down-regulated |
| other | Down-regulated |
| other | Down-regulated |
| other | Down-regulated |
| enzyme | Down-regulated |
| other | Down-regulated |
| transcription | Down-regulated |
| other | Down-regulated |
| other | Down-regulated |
| enzyme | Up-regulated |
| enzyme | Down-regulated |
| kinase | Down-regulated |
| other | Down-regulated |
| other | Down-regulated |
| other | Down-regulated |
| peptidase | Down-regulated |
| enzyme | Up-regulated |
| peptidase | Down-regulated |
| other | Down-regulated |
| kinase | Down-regulated |
| enzyme | Down-regulated |
| peptidase | Down-regulated |
| enzyme | Up-regulated |
| transporter | Down-regulated |
| enzyme | Down-regulated |
| other | Down-regulated |
| other | Down-regulated |
| other | Down-regulated |
| enzyme | Down-regulated |
| transcription | Down-regulated |
| transporter | Up-regulated |
| other | Down-regulated |
| peptidase | Down-regulated |
| other | Down-regulated |
| other | Down-regulated |
| other | Down-regulated |
| other | Down-regulated |
| enzyme | Down-regulated |
| transcription | Down-regulated |
| other | Down-regulated |
| enzyme | Down-regulated |
| peptidase | Down-regulated |
| other | Down-regulated |
| other | Down-regulated |
| enzyme | Up-regulated |
| other | Down-regulated |
| other | Down-regulated |
| enzyme | Down-regulated |

| | |
|---|---|
| other | Down-regulated |
| enzyme | Down-regulated |
| enzyme | Down-regulated |
| other | Down-regulated |
| enzyme | Down-regulated |
| enzyme | Up-regulated |
| transcription | Down-regulated |
| enzyme | Up-regulated |
| peptidase | Up-regulated |
| other | Up-regulated |
| enzyme | Up-regulated |
| other | Down-regulated |
| other | Down-regulated |
| enzyme | Down-regulated |
| phosphatase | Down-regulated |
| other | Down-regulated |
| other | Down-regulated |
| translation re | Down-regulated |
| other | Down-regulated |
| transcription | Down-regulated |
| enzyme | Down-regulated |
| other | Down-regulated |
| other | Down-regulated |
| other | Down-regulated |
| enzyme | Up-regulated |
| transporter | Down-regulated |
| enzyme | Down-regulated |
| other | Down-regulated |
| other | Down-regulated |
| enzyme | Down-regulated |
| other | Down-regulated |
| other | Down-regulated |
| transcription | Down-regulated |
| enzyme | Down-regulated |
| other | Down-regulated |
| enzyme | Down-regulated |
| enzyme | Up-regulated |
| other | Down-regulated |
| enzyme | Up-regulated |
| enzyme | Down-regulated |
| enzyme | Down-regulated |
| enzyme | Down-regulated |
| other | Down-regulated |
| enzyme | Down-regulated |
| transporter | Down-regulated |
| enzyme | Down-regulated |
| other | Down-regulated |
| other | Down-regulated |
| enzyme | Down-regulated |
| transporter | Up-regulated |
| transporter | Down-regulated |
| enzyme | Down-regulated |
| other | Down-regulated |
| translation re | Down-regulated |
| enzyme | Down-regulated |
| other | Down-regulated |
| enzyme | Down-regulated |
| peptidase | Down-regulated |

| | |
|---|---|
| other | Down-regulated |
| transporter | Down-regulated |
| other | Down-regulated |
| enzyme | Up-regulated |
| other | Down-regulated |
| other | Down-regulated |
| translation re | Down-regulated |
| other | Down-regulated |
| other | Down-regulated |
| kinase | Down-regulated |
| other | Down-regulated |
| other | Down-regulated |
| enzyme | Up-regulated |
| other | Down-regulated |
| other | Down-regulated |
| other | Down-regulated |
| enzyme | Down-regulated |
| transporter | Up-regulated |
| transporter | Down-regulated |
| other | Down-regulated |
| transcription | Down-regulated |
| kinase | Down-regulated |
| enzyme | Down-regulated |
| other | Down-regulated |
| enzyme | Down-regulated |
| other | Down-regulated |
| other | Down-regulated |
| enzyme | Down-regulated |
| other | Down-regulated |
| transporter | Down-regulated |
| enzyme | Down-regulated |
| other | Down-regulated |
| enzyme | Down-regulated |
| other | Up-regulated |
| transcription | Up-regulated |
| other | Down-regulated |
| transporter | Down-regulated |
| phosphatase | Down-regulated |
| other | Down-regulated |
| transcription | Down-regulated |
| other | Down-regulated |
| peptidase | Up-regulated |
| translation re | Down-regulated |
| enzyme | Down-regulated |
| other | Down-regulated |
| other | Down-regulated |
| other | Up-regulated |
| peptidase | Up-regulated |
| ion channel | Up-regulated |
| transcription | Down-regulated |
| enzyme | Down-regulated |
| transporter | Up-regulated |
| enzyme | Down-regulated |
| other | Down-regulated |
| transmembra | Up-regulated |
| other | Down-regulated |
| other | Down-regulated |
| other | Down-regulated |

| | |
|---|---|
| G-protein co | Up-regulated |
| translation re | Down-regulated |
| other | Down-regulated |
| other | Down-regulated |
| enzyme | Down-regulated |
| enzyme | Down-regulated |
| G-protein co | Up-regulated |
| enzyme | Up-regulated |
| enzyme | Down-regulated |
| enzyme | Down-regulated |
| transcription | Down-regulated |
| other | Down-regulated |
| enzyme | Down-regulated |
| other | Down-regulated |
| other | Down-regulated |
| enzyme | Up-regulated |
| other | Down-regulated |
| enzyme | Up-regulated |
| other | Down-regulated |
| other | Down-regulated |
| enzyme | Down-regulated |
| enzyme | Down-regulated |
| other | Down-regulated |
| other | Down-regulated |
| enzyme | Down-regulated |
| phosphatase | Down-regulated |
| other | Down-regulated |
| other | Down-regulated |
| enzyme | Down-regulated |
| other | Down-regulated |
| other | Down-regulated |
| other | Down-regulated |
| other | Down-regulated |
| other | Down-regulated |
| transcription | Down-regulated |
| transcription | Down-regulated |
| translation re | Down-regulated |
| enzyme | Up-regulated |
| transcription | Down-regulated |
| kinase | Down-regulated |
| transporter | Down-regulated |
| translation re | Down-regulated |
| other | Down-regulated |
| other | Down-regulated |
| phosphatase | Down-regulated |
| phosphatase | Down-regulated |
| transcription | Down-regulated |
| enzyme | Up-regulated |
| other | Down-regulated |
| transcription | Down-regulated |
| other | Down-regulated |
| enzyme | Down-regulated |
| other | Down-regulated |
| other | Down-regulated |
| other | Up-regulated |
| other | Down-regulated |
| enzyme | Up-regulated |
| transcription | Down-regulated |

| | |
|---|---|
| other | Down-regulated |
| other | Down-regulated |
| enzyme | Down-regulated |
| translation re | Down-regulated |
| enzyme | Down-regulated |
| ligand-deper | Up-regulated |
| enzyme | Down-regulated |
| kinase | Down-regulated |
| enzyme | Down-regulated |
| other | Down-regulated |
| other | Down-regulated |
| phosphatase | Down-regulated |
| enzyme | Up-regulated |
| other | Down-regulated |
| kinase | Down-regulated |
| peptidase | Down-regulated |
| other | Down-regulated |
| transporter | Up-regulated |
| transcription | Down-regulated |
| enzyme | Down-regulated |
| other | Down-regulated |
| other | Down-regulated |
| phosphatase | Down-regulated |
| enzyme | Down-regulated |
| other | Down-regulated |
| enzyme | Down-regulated |
| transcription | Down-regulated |
| other | Down-regulated |
| enzyme | Up-regulated |
| translation re | Down-regulated |
| other | Down-regulated |
| other | Down-regulated |
| other | Down-regulated |
| other | Down-regulated |
| enzyme | Down-regulated |
| other | Down-regulated |
| enzyme | Down-regulated |
| enzyme | Down-regulated |
| other | Down-regulated |
| other | Down-regulated |
| other | Down-regulated |
| enzyme | Up-regulated |
| transcription | Down-regulated |
| other | Down-regulated |
| transcription | Down-regulated |
| enzyme | Up-regulated |
| enzyme | Down-regulated |
| transmembra | Up-regulated |
| transcription | Down-regulated |
| other | Up-regulated |
| enzyme | Down-regulated |
| transcription | Down-regulated |
| transporter | Down-regulated |
| transcription | Down-regulated |
| other | Down-regulated |
| other | Down-regulated |
| phosphatase | Down-regulated |
| transporter | Up-regulated |

| | |
|---|---|
| other | Down-regulated |
| enzyme | Up-regulated |
| other | Down-regulated |
| other | Down-regulated |
| other | Down-regulated |
| enzyme | Down-regulated |
| other | Down-regulated |
| other | Down-regulated |
| other | Up-regulated |
| enzyme | Down-regulated |
| enzyme | Up-regulated |
| enzyme | Down-regulated |
| enzyme | Up-regulated |
| transporter | Up-regulated |
| enzyme | Up-regulated |
| other | Down-regulated |
| other | Down-regulated |
| transcription | Down-regulated |
| enzyme | Down-regulated |
| other | Down-regulated |
| other | Down-regulated |
| other | Down-regulated |
| enzyme | Down-regulated |
| other | Down-regulated |
| transporter | Up-regulated |
| other | Down-regulated |
| other | Down-regulated |
| transcription | Down-regulated |
| other | Down-regulated |
| transcription | Down-regulated |
| other | Down-regulated |
| other | Down-regulated |
| other | Down-regulated |
| transmembra | Up-regulated |
| enzyme | Down-regulated |
| transporter | Down-regulated |
| enzyme | Up-regulated |
| other | Down-regulated |
| enzyme | Up-regulated |
| other | Down-regulated |
| other | Down-regulated |
| other | Down-regulated |
| other | Down-regulated |
| transcription | Down-regulated |
| other | Down-regulated |
| enzyme | Down-regulated |
| other | Down-regulated |
| other | Down-regulated |
| enzyme | Down-regulated |
| enzyme | Down-regulated |
| enzyme | Down-regulated |
| transcription | Down-regulated |
| other | Up-regulated |
| other | Down-regulated |
| transporter | Down-regulated |
| other | Down-regulated |
| other | Down-regulated |
| phosphatase | Down-regulated |

| | |
|---|---|
| kinase | Down-regulated |
| kinase | Down-regulated |
| peptidase | Down-regulated |
| peptidase | Down-regulated |
| other | Up-regulated |
| transcription | Down-regulated |
| enzyme | Up-regulated |
| other | Down-regulated |
| other | Down-regulated |
| enzyme | Down-regulated |
| enzyme | Down-regulated |
| enzyme | Down-regulated |
| enzyme | Up-regulated |
| peptidase | Down-regulated |
| other | Down-regulated |
| transcription | Down-regulated |
| transcription | Down-regulated |
| enzyme | Down-regulated |
| other | Down-regulated |
| enzyme | Down-regulated |
| enzyme | Down-regulated |
| transporter | Down-regulated |
| enzyme | Up-regulated |
| translation re | Down-regulated |
| other | Down-regulated |
| ion channel | Down-regulated |
| enzyme | Up-regulated |
| enzyme | Up-regulated |
| enzyme | Down-regulated |
| enzyme | Down-regulated |
| other | Down-regulated |
| G-protein co | Down-regulated |
| transporter | Down-regulated |
| other | Down-regulated |
| enzyme | Down-regulated |
| enzyme | Up-regulated |
| enzyme | Up-regulated |
| peptidase | Up-regulated |
| other | Down-regulated |
| transcription | Down-regulated |
| enzyme | Down-regulated |
| other | Down-regulated |
| enzyme | Down-regulated |
| other | Down-regulated |
| other | Down-regulated |
| other | Down-regulated |
| enzyme | Down-regulated |
| enzyme | Down-regulated |
| other | Down-regulated |
| transporter | Down-regulated |
| other | Down-regulated |
| other | Down-regulated |
| other | Down-regulated |
| enzyme | Down-regulated |
| enzyme | Down-regulated |
| enzyme | Down-regulated |
| other | Down-regulated |
| other | Down-regulated |

| | |
|---|---|
| kinase | Down-regulated |
| enzyme | Up-regulated |
| peptidase | Down-regulated |
| other | Down-regulated |
| other | Up-regulated |
| other | Down-regulated |
| enzyme | Up-regulated |
| transporter | Up-regulated |
| transcription | Down-regulated |
| other | Down-regulated |
| peptidase | Down-regulated |
| enzyme | Up-regulated |
| enzyme | Down-regulated |
| enzyme | Down-regulated |
| other | Down-regulated |
| other | Down-regulated |
| other | Down-regulated |
| enzyme | Up-regulated |
| enzyme | Down-regulated |
| other | Down-regulated |
| other | Down-regulated |
| other | Down-regulated |
| other | Down-regulated |
| other | Down-regulated |
| other | Down-regulated |
| other | Down-regulated |
| enzyme | Down-regulated |
| other | Up-regulated |
| transcription | Down-regulated |
| enzyme | Down-regulated |
| other | Down-regulated |
| other | Down-regulated |
| enzyme | Down-regulated |
| other | Down-regulated |
| other | Up-regulated |
| transcription | Down-regulated |
| kinase | Up-regulated |
| other | Down-regulated |
| enzyme | Down-regulated |
| translation re | Down-regulated |
| enzyme | Up-regulated |
| translation re | Down-regulated |
| transporter | Down-regulated |
| enzyme | Down-regulated |
| other | Down-regulated |
| transporter | Down-regulated |
| other | Down-regulated |
| other | Down-regulated |
| transporter | Down-regulated |
| enzyme | Up-regulated |
| transcription | Down-regulated |
| other | Down-regulated |
| other | Down-regulated |
| cytokine | Up-regulated |
| other | Down-regulated |
| other | Down-regulated |
| transcription | Down-regulated |
| other | Down-regulated |

| | |
|---|---|
| other | Up-regulated |
| other | Down-regulated |
| transporter | Down-regulated |
| peptidase | Down-regulated |
| other | Down-regulated |
| other | Down-regulated |
| enzyme | Down-regulated |
| enzyme | Down-regulated |
| phosphatase | Down-regulated |
| enzyme | Down-regulated |
| translation re | Down-regulated |
| transcription | Down-regulated |
| other | Down-regulated |
| other | Down-regulated |
| phosphatase | Down-regulated |
| transporter | Down-regulated |
| other | Down-regulated |
| other | Down-regulated |
| other | Down-regulated |
| kinase | Down-regulated |
| transporter | Down-regulated |
| other | Down-regulated |
| transcription | Down-regulated |
| other | Down-regulated |
| other | Down-regulated |
| other | Down-regulated |
| transcription | Down-regulated |
| translation re | Down-regulated |
| other | Up-regulated |
| other | Down-regulated |
| other | Down-regulated |
| other | Down-regulated |
| peptidase | Up-regulated |
| other | Down-regulated |
| other | Down-regulated |
| other | Down-regulated |
| transcription | Down-regulated |
| other | Down-regulated |
| transcription | Down-regulated |
| other | Down-regulated |
| transcription | Down-regulated |
| other | Down-regulated |
| enzyme | Down-regulated |
| enzyme | Down-regulated |
| transcription | Down-regulated |
| other | Down-regulated |
| enzyme | Down-regulated |
| other | Down-regulated |
| phosphatase | Down-regulated |
| enzyme | Down-regulated |
| enzyme | Down-regulated |
| other | Down-regulated |
| transcription | Down-regulated |
| transcription | Down-regulated |
| transcription | Down-regulated |
| transcription | Down-regulated |
| transporter | Up-regulated |
| transcription | Down-regulated |

| | |
|---|---|
| other | Down-regulated |
| enzyme | Down-regulated |
| other | Down-regulated |
| transcription | Down-regulated |
| enzyme | Down-regulated |
| kinase | Down-regulated |
| other | Down-regulated |
| enzyme | Down-regulated |
| enzyme | Up-regulated |
| enzyme | Up-regulated |
| transporter | Down-regulated |
| transcription | Down-regulated |
| enzyme | Down-regulated |
| transcription | Down-regulated |
| other | Down-regulated |
| other | Down-regulated |
| enzyme | Down-regulated |
| other | Down-regulated |
| other | Down-regulated |
| other | Down-regulated |
| other | Down-regulated |
| other | Down-regulated |
| other | Down-regulated |
| other | Down-regulated |
| enzyme | Down-regulated |
| transporter | Down-regulated |
| transcription | Down-regulated |
| enzyme | Up-regulated |
| translation re | Down-regulated |
| other | Down-regulated |
| other | Down-regulated |
| other | Down-regulated |
| other | Down-regulated |
| other | Down-regulated |
| transcription | Down-regulated |
| enzyme | Down-regulated |
| peptidase | Down-regulated |
| other | Down-regulated |
| other | Up-regulated |
| other | Down-regulated |
| other | Up-regulated |
| enzyme | Up-regulated |
| enzyme | Down-regulated |
| enzyme | Down-regulated |
| enzyme | Up-regulated |
| other | Up-regulated |
| other | Down-regulated |
| other | Down-regulated |
| other | Down-regulated |
| enzyme | Up-regulated |
| other | Down-regulated |
| other | Down-regulated |
| other | Down-regulated |
| other | Down-regulated |
| transcription | Down-regulated |
| other | Down-regulated |
| transporter | Up-regulated |
| peptidase | Up-regulated |

| | |
|---|---|
| enzyme | Down-regulated |
| transporter | Down-regulated |
| enzyme | Down-regulated |
| translation re | Down-regulated |
| enzyme | Up-regulated |
| enzyme | Down-regulated |
| enzyme | Down-regulated |
| kinase | Down-regulated |
| peptidase | Down-regulated |
| enzyme | Down-regulated |
| other | Down-regulated |
| other | Up-regulated |
| ion channel | Down-regulated |
| enzyme | Up-regulated |
| other | Down-regulated |
| other | Down-regulated |
| other | Down-regulated |
| enzyme | Up-regulated |
| transcription | Down-regulated |
| other | Down-regulated |
| other | Down-regulated |
| transporter | Down-regulated |
| other | Down-regulated |
| transporter | Down-regulated |
| enzyme | Down-regulated |
| other | Down-regulated |
| other | Down-regulated |
| enzyme | Down-regulated |
| transcription | Down-regulated |
| other | Down-regulated |
| other | Up-regulated |
| other | Up-regulated |
| translation re | Down-regulated |
| other | Down-regulated |
| enzyme | Down-regulated |
| other | Down-regulated |
| translation re | Down-regulated |
| kinase | Down-regulated |
| transporter | Down-regulated |
| transcription | Down-regulated |
| transcription | Down-regulated |
| other | Down-regulated |
| enzyme | Down-regulated |
| enzyme | Down-regulated |
| transcription | Down-regulated |
| transcription | Down-regulated |
| enzyme | Up-regulated |
| transmembra | Down-regulated |
| other | Down-regulated |
| enzyme | Down-regulated |
| translation re | Down-regulated |
| enzyme | Down-regulated |
| other | Down-regulated |
| transporter | Down-regulated |
| transcription | Down-regulated |
| other | Down-regulated |
| enzyme | Down-regulated |
| other | Down-regulated |

| | |
|---|---|
| other | Up-regulated |
| other | Down-regulated |
| transcription | Down-regulated |
| other | Down-regulated |
| peptidase | Down-regulated |
| other | Down-regulated |
| enzyme | Up-regulated |
| other | Down-regulated |
| enzyme | Down-regulated |
| transcription | Down-regulated |
| transporter | Down-regulated |
| enzyme | Up-regulated |
| other | Down-regulated |
| other | Down-regulated |
| other | Down-regulated |
| enzyme | Down-regulated |
| other | Down-regulated |
| other | Down-regulated |
| other | Down-regulated |
| enzyme | Up-regulated |
| other | Down-regulated |
| other | Down-regulated |
| other | Down-regulated |
| other | Down-regulated |
| enzyme | Down-regulated |
| other | Down-regulated |
| peptidase | Up-regulated |
| phosphatase | Down-regulated |
| other | Down-regulated |
| transcription | Down-regulated |
| translation re | Down-regulated |
| other | Down-regulated |
| other | Down-regulated |
| other | Up-regulated |
| other | Down-regulated |
| ion channel | Down-regulated |
| other | Down-regulated |
| enzyme | Down-regulated |
| transcription | Down-regulated |
| cytokine | Down-regulated |
| enzyme | Up-regulated |
| enzyme | Down-regulated |
| enzyme | Down-regulated |
| other | Down-regulated |
| peptidase | Up-regulated |
| other | Down-regulated |
| other | Down-regulated |
| other | Down-regulated |
| other | Down-regulated |
| other | Down-regulated |
| peptidase | Up-regulated |
| transmembra | Down-regulated |
| transporter | Up-regulated |
| translation re | Down-regulated |
| transcription | Down-regulated |
| transporter | Down-regulated |
| enzyme | Down-regulated |
| other | Down-regulated |

| | |
|---|---|
| other | Down-regulated |
| enzyme | Up-regulated |
| kinase | Down-regulated |
| other | Down-regulated |
| other | Down-regulated |
| other | Down-regulated |
| other | Down-regulated |
| enzyme | Down-regulated |
| other | Down-regulated |
| transporter | Up-regulated |
| other | Down-regulated |
| other | Down-regulated |
| enzyme | Down-regulated |
| other | Down-regulated |
| enzyme | Up-regulated |
| translation re | Down-regulated |
| other | Down-regulated |
| other | Down-regulated |
| other | Down-regulated |
| other | Down-regulated |
| phosphatase | Down-regulated |
| growth facto | Down-regulated |
| other | Down-regulated |
| kinase | Up-regulated |
| peptidase | Up-regulated |
| other | Down-regulated |
| peptidase | Down-regulated |
| other | Down-regulated |
| other | Down-regulated |
| phosphatase | Down-regulated |
| peptidase | Down-regulated |
| transcription | Down-regulated |
| other | Down-regulated |
| enzyme | Down-regulated |
| transcription | Down-regulated |
| enzyme | Down-regulated |
| kinase | Down-regulated |
| transmembra | Down-regulated |
| other | Down-regulated |
| kinase | Down-regulated |
| other | Down-regulated |
| enzyme | Down-regulated |
| enzyme | Down-regulated |
| other | Down-regulated |
| other | Down-regulated |
| transcription | Down-regulated |
| other | Down-regulated |
| peptidase | Down-regulated |
| enzyme | Up-regulated |
| cytokine | Down-regulated |
| other | Down-regulated |
| transporter | Down-regulated |
| other | Down-regulated |
| enzyme | Down-regulated |
| enzyme | Down-regulated |
| transcription | Down-regulated |
| enzyme | Down-regulated |
| enzyme | Down-regulated |

| | |
|---|---|
| other | Up-regulated |
| kinase | Down-regulated |
| other | Down-regulated |
| other | Down-regulated |
| other | Down-regulated |
| other | Down-regulated |
| other | Down-regulated |
| other | Down-regulated |
| other | Down-regulated |
| transporter | Down-regulated |
| transcription | Down-regulated |
| enzyme | Down-regulated |
| other | Down-regulated |
| transcription | Down-regulated |
| enzyme | Up-regulated |
| phosphatase | Down-regulated |
| transporter | Down-regulated |
| enzyme | Down-regulated |
| kinase | Down-regulated |
| transporter | Down-regulated |
| enzyme | Up-regulated |
| other | Down-regulated |
| transcription | Down-regulated |
| other | Down-regulated |
| other | Down-regulated |
| enzyme | Down-regulated |
| other | Down-regulated |
| other | Down-regulated |
| peptidase | Up-regulated |
| transcription | Up-regulated |
| enzyme | Up-regulated |
| other | Down-regulated |
| transporter | Down-regulated |
| other | Down-regulated |
| enzyme | Down-regulated |
| other | Down-regulated |
| transporter | Down-regulated |
| kinase | Down-regulated |
| enzyme | Up-regulated |
| other | Down-regulated |
| enzyme | Down-regulated |
| enzyme | Down-regulated |
| enzyme | Down-regulated |
| other | Down-regulated |
| enzyme | Down-regulated |
| enzyme | Down-regulated |
| transporter | Down-regulated |
| other | Down-regulated |
| other | Down-regulated |
| transcription | Down-regulated |
| other | Down-regulated |
| other | Down-regulated |
| enzyme | Down-regulated |
| other | Down-regulated |
| other | Down-regulated |
| translation re | Down-regulated |
| other | Down-regulated |
| transporter | Down-regulated |

| | |
|---|---|
| transporter | Up-regulated |
| other | Down-regulated |
| transcription | Down-regulated |
| transcription | Up-regulated |
| transcription | Down-regulated |
| other | Up-regulated |
| other | Down-regulated |
| enzyme | Down-regulated |
| other | Down-regulated |
| other | Down-regulated |
| enzyme | Down-regulated |
| enzyme | Down-regulated |
| transmembra | Down-regulated |
| enzyme | Down-regulated |
| enzyme | Down-regulated |
| other | Down-regulated |
| other | Down-regulated |
| other | Down-regulated |
| other | Up-regulated |
| enzyme | Down-regulated |
| other | Down-regulated |
| other | Down-regulated |
| other | Down-regulated |
| transporter | Down-regulated |
| enzyme | Up-regulated |
| enzyme | Down-regulated |
| other | Up-regulated |
| kinase | Down-regulated |
| other | Down-regulated |
| enzyme | Down-regulated |
| translation re | Down-regulated |
| enzyme | Down-regulated |
| enzyme | Up-regulated |
| enzyme | Down-regulated |
| enzyme | Up-regulated |
| other | Down-regulated |
| enzyme | Down-regulated |
| enzyme | Down-regulated |
| transcription | Down-regulated |
| enzyme | Up-regulated |
| G-protein co | Down-regulated |
| enzyme | Down-regulated |
| other | Down-regulated |
| enzyme | Down-regulated |
| peptidase | Down-regulated |
| other | Down-regulated |
| other | Down-regulated |
| other | Down-regulated |
| enzyme | Down-regulated |
| enzyme | Up-regulated |
| other | Up-regulated |
| peptidase | Up-regulated |
| other | Up-regulated |
| transmembra | Up-regulated |
| enzyme | Up-regulated |
| transcription | Down-regulated |
| enzyme | Up-regulated |
| transcription | Down-regulated |

| | |
|---|---|
| other | Down-regulated |
| other | Up-regulated |
| other | Down-regulated |
| kinase | Down-regulated |
| enzyme | Down-regulated |
| peptidase | Down-regulated |
| other | Down-regulated |
| transporter | Up-regulated |
| other | Down-regulated |
| enzyme | Up-regulated |
| other | Down-regulated |
| kinase | Down-regulated |
| transcription | Down-regulated |
| other | Down-regulated |
| other | Down-regulated |
| G-protein co | Down-regulated |
| transcription | Down-regulated |
| other | Down-regulated |
| transporter | Down-regulated |
| transcription | Down-regulated |
| other | Up-regulated |
| other | Up-regulated |
| other | Down-regulated |
| other | Down-regulated |
| other | Down-regulated |
| other | Down-regulated |
| other | Down-regulated |
| other | Down-regulated |
| other | Down-regulated |
| kinase | Down-regulated |
| transmembra | Down-regulated |
| enzyme | Down-regulated |
| other | Down-regulated |
| enzyme | Up-regulated |
| other | Down-regulated |
| other | Down-regulated |
| enzyme | Down-regulated |
| other | Down-regulated |
| translation re | Down-regulated |
| other | Down-regulated |
| other | Down-regulated |
| other | Down-regulated |
| other | Down-regulated |
| enzyme | Down-regulated |
| other | Down-regulated |
| phosphatase | Down-regulated |
| other | Down-regulated |
| other | Down-regulated |
| transporter | Down-regulated |
| peptidase | Down-regulated |
| other | Down-regulated |
| kinase | Down-regulated |
| enzyme | Down-regulated |
| growth facto | Up-regulated |
| other | Down-regulated |
| other | Down-regulated |
| other | Down-regulated |
| enzyme | Down-regulated |

| | |
|---|---|
| enzyme | Down-regulated |
| transcription | Down-regulated |
| transporter | Down-regulated |
| other | Down-regulated |
| enzyme | Up-regulated |
| other | Down-regulated |
| transcription | Down-regulated |
| other | Down-regulated |
| transporter | Down-regulated |
| other | Down-regulated |
| growth facto | Down-regulated |
| other | Down-regulated |
| other | Down-regulated |
| transcription | Down-regulated |
| other | Down-regulated |
| kinase | Down-regulated |
| other | Down-regulated |
| peptidase | Down-regulated |
| other | Down-regulated |
| enzyme | Up-regulated |
| enzyme | Up-regulated |
| other | Up-regulated |
| other | Down-regulated |
| transporter | Up-regulated |
| other | Down-regulated |
| enzyme | Down-regulated |
| translation re | Down-regulated |
| cytokine | Up-regulated |
| other | Down-regulated |
| kinase | Down-regulated |
| enzyme | Up-regulated |
| other | Down-regulated |
| enzyme | Down-regulated |
| other | Down-regulated |
| enzyme | Down-regulated |
| phosphatase | Down-regulated |
| transcription | Down-regulated |
| other | Down-regulated |
| enzyme | Up-regulated |
| other | Down-regulated |
| transporter | Down-regulated |
| peptidase | Down-regulated |
| other | Down-regulated |
| enzyme | Down-regulated |
| other | Down-regulated |
| other | Down-regulated |
| enzyme | Down-regulated |
| enzyme | Down-regulated |
| other | Down-regulated |
| other | Down-regulated |
| enzyme | Down-regulated |
| enzyme | Down-regulated |
| other | Down-regulated |
| other | Down-regulated |
| other | Up-regulated |
| enzyme | Up-regulated |
| other | Down-regulated |
| other | Down-regulated |

| | |
|---|---|
| enzyme | Down-regulated |
| other | Down-regulated |
| transporter | Down-regulated |
| transcription | Down-regulated |
| other | Down-regulated |
| other | Down-regulated |
| enzyme | Down-regulated |
| transcription | Down-regulated |
| ion channel | Up-regulated |
| other | Down-regulated |
| other | Down-regulated |
| enzyme | Down-regulated |
| transcription | Down-regulated |
| other | Down-regulated |
| other | Down-regulated |
| transcription | Down-regulated |
| other | Down-regulated |
| peptidase | Up-regulated |
| phosphatase | Down-regulated |
| peptidase | Down-regulated |
| enzyme | Down-regulated |
| other | Down-regulated |
| other | Down-regulated |
| other | Down-regulated |
| enzyme | Up-regulated |
| other | Down-regulated |
| kinase | Down-regulated |
| enzyme | Down-regulated |
| enzyme | Up-regulated |
| transcription | Down-regulated |
| enzyme | Down-regulated |
| other | Down-regulated |
| enzyme | Down-regulated |
| peptidase | Down-regulated |
| other | Down-regulated |
| other | Down-regulated |
| other | Down-regulated |
| other | Down-regulated |
| enzyme | Down-regulated |
| other | Up-regulated |
| other | Down-regulated |
| other | Down-regulated |
| other | Down-regulated |
| other | Down-regulated |
| other | Down-regulated |
| other | Down-regulated |
| other | Down-regulated |
| other | Down-regulated |
| enzyme | Down-regulated |
| enzyme | Down-regulated |
| other | Down-regulated |
| enzyme | Up-regulated |
| other | Down-regulated |
| enzyme | Down-regulated |
| transporter | Down-regulated |
| kinase | Down-regulated |
| other | Down-regulated |
| enzyme | Down-regulated |

| | |
|---|---|
| transcription | Down-regulated |
| transcription | Down-regulated |
| transcription | Down-regulated |
| other | Down-regulated |
| peptidase | Down-regulated |
| other | Down-regulated |
| other | Down-regulated |
| other | Down-regulated |
| other | Down-regulated |
| other | Down-regulated |
| kinase | Down-regulated |
| other | Down-regulated |
| other | Down-regulated |
| other | Down-regulated |
| enzyme | Down-regulated |
| other | Down-regulated |
| other | Down-regulated |
| other | Down-regulated |
| other | Down-regulated |
| enzyme | Down-regulated |
| transporter | Down-regulated |
| enzyme | Down-regulated |
| enzyme | Down-regulated |
| transporter | Up-regulated |
| other | Down-regulated |
| ion channel | Down-regulated |
| other | Down-regulated |
| other | Down-regulated |
| other | Down-regulated |
| other | Up-regulated |
| enzyme | Down-regulated |
| enzyme | Down-regulated |
| enzyme | Down-regulated |
| other | Up-regulated |
| transcription | Down-regulated |
| other | Down-regulated |
| enzyme | Up-regulated |
| enzyme | Down-regulated |
| peptidase | Up-regulated |
| other | Down-regulated |
| other | Down-regulated |
| enzyme | Down-regulated |
| transcription | Down-regulated |
| other | Down-regulated |
| other | Down-regulated |
| enzyme | Down-regulated |
| enzyme | Down-regulated |
| other | Down-regulated |
| enzyme | Down-regulated |
| other | Down-regulated |
| other | Down-regulated |
| enzyme | Down-regulated |
| enzyme | Down-regulated |
| other | Down-regulated |
| other | Down-regulated |
| transcription | Down-regulated |
| other | Down-regulated |
| other | Down-regulated |

| | |
|---|---|
| other | Down-regulated |
| transcription | Down-regulated |
| other | Down-regulated |
| enzyme | Down-regulated |
| transcription | Down-regulated |
| transcription | Down-regulated |
| enzyme | Up-regulated |
| other | Down-regulated |
| enzyme | Down-regulated |
| transcription | Down-regulated |
| transporter | Up-regulated |
| phosphatase | Down-regulated |
| other | Down-regulated |
| transcription | Down-regulated |
| peptidase | Up-regulated |
| other | Down-regulated |
| other | Down-regulated |
| kinase | Down-regulated |
| enzyme | Down-regulated |
| enzyme | Down-regulated |
| transporter | Up-regulated |
| kinase | Down-regulated |
| transporter | Up-regulated |
| other | Down-regulated |
| other | Down-regulated |
| enzyme | Down-regulated |
| enzyme | Down-regulated |
| transcription | Down-regulated |
| peptidase | Down-regulated |
| enzyme | Up-regulated |
| transcription | Down-regulated |
| other | Down-regulated |
| transcription | Down-regulated |
| other | Down-regulated |
| other | Down-regulated |
| other | Down-regulated |
| enzyme | Down-regulated |
| enzyme | Up-regulated |
| other | Down-regulated |
| other | Down-regulated |
| other | Up-regulated |
| transcription | Down-regulated |
| other | Down-regulated |
| other | Down-regulated |
| enzyme | Down-regulated |
| other | Down-regulated |
| other | Down-regulated |
| other | Down-regulated |
| other | Down-regulated |
| other | Down-regulated |
| other | Down-regulated |
| other | Down-regulated |
| other | Down-regulated |
| enzyme | Down-regulated |
| enzyme | Down-regulated |
| other | Down-regulated |
| enzyme | Down-regulated |
| other | Down-regulated |

| | |
|---|---|
| other | Down-regulated |
| enzyme | Down-regulated |
| transcription | Down-regulated |
| kinase | Down-regulated |
| other | Down-regulated |
| other | Down-regulated |
| kinase | Down-regulated |
| enzyme | Down-regulated |
| enzyme | Down-regulated |
| other | Down-regulated |
| other | Down-regulated |
| kinase | Down-regulated |
| other | Down-regulated |
| transporter | Down-regulated |
| other | Down-regulated |
| transcription | Down-regulated |
| other | Up-regulated |
| enzyme | Down-regulated |
| enzyme | Down-regulated |
| other | Down-regulated |
| enzyme | Up-regulated |
| enzyme | Down-regulated |
| transporter | Up-regulated |
| other | Up-regulated |
| other | Down-regulated |
| other | Down-regulated |
| other | Down-regulated |
| other | Down-regulated |
| other | Down-regulated |
| enzyme | Down-regulated |
| other | Down-regulated |
| other | Down-regulated |
| kinase | Down-regulated |
| enzyme | Down-regulated |
| transporter | Down-regulated |
| other | Down-regulated |
| other | Down-regulated |
| transcription | Down-regulated |
| enzyme | Up-regulated |
| other | Down-regulated |
| other | Down-regulated |
| other | Up-regulated |
| other | Down-regulated |
| transporter | Down-regulated |
| other | Down-regulated |
| other | Down-regulated |
| kinase | Down-regulated |
| enzyme | Up-regulated |
| kinase | Down-regulated |
| other | Down-regulated |
| enzyme | Down-regulated |
| other | Down-regulated |
| enzyme | Down-regulated |
| other | Down-regulated |
| other | Down-regulated |
| enzyme | Down-regulated |
| other | Down-regulated |
| other | Down-regulated |

| | |
|---|---|
| enzyme | Down-regulated |
| transcription | Down-regulated |
| other | Down-regulated |
| transcription | Down-regulated |
| transcription | Down-regulated |
| other | Down-regulated |
| transcription | Down-regulated |
| enzyme | Down-regulated |
| transcription | Down-regulated |
| other | Down-regulated |
| other | Down-regulated |
| other | Down-regulated |
| other | Down-regulated |
| enzyme | Down-regulated |
| enzyme | Down-regulated |
| other | Down-regulated |
| transporter | Down-regulated |
| other | Down-regulated |
| transcription | Down-regulated |
| other | Down-regulated |
| other | Down-regulated |
| other | Down-regulated |
| enzyme | Down-regulated |
| enzyme | Up-regulated |
| enzyme | Down-regulated |
| enzyme | Down-regulated |
| other | Down-regulated |
| phosphatase | Down-regulated |
| other | Down-regulated |
| other | Down-regulated |
| other | Down-regulated |
| enzyme | Down-regulated |
| other | Down-regulated |
| other | Down-regulated |
| peptidase | Down-regulated |
| other | Down-regulated |
| other | Down-regulated |
| other | Down-regulated |
| transporter | Down-regulated |
| other | Down-regulated |
| transcription | Down-regulated |
| other | Down-regulated |
| enzyme | Down-regulated |
| transporter | Down-regulated |
| transcription | Down-regulated |
| other | Down-regulated |
| other | Down-regulated |
| other | Down-regulated |
| enzyme | Down-regulated |
| other | Up-regulated |
| other | Down-regulated |
| other | Down-regulated |
| transcription | Down-regulated |
| other | Down-regulated |
| enzyme | Up-regulated |
| enzyme | Down-regulated |
| enzyme | Down-regulated |
| other | Down-regulated |

| | |
|---|---|
| kinase | Up-regulated |
| other | Down-regulated |
| enzyme | Up-regulated |
| other | Down-regulated |
| other | Down-regulated |
| other | Down-regulated |
| enzyme | Down-regulated |
| transcription | Down-regulated |
| other | Down-regulated |
| other | Down-regulated |
| phosphatase | Down-regulated |
| other | Down-regulated |
| other | Up-regulated |
| other | Up-regulated |
| enzyme | Down-regulated |
| enzyme | Up-regulated |
| enzyme | Down-regulated |
| transcription | Down-regulated |
| transporter | Down-regulated |
| enzyme | Down-regulated |
| enzyme | Up-regulated |
| phosphatase | Down-regulated |
| enzyme | Down-regulated |
| transcription | Down-regulated |
| peptidase | Down-regulated |
| other | Down-regulated |
| other | Up-regulated |
| enzyme | Down-regulated |
| enzyme | Up-regulated |
| transmembra | Down-regulated |
| other | Down-regulated |
| other | Down-regulated |
| other | Down-regulated |
| transcription | Down-regulated |
| enzyme | Down-regulated |
| other | Down-regulated |
| peptidase | Down-regulated |
| enzyme | Down-regulated |
| enzyme | Up-regulated |
| enzyme | Up-regulated |
| transcription | Down-regulated |
| other | Down-regulated |
| other | Down-regulated |
| other | Down-regulated |
| transcription | Up-regulated |
| other | Down-regulated |
| enzyme | Down-regulated |
| enzyme | Down-regulated |
| other | Down-regulated |
| other | Down-regulated |
| phosphatase | Down-regulated |
| enzyme | Down-regulated |
| enzyme | Down-regulated |
| other | Down-regulated |
| other | Down-regulated |
| other | Down-regulated |
| other | Up-regulated |
| other | Down-regulated |

| | |
|---|---|
| other | Down-regulated |
| enzyme | Down-regulated |
| other | Down-regulated |
| other | Down-regulated |
| enzyme | Down-regulated |
| other | Down-regulated |
| enzyme | Down-regulated |
| transporter | Up-regulated |
| other | Down-regulated |
| other | Down-regulated |
| other | Down-regulated |
| transcription | Down-regulated |
| other | Down-regulated |
| other | Down-regulated |
| other | Down-regulated |
| other | Down-regulated |
| enzyme | Down-regulated |
| other | Down-regulated |
| other | Down-regulated |
| enzyme | Down-regulated |
| other | Down-regulated |
| enzyme | Down-regulated |
| other | Down-regulated |
| enzyme | Down-regulated |
| enzyme | Up-regulated |
| transcription | Down-regulated |
| peptidase | Down-regulated |
| other | Down-regulated |
| enzyme | Down-regulated |
| other | Down-regulated |
| other | Down-regulated |
| transcription | Down-regulated |
| kinase | Down-regulated |
| other | Down-regulated |
| transcription | Down-regulated |
| transcription | Down-regulated |
| enzyme | Down-regulated |
| transcription | Down-regulated |
| enzyme | Down-regulated |
| other | Down-regulated |
| other | Down-regulated |
| transcription | Down-regulated |
| other | Up-regulated |
| enzyme | Down-regulated |
| other | Down-regulated |
| other | Down-regulated |
| other | Down-regulated |
| other | Down-regulated |
| other | Down-regulated |
| peptidase | Down-regulated |
| enzyme | Down-regulated |
| peptidase | Down-regulated |
| other | Down-regulated |
| other | Down-regulated |
| other | Down-regulated |
| other | Down-regulated |
| kinase | Up-regulated |
| other | Down-regulated |

| | |
|---|---|
| kinase | Down-regulated |
| transporter | Up-regulated |
| other | Down-regulated |
| transcription | Down-regulated |
| transcription | Down-regulated |
| other | Down-regulated |
| transcription | Down-regulated |
| enzyme | Down-regulated |
| transporter | Up-regulated |
| kinase | Down-regulated |
| transcription | Down-regulated |
| other | Down-regulated |
| other | Down-regulated |
| kinase | Down-regulated |
| other | Up-regulated |
| other | Down-regulated |
| enzyme | Down-regulated |
| other | Down-regulated |
| other | Down-regulated |
| other | Down-regulated |
| transporter | Down-regulated |
| transcription | Down-regulated |
| kinase | Down-regulated |
| enzyme | Down-regulated |
| other | Down-regulated |
| other | Down-regulated |
| transcription | Down-regulated |
| enzyme | Up-regulated |
| other | Down-regulated |
| transporter | Down-regulated |
| other | Down-regulated |
| other | Down-regulated |
| enzyme | Down-regulated |
| other | Down-regulated |
| other | Down-regulated |
| phosphatase | Down-regulated |
| other | Down-regulated |
| enzyme | Down-regulated |
| other | Down-regulated |
| enzyme | Down-regulated |
| kinase | Up-regulated |
| enzyme | Down-regulated |
| transcription | Down-regulated |
| other | Down-regulated |
| transporter | Down-regulated |
| transcription | Down-regulated |
| enzyme | Down-regulated |
| other | Down-regulated |
| enzyme | Up-regulated |
| other | Down-regulated |
| transcription | Down-regulated |
| transcription | Down-regulated |
| transcription | Down-regulated |
| other | Down-regulated |
| enzyme | Down-regulated |
| enzyme | Down-regulated |
| enzyme | Down-regulated |
| transmembra | Down-regulated |

| | |
|---|---|
| transporter | Down-regulated |
| phosphatase | Down-regulated |
| enzyme | Down-regulated |
| other | Down-regulated |
| other | Down-regulated |
| other | Up-regulated |
| transcription | Down-regulated |
| other | Down-regulated |
| other | Down-regulated |
| other | Down-regulated |
| other | Down-regulated |
| enzyme | Down-regulated |
| enzyme | Down-regulated |
| other | Down-regulated |
| enzyme | Down-regulated |
| other | Down-regulated |
| other | Down-regulated |
| other | Up-regulated |
| enzyme | Down-regulated |
| other | Down-regulated |
| peptidase | Down-regulated |
| other | Up-regulated |
| other | Down-regulated |
| other | Down-regulated |
| enzyme | Up-regulated |
| other | Up-regulated |
| translation re | Down-regulated |
| translation re | Down-regulated |
| other | Down-regulated |
| transcription | Down-regulated |
| other | Down-regulated |
| enzyme | Down-regulated |
| other | Down-regulated |
| other | Down-regulated |
| other | Down-regulated |
| enzyme | Down-regulated |
| other | Down-regulated |
| transcription | Down-regulated |
| ion channel | Up-regulated |
| enzyme | Down-regulated |
| other | Down-regulated |
| enzyme | Down-regulated |
| enzyme | Up-regulated |
| enzyme | Down-regulated |
| other | Down-regulated |
| enzyme | Up-regulated |
| transcription | Down-regulated |
| transcription | Down-regulated |
| other | Down-regulated |
| other | Down-regulated |
| other | Down-regulated |
| other | Down-regulated |
| other | Up-regulated |
| other | Down-regulated |
| other | Down-regulated |
| other | Down-regulated |
| other | Down-regulated |
| other | Down-regulated |

| | |
|---|---|
| enzyme | Down-regulated |
| enzyme | Down-regulated |
| other | Down-regulated |
| transporter | Down-regulated |
| growth facto | Up-regulated |
| cytokine | Down-regulated |
| other | Down-regulated |
| other | Down-regulated |
| enzyme | Down-regulated |
| other | Down-regulated |
| other | Down-regulated |
| enzyme | Down-regulated |
| other | Down-regulated |
| other | Down-regulated |
| other | Down-regulated |
| enzyme | Down-regulated |
| transporter | Down-regulated |
| other | Up-regulated |
| cytokine | Up-regulated |
| kinase | Up-regulated |
| other | Down-regulated |
| ligand-deper | Up-regulated |
| other | Down-regulated |
| other | Down-regulated |
| other | Down-regulated |
| other | Down-regulated |
| other | Down-regulated |
| enzyme | Up-regulated |
| other | Down-regulated |
| other | Down-regulated |
| enzyme | Up-regulated |
| other | Down-regulated |
| transporter | Down-regulated |
| other | Down-regulated |
| other | Down-regulated |
| other | Down-regulated |
| enzyme | Up-regulated |
| other | Down-regulated |
| enzyme | Up-regulated |
| enzyme | Down-regulated |
| transcription | Down-regulated |
| other | Down-regulated |
| transcription | Down-regulated |
| other | Down-regulated |
| other | Down-regulated |
| other | Down-regulated |
| other | Down-regulated |
| other | Down-regulated |
| other | Down-regulated |
| enzyme | Up-regulated |
| other | Down-regulated |
| enzyme | Down-regulated |
| other | Down-regulated |
| enzyme | Up-regulated |
| enzyme | Up-regulated |
| transcription | Down-regulated |
| enzyme | Up-regulated |
| enzyme | Down-regulated |

| | |
|---|---|
| enzyme | Up-regulated |
| other | Down-regulated |
| other | Down-regulated |
| transcription | Down-regulated |
| enzyme | Down-regulated |
| transcription | Down-regulated |
| ion channel | Down-regulated |
| other | Down-regulated |
| growth facto | Down-regulated |
| transcription | Down-regulated |
| other | Down-regulated |
| other | Down-regulated |
| other | Down-regulated |
| peptidase | Down-regulated |
| other | Down-regulated |
| transcription | Down-regulated |
| other | Up-regulated |
| other | Down-regulated |
| enzyme | Down-regulated |
| other | Down-regulated |
| other | Down-regulated |
| other | Down-regulated |
| peptidase | Down-regulated |
| transcription | Down-regulated |
| transcription | Down-regulated |
| other | Down-regulated |
| enzyme | Down-regulated |
| other | Down-regulated |
| other | Down-regulated |
| enzyme | Down-regulated |
| transcription | Down-regulated |
| transcription | Down-regulated |
| other | Down-regulated |
| kinase | Down-regulated |
| growth facto | Down-regulated |
| other | Down-regulated |
| other | Down-regulated |
| transporter | Down-regulated |
| transporter | Down-regulated |
| transcription | Down-regulated |
| other | Down-regulated |
| kinase | Down-regulated |
| other | Down-regulated |
| other | Down-regulated |
| other | Down-regulated |
| kinase | Down-regulated |
| transcription | Down-regulated |
| enzyme | Up-regulated |
| other | Down-regulated |
| other | Down-regulated |
| peptidase | Down-regulated |
| other | Down-regulated |
| enzyme | Down-regulated |
| other | Down-regulated |
| phosphatase | Down-regulated |
| other | Up-regulated |
| transporter | Down-regulated |
| transcription | Down-regulated |

| | |
|---|---|
| enzyme | Down-regulated |
| other | Down-regulated |
| enzyme | Down-regulated |
| enzyme | Down-regulated |
| enzyme | Up-regulated |
| enzyme | Down-regulated |
| transporter | Down-regulated |
| other | Down-regulated |
| other | Down-regulated |
| transporter | Down-regulated |
| other | Down-regulated |
| other | Down-regulated |
| other | Down-regulated |
| other | Down-regulated |
| other | Down-regulated |
| other | Down-regulated |
| peptidase | Down-regulated |
| phosphatase | Down-regulated |
| phosphatase | Down-regulated |
| enzyme | Down-regulated |
| enzyme | Down-regulated |
| other | Down-regulated |
| kinase | Down-regulated |
| other | Down-regulated |
| enzyme | Down-regulated |
| transcription | Down-regulated |
| enzyme | Up-regulated |
| peptidase | Down-regulated |
| other | Down-regulated |
| other | Down-regulated |
| enzyme | Down-regulated |
| other | Down-regulated |
| other | Down-regulated |
| other | Down-regulated |
| other | Down-regulated |
| enzyme | Down-regulated |
| transporter | Up-regulated |
| enzyme | Down-regulated |
| transporter | Down-regulated |
| transcription | Down-regulated |
| transporter | Down-regulated |
| peptidase | Down-regulated |
| other | Up-regulated |
| enzyme | Down-regulated |
| other | Down-regulated |
| enzyme | Down-regulated |
| kinase | Down-regulated |
| other | Down-regulated |
| other | Down-regulated |
| transporter | Down-regulated |
| other | Down-regulated |
| transporter | Down-regulated |
| other | Down-regulated |
| other | Down-regulated |
| other | Down-regulated |
| other | Up-regulated |
| enzyme | Down-regulated |
| other | Down-regulated |

| | |
|---|---|
| other | Down-regulated |
| other | Down-regulated |
| other | Down-regulated |
| other | Down-regulated |
| other | Down-regulated |
| translation re | Down-regulated |
| enzyme | Down-regulated |
| enzyme | Down-regulated |
| transcription | Down-regulated |
| translation re | Down-regulated |
| peptidase | Down-regulated |
| other | Down-regulated |
| other | Down-regulated |
| other | Down-regulated |
| enzyme | Up-regulated |
| transcription | Down-regulated |
| other | Down-regulated |
| enzyme | Down-regulated |
| other | Down-regulated |
| other | Down-regulated |
| other | Up-regulated |
| other | Down-regulated |
| other | Down-regulated |
| transcription | Up-regulated |
| transcription | Down-regulated |
| other | Down-regulated |
| transcription | Down-regulated |
| other | Down-regulated |
| other | Down-regulated |
| other | Down-regulated |
| other | Down-regulated |
| other | Down-regulated |
| enzyme | Up-regulated |
| enzyme | Down-regulated |
| enzyme | Up-regulated |
| transporter | Down-regulated |
| enzyme | Down-regulated |
| other | Down-regulated |
| other | Down-regulated |
| transcription | Down-regulated |
| other | Down-regulated |
| ion channel | Down-regulated |
| other | Down-regulated |
| transporter | Up-regulated |
| translation re | Down-regulated |
| other | Down-regulated |
| enzyme | Down-regulated |
| transporter | Down-regulated |
| other | Down-regulated |
| enzyme | Down-regulated |
| transporter | Down-regulated |
| other | Down-regulated |
| transporter | Down-regulated |
| other | Down-regulated |
| transcription | Down-regulated |
| other | Down-regulated |
| other | Down-regulated |
| other | Down-regulated |

| | |
|---|---|
| other | Down-regulated |
| enzyme | Down-regulated |
| transcription | Down-regulated |
| enzyme | Down-regulated |
| transcription | Down-regulated |
| other | Down-regulated |
| other | Down-regulated |
| other | Down-regulated |
| enzyme | Down-regulated |
| enzyme | Down-regulated |
| other | Down-regulated |
| peptidase | Down-regulated |
| other | Down-regulated |
| other | Down-regulated |
| other | Down-regulated |
| other | Down-regulated |
| peptidase | Up-regulated |
| enzyme | Down-regulated |
| other | Down-regulated |
| other | Down-regulated |
| kinase | Down-regulated |
| other | Down-regulated |
| kinase | Down-regulated |
| enzyme | Down-regulated |
| phosphatase | Down-regulated |
| transcription | Down-regulated |
| enzyme | Down-regulated |
| other | Down-regulated |
| other | Down-regulated |
| transcription | Down-regulated |
| transcription | Down-regulated |
| other | Down-regulated |
| other | Down-regulated |
| kinase | Down-regulated |
| other | Down-regulated |
| other | Up-regulated |
| transporter | Down-regulated |
| other | Down-regulated |
| peptidase | Down-regulated |
| enzyme | Down-regulated |
| other | Down-regulated |
| enzyme | Down-regulated |
| enzyme | Down-regulated |
| other | Down-regulated |
| peptidase | Up-regulated |
| other | Down-regulated |
| other | Down-regulated |
| other | Down-regulated |
| ion channel | Down-regulated |
| other | Down-regulated |
| enzyme | Down-regulated |
| other | Down-regulated |
| kinase | Down-regulated |
| peptidase | Down-regulated |
| enzyme | Up-regulated |
| other | Down-regulated |
| other | Down-regulated |
| other | Down-regulated |

| | |
|---|---|
| transporter | Down-regulated |
| other | Down-regulated |
| enzyme | Down-regulated |
| enzyme | Up-regulated |
| other | Down-regulated |
| other | Down-regulated |
| other | Up-regulated |
| enzyme | Down-regulated |
| other | Down-regulated |
| other | Down-regulated |
| enzyme | Down-regulated |
| transcription | Down-regulated |
| enzyme | Down-regulated |
| peptidase | Down-regulated |
| enzyme | Down-regulated |
| other | Down-regulated |
| other | Down-regulated |
| transcription | Down-regulated |
| kinase | Down-regulated |
| peptidase | Down-regulated |
| enzyme | Down-regulated |
| other | Down-regulated |
| transmembra | Down-regulated |
| transporter | Down-regulated |
| enzyme | Up-regulated |
| transporter | Down-regulated |
| transcription | Down-regulated |
| G-protein co | Down-regulated |
| other | Down-regulated |
| transcription | Down-regulated |
| kinase | Down-regulated |
| other | Up-regulated |
| other | Down-regulated |
| cytokine | Down-regulated |
| enzyme | Down-regulated |
| enzyme | Up-regulated |
| other | Down-regulated |
| other | Down-regulated |
| other | Down-regulated |
| other | Down-regulated |
| phosphatase | Down-regulated |
| other | Down-regulated |
| transcription | Down-regulated |
| enzyme | Up-regulated |
| enzyme | Up-regulated |
| enzyme | Down-regulated |
| other | Down-regulated |
| other | Down-regulated |
| kinase | Down-regulated |
| enzyme | Down-regulated |
| transporter | Down-regulated |
| other | Down-regulated |
| other | Down-regulated |
| enzyme | Down-regulated |
| enzyme | Down-regulated |
| other | Down-regulated |
| transporter | Up-regulated |
| transcription | Down-regulated |

| | |
|---|---|
| other | Down-regulated |
| other | Down-regulated |
| kinase | Up-regulated |
| transmembra | Up-regulated |
| other | Down-regulated |
| other | Down-regulated |
| other | Down-regulated |
| other | Down-regulated |
| kinase | Down-regulated |
| enzyme | Down-regulated |
| other | Down-regulated |
| other | Down-regulated |
| enzyme | Down-regulated |
| other | Down-regulated |
| other | Down-regulated |
| other | Down-regulated |
| enzyme | Down-regulated |
| other | Down-regulated |
| other | Down-regulated |
| other | Down-regulated |
| other | Down-regulated |
| kinase | Down-regulated |
| enzyme | Down-regulated |
| other | Down-regulated |
| other | Down-regulated |
| enzyme | Down-regulated |
| other | Down-regulated |
| transcription | Down-regulated |
| other | Down-regulated |
| other | Down-regulated |
| other | Down-regulated |
| transporter | Down-regulated |
| enzyme | Up-regulated |
| enzyme | Down-regulated |
| transporter | Down-regulated |
| other | Down-regulated |
| other | Down-regulated |
| other | Down-regulated |
| other | Down-regulated |
| kinase | Down-regulated |
| kinase | Down-regulated |
| other | Down-regulated |
| transcription | Down-regulated |
| transcription | Down-regulated |
| enzyme | Down-regulated |
| ion channel | Down-regulated |
| other | Down-regulated |
| other | Down-regulated |
| other | Down-regulated |
| transcription | Down-regulated |
| transcription | Down-regulated |
| enzyme | Down-regulated |
| other | Down-regulated |
| other | Down-regulated |
| other | Up-regulated |
| enzyme | Down-regulated |
| other | Down-regulated |
| ion channel | Down-regulated |

| | |
|---|---|
| transporter | Up-regulated |
| other | Down-regulated |
| other | Down-regulated |
| other | Up-regulated |
| other | Down-regulated |
| other | Down-regulated |
| enzyme | Down-regulated |
| ion channel | Down-regulated |
| other | Up-regulated |
| other | Down-regulated |
| transporter | Down-regulated |
| other | Down-regulated |
| other | Up-regulated |
| transcription | Down-regulated |
| other | Down-regulated |
| other | Down-regulated |
| enzyme | Down-regulated |
| other | Down-regulated |
| ion channel | Down-regulated |
| other | Down-regulated |
| enzyme | Down-regulated |
| other | Down-regulated |
| transcription | Down-regulated |
| enzyme | Down-regulated |
| other | Down-regulated |
| enzyme | Down-regulated |
| other | Up-regulated |
| other | Down-regulated |
| other | Down-regulated |
| other | Down-regulated |
| enzyme | Up-regulated |
| enzyme | Down-regulated |
| transcription | Down-regulated |
| transcription | Down-regulated |
| enzyme | Down-regulated |
| other | Down-regulated |
| other | Down-regulated |
| other | Down-regulated |
| other | Down-regulated |
| other | Down-regulated |
| kinase | Down-regulated |
| other | Down-regulated |
| enzyme | Down-regulated |
| enzyme | Up-regulated |
| enzyme | Up-regulated |
| other | Down-regulated |
| other | Down-regulated |
| peptidase | Down-regulated |
| phosphatase | Down-regulated |
| other | Down-regulated |
| enzyme | Up-regulated |
| other | Down-regulated |
| enzyme | Down-regulated |
| transporter | Down-regulated |
| other | Down-regulated |
| other | Down-regulated |
| translation re | Down-regulated |
| enzyme | Down-regulated |

| | |
|---|---|
| other | Down-regulated |
| ion channel | Down-regulated |
| other | Down-regulated |
| transmembra | Up-regulated |
| transcription | Down-regulated |
| enzyme | Down-regulated |
| peptidase | Down-regulated |
| other | Down-regulated |
| transcription | Down-regulated |
| enzyme | Down-regulated |
| enzyme | Down-regulated |
| other | Down-regulated |
| enzyme | Down-regulated |
| enzyme | Up-regulated |
| other | Down-regulated |
| enzyme | Down-regulated |
| enzyme | Down-regulated |
| transporter | Down-regulated |
| other | Down-regulated |
| other | Down-regulated |
| other | Down-regulated |
| transporter | Down-regulated |
| other | Down-regulated |
| G-protein co | Up-regulated |
| phosphatase | Down-regulated |
| transmembra | Up-regulated |
| enzyme | Down-regulated |
| phosphatase | Down-regulated |
| enzyme | Up-regulated |
| cytokine | Down-regulated |
| enzyme | Down-regulated |
| other | Down-regulated |
| enzyme | Down-regulated |
| other | Down-regulated |
| enzyme | Down-regulated |
| other | Down-regulated |
| other | Down-regulated |
| other | Up-regulated |
| other | Up-regulated |
| other | Up-regulated |
| translation re | Down-regulated |
| enzyme | Up-regulated |
| enzyme | Down-regulated |
| other | Down-regulated |
| other | Down-regulated |
| transcription | Down-regulated |
| other | Down-regulated |
| transcription | Down-regulated |
| transcription | Down-regulated |
| other | Down-regulated |
| other | Down-regulated |
| other | Down-regulated |
| other | Down-regulated |
| translation re | Down-regulated |
| enzyme | Down-regulated |
| other | Down-regulated |
| enzyme | Down-regulated |
| other | Down-regulated |

| | |
|---|---|
| transcription | Down-regulated |
| enzyme | Down-regulated |
| other | Down-regulated |
| enzyme | Down-regulated |
| transcription | Down-regulated |
| other | Down-regulated |
| other | Down-regulated |
| transcription | Down-regulated |
| enzyme | Down-regulated |
| other | Down-regulated |
| enzyme | Down-regulated |
| other | Down-regulated |
| other | Down-regulated |
| other | Down-regulated |
| enzyme | Down-regulated |
| other | Down-regulated |
| transcription | Down-regulated |
| other | Down-regulated |
| other | Down-regulated |
| transcription | Down-regulated |
| other | Up-regulated |
| kinase | Down-regulated |
| enzyme | Down-regulated |
| other | Down-regulated |
| other | Down-regulated |
| peptidase | Down-regulated |
| cytokine | Down-regulated |
| other | Down-regulated |
| enzyme | Down-regulated |
| other | Down-regulated |
| other | Down-regulated |
| other | Down-regulated |
| other | Down-regulated |
| transcription | Down-regulated |
| transcription | Down-regulated |
| enzyme | Down-regulated |
| peptidase | Down-regulated |
| enzyme | Down-regulated |
| other | Down-regulated |
| other | Down-regulated |
| other | Up-regulated |
| transcription | Down-regulated |
| enzyme | Down-regulated |
| other | Down-regulated |
| enzyme | Down-regulated |
| enzyme | Down-regulated |
| peptidase | Down-regulated |
| transcription | Down-regulated |
| transcription | Down-regulated |
| transcription | Down-regulated |
| transporter | Down-regulated |
| other | Down-regulated |
| enzyme | Up-regulated |
| other | Down-regulated |
| peptidase | Down-regulated |
| other | Down-regulated |
| other | Down-regulated |
| kinase | Down-regulated |

| | |
|---|---|
| other | Down-regulated |
| other | Down-regulated |
| enzyme | Down-regulated |
| other | Down-regulated |
| enzyme | Up-regulated |
| enzyme | Down-regulated |
| enzyme | Down-regulated |
| other | Down-regulated |
| other | Down-regulated |
| other | Down-regulated |
| phosphatase | Down-regulated |
| enzyme | Down-regulated |
| other | Down-regulated |
| transmembra | Down-regulated |
| other | Down-regulated |
| other | Up-regulated |
| other | Down-regulated |
| other | Down-regulated |
| enzyme | Down-regulated |
| transporter | Down-regulated |
| kinase | Down-regulated |
| other | Down-regulated |
| transporter | Down-regulated |
| other | Down-regulated |
| other | Down-regulated |
| other | Down-regulated |
| enzyme | Down-regulated |
| enzyme | Down-regulated |
| other | Down-regulated |
| enzyme | Down-regulated |
| other | Down-regulated |
| other | Down-regulated |
| other | Down-regulated |
| other | Down-regulated |
| enzyme | Down-regulated |
| kinase | Up-regulated |
| other | Down-regulated |
| kinase | Down-regulated |
| enzyme | Down-regulated |
| transporter | Down-regulated |
| other | Down-regulated |
| enzyme | Down-regulated |
| other | Down-regulated |
| transcription | Down-regulated |
| other | Down-regulated |
| enzyme | Down-regulated |
| enzyme | Down-regulated |
| enzyme | Down-regulated |
| other | Down-regulated |
| ion channel | Up-regulated |
| enzyme | Down-regulated |
| other | Down-regulated |
| other | Down-regulated |
| enzyme | Down-regulated |
| transcription | Down-regulated |
| enzyme | Down-regulated |
| enzyme | Down-regulated |
| kinase | Down-regulated |

| | |
|---|---|
| enzyme | Down-regulated |
| enzyme | Down-regulated |
| enzyme | Up-regulated |
| other | Down-regulated |
| transcription | Down-regulated |
| transporter | Up-regulated |
| transmembra | Up-regulated |
| transcription | Down-regulated |
| other | Down-regulated |
| transcription | Up-regulated |
| transcription | Down-regulated |
| peptidase | Down-regulated |
| other | Down-regulated |
| enzyme | Up-regulated |
| other | Down-regulated |
| other | Down-regulated |
| other | Down-regulated |
| enzyme | Down-regulated |
| other | Down-regulated |
| enzyme | Down-regulated |
| other | Down-regulated |
| other | Down-regulated |
| enzyme | Down-regulated |
| transcription | Down-regulated |
| peptidase | Down-regulated |
| transcription | Down-regulated |
| phosphatase | Down-regulated |
| other | Down-regulated |
| enzyme | Up-regulated |
| other | Down-regulated |
| other | Down-regulated |
| enzyme | Down-regulated |
| transporter | Up-regulated |
| enzyme | Down-regulated |
| enzyme | Down-regulated |
| translation re | Down-regulated |
| other | Down-regulated |
| other | Down-regulated |
| enzyme | Down-regulated |
| enzyme | Up-regulated |
| other | Down-regulated |
| other | Down-regulated |
| other | Down-regulated |
| other | Down-regulated |
| other | Down-regulated |
| other | Up-regulated |
| translation re | Down-regulated |
| other | Down-regulated |
| other | Down-regulated |
| other | Down-regulated |
| transcription | Down-regulated |
| enzyme | Down-regulated |
| transcription | Down-regulated |
| enzyme | Down-regulated |
| transmembra | Up-regulated |
| other | Down-regulated |
| phosphatase | Down-regulated |
| other | Down-regulated |

| | |
|---|---|
| transporter | Down-regulated |
| other | Down-regulated |
| enzyme | Down-regulated |
| other | Down-regulated |
| transcription | Down-regulated |
| enzyme | Down-regulated |
| enzyme | Down-regulated |
| enzyme | Down-regulated |
| enzyme | Down-regulated |
| kinase | Down-regulated |
| peptidase | Down-regulated |
| transcription | Down-regulated |
| enzyme | Up-regulated |
| other | Down-regulated |
| transcription | Down-regulated |
| other | Down-regulated |
| transcription | Down-regulated |
| enzyme | Down-regulated |
| other | Down-regulated |
| other | Down-regulated |
| transcription | Down-regulated |
| enzyme | Down-regulated |
| other | Down-regulated |
| transcription | Down-regulated |
| other | Down-regulated |
| other | Down-regulated |
| enzyme | Down-regulated |
| other | Down-regulated |
| transcription | Down-regulated |
| other | Down-regulated |
| other | Up-regulated |
| other | Down-regulated |
| other | Down-regulated |
| transcription | Down-regulated |
| other | Down-regulated |
| other | Down-regulated |
| enzyme | Down-regulated |
| enzyme | Down-regulated |
| other | Down-regulated |
| other | Down-regulated |
| transcription | Down-regulated |
| transcription | Down-regulated |
| other | Down-regulated |
| enzyme | Down-regulated |
| enzyme | Down-regulated |
| other | Down-regulated |
| other | Down-regulated |
| other | Down-regulated |
| enzyme | Down-regulated |
| enzyme | Down-regulated |
| kinase | Down-regulated |
| ion channel | Up-regulated |
| other | Down-regulated |
| other | Down-regulated |
| enzyme | Down-regulated |
| transcription | Down-regulated |
| other | Up-regulated |
| other | Down-regulated |

| | |
|---|---|
| transcription | Down-regulated |
| other | Down-regulated |
| kinase | Down-regulated |
| other | Down-regulated |
| transcription | Down-regulated |
| transcription | Down-regulated |
| enzyme | Down-regulated |
| other | Down-regulated |
| other | Down-regulated |
| transcription | Down-regulated |
| other | Down-regulated |
| enzyme | Up-regulated |
| transcription | Down-regulated |
| transcription | Down-regulated |
| other | Down-regulated |
| other | Down-regulated |
| enzyme | Down-regulated |
| other | Down-regulated |
| transcription | Down-regulated |
| transcription | Down-regulated |
| transcription | Down-regulated |
| enzyme | Down-regulated |
| enzyme | Up-regulated |
| other | Down-regulated |
| transcription | Down-regulated |
| other | Down-regulated |
| phosphatase | Down-regulated |
| phosphatase | Down-regulated |
| enzyme | Down-regulated |
| kinase | Down-regulated |
| enzyme | Down-regulated |
| other | Down-regulated |
| other | Down-regulated |
| other | Down-regulated |
| enzyme | Down-regulated |
| other | Up-regulated |
| transcription | Down-regulated |
| other | Down-regulated |
| other | Down-regulated |
| enzyme | Down-regulated |
| other | Down-regulated |
| enzyme | Down-regulated |
| other | Down-regulated |
| other | Down-regulated |
| other | Down-regulated |
| other | Down-regulated |
| other | Down-regulated |
| other | Down-regulated |
| phosphatase | Down-regulated |
| other | Down-regulated |
| transcription | Down-regulated |
| transporter | Up-regulated |
| enzyme | Down-regulated |
| enzyme | Down-regulated |
| transcription | Down-regulated |
| other | Down-regulated |
| kinase | Down-regulated |
| translation re | Down-regulated |

| | |
|---|---|
| other | Down-regulated |
| transporter | Up-regulated |
| other | Down-regulated |
| other | Down-regulated |
| other | Down-regulated |
| other | Down-regulated |
| other | Down-regulated |
| enzyme | Down-regulated |
| enzyme | Down-regulated |
| transporter | Up-regulated |
| transporter | Up-regulated |
| other | Down-regulated |
| other | Down-regulated |
| enzyme | Down-regulated |
| transcription | Down-regulated |
| enzyme | Down-regulated |
| other | Down-regulated |
| other | Down-regulated |
| phosphatase | Up-regulated |
| kinase | Down-regulated |
| transporter | Up-regulated |
| other | Down-regulated |
| other | Up-regulated |
| other | Down-regulated |
| G-protein co | Down-regulated |
| other | Down-regulated |
| other | Down-regulated |
| other | Down-regulated |
| transcription | Down-regulated |
| enzyme | Up-regulated |
| transmembra | Up-regulated |
| other | Down-regulated |
| other | Down-regulated |
| transporter | Up-regulated |
| other | Down-regulated |
| other | Down-regulated |
| other | Down-regulated |
| kinase | Down-regulated |
| enzyme | Down-regulated |
| other | Down-regulated |
| enzyme | Down-regulated |
| other | Down-regulated |
| transporter | Down-regulated |
| other | Down-regulated |
| other | Down-regulated |
| other | Down-regulated |
| phosphatase | Down-regulated |
| other | Up-regulated |
| other | Down-regulated |
| other | Down-regulated |
| other | Down-regulated |
| enzyme | Down-regulated |
| peptidase | Down-regulated |
| enzyme | Down-regulated |
| transporter | Down-regulated |
| enzyme | Down-regulated |
| ligand-deper | Down-regulated |
| enzyme | Down-regulated |

| | |
|---|---|
| other | Down-regulated |
| kinase | Down-regulated |
| transcription | Down-regulated |
| other | Down-regulated |
| kinase | Down-regulated |
| kinase | Down-regulated |
| kinase | Down-regulated |
| other | Down-regulated |
| transporter | Down-regulated |
| transcription | Down-regulated |
| other | Down-regulated |
| other | Down-regulated |
| other | Down-regulated |
| phosphatase | Down-regulated |
| transcription | Down-regulated |
| transporter | Up-regulated |
| other | Down-regulated |
| other | Down-regulated |
| peptidase | Down-regulated |
| other | Down-regulated |
| enzyme | Down-regulated |
| other | Down-regulated |
| other | Down-regulated |
| kinase | Up-regulated |
| other | Down-regulated |
| peptidase | Down-regulated |
| other | Down-regulated |
| transcription | Down-regulated |
| enzyme | Down-regulated |
| other | Down-regulated |
| other | Down-regulated |
| transcription | Down-regulated |
| enzyme | Down-regulated |
| other | Down-regulated |
| transcription | Down-regulated |
| other | Down-regulated |
| enzyme | Down-regulated |
| other | Down-regulated |
| ligand-deper | Up-regulated |
| other | Down-regulated |
| enzyme | Down-regulated |
| phosphatase | Down-regulated |
| other | Down-regulated |
| enzyme | Down-regulated |
| transcription | Down-regulated |
| other | Down-regulated |
| transcription | Down-regulated |
| other | Down-regulated |
| other | Down-regulated |
| other | Down-regulated |
| kinase | Up-regulated |
| transmembra | Up-regulated |
| other | Down-regulated |
| enzyme | Down-regulated |
| other | Up-regulated |
| enzyme | Down-regulated |
| other | Down-regulated |
| other | Down-regulated |

| | |
|---|---|
| transcription | Down-regulated |
| other | Down-regulated |
| other | Down-regulated |
| transporter | Down-regulated |
| other | Down-regulated |
| other | Down-regulated |
| other | Down-regulated |
| phosphatase | Down-regulated |
| transcription | Down-regulated |
| peptidase | Down-regulated |
| transporter | Down-regulated |
| other | Down-regulated |
| peptidase | Down-regulated |
| other | Down-regulated |
| other | Down-regulated |
| peptidase | Down-regulated |
| other | Down-regulated |
| other | Down-regulated |
| transcription | Down-regulated |
| transcription | Down-regulated |
| other | Down-regulated |
| other | Down-regulated |
| transcription | Down-regulated |
| enzyme | Down-regulated |
| other | Down-regulated |
| kinase | Down-regulated |
| other | Down-regulated |
| transcription | Down-regulated |
| enzyme | Up-regulated |
| other | Up-regulated |
| other | Down-regulated |
| enzyme | Down-regulated |
| kinase | Down-regulated |
| other | Down-regulated |
| peptidase | Down-regulated |
| other | Down-regulated |
| other | Down-regulated |
| other | Down-regulated |
| other | Down-regulated |
| enzyme | Down-regulated |
| enzyme | Down-regulated |
| other | Down-regulated |
| other | Down-regulated |
| enzyme | Down-regulated |
| other | Up-regulated |
| kinase | Down-regulated |
| other | Down-regulated |
| enzyme | Down-regulated |
| other | Down-regulated |
| kinase | Down-regulated |
| phosphatase | Down-regulated |
| other | Down-regulated |
| other | Down-regulated |
| peptidase | Down-regulated |
| other | Down-regulated |
| other | Down-regulated |
| enzyme | Down-regulated |
| enzyme | Up-regulated |

| | |
|---|---|
| enzyme | Down-regulated |
| enzyme | Down-regulated |
| enzyme | Down-regulated |
| transcription | Down-regulated |
| enzyme | Down-regulated |
| other | Down-regulated |
| peptidase | Down-regulated |
| other | Down-regulated |
| transcription | Down-regulated |
| other | Down-regulated |
| other | Down-regulated |
| transporter | Down-regulated |
| peptidase | Down-regulated |
| cytokine | Down-regulated |
| other | Up-regulated |
| other | Down-regulated |
| kinase | Down-regulated |
| enzyme | Up-regulated |
| other | Down-regulated |
| other | Down-regulated |
| other | Down-regulated |
| other | Down-regulated |
| other | Down-regulated |
| enzyme | Down-regulated |
| other | Down-regulated |
| other | Down-regulated |
| other | Up-regulated |
| other | Down-regulated |
| enzyme | Down-regulated |
| peptidase | Down-regulated |
| transcription | Down-regulated |
| transcription | Down-regulated |
| other | Down-regulated |
| transcription | Down-regulated |
| other | Down-regulated |
| transcription | Down-regulated |
| transcription | Down-regulated |
| other | Down-regulated |
| other | Down-regulated |
| enzyme | Down-regulated |
| other | Down-regulated |
| kinase | Down-regulated |
| enzyme | Down-regulated |
| transporter | Down-regulated |
| peptidase | Up-regulated |
| transcription | Down-regulated |
| transporter | Down-regulated |
| other | Down-regulated |
| kinase | Down-regulated |
| other | Down-regulated |
| other | Down-regulated |
| transporter | Down-regulated |
| transcription | Down-regulated |
| other | Down-regulated |
| other | Down-regulated |
| other | Down-regulated |
| other | Down-regulated |
| enzyme | Down-regulated |

| | |
|---|---|
| other | Down-regulated |
| other | Down-regulated |
| other | Down-regulated |
| other | Down-regulated |
| other | Down-regulated |
| transcription | Down-regulated |
| other | Up-regulated |
| other | Down-regulated |
| transporter | Up-regulated |
| enzyme | Down-regulated |
| other | Down-regulated |
| other | Down-regulated |
| kinase | Down-regulated |
| enzyme | Down-regulated |
| enzyme | Down-regulated |
| enzyme | Down-regulated |
| other | Down-regulated |
| transporter | Down-regulated |
| other | Down-regulated |
| other | Down-regulated |
| transcription | Down-regulated |
| transcription | Down-regulated |
| transporter | Down-regulated |
| other | Down-regulated |
| transporter | Down-regulated |
| other | Down-regulated |
| other | Down-regulated |
| other | Down-regulated |
| transcription | Up-regulated |
| transporter | Up-regulated |
| kinase | Down-regulated |
| enzyme | Down-regulated |
| transcription | Down-regulated |
| other | Down-regulated |
| other | Down-regulated |
| enzyme | Up-regulated |
| enzyme | Down-regulated |
| other | Down-regulated |
| other | Down-regulated |
| transmembra | Up-regulated |
| other | Down-regulated |
| enzyme | Down-regulated |
| enzyme | Down-regulated |
| enzyme | Up-regulated |
| other | Down-regulated |
| other | Down-regulated |
| other | Down-regulated |
| other | Up-regulated |
| enzyme | Down-regulated |
| other | Down-regulated |
| other | Down-regulated |
| enzyme | Up-regulated |
| enzyme | Up-regulated |
| other | Down-regulated |
| enzyme | Down-regulated |
| transcription | Down-regulated |
| kinase | Down-regulated |
| enzyme | Up-regulated |

| | |
|---|---|
| kinase | Down-regulated |
| enzyme | Down-regulated |
| enzyme | Down-regulated |
| transcription | Down-regulated |
| enzyme | Down-regulated |
| transmembra | Down-regulated |
| other | Down-regulated |
| other | Down-regulated |
| transcription | Down-regulated |
| other | Down-regulated |
| other | Down-regulated |
| peptidase | Down-regulated |
| phosphatase | Down-regulated |
| enzyme | Down-regulated |
| other | Down-regulated |
| transcription | Down-regulated |
| transporter | Down-regulated |
| enzyme | Down-regulated |
| transcription | Down-regulated |
| enzyme | Down-regulated |
| other | Down-regulated |
| other | Down-regulated |
| other | Down-regulated |
| other | Down-regulated |
| other | Down-regulated |
| other | Down-regulated |
| enzyme | Down-regulated |
| transcription | Down-regulated |
| enzyme | Down-regulated |
| enzyme | Down-regulated |
| enzyme | Up-regulated |
| enzyme | Down-regulated |
| transcription | Down-regulated |
| other | Down-regulated |
| enzyme | Down-regulated |
| enzyme | Down-regulated |
| other | Down-regulated |
| enzyme | Down-regulated |
| enzyme | Up-regulated |
| transmembra | Up-regulated |
| other | Down-regulated |
| transcription | Down-regulated |
| other | Down-regulated |
| other | Down-regulated |
| enzyme | Down-regulated |
| other | Down-regulated |
| other | Down-regulated |
| enzyme | Up-regulated |
| other | Up-regulated |
| transmembra | Up-regulated |
| other | Down-regulated |
| other | Down-regulated |
| ligand-deper | Down-regulated |
| transcription | Down-regulated |
| enzyme | Up-regulated |
| other | Down-regulated |
| other | Down-regulated |
| transcription | Down-regulated |

| | |
|---|---|
| enzyme | Down-regulated |
| enzyme | Up-regulated |
| other | Down-regulated |
| phosphatase | Up-regulated |
| other | Down-regulated |
| phosphatase | Down-regulated |
| phosphatase | Down-regulated |
| other | Down-regulated |
| other | Down-regulated |
| enzyme | Down-regulated |
| other | Down-regulated |
| enzyme | Down-regulated |
| other | Down-regulated |
| enzyme | Down-regulated |
| transporter | Down-regulated |
| other | Down-regulated |
| peptidase | Down-regulated |
| phosphatase | Up-regulated |
| peptidase | Up-regulated |
| other | Up-regulated |
| other | Down-regulated |
| transcription | Down-regulated |
| other | Down-regulated |
| enzyme | Down-regulated |
| transcription | Down-regulated |
| enzyme | Down-regulated |
| other | Down-regulated |
| other | Down-regulated |
| enzyme | Down-regulated |
| other | Down-regulated |
| enzyme | Down-regulated |
| transcription | Down-regulated |
| other | Down-regulated |
| other | Down-regulated |
| peptidase | Down-regulated |
| enzyme | Up-regulated |
| transcription | Down-regulated |
| transporter | Down-regulated |
| other | Down-regulated |
| other | Down-regulated |
| enzyme | Up-regulated |
| transcription | Down-regulated |
| transcription | Down-regulated |
| other | Down-regulated |
| other | Down-regulated |
| peptidase | Down-regulated |
| kinase | Down-regulated |
| transcription | Down-regulated |
| phosphatase | Down-regulated |
| kinase | Down-regulated |
| other | Up-regulated |
| enzyme | Down-regulated |
| other | Down-regulated |
| other | Up-regulated |
| other | Down-regulated |
| phosphatase | Up-regulated |
| transcription | Down-regulated |
| transcription | Down-regulated |

| | |
|---|---|
| enzyme | Down-regulated |
| enzyme | Down-regulated |
| transcription | Down-regulated |
| other | Down-regulated |
| other | Down-regulated |
| transcription | Down-regulated |
| enzyme | Down-regulated |
| transcription | Down-regulated |
| transmembra | Down-regulated |
| other | Up-regulated |
| enzyme | Down-regulated |
| other | Down-regulated |
| enzyme | Down-regulated |
| other | Down-regulated |
| other | Down-regulated |
| enzyme | Down-regulated |
| other | Down-regulated |
| enzyme | Down-regulated |
| transcription | Down-regulated |
| transporter | Down-regulated |
| enzyme | Down-regulated |
| enzyme | Up-regulated |
| other | Down-regulated |
| other | Down-regulated |
| enzyme | Down-regulated |
| enzyme | Up-regulated |
| enzyme | Down-regulated |
| kinase | Down-regulated |
| other | Down-regulated |
| other | Down-regulated |
| kinase | Down-regulated |
| other | Down-regulated |
| phosphatase | Down-regulated |
| enzyme | Down-regulated |
| transcription | Down-regulated |
| other | Down-regulated |
| transporter | Down-regulated |
| enzyme | Down-regulated |
| other | Down-regulated |
| other | Down-regulated |
| enzyme | Down-regulated |
| enzyme | Down-regulated |
| transcription | Down-regulated |
| enzyme | Down-regulated |
| transcription | Down-regulated |
| other | Down-regulated |
| other | Down-regulated |
| other | Down-regulated |
| transporter | Down-regulated |
| other | Down-regulated |
| transcription | Down-regulated |
| transcription | Down-regulated |
| peptidase | Down-regulated |
| kinase | Down-regulated |
| enzyme | Up-regulated |
| other | Down-regulated |
| transcription | Down-regulated |
| transcription | Down-regulated |

| | |
|---|---|
| other | Down-regulated |
| other | Down-regulated |
| enzyme | Up-regulated |
| kinase | Down-regulated |
| other | Up-regulated |
| G-protein co | Down-regulated |
| other | Down-regulated |
| kinase | Down-regulated |
| other | Down-regulated |
| enzyme | Down-regulated |
| transporter | Down-regulated |
| transcription | Down-regulated |
| other | Down-regulated |
| other | Down-regulated |
| other | Down-regulated |
| phosphatase | Down-regulated |
| peptidase | Down-regulated |
| enzyme | Down-regulated |
| transporter | Down-regulated |
| enzyme | Down-regulated |
| enzyme | Down-regulated |
| phosphatase | Down-regulated |
| enzyme | Down-regulated |
| other | Down-regulated |
| other | Down-regulated |
| kinase | Down-regulated |
| other | Down-regulated |
| other | Down-regulated |
| enzyme | Down-regulated |
| enzyme | Down-regulated |
| other | Up-regulated |
| other | Down-regulated |
| other | Down-regulated |
| transcription | Down-regulated |
| other | Down-regulated |
| transcription | Down-regulated |
| transcription | Down-regulated |
| other | Down-regulated |
| other | Down-regulated |
| enzyme | Down-regulated |
| translation re | Down-regulated |
| transcription | Down-regulated |
| other | Down-regulated |
| other | Down-regulated |
| kinase | Down-regulated |
| transporter | Up-regulated |
| enzyme | Down-regulated |
| other | Down-regulated |
| other | Down-regulated |
| G-protein co | Down-regulated |
| transcription | Down-regulated |
| other | Down-regulated |
| enzyme | Down-regulated |
| other | Down-regulated |
| translation re | Down-regulated |
| enzyme | Down-regulated |
| transporter | Down-regulated |
| other | Down-regulated |

| | |
|---|---|
| other | Down-regulated |
| enzyme | Down-regulated |
| enzyme | Down-regulated |
| transcription | Down-regulated |
| enzyme | Down-regulated |
| other | Up-regulated |
| other | Down-regulated |
| other | Down-regulated |
| transcription | Down-regulated |
| other | Down-regulated |
| enzyme | Down-regulated |
| transporter | Down-regulated |
| other | Down-regulated |
| enzyme | Down-regulated |
| other | Down-regulated |
| other | Down-regulated |
| other | Down-regulated |
| other | Down-regulated |
| other | Down-regulated |
| transporter | Up-regulated |
| other | Down-regulated |
| other | Down-regulated |
| enzyme | Up-regulated |
| enzyme | Down-regulated |
| other | Down-regulated |
| phosphatase | Down-regulated |
| other | Down-regulated |
| other | Down-regulated |
| kinase | Down-regulated |
| enzyme | Up-regulated |
| enzyme | Down-regulated |
| enzyme | Down-regulated |
| other | Down-regulated |
| enzyme | Down-regulated |
| enzyme | Down-regulated |
| other | Down-regulated |
| enzyme | Down-regulated |
| other | Down-regulated |
| other | Down-regulated |
| transcription | Down-regulated |
| transcription | Down-regulated |
| peptidase | Down-regulated |
| other | Down-regulated |
| enzyme | Down-regulated |
| kinase | Down-regulated |
| enzyme | Down-regulated |
| phosphatase | Down-regulated |
| other | Down-regulated |
| enzyme | Down-regulated |
| other | Down-regulated |
| other | Down-regulated |
| enzyme | Down-regulated |
| other | Down-regulated |
| transcription | Down-regulated |
| enzyme | Down-regulated |
| transcription | Down-regulated |
| other | Down-regulated |
| transporter | Down-regulated |

| | |
|---|---|
| other | Down-regulated |
| other | Down-regulated |
| kinase | Up-regulated |
| kinase | Down-regulated |
| other | Down-regulated |
| other | Down-regulated |
| ion channel | Down-regulated |
| enzyme | Down-regulated |
| other | Down-regulated |
| enzyme | Down-regulated |
| transporter | Down-regulated |
| enzyme | Down-regulated |
| peptidase | Down-regulated |
| other | Down-regulated |
| enzyme | Up-regulated |
| other | Down-regulated |
| transcription | Down-regulated |
| other | Down-regulated |
| other | Down-regulated |
| other | Down-regulated |
| enzyme | Down-regulated |
| enzyme | Up-regulated |
| enzyme | Down-regulated |
| transcription | Down-regulated |
| other | Down-regulated |
| transporter | Down-regulated |
| transporter | Up-regulated |
| transcription | Down-regulated |
| other | Down-regulated |
| other | Down-regulated |
| ion channel | Down-regulated |
| other | Down-regulated |
| enzyme | Down-regulated |
| enzyme | Down-regulated |
| other | Up-regulated |
| other | Down-regulated |
| other | Down-regulated |
| other | Down-regulated |
| transporter | Down-regulated |
| other | Down-regulated |
| enzyme | Down-regulated |
| enzyme | Up-regulated |
| transcription | Down-regulated |
| transporter | Down-regulated |
| transcription | Down-regulated |
| transcription | Down-regulated |
| other | Down-regulated |
| G-protein co | Up-regulated |
| transcription | Down-regulated |
| other | Down-regulated |
| other | Down-regulated |
| enzyme | Down-regulated |
| enzyme | Down-regulated |
| other | Down-regulated |
| G-protein co | Down-regulated |
| transcription | Down-regulated |
| other | Down-regulated |
| other | Down-regulated |

| | |
|---|---|
| other | Down-regulated |
| other | Down-regulated |
| other | Down-regulated |
| other | Down-regulated |
| growth facto | Down-regulated |
| enzyme | Down-regulated |
| other | Down-regulated |
| other | Down-regulated |
| transporter | Down-regulated |
| other | Down-regulated |
| other | Down-regulated |
| enzyme | Down-regulated |
| transporter | Down-regulated |
| enzyme | Down-regulated |
| other | Down-regulated |
| other | Down-regulated |
| enzyme | Up-regulated |
| enzyme | Down-regulated |
| enzyme | Down-regulated |
| enzyme | Down-regulated |
| enzyme | Down-regulated |
| other | Down-regulated |
| enzyme | Down-regulated |
| other | Down-regulated |
| other | Up-regulated |
| enzyme | Up-regulated |
| other | Down-regulated |
| other | Down-regulated |
| other | Down-regulated |
| other | Down-regulated |
| kinase | Down-regulated |
| enzyme | Down-regulated |
| other | Down-regulated |
| kinase | Down-regulated |
| other | Down-regulated |
| enzyme | Down-regulated |
| enzyme | Down-regulated |
| other | Down-regulated |
| other | Down-regulated |
| other | Down-regulated |
| enzyme | Up-regulated |
| other | Down-regulated |
| enzyme | Down-regulated |
| transcription | Down-regulated |
| kinase | Down-regulated |
| cytokine | Down-regulated |
| other | Up-regulated |
| enzyme | Down-regulated |
| transcription | Down-regulated |
| enzyme | Up-regulated |
| other | Down-regulated |
| other | Down-regulated |
| enzyme | Down-regulated |
| enzyme | Down-regulated |
| enzyme | Down-regulated |
| other | Down-regulated |
| other | Down-regulated |
| transcription | Down-regulated |

| | |
|---|---|
| other | Down-regulated |
| enzyme | Down-regulated |
| other | Down-regulated |
| other | Down-regulated |
| other | Down-regulated |
| other | Down-regulated |
| other | Down-regulated |
| transporter | Down-regulated |
| enzyme | Down-regulated |
| other | Down-regulated |
| enzyme | Up-regulated |
| kinase | Down-regulated |
| other | Down-regulated |
| other | Down-regulated |
| enzyme | Down-regulated |
| transporter | Down-regulated |
| kinase | Up-regulated |
| other | Down-regulated |
| enzyme | Down-regulated |
| other | Down-regulated |
| other | Down-regulated |
| other | Down-regulated |
| other | Up-regulated |
| other | Down-regulated |
| enzyme | Down-regulated |
| other | Down-regulated |
| enzyme | Down-regulated |
| other | Down-regulated |
| other | Down-regulated |
| transcription | Up-regulated |
| enzyme | Down-regulated |
| transcription | Down-regulated |
| other | Down-regulated |
| enzyme | Down-regulated |
| enzyme | Down-regulated |
| enzyme | Down-regulated |
| other | Down-regulated |
| other | Down-regulated |
| other | Down-regulated |
| transcription | Down-regulated |
| peptidase | Down-regulated |
| enzyme | Down-regulated |
| kinase | Down-regulated |
| other | Down-regulated |
| enzyme | Down-regulated |
| other | Down-regulated |
| transcription | Down-regulated |
| other | Down-regulated |
| other | Down-regulated |
| other | Down-regulated |
| other | Down-regulated |
| phosphatase | Up-regulated |
| other | Down-regulated |
| enzyme | Down-regulated |
| other | Down-regulated |
| enzyme | Up-regulated |
| other | Down-regulated |
| enzyme | Down-regulated |

| | |
|---|---|
| kinase | Down-regulated |
| other | Down-regulated |
| transcription | Down-regulated |
| enzyme | Down-regulated |
| enzyme | Down-regulated |
| enzyme | Down-regulated |
| enzyme | Down-regulated |
| other | Up-regulated |
| transmembra | Up-regulated |
| enzyme | Up-regulated |
| transcription | Down-regulated |
| transcription | Down-regulated |
| transcription | Down-regulated |
| enzyme | Down-regulated |
| other | Down-regulated |
| other | Down-regulated |
| other | Down-regulated |
| enzyme | Down-regulated |
| other | Down-regulated |
| other | Down-regulated |
| transcription | Down-regulated |
| other | Down-regulated |
| other | Down-regulated |
| other | Down-regulated |
| other | Up-regulated |
| enzyme | Down-regulated |
| transcription | Down-regulated |
| other | Down-regulated |
| other | Down-regulated |
| other | Down-regulated |
| enzyme | Up-regulated |
| other | Down-regulated |
| transcription | Down-regulated |
| transcription | Down-regulated |
| other | Up-regulated |
| enzyme | Up-regulated |
| other | Down-regulated |
| other | Down-regulated |
| other | Down-regulated |
| transcription | Down-regulated |
| enzyme | Down-regulated |
| other | Down-regulated |
| transcription | Down-regulated |
| other | Down-regulated |
| other | Down-regulated |
| enzyme | Down-regulated |
| transporter | Up-regulated |
| transcription | Down-regulated |
| other | Down-regulated |
| enzyme | Down-regulated |
| other | Down-regulated |
| other | Down-regulated |
| enzyme | Down-regulated |
| phosphatase | Down-regulated |
| kinase | Down-regulated |
| other | Down-regulated |
| other | Down-regulated |
| kinase | Down-regulated |

| | |
|---|---|
| other | Down-regulated |
| other | Down-regulated |
| other | Down-regulated |
| other | Down-regulated |
| other | Down-regulated |
| enzyme | Down-regulated |
| transcription | Down-regulated |
| transcription | Down-regulated |
| other | Down-regulated |
| phosphatase | Down-regulated |
| other | Down-regulated |
| transporter | Down-regulated |
| enzyme | Down-regulated |
| other | Down-regulated |
| enzyme | Down-regulated |
| enzyme | Down-regulated |
| other | Down-regulated |
| other | Down-regulated |
| other | Down-regulated |
| other | Down-regulated |
| ion channel | Down-regulated |
| other | Down-regulated |
| other | Down-regulated |
| enzyme | Down-regulated |
| other | Down-regulated |
| transcription | Down-regulated |
| other | Down-regulated |
| other | Up-regulated |
| enzyme | Down-regulated |
| enzyme | Down-regulated |
| other | Down-regulated |
| enzyme | Up-regulated |
| other | Down-regulated |
| enzyme | Down-regulated |
| enzyme | Down-regulated |
| transporter | Up-regulated |
| enzyme | Down-regulated |
| other | Up-regulated |
| enzyme | Down-regulated |
| transporter | Down-regulated |
| enzyme | Down-regulated |
| other | Down-regulated |
| kinase | Down-regulated |
| other | Down-regulated |
| other | Down-regulated |
| kinase | Down-regulated |
| other | Down-regulated |
| enzyme | Down-regulated |
| enzyme | Up-regulated |
| enzyme | Down-regulated |
| other | Up-regulated |
| other | Down-regulated |
| other | Down-regulated |
| other | Down-regulated |
| enzyme | Down-regulated |
| other | Down-regulated |
| other | Down-regulated |
| transporter | Down-regulated |

| | |
|---|---|
| other | Down-regulated |
| other | Down-regulated |
| other | Up-regulated |
| other | Down-regulated |
| other | Down-regulated |
| enzyme | Down-regulated |
| transcription | Down-regulated |
| kinase | Down-regulated |
| other | Down-regulated |
| transcription | Down-regulated |
| other | Down-regulated |
| transporter | Down-regulated |
| cytokine | Up-regulated |
| transcription | Down-regulated |
| transcription | Down-regulated |
| translation re | Down-regulated |
| other | Up-regulated |
| enzyme | Down-regulated |
| other | Down-regulated |
| enzyme | Up-regulated |
| transcription | Down-regulated |
| transporter | Down-regulated |
| other | Down-regulated |
| G-protein co | Down-regulated |
| transporter | Down-regulated |
| other | Down-regulated |
| enzyme | Down-regulated |
| other | Down-regulated |
| other | Up-regulated |
| other | Down-regulated |
| enzyme | Down-regulated |
| kinase | Down-regulated |
| ion channel | Down-regulated |
| other | Down-regulated |
| G-protein co | Up-regulated |
| other | Down-regulated |
| transporter | Down-regulated |
| enzyme | Down-regulated |
| transcription | Down-regulated |
| other | Down-regulated |
| other | Down-regulated |
| other | Down-regulated |
| other | Down-regulated |
| other | Down-regulated |
| other | Down-regulated |
| enzyme | Down-regulated |
| kinase | Down-regulated |
| transcription | Down-regulated |
| enzyme | Up-regulated |
| transporter | Up-regulated |
| other | Down-regulated |
| other | Down-regulated |
| enzyme | Down-regulated |
| transporter | Down-regulated |
| enzyme | Down-regulated |
| enzyme | Down-regulated |
| enzyme | Up-regulated |
| other | Down-regulated |

| | |
|---|---|
| other | Up-regulated |
| other | Down-regulated |
| enzyme | Up-regulated |
| enzyme | Down-regulated |
| kinase | Down-regulated |
| other | Down-regulated |
| transporter | Down-regulated |
| kinase | Down-regulated |
| kinase | Down-regulated |
| transcription | Down-regulated |
| other | Down-regulated |
| other | Down-regulated |
| other | Down-regulated |
| other | Down-regulated |
| transporter | Down-regulated |
| other | Down-regulated |
| transcription | Down-regulated |
| enzyme | Down-regulated |
| translation re | Down-regulated |
| other | Down-regulated |
| ligand-deper | Down-regulated |
| other | Down-regulated |
| other | Down-regulated |
| other | Up-regulated |
| enzyme | Down-regulated |
| other | Down-regulated |
| kinase | Down-regulated |
| other | Up-regulated |
| kinase | Down-regulated |
| other | Down-regulated |
| peptidase | Down-regulated |
| other | Down-regulated |
| other | Up-regulated |
| other | Down-regulated |
| other | Down-regulated |
| enzyme | Up-regulated |
| enzyme | Down-regulated |
| other | Down-regulated |
| other | Down-regulated |
| enzyme | Down-regulated |
| other | Down-regulated |
| other | Down-regulated |
| transporter | Down-regulated |
| enzyme | Up-regulated |
| other | Down-regulated |
| enzyme | Down-regulated |
| other | Down-regulated |
| enzyme | Up-regulated |
| other | Down-regulated |
| other | Down-regulated |
| enzyme | Down-regulated |
| other | Down-regulated |
| ion channel | Down-regulated |
| peptidase | Down-regulated |
| other | Down-regulated |
| transporter | Up-regulated |
| other | Down-regulated |
| enzyme | Up-regulated |

| | |
|---|---|
| other | Down-regulated |
| enzyme | Up-regulated |
| other | Down-regulated |
| other | Down-regulated |
| other | Down-regulated |
| other | Down-regulated |
| enzyme | Down-regulated |
| enzyme | Down-regulated |
| enzyme | Down-regulated |
| enzyme | Down-regulated |
| other | Down-regulated |
| transcription | Down-regulated |
| other | Down-regulated |
| other | Down-regulated |
| other | Down-regulated |
| other | Down-regulated |
| transporter | Down-regulated |
| other | Up-regulated |
| other | Down-regulated |
| transmembra | Up-regulated |
| enzyme | Up-regulated |
| other | Down-regulated |
| enzyme | Down-regulated |
| other | Down-regulated |
| other | Down-regulated |
| enzyme | Down-regulated |
| transporter | Down-regulated |
| growth facto | Up-regulated |
| enzyme | Down-regulated |
| other | Down-regulated |
| other | Down-regulated |
| other | Down-regulated |
| other | Up-regulated |
| transcription | Down-regulated |
| transporter | Down-regulated |
| other | Down-regulated |
| transporter | Down-regulated |
| other | Down-regulated |
| kinase | Up-regulated |
| phosphatase | Down-regulated |
| other | Up-regulated |
| peptidase | Down-regulated |
| other | Down-regulated |
| transporter | Down-regulated |
| other | Up-regulated |
| phosphatase | Down-regulated |
| kinase | Up-regulated |
| G-protein co | Down-regulated |
| other | Down-regulated |
| other | Down-regulated |
| enzyme | Down-regulated |
| enzyme | Up-regulated |
| other | Down-regulated |
| enzyme | Down-regulated |
| transporter | Down-regulated |
| other | Down-regulated |
| transporter | Down-regulated |
| other | Down-regulated |

| | |
|---|---|
| other | Down-regulated |
| other | Down-regulated |
| transcription | Down-regulated |
| kinase | Down-regulated |
| other | Up-regulated |
| enzyme | Down-regulated |
| transcription | Down-regulated |
| enzyme | Down-regulated |
| enzyme | Up-regulated |
| enzyme | Down-regulated |
| other | Down-regulated |
| other | Down-regulated |
| transporter | Up-regulated |
| other | Down-regulated |
| enzyme | Down-regulated |
| other | Down-regulated |
| other | Down-regulated |
| transporter | Up-regulated |
| enzyme | Up-regulated |
| other | Down-regulated |
| enzyme | Up-regulated |
| other | Down-regulated |
| other | Down-regulated |
| other | Down-regulated |
| other | Down-regulated |
| other | Down-regulated |
| other | Down-regulated |
| transcription | Down-regulated |
| transcription | Down-regulated |
| other | Up-regulated |
| other | Down-regulated |
| enzyme | Down-regulated |
| transcription | Down-regulated |
| transcription | Down-regulated |
| enzyme | Down-regulated |
| transporter | Down-regulated |
| other | Down-regulated |
| transcription | Down-regulated |
| transporter | Down-regulated |
| other | Down-regulated |
| other | Down-regulated |
| transcription | Down-regulated |
| transcription | Down-regulated |
| enzyme | Down-regulated |
| peptidase | Down-regulated |
| transcription | Down-regulated |
| kinase | Down-regulated |
| phosphatase | Down-regulated |
| other | Down-regulated |
| other | Down-regulated |
| other | Down-regulated |
| other | Down-regulated |
| transmembra | Down-regulated |
| other | Down-regulated |
| other | Down-regulated |
| enzyme | Down-regulated |
| other | Down-regulated |
| other | Down-regulated |

| | |
|---|---|
| other | Up-regulated |
| other | Down-regulated |
| enzyme | Down-regulated |
| other | Down-regulated |
| transcription | Down-regulated |
| peptidase | Down-regulated |
| enzyme | Down-regulated |
| kinase | Up-regulated |
| other | Down-regulated |
| transcription | Down-regulated |
| other | Down-regulated |
| other | Down-regulated |
| kinase | Down-regulated |
| transcription | Down-regulated |
| enzyme | Down-regulated |
| other | Down-regulated |
| kinase | Down-regulated |
| enzyme | Down-regulated |
| other | Down-regulated |
| enzyme | Down-regulated |
| transporter | Up-regulated |
| kinase | Down-regulated |
| transcription | Down-regulated |
| other | Down-regulated |
| other | Down-regulated |
| other | Down-regulated |
| other | Down-regulated |
| transcription | Down-regulated |
| other | Down-regulated |
| other | Down-regulated |
| other | Down-regulated |
| enzyme | Down-regulated |
| transmembra | Down-regulated |
| transcription | Down-regulated |
| other | Up-regulated |
| other | Down-regulated |
| transcription | Down-regulated |
| enzyme | Up-regulated |
| other | Down-regulated |
| other | Down-regulated |
| phosphatase | Down-regulated |
| peptidase | Down-regulated |
| other | Down-regulated |
| enzyme | Down-regulated |
| other | Down-regulated |
| enzyme | Down-regulated |
| enzyme | Down-regulated |
| transcription | Down-regulated |
| enzyme | Down-regulated |
| other | Down-regulated |
| enzyme | Down-regulated |
| cytokine | Down-regulated |
| transporter | Up-regulated |
| transporter | Down-regulated |
| other | Down-regulated |
| other | Down-regulated |
| other | Down-regulated |
| other | Down-regulated |

| | |
|---|---|
| transcription | Down-regulated |
| enzyme | Down-regulated |
| other | Down-regulated |
| enzyme | Down-regulated |
| transcription | Down-regulated |
| enzyme | Down-regulated |
| other | Down-regulated |
| transcription | Down-regulated |
| transcription | Down-regulated |
| enzyme | Down-regulated |
| other | Down-regulated |
| other | Down-regulated |
| other | Down-regulated |
| other | Down-regulated |
| transcription | Down-regulated |
| transporter | Down-regulated |
| transcription | Down-regulated |
| transporter | Down-regulated |
| other | Down-regulated |
| kinase | Down-regulated |
| transporter | Down-regulated |
| other | Down-regulated |
| enzyme | Down-regulated |
| other | Down-regulated |
| other | Down-regulated |
| enzyme | Up-regulated |
| other | Down-regulated |
| other | Down-regulated |
| kinase | Down-regulated |
| other | Down-regulated |
| enzyme | Down-regulated |
| transcription | Down-regulated |
| other | Down-regulated |
| transcription | Down-regulated |
| other | Down-regulated |
| cytokine | Up-regulated |
| other | Down-regulated |
| enzyme | Down-regulated |
| phosphatase | Down-regulated |
| other | Down-regulated |
| other | Down-regulated |
| other | Down-regulated |
| kinase | Down-regulated |
| peptidase | Up-regulated |
| other | Down-regulated |
| transcription | Down-regulated |
| other | Down-regulated |
| transcription | Down-regulated |
| transcription | Up-regulated |
| other | Down-regulated |
| other | Down-regulated |
| other | Up-regulated |
| transporter | Down-regulated |
| kinase | Down-regulated |
| enzyme | Up-regulated |
| kinase | Down-regulated |
| other | Up-regulated |
| transcription | Down-regulated |

| | |
|---|---|
| kinase | Down-regulated |
| transporter | Down-regulated |
| transcription | Down-regulated |
| transporter | Down-regulated |
| enzyme | Up-regulated |
| other | Down-regulated |
| phosphatase | Down-regulated |
| transporter | Down-regulated |
| phosphatase | Down-regulated |
| phosphatase | Down-regulated |
| other | Down-regulated |
| other | Down-regulated |
| other | Down-regulated |
| transporter | Down-regulated |
| other | Down-regulated |
| enzyme | Down-regulated |
| enzyme | Down-regulated |
| enzyme | Up-regulated |
| enzyme | Down-regulated |
| other | Down-regulated |
| enzyme | Down-regulated |
| enzyme | Down-regulated |
| transporter | Up-regulated |
| other | Down-regulated |
| other | Down-regulated |
| other | Up-regulated |
| transporter | Up-regulated |
| other | Down-regulated |
| kinase | Down-regulated |
| peptidase | Down-regulated |
| other | Down-regulated |
| peptidase | Down-regulated |
| other | Down-regulated |
| transcription | Down-regulated |
| enzyme | Up-regulated |
| peptidase | Up-regulated |
| other | Down-regulated |
| transcription | Down-regulated |
| transcription | Down-regulated |
| other | Down-regulated |
| other | Down-regulated |
| other | Down-regulated |
| ion channel | Down-regulated |
| other | Down-regulated |
| transcription | Down-regulated |
| kinase | Down-regulated |
| enzyme | Up-regulated |
| enzyme | Down-regulated |
| other | Down-regulated |
| transporter | Down-regulated |
| other | Down-regulated |
| enzyme | Up-regulated |
| other | Down-regulated |
| enzyme | Down-regulated |
| transporter | Up-regulated |
| kinase | Down-regulated |
| other | Down-regulated |
| other | Down-regulated |

| | |
|---|---|
| G-protein co | Up-regulated |
| other | Down-regulated |
| other | Down-regulated |
| transporter | Up-regulated |
| transcription | Down-regulated |
| other | Down-regulated |
| transporter | Up-regulated |
| kinase | Down-regulated |
| other | Down-regulated |
| other | Down-regulated |
| peptidase | Down-regulated |
| other | Down-regulated |
| other | Down-regulated |
| kinase | Down-regulated |
| other | Down-regulated |
| enzyme | Up-regulated |
| enzyme | Down-regulated |
| transporter | Down-regulated |
| phosphatase | Down-regulated |
| enzyme | Down-regulated |
| transcription | Down-regulated |
| other | Down-regulated |
| other | Down-regulated |
| other | Down-regulated |
| enzyme | Down-regulated |
| transcription | Down-regulated |
| enzyme | Down-regulated |
| phosphatase | Down-regulated |
| other | Down-regulated |
| transcription | Down-regulated |
| enzyme | Down-regulated |
| other | Down-regulated |
| kinase | Up-regulated |
| transcription | Down-regulated |
| other | Up-regulated |
| other | Down-regulated |
| other | Down-regulated |
| kinase | Down-regulated |
| transcription | Up-regulated |
| other | Down-regulated |
| other | Down-regulated |
| ion channel | Down-regulated |
| other | Down-regulated |
| other | Down-regulated |
| other | Down-regulated |
| other | Down-regulated |
| other | Down-regulated |
| transporter | Down-regulated |
| other | Down-regulated |
| kinase | Down-regulated |
| transcription | Down-regulated |
| peptidase | Down-regulated |
| other | Down-regulated |
| other | Down-regulated |
| other | Up-regulated |
| transmembra | Down-regulated |
| other | Down-regulated |
| G-protein co | Up-regulated |

| | |
|---|---|
| transcription | Down-regulated |
| peptidase | Up-regulated |
| enzyme | Down-regulated |
| enzyme | Down-regulated |
| transmembra | Up-regulated |
| other | Down-regulated |
| enzyme | Down-regulated |
| enzyme | Up-regulated |
| kinase | Down-regulated |
| peptidase | Down-regulated |
| enzyme | Down-regulated |
| other | Down-regulated |
| other | Down-regulated |
| transmembra | Down-regulated |
| other | Down-regulated |
| enzyme | Down-regulated |
| enzyme | Down-regulated |
| transmembra | Down-regulated |
| other | Down-regulated |
| other | Down-regulated |
| transcription | Down-regulated |
| enzyme | Up-regulated |
| kinase | Down-regulated |
| other | Down-regulated |
| other | Up-regulated |
| other | Down-regulated |
| enzyme | Up-regulated |
| other | Down-regulated |
| other | Down-regulated |
| other | Down-regulated |
| enzyme | Up-regulated |
| enzyme | Down-regulated |
| other | Down-regulated |
| enzyme | Up-regulated |
| other | Down-regulated |
| other | Down-regulated |
| transcription | Down-regulated |
| other | Down-regulated |
| other | Up-regulated |
| translation re | Down-regulated |
| phosphatase | Down-regulated |
| other | Down-regulated |
| enzyme | Down-regulated |
| other | Down-regulated |
| transporter | Down-regulated |
| other | Down-regulated |
| other | Down-regulated |
| transcription | Down-regulated |
| enzyme | Down-regulated |
| other | Up-regulated |
| other | Down-regulated |
| transporter | Up-regulated |
| enzyme | Down-regulated |
| other | Down-regulated |
| other | Down-regulated |
| kinase | Down-regulated |
| transcription | Down-regulated |
| peptidase | Down-regulated |

| | |
|---|---|
| peptidase | Down-regulated |
| peptidase | Up-regulated |
| enzyme | Down-regulated |
| enzyme | Down-regulated |
| transporter | Up-regulated |
| other | Down-regulated |
| other | Down-regulated |
| enzyme | Down-regulated |
| transcription | Down-regulated |
| other | Down-regulated |
| other | Down-regulated |
| enzyme | Down-regulated |
| other | Down-regulated |
| enzyme | Up-regulated |
| other | Down-regulated |
| transcription | Down-regulated |
| enzyme | Up-regulated |
| enzyme | Down-regulated |
| enzyme | Down-regulated |
| other | Down-regulated |
| transcription | Down-regulated |
| other | Up-regulated |
| other | Down-regulated |
| other | Down-regulated |
| other | Down-regulated |
| transcription | Up-regulated |
| other | Down-regulated |
| transporter | Down-regulated |
| other | Up-regulated |
| other | Down-regulated |
| other | Down-regulated |
| other | Down-regulated |
| kinase | Down-regulated |
| other | Down-regulated |
| enzyme | Down-regulated |
| other | Down-regulated |
| other | Down-regulated |
| other | Down-regulated |
| other | Down-regulated |
| enzyme | Up-regulated |
| transcription | Down-regulated |
| other | Down-regulated |
| transporter | Down-regulated |
| other | Down-regulated |
| phosphatase | Down-regulated |
| other | Down-regulated |
| other | Down-regulated |
| enzyme | Down-regulated |
| enzyme | Up-regulated |
| other | Down-regulated |
| other | Down-regulated |
| enzyme | Down-regulated |
| enzyme | Down-regulated |
| other | Down-regulated |
| peptidase | Down-regulated |
| other | Down-regulated |
| other | Up-regulated |
| other | Down-regulated |

| | |
|---|---|
| transcription | Down-regulated |
| enzyme | Down-regulated |
| other | Down-regulated |
| transporter | Up-regulated |
| other | Down-regulated |
| other | Down-regulated |
| other | Down-regulated |
| kinase | Down-regulated |
| other | Down-regulated |
| transporter | Down-regulated |
| other | Down-regulated |
| other | Down-regulated |
| other | Down-regulated |
| other | Down-regulated |
| transcription | Down-regulated |
| other | Down-regulated |
| other | Up-regulated |
| transporter | Up-regulated |
| enzyme | Up-regulated |
| other | Down-regulated |
| phosphatase | Down-regulated |
| other | Down-regulated |
| enzyme | Down-regulated |
| other | Down-regulated |
| enzyme | Down-regulated |
| other | Down-regulated |
| transporter | Down-regulated |
| other | Down-regulated |
| enzyme | Up-regulated |
| other | Down-regulated |
| enzyme | Down-regulated |
| other | Down-regulated |
| transmembra | Down-regulated |
| transporter | Down-regulated |
| other | Down-regulated |
| other | Up-regulated |
| other | Down-regulated |
| G-protein co | Up-regulated |
| other | Down-regulated |
| other | Down-regulated |
| enzyme | Down-regulated |
| other | Down-regulated |
| other | Down-regulated |
| other | Down-regulated |
| G-protein co | Down-regulated |
| other | Down-regulated |
| other | Down-regulated |
| transcription | Down-regulated |
| enzyme | Down-regulated |
| other | Down-regulated |
| enzyme | Up-regulated |
| other | Down-regulated |
| enzyme | Down-regulated |
| transcription | Down-regulated |
| enzyme | Down-regulated |
| other | Down-regulated |
| other | Down-regulated |
| transporter | Up-regulated |

| | |
|---|---|
| transporter | Down-regulated |
| transporter | Up-regulated |
| enzyme | Down-regulated |
| transporter | Down-regulated |
| enzyme | Down-regulated |
| other | Down-regulated |
| other | Down-regulated |
| transcription | Down-regulated |
| kinase | Down-regulated |
| other | Down-regulated |
| other | Up-regulated |
| other | Down-regulated |
| other | Down-regulated |
| other | Down-regulated |
| other | Down-regulated |
| enzyme | Down-regulated |
| enzyme | Down-regulated |
| other | Down-regulated |
| other | Down-regulated |
| other | Down-regulated |
| ion channel | Up-regulated |
| other | Down-regulated |
| other | Down-regulated |
| other | Down-regulated |
| other | Down-regulated |
| enzyme | Down-regulated |
| other | Down-regulated |
| enzyme | Down-regulated |
| other | Down-regulated |
| enzyme | Down-regulated |
| transporter | Down-regulated |
| other | Down-regulated |
| transporter | Down-regulated |
| transcription | Down-regulated |
| other | Down-regulated |
| other | Down-regulated |
| kinase | Down-regulated |
| enzyme | Up-regulated |
| other | Down-regulated |
| other | Down-regulated |
| other | Down-regulated |
| enzyme | Down-regulated |
| transmembra | Up-regulated |
| transporter | Up-regulated |
| enzyme | Down-regulated |
| other | Down-regulated |
| enzyme | Down-regulated |
| other | Down-regulated |
| other | Down-regulated |
| other | Down-regulated |
| other | Down-regulated |
| enzyme | Up-regulated |
| enzyme | Down-regulated |
| other | Up-regulated |
| enzyme | Down-regulated |
| other | Down-regulated |
| enzyme | Down-regulated |
| ligand-deper | Down-regulated |

| | |
|---|---|
| enzyme | Up-regulated |
| kinase | Down-regulated |
| other | Down-regulated |
| transcription | Down-regulated |
| transcription | Down-regulated |
| ligand-deper | Down-regulated |
| enzyme | Down-regulated |
| peptidase | Down-regulated |
| other | Down-regulated |
| other | Down-regulated |
| enzyme | Down-regulated |
| transcription | Down-regulated |
| other | Down-regulated |
| other | Down-regulated |
| transporter | Down-regulated |
| transporter | Down-regulated |
| transcription | Down-regulated |
| enzyme | Down-regulated |
| enzyme | Down-regulated |
| transporter | Up-regulated |
| other | Down-regulated |
| enzyme | Down-regulated |
| other | Down-regulated |
| other | Down-regulated |
| transcription | Up-regulated |
| other | Down-regulated |
| transporter | Down-regulated |
| transporter | Down-regulated |
| transcription | Down-regulated |
| kinase | Down-regulated |
| other | Up-regulated |
| phosphatase | Up-regulated |
| peptidase | Down-regulated |
| transcription | Down-regulated |
| other | Down-regulated |
| growth facto | Up-regulated |
| other | Down-regulated |
| translation re | Down-regulated |
| transmembra | Down-regulated |
| transcription | Down-regulated |
| enzyme | Down-regulated |
| other | Down-regulated |
| transporter | Down-regulated |
| other | Up-regulated |
| other | Down-regulated |
| enzyme | Down-regulated |
| other | Down-regulated |
| other | Down-regulated |
| enzyme | Down-regulated |
| transcription | Down-regulated |
| other | Down-regulated |
| other | Down-regulated |
| other | Down-regulated |
| other | Down-regulated |
| other | Down-regulated |
| other | Down-regulated |
| other | Down-regulated |
| other | Down-regulated |

| | |
|---|---|
| G-protein co | Up-regulated |
| enzyme | Down-regulated |
| other | Down-regulated |
| enzyme | Down-regulated |
| other | Down-regulated |
| other | Down-regulated |
| other | Down-regulated |
| enzyme | Down-regulated |
| other | Down-regulated |
| other | Up-regulated |
| transporter | Down-regulated |
| enzyme | Down-regulated |
| other | Down-regulated |
| enzyme | Down-regulated |
| other | Down-regulated |
| other | Down-regulated |
| enzyme | Down-regulated |
| enzyme | Down-regulated |
| enzyme | Down-regulated |
| other | Down-regulated |
| transporter | Down-regulated |
| transcription | Down-regulated |
| other | Down-regulated |
| other | Down-regulated |
| other | Down-regulated |
| peptidase | Down-regulated |
| enzyme | Down-regulated |
| enzyme | Up-regulated |
| enzyme | Down-regulated |
| transcription | Down-regulated |
| other | Down-regulated |
| enzyme | Up-regulated |
| other | Down-regulated |
| other | Up-regulated |
| G-protein co | Down-regulated |
| transporter | Down-regulated |
| enzyme | Down-regulated |
| other | Down-regulated |
| other | Down-regulated |
| other | Down-regulated |
| transcription | Down-regulated |
| other | Up-regulated |
| transporter | Down-regulated |
| enzyme | Down-regulated |
| transcription | Down-regulated |
| enzyme | Down-regulated |
| other | Down-regulated |
| enzyme | Down-regulated |
| enzyme | Down-regulated |
| other | Down-regulated |
| enzyme | Down-regulated |
| other | Up-regulated |
| transcription | Down-regulated |
| other | Down-regulated |
| cytokine | Down-regulated |
| other | Down-regulated |
| other | Down-regulated |
| transporter | Down-regulated |

| | |
|---|---|
| transmembra | Down-regulated |
| other | Down-regulated |
| kinase | Down-regulated |
| other | Down-regulated |
| other | Down-regulated |
| other | Down-regulated |
| enzyme | Down-regulated |
| kinase | Down-regulated |
| other | Down-regulated |
| transcription | Down-regulated |
| other | Up-regulated |
| other | Down-regulated |
| other | Down-regulated |
| enzyme | Down-regulated |
| other | Down-regulated |
| other | Down-regulated |
| transcription | Down-regulated |
| enzyme | Down-regulated |
| other | Down-regulated |
| other | Down-regulated |
| enzyme | Up-regulated |
| kinase | Down-regulated |
| other | Up-regulated |
| other | Down-regulated |
| transporter | Down-regulated |
| enzyme | Down-regulated |
| other | Down-regulated |
| other | Down-regulated |
| other | Down-regulated |
| transcription | Down-regulated |
| other | Up-regulated |
| transcription | Down-regulated |
| other | Down-regulated |
| enzyme | Up-regulated |
| enzyme | Up-regulated |
| transcription | Down-regulated |
| transcription | Down-regulated |
| transporter | Down-regulated |
| transporter | Down-regulated |
| kinase | Down-regulated |
| other | Down-regulated |
| transcription | Down-regulated |
| other | Down-regulated |
| other | Down-regulated |
| kinase | Down-regulated |
| enzyme | Down-regulated |
| enzyme | Down-regulated |
| other | Down-regulated |
| other | Down-regulated |
| enzyme | Down-regulated |
| enzyme | Up-regulated |
| transporter | Down-regulated |
| phosphatase | Down-regulated |
| enzyme | Down-regulated |
| kinase | Down-regulated |
| transmembra | Down-regulated |
| other | Down-regulated |
| other | Down-regulated |

| | |
|---|---|
| other | Down-regulated |
| ion channel | Down-regulated |
| other | Down-regulated |
| peptidase | Down-regulated |
| other | Down-regulated |
| enzyme | Up-regulated |
| kinase | Down-regulated |
| other | Down-regulated |
| other | Up-regulated |
| transcription | Down-regulated |
| other | Down-regulated |
| other | Down-regulated |
| enzyme | Up-regulated |
| enzyme | Up-regulated |
| transcription | Down-regulated |
| transcription | Up-regulated |
| transcription | Down-regulated |
| phosphatase | Up-regulated |
| transcription | Down-regulated |
| transcription | Down-regulated |
| other | Down-regulated |
| other | Down-regulated |
| transcription | Down-regulated |
| enzyme | Down-regulated |
| enzyme | Up-regulated |
| enzyme | Down-regulated |
| enzyme | Down-regulated |
| transporter | Down-regulated |
| other | Down-regulated |
| kinase | Down-regulated |
| other | Down-regulated |
| other | Down-regulated |
| enzyme | Down-regulated |
| enzyme | Down-regulated |
| transcription | Down-regulated |
| enzyme | Down-regulated |
| other | Down-regulated |
| transcription | Down-regulated |
| other | Up-regulated |
| transporter | Down-regulated |
| phosphatase | Down-regulated |
| other | Down-regulated |
| ion channel | Up-regulated |
| transcription | Up-regulated |
| other | Down-regulated |
| other | Down-regulated |
| enzyme | Down-regulated |
| other | Down-regulated |
| other | Down-regulated |
| enzyme | Down-regulated |
| transporter | Down-regulated |
| other | Down-regulated |
| enzyme | Down-regulated |
| other | Down-regulated |
| peptidase | Up-regulated |
| other | Down-regulated |
| other | Down-regulated |
| other | Down-regulated |

| | |
|---|---|
| other | Down-regulated |
| enzyme | Down-regulated |
| other | Down-regulated |
| enzyme | Up-regulated |
| enzyme | Down-regulated |
| phosphatase | Down-regulated |
| other | Down-regulated |
| enzyme | Down-regulated |
| other | Down-regulated |
| kinase | Down-regulated |
| transcription | Down-regulated |
| other | Up-regulated |
| other | Down-regulated |
| enzyme | Up-regulated |
| other | Down-regulated |
| other | Down-regulated |
| kinase | Down-regulated |
| other | Down-regulated |
| other | Down-regulated |
| other | Down-regulated |
| enzyme | Down-regulated |
| other | Down-regulated |
| enzyme | Down-regulated |
| transmembra | Up-regulated |
| peptidase | Down-regulated |
| other | Down-regulated |
| other | Down-regulated |
| transcription | Down-regulated |
| other | Up-regulated |
| kinase | Up-regulated |
| peptidase | Down-regulated |
| transcription | Down-regulated |
| peptidase | Down-regulated |
| enzyme | Down-regulated |
| other | Down-regulated |
| enzyme | Down-regulated |
| enzyme | Down-regulated |
| phosphatase | Down-regulated |
| phosphatase | Down-regulated |
| other | Down-regulated |
| other | Down-regulated |
| transcription | Down-regulated |
| other | Down-regulated |
| transporter | Down-regulated |
| peptidase | Down-regulated |
| transporter | Down-regulated |
| other | Down-regulated |
| other | Down-regulated |
| other | Down-regulated |
| transcription | Down-regulated |
| transporter | Down-regulated |
| enzyme | Down-regulated |
| transcription | Down-regulated |
| other | Down-regulated |
| enzyme | Down-regulated |
| enzyme | Down-regulated |
| other | Down-regulated |
| enzyme | Up-regulated |

| | |
|---|---|
| peptidase | Down-regulated |
| peptidase | Down-regulated |
| enzyme | Down-regulated |
| transcription | Down-regulated |
| other | Down-regulated |
| other | Down-regulated |
| other | Up-regulated |
| enzyme | Down-regulated |
| other | Up-regulated |
| other | Down-regulated |
| transcription | Up-regulated |
| peptidase | Down-regulated |
| other | Down-regulated |
| other | Down-regulated |
| other | Down-regulated |
| enzyme | Down-regulated |
| other | Up-regulated |
| other | Up-regulated |
| other | Down-regulated |
| enzyme | Down-regulated |
| transcription | Down-regulated |
| other | Down-regulated |
| other | Down-regulated |
| kinase | Down-regulated |
| enzyme | Up-regulated |
| other | Up-regulated |
| enzyme | Down-regulated |
| other | Up-regulated |
| transporter | Down-regulated |
| transporter | Down-regulated |
| transcription | Down-regulated |
| transporter | Down-regulated |
| transporter | Down-regulated |
| other | Down-regulated |
| other | Down-regulated |
| enzyme | Down-regulated |
| other | Down-regulated |
| other | Down-regulated |
| enzyme | Up-regulated |
| other | Down-regulated |
| other | Down-regulated |
| enzyme | Down-regulated |
| other | Down-regulated |
| other | Down-regulated |
| enzyme | Up-regulated |
| transcription | Up-regulated |
| transporter | Down-regulated |
| enzyme | Down-regulated |
| other | Up-regulated |
| other | Down-regulated |
| other | Down-regulated |
| other | Down-regulated |
| transcription | Down-regulated |
| other | Down-regulated |
| transcription | Down-regulated |
| enzyme | Down-regulated |
| enzyme | Up-regulated |
| other | Down-regulated |

| | |
|---|---|
| enzyme | Down-regulated |
| transcription | Down-regulated |
| other | Down-regulated |
| other | Down-regulated |
| transcription | Down-regulated |
| other | Down-regulated |
| other | Down-regulated |
| other | Down-regulated |
| other | Up-regulated |
| other | Up-regulated |
| translation re | Down-regulated |
| enzyme | Down-regulated |
| enzyme | Down-regulated |
| enzyme | Down-regulated |
| peptidase | Down-regulated |
| enzyme | Down-regulated |
| enzyme | Down-regulated |
| other | Down-regulated |
| other | Down-regulated |
| other | Down-regulated |
| other | Down-regulated |
| other | Down-regulated |
| other | Down-regulated |
| other | Down-regulated |
| phosphatase | Up-regulated |
| transmembra | Down-regulated |
| enzyme | Down-regulated |
| other | Down-regulated |
| enzyme | Up-regulated |
| enzyme | Up-regulated |
| other | Down-regulated |
| transcription | Up-regulated |
| enzyme | Down-regulated |
| enzyme | Down-regulated |
| other | Down-regulated |
| growth facto | Down-regulated |
| other | Down-regulated |
| transporter | Down-regulated |
| other | Down-regulated |
| other | Down-regulated |
| other | Down-regulated |
| enzyme | Down-regulated |
| enzyme | Down-regulated |
| other | Down-regulated |
| other | Down-regulated |
| enzyme | Down-regulated |
| other | Down-regulated |
| transcription | Down-regulated |
| enzyme | Down-regulated |
| other | Down-regulated |
| transcription | Down-regulated |
| kinase | Down-regulated |
| other | Down-regulated |
| enzyme | Down-regulated |
| other | Down-regulated |
| enzyme | Down-regulated |
| transcription | Down-regulated |
| other | Down-regulated |

| | |
|---|---|
| other | Down-regulated |
| other | Down-regulated |
| other | Down-regulated |
| other | Down-regulated |
| other | Down-regulated |
| other | Down-regulated |
| other | Up-regulated |
| enzyme | Down-regulated |
| transporter | Down-regulated |
| other | Down-regulated |
| other | Down-regulated |
| transcription | Down-regulated |
| G-protein co | Up-regulated |
| kinase | Down-regulated |
| other | Down-regulated |
| other | Down-regulated |
| transcription | Down-regulated |
| other | Down-regulated |
| transcription | Down-regulated |
| other | Down-regulated |
| enzyme | Down-regulated |
| other | Down-regulated |
| other | Down-regulated |
| other | Down-regulated |
| other | Down-regulated |
| transporter | Down-regulated |
| peptidase | Up-regulated |
| other | Down-regulated |
| transporter | Down-regulated |
| enzyme | Down-regulated |
| other | Down-regulated |
| enzyme | Down-regulated |
| other | Down-regulated |
| enzyme | Down-regulated |
| growth facto | Up-regulated |
| other | Down-regulated |
| enzyme | Up-regulated |
| transporter | Up-regulated |
| other | Down-regulated |
| kinase | Down-regulated |
| other | Down-regulated |
| other | Down-regulated |
| kinase | Down-regulated |
| other | Down-regulated |
| enzyme | Down-regulated |
| enzyme | Down-regulated |
| other | Down-regulated |
| other | Up-regulated |
| other | Down-regulated |
| transcription | Down-regulated |
| peptidase | Down-regulated |
| other | Down-regulated |
| kinase | Down-regulated |
| kinase | Down-regulated |
| enzyme | Down-regulated |
| other | Down-regulated |
| other | Down-regulated |
| transcription | Down-regulated |

| | |
|---|---|
| other | Down-regulated |
| enzyme | Down-regulated |
| enzyme | Down-regulated |
| other | Down-regulated |
| transcription | Down-regulated |
| enzyme | Down-regulated |
| ligand-deper | Down-regulated |
| enzyme | Up-regulated |
| enzyme | Down-regulated |
| other | Down-regulated |
| other | Down-regulated |
| other | Down-regulated |
| enzyme | Down-regulated |
| peptidase | Down-regulated |
| transcription | Down-regulated |
| other | Down-regulated |
| other | Down-regulated |
| transporter | Down-regulated |
| other | Down-regulated |
| other | Down-regulated |
| enzyme | Down-regulated |
| other | Down-regulated |
| translation re | Down-regulated |
| enzyme | Down-regulated |
| enzyme | Down-regulated |
| other | Down-regulated |
| transporter | Up-regulated |
| transcription | Down-regulated |
| enzyme | Down-regulated |
| other | Down-regulated |
| enzyme | Up-regulated |
| transporter | Down-regulated |
| transcription | Down-regulated |
| other | Down-regulated |
| transcription | Down-regulated |
| other | Down-regulated |
| translation re | Down-regulated |
| other | Down-regulated |
| transcription | Down-regulated |
| other | Down-regulated |
| other | Down-regulated |
| cytokine | Down-regulated |
| transporter | Down-regulated |
| kinase | Up-regulated |
| other | Down-regulated |
| kinase | Up-regulated |
| transporter | Down-regulated |
| transmembra | Up-regulated |
| other | Down-regulated |
| other | Down-regulated |
| transcription | Down-regulated |
| phosphatase | Down-regulated |
| other | Down-regulated |
| enzyme | Down-regulated |
| enzyme | Down-regulated |
| other | Up-regulated |
| kinase | Down-regulated |
| peptidase | Down-regulated |

| | |
|---|---|
| other | Down-regulated |
| kinase | Down-regulated |
| other | Down-regulated |
| enzyme | Down-regulated |
| transporter | Down-regulated |
| transcription | Down-regulated |
| other | Down-regulated |
| peptidase | Down-regulated |
| other | Up-regulated |
| cytokine | Down-regulated |
| kinase | Down-regulated |
| other | Down-regulated |
| enzyme | Down-regulated |
| enzyme | Down-regulated |
| other | Down-regulated |
| other | Up-regulated |
| other | Down-regulated |
| transcription | Down-regulated |
| transcription | Up-regulated |
| kinase | Down-regulated |
| other | Down-regulated |
| kinase | Up-regulated |
| other | Down-regulated |
| other | Down-regulated |
| transcription | Down-regulated |
| other | Down-regulated |
| transcription | Down-regulated |
| enzyme | Down-regulated |
| transporter | Down-regulated |
| other | Down-regulated |
| other | Down-regulated |
| transporter | Up-regulated |
| other | Up-regulated |
| other | Down-regulated |
| other | Down-regulated |
| other | Up-regulated |
| enzyme | Down-regulated |
| other | Up-regulated |
| enzyme | Down-regulated |
| enzyme | Up-regulated |
| other | Down-regulated |
| enzyme | Down-regulated |
| transcription | Down-regulated |
| enzyme | Up-regulated |
| other | Down-regulated |
| other | Down-regulated |
| enzyme | Up-regulated |
| translation re | Down-regulated |
| other | Down-regulated |
| phosphatase | Down-regulated |
| enzyme | Up-regulated |
| transporter | Up-regulated |
| transcription | Down-regulated |
| other | Down-regulated |
| other | Down-regulated |
| enzyme | Down-regulated |
| enzyme | Down-regulated |
| peptidase | Down-regulated |

| | |
|---|---|
| kinase | Down-regulated |
| transporter | Down-regulated |
| transcription | Down-regulated |
| other | Down-regulated |
| enzyme | Up-regulated |
| enzyme | Up-regulated |
| transcription | Down-regulated |
| other | Down-regulated |
| enzyme | Down-regulated |
| other | Down-regulated |
| ion channel | Up-regulated |
| other | Down-regulated |
| transporter | Down-regulated |
| other | Down-regulated |
| other | Down-regulated |
| ion channel | Down-regulated |
| transporter | Down-regulated |
| enzyme | Down-regulated |
| transporter | Up-regulated |
| other | Down-regulated |
| enzyme | Down-regulated |
| transcription | Down-regulated |
| kinase | Down-regulated |
| enzyme | Down-regulated |
| enzyme | Down-regulated |
| kinase | Up-regulated |
| other | Down-regulated |
| other | Down-regulated |
| other | Down-regulated |
| other | Down-regulated |
| enzyme | Down-regulated |
| other | Up-regulated |
| kinase | Down-regulated |
| transcription | Down-regulated |
| other | Down-regulated |
| transcription | Down-regulated |
| other | Down-regulated |
| transcription | Up-regulated |
| other | Down-regulated |
| other | Down-regulated |
| transcription | Down-regulated |
| transcription | Down-regulated |
| other | Up-regulated |
| transcription | Down-regulated |
| other | Down-regulated |
| other | Up-regulated |
| enzyme | Down-regulated |
| transporter | Down-regulated |
| enzyme | Down-regulated |
| enzyme | Down-regulated |
| other | Up-regulated |
| enzyme | Down-regulated |
| other | Down-regulated |
| G-protein co | Down-regulated |
| enzyme | Down-regulated |
| transcription | Down-regulated |
| transporter | Down-regulated |
| enzyme | Down-regulated |

| | |
|---|---|
| enzyme | Down-regulated |
| transcription | Down-regulated |
| other | Down-regulated |
| enzyme | Down-regulated |
| other | Down-regulated |
| other | Down-regulated |
| transcription | Down-regulated |
| enzyme | Down-regulated |
| enzyme | Down-regulated |
| other | Down-regulated |
| transcription | Down-regulated |
| kinase | Down-regulated |
| transcription | Down-regulated |
| enzyme | Down-regulated |
| other | Down-regulated |
| enzyme | Up-regulated |
| enzyme | Down-regulated |
| other | Down-regulated |
| other | Down-regulated |
| enzyme | Down-regulated |
| other | Down-regulated |
| enzyme | Down-regulated |
| other | Down-regulated |
| enzyme | Down-regulated |
| other | Down-regulated |
| transcription | Down-regulated |
| peptidase | Down-regulated |
| transporter | Down-regulated |
| enzyme | Down-regulated |
| enzyme | Down-regulated |
| other | Down-regulated |
| enzyme | Up-regulated |
| kinase | Down-regulated |
| enzyme | Down-regulated |
| transcription | Down-regulated |
| enzyme | Down-regulated |
| other | Down-regulated |
| kinase | Down-regulated |
| other | Down-regulated |
| enzyme | Down-regulated |
| enzyme | Down-regulated |
| phosphatase | Down-regulated |
| enzyme | Up-regulated |
| other | Up-regulated |
| transporter | Down-regulated |
| kinase | Up-regulated |
| transcription | Down-regulated |
| other | Down-regulated |
| enzyme | Down-regulated |
| other | Down-regulated |
| other | Down-regulated |
| enzyme | Down-regulated |
| transcription | Down-regulated |
| transporter | Down-regulated |
| phosphatase | Down-regulated |
| other | Down-regulated |
| transporter | Down-regulated |
| enzyme | Down-regulated |

| | |
|---|---|
| other | Down-regulated |
| enzyme | Up-regulated |
| transporter | Down-regulated |
| enzyme | Down-regulated |
| kinase | Down-regulated |
| other | Down-regulated |
| other | Down-regulated |
| kinase | Down-regulated |
| phosphatase | Down-regulated |
| enzyme | Down-regulated |
| other | Down-regulated |
| transporter | Down-regulated |
| transcription | Up-regulated |
| enzyme | Down-regulated |
| other | Down-regulated |
| other | Down-regulated |
| transporter | Down-regulated |
| other | Down-regulated |
| transcription | Down-regulated |
| other | Down-regulated |
| enzyme | Up-regulated |
| other | Down-regulated |
| enzyme | Down-regulated |
| other | Down-regulated |
| other | Down-regulated |
| other | Up-regulated |
| transcription | Down-regulated |
| enzyme | Down-regulated |
| other | Down-regulated |
| other | Down-regulated |
| transporter | Down-regulated |
| phosphatase | Up-regulated |
| other | Down-regulated |
| transcription | Down-regulated |
| transmembra | Up-regulated |
| peptidase | Down-regulated |
| other | Down-regulated |
| transcription | Down-regulated |
| other | Down-regulated |
| other | Down-regulated |
| other | Down-regulated |
| kinase | Down-regulated |
| G-protein co | Down-regulated |
| other | Down-regulated |
| transcription | Down-regulated |
| other | Down-regulated |
| kinase | Down-regulated |
| other | Down-regulated |
| other | Down-regulated |
| other | Down-regulated |
| transporter | Down-regulated |
| phosphatase | Up-regulated |
| enzyme | Down-regulated |
| other | Down-regulated |
| transcription | Down-regulated |
| kinase | Down-regulated |
| other | Down-regulated |
| other | Down-regulated |

| | |
|---|---|
| other | Down-regulated |
| other | Down-regulated |
| transcription | Down-regulated |
| transporter | Down-regulated |
| enzyme | Down-regulated |
| enzyme | Up-regulated |
| phosphatase | Down-regulated |
| enzyme | Up-regulated |
| enzyme | Up-regulated |
| enzyme | Down-regulated |
| kinase | Down-regulated |
| enzyme | Down-regulated |
| transcription | Down-regulated |
| enzyme | Down-regulated |
| other | Up-regulated |
| kinase | Down-regulated |
| phosphatase | Up-regulated |
| other | Down-regulated |
| peptidase | Down-regulated |
| other | Down-regulated |
| enzyme | Down-regulated |
| other | Down-regulated |
| enzyme | Down-regulated |
| other | Down-regulated |
| enzyme | Down-regulated |
| enzyme | Down-regulated |
| ligand-deper | Up-regulated |
| G-protein co | Down-regulated |
| other | Down-regulated |
| other | Up-regulated |
| other | Down-regulated |
| transporter | Down-regulated |
| other | Down-regulated |
| other | Down-regulated |
| transporter | Down-regulated |
| other | Down-regulated |
| other | Down-regulated |
| transcription | Down-regulated |
| other | Down-regulated |
| enzyme | Down-regulated |
| other | Down-regulated |
| transcription | Down-regulated |
| enzyme | Down-regulated |
| peptidase | Up-regulated |
| ligand-deper | Down-regulated |
| other | Down-regulated |
| other | Down-regulated |
| other | Down-regulated |
| kinase | Down-regulated |
| enzyme | Down-regulated |
| other | Down-regulated |
| other | Down-regulated |
| enzyme | Down-regulated |
| other | Down-regulated |
| transcription | Down-regulated |
| transcription | Down-regulated |
| enzyme | Up-regulated |
| enzyme | Down-regulated |

| | |
|---|---|
| enzyme | Down-regulated |
| other | Down-regulated |
| enzyme | Up-regulated |
| other | Down-regulated |
| kinase | Down-regulated |
| enzyme | Down-regulated |
| other | Down-regulated |
| kinase | Down-regulated |
| enzyme | Up-regulated |
| transcription | Down-regulated |
| kinase | Down-regulated |
| other | Down-regulated |
| transcription | Down-regulated |
| other | Down-regulated |
| other | Down-regulated |
| enzyme | Up-regulated |
| enzyme | Down-regulated |
| other | Down-regulated |
| transcription | Up-regulated |
| peptidase | Down-regulated |
| kinase | Down-regulated |
| kinase | Up-regulated |
| other | Down-regulated |
| transporter | Down-regulated |
| kinase | Down-regulated |
| phosphatase | Down-regulated |
| enzyme | Down-regulated |
| enzyme | Down-regulated |
| transcription | Down-regulated |
| other | Down-regulated |
| transcription | Down-regulated |
| other | Down-regulated |
| phosphatase | Down-regulated |
| enzyme | Down-regulated |
| transcription | Down-regulated |
| other | Down-regulated |
| kinase | Down-regulated |
| enzyme | Down-regulated |
| enzyme | Up-regulated |
| other | Down-regulated |
| other | Down-regulated |
| other | Up-regulated |
| other | Down-regulated |
| transmembra | Down-regulated |
| transporter | Down-regulated |
| other | Down-regulated |
| other | Down-regulated |
| enzyme | Up-regulated |
| other | Down-regulated |
| transmembra | Down-regulated |
| other | Down-regulated |
| enzyme | Down-regulated |
| other | Down-regulated |
| peptidase | Up-regulated |
| kinase | Down-regulated |
| enzyme | Down-regulated |
| peptidase | Down-regulated |
| enzyme | Down-regulated |

| | |
|---|---|
| other | Down-regulated |
| transcription | Down-regulated |
| transporter | Down-regulated |
| phosphatase | Down-regulated |
| other | Down-regulated |
| other | Down-regulated |
| enzyme | Down-regulated |
| transcription | Down-regulated |
| other | Down-regulated |
| enzyme | Up-regulated |
| other | Down-regulated |
| translation re | Down-regulated |
| transporter | Down-regulated |
| transcription | Down-regulated |
| enzyme | Up-regulated |
| enzyme | Down-regulated |
| other | Down-regulated |
| other | Up-regulated |
| other | Down-regulated |
| transcription | Down-regulated |
| transmembra | Up-regulated |
| enzyme | Down-regulated |
| other | Down-regulated |
| transmembra | Up-regulated |
| transporter | Down-regulated |
| peptidase | Up-regulated |
| other | Down-regulated |
| other | Up-regulated |
| other | Down-regulated |
| other | Down-regulated |
| ion channel | Down-regulated |
| enzyme | Down-regulated |
| other | Up-regulated |
| enzyme | Down-regulated |
| enzyme | Down-regulated |
| other | Down-regulated |
| enzyme | Down-regulated |
| other | Down-regulated |
| enzyme | Down-regulated |
| enzyme | Down-regulated |
| enzyme | Up-regulated |
| other | Down-regulated |
| G-protein co | Down-regulated |
| phosphatase | Down-regulated |
| enzyme | Down-regulated |
| other | Down-regulated |
| enzyme | Down-regulated |
| enzyme | Down-regulated |
| other | Down-regulated |
| ion channel | Down-regulated |
| enzyme | Up-regulated |
| other | Down-regulated |
| enzyme | Down-regulated |
| other | Down-regulated |
| other | Down-regulated |
| enzyme | Down-regulated |
| transporter | Down-regulated |
| transcription | Up-regulated |

| | |
|---|---|
| other | Up-regulated |
| other | Up-regulated |
| other | Down-regulated |
| transporter | Down-regulated |
| other | Down-regulated |
| other | Down-regulated |
| other | Down-regulated |
| transporter | Down-regulated |
| enzyme | Down-regulated |
| other | Down-regulated |
| transporter | Down-regulated |
| other | Down-regulated |
| other | Down-regulated |
| enzyme | Up-regulated |
| enzyme | Down-regulated |
| enzyme | Down-regulated |
| enzyme | Down-regulated |
| enzyme | Down-regulated |
| transporter | Up-regulated |
| transporter | Down-regulated |
| other | Down-regulated |
| enzyme | Up-regulated |
| enzyme | Down-regulated |
| phosphatase | Down-regulated |
| other | Up-regulated |
| other | Down-regulated |
| transporter | Down-regulated |
| other | Down-regulated |
| transporter | Down-regulated |
| enzyme | Down-regulated |
| transcription | Down-regulated |
| other | Down-regulated |
| other | Down-regulated |
| enzyme | Down-regulated |
| other | Down-regulated |
| enzyme | Down-regulated |
| other | Down-regulated |
| enzyme | Down-regulated |
| transporter | Down-regulated |
| transcription | Down-regulated |
| transmembra | Down-regulated |
| enzyme | Down-regulated |
| enzyme | Down-regulated |
| enzyme | Down-regulated |
| other | Down-regulated |
| transcription | Down-regulated |
| phosphatase | Up-regulated |
| peptidase | Up-regulated |
| other | Down-regulated |
| transcription | Down-regulated |
| ion channel | Down-regulated |
| enzyme | Up-regulated |
| transcription | Down-regulated |
| other | Up-regulated |
| transcription | Up-regulated |
| enzyme | Down-regulated |
| enzyme | Down-regulated |
| transcription | Down-regulated |

| | |
|---|---|
| transcription | Down-regulated |
| transporter | Down-regulated |
| other | Down-regulated |
| enzyme | Up-regulated |
| other | Up-regulated |
| transporter | Down-regulated |
| enzyme | Down-regulated |
| enzyme | Down-regulated |
| other | Up-regulated |
| ligand-deper | Up-regulated |
| enzyme | Down-regulated |
| transporter | Up-regulated |
| enzyme | Up-regulated |
| enzyme | Down-regulated |
| peptidase | Down-regulated |
| enzyme | Down-regulated |
| other | Down-regulated |
| other | Down-regulated |
| transporter | Down-regulated |
| enzyme | Down-regulated |
| enzyme | Down-regulated |
| other | Down-regulated |
| phosphatase | Down-regulated |
| other | Down-regulated |
| enzyme | Up-regulated |
| enzyme | Up-regulated |
| enzyme | Down-regulated |
| other | Down-regulated |
| enzyme | Down-regulated |
| enzyme | Down-regulated |
| transporter | Down-regulated |
| transporter | Down-regulated |
| other | Down-regulated |
| kinase | Down-regulated |
| peptidase | Down-regulated |
| transmembra | Up-regulated |
| enzyme | Down-regulated |
| other | Down-regulated |
| other | Down-regulated |
| other | Down-regulated |
| enzyme | Down-regulated |
| other | Down-regulated |
| other | Down-regulated |
| transcription | Down-regulated |
| kinase | Up-regulated |
| other | Down-regulated |
| other | Down-regulated |
| other | Down-regulated |
| other | Down-regulated |
| transporter | Down-regulated |
| enzyme | Down-regulated |
| other | Down-regulated |
| growth facto | Up-regulated |
| other | Down-regulated |
| other | Down-regulated |
| enzyme | Up-regulated |
| enzyme | Up-regulated |
| transcription | Down-regulated |

| | |
|---|---|
| enzyme | Down-regulated |
| enzyme | Down-regulated |
| enzyme | Down-regulated |
| peptidase | Down-regulated |
| enzyme | Up-regulated |
| other | Down-regulated |
| other | Down-regulated |
| other | Down-regulated |
| other | Down-regulated |
| transcription | Down-regulated |
| other | Down-regulated |
| enzyme | Up-regulated |
| other | Down-regulated |
| transcription | Up-regulated |
| transcription | Down-regulated |
| transporter | Down-regulated |
| transporter | Up-regulated |
| enzyme | Down-regulated |
| other | Down-regulated |
| enzyme | Down-regulated |
| other | Down-regulated |
| other | Down-regulated |
| other | Down-regulated |
| enzyme | Down-regulated |
| transcription | Up-regulated |
| enzyme | Down-regulated |
| transcription | Down-regulated |
| other | Up-regulated |
| transporter | Down-regulated |
| transcription | Down-regulated |
| other | Up-regulated |
| enzyme | Down-regulated |
| enzyme | Down-regulated |
| other | Down-regulated |
| enzyme | Up-regulated |
| enzyme | Down-regulated |
| transcription | Down-regulated |
| ion channel | Down-regulated |
| other | Down-regulated |
| other | Down-regulated |
| other | Down-regulated |
| G-protein co | Down-regulated |
| other | Down-regulated |
| other | Down-regulated |
| transporter | Down-regulated |
| transcription | Down-regulated |
| transcription | Down-regulated |
| transporter | Down-regulated |
| other | Down-regulated |
| transporter | Down-regulated |
| peptidase | Down-regulated |
| other | Down-regulated |
| phosphatase | Down-regulated |
| other | Down-regulated |
| other | Down-regulated |
| enzyme | Down-regulated |
| enzyme | Down-regulated |
| enzyme | Up-regulated |

| | |
|---|---|
| transporter | Down-regulated |
| transcription | Down-regulated |
| transporter | Down-regulated |
| other | Down-regulated |
| other | Down-regulated |
| other | Up-regulated |
| enzyme | Up-regulated |
| enzyme | Down-regulated |
| kinase | Down-regulated |
| enzyme | Up-regulated |
| kinase | Down-regulated |
| transcription | Down-regulated |
| peptidase | Down-regulated |
| other | Down-regulated |
| transporter | Down-regulated |
| peptidase | Down-regulated |
| enzyme | Down-regulated |
| other | Down-regulated |
| transcription | Down-regulated |
| enzyme | Down-regulated |
| other | Down-regulated |
| transporter | Down-regulated |
| enzyme | Up-regulated |
| enzyme | Down-regulated |
| other | Down-regulated |
| enzyme | Up-regulated |
| other | Down-regulated |
| transcription | Down-regulated |
| other | Down-regulated |
| enzyme | Down-regulated |
| peptidase | Down-regulated |
| other | Down-regulated |
| enzyme | Down-regulated |
| enzyme | Down-regulated |
| enzyme | Down-regulated |
| enzyme | Down-regulated |
| other | Down-regulated |
| transcription | Down-regulated |
| transcription | Down-regulated |
| enzyme | Down-regulated |
| enzyme | Down-regulated |
| enzyme | Down-regulated |
| transporter | Down-regulated |
| enzyme | Down-regulated |
| other | Down-regulated |
| other | Down-regulated |
| transcription | Down-regulated |
| other | Down-regulated |
| cytokine | Down-regulated |
| transporter | Down-regulated |
| other | Down-regulated |
| other | Down-regulated |
| transcription | Down-regulated |
| other | Down-regulated |
| enzyme | Down-regulated |
| other | Up-regulated |
| kinase | Down-regulated |
| translation re | Down-regulated |

| | |
|---|---|
| other | Down-regulated |
| other | Down-regulated |
| other | Up-regulated |
| enzyme | Up-regulated |
| other | Down-regulated |
| enzyme | Down-regulated |
| other | Down-regulated |
| enzyme | Down-regulated |
| enzyme | Up-regulated |
| translation re | Down-regulated |
| enzyme | Up-regulated |
| enzyme | Down-regulated |
| transcription | Down-regulated |
| other | Down-regulated |
| enzyme | Down-regulated |
| other | Down-regulated |
| transporter | Down-regulated |
| kinase | Down-regulated |
| other | Down-regulated |
| other | Up-regulated |
| other | Down-regulated |
| kinase | Down-regulated |
| transcription | Down-regulated |
| other | Up-regulated |
| transporter | Down-regulated |
| enzyme | Up-regulated |
| other | Down-regulated |
| transporter | Up-regulated |
| transporter | Up-regulated |
| enzyme | Down-regulated |
| enzyme | Down-regulated |
| enzyme | Down-regulated |
| phosphatase | Down-regulated |
| enzyme | Down-regulated |
| enzyme | Down-regulated |
| other | Down-regulated |
| other | Down-regulated |
| transcription | Down-regulated |
| other | Down-regulated |
| ion channel | Down-regulated |
| other | Down-regulated |
| kinase | Down-regulated |
| transcription | Down-regulated |
| transporter | Down-regulated |
| other | Down-regulated |
| other | Down-regulated |
| other | Down-regulated |
| other | Down-regulated |
| other | Down-regulated |
| other | Down-regulated |
| transcription | Down-regulated |
| other | Down-regulated |
| enzyme | Down-regulated |
| enzyme | Down-regulated |
| other | Down-regulated |
| transcription | Down-regulated |
| other | Down-regulated |
| other | Down-regulated |

| | |
|---|---|
| transcription | Down-regulated |
| other | Down-regulated |
| transporter | Down-regulated |
| other | Down-regulated |
| enzyme | Down-regulated |
| enzyme | Down-regulated |
| other | Down-regulated |
| kinase | Down-regulated |
| other | Down-regulated |
| other | Down-regulated |
| other | Down-regulated |
| enzyme | Up-regulated |
| transporter | Down-regulated |
| other | Up-regulated |
| enzyme | Down-regulated |
| other | Down-regulated |
| enzyme | Down-regulated |
| enzyme | Down-regulated |
| enzyme | Down-regulated |
| enzyme | Down-regulated |
| enzyme | Up-regulated |
| enzyme | Down-regulated |
| other | Down-regulated |
| enzyme | Down-regulated |
| kinase | Down-regulated |
| kinase | Down-regulated |
| other | Down-regulated |
| peptidase | Down-regulated |
| other | Down-regulated |
| other | Down-regulated |
| transporter | Down-regulated |
| other | Down-regulated |
| enzyme | Down-regulated |
| enzyme | Down-regulated |
| other | Down-regulated |
| enzyme | Down-regulated |
| enzyme | Down-regulated |
| other | Down-regulated |
| enzyme | Up-regulated |
| other | Down-regulated |
| other | Down-regulated |
| other | Down-regulated |
| other | Down-regulated |
| transporter | Up-regulated |
| enzyme | Down-regulated |
| enzyme | Up-regulated |
| transporter | Up-regulated |
| kinase | Up-regulated |
| other | Down-regulated |
| phosphatase | Down-regulated |
| enzyme | Down-regulated |
| transporter | Down-regulated |
| phosphatase | Up-regulated |
| other | Up-regulated |
| enzyme | Down-regulated |
| other | Down-regulated |
| enzyme | Down-regulated |
| phosphatase | Up-regulated |

| | |
|---|---|
| transporter | Down-regulated |
| enzyme | Up-regulated |
| phosphatase | Down-regulated |
| other | Down-regulated |
| kinase | Down-regulated |
| transcription | Down-regulated |
| enzyme | Down-regulated |
| other | Down-regulated |
| other | Down-regulated |
| enzyme | Up-regulated |
| transcription | Down-regulated |
| transporter | Down-regulated |
| enzyme | Up-regulated |
| other | Up-regulated |
| transporter | Down-regulated |
| other | Down-regulated |
| other | Up-regulated |
| other | Down-regulated |
| transporter | Up-regulated |
